# FGFR3 activating mutations induce luminal-like papillary bladder tumor formation and favor a male gender bias

**DOI:** 10.1101/2021.09.02.458778

**Authors:** Ming-Jun Shi, Aura Moreno-Vega, Jacqueline Fontugne, Xiang-Yu Meng, Florent Dufour, Aurélie Kamoun, Sia Viborg Lindskrog, Audrey Rapinat, Claire Dunois-Larde, May-Linda Lepage, Elodie Chapeaublanc, Olivier Levrel, Thierry Lebret, Anna Almeida, Aurélien De Reynies, Lars Dyrskjøt, Yves Allory, François Radvanyi, Isabelle Bernard-Pierrot

## Abstract

**Background:** FGFR3 mutations are among the most frequent genetic alterations in bladder cancer and are enriched in the luminal papillary subtype of muscle-invasive tumors (MIBC) and luminal-like classes 1 and 3 of non-MIBC. To study their oncogenic properties *in vivo*, we developed here a genetically engineered mouse (GEM) model expressing the most frequent FGFR3 mutation, FGFR3-S249C, in urothelial cells.

**Methods:** Bladder tumorigenesis was monitored in FGFR3-S249C mice. FGFR3 expression was assessed by RT-qPCR in the transgenic mice urothelium and in various human epithelia. Transcriptomic data were obtained from mouse bladder tumors and crossspecies comparisons were performed. Sex bias in FGFR3-mutated tumors was evaluated in our GEM model and in the TCGA and UROMOL cohorts of patients including 408 MIBC and 419 NMIBC, respectively. The association of androgen receptor (AR) activity, based on the expression of its target genes, with FGFR3 mutations was examined in these two cohorts. Binding of AR to its response element and AR phosphorylation in FGFR3-dependent cell lines were evaluated.

**Results:** FGFR3-S249C expression in the urothelium of mice induced spontaneous low-grade papillary bladder tumors resembling the human counterpart at the histological and transcriptomic levels. Mutant-FGFR3 expression levels impacted tumor formation incidence in mice and mutant-FGFR3-driven human tumors were restricted to epithelia presenting high normal expression levels of FGFR3. The known bladder cancer male gender bias, also found in our model, was even higher in human FGFR3-mutated compared to wild-type tumors and associated with a higher AR regulon activity considering gender adjustment. AR phosphorylation and regulon activity were modulated by FGFR3 in FGFR3-dependent models.

**Conclusions:** Mutant-FGFR3 is an oncogene per se, inducing bladder tumorigenesis. Patients with early stage bladder lesions could thus potentially benefit from FGFR3 targeting. Our results also reinforce the interest in elucidating the role of AR in bladder carcinogenesis, specifically in FGFR3-mutated driven tumors. Finally, our results suggest FGFR3 expression level in epithelium as a determinant for the FGFR3-driven tumors tissue specificity.

## Introduction

Bladder cancer (BCa) is the sixth most common cancer in men worldwide, with an even higher incidence in Western Europe and North America (4th most common cancer in men)[1]. At diagnosis, the majority of tumors are non-muscle-invasive urothelial carcinomas (NMIBC) (75%). In spite of their favorable prognosis, NMIBCs have a high recurrence rate (70%) and a subset progresses (10-15%) to the more aggressive form of disease, muscle-invasive bladder cancer (MIBC). Molecular classifications have been established in both NMIBC and MIBC in order to identify different biological processes to support patient stratification for more adapted therapies [2–7].

Fibroblast growth factor receptor 3 (FGFR3) is a receptor tyrosine kinase with frequent genetic alterations in BCa [3,5,8,9]. Point mutations (observed in 70 % of NMIBCs and 15% of MIBCs) or chromosomal translocations resulting in protein fusions (affecting 5% of MIBCs), lead to a constitutively active FGFR3. The oncogenic properties of an altered FGFR3 have been shown *in vitro* and an FGFR3 oncogenic dependency for tumor growth has been demonstrated both *in vitro* and *in vivo* (cell lines or patient-derived xenografts) [10–14]. Recently, a phase II study investigating the efficacy of Erdafinitib (Balversa™), an FGFR inhibitor, showed a 40% objective response rate in eligible patients with locally advanced or metastatic BCa with FGFR alterations [15], resulting in the FDA approval in this setting.

To determine the functional role of a mutant FGFR3 in bladder cancer *in vivo*, several teams have developed FGFR3-altered genetically engineered mice (GEM) through transgene insertion. Two studies were based on the insertion of a mouse mutant FGFR3 (K644E[16] or S243C[17]), whereas another study used a human FGFR3-S249C transgene targeted to the urothelium [18]. So far, results suggest that although FGFR3 activation alone is not sufficient to induce tumorigenesis [16,18,19], it can promote tumor formation when associated with other molecular alterations (p53/pRB deficiency [17] or PTEN loss [19]) or with a carcinogen treatment [18].

In this study, we report for the first time a GEM model overexpressing the human FGFR3b-S249C (hFGFR3-S249C) mutant in the urothelium, in which mice develop low-grade papillary bladder carcinomas presenting genomic alterations and resembling the human counterpart at the histological and transcriptomic levels. The impact of FGFR3-S249C expression on the penetrance of mice tumor formation led us to investigate FGFR3 expression level in different normal human epithelia and its role in the tissue specificity of FGFR3-driven tumors. The higher incidence of tumors in male mice prompted us to explore sex distribution and androgen receptor (AR) activation in FGFR3-mutated human bladder tumors. The higher male/female ratio in FGFR3-mutated tumors, associated with a higher activation of AR compared to wildtype tumors considering gender adjustment, reinforces the hypothesis of a role for AR in bladder carcinogenesis and stresses the importance of elucidating its role specifically in an FGFR3-mutated context.

## Methods

### Mouse models

Animal care and use for this study were performed in accordance with the recommendations of the European Community (2010/63/UE) for the care and use of laboratory animals. Experimental procedures were specifically approved by the ethics committee of the Institut Curie CEEA-IC #118 (Authorization APAFiS#26671-2020072108262352-v2 given by National Authority) in compliance with the international guidelines.

### Generation of UII-hFGFR3-S249C transgenic mice

The expression of a human FGFR3-IIIb carrying the S249C mutation was targeted to the urothelium of mice by using the 5’ regulatory region of the mouse uroplakin II promoter. The UII-hFGFR3b-S249C construct was obtained by inserting the 3.6 kb murine Uroplakin II promoter (UII) [20] excised with SalI and BamHI into the same restriction sites of the vector containing the β-globin intron 2 and the 3’ polyadenylation sequences of SV40 [21] followed by the insertion of a human S249C mutated FGFR3 cDNA excised with XbaI and HindIII into the SmaI site of this vector. All PCR-generated segments were verified by sequencing both strands. The pUII-hFGFR3b-S249C constructs excised with KpnI were purified and microinjected into fertilized B6D2 oocytes. Genomic DNA was extracted from mouse tails and screened by PCR to identify transgene integration. Two lines were selected, L569 and L538, and mice were back-crossed five times to C57BL/6J mice. Mice were of a mixed background and littermates were used as control. Bladders from mice aged 1 to 24 months were examined for macroscopic lesions followed by a histopathological analysis when required. Mice were then intercrossed to obtain hetero-and homozygous mice for the transgene.

### Carcinogen treatment

BBN (N-butyl-N-(4-hydroxybutyl)-nitrosamine) was purchased from Tokyo Kasei Kogyo (Tokyo, Japan). Animals were housed in plastic cages in a controlled-environment room maintained at 22°C ±1°C with 12h light-12h dark cycles. All animals received food *ad libitum*. The UII-hFGFR3-S249C mice and control mice were aged 8-10 weeks old at the time of first carcinogen administration and were randomized within the different cages. The BBN was diluted at 0.05% in drinking water (ad libitum) for 8 weeks (the BBN solution was freshly prepared every 2-3 days). After withdrawal of BBN administration, drinking water without added chemicals was available *ad libitum*. Tumor formation and progression was followed weekly by echography in a blind way concerning mice genotype. Mice were sacrificed when tumors reached 80% of bladder volume or when weight loss was greater than 20% of body weight.

### RT-qPCR analyses

Total RNA from mouse urothelium was obtained using the RNeasy mini kit (Qiagen, Courtaboeuf, France) according to the manufacturer’s instructions. One μg of total RNA was reverse transcribed using random hexamers (20 pmol) and 200 units of MMLV reverse transcriptase. The expression levels of the human FGFR3 transgene in urothelium and other tissues of transgenic mice were determined by real time PCR analysis. The mouse Ef1a gene was used as a control gene. Quantitative real time PCR was performed using a SYBR green PCR master Mix according to the manufacturer’s instructions (Applied Biosystems, Foster city, CA, USA), on an ABI prism 7900 sequence detection system (Applied Biosystems). *FGFR3* expression levels were calculated using the comparative Ct method normalized to *Ef1a* mRNA expression levels. The sequence of these primers used were as follow:

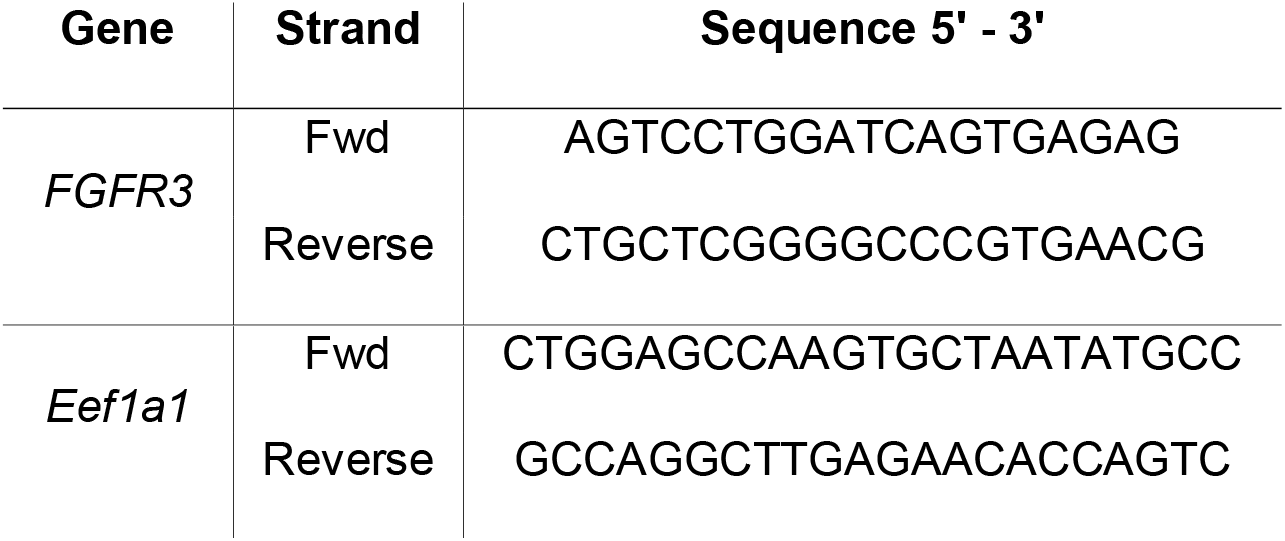

### Radioactive PCR

To compare the relative expression of the FGFR3 transgene to that of the endogenous murine *Fgfr3* in urothelium, transgenic urothelium cDNA was amplified in presence of 32P dCTP using the forward 5’-GCAGGCATCCTCAGCTAC-3’ and reverse 5’-TGGACTCGACCGGAGCGTA-3’ primers which recognize both human and mouse *FGFR3*. The 107 bp amplified products were then digested with RsAI and HinP1I. The human amplified product possesses a RsAI restriction site and the mouse amplified product a HinP1I restriction site. After digestion, two fragments of 88 bp and 19 bp were obtained from the amplified human FGFR3 cDNA and two fragments of 59 bp and 48 bp were obtained from the amplified mouse Fgfr3 cDNA. The digested products were subjected to polyacrylamide gel electrophoresis and the intensities of the bands were quantified with a Molecular Dynamics Storm PhosphorImager (Molecular Dynamics/Amersham, Sunnyvale, CA, USA).

### Histological analyses

UII-hFGFR3-S249C mutant and control mice bladders were fixed in 10 % formalin, embedded in paraffin and cut at 4-μm thick slides for histological and immunohistochemical analyses. Histological hematoxylin and eosin-stained (H&E) slides for tumors and hyperplasia were reviewed by two genito-urinary pathologists. Proliferation rate was estimated by immunohistochemistry with an anti-Ki67 polyclonal rabbit antibody (Abcam, ab15580, pH6, 1:20 000 dilution) using the Autostainer 480 (Lab vision) system.

### Whole exome sequencing and identification of copy number alterations

DNA from UII-hFGFR3-S249C mouse normal urothelia, hyperplastic urothelia and urothelial carcinomas was extracted using phenol-chloroform. Whole exome libraries were prepared by Integragen (Evry, France). Raw sequence alignment and variant calling were carried out using Illumina CASAVA 1.8 software (mm10 mouse reference genome). Each variant was annotated according to its presence in the 1000Genome, Exome Variant Server (EVS) or Integragen database, and according to its functional category (synonymous, missense, nonsense, splice variant, frameshift or in-frame indels). Reliable somatic variants were identified as those having a sequencing depth in ≥10 reads in tumor and normal urothelium samples, with ≥3 variant calls representing ≥15% total reads in the tumor, ≤1 variant calls representing <5% total reads in the normal urothelium, and a QPHRED score ≥20 for both SNP detection and genotype calling (≥30 for indels).

Copy number alterations (CNAs) were identified using coverage data to calculate the log ratio of the coverage in each tumor sample as compared to a normal urothelium sample. Log-ratio profiles were then smoothed using the circular binary segmentation algorithm as implemented in the Bioconductor package DNAcopy. The most frequent smoothed value was considered to be the zero level of each sample. Segments with a smoothed log ratio above zero + 0.15 or below zero – 0.15 were considered to have gains and deletions, respectively. High-level amplification and homozygous deletion thresholds were defined as the mean +7 SD of smoothed log ratios in regions with gains and deletions, respectively.

The identified frequent chromosomal gains were further validated by qPCR using genomic DNA. Primers targeting exonic regions from different genes found in the most frequently altered chromosomes were designed. A Taqman qPCR (Applied Biosystems) was carried out on gDNA to compare expression levels between normal urothelium and tumors from UII-hFGFR3-S249C mice. Normalization was performed using genes present on chromosomes without genomic alteration. The designed primers were the following:

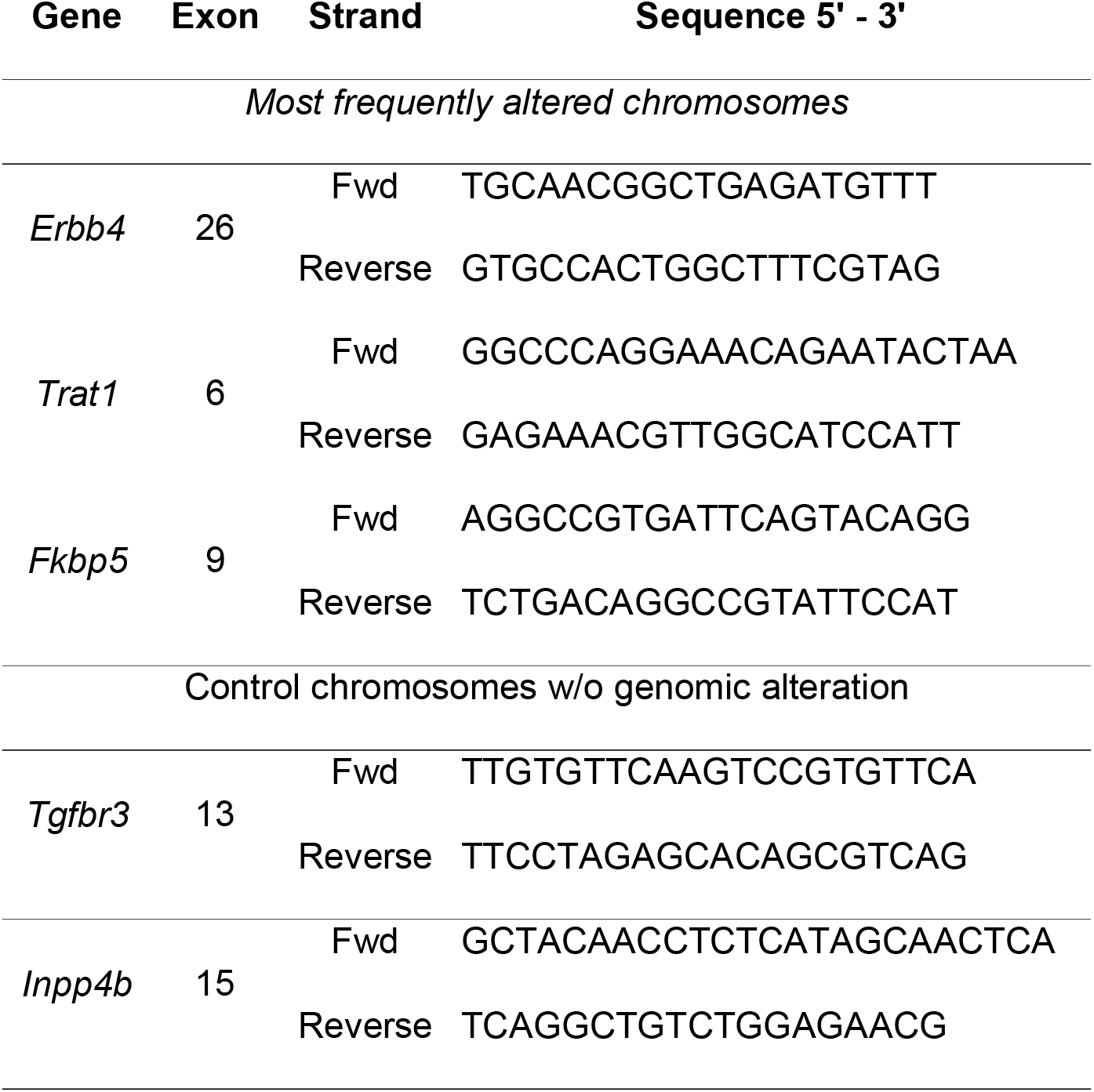

### Microarray transcriptome profiling

Total RNA (200ng) from UII-hFGFR3-S249C mouse normal urothelium, hyperplastic lesions and urothelial carcinomas was analyzed with the Affymetrix Mouse Exon 1.0 ST. Array gene expression was RMA normalized and annotated to the GRCm38 genome version using Brain-Array annotation provided one single value per gene. The LIMMA algorithm was applied to calculate the genes having a significant change of expression between urothelial carcinoma, hyperplasia and normal urothelium. Genes were considered to be differentially expressed when they presented an absolute log2FC ≥0.58 and an adjusted p-value <0.05. P-values were adjusted for multiple comparisons using the Benjamini-Hochberg correction. Upstream regulator analysis based on the differentially expressed genes was performed with the Ingenuity Pathway Analysis (IPA) software to identify key TFs as well as predict their transcriptional activities. FGFR3 regulated transcriptomes (FGFR3 knockdown versus control) from three BCa-derived cell lines (UM-UC-14, MGH-U3 and RT112) were previously published by host lab [8,14]. Since all these cell lines were FGFR3 dependent, we considered them as one group, treated and non-treated, and performed similar analyses as above to double confirm TFs identified from UII-hFGFR3-S249C mouse model.

### Pathway Enrichment

Genes with an adjusted p-value inferior to 0.05 were used to carry out an enrichment analysis of Pathways. The enrichment analysis was done using Gene Set Enrichment Analysis (GSEA) method with *amus musculus* Affy Exon 1.0 ST genomic background. Significantly enriched pathways were defined as having an adjusted p-value (Benjamini and Hochberg) inferior to 0.05.

### Cross-species hierarchical clustering

Microarray transcriptomic data from UII-hFGFR3-S249C and BBN mice was combined with transcriptomic array data from human bladder tumors (CIT; Affymetrix Exon 1.0 ST; 96 MIBC and 99 NMIBC). Batch effects due to data combination were corrected using the surrogate variable analysis R package. The protocol used for coclustering of the two species was that of previously described [22]. Hierarchical clustering was done using a gene signature derived from the consensus molecular classification of MIBC [2].

### Transcriptome classifier

Subtype calls were done on murine hFGFR3-S249C and previously established murine BBN induced tumor transcriptomes. Samples were classified using a 4-class classifier for NMIBC [7] or the BASE47 classification algorithm[23] and the median centered expression of the murine orthologues found in the BASE47 signature, as previously described [2,22].

### Gender bias in tumors with FGFR3 mutations

We established a merged cohort of 1,220 BCa subjects with both FGFR3 mutation and gender information available, based on data from our Carte d’Identité des Tumeurs (CIT) database and public sources [2,3,5,24,25]. For MIBC subjects from the TCGA dataset, transcriptome-derived molecular classification was determined as previously described [2]. Molecular classification for NMIBC samples included in the UROMOL study was extracted from supplementary data of the associated publication [5,7]. We compared the gender distribution (male vs. female) between bladder cancers harboring or not an FGFR3 mutation in the overall cohort and in the different subgroups of MIBC and NMIBC. Odds ratios (ORs), corresponding to 95% confidence intervals (95% CIs), and Z-test based *P* values were calculated. An OR > 1 indicates a higher proportion of males in FGFR3 mutated tumors, and a 95% CI not covering 1 or P < 0.05 indicates a statistically significant difference.

### AR regulon activity

RNA-seq derived transcriptome data (fragments per kilobase of transcript per million, FPKM normalization with log2 transformation) of the UROMOL NMIBC (n = 476) [5,7] and TCGA MIBC (n = 408) [3] samples were downloaded from the ArrayExpress (https://www.ebi.ac.uk/arrayexpress/, accession number E-MTAB-4321) and UCSC Xena (https://xenabrowser.net/) databases, respectively. Computationally predicted AR regulon genes were extracted from supplementary data of TCGA MIBC [3] and for UROMOL NMIBC they were predicted using the same method than for TCGA [7] (Supplementary Table 3). We calculated for each of the above samples an AR regulon activity score as the difference of the sample-specific enrichment score of positive targets and that of negative targets obtained using the Gene Set Variation Analysis (GSVA) algorithm [26]. We compared AR regulon activity between tumors from the two subgroups identified as presenting a higher male gender bias in FGFR3-mutated tumors and those from other subgroups, respectively for NMIBC (Class 3 vs. others) and MIBC (LumP vs. others), using linear regression analysis with gender as a covariate for adjustment. The same analysis was applied to the comparison of FGFR3-mutated and wild-type tumors within these two subgroups. Gender-stratified non-parametric comparison of AR regulon activity between FGFR3 mutated and non-muted tumors was also performed, respectively for NMIBC and MIBC.

### Cell culture

The human bladder cancer derived cell line UM-UC-14 was obtained from DSMZ (Heidelberg, Germany) and cells were cultured at 37°C in an atmosphere of 5% CO2 in DMEM medium supplemented with 10% fetal bovine serum (FBS). Cells were routinely tested for mycoplasma contamination.

### Small molecule inhibitors

The PD173074 inhibitor was purchased from Calbiochem (Merck Eurolab, Fontenay Sous Bois, France).

### Transcription factor activity analysis

UM-UC-14 cells were seeded in 100mm plates at a density of 3.0×10^6^cells/dish. Cells were plated and left to adhere overnight. Afterwards, cells were treated for 40 hours with the pan-FGFR inhibitor PD173074 [100nM] (Calbiochem, Merck Eurolab, France). Control cells were treated with DMSO vehicle diluted proportionally to the inhibitor. After the 40h of treatment, nuclear fractions were isolated for analysis using the TF Activation Profilin Plate Array I from Signosis (according to the manufacturer’s protocol). Cellular fractions were recovered using the Thermo Fisher NE-PER nuclear and cytoplasmic extraction kit (ref 78833), following the manufacturer’s instructions.

### Immunoblotting

UM-UC-14 cells were resuspended in Laemmli lysis buffer (50 mM Tris-HCl pH 6.8, 2 mM DTT, 2.5 mM EDTA, 2.5 mM EGTA, 2% SDS, 5% glycerol with protease inhibitors and phosphatase inhibitors) (Roche). Obtained lysates were clarified by centrifugation. Protein concentration was determined by the BCA method (Thermo Scientific) and 20μg of protein per sample were resolved by SDS-PAGE in 7.5 or 10% polyacrylamide gels. Following electrotransfer of proteins to a nitrocellulose membrane, membranes were probed with antibodies against p-AR (Invitrogen ref: PA5-64643, diluted 1/1000), AR (Cell Signaling Technology ref:5153, diluted 1/500), MYC (Cell Signaling Technology ref: 9402, diluted 1/500) and loading control protein □-Tubulin (Sigma Aldrich ref: ref T9026, diluted 1/20,000). The secondary antibodies used were anti-mouse IgG, HRP-linked, and anti-rabbit IgG, HRP-linked antibody (Cell Signaling Technology ref: 7076 and ref 7074, diluted 1/3000-respectively). Signal was detected using SuperSignal West Femto (ThermoFisher) or Clarity Western ECL (BioRad) substrates followed by exposure on X-ray film (ThermoFischer).

### Statistical analysis

Linear Models for Microarray Data (LIMMA) was used to analyze microarray experiments involving simultaneous comparisons between large numbers of RNA targets [27]. Benjamini and Hochberg adjustment was performed for multiple testing corrections. Unless otherwise specified, continuous variables were shown as mean ± standard deviation, and categorical variables shown as counts by factor levels. Fisher’s exact test was used to test dependency between two categorical variables, and Wilcoxon rank sum test was used for comparison between two independent groups regarding the distribution of a continuous variable.

## Results

### FGFR3-S249C expression in Uroplakin II-expressing cells induces urothelial hyperplasia and non-muscle-invasive low-grade papillary urothelial carcinoma

To determine the role of a constitutively activated FGFR3 mutant in bladder tumorigenesis, we generated transgenic mice expressing a human mutant receptor in the urothelium. We focused on the hFGFR3-S249C mutation, the most common FGFR3 mutation in both NMIBC and MIBC [8], and used the mouse Uroplakin II (UII) gene promoter to specifically target its expression to urothelial cells [28] (Fig. 1A). We selected two founders for our UII-hFGFR3-S249C model, numbers 569 and 538, which expressed high levels of the human FGFR3 transgene in the urothelium as evidenced by RT-qPCR (Supplementary Fig 1A) and transmitted the transgene to their offspring. *In situ* hybridization using a human FGFR3-specific probe showed hFGFR3 mRNA expression in the supra-basal and intermediate cell layers and in very few basal cells of the urothelium (Supplementary Fig. 1B). Moreover, human FGFR3 mRNA expression levels in the urothelium were 4 and 1.5-fold higher than that of endogenous mouse Fgfr3 in founders 569 and 538, respectively, as assessed by radioactive PCR (Supplementary Fig. 1C). These two founders were viable and fertile and transmitted the transgene to their offspring in a Mendelian fashion. Following propagation of founder lines, we examined the bladder of transgenic mice aged 1 to 18 months old. Histological analysis of UII-hFGFR3-S249C mice bladders identified hyperplastic lesions, defined by a thickened urothelium without cytological atypia, with an increase to seven to ten cell layers and focally more (ten to twenty) at 18 months (Fig. 1B). The penetrance of the phenotype was complete from 6 months of age in both lines. Macroscopically, focal papillary lesions were observed after 15 months with a low penetrance in both lines (~10% and 4% for 569 and 538, respectively). Histological analysis of these lesions revealed they were non-invasive urothelial carcinomas (stage pTa), characterized by a papillary tumor architecture with either exophytic or mixed (exophytic and inverted) growth patterns, low-grade cytologic atypia with homogeneous nuclei size and scarce mitotic figures (Fig. 1B). We also observed, as previously described [18], that hFGFR3-S249C expression promoted N-Butyl-N(4-hydroxybutyl) nitrosamine (BBN)-induced tumor growth (Supplementary Fig. 2A). We then selected the 569 line, presenting a higher expression of hFGFR3-S249C and a higher penetrance of the phenotype, for further transcriptomic characterization, using an Affymetrix mouse exon array. Hyperplastic lesions were similar to normal urothelium in terms of both proliferation rate and transcriptomic profile, respectively determined by mKi67 expression levels (Fig. 1C) and principal component analysis (Fig. 1D). In contrast, the tumors presented a significantly higher proliferation rate (Fig. 1C) and principal component analysis highlighted a distinct transcriptomic profile compared to normal and hyperplasia samples (Fig. 1D). Similar to human low-grade pTa urothelial papillary carcinoma, the proliferation rate in mouse tumors was low, with <10% of Ki67-labelled tumor nuclei by immunohistochemistry (Supplementary Fig. 2B). Whole exome sequencing analysis of 7 tumors did not reveal any recurrent mutations induced by hFGFR3-249C expression but showed recurrent copy number alterations, the most common being chromosome 16 gains in 5 out of 7 tumors (Fig. 1E). We selected 3 genes (*Trat1, Erbb4, Fkbp5*) located in 3 gained regions (chr16, chr1 and chr17, respectively) and verified their gain by qPCR on genomic DNA, in 6 of the tumors previously analyzed by whole exome sequencing and in 4 additional tumors (Supplementary Fig. 2C).

**Figure 1.**
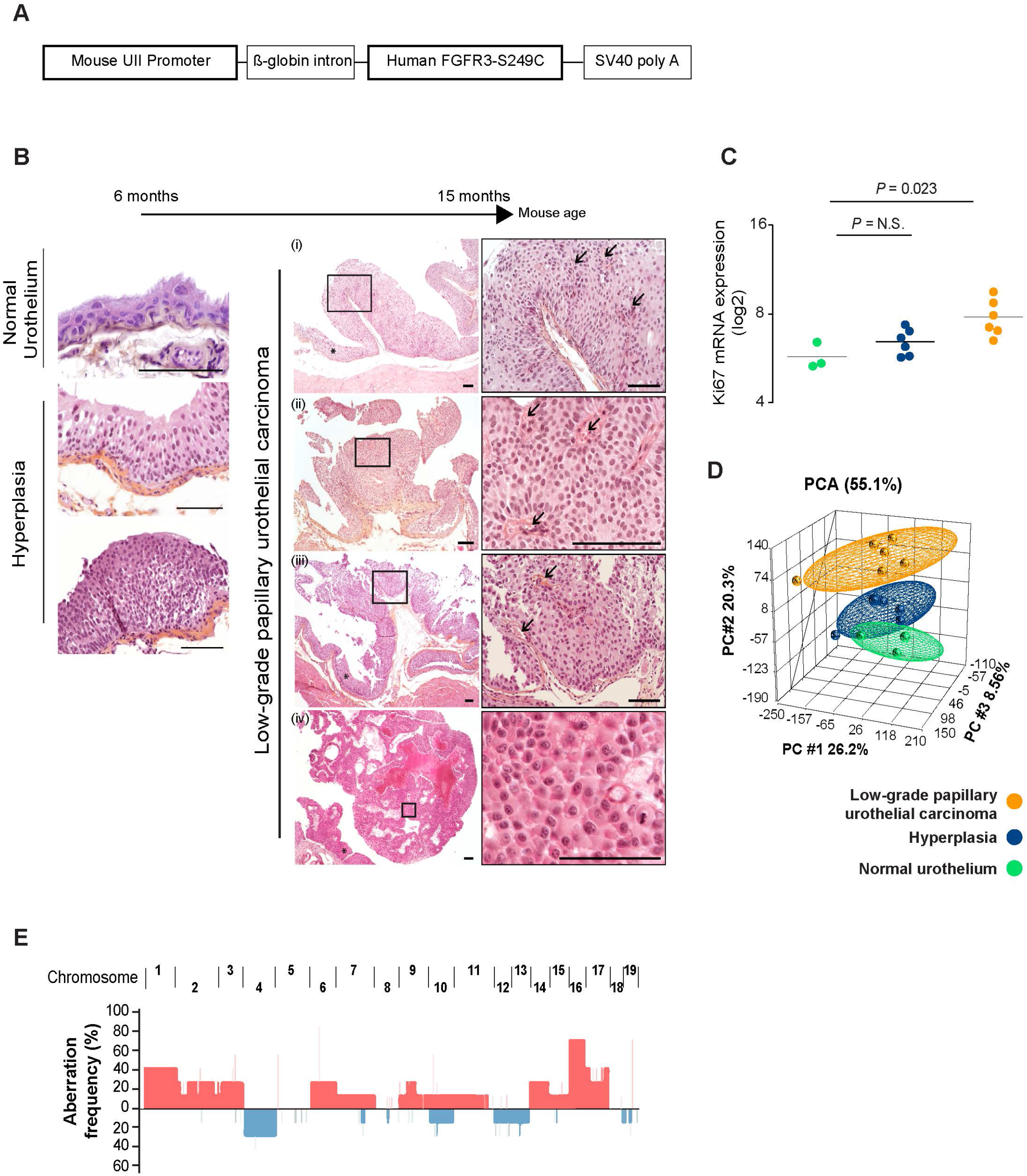
UPII-FGFR3-S249C transgenic mice develop urothelial hyperplasia and non-muscle-invasive low-grade urothelial carcinoma. A) Chimeric construct used to generate transgenic mice, consisting of a 3.6-kb mouse UPII gene promoter and a 2.1-kb human FGFR3b cDNA carrying the mutation S249C. B) Representative H&E histology of urothelial lesions in hFGFR3-S249C mice compared to normal mouse urothelium. Hyperplastic lesions (left panel) or low grade papillary urothelial carcinomas (right panel) developed in hFGFR3 S249C mice from 6 months and 15 months of age, respectively. Stars show tumor-adjacent urothelial hyperplasia. Arrows point to papillae fibrovascular cores. Scale bar: 100μm. C) mKi67 mRNA expression levels (Affymetrix Mouse Exon 1.0 ST. Array signal) in tumor and hyperplastic urothelium from UPII-hFGFR3-S249C mice and in normal urothelium from control littermates. D) Principal component analysis of all genes expressed on the Affymetrix Mouse Exon 1.0 ST. Array from tumor and hyperplastic urothelium from UPII-hFGFR3-S249C mice and from normal urothelial samples from control littermate mice (n= 6 tumors, 6 hyperplastic lesions, 3 control urothelium). E) Frequency of chromosomal copy number alterations in tumors from UII-hFGFR3 S249C. (red = gain; blue= loss).

Taken together, these results showed that hFGFR3-S249C is oncogenic in the bladder urothelium *in vivo*, inducing tumors with genomic alterations.

### The UII-hFGFR3-S249C model is a model of human luminal papillary BCa

We further explored the transcriptomes of our newly generated model to determine if mouse tumors presented known features of human FGFR3-mutated tumors. In line with the PCA (Fig. 1D), differential gene expression analysis revealed very few statistically significant dysregulated genes between hyperplasia and normal urothelium from littermates (controls) (Fig. 2A, right panel and supplementary Table 1), whereas 989 differentially expressed genes (DEGs) were identified between hFGFR3-S249C mouse tumors and controls (Fig. 2A, left panel and supplementary Table 1). GSEA analysis of the DEGs using Reactome and KEGG databases highlighted, as previsouly demonstrated upon FGFR3 modulation in human bladder cancer-derived cell lines [29,30], the increase of “cell cycle” and “DNA replication” and a decrease of “focal adhesion” and “cell adhesion” terms (Fig. 2B). Contrary to what has been reported after FGFR3-BAIAP2L1 fusion gene overexpression in Rat-2 cells [11], FGFR3-S249C expression induced an up-regulation of the P53 signalling pathway (Fig. 2B) We estimated the immune cell infiltration abundance in hFGFR3-S249C tumors by applying the mouse Microenvironment Cell Populations-counter method (mMCP-counter) [31] to our UII-FGFR3-S249C mouse transcriptomic data. We estimated a weak infiltration of hFGFR3-S249C low-grade luminal papillary tumors as compared to normal urothelium for all types of immune cells (Fig. 2C) consistently with former studies that have shown that human molecular subtypes enriched in FGFR3-mutated tumors are characterized by a lower immune response and infiltrating immune cell activity compared to other subtypes [2,5,7]

**Figure 2:**
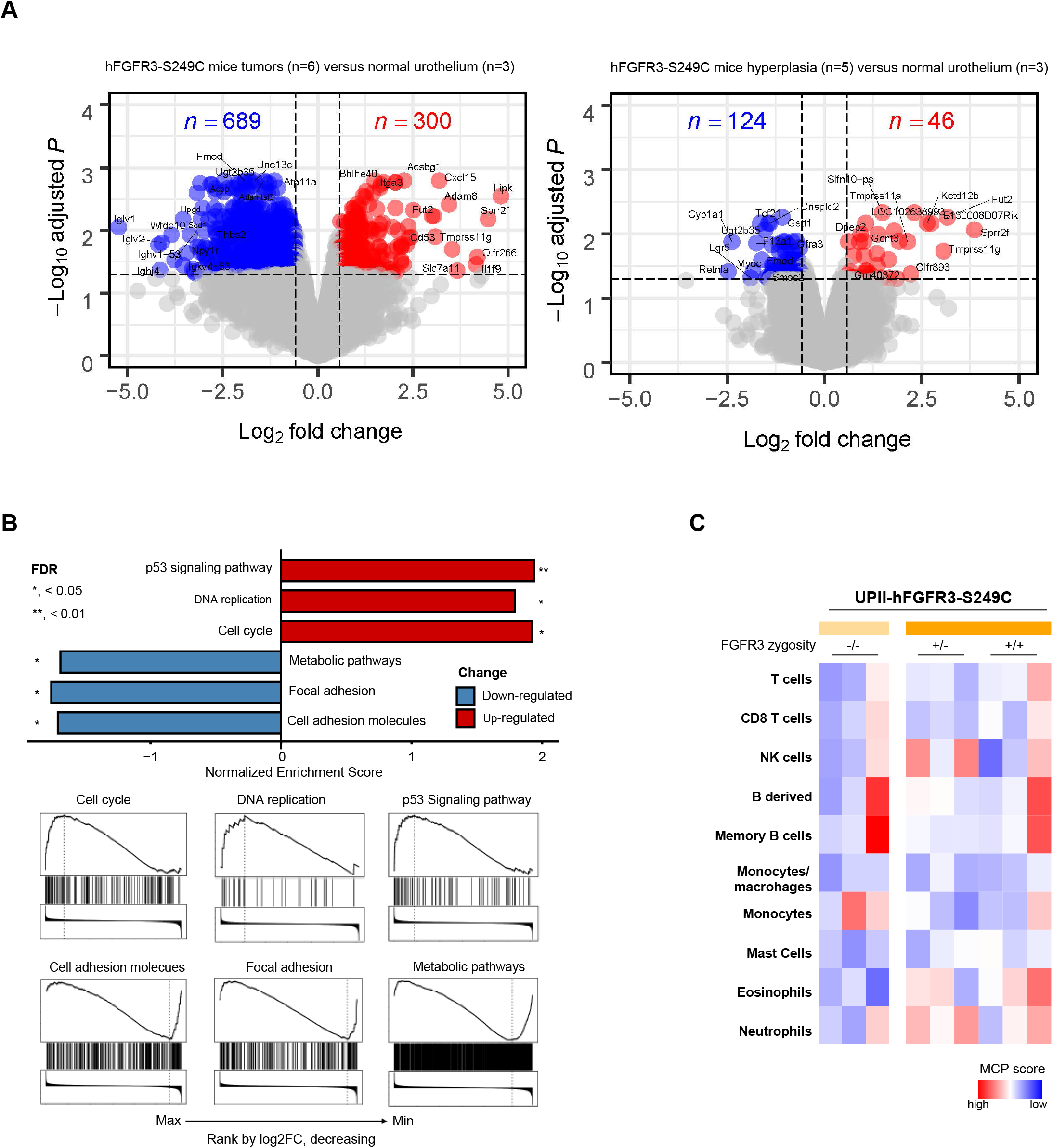
Mouse hFGFR3-S249C tumors present known FGFR3-induced pathway activation and are poorly infiltrated by immune cells. A) Volcano plot of the differentially expressed genes (|log2FC|>0.58;) between hFGFR3-S249C mice tumors and littermates normal urothelium (controls) (DEGs: 989 genes considering adj.*P*-val<0.05, left panel) and between hFGFR3-S249C mice hyperplasia and controls (DEGs:170 genes considering adj.*P*-val<0.05, right panel). B) Murine Gene Set Enrichment Analysis (GSEA) of deregulated genes in hFGFR3-S249C tumors as compared to normal urothelium from littermates. The significantly enriched terms, up- (in red) or downregulated (in blue), and their FDRs are displayed. C) Heatmap of mMCP counter scores for estimation of tumor immune infiltration abundance based on transcriptomic data from hFGFR3-S249C mice tumors (n=6), hyperplasia (n=5) and control urothelium from littermates (n=3).

Given the papillary nature of hFGFR3-S249C-induced murine tumors, we hypothesized that they recapitulate the luminal papillary molecular phenotype of human bladder cancer. Conversely, we and others have previously shown that BBN-induced bladder tumors in mice represent a model of the basal subgroup of muscle-invasive BCa [22,32]. To classify hFGFR3-S249C tumors, we first applied a molecular classifier allowing to distinguish between four classes of NMIBC [7]. The six hFGFR3-S249C-induced tumors showed high and similar correlations to the NMIBC classes 1 and 3 centroids of gene expression, which correspond to two classes that are enriched in FGFR3-mutated tumors in humans [5,7] (Fig. 3A). Next, we applied the BASE47 classifier to distinguish between luminal and basal BCa subtypes [23]. According to this classifier, FGFR3-induced bladder mice tumors were defined as luminal subtype, whereas our previously obtained BBN-induced tumors [32] classified as basal (Fig. 3B). To further validate these findings, we performed a cross-species comparison by co-clustering the hFGFR3-S249C and BBN mice tumors with human tumors from our CIT cohort (n = 96 MIBCs and 99 NMIBCs) [33], taking genes from a recently developed consensus classifier for basal, luminal-papillary and neuroendocrine-like human BCas [2] and employing the corresponding orthologues across the species. We found that BBN mice tumors co-clustered with human basal tumors and hFGFR3-S249C co-clustered with luminal papillary tumors, a subgroup enriched in *FGFR3* mutations (Fig. 3C). Interestingly, contrasting the mouse tumors’ microenvironment infiltration constitution estimated with mMCP-counter to those of TCGA MIBC tumors (n = 408) estimated with MCP-counter, we found similar clustering results suggesting that both BBN and FGFR3 models are relevant models to decipher the role of tumor microenvironment in bladder basal and luminal tumors, respectively (Fig. 3D).

**Figure 3:**
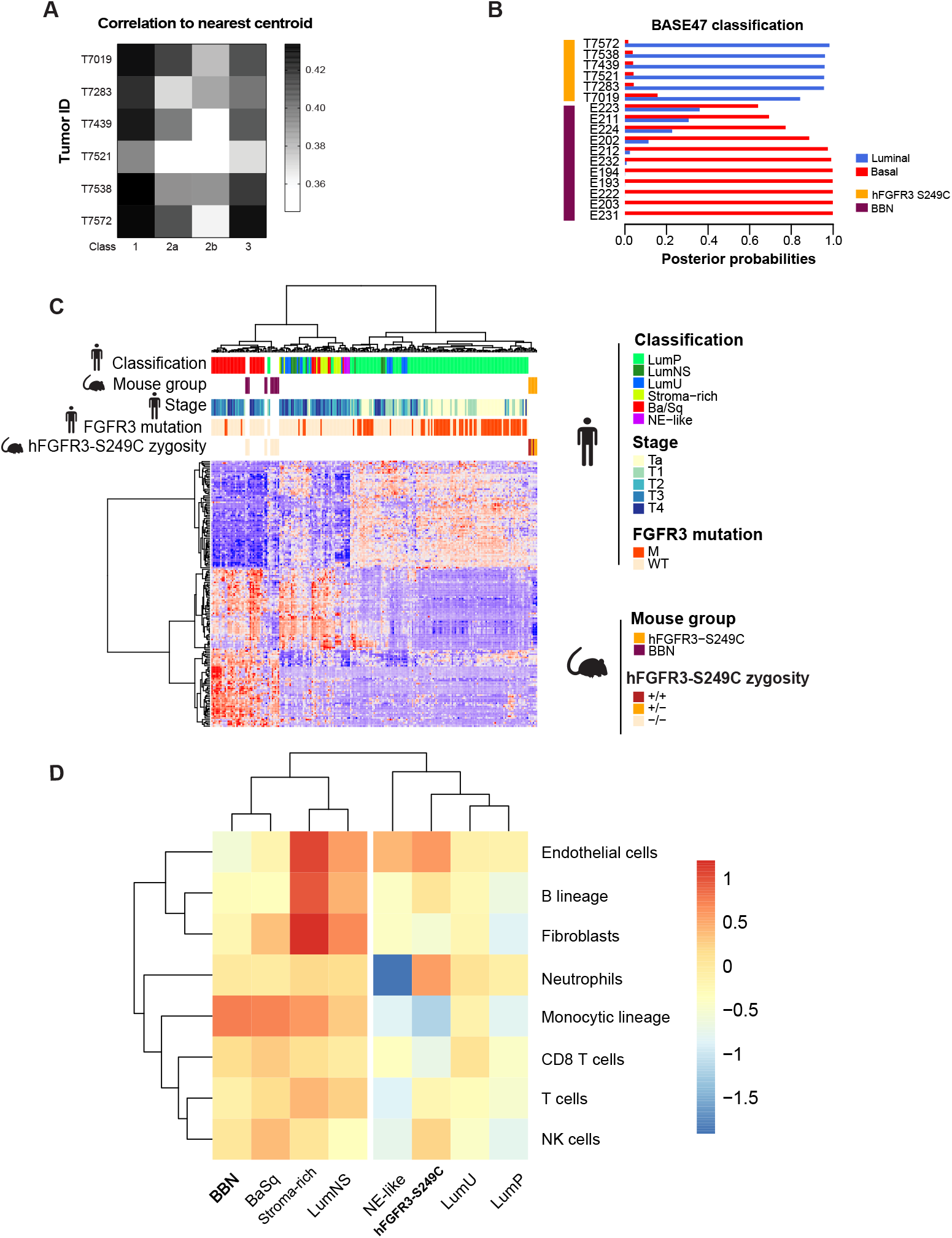
Mouse hFGFR3-S249C bladder tumors resemble human luminal papillary tumors at the transcriptomic level. A) Correlation to the Hedegaard NonMuscle Invasive Carcinoma classifier [5,7] and B) the BASE47 classifier [23] for tumors of hFGFR3-S249C mice. Muscle-invasive tumors derived from BBN treated mice are used for comparison [32]. C) Cross-species, unsupervised hierarchical clustering of hFGFR3-S249C mice tumors (n=6), BBN-induced tumors (n= 11) and human bladder tumors (n= 193 from the CIT series) considering genes from a recently developed consensus classifier for basal and luminal-papillary human BCas [2]. D) Cross-species unsupervised hierarchical clustering regarding the microenvironment cellular compositions estimated with mMCP-counter and MCP-counter, for BBN and hFGFR3-S249C models as well as TCGA MIBC tumors (n = 408).

Taken together, our data suggest that hFGFR3-S249C mouse provide a relevant and potentially useful model to decipher the role of FGFR3 in BCa formation, including in the cross-talk with stroma.

### FGFR3 expression levels impact tumor formation in UII-hFGFR3-S249C mice and could account for the tissue specificity of mutant FGFR3-induced tumors

To test if the mutated *FGFR3* transgene expression levels influence tumorigenesis, we studied a large series of mice (n = 402) and compared the frequency of tumors in 18-month-old UII-hFGFR3-S249C heterozygous (one allele of the transgene) and homozygous (two alleles of the transgene) mice. The frequency of tumor formation, was significantly higher (*P* = 4.8-E9) in homozygous compared to heterozygous mice (~ 40% and 10%, respectively) (Fig. 4A). Strikingly, multifocal tumors were specifically identified in homozygous mice, whereas heterozygous mice only developed unifocal tumors. We hypothesized that the increased sensitivity to tumor development could be linked to the significantly higher expression level of hFGFR3 in the urothelium of homozygous compared to heterozygous mice, as assessed by RT-qPCR (Fig. 4B). Following this hypothesis, we measured FGFR3b (main isoform expressed in epithelial cells) expression levels in different normal human epithelia, including urothelium, obtained after laser microdissection. Interestingly, epithelia presenting high expression levels of FGFR3b were those in which FGFR3-mutated tumors are described (bladder, skin, exocervix) (Fig. 4C) [3,34–36]. Our data suggest that *FGFR3* mutations require an epithelium with a high expression of FGFR3 to induce tumor formation. Nevertheless, although *FGFR3* gene dosage in mice influenced tumor frequency, it did not reduce tumor development latency or induce progression towards muscle-invasive BCa. No histopathological difference was observed in hFGFR3-S249C-induced tumors between heterozygous and homozygous mice.

**Figure 4:**
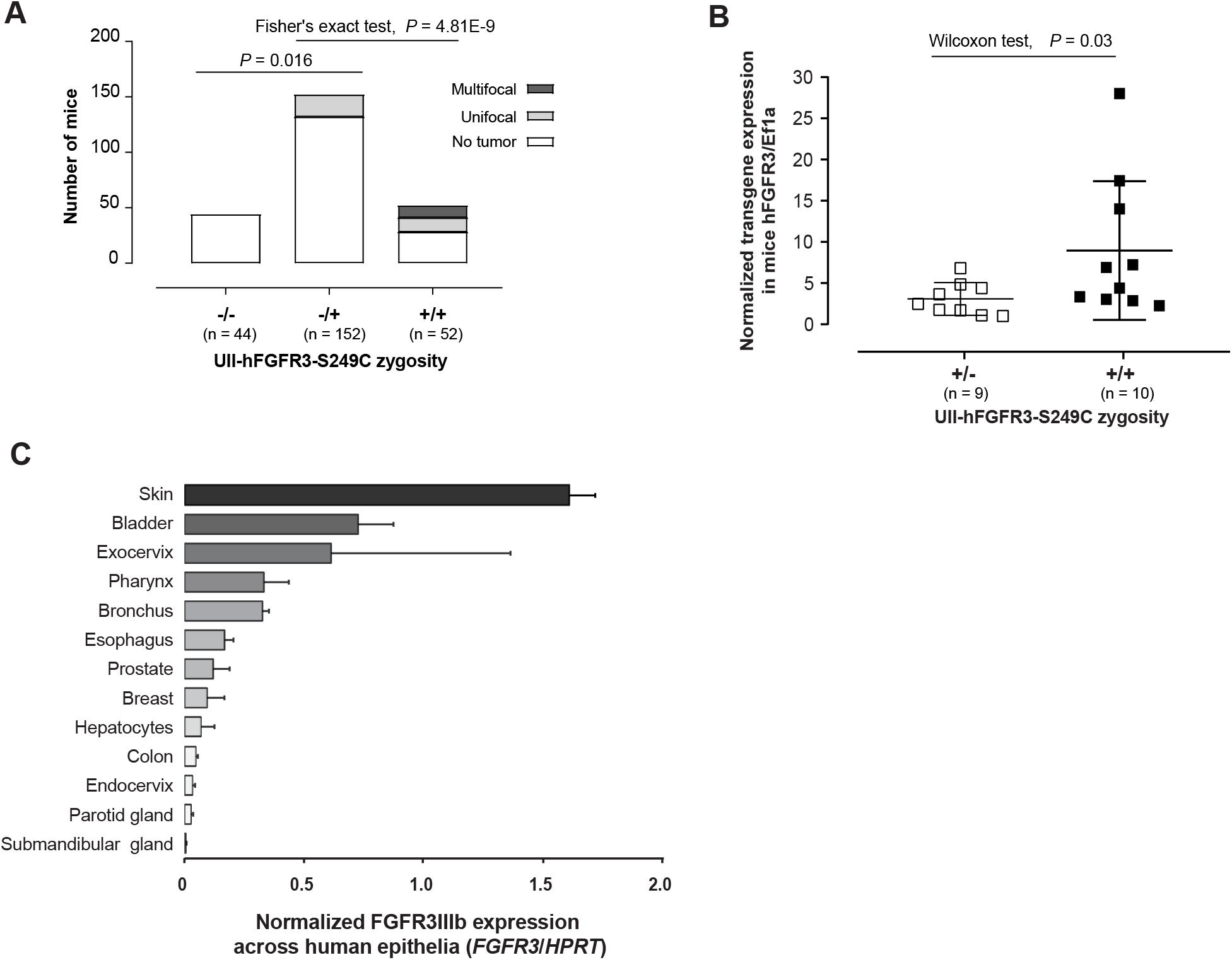
FGFR3-induced tumor development is dependent on FGFR3 expression levels. A) Frequency of unifocal or plurifocal bladder tumor development in hFGFR3-S249C homozygous (+/+) or heterozygous (+/-) mice versus control littermates (-/-). Proportions were compared using Fisher’s exact test. B) hFGFR3-IIIb mRNA expression evaluated by RT-qPCR in hFGFR3-S249C homozygous (+/+) or heterozygous (+/-) mice. Results were normalized using Eef1a expression levels. The statistical significance of differences was assessed using Wilcoxon test. C) FGFR3-IIIb expression levels as assessed by RT-qPCR in different human epithelia obtained after laser-microdissection. Results were normalized using HPRT expression levels.

### Mutant-FGFR3 induce a higher male gender bias in bladder tumor formation and a higher androgen receptor activation compared to wild type tumors

In humans, it is well known that bladder cancer is a sex-biased cancer with men being three times more susceptible to BCa than women [37]. It has recently been proposed that sex specificity could drive the distribution of the different molecular subtypes, males being enriched with tumors of luminal papillary (LumP) and neuroendocrine-like subtypes and females with the basal/squamous (Ba/Sq) subtype [38]. However, the role of sex-related pathways in bladder tumor development, and that of the androgen receptor in particular, has not been elucidated. Consequently, we further analyzed the gender of animals that developed tumors in our mutant FGFR3-induced model. We observed a significantly higher proportion of tumors in male compared to female mice, considering tumor zygosity as stratification variable (Fig. 5A). Prompted by this observation, we explored the relationship between gender and *FGFR3* mutation status in human tumors. Since recent bioinformatics analyses and functional *in vitro* experiments suggested that other *FGFR3* mutations than S249C display same or higher oncogenic potential [39,40], we considered all *FGFR3* mutations and not only S249C for further analysis. Considering all tumors, we observed a significantly higher male to female ratio in *FGFR3*-mutated tumors compared to *FGFR3* wild-type tumors, suggesting that sex-specific pathways could synergize with *FGFR3* mutations to induce bladder tumor formation. We then investigated AR activation as one of these potential pathways. Using AR regulon gene sets defined for MIBC and NMIBC, we calculated AR regulon activity in both NMIBC from the UROMOL cohort and MIBC from the TCGA cohort using gene set variation analysis (GSVA). Considering gender as a covariate for adjustment, we observed a significantly higher AR regulon activity in *FGFR3*-mutated tumors compared to *FGFR3* wild-types in both NMIBC and MIBC (Fig. 5C), independently of the molecular subtypes (Supplementary Fig. 3A). Nonetheless, the male to female ratio was significantly higher (around 3.5-fold) in *FGFR3*-mutated tumors compared to *FGFR3* wild-type tumors only in 2 subtypes: the NMIBC class 3 (UROMOL cohort) and the MIBC LumP (TCGA cohort) tumors (Fig. 5B). Interestingly, these 2 subgroups were enriched for *FGFR3* mutations and more likely to rely on AR activity, given their higher AR regulon activity, as previously described [2,7] and when considering gender for adjustment (Supplementary Fig. 3B) or sex-stratification [38]. We finally investigated whether the higher AR regulon activity observed in *FGFR3*-mutated tumors was linked to FGFR3 activity. Using the Ingenuity Pathway Analysis software (IPA, Qiagen), we searched for upstream transcriptional regulators that could induce the gene expression changes upon FGFR3 up-regulation in our mouse model. We identified that mutant hFGFR3 expression in mice induced a significant increase in AR activity compared to control mice (Fig. 5D). Validating our approach, mutant FGFR3 also induced a high MYC activity in mouse BCas, in accordance with what we have previously reported in human FGFR3-dependent models *in vitro* and *in vivo* [14]. To further corroborate the regulation of AR activity by FGFR3 in humans, we analyzed the transcriptomic data from 3 FGFR3-dependent cell lines (MGH-U3, FGFR3-Y375C mutation; UM-UC-14, FGFR3-S249C mutation; RT112, FGFR3-TACC3 fusion), and carried out an upstream regulator I PA analysis on the DEGs upon FGFR3-knockout (Supplementary Table 2). Depletion of FGFR3 led to a significant decrease of AR and MYC activity (Fig. 5D). To validate our transcriptomic-based predictions, we used a TF activation profiling array to measure the binding of AR to its DNA target sequence, comparing FGFR3-S249C inhibition to control conditions in UM-UC-14 cells. We used MYC as control and confirmed that the inhibition of FGFR3-S249C led to a significant decrease of the binding of both TFs to their DNA-target sequence (Fig. 5E). Western-blot analysis further showed that the down-regulation of MYC activity upon FGFR3 inhibition in UM-UC-14 cells was associated, as previously reported [14], to a decrease of MYC expression whereas inhibition of AR activity was mostly associated to a decrease in AR phosphorylation (Fig. 5F).

**Figure 5:**
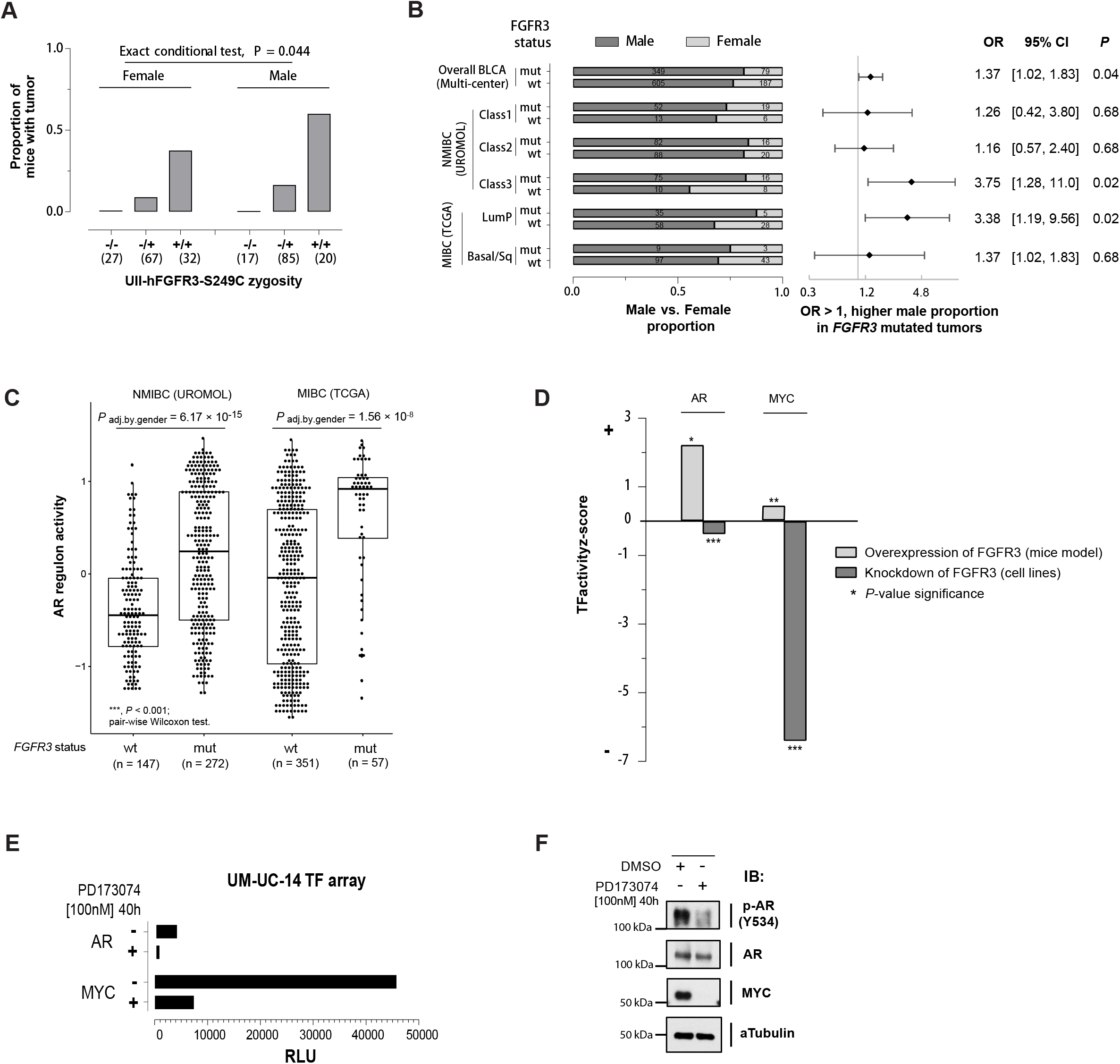
*FGFR3* mutation induces AR activation and is associated with a higher male gender bias in human tumors and with a higher AR activity compared to *FGFR3* wild-type tumors. A) Frequency of bladder tumors in male and female UII-hFGFR3-S249C mice. Gender bias in tumor occurrence, stratified by UII-hFGFR3-S249C zygosity. A Higher proportion of male mice developed a tumor (tumor occurrence rate = 21% vs. 14%, respectively in male and female animals; Zelen’s exact conditional test with UII-hFGFR3-S249C zygosity as stratification, twosided, *P* = 0.044; common odds ratio estimate = 2.19, 95% confidence interval = [1.02, 4.70]). (B) Comparison of gender distribution between *FGFR3*-mutated and wild-type human bladder tumors of different subgroups. Molecular classifications for both NMIBC (UROMOL) and MIBC (TCGA) as described previously, in which NMIBC Class 3 and MIBC luminal papillary (MIBC LumP) subtypes are known as to be enriched for *FGFR3* mutations [3,5,7]. NMIBC, non-muscle-invasive bladder cancer; MIBC, muscle-invasive bladder cancer. Odds ratios (ORs), corresponding 95% confidence intervals (95% CIs), and Z-test based P values were calculated (Methods). OR > 1, represents higher male proportion in *FGFR3* mutated tumors. (C) AR activity scores for NMIBC and MIBC, calculated via GSVA (Gene Set Variation Analysis), using defined AR regulon target gene sets. Comparison of AR regulon activity between *FGFR3*-mutated and wild-type NMIBC or MIBC human bladder tumors, *P* values: Linear regression with gender as covariate for adjustment. (D) Statistically significant activation states of AR and MYC inferred using the Ingenuity Pathway Analysis (IPA) software. Activation z-scores were calculated using the expression levels of transcription factor target genes in UPII-hFGFR3-S249C mouse bladder tumors as compared to control urothelium from littermates and in FGFR3-dependant bladder cancer derived cell lines (UM-UC-14 (FGFR3-S249C), MGH-U3 (FGFR3-Y375C) and RT112 (FGFR3-TACC3) cells) upon FGFR3 knockdown[8,14]. MYC is known to be regulated by mutated FGFR3 thus being a positive control. (E) Activation levels of AR and MYC in UM-UC-14 cells treated with the pan-FGFR inhibitor PD173074 [100nM,40h]. Activity levels were assayed using a TF Activation Profiling Array (RLU: Relative Luminescence Units). (F) Modulation of phosphorylated AR and total AR protein level in UM-UC-14 cells after FGFR3 inhibition (PD173074, 100nM for 40 hours). MYC represents positive control and tubulin was used as loading control.

## Discussion

We describe here the first transgenic mouse model demonstrating the tumorigenic role of a mutant FGFR3 in the bladder *in vivo*. Expression of hFGFR3-S249C in Uroplakin-II expressing cells induced spontaneous low-grade papillary tumor formation and favored BBN carcinogen-induced tumor development. We observed hFGFR3-S249C-induced tumorigenesis in two different transgenic lines, indicating that the observed effect was likely induced by the expression of the transgene itself rather than to the alteration of an endogenous key gene resulting from the nonspecific insertion of the transgene. Surprisingly, a mutated FGFR3 has already been targeted to urothelial cells using the same promoter without any spontaneous tumor formation being reported [16–19]. Nonetheless, in some of these studies, expression of the mutated receptor did promote bladder tumor development but only when combined to carcinogen exposure [18] or in association with *Pten* loss [19] or P53/pRB deficiency [17]. The discrepancy between the previously developed GEM models and ours could be linked to the specifically studied *FGFR3* mutation (S249C here, K644E in two previous studies [16,19]) or to the use of an inducible model for the expression of FGFR3-S249C in other studies [17,18]. We additionally showed that FGFR3-S249C expression levels impact the frequency of tumor formation, suggesting that a lower expression of the transgene in the former GEM models could also account for the absence of tumorigenesis.

We used here the most frequent mutation of FGFR3 in BCa but we have recently shown that the higher frequency of this mutation (FGFR3-S249C) is likely due to APOBEC mutagenesis rather than an increased tumorigenicity this mutation compared to other recurrent *FGFR3* mutations [8]. In line with our conclusions, a bioinformatics prediction of mutation tumorigenicity also revealed other FGFR3 mutations with high oncogenic potential [40]. We therefore hypothesize that other FGFR3 mutants, if expressed at sufficiently high levels, would also induce BCa formation.

*FGFR3* mutations are found in very few tumor types. FGFR3-S249C in particular, which is likely induced by APOBEC mutagenesis, is not frequently found in other APOBEC-related cancer types, such as breast or lung cancers, which suggests that the mutation brings little selective advantage to these epithelial cells. Our model points out that a sufficient level of expression of a mutant FGFR3 is critical to induce tumor formation, and that the weak expression level of FGFR3 observed in lung and breast normal epithelial cells could explain the absence of *FGFR3* mutation-driven tumor formation in these tissues.

Recently, the first pan-FGFR inhibitor – Erdafitinib/Balversa™ – has been approved by the FDA for patients with locally advanced or metastatic BCa presenting FGFR alterations. Considering the increasing interest of targeting FGFR3 for BCa treatment, having a model that resembles the human counterparts at the histological and transcriptomic levels such as ours, may have clinical translational value to evaluate drug response and to understand acquired drug resistance mechanisms. In particular, the model we present here is the first immunocompetent model of mutant FGFR3-induced low-grade luminal papillary carcinomas and we showed that the tumor microenvironment resembles the human counterparts. FGFR3-mutated tumors are non-T cell inflamed and have been associated to a poor immune-infiltrated microenvironment, being therefore less prone to respond to immunotherapy [3,41,42]. To confirm this hypothesis, a phase 1b/2 clinical trial (NCT03123055) comparing the efficacy of an anti-FGFR3 therapy (B-701, specific monoclonal antibody targeting FGFR3) coupled with immunotherapy (pembrolizumab) in advanced BCa patients harboring an altered FGFR3 is ongoing. Allografts obtained from this model (latency and penetrance of the phenotype won’t allow a direct use of the model), should provide a better understanding of immune-escape or immune-suppression mechanisms driven by a mutated/active FGFR3 and allow the evaluation of combined therapies using FGFR and checkpoint inhibitors.

This model of FGFR3-induced tumors should also allow for a better understanding of the signaling pathways activated by FGFR3 during tumor progression including the cross-talk with the tumor microenvironment, at least in NMIBC tumors. Our GEM model confirmed the activation of MYC by FGFR3 [14], which could contribute to an FGFR3-induced tumorigenesis through the promotion of cell hyperproliferation. Our model also highlighted an activation of the androgen receptor by a mutant FGFR3. Such results were further corroborated in human-derived preclinical models and supported by a higher activity of AR in *FGFR3*-mutated tumors compared to wild-type ones. Additional analysis of the signaling pathway leading to this ligand-independent, but FGFR3-induced activation of AR is worth further investigation. Since we did not observe any transcriptomic or post-transcriptomic regulation of AR expression levels but rather a decrease of its phosphorylation, we could assume that, as reported for EGFR [43], FGFR3 could modulate AR activity through its phosphorylation. AR activity in LumP MIBC has recently been proposed to favor luminal differentiation and to account for the enrichment of males in LumP MIBC [38]. Therefore, the hyperactivation of AR in FGFR3-mutated tumors could likely contribute to the obvious biased ratio of FGFR3-induced tumors in males versus females in LumP MIBC and maybe also in class 3 NMIBC [7]. Due to the globally higher proportion of males presenting BCa, the interest in understanding the role of AR during bladder tumor development and evaluating AR as a therapeutic target has been the focus of several studies and clinical trials [44,45]. Our results suggest that this interest should be particularly important in an *FGFR3*-mutated context.

## Conclusion

Our work demonstrates the driver oncogenic role of a mutated FGFR3 in bladder, bladder tumor formation *in vivo*, which could support the indication extension of FGFR inhibitors to early stages of bladder tumor development. Our model also sheds light on a more prevalent male gender-bias in human mutated FGFR3-induced tumors compared to wild-type ones suggesting a role of mutant-FGFR3 in the gender disparity of bladder cancer. Finally, our results allow to propose FGFR3 expression level in normal epithelium as a determinant of the FGFR3-driven tumors tissue specificity.

## Abbreviations

(FGFR): fibroblast growth factor receptor
(AR): Androgen receptor
(FCS): fetal calf serum
(NMIBC): non-muscle-invasive bladder cancer
(MIBC): muscle-invasive bladder cancer
(RT-qPCR): reverse transcription-quantitative polymerase chain reaction

## DECLARATIONS

### Ethics approval and consent to participate

Not relevant here

### Consent for publication

Not relevant here

### Data Availability

RNAseq data for a cohort of 476 tumors including 460 NMIBC was obtained from ArrayExpress E-MTAB-4321 [5]. RNAseq data and *FGFR3* mutational status for 407 TCGA MIBC were downloaded from cBioPortal for Cancer Genomics:http://www.cbioportal.org/. The microarray for MGH-U3, RT112 and UM-UC-14 cells treated with FGFR3 siRNA are available from GEO (https://www.ncbi.nlm.nih.gov/geo/) under accession number GSE84733 and GSE125547.

All other publicly available data used in the current study can be found as described in the Methods. Other data that support the findings of this study are available from the corresponding author upon reasonable request.

The microarray for mouse tumors and hyperplasia and urothelium from control littermate have been deposit on GEO under reference GSE151888

### Conflict of interest disclosures

The authors have no potential conflict of interest to declare.

### Fundings

This work was supported by a grant from *Ligue Nationale Contre le Cancer* (AMV, MS, JF, XM, FD, CDL, EC, ML, YA, FR, IBP) as an associated team (*Equipe labellisée*), the *“Carte d’Identité des Tumeurs”* program initiated, developed and funded by *Ligue Nationale Contre le Cancer*. MS was supported by National Natural Science Foundation of China (NSFC, 82002672). AMV was supported by a scholarship from the French ministry of research. XM was supported by a fellowship from ITMO Cancer AVIESAN, within the framework of Cancer Plan. JF was supported by the *Fondation ARC pour la recherche sur le cancer*.

## Author’s contribution

MS, AMV, JF, XM designed, performed experiments or bioinformatics analyses, analyzed and interpreted the data. AMV and XM performed most of the bioinformatics analyses together with help of AK and SVL for tumor’s classification and of SVL for Androgen Receptor regulon analysis. MS, AMV and FD performed the TF array, genotyping of mice colonies and analysis of Androgen Receptor activation by FGFR3. JF performed immuno-histopathological analysis of the mouse tumors and hyperplasia together with OL for hyperplasia under YA supervision. AR analyzed FGFR3 expression in different epithelia under supervision of AA. CDL established the mouse model and select the founders based of FGFR3 expression. MLL and IBP followed tumor formation in mice by ultrasound echography and performed tumor resection before DNA/RNA extraction. EC normalized and annotated Affymetrix mouse array data and centralized the data for bioinformatics analyses. TL provided samples and help with clinical data analysis for the CIT series. ADR supervised AK for classification of tumors. LD supervised SVL for tumors’ classification and Androgen Receptor regulon analysis. FR and IBP designed and supervised research, analyzed and interpreted the data. IBP coordinated the study. IBP, MS, AMV, JF and XM wrote the paper. All authors critically read and contributed to the final version of the manuscript.

## Acknowledgments

We thank Xue-Ru Wu and Tung-Tien Sun (NYU), for providing the plasmid with the murine uroplakin II promoter, Jeanne-Marie Girault for the hFGFR3-S249C plasmid construct, Martine Blanche for establishing the UII-hFGFR3-S249C transgenic mice, David Gentien from the genomics platforms and the animal facilities members of Institut Curie particularly Celine Daviaud and Sonia Jannet. We thank Eric Letouzé and Geco (Integragen, Evry, France) for their analysis of mutations and copy number variation in mouse tumors. We thank Ellen Zwarthoff (Erasmus MC Cancer Institute, Netherlands) for providing FGFR3 mutation status for her cohort of tumors for which transcriptomic data were publicly available.

---

**Supplementary Figure 1.** A) Validation of the relative mRNA expression levels of the human FGFR3 transgene in the urothelium of transgenic UPII-FGFR3-S249C mice. B) In situ hybridization showing expression of the human FGFR3 transgene at the supra-basal and intermediate cell layers of UPII-FGFR3-S249C mouse urothelium (line 569) (4 months of age). Magnificationx100. C) Radioactive PCR showing the expression of both human and mouse FGFR3 digested amplicons (cDNA) in a control littermate and UPII-FGFR3-S249C (line 569 and 538) mice. The bands of 59 and 48 bp correspond to mouse endogenous FGFR3, and the band of 88 bp to the human FGFR3 transgene.

**Supplementary Figure 2**. A) Survival plot of UPII-hFGFR3-S249C mice (line 569) (FGFR3 +/- or +/+) versus control mice from littermates (FGFR3 -/-) following treatment with 0.05% BBN in drinking water for 8 weeks. B) Representative immunohistochemistry staining showing Ki67 expression in hyperplasia (middle panel) and papillary urothelial carcinoma (right panel) of UPII-hFGFR3-S249C mice (line 569) and in the normal urothelium from control littermates (left panel). Scale bars: 100μm. C) Genomic DNA qPCR validation of genes found in frequently amplified regions (chromosomes 1, 16 and 17) of tumors from UPII-hFGFR3-S249C mice (line 569). Shown is the ratio of relative expressions of exonic regions of genes found in amplified chromosomes (Trat1, Erbb4, Fkbp5) against the genes found in stable chromosomes (Tgfbr3, Inpp4b). Each relative expression value was calculated using the 2-^ΔΔ^Ct method and values were normalized to control urothelia for each sample.

**Supplementary Figure 3**. A-B) Molecular classifications for both NMIBC (UROMOL) and MIBC (TCGA) cohorts as described previously [3,5,7]. A) Comparison of AR regulon activity in human bladder cancers bearing *FGFR3* mutation against tumors with *FGFR3* wild-type, separated by different molecular groups within NMIBC UROMOL cohort (left panel) and MIBC TCGA cohort (right panel), respectively. Statistical test for each comparison was adjusted by gender. B) Distribution of AR regulon activity among NMIBC classes (upper left panel) and MIBC subtypes (lower left panel). Statistical test for each comparison was adjusted by gender. NMIBC, nonmuscle-invasive bladder cancer; MIBC, muscle-invasive bladder cancer.

**Supplementary Table 1.** Differential gene expression analysis comparing hFGFR3-S249C mice hyperplasia samples (n=5) to normal urothelium (n=3) from control mouse littermates **(A)**; Differential gene expression analysis comparing hFGFR3-S249C mice tumor samples (n=6) to normal urothelium (n=3) from control mouse littermates **(B)**.

**Supplementary Table 2.** Differential gene expression analysis comparing siFGFR3 vs lipofectamine treated MGH-U3, UM-UC-14 and RT112 cells.

**Supplementary Table 3.** AR regulon gene set in NMIBC (UROMOL cohort).

## Notes

### Competing Interest Statement

The authors have declared no competing interest.

