## Supplemental figures and tables for "FGFR3 activating mutations induce luminal-like papillary bladder tumor formation and favor a male gender bias"

Sup.Fig1

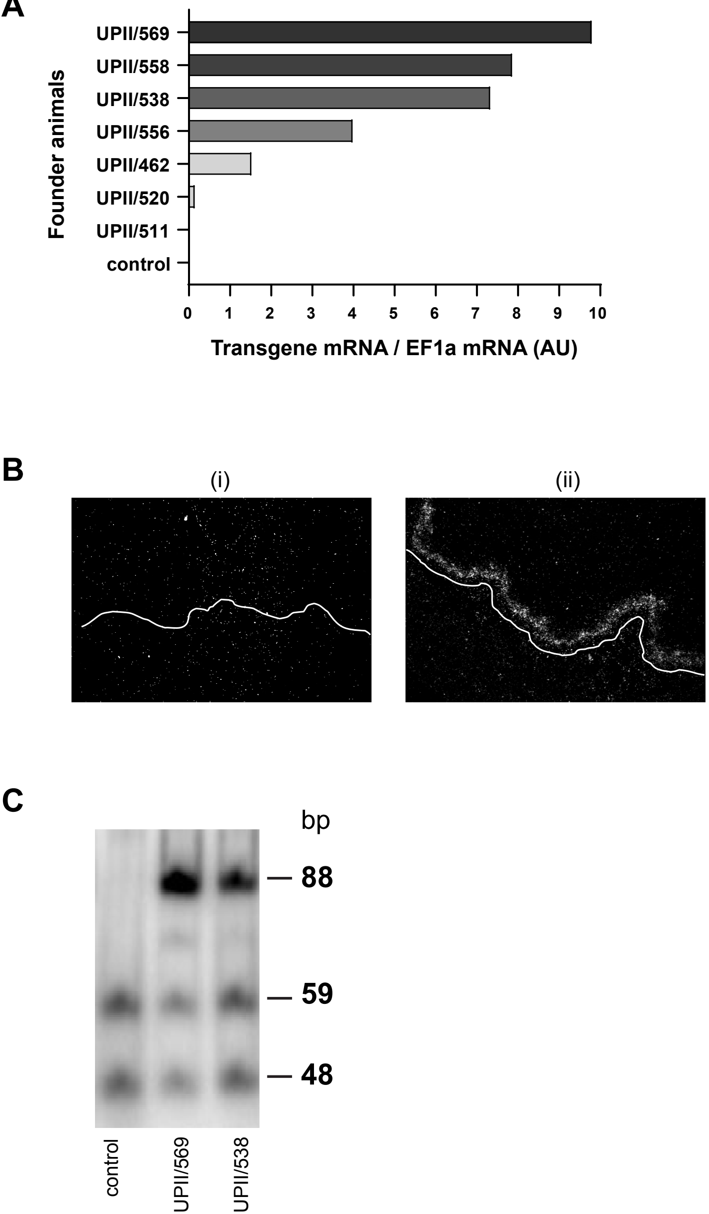
A
UPII/569
UPII/558
UPII/538
UPII/556
UPII/462
UPII/520
UPII/511
control
Founder animals
0 1 2 3 4 5 6 7 8 9 10
Transgene mRNA / EF1a mRNA (AU)
B
(i)
(ii)
C
bp
88
59
48
control
UPII/569
UPII/538

Sup.Fig2

**A**

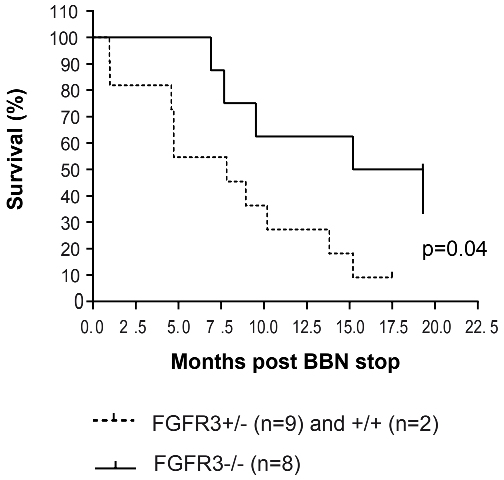
Survival (%)
110
100
90
80
70
60
50
40
30
20
10
0
0.0
2 .5
5.0
7 .5
10.0
12.5
15.0
17.5
20.0
22. 5
Months post BBN stop
p=0.04
FGFR3+/- (n=9) and +/+ (n=2)
FGFR3-/- (n=8)

**B**

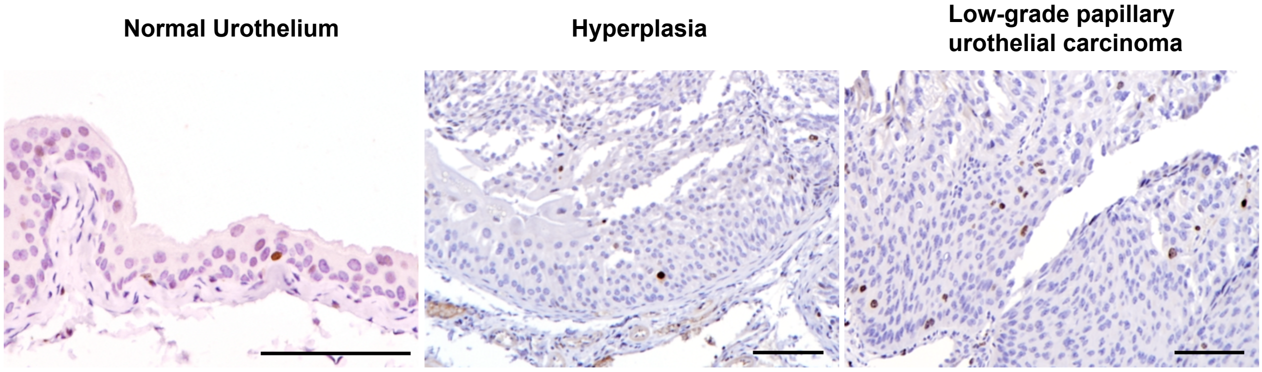
Normal Urothelium
Hyperplasia
Low-grade papillary urothelial carcinoma

**C**

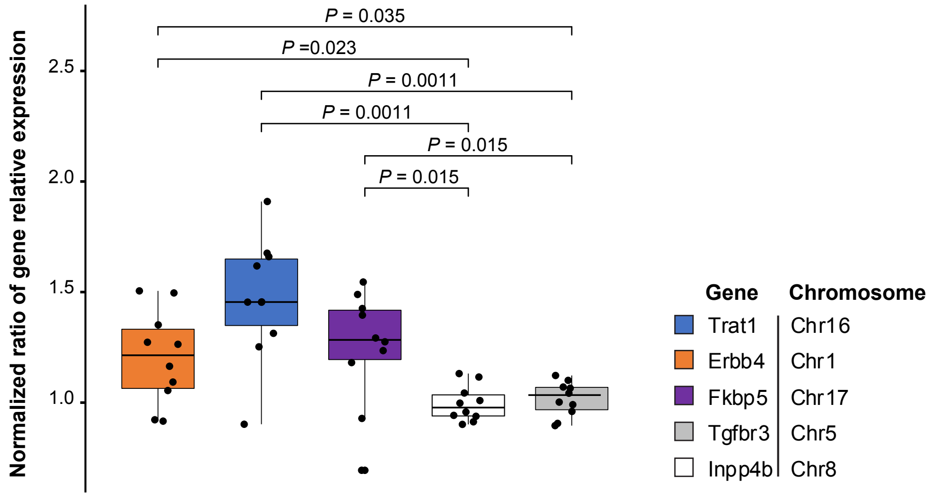
Normalized ratio of gene relative expression
2.5
2.0
1.5
1.0
P = 0.035
P =0.023
P = 0.0011
P = 0.0011
P = 0.015
P = 0.015
Gene
Chromosome
Trat1
Chr16
Erbb4
Chr1
Fkbp5
Chr17
Tgfbr3
Chr5
Inpp4b
Chr8

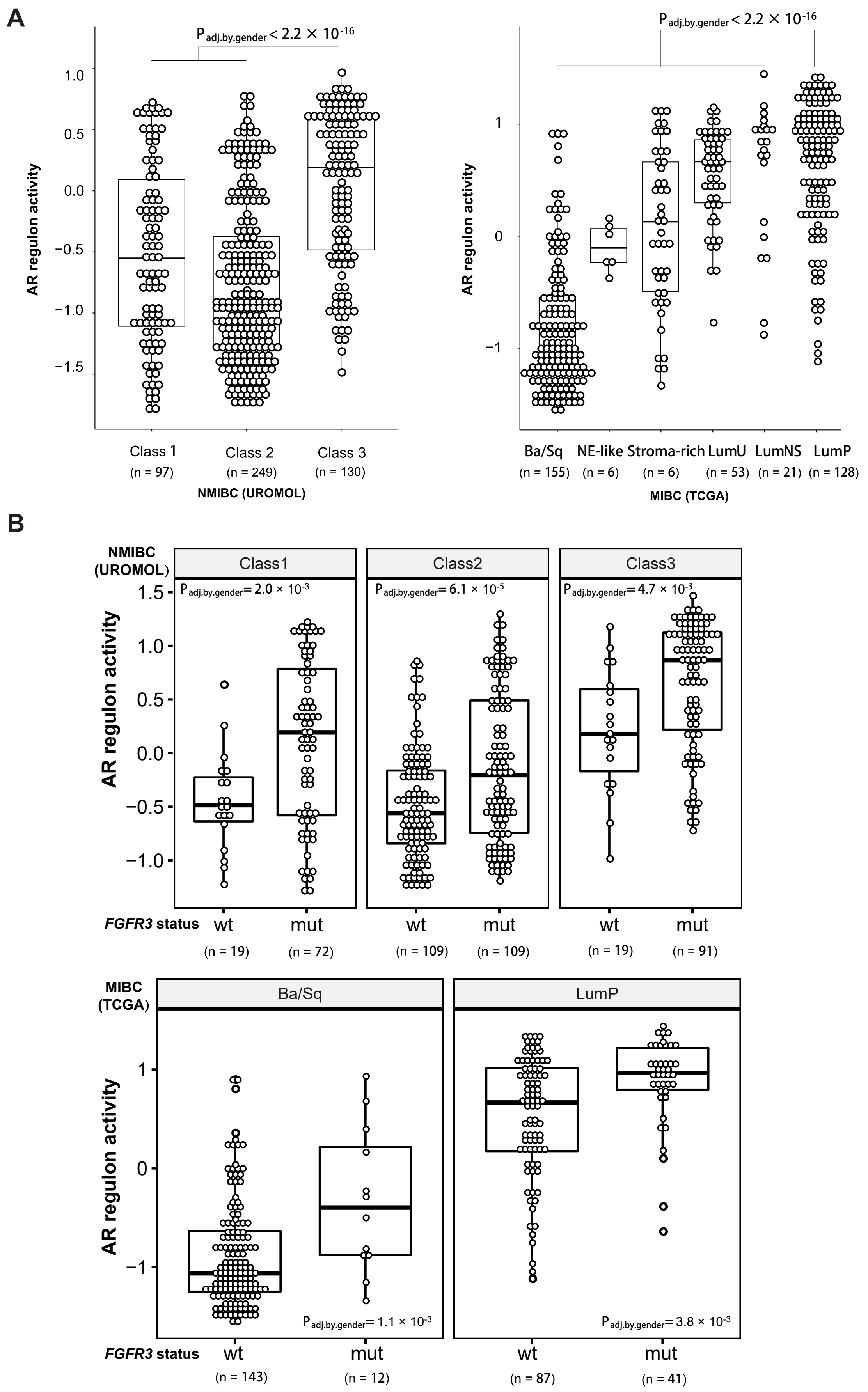

A
$P_{adj.by.gender} < 2.2 \times 10^{-16}$
AR regulon activity
1.0
0.5
0.0
−0.5
−1.0
−1.5
Class 1
(n = 97)
Class 2
(n = 249)
Class 3
(n = 130)
NMIBC (UROMOL)
$P_{adj.by.gender} < 2.2 \times 10^{-16}$
AR regulon activity
1
0
−1
Ba/Sq
(n = 155)
NE-like
(n = 6)
Stroma-rich
(n = 6)
LumU
(n = 53)
LumNS
(n = 21)
LumP
(n = 128)
MIBC (TCGA)
B
NMIBC (UROMOL)
Class1
Class2
Class3
$P_{adj.by.gender} = 2.0 \times 10^{-3}$
$P_{adj.by.gender} = 6.1 \times 10^{-5}$
$P_{adj.by.gender} = 4.7 \times 10^{-3}$
AR regulon activity
1.5
1.0
0.5
0.0
−0.5
−1.0
FGFR3 status
wt
mut
(n = 19)
(n = 72)
(n = 109)
(n = 109)
(n = 19)
(n = 91)
MIBC (TCGA)
Ba/Sq
LumP
AR regulon activity
1
0
−1
$P_{adj.by.gender} = 1.1 \times 10^{-3}$
$P_{adj.by.gender} = 3.8 \times 10^{-3}$
FGFR3 status
wt
mut
(n = 143)
(n = 12)
(n = 87)
(n = 41)

Supplementary Table 2. Differential gene expression analysis comparing siFGFR3 vs lipofectamine treated MGH-U3, UM-UC-14 and RT112 cells.

| Probeset_ID | GeneID | logFC | AveExpr | t | P.Value | adj.P.Val |
|---|---|---|---|---|---|---|
| 25805_at | BAMBI | 1,44756916 | 7,78502 | 17,7561943 | 2,97E-11 | 5,72E-07 |
| 2118_at | ETV4 | -1,7361277 | 6,18553909 | -13,2515518 | 1,65E-09 | 1,09E-05 |
| 200958_at | MUC20 | 1,55188641 | 10,1520198 | 13,2211278 | 1,70E-09 | 1,09E-05 |
| 7277_at | TUBA4A | -1,44728337 | 10,1560711 | -12,6904278 | 2,95E-09 | 1,21E-05 |
| 7832_at | BTG2 | 1,2120188 | 9,06507693 | 12,4777424 | 3,69E-09 | 1,21E-05 |
| 23584_at | VSIG2 | 1,50697457 | 8,56623456 | 12,3566477 | 4,21E-09 | 1,21E-05 |
| 25902_at | MTHFD1L | -1,22265365 | 7,49669121 | -12,3141287 | 4,40E-09 | 1,21E-05 |
| 744_at | MPPED2 | 1,41861673 | 6,57969428 | 11,3030638 | 1,37E-08 | 3,30E-05 |
| 54733_at | SLC35F2 | -0,99666769 | 8,45595348 | -11,171446 | 1,60E-08 | 3,39E-05 |
| 2896_at | GRN | 0,98563951 | 10,5274043 | 11,087132 | 1,76E-08 | 3,39E-05 |
| 121506_at | ERP27 | 1,87397406 | 8,90354501 | 10,9623076 | 2,04E-08 | 3,39E-05 |
| 23710_at | GABARAPL1 | 1,19806476 | 7,1210769 | 10,9352398 | 2,11E-08 | 3,39E-05 |
| 23158_at | TBC1D9 | 1,20501488 | 8,24157189 | 10,7130027 | 2,76E-08 | 3,62E-05 |
| 80273_at | GRPEL1 | -0,77907535 | 9,32668316 | -10,6977014 | 2,81E-08 | 3,62E-05 |
| 617_at | BCS1L | -0,76568706 | 8,15776675 | -10,6945126 | 2,82E-08 | 3,62E-05 |
| 9816_at | URB2 | -0,91287767 | 7,49767133 | -10,6245835 | 3,07E-08 | 3,70E-05 |
| 10608_at | MXD4 | 1,10401689 | 7,70059983 | 10,4569483 | 3,77E-08 | 4,27E-05 |
| 10940_at | POP1 | -0,79518869 | 7,22653367 | -10,3942183 | 4,08E-08 | 4,36E-05 |
| 2119_at | ETV5 | -1,59094177 | 5,72351029 | -10,2102193 | 5,13E-08 | 5,09E-05 |
| 9933_at | KIAA0020 | -1,06623581 | 7,8619509 | -10,1874768 | 5,28E-08 | 5,09E-05 |
| 441108_at | C5orf56 | 1,33373325 | 7,49770488 | 10,1030653 | 5,88E-08 | 5,10E-05 |
| 26579_at | MYEOV | -2,06914577 | 8,05937618 | -10,0958464 | 5,93E-08 | 5,10E-05 |
| 23223_at | RRP12 | -0,81158894 | 6,88746737 | -10,0592241 | 6,21E-08 | 5,10E-05 |
| 4869_at | NPM1 | -0,75938069 | 10,414159 | -10,0413097 | 6,36E-08 | 5,10E-05 |
| 8607_at | RUVBL1 | -0,95943401 | 8,88176967 | -9,92965402 | 7,33E-08 | 5,33E-05 |
| 10808_at | HSPH1 | -0,56145181 | 9,53648571 | -9,92162421 | 7,41E-08 | 5,33E-05 |
| 119559_at | SFXN4 | -0,6957159 | 7,96802104 | -9,90381464 | 7,58E-08 | 5,33E-05 |
| 9188_at | DDX21 | -1,13324807 | 9,94634161 | -9,8868246 | 7,75E-08 | 5,33E-05 |
| 29090_at | TIMM21 | -0,73613288 | 7,34554508 | -9,85531331 | 8,07E-08 | 5,36E-05 |
| 64759_at | TNS3 | 1,05741082 | 9,92027561 | 9,7074077 | 9,78E-08 | 6,28E-05 |
| 5708_at | PSMD2 | -0,55709093 | 10,4994066 | -9,66582214 | 1,03E-07 | 6,31E-05 |
| 11340_at | EXOSC8 | -0,99576479 | 8,23719854 | -9,65426335 | 1,05E-07 | 6,31E-05 |
| 10133_at | OPTN | 1,29721896 | 8,01434359 | 9,61262511 | 1,11E-07 | 6,47E-05 |
| 23327_at | NEDD4L | 0,78317109 | 7,87533727 | 9,57472938 | 1,16E-07 | 6,60E-05 |
| 1662_at | DDX10 | -0,809869 | 7,46292014 | -9,53644225 | 1,23E-07 | 6,70E-05 |
| 151963_at | MB21D2 | 0,96990067 | 7,82185411 | 9,50626024 | 1,28E-07 | 6,70E-05 |
| 653602_at | LOC653602 | 1,01835191 | 6,87849893 | 9,49900395 | 1,29E-07 | 6,70E-05 |
| 10906_at | TRAFD1 | 0,82881842 | 9,20931755 | 9,47745732 | 1,33E-07 | 6,72E-05 |
| 10514_at | MYBBP1A | -0,63908908 | 6,88455044 | -9,39772377 | 1,47E-07 | 7,05E-05 |
| 10419_at | PRMT5 | -0,80730291 | 9,77854869 | -9,39508138 | 1,48E-07 | 7,05E-05 |
| 960_at | CD44 | -1,16092142 | 9,34722028 | -9,38428487 | 1,50E-07 | 7,05E-05 |
| 6472_at | SHMT2 | -0,71091374 | 9,24038532 | -9,32491312 | 1,63E-07 | 7,46E-05 |
| 2043_at | EPHA4 | 1,41809387 | 8,53611307 | 9,25887993 | 1,78E-07 | 7,87E-05 |
| 2717_at | GLA | -0,86093906 | 8,72034193 | -9,23718755 | 1,83E-07 | 7,87E-05 |

| | | | | | | |
|---|---|---|---|---|---|---|
| 219927_at | MRPL21 | -0,64217646 | 9,37452804 | -9,23394723 | 1,84E-07 | 7,87E-05 |
| 113791_at | PIK3IP1 | 1,14126459 | 6,92309239 | 9,13671823 | 2,10E-07 | 8,79E-05 |
| 10969_at | EBNA1BP2 | -0,93180926 | 9,46925927 | -9,08023722 | 2,27E-07 | 9,30E-05 |
| 5704_at | PSMC4 | -0,56893112 | 10,185036 | -9,02969716 | 2,43E-07 | 9,72E-05 |
| 81034_at | SLC25A32 | -0,87695699 | 9,13787612 | -8,99563219 | 2,55E-07 | 9,72E-05 |
| 10131_at | TRAP1 | -0,87802812 | 9,39832761 | -8,99475105 | 2,55E-07 | 9,72E-05 |
| 219_at | ALDH1B1 | -0,93046883 | 7,72283469 | -8,98441551 | 2,59E-07 | 9,72E-05 |
| 18_at | ABAT | 1,07221297 | 7,63323252 | 8,97453059 | 2,62E-07 | 9,72E-05 |
| 144165_at | PRICKLE1 | 0,93012621 | 6,99773973 | 8,94291056 | 2,74E-07 | 9,90E-05 |
| 26524_at | LATS2 | 0,84952945 | 7,70565912 | 8,93418601 | 2,78E-07 | 9,90E-05 |
| 10602_at | CDC42EP3 | 1,10970381 | 9,60869474 | 8,90299286 | 2,90E-07 | 0,00010154 |
| 51154_at | MRTO4 | -1,22192825 | 7,67421271 | -8,87207846 | 3,03E-07 | 0,00010276 |
| 677_at | ZFP36L1 | 0,64194965 | 8,15454646 | 8,86882233 | 3,04E-07 | 0,00010276 |
| 23481_at | PES1 | -0,53322771 | 8,3643011 | -8,8560224 | 3,10E-07 | 0,00010281 |
| 1200_at | TPP1 | 0,64277669 | 8,08905051 | 8,82500913 | 3,23E-07 | 0,00010288 |
| 83858_at | ATAD3B | -0,63787022 | 8,35895984 | -8,82072623 | 3,25E-07 | 0,00010288 |
| 7443_at | VRK1 | -1,24738805 | 8,11784834 | -8,81619814 | 3,27E-07 | 0,00010288 |
| 6038_at | RNASE4 | 1,54060233 | 7,23980726 | 8,80796598 | 3,31E-07 | 0,00010288 |
| 92856_at | IMP4 | -0,80856858 | 8,86869032 | -8,78614397 | 3,41E-07 | 0,0001044 |
| 58476_at | TP53INP2 | 1,70302576 | 7,35600665 | 8,77273904 | 3,48E-07 | 0,00010473 |
| 7088_at | TLE1 | 0,86649747 | 7,43447291 | 8,74235591 | 3,63E-07 | 0,00010763 |
| 11099_at | PTPN21 | 1,17227231 | 6,72725111 | 8,72330218 | 3,73E-07 | 0,0001089 |
| 84916_at | CIRH1A | -0,96479592 | 9,86821347 | -8,64817388 | 4,15E-07 | 0,0001166 |
| 51257_at | MARCH2 | 0,5807422 | 7,25474529 | 8,64532663 | 4,17E-07 | 0,0001166 |
| 10963_at | STIP1 | -0,73327853 | 10,2079995 | -8,63781498 | 4,21E-07 | 0,0001166 |
| 3853_at | KRT6A | -1,69346943 | 6,82843427 | -8,63369755 | 4,24E-07 | 0,0001166 |
| 5660_at | PSAP | 0,69070497 | 12,2425493 | 8,56053561 | 4,70E-07 | 0,00012762 |
| 56852_at | RAD18 | -0,70440434 | 5,97628361 | -8,53886849 | 4,85E-07 | 0,00012982 |
| 6949_at | TCOF1 | -0,76885586 | 7,26755723 | -8,52042576 | 4,98E-07 | 0,00013148 |
| 83595_at | SOX7 | -0,90815025 | 7,36241327 | -8,50890039 | 5,07E-07 | 0,00013187 |
| 10244_at | RABEPK | -0,48969654 | 7,91125255 | -8,47551966 | 5,32E-07 | 0,00013652 |
| 8602_at | NOP14 | -0,82766239 | 8,92333785 | -8,4552695 | 5,47E-07 | 0,00013835 |
| 55715_at | DOK4 | 0,44250174 | 6,59436376 | 8,44587358 | 5,55E-07 | 0,00013835 |
| 51062_at | ATL1 | 1,5026221 | 6,04869865 | 8,43913621 | 5,60E-07 | 0,00013835 |
| 3376_at | IARS | -0,8007755 | 9,9459394 | -8,3765022 | 6,14E-07 | 0,0001496 |
| 64081_at | PBLD | 0,87653837 | 6,32701638 | 8,33808375 | 6,49E-07 | 0,00015623 |
| 80824_at | DUSP16 | 0,96724818 | 7,43911324 | 8,30341303 | 6,83E-07 | 0,00016232 |
| 25840_at | METTL7A | 1,00410025 | 8,76284435 | 8,29028333 | 6,96E-07 | 0,00016345 |
| 5699_at | PSMB10 | 0,77372443 | 9,06674768 | 8,2715882 | 7,15E-07 | 0,00016348 |
| 79731_at | NARS2 | -0,57342864 | 8,18276327 | -8,26980115 | 7,17E-07 | 0,00016348 |
| 669_at | BPGM | 1,21904539 | 9,43410006 | 8,25886168 | 7,29E-07 | 0,00016348 |
| 2548_at | GAA | 0,78893295 | 8,03035749 | 8,25730311 | 7,30E-07 | 0,00016348 |
| 55651_at | NHP2 | -0,70083669 | 9,61312058 | -8,24979877 | 7,38E-07 | 0,00016348 |
| 25929_at | GEMIN5 | -0,75354141 | 8,13861545 | -8,23132596 | 7,59E-07 | 0,00016372 |
| 10469_at | TIMM44 | -0,52767002 | 7,78859429 | -8,22983269 | 7,60E-07 | 0,00016372 |
| 57509_at | MTUS1 | 0,92408869 | 7,72211128 | 8,22575809 | 7,65E-07 | 0,00016372 |
| 1831_at | TSC22D3 | 1,03889956 | 8,17345792 | 8,20887651 | 7,84E-07 | 0,000166 |
| 6950_at | TCP1 | -0,64283786 | 9,50345393 | -8,19701814 | 7,98E-07 | 0,00016709 |
| 23659_at | PLA2G15 | 0,62547714 | 7,67659116 | 8,17667331 | 8,22E-07 | 0,00017034 |
| 55706_at | NDC1 | -1,15875444 | 8,7456503 | -8,1687994 | 8,32E-07 | 0,0001705 |

| | | | | | | |
|---|---|---|---|---|---|---|
| 79711_at | IPO4 | -1,13574887 | 8,15075111 | -8,15779968 | 8,46E-07 | 0,00017131 |
| 64434_at | NOM1 | -0,69122716 | 6,89883015 | -8,14477153 | 8,62E-07 | 0,00017131 |
| 1050_at | CEBPA | 0,85066187 | 8,78726355 | 8,14436754 | 8,63E-07 | 0,00017131 |
| 9982_at | FGFBP1 | -2,03014229 | 8,38060614 | -8,13237652 | 8,78E-07 | 0,0001726 |
| 7203_at | CCT3 | -0,55909082 | 11,0418385 | -8,11302748 | 9,04E-07 | 0,00017584 |
| 81570_at | CLPB | -0,54397312 | 6,96844464 | -8,08451694 | 9,43E-07 | 0,00018162 |
| 51096_at | UTP18 | -0,71878659 | 9,09206183 | -8,0470492 | 9,97E-07 | 0,00018715 |
| 5198_at | PFAS | -1,05362357 | 7,41446166 | -8,04379127 | 1,00E-06 | 0,00018715 |
| 55379_at | LRRC59 | -0,4756368 | 10,6456751 | -8,0425028 | 1,00E-06 | 0,00018715 |
| 3145_at | HMBS | -0,75079571 | 8,47834525 | -8,036463 | 1,01E-06 | 0,00018715 |
| 1848_at | DUSP6 | -2,60936043 | 7,67069864 | -8,02647184 | 1,03E-06 | 0,00018715 |
| 55157_at | DARS2 | -0,7489429 | 7,51724061 | -8,02053514 | 1,04E-06 | 0,00018715 |
| 6609_at | SMPD1 | 0,63687089 | 6,51379571 | 8,0153998 | 1,05E-06 | 0,00018715 |
| 23170_at | TTLL12 | -0,58789339 | 8,84164206 | -8,01293768 | 1,05E-06 | 0,00018715 |
| 9688_at | NUP93 | -0,82417148 | 9,14306258 | -8,00594708 | 1,06E-06 | 0,00018738 |
| 9631_at | NUP155 | -0,90804235 | 8,5341172 | -7,99537806 | 1,08E-06 | 0,00018865 |
| 8764_at | TNFRSF14 | 0,84309129 | 7,11078705 | 7,97494352 | 1,11E-06 | 0,00019165 |
| 57522_at | SRGAP1 | 0,54952818 | 6,91743122 | 7,9720703 | 1,12E-06 | 0,00019165 |
| 54606_at | DDX56 | -0,59910674 | 9,01908766 | -7,96186404 | 1,13E-06 | 0,00019165 |
| 23160_at | WDR43 | -1,00046993 | 8,05839153 | -7,95692862 | 1,14E-06 | 0,00019165 |
| 6183_at | MRPS12 | -0,86875263 | 7,41351888 | -7,95400778 | 1,15E-06 | 0,00019165 |
| 5663_at | PSEN1 | 0,46361613 | 8,77958296 | 7,94470941 | 1,16E-06 | 0,00019165 |
| 51231_at | VRK3 | 0,49222464 | 8,1692358 | 7,94169208 | 1,17E-06 | 0,00019165 |
| 165055_at | CCDC138 | -0,50528795 | 6,00433104 | -7,93817585 | 1,17E-06 | 0,00019165 |
| 2193_at | FARSA | -0,58937308 | 9,23315585 | -7,90289617 | 1,24E-06 | 0,0002002 |
| 8458_at | TTF2 | -1,14233825 | 7,75745912 | -7,89813464 | 1,25E-06 | 0,0002002 |
| 158_at | ADSL | -0,51883763 | 9,59691643 | -7,87706372 | 1,29E-06 | 0,00020431 |
| 768_at | CA9 | -0,8038967 | 5,87614174 | -7,87381475 | 1,29E-06 | 0,00020431 |
| 4839_at | NOP2 | -0,86956846 | 8,47308225 | -7,86527471 | 1,31E-06 | 0,00020529 |
| 10598_at | AHSA1 | -0,76613334 | 9,98461139 | -7,84917554 | 1,34E-06 | 0,00020867 |
| 4082_at | MARCKS | 0,87815535 | 10,2970153 | 7,81416948 | 1,42E-06 | 0,00021821 |
| 54801_at | HAUS6 | -0,87472461 | 6,62274382 | -7,80925436 | 1,43E-06 | 0,00021821 |
| 55716_at | LMBR1L | 0,91204581 | 7,00350722 | 7,80402316 | 1,44E-06 | 0,00021822 |
| 26137_at | ZBTB20 | 1,18433818 | 5,15276162 | 7,79683618 | 1,45E-06 | 0,0002189 |
| 1737_at | DLAT | -0,56573888 | 8,49804932 | -7,77339402 | 1,51E-06 | 0,00022512 |
| 3326_at | HSP90AB1 | -0,53236135 | 11,9790408 | -7,76190492 | 1,53E-06 | 0,00022696 |
| 9790_at | BMS1 | -0,61201771 | 8,15598897 | -7,75800487 | 1,54E-06 | 0,00022696 |
| 23654_at | PLXNB2 | 0,61981391 | 10,995074 | 7,72729933 | 1,62E-06 | 0,00023609 |
| 10807_at | SDCCAG3 | -0,69312374 | 8,37078686 | -7,70861982 | 1,66E-06 | 0,00023953 |
| 25833_at | POU2F3 | 1,46837247 | 6,01085134 | 7,70806019 | 1,67E-06 | 0,00023953 |
| 57291_at | DANCR | -0,75586991 | 9,45587973 | -7,70311393 | 1,68E-06 | 0,00023957 |
| 11222_at | MRPL3 | -0,61778617 | 10,7906324 | -7,69411573 | 1,70E-06 | 0,00024112 |
| 55281_at | TMEM140 | 1,79211648 | 7,04710532 | 7,6495624 | 1,82E-06 | 0,00025637 |
| 113174_at | SAAL1 | -0,81685242 | 7,32873989 | -7,6256703 | 1,89E-06 | 0,0002616 |
| 891_at | CCNB1 | -1,25660948 | 8,91515341 | -7,62251628 | 1,90E-06 | 0,0002616 |
| 219285_at | SAMD9L | 1,55191658 | 8,70214057 | 7,62247296 | 1,90E-06 | 0,0002616 |
| 10957_at | PNRC1 | 1,3571165 | 9,58709351 | 7,61605466 | 1,92E-06 | 0,00026234 |
| 64285_at | RHBDF1 | 0,48758543 | 7,54647127 | 7,60630168 | 1,95E-06 | 0,00026446 |
| 59342_at | SCPEP1 | 0,78282261 | 11,0869123 | 7,59291607 | 1,99E-06 | 0,00026527 |
| 4927_at | NUP88 | -0,78801004 | 8,74136946 | -7,58839891 | 2,00E-06 | 0,00026527 |

| | | | | | | |
|---|---|---|---|---|---|---|
| 11329_at | STK38 | 0,74013103 | 9,15285246 | 7,58785289 | 2,01E-06 | 0,00026527 |
| 4733_at | DRG1 | -0,42848656 | 9,26202419 | -7,5834025 | 2,02E-06 | 0,00026527 |
| 10534_at | SSSCA1 | -0,49006212 | 8,60012343 | -7,58199415 | 2,02E-06 | 0,00026527 |
| 10885_at | WDR3 | -0,88805062 | 8,55034533 | -7,552786 | 2,12E-06 | 0,00027572 |
| 643008_at | SMIM5 | 1,04331178 | 6,92930839 | 7,53053229 | 2,19E-06 | 0,00028263 |
| 124454_at | EARS2 | -0,39194108 | 6,86149673 | -7,52827757 | 2,20E-06 | 0,00028263 |
| 255488_at | RNF144B | 0,77018591 | 7,02151789 | 7,51612488 | 2,24E-06 | 0,00028613 |
| 169714_at | QSOX2 | -0,55706276 | 7,18457122 | -7,51115808 | 2,26E-06 | 0,00028646 |
| 80221_at | ACSF2 | 0,79258827 | 8,80873753 | 7,50580569 | 2,28E-06 | 0,00028697 |
| 8891_at | EIF2B3 | -0,76370973 | 7,56624057 | -7,49366487 | 2,32E-06 | 0,00029045 |
| 29902_at | FAM216A | -1,14902823 | 6,17096489 | -7,48770888 | 2,34E-06 | 0,00029045 |
| 57659_at | ZBTB4 | 0,66245087 | 8,56804479 | 7,48568851 | 2,35E-06 | 0,00029045 |
| 471_at | ATIC | -0,65706273 | 10,3569258 | -7,47487554 | 2,39E-06 | 0,00029352 |
| 114034_at | TOE1 | -0,56306866 | 7,6804029 | -7,43222767 | 2,56E-06 | 0,00031058 |
| 901_at | CCNG2 | 1,17135464 | 7,49648988 | 7,43080907 | 2,56E-06 | 0,00031058 |
| 116985_at | ARAP1 | 0,52216108 | 7,9938459 | 7,41914197 | 2,61E-06 | 0,00031334 |
| 84955_at | NUDCD1 | -0,72215518 | 7,56676294 | -7,41655178 | 2,62E-06 | 0,00031334 |
| 4234_at | METTL1 | -0,85802953 | 7,19331292 | -7,4132936 | 2,64E-06 | 0,00031334 |
| 56915_at | EXOSC5 | -0,75476811 | 8,32129057 | -7,40920605 | 2,65E-06 | 0,00031343 |
| 283537_at | SLC46A3 | 1,09673624 | 5,24513076 | 7,38999136 | 2,73E-06 | 0,00032109 |
| 9037_at | SEMA5A | 0,84542845 | 9,02910871 | 7,37991832 | 2,78E-06 | 0,00032426 |
| 2288_at | FKBP4 | -0,64754743 | 6,8966337 | -7,36783138 | 2,83E-06 | 0,00032852 |
| 23075_at | SWAP70 | 0,51364739 | 8,66730278 | 7,34011974 | 2,96E-06 | 0,00034118 |
| 6157_at | RPL27A | -0,72852567 | 5,50816495 | -7,33449099 | 2,98E-06 | 0,00034219 |
| 6249_at | CLIP1 | 0,56891417 | 8,18865913 | 7,32986286 | 3,01E-06 | 0,00034266 |
| 22984_at | PDCD11 | -0,53002559 | 7,58515414 | -7,32610474 | 3,02E-06 | 0,00034268 |
| 114800_at | CCDC85A | 1,3714257 | 6,70839647 | 7,31584989 | 3,07E-06 | 0,00034627 |
| 6778_at | STAT6 | 0,48686306 | 9,32067595 | 7,29586382 | 3,17E-06 | 0,00035336 |
| 4715_at | NDUFB9 | -0,51326038 | 10,4162327 | -7,29575257 | 3,17E-06 | 0,00035336 |
| 53349_at | ZFYVE1 | 0,81514018 | 6,72747016 | 7,27778782 | 3,27E-06 | 0,00036042 |
| 51491_at | NOP16 | -1,23529448 | 7,84902526 | -7,27297271 | 3,29E-06 | 0,00036042 |
| 51253_at | MRPL37 | -0,71170688 | 9,77032392 | -7,27251203 | 3,29E-06 | 0,00036042 |
| 64581_at | CLEC7A | 1,87852686 | 6,67743127 | 7,26162144 | 3,35E-06 | 0,00036466 |
| 84319_at | CMSS1 | -1,32815193 | 8,92566097 | -7,24257913 | 3,45E-06 | 0,00037379 |
| 6402_at | SELL | 1,0639441 | 5,07620196 | 7,23309781 | 3,51E-06 | 0,00037737 |
| 22863_at | ATG14 | 0,59681172 | 7,00019876 | 7,22838875 | 3,53E-06 | 0,00037811 |
| 2634_at | GBP2 | 1,17915139 | 8,0256135 | 7,22012986 | 3,58E-06 | 0,00038102 |
| 1723_at | DHODH | -0,88462885 | 6,12872455 | -7,19574898 | 3,72E-06 | 0,00039401 |
| 57122_at | NUP107 | -0,96053618 | 8,75911217 | -7,1910894 | 3,75E-06 | 0,00039479 |
| 3959_at | LGALS3BP | 0,52573378 | 11,5712066 | 7,16760387 | 3,89E-06 | 0,00040772 |
| 10726_at | NUDC | -0,75034914 | 9,74313497 | -7,16286935 | 3,92E-06 | 0,00040862 |
| 54663_at | WDR74 | -0,83114854 | 8,72795916 | -7,15331886 | 3,99E-06 | 0,00040915 |
| 29889_at | GNL2 | -0,64013987 | 8,61757731 | -7,15255823 | 3,99E-06 | 0,00040915 |
| 54677_at | CROT | 0,69832278 | 7,22578846 | 7,15204355 | 3,99E-06 | 0,00040915 |
| 4924_at | NUCB1 | 0,57395442 | 8,17237284 | 7,14560424 | 4,03E-06 | 0,00041123 |
| 10868_at | USP20 | 0,4103657 | 6,67857001 | 7,13995003 | 4,07E-06 | 0,0004128 |
| 9533_at | POLR1C | -0,50196622 | 7,98375305 | -7,11694976 | 4,23E-06 | 0,00042615 |
| 23642_at | SNHG1 | -1,19517689 | 9,06754502 | -7,11043853 | 4,27E-06 | 0,0004284 |
| 8624_at | PSMG1 | -0,7120623 | 10,4497908 | -7,10067454 | 4,34E-06 | 0,00043295 |
| 4691_at | NCL | -0,62524904 | 12,0860757 | -7,09502011 | 4,38E-06 | 0,00043468 |

| | | | | | | |
|---|---|---|---|---|---|---|
| 63874_at | ABHD4 | 0,57639978 | 8,85877256 | 7,07158505 | 4,55E-06 | 0,00044915 |
| 5164_at | PDK2 | 0,41683041 | 6,8023345 | 7,05850951 | 4,64E-06 | 0,00045554 |
| 6692_at | SPINT1 | 0,71246432 | 10,4016363 | 7,05656633 | 4,66E-06 | 0,00045554 |
| 54069_at | MIS18A | -0,92510938 | 8,29929775 | -7,02755814 | 4,88E-06 | 0,00047509 |
| 55690_at | PACS1 | 0,48921463 | 7,10003068 | 7,02377241 | 4,91E-06 | 0,00047562 |
| 55732_at | C1orf112 | -1,03412114 | 6,77954621 | -7,01440558 | 4,99E-06 | 0,0004805 |
| 53838_at | C11orf24 | -0,66911078 | 8,7311625 | -7,01060713 | 5,02E-06 | 0,00048107 |
| 10813_at | UTP14A | -0,63385755 | 7,16258005 | -6,99049458 | 5,19E-06 | 0,0004929 |
| 54517_at | PUS7 | -1,28867078 | 8,14454817 | -6,98961221 | 5,19E-06 | 0,0004929 |
| 55759_at | WDR12 | -1,17668319 | 8,71984589 | -6,9838741 | 5,24E-06 | 0,0004951 |
| 79654_at | HECTD3 | 0,64729349 | 7,72476799 | 6,97911535 | 5,28E-06 | 0,00049652 |
| 113115_at | MTFR2 | -1,13719868 | 6,82006153 | -6,97013093 | 5,36E-06 | 0,00050141 |
| 10472_at | ZBTB18 | 0,59833931 | 5,43352115 | 6,96266018 | 5,43E-06 | 0,00050511 |
| 60491_at | NIF3L1 | -0,67308448 | 8,73363982 | -6,94232182 | 5,61E-06 | 0,00051968 |
| 57696_at | DDX55 | -0,5660353 | 7,34158569 | -6,93328516 | 5,70E-06 | 0,0005249 |
| 84967_at | LSM10 | -0,47880189 | 7,92661394 | -6,93028775 | 5,72E-06 | 0,00052497 |
| 9764_at | KIAA0513 | 0,49107972 | 6,28441762 | 6,92548769 | 5,77E-06 | 0,00052587 |
| 3916_at | LAMP1 | 0,40609406 | 11,271716 | 6,92345063 | 5,79E-06 | 0,00052587 |
| 7347_at | UCHL3 | -0,65489851 | 9,07748566 | -6,91694669 | 5,85E-06 | 0,00052902 |
| 55720_at | TSR1 | -0,70748802 | 7,71118654 | -6,90992589 | 5,92E-06 | 0,00053063 |
| 10713_at | USP39 | -0,50685751 | 8,44742538 | -6,90938447 | 5,92E-06 | 0,00053063 |
| 5831_at | PYCR1 | -0,48374963 | 8,22083142 | -6,89489208 | 6,07E-06 | 0,00053871 |
| 23590_at | PDSS1 | -1,21280888 | 6,67597312 | -6,89454947 | 6,07E-06 | 0,00053871 |
| 88455_at | ANKRD13A | 0,58711725 | 9,73744088 | 6,88828935 | 6,13E-06 | 0,00054178 |
| 10056_at | FARSB | -1,09306236 | 9,41730733 | -6,87989878 | 6,22E-06 | 0,0005468 |
| 80700_at | UBXN6 | 0,49008435 | 8,3237665 | 6,86843505 | 6,33E-06 | 0,00055468 |
| 1628_at | DBP | 1,0821764 | 7,73580573 | 6,86326926 | 6,39E-06 | 0,00055539 |
| 3313_at | HSPA9 | -0,64787722 | 9,74099221 | -6,86215752 | 6,40E-06 | 0,00055539 |
| 5036_at | PA2G4 | -0,78559558 | 8,93959499 | -6,85484388 | 6,48E-06 | 0,00055937 |
| 200734_at | SPRED2 | -1,03878693 | 6,55648807 | -6,85237805 | 6,50E-06 | 0,00055937 |
| 56919_at | DHX33 | -0,47913843 | 7,77892312 | -6,84322298 | 6,60E-06 | 0,00056536 |
| 58477_at | SRPRB | -0,65394143 | 8,7724939 | -6,83752411 | 6,67E-06 | 0,00056817 |
| 1601_at | DAB2 | 1,02187884 | 6,13256216 | 6,83353855 | 6,71E-06 | 0,0005694 |
| 51602_at | NOP58 | -0,81542274 | 10,0230711 | -6,81244245 | 6,95E-06 | 0,00058702 |
| 57559_at | STAMBPL1 | -1,096177 | 6,10202075 | -6,7889717 | 7,22E-06 | 0,00060761 |
| 9136_at | RRP9 | -0,69859258 | 7,30424767 | -6,7841478 | 7,28E-06 | 0,00060789 |
| 54869_at | EPS8L1 | 0,67705808 | 7,12599607 | 6,78108002 | 7,32E-06 | 0,00060789 |
| 104797538_a | KCNMB2-AS1 | 0,93937188 | 4,59758748 | 6,77947518 | 7,34E-06 | 0,00060789 |
| 57650_at | KIAA1524 | -1,20157762 | 5,70011323 | -6,7782398 | 7,35E-06 | 0,00060789 |
| 55304_at | SPTLC3 | 1,69368766 | 7,16768964 | 6,76737432 | 7,49E-06 | 0,00061631 |
| 3312_at | HSPA8 | -0,43771866 | 12,7760888 | -6,75098647 | 7,69E-06 | 0,00063037 |
| 8662_at | EIF3B | -0,40383078 | 9,51021248 | -6,74867223 | 7,72E-06 | 0,00063037 |
| 10916_at | MAGED2 | 0,88246178 | 8,84773753 | 6,74493615 | 7,77E-06 | 0,00063162 |
| 79077_at | DCTPP1 | -0,86160689 | 10,2308755 | -6,73590001 | 7,89E-06 | 0,00063849 |
| 79866_at | BORA | -0,91410728 | 7,59763783 | -6,72487953 | 8,03E-06 | 0,00064759 |
| 3419_at | IDH3A | -0,93957865 | 9,34214802 | -6,7003222 | 8,37E-06 | 0,00067185 |
| 51646_at | YPEL5 | 1,30467162 | 9,02673673 | 6,68610364 | 8,57E-06 | 0,00068461 |
| 80324_at | PUS1 | -0,92784061 | 7,49333232 | -6,68139979 | 8,64E-06 | 0,00068461 |
| 64946_at | CENPH | -1,39130673 | 7,31100616 | -6,68047772 | 8,65E-06 | 0,00068461 |
| 3300_at | DNAJB2 | 0,49764334 | 8,15514815 | 6,67916045 | 8,67E-06 | 0,00068461 |

| | | | | | | |
|---|---|---|---|---|---|---|
| 9093_at | DNAJA3 | -0,55126196 | 9,17341446 | -6,67626139 | 8,71E-06 | 0,00068513 |
| 10492_at | SYNCRIP | -0,51449304 | 8,87233617 | -6,67367652 | 8,75E-06 | 0,0006853 |
| 26135_at | SERBP1 | -0,59841509 | 8,1715532 | -6,67044904 | 8,80E-06 | 0,00068622 |
| 4016_at | LOXL1 | -2,65651767 | 7,6320302 | -6,64927243 | 9,12E-06 | 0,00070813 |
| 201895_at | SMIM14 | 1,13653979 | 8,66975769 | 6,63701056 | 9,31E-06 | 0,0007183 |
| 54800_at | KLHL24 | 1,55567364 | 6,19122696 | 6,6358326 | 9,32E-06 | 0,0007183 |
| 7716_at | VEZF1 | 0,40141524 | 8,91682896 | 6,63359695 | 9,36E-06 | 0,0007183 |
| 57326_at | PBXIP1 | 1,28032192 | 6,92986475 | 6,62699291 | 9,46E-06 | 0,00072343 |
| 23082_at | PPRC1 | -0,75193395 | 7,92594235 | -6,6122491 | 9,70E-06 | 0,00073864 |
| 11260_at | XPOT | -0,62424791 | 9,06762491 | -6,60265886 | 9,86E-06 | 0,00074394 |
| 55622_at | TTC27 | -0,62858787 | 7,57155943 | -6,60181678 | 9,87E-06 | 0,00074394 |
| 1984_at | EIF5A | -0,66180915 | 9,34812102 | -6,60099072 | 9,89E-06 | 0,00074394 |
| 129401_at | NUP35 | -0,9744554 | 7,89017566 | -6,59445401 | 1,00E-05 | 0,00074924 |
| 57655_at | GRAMD1A | 0,63655281 | 7,28688454 | 6,58843875 | 1,01E-05 | 0,00075349 |
| 159371_at | SLC35G1 | -0,54117692 | 6,62025017 | -6,58648872 | 1,01E-05 | 0,00075349 |
| 3730_at | ANOS1 | 1,71707346 | 7,75363905 | 6,57788262 | 1,03E-05 | 0,00076156 |
| 23503_at | ZFYVE26 | 0,85559622 | 7,41803365 | 6,56596778 | 1,05E-05 | 0,00077404 |
| 9238_at | TBRG4 | -0,62994302 | 8,60522156 | -6,56000278 | 1,06E-05 | 0,00077681 |
| 151648_at | SGOL1 | -0,96311817 | 6,12222539 | -6,55794528 | 1,06E-05 | 0,00077681 |
| 8668_at | EIF3I | -0,39806819 | 11,5477136 | -6,55708009 | 1,06E-05 | 0,00077681 |
| 5702_at | PSMC3 | -0,67210526 | 9,6245391 | -6,54855356 | 1,08E-05 | 0,00078511 |
| 6414_at | SEPP1 | 2,90803412 | 6,3642669 | 6,53657835 | 1,10E-05 | 0,00079815 |
| 5723_at | PSPH | -0,46945863 | 7,68314358 | -6,53257847 | 1,11E-05 | 0,00079903 |
| 10622_at | POLR3G | -0,62827663 | 4,63216174 | -6,53150166 | 1,11E-05 | 0,00079903 |
| 149076_at | ZNF362 | 0,63363069 | 9,10658657 | 6,51393156 | 1,15E-05 | 0,0008201 |
| 30968_at | STOML2 | -0,70800087 | 9,78288345 | -6,50267396 | 1,17E-05 | 0,0008328 |
| 84247_at | LDOC1L | 0,49710534 | 8,58992601 | 6,4999085 | 1,17E-05 | 0,00083363 |
| 284106_at | CISD3 | -0,44283915 | 7,70182045 | -6,49621639 | 1,18E-05 | 0,00083579 |
| 10856_at | RUVBL2 | -0,70857415 | 10,0143895 | -6,48805779 | 1,20E-05 | 0,00084434 |
| 776_at | CACNA1D | 0,7664977 | 5,43707736 | 6,46796059 | 1,24E-05 | 0,00087049 |
| 57630_at | SH3RF1 | 0,3712902 | 7,53009614 | 6,46344749 | 1,25E-05 | 0,00087134 |
| 55876_at | GSDMB | 1,00694185 | 6,90195502 | 6,46233316 | 1,25E-05 | 0,00087134 |
| 51185_at | CRBN | 0,5733321 | 8,3966831 | 6,46098651 | 1,25E-05 | 0,00087134 |
| 123263_at | MTFMT | -0,54333164 | 7,85482197 | -6,45707103 | 1,26E-05 | 0,00087401 |
| 3751_at | KCND2 | 1,76576672 | 5,67335829 | 6,45099026 | 1,27E-05 | 0,00087994 |
| 26520_at | TIMM9 | -0,79228058 | 7,58070104 | -6,44402383 | 1,29E-05 | 0,0008827 |
| 54888_at | NSUN2 | -0,52040719 | 8,67749741 | -6,442517 | 1,29E-05 | 0,0008827 |
| 5709_at | PSMD3 | -0,53391893 | 10,0194741 | -6,44232325 | 1,29E-05 | 0,0008827 |
| 8821_at | INPP4B | -0,90680949 | 7,65850018 | -6,44079937 | 1,30E-05 | 0,0008827 |
| 10575_at | CCT4 | -0,65471645 | 11,1463618 | -6,43747102 | 1,30E-05 | 0,00088459 |
| 55793_at | FAM63A | 0,54118599 | 7,01510558 | 6,42192076 | 1,34E-05 | 0,00090519 |
| 54828_at | BCAS3 | 0,54027966 | 6,6506811 | 6,41793154 | 1,35E-05 | 0,00090819 |
| 79767_at | ELMO3 | 0,42547746 | 7,18711085 | 6,41228972 | 1,36E-05 | 0,00090943 |
| 9550_at | ATP6V1G1 | 0,42942328 | 8,074655 | 6,41029769 | 1,37E-05 | 0,00090943 |
| 3396_at | ICT1 | -0,45841462 | 9,10028831 | -6,4093864 | 1,37E-05 | 0,00090943 |
| 7804_at | LRP8 | -2,00071926 | 7,91057248 | -6,4089664 | 1,37E-05 | 0,00090943 |
| 90417_at | KNSTRN | -1,11686724 | 8,13986377 | -6,40535925 | 1,38E-05 | 0,00090943 |
| 6790_at | AURKA | -1,26435404 | 9,43154207 | -6,40385763 | 1,38E-05 | 0,00090943 |
| 89846_at | FGD3 | 1,17357599 | 8,13974641 | 6,4020911 | 1,39E-05 | 0,00090943 |
| 84250_at | SLF1 | -0,69901691 | 5,23064049 | -6,39960135 | 1,39E-05 | 0,00090943 |

| | | | | | | |
|---|---|---|---|---|---|---|
| 55559_at | HAUS7 | -0,8589843 | 6,81760012 | -6,39900219 | 1,39E-05 | 0,00090943 |
| 152006_at | RNF38 | 0,75210127 | 8,46733453 | 6,39221512 | 1,41E-05 | 0,00091695 |
| 4522_at | MTHFD1 | -0,97236005 | 10,1229174 | -6,38293144 | 1,43E-05 | 0,0009285 |
| 23236_at | PLCB1 | 0,95964328 | 6,66540761 | 6,37476359 | 1,45E-05 | 0,00093842 |
| 6838_at | SURF6 | -0,38215844 | 7,52814455 | -6,37140605 | 1,46E-05 | 0,00094068 |
| 374395_at | TMEM179B | 0,42541196 | 10,2458404 | 6,36492693 | 1,48E-05 | 0,00094684 |
| 55726_at | ASUN | -0,49889076 | 9,33531811 | -6,36055672 | 1,49E-05 | 0,00094684 |
| 130557_at | ZNF513 | 0,49768334 | 7,60835646 | 6,36007845 | 1,49E-05 | 0,00094684 |
| 687_at | KLF9 | 0,89368592 | 6,4321989 | 6,35985362 | 1,49E-05 | 0,00094684 |
| 5707_at | PSMD1 | -0,62867402 | 9,82433333 | -6,35600063 | 1,50E-05 | 0,00094998 |
| 3068_at | HDGF | -0,42603236 | 10,5794453 | -6,34556533 | 1,53E-05 | 0,00096141 |
| 3146_at | HMGB1 | -0,75603107 | 8,48541839 | -6,34521802 | 1,53E-05 | 0,00096141 |
| 253461_at | ZBTB38 | 0,46456251 | 6,33238211 | 6,33227213 | 1,56E-05 | 0,00097984 |
| 266722_at | HS6ST3 | 0,88767992 | 7,38567237 | 6,32361354 | 1,59E-05 | 0,00099131 |
| 4758_at | NEU1 | 0,68055406 | 8,85329906 | 6,30664815 | 1,63E-05 | 0,00101739 |
| 10465_at | PPIH | -0,76930011 | 7,72341131 | -6,30238294 | 1,64E-05 | 0,00102159 |
| 9710_at | KIAA0355 | 0,63834808 | 6,96605845 | 6,29581393 | 1,66E-05 | 0,00102598 |
| 375790_at | AGRN | 0,45249651 | 8,69948018 | 6,29528567 | 1,66E-05 | 0,00102598 |
| 163786_at | SASS6 | -0,93386564 | 7,67205369 | -6,29431206 | 1,67E-05 | 0,00102598 |
| 7372_at | UMPS | -0,44757871 | 7,28993475 | -6,28491407 | 1,69E-05 | 0,00103943 |
| 3338_at | DNAJC4 | 0,62259454 | 7,5798571 | 6,27898921 | 1,71E-05 | 0,00104598 |
| 708_at | C1QBP | -0,90548622 | 10,6053722 | -6,27759886 | 1,72E-05 | 0,00104598 |
| 79902_at | NUP85 | -0,71417339 | 9,12899307 | -6,26993841 | 1,74E-05 | 0,00105657 |
| 60487_at | TRMT11 | -0,63982804 | 7,38345167 | -6,26598372 | 1,75E-05 | 0,00106048 |
| 55388_at | MCM10 | -1,56602779 | 6,87413032 | -6,26200883 | 1,76E-05 | 0,00106444 |
| 678_at | ZFP36L2 | 0,73203557 | 9,92847089 | 6,25242079 | 1,79E-05 | 0,00107636 |
| 23246_at | BOP1 | -1,16202176 | 8,13743812 | -6,25196006 | 1,79E-05 | 0,00107636 |
| 23212_at | RRS1 | -0,92632033 | 9,52412999 | -6,24993403 | 1,80E-05 | 0,00107679 |
| 10576_at | CCT2 | -0,65065132 | 10,766327 | -6,2477486 | 1,81E-05 | 0,00107752 |
| 94241_at | TP53INP1 | 2,06314121 | 8,18241949 | 6,24097073 | 1,83E-05 | 0,00108603 |
| 6016_at | RIT1 | 0,72718474 | 8,39431357 | 6,2396393 | 1,83E-05 | 0,00108603 |
| 51042_at | ZNF593 | -0,63539757 | 8,62016281 | -6,2336354 | 1,85E-05 | 0,00109403 |
| 147798_at | TMC4 | 1,59124676 | 8,46865965 | 6,23125752 | 1,86E-05 | 0,00109518 |
| 84154_at | RPF2 | -0,90741199 | 8,8469401 | -6,22600295 | 1,88E-05 | 0,00109975 |
| 5688_at | PSMA7 | -0,44588509 | 10,8504836 | -6,22534077 | 1,88E-05 | 0,00109975 |
| 100507567_a | RAB11B-AS1 | 0,96845298 | 6,50748603 | 6,22237412 | 1,89E-05 | 0,00110208 |
| 100499467_a | LINC00673 | 0,49853544 | 8,32429585 | 6,21683529 | 1,91E-05 | 0,00110936 |
| 112483_at | SAT2 | 0,48137195 | 7,92654051 | 6,20884936 | 1,93E-05 | 0,00112146 |
| 126382_at | NR2C2AP | -0,92195485 | 7,55342053 | -6,20626073 | 1,94E-05 | 0,00112313 |
| 10799_at | RPP40 | -0,97589497 | 7,8825172 | -6,20293962 | 1,95E-05 | 0,00112624 |
| 51002_at | TPRKB | -0,46357051 | 8,52142155 | -6,18890925 | 2,00E-05 | 0,0011506 |
| 285943_at | HOXA-AS2 | 0,52539462 | 6,5724166 | 6,18341512 | 2,02E-05 | 0,00115513 |
| 7586_at | ZKSCAN1 | 0,68549948 | 8,41126739 | 6,18323273 | 2,02E-05 | 0,00115513 |
| 5437_at | POLR2H | -0,41978707 | 9,96571658 | -6,16999954 | 2,07E-05 | 0,00117855 |
| 4189_at | DNAJB9 | 0,62266891 | 7,11822743 | 6,16685717 | 2,08E-05 | 0,00118153 |
| 117246_at | FTSJ3 | -0,44615613 | 8,57038905 | -6,1503304 | 2,14E-05 | 0,00121249 |
| 84102_at | SLC41A2 | 1,0779876 | 6,06821296 | 6,14247881 | 2,17E-05 | 0,0012244 |
| 123228_at | SENP8 | 0,41880043 | 6,11360171 | 6,14136929 | 2,17E-05 | 0,0012244 |
| 80218_at | NAA50 | -0,48218124 | 9,75185531 | -6,12768803 | 2,23E-05 | 0,00124694 |
| 58480_at | RHOU | 1,11212617 | 7,68269382 | 6,12758811 | 2,23E-05 | 0,00124694 |

| | | | | | | |
|---|---|---|---|---|---|---|
| 400916_at | CHCHD10 | -0,54718565 | 9,70001693 | -6,11902667 | 2,26E-05 | 0,0012598 |
| 84818_at | IL17RC | 0,4431662 | 7,0256106 | 6,11840687 | 2,26E-05 | 0,0012598 |
| 3931_at | LCAT | 0,43964618 | 5,85698154 | 6,10741207 | 2,31E-05 | 0,00128057 |
| 51434_at | ANAPC7 | -0,5407235 | 8,18964335 | -6,09943563 | 2,34E-05 | 0,00129204 |
| 5690_at | PSMB2 | -0,5444156 | 8,50718069 | -6,09903873 | 2,34E-05 | 0,00129204 |
| 79080_at | CCDC86 | -1,21156632 | 8,58054053 | -6,09538907 | 2,36E-05 | 0,00129426 |
| 8260_at | NAA10 | -0,50544439 | 8,85583876 | -6,09426526 | 2,36E-05 | 0,00129426 |
| 51537_at | MTFP1 | -1,08902916 | 7,82085802 | -6,09317272 | 2,37E-05 | 0,00129426 |
| 9056_at | SLC7A7 | 0,8058007 | 5,68190445 | 6,08589336 | 2,40E-05 | 0,00130717 |
| 6655_at | SOS2 | 0,6083523 | 7,68966991 | 6,08395492 | 2,40E-05 | 0,00130792 |
| 10166_at | SLC25A15 | -0,69966379 | 6,74671701 | -6,0775542 | 2,43E-05 | 0,0013158 |
| 10607_at | TBL3 | -0,44446912 | 7,93748915 | -6,07632048 | 2,44E-05 | 0,0013158 |
| 64222_at | TOR3A | -0,58266377 | 8,66559445 | -6,07507534 | 2,44E-05 | 0,0013158 |
| 27000_at | DNAJC2 | -0,82427601 | 8,78691336 | -6,07412622 | 2,45E-05 | 0,0013158 |
| 55337_at | C19orf66 | 0,84174824 | 7,76903364 | 6,06627963 | 2,48E-05 | 0,00132709 |
| 7915_at | ALDH5A1 | 0,63758404 | 8,14824925 | 6,0660887 | 2,48E-05 | 0,00132709 |
| 10023_at | FRAT1 | 0,58012643 | 5,55601713 | 6,06328664 | 2,49E-05 | 0,00132786 |
| 6832_at | SUPV3L1 | -0,43819143 | 7,91721892 | -6,06214466 | 2,50E-05 | 0,00132786 |
| 23670_at | TMEM2 | 0,9259366 | 8,3881057 | 6,06103084 | 2,50E-05 | 0,00132786 |
| 4094_at | MAF | 0,67007947 | 4,49446156 | 6,05496965 | 2,53E-05 | 0,00133643 |
| 727910_at | TLCD2 | 0,45384775 | 6,30213362 | 6,05424764 | 2,53E-05 | 0,00133643 |
| 131474_at | CHCHD4 | -0,77869301 | 7,64547154 | -6,05103185 | 2,55E-05 | 0,00133717 |
| 23016_at | EXOSC7 | -0,5085722 | 7,14144444 | -6,05082144 | 2,55E-05 | 0,00133717 |
| 55572_at | FOXRED1 | -0,45760597 | 8,04749256 | -6,04755052 | 2,56E-05 | 0,00133767 |
| 57169_at | ZNFX1 | 0,82616849 | 9,26113852 | 6,04752107 | 2,56E-05 | 0,00133767 |
| 1802_at | DPH2 | -0,51238478 | 7,71659798 | -6,04470706 | 2,58E-05 | 0,00134019 |
| 58191_at | CXCL16 | 0,52183337 | 7,92451672 | 6,04337663 | 2,58E-05 | 0,00134019 |
| 64328_at | XPO4 | -0,51932357 | 7,31499739 | -6,03787737 | 2,61E-05 | 0,00134959 |
| 10539_at | GLRX3 | -0,43614272 | 9,67697066 | -6,02555859 | 2,66E-05 | 0,00137532 |
| 26960_at | NBEA | 0,80220215 | 6,41431735 | 6,02283996 | 2,68E-05 | 0,00137532 |
| 8019_at | BRD3 | 0,51706586 | 8,51723138 | 6,02259268 | 2,68E-05 | 0,00137532 |
| 4552_at | MTRR | -0,65620536 | 8,4209058 | -6,0168575 | 2,70E-05 | 0,0013856 |
| 23053_at | ZSWIM8 | 0,57318759 | 6,20851771 | 6,01178161 | 2,73E-05 | 0,00139107 |
| 6573_at | SLC19A1 | -0,75539747 | 6,70454475 | -6,0116168 | 2,73E-05 | 0,00139107 |
| 84172_at | POLR1B | -0,767853 | 7,42076068 | -6,00475239 | 2,76E-05 | 0,0014043 |
| 11119_at | BTN3A1 | 0,85612544 | 7,18479471 | 6,00090677 | 2,78E-05 | 0,00140694 |
| 56902_at | PNO1 | -0,84992756 | 8,4855541 | -6,00070667 | 2,78E-05 | 0,00140694 |
| 90427_at | BMF | 1,18797991 | 6,56696274 | 5,99354752 | 2,82E-05 | 0,00142111 |
| 10291_at | SF3A1 | -0,29641111 | 8,99952298 | -5,9840822 | 2,87E-05 | 0,00144131 |
| 64216_at | TFB2M | -0,48011877 | 8,37277019 | -5,97473667 | 2,91E-05 | 0,00146151 |
| 54206_at | ERRFI1 | -0,63097425 | 8,90246184 | -5,96483542 | 2,96E-05 | 0,00148348 |
| 1736_at | DKC1 | -0,99512061 | 9,18724937 | -5,9633377 | 2,97E-05 | 0,00148356 |
| 9319_at | TRIP13 | -1,64603328 | 8,3713552 | -5,96042828 | 2,99E-05 | 0,00148737 |
| 23210_at | JMJD6 | -0,35816225 | 6,90090556 | -5,95751971 | 3,00E-05 | 0,0014912 |
| 4610_at | MYCL | 0,63409614 | 6,54086217 | 5,95605171 | 3,01E-05 | 0,00149124 |
| 79875_at | THSD4 | -0,5483298 | 6,12276353 | -5,94724168 | 3,06E-05 | 0,00151014 |
| 51377_at | UCHL5 | -0,79319531 | 7,12270558 | -5,9460505 | 3,07E-05 | 0,00151014 |
| 388796_at | SNHG17 | -1,05622084 | 9,37019355 | -5,94181968 | 3,09E-05 | 0,00151763 |
| 84861_at | KLHL22 | 0,54024892 | 6,81152063 | 5,93456975 | 3,13E-05 | 0,00153337 |
| 55646_at | LYAR | -1,59605612 | 8,31366625 | -5,93139073 | 3,15E-05 | 0,00153813 |

| | | | | | | |
|---|---|---|---|---|---|---|
| 641_at | BLM | -1,19747453 | 6,99362233 | -5,92906264 | 3,16E-05 | 0,00154059 |
| 25888_at | ZNF473 | -0,38443997 | 6,59882411 | -5,92523901 | 3,18E-05 | 0,00154716 |
| 29914_at | UBIAD1 | -0,42547422 | 7,05313242 | -5,92136753 | 3,20E-05 | 0,00155392 |
| 81892_at | SLIRP | -0,7528098 | 11,5385627 | -5,91916389 | 3,22E-05 | 0,00155609 |
| 1263_at | PLK3 | -0,48609519 | 6,12730249 | -5,91161134 | 3,26E-05 | 0,00157317 |
| 5025_at | P2RX4 | 0,58832983 | 7,65409801 | 5,91017964 | 3,27E-05 | 0,00157323 |
| 694_at | BTG1 | 0,75179573 | 9,28420331 | 5,90569574 | 3,29E-05 | 0,00158069 |
| 11073_at | TOPBP1 | -0,75136557 | 9,19323653 | -5,90471654 | 3,30E-05 | 0,00158069 |
| 160364_at | CLEC12A | 1,50543613 | 4,95320246 | 5,90326766 | 3,31E-05 | 0,00158083 |
| 10989_at | IMMT | -0,33008047 | 9,93935291 | -5,90128702 | 3,32E-05 | 0,00158249 |
| 8879_at | SGPL1 | 0,70099152 | 9,10543351 | 5,89704696 | 3,34E-05 | 0,00159053 |
| 6427_at | SRSF2 | -0,42732087 | 10,6731157 | -5,89041965 | 3,38E-05 | 0,00160482 |
| 101927746_a | LOC10192774 | -0,83007089 | 4,19367353 | -5,88925516 | 3,39E-05 | 0,00160482 |
| 79023_at | NUP37 | -0,69754785 | 9,06533812 | -5,88630795 | 3,41E-05 | 0,0016061 |
| 55210_at | ATAD3A | -0,71197677 | 8,40390119 | -5,88605643 | 3,41E-05 | 0,0016061 |
| 23112_at | TNRC6B | 0,47759757 | 7,50610035 | 5,88219394 | 3,43E-05 | 0,00161324 |
| 2618_at | GART | -0,8485233 | 8,52403111 | -5,87794813 | 3,46E-05 | 0,00162154 |
| 1847_at | DUSP5 | -0,87291859 | 8,13700622 | -5,87399214 | 3,48E-05 | 0,00162905 |
| 11152_at | WDR45 | 0,57446492 | 7,73427641 | 5,87150947 | 3,50E-05 | 0,00163232 |
| 64743_at | WDR13 | 0,43529217 | 8,4838056 | 5,86902687 | 3,52E-05 | 0,0016356 |
| 8705_at | B3GALT4 | 0,59021711 | 7,62088493 | 5,86623429 | 3,53E-05 | 0,00163981 |
| 6877_at | TAF5 | -0,68830333 | 6,96536393 | -5,85975261 | 3,57E-05 | 0,0016549 |
| 56339_at | METTL3 | -0,75432101 | 8,75341864 | -5,84272088 | 3,68E-05 | 0,00170022 |
| 23649_at | POLA2 | -0,66510731 | 6,68980609 | -5,84193919 | 3,69E-05 | 0,00170022 |
| 1951_at | CELSR3 | 0,47232739 | 7,20044175 | 5,83892738 | 3,71E-05 | 0,00170532 |
| 4077_at | NBR1 | 0,7186472 | 8,76679821 | 5,83402095 | 3,74E-05 | 0,00171625 |
| 134637_at | ADAT2 | -0,5719745 | 5,42710825 | -5,83187754 | 3,76E-05 | 0,00171875 |
| 283450_at | HECTD4 | 0,46608924 | 7,36087608 | 5,8241024 | 3,81E-05 | 0,00173753 |
| 6773_at | STAT2 | 1,00125199 | 7,77464867 | 5,82234841 | 3,82E-05 | 0,00173753 |
| 339448_at | C1orf174 | -0,36974215 | 8,45596191 | -5,82183453 | 3,82E-05 | 0,00173753 |
| 23179_at | RGL1 | 0,94561758 | 6,81522428 | 5,8178781 | 3,85E-05 | 0,00174049 |
| 59084_at | ENPP5 | 1,55387466 | 4,96022176 | 5,81772814 | 3,85E-05 | 0,00174049 |
| 60386_at | SLC25A19 | -0,91506741 | 7,08781434 | -5,81694734 | 3,86E-05 | 0,00174049 |
| 79629_at | OCEL1 | 0,58831151 | 7,80365954 | 5,80913859 | 3,91E-05 | 0,00176088 |
| 57673_at | BEND3 | -0,63394361 | 6,73695082 | -5,8078034 | 3,92E-05 | 0,00176098 |
| 23753_at | SDF2L1 | -0,69319109 | 8,00448162 | -5,80453609 | 3,94E-05 | 0,00176439 |
| 2166_at | FAAH | 0,37929296 | 6,91682051 | 5,8041282 | 3,95E-05 | 0,00176439 |
| 10171_at | RCL1 | -0,44759222 | 7,39892169 | -5,80055328 | 3,97E-05 | 0,00177163 |
| 55030_at | FBXO34 | 0,41752489 | 9,10509131 | 5,79323276 | 4,03E-05 | 0,00177812 |
| 7318_at | UBA7 | 1,12124705 | 8,39247456 | 5,79309 | 4,03E-05 | 0,00177812 |
| 221150_at | SKA3 | -1,33668623 | 7,86724137 | -5,79231817 | 4,03E-05 | 0,00177812 |
| 7039_at | TGFA | -0,53123292 | 6,77700945 | -5,7916031 | 4,04E-05 | 0,00177812 |
| 10915_at | TCERG1 | -0,5084009 | 9,28054749 | -5,79099707 | 4,04E-05 | 0,00177812 |
| 254827_at | NAALADL2 | 1,75195578 | 5,11431102 | 5,79082175 | 4,04E-05 | 0,00177812 |
| 5324_at | PLAG1 | 0,96938911 | 6,76520578 | 5,78633919 | 4,08E-05 | 0,00178623 |
| 84515_at | MCM8 | -1,114563 | 6,79991746 | -5,78574808 | 4,08E-05 | 0,00178623 |
| 26224_at | FBXL3 | 0,39213447 | 8,99854705 | 5,77464398 | 4,16E-05 | 0,00181807 |
| 3182_at | HNRNPAB | -0,68565608 | 11,9150871 | -5,76340479 | 4,25E-05 | 0,00185098 |
| 79089_at | TMUB2 | 0,30262263 | 7,66710749 | 5,76092567 | 4,27E-05 | 0,00185505 |
| 2706_at | GJB2 | -1,2001084 | 8,24396185 | -5,75680506 | 4,30E-05 | 0,00186464 |

| | | | | | | |
|---|---|---|---|---|---|---|
| 9656_at | MDC1 | -0,49188339 | 6,96952132 | -5,75429643 | 4,32E-05 | 0,00186886 |
| 1434_at | CSE1L | -0,80791783 | 10,7013205 | -5,75036138 | 4,35E-05 | 0,00187793 |
| 54874_at | FNBP1L | 0,75551789 | 8,73973787 | 5,74089866 | 4,42E-05 | 0,00190592 |
| 5805_at | PTS | -0,5606788 | 9,08229344 | -5,73638131 | 4,46E-05 | 0,0019135 |
| 5496_at | PPM1G | -0,63437262 | 9,37531136 | -5,73621763 | 4,46E-05 | 0,0019135 |
| 9277_at | WDR46 | -0,6469745 | 7,83615271 | -5,73172569 | 4,50E-05 | 0,00192109 |
| 3420_at | IDH3B | -0,29246435 | 9,50827621 | -5,73155473 | 4,50E-05 | 0,00192109 |
| 79735_at | TBC1D17 | 0,43677576 | 7,02839411 | 5,72795591 | 4,53E-05 | 0,00192931 |
| 11257_at | TP53TG1 | 0,52580763 | 8,83798606 | 5,7207858 | 4,59E-05 | 0,00194414 |
| 4070_at | TACSTD2 | 0,38147201 | 12,6401183 | 5,72048086 | 4,59E-05 | 0,00194414 |
| 51_at | ACOX1 | 0,61243145 | 9,10723358 | 5,72004287 | 4,59E-05 | 0,00194414 |
| 90381_at | TICRR | -0,98229557 | 6,70990804 | -5,71477935 | 4,64E-05 | 0,00195839 |
| 2648_at | KAT2A | -0,44035689 | 8,45431662 | -5,70943217 | 4,68E-05 | 0,00197305 |
| 516_at | ATP5G1 | -0,8643944 | 10,2927271 | -5,70498984 | 4,72E-05 | 0,00198345 |
| 199692_at | ZNF627 | 0,5971874 | 7,65604901 | 5,70410498 | 4,73E-05 | 0,00198345 |
| 51111_at | SUV420H1 | 0,48845302 | 7,72924087 | 5,70209727 | 4,74E-05 | 0,00198632 |
| 10460_at | TACC3 | -1,10598023 | 8,85302331 | -5,69769064 | 4,78E-05 | 0,00199393 |
| 64112_at | MOAP1 | 0,35812985 | 7,93253808 | 5,6975811 | 4,78E-05 | 0,00199393 |
| 8892_at | EIF2B2 | -0,46957656 | 8,86892291 | -5,69269155 | 4,82E-05 | 0,00200728 |
| 8882_at | ZPR1 | -0,5560539 | 7,73470349 | -5,6899289 | 4,85E-05 | 0,00201299 |
| 1389_at | CREBL2 | 0,48159279 | 8,25987865 | 5,68240581 | 4,92E-05 | 0,00203617 |
| 10946_at | SF3A3 | -0,63566166 | 8,98248524 | -5,67977062 | 4,94E-05 | 0,00204151 |
| 81929_at | SEH1L | -0,80696052 | 8,64604044 | -5,66147624 | 5,11E-05 | 0,00210201 |
| 64858_at | DCLRE1B | -0,89714418 | 6,86234872 | -5,66127692 | 5,11E-05 | 0,00210201 |
| 79828_at | METTL8 | -0,47161026 | 6,08697373 | -5,65884048 | 5,13E-05 | 0,00210612 |
| 1287_at | COL4A5 | 0,55202816 | 6,9222156 | 5,65784662 | 5,14E-05 | 0,00210612 |
| 79830_at | ZMYM1 | -0,45528692 | 6,77508829 | -5,64685329 | 5,24E-05 | 0,00214284 |
| 7134_at | TNNC1 | 0,70656738 | 6,45431177 | 5,64597269 | 5,25E-05 | 0,00214284 |
| 22903_at | BTBD3 | 0,59749559 | 9,26139109 | 5,64381014 | 5,27E-05 | 0,00214672 |
| 79733_at | E2F8 | -0,9848413 | 7,87290687 | -5,64110276 | 5,30E-05 | 0,00215274 |
| 4957_at | ODF2 | -0,53511213 | 7,4862863 | -5,63548644 | 5,35E-05 | 0,00217022 |
| 11069_at | RAPGEF4 | 0,99322194 | 4,26080345 | 5,63054659 | 5,40E-05 | 0,00218517 |
| 80832_at | APOL4 | 1,06301951 | 6,88666351 | 5,62141596 | 5,49E-05 | 0,00221706 |
| 124446_at | TMEM219 | 0,64521647 | 8,3421102 | 5,6175259 | 5,53E-05 | 0,00222812 |
| 51633_at | OTUD6B | -0,74916262 | 8,67406109 | -5,61555313 | 5,55E-05 | 0,00223146 |
| 64787_at | EPS8L2 | 0,55515829 | 8,26919183 | 5,60787679 | 5,63E-05 | 0,00225812 |
| 23426_at | GRIP1 | 0,39699247 | 4,6297445 | 5,60570576 | 5,65E-05 | 0,00226156 |
| 27250_at | PDCD4 | 1,95212692 | 8,14401868 | 5,60475432 | 5,66E-05 | 0,00226156 |
| 23560_at | GTPBP4 | -0,74098773 | 8,0885461 | -5,60204016 | 5,69E-05 | 0,00226403 |
| 55238_at | SLC38A7 | 0,34860916 | 7,08530342 | 5,6018785 | 5,69E-05 | 0,00226403 |
| 6196_at | RPS6KA2 | 0,53253531 | 6,48933465 | 5,59360936 | 5,77E-05 | 0,00229364 |
| 64794_at | DDX31 | -0,35917454 | 6,59726392 | -5,5891277 | 5,82E-05 | 0,00230473 |
| 5176_at | SERPINF1 | 0,68969079 | 6,53717267 | 5,58870122 | 5,83E-05 | 0,00230473 |
| 27339_at | PRPF19 | -0,49052732 | 9,886838 | -5,56762789 | 6,05E-05 | 0,0023901 |
| 55329_at | MNS1 | -1,31287961 | 6,17121267 | -5,56251534 | 6,11E-05 | 0,00240757 |
| 440145_at | MZT1 | -0,8544743 | 8,80075568 | -5,55721706 | 6,17E-05 | 0,00242601 |
| 81624_at | DIAPH3 | -0,5367678 | 6,41256464 | -5,55481755 | 6,20E-05 | 0,0024317 |
| 292_at | SLC25A5 | -0,61531903 | 11,978759 | -5,5535926 | 6,21E-05 | 0,0024322 |
| 3590_at | IL11RA | 0,52032023 | 6,65442873 | 5,55193821 | 6,23E-05 | 0,00243461 |
| 55014_at | STX17 | 0,58586729 | 7,4047799 | 5,5465138 | 6,29E-05 | 0,00244798 |

| | | | | | | |
|---|---|---|---|---|---|---|
| 6041_at | RNASEL | 0,68272092 | 7,38017645 | 5,5461273 | 6,30E-05 | 0,00244798 |
| 10471_at | PFDN6 | -0,42404858 | 8,94549513 | -5,54562147 | 6,30E-05 | 0,00244798 |
| 51660_at | MPC1 | -0,61973598 | 8,32228365 | -5,54077634 | 6,36E-05 | 0,00246479 |
| 51134_at | CEP83 | -0,65936638 | 5,96908302 | -5,53947643 | 6,37E-05 | 0,00246569 |
| 3735_at | KARS | -0,30358062 | 11,2113594 | -5,53548868 | 6,42E-05 | 0,00247876 |
| 79660_at | PPP1R3B | 1,51706341 | 6,87842749 | 5,53434773 | 6,43E-05 | 0,00247897 |
| 375341_at | C3orf62 | 0,64517048 | 6,04775953 | 5,53135661 | 6,47E-05 | 0,00248315 |
| 100506314_a | LOC10050631 | 0,48938053 | 5,77843702 | 5,53124378 | 6,47E-05 | 0,00248315 |
| 23471_at | TRAM1 | 0,60875718 | 11,1301972 | 5,52071691 | 6,60E-05 | 0,00252643 |
| 55957_at | LIN37 | 0,37098546 | 7,9585982 | 5,51884494 | 6,62E-05 | 0,00253007 |
| 159090_at | FAM122B | -0,53612894 | 8,58808346 | -5,51743431 | 6,64E-05 | 0,0025316 |
| 29078_at | NDUFAF4 | -0,85326752 | 7,74665454 | -5,50750612 | 6,76E-05 | 0,00256826 |
| 59272_at | ACE2 | 1,623036 | 5,71632323 | 5,50742798 | 6,76E-05 | 0,00256826 |
| 11047_at | ADRM1 | -0,56870505 | 10,523544 | -5,50278308 | 6,82E-05 | 0,00258513 |
| 100130449_a | PP14571 | 1,28214767 | 6,83725692 | 5,50168828 | 6,83E-05 | 0,00258523 |
| 121512_at | FGD4 | 0,38471213 | 5,53706695 | 5,49950522 | 6,86E-05 | 0,00259051 |
| 79075_at | DSCC1 | -1,58781435 | 7,19175711 | -5,49543295 | 6,91E-05 | 0,00260483 |
| 51073_at | MRPL4 | -0,60676136 | 8,59055308 | -5,49072261 | 6,97E-05 | 0,0026223 |
| 29920_at | PYCR2 | -0,55935203 | 8,77803646 | -5,48933597 | 6,99E-05 | 0,00262386 |
| 54928_at | IMPAD1 | -0,4922372 | 8,6433904 | -5,48687269 | 7,02E-05 | 0,00263062 |
| 25926_at | NOL11 | -0,67377767 | 9,06979281 | -5,48504621 | 7,04E-05 | 0,00263433 |
| 6302_at | TSPAN31 | 0,72091705 | 9,42027089 | 5,48134827 | 7,09E-05 | 0,00264714 |
| 54981_at | NMRK1 | 0,8353323 | 7,45645442 | 5,47980931 | 7,11E-05 | 0,0026495 |
| 23107_at | MRPS27 | -0,52569439 | 8,85354677 | -5,47647329 | 7,15E-05 | 0,00266063 |
| 84326_at | C16orf13 | -0,48006508 | 8,95199272 | -5,46803144 | 7,27E-05 | 0,00269702 |
| 3954_at | LETM1 | -0,38246296 | 8,2833025 | -5,4662868 | 7,29E-05 | 0,00270048 |
| 79017_at | GGCT | -0,64045372 | 10,3392197 | -5,46348271 | 7,33E-05 | 0,00270923 |
| 154743_at | C7orf60 | 0,79033518 | 7,71153458 | 5,4500166 | 7,51E-05 | 0,00277186 |
| 84245_at | MRI1 | -0,41888869 | 8,21126779 | -5,44459838 | 7,59E-05 | 0,00279429 |
| 100507459_a | NRSN2-AS1 | 0,9518328 | 5,14939251 | 5,44355261 | 7,60E-05 | 0,00279434 |
| 9267_at | CYTH1 | 0,33492224 | 6,89982034 | 5,44157747 | 7,63E-05 | 0,00279918 |
| 81848_at | SPRY4 | -0,82510656 | 5,27092569 | -5,43829756 | 7,68E-05 | 0,00280523 |
| 80777_at | CYB5B | -0,71219786 | 8,57694511 | -5,43813282 | 7,68E-05 | 0,00280523 |
| 23401_at | FRAT2 | 0,59093491 | 8,13703585 | 5,43654748 | 7,70E-05 | 0,00280523 |
| 440957_at | SMIM4 | -0,58953013 | 6,57055368 | -5,43628306 | 7,70E-05 | 0,00280523 |
| 23417_at | MLYCD | 0,43803877 | 6,64827536 | 5,43424779 | 7,73E-05 | 0,00280793 |
| 153222_at | CREBRF | 1,31432395 | 5,68760804 | 5,43371356 | 7,74E-05 | 0,00280793 |
| 83636_at | C19orf12 | 0,38724466 | 7,33163662 | 5,42743866 | 7,83E-05 | 0,00283221 |
| 81887_at | LAS1L | -0,54628371 | 7,43198004 | -5,42484523 | 7,87E-05 | 0,00283221 |
| 55119_at | PRPF38B | -0,4973807 | 8,16840512 | -5,42444308 | 7,87E-05 | 0,00283221 |
| 26155_at | NOC2L | -0,62437302 | 8,15372211 | -5,42298435 | 7,89E-05 | 0,00283221 |
| 84844_at | PHF5A | -0,56168445 | 8,71804442 | -5,42268786 | 7,90E-05 | 0,00283221 |
| 8669_at | EIF3J | -0,48995047 | 8,23362317 | -5,42219411 | 7,91E-05 | 0,00283221 |
| 28982_at | FLVCR1 | -0,84160425 | 7,87147604 | -5,42193988 | 7,91E-05 | 0,00283221 |
| 55718_at | POLR3E | -0,40804222 | 8,06355244 | -5,41959062 | 7,94E-05 | 0,0028366 |
| 57185_at | NIPAL3 | 0,43769641 | 8,15833325 | 5,41787284 | 7,97E-05 | 0,0028366 |
| 81875_at | ISG20L2 | -0,44729867 | 9,25539242 | -5,41767718 | 7,97E-05 | 0,0028366 |
| 4173_at | MCM4 | -0,96021988 | 9,09508977 | -5,41672934 | 7,99E-05 | 0,0028366 |
| 51373_at | MRPS17 | -0,65275514 | 8,82748571 | -5,41608488 | 8,00E-05 | 0,0028366 |
| 51605_at | TRMT6 | -0,72288949 | 8,64452514 | -5,41236287 | 8,05E-05 | 0,0028509 |

| | | | | | | |
|---|---|---|---|---|---|---|
| 159091_at | FAM122C | 0,32396028 | 4,3919431 | 5,40985745 | 8,09E-05 | 0,00285364 |
| 100131096_a | TNRC6C-AS1 | 0,61436936 | 6,18422648 | 5,4092168 | 8,10E-05 | 0,00285364 |
| 11276_at | SYNRG | 0,59912931 | 7,41167391 | 5,4079159 | 8,12E-05 | 0,00285364 |
| 348180_at | CTU2 | -0,63327213 | 6,7471621 | -5,40703354 | 8,13E-05 | 0,00285364 |
| 54908_at | SPDL1 | -0,93951334 | 7,07982935 | -5,40688535 | 8,13E-05 | 0,00285364 |
| 23097_at | CDK19 | 0,46742225 | 7,38568608 | 5,40450136 | 8,17E-05 | 0,00285691 |
| 55144_at | LRRC8D | -0,41686103 | 9,37677533 | -5,40429476 | 8,17E-05 | 0,00285691 |
| 55239_at | OGFOD1 | -0,62527954 | 8,4110174 | -5,39503027 | 8,31E-05 | 0,00290096 |
| 100874032_a | PRRT3-AS1 | -0,98474412 | 6,16904798 | -5,39251157 | 8,35E-05 | 0,00290923 |
| 10520_at | ZNF211 | 0,39425365 | 7,41194126 | 5,38507187 | 8,47E-05 | 0,00294419 |
| 57590_at | WDFY1 | 0,3878656 | 8,79234874 | 5,38357961 | 8,49E-05 | 0,00294701 |
| 26517_at | TIMM13 | -0,56186869 | 8,61272428 | -5,38214714 | 8,51E-05 | 0,00294951 |
| 56993_at | TOMM22 | -0,52096411 | 9,0673504 | -5,37626591 | 8,61E-05 | 0,00297642 |
| 8720_at | MBTPS1 | 0,34421526 | 8,52610524 | 5,37236682 | 8,67E-05 | 0,0029926 |
| 11331_at | PHB2 | -0,36255059 | 11,3647379 | -5,37135211 | 8,69E-05 | 0,00299287 |
| 100507246_a | SNHG16 | -0,60529095 | 10,2140058 | -5,36815211 | 8,74E-05 | 0,00300348 |
| 79573_at | TTC13 | -0,65235189 | 7,55530993 | -5,36751036 | 8,75E-05 | 0,00300348 |
| 85414_at | SLC45A3 | -0,69675498 | 6,28085437 | -5,36470626 | 8,79E-05 | 0,00300918 |
| 5245_at | PHB | -0,63096589 | 9,64602154 | -5,36456545 | 8,79E-05 | 0,00300918 |
| 4312_at | MMP1 | -0,93046781 | 3,85521151 | -5,36337343 | 8,81E-05 | 0,00301048 |
| 3608_at | ILF2 | -0,44971526 | 11,0741695 | -5,36149942 | 8,85E-05 | 0,0030156 |
| 83719_at | YPEL3 | 1,32249015 | 7,22381218 | 5,35746777 | 8,91E-05 | 0,00303284 |
| 55276_at | PGM2 | -0,39884137 | 9,37327421 | -5,35631321 | 8,93E-05 | 0,00303397 |
| 7031_at | TFF1 | -1,42982043 | 7,14503892 | -5,34596518 | 9,10E-05 | 0,00308074 |
| 23310_at | NCAPD3 | -1,0355407 | 8,22399876 | -5,34535437 | 9,11E-05 | 0,00308074 |
| 26519_at | TIMM10 | -0,88940033 | 8,85212571 | -5,34487947 | 9,12E-05 | 0,00308074 |
| 8666_at | EIF3G | -0,31974616 | 10,2984678 | -5,34378529 | 9,14E-05 | 0,00308074 |
| 4343_at | MOV10 | 0,73373379 | 8,27077386 | 5,34332652 | 9,15E-05 | 0,00308074 |
| 2189_at | FANCG | -0,87364805 | 7,73914185 | -5,34181873 | 9,17E-05 | 0,00308397 |
| 5218_at | CDK14 | 0,74806813 | 7,39217518 | 5,33698419 | 9,26E-05 | 0,0031039 |
| 7644_at | ZNF91 | 0,58109229 | 6,92866339 | 5,33646846 | 9,27E-05 | 0,0031039 |
| 2036_at | EPB41L1 | 0,33212699 | 7,99105746 | 5,33467868 | 9,30E-05 | 0,00310738 |
| 79589_at | RNF128 | 0,71989576 | 10,3485171 | 5,33399345 | 9,31E-05 | 0,00310738 |
| 494143_at | CHAC2 | -1,49479877 | 7,98701049 | -5,33274171 | 9,33E-05 | 0,00310745 |
| 121053_at | C12orf45 | -0,60234923 | 6,82929847 | -5,33211774 | 9,34E-05 | 0,00310745 |
| 84946_at | LTV1 | -0,76441239 | 7,91076328 | -5,33037846 | 9,37E-05 | 0,00311212 |
| 97_at | ACYP1 | -0,60054092 | 7,72086101 | -5,32892187 | 9,40E-05 | 0,00311518 |
| 25914_at | RTTN | -0,4379207 | 6,21206257 | -5,32665284 | 9,44E-05 | 0,00312295 |
| 10694_at | CCT8 | -0,51301923 | 11,2091517 | -5,32413926 | 9,48E-05 | 0,00313218 |
| 23165_at | NUP205 | -0,84995374 | 8,57904209 | -5,3166847 | 9,61E-05 | 0,00317042 |
| 1104_at | RCC1 | -0,92887054 | 8,62718251 | -5,31176177 | 9,70E-05 | 0,00319409 |
| 93594_at | TBC1D31 | -0,62936023 | 6,80799553 | -5,3088794 | 9,75E-05 | 0,00320448 |
| 9183_at | ZW10 | -0,52122192 | 7,99850101 | -5,30781368 | 9,77E-05 | 0,00320448 |
| 92181_at | UBTD2 | 0,46506998 | 9,06204698 | 5,30726372 | 9,78E-05 | 0,00320448 |
| 25813_at | SAMM50 | -0,30555161 | 10,1303642 | -5,3028484 | 9,86E-05 | 0,00322163 |
| 55359_at | STYK1 | 0,3821098 | 6,68691286 | 5,30256768 | 9,87E-05 | 0,00322163 |
| 84930_at | MASTL | -0,79301336 | 6,9572997 | -5,30129809 | 9,89E-05 | 0,00322379 |
| 65083_at | NOL6 | -0,5164187 | 6,81553322 | -5,29814285 | 9,95E-05 | 0,00323729 |
| 27335_at | EIF3K | -0,32602086 | 10,8955979 | -5,29628935 | 9,98E-05 | 0,003243 |
| 2224_at | FDPS | -0,6440454 | 9,80533451 | -5,29332831 | 0,00010039 | 0,00325068 |

| | | | | | | |
|---|---|---|---|---|---|---|
| 55009_at | C19orf24 | -0,4086639 | 8,26530007 | -5,29320801 | 0,00010041 | 0,00325068 |
| 9824_at | ARHGAP11A | -1,1859388 | 6,01967986 | -5,28855035 | 0,00010128 | 0,00327349 |
| 6945_at | MLX | -0,4655082 | 9,04809552 | -5,28716437 | 0,00010154 | 0,00327601 |
| 10451_at | VAV3 | 0,82947337 | 7,00073861 | 5,28633708 | 0,0001017 | 0,00327601 |
| 54931_at | TRMT10C | -0,46713801 | 8,5261686 | -5,285223 | 0,00010191 | 0,00327734 |
| 22822_at | PHLDA1 | -1,22804495 | 8,30023339 | -5,28275147 | 0,00010238 | 0,0032786 |
| 23214_at | XPO6 | -0,42377819 | 9,44366177 | -5,28195501 | 0,00010253 | 0,0032786 |
| 1551_at | CYP3A7 | 0,92074487 | 5,5138135 | 5,2817904 | 0,00010257 | 0,0032786 |
| 7298_at | TYMS | -1,4872989 | 9,85514139 | -5,28144205 | 0,00010263 | 0,0032786 |
| 8027_at | STAM | -0,59303821 | 8,36031314 | -5,27327529 | 0,00010421 | 0,00332337 |
| 6839_at | SUV39H1 | -0,80314027 | 7,30573675 | -5,27129773 | 0,00010459 | 0,00333013 |
| 5230_at | PGK1 | -0,36832951 | 10,3657966 | -5,27036061 | 0,00010477 | 0,00333044 |
| 92609_at | TIMM50 | -0,47705967 | 7,34344325 | -5,26677723 | 0,00010548 | 0,00334725 |
| 1894_at | ECT2 | -1,18823248 | 8,46002916 | -5,26399915 | 0,00010602 | 0,0033591 |
| 64393_at | ZMAT3 | 0,59206654 | 7,25791151 | 5,25505826 | 0,00010781 | 0,00340999 |
| 285958_at | SNHG15 | -0,98654241 | 7,92602504 | -5,24830612 | 0,00010917 | 0,00344758 |
| 4909_at | NTF4 | 0,8798151 | 7,45367799 | 5,2469871 | 0,00010944 | 0,00345043 |
| 4121_at | MAN1A1 | 1,29197809 | 8,86532866 | 5,24329957 | 0,0001102 | 0,0034686 |
| 28998_at | MRPL13 | -0,44867388 | 7,80168718 | -5,24020174 | 0,00011084 | 0,00348244 |
| 27086_at | FOXP1 | 0,72433186 | 7,00262185 | 5,23941985 | 0,000111 | 0,00348244 |
| 3673_at | ITGA2 | -1,32920708 | 8,93390334 | -5,23786048 | 0,00011133 | 0,00348693 |
| 2625_at | GATA3 | 0,61602833 | 9,99800819 | 5,22456593 | 0,00011413 | 0,00355649 |
| 10974_at | ADIRF | 0,49032893 | 11,7168294 | 5,22385599 | 0,00011428 | 0,00355649 |
| 55813_at | UTP6 | -0,49368405 | 8,65299443 | -5,22370899 | 0,00011431 | 0,00355649 |
| 79135_at | APOO | -0,51662574 | 8,66491374 | -5,22301033 | 0,00011446 | 0,00355649 |
| 80179_at | MYO19 | -0,60777434 | 7,87549752 | -5,22296261 | 0,00011447 | 0,00355649 |
| 64105_at | CENPK | -1,56277142 | 7,35755343 | -5,21929241 | 0,00011526 | 0,00357523 |
| 114625_at | ERMAP | 0,60980839 | 6,99375015 | 5,21293657 | 0,00011664 | 0,00361219 |
| 23421_at | ITGB3BP | -0,9542494 | 7,76316287 | -5,20934771 | 0,00011742 | 0,0036201 |
| 7126_at | TNFAIP1 | 0,43216399 | 8,4500571 | 5,20919726 | 0,00011746 | 0,0036201 |
| 60528_at | ELAC2 | -0,47193245 | 8,33245975 | -5,20836134 | 0,00011764 | 0,0036201 |
| 10279_at | PRSS16 | 0,64151138 | 7,65245039 | 5,20788564 | 0,00011775 | 0,0036201 |
| 84958_at | SYTL1 | 0,61714731 | 8,76973173 | 5,20749017 | 0,00011783 | 0,0036201 |
| 1652_at | DDT | -0,46307152 | 10,3875235 | -5,20588651 | 0,00011819 | 0,00362521 |
| 5920_at | RARRES3 | 1,74858303 | 8,29398805 | 5,20283634 | 0,00011886 | 0,00364018 |
| 56992_at | KIF15 | -1,43205141 | 6,69772338 | -5,20091012 | 0,00011929 | 0,00364754 |
| 220992_at | ZNF485 | -0,43439362 | 4,6338495 | -5,19840576 | 0,00011985 | 0,00365889 |
| 348995_at | NUP43 | -0,49156672 | 7,56567877 | -5,19664178 | 0,00012025 | 0,00366519 |
| 79590_at | MRPL24 | -0,73288931 | 9,73765261 | -5,19508541 | 0,0001206 | 0,00366608 |
| 54443_at | ANLN | -1,23567711 | 9,49084697 | -5,19482625 | 0,00012066 | 0,00366608 |
| 4600_at | MX2 | 1,65458669 | 9,2525915 | 5,19117273 | 0,00012149 | 0,00368546 |
| 11342_at | RNF13 | 0,53246094 | 9,50774678 | 5,18584353 | 0,00012271 | 0,00371661 |
| 1854_at | DUT | -0,65635883 | 8,158219 | -5,18099481 | 0,00012383 | 0,00374468 |
| 23548_at | TTC33 | 0,74359435 | 6,85643279 | 5,17775448 | 0,00012459 | 0,00376161 |
| 131076_at | CCDC58 | -0,93323815 | 7,52144996 | -5,17380825 | 0,00012551 | 0,00378363 |
| 5869_at | RAB5B | 0,61126559 | 8,924603 | 5,17070384 | 0,00012625 | 0,0037998 |
| 92935_at | MARS2 | -0,66822173 | 6,82049223 | -5,16801587 | 0,00012688 | 0,00380743 |
| 6340_at | SCNN1G | 1,46757622 | 6,13227808 | 5,16797341 | 0,00012689 | 0,00380743 |
| 26973_at | CHORDC1 | -0,55973026 | 6,29798241 | -5,16375413 | 0,0001279 | 0,00383076 |
| 55341_at | LSG1 | -0,44694958 | 8,58094178 | -5,16306409 | 0,00012807 | 0,00383076 |

| | | | | | | |
|---|---|---|---|---|---|---|
| 10293_at | TRAIP | -0,85872283 | 6,12539595 | -5,16135027 | 0,00012848 | 0,00383715 |
| 51078_at | THAP4 | -0,41918684 | 8,31812617 | -5,15489395 | 0,00013005 | 0,00387796 |
| 151354_at | FAM84A | 0,79095932 | 7,2273679 | 5,15252532 | 0,00013063 | 0,00388645 |
| 254048_at | UBN2 | 0,46846151 | 6,41861455 | 5,15208577 | 0,00013074 | 0,00388645 |
| 10574_at | CCT7 | -0,62136079 | 11,0571473 | -5,15047016 | 0,00013114 | 0,00389227 |
| 90627_at | STARD13 | 1,06380566 | 6,91088552 | 5,14889836 | 0,00013152 | 0,00389434 |
| 55577_at | NAGK | 0,67228935 | 10,0706392 | 5,14834585 | 0,00013166 | 0,00389434 |
| 2023_at | ENO1 | -0,56135573 | 11,417873 | -5,14745373 | 0,00013188 | 0,00389434 |
| 6426_at | SRSF1 | -0,51368745 | 10,1240844 | -5,14680094 | 0,00013204 | 0,00389434 |
| 55687_at | TRMU | -0,34188661 | 7,36310615 | -5,14610412 | 0,00013222 | 0,00389434 |
| 64425_at | POLR1E | -0,98064803 | 7,21365468 | -5,14467768 | 0,00013257 | 0,00389884 |
| 6835_at | SURF2 | -0,67044195 | 8,22960138 | -5,13982231 | 0,00013379 | 0,00392861 |
| 25953_at | PNKD | -0,43055787 | 8,21875605 | -5,12518815 | 0,00013752 | 0,00403217 |
| 825_at | CAPN3 | 0,63675681 | 4,81259682 | 5,12412834 | 0,0001378 | 0,00403409 |
| 23396_at | PIP5K1C | 0,39437002 | 7,43438695 | 5,12158463 | 0,00013846 | 0,00403968 |
| 6310_at | ATXN1 | 1,18622962 | 7,75097716 | 5,12101182 | 0,00013861 | 0,00403968 |
| 27349_at | MCAT | -0,35963511 | 8,04002147 | -5,12045093 | 0,00013876 | 0,00403968 |
| 90246_at | LOC90246 | 0,55121845 | 6,83021805 | 5,12017495 | 0,00013883 | 0,00403968 |
| 64318_at | NOC3L | -0,93797646 | 7,92076778 | -5,11743538 | 0,00013955 | 0,00405445 |
| 10054_at | UBA2 | -0,45970614 | 10,8420771 | -5,11626123 | 0,00013986 | 0,00405702 |
| 64089_at | SNX16 | 0,53113764 | 6,44729691 | 5,11482042 | 0,00014024 | 0,00405702 |
| 54107_at | POLE3 | -0,6398917 | 9,08538662 | -5,11470362 | 0,00014027 | 0,00405702 |
| 9772_at | KIAA0195 | 0,50495171 | 8,40742734 | 5,11043866 | 0,0001414 | 0,00408296 |
| 1763_at | DNA2 | -0,80839585 | 6,99465505 | -5,10960726 | 0,00014162 | 0,00408296 |
| 125061_at | AFMID | -0,47717999 | 6,16865174 | -5,10853481 | 0,00014191 | 0,00408296 |
| 10528_at | NOP56 | -0,81975845 | 9,7360895 | -5,10814414 | 0,00014201 | 0,00408296 |
| 55226_at | NAT10 | -0,80576077 | 8,05954973 | -5,10497643 | 0,00014286 | 0,0041013 |
| 23255_at | MTCL1 | 0,65472172 | 7,62003655 | 5,09927325 | 0,00014441 | 0,00413517 |
| 84961_at | FBXL20 | 0,52660767 | 6,57313291 | 5,0990379 | 0,00014447 | 0,00413517 |
| 79892_at | MCMBP | -0,71124879 | 9,17863638 | -5,09708374 | 0,00014501 | 0,00414428 |
| 255104_at | TMCO4 | 0,44309799 | 6,81811813 | 5,09267261 | 0,00014622 | 0,00417273 |
| 57466_at | SCAF4 | -0,3055143 | 7,99150625 | -5,0838878 | 0,00014866 | 0,00422727 |
| 54512_at | EXOSC4 | -0,59858434 | 8,6562631 | -5,08312346 | 0,00014888 | 0,00422727 |
| 1512_at | CTSH | 0,3086593 | 12,4410794 | 5,08283717 | 0,00014896 | 0,00422727 |
| 2592_at | GALT | 0,32722927 | 8,10908417 | 5,0826632 | 0,00014901 | 0,00422727 |
| 84128_at | WDR75 | -0,65263881 | 8,09418369 | -5,07882441 | 0,00015009 | 0,00424664 |
| 6237_at | RRAS | 0,66891192 | 9,33969197 | 5,07868508 | 0,00015013 | 0,00424664 |
| 10213_at | PSMD14 | -0,66363154 | 10,5754415 | -5,07423469 | 0,0001514 | 0,00427622 |
| 8870_at | IER3 | -0,79165119 | 9,7302693 | -5,07190029 | 0,00015207 | 0,00428883 |
| 25844_at | YIPF3 | 0,6346027 | 9,1705746 | 5,0666674 | 0,00015358 | 0,00432513 |
| 64115_at | C10orf54 | 0,76056888 | 7,47206649 | 5,06494438 | 0,00015408 | 0,00433291 |
| 51696_at | HECA | 0,48499116 | 8,39340741 | 5,06217387 | 0,00015489 | 0,00434557 |
| 10436_at | EMG1 | -0,73067208 | 8,92850329 | -5,06102843 | 0,00015522 | 0,00434557 |
| 84572_at | GNPTG | 0,56589335 | 8,16898512 | 5,06047289 | 0,00015539 | 0,00434557 |
| 51320_at | MEX3C | 0,40903334 | 7,90191186 | 5,06032152 | 0,00015543 | 0,00434557 |
| 7753_at | ZNF202 | -0,29590185 | 6,52619651 | -5,04884461 | 0,00015884 | 0,00443173 |
| 9249_at | DHRS3 | 0,40778019 | 10,6796243 | 5,04841187 | 0,00015897 | 0,00443173 |
| 6947_at | TCN1 | -1,26857374 | 6,02227847 | -5,04348286 | 0,00016046 | 0,00446681 |
| 5580_at | PRKCD | 0,6561328 | 9,00500412 | 5,04060462 | 0,00016134 | 0,00448474 |
| 54861_at | SNRK | 0,43309507 | 8,15511545 | 5,03925518 | 0,00016175 | 0,00448974 |

| | | | | | | |
|---|---|---|---|---|---|---|
| 9500_at | MAGED1 | 0,71969797 | 10,3818166 | 5,03499163 | 0,00016307 | 0,00451629 |
| 84441_at | MAML2 | 0,68427866 | 7,83226622 | 5,03462257 | 0,00016318 | 0,00451629 |
| 51010_at | EXOSC3 | -0,50588183 | 7,77968457 | -5,02909073 | 0,0001649 | 0,00455427 |
| 6892_at | TAPBP | 0,74386593 | 9,91040131 | 5,02859986 | 0,00016505 | 0,00455427 |
| 25796_at | PGLS | 0,3914984 | 8,87292852 | 5,02793352 | 0,00016526 | 0,00455427 |
| 55149_at | MTPAP | -0,4996958 | 7,38268543 | -5,02696473 | 0,00016557 | 0,00455612 |
| 84879_at | MFSD2A | -0,75916354 | 7,66992812 | -5,02569071 | 0,00016597 | 0,00456062 |
| 5591_at | PRKDC | -0,61308435 | 9,09097673 | -5,02427759 | 0,00016641 | 0,00456164 |
| 10538_at | BATF | 0,40286455 | 8,43670752 | 5,02406969 | 0,00016648 | 0,00456164 |
| 10038_at | PARP2 | -0,64978085 | 8,58647113 | -5,01862715 | 0,0001682 | 0,00460016 |
| 201254_at | STRA13 | -0,72070355 | 9,84308038 | -5,01813689 | 0,00016836 | 0,00460016 |
| 140738_at | TMEM37 | 0,54946252 | 6,28048493 | 5,0169303 | 0,00016875 | 0,00460417 |
| 131601_at | TPRA1 | 0,31254393 | 8,09581396 | 5,01439545 | 0,00016956 | 0,00461257 |
| 158358_at | KIAA2026 | 0,42012141 | 6,70371238 | 5,01383627 | 0,00016974 | 0,00461257 |
| 200315_at | APOBEC3A | 2,09074513 | 5,2483111 | 5,01373379 | 0,00016977 | 0,00461257 |
| 284161_at | GDPD1 | 0,94235597 | 4,65023029 | 5,00753491 | 0,00017178 | 0,00466057 |
| 28970_at | C11orf54 | 0,72504852 | 7,60904652 | 5,00586418 | 0,00017233 | 0,00466285 |
| 8565_at | YARS | -0,85904045 | 9,43178629 | -5,00579455 | 0,00017235 | 0,00466285 |
| 7407_at | VARS | -0,7145902 | 7,79621877 | -5,00415602 | 0,00017289 | 0,00466995 |
| 5590_at | PRKCZ | 0,42238928 | 8,66954176 | 5,00351497 | 0,0001731 | 0,00466995 |
| 79922_at | MRM1 | -0,4782001 | 6,87029979 | -5,00063507 | 0,00017404 | 0,00468899 |
| 78987_at | CRELD1 | 0,56403793 | 6,49277723 | 4,9991988 | 0,00017452 | 0,00469152 |
| 79723_at | SUV39H2 | -0,59824885 | 6,49904637 | -4,99888002 | 0,00017463 | 0,00469152 |
| 790_at | CAD | -0,54515656 | 7,7294139 | -4,99635379 | 0,00017547 | 0,00470751 |
| 85456_at | TNKS1BP1 | 0,47144833 | 6,82379029 | 4,99429925 | 0,00017615 | 0,00471935 |
| 54453_at | RIN2 | 0,41161166 | 9,42204028 | 4,99169968 | 0,00017702 | 0,00473612 |
| 166336_at | PRICKLE2 | 0,9491417 | 5,48207819 | 4,98992758 | 0,00017762 | 0,0047455 |
| 284058_at | KANSL1 | 0,3527495 | 8,94489758 | 4,98549551 | 0,00017912 | 0,00477901 |
| 100132707_a | PAXIP1-AS2 | 0,61613409 | 6,7208046 | 4,98367273 | 0,00017974 | 0,00478287 |
| 55299_at | BRIX1 | -1,07204251 | 8,1810964 | -4,98361507 | 0,00017976 | 0,00478287 |
| 210_at | ALAD | 0,43155817 | 7,18883944 | 4,9784534 | 0,00018154 | 0,00482337 |
| 11337_at | GABARAP | 0,57283555 | 10,7613957 | 4,97705464 | 0,00018202 | 0,0048257 |
| 10142_at | AKAP9 | 0,75805094 | 6,89534409 | 4,97675057 | 0,00018213 | 0,0048257 |
| 5547_at | PRCP | 0,8737258 | 9,59795975 | 4,97454966 | 0,00018289 | 0,0048261 |
| 124220_at | ZG16B | 0,5094473 | 8,68277973 | 4,97450617 | 0,0001829 | 0,0048261 |
| 11171_at | STRAP | -0,32552388 | 10,7780484 | -4,97440182 | 0,00018294 | 0,0048261 |
| 5710_at | PSMD4 | -0,33663546 | 10,3663903 | -4,97364853 | 0,0001832 | 0,0048261 |
| 1665_at | DHX15 | -0,47314507 | 10,5846285 | -4,97279383 | 0,0001835 | 0,0048261 |
| 56995_at | TULP4 | 0,24782705 | 7,71551631 | 4,97203023 | 0,00018377 | 0,0048261 |
| 59307_at | SIGIRR | 0,38133976 | 7,42395086 | 4,97166912 | 0,00018389 | 0,0048261 |
| 57486_at | NLN | -0,57174786 | 7,07023474 | -4,9702614 | 0,00018439 | 0,00483246 |
| 65260_at | COA7 | -0,73246066 | 7,47013059 | -4,96808082 | 0,00018515 | 0,00483257 |
| 5734_at | PTGER4 | 0,72866439 | 8,35213953 | 4,96789377 | 0,00018522 | 0,00483257 |
| 55787_at | TXLNG | -0,54999418 | 7,41353972 | -4,96682607 | 0,0001856 | 0,00483257 |
| 25901_at | CCDC28A | 0,59920142 | 7,20341883 | 4,9667535 | 0,00018562 | 0,00483257 |
| 7283_at | TUBG1 | -1,07531119 | 10,4434906 | -4,96626792 | 0,00018579 | 0,00483257 |
| 57761_at | TRIB3 | -0,59202794 | 7,13215771 | -4,96597567 | 0,0001859 | 0,00483257 |
| 4809_at | SNU13 | -0,45689434 | 9,27404699 | -4,96262449 | 0,00018709 | 0,00485694 |
| 151050_at | KANSL1L | 0,70878058 | 5,78079415 | 4,95914993 | 0,00018833 | 0,00488259 |
| 92106_at | OXNAD1 | -0,66416266 | 7,022488 | -4,95639456 | 0,00018932 | 0,00490167 |

| | | | | | | |
|---|---|---|---|---|---|---|
| 3716_at | JAK1 | 0,46234765 | 9,17239487 | 4,95560554 | 0,0001896 | 0,00490245 |
| 91452_at | ACBD5 | 0,53183027 | 8,22359146 | 4,95222652 | 0,00019083 | 0,00492749 |
| 4953_at | ODC1 | -1,02236954 | 10,119975 | -4,9465905 | 0,00019289 | 0,00497401 |
| 3659_at | IRF1 | 1,0943914 | 8,53409337 | 4,94530648 | 0,00019336 | 0,00497952 |
| 25938_at | HEATR5A | 0,64076256 | 7,84640635 | 4,94267321 | 0,00019433 | 0,00499789 |
| 136895_at | C7orf31 | 0,90367295 | 5,70746695 | 4,94158657 | 0,00019474 | 0,00500158 |
| 64063_at | PRSS22 | 0,63128427 | 7,61928227 | 4,94073333 | 0,00019505 | 0,00500305 |
| 22948_at | CCT5 | -0,88004366 | 8,41268168 | -4,9380763 | 0,00019604 | 0,00501706 |
| 57492_at | ARID1B | 0,30741554 | 7,68213991 | 4,93716792 | 0,00019638 | 0,00501706 |
| 55070_at | DET1 | 0,51774644 | 6,8066764 | 4,93655227 | 0,00019661 | 0,00501706 |
| 4645_at | MYO5B | 0,6256047 | 7,40501667 | 4,93551892 | 0,000197 | 0,00501706 |
| 7376_at | NR1H2 | 0,3831491 | 8,21403241 | 4,93540911 | 0,00019704 | 0,00501706 |
| 25879_at | DCAF13 | -0,41753898 | 9,38838415 | -4,93509194 | 0,00019716 | 0,00501706 |
| 3320_at | HSP90AA1 | -0,23109836 | 12,3012744 | -4,93382265 | 0,00019764 | 0,00502258 |
| 6482_at | ST3GAL1 | 0,62216793 | 8,81635999 | 4,93286932 | 0,000198 | 0,00502509 |
| 81611_at | ANP32E | -0,69795675 | 8,70842304 | -4,92994249 | 0,00019911 | 0,00504657 |
| 51278_at | IER5 | -0,42900497 | 9,00138745 | -4,92669332 | 0,00020034 | 0,00507127 |
| 3425_at | IDUA | 0,41947006 | 6,59542365 | 4,92250494 | 0,00020195 | 0,00510482 |
| 79073_at | TMEM109 | -0,4210885 | 9,64908 | -4,92186229 | 0,0002022 | 0,00510482 |
| 10721_at | POLQ | -1,15583366 | 6,91340694 | -4,92104859 | 0,00020251 | 0,00510606 |
| 7284_at | TUFM | -0,42145805 | 7,90119201 | -4,91750792 | 0,00020389 | 0,00513396 |
| 171568_at | POLR3H | -0,34660514 | 8,00329746 | -4,90666248 | 0,00020815 | 0,0052199 |
| 10478_at | SLC25A17 | -0,2980607 | 7,06949461 | -4,90652824 | 0,00020821 | 0,0052199 |
| 53827_at | FXYD5 | -0,47713952 | 8,26129727 | -4,90613698 | 0,00020836 | 0,0052199 |
| 140461_at | ASB8 | 0,35223286 | 7,81381273 | 4,90608297 | 0,00020838 | 0,0052199 |
| 5987_at | TRIM27 | -0,52351129 | 8,9899062 | -4,90020685 | 0,00021074 | 0,00526511 |
| 5717_at | PSMD11 | -0,61783626 | 8,89403656 | -4,89965152 | 0,00021096 | 0,00526511 |
| 375287_at | RBM43 | 0,72096176 | 5,94553587 | 4,89953189 | 0,00021101 | 0,00526511 |
| 51692_at | CPSF3 | -0,52557566 | 8,84291381 | -4,89847877 | 0,00021143 | 0,00526889 |
| 5819_at | PVRL2 | 0,31746197 | 8,14403937 | 4,89578366 | 0,00021253 | 0,00528926 |
| 2935_at | GSPT1 | -0,41024027 | 8,54396916 | -4,89464957 | 0,00021299 | 0,0052939 |
| 4739_at | NEDD9 | 0,50593767 | 8,27862751 | 4,8926978 | 0,00021378 | 0,00530684 |
| 51575_at | ESF1 | -0,62542519 | 7,0165519 | -4,89173194 | 0,00021418 | 0,00530981 |
| 10592_at | SMC2 | -0,88262639 | 7,44393269 | -4,88801644 | 0,00021571 | 0,00534079 |
| 90102_at | PHLDB2 | 0,43067483 | 7,8989392 | 4,88700965 | 0,00021612 | 0,00534421 |
| 6742_at | SSBP1 | -0,48425191 | 9,81718252 | -4,8825029 | 0,00021799 | 0,00538357 |
| 908_at | CCT6A | -0,77667943 | 9,62243952 | -4,8816532 | 0,00021835 | 0,00538543 |
| 55968_at | NSFL1C | -0,28225944 | 9,14105438 | -4,87865018 | 0,00021961 | 0,00540857 |
| 29028_at | ATAD2 | -1,24976468 | 8,52981405 | -4,87807575 | 0,00021985 | 0,00540857 |
| 84816_at | RTN4IP1 | -0,33493044 | 6,76387199 | -4,87596483 | 0,00022074 | 0,00542353 |
| 6723_at | SRM | -1,06855388 | 10,1210042 | -4,87522551 | 0,00022105 | 0,00542429 |
| 64121_at | RRAGC | 0,41382287 | 8,75922836 | 4,87308784 | 0,00022196 | 0,0054396 |
| 81631_at | MAP1LC3B | 0,61521338 | 9,49085127 | 4,87170575 | 0,00022254 | 0,00544708 |
| 55135_at | WRAP53 | -0,52446891 | 7,62978083 | -4,86997692 | 0,00022328 | 0,00545821 |
| 63967_at | CLSPN | -0,70628027 | 5,14398065 | -4,86914243 | 0,00022364 | 0,00546001 |
| 93323_at | HAUS8 | -0,77354853 | 6,53927181 | -4,8678302 | 0,0002242 | 0,00546559 |
| 79641_at | ROGDI | 0,54954252 | 8,42714247 | 4,86728603 | 0,00022443 | 0,00546559 |
| 9521_at | EEF1E1 | -0,54584626 | 8,09231787 | -4,86559461 | 0,00022516 | 0,00547231 |
| 54361_at | WNT4 | 0,46810749 | 6,15412123 | 4,86532648 | 0,00022528 | 0,00547231 |
| 55695_at | NSUN5 | -0,44100268 | 8,44760501 | -4,86213137 | 0,00022666 | 0,00548557 |

| | | | | | | |
|---|---|---|---|---|---|---|
| 54962_at | TIPIN | -1,01399735 | 7,20933522 | -4,86085876 | 0,00022721 | 0,00548557 |
| 9128_at | PRPF4 | -0,60221745 | 8,56758212 | -4,86055086 | 0,00022735 | 0,00548557 |
| 84131_at | CEP78 | -0,90188563 | 7,46324046 | -4,85996655 | 0,0002276 | 0,00548557 |
| 8760_at | CDS2 | 0,46633051 | 8,51194087 | 4,85944679 | 0,00022783 | 0,00548557 |
| 9343_at | EFTUD2 | -0,33834182 | 9,73426136 | -4,85897023 | 0,00022804 | 0,00548557 |
| 26031_at | OSBPL3 | -0,58305797 | 7,04430452 | -4,85896963 | 0,00022804 | 0,00548557 |
| 25855_at | BRMS1 | -0,31915521 | 8,77319811 | -4,85882251 | 0,0002281 | 0,00548557 |
| 55131_at | RBM28 | -0,67904725 | 8,2875007 | -4,8579953 | 0,00022846 | 0,00548742 |
| 79707_at | NOL9 | -0,37290203 | 7,70968336 | -4,85554375 | 0,00022954 | 0,00550639 |
| 6567_at | SLC16A2 | 0,65711898 | 5,33871186 | 4,853052 | 0,00023064 | 0,00552587 |
| 4616_at | GADD45B | 0,6992363 | 7,94252758 | 4,8519914 | 0,00023111 | 0,00552757 |
| 23625_at | FAM89B | 0,44995223 | 9,65655752 | 4,85159481 | 0,00023128 | 0,00552757 |
| 57510_at | XPO5 | -0,49777547 | 7,99097032 | -4,84857207 | 0,00023263 | 0,00555281 |
| 168455_at | CCDC71L | 0,49088894 | 6,67149537 | 4,8446927 | 0,00023436 | 0,00558734 |
| 705_at | BYSL | -0,95172372 | 8,33164197 | -4,83897198 | 0,00023695 | 0,005642 |
| 122769_at | LRR1 | -1,06546657 | 7,81153513 | -4,82945915 | 0,00024132 | 0,00573406 |
| 92667_at | MGME1 | -0,42881233 | 9,03138781 | -4,82925031 | 0,00024141 | 0,00573406 |
| 81558_at | FAM117A | 0,45766499 | 7,33022917 | 4,82809925 | 0,00024195 | 0,00573869 |
| 92747_at | BPIFB1 | 1,49163131 | 6,92536992 | 4,82754584 | 0,0002422 | 0,00573869 |
| 100505839_a | SH3PXD2A-AS | -0,66427426 | 4,78002581 | -4,82670388 | 0,00024259 | 0,00574092 |
| 7005_at | TEAD3 | 0,45493047 | 7,37674612 | 4,82108584 | 0,00024523 | 0,00579606 |
| 51142_at | CHCHD2 | -0,42850812 | 10,3292685 | -4,81846758 | 0,00024646 | 0,00581422 |
| 2192_at | FBLN1 | 0,72117285 | 10,3430297 | 4,81818019 | 0,0002466 | 0,00581422 |
| 3416_at | IDE | -0,35029895 | 9,20031483 | -4,81349354 | 0,00024883 | 0,00585963 |
| 7866_at | IFRD2 | -1,00970825 | 8,71899073 | -4,81214151 | 0,00024948 | 0,0058677 |
| 9189_at | ZBED1 | 0,54656154 | 7,72537458 | 4,81073395 | 0,00025015 | 0,00587642 |
| 9682_at | KDM4A | 0,58971153 | 7,97963171 | 4,80270699 | 0,00025404 | 0,00594967 |
| 10921_at | RNPS1 | -0,29777267 | 9,2220195 | -4,80269035 | 0,00025405 | 0,00594967 |
| 10126_at | DNAL4 | 0,39317541 | 7,61591253 | 4,80235722 | 0,00025421 | 0,00594967 |
| 54919_at | DNAAF5 | -0,39417497 | 8,67371878 | -4,8017574 | 0,0002545 | 0,00594967 |
| 51651_at | PTRH2 | -0,62777495 | 8,03305875 | -4,80008279 | 0,00025533 | 0,00596162 |
| 10521_at | DDX17 | 0,30543363 | 8,5012249 | 4,79329998 | 0,00025868 | 0,00602855 |
| 79693_at | YRDC | -0,45999558 | 8,44964335 | -4,7930193 | 0,00025882 | 0,00602855 |
| 79053_at | ALG8 | -0,44692322 | 9,54099753 | -4,79012789 | 0,00026026 | 0,00605485 |
| 5714_at | PSMD8 | -0,34300205 | 10,6413203 | -4,78893321 | 0,00026086 | 0,00606147 |
| 10432_at | RBM14 | -0,55607735 | 7,79281498 | -4,78724191 | 0,00026171 | 0,00606883 |
| 9185_at | REPS2 | 0,47608193 | 6,27329616 | 4,78705037 | 0,00026181 | 0,00606883 |
| 5608_at | MAP2K6 | 0,44237994 | 6,22898636 | 4,78443729 | 0,00026313 | 0,00607626 |
| 7562_at | ZNF708 | 0,72403329 | 6,16130987 | 4,78389792 | 0,0002634 | 0,00607626 |
| 65263_at | PYCRL | -0,70760491 | 6,96340669 | -4,7835712 | 0,00026357 | 0,00607626 |
| 8673_at | VAMP8 | 0,41673807 | 10,9593789 | 4,78282328 | 0,00026395 | 0,00607626 |
| 10243_at | GPHN | -0,36495597 | 7,33639548 | -4,78273988 | 0,00026399 | 0,00607626 |
| 84216_at | TMEM117 | 0,50868925 | 7,68713648 | 4,78213868 | 0,00026429 | 0,00607626 |
| 4176_at | MCM7 | -1,12322096 | 9,60198198 | -4,782056 | 0,00026434 | 0,00607626 |
| 80312_at | TET1 | 0,81704834 | 5,33902906 | 4,77983659 | 0,00026547 | 0,00608798 |
| 23397_at | NCAPH | -1,34209059 | 8,22839609 | -4,77981654 | 0,00026548 | 0,00608798 |
| 26784_at | SNORA64 | -0,88269002 | 6,37098132 | -4,77660099 | 0,00026713 | 0,00611239 |
| 55010_at | PARPBP | -0,99991684 | 6,29839102 | -4,7765036 | 0,00026718 | 0,00611239 |
| 23543_at | RBFOX2 | 0,42971188 | 9,51426537 | 4,7706288 | 0,00027022 | 0,00617461 |
| 6730_at | SRP68 | -0,34523253 | 9,65739269 | -4,76890321 | 0,00027112 | 0,00618782 |

| | | | | | | |
|---|---|---|---|---|---|---|
| 1841_at | DTYMK | -1,06622926 | 8,94197106 | -4,76785712 | 0,00027166 | 0,00619297 |
| 7818_at | DAP3 | -0,38602338 | 9,63459899 | -4,76597142 | 0,00027265 | 0,00620816 |
| 5982_at | RFC2 | -1,09227401 | 8,77722492 | -4,76318354 | 0,00027412 | 0,00623424 |
| 6867_at | TACC1 | 0,80130768 | 8,87656952 | 4,75830006 | 0,00027671 | 0,0062825 |
| 407975_at | MIR17HG | -0,90283065 | 4,76299804 | -4,75748525 | 0,00027715 | 0,0062825 |
| 8872_at | CDC123 | -0,48892886 | 9,65783683 | -4,75719963 | 0,0002773 | 0,0062825 |
| 9221_at | NOLC1 | -0,84283157 | 8,69333835 | -4,75673807 | 0,00027755 | 0,0062825 |
| 27166_at | PRELID1 | -0,65870917 | 9,8556812 | -4,75273357 | 0,0002797 | 0,00632377 |
| 55789_at | DEPDC1B | -0,84822052 | 8,23286374 | -4,74889356 | 0,00028178 | 0,00636331 |
| 64208_at | POPDC3 | -0,52242511 | 4,28064227 | -4,74720018 | 0,0002827 | 0,00637665 |
| 79441_at | HAUS3 | -0,55267699 | 7,08651976 | -4,74550696 | 0,00028363 | 0,00639003 |
| 55035_at | NOL8 | -0,39561849 | 8,06601339 | -4,74428674 | 0,00028429 | 0,00639761 |
| 11140_at | CDC37 | -0,31995155 | 9,73270578 | -4,74286607 | 0,00028507 | 0,00640768 |
| 84331_at | FAM195A | -0,51155203 | 7,96615409 | -4,74054616 | 0,00028635 | 0,00642893 |
| 1508_at | CTSB | 0,61581874 | 8,62679912 | 4,73860407 | 0,00028743 | 0,00644556 |
| 51202_at | DDX47 | -0,32730078 | 10,195996 | -4,73734496 | 0,00028813 | 0,00644721 |
| 81610_at | FAM83D | -1,25224093 | 9,76807894 | -4,73726648 | 0,00028817 | 0,00644721 |
| 25886_at | POC1A | -0,77977859 | 6,51930858 | -4,7343416 | 0,0002898 | 0,00647619 |
| 9585_at | KIF20B | -0,94534356 | 6,76165392 | -4,73117214 | 0,00029158 | 0,00650839 |
| 5639_at | PRRG2 | 0,35996525 | 6,73264885 | 4,72781804 | 0,00029348 | 0,0065431 |
| 152926_at | PPM1K | 0,89839996 | 5,41928123 | 4,72662815 | 0,00029415 | 0,00655057 |
| 23637_at | RABGAP1 | 0,37316673 | 8,84854984 | 4,72425107 | 0,00029551 | 0,00657311 |
| 51729_at | WBP11 | -0,37258772 | 8,98616488 | -4,72309833 | 0,00029616 | 0,00657393 |
| 672_at | BRCA1 | -0,9783164 | 7,10595839 | -4,72299259 | 0,00029622 | 0,00657393 |
| 84283_at | TMEM79 | 0,63605841 | 6,68068641 | 4,72078023 | 0,00029749 | 0,00659293 |
| 4605_at | MYBL2 | -1,20172699 | 8,79291775 | -4,72030762 | 0,00029777 | 0,00659293 |
| 284992_at | CCDC150 | -0,50282332 | 4,83573715 | -4,71912772 | 0,00029844 | 0,00660039 |
| 9931_at | HELZ | 0,31483666 | 8,1358822 | 4,71764077 | 0,0002993 | 0,00661179 |
| 9052_at | GPRC5A | 0,47485792 | 11,1245685 | 4,71676413 | 0,00029981 | 0,00661541 |
| 4125_at | MAN2B1 | 0,5158613 | 8,81628296 | 4,71511711 | 0,00030077 | 0,00662748 |
| 26574_at | AATF | -0,6256668 | 9,96224938 | -4,71463708 | 0,00030105 | 0,00662748 |
| 51182_at | HSPA14 | -0,39747084 | 7,3445217 | -4,71367673 | 0,00030161 | 0,00662945 |
| 5705_at | PSMC5 | -0,34175198 | 9,81026751 | -4,71305888 | 0,00030197 | 0,00662945 |
| 5693_at | PSMB5 | -0,42034477 | 9,9794006 | -4,71271273 | 0,00030217 | 0,00662945 |
| 60488_at | MRPS35 | -0,34257967 | 9,28593231 | -4,7105596 | 0,00030343 | 0,00664952 |
| 79954_at | NOL10 | -0,52394134 | 7,97987684 | -4,70095142 | 0,00030912 | 0,00676653 |
| 10105_at | PPIF | -0,5986464 | 10,1891406 | -4,69943042 | 0,00031003 | 0,00677876 |
| 84769_at | MPV17L2 | -0,40966361 | 7,99332169 | -4,69764013 | 0,0003111 | 0,00679457 |
| 5983_at | RFC3 | -1,2142795 | 7,64578563 | -4,69322629 | 0,00031377 | 0,00684508 |
| 10404_at | CPQ | 0,87745376 | 6,6784357 | 4,68678283 | 0,00031771 | 0,00692314 |
| 10956_at | OS9 | 0,40290701 | 9,4567444 | 4,68612075 | 0,00031812 | 0,00692419 |
| 134492_at | NUDCD2 | -0,45508803 | 8,45199039 | -4,68446465 | 0,00031914 | 0,00693856 |
| 1207_at | CLNS1A | -0,38563861 | 10,2766789 | -4,68388363 | 0,0003195 | 0,00693856 |
| 57171_at | DOLPP1 | -0,56088689 | 8,82582457 | -4,68313184 | 0,00031996 | 0,00694085 |
| 84078_at | KBTBD7 | 0,42961955 | 7,57553424 | 4,68071715 | 0,00032146 | 0,00696553 |
| 64839_at | FBXL17 | 0,38495245 | 6,19757734 | 4,67917837 | 0,00032242 | 0,00697413 |
| 6257_at | RXRB | 0,37988914 | 7,42525533 | 4,67892011 | 0,00032258 | 0,00697413 |
| 1967_at | EIF2B1 | -0,42276988 | 9,45031226 | -4,67739377 | 0,00032354 | 0,00698693 |
| 1983_at | EIF5 | -0,47146996 | 10,6068336 | -4,67526845 | 0,00032487 | 0,0069996 |
| 158586_at | ZXDB | 0,41980695 | 6,15138748 | 4,67498334 | 0,00032505 | 0,0069996 |

| | | | | | | |
|---|---|---|---|---|---|---|
| 8816_at | DCAF5 | 0,43190216 | 8,02926249 | 4,67472534 | 0,00032522 | 0,0069996 |
| 3948_at | LDHC | -0,53248281 | 4,22571872 | -4,6721589 | 0,00032684 | 0,00702664 |
| 2588_at | GALNS | 0,43377353 | 8,87641034 | 4,67011741 | 0,00032813 | 0,00704139 |
| 1326_at | MAP3K8 | 0,50958606 | 5,49909232 | 4,66992615 | 0,00032825 | 0,00704139 |
| 10090_at | UST | 1,15564294 | 7,0476529 | 4,66846409 | 0,00032919 | 0,00705351 |
| 1777_at | DNASE2 | 0,51205205 | 8,13490147 | 4,66761446 | 0,00032973 | 0,00705507 |
| 8417_at | STX7 | 0,62374602 | 7,5445853 | 4,66720361 | 0,00032999 | 0,00705507 |
| 5087_at | PBX1 | 0,28657501 | 9,93281275 | 4,6659783 | 0,00033078 | 0,007064 |
| 55795_at | PCID2 | -0,41857551 | 7,87889265 | -4,66166126 | 0,00033356 | 0,00711547 |
| 9493_at | KIF23 | -1,1117286 | 8,15166564 | -4,65978331 | 0,00033477 | 0,00713003 |
| 7673_at | ZNF222 | 0,4652281 | 5,81133851 | 4,65946612 | 0,00033498 | 0,00713003 |
| 29107_at | NXT1 | -0,56484709 | 9,35017211 | -4,65841593 | 0,00033566 | 0,00713668 |
| 94103_at | ORMDL3 | 0,54498664 | 7,53286986 | 4,656976 | 0,0003366 | 0,00714874 |
| 51318_at | MRPL35 | -0,63777791 | 7,64027252 | -4,65320286 | 0,00033907 | 0,00719331 |
| 57515_at | SERINC1 | 0,59157548 | 10,2423733 | 4,64949277 | 0,00034152 | 0,0072373 |
| 56906_at | THAP10 | -0,57129635 | 7,75039325 | -4,6458194 | 0,00034396 | 0,00728106 |
| 54558_at | SPATA6 | 0,67650168 | 4,99518888 | 4,64204313 | 0,00034649 | 0,00732656 |
| 64847_at | SPATA20 | 0,38923517 | 8,97235644 | 4,63937365 | 0,00034829 | 0,00735655 |
| 728568_at | C12orf73 | -0,28622998 | 6,85602592 | -4,63600018 | 0,00035058 | 0,00739677 |
| 284390_at | ZNF763 | 0,68513141 | 3,60021899 | 4,63441978 | 0,00035166 | 0,00741139 |
| 51192_at | CKLF | -0,82199049 | 8,42154477 | -4,63340716 | 0,00035235 | 0,00741786 |
| 23277_at | CLUH | -0,43874738 | 8,49275405 | -4,63063819 | 0,00035425 | 0,00744971 |
| 1540_at | CYLD | 0,56778973 | 6,09021968 | 4,6289658 | 0,0003554 | 0,00746576 |
| 5777_at | PTPN6 | 0,64108359 | 8,5011937 | 4,62833763 | 0,00035584 | 0,00746576 |
| 81847_at | RNF146 | 0,54650875 | 7,45879403 | 4,62784691 | 0,00035618 | 0,00746576 |
| 4436_at | MSH2 | -0,57432716 | 8,09049832 | -4,62636696 | 0,0003572 | 0,00746673 |
| 91949_at | COG7 | 0,25952458 | 7,57610169 | 4,62615308 | 0,00035735 | 0,00746673 |
| 55837_at | EAPP | 0,38366743 | 8,1654548 | 4,62610237 | 0,00035739 | 0,00746673 |
| 27436_at | EML4 | -0,43911398 | 8,39674551 | -4,62274825 | 0,00035972 | 0,0075074 |
| 64151_at | NCAPG | -1,42373431 | 8,34908976 | -4,61835348 | 0,00036281 | 0,00756359 |
| 2967_at | GTF2H3 | -0,28975861 | 7,39474147 | -4,6147583 | 0,00036535 | 0,00760839 |
| 79415_at | C17orf62 | 0,30474731 | 8,29789377 | 4,6124647 | 0,00036698 | 0,00763414 |
| 60412_at | EXOC4 | 0,37914241 | 7,50041394 | 4,60950579 | 0,0003691 | 0,0076699 |
| 5817_at | PVR | -0,41654068 | 7,25569715 | -4,6083756 | 0,00036991 | 0,00767849 |
| 84749_at | USP30 | 0,36406635 | 7,64350393 | 4,60550986 | 0,00037198 | 0,0077131 |
| 701_at | BUB1B | -1,27161628 | 8,93874456 | -4,6032603 | 0,00037361 | 0,00773314 |
| 51611_at | DPH5 | -0,40947431 | 6,98022177 | -4,60306995 | 0,00037375 | 0,00773314 |
| 81607_at | PVRL4 | 0,58760499 | 7,26814885 | 4,60163515 | 0,00037479 | 0,00774643 |
| 9534_at | ZNF254 | 0,43502242 | 5,15921894 | 4,59861599 | 0,000377 | 0,00778371 |
| 4200_at | ME2 | -0,42918835 | 8,00030177 | -4,59748626 | 0,00037783 | 0,00779249 |
| 10420_at | TESK2 | 0,73918797 | 6,76503723 | 4,59679499 | 0,00037834 | 0,00779463 |
| 8061_at | FOSL1 | -1,87071803 | 6,9129669 | -4,59430071 | 0,00038018 | 0,00782419 |
| 23395_at | LARS2 | -0,38763426 | 7,65390165 | -4,59222526 | 0,00038172 | 0,00784747 |
| 11118_at | BTN3A2 | 1,3975228 | 7,82477299 | 4,59151718 | 0,00038225 | 0,00784992 |
| 114049_at | WBSCR22 | -0,28323211 | 9,54884911 | -4,58997785 | 0,00038339 | 0,00786509 |
| 3385_at | ICAM3 | -0,53226295 | 7,49051458 | -4,58757293 | 0,00038519 | 0,00788721 |
| 10280_at | SIGMAR1 | -0,50679982 | 10,0006264 | -4,58744184 | 0,00038529 | 0,00788721 |
| 55352_at | COPRS | -0,62833023 | 9,13720453 | -4,58420712 | 0,00038773 | 0,00792862 |
| 89845_at | ABCC10 | 0,54315342 | 8,24711184 | 4,58346763 | 0,00038828 | 0,00793162 |
| 23446_at | SLC44A1 | 0,46456012 | 8,22585492 | 4,58218025 | 0,00038926 | 0,00794311 |

| | | | | | | |
|---|---|---|---|---|---|---|
| 79009_at | DDX50 | -0,31846113 | 8,57244397 | -4,58105216 | 0,00039012 | 0,00795215 |
| 83464_at | APH1B | 0,5402762 | 5,56538289 | 4,57944771 | 0,00039134 | 0,00796861 |
| 79137_at | FAM134A | 0,33063405 | 8,53970432 | 4,57502127 | 0,00039472 | 0,00802913 |
| 5691_at | PSMB3 | -0,42989127 | 11,0965456 | -4,5740748 | 0,00039545 | 0,00803042 |
| 79989_at | TTC26 | -0,48892267 | 5,65231784 | -4,57385571 | 0,00039562 | 0,00803042 |
| 23195_at | MDN1 | -0,33716704 | 5,89301928 | -4,57101008 | 0,00039782 | 0,00806564 |
| 53344_at | CHIC1 | 0,37894086 | 5,46281905 | 4,57052906 | 0,00039819 | 0,00806564 |
| 51022_at | GLRX2 | -0,57406465 | 8,36893432 | -4,56887032 | 0,00039948 | 0,00808326 |
| 9990_at | SLC12A6 | 0,38139526 | 7,32086201 | 4,56793484 | 0,00040021 | 0,00808951 |
| 55143_at | CDCA8 | -1,21225696 | 8,58040424 | -4,56449896 | 0,0004029 | 0,00813533 |
| 387103_at | CENPW | -1,10291904 | 8,00403128 | -4,56286374 | 0,00040419 | 0,00815276 |
| 7936_at | NELFE | -0,26615565 | 9,28689183 | -4,56102373 | 0,00040564 | 0,0081735 |
| 143872_at | ARHGAP42 | 0,77237874 | 6,75256746 | 4,55985129 | 0,00040657 | 0,00818364 |
| 11004_at | KIF2C | -1,24371042 | 8,27995422 | -4,55513962 | 0,00041032 | 0,00825056 |
| 9665_at | KIAA0430 | 0,56731628 | 7,86041786 | 4,5537038 | 0,00041147 | 0,00826507 |
| 79004_at | CUEDC2 | 0,33131367 | 8,79125803 | 4,55244128 | 0,00041249 | 0,00827682 |
| 57647_at | DHX37 | -0,62219876 | 7,29592895 | -4,55124252 | 0,00041345 | 0,00828689 |
| 7371_at | UCK2 | -0,96248483 | 8,16429677 | -4,55075118 | 0,00041385 | 0,00828689 |
| 23038_at | WDTC1 | 0,3749691 | 7,12819413 | 4,54985311 | 0,00041458 | 0,00828771 |
| 22856_at | CHSY1 | -0,4739106 | 8,67076531 | -4,54963531 | 0,00041475 | 0,00828771 |
| 55852_at | TEX2 | -0,33628547 | 7,59725237 | -4,54705226 | 0,00041685 | 0,00832095 |
| 100862671_a | TMEM265 | 0,32308684 | 6,4008967 | 4,54437718 | 0,00041903 | 0,00835584 |
| 55159_at | RFWD3 | -0,64057983 | 8,22879501 | -4,54354093 | 0,00041971 | 0,00836084 |
| 29959_at | NRBP1 | 0,36328748 | 9,02218985 | 4,54221377 | 0,0004208 | 0,00837386 |
| 55621_at | TRMT1 | -0,63238787 | 7,53651537 | -4,54107979 | 0,00042173 | 0,00838375 |
| 9928_at | KIF14 | -1,1173382 | 7,12459137 | -4,54015107 | 0,0004225 | 0,00838747 |
| 9040_at | UBE2M | -0,49226865 | 9,19344225 | -4,53979636 | 0,00042279 | 0,00838747 |
| 51379_at | CRLF3 | -0,39971099 | 8,28208198 | -4,53418409 | 0,00042745 | 0,00847115 |
| 9270_at | ITGB1BP1 | -0,49071044 | 7,6705258 | -4,53169449 | 0,00042953 | 0,00850368 |
| 8318_at | CDC45 | -1,46067215 | 7,50913667 | -4,52976025 | 0,00043116 | 0,0085271 |
| 6510_at | SLC1A5 | -0,59935173 | 9,26074561 | -4,52684463 | 0,00043362 | 0,00856329 |
| 57530_at | CGN | 0,55457536 | 6,96636527 | 4,52654138 | 0,00043388 | 0,00856329 |
| 9791_at | PTDSS1 | -0,64791349 | 10,7512881 | -4,52536546 | 0,00043487 | 0,0085738 |
| 120114_at | FAT3 | 0,38479291 | 3,74457672 | 4,5248658 | 0,0004353 | 0,0085738 |
| 23594_at | ORC6 | -1,32114403 | 8,47545992 | -4,52401305 | 0,00043603 | 0,00857932 |
| 54475_at | NLE1 | -0,79781848 | 6,68887617 | -4,51993188 | 0,00043952 | 0,00863918 |
| 5425_at | POLD2 | -0,74125462 | 9,63609658 | -4,51791597 | 0,00044125 | 0,00866397 |
| 2934_at | GSN | 1,41031277 | 8,0148481 | 4,51742185 | 0,00044168 | 0,00866397 |
| 374819_at | LRRC37A3 | 1,122041 | 4,21769527 | 4,51489378 | 0,00044386 | 0,00869803 |
| 6418_at | SET | -0,35500147 | 11,2686108 | -4,51383427 | 0,00044478 | 0,0087072 |
| 10379_at | IRF9 | 1,42218031 | 8,70135742 | 4,51210852 | 0,00044629 | 0,0087236 |
| 342908_at | ZNF404 | 0,82782414 | 4,98553717 | 4,5118008 | 0,00044656 | 0,0087236 |
| 84937_at | ZNRF1 | 0,41564978 | 8,39363984 | 4,51129415 | 0,000447 | 0,0087236 |
| 124540_at | MSI2 | 0,26825547 | 7,56115125 | 4,51063199 | 0,00044758 | 0,0087236 |
| 5203_at | PFDN4 | -0,422369 | 7,08539241 | -4,51027917 | 0,00044789 | 0,0087236 |
| 5889_at | RAD51C | -0,66854082 | 7,25960829 | -4,50967844 | 0,00044841 | 0,00872503 |
| 84705_at | GTPBP3 | -0,44126014 | 6,84625692 | -4,50855342 | 0,0004494 | 0,00873542 |
| 5719_at | PSMD13 | -0,38111001 | 8,22757296 | -4,50763951 | 0,0004502 | 0,00874222 |
| 150051_at | LOC150051 | -0,80413071 | 6,31977724 | -4,50636434 | 0,00045133 | 0,00874308 |
| 169270_at | ZNF596 | 0,49510856 | 4,25816635 | 4,50614203 | 0,00045152 | 0,00874308 |

| | | | | | | |
|---|---|---|---|---|---|---|
| 4942_at | OAT | -0,33955533 | 10,4837638 | -4,50573445 | 0,00045188 | 0,00874308 |
| 55917_at | CTTNBP2NL | 0,59688535 | 7,8764231 | 4,50553141 | 0,00045206 | 0,00874308 |
| 10043_at | TOM1 | 0,4383928 | 7,21751604 | 4,50364242 | 0,00045374 | 0,00876663 |
| 55620_at | STAP2 | 0,39462358 | 8,84046746 | 4,50092952 | 0,00045615 | 0,00879513 |
| 388403_at | YPEL2 | 0,872428 | 5,45979472 | 4,50048453 | 0,00045655 | 0,00879513 |
| 3843_at | IPO5 | -0,65665638 | 8,7304291 | -4,50042751 | 0,0004566 | 0,00879513 |
| 79691_at | QTRTD1 | -0,47726651 | 7,67245563 | -4,49960568 | 0,00045733 | 0,00879513 |
| 28977_at | MRPL42 | -0,61417842 | 8,63486478 | -4,49942571 | 0,00045749 | 0,00879513 |
| 79095_at | C9orf16 | 0,53083652 | 9,13044251 | 4,4942111 | 0,00046219 | 0,00886546 |
| 55200_at | PLEKHG6 | 0,60529609 | 7,6164622 | 4,49401973 | 0,00046236 | 0,00886546 |
| 506_at | ATP5B | -0,30896974 | 12,098279 | -4,49359352 | 0,00046274 | 0,00886546 |
| 537_at | ATP6AP1 | 0,25981429 | 10,3242126 | 4,49287972 | 0,00046339 | 0,00886546 |
| 8743_at | TNFSF10 | 2,09996892 | 9,9490868 | 4,49281013 | 0,00046345 | 0,00886546 |
| 8837_at | CFLAR | 0,50451843 | 6,60938666 | 4,48671438 | 0,00046902 | 0,0089546 |
| 51726_at | DNAJB11 | -0,4934567 | 10,2642414 | -4,48668461 | 0,00046904 | 0,0089546 |
| 5885_at | RAD21 | -0,31999252 | 9,73595511 | -4,48559143 | 0,00047005 | 0,00895571 |
| 246243_at | RNASEH1 | -0,42481122 | 8,0770292 | -4,48513715 | 0,00047047 | 0,00895571 |
| 57658_at | CALCOCO1 | 0,60961587 | 6,45720464 | 4,48510457 | 0,0004705 | 0,00895571 |
| 874_at | CBR3 | -0,78466922 | 7,40011385 | -4,48450581 | 0,00047105 | 0,00895737 |
| 8825_at | LIN7A | 0,53587442 | 4,98750564 | 4,48378592 | 0,00047171 | 0,00896021 |
| 5888_at | RAD51 | -0,87247175 | 6,86809483 | -4,48330643 | 0,00047216 | 0,00896021 |
| 149175_at | MANEAL | -0,50972502 | 9,3342017 | -4,48283317 | 0,00047259 | 0,00896021 |
| 57405_at | SPC25 | -1,48108079 | 7,25982538 | -4,48054922 | 0,00047471 | 0,00899152 |
| 23596_at | OPN3 | 0,77359521 | 8,72753069 | 4,47956512 | 0,00047563 | 0,00900001 |
| 109_at | ADCY3 | -0,71119252 | 7,79796716 | -4,47770227 | 0,00047737 | 0,00902404 |
| 54892_at | NCAPG2 | -1,17768823 | 7,81576997 | -4,47675181 | 0,00047826 | 0,00903199 |
| 991_at | CDC20 | -1,49864067 | 10,4397234 | -4,47609112 | 0,00047887 | 0,00903407 |
| 9730_at | VPRBP | -0,31272312 | 6,48931814 | -4,47536443 | 0,00047956 | 0,00903407 |
| 54552_at | GNL3L | -0,32877304 | 6,68516744 | -4,47513499 | 0,00047977 | 0,00903407 |
| 284184_at | C17orf89 | -0,49992114 | 9,24911906 | -4,47401612 | 0,00048082 | 0,00904505 |
| 140809_at | SRXN1 | -0,39400898 | 10,1593556 | -4,4734515 | 0,00048136 | 0,00904622 |
| 401466_at | C8orf59 | -0,31095926 | 7,35240868 | -4,46761881 | 0,00048689 | 0,00914126 |
| 4487_at | MSX1 | -0,52358856 | 7,5404611 | -4,4661872 | 0,00048826 | 0,00915478 |
| 55835_at | CENPJ | -0,61743025 | 6,46030458 | -4,46587079 | 0,00048856 | 0,00915478 |
| 10728_at | PTGES3 | -0,33488798 | 10,4383495 | -4,46388187 | 0,00049047 | 0,0091816 |
| 11137_at | PWP1 | -0,39060006 | 9,44754831 | -4,46323921 | 0,00049109 | 0,00918425 |
| 56898_at | BDH2 | 1,00732095 | 5,79375232 | 4,46210013 | 0,00049218 | 0,00919585 |
| 51188_at | SS18L2 | -0,43924111 | 8,58998333 | -4,45889698 | 0,00049528 | 0,00924247 |
| 51118_at | UTP11L | -0,58863175 | 9,05217111 | -4,45853179 | 0,00049564 | 0,00924247 |
| 79667_at | KLF3-AS1 | 0,35756345 | 3,82206149 | 4,45734324 | 0,00049679 | 0,00925507 |
| 10567_at | RABAC1 | 0,35344498 | 9,95845151 | 4,4550439 | 0,00049904 | 0,0092879 |
| 10360_at | NPM3 | -0,78335093 | 8,07308829 | -4,45440953 | 0,00049966 | 0,00929048 |
| 2872_at | MKNK2 | 0,48599311 | 10,1789388 | 4,45265959 | 0,00050138 | 0,00931343 |
| 11041_at | B4GAT1 | 0,42210679 | 7,87619475 | 4,44736411 | 0,00050661 | 0,00940159 |
| 221037_at | JMJD1C | 0,29898554 | 7,14618772 | 4,4440292 | 0,00050994 | 0,00945419 |
| 79613_at | TANGO6 | -0,32756577 | 6,42830087 | -4,43829239 | 0,00051571 | 0,00955201 |
| 79842_at | ZBTB3 | 0,3775964 | 6,92299073 | 4,43662535 | 0,0005174 | 0,00956384 |
| 84135_at | UTP15 | -0,82606587 | 6,24351599 | -4,43656193 | 0,00051746 | 0,00956384 |
| 9706_at | ULK2 | 0,30854215 | 5,34280445 | 4,43619394 | 0,00051784 | 0,00956384 |
| 3833_at | KIFC1 | -0,88360783 | 8,06873219 | -4,4330143 | 0,00052108 | 0,0096145 |

| | | | | | | |
|---|---|---|---|---|---|---|
| 9675_at | TTI1 | -0,38152243 | 8,22010629 | -4,43251163 | 0,00052159 | 0,00961478 |
| 1839_at | HBEGF | -0,99765933 | 7,43000458 | -4,4315596 | 0,00052257 | 0,00962098 |
| 81790_at | RNF170 | 0,39758169 | 6,71565172 | 4,43120882 | 0,00052293 | 0,00962098 |
| 2261_at | FGFR3 | -0,97595866 | 8,48330347 | -4,42888532 | 0,00052532 | 0,00965182 |
| 1286_at | COL4A4 | 1,36620281 | 6,25189709 | 4,42860697 | 0,00052561 | 0,00965182 |
| 1106_at | CHD2 | 0,47813566 | 7,77134376 | 4,4265214 | 0,00052776 | 0,00967638 |
| 51635_at | DHRS7 | 0,691602 | 9,27365533 | 4,42634254 | 0,00052795 | 0,00967638 |
| 9184_at | BUB3 | -0,48091158 | 9,89402891 | -4,42504975 | 0,00052929 | 0,00969175 |
| 55215_at | FANCI | -1,29431152 | 8,2843814 | -4,42191988 | 0,00053255 | 0,00973884 |
| 10809_at | STARD10 | 0,32377319 | 8,79699971 | 4,42161441 | 0,00053287 | 0,00973884 |
| 54832_at | VPS13C | 0,52839697 | 7,02087094 | 4,42090104 | 0,00053362 | 0,00974325 |
| 51388_at | NIP7 | -0,76646225 | 8,75731026 | -4,41895664 | 0,00053566 | 0,00977127 |
| 27244_at | SESN1 | 0,71902275 | 7,70958371 | 4,41799844 | 0,00053667 | 0,00978042 |
| 123720_at | WHAMM | 0,31011018 | 8,22680774 | 4,4142074 | 0,00054068 | 0,00984422 |
| 83461_at | CDCA3 | -1,1012923 | 8,19069433 | -4,41239532 | 0,00054261 | 0,00986757 |
| 7266_at | DNAJC7 | -0,25897443 | 9,13146173 | -4,41196042 | 0,00054308 | 0,00986757 |
| 4998_at | ORC1 | -1,79334005 | 6,60105692 | -4,41156045 | 0,0005435 | 0,00986757 |
| 114791_at | TUBGCP5 | -0,33289723 | 6,33197299 | -4,41064352 | 0,00054448 | 0,00987606 |
| 23673_at | STX12 | 0,43906411 | 8,86926666 | 4,40984895 | 0,00054533 | 0,00988219 |
| 8503_at | PIK3R3 | 0,81563044 | 7,1476881 | 4,40423535 | 0,00055138 | 0,00997635 |
| 51095_at | TRNT1 | -0,43745133 | 7,3680533 | -4,40406745 | 0,00055157 | 0,00997635 |
| 169166_at | SNX31 | 1,61572045 | 5,29592112 | 4,40310672 | 0,00055261 | 0,00998584 |
| 55093_at | WDYHV1 | -0,50206661 | 7,12308802 | -4,40083917 | 0,00055508 | 0,01002105 |
| 55127_at | HEATR1 | -0,60606776 | 7,75066376 | -4,38858779 | 0,00056861 | 0,0102521 |
| 2177_at | FANCD2 | -1,00425632 | 7,43851669 | -4,38829686 | 0,00056894 | 0,0102521 |
| 515_at | ATP5F1 | -0,3122143 | 10,4407962 | -4,38739739 | 0,00056995 | 0,01026066 |
| 5832_at | ALDH18A1 | -0,43977359 | 8,96801277 | -4,38610239 | 0,0005714 | 0,01027723 |
| 10444_at | ZER1 | 0,2854572 | 7,01143166 | 4,38497614 | 0,00057267 | 0,01029042 |
| 55603_at | FAM46A | 0,82739438 | 7,91583438 | 4,38259561 | 0,00057536 | 0,01032568 |
| 83886_at | PRSS27 | 0,64414045 | 6,49946283 | 4,38229053 | 0,0005757 | 0,01032568 |
| 10062_at | NR1H3 | -0,34353424 | 7,15033482 | -4,38058679 | 0,00057764 | 0,0103346 |
| 283209_at | PGM2L1 | 0,81738881 | 7,25820091 | 4,38053497 | 0,0005777 | 0,0103346 |
| 10189_at | ALYREF | -0,83893554 | 10,6555078 | -4,38043419 | 0,00057781 | 0,0103346 |
| 63875_at | MRPL17 | -0,49537225 | 8,71742793 | -4,37983318 | 0,00057849 | 0,01033724 |
| 51018_at | RRP15 | -0,90857675 | 6,85025641 | -4,37565379 | 0,00058327 | 0,01040456 |
| 6936_at | GCFC2 | -0,53551499 | 6,55970408 | -4,37558466 | 0,00058335 | 0,01040456 |
| 64782_at | AEN | -0,52810499 | 7,25120977 | -4,37512357 | 0,00058388 | 0,01040456 |
| 64395_at | GMCL1 | 0,29647741 | 8,99225012 | 4,37313489 | 0,00058617 | 0,01043572 |
| 25824_at | PRDX5 | 0,3946021 | 11,2485195 | 4,37171586 | 0,00058781 | 0,01045525 |
| 64969_at | MRPS5 | -0,49297508 | 8,47682752 | -4,36853129 | 0,00059151 | 0,01051131 |
| 89944_at | GLB1L2 | -0,65110741 | 6,82346398 | -4,36632535 | 0,00059409 | 0,01054734 |
| 79269_at | DCAF10 | 0,32541356 | 7,71945414 | 4,36403065 | 0,00059678 | 0,0105798 |
| 79836_at | LONRF3 | -0,72750749 | 6,07962114 | -4,36383028 | 0,00059701 | 0,0105798 |
| 7272_at | TTK | -1,1964475 | 7,72584688 | -4,36334461 | 0,00059758 | 0,01058019 |
| 91057_at | CCDC34 | -0,76846577 | 6,02584339 | -4,36126391 | 0,00060004 | 0,01061389 |
| 84458_at | LCOR | 0,58798798 | 8,70799606 | 4,35721263 | 0,00060485 | 0,01068913 |
| 7965_at | AIMP2 | -0,52491021 | 9,59136501 | -4,35455971 | 0,00060802 | 0,0107353 |
| 2305_at | FOXM1 | -1,3473484 | 8,62666449 | -4,35303682 | 0,00060984 | 0,01075054 |
| 23265_at | EXOC7 | 0,30937587 | 8,52380046 | 4,35291065 | 0,00061 | 0,01075054 |
| 6390_at | SDHB | -0,49732602 | 10,0735952 | -4,35178148 | 0,00061135 | 0,01076464 |

| | | | | | | |
|---|---|---|---|---|---|---|
| 1786_at | DNMT1 | -0,71597284 | 9,98091277 | -4,35052044 | 0,00061288 | 0,01078157 |
| 1468_at | SLC25A10 | -0,65711012 | 7,65862071 | -4,34944555 | 0,00061418 | 0,01079458 |
| 138241_at | C9orf85 | 0,44446397 | 7,56352561 | 4,3471763 | 0,00061693 | 0,01083309 |
| 990_at | CDC6 | -1,78605658 | 8,21467878 | -4,34580277 | 0,0006186 | 0,01084551 |
| 151242_at | PPP1R1C | 0,23555641 | 4,47064601 | 4,34567068 | 0,00061876 | 0,01084551 |
| 90594_at | ZNF439 | 0,67068212 | 5,073711 | 4,34392116 | 0,0006209 | 0,01087179 |
| 9531_at | BAG3 | -0,50291677 | 9,84818095 | -4,34352116 | 0,00062139 | 0,01087179 |
| 23244_at | PDS5A | -0,29585417 | 8,82015705 | -4,342162 | 0,00062306 | 0,01089043 |
| 27175_at | TUBG2 | -0,33540208 | 7,509094 | -4,34173161 | 0,00062359 | 0,01089043 |
| 27079_at | RPUSD2 | -0,29723293 | 7,03616116 | -4,34074229 | 0,0006248 | 0,01089361 |
| 7476_at | WNT7A | -0,59371319 | 5,81874032 | -4,34063593 | 0,00062494 | 0,01089361 |
| 25788_at | RAD54B | -1,2606327 | 6,65502747 | -4,34020613 | 0,00062547 | 0,01089361 |
| 84365_at | NIFK | -0,57576322 | 8,91693063 | -4,33960663 | 0,0006262 | 0,01089664 |
| 1666_at | DECR1 | 0,49852102 | 8,98521517 | 4,33635646 | 0,00063023 | 0,01095681 |
| 10116_at | FEM1B | 0,25226278 | 8,4029166 | 4,33582839 | 0,00063089 | 0,01095834 |
| 54813_at | KLHL28 | 0,69587371 | 7,34438005 | 4,33520868 | 0,00063166 | 0,01096186 |
| 81603_at | TRIM8 | 0,33684427 | 7,32189051 | 4,33368082 | 0,00063357 | 0,01096533 |
| 7321_at | UBE2D1 | 0,63817855 | 9,01846802 | 4,33356353 | 0,00063371 | 0,01096533 |
| 51700_at | CYB5R2 | 0,8781496 | 5,34425884 | 4,33340607 | 0,00063391 | 0,01096533 |
| 6421_at | SFPQ | -0,35339668 | 9,31206568 | -4,33292532 | 0,00063451 | 0,01096533 |
| 5163_at | PDK1 | -0,6252816 | 6,88318604 | -4,33276977 | 0,00063471 | 0,01096533 |
| 10535_at | RNASEH2A | -0,95500625 | 9,02538853 | -4,33164311 | 0,00063612 | 0,01097438 |
| 100287616_a | LOXL1-AS1 | -1,36363382 | 6,37094289 | -4,33105066 | 0,00063686 | 0,01097438 |
| 3213_at | HOXB3 | 0,39073213 | 7,53538159 | 4,33098952 | 0,00063694 | 0,01097438 |
| 410_at | ARSA | 0,30750931 | 6,48051575 | 4,32986138 | 0,00063836 | 0,010989 |
| 152100_at | CMC1 | -0,36428418 | 7,43187378 | -4,32869361 | 0,00063983 | 0,01100451 |
| 90634_at | N4BP2L1 | 0,56824918 | 4,85554141 | 4,32739549 | 0,00064147 | 0,01102289 |
| 65986_at | ZBTB10 | 0,56266843 | 5,85415777 | 4,32604124 | 0,00064319 | 0,01104254 |
| 5984_at | RFC4 | -1,1941709 | 9,1429621 | -4,32456839 | 0,00064506 | 0,01106482 |
| 58508_at | KMT2C | 0,39014377 | 8,06274548 | 4,32398525 | 0,0006458 | 0,01106536 |
| 10226_at | PLIN3 | 0,3345306 | 9,22887751 | 4,32364162 | 0,00064624 | 0,01106536 |
| 3615_at | IMPDH2 | -0,92536591 | 10,6038326 | -4,32227611 | 0,00064798 | 0,01108045 |
| 64981_at | MRPL34 | -0,30087679 | 8,9687263 | -4,32205115 | 0,00064827 | 0,01108045 |
| 10523_at | CHERP | -0,30526495 | 7,68717615 | -4,32154613 | 0,00064892 | 0,01108167 |
| 112616_at | CMTM7 | -0,61462172 | 8,7554925 | -4,31950004 | 0,00065154 | 0,01111666 |
| 22921_at | MSRB2 | 0,29312055 | 6,39417215 | 4,31652163 | 0,00065539 | 0,01117231 |
| 29126_at | CD274 | 0,67046911 | 5,40970424 | 4,31454689 | 0,00065795 | 0,01120604 |
| 6749_at | SSRP1 | -0,67365994 | 9,95968821 | -4,31356811 | 0,00065922 | 0,01121779 |
| 26100_at | WIPI2 | 0,29272889 | 7,77981386 | 4,31293427 | 0,00066004 | 0,01122193 |
| 55196_at | KIAA1551 | 0,45880012 | 7,16922538 | 4,31035565 | 0,00066341 | 0,01126927 |
| 282809_at | POC1B | -0,35048589 | 7,25251559 | -4,3092206 | 0,0006649 | 0,0112846 |
| 3276_at | PRMT1 | -0,38500542 | 6,46331986 | -4,3086061 | 0,00066571 | 0,01128836 |
| 3482_at | IGF2R | 0,40449901 | 9,00109008 | 4,30689573 | 0,00066796 | 0,01130258 |
| 51805_at | COQ3 | -0,71258678 | 7,75725092 | -4,30666441 | 0,00066827 | 0,01130258 |
| 9271_at | PIWIL1 | -0,50936053 | 3,76866917 | -4,30663345 | 0,00066831 | 0,01130258 |
| 6240_at | RRM1 | -0,90795721 | 9,38943482 | -4,30565107 | 0,00066961 | 0,01131459 |
| 29760_at | BLNK | 1,84988755 | 6,2062278 | 4,30443098 | 0,00067122 | 0,01133195 |
| 9156_at | EXO1 | -1,33660626 | 7,01508346 | -4,30159807 | 0,00067499 | 0,01138557 |
| 91419_at | XRCC6BP1 | -0,37246324 | 8,27171882 | -4,29820308 | 0,00067953 | 0,01145216 |
| 23329_at | TBC1D30 | -0,33742728 | 5,6280343 | -4,29719829 | 0,00068088 | 0,01145567 |

| | | | | | | |
|---|---|---|---|---|---|---|
| 51021_at | MRPS16 | -0,38835815 | 9,09730725 | -4,29716336 | 0,00068093 | 0,01145567 |
| 10531_at | PITRM1 | -0,52997806 | 9,08953518 | -4,29508764 | 0,00068373 | 0,01148581 |
| 3985_at | LIMK2 | 0,49096559 | 8,9933954 | 4,29495028 | 0,00068391 | 0,01148581 |
| 463_at | ZFHX3 | 0,68003911 | 7,17174072 | 4,29222961 | 0,0006876 | 0,01153767 |
| 2186_at | BPTF | 0,31400958 | 8,5990539 | 4,29088385 | 0,00068943 | 0,01155833 |
| 29994_at | BAZ2B | 0,76656181 | 7,9936991 | 4,29032575 | 0,00069019 | 0,01156103 |
| 11005_at | SPINK5 | -0,88535934 | 4,60544 | -4,28823543 | 0,00069305 | 0,01159881 |
| 4524_at | MTHFR | 0,32329442 | 6,25860563 | 4,28522074 | 0,00069719 | 0,0116448 |
| 6634_at | SNRPD3 | -0,35182366 | 10,6409784 | -4,284845 | 0,00069771 | 0,0116448 |
| 4521_at | NUDT1 | -0,72765732 | 7,87482869 | -4,28439285 | 0,00069834 | 0,0116448 |
| 79639_at | TMEM53 | 0,40211577 | 7,74655636 | 4,28410506 | 0,00069873 | 0,0116448 |
| 394_at | ARHGAP5 | 0,28156529 | 8,20782176 | 4,28404094 | 0,00069882 | 0,0116448 |
| 79174_at | CRELD2 | -0,30058687 | 8,92388726 | -4,28357404 | 0,00069947 | 0,01164548 |
| 5289_at | PIK3C3 | 0,48720894 | 5,40652436 | 4,28223104 | 0,00070133 | 0,01166636 |
| 9088_at | PKMYT1 | -0,63527898 | 7,67879577 | -4,2802278 | 0,00070411 | 0,01169594 |
| 51603_at | METTL13 | -0,37253432 | 8,03075502 | -4,28007803 | 0,00070432 | 0,01169594 |
| 9055_at | PRC1 | -1,15406356 | 9,89004693 | -4,27924593 | 0,00070548 | 0,01170511 |
| 221935_at | SDK1 | 0,54228809 | 6,41883724 | 4,27743004 | 0,00070802 | 0,01173711 |
| 1891_at | ECH1 | 0,44959213 | 10,2691229 | 4,2757778 | 0,00071033 | 0,01176346 |
| 55733_at | HHAT | 0,82331522 | 6,4555422 | 4,27540702 | 0,00071085 | 0,01176346 |
| 3913_at | LAMB2 | 0,84542007 | 7,46744443 | 4,27499238 | 0,00071144 | 0,01176346 |
| 29899_at | GPSM2 | -0,7955082 | 8,06785764 | -4,27170905 | 0,00071607 | 0,01182996 |
| 166785_at | MMAA | 0,3754979 | 4,7413809 | 4,26865001 | 0,00072042 | 0,01189157 |
| 8082_at | SSPN | 0,53526238 | 5,48670055 | 4,26662937 | 0,00072331 | 0,01192898 |
| 10248_at | POP7 | -0,53795912 | 8,76806477 | -4,26536884 | 0,00072511 | 0,01194854 |
| 10556_at | RPP30 | -0,41744972 | 6,16901916 | -4,26408905 | 0,00072695 | 0,0119686 |
| 4128_at | MAOA | 0,77750874 | 9,17068327 | 4,26339614 | 0,00072795 | 0,01197479 |
| 4141_at | MARS | -0,38445389 | 9,48238061 | -4,26253393 | 0,00072919 | 0,01198501 |
| 30836_at | DNTTIP2 | -0,48809007 | 9,68260038 | -4,26154791 | 0,00073062 | 0,01199818 |
| 8562_at | DENR | -0,37935504 | 9,73598188 | -4,26078155 | 0,00073173 | 0,01200498 |
| 1523_at | CUX1 | 0,38025608 | 7,91971909 | 4,26028765 | 0,00073244 | 0,01200498 |
| 92399_at | MRRF | -0,32713758 | 6,75773629 | -4,25997117 | 0,0007329 | 0,01200498 |
| 8495_at | PPFIBP2 | 0,64661978 | 7,80520967 | 4,25941785 | 0,0007337 | 0,01200757 |
| 9895_at | TECPR2 | 0,35416491 | 7,07315964 | 4,25900353 | 0,00073431 | 0,01200757 |
| 2801_at | GOLGA2 | 0,26785236 | 7,67105533 | 4,25428146 | 0,0007412 | 0,01210064 |
| 2597_at | GAPDH | -0,22121245 | 12,4229141 | -4,25424643 | 0,00074125 | 0,01210064 |
| 10849_at | CD3EAP | -0,45209426 | 7,57718874 | -4,25132257 | 0,00074556 | 0,01215311 |
| 7249_at | TSC2 | 0,31545098 | 7,62165545 | 4,25120545 | 0,00074573 | 0,01215311 |
| 9120_at | SLC16A6 | -0,46129623 | 4,38848733 | -4,25066514 | 0,00074653 | 0,01215584 |
| 1351_at | COX8A | -0,29823258 | 11,6572823 | -4,24989884 | 0,00074766 | 0,01216402 |
| 57697_at | FANCM | -0,57531621 | 5,11848417 | -4,24776831 | 0,00075082 | 0,0122042 |
| 23357_at | ANGEL1 | -0,2617229 | 7,58650329 | -4,24715037 | 0,00075174 | 0,0122042 |
| 10198_at | MPHOSPH9 | -0,55292918 | 6,39143492 | -4,24695566 | 0,00075203 | 0,0122042 |
| 29088_at | MRPL15 | -0,57976561 | 9,54354659 | -4,24398539 | 0,00075647 | 0,01225374 |
| 5427_at | POLE2 | -1,49581024 | 7,53696647 | -4,24390684 | 0,00075659 | 0,01225374 |
| 7626_at | ZNF75D | 0,42068407 | 5,62150642 | 4,24363544 | 0,00075699 | 0,01225374 |
| 7248_at | TSC1 | 0,45237016 | 8,46794722 | 4,23873074 | 0,00076438 | 0,01236299 |
| 9205_at | ZMYM5 | 0,30905577 | 5,44605519 | 4,23788993 | 0,00076566 | 0,01237321 |
| 283349_at | RASSF3 | -0,49362272 | 7,40606518 | -4,23694393 | 0,00076709 | 0,01238051 |
| 306_at | ANXA3 | 0,90895649 | 7,97037581 | 4,23646571 | 0,00076782 | 0,01238051 |

| | | | | | | |
|---|---|---|---|---|---|---|
| 9837_at | GINS1 | -1,10372885 | 9,50440075 | -4,23632362 | 0,00076804 | 0,01238051 |
| 2395_at | FXN | -0,73699695 | 6,15374836 | -4,23512452 | 0,00076986 | 0,0123899 |
| 55632_at | G2E3 | -0,34958674 | 7,30995923 | -4,23472146 | 0,00077048 | 0,0123899 |
| 121642_at | ALKBH2 | -0,429801 | 7,08241638 | -4,23429908 | 0,00077112 | 0,0123899 |
| 11065_at | UBE2C | -1,08870817 | 10,0610705 | -4,23425455 | 0,00077119 | 0,0123899 |
| 93622_at | LOC93622 | -0,40759384 | 8,6931652 | -4,23378985 | 0,0007719 | 0,01239098 |
| 101928000_a | LOC10192800 | -0,25880075 | 4,59564612 | -4,23185009 | 0,00077488 | 0,01242835 |
| 7402_at | UTRN | 0,41432749 | 6,14061302 | 4,23102602 | 0,00077614 | 0,0124383 |
| 79622_at | SNRNP25 | -0,79109614 | 9,82295938 | -4,22990672 | 0,00077787 | 0,01245556 |
| 29063_at | ZCCHC4 | -0,27739928 | 5,70109655 | -4,2287409 | 0,00077967 | 0,01247401 |
| 5863_at | RGL2 | 0,43068692 | 8,46940262 | 4,22671142 | 0,00078281 | 0,01251283 |
| 23286_at | WWC1 | 0,42928178 | 7,40616584 | 4,22616479 | 0,00078366 | 0,01251283 |
| 22976_at | PAXIP1 | -0,27304394 | 8,5336076 | -4,22591771 | 0,00078404 | 0,01251283 |
| 4891_at | SLC11A2 | 0,49497214 | 8,11775862 | 4,2248841 | 0,00078565 | 0,01252811 |
| 9141_at | PDCD5 | -0,6166407 | 8,50513403 | -4,22373858 | 0,00078744 | 0,01254621 |
| 84262_at | PSMG3 | -0,54635549 | 9,21935571 | -4,22218665 | 0,00078986 | 0,01257447 |
| 8848_at | TSC22D1 | 0,31647122 | 5,39353585 | 4,21621859 | 0,00079926 | 0,01268891 |
| 9787_at | DLGAP5 | -1,16250683 | 9,33080281 | -4,21590486 | 0,00079976 | 0,01268891 |
| 10628_at | TXNIP | 0,66187018 | 10,5661296 | 4,21573061 | 0,00080004 | 0,01268891 |
| 7696_at | ZNF137P | 0,4452931 | 5,13688096 | 4,21561687 | 0,00080022 | 0,01268891 |
| 8131_at | NPRL3 | -0,29423255 | 7,7184352 | -4,21553805 | 0,00080034 | 0,01268891 |
| 79000_at | AUNIP | -1,01160908 | 6,46740594 | -4,21468517 | 0,0008017 | 0,01269993 |
| 54464_at | XRN1 | 0,59755956 | 6,6789523 | 4,21223047 | 0,00080561 | 0,01275143 |
| 80746_at | TSEN2 | -0,67569729 | 7,7058302 | -4,211531 | 0,00080673 | 0,01275865 |
| 11094_at | CACFD1 | 0,38378618 | 7,51486538 | 4,21011225 | 0,000809 | 0,01277961 |
| 93343_at | MVB12A | 0,24967948 | 8,35894581 | 4,20987613 | 0,00080938 | 0,01277961 |
| 1743_at | DLST | -0,2877702 | 9,37414057 | -4,20908237 | 0,00081066 | 0,01278549 |
| 2066_at | ERBB4 | 0,3508235 | 4,39636294 | 4,20881877 | 0,00081108 | 0,01278549 |
| 200916_at | RPL22L1 | -0,73255393 | 7,77987422 | -4,20774461 | 0,00081281 | 0,01280228 |
| 4830_at | NME1 | -1,08722083 | 11,0540482 | -4,20720149 | 0,00081369 | 0,01280421 |
| 23640_at | HSPBP1 | -0,53956365 | 8,52107735 | -4,20684507 | 0,00081426 | 0,01280421 |
| 5725_at | PTBP1 | -0,34825874 | 11,3872226 | -4,20572735 | 0,00081607 | 0,01282217 |
| 2010_at | EMD | -0,25318524 | 8,87907364 | -4,20415839 | 0,00081862 | 0,01283282 |
| 54625_at | PARP14 | 1,07428974 | 8,52603099 | 4,20374667 | 0,00081928 | 0,01283282 |
| 79027_at | ZNF655 | 0,40629252 | 7,6137651 | 4,20354248 | 0,00081962 | 0,01283282 |
| 10714_at | POLD3 | -0,42307667 | 7,81928035 | -4,202915 | 0,00082064 | 0,01283282 |
| 8399_at | PLA2G10 | 1,58703008 | 6,81352134 | 4,20258262 | 0,00082118 | 0,01283282 |
| 126295_at | ZNF57 | -0,39589181 | 6,94120267 | -4,20241276 | 0,00082146 | 0,01283282 |
| 23258_at | DENND5A | 0,25621082 | 7,96233519 | 4,20219147 | 0,00082182 | 0,01283282 |
| 10376_at | TUBA1B | -0,70390839 | 11,2518579 | -4,20203091 | 0,00082208 | 0,01283282 |
| 2175_at | FANCA | -0,65023249 | 6,63343794 | -4,20101251 | 0,00082374 | 0,01284715 |
| 7259_at | TSPYL1 | 0,28405715 | 7,33996975 | 4,20065213 | 0,00082433 | 0,01284715 |
| 2635_at | GBP3 | 0,81036481 | 9,6411411 | 4,19904684 | 0,00082696 | 0,01287773 |
| 55038_at | CDCA4 | -0,65623516 | 8,9474926 | -4,19601194 | 0,00083196 | 0,01294506 |
| 115_at | ADCY9 | 0,45333762 | 6,66849575 | 4,19552148 | 0,00083277 | 0,01294721 |
| 11104_at | KATNA1 | -0,31188495 | 7,84298614 | -4,193952 | 0,00083537 | 0,01297714 |
| 84317_at | CCDC115 | 0,19066303 | 8,377751 | 4,19273375 | 0,00083739 | 0,01299455 |
| 10346_at | TRIM22 | 2,61718246 | 7,5453773 | 4,19246458 | 0,00083784 | 0,01299455 |
| 6233_at | RPS27A | -0,42590334 | 6,4697354 | -4,18984938 | 0,0008422 | 0,01305168 |
| 10153_at | CEBPZ | -0,56284088 | 8,6173776 | -4,18867112 | 0,00084417 | 0,01307173 |

| | | | | | | |
|---|---|---|---|---|---|---|
| 5801_at | PTPRR | 0,86347311 | 6,52390317 | 4,18528085 | 0,00084987 | 0,01314946 |
| 7353_at | UFD1L | -0,40763609 | 9,52452488 | -4,18382285 | 0,00085234 | 0,01316289 |
| 222236_at | NAPEPLD | 0,33484036 | 6,51546917 | 4,18374944 | 0,00085246 | 0,01316289 |
| 3554_at | IL1R1 | 0,80705465 | 4,96987806 | 4,18344567 | 0,00085297 | 0,01316289 |
| 27341_at | RRP7A | -0,44905211 | 7,86651647 | -4,18315114 | 0,00085347 | 0,01316289 |
| 93081_at | TEX30 | -1,07591952 | 6,8015689 | -4,181736 | 0,00085588 | 0,01318938 |
| 51701_at | NLK | 0,43325346 | 7,55322503 | 4,18102292 | 0,00085709 | 0,01319751 |
| 53_at | ACP2 | 0,36246047 | 8,66055465 | 4,18010685 | 0,00085865 | 0,01319924 |
| 51196_at | PLCE1 | 0,67254643 | 8,09103039 | 4,17991029 | 0,00085898 | 0,01319924 |
| 81844_at | TRIM56 | 0,45024122 | 8,04485707 | 4,17975112 | 0,00085926 | 0,01319924 |
| 26298_at | EHF | 0,40877191 | 8,9207691 | 4,17869097 | 0,00086107 | 0,01321652 |
| 6182_at | MRPL12 | -0,68700539 | 9,94793651 | -4,17811612 | 0,00086205 | 0,01322109 |
| 96459_at | FNIP1 | 0,3373247 | 6,55977924 | 4,17761106 | 0,00086292 | 0,01322383 |
| 64866_at | CDCP1 | -0,40250646 | 7,33511629 | -4,17606752 | 0,00086557 | 0,01325389 |
| 79726_at | WDR59 | 0,24115315 | 7,24253179 | 4,17509858 | 0,00086723 | 0,01326888 |
| 80351_at | TNKS2 | 0,51877823 | 8,37106035 | 4,17468248 | 0,00086795 | 0,01326932 |
| 9187_at | SLC24A1 | 0,45596716 | 5,62322361 | 4,17355076 | 0,0008699 | 0,01328864 |
| 100506753_a | AZIN1-AS1 | 0,69891432 | 5,0153038 | 4,17306615 | 0,00087074 | 0,0132909 |
| 3838_at | KPNA2 | -0,47156559 | 11,9950287 | -4,17155892 | 0,00087335 | 0,0133202 |
| 2491_at | CENPI | -1,02254461 | 6,32548329 | -4,17088179 | 0,00087453 | 0,01332758 |
| 284098_at | PIGW | -0,73345265 | 8,07425555 | -4,16978631 | 0,00087644 | 0,01334607 |
| 11046_at | SLC35D2 | 0,64614411 | 6,51615785 | 4,16929136 | 0,0008773 | 0,01334865 |
| 51136_at | RNFT1 | 0,56973861 | 7,89977093 | 4,16572829 | 0,00088353 | 0,01341398 |
| 284129_at | SLC26A11 | 0,61667226 | 7,26412962 | 4,16565388 | 0,00088366 | 0,01341398 |
| 2821_at | GPI | -0,6235458 | 11,3566483 | -4,16535436 | 0,00088419 | 0,01341398 |
| 9975_at | NR1D2 | 0,60568344 | 8,40972923 | 4,16524688 | 0,00088438 | 0,01341398 |
| 8487_at | GEMIN2 | -0,55963255 | 6,87279002 | -4,16294425 | 0,00088843 | 0,01346491 |
| 83540_at | NUF2 | -1,21574647 | 7,66420588 | -4,16163073 | 0,00089076 | 0,01348049 |
| 8192_at | CLPP | -0,35481528 | 8,99446467 | -4,16157151 | 0,00089086 | 0,01348049 |
| 10615_at | SPAG5 | -1,13856491 | 8,3332049 | -4,15997563 | 0,00089369 | 0,0135127 |
| 6015_at | RING1 | 0,21123967 | 8,73641339 | 4,15782525 | 0,00089752 | 0,01355659 |
| 39_at | ACAT2 | -0,7926152 | 9,36749905 | -4,15755552 | 0,000898 | 0,01355659 |
| 8802_at | SUCLG1 | -0,41796538 | 10,2096264 | -4,15597957 | 0,00090082 | 0,01358848 |
| 10195_at | ALG3 | -0,30801915 | 10,0710806 | -4,15482993 | 0,00090288 | 0,01360892 |
| 63922_at | CHTF18 | -0,82463816 | 7,28982346 | -4,15169393 | 0,00090853 | 0,01367839 |
| 5934_at | RBL2 | 0,42118627 | 7,54976657 | 4,1514825 | 0,00090891 | 0,01367839 |
| 90871_at | TMEM261 | -0,36494205 | 7,58092764 | -4,14883977 | 0,0009137 | 0,01373515 |
| 57704_at | GBA2 | 0,31782388 | 7,45756923 | 4,14861505 | 0,00091411 | 0,01373515 |
| 150962_at | PUS10 | 0,98674784 | 6,67834105 | 4,14821698 | 0,00091483 | 0,01373532 |
| 23404_at | EXOSC2 | -0,83127215 | 8,61649392 | -4,14525965 | 0,00092023 | 0,01380558 |
| 9658_at | ZNF516 | 0,35951717 | 5,9237319 | 4,14401415 | 0,00092251 | 0,01382906 |
| 10898_at | CPSF4 | -0,40304127 | 8,55138335 | -4,1419467 | 0,00092631 | 0,01387525 |
| 78995_at | C17orf53 | -0,46982918 | 6,42435358 | -4,13944878 | 0,00093093 | 0,01392804 |
| 11230_at | PRAF2 | 0,4001815 | 7,68107279 | 4,13904683 | 0,00093167 | 0,01392804 |
| 7768_at | ZNF225 | 0,42429738 | 4,6689317 | 4,13874884 | 0,00093222 | 0,01392804 |
| 118932_at | ANKRD22 | -0,75712487 | 6,98060903 | -4,13847693 | 0,00093273 | 0,01392804 |
| 22995_at | CEP152 | -0,67561045 | 5,9148732 | -4,13559295 | 0,0009381 | 0,01399734 |
| 6229_at | RPS24 | -0,25019979 | 11,5150829 | -4,13136936 | 0,00094601 | 0,01410455 |
| 26040_at | SETBP1 | 0,52494002 | 5,92711927 | 4,12973818 | 0,00094909 | 0,01413947 |
| 23334_at | SZT2 | 0,34075253 | 6,25721006 | 4,12892308 | 0,00095063 | 0,01414082 |

| | | | | | | |
|---|---|---|---|---|---|---|
| 4477_at | MSMB | 0,61600754 | 3,35487666 | 4,12875417 | 0,00095095 | 0,01414082 |
| 6659_at | SOX4 | 0,75475158 | 9,70993884 | 4,12852599 | 0,00095138 | 0,01414082 |
| 10509_at | SEMA4B | -0,52311181 | 9,63500874 | -4,12652754 | 0,00095518 | 0,01418321 |
| 57175_at | CORO1B | 0,31129484 | 9,74831212 | 4,12624721 | 0,00095571 | 0,01418321 |
| 85395_at | FAM207A | -0,66451057 | 7,23516014 | -4,12425861 | 0,0009595 | 0,01422851 |
| 8195_at | MKKS | -0,35370175 | 7,99407963 | -4,12181753 | 0,00096417 | 0,01428682 |
| 83658_at | DYNLRB1 | 0,29099319 | 10,9673863 | 4,12030074 | 0,00096709 | 0,01429647 |
| 56652_at | C10orf2 | -0,67931887 | 7,08201333 | -4,12025654 | 0,00096717 | 0,01429647 |
| 80178_at | C16orf59 | -0,64011507 | 7,29075887 | -4,11998742 | 0,00096769 | 0,01429647 |
| 480_at | ATP1A4 | 0,49374132 | 4,83914654 | 4,11993553 | 0,00096779 | 0,01429647 |
| 5966_at | REL | 0,75649421 | 8,20548304 | 4,11847192 | 0,00097062 | 0,0143272 |
| 389072_at | PLEKHM3 | 0,67719712 | 6,16599054 | 4,11431159 | 0,00097869 | 0,01442447 |
| 10610_at | ST6GALNAC2 | 0,87278884 | 7,92608866 | 4,11430517 | 0,0009787 | 0,01442447 |
| 51184_at | GPN3 | -0,77013906 | 8,11027837 | -4,11364798 | 0,00097999 | 0,01443232 |
| 23203_at | PMPCA | -0,32482741 | 8,2494361 | -4,11068601 | 0,00098578 | 0,01450662 |
| 3032_at | HADHB | 0,46458724 | 10,1385487 | 4,10962461 | 0,00098787 | 0,01452622 |
| 51654_at | CDK5RAP1 | -0,40695397 | 7,79243722 | -4,10922145 | 0,00098866 | 0,0145268 |
| 8504_at | PEX3 | -0,42690691 | 7,65818859 | -4,10838875 | 0,00099031 | 0,01453121 |
| 10426_at | TUBGCP3 | -0,59820177 | 6,69424419 | -4,10830398 | 0,00099047 | 0,01453121 |
| 8613_at | PPAP2B | 0,59745669 | 9,54346513 | 4,10692446 | 0,0009932 | 0,0145601 |
| 93436_at | ARMC6 | -0,27066821 | 7,39077189 | -4,10615929 | 0,00099471 | 0,01457123 |
| 27324_at | TOX3 | 1,19784163 | 5,43771816 | 4,10516544 | 0,00099668 | 0,0145827 |
| 56164_at | STK31 | 0,45532876 | 3,49220259 | 4,10500116 | 0,00099701 | 0,0145827 |
| 23339_at | VPS39 | 0,29314829 | 7,69265287 | 4,10325406 | 0,00100049 | 0,01461855 |
| 51728_at | POLR3K | -0,83451512 | 9,50849996 | -4,10268719 | 0,00100162 | 0,01461855 |
| 3692_at | EIF6 | -0,36283159 | 11,141126 | -4,10262693 | 0,00100174 | 0,01461855 |
| 23350_at | U2SURP | -0,48703838 | 8,71127622 | -4,1009629 | 0,00100506 | 0,01465599 |
| 23076_at | RRP1B | -0,50560407 | 7,84476618 | -4,09932862 | 0,00100834 | 0,01468515 |
| 23461_at | ABCA5 | 0,85897844 | 5,40732491 | 4,09901803 | 0,00100897 | 0,01468515 |
| 28983_at | TMPRSS11E | 1,68364538 | 5,71474922 | 4,0978473 | 0,00101132 | 0,01468515 |
| 192111_at | PGAM5 | -0,58983694 | 7,77443767 | -4,09782577 | 0,00101137 | 0,01468515 |
| 51762_at | RAB8B | 0,48586095 | 7,25135468 | 4,09760042 | 0,00101182 | 0,01468515 |
| 118987_at | PDZD8 | -0,2493095 | 7,27345131 | -4,09706213 | 0,00101291 | 0,01468515 |
| 199953_at | TMEM201 | -0,4744732 | 7,50229758 | -4,09693697 | 0,00101316 | 0,01468515 |
| 80823_at | BHLHB9 | 0,71690394 | 6,26425447 | 4,0969356 | 0,00101316 | 0,01468515 |
| 79832_at | QSER1 | -0,43528364 | 7,64511188 | -4,09478215 | 0,00101752 | 0,01473722 |
| 7762_at | ZNF215 | -0,37308492 | 5,60297816 | -4,09383153 | 0,00101945 | 0,01475407 |
| 84708_at | LNX1 | 0,89921502 | 6,9024812 | 4,09308125 | 0,00102098 | 0,01476506 |
| 9371_at | KIF3B | 0,31660498 | 8,4803754 | 4,09139746 | 0,00102441 | 0,01480358 |
| 2810_at | SFN | -0,579136 | 12,1542469 | -4,09024893 | 0,00102676 | 0,01482568 |
| 81846_at | SBF2 | 0,42785061 | 6,98580765 | 4,08989688 | 0,00102748 | 0,01482568 |
| 7015_at | TERT | -0,48229344 | 5,80456422 | -4,08821156 | 0,00103093 | 0,01485355 |
| 23483_at | TGDS | -0,41718264 | 6,62001463 | -4,0882038 | 0,00103095 | 0,01485355 |
| 113263_at | GLCCI1 | 0,97396781 | 7,54241115 | 4,08715255 | 0,00103311 | 0,01486812 |
| 10985_at | GCN1 | -0,23421432 | 8,87039532 | -4,08696241 | 0,00103351 | 0,01486812 |
| 220042_at | DDIAS | -1,33425846 | 7,58459289 | -4,08616173 | 0,00103516 | 0,01488075 |
| 65980_at | BRD9 | -0,26288812 | 7,90968102 | -4,08307382 | 0,00104155 | 0,01496149 |
| 5927_at | KDM5A | 0,37314301 | 7,01498504 | 4,08234707 | 0,00104306 | 0,01497202 |
| 1958_at | EGR1 | 0,63912918 | 6,39668593 | 4,08054575 | 0,00104681 | 0,0149995 |
| 28231_at | SLCO4A1 | -1,81543433 | 7,63297606 | -4,08010261 | 0,00104774 | 0,0149995 |

| | | | | | | |
|---|---|---|---|---|---|---|
| 10212_at | DDX39A | -0,89278319 | 9,88292722 | -4,07983471 | 0,0010483 | 0,0149995 |
| 7026_at | NR2F2 | 0,28102837 | 9,29162077 | 4,07965077 | 0,00104868 | 0,0149995 |
| 26354_at | GNL3 | -0,64280068 | 9,15164673 | -4,07956254 | 0,00104887 | 0,0149995 |
| 134430_at | WDR36 | -0,61425905 | 7,45800634 | -4,07889718 | 0,00105026 | 0,01500827 |
| 51263_at | MRPL30 | -0,37089558 | 7,9351086 | -4,0785099 | 0,00105107 | 0,01500873 |
| 9746_at | CLSTN3 | 0,3340737 | 7,16168202 | 4,07765099 | 0,00105287 | 0,01502333 |
| 286826_at | LIN9 | -0,82717313 | 6,2433382 | -4,07694346 | 0,00105436 | 0,0150334 |
| 5494_at | PPM1A | 0,48091954 | 7,44755805 | 4,07616485 | 0,001056 | 0,01504563 |
| 8564_at | KMO | 0,57959332 | 3,98785138 | 4,07498799 | 0,00105848 | 0,01506984 |
| 25880_at | TMEM186 | -0,2957741 | 7,34702265 | -4,07344834 | 0,00106174 | 0,01509723 |
| 11214_at | AKAP13 | 0,32281041 | 6,83063625 | 4,07316898 | 0,00106233 | 0,01509723 |
| 1509_at | CTSD | 0,43136707 | 9,81769411 | 4,07296709 | 0,00106275 | 0,01509723 |
| 128637_at | TBC1D20 | 0,35023096 | 8,52904908 | 4,07250584 | 0,00106373 | 0,01509999 |
| 25809_at | TTLL1 | 0,56620973 | 6,07796293 | 4,07128641 | 0,00106632 | 0,01512125 |
| 442075_at | EMC3-AS1 | -0,58952845 | 6,13297293 | -4,07106212 | 0,0010668 | 0,01512125 |
| 5426_at | POLE | -0,71592957 | 7,54239665 | -4,06867619 | 0,00107189 | 0,01517267 |
| 401505_at | TOMM5 | -0,67648213 | 10,3015829 | -4,06862177 | 0,00107201 | 0,01517267 |
| 5373_at | PMM2 | -0,37399208 | 7,42185675 | -4,06790195 | 0,00107355 | 0,01517267 |
| 23130_at | ATG2A | 0,32404313 | 8,36129193 | 4,06788747 | 0,00107358 | 0,01517267 |
| 51433_at | ANAPC5 | -0,24546156 | 9,77790683 | -4,06425564 | 0,00108139 | 0,0152718 |
| 84752_at | B3GNT9 | 0,45726622 | 6,45660424 | 4,06093831 | 0,00108857 | 0,01535189 |
| 84299_at | MIEN1 | 0,281474 | 9,55659054 | 4,06063418 | 0,00108923 | 0,01535189 |
| 113026_at | PLCD3 | 0,30509456 | 8,4068133 | 4,06036995 | 0,0010898 | 0,01535189 |
| 115708_at | TRMT61A | -0,38985187 | 7,3079186 | -4,06016708 | 0,00109024 | 0,01535189 |
| 57224_at | NHSL1 | -0,29280303 | 6,94528365 | -4,05839381 | 0,00109411 | 0,01539506 |
| 7167_at | TPI1 | -0,49809209 | 10,8953861 | -4,05768742 | 0,00109565 | 0,01540553 |
| 6124_at | RPL4 | -0,21618545 | 12,1649138 | -4,05654846 | 0,00109815 | 0,01542932 |
| 5621_at | PRNP | -0,69844589 | 10,7746218 | -4,05440319 | 0,00110286 | 0,01548424 |
| 8438_at | RAD54L | -1,26101267 | 7,00022569 | -4,05081116 | 0,0011108 | 0,0155843 |
| 8568_at | RRP1 | -0,79651307 | 8,08436941 | -4,05041898 | 0,00111167 | 0,01558516 |
| 286343_at | LURAP1L | 0,95051904 | 7,90344236 | 4,0473955 | 0,0011184 | 0,01566811 |
| 22900_at | CARD8 | 0,3083425 | 5,77937592 | 4,046345 | 0,00112074 | 0,0156896 |
| 1303_at | COL12A1 | 0,68918972 | 9,27427133 | 4,04301799 | 0,00112821 | 0,01578271 |
| 23060_at | ZNF609 | 0,25989498 | 7,13382672 | 4,04227885 | 0,00112988 | 0,01579455 |
| 2114_at | ETS2 | -0,40770988 | 8,70922024 | -4,04086002 | 0,00113309 | 0,01582788 |
| 25943_at | C20orf194 | 0,65771572 | 7,27299268 | 4,04014052 | 0,00113472 | 0,01583915 |
| 64598_at | MOSPD3 | 0,46351769 | 7,70781842 | 4,03906762 | 0,00113715 | 0,01584835 |
| 29934_at | SNX12 | 0,24489097 | 8,64757233 | 4,03862644 | 0,00113815 | 0,01584835 |
| 844_at | CASQ1 | 1,46817783 | 5,83679915 | 4,03840072 | 0,00113867 | 0,01584835 |
| 83440_at | ADPGK | -0,27232902 | 7,80664995 | -4,0384006 | 0,00113867 | 0,01584835 |
| 138050_at | HGSNAT | 0,46262164 | 7,46190846 | 4,03701252 | 0,00114183 | 0,01588087 |
| 9367_at | RAB9A | 0,30061162 | 7,94571162 | 4,03639166 | 0,00114324 | 0,0158891 |
| 57153_at | SLC44A2 | 0,33980724 | 9,6275478 | 4,03562081 | 0,00114501 | 0,0158998 |
| 9868_at | TOMM70A | -0,60284711 | 8,80421051 | -4,03533291 | 0,00114566 | 0,0158998 |
| 79132_at | DHX58 | 0,72296078 | 6,51260836 | 4,03439192 | 0,00114782 | 0,01591824 |
| 101927279_a | EP300-AS1 | -0,48409184 | 5,40910422 | -4,03278815 | 0,0011515 | 0,01595783 |
| 2271_at | FH | -0,41263705 | 8,03729961 | -4,03182922 | 0,00115371 | 0,01596419 |
| 8187_at | ZNF239 | -0,86101899 | 5,29188074 | -4,03173228 | 0,00115393 | 0,01596419 |
| 1994_at | ELAVL1 | -0,22877314 | 7,89537983 | -4,03150936 | 0,00115445 | 0,01596419 |
| 138151_at | NACC2 | 0,6431832 | 7,97418787 | 4,03085751 | 0,00115595 | 0,01597353 |

| | | | | | | |
|---|---|---|---|---|---|---|
| 9709_at | HERPUD1 | 0,46827629 | 10,201228 | 4,02998402 | 0,00115797 | 0,01598996 |
| 9045_at | RPL14 | -0,3339444 | 10,9594232 | -4,02910624 | 0,00116 | 0,0159966 |
| 3643_at | INSR | 0,43708982 | 6,3671107 | 4,02901371 | 0,00116022 | 0,0159966 |
| 64949_at | MRPS26 | -0,39309125 | 9,37208028 | -4,02870081 | 0,00116094 | 0,0159966 |
| 83879_at | CDCA7 | -1,61134957 | 8,24444086 | -4,02721111 | 0,0011644 | 0,01603282 |
| 7874_at | USP7 | -0,23542458 | 8,94686895 | -4,02606741 | 0,00116707 | 0,01604791 |
| 25853_at | DCAF12 | 0,22203691 | 9,12679432 | 4,02568201 | 0,00116797 | 0,01604791 |
| 54762_at | GRAMD1C | 1,33326803 | 6,21343594 | 4,02563242 | 0,00116808 | 0,01604791 |
| 55635_at | DEPDC1 | -1,01584527 | 7,30964647 | -4,02531107 | 0,00116883 | 0,01604791 |
| 6009_at | RHEB | -0,36346905 | 6,87373447 | -4,02483743 | 0,00116994 | 0,01605167 |
| 1107_at | CHD3 | 0,42971077 | 7,70178181 | 4,02377614 | 0,00117242 | 0,01606701 |
| 57819_at | LSM2 | -0,79086363 | 8,41229843 | -4,02364694 | 0,00117273 | 0,01606701 |
| 79690_at | GAL3ST4 | 0,5423034 | 7,01511417 | 4,0225316 | 0,00117534 | 0,01609142 |
| 64225_at | ATL2 | -0,38460319 | 7,13475334 | -4,02169111 | 0,00117732 | 0,01610702 |
| 1453_at | CSNK1D | 0,29231737 | 9,44295645 | 4,02127299 | 0,0011783 | 0,01610904 |
| 55055_at | ZWILCH | -0,96141458 | 8,673828 | -4,0196866 | 0,00118204 | 0,01613333 |
| 10440_at | TIMM17A | -0,62072507 | 8,73841841 | -4,01962049 | 0,0011822 | 0,01613333 |
| 2617_at | GARS | -0,60258487 | 10,4147461 | -4,01945497 | 0,00118259 | 0,01613333 |
| 84733_at | CBX2 | -0,34550381 | 6,48232906 | -4,0152273 | 0,00119263 | 0,01625871 |
| 8884_at | SLC5A6 | -0,35916407 | 8,39230776 | -4,01481624 | 0,00119361 | 0,01626057 |
| 55841_at | WWC3 | 0,37392524 | 8,56226071 | 4,01444987 | 0,00119448 | 0,01626098 |
| 57602_at | USP36 | -0,27513164 | 7,00476981 | -4,01155163 | 0,00120142 | 0,01633306 |
| 55280_at | CWF19L1 | -0,26551985 | 7,30026256 | -4,01153089 | 0,00120147 | 0,01633306 |
| 9410_at | SNRNP40 | -1,01598135 | 8,85588146 | -4,01090525 | 0,00120298 | 0,01634197 |
| 51710_at | ZNF44 | 0,32547007 | 6,43058015 | 4,00832042 | 0,00120921 | 0,01641506 |
| 9374_at | PPT2 | 0,46081901 | 7,45005188 | 4,00769098 | 0,00121073 | 0,01642416 |
| 607_at | BCL9 | 0,44563389 | 7,62784854 | 4,00593041 | 0,001215 | 0,01647048 |
| 79675_at | FASTKD1 | -0,4267119 | 7,94376656 | -4,00341397 | 0,00122113 | 0,01653275 |
| 149473_at | CCDC24 | 0,27789395 | 6,42782134 | 4,00333976 | 0,00122131 | 0,01653275 |
| 5501_at | PPP1CC | -0,26024418 | 11,3911544 | -4,00201385 | 0,00122455 | 0,016565 |
| 57677_at | ZFP14 | 0,3380501 | 3,79949734 | 4,00095856 | 0,00122714 | 0,01656935 |
| 93109_at | TMEM44 | 0,38501423 | 6,83454038 | 4,00094851 | 0,00122716 | 0,01656935 |
| 55744_at | COA1 | -0,34843148 | 6,76253079 | -4,00083034 | 0,00122745 | 0,01656935 |
| 122953_at | JDP2 | 0,71509244 | 6,30990572 | 3,99937518 | 0,00123103 | 0,016606 |
| 84790_at | TUBA1C | -0,75565472 | 10,8767964 | -3,99834941 | 0,00123356 | 0,01661758 |
| 65057_at | ACD | -0,5029328 | 8,59464687 | -3,99832702 | 0,00123362 | 0,01661758 |
| 7789_at | ZXDA | 0,64687902 | 5,71330052 | 3,99680508 | 0,00123738 | 0,01665659 |
| 54906_at | FAM208B | -0,38233513 | 7,34730127 | -3,99462491 | 0,00124278 | 0,0167177 |
| 51088_at | KLHL5 | 0,39169811 | 6,56083905 | 3,99337175 | 0,0012459 | 0,01674796 |
| 2799_at | GNS | 0,42470028 | 9,37841629 | 3,99280674 | 0,00124731 | 0,01674964 |
| 80195_at | TMEM254 | 0,4611382 | 6,64111735 | 3,99262468 | 0,00124777 | 0,01674964 |
| 84310_at | C7orf50 | -0,30176532 | 8,35521217 | -3,991884 | 0,00124962 | 0,01676279 |
| 79172_at | CENPO | -0,68640669 | 7,16196478 | -3,99104766 | 0,00125171 | 0,01677917 |
| 51367_at | POP5 | -0,3803565 | 8,48726363 | -3,98701454 | 0,00126185 | 0,01689192 |
| 57161_at | PELI2 | 0,70589973 | 6,59218169 | 3,98700516 | 0,00126187 | 0,01689192 |
| 6188_at | RPS3 | -0,22980957 | 11,3293199 | -3,98514264 | 0,00126659 | 0,01694321 |
| 51643_at | TMBIM4 | 0,40502474 | 10,1345891 | 3,98265375 | 0,00127291 | 0,01701204 |
| 56647_at | BCCIP | -0,59922688 | 8,15366881 | -3,98189237 | 0,00127485 | 0,01701204 |
| 27346_at | TMEM97 | -0,76396847 | 9,33991594 | -3,98176175 | 0,00127518 | 0,01701204 |
| 54555_at | DDX49 | -0,27401076 | 8,40347855 | -3,98173052 | 0,00127526 | 0,01701204 |

| | | | | | | |
|---|---|---|---|---|---|---|
| 23424_at | TDRD7 | 0,76453539 | 7,71799898 | 3,98079743 | 0,00127765 | 0,01703204 |
| 4288_at | MKI67 | -1,14446501 | 8,30312401 | -3,98014921 | 0,0012793 | 0,01703403 |
| 2064_at | ERBB2 | 0,44367656 | 6,6217479 | 3,97991471 | 0,00127991 | 0,01703403 |
| 10207_at | INADL | 0,51728755 | 6,29773024 | 3,97970271 | 0,00128045 | 0,01703403 |
| 55320_at | MIS18BP1 | -0,52896709 | 7,66574243 | -3,9791265 | 0,00128193 | 0,01704192 |
| 389362_at | PSMG4 | -0,33383046 | 7,72052363 | -3,97874441 | 0,00128291 | 0,01704319 |
| 57162_at | PELI1 | 0,54878027 | 6,05899223 | 3,97271675 | 0,00129848 | 0,01723817 |
| 80345_at | ZSCAN16 | 0,58116364 | 5,62412539 | 3,97127795 | 0,00130222 | 0,01727598 |
| 5791_at | PTPRE | -0,37857938 | 6,25767105 | -3,97076357 | 0,00130357 | 0,01728188 |
| 50848_at | F11R | 0,40546607 | 10,0974319 | 3,96809865 | 0,00131054 | 0,01736238 |
| 11108_at | PRDM4 | 0,22525307 | 8,17527285 | 3,96732009 | 0,00131258 | 0,01736577 |
| 55683_at | KANSL3 | 0,25668442 | 7,02902063 | 3,9673144 | 0,0013126 | 0,01736577 |
| 6672_at | SP100 | 0,81320882 | 7,42946418 | 3,96657004 | 0,00131456 | 0,01737974 |
| 81890_at | QTRT1 | -0,46614615 | 7,99021829 | -3,96601677 | 0,00131601 | 0,01738707 |
| 9833_at | MELK | -1,22287533 | 8,9951612 | -3,96463007 | 0,00131967 | 0,01742346 |
| 1201_at | CLN3 | 0,48001086 | 8,02319965 | 3,96400971 | 0,00132131 | 0,01743317 |
| 5089_at | PBX2 | 0,39185339 | 7,47192838 | 3,96336847 | 0,00132301 | 0,01744362 |
| 3609_at | ILF3 | -0,44249002 | 9,2098553 | -3,96201017 | 0,00132661 | 0,01747916 |
| 89932_at | PAPLN | 0,489539 | 6,13959442 | 3,95955954 | 0,00133314 | 0,01755315 |
| 10525_at | HYOU1 | -0,32284184 | 9,35777411 | -3,95851373 | 0,00133593 | 0,01757794 |
| 120_at | ADD3 | 0,57984141 | 9,00469801 | 3,95794347 | 0,00133746 | 0,01758601 |
| 3636_at | INPPL1 | 0,39356672 | 8,32984989 | 3,95607992 | 0,00134246 | 0,01763973 |
| 154810_at | AMOTL1 | 0,62784597 | 6,30564278 | 3,95438801 | 0,00134702 | 0,01767596 |
| 4172_at | MCM3 | -1,04147312 | 9,62016046 | -3,95437469 | 0,00134705 | 0,01767596 |
| 50640_at | PNPLA8 | 0,66388181 | 8,22345702 | 3,95148006 | 0,00135489 | 0,01775684 |
| 134266_at | GRPEL2 | -0,2930176 | 8,66360142 | -3,95140868 | 0,00135508 | 0,01775684 |
| 89853_at | MVB12B | 0,36823911 | 6,78750374 | 3,95107638 | 0,00135598 | 0,01775684 |
| 51493_at | RTCB | -0,23637976 | 9,48780353 | -3,9506624 | 0,00135711 | 0,0177595 |
| 3486_at | IGFBP3 | 0,65483382 | 12,538115 | 3,9475026 | 0,00136573 | 0,01785138 |
| 6241_at | RRM2 | -1,63792895 | 10,174383 | -3,94740896 | 0,00136598 | 0,01785138 |
| 63979_at | FIGNL1 | -0,91503702 | 7,49583955 | -3,94654292 | 0,00136836 | 0,01787025 |
| 4303_at | FOXO4 | 0,4269305 | 5,70303746 | 3,94573425 | 0,00137057 | 0,0178871 |
| 2187_at | FANCB | -0,83234442 | 4,43656817 | -3,9444891 | 0,001374 | 0,01791964 |
| 84133_at | ZNRF3 | -0,34419145 | 7,09640387 | -3,94354785 | 0,00137659 | 0,01792974 |
| 54840_at | APTX | -0,34153436 | 7,35910262 | -3,94353269 | 0,00137663 | 0,01792974 |
| 55240_at | STEAP3 | -0,54205101 | 7,43238638 | -3,94230133 | 0,00138003 | 0,01795163 |
| 5422_at | POLA1 | -0,72051615 | 7,36477316 | -3,94224924 | 0,00138018 | 0,01795163 |
| 28969_at | BZW2 | -0,45764868 | 10,5502968 | -3,94126097 | 0,00138291 | 0,01797508 |
| 283871_at | PGP | -0,72313119 | 8,8077519 | -3,94009213 | 0,00138616 | 0,01800508 |
| 64400_at | AKTIP | 0,40608782 | 8,36035013 | 3,93953703 | 0,0013877 | 0,01801297 |
| 9749_at | PHACTR2 | 0,85531981 | 8,16660885 | 3,93908705 | 0,00138895 | 0,01801708 |
| 9252_at | RPS6KA5 | 0,41796266 | 6,18861345 | 3,93796262 | 0,00139208 | 0,01804557 |
| 64785_at | GINS3 | -1,11480252 | 7,67470523 | -3,93718575 | 0,00139425 | 0,01806154 |
| 79728_at | PALB2 | -0,41986612 | 8,18675234 | -3,93517751 | 0,00139988 | 0,01811414 |
| 115110_at | LOC115110 | 0,41295003 | 6,77792475 | 3,93500105 | 0,00140037 | 0,01811414 |
| 142_at | PARP1 | -0,54445548 | 9,99755451 | -3,93472901 | 0,00140114 | 0,01811414 |
| 100128881_a | VPS9D1-AS1 | -0,35864936 | 6,18377636 | -3,93285764 | 0,0014064 | 0,01817001 |
| 10227_at | MFSD10 | 0,42713489 | 9,54865678 | 3,93221452 | 0,00140821 | 0,01818126 |
| 29980_at | DONSON | -0,75770664 | 9,50340851 | -3,9312097 | 0,00141105 | 0,0181998 |
| 8243_at | SMC1A | -0,23251367 | 7,00514529 | -3,93084965 | 0,00141207 | 0,0181998 |

| | | | | | | |
|---|---|---|---|---|---|---|
| 6574_at | SLC20A1 | -0,75911407 | 10,1340205 | -3,9307038 | 0,00141249 | 0,0181998 |
| 51659_at | GINS2 | -1,37465654 | 8,74806981 | -3,93008007 | 0,00141425 | 0,01821039 |
| 440854_at | CAPN14 | 0,52347377 | 3,87906392 | 3,92672377 | 0,0014238 | 0,01832108 |
| 724102_at | SNHG4 | -0,72162694 | 3,73315833 | -3,92173154 | 0,00143812 | 0,01849302 |
| 10155_at | TRIM28 | -0,41754612 | 9,73710776 | -3,92038432 | 0,00144201 | 0,01853067 |
| 59348_at | ZNF350 | 0,57788482 | 5,83201553 | 3,91967645 | 0,00144406 | 0,01854462 |
| 2805_at | GOT1 | -0,48422682 | 9,38962729 | -3,91917486 | 0,00144551 | 0,01854762 |
| 79036_at | KXD1 | -0,3225175 | 9,01546348 | -3,91893128 | 0,00144622 | 0,01854762 |
| 51067_at | YARS2 | -0,34637454 | 8,3511975 | -3,91634012 | 0,00145375 | 0,01863184 |
| 135932_at | TMEM139 | 0,9059023 | 6,45429411 | 3,91493948 | 0,00145784 | 0,01866649 |
| 949_at | SCARB1 | -0,37786534 | 8,48073143 | -3,91445654 | 0,00145926 | 0,01866649 |
| 51776_at | ZAK | -0,5610147 | 6,80392073 | -3,91441951 | 0,00145936 | 0,01866649 |
| 55034_at | MOCOS | -0,565284 | 7,82119592 | -3,91199361 | 0,00146648 | 0,01874508 |
| 28951_at | TRIB2 | -0,545192 | 8,06925891 | -3,91146691 | 0,00146803 | 0,01875245 |
| 352954_at | GATS | 0,57132035 | 5,79336407 | 3,90978977 | 0,00147298 | 0,01880317 |
| 84440_at | RAB11FIP4 | 0,64051066 | 6,7905637 | 3,90861315 | 0,00147646 | 0,01883463 |
| 6491_at | STIL | -1,0857169 | 8,29300752 | -3,90829603 | 0,0014774 | 0,01883463 |
| 10009_at | ZBTB33 | 0,31854802 | 9,21448854 | 3,90762937 | 0,00147938 | 0,01884736 |
| 10430_at | TMEM147 | -0,24989856 | 10,7293563 | -3,90505769 | 0,00148703 | 0,01891687 |
| 4860_at | PNP | -1,13417788 | 8,61652661 | -3,90497231 | 0,00148728 | 0,01891687 |
| 55667_at | DENND4C | 0,68854167 | 9,04615572 | 3,90480593 | 0,00148778 | 0,01891687 |
| 83986_at | FAM234A | 0,39014314 | 8,64363915 | 3,90321528 | 0,00149253 | 0,01895914 |
| 1503_at | CTPS1 | -1,07100864 | 8,37761946 | -3,90303572 | 0,00149307 | 0,01895914 |
| 84181_at | CHD6 | 0,51518175 | 6,66649326 | 3,90171957 | 0,00149702 | 0,01899675 |
| 3631_at | INPP4A | 0,39245278 | 6,70478396 | 3,90033637 | 0,00150118 | 0,019037 |
| 127943_at | FCRLB | -0,37260261 | 6,65775839 | -3,89823181 | 0,00150753 | 0,01909412 |
| 23405_at | DICER1 | 0,4600708 | 8,9722494 | 3,89818733 | 0,00150767 | 0,01909412 |
| 8910_at | SGCE | 0,35900958 | 7,2504359 | 3,89772549 | 0,00150906 | 0,01909927 |
| 146909_at | KIF18B | -1,05764546 | 8,55588803 | -3,89698647 | 0,0015113 | 0,01911505 |
| 51773_at | RSF1 | 0,39946447 | 7,0528602 | 3,89626258 | 0,0015135 | 0,01913027 |
| 83657_at | DYNLRB2 | 0,71344779 | 4,75489623 | 3,89569784 | 0,00151522 | 0,01913338 |
| 56897_at | WRNIP1 | -0,26021616 | 9,48574113 | -3,89552811 | 0,00151573 | 0,01913338 |
| 9891_at | NUAK1 | 0,65931425 | 7,67532318 | 3,89480258 | 0,00151794 | 0,01914871 |
| 3433_at | IFIT2 | 1,58189529 | 7,23035755 | 3,89441156 | 0,00151913 | 0,0191512 |
| 57026_at | PDXP | -0,3411071 | 7,74442878 | -3,89396084 | 0,00152051 | 0,01915599 |
| 79649_at | MAP7D3 | -0,24931514 | 6,12579179 | -3,89270095 | 0,00152436 | 0,01919193 |
| 23230_at | VPS13A | 0,33652257 | 6,7685189 | 3,89205531 | 0,00152633 | 0,01920426 |
| 9618_at | TRAF4 | 0,32288785 | 8,35742489 | 3,89023738 | 0,00153191 | 0,01925982 |
| 57129_at | MRPL47 | -0,49711221 | 10,123564 | -3,88996527 | 0,00153275 | 0,01925982 |
| 286257_at | C9orf142 | -0,49610292 | 8,40484435 | -3,88938101 | 0,00153455 | 0,01926985 |
| 2091_at | FBL | -0,87532512 | 10,9259484 | -3,88804102 | 0,00153868 | 0,01930903 |
| 81031_at | SLC2A10 | 1,1423709 | 8,14572316 | 3,88768954 | 0,00153976 | 0,01930903 |
| 443_at | ASPA | 1,07857438 | 3,97718295 | 3,88739558 | 0,00154067 | 0,01930903 |
| 91689_at | SMDT1 | 0,21531006 | 10,0278592 | 3,88645895 | 0,00154357 | 0,01933279 |
| 6809_at | STX3 | 0,30697374 | 7,48040945 | 3,88218984 | 0,00155686 | 0,0194865 |
| 60561_at | RINT1 | -0,40438011 | 7,57817539 | -3,8818416 | 0,00155795 | 0,01948746 |
| 9648_at | GCC2 | 0,62892192 | 7,7870634 | 3,88115541 | 0,00156009 | 0,01949321 |
| 4293_at | MAP3K9 | 0,25414462 | 7,02678068 | 3,88077798 | 0,00156128 | 0,01949321 |
| 9693_at | RAPGEF2 | 0,71657941 | 5,96676786 | 3,88072522 | 0,00156144 | 0,01949321 |
| 25821_at | MTO1 | -0,42403377 | 8,15037847 | -3,87741738 | 0,00157184 | 0,01961039 |

| | | | | | | |
|---|---|---|---|---|---|---|
| 220134_at | SKA1 | -1,29887532 | 6,3967239 | -3,87614196 | 0,00157587 | 0,01964794 |
| 10384_at | BTN3A3 | 1,87757733 | 7,70748559 | 3,874438 | 0,00158128 | 0,01970252 |
| 23463_at | ICMT | -0,41455932 | 9,95069244 | -3,87269224 | 0,00158683 | 0,01975893 |
| 53832_at | IL20RA | -0,53214817 | 7,05276078 | -3,87055943 | 0,00159364 | 0,01981378 |
| 26286_at | ARFGAP3 | 0,5057409 | 8,54050674 | 3,87021934 | 0,00159473 | 0,01981378 |
| 100289635_a | ZNF605 | 0,45598837 | 6,64034235 | 3,87015421 | 0,00159494 | 0,01981378 |
| 79668_at | PARP8 | 0,67415129 | 6,76506562 | 3,87002571 | 0,00159535 | 0,01981378 |
| 79048_at | SECISBP2 | 0,55187512 | 7,87317918 | 3,86963487 | 0,0015966 | 0,01981656 |
| 60314_at | C12orf10 | -0,35673016 | 8,71477882 | -3,86909306 | 0,00159834 | 0,01982536 |
| 7589_at | ZSCAN21 | 0,3190715 | 7,13728192 | 3,86729162 | 0,00160413 | 0,01987209 |
| 4488_at | MSX2 | 0,29838875 | 8,89937223 | 3,86727978 | 0,00160417 | 0,01987209 |
| 8226_at | PUDP | 0,3928657 | 8,40556493 | 3,86650455 | 0,00160667 | 0,01989027 |
| 90075_at | ZNF30 | -0,32949286 | 5,50182352 | -3,86501603 | 0,00161148 | 0,019937 |
| 10785_at | WDR4 | -1,13092342 | 7,40104882 | -3,86443942 | 0,00161335 | 0,0199473 |
| 5902_at | RANBP1 | -0,68519667 | 6,49780199 | -3,86191458 | 0,00162155 | 0,02003585 |
| 8563_at | THOC5 | -0,41249898 | 8,41656753 | -3,85841988 | 0,00163297 | 0,02016406 |
| 29979_at | UBQLN1 | -0,44998827 | 9,58533194 | -3,85711644 | 0,00163725 | 0,02018715 |
| 56977_at | STOX2 | 0,67780123 | 6,7331961 | 3,85710391 | 0,00163729 | 0,02018715 |
| 256643_at | CXorf23 | 0,56525493 | 5,86230069 | 3,85689361 | 0,00163799 | 0,02018715 |
| 79774_at | GRTP1 | 0,36310948 | 6,30728743 | 3,85611822 | 0,00164054 | 0,02019349 |
| 27072_at | VPS41 | 0,43316724 | 6,82854397 | 3,85610077 | 0,0016406 | 0,02019349 |
| 79677_at | SMC6 | -0,48910809 | 7,03499865 | -3,85537704 | 0,00164298 | 0,02020996 |
| 64968_at | MRPS6 | -0,27815369 | 10,3097482 | -3,85425977 | 0,00164668 | 0,02024245 |
| 81930_at | KIF18A | -1,02327729 | 6,62255552 | -3,85367311 | 0,00164862 | 0,02025339 |
| 85465_at | EPT1 | -0,61618049 | 8,40962792 | -3,85231625 | 0,00165312 | 0,02027852 |
| 7014_at | TERF2 | -0,24792762 | 7,65931832 | -3,8521834 | 0,00165356 | 0,02027852 |
| 55771_at | PRR11 | -0,91473808 | 8,07971571 | -3,85210454 | 0,00165382 | 0,02027852 |
| 622_at | BDH1 | -0,44772887 | 6,17689857 | -3,85037513 | 0,00165958 | 0,02032548 |
| 79618_at | HMBOX1 | 0,49617033 | 6,87520697 | 3,85032004 | 0,00165976 | 0,02032548 |
| 2958_at | GTF2A2 | -0,36982627 | 6,73706942 | -3,84928666 | 0,00166321 | 0,02035479 |
| 7112_at | TMPO | -0,94605855 | 8,67878362 | -3,84667988 | 0,00167195 | 0,0204487 |
| 6741_at | SSB | -0,50949156 | 9,66059803 | -3,84609474 | 0,00167391 | 0,02045976 |
| 6218_at | RPS17 | -0,30073485 | 11,3359305 | -3,84550369 | 0,0016759 | 0,02047109 |
| 5558_at | PRIM2 | -0,31231556 | 4,07585592 | -3,84506193 | 0,00167739 | 0,02047629 |
| 55080_at | TAPBPL | 0,8462041 | 7,8611379 | 3,84329849 | 0,00168335 | 0,02053596 |
| 91687_at | CENPL | -0,78123362 | 6,88558605 | -3,84165145 | 0,00168893 | 0,02059101 |
| 101928578_a | RAP2C-AS1 | 0,38661529 | 4,69710071 | 3,84034641 | 0,00169336 | 0,02062143 |
| 9015_at | TAF1A | -0,93851998 | 6,50270985 | -3,84028738 | 0,00169356 | 0,02062143 |
| 3832_at | KIF11 | -1,03265908 | 8,43144961 | -3,83611003 | 0,00170784 | 0,02078216 |
| 1520_at | CTSS | 1,76062446 | 6,8146929 | 3,8340103 | 0,00171507 | 0,0208314 |
| 56342_at | PPAN | -0,62440917 | 8,33548394 | -3,83399953 | 0,0017151 | 0,0208314 |
| 79915_at | ATAD5 | -0,81274171 | 6,28836299 | -3,83399093 | 0,00171513 | 0,0208314 |
| 51104_at | ABHD17B | 0,46613636 | 7,64763537 | 3,83275779 | 0,00171939 | 0,02086994 |
| 56892_at | C8orf4 | 0,87097399 | 8,08295097 | 3,83204515 | 0,00172185 | 0,0208867 |
| 8480_at | RAE1 | -0,30627906 | 9,16860609 | -3,83073259 | 0,0017264 | 0,02090375 |
| 158584_at | FAAH2 | 0,46964447 | 5,73062177 | 3,83014023 | 0,00172846 | 0,02090375 |
| 8609_at | KLF7 | 0,329565 | 6,83473183 | 3,83000044 | 0,00172895 | 0,02090375 |
| 84230_at | LRRC8C | -1,1275536 | 6,76981559 | -3,829966 | 0,00172907 | 0,02090375 |
| 1678_at | TIMM8A | -1,07506167 | 7,14822878 | -3,82995163 | 0,00172912 | 0,02090375 |
| 203068_at | TUBB | -0,67261435 | 11,8242141 | -3,82934531 | 0,00173122 | 0,02090375 |

| | | | | | | |
|---|---|---|---|---|---|---|
| 375444_at | C5orf34 | -1,02583602 | 6,24866098 | -3,82914855 | 0,00173191 | 0,02090375 |
| 112770_at | GLMP | 0,70673741 | 8,22108968 | 3,82866896 | 0,00173358 | 0,02090375 |
| 55262_at | C7orf43 | 0,22001686 | 7,24335313 | 3,82838447 | 0,00173457 | 0,02090375 |
| 90338_at | ZNF160 | 0,2069044 | 5,35700676 | 3,82835024 | 0,00173469 | 0,02090375 |
| 2590_at | GALNT2 | 0,27716028 | 9,34910048 | 3,82788338 | 0,00173632 | 0,02090375 |
| 79370_at | BCL2L14 | 0,89074076 | 5,22487104 | 3,82742799 | 0,00173791 | 0,02090375 |
| 51593_at | SRRT | -0,3982215 | 9,1838556 | -3,82704534 | 0,00173925 | 0,02090375 |
| 283927_at | NUDT7 | 0,59255786 | 5,20461473 | 3,8270037 | 0,00173939 | 0,02090375 |
| 25909_at | AHCTF1 | -0,44101737 | 8,00553505 | -3,82687406 | 0,00173985 | 0,02090375 |
| 699_at | BUB1 | -1,08240171 | 7,98867678 | -3,82653172 | 0,00174105 | 0,02090375 |
| 80212_at | CCDC92 | 0,95310331 | 7,97954324 | 3,8260202 | 0,00174284 | 0,02090375 |
| 51759_at | C9orf78 | -0,27274193 | 8,01005132 | -3,82575966 | 0,00174375 | 0,02090375 |
| 51258_at | MRPL51 | -0,38910865 | 10,784819 | -3,82572309 | 0,00174388 | 0,02090375 |
| 23328_at | SASH1 | 0,86769214 | 8,33589595 | 3,82276084 | 0,0017543 | 0,02101555 |
| 55000_at | TUG1 | 0,34759509 | 9,90218932 | 3,82031369 | 0,00176295 | 0,02110609 |
| 53836_at | GPR87 | 0,5747814 | 9,44396001 | 3,81889606 | 0,00176798 | 0,02115319 |
| 518_at | ATP5G3 | -0,2071984 | 10,1729305 | -3,81841415 | 0,0017697 | 0,02116056 |
| 3162_at | HMOX1 | 0,48757493 | 8,20035053 | 3,81683653 | 0,00177532 | 0,02121463 |
| 166379_at | BBS12 | 0,79786927 | 5,00827141 | 3,81616333 | 0,00177772 | 0,02121738 |
| 50814_at | NSDHL | -0,37074902 | 8,49264912 | -3,81605981 | 0,00177809 | 0,02121738 |
| 10950_at | BTG3 | -0,68506007 | 8,81042461 | -3,81568184 | 0,00177945 | 0,02121738 |
| 56942_at | CMC2 | -0,71212859 | 9,33008825 | -3,81550471 | 0,00178008 | 0,02121738 |
| 10907_at | TXNL4A | -0,41208141 | 7,67471389 | -3,81523208 | 0,00178106 | 0,02121738 |
| 638_at | BIK | 0,63130609 | 7,59445079 | 3,81487139 | 0,00178235 | 0,02121966 |
| 8897_at | MTMR3 | 0,32417836 | 6,85074181 | 3,812895 | 0,00178945 | 0,02127813 |
| 23378_at | RRP8 | -0,30906014 | 5,99139569 | -3,81288898 | 0,00178947 | 0,02127813 |
| 59277_at | NTN4 | 0,50980306 | 9,30751345 | 3,81242893 | 0,00179113 | 0,02128468 |
| 9968_at | MED12 | 0,33399358 | 7,47411508 | 3,81055303 | 0,0017979 | 0,02135197 |
| 139221_at | MUM1L1 | 0,79825701 | 5,42873229 | 3,80994825 | 0,00180009 | 0,02136479 |
| 51399_at | TRAPPC4 | -0,23793082 | 9,10486825 | -3,80916892 | 0,00180291 | 0,02138513 |
| 151525_at | WDSUB1 | 0,50857132 | 6,5635207 | 3,80780571 | 0,00180786 | 0,02143065 |
| 6732_at | SRPK1 | -0,48475687 | 8,85311686 | -3,80538613 | 0,00181668 | 0,02152197 |
| 55486_at | PARL | -0,21440354 | 9,11505498 | -3,8044422 | 0,00182013 | 0,02154962 |
| 9133_at | CCNB2 | -0,94142174 | 9,69301226 | -3,80360356 | 0,00182321 | 0,02157275 |
| 153684_at | LOC153684 | 0,34841961 | 5,68368267 | 3,8025277 | 0,00182716 | 0,02160622 |
| 7056_at | THBD | -0,94186545 | 8,45694767 | -3,80215659 | 0,00182852 | 0,02160909 |
| 100130705_a | LOC10013070 | 0,31992555 | 5,25251066 | 3,80121818 | 0,00183198 | 0,02163055 |
| 54809_at | SAMD9 | 0,81680814 | 9,63590521 | 3,80105369 | 0,00183258 | 0,02163055 |
| 284207_at | METRNL | 0,48374278 | 8,25180976 | 3,8004359 | 0,00183486 | 0,02164419 |
| 9735_at | KNTC1 | -0,99613252 | 7,03721611 | -3,79912306 | 0,00183972 | 0,02168815 |
| 6120_at | RPE | -0,39802525 | 8,34946568 | -3,79825403 | 0,00184293 | 0,02169029 |
| 57688_at | ZSWIM6 | 0,39866725 | 8,65860598 | 3,79797863 | 0,00184396 | 0,02169029 |
| 3704_at | ITPA | -0,21668411 | 8,76274617 | -3,79794818 | 0,00184407 | 0,02169029 |
| 9966_at | TNFSF15 | 0,66395528 | 5,90424983 | 3,79785871 | 0,0018444 | 0,02169029 |
| 55728_at | N4BP2 | 0,44423787 | 6,35359675 | 3,79734869 | 0,00184629 | 0,02169931 |
| 3898_at | LAD1 | -0,73480241 | 8,63497596 | -3,79674171 | 0,00184855 | 0,02170133 |
| 9793_at | CKAP5 | -0,55303705 | 8,2875081 | -3,79664872 | 0,0018489 | 0,02170133 |
| 5718_at | PSMD12 | -0,46329564 | 9,46527083 | -3,79639357 | 0,00184985 | 0,02170133 |
| 167691_at | LCA5 | 0,46013133 | 4,83647277 | 3,79564014 | 0,00185265 | 0,02172103 |
| 728621_at | CCDC30 | 0,19841526 | 3,04108852 | 3,79351471 | 0,00186059 | 0,02180085 |

| | | | | | | |
|---|---|---|---|---|---|---|
| 10051_at | SMC4 | -0,82139786 | 9,78292806 | -3,7929667 | 0,00186265 | 0,02181164 |
| 2556_at | GABRA3 | 0,40684279 | 4,66370794 | 3,79166863 | 0,00186752 | 0,0218554 |
| 90407_at | TMEM41A | -0,37817143 | 7,71782093 | -3,79094476 | 0,00187024 | 0,0218632 |
| 7516_at | XRCC2 | -0,45839327 | 6,98322009 | -3,79088797 | 0,00187045 | 0,0218632 |
| 4088_at | SMAD3 | 0,44135682 | 8,60394002 | 3,78930254 | 0,00187643 | 0,02191976 |
| 9232_at | PTTG1 | -0,92064752 | 9,46762213 | -3,78878815 | 0,00187837 | 0,02192 |
| 283989_at | TSEN54 | -0,31660394 | 8,19050829 | -3,78869478 | 0,00187873 | 0,02192 |
| 51479_at | ANKFY1 | 0,37421329 | 7,85502134 | 3,78814393 | 0,00188081 | 0,02193103 |
| 146956_at | EME1 | -0,63998623 | 6,68299866 | -3,78641816 | 0,00188735 | 0,02199402 |
| 10256_at | CNKSR1 | 0,5140765 | 7,69247013 | 3,78589427 | 0,00188935 | 0,02199418 |
| 81037_at | CLPTM1L | -0,27524231 | 9,94184083 | -3,78581378 | 0,00188965 | 0,02199418 |
| 100507577_a | LOC10050757 | 0,52580671 | 5,94678224 | 3,78490904 | 0,0018931 | 0,02202096 |
| 114781_at | BTBD9 | 0,26491366 | 5,66387044 | 3,78349972 | 0,00189847 | 0,02207018 |
| 969_at | CD69 | 0,43017089 | 3,36045752 | 3,7826971 | 0,00190154 | 0,02209253 |
| 57523_at | NYNRIN | 0,58414442 | 4,88159507 | 3,78129953 | 0,0019069 | 0,02214141 |
| 9879_at | DDX46 | -0,40864498 | 7,99510738 | -3,7797013 | 0,00191304 | 0,02219862 |
| 7920_at | ABHD16A | 0,27035297 | 8,22043094 | 3,77941893 | 0,00191413 | 0,02219862 |
| 4999_at | ORC2 | -0,29183238 | 8,2587877 | -3,77889865 | 0,00191614 | 0,02220851 |
| 5577_at | PRKAR2B | -0,47120319 | 7,03462948 | -3,77845771 | 0,00191784 | 0,02221486 |
| 134548_at | SOWAHA | 0,50879027 | 5,56463623 | 3,77786334 | 0,00192013 | 0,02222809 |
| 10550_at | ARL6IP5 | 0,6468185 | 10,6367732 | 3,77593486 | 0,0019276 | 0,02230115 |
| 54865_at | GPATCH4 | -0,88398602 | 8,25935532 | -3,7754177 | 0,00192961 | 0,02230776 |
| 64928_at | MRPL14 | -0,39397587 | 9,3050148 | -3,77519115 | 0,00193049 | 0,02230776 |
| 353_at | APRT | -0,35432467 | 9,18155546 | -3,77458518 | 0,00193285 | 0,02232034 |
| 10988_at | METAP2 | -0,47369972 | 8,75604379 | -3,7743156 | 0,0019339 | 0,02232034 |
| 51649_at | MRPS23 | -0,57379372 | 8,56655896 | -3,77261591 | 0,00194053 | 0,02238343 |
| 23126_at | POGZ | 0,23667861 | 8,74404544 | 3,7700044 | 0,00195075 | 0,02248796 |
| 116844_at | LRG1 | 0,52815616 | 6,3377669 | 3,76940673 | 0,0019531 | 0,02250157 |
| 80216_at | ALPK1 | 0,45000052 | 6,49886179 | 3,76896005 | 0,00195486 | 0,02250836 |
| 63926_at | ANKEF1 | -0,42363189 | 6,3080235 | -3,76848705 | 0,00195672 | 0,02251634 |
| 5682_at | PSMA1 | -0,44740981 | 10,5632206 | -3,76805046 | 0,00195844 | 0,02252269 |
| 27085_at | MTBP | -0,64245847 | 5,08369227 | -3,76312214 | 0,00197798 | 0,02273373 |
| 3091_at | HIF1A | -0,33950296 | 12,3891958 | -3,76198227 | 0,00198252 | 0,02277239 |
| 10591_at | DNPH1 | -0,53549385 | 8,96499189 | -3,75972764 | 0,00199154 | 0,02286238 |
| 902_at | CCNH | -0,5579257 | 7,46714917 | -3,75933479 | 0,00199312 | 0,02286685 |
| 23139_at | MAST2 | -0,26034626 | 7,0430883 | -3,75838627 | 0,00199693 | 0,02289694 |
| 81706_at | PPP1R14C | -0,5971452 | 7,0574432 | -3,75751609 | 0,00200043 | 0,02291596 |
| 4686_at | NCBP1 | -0,38203328 | 7,37055392 | -3,75734786 | 0,00200111 | 0,02291596 |
| 9973_at | CCS | 0,25318793 | 8,3464878 | 3,7570879 | 0,00200216 | 0,02291596 |
| 56649_at | TMPRSS4 | -0,55108106 | 9,69451716 | -3,75626991 | 0,00200546 | 0,02294012 |
| 10693_at | CCT6B | 0,53477295 | 4,26074354 | 3,75562287 | 0,00200807 | 0,0229564 |
| 55664_at | CDC37L1 | 0,31953954 | 6,60001804 | 3,75411791 | 0,00201417 | 0,02301243 |
| 65265_at | C8orf33 | -0,58818714 | 7,98291762 | -3,75256566 | 0,00202048 | 0,0230708 |
| 9742_at | IFT140 | 0,30998612 | 5,62130771 | 3,75178825 | 0,00202364 | 0,02309327 |
| 3965_at | LGALS9 | 0,93842636 | 7,17439132 | 3,7500804 | 0,00203062 | 0,02315912 |
| 1312_at | COMT | 0,25457851 | 10,1244875 | 3,74831858 | 0,00203784 | 0,02322771 |
| 26974_at | ZNF285 | 0,57043957 | 5,19392683 | 3,74690924 | 0,00204363 | 0,02327997 |
| 10322_at | SMYD5 | -0,40251894 | 7,19455565 | -3,74637363 | 0,00204583 | 0,02328962 |
| 5347_at | PLK1 | -0,99751675 | 8,12013867 | -3,74580095 | 0,0020482 | 0,02328962 |
| 7771_at | ZNF112 | 0,47869992 | 5,75700182 | 3,7457778 | 0,00204829 | 0,02328962 |

| | | | | | | |
|---|---|---|---|---|---|---|
| 54927_at | CHCHD3 | -0,56283091 | 7,96384289 | -3,74528627 | 0,00205032 | 0,02328962 |
| 163590_at | TOR1AIP2 | 0,39142626 | 7,1165213 | 3,74511463 | 0,00205103 | 0,02328962 |
| 10616_at | RBCK1 | 0,43295288 | 8,2887082 | 3,74494518 | 0,00205173 | 0,02328962 |
| 64224_at | HERPUD2 | 0,36672508 | 8,51509136 | 3,74440294 | 0,00205397 | 0,02330134 |
| 3422_at | IDI1 | -0,34160563 | 8,20276241 | -3,74395423 | 0,00205583 | 0,02330869 |
| 6941_at | TCF19 | -0,95723797 | 7,72928386 | -3,74217524 | 0,00206321 | 0,02337861 |
| 7098_at | TLR3 | 1,32819135 | 8,26001681 | 3,74163806 | 0,00206545 | 0,02339017 |
| 64854_at | USP46 | 0,50987752 | 7,46090831 | 3,74090319 | 0,00206851 | 0,02341106 |
| 157247_at | MGC27345 | -0,39075029 | 6,81664167 | -3,74047554 | 0,00207029 | 0,02341748 |
| 4706_at | NDUFAB1 | -0,44335682 | 10,662723 | -3,73995047 | 0,00207248 | 0,02342851 |
| 1112_at | FOXN3 | 0,48457712 | 8,07796585 | 3,73920055 | 0,00207561 | 0,02343983 |
| 91869_at | RFT1 | -0,65305836 | 7,07740817 | -3,73912845 | 0,00207592 | 0,02343983 |
| 30000_at | TNPO2 | -0,32347487 | 8,00836288 | -3,73837508 | 0,00207907 | 0,02344953 |
| 127253_at | TYW3 | -0,32424698 | 8,13133506 | -3,73811885 | 0,00208014 | 0,02344953 |
| 27347_at | STK39 | -0,3804142 | 8,89619954 | -3,7380511 | 0,00208043 | 0,02344953 |
| 1183_at | CLCN4 | -0,27632338 | 5,21008449 | -3,73774285 | 0,00208172 | 0,02345037 |
| 83861_at | RSPH3 | 0,33023564 | 6,64813392 | 3,7348374 | 0,00209394 | 0,02357427 |
| 9528_at | TMEM59 | 0,57969298 | 11,0459329 | 3,73431644 | 0,00209614 | 0,02358525 |
| 900_at | CCNG1 | 0,44591404 | 11,0112232 | 3,73315441 | 0,00210105 | 0,02362378 |
| 24137_at | KIF4A | -1,02226477 | 8,18938501 | -3,73292688 | 0,00210202 | 0,02362378 |
| 7514_at | XPO1 | -0,45303834 | 10,4706592 | -3,73146568 | 0,00210822 | 0,02367962 |
| 51076_at | CUTC | -0,46287214 | 7,54069093 | -3,73099069 | 0,00211023 | 0,02368849 |
| 57822_at | GRHL3 | 0,47044917 | 7,22814953 | 3,7286474 | 0,00212022 | 0,02378675 |
| 119385_at | AGAP11 | 0,55019519 | 5,19559437 | 3,72718531 | 0,00212648 | 0,02384305 |
| 26528_at | DAZAP1 | -0,40457024 | 9,42890602 | -3,72620889 | 0,00213067 | 0,02387531 |
| 3174_at | HNF4G | 0,35951014 | 4,60007445 | 3,72593716 | 0,00213183 | 0,02387531 |
| 55630_at | SLC39A4 | -0,30561463 | 8,010391 | -3,72424998 | 0,0021391 | 0,02394271 |
| 57447_at | NDRG2 | 0,56564481 | 8,07527497 | 3,7227355 | 0,00214563 | 0,02400195 |
| 5434_at | POLR2E | -0,39782691 | 10,5495594 | -3,72197466 | 0,00214893 | 0,02401102 |
| 79833_at | GEMIN6 | -0,71012034 | 7,74268471 | -3,72197214 | 0,00214894 | 0,02401102 |
| 405_at | ARNT | 0,45935118 | 7,32544947 | 3,72133782 | 0,00215169 | 0,02402651 |
| 55722_at | CEP72 | -0,59439329 | 6,80868508 | -3,72107695 | 0,00215282 | 0,02402651 |
| 10026_at | PIGK | -0,59378554 | 7,85306041 | -3,71840824 | 0,00216443 | 0,02414211 |
| 150223_at | YDJC | -0,43775909 | 8,75420226 | -3,71674196 | 0,00217171 | 0,02420288 |
| 25907_at | TMEM158 | -0,66158492 | 6,65268972 | -3,71658669 | 0,00217239 | 0,02420288 |
| 54993_at | ZSCAN2 | 0,38309864 | 6,21711386 | 3,71578738 | 0,00217589 | 0,024221 |
| 7869_at | SEMA3B | -0,51078767 | 6,54314246 | -3,71554538 | 0,00217696 | 0,024221 |
| 9683_at | N4BP1 | 0,41007086 | 8,45758547 | 3,71510967 | 0,00217887 | 0,024221 |
| 55739_at | CARKD | 0,22009511 | 9,12247081 | 3,71506915 | 0,00217905 | 0,024221 |
| 9798_at | IST1 | 0,35973707 | 5,67954983 | 3,71329596 | 0,00218685 | 0,02428656 |
| 1827_at | RCAN1 | 0,33776211 | 7,24210701 | 3,71315602 | 0,00218747 | 0,02428656 |
| 3161_at | HMMR | -1,03710403 | 8,00791791 | -3,71184483 | 0,00219326 | 0,02433681 |
| 91368_at | CDKN2AIPNL | -0,32146427 | 5,16407543 | -3,71111242 | 0,0021965 | 0,02435874 |
| 347_at | APOD | 0,68249294 | 6,13978368 | 3,7088216 | 0,00220666 | 0,02445741 |
| 84260_at | TCHP | -0,26087293 | 7,11260773 | -3,70635856 | 0,00221765 | 0,02456502 |
| 27440_at | CECR5 | -0,44508583 | 8,14050293 | -3,70508662 | 0,00222334 | 0,02461395 |
| 55735_at | DNAJC11 | -0,45858631 | 8,35272758 | -3,7042654 | 0,00222703 | 0,02464057 |
| 2335_at | FN1 | 0,53100441 | 8,67170198 | 3,70263566 | 0,00223436 | 0,02469536 |
| 4067_at | LYN | 0,64755845 | 9,38296549 | 3,70259433 | 0,00223454 | 0,02469536 |
| 9557_at | CHD1L | -0,56887062 | 6,90848463 | -3,7022503 | 0,00223609 | 0,02469833 |

| | | | | | | |
|---|---|---|---|---|---|---|
| 9918_at | NCAPD2 | -0,72451096 | 8,98466418 | -3,7016648 | 0,00223874 | 0,02471333 |
| 55168_at | MRPS18A | -0,33203448 | 8,94506555 | -3,70074431 | 0,00224289 | 0,02474507 |
| 1058_at | CENPA | -1,07077754 | 8,51459714 | -3,69988179 | 0,0022468 | 0,02477395 |
| 25885_at | POLR1A | -0,37456875 | 7,25123173 | -3,69950701 | 0,0022485 | 0,02477849 |
| 10475_at | TRIM38 | 0,73399939 | 8,32893117 | 3,69803522 | 0,00225518 | 0,02483793 |
| 11169_at | WDHD1 | -1,18328945 | 6,70653352 | -3,6973751 | 0,00225818 | 0,0248568 |
| 64979_at | MRPL36 | -0,46989163 | 9,49361284 | -3,69590223 | 0,0022649 | 0,0249165 |
| 10884_at | MRPS30 | -0,44459108 | 9,04729765 | -3,69549532 | 0,00226676 | 0,02492272 |
| 11315_at | PARK7 | -0,27423539 | 11,5304609 | -3,69470014 | 0,0022704 | 0,0249383 |
| 5166_at | PDK4 | 0,25175362 | 3,6566026 | 3,69461968 | 0,00227077 | 0,0249383 |
| 7032_at | TFF2 | -0,54374503 | 5,5567076 | -3,69290143 | 0,00227865 | 0,02501023 |
| 283_at | ANG | 0,55143987 | 5,78221917 | 3,69262626 | 0,00227991 | 0,02501023 |
| 440993_at | LINC00969 | 0,338945 | 5,29565961 | 3,69164393 | 0,00228443 | 0,02504557 |
| 4174_at | MCM5 | -1,21959 | 8,75678867 | -3,69026658 | 0,00229079 | 0,02510095 |
| 22974_at | TPX2 | -1,12655709 | 9,57135586 | -3,68969166 | 0,00229345 | 0,02510134 |
| 5536_at | PPP5C | -0,4314403 | 7,9577177 | -3,68966841 | 0,00229355 | 0,02510134 |
| 10926_at | DBF4 | -1,04359473 | 8,30601343 | -3,68941337 | 0,00229473 | 0,02510134 |
| 79980_at | DSN1 | -0,79489847 | 8,29628299 | -3,6886772 | 0,00229814 | 0,0251201 |
| 57506_at | MAVS | 0,29510855 | 8,4011701 | 3,68848018 | 0,00229906 | 0,0251201 |
| 8416_at | ANXA9 | 0,65966246 | 7,56059401 | 3,68791308 | 0,00230169 | 0,02513459 |
| 9442_at | MED27 | -0,35398593 | 6,86379387 | -3,68759403 | 0,00230317 | 0,02513652 |
| 1192_at | CLIC1 | -0,20686511 | 11,2767343 | -3,68716739 | 0,00230515 | 0,02514391 |
| 23646_at | PLD3 | 0,75349912 | 8,79152775 | 3,68614453 | 0,00230991 | 0,02518158 |
| 4528_at | MTIF2 | -0,46490239 | 8,82573036 | -3,68454676 | 0,00231737 | 0,02523586 |
| 1495_at | CTNNA1 | 0,19460348 | 10,1862458 | 3,68427908 | 0,00231862 | 0,02523586 |
| 5926_at | ARID4A | 0,47231704 | 6,41669051 | 3,68418795 | 0,00231905 | 0,02523586 |
| 5792_at | PTPRF | 0,64510417 | 9,28387269 | 3,68395582 | 0,00232013 | 0,02523586 |
| 23239_at | PHLPP1 | 0,29714933 | 7,02349495 | 3,68315887 | 0,00232386 | 0,02526219 |
| 3183_at | HNRNPC | -0,64840003 | 8,50332412 | -3,68089658 | 0,00233449 | 0,02536342 |
| 5565_at | PRKAB2 | 0,53521216 | 7,91775678 | 3,68012919 | 0,00233811 | 0,02538839 |
| 388569_at | ZNF324B | -0,30889547 | 6,09801733 | -3,67972801 | 0,00234 | 0,02539463 |
| 100505715_a | LOC10050571 | 0,52615026 | 5,92383829 | 3,67841455 | 0,00234621 | 0,02543932 |
| 27433_at | TOR2A | 0,31115752 | 7,24485481 | 3,67819785 | 0,00234724 | 0,02543932 |
| 25_at | ABL1 | 0,22561128 | 8,86276961 | 3,67801914 | 0,00234808 | 0,02543932 |
| 144233_at | BCDIN3D | 0,25820301 | 6,64362321 | 3,67662151 | 0,00235471 | 0,02547259 |
| 57474_at | ZNF490 | 0,5004142 | 7,39914349 | 3,6765887 | 0,00235487 | 0,02547259 |
| 390980_at | ZNF805 | 0,29851131 | 6,19984197 | 3,67653549 | 0,00235512 | 0,02547259 |
| 6304_at | SATB1 | 0,81500206 | 6,46081273 | 3,67571324 | 0,00235903 | 0,02550056 |
| 2752_at | GLUL | 0,5708969 | 9,42118166 | 3,6752098 | 0,00236143 | 0,02551216 |
| 55755_at | CDK5RAP2 | -0,32847539 | 7,187145 | -3,67414346 | 0,00236651 | 0,02555276 |
| 51204_at | TACO1 | -0,29347657 | 7,79996917 | -3,6725057 | 0,00237434 | 0,02562297 |
| 51699_at | VPS29 | -0,31112682 | 10,1619028 | -3,67165538 | 0,00237842 | 0,02565259 |
| 898_at | CCNE1 | -0,79179201 | 7,40188222 | -3,67030263 | 0,00238492 | 0,02569521 |
| 7444_at | VRK2 | -0,35579661 | 7,72371325 | -3,67027785 | 0,00238504 | 0,02569521 |
| 222584_at | FAM83B | 0,38456954 | 8,17569307 | 3,66979768 | 0,00238735 | 0,02570573 |
| 80315_at | CPEB4 | 0,58983884 | 6,8379279 | 3,66901275 | 0,00239114 | 0,02573209 |
| 8883_at | NAE1 | -0,38732003 | 10,0449378 | -3,6685528 | 0,00239336 | 0,0257416 |
| 55257_at | MRGBP | -0,36716726 | 9,5458287 | -3,66730217 | 0,0023994 | 0,02578526 |
| 64773_at | PCED1A | 0,5066124 | 8,73627744 | 3,66715956 | 0,00240009 | 0,02578526 |
| 1022_at | CDK7 | -0,50507538 | 7,34015241 | -3,665948 | 0,00240597 | 0,02583396 |

| | | | | | | |
|---|---|---|---|---|---|---|
| 83543_at | AIF1L | -0,32340823 | 6,13226274 | -3,66460714 | 0,00241248 | 0,02588952 |
| 27018_at | NGFRAP1 | 0,21352174 | 10,4272801 | 3,66373368 | 0,00241674 | 0,02592075 |
| 3476_at | IGBP1 | 0,3453853 | 9,30553111 | 3,66153081 | 0,00242751 | 0,02600762 |
| 119032_at | BORCS7 | 0,48354834 | 7,28979072 | 3,66152389 | 0,00242754 | 0,02600762 |
| 6880_at | TAF9 | -0,59253306 | 8,43686057 | -3,66054693 | 0,00243233 | 0,02604001 |
| 84064_at | HDHD2 | 0,26221772 | 8,48721343 | 3,66035616 | 0,00243327 | 0,02604001 |
| 4287_at | ATXN3 | 0,34067959 | 6,710547 | 3,659869 | 0,00243566 | 0,02605114 |
| 27235_at | COQ2 | -0,31546455 | 7,72172112 | -3,65896617 | 0,0024401 | 0,02608416 |
| 79818_at | ZNF552 | 0,73370974 | 7,67837828 | 3,65859785 | 0,00244192 | 0,02608908 |
| 54799_at | MBTD1 | 0,65556026 | 6,13191699 | 3,65818936 | 0,00244393 | 0,02609612 |
| 23345_at | SYNE1 | 0,29026642 | 4,32542661 | 3,65552563 | 0,0024571 | 0,02622224 |
| 3720_at | JARID2 | 0,56597301 | 7,86029165 | 3,65459215 | 0,00246173 | 0,02625713 |
| 26112_at | CCDC69 | 0,52333882 | 5,20814584 | 3,65402091 | 0,00246457 | 0,02627287 |
| 9813_at | EFCAB14 | 0,43769082 | 9,71228797 | 3,6528833 | 0,00247024 | 0,0263165 |
| 649_at | BMP1 | 0,22290196 | 7,18340084 | 3,65242753 | 0,00247251 | 0,0263165 |
| 10403_at | NDC80 | -1,18323609 | 7,8848521 | -3,65237686 | 0,00247276 | 0,0263165 |
| 2030_at | SLC29A1 | -0,51742281 | 8,62968459 | -3,6488207 | 0,00249058 | 0,02649141 |
| 113130_at | CDCA5 | -1,20141156 | 8,01215757 | -3,64793776 | 0,00249502 | 0,02652402 |
| 10229_at | COQ7 | -0,22006968 | 6,72509665 | -3,64665423 | 0,00250149 | 0,02656817 |
| 6626_at | SNRPA | -0,57304337 | 9,95175139 | -3,64656713 | 0,00250193 | 0,02656817 |
| 3202_at | HOXA5 | 0,76219157 | 8,3546896 | 3,64515228 | 0,00250908 | 0,02662946 |
| 23658_at | LSM5 | -0,60585784 | 7,49865894 | -3,64421914 | 0,00251381 | 0,02664913 |
| 3251_at | HPRT1 | -0,85813282 | 9,72870342 | -3,64395271 | 0,00251517 | 0,02664913 |
| 2116_at | ETV2 | 0,38109279 | 5,72830103 | 3,64360472 | 0,00251693 | 0,02664913 |
| 10196_at | PRMT3 | -0,84764013 | 7,29982693 | -3,64341792 | 0,00251788 | 0,02664913 |
| 145957_at | NRG4 | 0,53373592 | 6,20147434 | 3,64319712 | 0,002519 | 0,02664913 |
| 2707_at | GJB3 | -0,48547425 | 8,41638991 | -3,64315137 | 0,00251924 | 0,02664913 |
| 5557_at | PRIM1 | -0,99895932 | 8,12622294 | -3,64283085 | 0,00252087 | 0,02665174 |
| 55165_at | CEP55 | -1,01106623 | 9,30938939 | -3,64254187 | 0,00252234 | 0,02665266 |
| 55355_at | HJURP | -1,09159492 | 8,90068521 | -3,64045639 | 0,00253298 | 0,02675042 |
| 201229_at | LYRM9 | 0,50384441 | 5,17624855 | 3,63996259 | 0,0025355 | 0,02676021 |
| 9881_at | TRANK1 | 0,88829936 | 5,59670509 | 3,63973215 | 0,00253668 | 0,02676021 |
| 57823_at | SLAMF7 | 1,51344756 | 6,1158338 | 3,63909714 | 0,00253994 | 0,02677987 |
| 29081_at | METTL5 | -0,30058624 | 9,92397941 | -3,63785773 | 0,0025463 | 0,02683227 |
| 7873_at | MANF | -0,61626615 | 9,88082543 | -3,63538027 | 0,00255907 | 0,02695205 |
| 151246_at | SGOL2 | -1,0270092 | 7,85054041 | -3,63484364 | 0,00256184 | 0,02696652 |
| 8859_at | STK19 | 0,39951853 | 7,8587782 | 3,63445807 | 0,00256384 | 0,02697278 |
| 1073_at | CFL2 | 0,51635587 | 7,9820088 | 3,63365962 | 0,00256797 | 0,02700154 |
| 8805_at | TRIM24 | 0,53158887 | 9,50657081 | 3,63201361 | 0,00257652 | 0,02707664 |
| 23031_at | MAST3 | 0,24040546 | 6,72862868 | 3,6313274 | 0,00258009 | 0,02709939 |
| 221322_at | TBC1D32 | 0,60439792 | 4,81564518 | 3,62963185 | 0,00258894 | 0,02717749 |
| 64960_at | MRPS15 | -0,47281506 | 10,0641013 | -3,62819535 | 0,00259646 | 0,02724158 |
| 27067_at | STAU2 | 0,26207076 | 7,61909481 | 3,62572734 | 0,00260943 | 0,02736275 |
| 338440_at | ANO9 | 0,36742922 | 6,87935484 | 3,62141799 | 0,00263223 | 0,02758685 |
| 1140_at | CHRNB1 | 0,46890292 | 5,5329601 | 3,62085534 | 0,00263522 | 0,02760319 |
| 22990_at | PCNX | 0,28414422 | 6,91651816 | 3,62053529 | 0,00263692 | 0,02760602 |
| 729830_at | FAM160A1 | 0,39187563 | 6,32219053 | 3,61975126 | 0,0026411 | 0,02763474 |
| 84159_at | ARID5B | 0,22819995 | 6,12025555 | 3,61891363 | 0,00264557 | 0,02766649 |
| 8661_at | EIF3A | -0,17986424 | 10,2825618 | -3,61816028 | 0,0026496 | 0,02769357 |
| 51439_at | FAM8A1 | 0,3759636 | 9,26850878 | 3,61683976 | 0,00265667 | 0,02775246 |

| | | | | | | |
|---|---|---|---|---|---|---|
| 2051_at | EPHB6 | -0,7792524 | 8,78184218 | -3,61598653 | 0,00266126 | 0,02777952 |
| 3575_at | IL7R | -0,45893243 | 4,10768673 | -3,61582026 | 0,00266215 | 0,02777952 |
| 84798_at | C19orf48 | -1,11222404 | 8,5119229 | -3,61311109 | 0,00267675 | 0,02791678 |
| 23228_at | PLCL2 | 0,55627545 | 5,95697228 | 3,6125524 | 0,00267978 | 0,02793318 |
| 7023_at | TFAP4 | -0,32364119 | 6,64459647 | -3,61191866 | 0,00268321 | 0,02795382 |
| 57232_at | ZNF630 | 0,41720521 | 5,65994085 | 3,60845341 | 0,00270205 | 0,02813491 |
| 144577_at | C12orf66 | -0,4635177 | 6,0047195 | -3,60788968 | 0,00270513 | 0,02815174 |
| 113177_at | IZUMO4 | 0,29021633 | 5,66767739 | 3,60718841 | 0,00270896 | 0,02817641 |
| 1285_at | COL4A3 | 0,39523579 | 4,44775489 | 3,6051126 | 0,00272034 | 0,02827951 |
| 79663_at | HSPBAP1 | 0,27350941 | 5,71643391 | 3,60450662 | 0,00272367 | 0,02829386 |
| 51236_at | HGH1 | -0,46883127 | 8,17670019 | -3,60432738 | 0,00272466 | 0,02829386 |
| 80213_at | TM2D3 | 0,29699931 | 8,89443808 | 3,60352181 | 0,0027291 | 0,02832466 |
| 28960_at | DCPS | -0,31142642 | 8,61166427 | -3,6028176 | 0,00273298 | 0,02832882 |
| 7100_at | TLR5 | 0,49048325 | 4,81970845 | 3,60272189 | 0,00273351 | 0,02832882 |
| 7746_at | ZSCAN9 | 0,39959095 | 6,87107877 | 3,60264933 | 0,00273391 | 0,02832882 |
| 51148_at | CERCAM | 0,30778569 | 6,49583003 | 3,60097873 | 0,00274315 | 0,02840929 |
| 55003_at | PAK1IP1 | -0,85690431 | 8,47151528 | -3,60016056 | 0,00274769 | 0,02844098 |
| 5593_at | PRKG2 | 0,26972982 | 4,25612895 | 3,59815394 | 0,00275884 | 0,02853867 |
| 2058_at | EPRS | -0,40818505 | 8,7657656 | -3,59793111 | 0,00276009 | 0,02853867 |
| 6813_at | STXBP2 | 0,32806672 | 8,0825698 | 3,59730733 | 0,00276357 | 0,02855643 |
| 9875_at | URB1 | -0,43890529 | 6,34453238 | -3,59709186 | 0,00276477 | 0,02855643 |
| 51559_at | NT5DC3 | -0,25145889 | 6,1818037 | -3,59572639 | 0,0027724 | 0,02861807 |
| 7799_at | PRDM2 | 0,23153408 | 6,90441971 | 3,59531693 | 0,0027747 | 0,02861807 |
| 9700_at | ESPL1 | -0,94205648 | 7,74407066 | -3,59522861 | 0,00277519 | 0,02861807 |
| 114789_at | SLC25A25 | -0,32876428 | 6,77127239 | -3,59318121 | 0,00278669 | 0,02871982 |
| 84875_at | PARP10 | 0,33322372 | 7,00297506 | 3,59294167 | 0,00278804 | 0,02871982 |
| 80095_at | ZNF606 | 0,45463636 | 6,83500863 | 3,59261045 | 0,00278991 | 0,02872368 |
| 56850_at | GRIPAP1 | 0,22242533 | 7,20781198 | 3,59147637 | 0,00279631 | 0,02877418 |
| 23366_at | KIAA0895 | 0,48844504 | 5,83547886 | 3,59111356 | 0,00279836 | 0,0287799 |
| 90416_at | C15orf57 | 0,39499931 | 6,4403977 | 3,58996921 | 0,00280483 | 0,02883111 |
| 51164_at | DCTN4 | 0,33709534 | 8,69771638 | 3,58887786 | 0,00281102 | 0,02887933 |
| 3980_at | LIG3 | -0,2656507 | 7,10857877 | -3,5882911 | 0,00281436 | 0,02889568 |
| 8648_at | NCOA1 | 0,3875634 | 7,76143659 | 3,58806997 | 0,00281561 | 0,02889568 |
| 147694_at | ZNF548 | 0,29443303 | 6,2557493 | 3,58764775 | 0,00281802 | 0,02890494 |
| 166378_at | SPATA5 | -0,50196495 | 5,05134265 | -3,58536961 | 0,00283102 | 0,02902063 |
| 80705_at | TSGA10 | 1,01403406 | 4,49321159 | 3,58507953 | 0,00283268 | 0,02902063 |
| 54821_at | ERCC6L | -1,02290039 | 6,33708968 | -3,58462486 | 0,00283528 | 0,02902063 |
| 100506860_a | LOC10050686 | -0,68121575 | 4,71390751 | -3,58461714 | 0,00283532 | 0,02902063 |
| 1955_at | MEGF9 | 0,48351823 | 8,31321404 | 3,58266253 | 0,00284654 | 0,02911997 |
| 1855_at | DVL1 | -0,27214494 | 8,10582105 | -3,58234088 | 0,00284839 | 0,02912344 |
| 26996_at | GPR160 | 0,50614418 | 8,00671702 | 3,58182953 | 0,00285133 | 0,02913807 |
| 51092_at | SIDT2 | 0,85475467 | 8,63122925 | 3,58105051 | 0,00285583 | 0,02916849 |
| 5160_at | PDHA1 | -0,23895526 | 9,27588094 | -3,58068569 | 0,00285793 | 0,02917453 |
| 25850_at | ZNF345 | 0,35744988 | 4,76006741 | 3,58014529 | 0,00286105 | 0,02917627 |
| 10092_at | ARPC5 | 0,212752 | 11,0898682 | 3,58013169 | 0,00286113 | 0,02917627 |
| 6631_at | SNRPC | -0,49504532 | 9,09928045 | -3,57955412 | 0,00286447 | 0,02919488 |
| 55027_at | HEATR3 | -0,30193082 | 5,36861407 | -3,57850913 | 0,00287053 | 0,02924111 |
| 81691_at | LOC81691 | -0,56771544 | 6,22296287 | -3,57605754 | 0,00288478 | 0,02936263 |
| 11124_at | FAF1 | -0,3530708 | 8,41252752 | -3,57593321 | 0,0028855 | 0,02936263 |
| 1729_at | DIAPH1 | -0,55623069 | 8,61201637 | -3,57557846 | 0,00288757 | 0,02936817 |

| | | | | | | |
|---|---|---|---|---|---|---|
| 84803_at | GPAT3 | -1,54914739 | 7,09214752 | -3,57429929 | 0,00289505 | 0,02942863 |
| 56983_at | POGLUT1 | -0,30468223 | 7,44147819 | -3,57388593 | 0,00289746 | 0,02943768 |
| 51144_at | HSD17B12 | -0,68615237 | 10,1873802 | -3,5730421 | 0,00290241 | 0,02947237 |
| 1616_at | DAXX | -0,32317685 | 8,21242159 | -3,57181673 | 0,0029096 | 0,02952789 |
| 51293_at | CD320 | -0,94589744 | 8,31559025 | -3,5715358 | 0,00291125 | 0,02952789 |
| 5526_at | PPP2R5B | 0,2932379 | 6,65223592 | 3,57132851 | 0,00291247 | 0,02952789 |
| 57685_at | CACHD1 | -0,55535281 | 6,98187718 | -3,57030503 | 0,0029185 | 0,02956102 |
| 6059_at | ABCE1 | -0,96298747 | 8,00425668 | -3,57025289 | 0,00291881 | 0,02956102 |
| 10797_at | MTHFD2 | -0,94386137 | 10,0753879 | -3,56924379 | 0,00292477 | 0,0295962 |
| 150384_at | GTSE1-AS1 | -0,41451123 | 4,16564038 | -3,56914409 | 0,00292536 | 0,0295962 |
| 3169_at | FOXA1 | 0,33777404 | 11,200798 | 3,56799175 | 0,00293218 | 0,02964962 |
| 25816_at | TNFAIP8 | 0,42966242 | 9,06646164 | 3,56575386 | 0,00294547 | 0,02976836 |
| 64783_at | RBM15 | -0,31831491 | 7,8589843 | -3,56418382 | 0,00295482 | 0,02984729 |
| 284403_at | WDR62 | -0,41287927 | 6,51343056 | -3,56327138 | 0,00296028 | 0,02988263 |
| 25804_at | LSM4 | -0,53295174 | 10,1424626 | -3,56307885 | 0,00296143 | 0,02988263 |
| 7756_at | ZNF207 | -0,28848039 | 10,0281418 | -3,56282033 | 0,00296298 | 0,02988263 |
| 114904_at | C1QTNF6 | 0,6361255 | 7,38082462 | 3,56233364 | 0,00296589 | 0,02989638 |
| 27258_at | LSM3 | -0,5318764 | 10,5346584 | -3,56187816 | 0,00296862 | 0,02990826 |
| 26160_at | IFT172 | 0,46552178 | 6,50218989 | 3,56139929 | 0,0029715 | 0,02992157 |
| 3920_at | LAMP2 | 0,38399703 | 8,69853266 | 3,5599579 | 0,00298017 | 0,02999317 |
| 6737_at | TRIM21 | 0,65322305 | 7,22282316 | 3,55912188 | 0,0029852 | 0,03002819 |
| 101927797_a | LOC10192779 | -0,4006949 | 3,01390552 | -3,55862708 | 0,00298819 | 0,03004254 |
| 7096_at | TLR1 | 0,30005793 | 3,37093282 | 3,55656644 | 0,00300066 | 0,03013963 |
| 5728_at | PTEN | 0,31452746 | 8,28148047 | 3,55640062 | 0,00300167 | 0,03013963 |
| 896_at | CCND3 | -0,36106794 | 9,48312087 | -3,55625619 | 0,00300254 | 0,03013963 |
| 100506621_a | LINC01279 | 0,75252463 | 4,65557862 | 3,55589831 | 0,00300471 | 0,03014012 |
| 4734_at | NEDD4 | 0,71121514 | 8,2440595 | 3,55573276 | 0,00300572 | 0,03014012 |
| 57140_at | RNPEPL1 | 0,25772644 | 8,272421 | 3,55504444 | 0,0030099 | 0,03016637 |
| 399665_at | FAM102A | 0,39325025 | 9,19912122 | 3,55451106 | 0,00301315 | 0,03018219 |
| 6119_at | RPA3 | -0,42832607 | 8,92854428 | -3,55395419 | 0,00301654 | 0,03018219 |
| 582_at | BBS1 | 0,67425206 | 7,54053555 | 3,55383319 | 0,00301728 | 0,03018219 |
| 101926944_a | LOC10192694 | 0,26377719 | 3,00570511 | 3,5537562 | 0,00301775 | 0,03018219 |
| 4796_at | TONSL | -0,58471593 | 6,50544057 | -3,55236492 | 0,00302625 | 0,03020925 |
| 8266_at | UBL4A | -0,39968841 | 8,36309515 | -3,5522086 | 0,0030272 | 0,03020925 |
| 11188_at | NISCH | 0,2174784 | 7,27229852 | 3,55214552 | 0,00302759 | 0,03020925 |
| 9857_at | CEP350 | 0,4807838 | 8,03413934 | 3,55185232 | 0,00302938 | 0,03020925 |
| 9985_at | REC8 | 0,41458597 | 6,64177436 | 3,55173452 | 0,0030301 | 0,03020925 |
| 6897_at | TARS | -0,55078993 | 9,52063772 | -3,55124479 | 0,0030331 | 0,03020925 |
| 483_at | ATP1B3 | -0,57563797 | 11,1020568 | -3,55102503 | 0,00303445 | 0,03020925 |
| 85481_at | PSKH2 | -0,27383843 | 4,36600601 | -3,55075337 | 0,00303612 | 0,03020925 |
| 400629_at | TEX19 | -0,30800463 | 5,06913498 | -3,55067032 | 0,00303663 | 0,03020925 |
| 79038_at | ZFYVE21 | 0,23430855 | 9,23863923 | 3,55055378 | 0,00303734 | 0,03020925 |
| 1997_at | ELF1 | 0,48479799 | 9,62570647 | 3,55049478 | 0,00303771 | 0,03020925 |
| 116151_at | FAM210B | 0,44247426 | 8,84351416 | 3,54903897 | 0,00304666 | 0,03028263 |
| 26269_at | FBXO8 | 0,5571459 | 7,70329367 | 3,54844217 | 0,00305033 | 0,03030354 |
| 51287_at | COA4 | -0,36399552 | 8,37815276 | -3,5468779 | 0,00305999 | 0,03037975 |
| #N/A | CH17-340M24 | 0,46897349 | 7,96171613 | 3,54668928 | 0,00306116 | 0,03037975 |
| 6584_at | SLC22A5 | 0,61206135 | 8,73318495 | 3,54596075 | 0,00306567 | 0,03040884 |
| 27297_at | CRCP | -0,21466174 | 7,33818811 | -3,545394 | 0,00306918 | 0,03042803 |
| 9961_at | MVP | 1,47881194 | 8,01948679 | 3,54491723 | 0,00307214 | 0,03044169 |

| | | | | | | |
|---|---|---|---|---|---|---|
| 9828_at | ARHGEF17 | 0,42706924 | 7,45649441 | 3,54415535 | 0,00307688 | 0,03047293 |
| 85476_at | GFM1 | -0,35773544 | 8,7119253 | -3,54327241 | 0,00308237 | 0,03051167 |
| 400506_at | KNOP1 | -0,37531846 | 6,90888152 | -3,5428524 | 0,00308499 | 0,0305219 |
| 6448_at | SGSH | 0,46959594 | 5,88563821 | 3,54207018 | 0,00308987 | 0,0305536 |
| 257019_at | FRMD3 | -0,9411845 | 6,02082752 | -3,54181746 | 0,00309145 | 0,0305536 |
| 5001_at | ORC5 | -0,37410484 | 6,77896771 | -3,54143755 | 0,00309382 | 0,0305536 |
| 79056_at | PRRG4 | 0,46454489 | 8,14309058 | 3,54132327 | 0,00309454 | 0,0305536 |
| 26073_at | POLDIP2 | -0,26304758 | 9,3675867 | -3,53990697 | 0,00310341 | 0,03062549 |
| 9439_at | MED23 | 0,42221383 | 7,62116367 | 3,5392126 | 0,00310777 | 0,0306528 |
| 55743_at | CHFR | 0,43524766 | 8,18318791 | 3,53850702 | 0,0031122 | 0,03068084 |
| 3093_at | UBE2K | -0,30314765 | 9,83883743 | -3,53719198 | 0,00312049 | 0,03074676 |
| 10903_at | MTMR11 | 0,41628596 | 6,49166632 | 3,53555871 | 0,0031308 | 0,03083266 |
| 10499_at | NCOA2 | 0,4421149 | 6,30790827 | 3,53499984 | 0,00313434 | 0,03085174 |
| 51566_at | ARMCX3 | 0,60341261 | 7,65400539 | 3,53353257 | 0,00314365 | 0,03092757 |
| 29127_at | RACGAP1 | -0,75121716 | 10,2410417 | -3,53221007 | 0,00315207 | 0,03099453 |
| 7008_at | TEF | 0,36444887 | 6,96828199 | 3,53017292 | 0,00316507 | 0,03110654 |
| 23233_at | EXOC6B | 0,29592351 | 7,89428409 | 3,52958061 | 0,00316887 | 0,0311156 |
| 23390_at | ZDHHC17 | 0,28598075 | 7,36318378 | 3,52952436 | 0,00316923 | 0,0311156 |
| 26959_at | HBP1 | 1,13056212 | 8,23122416 | 3,52630327 | 0,00318993 | 0,03129456 |
| 4677_at | NARS | -0,40648314 | 10,8299446 | -3,52618319 | 0,0031907 | 0,03129456 |
| 55071_at | C9orf40 | -0,63803501 | 7,01282694 | -3,52450086 | 0,00320157 | 0,03138519 |
| 55270_at | NUDT15 | -0,47959587 | 8,76237908 | -3,52374627 | 0,00320646 | 0,03140827 |
| 9819_at | TSC22D2 | 0,30743889 | 7,66214808 | 3,52363397 | 0,00320719 | 0,03140827 |
| 3872_at | KRT17 | 0,38109397 | 12,2027236 | 3,52252289 | 0,0032144 | 0,03146289 |
| 79819_at | WDR78 | 0,39661817 | 3,61184516 | 3,52194801 | 0,00321814 | 0,03148348 |
| 25778_at | DSTYK | 0,34105293 | 6,55602273 | 3,52067118 | 0,00322645 | 0,03154883 |
| 6908_at | TBP | -0,28927059 | 8,47189557 | -3,5201439 | 0,0032299 | 0,03156645 |
| 283991_at | UBALD2 | -0,39443399 | 8,08700096 | -3,51830453 | 0,00324193 | 0,03166199 |
| 5320_at | PLA2G2A | 0,76398211 | 6,81390276 | 3,51814732 | 0,00324296 | 0,03166199 |
| 10149_at | ADGRG2 | 0,50342575 | 3,98328548 | 3,51549051 | 0,00326042 | 0,03181638 |
| 8449_at | DHX16 | -0,21916149 | 8,59205357 | -3,5145993 | 0,0032663 | 0,03185762 |
| 26090_at | ABHD12 | 0,36138546 | 8,86576357 | 3,5139359 | 0,00327069 | 0,0318835 |
| 85028_at | SNHG12 | -0,61171497 | 8,09830593 | -3,51369692 | 0,00327227 | 0,0318835 |
| 55701_at | ARHGEF40 | 0,33305916 | 7,43380902 | 3,51280083 | 0,0032782 | 0,03192239 |
| 5152_at | PDE9A | -0,41299095 | 6,84837472 | -3,51259383 | 0,00327957 | 0,03192239 |
| 64761_at | PARP12 | 0,57333231 | 8,9345884 | 3,51231599 | 0,00328142 | 0,0319242 |
| 91107_at | TRIM47 | -0,36338235 | 6,87033523 | -3,51180545 | 0,0032848 | 0,03194103 |
| 23105_at | FSTL4 | 0,36729328 | 6,71500939 | 3,51101795 | 0,00329004 | 0,03197579 |
| 2176_at | FANCC | -0,45429136 | 6,30586241 | -3,51056616 | 0,00329304 | 0,03197767 |
| 126792_at | B3GALT6 | -0,28300689 | 8,43007303 | -3,51028601 | 0,00329491 | 0,03197767 |
| 23001_at | WDFY3 | 0,49075744 | 6,69195082 | 3,51024062 | 0,00329521 | 0,03197767 |
| 57393_at | TMEM27 | 0,95064712 | 4,71139935 | 3,50996974 | 0,00329702 | 0,03197908 |
| 348793_at | WDR53 | -0,43720358 | 6,81224697 | -3,50870567 | 0,00330545 | 0,03203775 |
| 10011_at | SRA1 | -0,20220811 | 8,37821046 | -3,50856534 | 0,00330639 | 0,03203775 |
| 124739_at | USP43 | 0,35601391 | 5,55506839 | 3,50710432 | 0,00331617 | 0,03211637 |
| 9738_at | CCP110 | -0,61934037 | 8,0689108 | -3,50643302 | 0,00332068 | 0,03214382 |
| 142678_at | MIB2 | 0,33317741 | 7,10655568 | 3,50513219 | 0,00332942 | 0,03221229 |
| 201161_at | CENPV | -1,0686837 | 7,85620804 | -3,50381665 | 0,00333829 | 0,03228187 |
| 22891_at | ZNF365 | -0,79851789 | 4,54545456 | -3,5031882 | 0,00334253 | 0,03230669 |
| 79709_at | COLGALT1 | -0,49284068 | 8,79769397 | -3,50245377 | 0,0033475 | 0,03233847 |

| | | | | | | |
|---|---|---|---|---|---|---|
| 152579_at | SCFD2 | -0,83628875 | 7,35726488 | -3,50098516 | 0,00335746 | 0,03241194 |
| 5993_at | RFX5 | 0,82555759 | 7,98795023 | 3,50083559 | 0,00335847 | 0,03241194 |
| 6341_at | SCO1 | -0,28762566 | 7,70749135 | -3,49914007 | 0,00337 | 0,03247574 |
| 84306_at | PDCD2L | -0,57876756 | 7,67481384 | -3,49912665 | 0,0033701 | 0,03247574 |
| 22906_at | TRAK1 | 0,47523462 | 8,69234498 | 3,49912025 | 0,00337014 | 0,03247574 |
| 163051_at | ZNF709 | 0,35359256 | 4,28793793 | 3,49764992 | 0,00338017 | 0,03255614 |
| 7884_at | SLBP | -0,43338083 | 10,6620349 | -3,49661686 | 0,00338724 | 0,03260791 |
| 8945_at | BTRC | 0,22900437 | 6,93413919 | 3,49569318 | 0,00339357 | 0,03265255 |
| 51752_at | ERAP1 | 0,56662308 | 7,49140374 | 3,49526004 | 0,00339655 | 0,03266484 |
| 254065_at | BRWD3 | 0,42660613 | 6,53330112 | 3,49417116 | 0,00340403 | 0,03272051 |
| 374393_at | FAM111B | -1,01366329 | 7,26654305 | -3,49374467 | 0,00340697 | 0,0327324 |
| 80308_at | FLAD1 | -0,32402099 | 7,63597244 | -3,49313542 | 0,00341117 | 0,03275641 |
| 126789_at | PUSL1 | -0,48600698 | 8,30804587 | -3,49275165 | 0,00341382 | 0,03276551 |
| 54976_at | C20orf27 | -0,62598109 | 8,46644364 | -3,49191567 | 0,00341959 | 0,03280459 |
| 11146_at | GLMN | -0,47448245 | 6,31177791 | -3,49140603 | 0,00342312 | 0,03280822 |
| 2065_at | ERBB3 | 0,88470837 | 8,38312642 | 3,49136864 | 0,00342338 | 0,03280822 |
| 6632_at | SNRPD1 | -0,55048645 | 8,5415106 | -3,49069618 | 0,00342804 | 0,0328365 |
| 1869_at | E2F1 | -0,48868685 | 7,43413019 | -3,49021423 | 0,00343138 | 0,0328365 |
| 7027_at | TFDP1 | -0,48903687 | 10,206491 | -3,49020495 | 0,00343144 | 0,0328365 |
| 79050_at | NOC4L | -0,45761774 | 7,80713745 | -3,48767729 | 0,00344903 | 0,03298837 |
| 100128927_a | ZBTB42 | 0,24569024 | 6,52907215 | 3,48731026 | 0,00345159 | 0,03299168 |
| 55703_at | POLR3B | -0,33115183 | 8,02426 | -3,48713692 | 0,0034528 | 0,03299168 |
| 64223_at | MLST8 | -0,24571102 | 8,35308478 | -3,48592184 | 0,00346129 | 0,03305644 |
| 101928689_a | LINC01424 | -0,67167509 | 4,37391554 | -3,4854356 | 0,0034647 | 0,03307256 |
| 27042_at | DIEXF | -0,5428234 | 7,31769242 | -3,4837435 | 0,00347657 | 0,03315443 |
| 5689_at | PSMB1 | -0,2483536 | 11,1783628 | -3,4837231 | 0,00347671 | 0,03315443 |
| 339263_at | C17orf51 | -0,25362355 | 5,37087154 | -3,48342136 | 0,00347884 | 0,03315825 |
| 8540_at | AGPS | -0,45554652 | 8,31781718 | -3,48259412 | 0,00348466 | 0,03319733 |
| 2108_at | ETFA | -0,37947619 | 10,2197596 | -3,48193641 | 0,0034893 | 0,03322508 |
| 10723_at | SLC12A7 | 0,24517242 | 8,961192 | 3,48089869 | 0,00349663 | 0,03327843 |
| 10137_at | RBM12 | -0,20983413 | 8,10027059 | -3,47911358 | 0,00350927 | 0,03338228 |
| 7029_at | TFDP2 | 0,60230296 | 6,75238165 | 3,47821532 | 0,00351565 | 0,03342646 |
| 11123_at | RCAN3 | 0,33171494 | 7,13475667 | 3,47551743 | 0,00353489 | 0,03359117 |
| 26262_at | TSPAN17 | 0,22911539 | 8,49287911 | 3,4747824 | 0,00354014 | 0,03359117 |
| 5694_at | PSMB6 | -0,85554875 | 10,5638271 | -3,47477924 | 0,00354017 | 0,03359117 |
| 23276_at | KLHL18 | -0,26838434 | 7,40421986 | -3,4746591 | 0,00354103 | 0,03359117 |
| 389136_at | VGLL3 | 0,70137008 | 8,64969674 | 3,47456586 | 0,00354169 | 0,03359117 |
| 9061_at | PAPSS1 | 0,41975823 | 9,21098048 | 3,47356534 | 0,00354887 | 0,03364263 |
| 3293_at | HSD17B3 | 0,51073804 | 6,47133301 | 3,47246451 | 0,00355678 | 0,03370103 |
| 3978_at | LIG1 | -0,60445999 | 8,06579813 | -3,47208953 | 0,00355947 | 0,03371001 |
| 50628_at | GEMIN4 | -0,62085893 | 7,61765346 | -3,47138032 | 0,00356458 | 0,0337418 |
| 54881_at | TEX10 | -0,53763129 | 7,75493545 | -3,47032122 | 0,00357223 | 0,03376914 |
| 23036_at | ZNF292 | 0,38377691 | 7,67651334 | 3,47029769 | 0,0035724 | 0,03376914 |
| 23765_at | IL17RA | 0,20720234 | 7,4222337 | 3,4702515 | 0,00357273 | 0,03376914 |
| 441869_at | ANKRD65 | 0,3735683 | 7,61270408 | 3,46963426 | 0,00357719 | 0,03379473 |
| 23595_at | ORC3 | -0,39407526 | 8,63710984 | -3,46920776 | 0,00358028 | 0,03380357 |
| 8935_at | SKAP2 | 0,55838742 | 9,11390056 | 3,46902012 | 0,00358164 | 0,03380357 |
| 378708_at | APITD1 | -0,52608697 | 6,67437014 | -3,46818228 | 0,00358771 | 0,03384432 |
| 9453_at | GGPS1 | 0,41963445 | 7,52961647 | 3,46748311 | 0,00359279 | 0,03386345 |
| 3140_at | MR1 | 0,83737755 | 7,7310924 | 3,46741869 | 0,00359326 | 0,03386345 |

| | | | | | | |
|---|---|---|---|---|---|---|
| 124975_at | GGT6 | 0,85410203 | 6,67839799 | 3,46678612 | 0,00359786 | 0,03389022 |
| 7629_at | ZNF76 | 0,24266326 | 7,62265673 | 3,46615812 | 0,00360243 | 0,0339167 |
| 54894_at | RNF43 | 0,46404836 | 5,7213547 | 3,4652949 | 0,00360872 | 0,03394815 |
| 80135_at | RPF1 | -0,28839803 | 9,39476156 | -3,46521677 | 0,00360929 | 0,03394815 |
| 101927204_a | LOC10192720 | 0,43261452 | 6,63544115 | 3,46486375 | 0,00361187 | 0,03395581 |
| 983_at | CDK1 | -1,17884674 | 8,96628895 | -3,4634879 | 0,00362193 | 0,03401506 |
| 57573_at | ZNF471 | 0,37687383 | 3,70756878 | 3,46337317 | 0,00362277 | 0,03401506 |
| 144363_at | LYRM5 | 0,45153376 | 7,38509623 | 3,46327823 | 0,00362347 | 0,03401506 |
| 7262_at | PHLDA2 | -0,70610712 | 8,90527117 | -3,46240799 | 0,00362985 | 0,03405838 |
| 5876_at | RABGGTB | -0,67978169 | 7,84233316 | -3,4615357 | 0,00363626 | 0,03410191 |
| 65993_at | MRPS34 | -0,45989512 | 9,39601816 | -3,45926551 | 0,003653 | 0,03423168 |
| 10383_at | TUBB4B | -0,80002351 | 12,2925224 | -3,45917644 | 0,00365365 | 0,03423168 |
| 642_at | BLMH | -0,54890995 | 8,00811627 | -3,45891558 | 0,00365558 | 0,03423309 |
| 10112_at | KIF20A | -0,93955532 | 9,75483622 | -3,45856416 | 0,00365818 | 0,03424079 |
| 84991_at | RBM17 | -0,3127961 | 7,41672784 | -3,45728371 | 0,00366767 | 0,0343129 |
| 79810_at | PTCD2 | -0,47429387 | 6,14922521 | -3,45542454 | 0,00368148 | 0,03442544 |
| 152137_at | CCDC50 | 0,26091012 | 7,14314969 | 3,45488644 | 0,00368549 | 0,03442608 |
| 8933_at | FAM127A | 0,41668672 | 9,343491 | 3,45482286 | 0,00368597 | 0,03442608 |
| 84289_at | ING5 | -0,28334581 | 6,63567847 | -3,45448024 | 0,00368852 | 0,03442608 |
| 27340_at | UTP20 | -0,73373954 | 7,78430874 | -3,45445621 | 0,0036887 | 0,03442608 |
| 1368_at | CPM | 0,33310277 | 3,83468702 | 3,45267855 | 0,00370199 | 0,03452589 |
| 9401_at | RECQL4 | -0,73863518 | 7,85124229 | -3,45237976 | 0,00370422 | 0,03452589 |
| 57348_at | TTYH1 | 0,61962687 | 5,41733255 | 3,45222571 | 0,00370538 | 0,03452589 |
| 4609_at | MYC | -1,06211157 | 9,81647786 | -3,45206743 | 0,00370656 | 0,03452589 |
| 55699_at | IARS2 | -0,30951379 | 9,86313561 | -3,4514063 | 0,00371152 | 0,03455538 |
| 64398_at | MPP5 | 0,38170896 | 8,18071192 | 3,45099494 | 0,00371461 | 0,03456743 |
| 165545_at | DQX1 | 0,49576872 | 5,86868028 | 3,45023995 | 0,00372029 | 0,03460354 |
| 63943_at | FKBPL | -0,26980058 | 6,86632019 | -3,44991621 | 0,00372273 | 0,0346047 |
| 7182_at | NR2C2 | 0,2079093 | 7,91868929 | 3,44974605 | 0,00372401 | 0,0346047 |
| 51083_at | GAL | -1,18569232 | 5,58259351 | -3,44656281 | 0,00374806 | 0,03481142 |
| 81618_at | ITM2C | 0,41346109 | 10,3161957 | 3,44544535 | 0,00375654 | 0,03485988 |
| 5202_at | PFDN2 | -0,57307825 | 9,4728919 | -3,44539834 | 0,0037569 | 0,03485988 |
| 54478_at | FAM64A | -0,74380136 | 7,57982169 | -3,44502657 | 0,00375972 | 0,03486931 |
| 11044_at | PAPD7 | -0,39757811 | 9,11745447 | -3,44369062 | 0,0037699 | 0,03494683 |
| 84265_at | POLR3GL | 0,58021985 | 6,89550323 | 3,4429852 | 0,00377528 | 0,03497514 |
| 51667_at | NUB1 | 0,44534029 | 9,29272799 | 3,44281455 | 0,00377658 | 0,03497514 |
| 200933_at | FBXO45 | -0,5358641 | 7,19077107 | -3,4423001 | 0,00378051 | 0,03499473 |
| 5985_at | RFC5 | -0,90667281 | 8,18521236 | -3,44125332 | 0,00378853 | 0,03502645 |
| 4603_at | MYBL1 | -0,82713946 | 5,96030819 | -3,44096961 | 0,0037907 | 0,03502645 |
| 51147_at | ING4 | 0,60885484 | 7,59289115 | 3,44095304 | 0,00379083 | 0,03502645 |
| 51444_at | RNF138 | -0,39469821 | 7,64449094 | -3,44090272 | 0,00379121 | 0,03502645 |
| 9063_at | PIAS2 | -0,18521716 | 5,76527046 | -3,43779528 | 0,00381512 | 0,03523039 |
| 7278_at | TUBA3C | -0,57324476 | 6,04231298 | -3,43641704 | 0,00382577 | 0,03527503 |
| 57570_at | TRMT5 | -0,34043039 | 7,39764316 | -3,43629939 | 0,00382668 | 0,03527503 |
| 23285_at | KIAA1107 | 0,63131613 | 4,98710679 | 3,43590224 | 0,00382975 | 0,03527503 |
| 5673_at | PSG5 | 0,98127839 | 4,59724299 | 3,43585804 | 0,00383009 | 0,03527503 |
| 1861_at | TOR1A | -0,25975884 | 9,2445857 | -3,4354407 | 0,00383333 | 0,03527503 |
| 3248_at | HPGD | 1,2340491 | 9,74295913 | 3,43535775 | 0,00383397 | 0,03527503 |
| 3108_at | HLA-DMA | 1,07527219 | 7,27340734 | 3,43512812 | 0,00383575 | 0,03527503 |
| 10274_at | STAG1 | -0,22918346 | 8,14043869 | -3,43490713 | 0,00383747 | 0,03527503 |

| | | | | | | |
|---|---|---|---|---|---|---|
| 57700_at | FAM160B1 | 0,64361698 | 7,61739717 | 3,43484481 | 0,00383795 | 0,03527503 |
| 84275_at | SLC25A33 | -0,52057961 | 6,62450795 | -3,43480469 | 0,00383826 | 0,03527503 |
| 51020_at | HDDC2 | -0,45401778 | 7,96838025 | -3,43436495 | 0,00384168 | 0,03528958 |
| 10005_at | ACOT8 | 0,32748425 | 8,59975632 | 3,43408906 | 0,00384382 | 0,03529245 |
| 51351_at | ZNF117 | 0,87898993 | 4,56809122 | 3,43289441 | 0,00385312 | 0,03536098 |
| 79974_at | CPED1 | -0,25496331 | 3,5248928 | -3,43185615 | 0,00386122 | 0,03541844 |
| 132321_at | C4orf33 | 0,80903198 | 6,31526218 | 3,43082552 | 0,00386928 | 0,03547546 |
| 81566_at | CSRNP2 | 0,47858785 | 7,3580878 | 3,43035027 | 0,003873 | 0,03549268 |
| 79134_at | TMEM185B | -0,2266823 | 8,19267223 | -3,42997506 | 0,00387594 | 0,03549387 |
| 56129_at | PCDHB7 | 0,33366781 | 5,00203054 | 3,42986352 | 0,00387682 | 0,03549387 |
| 440335_at | SMIM22 | 0,34505444 | 8,0032605 | 3,4291272 | 0,00388259 | 0,03552739 |
| 54704_at | PDP1 | 0,35675454 | 7,9559526 | 3,42892714 | 0,00388417 | 0,03552739 |
| 5583_at | PRKCH | -0,37542312 | 8,56123974 | -3,42746808 | 0,00389564 | 0,03561533 |
| 51279_at | C1RL | 0,643029 | 7,03666485 | 3,42702815 | 0,00389911 | 0,03561533 |
| 1606_at | DGKA | 0,30612512 | 8,20191191 | 3,42700105 | 0,00389933 | 0,03561533 |
| 151176_at | FAM132B | -0,37215086 | 5,99245554 | -3,42539293 | 0,00391203 | 0,03569056 |
| 10061_at | ABCF2 | -0,39765432 | 6,95195662 | -3,42533671 | 0,00391248 | 0,03569056 |
| 23385_at | NCSTN | 0,36933769 | 8,89774446 | 3,42525508 | 0,00391312 | 0,03569056 |
| 11116_at | FGFR1OP | -0,52992741 | 7,44258688 | -3,42418809 | 0,00392158 | 0,03575074 |
| 8871_at | SYNJ2 | 0,19727847 | 7,49209448 | 3,42387631 | 0,00392405 | 0,03575637 |
| 122525_at | C14orf28 | 0,68410086 | 4,96675835 | 3,42336597 | 0,0039281 | 0,03577638 |
| 147841_at | SPC24 | -0,98189382 | 7,63485771 | -3,42268585 | 0,00393351 | 0,0358087 |
| 83451_at | ABHD11 | -0,30041381 | 6,96598396 | -3,42023693 | 0,00395304 | 0,03595052 |
| 27090_at | ST6GALNAC4 | -0,35042076 | 7,28256472 | -3,42022559 | 0,00395313 | 0,03595052 |
| 284443_at | ZNF493 | 0,31915506 | 4,48798132 | 3,42003112 | 0,00395469 | 0,03595052 |
| 5874_at | RAB27B | 0,53412069 | 7,05431842 | 3,41944254 | 0,0039594 | 0,03597637 |
| 151636_at | DTX3L | 0,70699123 | 9,62110121 | 3,41811381 | 0,00397005 | 0,03605618 |
| 751071_at | METTL12 | -0,36633354 | 6,84503086 | -3,41780658 | 0,00397252 | 0,0360616 |
| 81553_at | FAM49A | 1,02942019 | 6,73733543 | 3,41719458 | 0,00397744 | 0,03608926 |
| 100113407_a | TMEM170B | 0,35254483 | 5,24349361 | 3,4166333 | 0,00398196 | 0,03611324 |
| 9200_at | HACD1 | -0,68773519 | 6,79294018 | -3,41591915 | 0,00398772 | 0,03613779 |
| 283951_at | C16orf91 | -0,33792373 | 7,63752417 | -3,415832 | 0,00398842 | 0,03613779 |
| 9365_at | KL | 0,83457775 | 3,88608444 | 3,41487223 | 0,00399617 | 0,03619099 |
| 81619_at | TSPAN14 | 0,25453057 | 9,89372626 | 3,41424623 | 0,00400123 | 0,03621982 |
| 132864_at | CPEB2 | 0,6050565 | 6,29546916 | 3,41360854 | 0,0040064 | 0,03624953 |
| 1164_at | CKS2 | -0,84809228 | 10,5164226 | -3,41161952 | 0,00402255 | 0,03637858 |
| 440311_at | LOC440311 | 0,50702121 | 9,36284485 | 3,41132099 | 0,00402498 | 0,03638347 |
| 127602_at | DNAH14 | -0,28108954 | 3,24837429 | -3,41108306 | 0,00402692 | 0,03638391 |
| 374_at | AREG | -0,74568657 | 5,73693715 | -3,41047003 | 0,00403191 | 0,03640427 |
| 288_at | ANK3 | 0,4920617 | 8,05307773 | 3,41020218 | 0,0040341 | 0,03640427 |
| 80765_at | STARD5 | 0,85407729 | 7,21763859 | 3,41011137 | 0,00403484 | 0,03640427 |
| 26150_at | RIBC2 | -0,46295447 | 5,21412273 | -3,40961203 | 0,00403892 | 0,036424 |
| 64780_at | MICAL1 | 0,57812532 | 6,90459545 | 3,40761645 | 0,00405525 | 0,03655421 |
| 56926_at | NCLN | -0,63330677 | 8,36419545 | -3,40719579 | 0,00405871 | 0,03656821 |
| 7357_at | UGCG | 0,36597565 | 8,26266995 | 3,40680797 | 0,00406189 | 0,0365798 |
| 30001_at | ERO1A | -0,40370224 | 10,0123683 | -3,40625895 | 0,0040664 | 0,03658897 |
| 65005_at | MRPL9 | -0,5120944 | 9,03812709 | -3,40622197 | 0,00406671 | 0,03658897 |
| 10765_at | KDM5B | 0,58262188 | 9,06208393 | 3,40559346 | 0,00407188 | 0,03661842 |
| 80086_at | TUBA4B | -0,33005054 | 6,0115303 | -3,40510427 | 0,00407591 | 0,03663756 |
| 105375355_a | LOC10537535 | 0,75303725 | 9,2110803 | 3,40408736 | 0,0040843 | 0,03668784 |

| | | | | | | |
|---|---|---|---|---|---|---|
| 222223_at | KIAA1324L | 0,46929847 | 7,10126031 | 3,40365705 | 0,00408786 | 0,03668784 |
| 10036_at | CHAF1A | -0,82339059 | 7,54271034 | -3,40357725 | 0,00408852 | 0,03668784 |
| 163732_at | CITED4 | 0,46430834 | 7,27128045 | 3,40340541 | 0,00408994 | 0,03668784 |
| 339766_at | MROH2A | 0,44043786 | 4,55741035 | 3,40327421 | 0,00409103 | 0,03668784 |
| 3053_at | SERPIND1 | -0,33882857 | 4,14309987 | -3,40243843 | 0,00409795 | 0,03673281 |
| 374918_at | IGFL1 | -1,17901456 | 6,51339241 | -3,40203022 | 0,00410134 | 0,03674606 |
| 7064_at | THOP1 | -0,80441474 | 7,99192431 | -3,40171511 | 0,00410395 | 0,03675239 |
| 29074_at | MRPL18 | -0,53021318 | 9,91142129 | -3,40120618 | 0,00410818 | 0,03675722 |
| 55013_at | CCDC109B | -0,62075494 | 6,20647188 | -3,40119067 | 0,00410831 | 0,03675722 |
| 51669_at | SARAF | 0,31248857 | 10,5708946 | 3,4000391 | 0,00411789 | 0,03682584 |
| 55057_at | AIM1L | 0,35535425 | 7,32147423 | 3,39917069 | 0,00412513 | 0,03685778 |
| 26747_at | NUFIP1 | -0,66420744 | 7,62449787 | -3,39910918 | 0,00412564 | 0,03685778 |
| 25780_at | RASGRP3 | 0,94578771 | 4,06333991 | 3,3989224 | 0,0041272 | 0,03685778 |
| 101410538_a | MMP24-AS1 | 0,41707777 | 8,34063778 | 3,39844278 | 0,00413121 | 0,03686749 |
| 140576_at | S100A16 | -0,28002673 | 11,0936632 | -3,39833403 | 0,00413211 | 0,03686749 |
| 5464_at | PPA1 | -0,24555368 | 11,5529611 | -3,39744146 | 0,00413958 | 0,03687024 |
| 8890_at | EIF2B4 | -0,18329916 | 8,77897788 | -3,3974197 | 0,00413976 | 0,03687024 |
| 1657_at | DMXL1 | 0,4691867 | 7,82842799 | 3,39725887 | 0,00414111 | 0,03687024 |
| 23295_at | MGRN1 | 0,25103209 | 8,2971702 | 3,39725072 | 0,00414118 | 0,03687024 |
| 675_at | BRCA2 | -1,06751988 | 5,65213041 | -3,39715351 | 0,00414199 | 0,03687024 |
| 47_at | ACLY | -0,33089868 | 9,49383868 | -3,39495279 | 0,00416047 | 0,03701763 |
| 64708_at | COPS7B | -0,29154704 | 7,39280328 | -3,39309532 | 0,00417613 | 0,03713982 |
| 2033_at | EP300 | 0,22200037 | 8,43892885 | 3,39279683 | 0,00417866 | 0,03714511 |
| 51629_at | SLC25A39 | -0,31357959 | 9,81044787 | -3,39099745 | 0,00419389 | 0,03725387 |
| 2517_at | FUCA1 | 0,74811874 | 10,1634402 | 3,39089535 | 0,00419476 | 0,03725387 |
| 9588_at | PRDX6 | -0,29266797 | 11,3544774 | -3,3905914 | 0,00419734 | 0,0372596 |
| 79039_at | DDX54 | -0,31421448 | 7,61174624 | -3,38994429 | 0,00420284 | 0,03729122 |
| 57707_at | TLDC1 | 0,26444033 | 7,74302611 | 3,38912007 | 0,00420985 | 0,03733624 |
| 4882_at | NPR2 | -0,2429103 | 5,05633063 | -3,38792893 | 0,00422 | 0,03740909 |
| 9646_at | CTR9 | 0,21615382 | 8,7097216 | 3,38729139 | 0,00422545 | 0,03742642 |
| 91801_at | ALKBH8 | -0,28528582 | 5,72240686 | -3,38724512 | 0,00422585 | 0,03742642 |
| 9777_at | TM9SF4 | 0,22353813 | 9,04356019 | 3,38638127 | 0,00423324 | 0,03747149 |
| 9841_at | ZBTB24 | -0,27547357 | 5,06332901 | -3,38612489 | 0,00423543 | 0,03747149 |
| 7556_at | ZNF10 | 0,23558328 | 4,90159608 | 3,38574193 | 0,00423871 | 0,03747149 |
| 80833_at | APOL3 | 0,83693311 | 7,12266031 | 3,38574173 | 0,00423872 | 0,03747149 |
| 4726_at | NDUFS6 | -0,34890379 | 10,0126159 | -3,38534168 | 0,00424215 | 0,03748463 |
| 23731_at | TMEM245 | 0,45985913 | 9,42451915 | 3,3850448 | 0,0042447 | 0,03748779 |
| 580_at | BARD1 | -0,64520044 | 7,44891774 | -3,38468242 | 0,00424781 | 0,03748779 |
| 868_at | CBLB | 0,67779245 | 7,52265611 | 3,3845657 | 0,00424881 | 0,03748779 |
| 126068_at | ZNF441 | 0,25216535 | 4,01805681 | 3,38433522 | 0,00425079 | 0,03748779 |
| 136306_at | SVOPL | 0,38057579 | 3,9877984 | 3,38395915 | 0,00425403 | 0,03748779 |
| 57639_at | CCDC146 | 0,78491397 | 4,79872997 | 3,38394121 | 0,00425418 | 0,03748779 |
| 735301_at | SNHG9 | -0,4775166 | 6,12608662 | -3,38195865 | 0,00427128 | 0,03759537 |
| 204219_at | CERS3 | 0,66626421 | 5,29629082 | 3,38186457 | 0,00427209 | 0,03759537 |
| 192670_at | AGO4 | 0,7209072 | 5,99495261 | 3,38184649 | 0,00427225 | 0,03759537 |
| 3107_at | HLA-C | 0,6111453 | 11,2889839 | 3,38160176 | 0,00427436 | 0,03759681 |
| 956_at | ENTPD3 | 0,71429384 | 6,65183452 | 3,38098628 | 0,00427969 | 0,03760134 |
| 9265_at | CYTH3 | 0,30818459 | 6,29150278 | 3,38093393 | 0,00428014 | 0,03760134 |
| 79968_at | WDR76 | -1,07844381 | 6,32084945 | -3,38086548 | 0,00428073 | 0,03760134 |
| 7276_at | TTR | -0,33042435 | 5,53446315 | -3,38059409 | 0,00428308 | 0,03760204 |

| | | | | | | |
|---|---|---|---|---|---|---|
| 10413_at | YAP1 | 0,35069098 | 9,64121287 | 3,38024141 | 0,00428614 | 0,03760204 |
| 8777_at | MPDZ | 0,4980895 | 7,3388596 | 3,3799009 | 0,00428909 | 0,03760204 |
| 57146_at | TMEM159 | 0,54108738 | 7,33437926 | 3,37971682 | 0,00429069 | 0,03760204 |
| 10682_at | EBP | -0,63730122 | 9,6132865 | -3,37938416 | 0,00429358 | 0,03760204 |
| 6697_at | SPR | -0,38421049 | 8,944645 | -3,37937075 | 0,00429369 | 0,03760204 |
| 27097_at | TAF5L | -0,48177842 | 7,47094977 | -3,37928068 | 0,00429448 | 0,03760204 |
| 11226_at | GALNT6 | 0,29484385 | 6,08421753 | 3,37878758 | 0,00429876 | 0,03762246 |
| 124045_at | SPATA33 | -0,41518052 | 6,73309988 | -3,37791043 | 0,0043064 | 0,03767216 |
| 23135_at | KDM6B | 0,21941778 | 6,96316245 | 3,37741733 | 0,00431069 | 0,03769264 |
| 56204_at | FAM214A | 0,91467144 | 9,08992405 | 3,37593633 | 0,00432363 | 0,03778857 |
| 64762_at | GAREM | 0,67605099 | 8,11708449 | 3,37568998 | 0,00432578 | 0,03779026 |
| 7726_at | TRIM26 | 0,3386665 | 8,56048497 | 3,37529265 | 0,00432926 | 0,0378035 |
| 91860_at | CALML4 | -0,70981603 | 6,01749462 | -3,37400052 | 0,00434059 | 0,03788526 |
| 29089_at | UBE2T | -1,22917827 | 9,00957058 | -3,3728783 | 0,00435045 | 0,03794142 |
| 54968_at | TMEM70 | -0,28009621 | 8,28007554 | -3,37274472 | 0,00435163 | 0,03794142 |
| 51451_at | LCMT1 | 0,21228743 | 8,83108318 | 3,37259656 | 0,00435293 | 0,03794142 |
| 7555_at | CNBP | -0,23250685 | 8,88075089 | -3,37091278 | 0,00436778 | 0,03805365 |
| 8500_at | PPFIA1 | -0,3102925 | 7,78713618 | -3,36918426 | 0,00438308 | 0,03816967 |
| 79918_at | SETD6 | -0,33115285 | 7,30044594 | -3,36844806 | 0,00438961 | 0,03819473 |
| 6643_at | SNX2 | 0,46115778 | 9,32390132 | 3,36837508 | 0,00439026 | 0,03819473 |
| 3099_at | HK2 | -0,66185778 | 8,04913694 | -3,36818976 | 0,00439191 | 0,03819473 |
| 11321_at | GPN1 | -0,40224024 | 8,48467556 | -3,36724393 | 0,00440032 | 0,0382506 |
| 157378_at | TMEM65 | -0,38967515 | 7,2691599 | -3,36665724 | 0,00440554 | 0,03827875 |
| 4218_at | RAB8A | -0,24624388 | 9,81364961 | -3,36613937 | 0,00441016 | 0,03830116 |
| 27302_at | BMP10 | -0,32293258 | 4,13049909 | -3,3659221 | 0,0044121 | 0,03830116 |
| 83475_at | DOHH | -0,64867336 | 6,87180839 | -3,36535661 | 0,00441715 | 0,03832772 |
| 60685_at | ZFAND3 | 0,43171879 | 8,13510351 | 3,36488453 | 0,00442137 | 0,03834467 |
| 2739_at | GLO1 | -0,38535666 | 10,7070393 | -3,36439748 | 0,00442573 | 0,03834467 |
| 440288_at | UBL7-AS1 | -0,46251101 | 6,11401522 | -3,36431035 | 0,00442651 | 0,03834467 |
| 9589_at | WTAP | -0,22490795 | 9,36271574 | -3,36424804 | 0,00442706 | 0,03834467 |
| 7009_at | TMBIM6 | 0,2456345 | 12,1568481 | 3,36370562 | 0,00443192 | 0,03836951 |
| 1259_at | CNGA1 | 0,84660447 | 5,83050534 | 3,36310299 | 0,00443733 | 0,03839905 |
| 404093_at | CUEDC1 | 0,3598699 | 7,03841703 | 3,36258704 | 0,00444196 | 0,03841037 |
| 7766_at | ZNF223 | 0,48681151 | 5,35619049 | 3,36251327 | 0,00444263 | 0,03841037 |
| 100507524_a | ARHGEF26-AS | 1,24041551 | 5,48474057 | 3,36122026 | 0,00445426 | 0,03849369 |
| 146050_at | ZSCAN29 | 0,23645268 | 7,31354553 | 3,35690485 | 0,00449331 | 0,03881375 |
| 22807_at | IKZF2 | 0,36262764 | 6,19910263 | 3,35624589 | 0,0044993 | 0,03884809 |
| 64760_at | FAM160B2 | 0,23601538 | 6,93452177 | 3,35567544 | 0,0045045 | 0,03887552 |
| 51447_at | IP6K2 | 0,42481494 | 8,10115475 | 3,35524072 | 0,00450846 | 0,03889229 |
| 3930_at | LBR | -0,61737691 | 10,2868535 | -3,3543458 | 0,00451663 | 0,03894532 |
| 25766_at | PRPF40B | 0,29450152 | 6,72298618 | 3,3532724 | 0,00452645 | 0,0390125 |
| 286437_at | LOC286437 | 0,22549662 | 6,09237014 | 3,35290597 | 0,0045298 | 0,03902397 |
| 79072_at | FASTKD3 | -0,27168964 | 7,82021994 | -3,35267708 | 0,0045319 | 0,03902458 |
| 9212_at | AURKB | -0,98674346 | 7,95687631 | -3,35219785 | 0,0045363 | 0,03904498 |
| 55272_at | IMP3 | -0,3899068 | 8,62255127 | -3,3512 | 0,00454546 | 0,03910639 |
| 9262_at | STK17B | -0,27910788 | 6,61388614 | -3,35057253 | 0,00455123 | 0,03913857 |
| 4670_at | HNRNPM | -0,33861947 | 10,7131812 | -3,35011958 | 0,0045554 | 0,03915696 |
| 7294_at | TXK | 0,49543266 | 5,70519154 | 3,34927156 | 0,00456322 | 0,03916284 |
| 84124_at | ZNF394 | 0,30577894 | 7,62255262 | 3,3491217 | 0,00456461 | 0,03916284 |
| 79258_at | MMEL1 | 0,57441008 | 7,09537347 | 3,34899028 | 0,00456582 | 0,03916284 |

| | | | | | | |
|---|---|---|---|---|---|---|
| 55858_at | TMEM165 | 0,38260006 | 9,37776101 | 3,34895628 | 0,00456614 | 0,03916284 |
| 4714_at | NDUFB8 | -0,1912075 | 8,91191885 | -3,3489436 | 0,00456625 | 0,03916284 |
| 427_at | ASAH1 | 0,63809695 | 9,95598483 | 3,34765947 | 0,00457813 | 0,03924721 |
| 79932_at | KIAA0319L | 0,44299639 | 8,38974593 | 3,34621874 | 0,00459149 | 0,03934424 |
| 5214_at | PFKP | -0,64941391 | 9,61759905 | -3,34554505 | 0,00459775 | 0,03938037 |
| 283219_at | KCTD21 | 0,58965384 | 6,32894215 | 3,3442659 | 0,00460966 | 0,03946484 |
| 4033_at | LRMP | 0,62752831 | 7,6719105 | 3,3429201 | 0,00462223 | 0,03955483 |
| 56133_at | PCDHB2 | 0,6100193 | 7,43566683 | 3,34203055 | 0,00463055 | 0,03957428 |
| 1138_at | CHRNA5 | -0,91415605 | 7,89176322 | -3,3418565 | 0,00463218 | 0,03957428 |
| 23177_at | CEP68 | 0,35483292 | 6,5671518 | 3,34181935 | 0,00463253 | 0,03957428 |
| 80310_at | PDGFD | 0,38486985 | 5,37575674 | 3,34179934 | 0,00463272 | 0,03957428 |
| 7379_at | UPK2 | 0,92801335 | 6,73041468 | 3,34097809 | 0,00464042 | 0,0396225 |
| 26994_at | RNF11 | 0,36046583 | 9,83448911 | 3,34046992 | 0,00464519 | 0,03964567 |
| 3070_at | HELLS | -1,10796771 | 7,44624575 | -3,33968132 | 0,0046526 | 0,03967919 |
| 57567_at | ZNF319 | 0,22042941 | 7,16024195 | 3,33961416 | 0,00465324 | 0,03967919 |
| 5933_at | RBL1 | -0,67491909 | 5,45086172 | -3,33814395 | 0,00466709 | 0,03977975 |
| 59338_at | PLEKHA1 | 0,30617861 | 8,35420697 | 3,33583717 | 0,00468892 | 0,0399481 |
| 57820_at | CCNB1IP1 | -0,32067684 | 7,98312049 | -3,33491297 | 0,00469769 | 0,04000515 |
| 4668_at | NAGA | 0,28577767 | 8,14845304 | 3,33434979 | 0,00470305 | 0,04003305 |
| 9601_at | PDIA4 | -0,29840968 | 10,4577087 | -3,33221988 | 0,00472335 | 0,04018703 |
| 1019_at | CDK4 | -0,59120627 | 10,7752218 | -3,33201487 | 0,00472531 | 0,04018703 |
| 373_at | TRIM23 | 0,44821565 | 5,55902523 | 3,3317546 | 0,0047278 | 0,04018778 |
| 55247_at | NEIL3 | -1,04976084 | 6,76412852 | -3,33156933 | 0,00472957 | 0,04018778 |
| 118980_at | SFXN2 | -0,39386604 | 7,45553586 | -3,33134174 | 0,00473175 | 0,04018855 |
| 4751_at | NEK2 | -0,86002883 | 8,05372789 | -3,32994209 | 0,00474516 | 0,04027652 |
| 53918_at | PELO | -0,38456399 | 8,48305089 | -3,32982487 | 0,00474629 | 0,04027652 |
| 2146_at | EZH2 | -0,77014444 | 8,50388497 | -3,32907127 | 0,00475353 | 0,04032019 |
| 55229_at | PANK4 | -0,22954748 | 7,65746096 | -3,32870643 | 0,00475703 | 0,0403322 |
| 23764_at | MAFF | -0,88753986 | 7,82239456 | -3,32815111 | 0,00476238 | 0,04035976 |
| 253980_at | KCTD13 | 0,37372155 | 6,24106248 | 3,32704854 | 0,00477301 | 0,04041681 |
| 10955_at | SERINC3 | 0,29383663 | 11,1227529 | 3,32701781 | 0,00477331 | 0,04041681 |
| 55163_at | PNPO | -0,25576475 | 8,15481904 | -3,32573175 | 0,00478574 | 0,04050427 |
| 51614_at | ERGIC3 | 0,26899418 | 11,4390891 | 3,32527423 | 0,00479017 | 0,04052396 |
| 6834_at | SURF1 | 0,18866226 | 8,78861027 | 3,32335041 | 0,00480885 | 0,04066408 |
| 100507557_a | LOC10050755 | 0,48520427 | 4,09240069 | 3,322494 | 0,00481718 | 0,0406872 |
| 267_at | AMFR | 0,2802346 | 9,0044951 | 3,32227385 | 0,00481933 | 0,0406872 |
| 9325_at | TRIP4 | 0,32095476 | 8,25870463 | 3,32220417 | 0,00482001 | 0,0406872 |
| 8718_at | TNFRSF25 | 0,23894869 | 6,66776228 | 3,32179038 | 0,00482404 | 0,0406872 |
| 27044_at | SND1 | -0,2042765 | 9,10519945 | -3,32176346 | 0,0048243 | 0,0406872 |
| 79801_at | SHCBP1 | -0,99449532 | 7,62188555 | -3,3215679 | 0,00482621 | 0,0406872 |
| 83985_at | SPNS1 | 0,28211083 | 8,77167116 | 3,32155233 | 0,00482636 | 0,0406872 |
| 51637_at | C14orf166 | -0,28918761 | 11,2597405 | -3,31933079 | 0,0048481 | 0,04085254 |
| 23234_at | DNAJC9 | -0,92569047 | 7,83268125 | -3,31649134 | 0,00487602 | 0,04106985 |
| 284004_at | HEXDC | 0,40853314 | 7,36141813 | 3,31612491 | 0,00487963 | 0,04108233 |
| 11130_at | ZWINT | -1,12060213 | 9,80636308 | -3,31528424 | 0,00488794 | 0,04113426 |
| 134145_at | FAM173B | -0,51404624 | 7,46340339 | -3,31393484 | 0,00490129 | 0,04122865 |
| 79101_at | TAF1D | -0,42855302 | 8,8842076 | -3,31329143 | 0,00490768 | 0,04126432 |
| 10513_at | APPBP2 | 0,40582993 | 8,19342149 | 3,3120045 | 0,00492046 | 0,0413538 |
| 219541_at | MED19 | 0,20024513 | 7,73084996 | 3,3110447 | 0,00493003 | 0,04140184 |
| 5476_at | CTSA | 0,34141275 | 10,7075964 | 3,31080829 | 0,00493238 | 0,04140184 |

| | | | | | | |
|---|---|---|---|---|---|---|
| 28971_at | AAMDC | 0,49920752 | 6,73446794 | 3,31078368 | 0,00493263 | 0,04140184 |
| 5695_at | PSMB7 | -0,33245392 | 10,7119794 | -3,30799059 | 0,00496057 | 0,04161822 |
| 266655_at | LINC00094 | 0,40850188 | 7,51296212 | 3,30699512 | 0,00497057 | 0,04166745 |
| 90488_at | TMEM263 | 0,46942009 | 9,17774946 | 3,30680809 | 0,00497245 | 0,04166745 |
| 51668_at | HSPB11 | -0,42225433 | 9,6242216 | -3,30649607 | 0,00497558 | 0,04166745 |
| 7695_at | ZNF136 | 0,33710755 | 6,84320045 | 3,30645256 | 0,00497602 | 0,04166745 |
| 147727_at | ILF3-AS1 | -0,51788585 | 7,15220194 | -3,30633041 | 0,00497725 | 0,04166745 |
| 83939_at | EIF2A | -0,34518447 | 10,3182115 | -3,30541531 | 0,00498647 | 0,04172649 |
| 444_at | ASPH | -0,3804531 | 9,20886409 | -3,30418468 | 0,0049989 | 0,0418123 |
| 89978_at | DPH6 | -0,29824937 | 5,36399644 | -3,30099177 | 0,00503128 | 0,04206489 |
| 23760_at | PITPNB | -0,30265163 | 10,0057804 | -3,30033627 | 0,00503795 | 0,04210241 |
| 83638_at | C11orf68 | 0,21816107 | 8,52991067 | 3,29990767 | 0,00504232 | 0,04211669 |
| 55142_at | HAUS2 | -0,45847431 | 6,8093551 | -3,2997397 | 0,00504403 | 0,04211669 |
| 5184_at | PEPD | 0,53900102 | 9,62716171 | 3,29924799 | 0,00504905 | 0,04214032 |
| 23023_at | TMCC1 | 0,58375049 | 7,61200038 | 3,29890751 | 0,00505253 | 0,04215108 |
| 25975_at | EGFL6 | 0,56988051 | 4,01282395 | 3,29521519 | 0,00509039 | 0,0424486 |
| 23558_at | WBP2 | 0,30187598 | 7,98670701 | 3,29265035 | 0,00511686 | 0,04265086 |
| 9637_at | FEZ2 | 0,47836342 | 9,54157745 | 3,29082446 | 0,00513579 | 0,04279011 |
| 54933_at | RHBDL2 | 0,44312205 | 6,96930761 | 3,28723909 | 0,00517316 | 0,04308282 |
| 83643_at | CCDC3 | -0,23291916 | 6,15829296 | -3,28677096 | 0,00517806 | 0,04310499 |
| 4976_at | OPA1 | -0,30608041 | 8,47806283 | -3,28638619 | 0,00518209 | 0,0431199 |
| 1349_at | COX7B | -0,27004597 | 10,6164899 | -3,28596306 | 0,00518652 | 0,04313818 |
| 10527_at | IPO7 | -0,22304302 | 9,414859 | -3,28537628 | 0,00519268 | 0,04317075 |
| 5129_at | CDK18 | 0,31181726 | 6,3132826 | 3,2842149 | 0,00520489 | 0,04324762 |
| 54468_at | MIOS | -0,37019327 | 7,65352167 | -3,28406969 | 0,00520642 | 0,04324762 |
| 262_at | AMD1 | -0,99617794 | 9,51577251 | -3,28323339 | 0,00521523 | 0,04326908 |
| 23303_at | KIF13B | 0,25294812 | 6,54607479 | 3,28320086 | 0,00521557 | 0,04326908 |
| 6709_at | SPTAN1 | 0,28493305 | 8,46330692 | 3,28318495 | 0,00521574 | 0,04326908 |
| 7311_at | UBA52 | -0,19882901 | 11,0328534 | -3,2807032 | 0,00524198 | 0,04346803 |
| 6117_at | RPA1 | -0,38086385 | 8,88666178 | -3,27997577 | 0,00524969 | 0,04351235 |
| 11339_at | OIP5 | -1,19081315 | 8,3820829 | -3,27962719 | 0,0052534 | 0,04351235 |
| 149466_at | C1orf210 | 0,65897579 | 6,5101363 | 3,27956092 | 0,0052541 | 0,04351235 |
| 51645_at | PPIL1 | -0,54764382 | 8,97100402 | -3,27884113 | 0,00526175 | 0,04355699 |
| 25911_at | DPCD | -0,28727704 | 8,38621278 | -3,27848931 | 0,0052655 | 0,04356926 |
| 54982_at | CLN6 | -0,47891395 | 8,35394962 | -3,27746048 | 0,00527646 | 0,04364124 |
| 84959_at | UBASH3B | -0,50990779 | 7,00134219 | -3,27721439 | 0,00527909 | 0,04364422 |
| 1676_at | DFFA | -0,23569621 | 8,27250207 | -3,27613059 | 0,00529067 | 0,04370848 |
| 2147_at | F2 | -0,37028719 | 4,88119131 | -3,27605283 | 0,0052915 | 0,04370848 |
| 5909_at | RAP1GAP | -0,23768812 | 7,14616075 | -3,27585041 | 0,00529367 | 0,04370848 |
| 199745_at | THAP8 | 0,33554003 | 6,80274929 | 3,27137194 | 0,00534182 | 0,04408714 |
| 100506190_a | LINC00963 | 0,37059876 | 6,56688043 | 3,27081978 | 0,00534778 | 0,04411748 |
| 92002_at | FAM58A | -0,43370417 | 8,07738459 | -3,27034157 | 0,00535295 | 0,04414125 |
| 54998_at | AURKAIP1 | -0,28880335 | 9,6164367 | -3,26956302 | 0,00536139 | 0,04419187 |
| 5321_at | PLA2G4A | -0,52686593 | 5,3270298 | -3,26912644 | 0,00536612 | 0,04421197 |
| 144100_at | PLEKHA7 | 0,48942699 | 8,72347247 | 3,268295 | 0,00537515 | 0,04423311 |
| 3009_at | HIST1H1B | -0,52596139 | 5,34960665 | -3,26828855 | 0,00537522 | 0,04423311 |
| 10039_at | PARP3 | 0,29674912 | 6,89635542 | 3,26825577 | 0,00537558 | 0,04423311 |
| 8727_at | CTNNAL1 | -0,77304757 | 8,36340066 | -3,26784223 | 0,00538007 | 0,0442512 |
| 254531_at | LPCAT4 | 0,17303731 | 8,82997306 | 3,2663505 | 0,00539632 | 0,04436592 |
| 57343_at | ZNF304 | 0,28021291 | 7,05585131 | 3,2657721 | 0,00540264 | 0,0443833 |

| | | | | | | |
|---|---|---|---|---|---|---|
| 54816_at | ZNF280D | 0,42391932 | 6,42848836 | 3,26573469 | 0,00540304 | 0,0443833 |
| 79616_at | CCNJL | 0,51028466 | 7,27198862 | 3,26465513 | 0,00541485 | 0,04443645 |
| 79158_at | GNPTAB | 0,54690611 | 8,82393556 | 3,26449527 | 0,0054166 | 0,04443645 |
| 101410533_a | YTHDF3-AS1 | -0,29851153 | 5,3197513 | -3,26447492 | 0,00541682 | 0,04443645 |
| 10865_at | ARID5A | -0,31218618 | 6,2938552 | -3,26429964 | 0,00541874 | 0,04443645 |
| 10209_at | EIF1 | -0,25954051 | 11,5818395 | -3,26262928 | 0,00543707 | 0,04456779 |
| 80024_at | SLC8B1 | 0,33830087 | 6,79249442 | 3,26196948 | 0,00544433 | 0,04460829 |
| 200407_at | CREG2 | -0,2837713 | 4,85931651 | -3,26130609 | 0,00545163 | 0,04463528 |
| 55173_at | MRPS10 | -0,38157316 | 9,358321 | -3,26115054 | 0,00545335 | 0,04463528 |
| 221584_at | ZSCAN12P1 | -0,29815434 | 4,79439326 | -3,26103948 | 0,00545457 | 0,04463528 |
| 5118_at | PCOLCE | 0,31424323 | 6,65219663 | 3,2606282 | 0,00545911 | 0,04465344 |
| 7107_at | GPR137B | 0,79105678 | 7,86200559 | 3,25946367 | 0,00547198 | 0,04473969 |
| 10635_at | RAD51AP1 | -0,99218309 | 7,94294839 | -3,25872707 | 0,00548013 | 0,04478735 |
| 79959_at | CEP76 | -0,44447403 | 6,72198818 | -3,25824947 | 0,00548543 | 0,0448116 |
| 136_at | ADORA2B | -1,13684612 | 8,75179711 | -3,25770063 | 0,00549152 | 0,04484233 |
| 376940_at | ZC3H6 | 0,46033508 | 5,60914659 | 3,25731165 | 0,00549583 | 0,04485025 |
| 23253_at | ANKRD12 | 0,41523763 | 7,17960323 | 3,25709894 | 0,0054982 | 0,04485025 |
| 114041_at | B3GALT5-AS1 | 0,2852182 | 5,36663487 | 3,25698458 | 0,00549947 | 0,04485025 |
| 284352_at | PPP1R37 | 0,3332924 | 7,26198399 | 3,25587539 | 0,00551181 | 0,04492326 |
| 865_at | CBFB | -0,3737041 | 9,60912826 | -3,25576127 | 0,00551309 | 0,04492326 |
| 5681_at | PSKH1 | 0,35899227 | 7,42487922 | 3,2528376 | 0,00554577 | 0,04517044 |
| 23450_at | SF3B3 | -0,35443428 | 8,51195692 | -3,2513474 | 0,0055625 | 0,04525743 |
| 54675_at | CRLS1 | -0,22502496 | 7,89255886 | -3,25114053 | 0,00556482 | 0,04525743 |
| 4302_at | MLLT6 | 0,35425758 | 7,71796022 | 3,25109292 | 0,00556536 | 0,04525743 |
| 4926_at | NUMA1 | 0,22594316 | 7,94631307 | 3,25104981 | 0,00556584 | 0,04525743 |
| 51143_at | DYNC1LI1 | -0,30010411 | 8,08200509 | -3,25052909 | 0,00557171 | 0,04528314 |
| 65018_at | PINK1 | 0,56642561 | 8,36946001 | 3,25035137 | 0,00557371 | 0,04528314 |
| 7079_at | TIMP4 | -0,98243732 | 6,55018205 | -3,24960941 | 0,00558207 | 0,04533198 |
| 1591_at | CYP24A1 | -1,02374532 | 6,20847743 | -3,24697235 | 0,00561191 | 0,04555506 |
| 9903_at | KLHL21 | -0,34435872 | 6,78076795 | -3,24361416 | 0,00565013 | 0,04584601 |
| 221477_at | C6orf89 | 0,1899965 | 8,11542547 | 3,24249102 | 0,00566297 | 0,04591428 |
| 7490_at | WT1 | -0,58282858 | 5,63693856 | -3,24246137 | 0,00566331 | 0,04591428 |
| 2034_at | EPAS1 | 0,38194643 | 10,7948833 | 3,24210991 | 0,00566733 | 0,04592758 |
| 310_at | ANXA7 | -0,30001161 | 9,89904999 | -3,2413678 | 0,00567584 | 0,04597718 |
| 5411_at | PNN | -0,4404681 | 9,07034599 | -3,24086001 | 0,00568167 | 0,04600504 |
| 9060_at | PAPSS2 | 0,56584369 | 7,08601912 | 3,23960626 | 0,00569609 | 0,04610239 |
| 54943_at | DNAJC28 | 0,65809797 | 4,44749038 | 3,2362925 | 0,00573436 | 0,04637589 |
| 2237_at | FEN1 | -1,13430415 | 8,47677011 | -3,23597503 | 0,00573804 | 0,04637589 |
| 3398_at | ID2 | 0,96750683 | 9,48077059 | 3,23592123 | 0,00573867 | 0,04637589 |
| 150465_at | TTL | -0,24529639 | 8,08840223 | -3,23584879 | 0,00573951 | 0,04637589 |
| 10606_at | PAICS | -1,03692505 | 10,2166261 | -3,23524596 | 0,0057465 | 0,04641296 |
| 647024_at | C6orf132 | 0,36007293 | 7,5334075 | 3,23455187 | 0,00575457 | 0,04645864 |
| 23389_at | MED13L | 0,30023244 | 8,09744985 | 3,23365669 | 0,00576499 | 0,04652327 |
| 51514_at | DTL | -1,43046655 | 8,46417131 | -3,23222286 | 0,00578172 | 0,04663874 |
| 8091_at | HMGA2 | -0,29910555 | 3,91084862 | -3,23196337 | 0,00578475 | 0,04664367 |
| 55746_at | NUP133 | -0,26830735 | 7,58167258 | -3,23174715 | 0,00578728 | 0,04664453 |
| 51315_at | KRCC1 | 0,62862482 | 8,0731965 | 3,23152444 | 0,00578989 | 0,04664601 |
| 23351_at | KHNYN | 0,31251014 | 8,24406483 | 3,23083075 | 0,00579801 | 0,04669193 |
| 89848_at | FCHSD1 | 0,22791738 | 6,11329384 | 3,23007511 | 0,00580687 | 0,04672855 |
| 100288637_a | LOC10028863 | -0,71662359 | 6,76992659 | -3,2299773 | 0,00580802 | 0,04672855 |

| | | | | | | |
|---|---|---|---|---|---|---|
| 3654_at | IRAK1 | -0,47317199 | 9,06084866 | -3,22971198 | 0,00581113 | 0,04672855 |
| 521_at | ATP5I | -0,33209096 | 10,768676 | -3,22961598 | 0,00581226 | 0,04672855 |
| 14_at | AAMP | -0,19327374 | 9,5179663 | -3,22938442 | 0,00581498 | 0,04673092 |
| 27292_at | DIMT1 | -0,61275508 | 8,523942 | -3,22851629 | 0,00582519 | 0,04676762 |
| 83903_at | GSG2 | -0,86522305 | 5,96330152 | -3,22823081 | 0,00582856 | 0,04676762 |
| 220988_at | HNRNPA3 | -0,3726409 | 8,09187524 | -3,22820614 | 0,00582885 | 0,04676762 |
| 124359_at | CDYL2 | 0,25947958 | 6,96759801 | 3,22817091 | 0,00582926 | 0,04676762 |
| 55110_at | MAGOHB | -0,34740876 | 5,92191913 | -3,22630354 | 0,0058513 | 0,04691665 |
| 55658_at | RNF126 | -0,34456135 | 8,65063293 | -3,22618459 | 0,00585271 | 0,04691665 |
| 4240_at | MFGE8 | 0,40612242 | 6,89860729 | 3,22587768 | 0,00585634 | 0,04692623 |
| 83872_at | HMCN1 | 0,87705987 | 4,20556637 | 3,2254419 | 0,0058615 | 0,04692702 |
| 64172_at | OSGEPL1 | -0,44835768 | 6,39602936 | -3,22542993 | 0,00586164 | 0,04692702 |
| 4191_at | MDH2 | -0,20405191 | 10,1589267 | -3,22505336 | 0,0058661 | 0,04692702 |
| 26278_at | SACS | -0,80947799 | 6,08537033 | -3,22500355 | 0,00586669 | 0,04692702 |
| 8869_at | ST3GAL5 | 0,47552148 | 8,64785945 | 3,22484114 | 0,00586862 | 0,04692702 |
| 150468_at | CKAP2L | -1,046251 | 7,39821919 | -3,22452123 | 0,00587241 | 0,04693788 |
| 338645_at | LUZP2 | 0,3289405 | 6,03305848 | 3,22422068 | 0,00587598 | 0,04694692 |
| 5329_at | PLAUR | -0,66021599 | 7,06222912 | -3,22371093 | 0,00588204 | 0,04697582 |
| 7620_at | ZNF69 | 0,53034479 | 6,60069731 | 3,22300229 | 0,00589047 | 0,04702364 |
| 64841_at | GNPNAT1 | -0,61202144 | 9,2432415 | -3,22261452 | 0,00589508 | 0,04704101 |
| 162394_at | SLFN5 | 0,89117979 | 7,83737758 | 3,22239047 | 0,00589775 | 0,04704283 |
| 3308_at | HSPA4 | -0,59894968 | 9,68743627 | -3,22195283 | 0,00590297 | 0,04705544 |
| 11270_at | NRM | -0,50458741 | 9,17703542 | -3,22184816 | 0,00590422 | 0,04705544 |
| 29952_at | DPP7 | 0,32379673 | 9,00233506 | 3,22159611 | 0,00590723 | 0,04705994 |
| 1519_at | CTSO | 1,13937611 | 6,33115782 | 3,22018252 | 0,00592413 | 0,04717505 |
| 5810_at | RAD1 | -0,39829911 | 7,57412761 | -3,21975868 | 0,0059292 | 0,04719596 |
| 1063_at | CENPF | -0,89705172 | 7,87371116 | -3,21875155 | 0,00594128 | 0,04727257 |
| 64963_at | MRPS11 | -0,16918329 | 7,12369977 | -3,21815639 | 0,00594843 | 0,04730991 |
| 283643_at | C14orf80 | -0,57880877 | 7,33629531 | -3,21743539 | 0,0059571 | 0,04735933 |
| 150297_at | C22orf42 | -0,28297235 | 3,87535221 | -3,21722563 | 0,00595963 | 0,04735986 |
| 92799_at | SHKBP1 | 0,25241266 | 7,77173184 | 3,21671161 | 0,00596582 | 0,04737732 |
| 3835_at | KIF22 | -0,73445232 | 8,41294878 | -3,21660649 | 0,00596709 | 0,04737732 |
| 54622_at | ARL15 | 0,64936691 | 6,6759939 | 3,21643117 | 0,0059692 | 0,04737732 |
| 197135_at | PATL2 | 0,19929594 | 4,89242643 | 3,21578437 | 0,00597701 | 0,04741974 |
| 53917_at | RAB24 | 0,28792476 | 7,84866252 | 3,21366247 | 0,00600269 | 0,04760387 |
| 8408_at | ULK1 | 0,84366187 | 7,59795523 | 3,21285138 | 0,00601253 | 0,04766233 |
| 23102_at | TBC1D2B | 0,253414 | 8,79511853 | 3,21253242 | 0,00601641 | 0,0476734 |
| 890_at | CCNA2 | -1,28736373 | 9,10468636 | -3,21232922 | 0,00601888 | 0,0476734 |
| 5412_at | UBL3 | 0,72239621 | 8,65708764 | 3,21185041 | 0,00602471 | 0,04769993 |
| 1033_at | CDKN3 | -1,09135695 | 9,15510642 | -3,21164598 | 0,0060272 | 0,04770003 |
| 51406_at | NOL7 | -0,42129224 | 9,58906571 | -3,21122172 | 0,00603237 | 0,04770707 |
| 2012_at | EMP1 | -1,23583531 | 9,32380942 | -3,21116642 | 0,00603304 | 0,04770707 |
| 51175_at | TUBE1 | -0,3975849 | 6,36970623 | -3,20979552 | 0,00604977 | 0,04781977 |
| 55839_at | CENPN | -0,90105745 | 7,41458806 | -3,20905473 | 0,00605883 | 0,04787175 |
| 23263_at | MCF2L | 0,19505381 | 5,82168418 | 3,2085657 | 0,00606482 | 0,04789942 |
| 79019_at | CENPM | -0,84931247 | 8,55765125 | -3,20709916 | 0,00608282 | 0,04802187 |
| 130814_at | PQLC3 | 0,62829467 | 8,01866464 | 3,20688735 | 0,00608542 | 0,04802274 |
| 4171_at | MCM2 | -1,12182859 | 9,6928585 | -3,20536313 | 0,00610419 | 0,04815113 |
| 57099_at | AVEN | -0,47641796 | 6,86449079 | -3,20384545 | 0,00612294 | 0,04827922 |
| 64149_at | C17orf75 | -0,21565997 | 7,4564313 | -3,20180019 | 0,00614829 | 0,04845928 |

| | | | | | | |
|---|---|---|---|---|---|---|
| 9581_at | PREPL | -0,38104897 | 7,88904973 | -3,20085476 | 0,00616004 | 0,04853207 |
| 29105_at | CFAP20 | -0,5368402 | 9,97579371 | -3,19920493 | 0,00618061 | 0,04867418 |
| 9718_at | ECE2 | -0,27846653 | 6,26468856 | -3,19869907 | 0,00618693 | 0,04868841 |
| 79800_at | CARF | 0,76037701 | 4,41467239 | 3,19865562 | 0,00618747 | 0,04868841 |
| 9547_at | CXCL14 | -0,33906268 | 4,61628404 | -3,19553316 | 0,00622662 | 0,04897649 |
| 6629_at | SNRPB2 | -0,21794894 | 10,0753484 | -3,19514301 | 0,00623153 | 0,04899509 |
| 2535_at | FZD2 | 0,72064984 | 6,96095098 | 3,19358634 | 0,00625116 | 0,04912935 |
| 284900_at | TTC28-AS1 | -0,22849507 | 5,70935627 | -3,19237011 | 0,00626653 | 0,04918857 |
| 22849_at | CPEB3 | 0,79515534 | 4,86494961 | 3,1921715 | 0,00626905 | 0,04918857 |
| 23522_at | KAT6B | 0,21212541 | 6,77557737 | 3,19202471 | 0,00627091 | 0,04918857 |
| 55900_at | ZNF302 | 0,30428861 | 7,2961827 | 3,19187493 | 0,0062728 | 0,04918857 |
| 54960_at | GEMIN8 | 0,25678496 | 5,4234811 | 3,19170808 | 0,00627492 | 0,04918857 |
| 5095_at | PCCA | 0,2345618 | 6,77154926 | 3,1915872 | 0,00627645 | 0,04918857 |
| 55593_at | OTUD5 | 0,16409634 | 7,27467908 | 3,19157799 | 0,00627657 | 0,04918857 |
| 57180_at | ACTR3B | -0,3668502 | 6,54553499 | -3,19019472 | 0,00629413 | 0,049291 |
| 5905_at | RANGAP1 | -0,61359146 | 8,40289774 | -3,18998366 | 0,00629681 | 0,049291 |
| 27241_at | BBS9 | 0,45218057 | 5,49108285 | 3,18994424 | 0,00629731 | 0,049291 |
| 112611_at | RWDD2A | 0,53753321 | 5,49153995 | 3,18935535 | 0,00630481 | 0,04932962 |
| 645212_at | EIF3J-AS1 | 0,3107061 | 7,22732189 | 3,18837885 | 0,00631726 | 0,04940695 |
| 7533_at | YWHAH | -0,71743431 | 9,28432695 | -3,18756565 | 0,00632764 | 0,04946808 |
| 83460_at | EMC6 | -0,26401806 | 8,53891134 | -3,18706371 | 0,00633406 | 0,04949817 |
| 55133_at | SRBD1 | -0,35846589 | 7,27026476 | -3,18669342 | 0,0063388 | 0,04951512 |
| 55922_at | NKRF | -0,52221562 | 7,86334882 | -3,18607771 | 0,00634669 | 0,04955663 |
| 54495_at | TMX3 | -0,31526726 | 7,69765989 | -3,18578665 | 0,00635042 | 0,04956568 |
| 256302_at | NATD1 | 0,49402765 | 5,84218202 | 3,18499494 | 0,00636058 | 0,0496249 |
| 125988_at | C19orf70 | -0,32518757 | 9,22319381 | -3,18406274 | 0,00637257 | 0,04969829 |
| 54682_at | MANSC1 | 0,7290126 | 7,67229571 | 3,18037781 | 0,00642017 | 0,05004927 |
| 1999_at | ELF3 | 0,51186967 | 9,97373148 | 3,17833905 | 0,00644666 | 0,05023543 |
| 85007_at | PHYKPL | 0,53968804 | 7,15403294 | 3,17658972 | 0,00646947 | 0,05039283 |
| 5713_at | PSMD7 | -0,20055178 | 10,8105296 | -3,17356575 | 0,0065091 | 0,050681 |
| 51239_at | ANKRD39 | -0,29570209 | 7,52564301 | -3,17305813 | 0,00651578 | 0,05071248 |
| 83478_at | ARHGAP24 | 0,23274946 | 4,45783853 | 3,17268091 | 0,00652074 | 0,05072326 |
| 389856_at | USP27X | 0,39494841 | 5,60260281 | 3,17255291 | 0,00652243 | 0,05072326 |
| 51001_at | MTERF3 | -0,52358277 | 8,02208872 | -3,16913469 | 0,0065676 | 0,05105396 |
| 1017_at | CDK2 | -0,55852076 | 9,07159414 | -3,16817531 | 0,00658034 | 0,05111921 |
| 83596_at | BCL2L12 | -0,62284236 | 7,63384689 | -3,1681026 | 0,0065813 | 0,05111921 |
| 119504_at | ANAPC16 | 0,16934278 | 9,52665921 | 3,16771042 | 0,00658652 | 0,05113909 |
| 805_at | CALM2 | -0,24715983 | 10,9415259 | -3,16649227 | 0,00660274 | 0,05124298 |
| 197370_at | NSMCE1 | -0,22650845 | 8,78344618 | -3,16620285 | 0,0066066 | 0,05124298 |
| 27303_at | RBMS3 | 0,63734816 | 5,42818136 | 3,16610677 | 0,00660788 | 0,05124298 |
| 899_at | CCNF | -0,71138127 | 7,54705782 | -3,16423526 | 0,00663289 | 0,05139808 |
| 27101_at | CACYBP | -0,43457061 | 8,90937722 | -3,16412041 | 0,00663443 | 0,05139808 |
| 8208_at | CHAF1B | -1,00389899 | 6,85858865 | -3,16401208 | 0,00663588 | 0,05139808 |
| 1460_at | CSNK2B | -0,21005743 | 11,1398107 | -3,16190304 | 0,0066642 | 0,05158212 |
| 80306_at | MED28 | 0,23858581 | 8,6301756 | 3,16178306 | 0,00666582 | 0,05158212 |
| 51116_at | MRPS2 | -0,59723561 | 8,35230154 | -3,16164469 | 0,00666768 | 0,05158212 |
| 118472_at | ZNF511 | -0,38935105 | 8,43073002 | -3,16098938 | 0,00667651 | 0,05162171 |
| 3705_at | ITPK1 | -0,26326913 | 7,08564505 | -3,16086702 | 0,00667815 | 0,05162171 |
| 8520_at | HAT1 | -0,56670517 | 9,70984913 | -3,15988556 | 0,0066914 | 0,05170335 |
| 11179_at | ZNF277 | 0,47014538 | 7,36481514 | 3,15689369 | 0,00673194 | 0,05199573 |

| | | | | | | |
|---|---|---|---|---|---|---|
| 497049_at | FLJ25758 | -0,29915765 | 4,53017077 | -3,15591062 | 0,00674531 | 0,05207752 |
| 1277_at | COL1A1 | 0,46462314 | 6,41945493 | 3,15571802 | 0,00674794 | 0,05207752 |
| 6513_at | SLC2A1 | -0,37505902 | 8,87315941 | -3,15500175 | 0,0067577 | 0,05213199 |
| 80008_at | TMEM156 | -0,63311182 | 4,08063343 | -3,15461807 | 0,00676294 | 0,0521515 |
| 550643_at | LINC01420 | 0,4286849 | 9,88338401 | 3,15438817 | 0,00676608 | 0,05215483 |
| 9445_at | ITM2B | 0,44707142 | 11,6699571 | 3,15416393 | 0,00676914 | 0,05215757 |
| 10933_at | MORF4L1 | 0,21642916 | 10,9148336 | 3,15376615 | 0,00677458 | 0,0521786 |
| 27327_at | TNRC6A | 0,25512695 | 8,1173611 | 3,15293823 | 0,00678591 | 0,05224499 |
| 29781_at | NCAPH2 | -0,27515148 | 6,82626576 | -3,15191624 | 0,00679992 | 0,05231736 |
| 100499466_a | LINC00674 | 0,38607005 | 7,01063265 | 3,1518566 | 0,00680074 | 0,05231736 |
| 677838_at | SNORA61 | -0,74602784 | 8,60722211 | -3,15138276 | 0,00680725 | 0,05234652 |
| 84498_at | FAM120B | 0,21509503 | 7,47217993 | 3,15118503 | 0,00680997 | 0,05234652 |
| 25851_at | TECPR1 | 0,53132959 | 7,00874004 | 3,15051737 | 0,00681915 | 0,05239621 |
| 7247_at | TSN | -0,4206162 | 8,16432372 | -3,15013303 | 0,00682444 | 0,05241597 |
| 80325_at | ABTB1 | 0,38416657 | 5,93910588 | 3,1496027 | 0,00683175 | 0,05245119 |
| 10253_at | SPRY2 | -0,86811881 | 4,8392512 | -3,14938672 | 0,00683473 | 0,05245316 |
| 4595_at | MUTYH | -0,27731395 | 7,65638695 | -3,14910136 | 0,00683867 | 0,05246248 |
| 23080_at | AVL9 | 0,27243507 | 7,25401621 | 3,14877643 | 0,00684316 | 0,052476 |
| 79447_at | PAGR1 | -0,628707 | 8,13612554 | -3,14750412 | 0,00686075 | 0,05259 |
| 7518_at | XRCC4 | -0,64036542 | 5,68592415 | -3,14655464 | 0,00687391 | 0,05266992 |
| 6118_at | RPA2 | -0,20216889 | 8,68007213 | -3,14601217 | 0,00688144 | 0,05270666 |
| 10585_at | POMT1 | 0,32713516 | 6,76680541 | 3,14225944 | 0,00693376 | 0,05308628 |
| 10128_at | LRPPRC | -0,45529067 | 9,35528048 | -3,14120511 | 0,00694853 | 0,05317687 |
| 2620_at | GAS2 | 0,30302412 | 3,61037063 | 3,14102086 | 0,00695112 | 0,05317687 |
| 140687_at | GCNT7 | -0,21557425 | 3,1239423 | -3,14018765 | 0,00696282 | 0,05324522 |
| 84847_at | LINC00525 | 0,44537884 | 6,31672062 | 3,13920795 | 0,0069766 | 0,05332943 |
| 5652_at | PRSS8 | 0,76340836 | 7,93383192 | 3,13875308 | 0,006983 | 0,05335724 |
| 90390_at | MED30 | -0,35571641 | 6,56807202 | -3,13761861 | 0,00699901 | 0,05344748 |
| 55727_at | BTBD7 | 0,19786527 | 8,4909842 | 3,13752275 | 0,00700036 | 0,05344748 |
| 2686_at | GGT7 | 0,51159903 | 6,30163173 | 3,13716712 | 0,00700539 | 0,05346466 |
| 389434_at | IYD | 0,40121442 | 3,3730551 | 3,13642449 | 0,0070159 | 0,05352364 |
| 51428_at | DDX41 | -0,21363629 | 8,8938568 | -3,13608457 | 0,00702071 | 0,05353916 |
| 2644_at | GCHFR | 0,33391117 | 7,90778604 | 3,13523003 | 0,00703283 | 0,05361034 |
| 129303_at | TMEM150A | 0,28297024 | 6,34356224 | 3,13503356 | 0,00703562 | 0,05361039 |
| 6698_at | SPRR1A | -0,59917024 | 6,33539985 | -3,13425137 | 0,00704673 | 0,05367384 |
| 4723_at | NDUFV1 | -0,22824844 | 10,5082449 | -3,13393846 | 0,00705118 | 0,05368652 |
| 9131_at | AIFM1 | -0,35441695 | 8,18378727 | -3,13100917 | 0,00709298 | 0,05396419 |
| 115416_at | MALSU1 | -0,30670717 | 10,254496 | -3,13099032 | 0,00709325 | 0,05396419 |
| 84823_at | LMNB2 | -1,06286567 | 9,08817301 | -3,13068813 | 0,00709758 | 0,05397579 |
| 9605_at | VPS9D1 | 0,20034086 | 6,81854893 | 3,12975156 | 0,007111 | 0,05405654 |
| 161582_at | DYX1C1 | 0,51860576 | 4,8641647 | 3,12936944 | 0,00711649 | 0,0540769 |
| 64976_at | MRPL40 | -0,33061599 | 8,95859932 | -3,12903053 | 0,00712136 | 0,05409255 |
| 105221694_a | BISPR | 0,74564616 | 7,59334407 | 3,12815065 | 0,00713401 | 0,05416731 |
| 10055_at | SAE1 | -0,32000989 | 9,42191339 | -3,12731274 | 0,00714608 | 0,05423759 |
| 4085_at | MAD2L1 | -1,30788039 | 9,26370572 | -3,12693324 | 0,00715156 | 0,05425775 |
| 391712_at | TRIM61 | -0,31975129 | 5,6510329 | -3,12621113 | 0,00716198 | 0,05431547 |
| 101927720_a | ZNF793-AS1 | 0,40290712 | 5,19680988 | 3,12531255 | 0,00717498 | 0,05439261 |
| 170690_at | ADAMTS16 | 0,30478068 | 5,06398885 | 3,12444997 | 0,00718748 | 0,05446592 |
| 9894_at | TELO2 | -0,29881246 | 7,27410288 | -3,12425333 | 0,00719033 | 0,05446611 |
| 57465_at | TBC1D24 | -0,3031371 | 6,89514641 | -3,12341704 | 0,00720247 | 0,05453664 |

| | | | | | | |
|---|---|---|---|---|---|---|
| 55333_at | SYNJ2BP | 0,27363052 | 8,70927195 | 3,12256448 | 0,00721487 | 0,05460906 |
| 54929_at | TMEM161A | -0,19686861 | 8,52448644 | -3,12200447 | 0,00722303 | 0,05464931 |
| 83851_at | SYT16 | 0,18493877 | 4,33624675 | 3,12123513 | 0,00723425 | 0,05471271 |
| 283337_at | ZNF740 | 0,16800111 | 6,68048181 | 3,1201147 | 0,00725062 | 0,054815 |
| 9878_at | TOX4 | 0,22420713 | 8,4176072 | 3,1199171 | 0,00725351 | 0,05481534 |
| 284080_at | LOC284080 | -0,24360986 | 4,91221994 | -3,11742932 | 0,00729 | 0,05506952 |
| 3663_at | IRF5 | 0,26056773 | 6,02532761 | 3,11640191 | 0,00730512 | 0,05516214 |
| 79071_at | ELOVL6 | -0,60379139 | 7,56066769 | -3,11540209 | 0,00731987 | 0,05525185 |
| 56927_at | GPR108 | 0,36670127 | 9,04806104 | 3,11502645 | 0,00732542 | 0,05527207 |
| 60672_at | MIIP | -0,21511397 | 7,46648337 | -3,11432487 | 0,0073358 | 0,05532867 |
| 57592_at | ZNF687 | 0,36402319 | 6,57886243 | 3,11404376 | 0,00733996 | 0,05533839 |
| 995_at | CDC25C | -0,68969354 | 6,34271055 | -3,11374277 | 0,00734441 | 0,05535033 |
| 88745_at | RRP36 | -0,35401166 | 9,16343619 | -3,11339105 | 0,00734963 | 0,05536795 |
| 5111_at | PCNA | -0,91728368 | 11,2044322 | -3,11282147 | 0,00735808 | 0,05540993 |
| 53635_at | PTOV1 | 0,41098969 | 9,16524228 | 3,11196334 | 0,00737082 | 0,05548423 |
| 9278_at | ZBTB22 | 0,28257285 | 7,09496282 | 3,10942115 | 0,00740871 | 0,0557267 |
| 5601_at | MAPK9 | -0,37700188 | 8,22162772 | -3,1094141 | 0,00740882 | 0,0557267 |
| 83592_at | AKR1E2 | -0,31978436 | 5,22170093 | -3,1070504 | 0,00744422 | 0,05596851 |
| 100170841_a | C17orf96 | -0,50189043 | 6,52762774 | -3,1068802 | 0,00744678 | 0,05596851 |
| 2879_at | GPX4 | 0,21296703 | 11,9303398 | 3,10656228 | 0,00745155 | 0,05597451 |
| 57184_at | FAM219B | 0,30902868 | 7,39982971 | 3,10644024 | 0,00745339 | 0,05597451 |
| 28985_at | MCTS1 | -0,26149946 | 5,9355708 | -3,10593032 | 0,00746106 | 0,05601027 |
| 7743_at | ZNF189 | 0,26275444 | 7,67652529 | 3,10536573 | 0,00746956 | 0,05604413 |
| 6621_at | SNAPC4 | -0,29714673 | 7,5652228 | -3,10524428 | 0,00747139 | 0,05604413 |
| 5364_at | PLXNB1 | 0,29546146 | 7,3420643 | 3,10321976 | 0,00750195 | 0,05625151 |
| 11067_at | C10orf10 | 0,23307115 | 5,78721317 | 3,10235201 | 0,00751509 | 0,05632811 |
| 10964_at | IFI44L | 2,4135576 | 7,34031514 | 3,10148882 | 0,00752819 | 0,0564043 |
| 51306_at | FAM13B | 0,72680612 | 8,32474198 | 3,10090209 | 0,0075371 | 0,05644911 |
| 8440_at | NCK2 | 0,45062497 | 9,11393197 | 3,10050169 | 0,00754319 | 0,05645367 |
| 130733_at | TMEM178A | 0,96446565 | 5,0655161 | 3,10047661 | 0,00754357 | 0,05645367 |
| 4673_at | NAP1L1 | -0,2871305 | 8,10245663 | -3,09914294 | 0,00756388 | 0,05658371 |
| 8664_at | EIF3D | -0,30153583 | 10,5777128 | -3,09875313 | 0,00756983 | 0,05660622 |
| 100289092_a | ATP2A1-AS1 | -0,53890846 | 4,92672883 | -3,0983597 | 0,00757584 | 0,05662916 |
| 58488_at | PCTP | 0,46417111 | 8,30137376 | 3,09712627 | 0,0075947 | 0,05674816 |
| 7384_at | UQCRC1 | -0,34149823 | 10,0667413 | -3,09608637 | 0,00761064 | 0,05684522 |
| 143458_at | LDLRAD3 | -0,95980195 | 6,94291901 | -3,09270037 | 0,00766278 | 0,05721245 |
| 23078_at | VWA8 | -0,23322187 | 7,69167184 | -3,0922882 | 0,00766915 | 0,05723782 |
| 55827_at | DCAF6 | 0,41964274 | 8,58576648 | 3,09158002 | 0,00768011 | 0,0572974 |
| 8704_at | B4GALT2 | -0,3277476 | 7,72729143 | -3,0913784 | 0,00768323 | 0,05729851 |
| 5754_at | PTK7 | 0,28074459 | 8,46474905 | 3,09041487 | 0,00769817 | 0,05737428 |
| 84904_at | ARHGEF39 | -0,70394154 | 6,26858925 | -3,09033903 | 0,00769935 | 0,05737428 |
| 56886_at | UGGT1 | -0,17696494 | 7,82806976 | -3,0896001 | 0,00771082 | 0,05743759 |
| 65082_at | VPS33A | 0,2344252 | 7,18454644 | 3,08870149 | 0,00772481 | 0,05751822 |
| 79745_at | CLIP4 | 0,66675458 | 7,19442471 | 3,08852081 | 0,00772762 | 0,05751822 |
| 8678_at | BECN1 | 0,27504656 | 9,11516733 | 3,0882599 | 0,00773169 | 0,05752625 |
| 498_at | ATP5A1 | -0,25970561 | 11,1336198 | -3,08805401 | 0,0077349 | 0,05752791 |
| 51368_at | TEX264 | 0,18955491 | 7,64276464 | 3,08781605 | 0,00773861 | 0,05752972 |
| 101927422_a | GNA14-AS1 | -0,27901609 | 3,68095446 | -3,0876555 | 0,00774111 | 0,05752972 |
| 27316_at | RBMX | -0,30516853 | 9,67872018 | -3,08708222 | 0,00775006 | 0,057573 |
| 9663_at | LPIN2 | 0,32309625 | 6,67745081 | 3,08689985 | 0,00775291 | 0,057573 |

| | | | | | | |
|---|---|---|---|---|---|---|
| 6300_at | MAPK12 | -0,31585003 | 6,89876549 | -3,08626507 | 0,00776284 | 0,05762451 |
| 9861_at | PSMD6 | -0,36544905 | 9,42869186 | -3,08506498 | 0,00778164 | 0,05774183 |
| 84888_at | SPPL2A | 0,30856606 | 10,1876737 | 3,08446994 | 0,00779098 | 0,05778887 |
| 54101_at | RIPK4 | 0,36679728 | 8,68960933 | 3,08415364 | 0,00779595 | 0,05780347 |
| 49856_at | WRAP73 | -0,23517009 | 7,61256819 | -3,08389356 | 0,00780004 | 0,05781153 |
| 29946_at | SERTAD3 | 0,4946459 | 6,94902958 | 3,08303705 | 0,00781352 | 0,05788915 |
| 550_at | AUP1 | -0,19285096 | 9,90538856 | -3,08246541 | 0,00782253 | 0,05791726 |
| 5634_at | PRPS2 | -0,48094978 | 9,63552381 | -3,0824147 | 0,00782333 | 0,05791726 |
| 9906_at | SLC35E2 | 0,70088227 | 6,66497868 | 3,08220544 | 0,00782663 | 0,05791944 |
| 9873_at | FCHSD2 | 0,30319711 | 8,06511894 | 3,08195575 | 0,00783057 | 0,05792634 |
| 6319_at | SCD | -0,71963502 | 9,86750724 | -3,08156591 | 0,00783672 | 0,05794962 |
| 148808_at | MFSD4 | 0,25923432 | 4,69493549 | 3,08000016 | 0,00786149 | 0,05811049 |
| 84951_at | TNS4 | -0,34952895 | 7,96078472 | -3,07911366 | 0,00787555 | 0,05819187 |
| 5337_at | PLD1 | 0,56884306 | 6,41088857 | 3,07892514 | 0,00787854 | 0,05819187 |
| 4282_at | MIF | -0,44496193 | 12,2032447 | -3,07697676 | 0,00790954 | 0,05837233 |
| 353355_at | ZNF233 | 0,3295038 | 4,14843562 | 3,07688047 | 0,00791108 | 0,05837233 |
| 4323_at | MMP14 | 0,23151627 | 8,57424589 | 3,07681843 | 0,00791207 | 0,05837233 |
| 58473_at | PLEKHB1 | 0,72771007 | 4,77664584 | 3,07551381 | 0,0079329 | 0,0585036 |
| 119392_at | SFR1 | -0,48513767 | 6,5672924 | -3,07485212 | 0,00794348 | 0,05852819 |
| 10363_at | HMG20A | 0,3952582 | 8,40352104 | 3,07482522 | 0,00794391 | 0,05852819 |
| 83480_at | PUS3 | -0,23571977 | 6,571339 | -3,07473571 | 0,00794535 | 0,05852819 |
| 10961_at | ERP29 | 0,22201715 | 10,524508 | 3,07232837 | 0,00798399 | 0,05879034 |
| 5175_at | PECAM1 | -0,40362642 | 4,76401694 | -3,07136569 | 0,00799949 | 0,058882 |
| 22870_at | PPP6R1 | -0,21344428 | 8,48856249 | -3,07111422 | 0,00800354 | 0,05888934 |
| 80305_at | TRABD | -0,19219848 | 7,94394665 | -3,07089313 | 0,00800711 | 0,05889308 |
| 7993_at | UBXN8 | -0,46523374 | 7,72995589 | -3,06989989 | 0,00802315 | 0,05898855 |
| 11346_at | SYNPO | 0,28038505 | 6,53754957 | 3,0675319 | 0,00806153 | 0,05924807 |
| 11043_at | MID2 | 0,29698462 | 6,9673226 | 3,06679038 | 0,00807358 | 0,05931403 |
| 144568_at | A2ML1 | 0,34987937 | 3,2280334 | 3,06517709 | 0,00809987 | 0,05946192 |
| 1062_at | CENPE | -1,12239455 | 7,40965261 | -3,06517602 | 0,00809988 | 0,05946192 |
| 23475_at | QPRT | 0,98493331 | 8,1395434 | 3,0648189 | 0,00810571 | 0,05948205 |
| 204_at | AK2 | -0,6651398 | 8,12317889 | -3,06426603 | 0,00811475 | 0,05952567 |
| 23032_at | USP33 | 0,36166457 | 8,41297974 | 3,0640242 | 0,0081187 | 0,05953201 |
| 91252_at | SLC39A13 | 0,24737842 | 7,83511079 | 3,06370574 | 0,00812391 | 0,05954755 |
| 440515_at | ZNF506 | 0,34497709 | 5,96091327 | 3,06345931 | 0,00812795 | 0,05955446 |
| 80235_at | PIGZ | 0,38507785 | 6,32323281 | 3,06303016 | 0,00813498 | 0,05958331 |
| 7498_at | XDH | -0,80734238 | 7,05379113 | -3,06274048 | 0,00813973 | 0,05959543 |
| 202781_at | PAXIP1-AS1 | -0,41545829 | 6,93259153 | -3,06179851 | 0,00815519 | 0,05968595 |
| 7328_at | UBE2H | 0,43643771 | 9,15477336 | 3,06087265 | 0,00817042 | 0,05977468 |
| 5932_at | RBBP8 | -0,56952801 | 8,24215206 | -3,06016665 | 0,00818204 | 0,05983703 |
| 84830_at | ADTRP | 0,40027903 | 4,69094535 | 3,05996431 | 0,00818538 | 0,0598387 |
| 282679_at | AQP11 | 0,35127769 | 5,21077377 | 3,05877603 | 0,008205 | 0,05995686 |
| 10197_at | PSME3 | -0,45858148 | 9,32867038 | -3,05860846 | 0,00820777 | 0,05995686 |
| 54815_at | GATAD2A | -0,18434304 | 8,61833031 | -3,05724469 | 0,00823035 | 0,06009901 |
| 7468_at | WHSC1 | -0,56952104 | 7,67554559 | -3,05622779 | 0,00824722 | 0,06019943 |
| 328_at | APEX1 | -0,24743248 | 11,5993505 | -3,05497755 | 0,00826802 | 0,06032836 |
| 7071_at | KLF10 | 0,35175253 | 9,83876032 | 3,05410392 | 0,00828258 | 0,06041173 |
| 6094_at | ROM1 | -0,33627064 | 5,19635554 | -3,0527016 | 0,00830601 | 0,06054955 |
| 967_at | CD63 | 0,22886768 | 10,6573247 | 3,05259672 | 0,00830776 | 0,06054955 |
| 10169_at | SERF2 | 0,49221537 | 7,89035417 | 3,05240415 | 0,00831099 | 0,06055013 |

| | | | | | | |
|---|---|---|---|---|---|---|
| 8420_at | SNHG3 | -1,00126354 | 5,52207538 | -3,05185526 | 0,00832018 | 0,06059419 |
| 652968_at | GATSL3 | 0,55422878 | 6,16006696 | 3,05079498 | 0,00833797 | 0,06067297 |
| 382_at | ARF6 | -0,19423659 | 10,0725665 | -3,05070022 | 0,00833956 | 0,06067297 |
| 54985_at | HCFC1R1 | 0,65383516 | 8,66314506 | 3,05048026 | 0,00834325 | 0,06067297 |
| 154807_at | VKORC1L1 | -0,30065153 | 7,5371128 | -3,05045989 | 0,0083436 | 0,06067297 |
| 3280_at | HES1 | 0,38675465 | 10,0333326 | 3,04938458 | 0,00836169 | 0,06078157 |
| 353497_at | POLN | -0,27468547 | 4,54366601 | -3,04903838 | 0,00836752 | 0,06080102 |
| 8893_at | EIF2B5 | -0,24766728 | 8,2989092 | -3,04880198 | 0,0083715 | 0,06080703 |
| 57094_at | CPA6 | 0,67711844 | 3,88353922 | 3,04852746 | 0,00837613 | 0,06081773 |
| 79717_at | PPCS | 0,2716614 | 8,58390465 | 3,04764358 | 0,00839106 | 0,06090313 |
| 140885_at | SIRPA | -0,41454963 | 6,39284126 | -3,04718932 | 0,00839874 | 0,06093591 |
| 175_at | AGA | 0,53512881 | 7,60227795 | 3,04604475 | 0,00841812 | 0,06105353 |
| 10999_at | SLC27A4 | -0,41322913 | 7,78069751 | -3,04492986 | 0,00843704 | 0,06116773 |
| 8914_at | TIMELESS | -0,62010552 | 6,634179 | -3,04470981 | 0,00844078 | 0,06117182 |
| 145483_at | FAM161B | 0,44092052 | 5,51914221 | 3,04441489 | 0,00844579 | 0,06118245 |
| 22876_at | INPP5F | -0,32079379 | 5,62717576 | -3,04424994 | 0,0084486 | 0,06118245 |
| 7105_at | TSPAN6 | 0,39505406 | 9,1208529 | 3,04286849 | 0,00847213 | 0,06132983 |
| 84240_at | ZCCHC9 | -0,29124472 | 8,38745314 | -3,04024835 | 0,00851695 | 0,06163109 |
| 23512_at | SUZ12 | -0,33145528 | 9,6677982 | -3,03773342 | 0,00856018 | 0,06192069 |
| 10328_at | EMC8 | -0,43832467 | 8,95895544 | -3,03711504 | 0,00857085 | 0,06197456 |
| 5720_at | PSME1 | 0,35346549 | 11,1619334 | 3,03632091 | 0,00858456 | 0,06205043 |
| 89958_at | SAPCD2 | -0,84502613 | 8,34598474 | -3,03489964 | 0,00860916 | 0,06220489 |
| 8000_at | PSCA | 1,1770882 | 10,2192949 | 3,03421749 | 0,00862099 | 0,06224642 |
| 9654_at | TTLL4 | -0,24509575 | 6,64479581 | -3,03419562 | 0,00862137 | 0,06224642 |
| 1054_at | CEBPG | -0,21894014 | 7,96081093 | -3,03394459 | 0,00862573 | 0,06224665 |
| 143879_at | KBTBD3 | 0,37751801 | 4,68966589 | 3,03377604 | 0,00862866 | 0,06224665 |
| 57593_at | EBF4 | 0,41314169 | 6,11559752 | 3,03363554 | 0,0086311 | 0,06224665 |
| 53944_at | CSNK1G1 | 0,28919249 | 7,78953905 | 3,03207047 | 0,00865833 | 0,0624197 |
| 84817_at | TXNDC17 | -0,4626106 | 7,64727795 | -3,03176332 | 0,00866369 | 0,06243494 |
| 4001_at | LMNB1 | -0,98710453 | 9,47488516 | -3,03134025 | 0,00867107 | 0,06246477 |
| 221806_at | VWDE | 0,30032664 | 4,11250973 | 3,03067167 | 0,00868275 | 0,06252551 |
| 3685_at | ITGAV | 0,46662197 | 10,8208506 | 3,0293723 | 0,00870549 | 0,06266583 |
| 8975_at | USP13 | -0,57160413 | 6,7588089 | -3,02870637 | 0,00871716 | 0,06272645 |
| 5002_at | SLC22A18 | 0,53336783 | 7,10459992 | 3,02833695 | 0,00872365 | 0,06273335 |
| 7637_at | ZNF84 | 0,37059485 | 7,17192761 | 3,02810175 | 0,00872778 | 0,06273335 |
| 5573_at | PRKAR1A | 0,29684331 | 10,6188138 | 3,02809408 | 0,00872791 | 0,06273335 |
| 147670_at | SMIM17 | 0,3877614 | 3,25344001 | 3,02786781 | 0,00873189 | 0,06273335 |
| 80174_at | DBF4B | -0,21937166 | 5,29080682 | -3,02772457 | 0,00873441 | 0,06273335 |
| 22853_at | LMTK2 | -0,2859977 | 6,75704741 | -3,0274892 | 0,00873854 | 0,06273969 |
| 100130275_a | TFAP2A-AS1 | 0,4790714 | 5,92563231 | 3,02601852 | 0,00876445 | 0,06290223 |
| 6875_at | TAF4B | -0,68828156 | 4,94014178 | -3,02542131 | 0,00877499 | 0,06295443 |
| 3035_at | HARS | -0,44311071 | 9,53744262 | -3,02502734 | 0,00878195 | 0,06298091 |
| 26156_at | RSL1D1 | -0,52997003 | 8,33343258 | -3,02329558 | 0,00881261 | 0,06316143 |
| 6636_at | SNRPF | -0,66230049 | 9,65359133 | -3,02323533 | 0,00881368 | 0,06316143 |
| 9937_at | DCLRE1A | -0,28749818 | 6,75816038 | -3,02288216 | 0,00881995 | 0,06318284 |
| 64423_at | INF2 | -0,20004011 | 7,29640764 | -3,02241252 | 0,00882829 | 0,06321907 |
| 100128191_a | TMPO-AS1 | -0,73113528 | 6,32851625 | -3,02203888 | 0,00883493 | 0,06324311 |
| 81873_at | ARPC5L | -0,257901 | 8,51434795 | -3,02179594 | 0,00883925 | 0,06325054 |
| 10325_at | RRAGB | 0,46474005 | 6,13596442 | 3,02151046 | 0,00884433 | 0,06326338 |
| 10483_at | SEC23B | -0,21069425 | 9,35520727 | -3,0205213 | 0,00886195 | 0,06336591 |

| | | | | | | |
|---|---|---|---|---|---|---|
| 8651_at | SOCS1 | 0,69706422 | 5,87512074 | 3,0193656 | 0,00888258 | 0,06348988 |
| 6322_at | SCML1 | -0,48404471 | 6,24054897 | -3,01884857 | 0,00889183 | 0,06353239 |
| 4649_at | MYO9A | 0,33017514 | 7,05086972 | 3,01821434 | 0,00890319 | 0,06356461 |
| 100101467_a | ZSCAN30 | 0,34456926 | 5,63967181 | 3,01808856 | 0,00890544 | 0,06356461 |
| 10395_at | DLC1 | 0,65345742 | 7,43867269 | 3,01804395 | 0,00890624 | 0,06356461 |
| 2730_at | GCLM | -0,64832439 | 9,27598286 | -3,01613468 | 0,00894052 | 0,06377281 |
| 8886_at | DDX18 | -0,54743805 | 8,89935039 | -3,0159193 | 0,0089444 | 0,06377281 |
| 51097_at | SCCPDH | -0,27809477 | 9,83696905 | -3,01586667 | 0,00894534 | 0,06377281 |
| 23586_at | DDX58 | 1,24180384 | 7,1223673 | 3,0151468 | 0,00895831 | 0,06384162 |
| 51421_at | AMOTL2 | 0,698088 | 8,7575969 | 3,01413158 | 0,00897663 | 0,06392822 |
| 3434_at | IFIT1 | 1,90689294 | 7,6346233 | 3,01410544 | 0,0089771 | 0,06392822 |
| 80004_at | ESRP2 | 0,42407662 | 9,61468344 | 3,0133626 | 0,00899052 | 0,06400017 |
| 3187_at | HNRNPH1 | -0,38410465 | 9,47849502 | -3,01229741 | 0,00900981 | 0,06411378 |
| 327_at | APEH | -0,29926196 | 8,95963652 | -3,0101819 | 0,00904824 | 0,06434313 |
| 26499_at | PLEK2 | -0,8965356 | 9,62570193 | -3,01015533 | 0,00904872 | 0,06434313 |
| 26821_at | SNORA74A | -0,26061988 | 4,00213128 | -3,00816862 | 0,00908496 | 0,06457696 |
| 9215_at | LARGE | 0,39021259 | 7,55323141 | 3,00732254 | 0,00910043 | 0,0646631 |
| 7083_at | TK1 | -1,08614819 | 9,51418247 | -3,00712773 | 0,009104 | 0,0646646 |
| 81614_at | NIPA2 | -0,25115935 | 8,90224994 | -3,00649299 | 0,00911563 | 0,06471981 |
| 114785_at | MBD6 | 0,26289678 | 7,36476979 | 3,00627747 | 0,00911959 | 0,06471981 |
| 57494_at | RIMKLB | 0,17721555 | 4,58671576 | 3,00615382 | 0,00912185 | 0,06471981 |
| 1629_at | DBT | 0,38535265 | 7,03654432 | 3,00581114 | 0,00912814 | 0,06474059 |
| 2071_at | ERCC3 | -0,20004944 | 8,38809066 | -3,00485814 | 0,00914566 | 0,06484093 |
| 83443_at | SF3B5 | -0,34106583 | 10,0359823 | -3,00447358 | 0,00915274 | 0,06486723 |
| 10553_at | HTATIP2 | 0,37584422 | 8,38539951 | 3,00399186 | 0,00916161 | 0,0648787 |
| 79679_at | VTCN1 | 1,81375252 | 4,75564702 | 3,00387376 | 0,00916378 | 0,0648787 |
| 26271_at | FBXO5 | -0,79036629 | 8,16271945 | -3,00383726 | 0,00916446 | 0,0648787 |
| 722_at | C4BPA | -0,21983978 | 4,95855573 | -3,00261935 | 0,00918693 | 0,06501392 |
| 146542_at | ZNF688 | 0,15325789 | 5,71328784 | 3,00233821 | 0,00919213 | 0,06502115 |
| 2889_at | RAPGEF1 | 0,22276487 | 6,81373005 | 3,00201742 | 0,00919806 | 0,06502115 |
| 1783_at | DYNC1LI2 | 0,18988083 | 9,03755838 | 3,00201642 | 0,00919808 | 0,06502115 |
| 26503_at | SLC17A5 | 0,99857647 | 7,85683056 | 3,00061027 | 0,00922413 | 0,06518137 |
| 55062_at | WIPI1 | 0,86863749 | 7,2510631 | 3,00014953 | 0,00923268 | 0,06521787 |
| 79003_at | MIS12 | -0,3459791 | 7,4564462 | -2,99872541 | 0,00925916 | 0,06538095 |
| 403313_at | PPAPDC2 | 0,36661115 | 7,60369541 | 2,99737282 | 0,00928438 | 0,065535 |
| 57826_at | RAP2C | 0,63594 | 8,29664784 | 2,99599534 | 0,00931013 | 0,06569271 |
| 83448_at | PUS7L | -0,37219235 | 6,32595405 | -2,99510041 | 0,0093269 | 0,06578694 |
| 9245_at | GCNT3 | -0,31036077 | 5,11489602 | -2,99444404 | 0,00933922 | 0,06584971 |
| 127703_at | C1orf216 | -0,40843348 | 7,36715554 | -2,99407189 | 0,00934621 | 0,06587489 |
| 92912_at | UBE2Q2 | 0,43904879 | 8,6589339 | 2,99383349 | 0,00935069 | 0,06588237 |
| 10892_at | MALT1 | -0,39288982 | 6,50325136 | -2,9933685 | 0,00935944 | 0,06591988 |
| 170626_at | XAGE3 | -0,29741322 | 6,27282395 | -2,99216931 | 0,00938203 | 0,06601673 |
| 26470_at | SEZ6L2 | 0,47600029 | 6,45798693 | 2,99216749 | 0,00938207 | 0,06601673 |
| 149373_at | LOC149373 | -0,37454604 | 4,34850159 | -2,99209312 | 0,00938347 | 0,06601673 |
| 4913_at | NTHL1 | -0,48832332 | 7,26609266 | -2,99056906 | 0,00941227 | 0,06615713 |
| 23623_at | RUSC1 | -0,30738676 | 9,01309868 | -2,99039089 | 0,00941564 | 0,06615713 |
| 26751_at | SH3YL1 | 0,42658988 | 7,85082405 | 2,99036835 | 0,00941607 | 0,06615713 |
| 29789_at | OLA1 | -0,26336881 | 5,90539077 | -2,99031029 | 0,00941717 | 0,06615713 |
| 79944_at | L2HGDH | -0,45145755 | 6,75636112 | -2,9898991 | 0,00942495 | 0,06616364 |
| 8863_at | PER3 | 0,36104219 | 6,14551626 | 2,98989873 | 0,00942496 | 0,06616364 |

| | | | | | | |
|---|---|---|---|---|---|---|
| 2956_at | MSH6 | -0,51432456 | 8,6752778 | -2,98920272 | 0,00943816 | 0,06623215 |
| 148932_at | MOB3C | 0,19884117 | 7,24588565 | 2,98848289 | 0,00945183 | 0,06630391 |
| 55644_at | OSGEP | -0,22756774 | 7,27853943 | -2,98821753 | 0,00945687 | 0,06631514 |
| 85363_at | TRIM5 | 0,42507413 | 5,32810148 | 2,98798226 | 0,00946134 | 0,06632237 |
| 8800_at | PEX11A | 0,43614436 | 6,59650137 | 2,98766387 | 0,0094674 | 0,06634069 |
| 2827_at | GPR3 | -0,3864115 | 4,90705444 | -2,98736643 | 0,00947306 | 0,06635623 |
| 6715_at | SRD5A1 | -0,16316213 | 7,74926329 | -2,98519012 | 0,0095146 | 0,06662294 |
| 10200_at | MPHOSPH6 | -0,51297051 | 7,87299355 | -2,98427018 | 0,00953221 | 0,06671471 |
| 996_at | CDC27 | -0,19635173 | 8,13500741 | -2,98414377 | 0,00953463 | 0,06671471 |
| 728224_at | KRTAP4-8 | -0,29357287 | 4,6140375 | -2,98325205 | 0,00955174 | 0,06677994 |
| 7773_at | ZNF230 | 0,26904889 | 4,23701126 | 2,98323405 | 0,00955208 | 0,06677994 |
| 56900_at | TMEM167B | 0,32043146 | 9,13820957 | 2,98311573 | 0,00955435 | 0,06677994 |
| 51186_at | WBP5 | 0,2563895 | 9,70053394 | 2,98288261 | 0,00955883 | 0,06678701 |
| 55194_at | EVA1B | 0,38077021 | 7,10499155 | 2,98169291 | 0,00958172 | 0,06691985 |
| 165215_at | FAM171B | 0,67781607 | 6,96801237 | 2,98153319 | 0,00958479 | 0,06691985 |
| 9804_at | TOMM20 | -0,24018235 | 10,7391831 | -2,9803187 | 0,00960822 | 0,06705909 |
| 191_at | AHCY | -0,47585312 | 10,0518161 | -2,97976956 | 0,00961883 | 0,06709166 |
| 23643_at | LY96 | 0,76073014 | 4,10276405 | 2,9797167 | 0,00961985 | 0,06709166 |
| 7569_at | ZNF182 | 0,3337078 | 5,90441399 | 2,97797714 | 0,00965354 | 0,06730226 |
| 55219_at | TMEM57 | 0,28075066 | 8,09960544 | 2,97738514 | 0,00966503 | 0,067358 |
| 51760_at | SYT17 | 0,45837887 | 6,33236219 | 2,97593589 | 0,00969322 | 0,06753003 |
| 51126_at | NAA20 | -0,36350838 | 9,36968312 | -2,97364982 | 0,00973785 | 0,06781643 |
| 29080_at | CCDC59 | -0,47251506 | 7,93278117 | -2,97327881 | 0,00974511 | 0,06784248 |
| 22955_at | SCMH1 | -0,26542615 | 7,88581981 | -2,97289617 | 0,00975261 | 0,06787013 |
| 5287_at | PIK3C2B | 0,52329252 | 9,27564522 | 2,97254542 | 0,00975949 | 0,06789345 |
| 134957_at | STXBP5 | 0,39737114 | 6,35396085 | 2,97170983 | 0,00977588 | 0,06793668 |
| 29763_at | PACSIN3 | -0,35658485 | 7,679665 | -2,97170894 | 0,0097759 | 0,06793668 |
| 23168_at | RTF1 | 0,18658802 | 7,95985601 | 2,97168974 | 0,00977628 | 0,06793668 |
| 286887_at | KRT6C | -0,26184126 | 3,08602781 | -2,96932798 | 0,00982278 | 0,06821374 |
| 9733_at | SART3 | -0,3861047 | 8,37792258 | -2,96924105 | 0,00982449 | 0,06821374 |
| 6720_at | SREBF1 | -0,47723745 | 8,23524985 | -2,96912565 | 0,00982677 | 0,06821374 |
| 100286938_a | VPS13A-AS1 | -0,30859841 | 5,18342284 | -2,96824225 | 0,00984423 | 0,0683103 |
| 56931_at | DUS3L | -0,66613604 | 7,25249691 | -2,96770253 | 0,00985491 | 0,06835978 |
| 90321_at | ZNF766 | 0,33549542 | 7,57909952 | 2,96696887 | 0,00986944 | 0,06843596 |
| 54620_at | FBXL19 | 0,31276202 | 7,63332486 | 2,9642239 | 0,00992402 | 0,06878961 |
| 1384_at | CRAT | 0,33322067 | 7,58232827 | 2,96395796 | 0,00992932 | 0,06880161 |
| 4356_at | MPP3 | -0,30716984 | 5,72858179 | -2,96319437 | 0,00994456 | 0,06888244 |
| 22868_at | FASTKD2 | -0,42737772 | 6,86951398 | -2,96204298 | 0,00996759 | 0,06901711 |
| 158471_at | PRUNE2 | 0,20712947 | 3,60726327 | 2,96153372 | 0,00997779 | 0,06906292 |
| 253635_at | GPATCH11 | -0,43510457 | 6,12993526 | -2,95805197 | 0,0100478 | 0,06949894 |
| 10190_at | TXNDC9 | -0,31499661 | 8,7954972 | -2,95792467 | 0,01005037 | 0,06949894 |
| 8402_at | SLC25A11 | -0,37800593 | 8,8182119 | -2,9576642 | 0,01005563 | 0,06949894 |
| 100652853_a | LINC01023 | -0,28580085 | 5,26083083 | -2,95723073 | 0,01006439 | 0,06949894 |
| 4598_at | MVK | -0,2966659 | 8,19431158 | -2,95721889 | 0,01006463 | 0,06949894 |
| 55072_at | RNF31 | 0,34844095 | 7,11321227 | 2,95705658 | 0,01006791 | 0,06949894 |
| 133522_at | PPARGC1B | -0,37810214 | 6,31076908 | -2,95702805 | 0,01006849 | 0,06949894 |
| 10102_at | TSFM | -0,226146 | 6,16481537 | -2,9569563 | 0,01006994 | 0,06949894 |
| 79815_at | NIPAL2 | 0,60913555 | 7,56048318 | 2,95679247 | 0,01007325 | 0,06949894 |
| 54984_at | PINX1 | -0,74482333 | 6,54439388 | -2,95609532 | 0,01008736 | 0,06957139 |
| 11320_at | MGAT4A | 0,81239728 | 5,62310415 | 2,95512871 | 0,01010696 | 0,06968161 |

| | | | | | | |
|---|---|---|---|---|---|---|
| 84661_at | DPY30 | -0,17334249 | 10,1740813 | -2,95479027 | 0,01011383 | 0,06970404 |
| 1663_at | DDX11 | -0,48201155 | 7,56042078 | -2,95439761 | 0,01012181 | 0,06973407 |
| 255758_at | TCTEX1D2 | -0,36806682 | 8,06765696 | -2,95388867 | 0,01013216 | 0,06978041 |
| 57475_at | PLEKHH1 | 0,34090034 | 6,43760048 | 2,95362319 | 0,01013756 | 0,06979267 |
| 7174_at | TPP2 | -0,21368984 | 7,9498131 | -2,95205781 | 0,01016948 | 0,06998737 |
| 28987_at | NOB1 | -0,4417854 | 8,69994903 | -2,95009453 | 0,01020964 | 0,0702387 |
| 6388_at | SDF2 | 0,2229528 | 8,56313398 | 2,9496927 | 0,01021788 | 0,07027029 |
| 151742_at | PPM1L | 0,44976687 | 5,82001937 | 2,94941447 | 0,01022359 | 0,07028447 |
| 23074_at | UHRF1BP1L | 0,1974654 | 5,55196791 | 2,94876199 | 0,01023699 | 0,07035149 |
| 10489_at | LRRC41 | 0,2204557 | 9,2749444 | 2,948009 | 0,01025248 | 0,07041367 |
| 2742_at | GLRA2 | -0,19365796 | 3,98604428 | -2,94789427 | 0,01025484 | 0,07041367 |
| 53630_at | BCO1 | 0,60342408 | 4,73106189 | 2,94778909 | 0,01025701 | 0,07041367 |
| 8317_at | CDC7 | -0,8403158 | 7,32389641 | -2,9458471 | 0,01029707 | 0,07066353 |
| 7378_at | UPP1 | -0,56961433 | 8,10009324 | -2,94525006 | 0,01030942 | 0,07072308 |
| 22887_at | FOXJ3 | -0,18631907 | 8,67661536 | -2,94454996 | 0,01032392 | 0,07079733 |
| 629_at | CFB | 1,61343423 | 5,65614493 | 2,94390235 | 0,01033735 | 0,07085861 |
| 23608_at | MKRN1 | 0,4152374 | 9,28539556 | 2,94376434 | 0,01034021 | 0,07085861 |
| 81786_at | TRIM7 | -0,58825489 | 5,79497999 | -2,94321368 | 0,01035165 | 0,07091175 |
| 22838_at | RNF44 | 0,35615279 | 8,44012386 | 2,94258929 | 0,01036463 | 0,07095958 |
| 6396_at | SEC13 | -0,35379247 | 10,0685226 | -2,94245322 | 0,01036746 | 0,07095958 |
| 473_at | RERE | 0,16507409 | 6,74208577 | 2,94234652 | 0,01036968 | 0,07095958 |
| 65003_at | MRPL11 | -0,56184379 | 9,07699274 | -2,94191797 | 0,01037861 | 0,07098361 |
| 6821_at | SUOX | 0,28932001 | 7,43852993 | 2,94182392 | 0,01038057 | 0,07098361 |
| 58517_at | RBM25 | -0,20302447 | 9,15279664 | -2,94148004 | 0,01038773 | 0,07100742 |
| 114796_at | PSMG3-AS1 | 0,27157363 | 5,29091374 | 2,94042591 | 0,01040973 | 0,07113256 |
| 653319_at | KIAA0895L | 0,42581832 | 7,8102965 | 2,93965247 | 0,01042591 | 0,07121781 |
| 285962_at | WEE2-AS1 | 0,27202781 | 4,26746571 | 2,9371044 | 0,01047936 | 0,07155755 |
| 84058_at | WDR54 | -0,30813037 | 9,05142534 | -2,93594437 | 0,01050378 | 0,0716989 |
| 100131998_a | RRN3P3 | 0,31265076 | 4,90358155 | 2,93550698 | 0,010513 | 0,07173644 |
| 221302_at | ZUFSP | -0,27181311 | 6,47570458 | -2,93406495 | 0,01054347 | 0,07191884 |
| 26227_at | PHGDH | -0,41193891 | 9,40309295 | -2,93364975 | 0,01055225 | 0,0719533 |
| 8818_at | DPM2 | -0,3733825 | 7,71843115 | -2,93128921 | 0,01060235 | 0,07226931 |
| 26608_at | TBL2 | -0,20346631 | 8,7347052 | -2,93030117 | 0,01062339 | 0,07236243 |
| 51012_at | PRELID3B | -0,3140137 | 8,99774021 | -2,93029478 | 0,01062352 | 0,07236243 |
| 4862_at | NPAS2 | -0,24744917 | 6,72169224 | -2,92927463 | 0,01064529 | 0,07248011 |
| 29901_at | SAC3D1 | -0,72929517 | 8,89081925 | -2,92913248 | 0,01064833 | 0,07248011 |
| 55450_at | CAMK2N1 | 0,32779997 | 10,8456494 | 2,92866136 | 0,01065839 | 0,07252302 |
| 64768_at | IPPK | -0,19749571 | 6,83797527 | -2,92730266 | 0,01068749 | 0,07269528 |
| 9552_at | SPAG7 | 0,2560696 | 8,09946841 | 2,9270157 | 0,01069364 | 0,07271146 |
| 64170_at | CARD9 | 0,32251999 | 5,94061635 | 2,92633693 | 0,01070821 | 0,07278484 |
| 58490_at | RPRD1B | -0,32435737 | 9,04162812 | -2,9259901 | 0,01071566 | 0,07280981 |
| 64118_at | DUS1L | -0,24678664 | 9,2884357 | -2,92556 | 0,01072491 | 0,07284696 |
| 64091_at | POPDC2 | 0,22514559 | 3,86991701 | 2,92452278 | 0,01074725 | 0,07297294 |
| 6434_at | TRA2B | -0,3963567 | 8,64363348 | -2,9242927 | 0,01075221 | 0,0729809 |
| 81620_at | CDT1 | -0,96982327 | 7,71535294 | -2,92380522 | 0,01076273 | 0,07302656 |
| 114882_at | OSBPL8 | 0,32729447 | 8,3004944 | 2,92314028 | 0,01077709 | 0,07309828 |
| 55680_at | RUFY2 | 0,18822502 | 5,24782134 | 2,92229729 | 0,01079533 | 0,0731962 |
| 112479_at | ERI2 | -0,26958051 | 5,8469943 | -2,92134879 | 0,01081588 | 0,07330977 |
| 51276_at | ZNF571 | 0,3394915 | 5,11738799 | 2,92102263 | 0,01082296 | 0,07333194 |
| 10211_at | FLOT1 | -0,42853992 | 8,7879064 | -2,92026287 | 0,01083946 | 0,07341794 |

| | | | | | | |
|---|---|---|---|---|---|---|
| 55770_at | EXOC2 | 0,30484119 | 7,52458967 | 2,91862574 | 0,01087511 | 0,07363348 |
| 8906_at | AP1G2 | 0,37281183 | 9,00006129 | 2,91835808 | 0,01088095 | 0,07364713 |
| 10992_at | SF3B2 | -0,17286593 | 9,92295833 | -2,91803264 | 0,01088805 | 0,07366932 |
| 4793_at | NFKBIB | -0,19016276 | 7,39287263 | -2,91770994 | 0,0108951 | 0,07369113 |
| 399717_at | GATA3-AS1 | 0,24309361 | 6,04477461 | 2,91654584 | 0,01092056 | 0,07383742 |
| 113540_at | CMTM1 | -0,29731461 | 4,77015792 | -2,91600061 | 0,01093251 | 0,07389225 |
| 54822_at | TRPM7 | 0,26849606 | 6,60333761 | 2,91556274 | 0,01094211 | 0,07391024 |
| 51319_at | RSRC1 | -0,59291064 | 7,85237521 | -2,91552937 | 0,01094284 | 0,07391024 |
| 3304_at | HSPA1B | -0,33003524 | 10,7961783 | -2,91465705 | 0,010962 | 0,07401368 |
| 84162_at | KIAA1109 | 0,3222888 | 6,11677296 | 2,91426936 | 0,01097053 | 0,07404528 |
| 339983_at | NAT8L | -0,2623363 | 6,08802844 | -2,91369111 | 0,01098325 | 0,07410521 |
| 10980_at | COPS6 | -0,37921655 | 10,1575296 | -2,91209342 | 0,01101849 | 0,07431694 |
| 100507463_a | PSMB8-AS1 | 1,01833328 | 5,80547383 | 2,91049986 | 0,01105375 | 0,07452864 |
| 8611_at | PPAP2A | 0,75919605 | 9,70126536 | 2,90855402 | 0,01109695 | 0,07479375 |
| 56139_at | PCDHA10 | -0,25182275 | 4,51574723 | -2,90826147 | 0,01110346 | 0,07481145 |
| 84518_at | CNFN | 0,41669036 | 6,24366469 | 2,90802375 | 0,01110875 | 0,07482093 |
| 54897_at | CASZ1 | 0,23644057 | 6,23256852 | 2,90733483 | 0,0111241 | 0,07489814 |
| 79669_at | C3orf52 | 0,54442179 | 7,08937414 | 2,90706318 | 0,01113016 | 0,07491275 |
| 84126_at | ATRIP | -0,49939152 | 6,34145778 | -2,90598534 | 0,01115423 | 0,07504854 |
| 6208_at | RPS14 | -0,18371797 | 10,6181146 | -2,90474233 | 0,01118206 | 0,07520948 |
| 102725022_a | LOC10272502 | -0,22241473 | 5,77013619 | -2,90379896 | 0,01120322 | 0,07532552 |
| 2140_at | EYA3 | -0,18782786 | 6,22273998 | -2,90295502 | 0,01122218 | 0,0754267 |
| 196294_at | IMMP1L | -0,26692152 | 6,16247424 | -2,90276748 | 0,0112264 | 0,07542874 |
| 27330_at | RPS6KA6 | 0,48797706 | 5,19970449 | 2,90251811 | 0,01123201 | 0,07544013 |
| 57088_at | PLSCR4 | 1,11405093 | 5,13549042 | 2,90190655 | 0,01124579 | 0,07550632 |
| 55890_at | GPRC5C | 0,20217324 | 7,31727814 | 2,90093617 | 0,01126768 | 0,07562693 |
| 150290_at | DUSP18 | 0,37504307 | 6,32631376 | 2,89969992 | 0,01129563 | 0,07573402 |
| 10613_at | ERLIN1 | -0,26504074 | 8,96839702 | -2,89966018 | 0,01129653 | 0,07573402 |
| 9925_at | ZBTB5 | 0,29255523 | 7,69463052 | 2,89959765 | 0,01129794 | 0,07573402 |
| 51524_at | TMEM138 | -0,38158745 | 9,04194756 | -2,8993868 | 0,01130272 | 0,07573402 |
| 79612_at | NAA16 | -0,2694058 | 6,43935951 | -2,89936141 | 0,01130329 | 0,07573402 |
| 57505_at | AARS2 | -0,26531326 | 7,12129067 | -2,89847432 | 0,0113234 | 0,07584238 |
| 1111_at | CHEK1 | -1,18995448 | 7,7705997 | -2,89771488 | 0,01134065 | 0,07593148 |
| 57479_at | PRR12 | 0,20365029 | 7,09985569 | 2,89686978 | 0,01135986 | 0,07600959 |
| 7327_at | UBE2G2 | -0,53296869 | 9,0265513 | -2,89685488 | 0,0113602 | 0,07600959 |
| 8554_at | PIAS1 | 0,39092637 | 8,29109847 | 2,89658345 | 0,01136638 | 0,07602201 |
| 1028_at | CDKN1C | 0,38506278 | 5,80256693 | 2,89632696 | 0,01137223 | 0,07602201 |
| 10780_at | ZNF234 | 0,19007394 | 5,60490309 | 2,89625353 | 0,0113739 | 0,07602201 |
| 8509_at | NDST2 | 0,21113787 | 7,3516607 | 2,89432766 | 0,01141787 | 0,07628943 |
| 4126_at | MANBA | 0,41356047 | 7,51586671 | 2,89402729 | 0,01142474 | 0,07628979 |
| 23122_at | CLASP2 | -0,31349608 | 7,24480403 | -2,89397914 | 0,01142585 | 0,07628979 |
| 11078_at | TRIOBP | 0,28460147 | 8,58708155 | 2,89270975 | 0,01145494 | 0,07645754 |
| 3749_at | KCNC4 | -0,28303751 | 5,84509295 | -2,89235753 | 0,01146302 | 0,07648501 |
| 7850_at | IL1R2 | -0,26171947 | 4,08921574 | -2,89171417 | 0,01147781 | 0,07655713 |
| 23362_at | PSD3 | 0,52526056 | 6,99129015 | 2,89131371 | 0,01148702 | 0,07659204 |
| 51218_at | GLRX5 | -0,32388605 | 10,5081117 | -2,88986802 | 0,01152033 | 0,07678759 |
| 252969_at | NEIL2 | -0,35431632 | 6,43484371 | -2,88942871 | 0,01153047 | 0,0768286 |
| 64283_at | ARHGEF28 | 0,50326987 | 6,24522149 | 2,88915852 | 0,01153671 | 0,07684361 |
| 2709_at | GJB5 | -0,42463398 | 8,57924254 | -2,88858656 | 0,01154994 | 0,0769051 |
| 23616_at | SH3BP1 | -0,20758668 | 7,86542963 | -2,88826013 | 0,01155749 | 0,07692731 |

| | | | | | | |
|---|---|---|---|---|---|---|
| 389898_at | UBE2NL | -0,20557333 | 5,72103308 | -2,88809738 | 0,01156126 | 0,07692731 |
| 161742_at | SPRED1 | -1,23743259 | 6,1177103 | -2,88653051 | 0,0115976 | 0,07712617 |
| 29058_at | TMEM230 | 0,13745919 | 10,5229716 | 2,88646351 | 0,01159915 | 0,07712617 |
| 226_at | ALDOA | -0,36337635 | 11,7834586 | -2,88511487 | 0,01163052 | 0,07728462 |
| 80205_at | CHD9 | 0,2909192 | 7,51781371 | 2,88506935 | 0,01163159 | 0,07728462 |
| 103_at | ADAR | 0,39384493 | 10,5923145 | 2,88485445 | 0,01163659 | 0,07728462 |
| 79176_at | FBXL15 | 0,25707616 | 7,23275413 | 2,88462042 | 0,01164205 | 0,07728462 |
| 55558_at | PLXNA3 | 0,22374202 | 6,87700923 | 2,88457766 | 0,01164304 | 0,07728462 |
| 133015_at | PACRGL | -0,32447521 | 6,63389983 | -2,88436428 | 0,01164802 | 0,07729101 |
| 90268_at | OTULIN | -0,23902016 | 6,91174966 | -2,88339837 | 0,01167057 | 0,07741399 |
| 57003_at | CCDC47 | -0,33324547 | 9,18097991 | -2,8830745 | 0,01167814 | 0,07743755 |
| 94056_at | SYAP1 | -0,23748103 | 9,44012117 | -2,88286073 | 0,01168314 | 0,07744405 |
| 128061_at | C1orf131 | -0,17588962 | 8,25266021 | -2,88201017 | 0,01170306 | 0,07754937 |
| 55676_at | SLC30A6 | -0,30298773 | 8,8215288 | -2,88175338 | 0,01170908 | 0,07756258 |
| 9520_at | NPEPPS | 0,34085891 | 10,0116314 | 2,88062605 | 0,01173554 | 0,07771113 |
| 7403_at | KDM6A | 0,65718918 | 7,67052291 | 2,87962476 | 0,01175909 | 0,07784032 |
| 57136_at | APMAP | -0,22236524 | 9,65469415 | -2,87942523 | 0,01176379 | 0,07784467 |
| 587_at | BCAT2 | 0,27657939 | 9,71000678 | 2,87906628 | 0,01177225 | 0,07787389 |
| 55748_at | CNDP2 | 0,21984489 | 8,09278745 | 2,87847277 | 0,01178624 | 0,07793971 |
| 23383_at | MAU2 | 0,23757185 | 8,0059097 | 2,878227 | 0,01179204 | 0,07795131 |
| 51490_at | C9orf114 | -0,2337147 | 7,53723956 | -2,87790461 | 0,01179966 | 0,07797488 |
| 9775_at | EIF4A3 | -0,26682124 | 10,6785406 | -2,87598486 | 0,01184509 | 0,07823528 |
| 84254_at | CAMKK1 | -0,27009814 | 5,96623001 | -2,87576819 | 0,01185023 | 0,07823528 |
| 1191_at | CLU | 1,33125863 | 9,33833894 | 2,8757254 | 0,01185125 | 0,07823528 |
| 26168_at | SENP3 | -0,22091971 | 6,76226128 | -2,8748435 | 0,01187219 | 0,07834667 |
| 54662_at | TBC1D13 | 0,31119705 | 7,68652666 | 2,87437613 | 0,0118833 | 0,07839315 |
| 79635_at | CCDC121 | 0,41594681 | 5,6028563 | 2,87360371 | 0,01190169 | 0,07848757 |
| 30844_at | EHD4 | 0,16380882 | 9,55981616 | 2,87331512 | 0,01190856 | 0,07850604 |
| 2784_at | GNB3 | -0,25192505 | 6,24734746 | -2,87312612 | 0,01191307 | 0,07850888 |
| 147179_at | WIPF2 | 0,15418595 | 8,02440602 | 2,87160137 | 0,01194948 | 0,07872191 |
| 91050_at | CCDC149 | 0,42671858 | 5,39690178 | 2,87104447 | 0,01196281 | 0,07878276 |
| 1645_at | AKR1C1 | -0,48707352 | 10,8765207 | -2,87063995 | 0,0119725 | 0,07881962 |
| 400451_at | FAM174B | 0,37506733 | 9,22025253 | 2,86866809 | 0,01201984 | 0,07909277 |
| 221981_at | THSD7A | 0,47406421 | 3,83983166 | 2,86856987 | 0,0120222 | 0,07909277 |
| 10398_at | MYL9 | 1,39120263 | 7,4271672 | 2,86774708 | 0,01204201 | 0,07917397 |
| 64284_at | RAB17 | 0,31702177 | 6,77931173 | 2,86764403 | 0,01204449 | 0,07917397 |
| 593_at | BCKDHA | 0,31282251 | 8,33958981 | 2,86748463 | 0,01204834 | 0,07917397 |
| 28989_at | NTMT1 | -0,61074761 | 8,09376755 | -2,86737495 | 0,01205098 | 0,07917397 |
| 9671_at | WSCD2 | 0,62102585 | 6,58354621 | 2,86706666 | 0,01205842 | 0,07919581 |
| 256227_at | STEAP1B | -0,23860698 | 3,58526962 | -2,86663309 | 0,01206889 | 0,07923754 |
| 221687_at | RNF182 | 0,54920635 | 4,91360144 | 2,866378 | 0,01207505 | 0,07925098 |
| 2356_at | FPGS | -0,27316948 | 7,73866448 | -2,86554382 | 0,01209522 | 0,07935634 |
| 8603_at | FAM193A | 0,20964223 | 7,99294176 | 2,86497267 | 0,01210905 | 0,0793719 |
| 7077_at | TIMP2 | 1,1053977 | 8,09095158 | 2,86483455 | 0,0121124 | 0,0793719 |
| 9122_at | SLC16A4 | 0,38445023 | 3,09134964 | 2,86478649 | 0,01211356 | 0,0793719 |
| 332_at | BIRC5 | -0,99197299 | 8,68984083 | -2,86476538 | 0,01211407 | 0,0793719 |
| 2107_at | ETF1 | -0,29583508 | 9,75792541 | -2,86450805 | 0,01212031 | 0,07938578 |
| 55187_at | VPS13D | 0,198346 | 7,18837208 | 2,863855 | 0,01213616 | 0,07946256 |
| 81606_at | LBH | 0,96164321 | 8,59009545 | 2,8635767 | 0,01214292 | 0,0794798 |
| 84928_at | TMEM209 | -0,32122492 | 8,51896839 | -2,86315988 | 0,01215305 | 0,07951909 |

| | | | | | | |
|---|---|---|---|---|---|---|
| 6468_at | FBXW4 | 0,35731683 | 8,73420231 | 2,86278809 | 0,01216209 | 0,07955124 |
| 84315_at | MON1A | -0,39720967 | 6,23397405 | -2,86232987 | 0,01217325 | 0,07959717 |
| 55218_at | EXD2 | 0,19246736 | 7,49277402 | 2,8617541 | 0,01218728 | 0,07966187 |
| 56704_at | JPH1 | -0,30153509 | 6,08704537 | -2,86103391 | 0,01220485 | 0,07974967 |
| 144404_at | TMEM120B | -0,200409 | 6,12545489 | -2,86059289 | 0,01221562 | 0,07976675 |
| 7170_at | TPM3 | -0,17118551 | 4,75778464 | -2,86058788 | 0,01221575 | 0,07976675 |
| 9231_at | DLG5 | 0,14538552 | 9,5409323 | 2,86022416 | 0,01222464 | 0,07979776 |
| 23767_at | FLRT3 | 1,17328565 | 8,41328302 | 2,85971128 | 0,01223719 | 0,07985261 |
| 4791_at | NFKB2 | 0,50829644 | 6,42707699 | 2,85879438 | 0,01225965 | 0,07997212 |
| 7754_at | ZNF204P | 0,62583599 | 5,58164149 | 2,85745104 | 0,01229264 | 0,08016015 |
| 4014_at | LOR | -0,22907675 | 5,05241448 | -2,85692586 | 0,01230556 | 0,08021724 |
| 54148_at | MRPL39 | -0,41729312 | 5,96837525 | -2,85661253 | 0,01231327 | 0,08024038 |
| 101927623_a | LOC10192762 | -0,34095201 | 3,86474712 | -2,85585428 | 0,01233196 | 0,08033498 |
| 8482_at | SEMA7A | 0,31380019 | 7,27038309 | 2,85416962 | 0,01237358 | 0,08057886 |
| 7094_at | TLN1 | 0,23700611 | 7,0462061 | 2,85384363 | 0,01238165 | 0,08060417 |
| 51442_at | VGLL1 | 0,91296896 | 8,89523361 | 2,85246469 | 0,01241584 | 0,08079944 |
| 9522_at | SCAMP1 | 0,34833128 | 8,33888435 | 2,85221486 | 0,01242205 | 0,08081252 |
| 135398_at | C6orf141 | -0,62005256 | 5,55826606 | -2,85194358 | 0,01242879 | 0,08082907 |
| 5074_at | PAWR | 0,28771421 | 7,97148216 | 2,85159773 | 0,01243739 | 0,0808577 |
| 10541_at | ANP32B | -0,38412583 | 10,9212274 | -2,85140539 | 0,01244217 | 0,08086151 |
| 11167_at | FSTL1 | 0,87610591 | 6,26093006 | 2,85098074 | 0,01245274 | 0,08087676 |
| 3358_at | HTR2C | 0,6982974 | 4,96772416 | 2,85097373 | 0,01245291 | 0,08087676 |
| 1604_at | CD55 | 0,53748305 | 7,36865679 | 2,85062997 | 0,01246148 | 0,08090445 |
| 26355_at | FAM162A | -0,42700023 | 10,1594238 | -2,85021888 | 0,01247172 | 0,08090445 |
| 7567_at | ZNF19 | 0,1702445 | 4,66204718 | 2,8501666 | 0,01247303 | 0,08090445 |
| 800_at | CALD1 | 0,66013635 | 6,39586718 | 2,85012857 | 0,01247398 | 0,08090445 |
| 3914_at | LAMB3 | 0,37468361 | 10,5180553 | 2,84897439 | 0,0125028 | 0,08106408 |
| 9690_at | UBE3C | -0,28665352 | 8,66036241 | -2,84879004 | 0,01250741 | 0,08106668 |
| 493856_at | CISD2 | -0,33845414 | 7,13038751 | -2,84773616 | 0,01253379 | 0,08121035 |
| 8533_at | COPS3 | -0,82243674 | 9,41284139 | -2,84749689 | 0,01253979 | 0,08122189 |
| 114879_at | OSBPL5 | 0,21338764 | 5,96566191 | 2,84730629 | 0,01254457 | 0,08122554 |
| 348654_at | GEN1 | -0,33203309 | 6,60961219 | -2,84702795 | 0,01255155 | 0,08124344 |
| 8775_at | NAPA | 0,44949908 | 8,26600402 | 2,84637487 | 0,01256795 | 0,08132226 |
| 57621_at | ZBTB2 | -0,38740454 | 7,88877375 | -2,84594839 | 0,01257867 | 0,08136429 |
| 84957_at | RELT | -0,35252245 | 7,45100192 | -2,84504716 | 0,01260135 | 0,08148366 |
| 64432_at | MRPS25 | -0,47248584 | 7,25359418 | -2,84399924 | 0,01262778 | 0,08159697 |
| 388962_at | BOLA3 | -0,67672148 | 10,3379848 | -2,84393652 | 0,01262936 | 0,08159697 |
| 2822_at | GPLD1 | 0,4884774 | 5,63170863 | 2,84384844 | 0,01263158 | 0,08159697 |
| 5253_at | PHF2 | 0,17244565 | 8,58060287 | 2,84226032 | 0,01267175 | 0,08182897 |
| 23172_at | FAM175B | -0,1966114 | 8,01022647 | -2,84181925 | 0,01268292 | 0,0818737 |
| 378938_at | MALAT1 | 0,37229493 | 10,5472747 | 2,84058432 | 0,01271427 | 0,08204854 |
| 283149_at | BCL9L | 0,19711387 | 7,60729643 | 2,84026571 | 0,01272236 | 0,08207331 |
| 5826_at | ABCD4 | 0,21370394 | 6,77887665 | 2,83970908 | 0,01273653 | 0,08211927 |
| 84908_at | FAM136A | -0,2678345 | 9,52238711 | -2,83965061 | 0,01273801 | 0,08211927 |
| 51295_at | ECSIT | -0,32386076 | 7,45341324 | -2,83934725 | 0,01274574 | 0,08212988 |
| 8241_at | RBM10 | -0,26911371 | 8,79991274 | -2,83925113 | 0,01274819 | 0,08212988 |
| 23225_at | NUP210 | -0,40171387 | 6,65897802 | -2,83905288 | 0,01275324 | 0,08213496 |
| 25974_at | MMACHC | -0,29168398 | 6,5941515 | -2,83863574 | 0,01276388 | 0,08217599 |
| 55573_at | CDV3 | -0,30233349 | 8,97889918 | -2,836873 | 0,01280892 | 0,08243842 |
| 492311_at | IGIP | 0,29131916 | 5,48860312 | 2,83631584 | 0,01282319 | 0,08250269 |

| | | | | | | |
|---|---|---|---|---|---|---|
| 100188893_a | TOMM6 | -0,23794611 | 10,6177013 | -2,83520177 | 0,01285177 | 0,0826514 |
| 3437_at | IFIT3 | 1,44036905 | 8,41730478 | 2,83500944 | 0,01285671 | 0,0826514 |
| 122773_at | KLHDC1 | 0,22080984 | 3,1695439 | 2,83484325 | 0,01286098 | 0,0826514 |
| 7351_at | UCP2 | 1,54888571 | 9,73921747 | 2,83474644 | 0,01286346 | 0,0826514 |
| 7415_at | VCP | -0,21681959 | 10,3427295 | -2,83432424 | 0,01287432 | 0,08269357 |
| 7867_at | MAPKAPK3 | -0,17432309 | 7,97495325 | -2,83315238 | 0,0129045 | 0,08285771 |
| 7019_at | TFAM | -0,60166238 | 7,77817891 | -2,83299814 | 0,01290848 | 0,08285771 |
| 137075_at | CLDN23 | 0,38921954 | 6,24208871 | 2,83282627 | 0,01291291 | 0,08285855 |
| 1437_at | CSF2 | -0,28615478 | 4,49941914 | -2,83245832 | 0,01292241 | 0,08289188 |
| 92714_at | ARRDC1 | 0,40834511 | 8,39094215 | 2,83043318 | 0,01297479 | 0,08320022 |
| 51102_at | MECR | -0,30877453 | 7,51052486 | -2,83011827 | 0,01298296 | 0,08322487 |
| 1660_at | DHX9 | -0,65166637 | 8,55850375 | -2,82910237 | 0,01300933 | 0,08336619 |
| 10623_at | POLR3C | -0,2645397 | 8,10686612 | -2,82880396 | 0,01301709 | 0,08338436 |
| 22841_at | RAB11FIP2 | 0,32657866 | 8,35398718 | 2,82866032 | 0,01302083 | 0,08338436 |
| 414927_at | MGC34796 | -0,23053352 | 5,28621027 | -2,82705323 | 0,01306269 | 0,0836219 |
| 55011_at | PIH1D1 | -0,22956848 | 8,70084123 | -2,82675294 | 0,01307053 | 0,0836219 |
| 64135_at | IFIH1 | 1,12167409 | 6,82088519 | 2,82673713 | 0,01307094 | 0,0836219 |
| 317649_at | EIF4E3 | 0,30235309 | 6,29856199 | 2,82635799 | 0,01308084 | 0,08365746 |
| 84986_at | ARHGAP19 | -0,38825159 | 7,05916793 | -2,82581773 | 0,01309497 | 0,08367214 |
| 3619_at | INCENP | -0,51320658 | 7,16844724 | -2,82575816 | 0,01309652 | 0,08367214 |
| 84918_at | LRP11 | 0,20635253 | 8,69384404 | 2,82566673 | 0,01309892 | 0,08367214 |
| 101927220_a | DEPDC1-AS1 | -0,29068114 | 3,63092128 | -2,82560566 | 0,01310051 | 0,08367214 |
| 84449_at | ZNF333 | 0,27569522 | 4,30990916 | 2,82541941 | 0,01310539 | 0,08367553 |
| 197257_at | LDHD | 0,23194425 | 4,81437005 | 2,82515405 | 0,01311233 | 0,08369215 |
| 80344_at | DCAF11 | 0,35486071 | 7,89095033 | 2,82310919 | 0,01316599 | 0,08400679 |
| 9541_at | CIR1 | 0,47718985 | 6,65956043 | 2,82277552 | 0,01317477 | 0,08402009 |
| 116842_at | LEAP2 | -0,42311434 | 5,84108375 | -2,82262088 | 0,01317884 | 0,08402009 |
| 8850_at | KAT2B | 1,09335997 | 8,03426225 | 2,8224622 | 0,01318301 | 0,08402009 |
| 993_at | CDC25A | -0,70610567 | 6,24197356 | -2,82236687 | 0,01318552 | 0,08402009 |
| 84336_at | TMEM101 | 0,18294545 | 7,65214331 | 2,82178825 | 0,01320077 | 0,08408941 |
| 7517_at | XRCC3 | -0,47969513 | 6,27330027 | -2,81981039 | 0,01325301 | 0,08439426 |
| 101927278_a | LOC10192727 | -0,20867399 | 4,10849375 | -2,81956914 | 0,01325939 | 0,08440702 |
| 78996_at | C7orf49 | -0,47248649 | 8,65384049 | -2,81872462 | 0,01328177 | 0,08452153 |
| 5440_at | POLR2K | -0,31230453 | 9,2499937 | -2,81825636 | 0,01329419 | 0,08457265 |
| 9173_at | IL1RL1 | -0,20241869 | 4,75872741 | -2,81789783 | 0,01330371 | 0,08459256 |
| 57118_at | CAMK1D | -0,21450953 | 5,97209264 | -2,81780776 | 0,0133061 | 0,08459256 |
| 10312_at | TCIRG1 | 0,36680485 | 8,53672392 | 2,8172872 | 0,01331994 | 0,0846499 |
| 7260_at | TSSC1 | -0,22022199 | 8,12328018 | -2,81713782 | 0,01332391 | 0,0846499 |
| 7390_at | UROS | -0,15966746 | 6,71559213 | -2,81664515 | 0,01333702 | 0,08468456 |
| 100129550_a | LOC10012955 | 0,60832009 | 5,84912872 | 2,81660245 | 0,01333816 | 0,08468456 |
| 643072_at | LOC643072 | 0,38703882 | 6,11827883 | 2,81594919 | 0,01335557 | 0,08476714 |
| 1846_at | DUSP4 | -0,79189629 | 5,90729893 | -2,8150887 | 0,01337853 | 0,0848849 |
| 93_at | ACVR2B | 0,29590624 | 6,6465852 | 2,8141675 | 0,01340315 | 0,08501314 |
| 51735_at | RAPGEF6 | -0,25167862 | 7,40134308 | -2,81363874 | 0,0134173 | 0,0850749 |
| 79752_at | ZFAND1 | -0,35036852 | 8,34159444 | -2,81343525 | 0,01342276 | 0,08508146 |
| 79697_at | C14orf169 | -0,26597349 | 8,19985996 | -2,81295339 | 0,01343567 | 0,08513531 |
| 81605_at | URM1 | -0,16530827 | 6,56503525 | -2,8124673 | 0,01344871 | 0,08518993 |
| 220382_at | FAM181B | 0,66545134 | 6,73911084 | 2,81139039 | 0,01347765 | 0,08534517 |
| 23175_at | LPIN1 | -0,21779909 | 7,08328457 | -2,81076955 | 0,01349436 | 0,0854229 |
| 26034_at | IPCEF1 | 0,26503713 | 3,9598292 | 2,81027697 | 0,01350763 | 0,08547883 |

| | | | | | | |
|---|---|---|---|---|---|---|
| 7594_at | ZNF43 | 0,23241028 | 7,2016612 | 2,80943606 | 0,01353032 | 0,08559428 |
| 79658_at | ARHGAP10 | 0,17493817 | 7,92757267 | 2,80906881 | 0,01354024 | 0,08562892 |
| 84936_at | ZFYVE19 | 0,36758736 | 7,49720959 | 2,80759031 | 0,01358025 | 0,08585375 |
| 5917_at | RARS | -0,29489377 | 10,4744763 | -2,80741702 | 0,01358495 | 0,08585526 |
| 58533_at | SNX6 | 0,24189672 | 9,78909106 | 2,80538181 | 0,01364023 | 0,08612914 |
| 54467_at | ANKIB1 | 0,32686214 | 9,53578756 | 2,80535402 | 0,01364098 | 0,08612914 |
| 96764_at | TGS1 | -0,329093 | 8,30272684 | -2,80532784 | 0,0136417 | 0,08612914 |
| 7297_at | TYK2 | 0,2282532 | 8,11533228 | 2,8041598 | 0,01367352 | 0,08626046 |
| 23511_at | NUP188 | -0,28484031 | 7,37553914 | -2,80409578 | 0,01367527 | 0,08626046 |
| 254251_at | LCORL | 0,34677905 | 6,13658884 | 2,80407161 | 0,01367593 | 0,08626046 |
| 9610_at | RIN1 | -0,24075782 | 6,73622145 | -2,80372731 | 0,01368533 | 0,08629148 |
| 147923_at | ZNF420 | 0,38219603 | 5,92330523 | 2,80294577 | 0,01370668 | 0,08638953 |
| 128872_at | HMGB3P1 | -0,29216503 | 5,22820688 | -2,80259903 | 0,01371617 | 0,08638953 |
| 282974_at | STK32C | -0,23277283 | 6,61944252 | -2,80251909 | 0,01371836 | 0,08638953 |
| 4935_at | GPR143 | 0,45108943 | 4,79327333 | 2,80244583 | 0,01372036 | 0,08638953 |
| 127700_at | OSCP1 | 0,52767046 | 6,07746595 | 2,80233837 | 0,0137233 | 0,08638953 |
| 2733_at | GLE1 | -0,27130567 | 7,68799559 | -2,80195574 | 0,01373378 | 0,08640418 |
| 81563_at | C1orf21 | 0,43352786 | 7,26809477 | 2,80192589 | 0,0137346 | 0,08640418 |
| 55617_at | TASP1 | -0,19084034 | 7,2673141 | -2,80090419 | 0,01376262 | 0,08655219 |
| 2063_at | NR2F6 | 0,20385708 | 8,59214002 | 2,8005833 | 0,01377143 | 0,08657917 |
| 6887_at | TAL2 | -0,25892597 | 4,70827598 | -2,80034344 | 0,01377802 | 0,08657917 |
| 81628_at | TSC22D4 | 0,26801014 | 6,83767948 | 2,79997596 | 0,01378813 | 0,08657917 |
| 1798_at | DPAGT1 | -0,36409345 | 9,70354939 | -2,79995796 | 0,01378862 | 0,08657917 |
| 26263_at | FBXO22 | -0,25184149 | 7,05118127 | -2,79986146 | 0,01379128 | 0,08657917 |
| 9556_at | C14orf2 | -0,35675905 | 9,51956183 | -2,7997669 | 0,01379388 | 0,08657917 |
| 4026_at | LPP | 0,34089086 | 7,83001338 | 2,79928599 | 0,01380712 | 0,08663403 |
| 27351_at | DESI1 | -0,53480274 | 8,79456501 | -2,79779154 | 0,01384833 | 0,08686436 |
| 55785_at | FGD6 | 0,22304237 | 6,1595468 | 2,79668708 | 0,01387887 | 0,08702757 |
| 162427_at | FAM134C | 0,32738687 | 9,26000467 | 2,79648872 | 0,01388436 | 0,08703368 |
| 221830_at | TWISTNB | -0,38127461 | 6,92510041 | -2,79558257 | 0,01390948 | 0,08715176 |
| 196374_at | KRT78 | -0,24762206 | 4,51223968 | -2,79548262 | 0,01391225 | 0,08715176 |
| 8930_at | MBD4 | -0,35568617 | 8,56729418 | -2,7949729 | 0,0139264 | 0,08721203 |
| 23176_at | SEPT8 | 0,43440011 | 8,74861824 | 2,79453329 | 0,01393861 | 0,08726015 |
| 3480_at | IGF1R | 0,26897469 | 8,06197899 | 2,79414023 | 0,01394954 | 0,0873002 |
| 84318_at | CCDC77 | -0,30515185 | 5,68799467 | -2,79314381 | 0,01397729 | 0,08744542 |
| 9842_at | PLEKHM1 | 0,33007853 | 7,9917513 | 2,79229123 | 0,01400107 | 0,08756577 |
| 4361_at | MRE11A | -0,26161826 | 6,23436703 | -2,79211337 | 0,01400603 | 0,0875684 |
| 140688_at | NOL4L | 0,30699166 | 6,61451412 | 2,79184877 | 0,01401343 | 0,08758618 |
| 29893_at | PSMC3IP | -0,93804016 | 7,16571949 | -2,79159615 | 0,01402049 | 0,08760188 |
| 2348_at | FOLR1 | 0,53402422 | 5,50601379 | 2,79067182 | 0,01404635 | 0,08773502 |
| 54885_at | TBC1D8B | 0,51947351 | 6,06060818 | 2,79026287 | 0,01405781 | 0,08777812 |
| 79665_at | DHX40 | 0,27468023 | 8,05252638 | 2,78946673 | 0,01408014 | 0,08788907 |
| 158405_at | KIAA1958 | 0,2119511 | 5,80939848 | 2,78872729 | 0,01410091 | 0,08799021 |
| 84934_at | RITA1 | -0,32755266 | 7,64479577 | -2,78847541 | 0,01410799 | 0,08799638 |
| 26531_at | OR11A1 | -0,2145913 | 3,72070694 | -2,78836722 | 0,01411103 | 0,08799638 |
| 403341_at | ZBTB34 | 0,46634628 | 7,64236646 | 2,78671766 | 0,01415751 | 0,08825764 |
| 780_at | DDR1 | 0,39485141 | 10,5390578 | 2,78568177 | 0,01418677 | 0,08839847 |
| 11333_at | PDAP1 | -0,37060951 | 6,32542961 | -2,78559316 | 0,01418928 | 0,08839847 |
| 1906_at | EDN1 | 0,81239384 | 8,20586824 | 2,7851358 | 0,01420222 | 0,08845048 |
| 80318_at | GKAP1 | 0,25864534 | 4,95411043 | 2,7844124 | 0,01422271 | 0,08854948 |

| | | | | | | |
|---|---|---|---|---|---|---|
| 57062_at | DDX24 | -0,28310568 | 9,55391556 | -2,78379936 | 0,0142401 | 0,08862909 |
| 55137_at | FIGN | 0,27115456 | 5,62834422 | 2,78323594 | 0,0142561 | 0,08867295 |
| 4072_at | EPCAM | -0,32714873 | 10,613978 | -2,78302907 | 0,01426198 | 0,08867295 |
| 8332_at | HIST1H2AL | -0,35887752 | 5,779394 | -2,7829727 | 0,01426358 | 0,08867295 |
| 23409_at | SIRT4 | 0,28279843 | 3,74760162 | 2,7826632 | 0,01427238 | 0,08867295 |
| 10555_at | AGPAT2 | 0,26901608 | 8,91965235 | 2,78262759 | 0,01427339 | 0,08867295 |
| 10233_at | LRRC23 | 0,42589874 | 5,58374652 | 2,7825218 | 0,0142764 | 0,08867295 |
| 54707_at | GPN2 | -0,20072 | 8,64915068 | -2,78241754 | 0,01427937 | 0,08867295 |
| 1474_at | CST6 | 0,61033738 | 8,88673986 | 2,78096598 | 0,01432073 | 0,08888774 |
| 1965_at | EIF2S1 | -0,39969045 | 9,99409385 | -2,78088008 | 0,01432319 | 0,08888774 |
| 84872_at | ZC3H10 | 0,25791406 | 6,38264434 | 2,77873685 | 0,01438449 | 0,08923524 |
| 55717_at | WDR11 | 0,26901508 | 8,39358507 | 2,77859868 | 0,01438845 | 0,08923524 |
| 55681_at | SCYL2 | 0,2664108 | 9,8121189 | 2,77771378 | 0,01441384 | 0,08936065 |
| 283464_at | GXYLT1 | 0,3058282 | 7,34140204 | 2,77757075 | 0,01441795 | 0,08936065 |
| 821_at | CANX | -0,22771958 | 10,7695394 | -2,77718688 | 0,01442898 | 0,08940025 |
| 9627_at | SNCAIP | 0,23097702 | 4,83202331 | 2,77691703 | 0,01443674 | 0,08941957 |
| 51109_at | RDH11 | -0,35108805 | 10,1616651 | -2,77659531 | 0,01444599 | 0,08944084 |
| 152877_at | FAM53A | 0,55969642 | 6,04745365 | 2,77647495 | 0,01444946 | 0,08944084 |
| 143384_at | CACUL1 | 0,271023 | 8,56977154 | 2,77614606 | 0,01445893 | 0,08947071 |
| 84612_at | PARD6B | 0,74740274 | 6,14868829 | 2,77539146 | 0,01448068 | 0,08957654 |
| 79913_at | ACTR5 | -0,34074534 | 6,97742451 | -2,77501485 | 0,01449155 | 0,08959144 |
| 79794_at | C12orf49 | 0,23415175 | 8,77804287 | 2,77498565 | 0,01449239 | 0,08959144 |
| 3028_at | HSD17B10 | -0,27222681 | 9,9948376 | -2,77359615 | 0,01453256 | 0,08981094 |
| 3799_at | KIF5B | -0,20730911 | 9,84972618 | -2,77262007 | 0,01456084 | 0,08995687 |
| 172_at | AFG3L1P | -0,25362902 | 6,56953423 | -2,77164332 | 0,0145892 | 0,09005978 |
| 79712_at | GTDC1 | 0,34399119 | 6,56342545 | 2,77160638 | 0,01459027 | 0,09005978 |
| 79047_at | KCTD15 | -0,64101248 | 5,88245938 | -2,77156321 | 0,01459153 | 0,09005978 |
| 2941_at | GSTA4 | 0,51234838 | 7,01803746 | 2,7708067 | 0,01461353 | 0,09016669 |
| 51542_at | VPS54 | 0,42730769 | 7,86561392 | 2,77017568 | 0,01463191 | 0,09025117 |
| 3098_at | HK1 | -0,20574962 | 9,97358789 | -2,76979054 | 0,01464313 | 0,09029151 |
| 11221_at | DUSP10 | 0,64392987 | 8,44559451 | 2,76945505 | 0,01465292 | 0,0903042 |
| 23321_at | TRIM2 | 0,33738263 | 7,93894857 | 2,7693986 | 0,01465457 | 0,0903042 |
| 3841_at | KPNA5 | 0,35029084 | 4,82373547 | 2,76884333 | 0,01467078 | 0,0903752 |
| 27069_at | GHITM | -0,20045026 | 10,5441064 | -2,76832562 | 0,01468591 | 0,0904395 |
| 2263_at | FGFR2 | 0,67813767 | 6,5668055 | 2,76804871 | 0,01469401 | 0,09046047 |
| 79882_at | ZC3H14 | -0,16441809 | 8,39368206 | -2,76776491 | 0,01470232 | 0,09047723 |
| 5786_at | PTPRA | 0,2701795 | 8,20283215 | 2,7676348 | 0,01470613 | 0,09047723 |
| 92249_at | LINC01278 | 0,20824894 | 6,90835399 | 2,7669249 | 0,01472693 | 0,09057629 |
| 5692_at | PSMB4 | -0,33436226 | 11,2238283 | -2,76637754 | 0,01474299 | 0,09061856 |
| 85477_at | SCIN | 1,46301109 | 5,07100396 | 2,76636997 | 0,01474322 | 0,09061856 |
| 23588_at | KLHDC2 | 0,60355865 | 9,02606234 | 2,76500023 | 0,01478348 | 0,09083705 |
| 23368_at | PPP1R13B | 0,33865759 | 7,43823087 | 2,76472521 | 0,01479158 | 0,09085782 |
| 23205_at | ACSBG1 | 0,59876634 | 4,97322954 | 2,7643155 | 0,01480365 | 0,09090298 |
| 6711_at | SPTBN1 | -0,24043088 | 7,5852375 | -2,76370384 | 0,01482169 | 0,09098475 |
| 65084_at | TMEM135 | 0,23757758 | 8,52551497 | 2,76314526 | 0,01483818 | 0,09105696 |
| 26255_at | PTTG3P | -0,57257481 | 6,31772994 | -2,76294403 | 0,01484412 | 0,09106444 |
| 83732_at | RIOK1 | -0,84266558 | 8,04410874 | -2,76218853 | 0,01486647 | 0,09117247 |
| 80820_at | EEPD1 | -0,22257967 | 6,3098869 | -2,76158733 | 0,01488427 | 0,09125259 |
| 7270_at | TTF1 | -0,17770866 | 6,7018046 | -2,76063912 | 0,01491239 | 0,09137885 |
| 10733_at | PLK4 | -1,02465344 | 6,36813083 | -2,76042694 | 0,01491869 | 0,09137885 |

| | | | | | | |
|---|---|---|---|---|---|---|
| 80311_at | KLHL15 | 0,47226268 | 7,19347389 | 2,76028468 | 0,01492291 | 0,09137885 |
| 390963_at | ZNF818P | 0,51138058 | 5,72444488 | 2,7602536 | 0,01492384 | 0,09137885 |
| 79903_at | NAA60 | 0,15107966 | 8,89903921 | 2,75970101 | 0,01494026 | 0,09145034 |
| 64840_at | PORCN | 0,24969179 | 7,84519109 | 2,75919456 | 0,01495533 | 0,0915135 |
| 10776_at | ARPP19 | -0,3120833 | 9,15104827 | -2,75826974 | 0,01498288 | 0,09165299 |
| 8537_at | BCAS1 | 0,9251819 | 7,00682936 | 2,7580592 | 0,01498916 | 0,09166229 |
| 203286_at | ANKS6 | -0,18945164 | 5,96887517 | -2,75726184 | 0,01501297 | 0,09177873 |
| 1890_at | TYMP | 0,45521742 | 7,85983284 | 2,75676877 | 0,01502771 | 0,09183969 |
| 81855_at | SFXN3 | 0,53399908 | 8,25634981 | 2,75647745 | 0,01503642 | 0,09184686 |
| 5409_at | PNMT | -0,3164892 | 5,24940282 | -2,75641082 | 0,01503841 | 0,09184686 |
| 92400_at | RBM18 | 0,20371584 | 7,94633241 | 2,75572288 | 0,01505902 | 0,09194353 |
| 11201_at | POLI | 0,54167566 | 5,63332106 | 2,7551292 | 0,01507682 | 0,09202305 |
| 8625_at | RFXANK | 0,23359188 | 8,26323002 | 2,75214514 | 0,0151666 | 0,09254175 |
| 7089_at | TLE2 | 0,55754441 | 6,90729723 | 2,75151601 | 0,0151856 | 0,09262831 |
| 8878_at | SQSTM1 | 0,16254078 | 8,15070461 | 2,75066935 | 0,0152112 | 0,09275141 |
| 94240_at | EPSTI1 | 1,28413063 | 5,79384004 | 2,75048774 | 0,0152167 | 0,09275141 |
| 57645_at | POGK | -0,19573819 | 8,80158782 | -2,75037117 | 0,01522023 | 0,09275141 |
| 100507217_a | LINC01578 | 0,24300121 | 9,27824054 | 2,75016538 | 0,01522646 | 0,09276005 |
| 8553_at | BHLHE40 | -0,44221935 | 8,63400861 | -2,74906176 | 0,01525992 | 0,09290444 |
| 55614_at | KIF16B | 0,37649705 | 6,25589175 | 2,74887315 | 0,01526565 | 0,09290444 |
| 728_at | C5AR1 | 0,2108137 | 4,05795014 | 2,74879628 | 0,01526799 | 0,09290444 |
| 5094_at | PCBP2 | -0,18377842 | 8,84143005 | -2,74874792 | 0,01526945 | 0,09290444 |
| 2963_at | GTF2F2 | -0,23077242 | 6,70342877 | -2,74858924 | 0,01527427 | 0,09290444 |
| 6274_at | S100A3 | -0,47956537 | 6,27039911 | -2,74799137 | 0,01529245 | 0,09298563 |
| 8570_at | KHSRP | -0,38237089 | 8,42716292 | -2,74774287 | 0,01530001 | 0,09300225 |
| 4149_at | MAX | 0,14410052 | 7,80391684 | 2,74722745 | 0,0153157 | 0,09304191 |
| 221442_at | ADCY10P1 | 0,428064 | 4,73985294 | 2,74721129 | 0,0153162 | 0,09304191 |
| 1727_at | CYB5R3 | 0,26915065 | 10,0905628 | 2,74694608 | 0,01532428 | 0,09306166 |
| 127262_at | TPRG1L | 0,30282823 | 9,27230876 | 2,74637323 | 0,01534175 | 0,09313839 |
| 163050_at | ZNF564 | 0,35432888 | 6,62764792 | 2,74601581 | 0,01535266 | 0,0931744 |
| 147912_at | SIX5 | 0,21792175 | 6,2412116 | 2,74575487 | 0,01536063 | 0,0931744 |
| 5827_at | PXMP2 | -0,47133079 | 9,07531648 | -2,74570372 | 0,01536219 | 0,0931744 |
| 7701_at | ZNF142 | -0,19853733 | 7,12608298 | -2,74490751 | 0,01538654 | 0,09329269 |
| 6428_at | SRSF3 | -0,26707392 | 9,70677906 | -2,74466591 | 0,01539393 | 0,09330815 |
| 1147_at | CHUK | -0,43293979 | 7,55045155 | -2,74437466 | 0,01540285 | 0,09333285 |
| 8508_at | NIPSNAP1 | 0,25759382 | 9,49037332 | 2,7437493 | 0,01542202 | 0,09339819 |
| 55654_at | TMEM127 | 0,20374476 | 8,52544741 | 2,74370511 | 0,01542337 | 0,09339819 |
| 116064_at | LRRC58 | -0,2235714 | 8,0382549 | -2,74354843 | 0,01542818 | 0,09339819 |
| 51499_at | TRIAP1 | -0,34281993 | 9,41242316 | -2,74277821 | 0,01545183 | 0,09351196 |
| 84296_at | GINS4 | -0,99294642 | 6,74255196 | -2,74212975 | 0,01547177 | 0,09360322 |
| 93627_at | TBCK | 0,38433941 | 8,02838312 | 2,74084282 | 0,01551141 | 0,09381358 |
| 54810_at | GIPC2 | 0,16334481 | 3,28402232 | 2,74032288 | 0,01552745 | 0,09388114 |
| 345462_at | ZNF879 | 0,35866449 | 5,77689165 | 2,73945099 | 0,01555439 | 0,09401452 |
| 55565_at | ZNF821 | 0,22171475 | 5,69881916 | 2,73815515 | 0,01559452 | 0,09422747 |
| 115361_at | GBP4 | 1,55794844 | 5,75792406 | 2,73768831 | 0,015609 | 0,09428538 |
| 10493_at | VAT1 | 0,34088502 | 8,65428639 | 2,73740243 | 0,01561787 | 0,09430941 |
| 162963_at | ZNF610 | 1,27635872 | 5,26694372 | 2,73575405 | 0,01566913 | 0,09458928 |
| 7366_at | UGT2B15 | 0,51213277 | 3,48554218 | 2,73531797 | 0,01568272 | 0,09464165 |
| 4190_at | MDH1 | -0,234255 | 10,1497787 | -2,73460895 | 0,01570483 | 0,09474544 |
| 387032_at | ZKSCAN4 | 0,20318218 | 6,58341095 | 2,73350884 | 0,01573921 | 0,09485459 |

| | | | | | | |
|---|---|---|---|---|---|---|
| 84674_at | CARD6 | 0,5910141 | 8,81233384 | 2,7334721 | 0,01574036 | 0,09485459 |
| 56981_at | PRDM11 | 0,18440116 | 4,93089141 | 2,73340051 | 0,0157426 | 0,09485459 |
| 57402_at | S100A14 | -0,26147956 | 10,7370188 | -2,73339974 | 0,01574262 | 0,09485459 |
| 843_at | CASP10 | 0,19383965 | 5,36771981 | 2,73322869 | 0,01574797 | 0,09485717 |
| 9635_at | CLCA2 | 0,79803664 | 6,80971341 | 2,73229111 | 0,01577735 | 0,09500438 |
| 83787_at | ARMC10 | -0,23104039 | 8,40590832 | -2,73176513 | 0,01579385 | 0,09506676 |
| 55088_at | CCDC186 | 0,34618829 | 6,77137213 | 2,73164632 | 0,01579758 | 0,09506676 |
| 23090_at | ZNF423 | 0,26204307 | 4,95987463 | 2,73120895 | 0,01581131 | 0,0950996 |
| 9139_at | CBFA2T2 | 0,34919352 | 7,10240869 | 2,73115822 | 0,01581291 | 0,0950996 |
| 4254_at | KITLG | 0,23272217 | 6,55902253 | 2,73075151 | 0,01582569 | 0,09514679 |
| 3291_at | HSD11B2 | -0,47440774 | 5,77222428 | -2,73052311 | 0,01583288 | 0,09516028 |
| 7398_at | USP1 | -0,56901652 | 9,32262531 | -2,73011689 | 0,01584566 | 0,09520742 |
| 605_at | BCL7A | 0,32639067 | 7,24526816 | 2,72990431 | 0,01585236 | 0,09521794 |
| 284613_at | CYB561D1 | 0,32957643 | 8,00357498 | 2,72951369 | 0,01586467 | 0,09526217 |
| 3423_at | IDS | 0,33427395 | 7,41472844 | 2,72790286 | 0,01591552 | 0,09553778 |
| 3570_at | IL6R | 0,63047984 | 6,1153133 | 2,72583844 | 0,01598093 | 0,09590053 |
| 25904_at | CNOT10 | -0,39187739 | 7,36629434 | -2,72501855 | 0,01600698 | 0,0960167 |
| 359845_at | FAM101B | -0,76580189 | 7,20812246 | -2,72491549 | 0,01601026 | 0,0960167 |
| 2181_at | ACSL3 | -0,74351264 | 9,81350745 | -2,72405096 | 0,01603778 | 0,09613236 |
| 26780_at | SNORA68 | -0,39476169 | 6,14601466 | -2,72390283 | 0,0160425 | 0,09613236 |
| 53346_at | TM6SF1 | -0,27501723 | 3,7058723 | -2,72383948 | 0,01604452 | 0,09613236 |
| 158747_at | MOSPD2 | 0,26202512 | 6,92020045 | 2,72325357 | 0,0160632 | 0,09621438 |
| 100128288_a | LOC10012828 | 0,37763591 | 4,41949865 | 2,72172589 | 0,01611201 | 0,09647676 |
| 84793_at | FOXD2-AS1 | -0,22898932 | 5,06511227 | -2,721369 | 0,01612344 | 0,09651517 |
| 120935_at | CCDC38 | -0,33172875 | 3,54759764 | -2,72114504 | 0,01613061 | 0,09652811 |
| 58528_at | RRAGD | 0,56411967 | 4,93745311 | 2,71999877 | 0,01616737 | 0,09671806 |
| 53353_at | LRP1B | 0,83852375 | 3,96474278 | 2,71980183 | 0,0161737 | 0,09672585 |
| 157313_at | CDCA2 | -1,06587115 | 7,50550346 | -2,71922639 | 0,01619219 | 0,09680639 |
| 1734_at | DIO2 | 0,44523466 | 7,27616341 | 2,718943 | 0,01620131 | 0,09683083 |
| 55287_at | TMEM40 | 0,24680069 | 8,5019601 | 2,7187811 | 0,01620652 | 0,09683192 |
| 139818_at | DOCK11 | 0,52636957 | 4,75779363 | 2,7181278 | 0,01622755 | 0,09692756 |
| 8968_at | HIST1H3F | -0,25488797 | 5,22952163 | -2,71743078 | 0,01625003 | 0,09703171 |
| 8763_at | CD164 | 0,37682617 | 11,028117 | 2,71582068 | 0,01630206 | 0,09731224 |
| 331_at | XIAP | 0,22128293 | 8,10556803 | 2,71498031 | 0,01632928 | 0,09744454 |
| 65983_at | GRAMD3 | 0,39024794 | 7,78896686 | 2,71322914 | 0,01638615 | 0,0977536 |
| 4133_at | MAP2 | 0,94247962 | 6,51242847 | 2,71247033 | 0,01641085 | 0,09787064 |
| 6541_at | SLC7A1 | -0,50870922 | 7,48330169 | -2,71230396 | 0,01641627 | 0,09787267 |
| 80314_at | EPC1 | 0,36288865 | 7,23309867 | 2,71204763 | 0,01642462 | 0,09789218 |
| 11062_at | DUS4L | -0,31445466 | 6,79982225 | -2,71112775 | 0,01645464 | 0,09804075 |
| 55750_at | AGK | -0,27726284 | 7,50590046 | -2,71062987 | 0,01647091 | 0,09810733 |
| 283335_at | LOC283335 | -0,21533299 | 5,10005381 | -2,70991178 | 0,0164944 | 0,09821689 |
| 8836_at | GGH | -0,47807651 | 8,76904579 | -2,7095276 | 0,01650698 | 0,09826143 |
| 64755_at | C16orf58 | 0,21350178 | 7,69976374 | 2,7080502 | 0,01655545 | 0,09851951 |
| 130752_at | MDH1B | 0,28974078 | 4,11042934 | 2,70769268 | 0,0165672 | 0,09855898 |
| 64066_at | MMP27 | -0,25320892 | 3,48727179 | -2,70740132 | 0,01657678 | 0,09858554 |
| 7536_at | SF1 | -0,22495832 | 9,6677606 | -2,70655569 | 0,01660462 | 0,09872062 |
| 10772_at | SRSF10 | -0,20928253 | 8,01954679 | -2,70566688 | 0,01663393 | 0,09884216 |
| 8165_at | AKAP1 | -0,44141766 | 8,20964439 | -2,70556827 | 0,01663719 | 0,09884216 |
| 79042_at | TSEN34 | -0,17469689 | 9,09302091 | -2,70512043 | 0,01665198 | 0,09884216 |
| 80144_at | FRAS1 | 0,55386072 | 6,30260126 | 2,70511589 | 0,01665213 | 0,09884216 |

| | | | | | | |
|---|---|---|---|---|---|---|
| 346157_at | ZNF391 | 0,17635137 | 4,5827979 | 2,70501243 | 0,01665554 | 0,09884216 |
| 1594_at | CYP27B1 | -0,31727347 | 5,22363941 | -2,70500315 | 0,01665585 | 0,09884216 |
| 4722_at | NDUFS3 | -0,24659611 | 10,061919 | -2,70449982 | 0,01667249 | 0,09891045 |
| 4837_at | NNMT | -0,17627752 | 4,32088461 | -2,70361932 | 0,01670164 | 0,09905288 |
| 51389_at | RWDD1 | -0,32260448 | 9,4008355 | -2,70328169 | 0,01671283 | 0,09908874 |
| 493753_at | COA5 | 0,2363792 | 6,90834157 | 2,70308424 | 0,01671938 | 0,09908955 |
| 2131_at | EXT1 | -0,29684968 | 9,58780333 | -2,70280914 | 0,01672851 | 0,09908955 |
| 6905_at | TBCE | -0,24690424 | 7,60938568 | -2,70277354 | 0,01672969 | 0,09908955 |
| 23509_at | POFUT1 | -0,16907905 | 6,83962625 | -2,70265735 | 0,01673355 | 0,09908955 |
| 84777_at | DLGAP1-AS2 | -0,45384015 | 4,82809577 | -2,70208903 | 0,01675242 | 0,09917085 |
| 201292_at | TRIM65 | -0,47178417 | 6,31328547 | -2,70047991 | 0,01680598 | 0,09942767 |
| 2009_at | EML1 | 0,16619139 | 3,97934777 | 2,70047541 | 0,01680613 | 0,09942767 |
| 80895_at | ILKAP | -0,33041136 | 7,96674555 | -2,70028598 | 0,01681245 | 0,0994345 |
| 29997_at | GLTSCR2 | 0,41037979 | 9,6955555 | 2,69964956 | 0,01683368 | 0,09952954 |
| 55631_at | LRRC40 | -0,34603191 | 8,5388248 | -2,69940816 | 0,01684175 | 0,09954665 |
| 64863_at | METTL4 | -0,42199691 | 7,040997 | -2,69892035 | 0,01685805 | 0,09961245 |
| 159686_at | CFAP58 | 0,18189047 | 3,39818748 | 2,69851534 | 0,01687159 | 0,09966192 |
| 83855_at | KLF16 | -0,24085481 | 7,15368255 | -2,698257 | 0,01688024 | 0,09968243 |
| 5019_at | OXCT1 | -0,43427836 | 6,43108627 | -2,69759621 | 0,01690238 | 0,09978256 |
| 3854_at | KRT6B | -0,24125275 | 5,24974291 | -2,69674214 | 0,01693103 | 0,09990988 |
| 5480_at | PPIC | 0,65898869 | 9,36700276 | 2,6966442 | 0,01693432 | 0,09990988 |
| 55114_at | ARHGAP17 | 0,17595787 | 8,44940028 | 2,69635691 | 0,01694397 | 0,09993621 |
| 3796_at | KIF2A | -0,26104033 | 7,97454453 | -2,69605986 | 0,01695395 | 0,09996449 |
| 5432_at | POLR2C | -0,269138 | 9,23080683 | -2,69563902 | 0,01696811 | 0,10001733 |
| 54741_at | LEPROT | 0,41209129 | 9,58057798 | 2,69510286 | 0,01698616 | 0,10007114 |
| 726_at | CAPN5 | 0,18039566 | 6,90257306 | 2,69500934 | 0,01698931 | 0,10007114 |
| 84259_at | DCUN1D5 | -0,18955104 | 5,38126568 | -2,69485437 | 0,01699453 | 0,10007114 |
| 2329_at | FMO4 | 0,45441506 | 4,49739395 | 2,69461894 | 0,01700246 | 0,10007114 |
| 10236_at | HNRNPR | -0,22428384 | 9,78597661 | -2,69459675 | 0,01700321 | 0,10007114 |
| 26595_at | OR8B2 | 0,23863427 | 3,1081669 | 2,69386506 | 0,0170279 | 0,10016307 |
| 9942_at | XYLB | -0,26673213 | 4,48815301 | -2,69364824 | 0,01703522 | 0,10016307 |
| 84311_at | MRPL45 | -0,22421606 | 9,09094722 | -2,69351342 | 0,01703977 | 0,10016307 |
| 4499_at | MT1M | -0,18091014 | 3,64878187 | -2,6934078 | 0,01704334 | 0,10016307 |
| 4144_at | MAT2A | -0,28590854 | 10,0481658 | -2,69332198 | 0,01704624 | 0,10016307 |
| 6338_at | SCNN1B | 0,75468292 | 5,67748051 | 2,69320994 | 0,01705003 | 0,10016307 |
| 3084_at | NRG1 | -0,21981112 | 5,51251135 | -2,69274825 | 0,01706564 | 0,10022423 |
| 256281_at | NUDT14 | 0,32150279 | 7,97705414 | 2,69245962 | 0,01707541 | 0,10025103 |
| 79725_at | THAP9 | 0,25980321 | 4,7619327 | 2,69198578 | 0,01709146 | 0,10031468 |
| 55872_at | PBK | -1,36203224 | 8,27017706 | -2,69127461 | 0,01711557 | 0,100392 |
| 728730_at | LOC728730 | 0,46342471 | 5,42331154 | 2,69116517 | 0,01711929 | 0,100392 |
| 128240_at | APOA1BP | -0,30315667 | 9,95283421 | -2,69113622 | 0,01712027 | 0,100392 |
| 80097_at | MZT2B | -0,41021661 | 10,4693663 | -2,69067771 | 0,01713584 | 0,10045271 |
| 54916_at | TMEM260 | 0,3130161 | 7,82912807 | 2,69045115 | 0,01714353 | 0,10046726 |
| 55775_at | TDP1 | -0,3045913 | 7,93182211 | -2,68959448 | 0,01717267 | 0,10060741 |
| 116123_at | FMO9P | 0,79059567 | 4,30658123 | 2,68844754 | 0,01721175 | 0,10080573 |
| 6876_at | TAGLN | 1,16769654 | 7,97082911 | 2,68816903 | 0,01722126 | 0,10083073 |
| 10920_at | COPS8 | -0,36834196 | 7,33598588 | -2,68757706 | 0,01724148 | 0,10089143 |
| 10247_at | HRSP12 | -0,29065643 | 8,09994872 | -2,68755876 | 0,0172421 | 0,10089143 |
| 1942_at | EFNA1 | 0,62912681 | 8,11965753 | 2,68680101 | 0,01726801 | 0,10101238 |
| 6586_at | SLIT3 | -0,26977543 | 5,3817564 | -2,68605766 | 0,01729347 | 0,10113058 |

| | | | | | | |
|---|---|---|---|---|---|---|
| 51003_at | MED31 | -0,27920462 | 6,54745257 | -2,68584619 | 0,01730072 | 0,10114227 |
| 23492_at | CBX7 | 0,44574608 | 6,91227317 | 2,68535267 | 0,01731765 | 0,10121052 |
| 128239_at | IQGAP3 | -0,39077223 | 5,22171673 | -2,68439087 | 0,01735069 | 0,10137284 |
| 8773_at | SNAP23 | 0,38255204 | 8,10945507 | 2,68376461 | 0,01737223 | 0,10146391 |
| 9342_at | SNAP29 | 0,15105789 | 7,61344326 | 2,68363164 | 0,01737681 | 0,10146391 |
| 54664_at | TMEM106B | 0,5499463 | 8,61180342 | 2,6832093 | 0,01739136 | 0,10151809 |
| 9289_at | ADGRG1 | 0,42658823 | 9,66862314 | 2,68290446 | 0,01740186 | 0,10154865 |
| 284029_at | LINC00324 | 0,43299807 | 4,66836828 | 2,68127091 | 0,01745827 | 0,10184699 |
| 9651_at | PLCH2 | 0,1940864 | 6,59428546 | 2,6809875 | 0,01746808 | 0,10187333 |
| 100288911_a | LOC10028891 | 0,70704765 | 6,3857916 | 2,68081975 | 0,01747389 | 0,10187635 |
| 3073_at | HEXA | 0,44070221 | 9,00066956 | 2,68035304 | 0,01749005 | 0,10193973 |
| 402682_at | UFSP1 | -0,4005072 | 6,69150524 | -2,67991477 | 0,01750524 | 0,10199742 |
| 100874309_a | EPN2-IT1 | -0,26881637 | 4,69306584 | -2,67893331 | 0,01753931 | 0,10216501 |
| 9768_at | KIAA0101 | -0,85150472 | 9,06720903 | -2,67863851 | 0,01754955 | 0,10219378 |
| 51400_at | PPME1 | -0,28742463 | 9,32938638 | -2,67772603 | 0,0175813 | 0,10234771 |
| 4814_at | NINJ1 | 0,32340079 | 8,68740786 | 2,67729991 | 0,01759614 | 0,10240318 |
| 80727_at | TTYH3 | 0,25590004 | 8,91266281 | 2,67692356 | 0,01760927 | 0,10244859 |
| 338557_at | FFAR4 | -0,29191878 | 3,90809939 | -2,67458224 | 0,01769111 | 0,10289366 |
| 200728_at | TMEM17 | 0,41234847 | 6,11630759 | 2,6740946 | 0,0177082 | 0,10294444 |
| 9568_at | GABBR2 | 0,39457985 | 6,83437118 | 2,67402823 | 0,01771053 | 0,10294444 |
| 866_at | SERPINA6 | -0,27656856 | 4,8144367 | -2,67377046 | 0,01771957 | 0,10296593 |
| 27037_at | TRMT2A | -0,16636208 | 7,57585027 | -2,67331389 | 0,0177356 | 0,10302798 |
| 112817_at | HOGA1 | 0,36339023 | 5,68349715 | 2,67252298 | 0,01776339 | 0,10315834 |
| 81789_at | TIGD6 | 0,31330068 | 4,23117358 | 2,67205505 | 0,01777986 | 0,10322284 |
| 9992_at | KCNE2 | -0,25130091 | 5,00617312 | -2,67098451 | 0,01781758 | 0,1034107 |
| 55277_at | FGGY | 0,62931774 | 6,95007333 | 2,67015752 | 0,01784678 | 0,10354894 |
| 54014_at | BRWD1 | 0,47875985 | 6,43876778 | 2,66941363 | 0,01787308 | 0,10367032 |
| 84451_at | KIAA1804 | -0,24969082 | 7,32544294 | -2,66897169 | 0,01788872 | 0,10371669 |
| 84285_at | EIF1AD | -0,17271924 | 8,84767749 | -2,66888362 | 0,01789184 | 0,10371669 |
| 5950_at | RBP4 | -0,3453512 | 4,81002667 | -2,66857879 | 0,01790264 | 0,10374807 |
| 546_at | ATRX | 0,45049995 | 7,57356566 | 2,66808227 | 0,01792025 | 0,10380205 |
| 54681_at | P4HTM | 0,29819785 | 7,37027272 | 2,66801215 | 0,01792273 | 0,10380205 |
| 10435_at | CDC42EP2 | -0,22378462 | 6,82153595 | -2,66684479 | 0,01796419 | 0,1040109 |
| 4907_at | NT5E | -0,71447892 | 3,96742352 | -2,66642339 | 0,01797918 | 0,10406641 |
| 123207_at | C15orf40 | 0,15345902 | 7,03588791 | 2,66612089 | 0,01798995 | 0,10409746 |
| 389123_at | IQCF2 | -0,24984524 | 3,95670842 | -2,66585962 | 0,01799926 | 0,10412002 |
| 57684_at | ZBTB26 | 0,27322248 | 5,86537066 | 2,66568994 | 0,0180053 | 0,10412373 |
| 3373_at | HYAL1 | -0,31580179 | 4,99379453 | -2,66545396 | 0,01801371 | 0,1041411 |
| 5521_at | PPP2R2B | 0,64009807 | 8,94897633 | 2,66437978 | 0,01805205 | 0,10433141 |
| 55561_at | CDC42BPG | 0,1849518 | 6,40956623 | 2,66370077 | 0,01807632 | 0,10444036 |
| 9555_at | H2AFY | -0,13927783 | 10,2295206 | -2,66325942 | 0,01809211 | 0,10450027 |
| 7286_at | TUFT1 | 0,36938696 | 9,43574392 | 2,66289196 | 0,01810528 | 0,10454494 |
| 28655_at | TRAV27 | -0,31747988 | 5,00817376 | -2,66265509 | 0,01811376 | 0,10456261 |
| 3156_at | HMGCR | -0,25050841 | 9,28100743 | -2,66249983 | 0,01811933 | 0,1045634 |
| 23064_at | SETX | 0,15061407 | 7,39460809 | 2,6623367 | 0,01812518 | 0,10456583 |
| 3693_at | ITGB5 | -1,00015756 | 8,41926262 | -2,6611894 | 0,01816637 | 0,10477209 |
| 23178_at | PASK | -0,46752498 | 6,52089028 | -2,66029408 | 0,01819858 | 0,10492644 |
| 171586_at | ABHD3 | -0,37294153 | 8,5990402 | -2,65948132 | 0,01822786 | 0,10506384 |
| 1969_at | EPHA2 | -0,4779249 | 9,54235585 | -2,6592311 | 0,01823689 | 0,10508442 |
| 23052_at | ENDOD1 | -0,30514449 | 7,09857165 | -2,65900741 | 0,01824496 | 0,10509949 |

| | | | | | | |
|---|---|---|---|---|---|---|
| 57205_at | ATP10D | 0,59980001 | 8,564604 | 2,65854839 | 0,01826154 | 0,10516352 |
| 401647_at | GOLGA7B | -0,71289677 | 5,55609576 | -2,65805979 | 0,0182792 | 0,10521694 |
| 250_at | ALPP | 0,91940267 | 6,59156907 | 2,6579895 | 0,01828174 | 0,10521694 |
| 28227_at | PPP2R3B | -0,38242363 | 6,26719204 | -2,65760353 | 0,0182957 | 0,10526585 |
| 7428_at | VHL | 0,23450327 | 7,73506163 | 2,65685497 | 0,01832281 | 0,10539035 |
| 64965_at | MRPS9 | -0,3539875 | 7,66713285 | -2,65593775 | 0,01835608 | 0,10553871 |
| 353299_at | RGSL1 | -0,20873532 | 4,47007822 | -2,65584193 | 0,01835956 | 0,10553871 |
| 26030_at | PLEKHG3 | -0,23433502 | 7,04158227 | -2,65459228 | 0,01840499 | 0,10576831 |
| 10159_at | ATP6AP2 | 0,27833975 | 10,516305 | 2,65384152 | 0,01843234 | 0,10589387 |
| 55821_at | ALLC | -0,28589926 | 4,4144159 | -2,6535945 | 0,01844135 | 0,10591402 |
| 7078_at | TIMP3 | -0,43559093 | 10,0044436 | -2,65274053 | 0,01847251 | 0,10603108 |
| 100996490_a | LINC01206 | -0,23291207 | 3,67449977 | -2,65269291 | 0,01847425 | 0,10603108 |
| 130940_at | CCDC148 | 0,15578691 | 3,81797236 | 2,65258377 | 0,01847824 | 0,10603108 |
| 4848_at | CNOT2 | 0,18055506 | 8,86552287 | 2,65229147 | 0,01848892 | 0,10606079 |
| 552889_at | ATXN7L3B | 0,47076397 | 9,06503585 | 2,65209786 | 0,018496 | 0,10606982 |
| 1513_at | CTSK | 0,66659289 | 5,54875086 | 2,65186257 | 0,01850461 | 0,10607229 |
| 150737_at | TTC30B | 0,4242716 | 6,23137945 | 2,65178502 | 0,01850745 | 0,10607229 |
| 5511_at | PPP1R8 | -0,25423255 | 9,17940239 | -2,65154448 | 0,01851625 | 0,10609119 |
| 100287808_a | LOC10028780 | -0,17311146 | 3,93390375 | -2,65138224 | 0,01852219 | 0,10609367 |
| 79133_at | NDUFAF5 | -0,18289134 | 6,38228352 | -2,65074516 | 0,01854554 | 0,10617842 |
| 148789_at | B3GALNT2 | -0,24670935 | 6,27991226 | -2,6506777 | 0,01854801 | 0,10617842 |
| 1371_at | CPOX | -0,34438462 | 7,86466853 | -2,65041436 | 0,01855767 | 0,10620216 |
| 1186_at | CLCN7 | 0,21290985 | 7,51245623 | 2,64999377 | 0,01857311 | 0,10625894 |
| 149685_at | ADIG | -0,26737081 | 5,51214775 | -2,64884248 | 0,01861543 | 0,10646944 |
| 3052_at | HCCS | -0,14572672 | 8,89700567 | -2,64847943 | 0,0186288 | 0,10649679 |
| 10024_at | TROAP | -0,62962506 | 7,97059587 | -2,64841227 | 0,01863127 | 0,10649679 |
| 83666_at | PARP9 | 0,73051691 | 7,83795354 | 2,64812883 | 0,01864171 | 0,10650215 |
| 54504_at | CPVL | -0,6623846 | 5,61793946 | -2,64808668 | 0,01864327 | 0,10650215 |
| 10025_at | MED16 | 0,29150981 | 8,423497 | 2,64769895 | 0,01865756 | 0,10655221 |
| 55920_at | RCC2 | -0,25622249 | 10,4835517 | -2,64671389 | 0,01869393 | 0,10672825 |
| 145853_at | C15orf61 | -0,32510622 | 7,25761391 | -2,64587329 | 0,01872501 | 0,10684881 |
| 29116_at | MYLIP | 0,72625437 | 8,69874838 | 2,64582687 | 0,01872673 | 0,10684881 |
| 26511_at | CHIC2 | 0,45335803 | 7,45635926 | 2,64569306 | 0,01873168 | 0,10684881 |
| 654434_at | SNHG20 | -0,56614551 | 5,31263002 | -2,6454041 | 0,01874239 | 0,1068782 |
| 10982_at | MAPRE2 | 0,18424121 | 7,95111575 | 2,64440189 | 0,01877955 | 0,1070387 |
| 4131_at | MAP1B | -0,16284363 | 4,4093999 | -2,64434535 | 0,01878164 | 0,1070387 |
| 2537_at | IFI6 | 1,47131819 | 10,4050141 | 2,64396326 | 0,01879583 | 0,10708788 |
| 8349_at | HIST2H2BE | 0,79209202 | 6,36580713 | 2,64344814 | 0,01881498 | 0,10711851 |
| 928_at | CD9 | 0,20882384 | 11,4376469 | 2,6434433 | 0,01881516 | 0,10711851 |
| 9529_at | BAG5 | -0,15515547 | 8,67992783 | -2,64332505 | 0,01881955 | 0,10711851 |
| 28957_at | MRPS28 | -0,2768717 | 8,85657518 | -2,64322024 | 0,01882345 | 0,10711851 |
| 60625_at | DHX35 | -0,27288158 | 7,06280611 | -2,64303023 | 0,01883052 | 0,10712709 |
| 8814_at | CDKL1 | -0,19313625 | 4,15274639 | -2,64115123 | 0,01890057 | 0,10749383 |
| 3123_at | HLA-DRB1 | 0,90974318 | 5,19949874 | 2,64077161 | 0,01891475 | 0,1075102 |
| 150684_at | COMMD1 | 0,18745076 | 7,69591952 | 2,64075658 | 0,01891531 | 0,1075102 |
| 6633_at | SNRPD2 | -0,32688759 | 10,6184154 | -2,64062608 | 0,01892019 | 0,1075102 |
| 25787_at | DGCR9 | -0,2073662 | 4,82820787 | -2,63979388 | 0,01895132 | 0,10763453 |
| 8455_at | ATRN | 0,24696242 | 8,35741059 | 2,63974257 | 0,01895324 | 0,10763453 |
| 22832_at | CEP162 | 0,30925963 | 5,1827437 | 2,63913944 | 0,01897584 | 0,10770864 |
| 23550_at | PSD4 | 0,38424861 | 6,29578798 | 2,63898515 | 0,01898163 | 0,10770864 |

| | | | | | | |
|---|---|---|---|---|---|---|
| 159_at | ADSS | -0,30818654 | 8,6766815 | -2,6389467 | 0,01898307 | 0,10770864 |
| 80222_at | TARS2 | -0,18174746 | 7,56697515 | -2,63827584 | 0,01900824 | 0,1077906 |
| 55334_at | SLC39A9 | 0,17400531 | 9,14042171 | 2,63826354 | 0,0190087 | 0,1077906 |
| 87178_at | PNPT1 | -0,82149292 | 8,53990699 | -2,63741104 | 0,01904074 | 0,10793296 |
| 11218_at | DDX20 | -0,39352453 | 7,63236746 | -2,63729744 | 0,01904502 | 0,10793296 |
| 4719_at | NDUFS1 | -0,23885041 | 7,74737107 | -2,63640861 | 0,01907848 | 0,10808946 |
| 89894_at | TMEM116 | 0,73945964 | 7,58932837 | 2,63621421 | 0,01908581 | 0,10808946 |
| 57787_at | MARK4 | 0,18143358 | 8,63618706 | 2,63611728 | 0,01908946 | 0,10808946 |
| 9134_at | CCNE2 | -0,88263483 | 7,50037342 | -2,63487056 | 0,01913653 | 0,10832411 |
| 1027_at | CDKN1B | 0,56372511 | 9,78024222 | 2,63458003 | 0,01914751 | 0,1083508 |
| 90693_at | CCDC126 | 0,32564012 | 6,0781138 | 2,63444831 | 0,01915249 | 0,1083508 |
| 5279_at | PIGC | 0,19560534 | 8,4206001 | 2,63299829 | 0,01920742 | 0,10862961 |
| 342979_at | PALM3 | 0,51608633 | 6,40075283 | 2,63217439 | 0,01923869 | 0,1086994 |
| 85459_at | CEP295 | -0,29712329 | 6,46272051 | -2,63214303 | 0,01923988 | 0,1086994 |
| 117178_at | SSX2IP | -0,23077826 | 6,63747145 | -2,63199006 | 0,0192457 | 0,1086994 |
| 259230_at | SGMS1 | 0,38663768 | 7,90886967 | 2,63189277 | 0,01924939 | 0,1086994 |
| 84952_at | CGNL1 | 0,67996 | 5,40302007 | 2,63188676 | 0,01924962 | 0,1086994 |
| 29775_at | CARD10 | -0,30411551 | 8,79845841 | -2,63178174 | 0,01925361 | 0,1086994 |
| 3776_at | KCNK2 | -0,25566505 | 5,04602024 | -2,6307806 | 0,01929171 | 0,10888258 |
| 10121_at | ACTR1A | 0,15508839 | 9,19707523 | 2,62923373 | 0,01935072 | 0,10913946 |
| 205428_at | C3orf58 | 0,41152318 | 7,04905759 | 2,62916573 | 0,01935332 | 0,10913946 |
| 4815_at | NINJ2 | -0,25786901 | 4,49968724 | -2,62914201 | 0,01935422 | 0,10913946 |
| 2595_at | GANC | 0,3343968 | 6,40427274 | 2,62860761 | 0,01937465 | 0,10922269 |
| 4931_at | NVL | -0,32091065 | 7,24503911 | -2,62725231 | 0,01942656 | 0,10947225 |
| 9984_at | THOC1 | -0,5117246 | 7,75694617 | -2,62715505 | 0,01943029 | 0,10947225 |
| 3763_at | KCNJ6 | -0,21669071 | 3,96185076 | -2,62683258 | 0,01944266 | 0,10950993 |
| 26999_at | CYFIP2 | 0,34880481 | 5,73430537 | 2,6255069 | 0,0194936 | 0,10976219 |
| 1892_at | ECHS1 | -0,30701829 | 10,3800918 | -2,62537064 | 0,01949884 | 0,10976219 |
| 5298_at | PI4KB | 0,26937917 | 7,98554643 | 2,62477257 | 0,01952187 | 0,10983012 |
| 4319_at | MMP10 | -0,52703901 | 4,44000024 | -2,62469181 | 0,01952499 | 0,10983012 |
| 129881_at | CCDC173 | -0,20943572 | 3,16338449 | -2,62461319 | 0,01952802 | 0,10983012 |
| 22902_at | RUFY3 | 0,28679834 | 5,89918245 | 2,62375589 | 0,01956108 | 0,10998399 |
| 9246_at | UBE2L6 | 0,88264001 | 9,17609437 | 2,62331403 | 0,01957815 | 0,11004781 |
| 33_at | ACADL | 0,30475696 | 4,99604953 | 2,62279166 | 0,01959834 | 0,11012918 |
| 85027_at | SMIM3 | -0,26284797 | 7,09578123 | -2,62197547 | 0,01962993 | 0,11027452 |
| 100131117_a | ZBTB20-AS1 | -0,25737344 | 3,84226223 | -2,62143112 | 0,01965103 | 0,11034868 |
| 54363_at | HAO1 | -0,19110748 | 3,39840844 | -2,62133925 | 0,01965459 | 0,11034868 |
| 84436_at | ZNF528 | 0,62862907 | 6,21321674 | 2,62045408 | 0,01968894 | 0,11050936 |
| 5445_at | PON2 | 0,4545168 | 9,72392347 | 2,62002949 | 0,01970545 | 0,11056976 |
| 22869_at | ZNF510 | 0,24990097 | 5,73475058 | 2,61954536 | 0,01972428 | 0,11064319 |
| 23077_at | MYCBP2 | 0,22591952 | 8,54044881 | 2,61924913 | 0,01973581 | 0,11067564 |
| 10165_at | SLC25A13 | -0,36260292 | 8,35974702 | -2,61902809 | 0,01974441 | 0,11069169 |
| 79751_at | SLC25A22 | -0,36813891 | 7,0016855 | -2,61876337 | 0,01975473 | 0,11071729 |
| 416_at | ARSF | -0,23029461 | 5,53248412 | -2,61803525 | 0,01978312 | 0,11084419 |
| 80119_at | PIF1 | -0,53074981 | 6,62180874 | -2,6178326 | 0,01979103 | 0,11084475 |
| 146057_at | TTBK2 | 0,16006394 | 4,77490104 | 2,61762454 | 0,01979916 | 0,11084475 |
| 6432_at | SRSF7 | -0,37995185 | 9,38826091 | -2,61749166 | 0,01980435 | 0,11084475 |
| 7692_at | ZNF133 | 0,19155646 | 6,93180121 | 2,61744321 | 0,01980624 | 0,11084475 |
| 253832_at | ZDHHC20 | 0,2066906 | 8,54963164 | 2,61692249 | 0,0198266 | 0,11092644 |
| 27291_at | R3HCC1L | 0,21703116 | 6,53735045 | 2,6165222 | 0,01984226 | 0,11097047 |

| | | | | | | |
|---|---|---|---|---|---|---|
| 200014_at | CC2D1B | 0,25535222 | 7,40728005 | 2,61642686 | 0,01984599 | 0,11097047 |
| 127428_at | TCEANC2 | -0,15294623 | 5,97130007 | -2,61623551 | 0,01985348 | 0,11098015 |
| 162655_at | ZNF519 | -0,22247638 | 4,06584669 | -2,6159649 | 0,01986408 | 0,11100718 |
| 84190_at | METTL25 | 0,21760707 | 6,68481577 | 2,61536117 | 0,01988775 | 0,11110721 |
| 10573_at | MRPL28 | -0,20298656 | 9,0298771 | -2,6150527 | 0,01989985 | 0,1111426 |
| 83593_at | RASSF5 | 0,24279732 | 7,13860143 | 2,61469618 | 0,01991385 | 0,11118853 |
| 84333_at | PCGF5 | 0,25451268 | 7,38505527 | 2,61437374 | 0,01992651 | 0,11122702 |
| 129138_at | ANKRD54 | -0,30979434 | 7,72789177 | -2,61329649 | 0,01996889 | 0,11143128 |
| 83743_at | GRWD1 | -0,37742397 | 8,2794462 | -2,61280685 | 0,01998818 | 0,11150662 |
| 51719_at | CAB39 | 0,30236906 | 9,37412016 | 2,61240452 | 0,02000405 | 0,11156281 |
| 545_at | ATR | -0,35569847 | 8,07299654 | -2,61200463 | 0,02001983 | 0,1116185 |
| 5284_at | PIGR | 0,20502001 | 5,46145638 | 2,61120602 | 0,02005138 | 0,11176205 |
| 10678_at | B3GNT2 | -0,26133117 | 7,32759253 | -2,61027685 | 0,02008814 | 0,1119346 |
| 84698_at | CAPS2 | 0,29725979 | 3,23713337 | 2,60915879 | 0,02013247 | 0,11214916 |
| 63891_at | RNF123 | 0,23894433 | 6,99394017 | 2,60864103 | 0,02015303 | 0,11223123 |
| 784_at | CACNB3 | 0,3547893 | 7,23580456 | 2,60797175 | 0,02017964 | 0,11234692 |
| 1543_at | CYP1A1 | 1,34249431 | 7,42506654 | 2,60732848 | 0,02020524 | 0,11245697 |
| 101060264_a | FOXP4-AS1 | -0,27009274 | 4,21001635 | -2,60653629 | 0,02023682 | 0,11260017 |
| 220213_at | OTUD1 | 0,69069198 | 9,03986572 | 2,60622104 | 0,02024939 | 0,11263762 |
| 54965_at | PIGX | -0,3131054 | 8,34714004 | -2,60558061 | 0,02027497 | 0,11274733 |
| 51550_at | CINP | -0,38895556 | 8,47521657 | -2,60530544 | 0,02028597 | 0,11277455 |
| 6303_at | SAT1 | 0,75080171 | 9,73038491 | 2,60516527 | 0,02029157 | 0,11277455 |
| 7767_at | ZNF224 | 0,52445343 | 6,0708226 | 2,60464243 | 0,02031249 | 0,11285825 |
| 27229_at | TUBGCP4 | -0,66211114 | 8,70864781 | -2,60394466 | 0,02034044 | 0,11298096 |
| 102723775_a | LINC01315 | -0,23389799 | 4,2826411 | -2,60372048 | 0,02034943 | 0,1129983 |
| 7511_at | XPNPEP1 | -0,31098697 | 9,02776693 | -2,6032269 | 0,02036923 | 0,11307566 |
| 8454_at | CUL1 | -0,24890202 | 9,05124129 | -2,60281606 | 0,02038573 | 0,11313464 |
| 650_at | BMP2 | 0,66056405 | 5,16165294 | 2,60260217 | 0,02039432 | 0,11314973 |
| 28996_at | HIPK2 | 0,21950003 | 7,35837336 | 2,60150662 | 0,0204384 | 0,11335583 |
| 5349_at | FXYD3 | 0,24361282 | 10,7154682 | 2,60138631 | 0,02044324 | 0,11335583 |
| 80071_at | CCDC15 | -0,31300769 | 5,35861864 | -2,60110814 | 0,02045445 | 0,11338534 |
| 6490_at | PMEL | 0,24921012 | 5,60148434 | 2,6008916 | 0,02046318 | 0,11339 |
| 84337_at | ELOF1 | -0,28703832 | 8,33355396 | -2,60079525 | 0,02046706 | 0,11339 |
| 147808_at | ZNF784 | 0,24674627 | 7,34419339 | 2,60061702 | 0,02047425 | 0,11339721 |
| 348262_at | FAM195B | 0,24966572 | 8,2373289 | 2,60013624 | 0,02049365 | 0,11347204 |
| 6228_at | RPS23 | -0,19166232 | 8,53126163 | -2,59920868 | 0,02053114 | 0,11364692 |
| 513_at | ATP5D | -0,24528339 | 9,21631567 | -2,59810313 | 0,0205759 | 0,11386197 |
| 55662_at | HIF1AN | 0,21800537 | 7,87300168 | 2,59736437 | 0,02060586 | 0,11399503 |
| 9927_at | MFN2 | -0,27794568 | 9,33138019 | -2,59667378 | 0,02063391 | 0,11411743 |
| 201725_at | C4orf46 | -0,89813658 | 7,24655546 | -2,59546829 | 0,02068296 | 0,1143281 |
| 10396_at | ATP8A1 | 0,34339712 | 3,96173856 | 2,59544584 | 0,02068387 | 0,1143281 |
| 56259_at | CTNNBL1 | -0,35381854 | 7,25780799 | -2,59520961 | 0,0206935 | 0,11434849 |
| 10559_at | SLC35A1 | 0,49934331 | 8,53402374 | 2,59499565 | 0,02070222 | 0,11435693 |
| 30834_at | ZNRD1 | -0,45020899 | 8,33422224 | -2,59488087 | 0,0207069 | 0,11435693 |
| 60678_at | EEFSEC | -0,28106411 | 7,39529777 | -2,59362253 | 0,02075827 | 0,1146078 |
| 1870_at | E2F2 | -0,34138748 | 6,17555361 | -2,59344683 | 0,02076546 | 0,11461461 |
| 23033_at | DOPEY1 | 0,2248393 | 7,06238943 | 2,59319301 | 0,02077584 | 0,11463906 |
| 7086_at | TKT | -0,20203361 | 10,9739617 | -2,59270773 | 0,0207957 | 0,11471579 |
| 10318_at | TNIP1 | 0,3235323 | 8,65802223 | 2,59239403 | 0,02080855 | 0,11475381 |
| 4089_at | SMAD4 | 0,16829722 | 6,55689476 | 2,58993165 | 0,02090967 | 0,11527848 |

| | | | | | | |
|---|---|---|---|---|---|---|
| 8407_at | TAGLN2 | -0,51021321 | 11,0194003 | -2,58942934 | 0,02093036 | 0,11533954 |
| 130576_at | LYPD6B | 0,62726968 | 9,10033151 | 2,58937193 | 0,02093272 | 0,11533954 |
| 104355292_a | LINC01525 | -0,23768586 | 4,35855306 | -2,58897045 | 0,02094927 | 0,11539773 |
| 9994_at | CASP8AP2 | -0,51921243 | 7,74027098 | -2,58793175 | 0,02099215 | 0,11560085 |
| 49855_at | SCAPER | 0,25522665 | 6,07876257 | 2,58766404 | 0,02100321 | 0,11562872 |
| 6047_at | RNF4 | -0,15550004 | 9,80660382 | -2,58696404 | 0,02103217 | 0,11575506 |
| 160760_at | PPTC7 | -0,24887838 | 8,40992839 | -2,58675283 | 0,02104092 | 0,11577011 |
| 3431_at | SP110 | 0,69609659 | 7,76713014 | 2,58640636 | 0,02105527 | 0,115816 |
| 9097_at | USP14 | -0,4745264 | 9,22224796 | -2,58594177 | 0,02107453 | 0,11588885 |
| 503693_at | LOH12CR2 | 0,37307093 | 4,6121116 | 2,58572858 | 0,02108337 | 0,1159044 |
| 10144_at | FAM13A | 0,28099875 | 7,70626987 | 2,58495761 | 0,02111539 | 0,11604726 |
| 83692_at | CD99L2 | 0,27955946 | 6,72986821 | 2,58473583 | 0,0211246 | 0,11606481 |
| 80347_at | COASY | -0,20730126 | 8,86778985 | -2,58428567 | 0,02114332 | 0,11613454 |
| 11113_at | CIT | -0,54434313 | 7,55460412 | -2,5834443 | 0,02117836 | 0,11627653 |
| 7184_at | HSP90B1 | -0,2168363 | 11,6453297 | -2,58328035 | 0,02118519 | 0,11627653 |
| 1981_at | EIF4G1 | -0,35673479 | 9,32320747 | -2,58323017 | 0,02118728 | 0,11627653 |
| 196541_at | METTL21C | 0,22206465 | 3,06837152 | 2,58276823 | 0,02120655 | 0,11634911 |
| 80271_at | ITPKC | 0,33603929 | 7,27917432 | 2,58215252 | 0,02123225 | 0,11645698 |
| 91351_at | DDX60L | 0,67257118 | 8,82930808 | 2,58186186 | 0,0212444 | 0,11649042 |
| 143279_at | HECTD2 | 0,23489319 | 4,88920037 | 2,58168814 | 0,02125166 | 0,11649708 |
| 54859_at | ELP6 | -0,24714855 | 7,46788988 | -2,58127983 | 0,02126874 | 0,11655753 |
| 8518_at | IKBKAP | -0,16425799 | 7,00944711 | -2,58068409 | 0,02129368 | 0,11666102 |
| 84527_at | ZNF559 | 0,28560133 | 7,54886406 | 2,57997013 | 0,02132361 | 0,11679177 |
| 102157402_a | AK6 | -0,54740773 | 8,25456814 | -2,57804012 | 0,02140471 | 0,11720267 |
| 27087_at | B3GAT1 | -0,25711336 | 5,2299116 | -2,57712161 | 0,02144342 | 0,11738122 |
| 8541_at | PPFIA3 | -0,20451679 | 6,40996443 | -2,5768609 | 0,02145441 | 0,11740806 |
| 192669_at | AGO3 | -0,16466572 | 6,42602246 | -2,57661481 | 0,0214648 | 0,11743153 |
| 55769_at | ZNF83 | 0,44039252 | 6,32659905 | 2,57606828 | 0,02148788 | 0,11752444 |
| 5465_at | PPARA | 0,53342173 | 6,11154777 | 2,57592341 | 0,021494 | 0,11752456 |
| 26175_at | TMEM251 | -0,327647 | 8,47782836 | -2,57536517 | 0,02151761 | 0,11762026 |
| 5627_at | PROS1 | 0,34906444 | 4,64599465 | 2,57392931 | 0,02157845 | 0,11791935 |
| 79098_at | C1orf116 | 0,38680011 | 8,1428614 | 2,57377286 | 0,02158509 | 0,11792218 |
| 2909_at | ARHGAP35 | 0,17720159 | 7,59808573 | 2,57328054 | 0,02160599 | 0,11800291 |
| 3155_at | HMGCL | 0,47538675 | 7,97447935 | 2,57283302 | 0,02162501 | 0,11804542 |
| 7584_at | ZNF35 | -0,20143009 | 6,67654796 | -2,572809 | 0,02162603 | 0,11804542 |
| 154214_at | RNF217 | 0,31874817 | 6,67289164 | 2,57178556 | 0,02166959 | 0,11820184 |
| 2806_at | GOT2 | -0,36222863 | 10,9925265 | -2,57156358 | 0,02167905 | 0,11820184 |
| 9786_at | KIAA0586 | -0,2303939 | 6,91115561 | -2,57147804 | 0,0216827 | 0,11820184 |
| 132320_at | SCLT1 | -0,70653978 | 5,54269384 | -2,57141967 | 0,02168518 | 0,11820184 |
| 8309_at | ACOX2 | -0,30744439 | 4,35831896 | -2,57141529 | 0,02168537 | 0,11820184 |
| 85444_at | LRRCC1 | -0,30400403 | 7,05517874 | -2,57124834 | 0,02169249 | 0,11820235 |
| 80124_at | VCPIP1 | 0,2368505 | 6,80715202 | 2,57112528 | 0,02169774 | 0,11820235 |
| 205327_at | C2orf69 | -0,42574496 | 8,02233528 | -2,57057193 | 0,02172135 | 0,11829754 |
| 4947_at | OAZ2 | 0,29601694 | 9,1419255 | 2,57025639 | 0,02173483 | 0,11833041 |
| 3551_at | IKBKB | 0,55291351 | 7,23753398 | 2,57014301 | 0,02173967 | 0,11833041 |
| 4676_at | NAP1L4 | -0,24047963 | 8,70659576 | -2,569958 | 0,02174758 | 0,11834001 |
| 2825_at | GPR1 | 0,25849974 | 3,10656557 | 2,56954552 | 0,02176522 | 0,11840255 |
| 51203_at | NUSAP1 | -0,7223811 | 9,78082701 | -2,56938296 | 0,02177218 | 0,11840695 |
| 84560_at | MT4 | -0,22028182 | 7,3765745 | -2,56917543 | 0,02178106 | 0,11842182 |
| 10269_at | ZMPSTE24 | -0,26048925 | 10,5612432 | -2,56856303 | 0,02180729 | 0,11853099 |

| | | | | | | |
|---|---|---|---|---|---|---|
| 3237_at | HOXD11 | -0,30743257 | 5,83314744 | -2,5681861 | 0,02182345 | 0,11858537 |
| 253558_at | LCLAT1 | -0,18850237 | 10,1564983 | -2,56643968 | 0,02189848 | 0,11891166 |
| 51523_at | CXXC5 | 0,54939737 | 9,28938001 | 2,56636024 | 0,0219019 | 0,11891166 |
| 86_at | ACTL6A | -0,40001877 | 9,52638163 | -2,56635747 | 0,02190202 | 0,11891166 |
| 149086_at | LINC01225 | -0,29119796 | 4,42585279 | -2,56580126 | 0,02192597 | 0,11900816 |
| 8797_at | TNFRSF10A | -0,92755956 | 7,56669152 | -2,5654754 | 0,02194001 | 0,11905084 |
| 3660_at | IRF2 | 0,6101728 | 8,99101709 | 2,56524183 | 0,02195009 | 0,11907195 |
| 6939_at | TCF15 | -0,42210184 | 6,08866295 | -2,5650193 | 0,02195968 | 0,11908474 |
| 57798_at | GATAD1 | 0,23014115 | 7,55793269 | 2,56464365 | 0,0219759 | 0,11908474 |
| 57491_at | AHRR | 0,32381749 | 5,13840073 | 2,5645097 | 0,02198168 | 0,11908474 |
| 100131454_a | DBIL5P | -0,26008912 | 4,48319287 | -2,56448385 | 0,0219828 | 0,11908474 |
| 166_at | AES | 0,42809196 | 9,90033272 | 2,56442256 | 0,02198545 | 0,11908474 |
| 144455_at | E2F7 | -0,84342132 | 8,0775526 | -2,56425274 | 0,02199278 | 0,11908474 |
| 83452_at | RAB33B | 1,15225319 | 6,68434226 | 2,56418486 | 0,02199572 | 0,11908474 |
| 6782_at | HSPA13 | -0,25685443 | 8,23802251 | -2,563834 | 0,02201089 | 0,11913338 |
| 6647_at | SOD1 | -0,25742942 | 11,3970288 | -2,56341253 | 0,02202912 | 0,1191775 |
| 51512_at | GTSE1 | -0,82700282 | 7,32561887 | -2,56335954 | 0,02203141 | 0,1191775 |
| 9815_at | GIT2 | 0,18639983 | 7,61546304 | 2,56309061 | 0,02204306 | 0,11920701 |
| 3015_at | H2AFZ | -0,80704608 | 11,5310979 | -2,56266767 | 0,02206138 | 0,11927262 |
| 100874291_a | ALMS1-IT1 | -0,37206613 | 4,33277387 | -2,56235048 | 0,02207513 | 0,11931347 |
| 152273_at | FGD5 | -0,32987222 | 5,09404103 | -2,56218778 | 0,02208219 | 0,11931814 |
| 6575_at | SLC20A2 | 0,88754833 | 8,08624458 | 2,56012748 | 0,02217173 | 0,1197684 |
| 84549_at | MAK16 | -0,72200607 | 8,38139041 | -2,55990998 | 0,02218121 | 0,11978598 |
| 57082_at | CASC5 | -1,02721975 | 6,30398171 | -2,55973501 | 0,02218883 | 0,11979357 |
| 8315_at | BRAP | 0,18795313 | 7,36776698 | 2,55940993 | 0,022203 | 0,11981309 |
| 5570_at | PKIB | 0,80728549 | 4,5644747 | 2,55936673 | 0,02220489 | 0,11981309 |
| 126731_at | CCSAP | -0,19353143 | 5,65973359 | -2,55908464 | 0,02221719 | 0,11984592 |
| 55286_at | C4orf19 | 0,46445199 | 7,98683401 | 2,55788556 | 0,02226957 | 0,12009484 |
| 84722_at | PSRC1 | -0,96692508 | 7,78904818 | -2,55683464 | 0,02231558 | 0,12030925 |
| 8479_at | HIRIP3 | -0,37500757 | 6,51406658 | -2,55596396 | 0,02235376 | 0,12048139 |
| 101929505_a | LOC10192950 | -0,19044596 | 3,67331255 | -2,55367813 | 0,0224543 | 0,12098944 |
| 128218_at | TMEM125 | 0,42205958 | 7,17108323 | 2,55251709 | 0,02250554 | 0,1212316 |
| 56922_at | MCCC1 | 0,54332029 | 8,60716029 | 2,55236307 | 0,02251234 | 0,12123435 |
| 54974_at | THG1L | -0,21735176 | 7,57030499 | -2,55173637 | 0,02254005 | 0,12134965 |
| 54058_at | C21orf58 | -0,39583273 | 5,95822477 | -2,5514544 | 0,02255253 | 0,12136195 |
| 2001_at | ELF5 | 0,67950497 | 6,04453813 | 2,55108394 | 0,02256894 | 0,12136195 |
| 26093_at | CCDC9 | -0,21084858 | 6,52524286 | -2,55106515 | 0,02256977 | 0,12136195 |
| 357_at | SHROOM2 | 0,21796894 | 7,93806916 | 2,55085576 | 0,02257905 | 0,12136195 |
| 101928200_a | MRGPRF-AS1 | -0,34163767 | 6,00080231 | -2,55083259 | 0,02258007 | 0,12136195 |
| 26118_at | WSB1 | 0,43953484 | 8,09051244 | 2,55083109 | 0,02258014 | 0,12136195 |
| 285971_at | ZNF775 | 0,3050465 | 6,28303731 | 2,55033966 | 0,02260193 | 0,12144518 |
| 6822_at | SULT2A1 | 1,15150331 | 4,30718265 | 2,54974031 | 0,02262853 | 0,1215542 |
| 124637_at | CYB5D1 | -0,25417015 | 6,72143404 | -2,54824265 | 0,02269513 | 0,12187799 |
| 100507475_a | IDH1-AS1 | 0,41288028 | 4,56902496 | 2,54790649 | 0,02271011 | 0,12190575 |
| 27247_at | NFU1 | -0,22101454 | 8,76430469 | -2,54784249 | 0,02271296 | 0,12190575 |
| 9491_at | PSMF1 | -0,22614609 | 9,25283135 | -2,54746541 | 0,02272977 | 0,12195771 |
| 10365_at | KLF2 | -0,32070148 | 6,1794697 | -2,54734141 | 0,0227353 | 0,12195771 |
| 152217_at | NCBP2-AS2 | -0,26158316 | 8,5352773 | -2,54577064 | 0,02280548 | 0,12229687 |
| 285148_at | IAH1 | 0,22426497 | 8,18330168 | 2,54564217 | 0,02281123 | 0,12229687 |
| 2549_at | GAB1 | 0,21600553 | 5,99495027 | 2,5450911 | 0,0228359 | 0,12238556 |

| | | | | | | |
|---|---|---|---|---|---|---|
| 7461_at | CLIP2 | 0,34006295 | 6,71026637 | 2,54498896 | 0,02284048 | 0,12238556 |
| 199746_at | U2AF1L4 | 0,23555336 | 6,61710277 | 2,54425817 | 0,02287324 | 0,12252706 |
| 202052_at | DNAJC18 | 0,20533518 | 5,66980942 | 2,54345636 | 0,02290925 | 0,12268214 |
| 9728_at | SECISBP2L | 0,67612227 | 6,79890764 | 2,54332986 | 0,02291493 | 0,12268214 |
| 7709_at | ZBTB17 | 0,15657972 | 7,09767063 | 2,54276316 | 0,02294042 | 0,12277484 |
| 64601_at | VPS16 | 0,15917881 | 8,01544341 | 2,54266149 | 0,02294499 | 0,12277484 |
| 828_at | CAPS | 0,59713684 | 8,12204012 | 2,54187508 | 0,02298041 | 0,12291173 |
| 3080_at | CFHR2 | 0,53351772 | 4,06926774 | 2,54181016 | 0,02298334 | 0,12291173 |
| 9429_at | ABCG2 | -0,86318049 | 6,51286416 | -2,54157797 | 0,02299381 | 0,12293358 |
| 90441_at | ZNF622 | -0,19059418 | 7,77193093 | -2,54127136 | 0,02300764 | 0,12295544 |
| 100505806_a | SNHG18 | 0,19817147 | 8,38098059 | 2,54120433 | 0,02301066 | 0,12295544 |
| 2026_at | ENO2 | -0,66882961 | 5,77030116 | -2,54063581 | 0,02303633 | 0,12305847 |
| 51031_at | GLOD4 | -0,18249075 | 8,08296448 | -2,53942124 | 0,02309126 | 0,12331772 |
| 60558_at | GUF1 | -0,31849592 | 7,49127799 | -2,53917245 | 0,02310253 | 0,12333672 |
| 51435_at | SCARA3 | 0,42808899 | 6,48098485 | 2,53898374 | 0,02311108 | 0,12333672 |
| 6563_at | SLC14A1 | 0,51939232 | 5,03288121 | 2,53891867 | 0,02311403 | 0,12333672 |
| 5471_at | PPAT | -0,94224025 | 7,53442471 | -2,53875954 | 0,02312124 | 0,12334104 |
| 113251_at | LARP4 | -0,2078212 | 8,00295706 | -2,53841984 | 0,02313665 | 0,12334239 |
| 65095_at | KRI1 | -0,21310245 | 6,94606277 | -2,53828813 | 0,02314262 | 0,12334239 |
| 6628_at | SNRPB | -0,65111653 | 11,6418131 | -2,53813055 | 0,02314978 | 0,12334239 |
| 440465_at | BAIAP2-AS1 | 0,2854628 | 6,69960353 | 2,53808039 | 0,02315205 | 0,12334239 |
| 10908_at | PNPLA6 | 0,30820578 | 7,35970635 | 2,5379887 | 0,02315621 | 0,12334239 |
| 58496_at | LY6G5B | -0,39502217 | 6,3360319 | -2,53769072 | 0,02316975 | 0,12334239 |
| 360023_at | ZBTB41 | 0,4525009 | 8,47682055 | 2,53767496 | 0,02317046 | 0,12334239 |
| 5654_at | HTRA1 | -0,41129842 | 7,42698178 | -2,53761493 | 0,02317319 | 0,12334239 |
| 204962_at | SLC44A5 | 0,61976841 | 7,17384966 | 2,53748436 | 0,02317912 | 0,12334239 |
| 100302740_a | FAS-AS1 | -0,19117532 | 3,37019189 | -2,53640635 | 0,02322816 | 0,1235692 |
| 116255_at | MOGAT1 | -0,31614511 | 4,70299221 | -2,5355903 | 0,02326535 | 0,12373287 |
| 23043_at | TNIK | -0,22918775 | 4,90819089 | -2,53529114 | 0,023279 | 0,12375527 |
| 7741_at | ZSCAN26 | 0,31272307 | 6,47680934 | 2,53521631 | 0,02328241 | 0,12375527 |
| 9179_at | AP4M1 | 0,3128519 | 7,83574821 | 2,5345424 | 0,02331319 | 0,12385431 |
| 6184_at | RPN1 | -0,29533985 | 11,0050676 | -2,53452675 | 0,0233139 | 0,12385431 |
| 8642_at | DCHS1 | -0,24378561 | 4,97332765 | -2,53420564 | 0,02332858 | 0,12388056 |
| 56910_at | STARD7 | -0,22070326 | 10,8080013 | -2,53413729 | 0,02333171 | 0,12388056 |
| 1586_at | CYP17A1 | -0,25154107 | 4,79001509 | -2,53357464 | 0,02335745 | 0,12395907 |
| 101928269_a | LOC10192826 | -0,26865222 | 3,93948754 | -2,53353284 | 0,02335936 | 0,12395907 |
| 10775_at | POP4 | -0,2115591 | 8,11873304 | -2,53165064 | 0,02344569 | 0,1243829 |
| 90784_at | LOC90784 | -0,29243689 | 7,14631976 | -2,53094399 | 0,02347818 | 0,12452096 |
| 5277_at | PIGA | -0,33878522 | 7,03381899 | -2,53058922 | 0,02349451 | 0,12457326 |
| 27131_at | SNX5 | -0,25223215 | 9,89872619 | -2,52922994 | 0,02355716 | 0,12484874 |
| 22821_at | RASA3 | -0,24540211 | 4,6575287 | -2,52918093 | 0,02355942 | 0,12484874 |
| 5307_at | PITX1 | -0,4720674 | 7,33094906 | -2,52880257 | 0,02357689 | 0,12490696 |
| 57478_at | USP31 | 0,35630103 | 9,21121267 | 2,52846849 | 0,02359233 | 0,12493679 |
| 3613_at | IMPA2 | -0,28114865 | 7,99087311 | -2,52833102 | 0,02359869 | 0,12493679 |
| 1267_at | CNP | 0,1828163 | 7,99865183 | 2,52814098 | 0,02360747 | 0,12493679 |
| 80012_at | PHC3 | 0,22072437 | 7,39937775 | 2,52811948 | 0,02360847 | 0,12493679 |
| 55092_at | TMEM51 | -0,30136592 | 7,66123549 | -2,52722658 | 0,0236498 | 0,12512114 |
| 283177_at | LOC283177 | -0,30904332 | 3,96095577 | -2,52529012 | 0,02373967 | 0,12556215 |
| 64342_at | HS1BP3 | 0,28144352 | 7,05830641 | 2,52498145 | 0,02375403 | 0,12560359 |
| 22861_at | NLRP1 | -0,18577389 | 4,91894585 | -2,52473293 | 0,0237656 | 0,12560518 |

| | | | | | | |
|---|---|---|---|---|---|---|
| 7862_at | BRPF1 | -0,21602461 | 7,33455389 | -2,52469477 | 0,02376737 | 0,12560518 |
| 5887_at | RAD23B | -0,17165035 | 9,6238206 | -2,52434736 | 0,02378355 | 0,1256562 |
| 79180_at | EFHD2 | -0,31062381 | 8,62707801 | -2,52286998 | 0,02385246 | 0,12598572 |
| 7103_at | TSPAN8 | 0,70525045 | 6,3317727 | 2,52223797 | 0,023882 | 0,12610715 |
| 83659_at | TEKT1 | -0,17381493 | 4,49399776 | -2,52123362 | 0,02392901 | 0,12632076 |
| 22826_at | DNAJC8 | -0,36808317 | 7,28685502 | -2,5202557 | 0,02397487 | 0,12652818 |
| 201973_at | PRIMPOL | -0,40324601 | 6,39314016 | -2,51897903 | 0,02403486 | 0,12680488 |
| 10262_at | SF3B4 | -0,29758994 | 8,43491177 | -2,51874369 | 0,02404594 | 0,12680488 |
| 8202_at | NCOA3 | 0,33043808 | 8,78428992 | 2,51872016 | 0,02404704 | 0,12680488 |
| 101929340_a | LOC10192934 | 0,32483829 | 4,74345116 | 2,5182526 | 0,02406906 | 0,12688625 |
| 9092_at | SART1 | -0,18810638 | 8,2841196 | -2,51737656 | 0,02411037 | 0,12706924 |
| 9130_at | FAM50A | 0,1763059 | 9,32252361 | 2,51715119 | 0,02412101 | 0,12709054 |
| 5467_at | PPARD | -0,11308933 | 7,54513828 | -2,51682565 | 0,02413638 | 0,12713676 |
| 26589_at | MRPL46 | -0,3224058 | 8,90237344 | -2,51665312 | 0,02414453 | 0,12714493 |
| 5912_at | RAP2B | 0,35729142 | 8,21607887 | 2,51643677 | 0,02415476 | 0,12716402 |
| 80142_at | PTGES2 | -0,31610252 | 7,7610237 | -2,51600489 | 0,02417519 | 0,127229 |
| 149951_at | COMMD7 | 0,30014209 | 8,6772307 | 2,51589655 | 0,02418031 | 0,127229 |
| 54930_at | HAUS4 | -0,43606267 | 8,45396835 | -2,51552763 | 0,02419778 | 0,1272853 |
| 5051_at | PAFAH2 | 0,33384556 | 6,2037835 | 2,5153914 | 0,02420423 | 0,1272853 |
| 57116_at | ZNF695 | -0,41663354 | 3,7772144 | -2,51514205 | 0,02421604 | 0,1273063 |
| 4792_at | NFKBIA | 0,65164842 | 10,2618516 | 2,5149978 | 0,02422288 | 0,1273063 |
| 3149_at | HMGB3 | -0,31458118 | 8,43753752 | -2,51488877 | 0,02422805 | 0,1273063 |
| 10250_at | SRRM1 | -0,24390572 | 9,74418753 | -2,51448422 | 0,02424724 | 0,12736515 |
| 147949_at | ZNF583 | 0,29175493 | 5,07088987 | 2,51437386 | 0,02425247 | 0,12736515 |
| 387893_at | SETD8 | -0,34139488 | 8,25409284 | -2,5141272 | 0,02426418 | 0,1273801 |
| 25890_at | ABI3BP | 0,65686996 | 4,56969648 | 2,5140353 | 0,02426854 | 0,1273801 |
| 23780_at | APOL2 | 0,35931846 | 7,32556684 | 2,51368008 | 0,02428542 | 0,12743395 |
| 64094_at | SMOC2 | -0,30151909 | 4,48956738 | -2,51317729 | 0,02430932 | 0,12745616 |
| 51626_at | DYNC2LI1 | 0,37391767 | 5,95641298 | 2,5131543 | 0,02431042 | 0,12745616 |
| 55326_at | AGPAT5 | -0,6100341 | 7,59230456 | -2,51297808 | 0,0243188 | 0,12745616 |
| 29102_at | DROSHA | -0,20029558 | 9,1517757 | -2,51295873 | 0,02431972 | 0,12745616 |
| 10505_at | SEMA4F | 0,19571268 | 6,92720925 | 2,51289536 | 0,02432274 | 0,12745616 |
| 387521_at | TMEM189 | -0,28297116 | 9,51402614 | -2,51200296 | 0,02436524 | 0,12764417 |
| 11030_at | RBPMS | 0,46594038 | 7,47944333 | 2,51183798 | 0,02437311 | 0,12765066 |
| 50506_at | DUOX2 | 0,82216191 | 5,6214837 | 2,51145571 | 0,02439134 | 0,12771144 |
| 10456_at | HAX1 | -0,46434166 | 9,27809827 | -2,51111055 | 0,02440782 | 0,12774718 |
| 10390_at | CEPT1 | -0,22151917 | 8,68144992 | -2,51103486 | 0,02441143 | 0,12774718 |
| 9662_at | CEP135 | -0,67757766 | 6,54105926 | -2,51053373 | 0,02443537 | 0,12778318 |
| 79572_at | ATP13A3 | -0,34389167 | 8,26529827 | -2,5105252 | 0,02443578 | 0,12778318 |
| 84532_at | ACSS1 | 0,62620124 | 7,20793021 | 2,51047437 | 0,02443821 | 0,12778318 |
| 27071_at | DAPP1 | 0,80807234 | 6,50920347 | 2,51032861 | 0,02444518 | 0,12778493 |
| 112849_at | L3HYPDH | -0,33146369 | 7,09865803 | -2,50918259 | 0,02450004 | 0,12802353 |
| 128178_at | EDARADD | 0,41735619 | 8,63791367 | 2,50909753 | 0,02450412 | 0,12802353 |
| 146198_at | ZFP90 | 0,21654398 | 7,21701899 | 2,5085526 | 0,02453025 | 0,12812531 |
| 654_at | BMP6 | -0,22881956 | 5,06418717 | -2,50827469 | 0,02454358 | 0,12813623 |
| 3113_at | HLA-DPA1 | 0,37736313 | 4,07278617 | 2,50819262 | 0,02454752 | 0,12813623 |
| 1548_at | CYP2A6 | -0,29169791 | 5,68423791 | -2,50809325 | 0,02455229 | 0,12813623 |
| 23065_at | EMC1 | -0,18408393 | 8,04725944 | -2,50745345 | 0,02458303 | 0,12823533 |
| 5986_at | RFNG | 0,16423473 | 7,67559433 | 2,50742096 | 0,0245846 | 0,12823533 |
| 116138_at | KLHDC3 | 0,16937035 | 8,47406261 | 2,50698142 | 0,02460574 | 0,12826337 |

| | | | | | | |
|---|---|---|---|---|---|---|
| 51517_at | NCKIPSD | -0,25447815 | 7,58020304 | -2,50666316 | 0,02462106 | 0,12826337 |
| 3775_at | KCNK1 | -0,22281181 | 7,33321332 | -2,50661089 | 0,02462357 | 0,12826337 |
| 9825_at | SPATA2 | 0,23408871 | 7,41403057 | 2,50658726 | 0,02462471 | 0,12826337 |
| 54938_at | SARS2 | -0,21632339 | 7,80168595 | -2,50652095 | 0,0246279 | 0,12826337 |
| 65117_at | RSRC2 | -0,15230407 | 8,98081118 | -2,50647906 | 0,02462992 | 0,12826337 |
| 7099_at | TLR4 | -1,06451733 | 6,57810947 | -2,50579298 | 0,02466299 | 0,12840085 |
| 26277_at | TINF2 | 0,27372036 | 8,61704042 | 2,50559826 | 0,02467238 | 0,12841504 |
| 1490_at | CTGF | 0,86021886 | 7,08544867 | 2,50544354 | 0,02467985 | 0,1284192 |
| 10572_at | SIVA1 | -0,54107698 | 8,97848472 | -2,50476668 | 0,02471253 | 0,12855454 |
| 401233_at | HTATSF1P2 | -0,58386567 | 5,22372597 | -2,50457348 | 0,02472187 | 0,12856021 |
| 23367_at | LARP1 | -0,30007862 | 8,78687423 | -2,50446796 | 0,02472697 | 0,12856021 |
| 7422_at | VEGFA | -0,64403638 | 8,32526545 | -2,50402495 | 0,02474839 | 0,12863689 |
| 80775_at | TMEM177 | -0,46723882 | 7,29209125 | -2,50368541 | 0,02476483 | 0,12868758 |
| 100506469_a | TMEM147-AS | -0,44327996 | 6,73487808 | -2,50142606 | 0,02487445 | 0,12919506 |
| 55723_at | ASF1B | -1,1340284 | 9,12043455 | -2,50139617 | 0,0248759 | 0,12919506 |
| 6894_at | TARBP1 | -0,53023861 | 7,27660242 | -2,50111245 | 0,0248897 | 0,12923189 |
| 10715_at | CERS1 | -0,28659056 | 5,0559988 | -2,50088593 | 0,02490072 | 0,12924563 |
| 2323_at | FLT3LG | 0,57462849 | 6,86755186 | 2,5006419 | 0,0249126 | 0,12924563 |
| 133_at | ADM | 0,87279811 | 9,99441396 | 2,50053434 | 0,02491784 | 0,12924563 |
| 3235_at | HOXD9 | 0,1770111 | 7,56946376 | 2,50050674 | 0,02491918 | 0,12924563 |
| 79596_at | RNF219 | -0,22889699 | 7,81452575 | -2,49981837 | 0,02495273 | 0,12934191 |
| 79713_at | IGFLR1 | -0,50074451 | 6,66004098 | -2,49972736 | 0,02495717 | 0,12934191 |
| 51301_at | GCNT4 | 0,30890153 | 5,03378333 | 2,49971259 | 0,02495789 | 0,12934191 |
| 24138_at | IFIT5 | 0,71113974 | 7,85919867 | 2,49923577 | 0,02498116 | 0,12941835 |
| 3017_at | HIST1H2BD | 0,58169196 | 8,07194008 | 2,49913503 | 0,02498608 | 0,12941835 |
| 7106_at | TSPAN4 | -0,28110281 | 6,71114418 | -2,49828567 | 0,02502759 | 0,12959849 |
| 63897_at | HEATR6 | 0,20514834 | 6,61035334 | 2,49813221 | 0,02503509 | 0,12960069 |
| 10801_at | SEPT9 | -0,17609832 | 8,78954854 | -2,49781118 | 0,0250508 | 0,12960069 |
| 54955_at | C1orf109 | -0,55849473 | 7,08116145 | -2,49776137 | 0,02505324 | 0,12960069 |
| 692227_at | SNORD104 | -0,50358344 | 8,76241031 | -2,49772703 | 0,02505492 | 0,12960069 |
| 1360_at | CPB1 | -0,28663709 | 3,66548168 | -2,49632352 | 0,02512373 | 0,12989329 |
| 942_at | CD86 | 0,99522208 | 5,65229015 | 2,49629806 | 0,02512498 | 0,12989329 |
| 101928869_a | GABRG3-AS1 | -0,20424754 | 4,37545623 | -2,4960709 | 0,02513613 | 0,12991608 |
| 80725_at | SRCIN1 | -0,37570813 | 5,93552339 | -2,49579708 | 0,02514958 | 0,12995074 |
| 84804_at | MFSD9 | 0,36581523 | 6,43206057 | 2,49524985 | 0,02517648 | 0,13004966 |
| 738_at | VPS51 | 0,17876945 | 9,64272455 | 2,49513304 | 0,02518223 | 0,13004966 |
| 6406_at | SEMG1 | -0,19603937 | 3,27839287 | -2,49485857 | 0,02519573 | 0,13008454 |
| 376267_at | RAB15 | 0,45703384 | 9,21902682 | 2,49456572 | 0,02521015 | 0,1301241 |
| 89970_at | RSPRY1 | 0,29084726 | 8,32171976 | 2,49436574 | 0,02522 | 0,13014007 |
| 23261_at | CAMTA1 | -0,16108894 | 5,62625782 | -2,49338109 | 0,02526856 | 0,13035571 |
| 3430_at | IFI35 | 1,22254654 | 8,32853637 | 2,49292118 | 0,02529127 | 0,13043793 |
| 3087_at | HHEX | -0,42609082 | 5,30807592 | -2,49262254 | 0,02530602 | 0,13047133 |
| 2971_at | GTF3A | -0,41975518 | 9,6008075 | -2,49248915 | 0,02531262 | 0,13047133 |
| 25801_at | GCA | 0,57630598 | 7,07157173 | 2,49229803 | 0,02532207 | 0,13047133 |
| 5378_at | PMS1 | -0,40327915 | 7,65698489 | -2,49224207 | 0,02532484 | 0,13047133 |
| 84954_at | MPND | 0,28702916 | 7,51889487 | 2,4911383 | 0,02537949 | 0,13071793 |
| 3945_at | LDHB | -0,31749683 | 12,0197003 | -2,49083761 | 0,02539439 | 0,13075975 |
| 11097_at | NUPL2 | -0,26853299 | 7,42402582 | -2,49059035 | 0,02540666 | 0,13078794 |
| 8811_at | GALR2 | -0,24156752 | 4,67579585 | -2,48986146 | 0,02544285 | 0,13089134 |
| 5110_at | PCMT1 | -0,38471582 | 9,71683621 | -2,48982102 | 0,02544486 | 0,13089134 |

| | | | | | | |
|---|---|---|---|---|---|---|
| 7113_at | TMPRSS2 | 0,78611171 | 6,09055032 | 2,48972924 | 0,02544942 | 0,13089134 |
| 55074_at | OXR1 | 0,4107223 | 8,03548438 | 2,48963851 | 0,02545393 | 0,13089134 |
| 3842_at | TNPO1 | -0,22623004 | 10,0755016 | -2,4891976 | 0,02547585 | 0,13096912 |
| 162681_at | C18orf54 | -0,55621822 | 5,37627614 | -2,4889477 | 0,02548828 | 0,13097815 |
| 64077_at | LHPP | 0,20037435 | 6,80577959 | 2,48888898 | 0,02549121 | 0,13097815 |
| 23247_at | KIAA0556 | 0,42724437 | 7,02812996 | 2,48867172 | 0,02550202 | 0,13099878 |
| 79623_at | GALNT14 | -0,63964742 | 7,70989838 | -2,48783183 | 0,02554387 | 0,13115393 |
| 83693_at | HSDL1 | 0,30667442 | 7,23822719 | 2,48779241 | 0,02554584 | 0,13115393 |
| 55812_at | SPATA7 | 0,62612063 | 7,02331459 | 2,48763403 | 0,02555374 | 0,13115953 |
| 51495_at | HACD3 | 0,2963013 | 10,9344549 | 2,48712898 | 0,02557895 | 0,13120094 |
| 5504_at | PPP1R2 | 0,16538078 | 9,50584089 | 2,48710321 | 0,02558024 | 0,13120094 |
| 387837_at | CLEC12B | 0,15731142 | 3,01625425 | 2,48706304 | 0,02558224 | 0,13120094 |
| 85441_at | HELZ2 | 0,37122795 | 6,4716357 | 2,48635686 | 0,02561753 | 0,13134697 |
| 5962_at | RDX | 0,43406625 | 8,15327949 | 2,48600516 | 0,02563513 | 0,1314022 |
| 3135_at | HLA-G | 0,23594077 | 7,69213904 | 2,48554627 | 0,0256581 | 0,13142305 |
| 100130967_a | C6orf99 | -0,26130999 | 4,13895499 | -2,48551735 | 0,02565955 | 0,13142305 |
| 3110_at | MNX1 | -0,23492121 | 6,42774081 | -2,48551506 | 0,02565966 | 0,13142305 |
| 80758_at | PRR7 | -0,59962839 | 6,44608674 | -2,48531894 | 0,02566949 | 0,13143842 |
| 147015_at | DHRS13 | -0,32372223 | 7,90825508 | -2,48475725 | 0,02569765 | 0,13154764 |
| 770_at | CA11 | 0,36257779 | 5,55543321 | 2,48447103 | 0,02571201 | 0,13158618 |
| 7153_at | TOP2A | -0,8427873 | 10,0892762 | -2,48431302 | 0,02571994 | 0,1315918 |
| 6165_at | RPL35A | -0,18498649 | 12,7604489 | -2,48392929 | 0,02573921 | 0,13165543 |
| 7074_at | TIAM1 | -0,33602713 | 7,24908735 | -2,48361988 | 0,02575475 | 0,13169998 |
| 8751_at | ADAM15 | -0,27478337 | 8,26585755 | -2,48151484 | 0,02586077 | 0,13213091 |
| 83591_at | THAP2 | 0,68991293 | 4,29004251 | 2,48146428 | 0,02586332 | 0,13213091 |
| 57369_at | GJD2 | -0,24557951 | 5,7838939 | -2,48137579 | 0,02586779 | 0,13213091 |
| 101927813_a | HMMR-AS1 | -0,60570573 | 4,49015984 | -2,48127274 | 0,02587299 | 0,13213091 |
| 4277_at | MICB | -0,59145134 | 7,52048778 | -2,48126615 | 0,02587332 | 0,13213091 |
| 80063_at | ATF7IP2 | 0,45147298 | 6,46845049 | 2,47988324 | 0,02594323 | 0,1324221 |
| 440757_at | LINC00851 | -0,25123628 | 5,12772668 | -2,47986618 | 0,02594409 | 0,1324221 |
| 5367_at | PMCH | -0,61498628 | 4,62206134 | -2,47931227 | 0,02597214 | 0,13253016 |
| 7559_at | ZNF12 | 0,26768546 | 7,48256367 | 2,47897938 | 0,02598901 | 0,13258114 |
| 376412_at | RNF126P1 | -0,20868443 | 6,03117217 | -2,47866678 | 0,02600487 | 0,13262689 |
| 284309_at | ZNF776 | 0,257631 | 7,95861255 | 2,47841412 | 0,02601769 | 0,13265715 |
| 11261_at | CHP1 | 0,15818635 | 10,3191549 | 2,47827657 | 0,02602467 | 0,13265764 |
| 116933_at | CLRN1-AS1 | -0,26859423 | 3,60715837 | -2,47746983 | 0,02606566 | 0,13281946 |
| 55186_at | SLC25A36 | 0,28100602 | 8,49657097 | 2,47738036 | 0,02607021 | 0,13281946 |
| 54958_at | TMEM160 | -0,24497192 | 8,7331529 | -2,47708864 | 0,02608504 | 0,13285991 |
| 6699_at | SPRR1B | -0,85447465 | 4,17483747 | -2,47602488 | 0,02613922 | 0,13310066 |
| 2117_at | ETV3 | 0,18978908 | 6,92055915 | 2,47569833 | 0,02615587 | 0,13315026 |
| 117247_at | SLC16A10 | 0,27808986 | 5,24479686 | 2,47525361 | 0,02617857 | 0,13323058 |
| 63916_at | ELMO2 | 0,24146785 | 7,85619425 | 2,47475427 | 0,02620407 | 0,13332516 |
| 114899_at | C1QTNF3 | 0,32859378 | 4,65001231 | 2,47460304 | 0,0262118 | 0,13332926 |
| 84984_at | CEP19 | 0,34971824 | 5,54853924 | 2,47328436 | 0,02627929 | 0,13363725 |
| 9698_at | PUM1 | 0,16253782 | 10,165979 | 2,47296743 | 0,02629553 | 0,13368457 |
| 55625_at | ZDHHC7 | 0,18934 | 9,79963118 | 2,47257756 | 0,02631553 | 0,13375093 |
| 10285_at | SMNDC1 | -0,35012518 | 9,86751463 | -2,472285 | 0,02633054 | 0,13379194 |
| 23564_at | DDAH2 | 0,32384177 | 6,54385738 | 2,47162456 | 0,02636447 | 0,13391222 |
| 9050_at | PSTPIP2 | 0,65665293 | 5,20190986 | 2,47155353 | 0,02636812 | 0,13391222 |
| 55300_at | PI4K2B | -0,37412832 | 8,52720226 | -2,47135181 | 0,02637849 | 0,13392959 |

| | | | | | | |
|---|---|---|---|---|---|---|
| 57520_at | HECW2 | 0,59879572 | 4,93026569 | 2,47092607 | 0,02640039 | 0,13400547 |
| 10170_at | DHRS9 | -1,58304894 | 6,41491103 | -2,46985945 | 0,02645534 | 0,13424901 |
| 84290_at | CAPNS2 | 0,43476021 | 8,74788322 | 2,46852263 | 0,02652436 | 0,13456382 |
| 222484_at | LNX2 | 0,25520006 | 8,70207984 | 2,4682562 | 0,02653814 | 0,13459826 |
| 284459_at | HKR1 | 0,18880809 | 6,6997253 | 2,46777278 | 0,02656315 | 0,13468496 |
| 11335_at | CBX3 | -0,47258316 | 10,1573296 | -2,46759296 | 0,02657247 | 0,13468496 |
| 2804_at | GOLGB1 | 0,54992911 | 8,37489575 | 2,46752064 | 0,02657621 | 0,13468496 |
| 10071_at | MUC12 | -0,23592887 | 3,89232534 | -2,46688519 | 0,02660914 | 0,13480471 |
| 11051_at | NUDT21 | -0,36750169 | 9,36214431 | -2,46679467 | 0,02661384 | 0,13480471 |
| 126410_at | CYP4F22 | -0,20840541 | 4,44275545 | -2,46662408 | 0,02662268 | 0,13481408 |
| 100287932_a | TIMM23 | -0,43350367 | 10,1347876 | -2,46516435 | 0,02669852 | 0,13516255 |
| 9377_at | COX5A | -0,17995188 | 11,6112217 | -2,46491161 | 0,02671167 | 0,1351936 |
| 9583_at | ENTPD4 | 0,34815817 | 7,68072011 | 2,46307706 | 0,02680731 | 0,13564201 |
| 23641_at | LDOC1 | 0,56889426 | 8,95536338 | 2,46282361 | 0,02682054 | 0,13566244 |
| 1161_at | ERCC8 | -0,16979738 | 5,38266075 | -2,462719 | 0,02682601 | 0,13566244 |
| 10240_at | MRPS31 | -0,18239251 | 7,8984307 | -2,46259538 | 0,02683247 | 0,13566244 |
| 8647_at | ABCB11 | -0,18043807 | 3,34582903 | -2,46206435 | 0,02686024 | 0,13576594 |
| 60494_at | CCDC81 | -0,19427556 | 3,87588963 | -2,46166118 | 0,02688134 | 0,13576594 |
| 145788_at | C15orf65 | 0,67993703 | 5,29237722 | 2,46158686 | 0,02688523 | 0,13576594 |
| 7532_at | YWHAG | -0,18379847 | 11,3304825 | -2,46156251 | 0,02688651 | 0,13576594 |
| 55758_at | RCOR3 | 0,33651299 | 7,61650479 | 2,46153056 | 0,02688818 | 0,13576594 |
| 2838_at | GPR15 | -0,16694218 | 4,42236008 | -2,4604215 | 0,02694633 | 0,13600476 |
| 29767_at | TMOD2 | 0,31822844 | 4,46132035 | 2,4603457 | 0,0269503 | 0,13600476 |
| 125704_at | FAM69C | -0,26542897 | 5,42421452 | -2,46022461 | 0,02695666 | 0,13600476 |
| 23174_at | ZCCHC14 | 0,16142353 | 6,9402262 | 2,45893547 | 0,02702442 | 0,13631092 |
| 57799_at | RAB40C | 0,22612041 | 7,4112202 | 2,45871161 | 0,0270362 | 0,13633466 |
| 4712_at | NDUFB6 | -0,44849991 | 9,11018633 | -2,45766387 | 0,02709142 | 0,13657733 |
| 22980_at | TCF25 | 0,19584395 | 9,2599315 | 2,45639288 | 0,02715854 | 0,13687989 |
| 79685_at | SAP30L | 0,32643286 | 8,13194675 | 2,45579332 | 0,02719026 | 0,13700391 |
| 93349_at | SP140L | 0,46629469 | 8,0549761 | 2,45550683 | 0,02720543 | 0,13702663 |
| 50650_at | ARHGEF3 | 0,3270017 | 7,20712798 | 2,45543949 | 0,02720899 | 0,13702663 |
| 102060282_a | RASSF1-AS1 | -0,20073837 | 4,1710411 | -2,45524505 | 0,02721929 | 0,13704267 |
| 84300_at | UQCC2 | -0,49398924 | 8,24368508 | -2,45408703 | 0,02728072 | 0,13731604 |
| 149345_at | SHISA4 | 0,42930498 | 6,86033296 | 2,45334214 | 0,0273203 | 0,13747935 |
| 10283_at | CWC27 | -0,20748539 | 7,70214661 | -2,45228415 | 0,02737661 | 0,13772674 |
| 222696_at | ZSCAN23 | 0,12958503 | 3,28987665 | 2,45201508 | 0,02739095 | 0,13774226 |
| 4728_at | NDUFS8 | -0,26915848 | 9,70352007 | -2,45188902 | 0,02739767 | 0,13774226 |
| 79966_at | SCD5 | 0,35152123 | 7,07732014 | 2,4518238 | 0,02740115 | 0,13774226 |
| 23403_at | FBXO46 | 0,23019222 | 7,6686511 | 2,45160313 | 0,02741292 | 0,13776547 |
| 1511_at | CTSG | -0,23325635 | 4,78352654 | -2,45131161 | 0,02742847 | 0,13780769 |
| 7226_at | TRPM2 | -0,58209619 | 6,97930691 | -2,45067601 | 0,02746242 | 0,13793618 |
| 10625_at | IVNS1ABP | -0,31510969 | 8,65658972 | -2,45042033 | 0,02747608 | 0,13793618 |
| 54739_at | XAF1 | 1,80797476 | 7,76802467 | 2,45036132 | 0,02747924 | 0,13793618 |
| 51230_at | PHF20 | 0,15658177 | 7,04271144 | 2,45025468 | 0,02748494 | 0,13793618 |
| 441212_at | RP9P | -0,26825665 | 8,0466131 | -2,45000037 | 0,02749855 | 0,13793618 |
| 344148_at | NCKAP5 | 0,49745035 | 3,62018905 | 2,44995182 | 0,02750114 | 0,13793618 |
| 26984_at | SEC22A | 0,31130616 | 6,34258073 | 2,44989521 | 0,02750417 | 0,13793618 |
| 5604_at | MAP2K1 | -0,3488546 | 10,09443 | -2,44919872 | 0,02754147 | 0,13808727 |
| 100131733_a | USP30-AS1 | 0,37546246 | 4,62618152 | 2,44852706 | 0,02757748 | 0,13823185 |
| 29943_at | PADI1 | -0,67149112 | 6,34503743 | -2,44803957 | 0,02760365 | 0,13832702 |

| | | | | | | |
|---|---|---|---|---|---|---|
| 90025_at | UBE3D | -0,46399396 | 6,33124074 | -2,4475954 | 0,02762751 | 0,13840699 |
| 253512_at | SLC25A30 | -0,21046757 | 5,55887228 | -2,44747504 | 0,02763398 | 0,13840699 |
| 22929_at | SEPHS1 | -0,37413839 | 8,0402742 | -2,44732048 | 0,02764229 | 0,13841262 |
| 221914_at | GPC2 | 0,2914285 | 6,02958965 | 2,44629685 | 0,02769738 | 0,13865174 |
| 23116_at | FAM179B | 0,43901562 | 7,63683382 | 2,44616587 | 0,02770444 | 0,13865174 |
| 6125_at | RPL5 | -0,19271785 | 12,949052 | -2,44519503 | 0,0277568 | 0,13887771 |
| 7070_at | THY1 | -0,18772343 | 5,23195991 | -2,44304158 | 0,02787328 | 0,13942429 |
| 64943_at | NT5DC2 | -0,55961515 | 8,89597781 | -2,44106488 | 0,02798061 | 0,13992482 |
| 9836_at | LCMT2 | -0,30350075 | 8,49113254 | -2,44088294 | 0,0279905 | 0,139938 |
| 101927391_a | LOC10192739 | 0,36154366 | 3,54335933 | 2,44014092 | 0,02803091 | 0,14010364 |
| 5638_at | PRRG1 | 0,2148027 | 7,53030236 | 2,43994605 | 0,02804153 | 0,14012036 |
| 23569_at | PADI4 | -0,56312751 | 4,92514295 | -2,43914086 | 0,02808545 | 0,14030344 |
| 23048_at | FNBP1 | 0,19239694 | 7,59782256 | 2,43884294 | 0,02810172 | 0,14034832 |
| 3467_at | IFNW1 | -0,2120527 | 3,89089053 | -2,43869743 | 0,02810967 | 0,14035162 |
| 10314_at | LANCL1 | 0,32578104 | 8,95972568 | 2,4385056 | 0,02812015 | 0,14035994 |
| 29799_at | YPEL1 | 0,36407371 | 4,84639815 | 2,43836651 | 0,02812775 | 0,14035994 |
| 30849_at | PIK3R4 | -0,2259251 | 8,51760611 | -2,43826703 | 0,02813319 | 0,14035994 |
| 9372_at | ZFYVE9 | 0,2220934 | 5,8905012 | 2,43804246 | 0,02814548 | 0,14038485 |
| 64282_at | PAPD5 | -0,22048333 | 7,99482458 | -2,43783815 | 0,02815665 | 0,14040425 |
| 374454_at | KRT77 | -0,32868841 | 4,60611482 | -2,43633303 | 0,02823913 | 0,14073742 |
| 54033_at | RBM11 | 0,44817247 | 4,49730697 | 2,43625632 | 0,02824334 | 0,14073742 |
| 100506473_a | LOC10050647 | -0,26105652 | 5,43388071 | -2,43609381 | 0,02825226 | 0,14073742 |
| 6526_at | SLC5A3 | -0,21585569 | 6,73075053 | -2,4359771 | 0,02825867 | 0,14073742 |
| 80206_at | FHOD3 | -0,40953376 | 6,84367858 | -2,43595291 | 0,02826 | 0,14073742 |
| 7156_at | TOP3A | -0,34308294 | 6,55786262 | -2,43559421 | 0,0282797 | 0,14079915 |
| 60686_at | C14orf93 | 0,19643017 | 5,30631018 | 2,43512961 | 0,02830524 | 0,14087614 |
| 9119_at | KRT75 | -0,35219122 | 4,90883736 | -2,43504687 | 0,02830979 | 0,14087614 |
| 6674_at | SPAG1 | -0,33288741 | 7,62937345 | -2,43402966 | 0,0283658 | 0,14111838 |
| 6604_at | SMARCD3 | 0,39560126 | 6,53223752 | 2,4337798 | 0,02837957 | 0,14115044 |
| 57097_at | PARP11 | 0,39634599 | 4,18356967 | 2,43327613 | 0,02840735 | 0,14125215 |
| 7003_at | TEAD1 | 0,15922552 | 8,0956876 | 2,43232316 | 0,02845999 | 0,14144821 |
| 123811_at | FOPNL | -0,27370235 | 8,73853174 | -2,43229639 | 0,02846147 | 0,14144821 |
| 140856_at | SCP2D1 | -0,2503385 | 3,90117599 | -2,43188979 | 0,02848396 | 0,14152346 |
| 553_at | AVPR1B | -0,29485324 | 7,06632769 | -2,43114214 | 0,02852535 | 0,14166657 |
| 51334_at | PRR16 | 0,37978403 | 3,65387885 | 2,43099851 | 0,02853331 | 0,14166657 |
| 5581_at | PRKCE | 0,1592844 | 5,99834093 | 2,43094349 | 0,02853636 | 0,14166657 |
| 157983_at | C9orf66 | 0,18353577 | 5,91238528 | 2,43083851 | 0,02854218 | 0,14166657 |
| 152110_at | NEK10 | 0,2686922 | 3,87285818 | 2,43060145 | 0,02855532 | 0,14168693 |
| 101928844_a | LOC10192884 | -0,18054742 | 5,95325875 | -2,43049923 | 0,02856099 | 0,14168693 |
| 5424_at | POLD1 | -0,55146076 | 7,71627596 | -2,42935141 | 0,02862473 | 0,14195077 |
| 9870_at | AREL1 | 0,24114032 | 8,97986171 | 2,42927611 | 0,02862891 | 0,14195077 |
| 7543_at | ZFX | 0,21254318 | 6,26470864 | 2,42896125 | 0,02864642 | 0,14200104 |
| 140456_at | ASB11 | -0,20631505 | 3,43356901 | -2,42851204 | 0,02867142 | 0,14208839 |
| 100505741_a | SPATA1 | 0,21045679 | 2,90896929 | 2,4281296 | 0,02869272 | 0,14215737 |
| 116225_at | ZMYND19 | -0,5657276 | 9,09012085 | -2,42798882 | 0,02870056 | 0,14215968 |
| 6624_at | FSCN1 | -0,82810376 | 9,43335286 | -2,42735111 | 0,02873612 | 0,14229921 |
| 9907_at | AP5Z1 | 0,18180045 | 6,72642117 | 2,42693244 | 0,02875949 | 0,14234874 |
| 9114_at | ATP6V0D1 | 0,23871434 | 10,7053437 | 2,42690714 | 0,0287609 | 0,14234874 |
| 10938_at | EHD1 | 0,19028362 | 8,00250213 | 2,42657142 | 0,02877966 | 0,14240496 |
| 6339_at | SCNN1D | 0,21789619 | 5,27358893 | 2,42612758 | 0,02880446 | 0,14249111 |

| | | | | | | |
|---|---|---|---|---|---|---|
| 1857_at | DVL3 | 0,18791341 | 8,9791766 | 2,42577418 | 0,02882423 | 0,14252523 |
| 59271_at | EVA1C | 0,30134502 | 5,86945034 | 2,42573977 | 0,02882616 | 0,14252523 |
| 55101_at | ATP5SL | -0,21664872 | 8,86209835 | -2,4250926 | 0,02886239 | 0,14266777 |
| 7942_at | TFEB | -0,27348682 | 6,87819589 | -2,42489348 | 0,02887355 | 0,14267275 |
| 6605_at | SMARCE1 | 0,21211075 | 10,0746985 | 2,42481028 | 0,02887822 | 0,14267275 |
| 57222_at | ERGIC1 | -0,1663936 | 9,13418578 | -2,42432775 | 0,02890528 | 0,14276224 |
| 11112_at | HIBADH | 0,20296477 | 8,11706019 | 2,42422303 | 0,02891115 | 0,14276224 |
| 100506233_a | RAB30-AS1 | -0,33377302 | 7,9243233 | -2,42331462 | 0,02896217 | 0,14295967 |
| 3163_at | HMOX2 | -0,22863812 | 7,98192296 | -2,42291068 | 0,02898488 | 0,14295967 |
| 154313_at | C6orf165 | 0,38604588 | 5,23642586 | 2,42279798 | 0,02899122 | 0,14295967 |
| 90780_at | PYGO2 | 0,17782247 | 7,40743712 | 2,42255882 | 0,02900468 | 0,14295967 |
| 23536_at | ADAT1 | -0,27861957 | 7,9444157 | -2,42253036 | 0,02900629 | 0,14295967 |
| 533_at | ATP6V0B | -0,15822856 | 9,63539221 | -2,42250656 | 0,02900762 | 0,14295967 |
| 7479_at | WNT8B | -0,17200749 | 5,37671798 | -2,42249945 | 0,02900803 | 0,14295967 |
| 28956_at | LAMTOR2 | 0,14550189 | 8,73992859 | 2,42245542 | 0,0290105 | 0,14295967 |
| 26133_at | TRPC4AP | 0,24308564 | 8,34040035 | 2,4220066 | 0,02903578 | 0,14304764 |
| 51504_at | TRMT112 | -0,24784636 | 11,0232231 | -2,42187006 | 0,02904348 | 0,14304896 |
| 11001_at | SLC27A2 | -0,38342552 | 7,30790962 | -2,42171523 | 0,0290522 | 0,14305537 |
| 54980_at | C2orf42 | 0,25357558 | 6,84229116 | 2,41963468 | 0,02916972 | 0,14359733 |
| 140846_at | FAM83C-AS1 | -0,1662295 | 4,40517409 | -2,41941257 | 0,02918229 | 0,14362252 |
| 54842_at | MFSD6 | 0,40065957 | 8,46056643 | 2,41909831 | 0,02920009 | 0,1436734 |
| 55148_at | UBR7 | -0,34848921 | 8,82405599 | -2,41827118 | 0,02924698 | 0,14386737 |
| 22885_at | ABLIM3 | 0,30838981 | 7,31869768 | 2,41796529 | 0,02926434 | 0,14391601 |
| 8021_at | NUP214 | -0,15110386 | 7,76555351 | -2,41653044 | 0,0293459 | 0,14428027 |
| 122809_at | SOCS4 | -0,26007938 | 7,10478397 | -2,41629649 | 0,02935922 | 0,14430892 |
| 494514_at | TYMSOS | -0,37756861 | 5,45473736 | -2,41481017 | 0,02944397 | 0,14463003 |
| 79644_at | SRD5A3 | 0,47535137 | 7,21286553 | 2,41461592 | 0,02945506 | 0,14463003 |
| 6903_at | TBCC | 0,21526528 | 8,41281205 | 2,41457944 | 0,02945715 | 0,14463003 |
| 79143_at | MBOAT7 | -0,17408943 | 9,27108334 | -2,41453967 | 0,02945942 | 0,14463003 |
| 4069_at | LYZ | 0,21151013 | 3,5380188 | 2,41438442 | 0,02946829 | 0,14463003 |
| 84968_at | PNMA6A | -0,35978703 | 4,93840421 | -2,41427868 | 0,02947433 | 0,14463003 |
| 90586_at | AOC4P | -0,19010651 | 4,92814097 | -2,41423013 | 0,02947711 | 0,14463003 |
| 203413_at | CT83 | -0,17336558 | 3,34024144 | -2,41406337 | 0,02948664 | 0,14463996 |
| 65010_at | SLC26A6 | -0,28278454 | 6,2036987 | -2,41346684 | 0,02952077 | 0,14477049 |
| 55471_at | NDUFAF7 | -0,19827426 | 6,2845994 | -2,41211303 | 0,02959835 | 0,14511404 |
| 1132_at | CHRM4 | -0,28978526 | 5,2507878 | -2,41158866 | 0,02962846 | 0,14522467 |
| 273_at | AMPH | 1,21605841 | 5,80578174 | 2,41133427 | 0,02964307 | 0,14525935 |
| 90933_at | TRIM41 | 0,20237181 | 8,02425835 | 2,41063177 | 0,02968347 | 0,1454203 |
| 84973_at | SNHG7 | -0,41167869 | 6,6389948 | -2,41030325 | 0,02970237 | 0,14546338 |
| 84548_at | TMEM185A | 0,36364326 | 7,79004808 | 2,41021662 | 0,02970736 | 0,14546338 |
| 2793_at | GNGT2 | -0,20858029 | 4,3032931 | -2,4089127 | 0,02978254 | 0,14579441 |
| 64429_at | ZDHHC6 | -0,41613652 | 8,70277165 | -2,40846793 | 0,02980822 | 0,14588306 |
| 83446_at | CCDC70 | -0,25401793 | 4,10700388 | -2,40821077 | 0,02982308 | 0,14591871 |
| 150084_at | IGSF5 | 0,43169152 | 4,5351417 | 2,40807698 | 0,02983081 | 0,14591948 |
| 11142_at | PKIG | 0,62257375 | 6,30765279 | 2,40784575 | 0,02984418 | 0,14594782 |
| 154091_at | SLC2A12 | 1,41785937 | 7,36125918 | 2,40643844 | 0,02992568 | 0,14630922 |
| 51264_at | MRPL27 | -0,23932831 | 9,5790899 | -2,40609022 | 0,02994588 | 0,14635815 |
| 10730_at | YME1L1 | -0,31686998 | 10,1696336 | -2,40600394 | 0,02995088 | 0,14635815 |
| 3836_at | KPNA1 | -0,16544321 | 8,41004607 | -2,40577023 | 0,02996445 | 0,14636896 |
| 92960_at | PEX11G | 0,32182103 | 5,99319175 | 2,40570405 | 0,02996829 | 0,14636896 |

| | | | | | | |
|---|---|---|---|---|---|---|
| 5711_at | PSMD5 | -0,17873896 | 9,12450643 | -2,4052235 | 0,0299962 | 0,14643493 |
| 6860_at | SYT4 | -0,16625795 | 3,83098574 | -2,40520979 | 0,029997 | 0,14643493 |
| 26017_at | FAM32A | 0,14081594 | 8,82110348 | 2,40418336 | 0,03005671 | 0,14668923 |
| 26260_at | FBXO25 | 0,23740705 | 7,09464699 | 2,4035602 | 0,03009301 | 0,14680835 |
| 125228_at | FAM210A | -0,52990532 | 6,87142703 | -2,40350279 | 0,03009636 | 0,14680835 |
| 10287_at | RGS19 | -0,20895207 | 8,44604664 | -2,40289371 | 0,03013189 | 0,14690875 |
| 554206_at | LOC554206 | 0,27979342 | 4,13492543 | 2,40288843 | 0,0301322 | 0,14690875 |
| 66004_at | LYNX1 | -0,2405659 | 6,20051599 | -2,40242304 | 0,03015937 | 0,14700403 |
| 29925_at | GMPPB | -0,37276265 | 8,20868035 | -2,40208926 | 0,03017887 | 0,14706188 |
| 84909_at | C9orf3 | 0,14438018 | 3,91860827 | 2,40178644 | 0,03019658 | 0,14711094 |
| 283902_at | HCCAT5 | -0,21354559 | 5,1806992 | -2,40132872 | 0,03022336 | 0,14720418 |
| 23545_at | ATP6V0A2 | -0,29730945 | 6,95516432 | -2,40106033 | 0,03023907 | 0,14724346 |
| 55707_at | NECAP2 | 0,26186857 | 8,54875225 | 2,40092987 | 0,03024671 | 0,14724346 |
| 101927513_a | LOC10192751 | -0,23135877 | 4,24052211 | -2,40077972 | 0,03025551 | 0,14724579 |
| 311_at | ANXA11 | 0,22276083 | 9,20866061 | 2,40063059 | 0,03026425 | 0,14724579 |
| 51430_at | SUCO | 0,34411516 | 8,84121193 | 2,40053035 | 0,03027012 | 0,14724579 |
| 84932_at | RAB2B | 0,2216356 | 8,35038009 | 2,40016787 | 0,03029138 | 0,14730842 |
| 58492_at | ZNF77 | -0,18834128 | 6,29168216 | -2,40004999 | 0,03029829 | 0,14730842 |
| 84811_at | BUD13 | -0,30495083 | 7,23729692 | -2,39912141 | 0,03035282 | 0,14753259 |
| 100506012_a | PPP5D1 | 0,40819306 | 3,79739954 | 2,39900403 | 0,03035972 | 0,14753259 |
| 84920_at | ALG10 | -0,36551436 | 5,39353738 | -2,39810619 | 0,03041254 | 0,14775168 |
| 54629_at | FAM63B | 0,41969291 | 7,73941021 | 2,39797702 | 0,03042014 | 0,14775168 |
| 81831_at | NETO2 | -1,20635141 | 8,84645471 | -2,39745481 | 0,03045091 | 0,14784575 |
| 23011_at | RAB21 | -0,29109441 | 9,1004365 | -2,39738785 | 0,03045486 | 0,14784575 |
| 160851_at | DGKH | -0,3828085 | 5,99306088 | -2,39724349 | 0,03046337 | 0,14784981 |
| 129831_at | RBM45 | -0,24152874 | 7,40754378 | -2,39706547 | 0,03047387 | 0,14786352 |
| 317_at | APAF1 | 0,34305088 | 6,50757564 | 2,39678711 | 0,03049029 | 0,14790596 |
| 3909_at | LAMA3 | -0,19537179 | 5,9896659 | -2,39636124 | 0,03051544 | 0,14799067 |
| 4700_at | NDUFA6 | -0,22814492 | 9,50059142 | -2,39591 | 0,0305421 | 0,14803666 |
| 91272_at | BOD1 | -0,31931512 | 9,12730854 | -2,39589325 | 0,0305431 | 0,14803666 |
| 54850_at | FBXL12 | 0,13885656 | 8,48954484 | 2,39581066 | 0,03054798 | 0,14803666 |
| 388591_at | RNF207 | 0,81051345 | 7,45399733 | 2,39528125 | 0,0305793 | 0,14815115 |
| 2122_at | MECOM | 0,48607157 | 9,51740727 | 2,39512861 | 0,03058833 | 0,14815767 |
| 55661_at | DDX27 | -0,40316595 | 9,06061274 | -2,39454074 | 0,03062315 | 0,14825487 |
| 7551_at | ZNF3 | 0,14481954 | 5,99910574 | 2,39452994 | 0,03062379 | 0,14825487 |
| 22993_at | HMGXB3 | -0,16097178 | 8,08825216 | -2,39436247 | 0,03063372 | 0,14826566 |
| 26234_at | FBXL5 | 0,27945191 | 8,76453717 | 2,39338756 | 0,03069157 | 0,14850834 |
| 152518_at | NFXL1 | -0,35920252 | 8,03809666 | -2,39274692 | 0,03072964 | 0,14856883 |
| 10621_at | POLR3F | -0,37529329 | 7,26901254 | -2,3924929 | 0,03074475 | 0,14856883 |
| 11007_at | CCDC85B | -0,37467847 | 7,0531768 | -2,39244408 | 0,03074765 | 0,14856883 |
| 8688_at | KRT37 | -0,25874559 | 4,89062024 | -2,39238011 | 0,03075146 | 0,14856883 |
| 221443_at | OARD1 | 0,26633696 | 6,97670543 | 2,39234545 | 0,03075352 | 0,14856883 |
| 80853_at | KDM7A | 0,42429689 | 4,99502278 | 2,39229792 | 0,03075635 | 0,14856883 |
| 57674_at | RNF213 | 0,70956712 | 8,51058768 | 2,39215086 | 0,0307651 | 0,14856883 |
| 54806_at | AHI1 | -0,17327181 | 5,89176263 | -2,39213959 | 0,03076577 | 0,14856883 |
| 64983_at | MRPL32 | -0,34619381 | 9,15450838 | -2,39164982 | 0,03079494 | 0,14867241 |
| 57561_at | ARRDC3 | 0,92426625 | 8,5234491 | 2,39131651 | 0,0308148 | 0,14870064 |
| 56903_at | PAPOLB | -0,12822287 | 3,21399709 | -2,39129268 | 0,03081623 | 0,14870064 |
| 25939_at | SAMHD1 | 0,91448584 | 7,73065031 | 2,39095833 | 0,03083617 | 0,14875959 |
| 70_at | ACTC1 | -0,17347419 | 4,41439152 | -2,39014761 | 0,03088457 | 0,14895578 |

| | | | | | | |
|---|---|---|---|---|---|---|
| 65982_at | ZSCAN18 | 0,40171059 | 7,11690231 | 2,38973949 | 0,03090896 | 0,14903611 |
| 283174_at | MIR4697HG | -0,23676828 | 3,32923849 | -2,38953804 | 0,03092101 | 0,14905404 |
| 135458_at | HUS1B | -0,17278475 | 4,0686497 | -2,38941857 | 0,03092815 | 0,14905404 |
| 2873_at | GPS1 | -0,20056782 | 9,29620002 | -2,3892702 | 0,03093703 | 0,14905953 |
| 100129654_a | TCF24 | 0,22824587 | 3,88170299 | 2,38859873 | 0,03097723 | 0,14921592 |
| 399959_at | MIR100HG | -0,13839956 | 2,90556473 | -2,38831945 | 0,03099397 | 0,14924796 |
| 5188_at | GATB | -0,18508828 | 6,43637337 | -2,38822921 | 0,03099938 | 0,14924796 |
| 9779_at | TBC1D5 | 0,23804221 | 7,56820518 | 2,38680679 | 0,03108478 | 0,14962169 |
| 9402_at | GRAP2 | -0,17891011 | 4,57153668 | -2,38620406 | 0,03112103 | 0,14975877 |
| 1356_at | CP | 1,16187987 | 7,76960095 | 2,38577827 | 0,03114666 | 0,14984468 |
| 51751_at | HIGD1B | -0,17441329 | 4,65971526 | -2,38523882 | 0,03117916 | 0,14996361 |
| 146540_at | ZNF785 | 0,30407871 | 6,00034809 | 2,38468355 | 0,03121266 | 0,15004409 |
| 4335_at | MNT | 0,21313676 | 8,30857061 | 2,38462727 | 0,03121605 | 0,15004409 |
| 9424_at | KCNK6 | 0,54601134 | 7,34354125 | 2,38457405 | 0,03121926 | 0,15004409 |
| 26115_at | TANC2 | 0,45020426 | 7,51969231 | 2,38412755 | 0,03124623 | 0,15009649 |
| 100506299_a | ID2-AS1 | 0,32531542 | 4,41093274 | 2,38402669 | 0,03125232 | 0,15009649 |
| 10013_at | HDAC6 | 0,25859973 | 7,17396634 | 2,38400641 | 0,03125354 | 0,15009649 |
| 3953_at | LEPR | 0,24008206 | 4,07585831 | 2,38386159 | 0,0312623 | 0,15010109 |
| 56970_at | ATXN7L3 | 0,18767628 | 7,73144349 | 2,3827175 | 0,03133151 | 0,15039595 |
| 23306_at | NEMP1 | -0,62958451 | 7,1450504 | -2,38104359 | 0,03143305 | 0,15084574 |
| 81502_at | HM13 | -0,17983777 | 7,88060083 | -2,38008438 | 0,03149137 | 0,15108799 |
| 10459_at | MAD2L2 | -0,42269408 | 8,98920334 | -2,37945265 | 0,03152984 | 0,15123487 |
| 2979_at | GUCA1B | -0,19281764 | 5,31999694 | -2,3791192 | 0,03155016 | 0,15129468 |
| 9679_at | FAM53B | 0,18355103 | 7,73279731 | 2,37881993 | 0,03156841 | 0,15134452 |
| 715_at | C1R | 0,98856476 | 7,60175839 | 2,37828915 | 0,0316008 | 0,15142759 |
| 400793_at | C1orf226 | 0,23664187 | 5,36885919 | 2,37827838 | 0,03160146 | 0,15142759 |
| 145508_at | CEP128 | -0,45061853 | 5,40129742 | -2,3775925 | 0,03164336 | 0,15159068 |
| 51642_at | MRPL48 | -0,32725618 | 9,2198769 | -2,37719751 | 0,03166752 | 0,15166868 |
| 113201_at | CASC4 | 0,39061346 | 10,7432346 | 2,37676748 | 0,03169384 | 0,151757 |
| 63923_at | TNN | -0,26208405 | 4,39966837 | -2,3765152 | 0,03170929 | 0,15177002 |
| 90701_at | SEC11C | -0,29285505 | 7,60200755 | -2,37639702 | 0,03171653 | 0,15177002 |
| 58485_at | TRAPPC1 | 0,46650502 | 8,39529009 | 2,3763372 | 0,03172019 | 0,15177002 |
| 4716_at | NDUFB10 | -0,27575973 | 10,5496062 | -2,37605027 | 0,03173778 | 0,15181646 |
| 78988_at | MRPL57 | -0,32565982 | 8,58115188 | -2,3753144 | 0,03178293 | 0,15199466 |
| 6477_at | SIAH1 | 0,18292777 | 7,69569485 | 2,3748199 | 0,0318133 | 0,15210214 |
| 155400_at | NSUN5P1 | -0,3797331 | 7,46603359 | -2,37426249 | 0,03184756 | 0,15222819 |
| 9938_at | ARHGAP25 | -0,2974452 | 4,82240595 | -2,37397364 | 0,03186533 | 0,15225741 |
| 8492_at | PRSS12 | 0,43394658 | 5,80806836 | 2,37390623 | 0,03186948 | 0,15225741 |
| 57508_at | INTS2 | -0,16810594 | 7,40473392 | -2,37345754 | 0,03189711 | 0,15235161 |
| 434_at | ASIP | -0,19731989 | 4,78721471 | -2,37303049 | 0,03192343 | 0,15243951 |
| 284312_at | ZSCAN1 | -0,23939216 | 4,81219302 | -2,37277237 | 0,03193934 | 0,15244008 |
| 6301_at | SARS | -0,21119441 | 8,73455492 | -2,37277187 | 0,03193937 | 0,15244008 |
| 56265_at | CPXM1 | -0,22641735 | 4,97212301 | -2,37202109 | 0,03198571 | 0,15262342 |
| 29104_at | N6AMT1 | -0,39870909 | 6,7472363 | -2,37100639 | 0,03204844 | 0,15288486 |
| 51475_at | CABP2 | -0,1995297 | 5,97144486 | -2,36936618 | 0,03215008 | 0,15333175 |
| 26548_at | ITGB1BP2 | -0,4887913 | 4,69890807 | -2,36868991 | 0,03219207 | 0,15349404 |
| 90637_at | ZFAND2A | -0,22132135 | 8,1215124 | -2,36834118 | 0,03221375 | 0,15355712 |
| 56904_at | SH3GLB2 | 0,33166414 | 7,71250664 | 2,36817821 | 0,03222388 | 0,15355712 |
| 1358_at | CPA2 | -0,19230086 | 4,30684488 | -2,36809247 | 0,03222922 | 0,15355712 |
| 2633_at | GBP1 | 1,04755627 | 8,93201782 | 2,36773239 | 0,03225162 | 0,1535808 |

| | | | | | |
|---|---|---|---|---|---|
| 2079_at | ERH | -0,17244135 | 12,1512004 | -2,36765669 | 0,03225634 | 0,1535808 |
| 401431_at | ATP6V0E2-AS: | -0,31337781 | 5,80394747 | -2,36758497 | 0,0322608 | 0,1535808 |
| 51081_at | MRPS7 | -0,31303701 | 9,43726609 | -2,3675002 | 0,03226608 | 0,1535808 |
| 399818_at | METTL10 | -0,46401675 | 6,18322361 | -2,36659139 | 0,03232272 | 0,1538124 |
| 4597_at | MVD | -0,31763234 | 7,61384099 | -2,36630233 | 0,03234076 | 0,15386021 |
| 7913_at | DEK | -0,48469345 | 11,0702978 | -2,36474074 | 0,03243835 | 0,15427104 |
| 94274_at | PPP1R14A | 0,20073034 | 4,25164373 | 2,36454458 | 0,03245063 | 0,15427104 |
| 22839_at | DLGAP4 | 0,26880168 | 7,37374472 | 2,3645365 | 0,03245114 | 0,15427104 |
| 374900_at | ZNF568 | 0,15111006 | 3,65769758 | 2,36401588 | 0,03248375 | 0,15438797 |
| 23729_at | SHPK | -0,28873731 | 5,68342814 | -2,36350706 | 0,03251565 | 0,15450148 |
| 79102_at | RNF26 | -0,41597431 | 8,47624645 | -2,36241995 | 0,03258391 | 0,15478765 |
| 3038_at | HAS3 | -0,75846591 | 7,00418511 | -2,36226193 | 0,03259385 | 0,15479666 |
| 790955_at | UQCC3 | -0,46733454 | 7,21019747 | -2,36185735 | 0,03261929 | 0,15486521 |
| 7187_at | TRAF3 | -0,22422817 | 7,29427804 | -2,36177682 | 0,03262436 | 0,15486521 |
| 143689_at | PIWIL4 | 0,28050891 | 3,51543624 | 2,36162603 | 0,03263385 | 0,15487209 |
| 137886_at | UBXN2B | -0,2441802 | 6,81819985 | -2,36140009 | 0,03264807 | 0,15489041 |
| 11066_at | SNRNP35 | 0,12850343 | 7,00553902 | 2,36130932 | 0,03265379 | 0,15489041 |
| 139212_at | PIH1D3 | -0,24369707 | 3,84789292 | -2,36035716 | 0,03271381 | 0,1551369 |
| 317781_at | DDX51 | -0,34619523 | 6,91783508 | -2,36006592 | 0,03273219 | 0,15518585 |
| 9014_at | TAF1B | -0,35104876 | 6,9766627 | -2,35972666 | 0,03275361 | 0,15521706 |
| 7073_at | TIAL1 | -0,21667955 | 7,57477392 | -2,35970646 | 0,03275489 | 0,15521706 |
| 9049_at | AIP | 0,16376436 | 8,58015334 | 2,3592576 | 0,03278325 | 0,15530952 |
| 2874_at | GPS2 | -0,14890157 | 8,64407928 | -2,35914258 | 0,03279052 | 0,15530952 |
| 286828_at | CSN1S2AP | -0,24856763 | 4,23525445 | -2,35890353 | 0,03280564 | 0,15532346 |
| 80223_at | RAB11FIP1 | 0,4670271 | 8,18535224 | 2,35884105 | 0,03280959 | 0,15532346 |
| 6502_at | SKP2 | -0,41649643 | 8,63168051 | -2,35866605 | 0,03282066 | 0,15533771 |
| 1954_at | MEGF8 | 0,24158317 | 6,86414371 | 2,35806168 | 0,03285893 | 0,15544588 |
| 1613_at | DAPK3 | 0,25661173 | 7,77100406 | 2,35779685 | 0,03287572 | 0,15544588 |
| 29086_at | BABAM1 | -0,16791436 | 9,17676007 | -2,35777508 | 0,0328771 | 0,15544588 |
| 84445_at | LZTS2 | 0,32616275 | 7,64621568 | 2,35774657 | 0,0328789 | 0,15544588 |
| 26254_at | OPTC | -0,264503 | 4,69526604 | -2,35766825 | 0,03288387 | 0,15544588 |
| 10450_at | PPIE | -0,15412785 | 7,58619111 | -2,35754024 | 0,03289198 | 0,1554461 |
| 23517_at | SKIV2L2 | -0,49418171 | 8,63278621 | -2,35718051 | 0,0329148 | 0,15551579 |
| 10040_at | TOM1L1 | 0,25163726 | 8,32994374 | 2,35694926 | 0,03292948 | 0,15554698 |
| 9667_at | SAFB2 | -0,14830843 | 8,71222773 | -2,35673285 | 0,03294322 | 0,15557374 |
| 23635_at | SSBP2 | 0,6769244 | 7,67017572 | 2,35607291 | 0,03298516 | 0,15569028 |
| 79807_at | GSTCD | -0,62417498 | 6,26408287 | -2,35606891 | 0,03298541 | 0,15569028 |
| 55156_at | ARMC1 | -0,15365093 | 9,11103846 | -2,35596302 | 0,03299215 | 0,15569028 |
| 4606_at | MYBPC2 | -0,23900229 | 4,2571982 | -2,35565482 | 0,03301175 | 0,15574465 |
| 1580_at | CYP4B1 | 1,23411822 | 7,76620837 | 2,35506238 | 0,03304947 | 0,15588442 |
| 728489_at | DNLZ | -0,51002847 | 8,22520099 | -2,35461992 | 0,03307767 | 0,15597922 |
| 134285_at | TMEM171 | -0,25263652 | 4,06930549 | -2,35412217 | 0,03310942 | 0,15609072 |
| 58513_at | EPS15L1 | -0,17847761 | 6,96405481 | -2,35293266 | 0,0331854 | 0,15641066 |
| 8925_at | HERC1 | 0,15815977 | 6,31353778 | 2,35174889 | 0,03326118 | 0,15670444 |
| 9971_at | NR1H4 | 0,33975445 | 3,47384052 | 2,35163938 | 0,0332682 | 0,15670444 |
| 25917_at | THUMPD3 | -0,29545613 | 8,104404 | -2,35157801 | 0,03327214 | 0,15670444 |
| 4494_at | MT1F | 0,5798725 | 8,8685054 | 2,35100678 | 0,03330878 | 0,15683866 |
| 10927_at | SPIN1 | 0,17112916 | 9,79783733 | 2,3507991 | 0,03332211 | 0,15686308 |
| 59082_at | CARD18 | -0,20717014 | 2,99596591 | -2,35038209 | 0,03334889 | 0,1569508 |
| 222229_at | LRWD1 | -0,24231684 | 7,5154852 | -2,35006207 | 0,03336945 | 0,15700923 |

| | | | | | | |
|---|---|---|---|---|---|---|
| 5338_at | PLD2 | 0,23009251 | 6,99381909 | 2,34945776 | 0,03340832 | 0,15715373 |
| 255919_at | CNEP1R1 | 0,25830213 | 8,03830149 | 2,34911524 | 0,03343037 | 0,15721906 |
| 7108_at | TM7SF2 | 0,57946049 | 8,88331642 | 2,34789415 | 0,03350909 | 0,1575508 |
| 8225_at | GTPBP6 | -0,32636771 | 7,76010452 | -2,34760759 | 0,03352759 | 0,15759931 |
| 567_at | B2M | 0,30858032 | 10,8984619 | 2,34633694 | 0,03360973 | 0,15792168 |
| 374491_at | TPTE2P6 | 0,73116148 | 3,27929294 | 2,34623584 | 0,03361628 | 0,15792168 |
| 112476_at | PRRT2 | 0,27869033 | 5,09498816 | 2,34603282 | 0,03362942 | 0,15792168 |
| 93164_at | HTR7P1 | 0,24371161 | 5,2372045 | 2,34598678 | 0,0336324 | 0,15792168 |
| 51277_at | DNAJC27 | 0,21202631 | 5,22284344 | 2,3459133 | 0,03363716 | 0,15792168 |
| 57691_at | KIAA1586 | -0,22108749 | 6,10456818 | -2,34476411 | 0,03371167 | 0,15820431 |
| 22909_at | FAN1 | -0,20538436 | 6,50495799 | -2,34473154 | 0,03371379 | 0,15820431 |
| 128153_at | SPATA17 | 0,68396348 | 4,83389402 | 2,34450879 | 0,03372825 | 0,15823363 |
| 54920_at | DUS2 | -0,22005202 | 7,6618155 | -2,34388163 | 0,033769 | 0,15838623 |
| 2948_at | GSTM4 | 0,58531512 | 7,81033853 | 2,3436408 | 0,03378466 | 0,1584211 |
| 2287_at | FKBP3 | -0,17874117 | 9,00826501 | -2,34333225 | 0,03380473 | 0,15847665 |
| 65250_at | C5orf42 | 0,2523437 | 4,04978502 | 2,34308517 | 0,03382081 | 0,15851347 |
| 6890_at | TAP1 | 0,69939928 | 9,30276563 | 2,34293788 | 0,03383041 | 0,15851985 |
| 1203_at | CLN5 | 0,27346203 | 7,52135084 | 2,34279588 | 0,03383965 | 0,15852463 |
| 25946_at | ZNF385A | 0,28243857 | 7,99384434 | 2,34231061 | 0,03387128 | 0,15862109 |
| 9781_at | RNF144A | 0,19626227 | 5,39802198 | 2,34208041 | 0,03388629 | 0,15862109 |
| 92715_at | DPH7 | -0,21840312 | 7,39747833 | -2,34186484 | 0,03390035 | 0,15862109 |
| 23162_at | MAPK8IP3 | 0,16991629 | 6,31894907 | 2,34182954 | 0,03390266 | 0,15862109 |
| 124583_at | CANT1 | 0,13236772 | 8,84695699 | 2,34171627 | 0,03391005 | 0,15862109 |
| 100505616_a | SNHG21 | -0,68380138 | 6,47737667 | -2,34166952 | 0,0339131 | 0,15862109 |
| 9140_at | ATG12 | 0,18123907 | 7,71151701 | 2,34142134 | 0,0339293 | 0,15862109 |
| 80078_at | LCAL1 | -0,17442913 | 4,13291041 | -2,34134402 | 0,03393435 | 0,15862109 |
| 85366_at | MYLK2 | -0,24069597 | 5,73134345 | -2,34134395 | 0,03393436 | 0,15862109 |
| 27201_at | GPR78 | 0,22545495 | 6,1619857 | 2,34080515 | 0,03396956 | 0,15874714 |
| 84075_at | FSCB | -0,18155697 | 2,88570534 | -2,3406734 | 0,03397818 | 0,15874888 |
| 55659_at | ZNF416 | 0,25955172 | 6,27928245 | 2,34029636 | 0,03400284 | 0,15875362 |
| 6296_at | ACSM3 | 0,27276982 | 3,92893921 | 2,34028182 | 0,03400379 | 0,15875362 |
| 7844_at | RNF103 | 0,33314382 | 7,71096293 | 2,34027995 | 0,03400392 | 0,15875362 |
| 55178_at | RNMTL1 | -0,2731396 | 6,95260987 | -2,33981384 | 0,03403443 | 0,15885757 |
| 63917_at | GALNT11 | 0,24773867 | 9,01503203 | 2,33919589 | 0,03407492 | 0,15898699 |
| 92675_at | DTD1 | -0,19539091 | 9,13521686 | -2,33913887 | 0,03407866 | 0,15898699 |
| 25983_at | NGDN | -0,26187861 | 8,37816855 | -2,33864411 | 0,03411112 | 0,15909988 |
| 4599_at | MX1 | 1,19552485 | 7,31996514 | 2,33813556 | 0,03414451 | 0,15921708 |
| 1795_at | DOCK3 | 0,35366237 | 4,3794652 | 2,33773952 | 0,03417054 | 0,15927577 |
| 1583_at | CYP11A1 | -0,38579226 | 5,03636304 | -2,33769241 | 0,03417364 | 0,15927577 |
| 873_at | CBR1 | -0,48254379 | 8,50296662 | -2,33748284 | 0,03418742 | 0,15930145 |
| 6747_at | SSR3 | -0,26544424 | 8,87868068 | -2,33694202 | 0,034223 | 0,15941645 |
| 171024_at | SYNPO2 | 0,38027019 | 4,600956 | 2,33685633 | 0,03422865 | 0,15941645 |
| 7764_at | ZNF217 | 0,31585655 | 9,48946069 | 2,33663066 | 0,03424351 | 0,15944711 |
| 199704_at | ZNF585A | 0,35341101 | 6,30813072 | 2,33624577 | 0,03426887 | 0,15952665 |
| 55823_at | VPS11 | 0,19812964 | 7,05696498 | 2,33516925 | 0,0343399 | 0,15973147 |
| 124491_at | TMEM170A | 0,24943855 | 8,17785076 | 2,33515493 | 0,03434085 | 0,15973147 |
| 8671_at | SLC4A4 | 0,32411093 | 4,14052298 | 2,33511202 | 0,03434368 | 0,15973147 |
| 140831_at | ZSWIM3 | 0,33790922 | 6,5017123 | 2,33507635 | 0,03434604 | 0,15973147 |
| 2677_at | GGCX | -0,17032644 | 8,37546621 | -2,33434558 | 0,03439435 | 0,15991752 |
| 114821_at | ZBED9 | 0,24570486 | 6,18313058 | 2,33416598 | 0,03440623 | 0,15993416 |

| | | | | | | |
|---|---|---|---|---|---|---|
| 59335_at | PRDM12 | -0,2156051 | 6,43347767 | -2,33399861 | 0,0344173 | 0,15994705 |
| 10524_at | KAT5 | 0,15707023 | 8,65352303 | 2,33330735 | 0,03446309 | 0,16012118 |
| 195828_at | ZNF367 | -0,89725651 | 8,13871821 | -2,33274279 | 0,03450052 | 0,16025645 |
| 8729_at | GBF1 | 0,15627513 | 7,6325642 | 2,33179928 | 0,03456316 | 0,16050874 |
| 3933_at | LCN1 | -0,24422085 | 4,50589153 | -2,3313404 | 0,03459367 | 0,16061168 |
| 553115_at | PEF1 | -0,15819639 | 8,19835894 | -2,33077915 | 0,03463102 | 0,16074633 |
| 283578_at | TMED8 | -0,32613318 | 7,9823538 | -2,33060549 | 0,03464258 | 0,16076127 |
| 274_at | BIN1 | -0,20598554 | 6,41472356 | -2,33044352 | 0,03465337 | 0,1607726 |
| 221927_at | BRAT1 | -0,18560944 | 7,52220086 | -2,33005902 | 0,03467899 | 0,16085274 |
| 3364_at | HUS1 | -0,17996008 | 7,49434587 | -2,32970593 | 0,03470254 | 0,1609145 |
| 79856_at | SNX22 | -0,18327123 | 5,18307586 | -2,32960883 | 0,03470901 | 0,1609145 |
| 387921_at | NHLRC3 | 0,40025095 | 7,29772244 | 2,32935836 | 0,03472573 | 0,16095325 |
| 9641_at | IKBKE | 0,28018536 | 7,03935473 | 2,32817128 | 0,03480505 | 0,16128209 |
| 56951_at | C5orf15 | 0,33482953 | 10,1747819 | 2,32747086 | 0,03485193 | 0,16141139 |
| 57703_at | CWC22 | -0,39510839 | 7,43232091 | -2,32735728 | 0,03485954 | 0,16141139 |
| 23196_at | FAM120A | -0,21418294 | 7,21303292 | -2,32729406 | 0,03486377 | 0,16141139 |
| 28976_at | ACAD9 | -0,31377494 | 8,89426455 | -2,32725384 | 0,03486647 | 0,16141139 |
| 6416_at | MAP2K4 | 0,21150993 | 7,74272863 | 2,32697924 | 0,03488487 | 0,16145778 |
| 9_at | NAT1 | 0,36886349 | 6,68287915 | 2,32630349 | 0,0349302 | 0,16158471 |
| 4678_at | NASP | -0,50706432 | 7,98750655 | -2,32622625 | 0,03493538 | 0,16158471 |
| 51106_at | TFB1M | -0,15879667 | 6,96469681 | -2,32603452 | 0,03494826 | 0,16158471 |
| 51742_at | ARID4B | 0,34394972 | 7,5653104 | 2,32586761 | 0,03495947 | 0,16158471 |
| 284889_at | MIF-AS1 | -0,37306705 | 6,3334123 | -2,32563604 | 0,03497502 | 0,16158471 |
| 81554_at | WBSCR16 | -0,19570019 | 5,71645615 | -2,3255135 | 0,03498326 | 0,16158471 |
| 80125_at | CCDC33 | -0,30535822 | 4,89641381 | -2,32550264 | 0,03498399 | 0,16158471 |
| 8824_at | CES2 | 0,15558825 | 8,55659521 | 2,32549143 | 0,03498474 | 0,16158471 |
| 7979_at | SHFM1 | -0,41311071 | 10,0236324 | -2,32544608 | 0,03498779 | 0,16158471 |
| 8788_at | DLK1 | -0,22726459 | 4,69754879 | -2,32274816 | 0,0351696 | 0,16238541 |
| 7846_at | TUBA1A | 0,76380389 | 7,38519888 | 2,32247641 | 0,03518796 | 0,16243125 |
| 1968_at | EIF2S3 | -0,17569701 | 10,6534406 | -2,32104906 | 0,03528455 | 0,16283811 |
| 23507_at | LRRC8B | -0,3097298 | 5,86546677 | -2,32058261 | 0,03531617 | 0,162945 |
| 340357_at | LOC340357 | -0,26556054 | 4,29329201 | -2,32025698 | 0,03533826 | 0,16300787 |
| 5901_at | RAN | -0,57162032 | 9,74760022 | -2,31950939 | 0,03538902 | 0,16320295 |
| 2197_at | FAU | -0,21854905 | 11,8154822 | -2,31932099 | 0,03540183 | 0,16322292 |
| 1545_at | CYP1B1 | 0,90985756 | 5,44325471 | 2,31871756 | 0,03544286 | 0,16337303 |
| 283011_at | FLJ37201 | -0,13109809 | 3,72751328 | -2,3182367 | 0,0354756 | 0,16345336 |
| 51016_at | EMC9 | -0,31669663 | 8,2546872 | -2,31821228 | 0,03547726 | 0,16345336 |
| 2543_at | GAGE1 | 0,20461153 | 3,49177462 | 2,3172953 | 0,03553977 | 0,16370218 |
| 84266_at | ALKBH7 | 0,2361151 | 6,93092053 | 2,31563383 | 0,03565329 | 0,16418581 |
| 115294_at | PCMTD1 | 0,92194252 | 7,58101269 | 2,31533478 | 0,03567375 | 0,1642408 |
| 6387_at | CXCL12 | -0,27634071 | 4,35930815 | -2,3150567 | 0,0356928 | 0,16428921 |
| 114769_at | CARD16 | 1,02324229 | 6,02093332 | 2,31445488 | 0,03573404 | 0,16443977 |
| 57095_at | PITHD1 | -0,2011193 | 8,01813965 | -2,31406184 | 0,035761 | 0,16452453 |
| 152485_at | ZNF827 | 0,93341894 | 7,49622321 | 2,31331345 | 0,03581239 | 0,16472162 |
| 11244_at | ZHX1 | 0,27838817 | 9,72450901 | 2,31270944 | 0,03585392 | 0,16485457 |
| 90678_at | LRSAM1 | 0,21894684 | 5,97317617 | 2,3126441 | 0,03585842 | 0,16485457 |
| 113_at | ADCY7 | -0,16008146 | 6,62621447 | -2,31108264 | 0,03596599 | 0,16530969 |
| 10333_at | TLR6 | -0,28290001 | 4,73696395 | -2,31039266 | 0,03601362 | 0,16548913 |
| 729220_at | FLJ45513 | -0,23793598 | 4,86125901 | -2,31001344 | 0,03603983 | 0,16557005 |
| 2530_at | FUT8 | 0,29153423 | 9,33054184 | 2,30982448 | 0,03605289 | 0,16559057 |

| | | | | | | |
|---|---|---|---|---|---|---|
| 5017_at | OVOL1 | -0,28400555 | 7,76716806 | -2,30965071 | 0,03606491 | 0,16560628 |
| 64108_at | RTP4 | 1,88697016 | 7,36747548 | 2,3091745 | 0,03609786 | 0,16569702 |
| 79753_at | SNIP1 | 0,16095337 | 6,79427483 | 2,30911657 | 0,03610187 | 0,16569702 |
| 84057_at | MND1 | -1,30158025 | 6,15858809 | -2,30886669 | 0,03611918 | 0,16573583 |
| 3976_at | LIF | -0,62763834 | 5,88939263 | -2,30871869 | 0,03612943 | 0,16573583 |
| 4899_at | NRF1 | -0,20537072 | 6,54003387 | -2,30862187 | 0,03613614 | 0,16573583 |
| 4175_at | MCM6 | -0,84734538 | 7,29993534 | -2,30731737 | 0,03622665 | 0,16611138 |
| 307_at | ANXA4 | 0,46156225 | 9,9083738 | 2,30614057 | 0,03630848 | 0,16644698 |
| 10210_at | TOPORS | 0,23114282 | 8,22842961 | 2,30542986 | 0,03635798 | 0,16663426 |
| 286436_at | H2BFM | -0,16744879 | 4,04337891 | -2,30484869 | 0,03639851 | 0,16678033 |
| 6382_at | SDC1 | -0,18388471 | 10,5553566 | -2,30332514 | 0,03650496 | 0,16719032 |
| 7520_at | XRCC5 | -0,19614287 | 11,097201 | -2,30331959 | 0,03650535 | 0,16719032 |
| 376497_at | SLC27A1 | 0,36076438 | 7,03232901 | 2,30272643 | 0,03654687 | 0,1673407 |
| 29937_at | NENF | -0,19305475 | 8,70256126 | -2,30228542 | 0,03657777 | 0,16744238 |
| 91543_at | RSAD2 | 1,94716165 | 5,69060455 | 2,3011077 | 0,03666041 | 0,16778082 |
| 63971_at | KIF13A | 0,3834025 | 7,12832422 | 2,3005056 | 0,03670273 | 0,16793459 |
| 1181_at | CLCN2 | -0,28074155 | 6,93261556 | -2,29998021 | 0,03673969 | 0,16806379 |
| 7561_at | ZNF14 | 0,25211214 | 6,1412456 | 2,29975764 | 0,03675536 | 0,16809555 |
| 1271_at | CNTFR | -0,24108774 | 6,20231177 | -2,29948541 | 0,03677453 | 0,168143 |
| 403150_at | FLJ31356 | 0,22788822 | 3,81650052 | 2,29936252 | 0,03678319 | 0,168143 |
| 56061_at | UBFD1 | -0,26670637 | 8,03781726 | -2,29903255 | 0,03680645 | 0,16818178 |
| 55753_at | OGDHL | -0,25936414 | 4,75755158 | -2,29899447 | 0,03680913 | 0,16818178 |
| 2734_at | GLG1 | 0,13443477 | 9,08118564 | 2,2988459 | 0,03681961 | 0,16818975 |
| 23480_at | SEC61G | -0,26848822 | 9,92820816 | -2,29849493 | 0,03684437 | 0,16826296 |
| 1978_at | EIF4EBP1 | -0,8728719 | 8,76326686 | -2,29812119 | 0,03687076 | 0,16834355 |
| 57464_at | STRIP2 | -0,37545383 | 5,26645495 | -2,2978743 | 0,0368882 | 0,16838327 |
| 79682_at | CENPU | -0,82381055 | 7,55370742 | -2,29726176 | 0,0369315 | 0,16850748 |
| 9057_at | SLC7A6 | 0,21477654 | 7,92325926 | 2,29714339 | 0,03693988 | 0,16850748 |
| 6256_at | RXRA | 0,25718764 | 9,02924464 | 2,29711825 | 0,03694165 | 0,16850748 |
| 3837_at | KPNB1 | -0,46948109 | 10,9313431 | -2,29688078 | 0,03695846 | 0,16851084 |
| 54858_at | PGPEP1 | 0,28842425 | 6,16495458 | 2,29686061 | 0,03695989 | 0,16851084 |
| 4795_at | NFKBIL1 | 0,12061974 | 7,08663726 | 2,29672294 | 0,03696963 | 0,16851539 |
| 10811_at | NOXA1 | 0,20157031 | 6,75384087 | 2,29547647 | 0,03705798 | 0,16887816 |
| 115426_at | UHRF2 | -0,18586949 | 8,78467172 | -2,29531841 | 0,0370692 | 0,16888932 |
| 5684_at | PSMA3 | -0,37698348 | 10,7251037 | -2,29419248 | 0,0371492 | 0,16921378 |
| 5358_at | PLS3 | 0,23338483 | 11,0687523 | 2,29333695 | 0,0372101 | 0,16944529 |
| 5509_at | PPP1R3D | 0,2988835 | 8,02118108 | 2,29311138 | 0,03722617 | 0,16944529 |
| 55454_at | CSGALNACT2 | -0,14749761 | 5,7473333 | -2,29310792 | 0,03722642 | 0,16944529 |
| 139538_at | VENTXP1 | -0,18469938 | 3,31171367 | -2,29290201 | 0,03724109 | 0,16945986 |
| 51000_at | SLC35B3 | 0,35517952 | 8,04603323 | 2,29269448 | 0,03725589 | 0,16945986 |
| 5664_at | PSEN2 | 0,34642919 | 6,53147872 | 2,29269282 | 0,03725601 | 0,16945986 |
| 10018_at | BCL2L11 | 0,4149452 | 7,08806078 | 2,29243363 | 0,0372745 | 0,16949188 |
| 101928909_a | LOC10192890 | -0,18187117 | 3,01991669 | -2,29234746 | 0,03728065 | 0,16949188 |
| 84740_at | AFAP1-AS1 | -0,25178875 | 3,93532294 | -2,29157754 | 0,03733563 | 0,16965699 |
| 7168_at | TPM1 | 0,27501438 | 8,78622948 | 2,29144813 | 0,03734488 | 0,16965699 |
| 4547_at | MTTP | 0,19171525 | 3,38341218 | 2,29137518 | 0,03735009 | 0,16965699 |
| 54808_at | DYM | -0,11757225 | 7,35915099 | -2,29134577 | 0,03735219 | 0,16965699 |
| 646450_at | ARIH2OS | 0,31874844 | 5,25088195 | 2,29109341 | 0,03737024 | 0,16966015 |
| 221504_at | ZBTB9 | -0,16468092 | 7,66581863 | -2,29108972 | 0,0373705 | 0,16966015 |
| 285512_at | FAM13A-AS1 | 0,25220421 | 5,6021752 | 2,2906556 | 0,03740156 | 0,16976116 |

| | | | | | | |
|---|---|---|---|---|---|---|
| 6169_at | RPL38 | -0,16401327 | 11,2267997 | -2,29028525 | 0,03742808 | 0,16983681 |
| 79780_at | CCDC82 | 0,21042892 | 6,32991866 | 2,29015723 | 0,03743725 | 0,16983681 |
| 6227_at | RPS21 | -0,2219073 | 12,1039539 | -2,28992768 | 0,0374537 | 0,16983681 |
| 285025_at | CCDC141 | 0,14761302 | 2,80892872 | 2,28982958 | 0,03746073 | 0,16983681 |
| 5340_at | PLG | -0,23074449 | 3,39514479 | -2,2898075 | 0,03746232 | 0,16983681 |
| 5602_at | MAPK10 | 0,20587351 | 5,55306472 | 2,28938733 | 0,03749245 | 0,16993343 |
| 6259_at | RYK | -0,3382779 | 9,42108164 | -2,2889023 | 0,03752726 | 0,1700512 |
| 51684_at | SUFU | 0,14576323 | 6,02597641 | 2,28862787 | 0,03754697 | 0,17009391 |
| 8532_at | CPZ | 0,4023423 | 7,98902716 | 2,28852523 | 0,03755435 | 0,17009391 |
| 2673_at | GFPT1 | 0,31090713 | 9,03154177 | 2,28808547 | 0,03758596 | 0,17019706 |
| 126282_at | TNFAIP8L1 | -0,47460088 | 6,79367205 | -2,28739863 | 0,03763538 | 0,17037454 |
| 27115_at | PDE7B | 0,25767874 | 5,12666549 | 2,28729499 | 0,03764284 | 0,17037454 |
| 216_at | ALDH1A1 | 0,38037607 | 4,44134503 | 2,28709692 | 0,03765711 | 0,17039908 |
| 285282_at | RABL3 | -0,2299239 | 7,64226874 | -2,28648521 | 0,0377012 | 0,17055852 |
| 2778_at | GNAS | 0,1181197 | 11,6634608 | 2,28602794 | 0,03773419 | 0,17066769 |
| 165679_at | SPTSSB | 0,6943365 | 6,32652224 | 2,28582294 | 0,03774899 | 0,17069455 |
| 9501_at | RPH3AL | -0,39390166 | 6,32568373 | -2,28504468 | 0,03780522 | 0,1709087 |
| 100507303_a | SNHG19 | -0,52075132 | 8,56893392 | -2,28481307 | 0,03782197 | 0,17091426 |
| 80709_at | AKNA | 0,20359557 | 6,48400338 | 2,28478229 | 0,0378242 | 0,17091426 |
| 8545_at | CGGBP1 | 0,18748884 | 8,80737395 | 2,28379862 | 0,03789542 | 0,17119593 |
| 27154_at | BRPF3 | 0,21458153 | 6,93332936 | 2,28355234 | 0,03791327 | 0,17123087 |
| 146862_at | UNC45B | -0,26308921 | 3,72379393 | -2,28343075 | 0,03792208 | 0,17123087 |
| 55128_at | TRIM68 | 0,31160692 | 7,36990656 | 2,2833241 | 0,03792982 | 0,17123087 |
| 11164_at | NUDT5 | -0,42976178 | 7,8193028 | -2,28246992 | 0,03799182 | 0,1714604 |
| 100533789_a | LRRC37A8P | -0,20749069 | 4,22821905 | -2,28237841 | 0,03799847 | 0,1714604 |
| 4898_at | NRD1 | -0,22633132 | 9,86208183 | -2,28148232 | 0,03806362 | 0,17171418 |
| 254042_at | METAP1D | -0,22645918 | 5,91307459 | -2,2803737 | 0,03814437 | 0,17203818 |
| 916_at | CD3E | -0,23894336 | 5,1034358 | -2,27980468 | 0,03818588 | 0,17218509 |
| 3728_at | JUP | 0,33437527 | 11,2297618 | 2,27904012 | 0,03824173 | 0,17239653 |
| 220064_at | ORAOV1 | -0,23830024 | 6,8247175 | -2,2785707 | 0,03827605 | 0,17251089 |
| 27336_at | HTATSF1 | -0,20974233 | 8,03626282 | -2,27814197 | 0,03830742 | 0,17261191 |
| 6744_at | SSFA2 | -0,26337777 | 8,45657225 | -2,27785669 | 0,03832831 | 0,17266564 |
| 3133_at | HLA-E | 0,42156493 | 9,58876646 | 2,27743193 | 0,03835944 | 0,17276545 |
| 100131897_a | FAM196B | -0,6191176 | 3,98874209 | -2,27626354 | 0,03844517 | 0,17310554 |
| 29915_at | HCFC2 | 0,28838366 | 6,42260534 | 2,27615802 | 0,03845292 | 0,17310554 |
| 6654_at | SOS1 | 0,33178813 | 7,69021432 | 2,27579539 | 0,03847957 | 0,17318504 |
| 9914_at | ATP2C2 | 0,37550738 | 6,97858333 | 2,27559655 | 0,03849419 | 0,17321037 |
| 254225_at | RNF169 | 0,12901024 | 8,11551001 | 2,27529347 | 0,03851648 | 0,17326805 |
| 10560_at | SLC19A2 | 0,4153117 | 8,1133985 | 2,27517777 | 0,038525 | 0,17326805 |
| 347252_at | IGFBPL1 | -0,38314073 | 5,57310566 | -2,27439348 | 0,03858276 | 0,17348733 |
| 1001_at | CDH3 | 0,20918417 | 10,3267184 | 2,27391979 | 0,03861769 | 0,17354939 |
| 1653_at | DDX1 | -0,34002631 | 10,3163513 | -2,27384657 | 0,03862309 | 0,17354939 |
| 54498_at | SMOX | -0,29263114 | 6,85633676 | -2,27383974 | 0,03862359 | 0,17354939 |
| 9364_at | RAB28 | -0,16606888 | 6,74568641 | -2,27329969 | 0,03866345 | 0,17368798 |
| 57589_at | RIC1 | 0,28511572 | 6,273705 | 2,27267855 | 0,03870934 | 0,1738536 |
| 5438_at | POLR2I | -0,3045434 | 9,05101357 | -2,27238333 | 0,03873118 | 0,1739111 |
| 51650_at | MRPS33 | -0,20866247 | 8,78988167 | -2,270578 | 0,03886492 | 0,17447099 |
| 8767_at | RIPK2 | -0,20878575 | 8,0440335 | -2,2702724 | 0,03888761 | 0,17453215 |
| 27350_at | APOBEC3C | 0,33431289 | 8,64839295 | 2,27014346 | 0,03889718 | 0,17453445 |
| 57221_at | ARFGEF3 | -0,19370261 | 6,0871268 | -2,26956191 | 0,03894039 | 0,17467654 |

| | | | | | | |
|---|---|---|---|---|---|---|
| 169981_at | SPIN3 | 0,14198963 | 5,94217148 | 2,26947326 | 0,03894698 | 0,17467654 |
| 100134368_a | LOC10013436 | -0,3089068 | 4,6368469 | -2,26921132 | 0,03896646 | 0,17472323 |
| 51520_at | LARS | -0,31741723 | 7,90456797 | -2,26907001 | 0,03897698 | 0,17472969 |
| 1537_at | CYC1 | -0,59945189 | 10,5459783 | -2,26817377 | 0,03904372 | 0,17498816 |
| 83690_at | CRISPLD1 | 0,22175191 | 3,47361485 | 2,26746156 | 0,03909683 | 0,17518543 |
| 50862_at | RNF141 | 0,27610267 | 8,31104257 | 2,26695648 | 0,03913453 | 0,17531361 |
| 11153_at | FICD | 0,16000513 | 6,58281929 | 2,26667181 | 0,0391558 | 0,17533063 |
| 79651_at | RHBDF2 | -0,22984671 | 7,1020019 | -2,26666197 | 0,03915654 | 0,17533063 |
| 116984_at | ARAP2 | 0,43695443 | 7,70126218 | 2,26625104 | 0,03918726 | 0,1754274 |
| 100505853_a | LOC10050585 | -0,20210565 | 4,83722914 | -2,26595302 | 0,03920955 | 0,17546988 |
| 100505761_a | RPARP-AS1 | -0,21545698 | 6,76650511 | -2,26588064 | 0,03921496 | 0,17546988 |
| 51091_at | SEPSECS | 0,38100516 | 5,51804772 | 2,2655465 | 0,03923998 | 0,17554103 |
| 120103_at | SLC36A4 | -0,35740681 | 5,59619536 | -2,26525755 | 0,03926162 | 0,17559706 |
| 2125_at | EVPL | -0,29568708 | 7,10254758 | -2,26461627 | 0,03930969 | 0,1757585 |
| 8364_at | HIST1H4C | -0,99190125 | 11,5079343 | -2,26447615 | 0,0393202 | 0,1757585 |
| 23582_at | CCNDBP1 | 0,39680911 | 8,53411003 | 2,26441099 | 0,03932509 | 0,1757585 |
| 7374_at | UNG | -0,51940923 | 9,24486677 | -2,26410297 | 0,0393482 | 0,1757913 |
| 151194_at | METTL21A | -0,27978322 | 6,94290025 | -2,26407002 | 0,03935068 | 0,1757913 |
| 790952_at | ESRG | -0,19069144 | 3,4179136 | -2,26329851 | 0,03940864 | 0,17600941 |
| 56126_at | PCDHB10 | 0,76612015 | 5,22940814 | 2,26313958 | 0,03942059 | 0,17602196 |
| 2300_at | FOXL1 | -0,31142221 | 5,11642674 | -2,26227972 | 0,0394853 | 0,17627006 |
| 9767_at | JADE3 | 0,19047363 | 7,53053891 | 2,26182663 | 0,03951944 | 0,17635463 |
| 27122_at | DKK3 | 0,20527727 | 4,43472828 | 2,26177042 | 0,03952367 | 0,17635463 |
| 826_at | CAPNS1 | 0,18891397 | 11,4161645 | 2,26166385 | 0,03953171 | 0,17635463 |
| 9373_at | PLAA | -0,16640414 | 7,23500838 | -2,26118687 | 0,03956769 | 0,17647427 |
| 124222_at | PAQR4 | -0,40004263 | 7,87406937 | -2,26105613 | 0,03957755 | 0,17647741 |
| 653784_at | MZT2A | -0,49208678 | 10,0097182 | -2,26010557 | 0,03964936 | 0,1767567 |
| 203562_at | TMEM31 | -0,21104836 | 3,54678661 | -2,25957622 | 0,03968941 | 0,17689427 |
| 27010_at | TPK1 | 0,45789676 | 4,05096992 | 2,25928101 | 0,03971175 | 0,1769338 |
| 55729_at | ATF7IP | 0,21382964 | 8,56365582 | 2,25921642 | 0,03971665 | 0,1769338 |
| 84976_at | DISP1 | 0,3932858 | 6,08812208 | 2,25883353 | 0,03974565 | 0,17695253 |
| 9805_at | SCRN1 | 0,29368944 | 8,09790899 | 2,25877058 | 0,03975042 | 0,17695253 |
| 1215_at | CMA1 | -0,25689732 | 3,68289612 | -2,2587631 | 0,03975099 | 0,17695253 |
| 54956_at | PARP16 | 0,16132386 | 7,26937457 | 2,25867595 | 0,03975759 | 0,17695253 |
| 202459_at | OSTCP1 | -0,18974962 | 3,70566976 | -2,2581924 | 0,03979426 | 0,17701522 |
| 10947_at | AP3M2 | -0,52083581 | 7,61621401 | -2,25806617 | 0,03980384 | 0,17701522 |
| 10199_at | MPHOSPH10 | -0,36925248 | 8,37531675 | -2,25804013 | 0,03980582 | 0,17701522 |
| 1056_at | CEL | 0,51193863 | 6,89929699 | 2,2580056 | 0,03980844 | 0,17701522 |
| 10370_at | CITED2 | 0,95699973 | 8,38106488 | 2,25775441 | 0,03982751 | 0,17705237 |
| 79581_at | SLC52A2 | -0,44997016 | 8,84121189 | -2,25765342 | 0,03983517 | 0,17705237 |
| 55691_at | FRMD4A | 0,30167988 | 6,17740548 | 2,25725679 | 0,03986531 | 0,17712582 |
| 51213_at | LUZP4 | -0,28721848 | 4,631247 | -2,25719385 | 0,03987009 | 0,17712582 |
| 126_at | ADH1C | 0,34068254 | 3,92063876 | 2,25697206 | 0,03988695 | 0,17715986 |
| 114044_at | MCM3AP-AS1 | -0,35140089 | 5,55153276 | -2,25677294 | 0,03990209 | 0,17716668 |
| 379025_at | PSMA3-AS1 | 0,26917197 | 7,47457978 | 2,25647713 | 0,0399246 | 0,17716668 |
| 200058_at | FLJ23867 | 0,32207413 | 8,44823383 | 2,25643791 | 0,03992758 | 0,17716668 |
| 6804_at | STX1A | -0,23583973 | 7,00679531 | -2,25637446 | 0,03993241 | 0,17716668 |
| 8237_at | USP11 | 0,28397045 | 9,58123623 | 2,2563474 | 0,03993447 | 0,17716668 |
| 56729_at | RETN | -0,28726508 | 5,99792776 | -2,25567839 | 0,03998543 | 0,17735189 |
| 782_at | CACNB1 | -0,18187726 | 5,51461241 | -2,25461472 | 0,04006656 | 0,17767086 |

| | | | | | | |
|---|---|---|---|---|---|---|
| 219402_at | MTIF3 | 0,22125754 | 8,40226481 | 2,25431469 | 0,04008948 | 0,17773156 |
| 80304_at | C2orf44 | -0,1603302 | 6,45661319 | -2,25413942 | 0,04010287 | 0,17773944 |
| 79971_at | WLS | 1,0876593 | 8,36626897 | 2,25404994 | 0,04010971 | 0,17773944 |
| 10537_at | UBD | 1,59693252 | 5,53373196 | 2,25382862 | 0,04012663 | 0,17777352 |
| 2253_at | FGF8 | -0,26461661 | 4,57038743 | -2,25243912 | 0,040233 | 0,1782038 |
| 101929621_a | LOC10192962 | -0,27255847 | 3,48416693 | -2,25211424 | 0,04025791 | 0,17827314 |
| 114801_at | TMEM200A | 0,39198794 | 4,39056826 | 2,25147523 | 0,04030695 | 0,17844927 |
| 57461_at | ISY1 | -0,15035312 | 6,88752196 | -2,25105873 | 0,04033894 | 0,17854987 |
| 54823_at | SWT1 | 0,40447501 | 5,45669335 | 2,25019081 | 0,04040568 | 0,17876185 |
| 8560_at | DEGS1 | -0,41095493 | 7,99305981 | -2,25014102 | 0,04040952 | 0,17876185 |
| 23555_at | TSPAN15 | 0,20079977 | 8,29044551 | 2,25007403 | 0,04041467 | 0,17876185 |
| 51053_at | GMNN | -0,6203985 | 9,701099 | -2,24979742 | 0,04043597 | 0,17881499 |
| 11035_at | RIPK3 | 0,39046091 | 6,05052448 | 2,24949699 | 0,04045911 | 0,17887627 |
| 204801_at | NLRP11 | -0,20057471 | 3,62195295 | -2,2491247 | 0,04048781 | 0,17896206 |
| 5446_at | PON3 | -0,19239751 | 3,57495237 | -2,24886992 | 0,04050746 | 0,17897238 |
| 5026_at | P2RX5 | -0,61029107 | 5,2277657 | -2,24885351 | 0,04050873 | 0,17897238 |
| 5435_at | POLR2F | -0,17582862 | 6,15961098 | -2,24830912 | 0,04055074 | 0,17911693 |
| 1211_at | CLTA | -0,13607062 | 10,5490541 | -2,24807899 | 0,04056852 | 0,17914939 |
| 84153_at | RNASEH2C | -0,34463484 | 8,04546185 | -2,24797317 | 0,04057669 | 0,17914939 |
| 375743_at | PTAR1 | 0,20215381 | 7,15354848 | 2,24765435 | 0,04060133 | 0,1792171 |
| 118611_at | C10orf90 | -0,13683306 | 3,99701156 | -2,24726865 | 0,04063116 | 0,17926863 |
| 2243_at | FGA | -0,13421947 | 3,30439047 | -2,24726272 | 0,04063162 | 0,17926863 |
| 100874167_a | LINC00423 | -0,216778 | 3,95655163 | -2,24670087 | 0,04067511 | 0,17941941 |
| 79443_at | FYCO1 | 0,29961371 | 6,85420883 | 2,24619758 | 0,0407141 | 0,17951319 |
| 144817_at | LINC00330 | -0,1800159 | 4,3343634 | -2,246078 | 0,04072337 | 0,17951319 |
| 57186_at | RALGAPA2 | 0,5193469 | 7,11191952 | 2,24606567 | 0,04072432 | 0,17951319 |
| 9209_at | LRRFIP2 | -0,32119049 | 7,1811934 | -2,24518736 | 0,04079247 | 0,17977244 |
| 148046_at | CIRBP-AS1 | -0,27289664 | 4,67635867 | -2,24488061 | 0,04081629 | 0,17983629 |
| 55335_at | NIPSNAP3B | 0,15048801 | 3,28543622 | 2,24475644 | 0,04082594 | 0,17983767 |
| 126308_at | MOB3A | 0,20461781 | 7,80143924 | 2,24457118 | 0,04084034 | 0,17985996 |
| 26128_at | KIF1BP | -0,22946136 | 8,87435933 | -2,24428738 | 0,04086241 | 0,17991601 |
| 23325_at | KIAA1033 | 0,33455245 | 8,24390937 | 2,2440511 | 0,04088079 | 0,17995581 |
| 55175_at | KLHL11 | -0,1426643 | 6,59270974 | -2,24333137 | 0,04093682 | 0,1801333 |
| 647309_at | GMNC | -0,21195085 | 3,85473134 | -2,2432205 | 0,04094546 | 0,1801333 |
| 6902_at | TBCA | -0,15322328 | 8,6880346 | -2,24312004 | 0,04095329 | 0,1801333 |
| 196513_at | DCP1B | 0,62856363 | 5,81965126 | 2,24305301 | 0,04095851 | 0,1801333 |
| 23506_at | GLTSCR1L | 0,26994032 | 6,99233832 | 2,24174162 | 0,04106085 | 0,18054214 |
| 257044_at | C1orf101 | 0,21910887 | 3,44883074 | 2,24161556 | 0,0410707 | 0,18054424 |
| 645431_at | FUT8-AS1 | 0,25073158 | 4,79575592 | 2,24095765 | 0,04112214 | 0,18069527 |
| 8193_at | DPF1 | -0,20329026 | 6,30221832 | -2,24093623 | 0,04112382 | 0,18069527 |
| 55204_at | GOLPH3L | 0,48943956 | 8,76277395 | 2,24016235 | 0,0411844 | 0,18088218 |
| 10399_at | GNB2L1 | -0,11611792 | 13,0722508 | -2,24015305 | 0,04118513 | 0,18088218 |
| 80117_at | ARL14 | 0,86709572 | 7,54206524 | 2,23987599 | 0,04120685 | 0,18093628 |
| 50485_at | SMARCAL1 | 0,16847242 | 7,8728032 | 2,23958363 | 0,04122977 | 0,18099568 |
| 51341_at | ZBTB7A | 0,14458733 | 7,81493512 | 2,23843623 | 0,04131985 | 0,18134979 |
| 7037_at | TFRC | -0,5812753 | 11,3468548 | -2,23827443 | 0,04133257 | 0,18136428 |
| 79784_at | MYH14 | 0,23801724 | 6,3252697 | 2,23764928 | 0,04138174 | 0,18151397 |
| 64147_at | KIF9 | -0,54203352 | 6,33104167 | -2,23760113 | 0,04138553 | 0,18151397 |
| 2959_at | GTF2B | 0,2226042 | 9,07588055 | 2,2368753 | 0,04144269 | 0,18169767 |
| 79064_at | TMEM223 | -0,29445459 | 9,28132784 | -2,23682979 | 0,04144627 | 0,18169767 |

| | | | | | | |
|---|---|---|---|---|---|---|
| 26610_at | ELP4 | -0,29225209 | 7,6167139 | -2,23640645 | 0,04147965 | 0,18180262 |
| 892_at | CCNC | -0,26778852 | 10,0459342 | -2,23621303 | 0,04149491 | 0,18182813 |
| 7442_at | TRPV1 | 0,24757742 | 5,39732871 | 2,23599378 | 0,04151221 | 0,18186257 |
| 23314_at | SATB2 | 0,36919441 | 6,41117051 | 2,23574146 | 0,04153213 | 0,18190847 |
| 100506686_a | IQCH-AS1 | 0,32823784 | 5,79510551 | 2,23518518 | 0,04157608 | 0,18205957 |
| 28973_at | MRPS18B | -0,13676502 | 8,94231516 | -2,23474555 | 0,04161084 | 0,1821095 |
| 54623_at | PAF1 | -0,13446062 | 8,30582156 | -2,23474027 | 0,04161126 | 0,1821095 |
| 8601_at | RGS20 | -0,21253651 | 5,95401588 | -2,23468232 | 0,04161584 | 0,1821095 |
| 80328_at | ULBP2 | -0,60939853 | 7,00511004 | -2,23437744 | 0,04163997 | 0,1821737 |
| 100130933_a | SMIM6 | 0,23656468 | 5,58585933 | 2,23425048 | 0,04165002 | 0,18217629 |
| 126299_at | ZNF428 | 0,21221095 | 6,99047786 | 2,23412734 | 0,04165977 | 0,18217758 |
| 29093_at | MRPL22 | -0,33708498 | 8,37478388 | -2,23378493 | 0,0416869 | 0,18221149 |
| 51013_at | EXOSC1 | -0,21808941 | 8,97592687 | -2,23367255 | 0,0416958 | 0,18221149 |
| 8434_at | RECK | 0,73484309 | 4,64849703 | 2,23363155 | 0,04169905 | 0,18221149 |
| 29960_at | FTSJ2 | -0,40810856 | 8,99428562 | -2,23355191 | 0,04170537 | 0,18221149 |
| 126353_at | MISP | 0,26495901 | 8,9139825 | 2,23241417 | 0,04179565 | 0,18256369 |
| 90233_at | ZNF551 | -0,32142416 | 6,30834345 | -2,23229732 | 0,04180493 | 0,18256369 |
| 26121_at | PRPF31 | -0,20878634 | 8,70828071 | -2,23214013 | 0,04181742 | 0,18257684 |
| 1755_at | DMBT1 | -0,28538446 | 5,8182524 | -2,23150255 | 0,04186812 | 0,1827478 |
| 8396_at | PIP4K2B | 0,20120431 | 7,67173153 | 2,2313025 | 0,04188404 | 0,1827478 |
| 55634_at | KRBOX4 | 0,24291049 | 7,33490734 | 2,23128994 | 0,04188504 | 0,1827478 |
| 114971_at | PTPMT1 | -0,18052871 | 8,06920358 | -2,23077707 | 0,04192588 | 0,18288456 |
| 91404_at | SESTD1 | 0,39039125 | 8,06824053 | 2,23058101 | 0,0419415 | 0,18291128 |
| 494513_at | DFNB59 | 0,22375182 | 4,25926163 | 2,23029772 | 0,04196408 | 0,18296834 |
| 222171_at | PRR15 | -0,34699097 | 8,14201836 | -2,23002488 | 0,04198584 | 0,18302179 |
| 81671_at | VMP1 | -0,15361166 | 10,551038 | -2,22982707 | 0,04200162 | 0,18304916 |
| 8581_at | LY6D | -0,52924185 | 11,4822345 | -2,22901234 | 0,04206669 | 0,18329124 |
| 405754_at | ERVFRD-1 | -0,21104098 | 4,23084406 | -2,22869143 | 0,04209234 | 0,18336154 |
| 55854_at | ZC3H15 | -0,36029965 | 10,0088968 | -2,22822429 | 0,04212971 | 0,18344952 |
| 79863_at | RBFA | -0,40459782 | 6,89435886 | -2,22820086 | 0,04213158 | 0,18344952 |
| 7165_at | TPD52L2 | -0,28172276 | 9,88399841 | -2,22791101 | 0,04215478 | 0,18350906 |
| 9587_at | MAD2L1BP | -0,16829832 | 9,02771717 | -2,22728628 | 0,04220483 | 0,18368543 |
| 84166_at | NLRC5 | 0,71349594 | 5,85793039 | 2,22708806 | 0,04222073 | 0,18371309 |
| 27151_at | CPAMD8 | 0,60185432 | 6,24110138 | 2,2262705 | 0,04228633 | 0,18395701 |
| 10113_at | PREB | -0,43569796 | 8,99785932 | -2,22579892 | 0,04232422 | 0,18408025 |
| 140838_at | NANP | -0,22407187 | 7,99295722 | -2,22542956 | 0,04235391 | 0,18416782 |
| 1806_at | DPYD | 0,29431997 | 6,51900273 | 2,2246658 | 0,04241538 | 0,18439348 |
| 2912_at | GRM2 | -0,18218768 | 4,97887137 | -2,22416355 | 0,04245585 | 0,18451762 |
| 10755_at | GIPC1 | 0,17113668 | 9,30947151 | 2,22407362 | 0,0424631 | 0,18451762 |
| 5970_at | RELA | 0,16167406 | 8,93518023 | 2,22355878 | 0,04250462 | 0,18463507 |
| 22890_at | ZBTB1 | 0,29924285 | 7,3632673 | 2,22350086 | 0,04250929 | 0,18463507 |
| 22881_at | ANKRD6 | 0,42530931 | 5,33789697 | 2,22330061 | 0,04252546 | 0,18466364 |
| 284359_at | IZUMO1 | -0,1739184 | 3,70693676 | -2,22249942 | 0,04259018 | 0,18490302 |
| 2194_at | FASN | -0,30713726 | 6,01159115 | -2,22217046 | 0,04261678 | 0,18497682 |
| 50488_at | MINK1 | 0,12430219 | 7,28306122 | 2,22144494 | 0,04267551 | 0,18518999 |
| 1369_at | CPN1 | -0,16710444 | 4,51687622 | -2,2209653 | 0,04271438 | 0,18531689 |
| 259266_at | ASPM | -0,79240438 | 7,3412395 | -2,22077672 | 0,04272966 | 0,18532143 |
| 25940_at | FAM98A | -0,1879398 | 9,44926087 | -2,2207151 | 0,04273466 | 0,18532143 |
| 27329_at | ANGPTL3 | -0,14959311 | 2,87863749 | -2,22049234 | 0,04275273 | 0,18533299 |
| 25897_at | RNF19A | 0,34786348 | 7,44959198 | 2,22044502 | 0,04275657 | 0,18533299 |

| | | | | | | |
|---|---|---|---|---|---|---|
| 7456_at | WIPF1 | 0,52335948 | 5,123397 | 2,22008114 | 0,0427861 | 0,18538665 |
| 602_at | BCL3 | 0,15987157 | 6,80565869 | 2,22005536 | 0,0427882 | 0,18538665 |
| 284680_at | C1orf111 | -0,29238549 | 4,46284284 | -2,21977879 | 0,04281066 | 0,18544226 |
| 154791_at | C7orf55 | -0,18642798 | 7,54617887 | -2,21932135 | 0,04284783 | 0,18555147 |
| 51313_at | FAM198B | 0,76865748 | 7,0946048 | 2,21912434 | 0,04286385 | 0,18555147 |
| 10154_at | PLXNC1 | 0,35416014 | 4,39173369 | 2,21911307 | 0,04286477 | 0,18555147 |
| 221035_at | REEP3 | 0,2520839 | 7,52045418 | 2,2189569 | 0,04287747 | 0,18555304 |
| 83881_at | MIXL1 | -0,2252771 | 4,41835113 | -2,21876344 | 0,04289321 | 0,18555304 |
| 8575_at | PRKRA | -0,15228527 | 8,50933574 | -2,218638 | 0,04290342 | 0,18555304 |
| 6527_at | SLC5A4 | -0,16653934 | 3,62828955 | -2,21863506 | 0,04290366 | 0,18555304 |
| 26285_at | CLDN17 | -0,2224893 | 4,79584675 | -2,21827458 | 0,04293301 | 0,18563831 |
| 23133_at | PHF8 | 0,18641213 | 6,05642531 | 2,21727474 | 0,04301452 | 0,18594901 |
| 1075_at | CTSC | 0,51587832 | 9,74690566 | 2,21686586 | 0,0430479 | 0,18605153 |
| 6612_at | SUMO3 | -0,23332966 | 10,1659472 | -2,21661073 | 0,04306874 | 0,18609984 |
| 389336_at | C5orf46 | -0,2450904 | 4,8662918 | -2,21640632 | 0,04308544 | 0,18613025 |
| 6811_at | STX5 | 0,22172927 | 7,03301123 | 2,21593871 | 0,04312367 | 0,18625363 |
| 26750_at | RPS6KC1 | 0,29296073 | 7,45099129 | 2,21565236 | 0,04314709 | 0,18631304 |
| 166824_at | RASSF6 | 0,76284767 | 6,57394689 | 2,21521181 | 0,04318316 | 0,18642697 |
| 84925_at | DIRC2 | 0,44806861 | 8,14768025 | 2,21478946 | 0,04321776 | 0,1865059 |
| 340719_at | NANOS1 | 0,53425268 | 7,59518944 | 2,21475231 | 0,0432208 | 0,1865059 |
| 5296_at | PIK3R2 | 0,20666107 | 7,4725565 | 2,21427247 | 0,04326015 | 0,18663029 |
| 23143_at | LRCH1 | -0,28015412 | 6,111454 | -2,21416448 | 0,04326901 | 0,18663029 |
| 64798_at | DEPTOR | -1,24472169 | 6,00135639 | -2,21392393 | 0,04328875 | 0,18666257 |
| 259217_at | HSPA12A | -0,42902774 | 5,67899897 | -2,21375381 | 0,04330271 | 0,18666257 |
| 132660_at | LIN54 | -0,34187747 | 5,50231462 | -2,21361137 | 0,04331441 | 0,18666257 |
| 9695_at | EDEM1 | -0,32718673 | 8,95485474 | -2,21360002 | 0,04331534 | 0,18666257 |
| 92579_at | G6PC3 | -0,1157492 | 9,12577016 | -2,21347543 | 0,04332557 | 0,18666257 |
| 10966_at | RAB40B | 0,53075971 | 6,20403222 | 2,21336519 | 0,04333463 | 0,18666257 |
| 79772_at | MCTP1 | -0,53032349 | 4,6265426 | -2,2125047 | 0,04340538 | 0,18692554 |
| 8828_at | NRP2 | -0,21754508 | 4,73315486 | -2,21132803 | 0,04350231 | 0,18724805 |
| 10948_at | STARD3 | 0,1500322 | 8,23781749 | 2,21122262 | 0,043511 | 0,18724805 |
| 56658_at | TRIM39 | 0,13615127 | 6,30420407 | 2,21115552 | 0,04351654 | 0,18724805 |
| 1262_at | CNGA4 | -0,28456595 | 4,03584984 | -2,21112272 | 0,04351924 | 0,18724805 |
| 152330_at | CNTN4 | 0,34072192 | 3,64717095 | 2,21093194 | 0,04353498 | 0,18724805 |
| 400073_at | C12orf76 | 0,35557782 | 6,33523582 | 2,21088812 | 0,0435386 | 0,18724805 |
| 23161_at | SNX13 | 0,28178169 | 7,68971768 | 2,21066395 | 0,0435571 | 0,18728581 |
| 51673_at | TPPP3 | -0,23752127 | 4,98169161 | -2,21046157 | 0,04357381 | 0,18731585 |
| 11266_at | DUSP12 | -0,37228044 | 8,58720284 | -2,21017217 | 0,04359772 | 0,18737275 |
| 9632_at | SEC24C | 0,190115 | 8,89611487 | 2,21006587 | 0,0436065 | 0,18737275 |
| 2028_at | ENPEP | 0,46552801 | 3,23088244 | 2,20925089 | 0,0436739 | 0,18762051 |
| 55588_at | MED29 | 0,2620775 | 8,27463059 | 2,20904634 | 0,04369083 | 0,1876514 |
| 3884_at | KRT33B | -0,28219017 | 5,01889797 | -2,20868086 | 0,0437211 | 0,18773954 |
| 101929897_a | LOC10192989 | -0,17332679 | 5,46959895 | -2,20808027 | 0,04377088 | 0,18791142 |
| 31_at | ACACA | -0,25402831 | 8,84754584 | -2,20794995 | 0,04378169 | 0,18791594 |
| 64175_at | P3H1 | 0,20918741 | 7,75803554 | 2,2077601 | 0,04379744 | 0,18794167 |
| 113000_at | RPUSD1 | -0,16200663 | 7,33648132 | -2,2070312 | 0,04385797 | 0,18815947 |
| 6671_at | SP4 | -0,26111447 | 6,05965531 | -2,20660695 | 0,04389323 | 0,18826883 |
| 4199_at | ME1 | -0,61155366 | 9,20866546 | -2,20542342 | 0,04399174 | 0,18864936 |
| 26586_at | CKAP2 | -0,34758459 | 6,98722877 | -2,20527822 | 0,04400384 | 0,18865925 |
| 93210_at | PGAP3 | 0,41826324 | 7,8933772 | 2,20414661 | 0,04409825 | 0,18902194 |

| | | | | | | |
|---|---|---|---|---|---|---|
| 27075_at | TSPAN13 | 0,50525973 | 10,652311 | 2,20336722 | 0,04416338 | 0,18925901 |
| 127829_at | ARL8A | 0,12140061 | 7,64860078 | 2,20284119 | 0,04420739 | 0,18940547 |
| 2553_at | GABPB1 | -0,3242528 | 7,87117871 | -2,20262612 | 0,0442254 | 0,18944048 |
| 16_at | AARS | -0,22120619 | 9,76395356 | -2,20231137 | 0,04425176 | 0,18949269 |
| 84276_at | NICN1 | 0,27562128 | 7,31906768 | 2,20224575 | 0,04425726 | 0,18949269 |
| 338661_at | TMEM225 | -0,21463251 | 3,95680075 | -2,20122768 | 0,04434264 | 0,18981607 |
| 51099_at | ABHD5 | 0,32472523 | 7,95114213 | 2,20108921 | 0,04435427 | 0,18982365 |
| 283303_at | MRGPRG-AS1 | -0,24098366 | 5,65691585 | -2,20032361 | 0,04441859 | 0,19005672 |
| 115939_at | TSR3 | -0,15823796 | 7,98229769 | -2,19989851 | 0,04445435 | 0,19016746 |
| 7220_at | TRPC1 | 0,37906566 | 3,79277263 | 2,1988688 | 0,04454107 | 0,19049612 |
| 10644_at | IGF2BP2 | -0,34801011 | 5,25941229 | -2,1984771 | 0,04457409 | 0,19055039 |
| 538_at | ATP7A | 0,32355448 | 7,44605853 | 2,1984289 | 0,04457816 | 0,19055039 |
| 79594_at | MUL1 | -0,15151933 | 8,38011458 | -2,1981606 | 0,0446008 | 0,19055039 |
| 51271_at | UBAP1 | 0,13391098 | 8,47222984 | 2,19812147 | 0,0446041 | 0,19055039 |
| 399664_at | MEX3D | 0,26619319 | 8,66888335 | 2,19810744 | 0,04460529 | 0,19055039 |
| 84305_at | WIBG | -0,17943921 | 8,11474399 | -2,1980148 | 0,0446131 | 0,19055039 |
| 9390_at | SLC22A13 | -0,2761938 | 4,43448288 | -2,19788054 | 0,04462444 | 0,19055655 |
| 116461_at | TSEN15 | -0,23877001 | 7,09631078 | -2,19742174 | 0,0446632 | 0,19067978 |
| 10686_at | CLDN16 | 0,27423847 | 6,61536031 | 2,19591781 | 0,04479046 | 0,19116052 |
| 25823_at | TPSG1 | -0,23960933 | 5,94407955 | -2,19585658 | 0,04479565 | 0,19116052 |
| 54790_at | TET2 | 0,38250152 | 7,01627862 | 2,19535253 | 0,04483838 | 0,19130049 |
| 23516_at | SLC39A14 | -0,42130443 | 7,11824359 | -2,194836 | 0,04488221 | 0,19144509 |
| 114883_at | OSBPL9 | 0,2444876 | 10,1369202 | 2,19434629 | 0,0449238 | 0,19158007 |
| 4116_at | MAGOH | -0,38836177 | 8,63748379 | -2,19335114 | 0,04500843 | 0,1918985 |
| 26509_at | MYOF | 0,57295906 | 10,5862308 | 2,19269645 | 0,04506419 | 0,19205578 |
| 133746_at | JMY | 0,52042224 | 6,34514242 | 2,19268387 | 0,04506526 | 0,19205578 |
| 28514_at | DLL1 | 0,402153 | 7,12999745 | 2,19226431 | 0,04510103 | 0,19216569 |
| 55008_at | HERC6 | 1,25673142 | 7,99358939 | 2,19196928 | 0,04512619 | 0,1922304 |
| 114571_at | SLC22A9 | -0,24682102 | 3,81792293 | -2,19149443 | 0,04516673 | 0,19235759 |
| 81622_at | UNC93B1 | 0,40235598 | 7,95583745 | 2,19138556 | 0,04517602 | 0,19235759 |
| 139189_at | DGKK | -0,31182987 | 4,15610551 | -2,19124033 | 0,04518843 | 0,19236789 |
| 2355_at | FOSL2 | 0,28382621 | 8,49048983 | 2,19010074 | 0,04528589 | 0,19274017 |
| 57096_at | RPGRIP1 | 0,27636526 | 4,79604433 | 2,18970416 | 0,04531985 | 0,19279977 |
| 387496_at | RASL11A | -0,29904188 | 5,42288949 | -2,18970345 | 0,04531991 | 0,19279977 |
| 51029_at | DESI2 | -0,23901589 | 8,16747001 | -2,18928929 | 0,0453554 | 0,19290816 |
| 23583_at | SMUG1 | 0,13587825 | 7,55131705 | 2,18898263 | 0,0453817 | 0,19294395 |
| 79714_at | CCDC51 | -0,22287994 | 7,41638634 | -2,18895052 | 0,04538445 | 0,19294395 |
| 84286_at | TMEM175 | 0,21530078 | 7,11345964 | 2,18865866 | 0,04540949 | 0,19294395 |
| 2572_at | GAD2 | -0,1919011 | 4,03632554 | -2,18863057 | 0,0454119 | 0,19294395 |
| 23030_at | KDM4B | 0,13998796 | 7,12592327 | 2,18860733 | 0,0454139 | 0,19294395 |
| 92552_at | ATXN3L | -0,15827558 | 2,97177547 | -2,18845785 | 0,04542673 | 0,19295591 |
| 54758_at | KLHDC4 | -0,32967181 | 5,72621403 | -2,1881615 | 0,04545218 | 0,1929964 |
| 92_at | ACVR2A | 0,41123995 | 7,33635586 | 2,18811353 | 0,0454563 | 0,1929964 |
| 148014_at | TTC9B | -0,20229429 | 5,34617522 | -2,187894 | 0,04547516 | 0,19303393 |
| 10803_at | CCR9 | -0,22557963 | 3,84430405 | -2,18772496 | 0,04548969 | 0,19305307 |
| 84132_at | USP42 | 0,24583599 | 7,46823979 | 2,18733813 | 0,04552295 | 0,19315168 |
| 26270_at | FBXO6 | 0,33869828 | 7,16674983 | 2,18678517 | 0,04557054 | 0,19328093 |
| 90480_at | GADD45GIP1 | -0,47686354 | 8,51120443 | -2,18654791 | 0,04559098 | 0,19328093 |
| 58493_at | INIP | -0,17788961 | 8,16276468 | -2,18653076 | 0,04559245 | 0,19328093 |
| 10480_at | EIF3M | -0,33419678 | 10,4462022 | -2,18651804 | 0,04559355 | 0,19328093 |

| | | | | | | |
|---|---|---|---|---|---|---|
| 55004_at | LAMTOR1 | 0,20860877 | 11,2913149 | 2,18633762 | 0,04560909 | 0,19329745 |
| 2642_at | GCGR | -0,23520691 | 6,08504749 | -2,18621377 | 0,04561977 | 0,19329745 |
| 27301_at | APEX2 | -0,29831383 | 8,24038498 | -2,18612349 | 0,04562755 | 0,19329745 |
| 83956_at | RACGAP1P | -0,14800741 | 3,17907202 | -2,18594898 | 0,0456426 | 0,19331867 |
| 55161_at | TMEM33 | -0,18437355 | 9,1618429 | -2,18564647 | 0,04566869 | 0,19338667 |
| 133383_at | SETD9 | -0,36061464 | 5,56144826 | -2,18511812 | 0,04571429 | 0,19353724 |
| 51816_at | CECR1 | 0,20423924 | 4,44380121 | 2,18479138 | 0,04574252 | 0,19361418 |
| 84981_at | MIR22HG | 0,49843991 | 8,31213218 | 2,18445615 | 0,04577149 | 0,19369427 |
| 388906_at | OGFRP1 | -0,2264199 | 6,31076928 | -2,18396769 | 0,04581374 | 0,19379004 |
| 100506697_a | LINC00327 | -0,14172667 | 3,56008574 | -2,18391702 | 0,04581813 | 0,19379004 |
| 90550_at | MCU | -0,27721336 | 8,16798659 | -2,18383973 | 0,04582482 | 0,19379004 |
| 56122_at | PCDHB14 | 1,03543397 | 6,20836262 | 2,18372941 | 0,04583437 | 0,19379004 |
| 2828_at | GPR4 | -0,18177714 | 5,14266424 | -2,18360311 | 0,0458453 | 0,19379374 |
| 130399_at | ACVR1C | 0,27156618 | 4,39283466 | 2,1832444 | 0,04587637 | 0,19388253 |
| 7728_at | ZNF175 | 0,2344321 | 6,39824933 | 2,18282802 | 0,04591246 | 0,19399249 |
| 386618_at | KCTD4 | -0,15253268 | 3,06736608 | -2,18209982 | 0,04597564 | 0,1941748 |
| 4097_at | MAFG | -0,29614139 | 7,98266607 | -2,18209833 | 0,04597577 | 0,1941748 |
| 4709_at | NDUFB3 | -0,17493233 | 9,62280205 | -2,18092117 | 0,04607807 | 0,19456422 |
| 285172_at | FAM126B | 0,16759554 | 6,17004473 | 2,18075463 | 0,04609256 | 0,19458275 |
| 3223_at | HOXC6 | 0,66262568 | 7,0628129 | 2,18040515 | 0,04612299 | 0,19462066 |
| 160897_at | GPR180 | -0,28065418 | 6,83962616 | -2,18039741 | 0,04612366 | 0,19462066 |
| 147372_at | CCBE1 | 0,23259217 | 4,50303298 | 2,18030331 | 0,04613186 | 0,19462066 |
| 27352_at | SGSM3 | 0,46689671 | 7,55360344 | 2,18017754 | 0,04614281 | 0,19462425 |
| 100505978_a | LOC10050597 | -0,16107418 | 4,19071744 | -2,1795281 | 0,04619942 | 0,19482035 |
| 3030_at | HADHA | 0,16715906 | 10,026513 | 2,1786717 | 0,04627416 | 0,19509282 |
| 2161_at | F12 | -0,5826903 | 7,03299774 | -2,1783315 | 0,04630388 | 0,19514168 |
| 84105_at | PCBD2 | -0,2169977 | 6,43870412 | -2,17830718 | 0,04630601 | 0,19514168 |
| 64422_at | ATG3 | -0,25960625 | 9,6933392 | -2,1770411 | 0,04641678 | 0,19552749 |
| 3109_at | HLA-DMB | 0,27086933 | 5,82878783 | 2,17702881 | 0,04641786 | 0,19552749 |
| 285375_at | LINC00620 | -0,16283812 | 4,46218961 | -2,17663705 | 0,04645219 | 0,19562788 |
| 5003_at | SLC22A18AS | -0,28643812 | 5,87180113 | -2,17652507 | 0,046462 | 0,19562788 |
| 100505989_a | LINC01207 | 0,86232582 | 3,73003783 | 2,17596939 | 0,04651074 | 0,1957903 |
| 11219_at | TREX2 | -0,28179093 | 4,95896088 | -2,17544788 | 0,04655653 | 0,19594023 |
| 1290_at | COL5A2 | 0,35883321 | 5,06921869 | 2,17501859 | 0,04659425 | 0,19605615 |
| 11080_at | DNAJB4 | 0,38176095 | 7,59091141 | 2,17474072 | 0,04661868 | 0,19611164 |
| 10146_at | G3BP1 | -0,16396141 | 8,08666444 | -2,17463706 | 0,0466278 | 0,19611164 |
| 55152_at | DALRD3 | 0,16619489 | 7,26530122 | 2,17422097 | 0,04666441 | 0,19614818 |
| 126402_at | CCDC105 | -0,27456114 | 5,34977755 | -2,17413516 | 0,04667197 | 0,19614818 |
| 3026_at | HABP2 | -0,20107837 | 4,67565027 | -2,17409605 | 0,04667541 | 0,19614818 |
| 27183_at | VPS4A | -0,22147778 | 8,63073471 | -2,17407551 | 0,04667722 | 0,19614818 |
| 54814_at | QPCTL | -0,13318137 | 5,84787737 | -2,17382673 | 0,04669913 | 0,19619745 |
| 4594_at | MUT | 0,26032361 | 8,13239174 | 2,17312019 | 0,0467614 | 0,19641625 |
| 22837_at | COBLL1 | 0,25502322 | 7,7270477 | 2,17274626 | 0,04679439 | 0,19644139 |
| 5029_at | P2RY2 | -0,3832436 | 7,68958859 | -2,17270538 | 0,046798 | 0,19644139 |
| 1609_at | DGKQ | 0,14415832 | 7,40900985 | 2,17263466 | 0,04680424 | 0,19644139 |
| 100505483_a | PRKAG2-AS1 | -0,38288259 | 4,08896847 | -2,17247203 | 0,0468186 | 0,19644139 |
| 5116_at | PCNT | -0,2959707 | 7,49779654 | -2,17243951 | 0,04682147 | 0,19644139 |
| 55907_at | CMAS | -0,26392021 | 8,41070203 | -2,17217174 | 0,04684512 | 0,19644139 |
| 102723526_a | LOC10272352 | -0,22523759 | 4,01184359 | -2,17214071 | 0,04684786 | 0,19644139 |
| 91662_at | NLRP12 | -0,21674606 | 4,88982668 | -2,17212817 | 0,04684897 | 0,19644139 |

| | | | | | | |
|---|---|---|---|---|---|---|
| 91010_at | FMNL3 | -0,2007459 | 4,25872444 | -2,17166674 | 0,04688976 | 0,19656961 |
| 4331_at | MNAT1 | -0,25059722 | 8,26356295 | -2,1712423 | 0,0469273 | 0,19667905 |
| 2150_at | F2RL1 | -0,48747779 | 7,8259895 | -2,17113308 | 0,04693696 | 0,19667905 |
| 403323_at | LOC403323 | 0,39750271 | 3,68005913 | 2,17102543 | 0,04694649 | 0,19667905 |
| 10336_at | PCGF3 | 0,16201305 | 7,71942202 | 2,17090659 | 0,04695701 | 0,19668035 |
| 6507_at | SLC1A3 | -0,31661387 | 6,65388127 | -2,17043473 | 0,04699881 | 0,19681262 |
| 150864_at | FAM117B | 0,24090922 | 6,48098916 | 2,17010839 | 0,04702773 | 0,19684505 |
| 85446_at | ZFHX2 | 0,18236701 | 5,53085303 | 2,1701046 | 0,04702807 | 0,19684505 |
| 3189_at | HNRNPH3 | -0,36910346 | 9,61729805 | -2,17000154 | 0,04703721 | 0,19684505 |
| 27345_at | KCNMB4 | 0,13998323 | 6,28399308 | 2,16956226 | 0,04707618 | 0,19696534 |
| 221491_at | C6orf1 | -0,16527114 | 6,797763 | -2,16925246 | 0,04710368 | 0,19700014 |
| 56945_at | MRPS22 | -0,37804308 | 8,68841638 | -2,16923817 | 0,04710495 | 0,19700014 |
| 3198_at | HOXA1 | 0,52532445 | 8,10358854 | 2,16883228 | 0,047141 | 0,19708368 |
| 1021_at | CDK6 | 0,35062725 | 6,57513942 | 2,16878294 | 0,04714539 | 0,19708368 |
| 3442_at | IFNA5 | 0,18418822 | 3,16024198 | 2,16847032 | 0,04717318 | 0,19715707 |
| 9451_at | EIF2AK3 | 0,28945321 | 8,50719853 | 2,16821032 | 0,0471963 | 0,19721093 |
| 54482_at | TRMT13 | -0,24559471 | 7,46981297 | -2,16732293 | 0,0472753 | 0,1974982 |
| 55763_at | EXOC1 | 0,29040977 | 8,66082045 | 2,16711606 | 0,04729374 | 0,19753237 |
| 3241_at | HPCAL1 | -0,44265888 | 7,93939119 | -2,16640831 | 0,04735686 | 0,19775313 |
| 51317_at | PHF21A | 0,24515889 | 6,98357722 | 2,16608417 | 0,04738579 | 0,19780031 |
| 23786_at | BCL2L13 | 0,10957945 | 7,46339196 | 2,16601916 | 0,0473916 | 0,19780031 |
| 201516_at | ZSCAN4 | 0,61560247 | 4,67153395 | 2,16592487 | 0,04740002 | 0,19780031 |
| 9352_at | TXNL1 | -0,23731752 | 9,61754775 | -2,16582171 | 0,04740923 | 0,19780031 |
| 22877_at | MLXIP | 0,24576288 | 7,90286177 | 2,16494453 | 0,04748765 | 0,19808459 |
| 55624_at | POMGNT1 | -0,11782779 | 8,04178877 | -2,16456667 | 0,04752147 | 0,19818274 |
| 89891_at | WDR34 | -0,38715797 | 9,18314323 | -2,16414799 | 0,04755897 | 0,19825764 |
| 57476_at | GRAMD1B | -0,19721231 | 5,28837532 | -2,16408881 | 0,04756427 | 0,19825764 |
| 57649_at | PHF12 | 0,26400981 | 7,35435383 | 2,16402143 | 0,04757031 | 0,19825764 |
| 55863_at | TMEM126B | -0,43124992 | 8,83844503 | -2,16291614 | 0,04766945 | 0,19862646 |
| 101928590_a | LOC10192859 | -0,13601886 | 2,91952099 | -2,16280506 | 0,04767942 | 0,19862646 |
| 2184_at | FAH | -0,37243198 | 8,04238546 | -2,16229243 | 0,04772548 | 0,19875062 |
| 4154_at | MBNL1 | -0,29582751 | 8,66054627 | -2,16220287 | 0,04773353 | 0,19875062 |
| 2747_at | GLUD2 | -0,12392238 | 7,23364519 | -2,16212889 | 0,04774018 | 0,19875062 |
| 7186_at | TRAF2 | -0,35852129 | 7,12516877 | -2,16173795 | 0,04777534 | 0,19878702 |
| 54534_at | MRPL50 | -0,44829257 | 8,20900703 | -2,16172708 | 0,04777632 | 0,19878702 |
| 115353_at | LRRC42 | -0,15646116 | 8,94076074 | -2,16168748 | 0,04777988 | 0,19878702 |
| 284439_at | SLC25A42 | 0,16739219 | 5,7470663 | 2,16140529 | 0,04780528 | 0,19884974 |
| 9818_at | NUP58 | -0,46523754 | 6,9306487 | -2,16109674 | 0,04783307 | 0,19892235 |
| 22927_at | HABP4 | 0,20692237 | 6,23568733 | 2,16047319 | 0,04788926 | 0,19911306 |
| 100507218_a | RNF157-AS1 | 0,35468824 | 4,20704558 | 2,16017942 | 0,04791576 | 0,19918024 |
| 7141_at | TNP1 | -0,23983292 | 5,45745705 | -2,1598243 | 0,0479478 | 0,19923385 |
| 57019_at | CIAPIN1 | -0,17627343 | 8,50764018 | -2,15980728 | 0,04794934 | 0,19923385 |
| 54209_at | TREM2 | -0,2336525 | 5,50958113 | -2,15958771 | 0,04796917 | 0,19927325 |
| 5436_at | POLR2G | -0,23608952 | 9,95597831 | -2,15936875 | 0,04798894 | 0,19930831 |
| 51616_at | TAF9B | 0,23614527 | 8,58727585 | 2,15917617 | 0,04800635 | 0,19930831 |
| 140825_at | NEURL2 | 0,17000937 | 5,00662093 | 2,15915071 | 0,04800865 | 0,19930831 |
| 114826_at | SMYD4 | -0,22124502 | 6,87359397 | -2,15896632 | 0,04802531 | 0,19933454 |
| 283551_at | LINC01588 | -0,14510654 | 4,61980331 | -2,15881052 | 0,0480394 | 0,19935006 |
| 402778_at | IFITM10 | 0,30939149 | 7,27808196 | 2,15856094 | 0,04806198 | 0,19940078 |
| 11313_at | LYPLA2 | 0,22068848 | 9,06767997 | 2,15836514 | 0,04807969 | 0,19943134 |

| | | | | | | |
|---|---|---|---|---|---|---|
| 83941_at | TM2D1 | 0,27063225 | 8,19475753 | 2,15778245 | 0,04813246 | 0,19960721 |
| 11174_at | ADAMTS6 | 0,14022652 | 3,59697312 | 2,15752942 | 0,04815538 | 0,19965931 |
| 644714_at | LIMD1-AS1 | -0,21816931 | 5,55886765 | -2,1571743 | 0,04818758 | 0,19974981 |
| 57717_at | PCDHB16 | 0,19679003 | 4,3089881 | 2,15696172 | 0,04820686 | 0,19978675 |
| 6667_at | SP1 | 0,27840195 | 7,48481312 | 2,15671297 | 0,04822944 | 0,19979603 |
| 55227_at | LRRC1 | 0,19247899 | 9,12515957 | 2,15670845 | 0,04822985 | 0,19979603 |
| 79175_at | ZNF343 | -0,21025673 | 6,68383957 | -2,15624175 | 0,04827222 | 0,19992858 |
| 146206_at | RLTPR | -0,24368567 | 5,61154632 | -2,15609825 | 0,04828526 | 0,19993959 |
| 7775_at | ZNF232 | -0,37388519 | 6,57277854 | -2,15595326 | 0,04829844 | 0,19995116 |
| 9770_at | RASSF2 | 0,79159778 | 6,2540956 | 2,15564312 | 0,04832663 | 0,19996691 |
| 80231_at | CXorf21 | -0,16106903 | 2,90046142 | -2,15562285 | 0,04832847 | 0,19996691 |
| 3115_at | HLA-DPB1 | 0,21007245 | 3,90678514 | 2,15555848 | 0,04833433 | 0,19996691 |
| 30815_at | ST6GALNAC6 | 0,21228419 | 7,03068706 | 2,15545475 | 0,04834376 | 0,19996691 |
| 26333_at | OR7A17 | -0,19445555 | 3,66881235 | -2,15470535 | 0,04841198 | 0,20020607 |
| 100507283_a | LOC10050728 | -0,17537537 | 3,54810011 | -2,15431821 | 0,04844725 | 0,2003024 |
| 90161_at | HS6ST2 | -0,43689325 | 8,113098 | -2,15419692 | 0,0484583 | 0,2003024 |
| 132001_at | TAMM41 | -0,34106236 | 6,1877569 | -2,15402826 | 0,04847368 | 0,2003024 |
| 9668_at | ZNF432 | 0,32130614 | 6,40204652 | 2,15399341 | 0,04847686 | 0,2003024 |
| 8394_at | PIP5K1A | 0,26198052 | 8,59835853 | 2,15361369 | 0,0485115 | 0,20040254 |
| 271_at | AMPD2 | -0,2114254 | 8,13775996 | -2,152964 | 0,04857082 | 0,20053853 |
| 79929_at | MAP6D1 | -0,26517486 | 6,16463173 | -2,1529478 | 0,0485723 | 0,20053853 |
| 6261_at | RYR1 | -0,30814537 | 4,20171511 | -2,15274984 | 0,04859039 | 0,20053853 |
| 23270_at | TSPYL4 | 0,3234341 | 9,04428567 | 2,15273835 | 0,04859144 | 0,20053853 |
| 966_at | CD59 | 0,17702643 | 9,62200022 | 2,15268332 | 0,04859647 | 0,20053853 |
| 157570_at | ESCO2 | -0,83531033 | 6,80552792 | -2,1524422 | 0,04861852 | 0,20058652 |
| 5334_at | PLCL1 | 0,21107192 | 3,17128734 | 2,151541 | 0,04870099 | 0,20088376 |
| 55666_at | NPLOC4 | -0,16263166 | 9,47538651 | -2,15132771 | 0,04872053 | 0,20092133 |
| 282997_at | PDCD4-AS1 | 0,23815885 | 4,02754489 | 2,15100396 | 0,0487502 | 0,20099548 |
| 92659_at | MAFG-AS1 | -0,2701195 | 7,23754279 | -2,15090386 | 0,04875938 | 0,20099548 |
| 84681_at | HINT2 | -0,20013398 | 8,91968329 | -2,1507127 | 0,04877691 | 0,20102472 |
| 27445_at | PCLO | 0,46533467 | 4,884278 | 2,15048588 | 0,04879771 | 0,20106746 |
| 23201_at | FAM168A | 0,24342344 | 7,34663079 | 2,14982722 | 0,04885819 | 0,20127357 |
| 5886_at | RAD23A | -0,35165103 | 9,57568569 | -2,14958451 | 0,04888048 | 0,20132238 |
| 6646_at | SOAT1 | 0,40085068 | 8,07597068 | 2,14928679 | 0,04890785 | 0,20139204 |
| 84324_at | SARNP | -0,18316589 | 6,92000116 | -2,14790093 | 0,04903543 | 0,20184856 |
| 100132074_a | FOXO6 | 0,6450702 | 7,96257012 | 2,14785491 | 0,04903967 | 0,20184856 |
| 196264_at | MPZL3 | 0,51026304 | 8,10660206 | 2,14770947 | 0,04905308 | 0,20185974 |
| 64793_at | CEP85 | -0,30679678 | 7,82479544 | -2,14749258 | 0,04907308 | 0,20185974 |
| 6503_at | SLA | -0,18185132 | 4,85378483 | -2,1474368 | 0,04907823 | 0,20185974 |
| 79631_at | EFTUD1 | 0,18192472 | 6,87000983 | 2,1473137 | 0,04908959 | 0,20185974 |
| 202333_at | CMYA5 | 0,53243634 | 4,73105676 | 2,14725733 | 0,04909479 | 0,20185974 |
| 56260_at | C8orf44 | 0,32081956 | 4,67894373 | 2,14664811 | 0,04915103 | 0,20204787 |
| 30008_at | EFEMP2 | 0,21020597 | 5,33907619 | 2,14650001 | 0,04916471 | 0,20206099 |
| 91544_at | UBXN11 | -0,23113286 | 6,66035249 | -2,14622046 | 0,04919055 | 0,20212405 |
| 494470_at | RNF165 | -0,21280483 | 6,06060534 | -2,14532662 | 0,04927324 | 0,20242065 |
| 60385_at | TSKS | -0,1834185 | 4,52203902 | -2,14502136 | 0,04930151 | 0,2024936 |
| 51170_at | HSD17B11 | 0,51846988 | 8,14188007 | 2,14456985 | 0,04934335 | 0,20262225 |
| 1412_at | CRYBA2 | -0,26576724 | 4,65438538 | -2,1440968 | 0,04938722 | 0,20273933 |
| 4292_at | MLH1 | -0,37182538 | 8,9240909 | -2,1439741 | 0,04939861 | 0,20273933 |
| 1478_at | CSTF2 | -0,60415209 | 7,46391648 | -2,14377765 | 0,04941684 | 0,20273933 |

| | | | | | | |
|---|---|---|---|---|---|---|
| 151887_at | CCDC80 | 0,48193583 | 6,75651489 | 2,14372838 | 0,04942142 | 0,20273933 |
| 1122_at | CHML | -0,31009622 | 6,40022199 | -2,14369533 | 0,04942449 | 0,20273933 |
| 9692_at | KIAA0391 | -0,25778507 | 8,41091476 | -2,14333743 | 0,04945773 | 0,20280999 |
| 3655_at | ITGA6 | -0,71673339 | 10,2922547 | -2,14328316 | 0,04946277 | 0,20280999 |
| 9704_at | DHX34 | -0,19696058 | 6,44481764 | -2,14302998 | 0,0494863 | 0,20286328 |
| 152185_at | SPICE1 | -0,2513746 | 4,45343477 | -2,14287951 | 0,04950029 | 0,20287746 |
| 84706_at | GPT2 | -0,53065649 | 6,34034491 | -2,14271525 | 0,04951556 | 0,20289689 |
| 51569_at | UFM1 | 0,22700134 | 8,28087541 | 2,14178581 | 0,04960208 | 0,20320816 |
| 256006_at | ANKRD31 | -0,11984406 | 3,20385024 | -2,14122101 | 0,04965472 | 0,20338057 |
| 6238_at | RRBP1 | 0,16722028 | 8,27457982 | 2,14083592 | 0,04969064 | 0,20346731 |
| 2526_at | FUT4 | 0,46745761 | 6,08513389 | 2,14073109 | 0,04970042 | 0,20346731 |
| 57541_at | ZNF398 | 0,12667383 | 7,58147546 | 2,14061452 | 0,0497113 | 0,20346731 |
| 6617_at | SNAPC1 | -0,63091851 | 7,91749065 | -2,1405412 | 0,04971814 | 0,20346731 |
| 3238_at | HOXD12 | -0,23368152 | 3,45223772 | -2,14007388 | 0,04976179 | 0,20360267 |
| 8522_at | GAS7 | -0,23539211 | 4,58293765 | -2,13903531 | 0,04985892 | 0,20395675 |
| 5595_at | MAPK3 | 0,13517958 | 8,51315043 | 2,13884088 | 0,04987712 | 0,2039684 |
| 6657_at | SOX2 | 0,52433323 | 5,53437662 | 2,13877869 | 0,04988294 | 0,2039684 |
| 57610_at | RANBP10 | 0,18671098 | 7,14976465 | 2,13862206 | 0,04989761 | 0,20398508 |
| 151516_at | ASPRV1 | -0,15771466 | 5,19182897 | -2,13793255 | 0,04996224 | 0,20420594 |
| 121278_at | TPH2 | -0,151976 | 3,1226414 | -2,13734368 | 0,05001749 | 0,20438841 |
| 139322_at | APOOL | 0,256342 | 7,43881606 | 2,13649191 | 0,05009751 | 0,20467199 |
| 203190_at | LGI3 | -0,1825776 | 5,47558875 | -2,13630364 | 0,05011522 | 0,2047009 |
| 132141_at | IQCF1 | -0,18905046 | 4,59178566 | -2,13562297 | 0,05017927 | 0,20491908 |
| 8484_at | GALR3 | -0,24645163 | 7,0799299 | -2,13544366 | 0,05019616 | 0,20493243 |
| 23012_at | STK38L | 0,29524782 | 7,73130985 | 2,13527713 | 0,05021185 | 0,20493243 |
| 51247_at | PAIP2 | 0,20865103 | 9,78568892 | 2,13520014 | 0,0502191 | 0,20493243 |
| 157285_at | SGK223 | -0,22200707 | 8,56554932 | -2,13510508 | 0,05022806 | 0,20493243 |
| 339221_at | ENPP7 | -0,25669619 | 5,87871978 | -2,13502365 | 0,05023573 | 0,20493243 |
| 100505975_a | ENO1-AS1 | 0,15744131 | 4,56581161 | 2,13479305 | 0,05025747 | 0,20497771 |
| 126074_at | SWSAP1 | 0,20991682 | 6,62055321 | 2,13429125 | 0,05030481 | 0,20512735 |
| 54729_at | NKX1-1 | -0,21778521 | 6,10617845 | -2,13403416 | 0,05032908 | 0,20518288 |
| 9149_at | DYRK1B | 0,29182008 | 6,36884344 | 2,13365175 | 0,0503652 | 0,20521566 |
| 117154_at | DACH2 | 0,5037499 | 4,19866967 | 2,13364588 | 0,05036576 | 0,20521566 |
| 79921_at | TCEAL4 | 0,17316447 | 8,94871988 | 2,13358278 | 0,05037172 | 0,20521566 |
| 6819_at | SULT1C2 | 0,13498996 | 3,59670465 | 2,13349793 | 0,05037974 | 0,20521566 |
| 10554_at | AGPAT1 | 0,17529061 | 8,65171082 | 2,13321823 | 0,05040618 | 0,20527995 |
| 284323_at | ZNF780A | 0,27075157 | 5,78522321 | 2,13290565 | 0,05043574 | 0,20535693 |
| 144406_at | WDR66 | 0,3961 | 4,58376567 | 2,13221795 | 0,05050084 | 0,20557165 |
| 5828_at | PEX2 | 0,241569 | 7,61785724 | 2,13212317 | 0,05050982 | 0,20557165 |
| 171425_at | CLYBL | 0,20200907 | 4,64616912 | 2,13187824 | 0,05053302 | 0,20562266 |
| 55174_at | INTS10 | -0,43642666 | 8,06745978 | -2,13146253 | 0,05057244 | 0,20573957 |
| 7699_at | ZNF140 | 0,36697298 | 7,28585897 | 2,13133702 | 0,05058434 | 0,20574455 |
| 1376_at | CPT2 | 0,22299283 | 8,0207182 | 2,1311342 | 0,05060359 | 0,20577937 |
| 10672_at | GNA13 | 0,28390157 | 8,40725669 | 2,1308911 | 0,05062666 | 0,20582975 |
| 440952_at | EIF1B-AS1 | -0,18835401 | 5,3085243 | -2,12949645 | 0,05075922 | 0,20630495 |
| 7508_at | XPC | 0,27927073 | 7,84866924 | 2,12943616 | 0,05076496 | 0,20630495 |
| 54780_at | NSMCE4A | -0,16847371 | 8,50331193 | -2,1288002 | 0,05082552 | 0,20650751 |
| 6603_at | SMARCD2 | -0,19292901 | 9,02812656 | -2,12829254 | 0,05087392 | 0,20666053 |
| 4646_at | MYO6 | 0,35260469 | 8,81244631 | 2,12772432 | 0,05092813 | 0,20683715 |
| 23240_at | KIAA0922 | -0,14054066 | 5,64531328 | -2,12746992 | 0,05095242 | 0,20689218 |

| | | | | | | |
|---|---|---|---|---|---|---|
| 5130_at | PCYT1A | -0,17648346 | 7,09789293 | -2,12677494 | 0,05101883 | 0,20711819 |
| 401093_at | MBNL1-AS1 | 0,50714857 | 5,94385205 | 2,12553638 | 0,05113739 | 0,20754153 |
| 1549_at | CYP2A7 | -0,50853323 | 4,280148 | -2,12546048 | 0,05114466 | 0,20754153 |
| 1416_at | CRYBB2P1 | -0,25181414 | 6,33965552 | -2,12520516 | 0,05116914 | 0,20759711 |
| 2768_at | GNA12 | -0,1361649 | 8,03190101 | -2,12477345 | 0,05121054 | 0,20768064 |
| 414328_at | IDNK | 0,36595868 | 5,86062508 | 2,12476569 | 0,05121129 | 0,20768064 |
| 101926960_a | LOC10192696 | 0,21083837 | 2,91527763 | 2,12443011 | 0,0512435 | 0,20776752 |
| 57595_at | PDZD4 | -0,19728963 | 5,72214913 | -2,12373134 | 0,05131062 | 0,2079959 |
| 121441_at | NEDD1 | -0,39040672 | 8,19397918 | -2,12311729 | 0,05136968 | 0,20819148 |
| 23309_at | SIN3B | 0,31711354 | 7,6315389 | 2,1226368 | 0,05141593 | 0,2083172 |
| 23216_at | TBC1D1 | 0,18236549 | 7,19076277 | 2,1225704 | 0,05142233 | 0,2083172 |
| 633_at | BGN | -0,20901312 | 5,2540177 | -2,12226157 | 0,05145208 | 0,20839391 |
| 84279_at | PRADC1 | -0,2574278 | 7,85442193 | -2,12209069 | 0,05146855 | 0,20841679 |
| 55833_at | UBAP2 | -0,14005596 | 9,62676551 | -2,12190179 | 0,05148676 | 0,20844672 |
| 10762_at | NUP50 | -0,22387865 | 7,36364605 | -2,12178033 | 0,05149848 | 0,20845033 |
| 8985_at | PLOD3 | -0,33045986 | 9,33660635 | -2,12027218 | 0,05164412 | 0,20899596 |
| 84950_at | PRPF38A | -0,2034326 | 8,97915545 | -2,12014756 | 0,05165618 | 0,20900082 |
| 5495_at | PPM1B | 0,1538646 | 8,60477668 | 2,11953159 | 0,05171578 | 0,20919806 |
| 79763_at | ISOC2 | -0,19691487 | 8,82447519 | -2,11934708 | 0,05173365 | 0,2092264 |
| 54460_at | MRPS21 | -0,22088466 | 10,1339305 | -2,11889599 | 0,05177736 | 0,20935921 |
| 401052_at | LOC401052 | 0,31522714 | 4,795665 | 2,11858437 | 0,05180757 | 0,20943741 |
| 2908_at | NR3C1 | 0,25746826 | 8,13813958 | 2,11827098 | 0,05183798 | 0,20946143 |
| 65065_at | NBEAL1 | 0,1518027 | 5,23502511 | 2,11815002 | 0,05184971 | 0,20946143 |
| 79595_at | SAP130 | -0,1708846 | 8,74139182 | -2,11808661 | 0,05185587 | 0,20946143 |
| 347273_at | MURC | 0,24493525 | 3,97273157 | 2,11807485 | 0,05185701 | 0,20946143 |
| 644186_at | SYCE3 | 0,16140382 | 5,0406724 | 2,1176199 | 0,05190119 | 0,20959593 |
| 89849_at | ATG16L2 | 0,3138229 | 5,83079805 | 2,11717332 | 0,05194459 | 0,20971201 |
| 8894_at | EIF2S2 | -0,33477641 | 7,5797088 | -2,11700903 | 0,05196056 | 0,20971201 |
| 2627_at | GATA6 | 0,52779262 | 6,9519475 | 2,11698815 | 0,05196259 | 0,20971201 |
| 114659_at | LRRC37B | 0,35452837 | 6,30131302 | 2,11628676 | 0,05203084 | 0,20994348 |
| 64744_at | SMAP2 | -0,17422032 | 7,96432374 | -2,11556311 | 0,05210135 | 0,21018394 |
| 5167_at | ENPP1 | 0,31063373 | 4,38512246 | 2,11507103 | 0,05214934 | 0,2103335 |
| 8519_at | IFITM1 | 1,49672382 | 7,32658984 | 2,11446027 | 0,05220897 | 0,21052991 |
| 100133461_a | LOC10013346 | -0,14974924 | 5,08343239 | -2,11406023 | 0,05224806 | 0,21064344 |
| 10046_at | MAMLD1 | -0,28176064 | 5,12223377 | -2,11352831 | 0,05230007 | 0,21080902 |
| 25770_at | C22orf31 | -0,17833763 | 4,10918871 | -2,11328137 | 0,05232424 | 0,2108623 |
| 85378_at | TUBGCP6 | 0,26923741 | 6,49429125 | 2,11289579 | 0,05236199 | 0,21091626 |
| 400684_at | LOC400684 | -0,26245359 | 4,81533767 | -2,11276137 | 0,05237516 | 0,21091626 |
| 80018_at | NAA25 | -0,32815911 | 8,15076835 | -2,11274167 | 0,05237709 | 0,21091626 |
| 118491_at | CFAP70 | 0,30478486 | 3,35549035 | 2,11269739 | 0,05238142 | 0,21091626 |
| 1717_at | DHCR7 | -0,47835588 | 10,0519936 | -2,11209468 | 0,0524405 | 0,21111002 |
| 51422_at | PRKAG2 | -0,31967181 | 7,35742437 | -2,11189817 | 0,05245978 | 0,2111435 |
| 5441_at | POLR2L | -0,33813416 | 9,16093942 | -2,11114806 | 0,05253342 | 0,21139572 |
| 1907_at | EDN2 | 0,66707174 | 6,76896373 | 2,11103104 | 0,05254492 | 0,21139782 |
| 25802_at | LMOD1 | -0,27180827 | 4,6165612 | -2,11020859 | 0,05262579 | 0,2116664 |
| 23533_at | PIK3R5 | -0,1799263 | 5,0676806 | -2,11012867 | 0,05263365 | 0,2116664 |
| 145845_at | LOC145845 | -0,14064727 | 3,99464111 | -2,10890105 | 0,05275459 | 0,21209447 |
| 26151_at | NAT9 | -0,12235835 | 7,45431395 | -2,10881101 | 0,05276347 | 0,21209447 |
| 3578_at | IL9 | -0,22242746 | 3,50353706 | -2,10871311 | 0,05277313 | 0,21209447 |
| 7265_at | TTC1 | -0,21226804 | 7,71917088 | -2,10847453 | 0,05279667 | 0,21214482 |

| | | | | | | |
|---|---|---|---|---|---|---|
| 84807_at | NFKBID | -0,169341 | 5,58469961 | -2,10805428 | 0,05283816 | 0,21221289 |
| 10548_at | TM9SF1 | 0,2309697 | 9,67157006 | 2,10798854 | 0,05284465 | 0,21221289 |
| 8801_at | SUCLG2 | 0,16860439 | 7,34912061 | 2,10796822 | 0,05284666 | 0,21221289 |
| 79670_at | ZCCHC6 | 0,23921337 | 6,26165259 | 2,10756958 | 0,05288605 | 0,21227832 |
| 6518_at | SLC2A5 | -0,18368753 | 5,69035347 | -2,10746252 | 0,05289663 | 0,21227832 |
| 23766_at | GABARAPL3 | 0,17777795 | 3,86935111 | 2,10739446 | 0,05290336 | 0,21227832 |
| 2054_at | STX2 | -0,22063725 | 6,54790806 | -2,10735733 | 0,05290703 | 0,21227832 |
| 662_at | BNIP1 | -0,17709801 | 6,26036085 | -2,10650929 | 0,05299095 | 0,21257074 |
| 51621_at | KLF13 | 0,19768513 | 6,40489033 | 2,10616267 | 0,05302529 | 0,21266419 |
| 89910_at | UBE3B | 0,10767494 | 7,65388592 | 2,10534282 | 0,05310658 | 0,21293038 |
| 79983_at | POF1B | 0,42038824 | 6,40041185 | 2,10521708 | 0,05311906 | 0,21293038 |
| 84766_at | CRACR2A | -0,21826141 | 4,83570098 | -2,10515903 | 0,05312482 | 0,21293038 |
| 162073_at | ITPRIPL2 | -0,15406921 | 8,85096318 | -2,10480301 | 0,05316017 | 0,21302773 |
| 143379_at | C10orf82 | -0,24948181 | 4,62110844 | -2,10458288 | 0,05318203 | 0,21303909 |
| 25841_at | ABTB2 | -0,2375205 | 6,23016576 | -2,10455183 | 0,05318512 | 0,21303909 |
| 84948_at | TIGD5 | -0,25460699 | 7,25348596 | -2,10369735 | 0,05327009 | 0,21333506 |
| 5209_at | PFKFB3 | 0,37407499 | 9,51342636 | 2,10325853 | 0,05331377 | 0,213453 |
| 22986_at | SORCS3 | -0,15718298 | 3,50235264 | -2,10317892 | 0,0533217 | 0,213453 |
| 51298_at | THEG | -0,26685675 | 5,13487357 | -2,10280992 | 0,05335846 | 0,21349581 |
| 23085_at | ERC1 | 0,30994713 | 6,01251267 | 2,10274818 | 0,05336461 | 0,21349581 |
| 10457_at | GPNMB | 1,27190758 | 7,52564629 | 2,10273786 | 0,05336564 | 0,21349581 |
| 57538_at | ALPK3 | 0,21272761 | 5,63917585 | 2,10261492 | 0,0533779 | 0,2135005 |
| 8877_at | SPHK1 | -0,33469934 | 6,12387549 | -2,10227831 | 0,05341146 | 0,21354649 |
| 116143_at | WDR92 | -0,18599277 | 7,48107825 | -2,10211141 | 0,05342812 | 0,21354649 |
| 285193_at | DUSP28 | 0,16184676 | 7,67811563 | 2,10201868 | 0,05343737 | 0,21354649 |
| 100132677_a | BSN-AS2 | -0,20864065 | 4,47244675 | -2,10196478 | 0,05344275 | 0,21354649 |
| 164781_at | DAW1 | -0,20277142 | 3,56357018 | -2,10193271 | 0,05344595 | 0,21354649 |
| 284996_at | RNF149 | -0,18823685 | 8,71743297 | -2,10175108 | 0,05346408 | 0,21354649 |
| 7627_at | ZNF75A | 0,28937088 | 6,4909205 | 2,10172188 | 0,053467 | 0,21354649 |
| 136647_at | MPLKIP | -0,1394545 | 8,80146705 | -2,10104668 | 0,05353445 | 0,21373989 |
| 5698_at | PSMB9 | 1,01452495 | 9,31847788 | 2,10097748 | 0,05354137 | 0,21373989 |
| 8366_at | HIST1H4B | -0,26677854 | 5,53058834 | -2,10090411 | 0,0535487 | 0,21373989 |
| 3008_at | HIST1H1E | -0,21228387 | 7,67031068 | -2,10030452 | 0,05360869 | 0,21393498 |
| 11186_at | RASSF1 | -0,33554995 | 6,86579345 | -2,10008544 | 0,05363062 | 0,21397817 |
| 9525_at | VPS4B | 0,24934899 | 8,78592595 | 2,09984388 | 0,05365481 | 0,21403036 |
| 2800_at | GOLGA1 | 0,25423042 | 7,38155947 | 2,09955131 | 0,05368413 | 0,21410296 |
| 29841_at | GRHL1 | 0,18115299 | 7,95397009 | 2,09928281 | 0,05371104 | 0,21416597 |
| 128866_at | CHMP4B | 0,23072404 | 10,5718761 | 2,09876153 | 0,05376333 | 0,21433011 |
| 4201_at | MEA1 | -0,24860468 | 10,4190129 | -2,09760363 | 0,05387965 | 0,21474938 |
| 8766_at | RAB11A | 0,16554121 | 9,42699415 | 2,09690285 | 0,05395016 | 0,21498594 |
| 8390_at | OR1G1 | -0,1956423 | 4,37629706 | -2,09652377 | 0,05398834 | 0,21509357 |
| 11319_at | ECD | -0,24719743 | 8,650379 | -2,09636711 | 0,05400413 | 0,21511197 |
| 401172_at | FLJ33360 | -0,20309764 | 3,79185758 | -2,09578366 | 0,05406295 | 0,21530026 |
| 89883_at | OR6W1P | -0,1648997 | 4,27832795 | -2,09565789 | 0,05407564 | 0,21530026 |
| 51501_at | C11orf73 | -0,15784148 | 6,37207354 | -2,09556583 | 0,05408493 | 0,21530026 |
| 5394_at | EXOSC10 | -0,21904057 | 8,69457506 | -2,09513166 | 0,05412876 | 0,21543022 |
| 84902_at | CEP89 | -0,19716477 | 6,26075911 | -2,09452713 | 0,05418984 | 0,21561272 |
| 118426_at | BORCS5 | -0,1398941 | 6,32012816 | -2,09445631 | 0,054197 | 0,21561272 |
| 85026_at | ARRDC1-AS1 | -0,13977114 | 7,16680677 | -2,09415262 | 0,05422771 | 0,21562359 |
| 54680_at | ZNHIT6 | -0,24979871 | 6,81426081 | -2,09413928 | 0,05422906 | 0,21562359 |

| | | | | | | |
|---|---|---|---|---|---|---|
| 1006_at | CDH8 | 0,20711988 | 4,1115141 | 2,09409725 | 0,05423331 | 0,21562359 |
| 59284_at | CACNG7 | -0,19230853 | 5,00840285 | -2,09386049 | 0,05425727 | 0,21565996 |
| 65251_at | ZNF649 | -0,21389863 | 6,1741934 | -2,09378558 | 0,05426485 | 0,21565996 |
| 27290_at | SPINK4 | -1,32775441 | 5,51639465 | -2,09296157 | 0,05434832 | 0,21594711 |
| 10014_at | HDAC5 | 0,24091089 | 6,58051255 | 2,09260485 | 0,05438449 | 0,21604627 |
| 57804_at | POLD4 | 0,39152733 | 8,86277497 | 2,09123047 | 0,05452405 | 0,21655603 |
| 10753_at | CAPN9 | 0,21876107 | 4,01333297 | 2,09070398 | 0,0545776 | 0,21669783 |
| 440087_at | SMCO3 | -0,1249474 | 3,21679262 | -2,09065829 | 0,05458225 | 0,21669783 |
| 2760_at | GM2A | 0,20986748 | 8,15126806 | 2,0904803 | 0,05460036 | 0,21672508 |
| 8444_at | DYRK3 | -0,17452524 | 6,15039538 | -2,08992554 | 0,05465686 | 0,21690465 |
| 283375_at | SLC39A5 | -0,25600035 | 6,06274735 | -2,08972088 | 0,05467772 | 0,21694273 |
| 343099_at | CCDC18 | -0,67046135 | 5,03092863 | -2,0891272 | 0,05473826 | 0,21711027 |
| 10746_at | MAP3K2 | 0,25504401 | 8,00146642 | 2,08895471 | 0,05475587 | 0,21711027 |
| 5644_at | PRSS1 | -0,17715446 | 3,76592131 | -2,08894869 | 0,05475648 | 0,21711027 |
| 90410_at | IFT20 | 0,41845138 | 8,30894208 | 2,0888192 | 0,0547697 | 0,21711027 |
| 23529_at | CLCF1 | -0,21923421 | 7,41276398 | -2,08868072 | 0,05478384 | 0,21711027 |
| 54583_at | EGLN1 | -0,21666982 | 7,84772764 | -2,08864416 | 0,05478757 | 0,21711027 |
| 4779_at | NFE2L1 | 0,19386159 | 9,35856284 | 2,08849529 | 0,05480278 | 0,21712035 |
| 6294_at | SAFB | -0,23438582 | 8,08696928 | -2,08832718 | 0,05481995 | 0,21712035 |
| 123722_at | FSD2 | -0,16151965 | 4,39713301 | -2,08828821 | 0,05482393 | 0,21712035 |
| 6470_at | SHMT1 | -0,83432222 | 8,04151103 | -2,08766301 | 0,05488785 | 0,21732881 |
| 10231_at | RCAN2 | 0,8910434 | 4,66671128 | 2,08647353 | 0,05500965 | 0,2177663 |
| 146395_at | GSG1L | -0,16128443 | 6,17016501 | -2,08576734 | 0,05508207 | 0,21800822 |
| 7072_at | TIA1 | 0,30284965 | 8,21775537 | 2,08522761 | 0,05513749 | 0,21807007 |
| 2564_at | GABRE | 0,7365285 | 8,91412628 | 2,08507987 | 0,05515267 | 0,21807007 |
| 8761_at | PABPC4 | -0,17551558 | 11,4862512 | -2,08493602 | 0,05516745 | 0,21807007 |
| 128553_at | TSHZ2 | 0,12914378 | 4,14977783 | 2,08474486 | 0,0551871 | 0,21807007 |
| 9643_at | MORF4L2 | -0,18216447 | 11,1949988 | -2,08458868 | 0,05520315 | 0,21807007 |
| 54948_at | MRPL16 | -0,4925429 | 8,86003296 | -2,08447598 | 0,05521475 | 0,21807007 |
| 22828_at | SCAF8 | 0,19810446 | 9,61371307 | 2,08444025 | 0,05521842 | 0,21807007 |
| 170685_at | NUDT10 | -0,17793963 | 5,65272605 | -2,08443408 | 0,05521906 | 0,21807007 |
| 6192_at | RPS4Y1 | -0,19323761 | 6,15418365 | -2,08430861 | 0,05523196 | 0,21807007 |
| 283914_at | LINC01566 | -0,201831 | 4,35805331 | -2,08430569 | 0,05523226 | 0,21807007 |
| 390064_at | OR51I2 | -0,18359824 | 4,33390887 | -2,08429326 | 0,05523354 | 0,21807007 |
| 64327_at | LMBR1 | -0,21048735 | 7,79761722 | -2,08429317 | 0,05523355 | 0,21807007 |
| 7050_at | TGIF1 | 0,14772256 | 7,58784211 | 2,08365788 | 0,05529895 | 0,21826393 |
| 284306_at | ZNF547 | 0,14568693 | 4,26301855 | 2,08359607 | 0,05530531 | 0,21826393 |
| 5757_at | PTMA | -0,22298169 | 5,52227343 | -2,08308298 | 0,05535819 | 0,21842785 |
| 10238_at | DCAF7 | 0,41512525 | 8,87564755 | 2,08281185 | 0,05538615 | 0,21843894 |
| 728431_at | LINC01137 | 0,16999239 | 6,67617691 | 2,08265886 | 0,05540193 | 0,21843894 |
| 101928099_a | PCAT7 | 0,68388961 | 5,74298626 | 2,08262809 | 0,0554051 | 0,21843894 |
| 201514_at | ZNF584 | -0,48730894 | 6,58580352 | -2,08261596 | 0,05540636 | 0,21843894 |
| 11345_at | GABARAPL2 | 0,12609417 | 10,367883 | 2,08201254 | 0,05546865 | 0,21863978 |
| 197407_at | ZNF48 | -0,14010231 | 6,59615869 | -2,08177438 | 0,05549325 | 0,21869201 |
| 222234_at | FAM185A | -0,33130026 | 4,21373457 | -2,08161605 | 0,05550961 | 0,21871174 |
| 9465_at | AKAP7 | 0,17794213 | 5,68599689 | 2,08147279 | 0,05552442 | 0,21872535 |
| 101929378_a | LOC10192937 | -0,13756839 | 3,72802954 | -2,08127005 | 0,05554539 | 0,2187632 |
| 23469_at | PHF3 | 0,19185856 | 7,11400533 | 2,08018472 | 0,05565773 | 0,21916086 |
| 6464_at | SHC1 | -0,41961658 | 10,2912887 | -2,07989717 | 0,05568754 | 0,21923339 |
| 23645_at | PPP1R15A | 0,25604674 | 7,58635564 | 2,07957674 | 0,05572076 | 0,21931879 |

| | | | | | | |
|---|---|---|---|---|---|---|
| 285768_at | LOC285768 | -0,15285235 | 4,04755642 | -2,07946841 | 0,055732 | 0,21931879 |
| 9880_at | ZBTB39 | 0,14804314 | 6,86759379 | 2,07907435 | 0,05577289 | 0,21943488 |
| 1145_at | CHRNE | -0,25912906 | 4,22861147 | -2,07886345 | 0,05579479 | 0,21947621 |
| 149563_at | C1orf64 | -0,16713291 | 4,73347418 | -2,07859006 | 0,05582318 | 0,21954308 |
| 54521_at | WDR44 | 0,21136283 | 7,05096542 | 2,07780394 | 0,05590491 | 0,21980016 |
| 161176_at | SYNE3 | -0,2082715 | 4,64382259 | -2,07767372 | 0,05591846 | 0,21980016 |
| 10942_at | PRSS21 | -0,25310602 | 4,74927842 | -2,07763218 | 0,05592278 | 0,21980016 |
| 10690_at | FUT9 | 0,6084797 | 3,81980917 | 2,07739225 | 0,05594776 | 0,2198336 |
| 254240_at | BPIFC | -0,19911052 | 3,77134392 | -2,07733117 | 0,05595412 | 0,2198336 |
| 64750_at | SMURF2 | 0,21718025 | 8,10780258 | 2,07699137 | 0,05598951 | 0,21992214 |
| 100287896_a | LOC10028789 | 0,3254019 | 6,84754165 | 2,07689558 | 0,05599949 | 0,21992214 |
| 2516_at | NR5A1 | -0,24201629 | 4,82493205 | -2,07676908 | 0,05601267 | 0,21992907 |
| 100506144_a | ZMYM6NB | 0,30087437 | 8,99495912 | 2,07652639 | 0,05603797 | 0,21998356 |
| 80139_at | ZNF703 | 0,26095803 | 7,84855281 | 2,07634806 | 0,05605656 | 0,21999006 |
| 102800314_a | MIR7515HG | -0,2065716 | 3,52492975 | -2,07629148 | 0,05606246 | 0,21999006 |
| 2720_at | GLB1 | 0,37268356 | 9,58010638 | 2,07490263 | 0,0562075 | 0,22051426 |
| 121665_at | SPPL3 | 0,14492448 | 8,89881137 | 2,07450517 | 0,05624907 | 0,22063241 |
| 176_at | ACAN | -0,16491432 | 4,92760406 | -2,07383228 | 0,05631951 | 0,22086373 |
| 3344_at | FOXN2 | -0,35504878 | 5,73816023 | -2,07284509 | 0,05642299 | 0,22122454 |
| 102724368_a | CELSR3-AS1 | -0,34449305 | 5,43854955 | -2,07247059 | 0,0564623 | 0,22124283 |
| 27076_at | LYPD3 | 0,35251862 | 9,54840998 | 2,07246813 | 0,05646256 | 0,22124283 |
| 11227_at | GALNT5 | -0,22549217 | 6,79894339 | -2,07238254 | 0,05647155 | 0,22124283 |
| 25775_at | C22orf24 | -0,24370759 | 4,43931257 | -2,07236294 | 0,0564736 | 0,22124283 |
| 10998_at | SLC27A5 | -0,39059864 | 6,47711765 | -2,0710744 | 0,05660906 | 0,2217284 |
| 6035_at | RNASE1 | 0,59398299 | 6,00825344 | 2,07079678 | 0,05663828 | 0,22179776 |
| 122876_at | GPHB5 | -0,17113162 | 4,66041329 | -2,06996671 | 0,05672574 | 0,22209511 |
| 5433_at | POLR2D | -0,40095799 | 7,95170279 | -2,06903338 | 0,05682423 | 0,22243549 |
| 400657_at | LINC00909 | 0,24560823 | 6,13814385 | 2,06872217 | 0,0568571 | 0,22251896 |
| 128486_at | FITM2 | -0,32196319 | 6,91223541 | -2,06809826 | 0,05692306 | 0,22273185 |
| 57106_at | NAT14 | 0,38232755 | 7,02679443 | 2,06765552 | 0,05696991 | 0,22286989 |
| 23335_at | WDR7 | 0,19264354 | 7,12074611 | 2,06685562 | 0,05705464 | 0,22315604 |
| 55777_at | MBD5 | 0,91599773 | 5,60056462 | 2,06673156 | 0,05706779 | 0,22316217 |
| 51506_at | UFC1 | 0,20737234 | 10,7981001 | 2,06656695 | 0,05708525 | 0,22317966 |
| 4289_at | MKLN1 | 0,26794579 | 7,06782731 | 2,06634627 | 0,05710865 | 0,22317966 |
| 100505738_a | MIR4458HG | -0,36508639 | 5,55484622 | -2,06633944 | 0,05710938 | 0,22317966 |
| 100131211_a | NEMP2 | -0,25051196 | 5,64269604 | -2,06618006 | 0,05712629 | 0,22317966 |
| 7299_at | TYR | -0,17013131 | 3,2998891 | -2,06614328 | 0,05713019 | 0,22317966 |
| 116496_at | FAM129A | -0,36822109 | 6,55430687 | -2,06539708 | 0,05720944 | 0,22344391 |
| 54187_at | NANS | -0,17843294 | 8,93778687 | -2,06465348 | 0,0572885 | 0,22368995 |
| 23324_at | MAN2B2 | 0,56614662 | 8,48029476 | 2,06455723 | 0,05729875 | 0,22368995 |
| 115196_at | ZNF554 | 0,30877044 | 5,442723 | 2,06447715 | 0,05730727 | 0,22368995 |
| 26995_at | TRUB2 | -0,16960277 | 8,23352717 | -2,06415404 | 0,05734167 | 0,22373494 |
| 10428_at | CFDP1 | 0,12910723 | 7,41113116 | 2,06415067 | 0,05734202 | 0,22373494 |
| 80830_at | APOL6 | 0,25839774 | 7,72580468 | 2,06399222 | 0,0573589 | 0,22374705 |
| 54433_at | GAR1 | -0,51398119 | 8,51573215 | -2,06390344 | 0,05736836 | 0,22374705 |
| 4781_at | NFIB | -0,29495409 | 7,09551742 | -2,06356601 | 0,05740432 | 0,22384197 |
| 219670_at | ENKUR | -0,18754169 | 3,32083715 | -2,062966 | 0,05746831 | 0,22402773 |
| 9230_at | RAB11B | 0,12100244 | 6,93757061 | 2,06290127 | 0,05747521 | 0,22402773 |
| 1833_at | EPYC | 0,42593585 | 3,25359305 | 2,06262687 | 0,05750451 | 0,22405892 |
| 26147_at | PHF19 | -0,8314777 | 7,96417556 | -2,06260839 | 0,05750648 | 0,22405892 |

| | | | | | | |
|---|---|---|---|---|---|---|
| 23304_at | UBR2 | 0,25049445 | 7,2610265 | 2,06244497 | 0,05752393 | 0,22408159 |
| 57498_at | KIDINS220 | 0,27094879 | 8,29295319 | 2,06233573 | 0,0575356 | 0,22408172 |
| 25950_at | RWDD3 | 0,26668942 | 7,41077365 | 2,0614086 | 0,05763471 | 0,22442237 |
| 100506124_a | LOC10050612 | 0,14933963 | 3,27240721 | 2,06119401 | 0,05765767 | 0,2244664 |
| 54825_at | CDHR2 | -0,21025636 | 5,98787226 | -2,06104917 | 0,05767318 | 0,22448139 |
| 11251_at | PTGDR2 | -0,17273759 | 5,89421683 | -2,06064521 | 0,05771644 | 0,22460439 |
| 404203_at | SPINK6 | -0,30767118 | 3,95389899 | -2,0604402 | 0,0577384 | 0,22464449 |
| 55761_at | TTC17 | 0,34377089 | 7,59497145 | 2,06023604 | 0,05776029 | 0,22468425 |
| 3620_at | IDO1 | 1,18367639 | 6,16330588 | 2,05969137 | 0,0578187 | 0,22486608 |
| 27429_at | HTRA2 | -0,1388873 | 8,31981212 | -2,05943765 | 0,05784594 | 0,22492657 |
| 9670_at | IPO13 | -0,18132608 | 7,86786285 | -2,05905312 | 0,05788723 | 0,22501055 |
| 3777_at | KCNK3 | -0,20068917 | 4,78852596 | -2,05898283 | 0,05789478 | 0,22501055 |
| 319089_at | TTC6 | 0,29502174 | 3,38640174 | 2,05891026 | 0,05790258 | 0,22501055 |
| 80208_at | SPG11 | 0,35361637 | 6,6675631 | 2,05855096 | 0,05794119 | 0,2251152 |
| 1593_at | CYP27A1 | -0,20104027 | 5,68423834 | -2,05843883 | 0,05795325 | 0,22511664 |
| 7940_at | LST1 | -0,12696879 | 4,95685064 | -2,05774857 | 0,05802752 | 0,22535969 |
| 2330_at | FMO5 | 0,11654871 | 3,80811287 | 2,05734189 | 0,05807131 | 0,22548432 |
| 55677_at | IWS1 | -0,17647513 | 8,32365761 | -2,05700273 | 0,05810786 | 0,22558077 |
| 257000_at | TINCR | 0,51509601 | 6,92777203 | 2,05544784 | 0,0582757 | 0,22615877 |
| 7941_at | PLA2G7 | 0,1140185 | 2,88912085 | 2,05535408 | 0,05828583 | 0,22615877 |
| 8804_at | CREG1 | 0,25641445 | 11,000821 | 2,05529724 | 0,05829197 | 0,22615877 |
| 196951_at | FAM227B | 0,15154383 | 2,95567821 | 2,05492901 | 0,0583318 | 0,2262677 |
| 60673_at | ATG101 | -0,2435076 | 8,49688476 | -2,05421236 | 0,05840937 | 0,22652298 |
| 53342_at | IL17D | -0,21145423 | 5,03746109 | -2,05291882 | 0,05854962 | 0,22702121 |
| 2104_at | ESRRG | 0,61958443 | 4,7207631 | 2,05251857 | 0,05859308 | 0,227144 |
| 51280_at | GOLM1 | 0,32294742 | 8,83901646 | 2,05200607 | 0,05864878 | 0,22731416 |
| 79901_at | CYBRD1 | 0,63447789 | 7,93081269 | 2,05176 | 0,05867553 | 0,22737211 |
| 130074_at | FAM168B | -0,31182187 | 8,89165296 | -2,05165068 | 0,05868742 | 0,22737245 |
| 3034_at | HAL | -0,13387924 | 4,41629903 | -2,05136568 | 0,05871843 | 0,22744685 |
| 5372_at | PMM1 | 0,35757729 | 7,0905637 | 2,05111626 | 0,05874558 | 0,22750627 |
| 101927379_a | LOC10192737 | 0,22742696 | 3,77917863 | 2,05077134 | 0,05878315 | 0,227606 |
| 1735_at | DIO3 | -0,17914592 | 5,68053585 | -2,05040907 | 0,05882263 | 0,22771309 |
| 3566_at | IL4R | -0,13915736 | 8,80366993 | -2,05024049 | 0,05884101 | 0,22773847 |
| 154661_at | RUNDC3B | 0,30815534 | 3,83881304 | 2,0500362 | 0,05886329 | 0,22777894 |
| 100130348_a | TMEM262 | -0,29634696 | 5,19678121 | -2,04959876 | 0,05891102 | 0,22790427 |
| 80854_at | SETD7 | -0,30136382 | 7,35468709 | -2,04945187 | 0,05892706 | 0,22790427 |
| 153443_at | SRFBP1 | -0,36265751 | 7,02929636 | -2,04931574 | 0,05894193 | 0,22790427 |
| 28984_at | RGCC | -0,25921447 | 3,44567358 | -2,04926108 | 0,0589479 | 0,22790427 |
| 115704_at | EVI5L | 0,12660642 | 6,06075891 | 2,04919756 | 0,05895483 | 0,22790427 |
| #N/A | CAND1,11 | -0,2163503 | 4,02171952 | -2,04890971 | 0,05898628 | 0,2279801 |
| 160_at | AP2A1 | 0,15065976 | 7,45628338 | 2,04818643 | 0,05906538 | 0,22824 |
| 5205_at | ATP8B1 | 0,29709869 | 10,2294116 | 2,04783181 | 0,05910419 | 0,22834418 |
| 84545_at | MRPL43 | -0,10624989 | 8,32972584 | -2,04754005 | 0,05913615 | 0,22842182 |
| 84792_at | FAM220A | 0,20355299 | 9,52814606 | 2,04700059 | 0,05919527 | 0,2285459 |
| 348801_at | LNP1 | -0,22808651 | 6,07415687 | -2,04696059 | 0,05919965 | 0,2285459 |
| 28672_at | TRAV12-3 | -0,17841902 | 2,83998299 | -2,046868 | 0,05920981 | 0,2285459 |
| 6753_at | SSTR3 | -0,22630132 | 6,04388034 | -2,04674085 | 0,05922375 | 0,2285459 |
| 146760_at | RTN4RL1 | -0,25475523 | 4,61752572 | -2,04667921 | 0,05923052 | 0,2285459 |
| 762_at | CA4 | -0,16372819 | 4,73399118 | -2,0465977 | 0,05923946 | 0,2285459 |
| 2533_at | FYB | 0,79961539 | 6,23119424 | 2,04643797 | 0,05925699 | 0,22855922 |

| | | | | | | |
|---|---|---|---|---|---|---|
| 51225_at | ABI3 | -0,23252266 | 5,89268795 | -2,04635001 | 0,05926664 | 0,22855922 |
| 11253_at | MAN1B1 | 0,15276439 | 8,52382871 | 2,04589916 | 0,05931615 | 0,22870435 |
| 162989_at | DEDD2 | -0,16735928 | 8,54869577 | -2,0446884 | 0,05944929 | 0,22917182 |
| 79673_at | ZNF329 | 0,6005093 | 5,99764409 | 2,04444478 | 0,05947611 | 0,22922934 |
| 9794_at | MAML1 | 0,11309456 | 9,41653648 | 2,04432309 | 0,05948951 | 0,22923513 |
| 55505_at | NOP10 | -0,28379032 | 10,9194736 | -2,04389471 | 0,05953671 | 0,22935179 |
| 54432_at | YIPF1 | 0,40024671 | 7,76391017 | 2,04383223 | 0,0595436 | 0,22935179 |
| 94239_at | H2AFV | -0,19652634 | 9,72309754 | -2,04292735 | 0,05964342 | 0,22969037 |
| 55308_at | DDX19A | -0,19579295 | 8,11794822 | -2,04259277 | 0,05968037 | 0,22978673 |
| 254013_at | METTL20 | 0,14326779 | 3,87448661 | 2,04218137 | 0,05972583 | 0,22989238 |
| 596_at | BCL2 | -0,16499749 | 4,94137202 | -2,04183401 | 0,05976424 | 0,22989238 |
| 101927018_a | LOC10192701 | -0,13262025 | 3,01302091 | -2,04182408 | 0,05976534 | 0,22989238 |
| 26119_at | LDLRAP1 | -0,26496943 | 7,522498 | -2,04181087 | 0,0597668 | 0,22989238 |
| 51304_at | ZDHHC3 | -0,18170572 | 7,68518648 | -2,04180472 | 0,05976748 | 0,22989238 |
| 823_at | CAPN1 | 0,26708441 | 10,0440899 | 2,04147276 | 0,05980421 | 0,22994941 |
| 9266_at | CYTH2 | 0,28819136 | 8,38497075 | 2,04145494 | 0,05980619 | 0,22994941 |
| 101928682_a | LOC10192868 | -0,1973877 | 4,81973339 | -2,04108144 | 0,05984754 | 0,23006249 |
| 402055_at | SRRD | -0,2212182 | 8,04343548 | -2,04066097 | 0,05989412 | 0,23019563 |
| 81704_at | DOCK8 | 0,72297995 | 6,80791715 | 2,04041754 | 0,05992111 | 0,2302534 |
| 84456_at | L3MBTL3 | 0,27304335 | 7,2621697 | 2,0399664 | 0,05997115 | 0,23039364 |
| 163859_at | SDE2 | -0,20588203 | 7,14161653 | -2,03987289 | 0,05998152 | 0,23039364 |
| 100996702_a | LINC01356 | -0,31639874 | 5,99688661 | -2,03920547 | 0,06005563 | 0,23050664 |
| 10097_at | ACTR2 | 0,18210846 | 10,9921612 | 2,03920342 | 0,06005586 | 0,23050664 |
| 11147_at | HHLA3 | 0,58687675 | 6,76106033 | 2,03920201 | 0,06005602 | 0,23050664 |
| 1413_at | CRYBA4 | -0,28722609 | 4,68545728 | -2,03917248 | 0,0600593 | 0,23050664 |
| 401491_at | VLDLR-AS1 | -0,20475062 | 4,12828598 | -2,0390692 | 0,06007078 | 0,23050664 |
| 339803_at | LOC339803 | -0,16362668 | 4,03181727 | -2,03837481 | 0,06014799 | 0,23075696 |
| 3577_at | CXCR1 | -0,21934648 | 6,04651056 | -2,03804955 | 0,06018419 | 0,23084302 |
| 9801_at | MRPL19 | -0,23686282 | 7,56171187 | -2,03795792 | 0,06019439 | 0,23084302 |
| 84890_at | ADO | -0,27484542 | 8,41092423 | -2,03740587 | 0,06025588 | 0,23102575 |
| 151188_at | ARL6IP6 | -0,20191238 | 7,45551136 | -2,03731487 | 0,06026602 | 0,23102575 |
| 10844_at | TUBGCP2 | -0,12758338 | 8,20953054 | -2,03700471 | 0,0603006 | 0,23111232 |
| 55737_at | VPS35 | -0,13763522 | 10,4196127 | -2,03663907 | 0,06034139 | 0,23122264 |
| 55332_at | DRAM1 | 1,18099363 | 8,8669697 | 2,03619749 | 0,06039068 | 0,2313655 |
| 55709_at | KBTBD4 | 0,19338824 | 7,49500825 | 2,03583303 | 0,0604314 | 0,23145349 |
| 9101_at | USP8 | 0,15939475 | 7,94561359 | 2,03577678 | 0,06043768 | 0,23145349 |
| 283159_at | OR8D1 | -0,20570371 | 3,91900922 | -2,03535531 | 0,0604848 | 0,23154536 |
| 10565_at | ARFGEF1 | 0,34950187 | 9,03369393 | 2,03534714 | 0,06048571 | 0,23154536 |
| 10289_at | EIF1B | -0,20190352 | 9,31041839 | -2,03493412 | 0,06053192 | 0,23167281 |
| 3359_at | HTR3A | -0,18474025 | 4,638275 | -2,03483458 | 0,06054306 | 0,23167281 |
| 120939_at | TMEM52B | 0,79015383 | 5,09971888 | 2,03392929 | 0,06064446 | 0,2320019 |
| 3594_at | IL12RB1 | -0,18195477 | 5,05602199 | -2,03376715 | 0,06066264 | 0,2320019 |
| 126526_at | C19orf47 | -0,1352559 | 6,07617851 | -2,03374441 | 0,06066519 | 0,2320019 |
| 51647_at | FAM96B | -0,2556844 | 10,3090587 | -2,03345195 | 0,06069799 | 0,23208128 |
| 7030_at | TFE3 | 0,28688592 | 7,71278794 | 2,03284289 | 0,06076636 | 0,23226586 |
| 1212_at | CLTB | -0,26388483 | 9,56776663 | -2,0327058 | 0,06078176 | 0,23226586 |
| 4259_at | MGST3 | 0,35130908 | 9,89864921 | 2,03269972 | 0,06078244 | 0,23226586 |
| 57017_at | COQ9 | -0,24268705 | 9,40403346 | -2,0325125 | 0,06080348 | 0,23230015 |
| 4161_at | MC5R | -0,22790989 | 3,95922704 | -2,03232834 | 0,06082417 | 0,2323237 |
| 4193_at | MDM2 | -0,13470938 | 6,66042176 | -2,03216326 | 0,06084273 | 0,2323237 |

| | | | | | | |
|---|---|---|---|---|---|---|
| 23433_at | RHOQ | 0,37616272 | 5,60682431 | 2,03209514 | 0,06085039 | 0,2323237 |
| 150677_at | OTOS | -0,19213803 | 5,83277992 | -2,03202852 | 0,06085788 | 0,2323237 |
| 25862_at | USP49 | 0,09625533 | 5,29828701 | 2,0319075 | 0,06087149 | 0,23232962 |
| 7128_at | TNFAIP3 | 0,6575933 | 6,47013274 | 2,03152962 | 0,06091401 | 0,23240259 |
| 8514_at | KCNAB2 | -0,16046646 | 6,23262471 | -2,03152316 | 0,06091474 | 0,23240259 |
| 10467_at | ZNHIT1 | 0,10410834 | 9,777189 | 2,03037592 | 0,061044 | 0,2328496 |
| 9422_at | ZNF264 | 0,29185223 | 7,02444377 | 2,03008124 | 0,06107724 | 0,23293028 |
| 255738_at | PCSK9 | -0,85961038 | 6,17894773 | -2,02980322 | 0,06110861 | 0,23300381 |
| 64207_at | IRF2BPL | 0,22522141 | 10,7594023 | 2,02954693 | 0,06113755 | 0,23306801 |
| 79090_at | TRAPPC6A | 0,71504369 | 7,73488992 | 2,02909357 | 0,06118877 | 0,23321712 |
| 60493_at | FASTKD5 | -0,16012944 | 8,30855916 | -2,02847661 | 0,06125854 | 0,23343684 |
| 23198_at | PSME4 | -0,16778663 | 8,62325361 | -2,02836574 | 0,06127108 | 0,23343846 |
| 1317_at | SLC31A1 | -0,21977722 | 9,46243978 | -2,02801733 | 0,06131052 | 0,23349147 |
| 51157_at | ZNF580 | 0,34134421 | 8,17234964 | 2,02792977 | 0,06132043 | 0,23349147 |
| 4534_at | MTM1 | 0,24271888 | 5,93895279 | 2,02792161 | 0,06132136 | 0,23349147 |
| 6565_at | SLC15A2 | 0,77207852 | 6,61450482 | 2,02690922 | 0,0614361 | 0,23388215 |
| 26220_at | DGCR5 | -0,23415938 | 5,94354172 | -2,02641449 | 0,06149225 | 0,23403044 |
| 181_at | AGRP | -0,20073264 | 6,5060082 | -2,02635189 | 0,06149936 | 0,23403044 |
| 285313_at | IGSF10 | 0,2149932 | 4,69593237 | 2,0260972 | 0,06152828 | 0,23407988 |
| 9723_at | SEMA3E | 0,85332914 | 5,10544393 | 2,02602352 | 0,06153665 | 0,23407988 |
| 9759_at | HDAC4 | -0,17454344 | 6,53547806 | -2,02591658 | 0,0615488 | 0,23407988 |
| 7280_at | TUBB2A | -0,62139004 | 8,56634181 | -2,02566767 | 0,06157709 | 0,23414124 |
| 4803_at | NGF | -0,21813317 | 5,83956209 | -2,02553979 | 0,06159163 | 0,2341503 |
| 283171_at | C11orf44 | -0,18877502 | 4,54360618 | -2,02534505 | 0,06161378 | 0,23418828 |
| 55006_at | TRMT61B | -0,236511 | 7,94158473 | -2,0251166 | 0,06163976 | 0,23424083 |
| 100505592_a | GAPLINC | -0,20185744 | 3,64562175 | -2,02466535 | 0,06169113 | 0,23436679 |
| 253982_at | ASPHD1 | -0,22056724 | 5,91271416 | -2,02461166 | 0,06169724 | 0,23436679 |
| 11160_at | ERLIN2 | 0,24065317 | 7,64450772 | 2,02405154 | 0,06176106 | 0,23456295 |
| 56271_at | BEX4 | 1,11324817 | 6,15041287 | 2,02371682 | 0,06179922 | 0,23466163 |
| 57572_at | DOCK6 | 0,17070312 | 6,84228462 | 2,02282012 | 0,06190157 | 0,23500394 |
| 197335_at | WDR90 | -0,34406923 | 7,14001587 | -2,02237478 | 0,06195246 | 0,23515078 |
| 10098_at | TSPAN5 | 0,23063701 | 6,22014996 | 2,02127261 | 0,06207857 | 0,23554544 |
| 388948_at | LINC01118 | -0,25341732 | 4,91973687 | -2,02125231 | 0,06208089 | 0,23554544 |
| 84433_at | CARD11 | -0,22486162 | 7,98531826 | -2,02098923 | 0,06211103 | 0,23561337 |
| 10010_at | TANK | 0,37631399 | 7,66336508 | 2,02075314 | 0,06213808 | 0,2356696 |
| 9820_at | CUL7 | 0,15596466 | 6,99993823 | 2,0202485 | 0,06219595 | 0,23584264 |
| 57109_at | REXO4 | -0,30880486 | 7,58123915 | -2,02002963 | 0,06222106 | 0,2358478 |
| 29928_at | TIMM22 | -0,30917145 | 7,94214026 | -2,02002323 | 0,0622218 | 0,2358478 |
| 2267_at | FGL1 | -0,15553854 | 4,34717667 | -2,01991351 | 0,06223439 | 0,23584912 |
| 23224_at | SYNE2 | 0,24514175 | 6,74806637 | 2,01888039 | 0,06235308 | 0,23625244 |
| 55284_at | UBE2W | 0,12964316 | 6,0968941 | 2,01825556 | 0,06242497 | 0,23647829 |
| 339324_at | ZNF260 | 0,39653447 | 7,92307343 | 2,01782445 | 0,06247461 | 0,23659178 |
| 6975_at | TECTB | -0,19487575 | 3,54965139 | -2,01762566 | 0,06249751 | 0,23659178 |
| 1962_at | EHHADH | 0,33349097 | 6,10405673 | 2,01761952 | 0,06249822 | 0,23659178 |
| 53407_at | STX18 | -0,12040665 | 7,90276843 | -2,01756889 | 0,06250405 | 0,23659178 |
| 26207_at | PITPNC1 | 0,24802392 | 6,67296423 | 2,01721421 | 0,06254494 | 0,23670003 |
| 26958_at | COPG2 | 0,179741 | 6,64175692 | 2,01664284 | 0,06261085 | 0,23684742 |
| 83955_at | NACAP1 | -0,18956646 | 9,6328377 | -2,0165204 | 0,06262499 | 0,23684742 |
| 353149_at | TBC1D26 | -0,23494389 | 5,41090272 | -2,01643493 | 0,06263485 | 0,23684742 |
| 9771_at | RAPGEF5 | -0,28345691 | 4,4833782 | -2,01640203 | 0,06263865 | 0,23684742 |

| | | | | | | |
|---|---|---|---|---|---|---|
| 28667_at | TRAV16 | -0,20520441 | 4,13967666 | -2,01634392 | 0,06264536 | 0,23684742 |
| 3783_at | KCNN4 | 0,24861299 | 8,98053558 | 2,01619894 | 0,06266211 | 0,23686424 |
| 29949_at | IL19 | -0,22455002 | 4,43798965 | -2,01578133 | 0,06271036 | 0,23700013 |
| 157556_at | BAALC-AS2 | -0,15592742 | 4,42384392 | -2,01542488 | 0,06275157 | 0,23708341 |
| 79659_at | DYNC2H1 | 0,14298909 | 3,69208058 | 2,01526709 | 0,06276983 | 0,23708341 |
| 114043_at | TSPEAR-AS2 | -0,23086361 | 5,12269493 | -2,0151631 | 0,06278186 | 0,23708341 |
| 101929210_a | LOC10192921 | 0,15813848 | 3,10188629 | 2,01512807 | 0,06278591 | 0,23708341 |
| 84197_at | POMK | -0,18388886 | 6,71025715 | -2,01496074 | 0,06280528 | 0,23708341 |
| 3297_at | HSF1 | -0,27004246 | 7,35506171 | -2,01488448 | 0,0628141 | 0,23708341 |
| 81_at | ACTN4 | 0,18461216 | 8,95502915 | 2,01482746 | 0,0628207 | 0,23708341 |
| 5991_at | RFX3 | 0,20628181 | 3,90358841 | 2,01473978 | 0,06283086 | 0,23708341 |
| 92140_at | MTDH | -0,43398751 | 9,64856133 | -2,01435625 | 0,06287528 | 0,23720457 |
| 79886_at | CAAP1 | -0,27109808 | 7,50215987 | -2,0142247 | 0,06289052 | 0,23721562 |
| 3703_at | STT3A | -0,14010266 | 10,9384697 | -2,01388653 | 0,06292972 | 0,23731701 |
| 100498859_a | LINC00881 | -0,28434153 | 4,14575063 | -2,01324009 | 0,06300472 | 0,23753328 |
| 50859_at | SPOCK3 | 0,37418999 | 3,49673977 | 2,0131797 | 0,06301174 | 0,23753328 |
| 57001_at | SDHAF3 | -0,46476584 | 7,34694737 | -2,01300565 | 0,06303194 | 0,23756297 |
| 23530_at | NNT | 0,21658447 | 7,70269214 | 2,01251745 | 0,06308866 | 0,23765299 |
| 58472_at | SQRDL | 0,49961541 | 8,19691545 | 2,01248329 | 0,06309263 | 0,23765299 |
| 114907_at | FBXO32 | 0,72060746 | 5,13941466 | 2,0124815 | 0,06309284 | 0,23765299 |
| 9013_at | TAF1C | 0,1289957 | 7,28088071 | 2,0111443 | 0,06324844 | 0,23817949 |
| 441476_at | C9orf173 | -0,3398079 | 5,97439422 | -2,01106783 | 0,06325735 | 0,23817949 |
| 84947_at | SERAC1 | 0,22472949 | 6,75508457 | 2,00940766 | 0,06345104 | 0,2388621 |
| 55776_at | SAYSD1 | 0,12786063 | 7,69307424 | 2,00808268 | 0,06360601 | 0,23936753 |
| 5630_at | PRPH | -0,21973002 | 6,26332708 | -2,00804733 | 0,06361015 | 0,23936753 |
| 200162_at | SPAG17 | 0,96543505 | 4,65148905 | 2,00793417 | 0,06362341 | 0,23937064 |
| 3818_at | KLKB1 | -0,16846796 | 4,06332162 | -2,00767049 | 0,0636543 | 0,23944009 |
| 5515_at | PPP2CA | -0,18080745 | 11,0326175 | -2,00750065 | 0,0636742 | 0,2394682 |
| 8722_at | CTSF | 0,26015662 | 5,98935272 | 2,00732078 | 0,06369528 | 0,23950074 |
| 79834_at | PEAK1 | 0,12204716 | 6,55948439 | 2,00711184 | 0,06371979 | 0,2395461 |
| 611_at | OPN1SW | -0,20379839 | 3,79973074 | -2,0068672 | 0,06374849 | 0,23958332 |
| 10479_at | SLC9A6 | 0,28908527 | 8,44632979 | 2,00681542 | 0,06375456 | 0,23958332 |
| 90529_at | STPG1 | 0,32675004 | 6,79872094 | 2,00609657 | 0,06383896 | 0,23972989 |
| 196913_at | LINC01599 | -0,20814323 | 5,53218193 | -2,00607474 | 0,06384153 | 0,23972989 |
| 101928797_a | LOC10192879 | -0,16317263 | 3,30105773 | -2,00602977 | 0,06384681 | 0,23972989 |
| 10232_at | MSLN | -0,22307627 | 4,25949527 | -2,00597571 | 0,06385316 | 0,23972989 |
| 83729_at | INHBE | -0,19519973 | 3,70624782 | -2,00595337 | 0,06385579 | 0,23972989 |
| 9538_at | EI24 | -0,28320094 | 10,2948129 | -2,00574897 | 0,06387981 | 0,23977335 |
| 84328_at | LZIC | -0,20297256 | 7,94532304 | -2,00535856 | 0,06392572 | 0,23989893 |
| 29954_at | POMT2 | -0,23145447 | 7,04979255 | -2,00516348 | 0,06394867 | 0,23993831 |
| 55177_at | RMDN3 | -0,21289398 | 8,77618132 | -2,00419822 | 0,06406235 | 0,24027406 |
| 9071_at | CLDN10 | -0,22096023 | 3,82667569 | -2,00419178 | 0,06406311 | 0,24027406 |
| 83744_at | ZNF484 | 0,17112873 | 4,85464122 | 2,00358599 | 0,06413454 | 0,24049516 |
| 401321_at | LINC00997 | -0,13512624 | 4,66938218 | -2,00317294 | 0,06418329 | 0,24063112 |
| 221823_at | PRPS1L1 | -0,19324879 | 3,81963844 | -2,00294007 | 0,06421079 | 0,24068737 |
| 85455_at | DISP2 | -0,41180315 | 6,63301488 | -2,00266979 | 0,06424272 | 0,24069666 |
| 4146_at | MATN1 | -0,18663174 | 4,71447314 | -2,00264074 | 0,06424615 | 0,24069666 |
| 80115_at | BAIAP2L2 | -0,18380659 | 5,87142594 | -2,00260181 | 0,06425075 | 0,24069666 |
| 149095_at | DCST1 | -0,16393757 | 4,81286175 | -2,00241359 | 0,064273 | 0,24073318 |
| 2938_at | GSTA1 | 1,35173358 | 4,69560231 | 2,00197744 | 0,06432458 | 0,24086093 |

| | | | | | | |
|---|---|---|---|---|---|---|
| 152098_at | ZCWPW2 | 0,17051502 | 3,03063385 | 2,00191374 | 0,06433211 | 0,24086093 |
| 89927_at | C16orf45 | 0,30846054 | 6,91205939 | 2,0010207 | 0,06443785 | 0,24120992 |
| 595_at | CCND1 | -0,87202134 | 9,85696372 | -2,00075911 | 0,06446885 | 0,24127909 |
| 6810_at | STX4 | 0,11853078 | 8,61534917 | 2,00022542 | 0,06453214 | 0,24146905 |
| 51232_at | CRIM1 | 0,35119064 | 8,61293507 | 2,00003127 | 0,06455518 | 0,24150835 |
| 89801_at | PPP1R3F | -0,18526619 | 5,24341784 | -1,99970333 | 0,06459412 | 0,24160708 |
| 27158_at | NDOR1 | -0,13350849 | 7,1064389 | -1,99883321 | 0,06469752 | 0,24194686 |
| 23169_at | SLC35D1 | 0,29638826 | 7,89435202 | 1,99871649 | 0,0647114 | 0,24195181 |
| 81493_at | SYNC | 0,22438985 | 4,97989666 | 1,99763653 | 0,06483998 | 0,2423855 |
| 4928_at | NUP98 | -0,29560108 | 7,94490836 | -1,99697555 | 0,06491878 | 0,24263301 |
| 9338_at | TCEAL1 | 0,29219184 | 8,99555812 | 1,99617553 | 0,06501429 | 0,24294282 |
| 285368_at | PRRT3 | 0,20110242 | 5,69724124 | 1,99568256 | 0,0650732 | 0,2431158 |
| 254228_at | FAM26E | -0,19293273 | 3,92825895 | -1,995347 | 0,06511333 | 0,24321854 |
| 339230_at | CCDC137 | -0,24712299 | 8,07557425 | -1,99501907 | 0,06515257 | 0,24327339 |
| 23587_at | ELP5 | -0,19451736 | 8,26748042 | -1,99501321 | 0,06515327 | 0,24327339 |
| 58489_at | ABHD17C | -0,25646237 | 9,25347122 | -1,9946097 | 0,06520158 | 0,24340659 |
| 202915_at | TMEM184A | 0,17661222 | 7,79018667 | 1,99405949 | 0,06526751 | 0,2436055 |
| 140453_at | MUC17 | -0,23365211 | 3,8165252 | -1,99382554 | 0,06529556 | 0,24366299 |
| 689_at | BTF3 | -0,28869544 | 11,1501225 | -1,99317558 | 0,06537355 | 0,24386581 |
| 8368_at | HIST1H4L | -0,24012041 | 6,41286211 | -1,99316158 | 0,06537523 | 0,24386581 |
| 375_at | ARF1 | -0,1277344 | 10,9252428 | -1,99304883 | 0,06538877 | 0,24386909 |
| 10761_at | PLAC1 | -0,21048407 | 4,65325581 | -1,9924086 | 0,06546569 | 0,24409274 |
| 3375_at | IAPP | -0,15712891 | 3,4081 | -1,99224207 | 0,06548571 | 0,24409274 |
| 6888_at | TALDO1 | -0,18126178 | 11,5653535 | -1,99223341 | 0,06548675 | 0,24409274 |
| 665_at | BNIP3L | 0,33123691 | 10,1741781 | 1,99210886 | 0,06550173 | 0,24410134 |
| 90809_at | TMEM55B | 0,2918494 | 8,66238697 | 1,99182954 | 0,06553533 | 0,24417932 |
| 148327_at | CREB3L4 | 0,27855656 | 6,69775888 | 1,99160028 | 0,06556292 | 0,24421692 |
| 9324_at | HMGN3 | 0,27061764 | 9,59140378 | 1,99150885 | 0,06557393 | 0,24421692 |
| 2202_at | EFEMP1 | 0,47285659 | 7,00371464 | 1,99130319 | 0,06559869 | 0,24421692 |
| 9899_at | SV2B | -0,15890622 | 4,05136373 | -1,99110747 | 0,06562227 | 0,24421692 |
| 26521_at | TIMM8B | -0,28673086 | 7,84078901 | -1,99108414 | 0,06562508 | 0,24421692 |
| 85364_at | ZCCHC3 | 0,26568687 | 7,64966867 | 1,99107494 | 0,06562619 | 0,24421692 |
| 26353_at | HSPB8 | 0,58036263 | 9,08446661 | 1,9910087 | 0,06563417 | 0,24421692 |
| 3925_at | STMN1 | -0,26854176 | 6,97532909 | -1,99084334 | 0,0656541 | 0,24424389 |
| 64083_at | GOLPH3 | 0,15417873 | 10,6315624 | 1,99070157 | 0,06567119 | 0,24426029 |
| 79169_at | C1orf35 | -0,18419174 | 7,66177454 | -1,99000122 | 0,06575566 | 0,24452729 |
| 93974_at | ATPIF1 | -0,15263114 | 8,51208023 | -1,98988427 | 0,06576978 | 0,24453258 |
| 207147_at | TRY2P | -0,1947865 | 4,33281384 | -1,98933278 | 0,06583639 | 0,24473299 |
| 3195_at | TLX1 | -0,19246047 | 4,48438499 | -1,98915508 | 0,06585786 | 0,24476558 |
| 6327_at | SCN2B | -0,16143764 | 4,34701433 | -1,98882698 | 0,06589753 | 0,24486576 |
| 80818_at | ZNF436 | 0,16778077 | 8,32667319 | 1,98856507 | 0,06592921 | 0,24493623 |
| 51069_at | MRPL2 | -0,32937012 | 7,45279473 | -1,9883465 | 0,06595566 | 0,24494288 |
| 85369_at | STRIP1 | 0,15839692 | 8,21695074 | 1,98834013 | 0,06595643 | 0,24494288 |
| 340547_at | VSIG1 | -0,17269857 | 3,77114559 | -1,98821023 | 0,06597215 | 0,24495404 |
| 3075_at | CFH | 0,58459847 | 8,66005757 | 1,98803299 | 0,06599361 | 0,2449865 |
| 8543_at | LMO4 | 0,31400907 | 5,83136665 | 1,98720272 | 0,06609422 | 0,24531272 |
| 2289_at | FKBP5 | -0,68303897 | 6,22878216 | -1,98694726 | 0,06612521 | 0,24538044 |
| 100507143_a | LINC00708 | -0,12088156 | 3,04084969 | -1,98667688 | 0,06615801 | 0,24541795 |
| 2830_at | GPR6 | -0,1482887 | 5,56930555 | -1,98665396 | 0,0661608 | 0,24541795 |
| 80835_at | TAS1R1 | -0,16663788 | 4,58722142 | -1,98649157 | 0,06618051 | 0,24544381 |

| | | | | | | |
|---|---|---|---|---|---|---|
| 56121_at | PCDHB15 | 0,458453 | 4,27557454 | 1,98553756 | 0,06629642 | 0,24582637 |
| 438_at | ASMT | -0,16451891 | 4,16691011 | -1,98505118 | 0,06635559 | 0,24596578 |
| 83667_at | SESN2 | 0,39030595 | 7,83095977 | 1,98494484 | 0,06636853 | 0,24596578 |
| 285766_at | LOC285766 | -0,16200683 | 3,73740095 | -1,98491367 | 0,06637233 | 0,24596578 |
| 79160_at | LOC79160 | -0,21162115 | 5,37764337 | -1,98439533 | 0,06643545 | 0,24615235 |
| 2773_at | GNAI3 | -0,16967507 | 9,45227598 | -1,98417397 | 0,06646242 | 0,24620493 |
| 6498_at | SKIL | 0,47756795 | 8,87008974 | 1,98315426 | 0,06658681 | 0,24660275 |
| 80155_at | NAA15 | -0,74726017 | 8,3476397 | -1,98308374 | 0,06659542 | 0,24660275 |
| 2583_at | B4GALNT1 | -0,40465588 | 6,36790932 | -1,98149237 | 0,06678998 | 0,24727567 |
| 1959_at | EGR2 | 0,2401537 | 3,93111243 | 1,98117346 | 0,06682903 | 0,24737271 |
| 8701_at | DNAH11 | -0,16223672 | 4,6055857 | -1,98101678 | 0,06684822 | 0,24739622 |
| 729991_at | BORCS8 | -0,15293812 | 7,25858067 | -1,98086462 | 0,06686687 | 0,24741769 |
| 4054_at | LTBP3 | 0,25890948 | 7,75195749 | 1,98052524 | 0,06690847 | 0,24752408 |
| 9258_at | MFHAS1 | 0,63874001 | 8,11040679 | 1,98020271 | 0,06694803 | 0,24762287 |
| 5054_at | SERPINE1 | -0,65275018 | 8,62857037 | -1,98009169 | 0,06696165 | 0,24762416 |
| 6713_at | SQLE | -0,42729653 | 9,86032747 | -1,9799624 | 0,06697752 | 0,24762416 |
| 9170_at | LPAR2 | 0,53300543 | 7,30416331 | 1,9798039 | 0,06699697 | 0,24762416 |
| 23539_at | SLC16A8 | -0,15688289 | 4,9911095 | -1,97968746 | 0,06701127 | 0,24762416 |
| 64802_at | NMNAT1 | 0,2276507 | 5,56271503 | 1,97957872 | 0,06702462 | 0,24762416 |
| 11198_at | SUPT16H | -0,48835639 | 9,14894917 | -1,97955817 | 0,06702715 | 0,24762416 |
| 23406_at | COTL1 | -0,71038193 | 11,0809425 | -1,97946686 | 0,06703836 | 0,24762416 |
| 55075_at | UACA | 0,18712052 | 5,14391071 | 1,97890561 | 0,06710734 | 0,24779578 |
| 57724_at | EPG5 | 0,14469014 | 5,83690099 | 1,97883936 | 0,06711548 | 0,24779578 |
| 401115_at | C4orf48 | 0,27551321 | 9,28126089 | 1,97877484 | 0,06712342 | 0,24779578 |
| 54661_at | PCDHB17P | -0,15093185 | 4,55938596 | -1,97829749 | 0,06718214 | 0,24796506 |
| 10151_at | HNRNPA3P1 | -0,11252251 | 3,95826275 | -1,97814408 | 0,06720103 | 0,24796971 |
| 9848_at | MFAP3L | 0,29610354 | 5,66612018 | 1,9780781 | 0,06720915 | 0,24796971 |
| 1874_at | E2F4 | -0,16334634 | 8,63308027 | -1,9778391 | 0,06723858 | 0,2480308 |
| 54826_at | GIN1 | -0,2033745 | 6,4113865 | -1,97678915 | 0,06736802 | 0,24845761 |
| 10237_at | SLC35B1 | -0,20420596 | 9,33941882 | -1,97669135 | 0,06738009 | 0,24845761 |
| 9910_at | RABGAP1L | 0,44546696 | 7,67589458 | 1,97635612 | 0,06742147 | 0,24856262 |
| 284353_at | NKPD1 | -0,23921739 | 6,06117611 | -1,97592736 | 0,06747443 | 0,2486467 |
| 4841_at | NONO | -0,26669116 | 10,5247961 | -1,97589214 | 0,06747878 | 0,2486467 |
| 55032_at | SLC35A5 | 0,43565012 | 8,82327669 | 1,975858 | 0,067483 | 0,2486467 |
| 5052_at | PRDX1 | -0,45669013 | 12,224594 | -1,97574447 | 0,06749703 | 0,24865084 |
| 92293_at | TMEM132C | -0,20768462 | 4,35972195 | -1,97563632 | 0,0675104 | 0,24865253 |
| 284716_at | RIMKLA | -0,12587814 | 4,55380968 | -1,97540546 | 0,06753894 | 0,24871012 |
| 9862_at | MED24 | -0,23201769 | 8,05524687 | -1,97490457 | 0,06760091 | 0,24886513 |
| 54386_at | TERF2IP | 0,20121859 | 8,68337338 | 1,97478614 | 0,06761557 | 0,24886513 |
| 64319_at | FBRS | 0,18156332 | 7,10189499 | 1,97475202 | 0,0676198 | 0,24886513 |
| 10569_at | SLU7 | 0,25860865 | 7,56227521 | 1,97420858 | 0,06768711 | 0,24904848 |
| 153129_at | SLC38A9 | 0,59090571 | 7,27361502 | 1,97414108 | 0,06769547 | 0,24904848 |
| 30061_at | SLC40A1 | 1,71130618 | 8,00379333 | 1,97323702 | 0,0678076 | 0,24941335 |
| 1282_at | COL4A1 | -0,49561471 | 7,07485058 | -1,97307598 | 0,06782759 | 0,24943925 |
| 100306951_a | PITPNA-AS1 | -0,17815614 | 7,66129623 | -1,97243957 | 0,06790664 | 0,2496823 |
| 407_at | ARR3 | -0,21073097 | 4,82053468 | -1,97223593 | 0,06793195 | 0,24969975 |
| 126298_at | IRGQ | 0,1316575 | 6,60771787 | 1,97201 | 0,06796004 | 0,24969975 |
| 196968_at | DNM1P46 | -0,15880129 | 5,05702176 | -1,97192878 | 0,06797015 | 0,24969975 |
| 6512_at | SLC1A7 | -0,28688474 | 4,84142738 | -1,9717118 | 0,06799714 | 0,24969975 |
| 23557_at | SNAPIN | 0,17727595 | 9,09388125 | 1,97155992 | 0,06801604 | 0,24969975 |

| | | | | | | |
|---|---|---|---|---|---|---|
| 27006_at | FGF22 | -0,23327861 | 6,28192885 | -1,97152655 | 0,0680202 | 0,24969975 |
| 84541_at | KBTBD8 | -0,40458782 | 4,8396056 | -1,97151072 | 0,06802217 | 0,24969975 |
| 100506844_a | LOC10050684 | 0,40857377 | 5,73517388 | 1,97147759 | 0,06802629 | 0,24969975 |
| 3809_at | KIR2DS4 | -0,20324992 | 3,76585666 | -1,97146348 | 0,06802805 | 0,24969975 |
| 5033_at | P4HA1 | -0,37484169 | 9,0011564 | -1,97119264 | 0,06806177 | 0,24975268 |
| 2582_at | GALE | -0,25723536 | 9,67561272 | -1,9711394 | 0,0680684 | 0,24975268 |
| 28992_at | MACROD1 | 0,18809472 | 6,43623443 | 1,97063016 | 0,06813185 | 0,24993789 |
| 643155_at | SMIM15 | -0,23818639 | 9,69686814 | -1,97014015 | 0,06819296 | 0,25007197 |
| 9506_at | PAGE4 | -0,18232058 | 3,88373849 | -1,97012889 | 0,06819436 | 0,25007197 |
| 100507266_a | STX18-AS1 | 0,32401139 | 4,40546346 | 1,96911066 | 0,06832151 | 0,25047032 |
| 3592_at | IL12A | 0,31510212 | 3,54935699 | 1,96895526 | 0,06834093 | 0,25047032 |
| 79591_at | C10orf76 | 0,18510951 | 7,19235109 | 1,96894669 | 0,068342 | 0,25047032 |
| 100287814_a | LINC00582 | -0,18973893 | 5,29391826 | -1,96883125 | 0,06835643 | 0,25047555 |
| 55783_at | CMTR2 | 0,26763715 | 8,01473033 | 1,9679134 | 0,06847128 | 0,25083554 |
| 4993_at | OR2C1 | -0,22518036 | 4,18451453 | -1,96778854 | 0,06848691 | 0,25083554 |
| 27343_at | POLL | 0,11565689 | 6,70662807 | 1,96773403 | 0,06849374 | 0,25083554 |
| 127687_at | C1orf122 | -0,24388557 | 9,76780012 | -1,96751229 | 0,06852152 | 0,25088958 |
| 659_at | BMPR2 | 0,46580565 | 8,6360084 | 1,96714429 | 0,06856765 | 0,25101076 |
| 9031_at | BAZ1B | -0,21972914 | 8,76314897 | -1,96658238 | 0,06863813 | 0,25120303 |
| 4615_at | MYD88 | 0,24974875 | 10,6630009 | 1,96651768 | 0,06864625 | 0,25120303 |
| 55794_at | DDX28 | -0,25293974 | 6,68780347 | -1,96609702 | 0,06869907 | 0,25134856 |
| 7164_at | TPD52L1 | -0,54664837 | 8,43055448 | -1,96524264 | 0,06880645 | 0,25169364 |
| 90050_at | FAM181A | -0,1969796 | 5,20444522 | -1,96499169 | 0,06883802 | 0,25176005 |
| 9972_at | NUP153 | -0,38171992 | 10,369764 | -1,96482126 | 0,06885947 | 0,25176005 |
| 5880_at | RAC2 | -0,33454886 | 8,93055469 | -1,96478674 | 0,06886382 | 0,25176005 |
| 105371371_a | LOC10537137 | -0,24676196 | 4,83662038 | -1,96455134 | 0,06889345 | 0,2518206 |
| 84874_at | ZNF514 | 0,15461682 | 6,19823533 | 1,96410352 | 0,06894986 | 0,25197898 |
| 8239_at | USP9X | 0,1900778 | 8,85191988 | 1,96380614 | 0,06898735 | 0,25206814 |
| 11322_at | TMC6 | 0,19851124 | 7,79861627 | 1,96355163 | 0,06901944 | 0,25213758 |
| 64102_at | TNMD | -0,14683666 | 3,78603418 | -1,96335043 | 0,06904482 | 0,25218248 |
| 54434_at | SSH1 | -0,122737 | 6,60943017 | -1,96282803 | 0,06911076 | 0,25237547 |
| 115098_at | CCDC124 | -0,3750108 | 8,42082238 | -1,96212252 | 0,06919991 | 0,25260951 |
| 51750_at | RTEL1 | -0,29627735 | 7,39798378 | -1,96211324 | 0,06920108 | 0,25260951 |
| 56894_at | AGPAT3 | 0,33804421 | 8,15492638 | 1,96198959 | 0,06921671 | 0,25261871 |
| 29109_at | FHOD1 | -0,35492633 | 7,14058257 | -1,96175851 | 0,06924594 | 0,25267751 |
| 55507_at | GPRC5D | -0,15676307 | 4,52761063 | -1,96152403 | 0,06927561 | 0,2526905 |
| 272_at | AMPD3 | 0,17736279 | 6,27884092 | 1,96136327 | 0,06929596 | 0,2526905 |
| 645676_at | ASH1L-AS1 | -0,20426257 | 7,39366536 | -1,96133496 | 0,06929954 | 0,2526905 |
| 219293_at | ATAD3C | -0,22705543 | 5,545887 | -1,96130216 | 0,06930369 | 0,2526905 |
| 55814_at | BDP1 | -0,256609 | 5,99413871 | -1,96121215 | 0,06931509 | 0,2526905 |
| 10446_at | LRRN2 | -0,26355244 | 5,43371892 | -1,96072127 | 0,06937727 | 0,25286932 |
| 2151_at | F2RL2 | -0,3866643 | 4,38005717 | -1,96009829 | 0,06945626 | 0,25310932 |
| 23047_at | PDS5B | 0,19614286 | 8,18091026 | 1,95976075 | 0,06949909 | 0,25311867 |
| 7045_at | TGFBI | 0,39299176 | 10,9314878 | 1,95974151 | 0,06950153 | 0,25311867 |
| 64651_at | CSRNP1 | 0,27075514 | 6,44245379 | 1,95967592 | 0,06950986 | 0,25311867 |
| 79883_at | PODNL1 | -0,18656996 | 5,43026055 | -1,95966388 | 0,06951138 | 0,25311867 |
| 55536_at | CDCA7L | -1,05471642 | 6,14600966 | -1,95945886 | 0,06953741 | 0,25312689 |
| 55601_at | DDX60 | 1,31664685 | 8,21434229 | 1,95941321 | 0,06954321 | 0,25312689 |
| 84881_at | RPUSD4 | -0,35125308 | 7,86758787 | -1,9592186 | 0,06956793 | 0,25312689 |
| 101927768_a | LOC10192776 | 0,19944597 | 4,19294368 | 1,9592035 | 0,06956985 | 0,25312689 |

| | | | | | | |
|---|---|---|---|---|---|---|
| 56683_at | C21orf59 | -0,33897268 | 8,72375233 | -1,95912877 | 0,06957934 | 0,25312689 |
| 56940_at | DUSP22 | 0,21306091 | 7,56017651 | 1,95861838 | 0,06964422 | 0,25331507 |
| 147991_at | DPY19L3 | 0,41854845 | 7,28879411 | 1,95829672 | 0,06968513 | 0,25341604 |
| 10683_at | DLL3 | -0,1594549 | 4,88827331 | -1,95721958 | 0,0698223 | 0,25383154 |
| 9296_at | ATP6V1F | 0,18123392 | 9,52707457 | 1,95719258 | 0,06982575 | 0,25383154 |
| 80070_at | ADAMTS20 | -0,16619929 | 2,98272227 | -1,95706199 | 0,06984239 | 0,25384415 |
| 9360_at | PPIG | -0,17937924 | 8,18005459 | -1,95613189 | 0,06996107 | 0,2542275 |
| 100128750_a | RBPMS-AS1 | 0,21926718 | 5,66773123 | 1,95599442 | 0,06997862 | 0,25424334 |
| 113675_at | SDSL | 0,34033434 | 6,71660257 | 1,95581136 | 0,070002 | 0,25428033 |
| 57460_at | PPM1H | 0,44859634 | 5,65774889 | 1,95519005 | 0,07008142 | 0,25452082 |
| 51360_at | MBTPS2 | -0,18152529 | 8,56210826 | -1,95424696 | 0,07020213 | 0,25491113 |
| 5478_at | PPIA | -0,32926957 | 8,72723969 | -1,95377471 | 0,07026264 | 0,25505072 |
| 7110_at | TMF1 | 0,30691181 | 6,75436466 | 1,9534668 | 0,07030212 | 0,25505072 |
| 80333_at | KCNIP4 | -0,13710119 | 4,12496452 | -1,95344485 | 0,07030493 | 0,25505072 |
| 5067_at | CNTN3 | 0,26566384 | 4,17756317 | 1,95343761 | 0,07030586 | 0,25505072 |
| 8502_at | PKP4 | -0,34761345 | 8,34781389 | -1,95343055 | 0,07030677 | 0,25505072 |
| 170082_at | TCEANC | 0,28047969 | 4,82905664 | 1,9529253 | 0,0703716 | 0,25522507 |
| 3797_at | KIF3C | 0,15228382 | 7,03683345 | 1,95284951 | 0,07038133 | 0,25522507 |
| 6625_at | SNRNP70 | -0,18352546 | 9,43170448 | -1,95257503 | 0,07041658 | 0,2552723 |
| 28797_at | IGLV3-19 | -0,24860782 | 4,36196735 | -1,9525417 | 0,07042086 | 0,2552723 |
| 23456_at | ABCB10 | -0,26064585 | 8,32467267 | -1,95238571 | 0,0704409 | 0,25527463 |
| 24139_at | EML2 | 0,14848796 | 7,18199842 | 1,95233039 | 0,07044801 | 0,25527463 |
| 10675_at | CSPG5 | -0,39236336 | 5,28980304 | -1,95209216 | 0,07047862 | 0,25533754 |
| 56245_at | C21orf62 | -0,20129917 | 4,18195127 | -1,95175297 | 0,07052224 | 0,25544751 |
| 399876_at | NAV2-AS4 | -0,15498086 | 4,48184129 | -1,95134118 | 0,07057522 | 0,25559136 |
| 84364_at | ARFGAP2 | 0,11160273 | 7,48759616 | 1,95008508 | 0,07073705 | 0,2561293 |
| 56101_at | PCDHGB5 | -0,15395749 | 5,81350349 | -1,94945741 | 0,07081805 | 0,25637437 |
| 92565_at | FANK1 | 0,44565763 | 5,8995396 | 1,94929929 | 0,07083846 | 0,2564001 |
| 317662_at | FAM149B1 | 0,16355656 | 7,15861113 | 1,94915864 | 0,07085663 | 0,25641767 |
| 9126_at | SMC3 | -0,4996999 | 8,69307693 | -1,94846361 | 0,07094646 | 0,25669452 |
| 80232_at | WDR26 | 0,27788138 | 8,16741731 | 1,9482585 | 0,07097299 | 0,25674228 |
| 6091_at | ROBO1 | 1,13746937 | 7,48754732 | 1,94808162 | 0,07099587 | 0,25674813 |
| 55160_at | ARHGEF10L | 0,25009276 | 8,01189063 | 1,94803997 | 0,07100126 | 0,25674813 |
| 161779_at | PGBD4 | -0,22390426 | 5,37576505 | -1,94789241 | 0,07102036 | 0,25676899 |
| 84892_at | POMGNT2 | -0,14851819 | 7,67440482 | -1,94765623 | 0,07105094 | 0,25683133 |
| 51107_at | APH1A | 0,16032904 | 9,35069477 | 1,94754605 | 0,07106521 | 0,25683472 |
| 10069_at | RWDD2B | -0,21994356 | 8,41243368 | -1,94732715 | 0,07109356 | 0,25688125 |
| 91319_at | DERL3 | -0,17148141 | 5,58715523 | -1,94724079 | 0,07110475 | 0,25688125 |
| 166815_at | TIGD2 | 0,53579911 | 7,82193056 | 1,94702971 | 0,07113211 | 0,25693189 |
| 100129845_a | PCOLCE-AS1 | -0,25642121 | 5,81005154 | -1,94634404 | 0,07122104 | 0,25720188 |
| 440498_at | HSBP1L1 | 0,30727117 | 7,96069262 | 1,94624757 | 0,07123356 | 0,25720188 |
| 79157_at | MFSD11 | 0,23423496 | 7,96497585 | 1,94601922 | 0,0712632 | 0,25725768 |
| 8654_at | PDE5A | 0,16514172 | 5,31442558 | 1,94592279 | 0,07127572 | 0,25725768 |
| 400655_at | LOC400655 | -0,21766226 | 4,06722935 | -1,94568313 | 0,07130685 | 0,25732182 |
| 1269_at | CNR2 | -0,20725728 | 4,37831415 | -1,9454271 | 0,07134012 | 0,25739366 |
| 54584_at | GNB1L | -0,28754193 | 6,63037848 | -1,94499575 | 0,0713962 | 0,25754776 |
| 3882_at | KRT32 | -0,16149089 | 5,0746853 | -1,94350617 | 0,07159017 | 0,2580854 |
| 56953_at | NT5M | -0,29559147 | 5,65741707 | -1,94333046 | 0,07161309 | 0,2580854 |
| 324_at | APC | 0,18644554 | 5,79933673 | 1,94328544 | 0,07161896 | 0,2580854 |
| 101927455_a | TMEM108-AS | -0,21642889 | 3,91186163 | -1,94326698 | 0,07162137 | 0,2580854 |

| | | | | | | |
|---|---|---|---|---|---|---|
| 121274_at | ZNF641 | 0,33739466 | 6,06580195 | 1,9430545 | 0,07164909 | 0,2580854 |
| 10321_at | CRISP3 | 1,64885669 | 3,93396193 | 1,94301522 | 0,07165421 | 0,2580854 |
| 261734_at | NPHP4 | -0,23121874 | 7,16504593 | -1,9429386 | 0,07166421 | 0,2580854 |
| 100874205_a | ATP11A-AS1 | -0,16653267 | 4,18012659 | -1,94292713 | 0,07166571 | 0,2580854 |
| 326340_at | ZAR1 | -0,1627461 | 4,45955992 | -1,94292621 | 0,07166583 | 0,2580854 |
| 54830_at | NUP62CL | -0,46856749 | 5,96724948 | -1,94274035 | 0,07169009 | 0,25812451 |
| 9669_at | EIF5B | -0,36231851 | 8,5191419 | -1,94248492 | 0,07172344 | 0,25819633 |
| 101927992_a | LINC01213 | 0,50995889 | 4,79072536 | 1,94194723 | 0,07179369 | 0,25835552 |
| 122704_at | MRPL52 | -0,42272929 | 8,66400459 | -1,94194117 | 0,07179448 | 0,25835552 |
| 146336_at | SSTR5-AS1 | -0,38472504 | 5,21872189 | -1,94164529 | 0,07183317 | 0,2584391 |
| 100507058_a | LINC00845 | -0,24421153 | 3,54781017 | -1,94155835 | 0,07184454 | 0,2584391 |
| 51174_at | TUBD1 | -0,13965504 | 6,21677788 | -1,94110885 | 0,07190336 | 0,25860238 |
| 56474_at | CTPS2 | 0,17128827 | 7,49188784 | 1,9409557 | 0,07192341 | 0,2586262 |
| 5133_at | PDCD1 | -0,21212593 | 5,92412249 | -1,94038782 | 0,07199779 | 0,25882472 |
| 7471_at | WNT1 | -0,17607996 | 4,94543016 | -1,94023421 | 0,07201793 | 0,25882472 |
| 80349_at | WDR61 | -0,1466512 | 8,83673476 | -1,9402266 | 0,07201892 | 0,25882472 |
| 63027_at | SLC22A23 | 0,18258894 | 7,02276496 | 1,93874446 | 0,07221345 | 0,25947541 |
| 631_at | BFSP1 | 0,42118422 | 4,63946491 | 1,93814976 | 0,07229164 | 0,2597079 |
| 2005_at | ELK4 | -0,24315621 | 6,98358005 | -1,93800034 | 0,07231129 | 0,25972141 |
| 4738_at | NEDD8 | -0,22767801 | 11,2283592 | -1,93783272 | 0,07233335 | 0,25972141 |
| 100507495_a | SDCBP2-AS1 | 0,20649925 | 4,88492363 | 1,93781375 | 0,07233584 | 0,25972141 |
| 9462_at | RASAL2 | 0,23795331 | 6,74998272 | 1,93735828 | 0,07239581 | 0,2598251 |
| 84101_at | USP44 | 0,45702107 | 3,43525584 | 1,93735491 | 0,07239625 | 0,2598251 |
| 146754_at | DNAH2 | -0,16971232 | 5,18615296 | -1,9372871 | 0,07240519 | 0,2598251 |
| 55289_at | ACOXL | 0,26376607 | 6,85695122 | 1,93673546 | 0,07247788 | 0,26003752 |
| 54554_at | WDR5B | 0,29676539 | 7,14077262 | 1,93578533 | 0,07260325 | 0,2604388 |
| 145773_at | FAM81A | -0,20629943 | 4,61358712 | -1,93565916 | 0,07261991 | 0,26045007 |
| 8450_at | CUL4B | 0,17090427 | 6,9988524 | 1,93547276 | 0,07264453 | 0,26048988 |
| 89782_at | LMLN | 0,25641886 | 7,22581804 | 1,93520093 | 0,07268045 | 0,26056507 |
| 9766_at | SUSD6 | 0,6090257 | 8,37595852 | 1,93510938 | 0,07269256 | 0,26056507 |
| 55752_at | SEPT11 | -0,47976698 | 7,62304247 | -1,93484678 | 0,07272728 | 0,26064104 |
| 11238_at | CA5B | -0,18600863 | 5,47121043 | -1,93438917 | 0,07278782 | 0,26079751 |
| 57469_at | PNMAL2 | -0,19340269 | 5,32581022 | -1,93430725 | 0,07279866 | 0,26079751 |
| 55762_at | ZNF701 | 0,18524512 | 4,63292915 | 1,93420986 | 0,07281156 | 0,26079751 |
| 4638_at | MYLK | 0,25978616 | 4,97947728 | 1,933866 | 0,07285709 | 0,26090012 |
| 83990_at | BRIP1 | -0,53638269 | 6,71207001 | -1,93378901 | 0,07286729 | 0,26090012 |
| 92979_at | MARCH9 | 0,2171651 | 6,46491673 | 1,93366746 | 0,0728834 | 0,26090313 |
| 9406_at | ZRANB2 | -0,25997136 | 9,12397351 | -1,93357824 | 0,07289522 | 0,26090313 |
| 101593348_a | LOC10159334 | -0,17280628 | 3,9941153 | -1,93321151 | 0,07294384 | 0,26102863 |
| 101927703_a | FOXCUT | -0,15993604 | 5,44223812 | -1,93260585 | 0,07302419 | 0,26126765 |
| 345778_at | MTX3 | -0,2988546 | 6,63082908 | -1,93232622 | 0,07306132 | 0,26130897 |
| 4163_at | MCC | 0,3632454 | 8,5359506 | 1,93231452 | 0,07306287 | 0,26130897 |
| 80224_at | NUBPL | -0,17483867 | 7,03221236 | -1,93190089 | 0,07311782 | 0,26145696 |
| 10924_at | SMPDL3A | 1,02046544 | 7,50369077 | 1,93179042 | 0,07313251 | 0,26146093 |
| 199800_at | ADM5 | -0,42649596 | 5,57752302 | -1,9315426 | 0,07316545 | 0,26153018 |
| 256815_at | C10orf67 | -0,16119373 | 4,62502238 | -1,93142795 | 0,0731807 | 0,26153614 |
| 6687_at | SPG7 | 0,11350338 | 7,65067416 | 1,93103937 | 0,0732324 | 0,26167235 |
| 80034_at | CSRNP3 | 0,16718186 | 4,08340153 | 1,93076275 | 0,07326922 | 0,26175537 |
| 27286_at | SRPX2 | -0,46539113 | 8,33714516 | -1,93055859 | 0,0732964 | 0,26180394 |
| 29097_at | CNIH4 | -0,18736484 | 8,3491312 | -1,93030601 | 0,07333005 | 0,26181302 |

| | | | | | | |
|---|---|---|---|---|---|---|
| 51125_at | GOLGA7 | 0,32521848 | 9,34813911 | 1,93030386 | 0,07333034 | 0,26181302 |
| 10656_at | KHDRBS3 | -0,16430413 | 5,71616765 | -1,93023344 | 0,07333972 | 0,26181302 |
| 80317_at | ZKSCAN3 | 0,19216892 | 5,33809213 | 1,930082 | 0,07335991 | 0,2618228 |
| 22994_at | CEP131 | -0,29022884 | 7,06458512 | -1,93000894 | 0,07336965 | 0,2618228 |
| 101927273_a | LINC01416 | -0,13914423 | 2,99077373 | -1,92936414 | 0,07345565 | 0,26208115 |
| 5326_at | PLAGL2 | 0,19496233 | 7,29409192 | 1,92909419 | 0,07349168 | 0,26214166 |
| 26098_at | EDRF1 | -0,11680295 | 7,54315747 | -1,92903319 | 0,07349982 | 0,26214166 |
| 26188_at | OR1C1 | -0,14367837 | 3,58229368 | -1,92869866 | 0,0735445 | 0,26225245 |
| 59285_at | CACNG6 | -0,24459829 | 5,99455956 | -1,92848125 | 0,07357355 | 0,26230749 |
| 9066_at | SYT7 | -0,18621821 | 5,39880602 | -1,92835248 | 0,07359076 | 0,2623203 |
| 54891_at | INO80D | 0,11240984 | 7,20495723 | 1,9276164 | 0,07368922 | 0,26262264 |
| 100874362_a | HOXB-AS1 | -0,27370893 | 4,73612222 | -1,92746845 | 0,07370902 | 0,26264462 |
| 399967_at | PATE2 | -0,16516892 | 3,39423292 | -1,92714566 | 0,07375224 | 0,26266344 |
| 2475_at | MTOR | -0,24383554 | 8,43733308 | -1,92714511 | 0,07375231 | 0,26266344 |
| 55830_at | GLT8D1 | 0,19880942 | 8,24162313 | 1,9271235 | 0,07375521 | 0,26266344 |
| 26206_at | SPAG8 | -0,1459508 | 3,92247086 | -1,92685741 | 0,07379086 | 0,26271691 |
| 8986_at | RPS6KA4 | -0,20851677 | 8,29652107 | -1,92680785 | 0,0737975 | 0,26271691 |
| 8813_at | DPM1 | -0,24611158 | 10,3329529 | -1,92657951 | 0,0738281 | 0,2627773 |
| 401548_at | SNX30 | 0,31586058 | 7,18787656 | 1,92576875 | 0,07393687 | 0,26300406 |
| 9615_at | GDA | -0,48970959 | 6,76359758 | -1,92571767 | 0,07394372 | 0,26300406 |
| 85439_at | STON2 | 0,44642954 | 7,42257082 | 1,92561038 | 0,07395813 | 0,26300406 |
| 3848_at | KRT1 | -0,44791556 | 4,06020582 | -1,92545984 | 0,07397835 | 0,26300406 |
| 8499_at | PPFIA2 | -0,1126702 | 3,3057802 | -1,92543951 | 0,07398108 | 0,26300406 |
| 9644_at | SH3PXD2A | -0,23648487 | 6,23184489 | -1,92542934 | 0,07398244 | 0,26300406 |
| 728192_at | LINC00460 | -0,24486763 | 4,52466717 | -1,92539253 | 0,07398739 | 0,26300406 |
| 123169_at | LEO1 | -0,16226775 | 8,10075699 | -1,92515126 | 0,0740198 | 0,26301718 |
| 731223_at | LINC01220 | 0,14354891 | 3,45219298 | 1,9251076 | 0,07402567 | 0,26301718 |
| 7004_at | TEAD4 | -0,55466419 | 7,67186797 | -1,92502074 | 0,07403734 | 0,26301718 |
| 2820_at | GPD2 | -0,41685903 | 8,2422012 | -1,92495493 | 0,07404618 | 0,26301718 |
| 140597_at | TCEAL2 | -0,19654575 | 3,4443724 | -1,92485703 | 0,07405935 | 0,26301718 |
| 101929034_a | LOC10192903 | -0,1555047 | 3,73239789 | -1,92442196 | 0,07411786 | 0,26317646 |
| 401398_at | POT1-AS1 | 0,26849996 | 3,08978935 | 1,92421261 | 0,07414603 | 0,26319075 |
| 285489_at | DOK7 | -0,52122074 | 7,98302305 | -1,92411902 | 0,07415862 | 0,26319075 |
| 9125_at | RQCD1 | -0,35011764 | 8,83417342 | -1,92394301 | 0,07418232 | 0,26319075 |
| 55183_at | RIF1 | -0,40380814 | 7,1251399 | -1,92382595 | 0,07419808 | 0,26319075 |
| 673_at | BRAF | 0,16833281 | 7,217615 | 1,92379747 | 0,07420192 | 0,26319075 |
| 55819_at | RNF130 | 0,33570234 | 7,72784354 | 1,92378303 | 0,07420386 | 0,26319075 |
| 8354_at | HIST1H3I | -0,14240434 | 6,25975571 | -1,92366979 | 0,07421911 | 0,26319639 |
| 79993_at | ELOVL7 | 0,35938278 | 8,18128697 | 1,92322737 | 0,07427873 | 0,26335931 |
| 1387_at | CREBBP | 0,14895972 | 8,03032694 | 1,92248939 | 0,07437827 | 0,26361861 |
| 2077_at | ERF | -0,16230222 | 6,7094634 | -1,92248225 | 0,07437923 | 0,26361861 |
| 23498_at | HAAO | -0,2269526 | 6,68544636 | -1,92220301 | 0,07441693 | 0,26365983 |
| 5582_at | PRKCG | -0,1728435 | 5,36962504 | -1,92219332 | 0,07441824 | 0,26365983 |
| 100128252_a | ZNF667-AS1 | -0,30775932 | 4,29801536 | -1,92196377 | 0,07444924 | 0,26372116 |
| 26037_at | SIPA1L1 | 0,14000896 | 7,28272447 | 1,92174554 | 0,07447872 | 0,26377709 |
| 140739_at | UBE2F | -0,29491751 | 7,16403604 | -1,9211928 | 0,07455344 | 0,2639932 |
| 64005_at | MYO1G | -0,24557828 | 4,36723287 | -1,92096862 | 0,07458377 | 0,26405204 |
| 93380_at | MMGT1 | -0,27206331 | 9,07329289 | -1,92072286 | 0,07461703 | 0,26412125 |
| 25789_at | TMEM59L | -0,26870435 | 5,60063637 | -1,92046986 | 0,07465128 | 0,26419394 |
| 55259_at | CASC1 | 0,22756803 | 3,00208013 | 1,91974168 | 0,07474994 | 0,26449451 |

| | | | | | | |
|---|---|---|---|---|---|---|
| 51371_at | POMP | -0,25234542 | 9,61617857 | -1,91930411 | 0,07480928 | 0,26465587 |
| 100131801_a | PET100 | 0,19980959 | 7,65769682 | 1,91912037 | 0,07483421 | 0,26469546 |
| 55198_at | APPL2 | 0,25284752 | 6,67647982 | 1,91887589 | 0,0748674 | 0,26476422 |
| 100505474_a | LINC01539 | -0,1752388 | 3,79638854 | -1,9173792 | 0,07507084 | 0,2653757 |
| 330_at | BIRC3 | 0,63006306 | 5,16757773 | 1,91732376 | 0,07507839 | 0,2653757 |
| 100130357_a | LOC10013035 | 0,21163837 | 4,10509559 | 1,9172999 | 0,07508164 | 0,2653757 |
| 149986_at | LSM14B | 0,15613395 | 7,36939747 | 1,91707709 | 0,07511197 | 0,2654342 |
| 79086_at | SMIM7 | 0,12279753 | 8,62425134 | 1,91695049 | 0,07512921 | 0,26543887 |
| 4929_at | NR4A2 | 0,1974153 | 4,76284185 | 1,91683566 | 0,07514485 | 0,26543887 |
| 1794_at | DOCK2 | -0,14967002 | 3,53541895 | -1,91676388 | 0,07515463 | 0,26543887 |
| 23352_at | UBR4 | -0,21837619 | 8,60773433 | -1,91653326 | 0,07518605 | 0,26550117 |
| 79755_at | ZNF750 | 0,65702629 | 5,75591821 | 1,91623596 | 0,07522658 | 0,2655899 |
| 7582_at | ZNF33B | 0,29790267 | 6,11224751 | 1,91608594 | 0,07524704 | 0,2655899 |
| 101927164_a | LOC10192716 | -0,26174459 | 4,33519469 | -1,91597862 | 0,07526167 | 0,2655899 |
| 915_at | CD3D | -0,1583285 | 4,42302634 | -1,91594451 | 0,07526633 | 0,2655899 |
| 57551_at | TAOK1 | 0,29025374 | 8,39710821 | 1,91538486 | 0,07534271 | 0,26581072 |
| 2162_at | F13A1 | -0,25619074 | 4,0718101 | -1,91455469 | 0,07545613 | 0,26608621 |
| 51754_at | TMEM8B | 0,33803203 | 6,27133969 | 1,91451948 | 0,07546095 | 0,26608621 |
| 64409_at | WBSCR17 | -0,20868768 | 4,66955184 | -1,91451011 | 0,07546223 | 0,26608621 |
| 112755_at | STX1B | -0,24014663 | 5,66437072 | -1,91406744 | 0,07552278 | 0,266248 |
| 4838_at | NODAL | -0,20009814 | 4,31500522 | -1,91397261 | 0,07553576 | 0,266248 |
| 51058_at | ZNF691 | 0,15844374 | 6,04851684 | 1,91376954 | 0,07556355 | 0,26629724 |
| 57718_at | PPP4R4 | -0,36651266 | 4,56091179 | -1,91362357 | 0,07558354 | 0,26631895 |
| 723809_at | LHFPL3-AS2 | -0,19735819 | 4,0838618 | -1,91341462 | 0,07561216 | 0,26637106 |
| 2022_at | ENG | -0,20083973 | 6,17278137 | -1,91326256 | 0,07563299 | 0,26639573 |
| 113419_at | TEX261 | -0,13332585 | 9,13422528 | -1,91315292 | 0,07564801 | 0,26639993 |
| 7750_at | ZMYM2 | 0,14917273 | 6,77630677 | 1,91295691 | 0,07567488 | 0,26644584 |
| 93099_at | DMKN | -0,23372472 | 8,33230239 | -1,91279515 | 0,07569706 | 0,26647521 |
| 100131561_a | FKSG29 | -0,18452825 | 4,50667544 | -1,9126034 | 0,07572335 | 0,26647589 |
| 285126_at | DNAJC5G | -0,20129889 | 4,79060837 | -1,91252066 | 0,0757347 | 0,26647589 |
| 25852_at | ARMC8 | -0,16113729 | 7,28868196 | -1,91245268 | 0,07574403 | 0,26647589 |
| 25836_at | NIPBL | 0,19359635 | 8,18170054 | 1,91239033 | 0,07575258 | 0,26647589 |
| 4299_at | AFF1 | 0,13453792 | 5,94930671 | 1,91199289 | 0,07580713 | 0,26661886 |
| 83445_at | GSG1 | -0,17217504 | 3,84840574 | -1,91189257 | 0,07582091 | 0,26661886 |
| 100996590_a | LINC01505 | -0,21351925 | 3,30749423 | -1,91139065 | 0,07588986 | 0,26669779 |
| 343641_at | TGM6 | -0,20586757 | 4,88594175 | -1,9112825 | 0,07590472 | 0,26669779 |
| 3998_at | LMAN1 | -0,16555007 | 7,89772778 | -1,91126243 | 0,07590748 | 0,26669779 |
| 154092_at | LINC01010 | -0,24517312 | 4,93895021 | -1,91122202 | 0,07591304 | 0,26669779 |
| 23580_at | CDC42EP4 | -0,20007467 | 7,59743941 | -1,91115772 | 0,07592188 | 0,26669779 |
| 27434_at | POLM | 0,2406243 | 6,45472792 | 1,91112466 | 0,07592642 | 0,26669779 |
| 161514_at | TBC1D21 | -0,2344 | 4,79748608 | -1,91088201 | 0,07595979 | 0,26676636 |
| 286410_at | ATP11C | 0,41845366 | 6,27030852 | 1,91072904 | 0,07598084 | 0,26679162 |
| 158293_at | FAM120AOS | 0,28647175 | 7,33539314 | 1,91050203 | 0,07601207 | 0,26685267 |
| 201501_at | ZBTB7C | 0,29801193 | 5,73681793 | 1,90925472 | 0,07618392 | 0,2673174 |
| 128308_at | MRPL55 | -0,25651947 | 7,51677475 | -1,90922715 | 0,07618772 | 0,2673174 |
| 3779_at | KCNMB1 | -0,27357449 | 4,92056704 | -1,90921094 | 0,07618996 | 0,2673174 |
| 5121_at | PCP4 | 0,21979378 | 4,59333212 | 1,90913842 | 0,07619996 | 0,2673174 |
| 23508_at | TTC9 | 0,39291488 | 7,06730934 | 1,9085994 | 0,07627435 | 0,26752963 |
| 9332_at | CD163 | -0,16086496 | 3,81257602 | -1,90822893 | 0,07632551 | 0,26766035 |
| 8793_at | TNFRSF10D | -0,1975197 | 5,56200673 | -1,90803205 | 0,07635271 | 0,26768775 |

| | | | | | | |
|---|---|---|---|---|---|---|
| 3684_at | ITGAM | 0,24930305 | 3,92491973 | 1,90784202 | 0,07637898 | 0,26768775 |
| 2744_at | GLS | 0,31269195 | 8,20570279 | 1,90782948 | 0,07638071 | 0,26768775 |
| 8996_at | NOL3 | -0,17579297 | 7,64450809 | -1,90777015 | 0,07638891 | 0,26768775 |
| 51690_at | LSM7 | -0,30863881 | 9,65929845 | -1,90746737 | 0,07643078 | 0,26778576 |
| 56605_at | ERO1B | 0,25708235 | 5,5494649 | 1,90732954 | 0,07644985 | 0,26780385 |
| 6309_at | SC5D | 0,29185858 | 8,09573435 | 1,90674703 | 0,07653047 | 0,26803755 |
| 101060200_a | ZNF891 | -0,215426 | 3,74219395 | -1,90635121 | 0,0765853 | 0,26818082 |
| 136227_at | COL26A1 | -0,20863394 | 5,76767209 | -1,90526375 | 0,07673612 | 0,26866011 |
| 101929926_a | LOC10192992 | -0,18029938 | 4,30801305 | -1,90353097 | 0,076977 | 0,26945446 |
| 10223_at | GPA33 | -0,26199728 | 5,2544467 | -1,90302814 | 0,07704702 | 0,26964539 |
| 84809_at | CROCCP2 | 0,16693657 | 8,5333162 | 1,90293832 | 0,07705954 | 0,26964539 |
| 644_at | BLVRA | 0,22479309 | 7,84660619 | 1,90262547 | 0,07710314 | 0,26973296 |
| 197258_at | FUK | 0,30665635 | 7,33926394 | 1,90255786 | 0,07711257 | 0,26973296 |
| 27180_at | SIGLEC9 | -0,18895347 | 4,62868161 | -1,9022975 | 0,07714888 | 0,2697765 |
| 2735_at | GLI1 | -0,24210352 | 4,97476754 | -1,90226775 | 0,07715303 | 0,2697765 |
| 90843_at | TCEAL8 | 0,38954765 | 6,15138175 | 1,90184829 | 0,07721156 | 0,26993217 |
| 23237_at | ARC | -0,20891663 | 5,06551062 | -1,90145331 | 0,07726671 | 0,27007597 |
| 2532_at | ACKR1 | -0,25249821 | 4,66266237 | -1,9008498 | 0,07735105 | 0,27032171 |
| 10579_at | TACC2 | 0,12668229 | 7,03460225 | 1,90036633 | 0,07741867 | 0,27050896 |
| 112744_at | IL17F | -0,2724363 | 4,26309541 | -1,90001597 | 0,07746771 | 0,27054098 |
| 22882_at | ZHX2 | 0,56731282 | 7,40296652 | 1,90001029 | 0,07746851 | 0,27054098 |
| 130589_at | GALM | 0,57053797 | 7,13463229 | 1,89999982 | 0,07746997 | 0,27054098 |
| 286333_at | FAM225A | -0,22268647 | 5,9796635 | -1,89942409 | 0,07755062 | 0,27077352 |
| 9117_at | SEC22C | -0,16672683 | 6,94798382 | -1,89904828 | 0,0776033 | 0,27090016 |
| 6011_at | GRK1 | -0,21721422 | 5,2089677 | -1,89896475 | 0,07761501 | 0,27090016 |
| 3263_at | HPX | -0,15812265 | 5,61245953 | -1,89876772 | 0,07764265 | 0,27094753 |
| 5393_at | EXOSC9 | -0,69651008 | 8,23043615 | -1,89857871 | 0,07766917 | 0,27099098 |
| 10841_at | FTCD | -0,15873364 | 6,3119013 | -1,89839654 | 0,07769474 | 0,27103109 |
| 80764_at | THAP7 | -0,27104162 | 7,24668201 | -1,89808757 | 0,07773812 | 0,27110041 |
| 1489_at | CTF1 | 0,20743733 | 6,32961084 | 1,89797286 | 0,07775423 | 0,27110041 |
| 148189_at | LINC00662 | 0,15977419 | 5,57762 | 1,89795438 | 0,07775683 | 0,27110041 |
| 5439_at | POLR2J | -0,13237624 | 8,21731759 | -1,89776977 | 0,07778276 | 0,27114176 |
| 101928280_a | LOC10192828 | -0,20478401 | 3,7651087 | -1,89723678 | 0,07785769 | 0,27135385 |
| 25822_at | DNAJB5 | 0,16262849 | 5,82461418 | 1,89689251 | 0,07790612 | 0,27143255 |
| 55061_at | SUSD4 | 0,18115562 | 4,39615622 | 1,89687593 | 0,07790845 | 0,27143255 |
| 9444_at | QKI | 0,33970578 | 7,6919363 | 1,89564565 | 0,07808176 | 0,27198713 |
| 27033_at | ZBTB32 | -0,21585813 | 4,51016111 | -1,8953863 | 0,07811833 | 0,27206534 |
| 23028_at | KDM1A | -0,41109715 | 9,84567075 | -1,89498802 | 0,07817453 | 0,27216953 |
| 3164_at | NR4A1 | -0,15967677 | 5,35723371 | -1,89497404 | 0,07817651 | 0,27216953 |
| 54855_at | FAM46C | 0,46537779 | 5,59494881 | 1,89480382 | 0,07820054 | 0,27219209 |
| 340277_at | FAM221A | 0,40422626 | 5,00697906 | 1,89472798 | 0,07821124 | 0,27219209 |
| 1749_at | DLX5 | 0,36925087 | 6,3717246 | 1,89450538 | 0,07824269 | 0,27224919 |
| 108_at | ADCY2 | -0,17732515 | 4,58765244 | -1,8944117 | 0,07825592 | 0,27224919 |
| 222659_at | PXT1 | -0,15752114 | 3,19915681 | -1,89326027 | 0,07841875 | 0,27276642 |
| 64764_at | CREB3L2 | 0,38950432 | 9,21376316 | 1,89315214 | 0,07843406 | 0,27277041 |
| 55860_at | ACTR10 | 0,18625253 | 9,98542404 | 1,89276296 | 0,07848918 | 0,27291281 |
| 101928687_a | LINC01468 | -0,2841691 | 2,9655899 | -1,8925915 | 0,07851347 | 0,27294801 |
| 8991_at | SELENBP1 | 0,35839839 | 5,22555126 | 1,89206074 | 0,07858871 | 0,2731603 |
| 55870_at | ASH1L | 0,17537978 | 8,16426058 | 1,89194295 | 0,07860542 | 0,27316891 |
| 1871_at | E2F3 | -0,43788856 | 7,30636572 | -1,89184332 | 0,07861955 | 0,27316891 |

| | | | | | | |
|---|---|---|---|---|---|---|
| 10667_at | FARS2 | -0,18750513 | 7,04428216 | -1,89153772 | 0,07866292 | 0,2732703 |
| 9480_at | ONECUT2 | -0,13581186 | 4,80577778 | -1,89092019 | 0,07875062 | 0,27352564 |
| 1105_at | CHD1 | -0,20409316 | 8,14230074 | -1,89015533 | 0,07885937 | 0,27385397 |
| 8178_at | ELL | 0,11143315 | 6,42094682 | 1,88967689 | 0,07892746 | 0,27401104 |
| 51297_at | BPIFA1 | -0,16551794 | 4,65534421 | -1,88963764 | 0,07893305 | 0,27401104 |
| 7111_at | TMOD1 | 0,12220841 | 5,57060704 | 1,88950836 | 0,07895146 | 0,27402557 |
| 9124_at | PDLIM1 | 0,1929523 | 12,2044298 | 1,88876576 | 0,07905728 | 0,27426452 |
| 29083_at | GTPBP8 | -0,15893588 | 8,24017432 | -1,88875299 | 0,0790591 | 0,27426452 |
| 55847_at | CISD1 | -0,24274898 | 9,19136521 | -1,88872454 | 0,07906315 | 0,27426452 |
| 64852_at | TUT1 | -0,1295252 | 6,39669665 | -1,88862566 | 0,07907725 | 0,27426452 |
| 100505893_a | SLIT2-IT1 | 0,11002962 | 3,20368908 | 1,88827971 | 0,07912661 | 0,27438629 |
| 56673_at | C11orf16 | -0,15607751 | 5,35672731 | -1,88785155 | 0,07918773 | 0,27450414 |
| 55422_at | ZNF331 | -0,21283815 | 6,64117428 | -1,88778134 | 0,07919775 | 0,27450414 |
| 7001_at | PRDX2 | -0,13647453 | 11,9432884 | -1,88774218 | 0,07920335 | 0,27450414 |
| 7301_at | TYRO3 | -0,40447585 | 7,0431049 | -1,88743415 | 0,07924735 | 0,27457361 |
| 54551_at | MAGEL2 | 0,1957596 | 4,37609646 | 1,88740234 | 0,0792519 | 0,27457361 |
| 1845_at | DUSP3 | -0,32801377 | 8,05161776 | -1,88695438 | 0,07931593 | 0,2747265 |
| 84852_at | ATP1A1-AS1 | 0,14191777 | 5,73867966 | 1,88689413 | 0,07932455 | 0,2747265 |
| 55002_at | TMCO3 | 0,34556453 | 8,40719208 | 1,88670273 | 0,07935193 | 0,27473214 |
| 255654_at | LOC255654 | -0,24783788 | 4,31348074 | -1,88668333 | 0,0793547 | 0,27473214 |
| 29940_at | DSE | 0,49177461 | 6,51705507 | 1,8865206 | 0,07937799 | 0,27476337 |
| 84444_at | DOT1L | -0,13409651 | 7,06007946 | -1,88617199 | 0,07942789 | 0,27488671 |
| 23294_at | ANKS1A | 0,16614209 | 9,08723383 | 1,88589641 | 0,07946736 | 0,2749739 |
| 2741_at | GLRA1 | -0,13922257 | 3,30467583 | -1,88579546 | 0,07948182 | 0,27497455 |
| 317705_at | VN1R5 | -0,1915772 | 3,4072691 | -1,88523927 | 0,07956154 | 0,27520093 |
| 10484_at | SEC23A | 0,27210744 | 9,79263608 | 1,88513605 | 0,07957635 | 0,27520272 |
| 23479_at | ISCU | -0,27242337 | 9,45046381 | -1,8849361 | 0,07960503 | 0,27521004 |
| 6429_at | SRSF4 | -0,14910123 | 9,60224123 | -1,8849221 | 0,07960704 | 0,27521004 |
| 57599_at | WDR48 | 0,25729639 | 7,15859139 | 1,88460226 | 0,07965294 | 0,27531709 |
| 5864_at | RAB3A | -0,14994577 | 6,13500227 | -1,88431118 | 0,07969473 | 0,27531709 |
| 85014_at | TMEM141 | -0,24327747 | 8,99038149 | -1,88391301 | 0,07975193 | 0,27531709 |
| 7633_at | ZNF79 | 0,14383668 | 6,23770385 | 1,8838924 | 0,0797549 | 0,27531709 |
| 101927449_a | LINC01282 | -0,16225579 | 3,96809488 | -1,88388037 | 0,07975663 | 0,27531709 |
| 26084_at | ARHGEF26 | 0,39031427 | 6,21041015 | 1,88386917 | 0,07975824 | 0,27531709 |
| 445_at | ASS1 | 0,30228389 | 9,53469251 | 1,88385777 | 0,07975987 | 0,27531709 |
| 27156_at | RSPH14 | -0,15406073 | 4,57486706 | -1,88382611 | 0,07976442 | 0,27531709 |
| 101929662_a | LOC10192966 | -0,18002985 | 3,62632474 | -1,88378308 | 0,07977061 | 0,27531709 |
| 8772_at | FADD | -0,22767152 | 9,54426755 | -1,88371131 | 0,07978093 | 0,27531709 |
| 23279_at | NUP160 | -0,18691094 | 6,665997 | -1,88316196 | 0,07985994 | 0,27549344 |
| 388695_at | LYSMD1 | 0,19712713 | 5,6133818 | 1,88315712 | 0,07986063 | 0,27549344 |
| 3491_at | CYR61 | 0,59608388 | 8,34537227 | 1,88276551 | 0,079917 | 0,27563852 |
| 116840_at | CNTROB | -0,15524599 | 6,08901267 | -1,88258893 | 0,07994242 | 0,27565387 |
| 91526_at | ANKRD44 | 0,10356304 | 3,9696038 | 1,88253585 | 0,07995007 | 0,27565387 |
| 6198_at | RPS6KB1 | -0,22678608 | 7,9511602 | -1,88239378 | 0,07997053 | 0,27567509 |
| 29121_at | CLEC2D | 0,20303605 | 4,37216974 | 1,88215926 | 0,08000433 | 0,27568487 |
| 115650_at | TNFRSF13C | -0,22613445 | 6,43596863 | -1,8821585 | 0,08000444 | 0,27568487 |
| 2823_at | GPM6A | 0,91628851 | 5,40190191 | 1,88204609 | 0,08002064 | 0,27568487 |
| 1208_at | CLPS | -0,19970014 | 5,52005671 | -1,88197686 | 0,08003062 | 0,27568487 |
| 23163_at | GGA3 | 0,13115597 | 7,76377887 | 1,88186976 | 0,08004606 | 0,27568876 |
| 160140_at | C11orf65 | 0,11276351 | 2,8447598 | 1,88173525 | 0,08006545 | 0,27570627 |

| | | | | | | |
|---|---|---|---|---|---|---|
| 11211_at | FZD10 | -0,16446903 | 7,88095072 | -1,88106239 | 0,08016255 | 0,27599127 |
| 10270_at | AKAP8 | -0,14441632 | 8,3005462 | -1,88063677 | 0,08022402 | 0,27615354 |
| 7566_at | ZNF18 | 0,22582836 | 5,79096778 | 1,88007712 | 0,08030491 | 0,27636973 |
| 8725_at | URI1 | -0,10335264 | 8,3863611 | -1,88000377 | 0,08031551 | 0,27636973 |
| 23185_at | LARP4B | -0,26146614 | 7,60404144 | -1,87967349 | 0,08036329 | 0,27648475 |
| 3337_at | DNAJB1 | -0,13241583 | 9,50213676 | -1,87928925 | 0,08041891 | 0,27659715 |
| 140731_at | ANKRD60 | -0,23140567 | 4,89523405 | -1,87924939 | 0,08042468 | 0,27659715 |
| 221545_at | C6orf136 | -0,15387333 | 7,51128455 | -1,87842652 | 0,08054391 | 0,27695776 |
| 23189_at | KANK1 | 0,12398516 | 7,44277411 | 1,87829795 | 0,08056256 | 0,27695885 |
| 6101_at | RP1 | -0,12338619 | 2,97509266 | -1,87822605 | 0,08057298 | 0,27695885 |
| 84303_at | CHCHD6 | -0,26444048 | 7,61475773 | -1,87781734 | 0,08063228 | 0,27711322 |
| 161424_at | NOP9 | -0,15183379 | 6,66038051 | -1,8776085 | 0,0806626 | 0,27716796 |
| 283417_at | DPY19L2 | 0,2183027 | 3,49414331 | 1,87747408 | 0,08068211 | 0,27717329 |
| 7421_at | VDR | -0,37339815 | 7,58926794 | -1,87731404 | 0,08070536 | 0,27717329 |
| 6666_at | SOX12 | 0,2917877 | 5,47534965 | 1,87718068 | 0,08072473 | 0,27717329 |
| 400604_at | TOB1-AS1 | 0,22063608 | 4,18374092 | 1,87713354 | 0,08073158 | 0,27717329 |
| 51393_at | TRPV2 | -0,14237691 | 4,90505252 | -1,87710248 | 0,08073609 | 0,27717329 |
| 285362_at | SUMF1 | 0,36437221 | 8,51496852 | 1,87698722 | 0,08075284 | 0,27718139 |
| 212_at | ALAS2 | -0,20272716 | 4,76144982 | -1,87671694 | 0,08079213 | 0,27721594 |
| 55166_at | CENPQ | -0,47730649 | 6,14546673 | -1,87671326 | 0,08079266 | 0,27721594 |
| 9901_at | SRGAP3 | 0,5197353 | 7,50018635 | 1,87662098 | 0,08080608 | 0,27721594 |
| 4973_at | OLR1 | 1,4893789 | 9,40118357 | 1,87585847 | 0,08091703 | 0,27751888 |
| 100133311_a | HOXA-AS3 | 0,27199039 | 4,92530307 | 1,8757476 | 0,08093317 | 0,27751888 |
| 51513_at | ETV7 | 0,45855843 | 5,39240198 | 1,87571716 | 0,0809376 | 0,27751888 |
| 4486_at | MST1R | -0,29551374 | 8,40570886 | -1,87560433 | 0,08095404 | 0,27752583 |
| 57862_at | ZNF410 | -0,15515142 | 8,81516106 | -1,87520603 | 0,08101207 | 0,27765871 |
| 3184_at | HNRNPD | -0,36023686 | 10,7843908 | -1,87514047 | 0,08102163 | 0,27765871 |
| 101927650_a | LOC10192765 | -0,13099093 | 4,57575091 | -1,8747811 | 0,08107402 | 0,27778885 |
| 5949_at | RBP3 | -0,17372069 | 5,58862355 | -1,87435104 | 0,08113677 | 0,2779544 |
| 9854_at | C2CD2L | -0,15892291 | 5,85405643 | -1,87421488 | 0,08115664 | 0,27797305 |
| 60682_at | SMAP1 | 0,1932545 | 9,05465294 | 1,87394638 | 0,08119585 | 0,27803157 |
| 23411_at | SIRT1 | 0,17649584 | 7,90189644 | 1,87390017 | 0,0812026 | 0,27803157 |
| 55500_at | ETNK1 | 0,25491091 | 6,66208566 | 1,87374457 | 0,08122532 | 0,27805997 |
| 338811_at | FAM19A2 | 0,21155017 | 4,23775885 | 1,87350715 | 0,08126002 | 0,27810418 |
| 5860_at | QDPR | -0,20406956 | 8,42826459 | -1,87345858 | 0,08126711 | 0,27810418 |
| 1761_at | DMRT1 | -0,8868349 | 5,02155997 | -1,87333924 | 0,08128456 | 0,27811447 |
| 8099_at | CDK2AP1 | -0,33036716 | 12,0446703 | -1,87304988 | 0,08132687 | 0,27812948 |
| 7431_at | VIM | -0,19263863 | 4,79892454 | -1,87304588 | 0,08132745 | 0,27812948 |
| 260293_at | CYP4X1 | 0,11004277 | 3,46866082 | 1,87301302 | 0,08133226 | 0,27812948 |
| 57613_at | FAM234B | 0,24630885 | 6,92493534 | 1,8717742 | 0,08151364 | 0,27870025 |
| 3159_at | HMGA1 | -0,60325659 | 7,69669333 | -1,87145455 | 0,08156049 | 0,27881097 |
| 101928841_a | LOC10192884 | -0,15075491 | 4,41956649 | -1,87135562 | 0,081575 | 0,27881108 |
| 10737_at | RFPL3S | 0,24779396 | 3,65813404 | 1,87051069 | 0,081699 | 0,27915089 |
| 266743_at | NPAS4 | -0,15213885 | 4,17376846 | -1,87048068 | 0,08170341 | 0,27915089 |
| 7436_at | VLDLR | -0,51664077 | 6,45717287 | -1,86978505 | 0,08180562 | 0,27943227 |
| 7143_at | TNR | -0,16668333 | 4,52861122 | -1,86972282 | 0,08181478 | 0,27943227 |
| 387263_at | C6orf120 | 0,40067178 | 8,92524817 | 1,86935502 | 0,08186887 | 0,27956747 |
| 10066_at | SCAMP2 | 0,12380029 | 9,59899826 | 1,86922556 | 0,08188792 | 0,27958296 |
| 27089_at | UQCRQ | -0,21081296 | 11,1138951 | -1,86875835 | 0,0819567 | 0,2797682 |
| 56125_at | PCDHB11 | 0,17342826 | 6,30976889 | 1,8686112 | 0,08197838 | 0,2797926 |

| | | | | | | |
|---|---|---|---|---|---|---|
| 375190_at | FAM228B | 0,67848923 | 5,70224386 | 1,8680522 | 0,08206076 | 0,28002415 |
| 375035_at | SFT2D2 | -0,21063024 | 7,81141211 | -1,86705485 | 0,08220792 | 0,28047665 |
| 131669_at | UROC1 | -0,16777281 | 4,19960532 | -1,86692177 | 0,08222757 | 0,28049403 |
| 51283_at | BFAR | -0,12921045 | 8,93522911 | -1,86654982 | 0,08228253 | 0,2806318 |
| 9240_at | PNMA1 | -0,21200535 | 7,44133801 | -1,86635397 | 0,08231148 | 0,28068084 |
| 4329_at | ALDH6A1 | 0,53758957 | 6,47124366 | 1,86586356 | 0,08238401 | 0,28087845 |
| 157680_at | VPS13B | 0,35033259 | 5,94845337 | 1,86568865 | 0,0824099 | 0,28091214 |
| 8359_at | HIST1H4A | -0,25550865 | 5,05225989 | -1,86545755 | 0,0824441 | 0,28091214 |
| 1917_at | EEF1A2 | -0,24869566 | 6,67773886 | -1,8654146 | 0,08245046 | 0,28091214 |
| 11338_at | U2AF2 | -0,21481233 | 8,12415679 | -1,86540268 | 0,08245223 | 0,28091214 |
| 6399_at | TRAPPC2 | 0,15108869 | 6,42078162 | 1,86524453 | 0,08247565 | 0,28094224 |
| 83641_at | FAM107B | -0,30187446 | 7,38442699 | -1,86500104 | 0,08251172 | 0,28100806 |
| 100381270_a | ZBED6 | 0,21014358 | 6,24307864 | 1,86491716 | 0,08252415 | 0,28100806 |
| 101929651_a | LINC01398 | -0,15831851 | 3,54652815 | -1,86478161 | 0,08254423 | 0,28102679 |
| 84975_at | MFSD5 | 0,10160604 | 8,34955889 | 1,86457715 | 0,08257454 | 0,28106878 |
| 6135_at | RPL11 | -0,1104475 | 12,3838258 | -1,86446313 | 0,08259145 | 0,28106878 |
| 79159_at | NOL12 | -0,25797747 | 7,25772921 | -1,86440318 | 0,08260034 | 0,28106878 |
| 85236_at | HIST1H2BK | 0,40413836 | 9,28498012 | 1,86427828 | 0,08261887 | 0,28108216 |
| 121355_at | GTSF1 | -0,25745254 | 3,51486477 | -1,86412891 | 0,08264103 | 0,28110791 |
| 6610_at | SMPD2 | -0,15613242 | 7,85471931 | -1,86375193 | 0,08269697 | 0,28124856 |
| 55629_at | PNRC2 | 0,31332281 | 8,21573058 | 1,86303922 | 0,08280284 | 0,2815589 |
| 128229_at | TSACC | -0,17531194 | 6,09781105 | -1,86252886 | 0,08287873 | 0,28176721 |
| 101929297_a | LOC10192929 | -0,1303906 | 3,09484807 | -1,8622999 | 0,0829128 | 0,28183328 |
| 40_at | ASIC2 | -0,17444292 | 4,17615624 | -1,86214241 | 0,08293623 | 0,28186321 |
| 1562_at | CYP2C18 | 0,37222685 | 4,46613757 | 1,8619169 | 0,08296981 | 0,28189254 |
| 50808_at | AK3 | 0,11763159 | 9,33176497 | 1,86188786 | 0,08297413 | 0,28189254 |
| 5646_at | PRSS3 | -0,14565964 | 5,88952614 | -1,861324 | 0,08305813 | 0,28206336 |
| 9987_at | HNRNPDL | -0,25928932 | 8,98091457 | -1,86131529 | 0,08305943 | 0,28206336 |
| 55103_at | RALGPS2 | 0,56280373 | 6,74281968 | 1,86125549 | 0,08306834 | 0,28206336 |
| 55313_at | CPPED1 | 0,3055983 | 6,08606781 | 1,86073224 | 0,08314637 | 0,28227856 |
| 23025_at | UNC13A | -0,15519517 | 4,43025179 | -1,86040958 | 0,08319452 | 0,28239224 |
| 8989_at | TRPA1 | -0,13683007 | 3,92047398 | -1,85967193 | 0,08330468 | 0,28267022 |
| 100506047_a | LOC10050604 | -0,20499972 | 4,32875076 | -1,85963241 | 0,08331059 | 0,28267022 |
| 1996_at | ELAVL4 | -0,1444796 | 3,36650663 | -1,85956654 | 0,08332043 | 0,28267022 |
| 27034_at | ACAD8 | 0,15225119 | 7,26083213 | 1,8592776 | 0,08336363 | 0,28276697 |
| 284948_at | SH2D6 | -0,25502709 | 4,69772458 | -1,85899121 | 0,08340647 | 0,28282377 |
| 10561_at | IFI44 | 1,51519816 | 7,27648482 | 1,85895516 | 0,08341186 | 0,28282377 |
| 284759_at | SIRPB2 | -0,21975372 | 3,62983952 | -1,85887118 | 0,08342442 | 0,28282377 |
| 223117_at | SEMA3D | 0,2115051 | 4,03726306 | 1,8583358 | 0,08350457 | 0,28301132 |
| 24147_at | FJX1 | -0,72260167 | 7,28262515 | -1,85830533 | 0,08350913 | 0,28301132 |
| 11019_at | LIAS | -0,33871585 | 6,84675658 | -1,85806597 | 0,08354498 | 0,28308303 |
| 283431_at | GAS2L3 | -0,41839708 | 6,40141845 | -1,85772086 | 0,0835967 | 0,28320846 |
| 157855_at | KCNU1 | -0,11847688 | 3,86812764 | -1,85758271 | 0,08361742 | 0,28322882 |
| 51244_at | CCDC174 | 0,1713611 | 6,65381598 | 1,85731446 | 0,08365765 | 0,28329014 |
| 125875_at | CLDND2 | -0,26875392 | 5,16246088 | -1,85723139 | 0,08367011 | 0,28329014 |
| 5528_at | PPP2R5D | -0,15210073 | 8,48617897 | -1,85713839 | 0,08368406 | 0,28329014 |
| 25865_at | PRKD2 | 0,17343289 | 8,71808773 | 1,85701786 | 0,08370215 | 0,28329014 |
| 127343_at | DMBX1 | -0,26545892 | 4,57866862 | -1,85697185 | 0,08370905 | 0,28329014 |
| 115584_at | SLC5A11 | -0,21083264 | 4,75897885 | -1,85667889 | 0,08375303 | 0,28335018 |
| 3045_at | HBD | -0,13536149 | 4,06575965 | -1,85657156 | 0,08376915 | 0,28335018 |

| | | | | | | |
|---|---|---|---|---|---|---|
| 9169_at | SCAF11 | 0,25457256 | 8,68228709 | 1,85655475 | 0,08377167 | 0,28335018 |
| 8089_at | YEATS4 | -0,23600746 | 7,47381376 | -1,85646113 | 0,08378573 | 0,28335018 |
| 79029_at | SPATA5L1 | -0,25495349 | 7,57778583 | -1,8563639 | 0,08380034 | 0,28335018 |
| 57719_at | ANO8 | 0,16423329 | 6,41591586 | 1,8557437 | 0,08389356 | 0,28354911 |
| 6764_at | ST5 | 0,21598411 | 7,16110906 | 1,85563468 | 0,08390996 | 0,28354911 |
| 9227_at | LRAT | -0,11761756 | 3,40195625 | -1,85562091 | 0,08391203 | 0,28354911 |
| 221895_at | JAZF1 | 0,3032858 | 6,35027481 | 1,85558086 | 0,08391805 | 0,28354911 |
| 55788_at | LMBRD1 | 0,54462642 | 9,79460233 | 1,85522013 | 0,08397233 | 0,28368274 |
| 284654_at | RSPO1 | -0,1851117 | 5,13303165 | -1,85478642 | 0,08403763 | 0,28385356 |
| 9046_at | DOK2 | -0,20811295 | 5,71060073 | -1,85461147 | 0,08406398 | 0,28388206 |
| 164127_at | CCDC185 | -0,18486899 | 4,38529275 | -1,85441846 | 0,08409307 | 0,28388206 |
| 5727_at | PTCH1 | 0,24668582 | 5,11681267 | 1,85438574 | 0,084098 | 0,28388206 |
| 100874182_a | LINC00424 | -0,18146121 | 5,24020438 | -1,85433918 | 0,08410501 | 0,28388206 |
| 174_at | AFP | -0,12683764 | 4,07269266 | -1,85348515 | 0,08423382 | 0,28423182 |
| 9489_at | PGS1 | 0,14071443 | 7,72010837 | 1,85340605 | 0,08424576 | 0,28423182 |
| 259239_at | WFDC11 | -0,16197969 | 3,96316718 | -1,85335873 | 0,0842529 | 0,28423182 |
| 301_at | ANXA1 | -0,28374566 | 10,9033574 | -1,85312017 | 0,08428892 | 0,28430354 |
| 163087_at | ZNF383 | 0,20144812 | 5,1299274 | 1,85264486 | 0,08436073 | 0,28445567 |
| 64747_at | MFSD1 | 0,43432399 | 10,3319484 | 1,85262614 | 0,08436356 | 0,28445567 |
| 64600_at | PLA2G2F | 0,43627231 | 6,29193099 | 1,85203225 | 0,08445336 | 0,28457925 |
| 90864_at | SPSB3 | 0,45690131 | 7,86998622 | 1,8519986 | 0,08445845 | 0,28457925 |
| 3706_at | ITPKA | -0,22085901 | 5,49573125 | -1,8519718 | 0,0844625 | 0,28457925 |
| 64376_at | IKZF5 | -0,15655171 | 6,45282458 | -1,85191225 | 0,08447152 | 0,28457925 |
| 60490_at | PPCDC | -0,16729187 | 6,99203864 | -1,85189531 | 0,08447408 | 0,28457925 |
| 389812_at | LCN15 | -0,17602606 | 5,74927412 | -1,85070893 | 0,08465376 | 0,28513471 |
| 421_at | ARVCF | -0,15409335 | 6,58088104 | -1,85056511 | 0,08467557 | 0,2851583 |
| 6924_at | TCEB3 | -0,22334636 | 8,67562717 | -1,85033208 | 0,08471091 | 0,28522745 |
| 57594_at | HOMEZ | 0,24402228 | 5,77013733 | 1,85022068 | 0,08472781 | 0,2852345 |
| 988_at | CDC5L | -0,27931067 | 9,23397189 | -1,84996953 | 0,08476592 | 0,28528614 |
| 221044_at | UCMA | -0,21728721 | 4,63828544 | -1,84992442 | 0,08477277 | 0,28528614 |
| 54925_at | ZSCAN32 | 0,10089411 | 6,8081818 | 1,84949664 | 0,08483772 | 0,28545486 |
| 10125_at | RASGRP1 | 0,17659389 | 3,48981431 | 1,84889998 | 0,0849284 | 0,28571004 |
| 140870_at | WFDC6 | -0,20791062 | 4,07248486 | -1,84878486 | 0,0849459 | 0,28571903 |
| 54939_at | COMMD4 | -0,13009164 | 8,36016586 | -1,84841625 | 0,08500197 | 0,28581601 |
| 5311_at | PKD2 | 0,3824642 | 8,07462945 | 1,8483038 | 0,08501908 | 0,28581601 |
| 55312_at | RFK | -0,15392896 | 8,63826021 | -1,84830273 | 0,08501924 | 0,28581601 |
| 727_at | C5 | 0,23565043 | 4,7850822 | 1,84791871 | 0,0850777 | 0,28596263 |
| 5201_at | PFDN1 | -0,14747959 | 7,92232054 | -1,84757096 | 0,08513067 | 0,28601799 |
| 4853_at | NOTCH2 | 0,32354027 | 8,15117862 | 1,84749718 | 0,08514192 | 0,28601799 |
| 4583_at | MUC2 | -0,31603705 | 5,97745927 | -1,84748467 | 0,08514382 | 0,28601799 |
| 55151_at | TMEM38B | -0,84348807 | 7,09823485 | -1,84736834 | 0,08516155 | 0,28601799 |
| 4879_at | NPPB | -0,22453146 | 5,21653649 | -1,84732329 | 0,08516842 | 0,28601799 |
| 100505495_a | PCAT19 | -0,18305121 | 4,07151014 | -1,84677294 | 0,08525234 | 0,28624993 |
| 55279_at | ZNF654 | 0,30601791 | 7,0391134 | 1,84654922 | 0,08528648 | 0,28627271 |
| 81562_at | LMAN2L | -0,15658279 | 8,30656636 | -1,84653368 | 0,08528885 | 0,28627271 |
| 340578_at | DCAF12L2 | 0,58829501 | 4,18447496 | 1,84561455 | 0,08542923 | 0,28669393 |
| 6727_at | SRP14 | 0,11821968 | 10,7093549 | 1,84525601 | 0,08548404 | 0,28682791 |
| 7158_at | TP53BP1 | 0,1541538 | 6,76547441 | 1,84504525 | 0,08551628 | 0,28688611 |
| 54795_at | TRPM4 | 0,21674744 | 7,01892556 | 1,8438988 | 0,08569182 | 0,28742496 |
| 2521_at | FUS | -0,27274093 | 9,76315512 | -1,84348467 | 0,08575531 | 0,28758784 |

| | | | | | | |
|---|---|---|---|---|---|---|
| 6002_at | RGS12 | 0,34494333 | 6,72409182 | 1,84311226 | 0,08581244 | 0,28772934 |
| 101928165_a | CASC17 | -0,24945447 | 3,79795778 | -1,84222134 | 0,08594925 | 0,28813791 |
| 55904_at | KMT2E | 0,24914962 | 9,32702486 | 1,84207748 | 0,08597136 | 0,28816188 |
| 83869_at | TTTY14 | -0,15464887 | 4,4887606 | -1,84164179 | 0,08603835 | 0,28833625 |
| 374868_at | ATP9B | 0,14929226 | 5,99240923 | 1,8415044 | 0,08605949 | 0,28835691 |
| 55095_at | SAMD4B | -0,10659037 | 6,96116238 | -1,84140368 | 0,08607498 | 0,28835868 |
| 101928433_a | LOC10192843 | 0,18551558 | 4,57317609 | 1,84124242 | 0,0860998 | 0,28839166 |
| 25987_at | TSKU | 0,13907459 | 9,55755301 | 1,84103113 | 0,08613232 | 0,28845045 |
| 26539_at | OR10H1 | -0,32959382 | 3,94397834 | -1,84070379 | 0,08618274 | 0,2885691 |
| 10106_at | CTDSP2 | 0,23588984 | 8,83568528 | 1,84048638 | 0,08621623 | 0,28863109 |
| 79587_at | CARS2 | -0,28027451 | 7,14927028 | -1,83996804 | 0,08629614 | 0,2888484 |
| 339487_at | ZBTB8OS | -0,23696553 | 8,21894628 | -1,83971257 | 0,08633555 | 0,28890264 |
| 80010_at | RMI1 | -0,19443912 | 8,13377355 | -1,83966855 | 0,08634234 | 0,28890264 |
| 50489_at | CD207 | -0,16541591 | 3,93421459 | -1,83898583 | 0,08644774 | 0,28920507 |
| 54961_at | SSH3 | 0,20903078 | 9,29424936 | 1,83872475 | 0,08648808 | 0,28928977 |
| 205147_at | AMER3 | -0,15586116 | 4,7075996 | -1,83852929 | 0,08651828 | 0,28934057 |
| 1535_at | CYBA | 0,27274236 | 10,2468681 | 1,83785472 | 0,08662261 | 0,2896392 |
| 101928403_a | LOC10192840 | 0,43544976 | 4,69560163 | 1,8375894 | 0,08666368 | 0,28972621 |
| 6615_at | SNAI1 | -0,21947356 | 4,81781027 | -1,83737318 | 0,08669716 | 0,28978785 |
| 5295_at | PIK3R1 | 0,42247119 | 8,09148605 | 1,83690707 | 0,08676937 | 0,289948 |
| 5979_at | RET | -0,14164422 | 4,84107482 | -1,83682129 | 0,08678266 | 0,289948 |
| 26122_at | EPC2 | 0,49220897 | 8,27453264 | 1,8367468 | 0,08679421 | 0,289948 |
| 1103_at | CHAT | -0,2627446 | 4,92420463 | -1,83667538 | 0,08680528 | 0,289948 |
| 9381_at | OTOF | 0,10522012 | 4,34374923 | 1,83648041 | 0,08683551 | 0,2899987 |
| 25977_at | NECAP1 | 0,16794692 | 8,77056951 | 1,83623486 | 0,0868736 | 0,29006865 |
| 375791_at | CYSRT1 | -0,45722667 | 7,41474832 | -1,83611942 | 0,08689151 | 0,29006865 |
| 3192_at | HNRNPU | -0,15507195 | 7,1938218 | -1,83605421 | 0,08690163 | 0,29006865 |
| 4661_at | MYT1 | -0,13792901 | 5,34263047 | -1,83592559 | 0,08692159 | 0,29008501 |
| 85300_at | ATCAY | -0,19555863 | 4,99798131 | -1,83531098 | 0,08701704 | 0,29028345 |
| 266_at | AMELY | -0,16428826 | 3,63728758 | -1,8352555 | 0,08702566 | 0,29028345 |
| 29967_at | LRP12 | -0,27973083 | 6,52801674 | -1,83512173 | 0,08704645 | 0,29028345 |
| 399671_at | HEATR4 | -0,16587356 | 3,68270476 | -1,83508779 | 0,08705173 | 0,29028345 |
| 27121_at | DKK4 | -0,18695159 | 3,93027612 | -1,83505769 | 0,0870564 | 0,29028345 |
| 28646_at | TRAV36DV7 | -0,21425031 | 3,57611198 | -1,83468714 | 0,08711402 | 0,29042529 |
| 686_at | BTD | 0,24122783 | 6,1327353 | 1,83437603 | 0,08716242 | 0,29053012 |
| 148066_at | ZNRF4 | -0,18876272 | 5,4824855 | -1,83429113 | 0,08717563 | 0,29053012 |
| 54474_at | KRT20 | -0,34527077 | 4,15430377 | -1,83379333 | 0,08725313 | 0,29073811 |
| 84320_at | ACBD6 | -0,18142395 | 7,27686937 | -1,83291451 | 0,0873901 | 0,2910125 |
| 791115_at | PWRN2 | -0,20035451 | 3,72847505 | -1,83288534 | 0,08739465 | 0,2910125 |
| 27043_at | PELP1 | -0,19801302 | 7,04305899 | -1,83285847 | 0,08739884 | 0,2910125 |
| 9743_at | ARHGAP32 | 0,43387855 | 8,08599607 | 1,83284951 | 0,08740024 | 0,2910125 |
| 240_at | ALOX5 | 0,98826525 | 7,99052265 | 1,83266801 | 0,08742855 | 0,2910125 |
| 148811_at | PM20D1 | 0,41209298 | 5,25597333 | 1,83265492 | 0,08743059 | 0,2910125 |
| 429_at | ASCL1 | -0,12603445 | 3,75756079 | -1,83253128 | 0,08744989 | 0,2910125 |
| 101928730_a | LOC10192873 | -0,16751315 | 4,12914252 | -1,83248998 | 0,08745634 | 0,2910125 |
| 9419_at | CRIPT | 0,23365199 | 6,8102259 | 1,83237597 | 0,08747413 | 0,29102145 |
| 388753_at | COA6 | -0,38402811 | 8,66881558 | -1,83211684 | 0,08751459 | 0,29110577 |
| 79632_at | FAM184A | -0,15055574 | 4,8229832 | -1,83195517 | 0,08753984 | 0,29113949 |
| 55176_at | SEC61A2 | -0,21357509 | 5,56517209 | -1,83180287 | 0,08756363 | 0,29116835 |
| 29785_at | CYP2S1 | -0,54972958 | 8,12647643 | -1,83143556 | 0,08762104 | 0,29130895 |

| | | | | | | |
|---|---|---|---|---|---|---|
| 10160_at | FARP1 | 0,47998686 | 7,76978965 | 1,83124471 | 0,08765088 | 0,29134549 |
| 5685_at | PSMA4 | -0,21662904 | 11,1032653 | -1,83117182 | 0,08766228 | 0,29134549 |
| 2260_at | FGFR1 | -0,12908222 | 5,4713952 | -1,83090326 | 0,08770429 | 0,2914305 |
| 3167_at | HMX2 | -0,2033831 | 4,35306599 | -1,83080575 | 0,08771955 | 0,2914305 |
| 6217_at | RPS16 | -0,10750935 | 12,4718588 | -1,83071824 | 0,08773324 | 0,2914305 |
| 29986_at | SLC39A2 | 0,41189516 | 5,9884041 | 1,83059471 | 0,08775258 | 0,29144447 |
| 6662_at | SOX9 | -0,60592432 | 6,61155304 | -1,83011157 | 0,08782824 | 0,29164547 |
| 56159_at | TEX11 | -0,11893294 | 3,06895837 | -1,82997882 | 0,08784904 | 0,29165333 |
| 2015_at | ADGRE1 | -0,18574084 | 4,42376898 | -1,82990322 | 0,08786089 | 0,29165333 |
| 23440_at | OTP | -0,18653352 | 3,91020686 | -1,82943974 | 0,08793355 | 0,29184423 |
| 101928525_a | LOC10192852 | -0,15841359 | 5,21378561 | -1,82914774 | 0,08797935 | 0,29194595 |
| 5457_at | POU4F1 | 0,31482366 | 5,38019 | 1,8281604 | 0,08813439 | 0,29238724 |
| 3849_at | KRT2 | -0,25404889 | 4,54669085 | -1,82810224 | 0,08814353 | 0,29238724 |
| 6205_at | RPS11 | 0,08990776 | 13,5027078 | 1,82801101 | 0,08815787 | 0,29238724 |
| 64377_at | CHST8 | -0,19191429 | 5,5677442 | -1,82751957 | 0,08823515 | 0,29259316 |
| 55893_at | ZNF395 | 0,21451415 | 7,65398161 | 1,82707653 | 0,08830487 | 0,29277396 |
| 9677_at | PPIP5K1 | 0,15446832 | 6,88112309 | 1,82675566 | 0,08835539 | 0,29279574 |
| 414301_at | DDI1 | -0,12586301 | 3,23825454 | -1,82670587 | 0,08836324 | 0,29279574 |
| 120863_at | DEPDC4 | -0,14018313 | 3,20947455 | -1,82666481 | 0,0883697 | 0,29279574 |
| 2444_at | FRK | 0,52012586 | 6,65237429 | 1,82655806 | 0,08838652 | 0,29279574 |
| 10549_at | PRDX4 | -0,25140616 | 9,90804131 | -1,82651258 | 0,08839369 | 0,29279574 |
| 57147_at | SCYL3 | 0,19108266 | 7,3755206 | 1,82637107 | 0,08841598 | 0,29279574 |
| 80789_at | INTS5 | -0,1340417 | 8,22237996 | -1,8263593 | 0,08841784 | 0,29279574 |
| 139596_at | UPRT | 0,28233808 | 6,48850346 | 1,82610158 | 0,08845846 | 0,29284544 |
| 2180_at | ACSL1 | 0,65197568 | 9,34694429 | 1,82600974 | 0,08847295 | 0,29284544 |
| 3754_at | KCNF1 | -0,21729519 | 5,08980864 | -1,82597481 | 0,08847845 | 0,29284544 |
| 83695_at | RHNO1 | -0,16421875 | 9,06308268 | -1,82557322 | 0,0885418 | 0,292945 |
| 54072_at | LINC00158 | -0,14580936 | 3,29127876 | -1,82553213 | 0,08854829 | 0,292945 |
| 95681_at | CEP41 | -0,12045818 | 7,221625 | -1,82549493 | 0,08855416 | 0,292945 |
| 6480_at | ST6GAL1 | 0,32562736 | 4,86160445 | 1,82493953 | 0,08864185 | 0,29318474 |
| 22835_at | ZFP30 | 0,13258918 | 7,00160465 | 1,82438824 | 0,08872897 | 0,2934225 |
| 84206_at | MEX3B | 0,44805583 | 4,74180239 | 1,82423111 | 0,08875381 | 0,29345428 |
| 64577_at | ALDH8A1 | 0,15132192 | 3,40289421 | 1,8241104 | 0,0887729 | 0,29346703 |
| 5929_at | RBBP5 | -0,21762889 | 7,58022635 | -1,82368358 | 0,08884043 | 0,29363988 |
| 439916_at | LINC00615 | -0,15913133 | 3,36120081 | -1,82348828 | 0,08887134 | 0,29369166 |
| 101928104_a | LINC01482 | -0,16522204 | 3,76926895 | -1,82324094 | 0,08891051 | 0,2937707 |
| 63973_at | NEUROG2 | -0,18783576 | 3,97952008 | -1,82301896 | 0,08894568 | 0,29382022 |
| 266629_at | SEC14L3 | -0,17374215 | 3,98417988 | -1,82290016 | 0,0889645 | 0,29382022 |
| 4607_at | MYBPC3 | -0,19245808 | 6,128699 | -1,82285751 | 0,08897126 | 0,29382022 |
| 8993_at | PGLYRP1 | -0,24217188 | 5,55147508 | -1,82244466 | 0,08903671 | 0,29397573 |
| 64599_at | GIGYF1 | 0,13113327 | 6,76474756 | 1,82236796 | 0,08904887 | 0,29397573 |
| 51471_at | NAT8B | -0,13132607 | 4,33957434 | -1,82154359 | 0,0891797 | 0,29427549 |
| 401145_at | CCSER1 | 0,26058606 | 4,34811264 | 1,82145761 | 0,08919336 | 0,29427549 |
| 51305_at | KCNK9 | -0,21403842 | 4,78315801 | -1,82136337 | 0,08920833 | 0,29427549 |
| 3769_at | KCNJ13 | 0,43081035 | 3,38709892 | 1,82134084 | 0,08921191 | 0,29427549 |
| 1089_at | CEACAM4 | -0,20956643 | 6,05195013 | -1,82131472 | 0,08921606 | 0,29427549 |
| 217_at | ALDH2 | 0,25007042 | 4,99290238 | 1,82110371 | 0,08924958 | 0,29431866 |
| 10130_at | PDIA6 | -0,18418894 | 11,000251 | -1,82103995 | 0,08925972 | 0,29431866 |
| 127540_at | HMGB4 | -0,14574302 | 3,64292284 | -1,82094393 | 0,08927498 | 0,29431866 |
| 9774_at | BCLAF1 | -0,23591389 | 6,93335718 | -1,82077343 | 0,08930208 | 0,29435764 |

| | | | | | | |
|---|---|---|---|---|---|---|
| 8692_at | HYAL2 | -0,13846498 | 7,82134596 | -1,82053962 | 0,08933927 | 0,29442982 |
| 8726_at | EED | -0,14983235 | 8,06625617 | -1,82038094 | 0,08936451 | 0,29446264 |
| 9741_at | LAPTM4A | 0,25796372 | 12,0546926 | 1,82007322 | 0,08941348 | 0,29451527 |
| 93649_at | MYOCD | 0,18987119 | 4,27290643 | 1,82004628 | 0,08941777 | 0,29451527 |
| 57584_at | ARHGAP21 | 0,19945745 | 4,80824723 | 1,81996701 | 0,08943039 | 0,29451527 |
| 2736_at | GLI2 | -0,16586241 | 4,92398445 | -1,81989635 | 0,08944164 | 0,29451527 |
| 8335_at | HIST1H2AB | -0,22988198 | 5,57679636 | -1,81958202 | 0,0894917 | 0,29462975 |
| 140730_at | RIMS4 | -0,12322411 | 4,55704617 | -1,81939264 | 0,08952187 | 0,29467872 |
| 27314_at | RAB30 | 0,40597267 | 5,53804286 | 1,81923124 | 0,0895476 | 0,29469472 |
| 257629_at | ANKS4B | -0,15849772 | 4,08286176 | -1,81917017 | 0,08955733 | 0,29469472 |
| 283194_at | LOC283194 | -0,21649783 | 4,45643056 | -1,81906356 | 0,08957433 | 0,2947003 |
| 4211_at | MEIS1 | 0,53375518 | 6,95574765 | 1,81893397 | 0,08959499 | 0,29471795 |
| 100526820_a | CAHM | -0,16512991 | 5,57374668 | -1,81873439 | 0,08962682 | 0,29477232 |
| 4064_at | CD180 | -0,18292176 | 3,83864333 | -1,81835426 | 0,08968748 | 0,29491037 |
| 3698_at | ITIH2 | -0,20868029 | 3,7936704 | -1,8182022 | 0,08971175 | 0,29491037 |
| 2167_at | FABP4 | -2,06994745 | 9,17247247 | -1,81818357 | 0,08971473 | 0,29491037 |
| 219527_at | LRRC55 | -0,18294507 | 4,11556959 | -1,81805781 | 0,08973481 | 0,29492605 |
| 23648_at | SSBP3 | 0,22922701 | 7,05118636 | 1,81779146 | 0,08977735 | 0,29501553 |
| 54848_at | ARHGEF38 | -0,21394172 | 3,87053278 | -1,81742404 | 0,08983606 | 0,29515812 |
| 3224_at | HOXC8 | -0,1735032 | 4,99974903 | -1,81666219 | 0,08995792 | 0,29550807 |
| 517_at | ATP5G2 | 0,12267771 | 9,97739151 | 1,81637297 | 0,09000421 | 0,29560975 |
| 338323_at | NLRP14 | -0,16758992 | 3,88922584 | -1,81623991 | 0,09002552 | 0,29562932 |
| 28410_at | IGHV3-72 | -0,18612286 | 3,81151379 | -1,81582029 | 0,09009274 | 0,29579964 |
| 374986_at | FAM73A | 0,2512074 | 7,62979695 | 1,81561651 | 0,09012541 | 0,29585645 |
| 23059_at | CLUAP1 | 0,44331811 | 6,40237541 | 1,81550227 | 0,09014372 | 0,29586616 |
| 114088_at | TRIM9 | 0,33885715 | 4,07885913 | 1,81513462 | 0,09020268 | 0,29600925 |
| 283152_at | CCDC153 | 0,22295701 | 5,02232787 | 1,81499329 | 0,09022536 | 0,29603323 |
| 834_at | CASP1 | 0,73059506 | 7,21686698 | 1,81431847 | 0,0903337 | 0,29633823 |
| 100996279_a | LINC00269 | -0,15611336 | 4,93113903 | -1,81416289 | 0,0903587 | 0,2963386 |
| 1820_at | ARID3A | 0,39750757 | 6,25488345 | 1,8140467 | 0,09037737 | 0,2963386 |
| 100129620_a | LOC10012962 | 0,1096935 | 3,16951661 | 1,81403053 | 0,09037997 | 0,2963386 |
| 4692_at | NDN | 0,56661346 | 7,35054366 | 1,81381525 | 0,09041457 | 0,29640161 |
| 140880_at | CST11 | -0,16538723 | 3,96469818 | -1,81365448 | 0,09044042 | 0,29642884 |
| 5898_at | RALA | -0,24816454 | 8,87505552 | -1,81357218 | 0,09045365 | 0,29642884 |
| 54509_at | RHOF | -0,32222109 | 5,46827876 | -1,81342069 | 0,09047802 | 0,29645825 |
| 8912_at | CACNA1H | -0,20934331 | 6,34424186 | -1,81223155 | 0,09066949 | 0,29699948 |
| 79109_at | MAPKAP1 | -0,11860843 | 8,08674125 | -1,81220334 | 0,09067403 | 0,29699948 |
| 729970_at | LOC729970 | 0,14598524 | 3,77923811 | 1,81203353 | 0,09070141 | 0,29703863 |
| 23621_at | BACE1 | 0,23038247 | 7,4055573 | 1,81147937 | 0,09079079 | 0,29725058 |
| 84243_at | ZDHHC18 | 0,11911948 | 8,39492708 | 1,81144092 | 0,09079699 | 0,29725058 |
| 5701_at | PSMC2 | -0,16191906 | 8,29941426 | -1,81094905 | 0,09087639 | 0,29742567 |
| 55063_at | ZCWPW1 | 0,13888241 | 5,58552034 | 1,81091834 | 0,09088135 | 0,29742567 |
| 56100_at | PCDHGB6 | -0,15765212 | 3,26915517 | -1,81050849 | 0,09094756 | 0,2975918 |
| 730091_at | LINC00886 | -0,23404361 | 4,89583234 | -1,81027283 | 0,09098566 | 0,29762839 |
| 101928565_a | LOC10192856 | -0,14416505 | 4,78279098 | -1,81024814 | 0,09098965 | 0,29762839 |
| 100130613_a | PRR32 | 0,20928325 | 4,49449866 | 1,80973693 | 0,09107233 | 0,29784415 |
| 3852_at | KRT5 | -1,10092073 | 10,259037 | -1,80956465 | 0,09110021 | 0,29784415 |
| 54756_at | IL17RD | 0,15050228 | 4,57651536 | 1,80955361 | 0,09110199 | 0,29784415 |
| 79183_at | TTPAL | 0,13972796 | 8,37418793 | 1,80930921 | 0,09114156 | 0,29787407 |
| 4296_at | MAP3K11 | 0,19176035 | 8,2830609 | 1,80930602 | 0,09114208 | 0,29787407 |

| | | | | | | |
|---|---|---|---|---|---|---|
| 3938_at | LCT | -0,16064319 | 4,27705298 | -1,80912745 | 0,09117099 | 0,29791804 |
| 7552_at | ZNF711 | 0,5490472 | 4,7797485 | 1,80891262 | 0,09120579 | 0,2979812 |
| 3709_at | ITPR2 | 0,25314546 | 5,48935354 | 1,80815221 | 0,09132906 | 0,29833334 |
| 51529_at | ANAPC11 | -0,1570674 | 7,90137603 | -1,80771694 | 0,09139969 | 0,29842493 |
| 51809_at | GALNT7 | 0,35053755 | 9,39616672 | 1,8076278 | 0,09141416 | 0,29842493 |
| 9488_at | PIGB | 0,26957672 | 7,81854425 | 1,80762407 | 0,09141476 | 0,29842493 |
| 10930_at | APOBEC2 | -0,17441702 | 4,7131117 | -1,80759756 | 0,09141907 | 0,29842493 |
| 55223_at | TRIM62 | 0,16027516 | 6,73265206 | 1,80706412 | 0,09150571 | 0,29865714 |
| 79033_at | ERI3 | -0,20686481 | 7,6889653 | -1,80620591 | 0,09164525 | 0,29906191 |
| 93650_at | ACPT | -0,19797579 | 6,4682629 | -1,80576844 | 0,09171645 | 0,29924357 |
| 56985_at | ADPRM | 0,11409818 | 5,46038162 | 1,80512748 | 0,09182087 | 0,2995335 |
| 8655_at | DYNLL1 | -0,09319051 | 11,9715556 | -1,80495019 | 0,09184976 | 0,29954814 |
| 10482_at | NXF1 | -0,17326418 | 8,82587663 | -1,80486265 | 0,09186404 | 0,29954814 |
| 1595_at | CYP51A1 | -0,38644984 | 9,04498299 | -1,8048026 | 0,09187383 | 0,29954814 |
| 8312_at | AXIN1 | -0,12719853 | 7,20773366 | -1,80471843 | 0,09188755 | 0,29954814 |
| 5292_at | PIM1 | -0,16462355 | 8,65622132 | -1,80421657 | 0,09196943 | 0,29968285 |
| 105378828_a | LOC10537882 | 0,34825113 | 5,62673151 | 1,80416589 | 0,09197771 | 0,29968285 |
| 55056_at | FLJ10038 | 0,17536121 | 5,87746442 | 1,80414657 | 0,09198086 | 0,29968285 |
| 9576_at | SPAG6 | -0,11628018 | 3,37581653 | -1,8040429 | 0,09199778 | 0,29968285 |
| 196446_at | MYRFL | 0,32128294 | 3,42669875 | 1,80394785 | 0,0920133 | 0,29968285 |
| 7803_at | PTP4A1 | -0,2641179 | 8,61605051 | -1,80389323 | 0,09202222 | 0,29968285 |
| 126321_at | MFSD12 | -0,24364694 | 7,77467257 | -1,80359047 | 0,09207167 | 0,29979322 |
| 23204_at | ARL6IP1 | -0,24503108 | 12,2352976 | -1,80328785 | 0,09212113 | 0,2998644 |
| 84219_at | WDR24 | 0,21108294 | 6,63214309 | 1,80321321 | 0,09213333 | 0,2998644 |
| 8372_at | HYAL3 | -0,25928383 | 5,75085366 | -1,80317095 | 0,09214024 | 0,2998644 |
| 3785_at | KCNQ2 | -0,1590341 | 5,04674789 | -1,80301715 | 0,09216539 | 0,29987695 |
| 23344_at | ESYT1 | -0,25047733 | 8,87277991 | -1,80291714 | 0,09218174 | 0,29987695 |
| 28954_at | REM1 | -0,2461628 | 5,38704754 | -1,80286177 | 0,0921908 | 0,29987695 |
| 11282_at | MGAT4B | 0,12783461 | 9,89904075 | 1,80225699 | 0,09228976 | 0,30014819 |
| 1032_at | CDKN2D | -0,21380666 | 7,07973101 | -1,80200501 | 0,09233103 | 0,3002233 |
| 79799_at | UGT2A3 | 0,15220085 | 2,93900166 | 1,80192562 | 0,09234403 | 0,3002233 |
| 10252_at | SPRY1 | 0,33597351 | 6,69199048 | 1,80169109 | 0,09238246 | 0,30029755 |
| 8875_at | VNN2 | 0,59573213 | 4,47865634 | 1,80135271 | 0,09243792 | 0,30042714 |
| 55578_at | SUPT20H | 0,18196885 | 8,92675833 | 1,80120557 | 0,09246205 | 0,30045486 |
| 1555_at | CYP2B6 | -0,53003379 | 4,64781318 | -1,80005489 | 0,09265092 | 0,30101782 |
| 55351_at | STK32B | -0,17224739 | 4,12902141 | -1,79875782 | 0,09286423 | 0,30165998 |
| 6504_at | SLAMF1 | -0,15671662 | 3,91161961 | -1,79858345 | 0,09289294 | 0,30170237 |
| 758_at | MPPED1 | -0,15098621 | 4,92641778 | -1,79831972 | 0,09293638 | 0,30179256 |
| 84665_at | MYPN | 0,3547809 | 5,0735908 | 1,7981692 | 0,09296118 | 0,30182221 |
| 137964_at | GPAT4 | 0,20488786 | 6,96529413 | 1,7978906 | 0,0930071 | 0,30192041 |
| 390874_at | ONECUT3 | -0,19726002 | 5,99690065 | -1,79759281 | 0,0930562 | 0,30201998 |
| 9789_at | SPCS2 | -0,1628347 | 4,10242503 | -1,79751443 | 0,09306913 | 0,30201998 |
| 203111_at | ERICH5 | 0,57580328 | 4,17595456 | 1,79689575 | 0,09317123 | 0,30230037 |
| 23467_at | NPTXR | 0,64720221 | 7,44271946 | 1,7966399 | 0,09321348 | 0,30238652 |
| 5156_at | PDGFRA | -0,12412397 | 3,30024751 | -1,79626569 | 0,0932753 | 0,30253615 |
| 112858_at | TP53RK | -0,09762206 | 7,76122526 | -1,79606897 | 0,09330782 | 0,3025436 |
| 3382_at | ICA1 | 0,1462671 | 5,5350016 | 1,79601258 | 0,09331714 | 0,3025436 |
| 808_at | CALM3 | 0,18953391 | 9,61458923 | 1,79596677 | 0,09332472 | 0,3025436 |
| 166793_at | ZBTB49 | 0,11088391 | 6,123389 | 1,79564385 | 0,09337813 | 0,30258483 |
| 11273_at | ATXN2L | -0,17706552 | 6,35752473 | -1,79563972 | 0,09337881 | 0,30258483 |

| | | | | | | |
|---|---|---|---|---|---|---|
| 337880_at | KRTAP11-1 | -0,22937372 | 5,29662005 | -1,79547722 | 0,0934057 | 0,30258483 |
| 80896_at | NPL | 0,37145367 | 6,35887579 | 1,79542826 | 0,0934138 | 0,30258483 |
| 11259_at | FILIP1L | 0,16323925 | 4,13170158 | 1,79541075 | 0,0934167 | 0,30258483 |
| 64978_at | MRPL38 | -0,22154058 | 7,94396542 | -1,79523059 | 0,09344651 | 0,30258483 |
| 5797_at | PTPRM | 0,72380751 | 9,46466991 | 1,79516313 | 0,09345768 | 0,30258483 |
| 388662_at | SLC6A17 | -0,23025859 | 6,25718729 | -1,7951304 | 0,0934631 | 0,30258483 |
| 11052_at | CPSF6 | -0,17529886 | 7,91476344 | -1,79488987 | 0,09350293 | 0,30266291 |
| 3007_at | HIST1H1D | -0,19550677 | 6,5588924 | -1,79449114 | 0,09356899 | 0,30282586 |
| 80228_at | ORAI2 | 0,15038461 | 6,78033424 | 1,79428453 | 0,09360324 | 0,30288581 |
| 114804_at | RNF157 | -0,24973858 | 4,63730418 | -1,79407904 | 0,09363731 | 0,30294518 |
| 3014_at | H2AFX | -0,67851674 | 8,88292678 | -1,7937831 | 0,0936864 | 0,3030531 |
| 54838_at | WBP1L | 0,21609276 | 8,23103491 | 1,79340812 | 0,09374864 | 0,30320349 |
| 84246_at | MED10 | -0,17864427 | 8,0233808 | -1,79316099 | 0,09378967 | 0,30328529 |
| 79611_at | ACSS3 | -0,13107745 | 3,39087984 | -1,79290393 | 0,09383237 | 0,30337245 |
| 11262_at | SP140 | 0,15592354 | 3,59403206 | 1,7927094 | 0,0938647 | 0,30342603 |
| 131831_at | ERICH6 | -0,1950355 | 3,93785595 | -1,79253191 | 0,0938942 | 0,30346322 |
| 64770_at | CCDC14 | -0,25863413 | 5,52762041 | -1,79245064 | 0,09390771 | 0,30346322 |
| 26190_at | FBXW2 | 0,22627914 | 8,31344101 | 1,79183242 | 0,09401055 | 0,30374459 |
| 55640_at | FLVCR2 | 0,26993322 | 5,77806142 | 1,79158157 | 0,09405231 | 0,30381289 |
| 2259_at | FGF14 | -0,12705927 | 3,61853816 | -1,79138896 | 0,09408438 | 0,30381289 |
| 26085_at | KLK13 | -0,17563226 | 5,11841542 | -1,79122092 | 0,09411237 | 0,30381289 |
| 22871_at | NLGN1 | 0,17707067 | 3,38832562 | 1,79116004 | 0,09412251 | 0,30381289 |
| 83538_at | TTC25 | 0,1837343 | 4,34451538 | 1,79115815 | 0,09412282 | 0,30381289 |
| 285548_at | LINC01096 | -0,22276463 | 3,9120591 | -1,79113717 | 0,09412632 | 0,30381289 |
| 1314_at | COPA | 0,17651915 | 9,03298479 | 1,79103375 | 0,09414355 | 0,3038176 |
| 57502_at | NLGN4X | -0,17300289 | 4,33406703 | -1,79085462 | 0,09417341 | 0,30385962 |
| 9047_at | SH2D2A | -0,2852395 | 6,20816195 | -1,79072455 | 0,09419509 | 0,30385962 |
| 6623_at | SNCG | 0,58109497 | 7,88827764 | 1,79067171 | 0,0942039 | 0,30385962 |
| 10825_at | NEU3 | -0,16396845 | 5,99210736 | -1,79047893 | 0,09423605 | 0,30390929 |
| 4035_at | LRP1 | 0,21720451 | 6,73235694 | 1,79039017 | 0,09425085 | 0,30390929 |
| 100132618_a | ZRANB2-AS1 | -0,14567728 | 3,94284137 | -1,78992621 | 0,09432827 | 0,30410802 |
| 7798_at | LUZP1 | 0,17469088 | 7,90875095 | 1,78955216 | 0,09439072 | 0,30422682 |
| 267002_at | PGBD2 | 0,17607748 | 5,59375692 | 1,78951635 | 0,0943967 | 0,30422682 |
| 84466_at | MEGF10 | 0,15940587 | 3,78015611 | 1,78866263 | 0,0945394 | 0,30460594 |
| 3200_at | HOXA3 | 0,10296054 | 6,30037225 | 1,78862337 | 0,09454596 | 0,30460594 |
| 80341_at | BPIFB2 | -0,19047528 | 5,05901788 | -1,78840581 | 0,09458236 | 0,30467224 |
| 201626_at | PDE12 | -0,35375645 | 7,00543235 | -1,78818373 | 0,09461952 | 0,304741 |
| 124411_at | ZNF720 | 0,22361618 | 7,91286072 | 1,78807505 | 0,09463772 | 0,30474864 |
| 92259_at | MRPS36 | -0,19842767 | 8,0934725 | -1,78795218 | 0,09465829 | 0,30476393 |
| 400713_at | ZNF880 | 0,60671453 | 5,36993451 | 1,78784577 | 0,09467611 | 0,30477037 |
| 23568_at | ARL2BP | 0,21806422 | 9,25896659 | 1,78762383 | 0,09471328 | 0,30483909 |
| 148109_at | FAM187B | -0,20026651 | 4,63261225 | -1,78619744 | 0,09495251 | 0,30555799 |
| 10079_at | ATP9A | 0,72395054 | 9,45885808 | 1,78610294 | 0,09496837 | 0,30555801 |
| 3780_at | KCNN1 | -0,21987458 | 5,34592259 | -1,78585112 | 0,09501067 | 0,30558526 |
| 4975_at | OMP | -0,18791758 | 3,77371881 | -1,78584447 | 0,09501179 | 0,30558526 |
| 201299_at | RDM1 | -0,46152809 | 6,12724335 | -1,78576918 | 0,09502444 | 0,30558526 |
| 55885_at | LMO3 | 0,21799567 | 4,82243646 | 1,78564175 | 0,09504585 | 0,3056031 |
| 54769_at | DIRAS2 | 0,1235678 | 3,77329027 | 1,78539588 | 0,09508717 | 0,30560388 |
| 323_at | APBB2 | 0,16320434 | 6,65344991 | 1,78535355 | 0,09509429 | 0,30560388 |
| 84820_at | POLR2J4 | -0,16949064 | 4,81070475 | -1,78531438 | 0,09510088 | 0,30560388 |

| | | | | | | |
|---|---|---|---|---|---|---|
| 1357_at | CPA1 | -0,20866467 | 3,83133639 | -1,78526279 | 0,09510955 | 0,30560388 |
| 103344929_a | BGLT3 | -0,23802539 | 4,2960119 | -1,78501078 | 0,09515193 | 0,30568908 |
| 26103_at | LRIT1 | -0,19863813 | 4,13704173 | -1,78465962 | 0,09521102 | 0,30582789 |
| 124093_at | CCDC78 | -0,19066469 | 4,3767688 | -1,78396743 | 0,09532758 | 0,30615125 |
| 6873_at | TAF2 | -0,19385073 | 8,64437209 | -1,78375179 | 0,09536392 | 0,3062169 |
| 57216_at | VANGL2 | 0,69713602 | 5,95069118 | 1,78337249 | 0,09542787 | 0,30635761 |
| 5375_at | PMP2 | -0,10323316 | 3,08408451 | -1,78330323 | 0,09543955 | 0,30635761 |
| 10544_at | PROCR | -0,51409881 | 7,6033703 | -1,7829742 | 0,09549506 | 0,30644029 |
| 130502_at | TTC32 | 0,14925962 | 6,60388192 | 1,78296198 | 0,09549712 | 0,30644029 |
| 5256_at | PHKA2 | 0,30205559 | 6,94074031 | 1,78280606 | 0,09552344 | 0,30647368 |
| 84294_at | UTP23 | -0,13640223 | 7,41984077 | -1,7825583 | 0,09556527 | 0,3065095 |
| 22937_at | SCAP | 0,13288054 | 8,60842114 | 1,78255143 | 0,09556643 | 0,3065095 |
| 23592_at | LEMD3 | -0,10340309 | 9,01576993 | -1,78213077 | 0,09563748 | 0,30666092 |
| 339287_at | MSL1 | -0,16686013 | 9,43613736 | -1,78208346 | 0,09564548 | 0,30666092 |
| 101060553_a | LOC10106055 | -0,18359259 | 4,5195132 | -1,78168497 | 0,09571284 | 0,30682582 |
| 157567_at | ANKRD46 | 0,51621746 | 8,04339072 | 1,7815163 | 0,09574136 | 0,30686619 |
| 51234_at | EMC4 | -0,29318196 | 11,1306069 | -1,78136087 | 0,09576765 | 0,3068994 |
| 219902_at | TMEM136 | 0,22111109 | 5,70538981 | 1,78113723 | 0,09580549 | 0,30691173 |
| 441151_at | TMEM151B | -0,16480884 | 5,15314123 | -1,78111479 | 0,09580929 | 0,30691173 |
| 389161_at | ANKUB1 | -0,12319399 | 4,56209797 | -1,78105565 | 0,0958193 | 0,30691173 |
| 100507527_a | THEG5 | -0,13869101 | 3,34632736 | -1,78069306 | 0,09588069 | 0,30705731 |
| 57609_at | DIP2B | 0,21999256 | 8,20511106 | 1,780198 | 0,09596456 | 0,30722881 |
| 9960_at | USP3 | 0,14187148 | 8,68011599 | 1,78018868 | 0,09596614 | 0,30722881 |
| 729438_at | GATSL2 | 0,28716872 | 7,43972369 | 1,77998431 | 0,09600078 | 0,30728865 |
| 134359_at | POC5 | -0,36384999 | 6,92970957 | -1,77963445 | 0,09606012 | 0,30742748 |
| 55108_at | BSDC1 | 0,29363344 | 8,12499724 | 1,7793827 | 0,09610283 | 0,30751309 |
| 26235_at | FBXL4 | 0,17153436 | 7,65035663 | 1,77907665 | 0,09615478 | 0,30762821 |
| 388419_at | BTBD17 | -0,18599281 | 4,73111213 | -1,77860211 | 0,09623537 | 0,30783494 |
| 79786_at | KLHL36 | 0,15376315 | 7,26637111 | 1,77826315 | 0,09629298 | 0,30796807 |
| 94104_at | PAXBP1 | -0,31953259 | 7,69532094 | -1,77791352 | 0,09635243 | 0,30806842 |
| 30009_at | TBX21 | -0,20934742 | 5,18199903 | -1,77789053 | 0,09635634 | 0,30806842 |
| 79575_at | ABHD8 | 0,23478504 | 6,05172711 | 1,77776939 | 0,09637695 | 0,30808317 |
| 80326_at | WNT10A | 0,88785111 | 8,36369142 | 1,77744131 | 0,09643278 | 0,30821049 |
| 90007_at | MIDN | 0,23409989 | 7,22439232 | 1,7772297 | 0,09646881 | 0,30827449 |
| 4210_at | MEFV | -0,24554348 | 5,30429232 | -1,77703203 | 0,09650247 | 0,30833091 |
| 92345_at | NAF1 | -0,28147747 | 6,80777943 | -1,77645131 | 0,09660142 | 0,30859589 |
| 51035_at | UBXN1 | 0,09486718 | 7,01617583 | 1,77620293 | 0,09664378 | 0,30867999 |
| 10767_at | HBS1L | -0,16478507 | 6,9220414 | -1,77553749 | 0,09675732 | 0,30899143 |
| 221143_at | N6AMT2 | -0,23066424 | 7,41923179 | -1,77517708 | 0,09681887 | 0,30908722 |
| 5301_at | PIN1P1 | -0,12424217 | 6,08334972 | -1,7751307 | 0,09682679 | 0,30908722 |
| 28786_at | IGLV4-3 | -0,13257348 | 5,06387374 | -1,77501367 | 0,09684679 | 0,30908722 |
| 1990_at | CELA1 | -0,15529729 | 5,14677606 | -1,77498607 | 0,0968515 | 0,30908722 |
| 340895_at | MALRD1 | -0,20338321 | 3,35638383 | -1,77460969 | 0,09691583 | 0,30924128 |
| 5683_at | PSMA2 | -0,26721718 | 10,0044364 | -1,77441493 | 0,09694913 | 0,30929632 |
| 10936_at | GPR75 | -0,17095739 | 4,5513574 | -1,77425995 | 0,09697564 | 0,30932965 |
| 7976_at | FZD3 | 0,60211807 | 6,65242707 | 1,77393794 | 0,09703074 | 0,30943345 |
| 266727_at | MDGA1 | 0,16455889 | 7,22585352 | 1,77388201 | 0,09704031 | 0,30943345 |
| 84419_at | C15orf48 | 0,36876657 | 4,76805488 | 1,77320708 | 0,09715589 | 0,30975074 |
| 11093_at | ADAMTS13 | 0,15714014 | 5,91978154 | 1,77216475 | 0,09733463 | 0,31026924 |
| 6123_at | RPL3L | -0,18435815 | 4,77537459 | -1,77164904 | 0,09742318 | 0,3105001 |

| | | | | | | |
|---|---|---|---|---|---|---|
| 5816_at | PVALB | -0,16696695 | 3,28826968 | -1,77150939 | 0,09744716 | 0,31052518 |
| 727699_at | LINC00163 | -0,19610631 | 5,23624695 | -1,77106458 | 0,09752361 | 0,31067009 |
| 9074_at | CLDN6 | -0,19261497 | 4,61995044 | -1,7710571 | 0,09752489 | 0,31067009 |
| 3191_at | HNRNPL | -0,1105324 | 9,9939103 | -1,77066097 | 0,09759302 | 0,31083569 |
| 23131_at | GPATCH8 | 0,1057023 | 8,03203751 | 1,77044464 | 0,09763024 | 0,31089845 |
| 9967_at | THRAP3 | -0,22361596 | 8,23738038 | -1,77035886 | 0,097645 | 0,31089845 |
| 27243_at | CHMP2A | 0,20637625 | 9,909413 | 1,76982684 | 0,0977366 | 0,31104162 |
| 4675_at | NAP1L3 | 0,61698796 | 4,12907494 | 1,76980471 | 0,09774041 | 0,31104162 |
| 27245_at | AHDC1 | -0,12267692 | 5,81723426 | -1,76977016 | 0,09774636 | 0,31104162 |
| 201625_at | DNAH12 | -0,13077816 | 3,2005967 | -1,76966791 | 0,09776398 | 0,31104162 |
| 10101_at | NUBP2 | -0,11588704 | 8,44366751 | -1,76962889 | 0,0977707 | 0,31104162 |
| 3306_at | HSPA2 | -0,2217495 | 5,65956947 | -1,76952289 | 0,09778896 | 0,31104835 |
| 55184_at | DZANK1 | 0,15771549 | 5,00619223 | 1,76923033 | 0,09783939 | 0,31112595 |
| 10296_at | MAEA | -0,20152906 | 9,71306555 | -1,76915172 | 0,09785295 | 0,31112595 |
| 100132815_a | IPO5P1 | 0,46280354 | 6,91369358 | 1,76910029 | 0,09786182 | 0,31112595 |
| 285464_at | CRIPAK | 0,225917 | 7,86005715 | 1,76893405 | 0,09789049 | 0,31113562 |
| 729230_at | CCR2 | -0,14599422 | 3,63985265 | -1,76889535 | 0,09789716 | 0,31113562 |
| 9022_at | CLIC3 | 1,1790888 | 7,75090942 | 1,76853771 | 0,09795887 | 0,31125458 |
| 7127_at | TNFAIP2 | 0,33211882 | 9,71236612 | 1,76849115 | 0,0979669 | 0,31125458 |
| 3958_at | LGALS3 | 0,61108745 | 10,4998027 | 1,76836169 | 0,09798925 | 0,31127424 |
| 127534_at | GJB4 | -0,24098664 | 5,65091906 | -1,76798314 | 0,09805462 | 0,31143054 |
| 10234_at | LRRC17 | 0,11784999 | 3,68269097 | 1,76779968 | 0,09808632 | 0,31147985 |
| 140875_at | LINC00028 | -0,22331512 | 5,08116483 | -1,76760955 | 0,09811918 | 0,31153284 |
| 7419_at | VDAC3 | -0,45352648 | 10,6734335 | -1,7674143 | 0,09815293 | 0,31155665 |
| 9551_at | ATP5J2 | -0,25096382 | 11,7172227 | -1,76731917 | 0,09816938 | 0,31155665 |
| 9255_at | AIMP1 | -0,25721234 | 8,18018214 | -1,76728551 | 0,0981752 | 0,31155665 |
| 1386_at | ATF2 | 0,11244858 | 7,3435766 | 1,76709757 | 0,0982077 | 0,31160847 |
| 64073_at | C19orf33 | 0,19333588 | 9,80546911 | 1,76666299 | 0,0982829 | 0,31174698 |
| 196394_at | AMN1 | 0,2296289 | 5,70150973 | 1,76658481 | 0,09829644 | 0,31174698 |
| 645_at | BLVRB | -0,11056703 | 10,1891685 | -1,76651455 | 0,0983086 | 0,31174698 |
| 58157_at | NGB | -0,1349918 | 5,53212869 | -1,76631854 | 0,09834254 | 0,31174698 |
| 8367_at | HIST1H4E | -0,32528733 | 7,24581662 | -1,76630853 | 0,09834428 | 0,31174698 |
| 7916_at | PRRC2A | -0,15584901 | 8,56473459 | -1,76628437 | 0,09834846 | 0,31174698 |
| 377677_at | CA13 | 0,29864192 | 5,68152586 | 1,76600404 | 0,09839702 | 0,3118496 |
| 3275_at | PRMT2 | 0,28687936 | 7,5357714 | 1,76567071 | 0,0984548 | 0,31198137 |
| 27294_at | DHDH | -0,15629537 | 5,54059198 | -1,76537375 | 0,09850629 | 0,3120932 |
| 51366_at | UBR5 | 0,19597364 | 8,74676669 | 1,76525094 | 0,09852759 | 0,31210936 |
| 653857_at | ACTR3C | 0,15867616 | 7,04248261 | 1,76500608 | 0,09857008 | 0,3121926 |
| 6509_at | SLC1A4 | -0,44833572 | 7,21950843 | -1,76484782 | 0,09859755 | 0,31220932 |
| 11180_at | WDR6 | 0,23834341 | 7,43152172 | 1,7647889 | 0,09860777 | 0,31220932 |
| 8405_at | SPOP | 0,33678469 | 8,05311585 | 1,76439031 | 0,09867699 | 0,31234635 |
| 83548_at | COG3 | 0,18572978 | 6,57602953 | 1,76435295 | 0,09868348 | 0,31234635 |
| 5522_at | PPP2R2C | -0,35467487 | 5,44076263 | -1,76385487 | 0,09877004 | 0,31256896 |
| 6578_at | SLCO2A1 | -0,21960681 | 4,59046298 | -1,76366337 | 0,09880334 | 0,31262298 |
| 285331_at | CCDC66 | -0,14006635 | 5,44460999 | -1,76310155 | 0,09890108 | 0,31288086 |
| 255394_at | TCP11L2 | 0,23522818 | 5,05327793 | 1,76289628 | 0,09893682 | 0,31294251 |
| 285084_at | LINC01305 | 0,11600772 | 3,59382135 | 1,76241539 | 0,09902058 | 0,31315603 |
| 64098_at | PARVG | -0,16685385 | 4,82043134 | -1,7617067 | 0,09914413 | 0,3134642 |
| 55269_at | PSPC1 | 0,10976792 | 7,24621361 | 1,76166978 | 0,09915057 | 0,3134642 |
| 644150_at | WIPF3 | -0,19428248 | 5,57341313 | -1,76153732 | 0,09917368 | 0,31347964 |

| | | | | | | |
|---|---|---|---|---|---|---|
| 5016_at | OVGP1 | 0,10200504 | 5,73276874 | 1,76140952 | 0,09919598 | 0,31347964 |
| 388939_at | C2orf71 | -0,21415562 | 4,82505799 | -1,761362 | 0,09920427 | 0,31347964 |
| 1946_at | EFNA5 | -0,20607132 | 7,90824457 | -1,76119738 | 0,09923301 | 0,31351901 |
| 201283_at | AMZ2P1 | 0,47963501 | 5,80378656 | 1,76108201 | 0,09925315 | 0,31353122 |
| 122258_at | SPACA7 | -0,21155531 | 3,79974933 | -1,76092795 | 0,09928005 | 0,31356309 |
| 4637_at | MYL6 | 0,12366826 | 12,7022977 | 1,76083427 | 0,09929641 | 0,31356309 |
| 390892_at | OR7A10 | -0,1126056 | 2,73001865 | -1,76074462 | 0,09931207 | 0,31356309 |
| 7783_at | ZP2 | -0,1585298 | 4,2949093 | -1,76049414 | 0,09935584 | 0,31364987 |
| 79065_at | ATG9A | 0,22772087 | 8,80294038 | 1,76029114 | 0,09939132 | 0,31371047 |
| 9152_at | SLC6A5 | -0,23679312 | 4,86558627 | -1,76001428 | 0,09943973 | 0,31374972 |
| 1479_at | CSTF3 | -0,38980137 | 7,78263164 | -1,75996041 | 0,09944915 | 0,31374972 |
| 4205_at | MEF2A | 0,11771477 | 7,50390225 | 1,75992859 | 0,09945472 | 0,31374972 |
| 7827_at | NPHS2 | -0,24378749 | 4,73344976 | -1,75984748 | 0,09946891 | 0,31374972 |
| 79927_at | FAM110D | -0,14553052 | 5,64041061 | -1,75958903 | 0,09951413 | 0,31384097 |
| 27127_at | SMC1B | -0,15302796 | 3,15111151 | -1,75940209 | 0,09954685 | 0,31389277 |
| 3037_at | HAS2 | -0,61969583 | 4,61614287 | -1,75930501 | 0,09956384 | 0,31389498 |
| 9407_at | TMPRSS11D | -0,1711295 | 4,80897512 | -1,75883483 | 0,0996462 | 0,31403995 |
| 84693_at | MCEE | 0,1781177 | 6,21717516 | 1,75876027 | 0,09965926 | 0,31403995 |
| 93979_at | CPA5 | -0,22938 | 4,81775585 | -1,75870821 | 0,09966838 | 0,31403995 |
| 8737_at | RIPK1 | 0,34261463 | 7,57731815 | 1,75864912 | 0,09967874 | 0,31403995 |
| 100505518_a | EDNRB-AS1 | -0,15567868 | 3,23457835 | -1,75857359 | 0,09969198 | 0,31403995 |
| 57336_at | ZNF287 | 0,4966752 | 4,46281629 | 1,75841104 | 0,09972047 | 0,31403995 |
| 112942_at | CFAP36 | -0,17080445 | 7,10691255 | -1,75832263 | 0,09973597 | 0,31403995 |
| 389332_at | LOC389332 | -0,1885294 | 5,07803363 | -1,75829824 | 0,09974025 | 0,31403995 |
| 7084_at | TK2 | -0,36833883 | 5,88303245 | -1,75801563 | 0,09978982 | 0,31414468 |
| 643641_at | ZNF862 | 0,33341723 | 6,15562441 | 1,75752972 | 0,0998751 | 0,31436176 |
| 6926_at | TBX3 | 0,77643015 | 9,20477851 | 1,75740011 | 0,09989785 | 0,31438202 |
| 126969_at | SLC44A3 | 0,2656985 | 9,14814052 | 1,75690197 | 0,09998536 | 0,31456416 |
| 64078_at | SLC28A3 | 0,16760451 | 3,55713282 | 1,75679947 | 0,10000338 | 0,31456416 |
| 10826_at | FAXDC2 | -0,12710088 | 5,07704513 | -1,75679182 | 0,10000472 | 0,31456416 |
| 100506421_a | LINC01158 | -0,2188527 | 4,7991392 | -1,75666469 | 0,10002707 | 0,31458309 |
| 2665_at | GDI2 | -0,32902836 | 11,013114 | -1,75638262 | 0,10007667 | 0,3146877 |
| 5539_at | PPY | -0,18126093 | 4,25985359 | -1,75527395 | 0,10027183 | 0,31524991 |
| 200894_at | ARL13B | -0,46220943 | 6,9545281 | -1,75517169 | 0,10028984 | 0,3152551 |
| 51599_at | LSR | 0,21198959 | 9,38661664 | 1,75497977 | 0,10032367 | 0,31530997 |
| 9526_at | MPDU1 | -0,12207227 | 9,71737689 | -1,75487417 | 0,10034228 | 0,31531703 |
| 22795_at | NID2 | -0,14232422 | 3,59897185 | -1,75468476 | 0,10037568 | 0,31537052 |
| 9182_at | RASSF9 | -0,13858506 | 2,93339468 | -1,75442524 | 0,10042145 | 0,31546288 |
| 200765_at | TIGD1 | -0,32357123 | 5,99116633 | -1,75427384 | 0,10044816 | 0,31548381 |
| 2295_at | FOXF2 | -0,28649716 | 8,53490135 | -1,75413009 | 0,10047353 | 0,31548381 |
| 10900_at | RUNDC3A | -0,1584858 | 5,73724973 | -1,75410904 | 0,10047725 | 0,31548381 |
| 9802_at | DAZAP2 | 0,19399693 | 10,6337343 | 1,75315737 | 0,10064534 | 0,31594668 |
| 84077_at | C3orf20 | -0,17283094 | 4,3493111 | -1,75305588 | 0,10066328 | 0,31594668 |
| 84467_at | FBN3 | -0,17083863 | 5,49724815 | -1,75299599 | 0,10067387 | 0,31594668 |
| 401324_at | LOC401324 | -0,13318226 | 4,200132 | -1,75286215 | 0,10069753 | 0,31595527 |
| 115024_at | NT5C3B | -0,16951372 | 9,14481264 | -1,752795 | 0,10070941 | 0,31595527 |
| 6601_at | SMARCC2 | 0,1531786 | 8,26951639 | 1,75264338 | 0,10073623 | 0,31598794 |
| 497634_at | LINC00293 | -0,25241237 | 3,95656621 | -1,75232684 | 0,10079224 | 0,31606834 |
| 51497_at | NELFCD | -0,17348171 | 10,2729712 | -1,75222531 | 0,10081021 | 0,31606834 |
| 402415_at | XKRX | -0,20028004 | 3,94695081 | -1,75215501 | 0,10082265 | 0,31606834 |

| | | | | | | |
|---|---|---|---|---|---|---|
| 54221_at | SNTG2 | -0,14705047 | 3,54008567 | -1,75205372 | 0,10084059 | 0,31606834 |
| 79781_at | IQCA1 | 0,4477212 | 5,59720024 | 1,752035 | 0,1008439 | 0,31606834 |
| 7059_at | THBS3 | 0,45079856 | 6,68301014 | 1,75113449 | 0,10100346 | 0,31649987 |
| 284422_at | SMIM24 | -0,24833037 | 4,87588565 | -1,75105658 | 0,10101728 | 0,31649987 |
| 64859_at | NABP1 | -0,21170959 | 7,5022364 | -1,7509799 | 0,10103087 | 0,31649987 |
| 8092_at | ALX1 | -0,11551417 | 4,44299599 | -1,75060285 | 0,10109776 | 0,31665793 |
| 171484_at | FAM9C | -0,13512952 | 3,40731007 | -1,75034894 | 0,10114283 | 0,31674209 |
| 55818_at | KDM3A | 0,20745958 | 8,31321452 | 1,7502662 | 0,10115752 | 0,31674209 |
| 55036_at | CCDC40 | -0,17137719 | 4,8992856 | -1,74975812 | 0,10124777 | 0,31697314 |
| 5790_at | PTPRCAP | 0,16576176 | 6,116174 | 1,74913153 | 0,10135916 | 0,3172445 |
| 9392_at | TGFBRAP1 | -0,1450919 | 7,16808949 | -1,7490853 | 0,10136738 | 0,3172445 |
| 375449_at | MAST4 | 0,25200926 | 7,44216279 | 1,74851059 | 0,10146965 | 0,31741225 |
| 11022_at | TDRKH | 0,12514936 | 4,72421412 | 1,74846635 | 0,10147753 | 0,31741225 |
| 23186_at | RCOR1 | -0,17919439 | 8,06295475 | -1,74843811 | 0,10148256 | 0,31741225 |
| 102723661_a | LOC10272366 | -0,15465288 | 4,36153372 | -1,74841376 | 0,1014869 | 0,31741225 |
| 84656_at | GLYR1 | 0,15662689 | 7,48232217 | 1,74810375 | 0,10154211 | 0,31751793 |
| 100289373_a | UBAC2-AS1 | -0,23896929 | 5,48292559 | -1,74803896 | 0,10155365 | 0,31751793 |
| 84889_at | SLC7A3 | -0,18234176 | 3,73882085 | -1,74788774 | 0,1015806 | 0,31755064 |
| 57210_at | SLC45A4 | -0,48215596 | 6,13774817 | -1,74698759 | 0,10174112 | 0,31800083 |
| 38_at | ACAT1 | -0,41403367 | 6,83965424 | -1,74657643 | 0,10181451 | 0,31817861 |
| 11079_at | RER1 | -0,1177838 | 9,3299701 | -1,74641766 | 0,10184287 | 0,31821559 |
| 79144_at | PPDPF | 0,18550319 | 10,6062344 | 1,74617437 | 0,10188633 | 0,31829976 |
| 255520_at | ELMOD2 | 0,35567839 | 7,08881241 | 1,74577981 | 0,10195685 | 0,31842298 |
| 10987_at | COPS5 | -0,20454997 | 9,37391365 | -1,74570345 | 0,1019705 | 0,31842298 |
| 9135_at | RABEP1 | 0,17028081 | 6,70472908 | 1,74567627 | 0,10197536 | 0,31842298 |
| 7104_at | TM4SF4 | -0,16560768 | 4,32529405 | -1,74543082 | 0,10201926 | 0,31850843 |
| 79829_at | NAA40 | -0,22052903 | 7,77449278 | -1,74510497 | 0,10207756 | 0,31863881 |
| 54461_at | FBXW5 | 0,12627093 | 8,52239305 | 1,74498927 | 0,10209827 | 0,31865181 |
| 56242_at | ZNF253 | 0,27912806 | 5,89280476 | 1,74467956 | 0,10215372 | 0,31872507 |
| 58498_at | MYL7 | -0,18165327 | 5,70752328 | -1,74463676 | 0,10216139 | 0,31872507 |
| 2258_at | FGF13 | 0,48614109 | 4,72566439 | 1,74458097 | 0,10217138 | 0,31872507 |
| 3185_at | HNRNPF | -0,27086878 | 9,56681135 | -1,74446829 | 0,10219156 | 0,31873642 |
| 643529_at | LINC00865 | 0,48055618 | 5,19124699 | 1,74380255 | 0,10231089 | 0,31905693 |
| 200316_at | APOBEC3F | 0,44139326 | 5,69862564 | 1,74303039 | 0,10244944 | 0,3194198 |
| 79228_at | THOC6 | -0,22389972 | 9,08386036 | -1,7429693 | 0,10246041 | 0,3194198 |
| 51249_at | TMEM69 | -0,17684857 | 9,74234941 | -1,74265764 | 0,10251639 | 0,31954259 |
| 10783_at | NEK6 | -0,51230596 | 7,61432706 | -1,74249038 | 0,10254644 | 0,31956211 |
| 285498_at | RNF212 | -0,22891001 | 5,4028696 | -1,74243812 | 0,10255583 | 0,31956211 |
| 3222_at | HOXC5 | -0,20848851 | 5,67519929 | -1,74202904 | 0,10262937 | 0,31973953 |
| 27185_at | DISC1 | -0,20146384 | 4,31290588 | -1,74187645 | 0,10265681 | 0,31975131 |
| 55972_at | SLC25A40 | -0,31319047 | 7,64085708 | -1,74182342 | 0,10266635 | 0,31975131 |
| 100506581_a | C16orf95 | -0,11340256 | 6,14885875 | -1,74165493 | 0,10269666 | 0,31979402 |
| 3761_at | KCNJ4 | -0,1615933 | 4,81949017 | -1,74144075 | 0,1027352 | 0,31984617 |
| 28425_at | IGHV3-47 | 0,17578943 | 4,14136022 | 1,74137734 | 0,10274662 | 0,31984617 |
| 7411_at | VBP1 | -0,31982147 | 9,402315 | -1,74127242 | 0,1027655 | 0,31985327 |
| 6343_at | SCT | -0,24227131 | 4,73115418 | -1,74114723 | 0,10278805 | 0,31987175 |
| 9508_at | ADAMTS3 | 0,13212672 | 3,45012346 | 1,7410502 | 0,10280552 | 0,31987444 |
| 8036_at | SHOC2 | 0,17431191 | 8,7765261 | 1,74084136 | 0,10284313 | 0,31993981 |
| 154881_at | KCTD7 | 0,17600823 | 5,43210707 | 1,74066654 | 0,10287463 | 0,31998612 |
| 5210_at | PFKFB4 | -0,14985877 | 6,61682051 | -1,74048841 | 0,10290674 | 0,3200343 |

| | | | | | | |
|---|---|---|---|---|---|---|
| 23413_at | NCS1 | -0,17622471 | 7,50690774 | -1,74032355 | 0,10293646 | 0,32007505 |
| 55052_at | MRPL20 | -0,25839738 | 10,5572289 | -1,74007453 | 0,10298136 | 0,320163 |
| 116071_at | BATF2 | 0,28192948 | 5,6741812 | 1,73990223 | 0,10301244 | 0,32020795 |
| 101928135_a | LOC10192813 | -0,12279687 | 3,22502459 | -1,73972667 | 0,10304412 | 0,32025473 |
| 347853_at | TBX10 | -0,14004712 | 5,01853487 | -1,73960263 | 0,1030665 | 0,32027264 |
| 55915_at | LANCL2 | -0,10002554 | 7,54219202 | -1,73939057 | 0,10310479 | 0,32033992 |
| 23136_at | EPB41L3 | 0,16775174 | 3,91285847 | 1,73893706 | 0,1031867 | 0,32054271 |
| 56180_at | MOSPD1 | 0,25075764 | 7,26196295 | 1,73807155 | 0,10334318 | 0,32097705 |
| 3972_at | LHB | -0,23372948 | 6,45762207 | -1,7378781 | 0,10337819 | 0,32103401 |
| 60492_at | CCDC90B | 0,16941314 | 6,45199589 | 1,7371658 | 0,10350716 | 0,32138273 |
| 101927869_a | LOC10192786 | -0,11019382 | 2,75035066 | -1,7370122 | 0,10353499 | 0,32141734 |
| 8754_at | ADAM9 | 0,36744911 | 10,2072543 | 1,73671794 | 0,10358833 | 0,3215311 |
| 135114_at | HINT3 | 0,28064283 | 6,95232644 | 1,73632677 | 0,10365927 | 0,32169945 |
| 5510_at | PPP1R7 | -0,10678687 | 9,29938444 | -1,73606538 | 0,1037067 | 0,3217668 |
| 55862_at | ECHDC1 | 0,19059845 | 9,24287838 | 1,73602306 | 0,10371438 | 0,3217668 |
| 100505573_a | INAFM2 | -0,3295468 | 7,86273443 | -1,73560458 | 0,10379035 | 0,32195065 |
| 4817_at | NIT1 | 0,17432975 | 7,62556111 | 1,73532587 | 0,10384098 | 0,32201547 |
| 51465_at | UBE2J1 | 0,12380799 | 8,41615774 | 1,73528728 | 0,10384799 | 0,32201547 |
| 5575_at | PRKAR1B | -0,19380563 | 6,27809398 | -1,73521348 | 0,1038614 | 0,32201547 |
| 8277_at | TKTL1 | -0,17800653 | 4,12915932 | -1,73493935 | 0,10391122 | 0,3221181 |
| 100506161_a | TAF1A-AS1 | 0,15036345 | 4,97449912 | 1,73481068 | 0,10393462 | 0,32212095 |
| 1195_at | CLK1 | 0,2843657 | 7,96952356 | 1,73475036 | 0,10394559 | 0,32212095 |
| 645832_at | SEBOX | -0,15271721 | 5,07258792 | -1,73448279 | 0,10399425 | 0,32215377 |
| 84557_at | MAP1LC3A | 0,12594934 | 5,82732519 | 1,73442305 | 0,10400512 | 0,32215377 |
| 51538_at | ZCCHC17 | -0,2104375 | 7,973761 | -1,7344163 | 0,10400635 | 0,32215377 |
| 58510_at | PRODH2 | -0,15362202 | 5,06051573 | -1,73418564 | 0,10404832 | 0,32219656 |
| 92344_at | GORAB | 0,2255889 | 6,98154093 | 1,73415656 | 0,10405361 | 0,32219656 |
| 7295_at | TXN | -0,23821603 | 11,2602678 | -1,73397266 | 0,10408709 | 0,32224841 |
| 5361_at | PLXNA1 | 0,16120519 | 8,76346688 | 1,73359532 | 0,10415581 | 0,32240935 |
| 1573_at | CYP2J2 | 0,2858981 | 8,63295374 | 1,73339046 | 0,10419313 | 0,32247306 |
| 2641_at | GCG | -0,15706284 | 3,25854987 | -1,73274191 | 0,10431138 | 0,32271187 |
| 10003_at | NAALAD2 | 0,12389272 | 2,98470767 | 1,73273175 | 0,10431323 | 0,32271187 |
| 253827_at | MSRB3 | 0,14230623 | 4,11279506 | 1,73269161 | 0,10432055 | 0,32271187 |
| 23063_at | WAPL | -0,14477257 | 8,61479951 | -1,73238586 | 0,10437634 | 0,32279825 |
| 8646_at | CHRD | -0,16778385 | 5,25118951 | -1,73235489 | 0,10438199 | 0,32279825 |
| 198437_at | LKAAEAR1 | -0,22573816 | 6,50995204 | -1,73173705 | 0,10449481 | 0,32309527 |
| 4294_at | MAP3K10 | -0,12400122 | 7,2239068 | -1,73148784 | 0,10454035 | 0,32315097 |
| 54487_at | DGCR8 | -0,14126561 | 7,48911697 | -1,73145485 | 0,10454638 | 0,32315097 |
| 6874_at | TAF4 | -0,1463068 | 7,89151358 | -1,73135583 | 0,10456448 | 0,32315506 |
| 7068_at | THRB | 0,61939403 | 7,31873876 | 1,73118428 | 0,10459584 | 0,32320013 |
| 2687_at | GGT5 | -0,16592114 | 5,31903223 | -1,73101129 | 0,10462747 | 0,32324603 |
| 1417_at | CRYBB3 | -0,18699167 | 5,28694067 | -1,73041495 | 0,10473659 | 0,32346439 |
| 27178_at | IL37 | -0,13250715 | 4,43499874 | -1,73037331 | 0,10474421 | 0,32346439 |
| 57189_at | KIAA1147 | 0,25844273 | 7,57311149 | 1,73033101 | 0,10475196 | 0,32346439 |
| 55705_at | IPO9 | -0,29796508 | 7,55678169 | -1,73025802 | 0,10476532 | 0,32346439 |
| 1264_at | CNN1 | -0,18291329 | 4,18999793 | -1,73013164 | 0,10478846 | 0,32348399 |
| 51303_at | FKBP11 | -0,32090231 | 7,49077321 | -1,72954993 | 0,10489504 | 0,32372904 |
| 92211_at | CDHR1 | -0,13688006 | 4,46615604 | -1,72951496 | 0,10490145 | 0,32372904 |
| 92285_at | ZNF585B | 0,18534372 | 6,51026048 | 1,72908947 | 0,10497947 | 0,32391793 |
| 128346_at | C1orf162 | -0,17111702 | 4,49351858 | -1,72886481 | 0,10502069 | 0,32399321 |

| | | | | | | |
|---|---|---|---|---|---|---|
| 5551_at | PRF1 | -0,18676149 | 5,68748601 | -1,72873477 | 0,10504456 | 0,32400029 |
| 9427_at | ECEL1 | -0,19094863 | 5,26056507 | -1,72851557 | 0,10508479 | 0,32400029 |
| 3646_at | EIF3E | -0,15139132 | 10,6275949 | -1,72850962 | 0,10508589 | 0,32400029 |
| 64799_at | IQCH | 0,09112029 | 3,76811426 | 1,72848576 | 0,10509027 | 0,32400029 |
| 253959_at | RALGAPA1 | 0,36064233 | 7,87993049 | 1,7272981 | 0,10530852 | 0,32455196 |
| 28676_at | TRAV10 | -0,15631109 | 3,62934598 | -1,72720849 | 0,10532501 | 0,32455196 |
| 55342_at | STRBP | -0,16227474 | 6,28502349 | -1,72719221 | 0,105328 | 0,32455196 |
| 55650_at | PIGV | 0,20866303 | 8,51057447 | 1,72714548 | 0,1053366 | 0,32455196 |
| 254295_at | PHYHD1 | -0,19060412 | 4,53592121 | -1,72657429 | 0,10544173 | 0,32477854 |
| 9033_at | PKD2L1 | -0,16381352 | 5,06066772 | -1,72656275 | 0,10544386 | 0,32477854 |
| 55039_at | TRMT12 | -0,15913506 | 7,62540828 | -1,72609763 | 0,10552953 | 0,32499047 |
| 57628_at | DPP10 | 0,1187335 | 2,91378218 | 1,72563851 | 0,10561417 | 0,32519912 |
| 4919_at | ROR1 | -0,41518171 | 4,99631155 | -1,72513171 | 0,10570766 | 0,32536896 |
| 9166_at | EBAG9 | -0,15062501 | 9,18637033 | -1,72506536 | 0,1057199 | 0,32536896 |
| 2053_at | EPHX2 | 0,26943721 | 4,4386259 | 1,72506485 | 0,10572 | 0,32536896 |
| 26009_at | ZZZ3 | -0,175729 | 8,29073726 | -1,72491082 | 0,10574843 | 0,32540448 |
| 83648_at | FAM167A | -0,11958543 | 4,8857732 | -1,72439476 | 0,10584374 | 0,32558005 |
| 56181_at | MTFR1L | 0,41433957 | 8,06251901 | 1,72438432 | 0,10584567 | 0,32558005 |
| 23438_at | HARS2 | 0,1791613 | 9,02198215 | 1,72432735 | 0,10585619 | 0,32558005 |
| 387885_at | CCDC42B | -0,29947839 | 5,61315474 | -1,72409022 | 0,10590002 | 0,32566283 |
| 57795_at | BRINP2 | -0,15847505 | 4,7845237 | -1,72348202 | 0,10601249 | 0,32580147 |
| 8538_at | BARX2 | -0,24769418 | 5,30744732 | -1,7234669 | 0,10601529 | 0,32580147 |
| 5971_at | RELB | 0,33884553 | 6,09588187 | 1,7234562 | 0,10601727 | 0,32580147 |
| 5148_at | PDE6G | -0,24986493 | 6,32624265 | -1,72343118 | 0,10602189 | 0,32580147 |
| 101928461_a | LOC10192846 | -0,19980808 | 4,11333426 | -1,7233892 | 0,10602966 | 0,32580147 |
| 1486_at | CTBS | 0,34944036 | 7,54455472 | 1,72326994 | 0,10605173 | 0,32581731 |
| 84969_at | TOX2 | -0,51548036 | 6,65051187 | -1,72314333 | 0,10607517 | 0,32583734 |
| 3429_at | IFI27 | 1,32984638 | 7,80962659 | 1,72291379 | 0,10611766 | 0,32591267 |
| 2954_at | GSTZ1 | 0,32632801 | 7,58583878 | 1,7227699 | 0,10614431 | 0,32591267 |
| 4326_at | MMP17 | -0,19598261 | 6,16361319 | -1,72256957 | 0,10618142 | 0,32591267 |
| 81629_at | TSSK3 | -0,1977797 | 6,27879214 | -1,72248651 | 0,10619681 | 0,32591267 |
| 55164_at | SHQ1 | -0,3508603 | 7,99259248 | -1,7224865 | 0,10619681 | 0,32591267 |
| 2566_at | GABRG2 | -0,12787222 | 3,65049281 | -1,7224628 | 0,1062012 | 0,32591267 |
| 550113_at | LOC550113 | -0,17251747 | 4,50550704 | -1,7223115 | 0,10622924 | 0,32594679 |
| 55024_at | BANK1 | 0,34291039 | 4,36333539 | 1,72203286 | 0,1062809 | 0,32605335 |
| 8744_at | TNFSF9 | -0,41425685 | 6,81696732 | -1,72176949 | 0,10632974 | 0,32615125 |
| 101928100_a | LOC10192810 | 0,43839721 | 7,63082585 | 1,72160691 | 0,10635991 | 0,32619183 |
| 1669_at | DEFA4 | -0,14190136 | 3,73376607 | -1,72130106 | 0,10641667 | 0,32631396 |
| 10225_at | CD96 | 0,69114905 | 4,81243364 | 1,72111677 | 0,10645088 | 0,32633796 |
| 282890_at | ZNF311 | 0,39241535 | 5,16130492 | 1,7210764 | 0,10645838 | 0,32633796 |
| 8330_at | HIST1H2AK | -0,32274774 | 6,65913665 | -1,72063518 | 0,10654034 | 0,32653724 |
| 10599_at | SLCO1B1 | -0,13150832 | 4,04535908 | -1,72018761 | 0,10662354 | 0,32674024 |
| 4286_at | MITF | 0,26014185 | 7,35874004 | 1,71970406 | 0,10671348 | 0,32696387 |
| 5867_at | RAB4A | 0,21085076 | 8,44616208 | 1,71950137 | 0,10675121 | 0,32702743 |
| 80149_at | ZC3H12A | 0,42060154 | 7,78876981 | 1,71928489 | 0,10679151 | 0,32709888 |
| 158257_at | MIRLET7DHG | -0,25104886 | 3,94067628 | -1,71883793 | 0,10687476 | 0,32730184 |
| 9659_at | PDE4DIP | 0,5073244 | 7,05282841 | 1,71847218 | 0,10694293 | 0,32745855 |
| 28755_at | TRAC | -0,13003204 | 4,0101091 | -1,71816475 | 0,10700026 | 0,32758201 |
| 26071_at | FAM127B | 0,25250526 | 8,70408092 | 1,71776115 | 0,10707557 | 0,32776047 |
| 127544_at | RNF19B | 0,11680022 | 8,58545187 | 1,71762343 | 0,10710127 | 0,32778707 |

| | | | | | | |
|---|---|---|---|---|---|---|
| 139324_at | HDX | 0,67420324 | 4,75017362 | 1,71740236 | 0,10714255 | 0,32784771 |
| 84905_at | ZNF341 | 0,12896532 | 6,65495401 | 1,71733196 | 0,10715569 | 0,32784771 |
| 283392_at | TRHDE-AS1 | 0,14860937 | 3,74699969 | 1,71716528 | 0,10718683 | 0,32784771 |
| 10136_at | CELA3A | -0,22371156 | 5,02633967 | -1,71715277 | 0,10718916 | 0,32784771 |
| 55327_at | LIN7C | -0,23358371 | 8,58550931 | -1,71629605 | 0,10734931 | 0,32828541 |
| 641384_at | TMEM75 | -0,1964216 | 3,85552339 | -1,71614656 | 0,10737728 | 0,32831882 |
| 98_at | ACYP2 | 0,37796862 | 7,49268724 | 1,71578435 | 0,10744507 | 0,32847394 |
| 54765_at | TRIM44 | 0,21547478 | 8,88134852 | 1,71546007 | 0,10750579 | 0,3285818 |
| 270_at | AMPD1 | -0,12770574 | 3,58649755 | -1,71541231 | 0,10751473 | 0,3285818 |
| 11196_at | SEC23IP | -0,14752227 | 8,0924144 | -1,71532269 | 0,10753152 | 0,3285818 |
| 414918_at | DENND6B | 0,23129152 | 5,45840091 | 1,71512587 | 0,1075684 | 0,32864235 |
| 10673_at | TNFSF13B | 0,85413859 | 5,03193345 | 1,71475732 | 0,10763748 | 0,32880126 |
| 52_at | ACP1 | -0,1615866 | 9,36184994 | -1,71452548 | 0,10768095 | 0,32886522 |
| 51564_at | HDAC7 | 0,14971311 | 7,45063413 | 1,7144636 | 0,10769256 | 0,32886522 |
| 994_at | CDC25B | -0,35632484 | 8,73972755 | -1,71425307 | 0,10773206 | 0,32893369 |
| 23245_at | ASTN2 | 0,28384317 | 4,63426634 | 1,71409738 | 0,10776128 | 0,32897036 |
| 5098_at | PCDHGC3 | -0,09721028 | 5,26849273 | -1,71400707 | 0,10777823 | 0,32897036 |
| 54531_at | MIER2 | 0,11474687 | 7,46864297 | 1,71387277 | 0,10780344 | 0,32899519 |
| 57414_at | RHBDD2 | 0,1954216 | 9,73635708 | 1,71367877 | 0,10783987 | 0,32905423 |
| 90737_at | PAGE5 | -0,23205727 | 5,00663143 | -1,71357065 | 0,10786017 | 0,32906407 |
| 8325_at | FZD8 | -0,1224359 | 5,69629376 | -1,71335662 | 0,10790038 | 0,32913461 |
| 118813_at | ZFYVE27 | -0,11567778 | 7,1282364 | -1,7129826 | 0,10797068 | 0,32929689 |
| 79841_at | AGBL2 | 0,36387457 | 3,92427953 | 1,71269933 | 0,10802394 | 0,32940719 |
| 962_at | CD48 | -0,17772746 | 4,76538797 | -1,7115342 | 0,10824328 | 0,33002379 |
| 56955_at | MEPE | -0,1740662 | 4,62863687 | -1,71097538 | 0,10834862 | 0,33024882 |
| 23137_at | SMC5 | -0,17970827 | 7,6118407 | -1,71096075 | 0,10835138 | 0,33024882 |
| 2767_at | GNA11 | -0,2555157 | 9,25166479 | -1,71072094 | 0,10839661 | 0,33027868 |
| 8216_at | LZTR1 | 0,11330724 | 6,9480835 | 1,71064539 | 0,10841086 | 0,33027868 |
| 150590_at | C2orf15 | 0,14170498 | 4,91931055 | 1,71063612 | 0,10841261 | 0,33027868 |
| 113802_at | HENMT1 | 0,19712229 | 7,34424822 | 1,71049172 | 0,10843986 | 0,33029558 |
| 23585_at | TMEM50A | 0,22578704 | 10,1325977 | 1,71033446 | 0,10846954 | 0,33029558 |
| 101927503_a | LOC10192750 | -0,19516593 | 5,94009404 | -1,71024683 | 0,10848608 | 0,33029558 |
| 22859_at | ADGRL1 | -0,18518097 | 7,01622848 | -1,710205 | 0,10849398 | 0,33029558 |
| 7043_at | TGFB3 | -0,15895457 | 5,94059435 | -1,7101525 | 0,10850389 | 0,33029558 |
| 101929478_a | LINGO1-AS2 | -0,1817101 | 4,60491469 | -1,70948401 | 0,10863017 | 0,33055847 |
| 1025_at | CDK9 | 0,14480969 | 7,40455779 | 1,70942905 | 0,10864056 | 0,33055847 |
| 283455_at | KSR2 | 0,22917609 | 4,380365 | 1,70942287 | 0,10864173 | 0,33055847 |
| 5451_at | POU2F1 | -0,13492861 | 7,13841602 | -1,70912064 | 0,10869887 | 0,3306801 |
| 10319_at | LAMC3 | -0,17935546 | 6,01729817 | -1,70899606 | 0,10872244 | 0,33068871 |
| 1457_at | CSNK2A1 | -0,1570089 | 8,83891621 | -1,70892414 | 0,10873604 | 0,33068871 |
| 56957_at | OTUD7B | 0,09154715 | 6,27512672 | 1,70878001 | 0,10876331 | 0,33071943 |
| 79369_at | B3GNT4 | -0,17656591 | 6,64290866 | -1,70848325 | 0,10881947 | 0,33083797 |
| 1230_at | CCR1 | 0,26298415 | 4,21803488 | 1,70831309 | 0,10885168 | 0,33088369 |
| 6795_at | AURKC | -0,17296423 | 5,63918524 | -1,70818914 | 0,10887515 | 0,33090282 |
| 5444_at | PON1 | 0,16520252 | 3,42033201 | 1,70780377 | 0,10894816 | 0,33107246 |
| 55171_at | TBCCD1 | -0,20853773 | 7,38879539 | -1,70760189 | 0,10898642 | 0,33113649 |
| 339751_at | MLK7-AS1 | -0,20126513 | 3,62108952 | -1,70735609 | 0,10903301 | 0,33122583 |
| 1793_at | DOCK1 | 0,15758981 | 6,83630282 | 1,70718623 | 0,10906523 | 0,33127144 |
| 259_at | AMBP | -0,1521432 | 5,41801753 | -1,70625621 | 0,10924175 | 0,33175528 |
| 10910_at | SUGT1 | -0,18106671 | 8,71976736 | -1,70609877 | 0,10927165 | 0,3317938 |

| | | | | | | |
|---|---|---|---|---|---|---|
| 7763_at | ZFAND5 | 0,15091036 | 9,91995933 | 1,70593852 | 0,1093021 | 0,33183395 |
| 102682016_a | LINC01159 | -0,14901464 | 4,8867455 | -1,70571659 | 0,10934428 | 0,33190969 |
| 55615_at | PRR5 | -0,1231103 | 7,39138245 | -1,70538323 | 0,10940767 | 0,33204977 |
| 100996539_a | ITPR1-AS1 | 0,24348357 | 4,25405811 | 1,70499762 | 0,10948103 | 0,33222007 |
| 81533_at | ITFG1 | 0,25315597 | 8,9536223 | 1,70426818 | 0,10961992 | 0,33258914 |
| 923_at | CD6 | -0,14956517 | 5,1666628 | -1,70394607 | 0,1096813 | 0,33265368 |
| 29035_at | C16orf72 | 0,12065097 | 8,65026698 | 1,70388627 | 0,10969269 | 0,33265368 |
| 5662_at | PSD | -0,1672782 | 5,21271055 | -1,70370788 | 0,1097267 | 0,33265368 |
| 100507530_a | MCPH1-AS1 | -0,17293048 | 4,71095163 | -1,70365545 | 0,1097367 | 0,33265368 |
| 100873954_a | SNRK-AS1 | 0,20176555 | 4,46249263 | 1,70362873 | 0,1097418 | 0,33265368 |
| 10158_at | PDZK1IP1 | 0,27356297 | 5,09197151 | 1,70357052 | 0,1097529 | 0,33265368 |
| 4778_at | NFE2 | -0,14422948 | 3,82405538 | -1,7035224 | 0,10976207 | 0,33265368 |
| 158506_at | ZNF645 | -0,11398329 | 3,93649003 | -1,70326374 | 0,10981142 | 0,33275088 |
| 3122_at | HLA-DRA | 0,83555107 | 5,23550985 | 1,70317212 | 0,1098289 | 0,33275151 |
| 303_at | ANXA2P1 | -0,15421231 | 6,09335544 | -1,70251786 | 0,10995381 | 0,33307757 |
| 56475_at | RPRM | 0,17078368 | 4,761415 | 1,70231401 | 0,10999276 | 0,33314315 |
| 26228_at | STAP1 | -0,11067879 | 3,05423383 | -1,70216712 | 0,11002083 | 0,33316668 |
| 54764_at | ZRANB1 | 0,09931389 | 8,09868589 | 1,70209237 | 0,11003511 | 0,33316668 |
| 8834_at | TMEM11 | -0,35631203 | 7,8907053 | -1,70171491 | 0,11010728 | 0,3333328 |
| 84447_at | SYVN1 | 0,1139491 | 8,21569887 | 1,70101318 | 0,11024156 | 0,33366698 |
| 4257_at | MGST1 | 0,15949771 | 11,5609531 | 1,70086708 | 0,11026953 | 0,33366698 |
| 9649_at | RALGPS1 | 0,16678539 | 5,92773699 | 1,70083942 | 0,11027483 | 0,33366698 |
| 221786_at | FAM200A | 0,27632182 | 6,24260743 | 1,70077609 | 0,11028696 | 0,33366698 |
| 127255_at | LRRIQ3 | 0,2253908 | 3,52437814 | 1,70024999 | 0,11038776 | 0,3339195 |
| 80851_at | SH3BP5L | 0,13883683 | 6,90176561 | 1,69991005 | 0,11045293 | 0,33405672 |
| 340526_at | RGAG4 | 0,19298888 | 5,37015887 | 1,69980629 | 0,11047283 | 0,33405672 |
| 2358_at | FPR2 | -0,14605976 | 4,43088466 | -1,69974209 | 0,11048515 | 0,33405672 |
| 2059_at | EPS8 | 0,24960393 | 10,7738434 | 1,69952415 | 0,11052696 | 0,3341307 |
| 4048_at | LTA4H | -0,1563285 | 10,074026 | -1,69919518 | 0,1105901 | 0,33422291 |
| 57045_at | TWSG1 | 0,13384426 | 8,48142674 | 1,69918443 | 0,11059216 | 0,33422291 |
| 642394_at | ADARB2-AS1 | -0,19350616 | 5,08344015 | -1,69898097 | 0,11063123 | 0,33428853 |
| 101927550_a | LOC10192755 | -0,13611661 | 4,68459441 | -1,69863849 | 0,11069702 | 0,33438681 |
| 23363_at | OBSL1 | 0,18152092 | 6,43349035 | 1,69856923 | 0,11071033 | 0,33438681 |
| 23307_at | FKBP15 | 0,09375701 | 7,83195378 | 1,69854058 | 0,11071583 | 0,33438681 |
| 222_at | ALDH3B2 | 0,17239047 | 5,86729726 | 1,69806997 | 0,1108063 | 0,33460759 |
| 51466_at | EVL | 0,24963988 | 6,83602007 | 1,69787458 | 0,11084388 | 0,33462132 |
| 23299_at | BICD2 | -0,15435239 | 8,06707463 | -1,69786569 | 0,11084559 | 0,33462132 |
| 27304_at | MOCS3 | -0,0936625 | 7,26662675 | -1,69767613 | 0,11088206 | 0,33467897 |
| 3117_at | HLA-DQA1 | -0,20467596 | 3,53677426 | -1,69726221 | 0,11096174 | 0,33486698 |
| 8732_at | RNGTT | -0,34320431 | 7,30060876 | -1,6969571 | 0,1110205 | 0,33495923 |
| 101928191_a | LOC10192819 | -0,1540793 | 4,19493634 | -1,6967729 | 0,11105599 | 0,33495923 |
| 11027_at | LILRA2 | -0,13263662 | 3,9080777 | -1,69675619 | 0,11105921 | 0,33495923 |
| 23252_at | OTUD3 | -0,38324124 | 6,40026414 | -1,69673852 | 0,11106261 | 0,33495923 |
| 133022_at | TRAM1L1 | 0,47533813 | 5,03415441 | 1,69665218 | 0,11107925 | 0,33495923 |
| 10329_at | TMEM5 | -0,30836684 | 8,01479193 | -1,69649379 | 0,11110978 | 0,33497645 |
| 8632_at | DNAH17 | -0,11092466 | 4,02172569 | -1,69644212 | 0,11111974 | 0,33497645 |
| 731_at | C8A | -0,12182818 | 3,30631454 | -1,69608133 | 0,11118931 | 0,33513374 |
| 5742_at | PTGS1 | -0,18346701 | 5,16500929 | -1,69581804 | 0,11124011 | 0,33523439 |
| 23650_at | TRIM29 | 0,13035661 | 10,9261539 | 1,69549528 | 0,1113024 | 0,33536966 |
| 282980_at | LINC00700 | -0,13898602 | 3,74878888 | -1,69520109 | 0,11135921 | 0,33541279 |

| | | | | | | |
|---|---|---|---|---|---|---|
| 283601_at | LINC00523 | -0,21019732 | 5,39123636 | -1,69517957 | 0,11136337 | 0,33541279 |
| 509_at | ATP5C1 | -0,29313007 | 10,7014026 | -1,69490822 | 0,11141579 | 0,33541279 |
| 64065_at | PERP | 0,18676048 | 11,5924173 | 1,69485646 | 0,11142579 | 0,33541279 |
| 1284_at | COL4A2 | -0,38130792 | 6,66309224 | -1,69483762 | 0,11142943 | 0,33541279 |
| 7711_at | ZNF155 | 0,14699776 | 4,97533788 | 1,69478245 | 0,1114401 | 0,33541279 |
| 131616_at | TMEM42 | 0,27318834 | 7,06800663 | 1,69476531 | 0,11144341 | 0,33541279 |
| 134728_at | IRAK1BP1 | 0,50033476 | 4,68353568 | 1,69470009 | 0,11145602 | 0,33541279 |
| 89_at | ACTN3 | -0,14854015 | 5,18184741 | -1,69444824 | 0,11150471 | 0,33550689 |
| 57092_at | PCNP | -0,14788035 | 10,1662831 | -1,69426071 | 0,11154097 | 0,3355636 |
| 79946_at | C10orf95 | -0,21773097 | 5,3383466 | -1,69388417 | 0,11161382 | 0,33573034 |
| 101928602_a | LOC10192860 | -0,18918297 | 5,37881083 | -1,69355741 | 0,11167708 | 0,33575725 |
| 90952_at | ESAM | 0,26220936 | 6,16054259 | 1,69351077 | 0,11168611 | 0,33575725 |
| 6505_at | SLC1A1 | -0,58266789 | 5,15585941 | -1,69343361 | 0,11170105 | 0,33575725 |
| 84450_at | ZNF512 | 0,16609854 | 7,4051798 | 1,6934285 | 0,11170204 | 0,33575725 |
| 400748_at | LOC400748 | -0,18706001 | 3,90189115 | -1,6933878 | 0,11170992 | 0,33575725 |
| 58529_at | MYOZ1 | 0,30147033 | 4,9226748 | 1,69278257 | 0,11182719 | 0,33601801 |
| 245972_at | ATP6V0D2 | -0,14882453 | 3,52239567 | -1,69276002 | 0,11183157 | 0,33601801 |
| 128954_at | GAB4 | -0,18874663 | 6,0433883 | -1,6922482 | 0,11193083 | 0,3362638 |
| 8747_at | ADAM21 | 0,22002796 | 3,76632085 | 1,69168706 | 0,11203974 | 0,33653851 |
| 147381_at | CBLN2 | -0,14715224 | 3,81606583 | -1,69158749 | 0,11205907 | 0,33654411 |
| 100272228_a | LINC00894 | 0,34739301 | 3,38381735 | 1,69146848 | 0,11208219 | 0,33656105 |
| 712_at | C1QA | -0,17237916 | 4,89739809 | -1,69115274 | 0,11214353 | 0,33668007 |
| 157_at | ADRBK2 | 0,18670188 | 6,35723577 | 1,69108457 | 0,11215678 | 0,33668007 |
| 9317_at | PTER | 0,24592748 | 7,27474281 | 1,6909762 | 0,11217784 | 0,33669083 |
| 214_at | ALCAM | -0,14741495 | 9,64285744 | -1,69074059 | 0,11222365 | 0,33677585 |
| 158035_at | LINC00032 | -0,16135881 | 3,70901612 | -1,69053026 | 0,11226455 | 0,33684612 |
| 9094_at | UNC119 | 0,1313502 | 7,55742354 | 1,6891401 | 0,11253524 | 0,3375868 |
| 374875_at | HSD11B1L | 0,15427807 | 5,97718343 | 1,68908256 | 0,11254646 | 0,3375868 |
| 101928324_a | LOC10192832 | -0,19304762 | 3,81837169 | -1,68897963 | 0,11256653 | 0,33759443 |
| 9957_at | HS3ST1 | -0,67566198 | 9,07908651 | -1,68875539 | 0,11261025 | 0,337673 |
| 100507423_a | MKNK1-AS1 | -0,22653706 | 4,75025825 | -1,68861847 | 0,11263696 | 0,33770051 |
| 55721_at | IQCC | -0,18290791 | 5,88204835 | -1,68842529 | 0,11267465 | 0,33776094 |
| 54766_at | BTG4 | -0,17231683 | 3,3858328 | -1,68819873 | 0,11271887 | 0,33782413 |
| 91833_at | WDR20 | 0,17178317 | 6,30476148 | 1,68813757 | 0,11273081 | 0,33782413 |
| 5020_at | OXT | -0,22341964 | 5,3087432 | -1,68793873 | 0,11276963 | 0,33788791 |
| 120071_at | GYLTL1B | -0,18164502 | 6,1972426 | -1,6877418 | 0,11280809 | 0,33795058 |
| 222537_at | HS3ST5 | -0,14934608 | 4,44967774 | -1,68759208 | 0,11283734 | 0,33798564 |
| 2353_at | FOS | -0,55433912 | 7,15816076 | -1,68735476 | 0,11288371 | 0,33807198 |
| 6915_at | TBXA2R | -0,15280706 | 5,52335745 | -1,68683187 | 0,11298595 | 0,33831302 |
| 1843_at | DUSP1 | 0,21737017 | 7,06044895 | 1,68676351 | 0,11299932 | 0,33831302 |
| 6901_at | TAZ | -0,11556994 | 7,31785204 | -1,68645111 | 0,11306045 | 0,33840357 |
| 8900_at | CCNA1 | -0,27551411 | 5,22579144 | -1,68642939 | 0,1130647 | 0,33840357 |
| 101926903_a | LSAMP-AS1 | -0,19118679 | 4,52046445 | -1,68623618 | 0,11310252 | 0,33846418 |
| 64710_at | NUCKS1 | -0,22445566 | 10,486488 | -1,68597045 | 0,11315456 | 0,3385673 |
| 84987_at | COX14 | 0,12345444 | 8,73049707 | 1,68559215 | 0,11322867 | 0,33863428 |
| 8880_at | FUBP1 | -0,22291415 | 8,26338466 | -1,68557946 | 0,11323116 | 0,33863428 |
| 255231_at | MCOLN2 | -0,14966664 | 4,17102708 | -1,68550145 | 0,11324645 | 0,33863428 |
| 245973_at | ATP6V1C2 | 0,29209864 | 4,42325032 | 1,68549728 | 0,11324726 | 0,33863428 |
| 136541_at | PRSS58 | -0,16274838 | 4,08773021 | -1,68527581 | 0,11329068 | 0,33871152 |
| 55851_at | PSENEN | 0,14122666 | 9,73124604 | 1,68455825 | 0,11343144 | 0,33906228 |

| | | | | | | |
|---|---|---|---|---|---|---|
| 9761_at | MLEC | -0,20708441 | 9,96919111 | -1,68449833 | 0,1134432 | 0,33906228 |
| 283150_at | FOXR1 | -0,11904621 | 3,7633612 | -1,6843417 | 0,11347395 | 0,33910157 |
| 152404_at | IGSF11 | 0,44815938 | 4,38279963 | 1,68403448 | 0,11353428 | 0,33911835 |
| 8470_at | SORBS2 | 0,63150709 | 5,01346046 | 1,68397413 | 0,11354614 | 0,33911835 |
| 9855_at | FARP2 | 0,17074073 | 5,65578104 | 1,68380831 | 0,11357872 | 0,33911835 |
| 408_at | ARRB1 | 0,39773376 | 6,46092913 | 1,68374594 | 0,11359097 | 0,33911835 |
| 79672_at | FN3KRP | -0,18437271 | 8,78338403 | -1,68372834 | 0,11359443 | 0,33911835 |
| 55692_at | LUC7L | -0,12742232 | 7,73926469 | -1,68372809 | 0,11359448 | 0,33911835 |
| 9204_at | ZMYM6 | 0,22316365 | 6,35301222 | 1,68360196 | 0,11361927 | 0,33911835 |
| 8557_at | TCAP | 0,14883363 | 5,97963422 | 1,6835962 | 0,1136204 | 0,33911835 |
| 100507008_a | LINC00844 | -0,12474387 | 2,73914313 | -1,68340169 | 0,11365864 | 0,33917992 |
| 79877_at | DCAKD | 0,1573085 | 7,9700157 | 1,6829865 | 0,1137403 | 0,33937103 |
| 25884_at | CHRDL2 | -0,21124749 | 4,85266172 | -1,68278742 | 0,11377947 | 0,33940753 |
| 55531_at | ELMOD1 | -0,12226202 | 3,05503186 | -1,68274525 | 0,11378777 | 0,33940753 |
| 58483_at | LINC00474 | -0,21306783 | 3,68917687 | -1,68255005 | 0,11382619 | 0,33943747 |
| 23636_at | NUP62 | -0,33288564 | 8,82701776 | -1,68251523 | 0,11383305 | 0,33943747 |
| 340252_at | ZNF680 | 0,22942041 | 6,80969412 | 1,68217902 | 0,11389926 | 0,33957322 |
| 619343_at | ERICH1-AS1 | -0,18662033 | 4,45315549 | -1,68210504 | 0,11391383 | 0,33957322 |
| 10750_at | GRAP | -0,10423677 | 5,29019163 | -1,68169483 | 0,11399467 | 0,33976161 |
| 200159_at | C1orf100 | -0,22216956 | 3,53970943 | -1,68103447 | 0,1141249 | 0,34009715 |
| 8526_at | DGKE | -0,24729821 | 4,38008064 | -1,68091979 | 0,11414753 | 0,34011198 |
| 113612_at | CYP2U1 | 0,3604122 | 5,1355248 | 1,68069563 | 0,11419178 | 0,34016548 |
| 148268_at | ZNF570 | 0,17858382 | 5,31818631 | 1,6806499 | 0,11420081 | 0,34016548 |
| 90199_at | WFDC8 | -0,18714677 | 3,94638239 | -1,6802175 | 0,1142862 | 0,34036722 |
| 64132_at | XYLT2 | -0,16060031 | 7,17909328 | -1,67992513 | 0,11434398 | 0,34042382 |
| 1175_at | AP2S1 | -0,23532207 | 9,85992058 | -1,67992105 | 0,11434478 | 0,34042382 |
| 10748_at | KLRAP1 | -0,22514668 | 3,37848549 | -1,67985304 | 0,11435823 | 0,34042382 |
| 348327_at | ZNF530 | -0,2564152 | 5,20830389 | -1,67969771 | 0,11438893 | 0,34046261 |
| 5533_at | PPP3CC | 0,19146259 | 6,2787919 | 1,67959156 | 0,11440992 | 0,34047248 |
| 84643_at | KIF2B | -0,26071178 | 3,95292709 | -1,67939679 | 0,11444844 | 0,34053111 |
| 56950_at | SMYD2 | -0,19160791 | 9,1156718 | -1,67931319 | 0,11446498 | 0,34053111 |
| 25988_at | HINFP | 0,11761936 | 7,28119046 | 1,67906679 | 0,11451373 | 0,34062354 |
| 53981_at | CPSF2 | -0,29718041 | 8,83037744 | -1,67894021 | 0,11453879 | 0,34064546 |
| 842_at | CASP9 | 0,14735133 | 7,15110644 | 1,67869865 | 0,11458661 | 0,34073509 |
| 50801_at | KCNK4 | -0,09939636 | 5,57151796 | -1,67835915 | 0,11465385 | 0,34080862 |
| 27319_at | BHLHE22 | -0,12738763 | 4,43554017 | -1,67831407 | 0,11466278 | 0,34080862 |
| 5300_at | PIN1 | -0,21045838 | 8,60680207 | -1,67830582 | 0,11466442 | 0,34080862 |
| 100630918_a | PARTICL | -0,19244379 | 5,60638814 | -1,6782115 | 0,11468311 | 0,34081158 |
| 102723886_a | LOC10272388 | -0,15222248 | 4,03686321 | -1,67768997 | 0,11478649 | 0,34099663 |
| 389641_at | LOC389641 | -0,41276989 | 7,08514601 | -1,6776368 | 0,11479704 | 0,34099663 |
| 253782_at | CERS6 | 0,19941272 | 8,7338498 | 1,67754501 | 0,11481524 | 0,34099663 |
| 142940_at | TRUB1 | -0,26613235 | 6,9773073 | -1,67754027 | 0,11481618 | 0,34099663 |
| 3249_at | HPN | -0,18020989 | 5,42649598 | -1,67736034 | 0,11485188 | 0,34105006 |
| 122481_at | AK7 | 0,20762554 | 3,75230995 | 1,67718804 | 0,11488607 | 0,341099 |
| 10603_at | SH2B2 | 0,1297444 | 6,68274271 | 1,67697262 | 0,11492883 | 0,34117337 |
| 57380_at | MRS2 | -0,16601145 | 7,87070945 | -1,67671065 | 0,11498085 | 0,34120699 |
| 57544_at | TXNDC16 | 0,1703543 | 6,94123276 | 1,67670349 | 0,11498227 | 0,34120699 |
| 51296_at | SLC15A3 | 0,16959318 | 5,28231459 | 1,67664798 | 0,11499329 | 0,34120699 |
| 25949_at | SYF2 | 0,19140556 | 7,90247298 | 1,67634693 | 0,1150531 | 0,34120986 |
| 51340_at | CRNKL1 | 0,18240155 | 7,9255355 | 1,67631867 | 0,11505872 | 0,34120986 |

| | | | | | | |
|---|---|---|---|---|---|---|
| 170575_at | GIMAP1 | -0,11765381 | 3,68674639 | -1,67626264 | 0,11506985 | 0,34120986 |
| 222161_at | DKFZP586I142 | 0,31302979 | 6,87241456 | 1,67624162 | 0,11507403 | 0,34120986 |
| 402665_at | IGLON5 | -0,2309488 | 5,64231815 | -1,67619736 | 0,11508283 | 0,34120986 |
| 9978_at | RBX1 | -0,15105976 | 10,5106018 | -1,67579042 | 0,11516373 | 0,3413972 |
| 5225_at | PGC | -0,16363943 | 4,9381812 | -1,67555485 | 0,11521059 | 0,34148355 |
| 4892_at | NRAP | -0,12218599 | 3,94620064 | -1,67542433 | 0,11523656 | 0,34149311 |
| 51806_at | CALML5 | -0,32605365 | 5,24225038 | -1,67536046 | 0,11524927 | 0,34149311 |
| 9871_at | SEC24D | 0,32979298 | 6,31350594 | 1,67505128 | 0,11531081 | 0,34162292 |
| 1128_at | CHRM1 | -0,15455466 | 4,40949085 | -1,67460911 | 0,11539888 | 0,34183124 |
| 140732_at | SUN5 | -0,10998974 | 4,70117912 | -1,67438875 | 0,11544279 | 0,34189475 |
| 102546294_at | LOC10254629 | -0,14113819 | 4,27960506 | -1,67432339 | 0,11545581 | 0,34189475 |
| 84335_at | AKT1S1 | -0,18057155 | 8,04211437 | -1,67419743 | 0,11548092 | 0,34191654 |
| 199675_at | MCEMP1 | -0,14392022 | 5,22844203 | -1,67297395 | 0,11572505 | 0,34251817 |
| 10263_at | CDK2AP2 | -0,24234227 | 9,00986147 | -1,67295233 | 0,11572937 | 0,34251817 |
| 84647_at | PLA2G12B | -0,17065774 | 4,5262036 | -1,6729118 | 0,11573746 | 0,34251817 |
| 340152_at | ZC3H12D | 0,23182978 | 4,39486309 | 1,67242749 | 0,11583424 | 0,3427519 |
| 3696_at | ITGB8 | 0,19763082 | 5,23475 | 1,67218993 | 0,11588173 | 0,34283977 |
| 51594_at | NBAS | 0,13520189 | 6,28623471 | 1,67204909 | 0,11590989 | 0,34287044 |
| 150197_at | LINC00896 | -0,19181869 | 5,45194014 | -1,67141688 | 0,11603639 | 0,34315005 |
| 51251_at | NT5C3A | 0,35020188 | 9,19971446 | 1,67139862 | 0,11604005 | 0,34315005 |
| 84864_at | MINA | -0,34451009 | 7,39029887 | -1,671207 | 0,11607841 | 0,3431645 |
| 80339_at | PNPLA3 | -0,21752342 | 5,72392481 | -1,67119628 | 0,11608056 | 0,3431645 |
| 161_at | AP2A2 | 0,11760847 | 7,80640382 | 1,67104791 | 0,11611028 | 0,34319967 |
| 160492_at | LMNTD1 | -0,17734491 | 4,73320961 | -1,67087103 | 0,11614571 | 0,34325174 |
| 25912_at | C1orf43 | -0,16968183 | 10,3989284 | -1,66990664 | 0,11633907 | 0,34372521 |
| 283297_at | OR10A4 | -0,14759784 | 4,14208063 | -1,66976656 | 0,11636717 | 0,34372521 |
| 124790_at | HEXIM2 | 0,17042381 | 6,48396063 | 1,66974002 | 0,1163725 | 0,34372521 |
| 54455_at | FBXO42 | 0,11784283 | 7,46762179 | 1,66971614 | 0,11637729 | 0,34372521 |
| 286204_at | CRB2 | -0,1526002 | 4,70990142 | -1,66916595 | 0,11648776 | 0,34399873 |
| 84189_at | SLITRK6 | 0,51970153 | 9,19050013 | 1,66864211 | 0,11659302 | 0,3442568 |
| 55209_at | SETD5 | 0,17270612 | 8,28637606 | 1,66853156 | 0,11661525 | 0,34426965 |
| 147744_at | TMEM190 | -0,15724129 | 5,70036089 | -1,66804236 | 0,11671364 | 0,3443206 |
| 2029_at | ENSA | -0,21455785 | 7,84578409 | -1,66804074 | 0,11671396 | 0,3443206 |
| 6548_at | SLC9A1 | 0,18004806 | 9,63781787 | 1,66802276 | 0,11671758 | 0,3443206 |
| 3094_at | HINT1 | -0,13153385 | 11,1154407 | -1,66801636 | 0,11671887 | 0,3443206 |
| 338707_at | B4GALNT4 | -0,24508681 | 5,04014148 | -1,667948 | 0,11673262 | 0,3443206 |
| 80975_at | TMPRSS5 | -0,21739155 | 5,24057681 | -1,66791258 | 0,11673975 | 0,3443206 |
| 168448_at | CDC14C | -0,15652025 | 2,74828094 | -1,66746306 | 0,11683024 | 0,34448509 |
| 57633_at | LRRN1 | 0,70375072 | 4,0468465 | 1,66745787 | 0,11683129 | 0,34448509 |
| 54462_at | CCSER2 | 0,10577431 | 7,2579751 | 1,66726098 | 0,11687094 | 0,34452347 |
| 130106_at | CIB4 | -0,23867803 | 4,00462163 | -1,66690162 | 0,11694335 | 0,34452347 |
| 2100_at | ESR2 | -0,13685858 | 4,20862514 | -1,66685442 | 0,11695286 | 0,34452347 |
| 93663_at | ARHGAP18 | -0,32774644 | 7,86885208 | -1,66685059 | 0,11695364 | 0,34452347 |
| 139886_at | SPIN4 | -0,42910726 | 9,11381501 | -1,66680498 | 0,11696283 | 0,34452347 |
| 2769_at | GNA15 | -0,26180298 | 9,29123427 | -1,66676592 | 0,1169707 | 0,34452347 |
| 1014_at | CDH16 | -0,25945222 | 4,82078002 | -1,66675894 | 0,11697211 | 0,34452347 |
| 3895_at | KTN1 | 0,17195488 | 10,4275332 | 1,66668316 | 0,11698739 | 0,34452347 |
| 5362_at | PLXNA2 | -0,23350062 | 7,81470145 | -1,66643936 | 0,11703655 | 0,34460594 |
| 54978_at | SLC35F6 | 0,07942263 | 8,41739357 | 1,66636687 | 0,11705117 | 0,34460594 |
| 64215_at | DNAJC1 | 0,12183095 | 8,50817141 | 1,66622994 | 0,11707879 | 0,34463459 |

| | | | | | | |
|---|---|---|---|---|---|---|
| 114036_at | LINC00310 | -0,16923676 | 3,99483044 | -1,6659291 | 0,1171395 | 0,3447606 |
| 51181_at | DCXR | -0,24022344 | 10,3905993 | -1,66567106 | 0,11719159 | 0,34486123 |
| 1544_at | CYP1A2 | -0,18681218 | 5,95337402 | -1,66541768 | 0,11724276 | 0,34495911 |
| 9941_at | EXOG | -0,15867681 | 6,19419583 | -1,66495476 | 0,1173363 | 0,34518161 |
| 140691_at | TRIM69 | 0,13643343 | 4,05100336 | 1,66471991 | 0,11738378 | 0,34526856 |
| 50834_at | TAS2R1 | -0,28481587 | 4,07247512 | -1,66460654 | 0,11740671 | 0,34527827 |
| 5955_at | RCN2 | -0,18350803 | 9,80618198 | -1,66452631 | 0,11742293 | 0,34527827 |
| 22981_at | NINL | 0,5852551 | 5,89106447 | 1,66430711 | 0,11746727 | 0,3453034 |
| 585_at | BBS4 | 0,26348512 | 6,80527978 | 1,66430684 | 0,11746733 | 0,3453034 |
| 2218_at | FKTN | -0,11867327 | 5,82353471 | -1,66411358 | 0,11750644 | 0,34534018 |
| 55345_at | ZGRF1 | -0,40100486 | 4,85407013 | -1,66399647 | 0,11753014 | 0,34534018 |
| 196_at | AHR | 0,34513524 | 10,483419 | 1,66397925 | 0,11753363 | 0,34534018 |
| 5830_at | PEX5 | 0,14315897 | 6,79230303 | 1,6635612 | 0,11761828 | 0,3455362 |
| 6760_at | SS18 | -0,21664192 | 7,5336321 | -1,6632286 | 0,11768566 | 0,34568145 |
| 79870_at | BAALC | -0,21882166 | 4,09559339 | -1,6630531 | 0,11772123 | 0,34568644 |
| 120526_at | DNAJC24 | -0,36547512 | 6,36768218 | -1,66302965 | 0,11772599 | 0,34568644 |
| 142891_at | SAMD8 | 0,23009866 | 6,34767277 | 1,66295462 | 0,1177412 | 0,34568644 |
| 245806_at | VGLL2 | -0,16754546 | 4,73294839 | -1,66272292 | 0,11778818 | 0,34577167 |
| 285205_at | LINC00636 | -0,14638009 | 3,8357331 | -1,66248432 | 0,11783658 | 0,3458143 |
| 9474_at | ATG5 | 0,19428576 | 8,43114503 | 1,66247433 | 0,1178386 | 0,3458143 |
| 25798_at | BRI3 | 0,1751587 | 9,30726746 | 1,66213912 | 0,11790663 | 0,34596122 |
| 253260_at | RICTOR | 0,30464465 | 7,36595709 | 1,66170676 | 0,11799442 | 0,34608847 |
| 6988_at | TCTA | 0,14318297 | 8,02584236 | 1,66168002 | 0,11799985 | 0,34608847 |
| 25934_at | NIPSNAP3A | 0,33364958 | 7,66368374 | 1,66161212 | 0,11801364 | 0,34608847 |
| 8988_at | HSPB3 | -0,14380902 | 4,25567683 | -1,66149749 | 0,11803693 | 0,34608847 |
| 6449_at | SGTA | -0,32035099 | 8,27607925 | -1,66146864 | 0,11804279 | 0,34608847 |
| 147807_at | ZNF524 | 0,14211437 | 6,15986063 | 1,66131825 | 0,11807336 | 0,34608847 |
| 51329_at | ARL6IP4 | -0,15519992 | 9,42570283 | -1,66130641 | 0,11807576 | 0,34608847 |
| 4659_at | PPP1R12A | 0,22183023 | 9,25917491 | 1,66108553 | 0,11812066 | 0,34616739 |
| 1522_at | CTSZ | 0,28499649 | 9,49792849 | 1,66043767 | 0,11825244 | 0,34650086 |
| 91851_at | CHRDL1 | -0,16328063 | 4,0690049 | -1,66015304 | 0,11831037 | 0,34661789 |
| 51193_at | ZNF639 | -0,14818273 | 7,55668433 | -1,65989449 | 0,11836302 | 0,3466739 |
| 2976_at | GTF3C2 | -0,16500884 | 9,27646507 | -1,65971756 | 0,11839906 | 0,3466739 |
| 58497_at | PRUNE | 0,14695239 | 7,54089035 | 1,65967944 | 0,11840682 | 0,3466739 |
| 57030_at | SLC17A7 | -0,13770336 | 5,93641532 | -1,65955756 | 0,11843165 | 0,3466739 |
| 6366_at | CCL21 | -0,14403782 | 5,34948 | -1,65955676 | 0,11843182 | 0,3466739 |
| 100506755_a | MIR497HG | -0,21933842 | 5,54261961 | -1,6595133 | 0,11844067 | 0,3466739 |
| 389177_at | TMEM212 | 0,15217189 | 3,64158897 | 1,65944073 | 0,11845546 | 0,3466739 |
| 93408_at | MYL10 | -0,11509476 | 5,21538417 | -1,65914962 | 0,1185148 | 0,34679488 |
| 27_at | ABL2 | 0,13344958 | 6,28458424 | 1,65873629 | 0,11859911 | 0,34698885 |
| 128822_at | CST9 | -0,14565215 | 4,62207802 | -1,6584491 | 0,11865771 | 0,34710758 |
| 57678_at | GPAM | -0,26210781 | 6,88481202 | -1,65823505 | 0,1187014 | 0,34713181 |
| 23054_at | NCOA6 | 0,1320023 | 9,19095908 | 1,65818147 | 0,11871234 | 0,34713181 |
| 5781_at | PTPN11 | -0,15367841 | 7,03158857 | -1,65807929 | 0,11873321 | 0,34713181 |
| 400084_at | LINC00939 | -0,13148629 | 3,52361024 | -1,65805548 | 0,11873807 | 0,34713181 |
| 101928790_a | PEX5L-AS2 | -0,18528998 | 3,62858886 | -1,65758632 | 0,11883392 | 0,34731646 |
| 3065_at | HDAC1 | -0,19263955 | 10,5394019 | -1,65747168 | 0,11885735 | 0,34731646 |
| 9256_at | BZRAP1 | -0,16230849 | 5,19383516 | -1,65740043 | 0,11887191 | 0,34731646 |
| 29842_at | TFCP2L1 | -0,18052073 | 5,15542734 | -1,65739339 | 0,11887335 | 0,34731646 |
| 10904_at | BLCAP | 0,1484605 | 11,098358 | 1,65726288 | 0,11890004 | 0,34733386 |

| | | | | | | |
|---|---|---|---|---|---|---|
| 9368_at | SLC9A3R1 | -0,30432115 | 10,023544 | -1,65718788 | 0,11891537 | 0,34733386 |
| 3227_at | HOXC11 | -0,25538262 | 4,34253118 | -1,65695104 | 0,11896381 | 0,34742266 |
| 23353_at | SUN1 | -0,17461982 | 7,6093562 | -1,65682443 | 0,11898971 | 0,34744563 |
| 6383_at | SDC2 | -0,44775797 | 6,43066063 | -1,65640838 | 0,11907486 | 0,34764157 |
| 4916_at | NTRK3 | -0,14372644 | 4,80414309 | -1,65581815 | 0,11919575 | 0,34794177 |
| 1291_at | COL6A1 | 0,15166194 | 6,35760062 | 1,65525645 | 0,11931089 | 0,34822511 |
| 3850_at | KRT3 | -0,16696792 | 4,28763344 | -1,65500709 | 0,11936204 | 0,34829368 |
| 84513_at | PPAPDC1B | 0,38016778 | 7,62523944 | 1,65496563 | 0,11937055 | 0,34829368 |
| 677769_at | SCARNA17 | 0,18240606 | 5,35817876 | 1,65482061 | 0,1194003 | 0,34832774 |
| 100129917_a | LOC10012991 | -0,2094398 | 6,24575074 | -1,65472909 | 0,11941909 | 0,34832978 |
| 83890_at | SPATA9 | 0,11186556 | 3,38256273 | 1,65456985 | 0,11945177 | 0,34835578 |
| 5696_at | PSMB8 | 0,48983758 | 9,37202413 | 1,65450268 | 0,11946556 | 0,34835578 |
| 9527_at | GOSR1 | 0,11817562 | 8,50804501 | 1,65442138 | 0,11948225 | 0,34835578 |
| 55331_at | ACER3 | -0,13623183 | 7,45063929 | -1,65406082 | 0,11955631 | 0,34850206 |
| 101928243_a | DARS-AS1 | -0,18715485 | 4,52342162 | -1,65397388 | 0,11957417 | 0,34850206 |
| 2035_at | EPB41 | 0,13119309 | 5,6559171 | 1,6539129 | 0,1195867 | 0,34850206 |
| 100505601_a | PARD3-AS1 | 0,21281003 | 3,84436424 | 1,653568 | 0,11965759 | 0,3486559 |
| 101927914_a | LOC10192791 | -0,16796885 | 4,76745692 | -1,65298416 | 0,11977768 | 0,34893672 |
| 79035_at | NABP2 | -0,3935266 | 8,23211616 | -1,65292332 | 0,1197902 | 0,34893672 |
| 149069_at | DCDC2B | -0,23943317 | 5,5259531 | -1,65279642 | 0,11981631 | 0,34896003 |
| 101929550_a | LOC10192955 | -0,19939059 | 4,00883435 | -1,65219094 | 0,11994099 | 0,34925998 |
| 8916_at | HERC3 | 0,17488394 | 6,53601837 | 1,65208376 | 0,11996308 | 0,34925998 |
| 254359_at | ZDHHC24 | 0,12535651 | 7,46902679 | 1,65203222 | 0,1199737 | 0,34925998 |
| 79736_at | TEFM | -0,24284511 | 7,70191079 | -1,65169172 | 0,12004388 | 0,34941149 |
| 773_at | CACNA1A | -0,14588888 | 4,80334718 | -1,65150189 | 0,12008302 | 0,34947261 |
| 79627_at | OGFRL1 | -0,20395759 | 5,12144495 | -1,65103075 | 0,12018022 | 0,34970265 |
| 60467_at | BPESC1 | -0,11657173 | 3,40198515 | -1,65087661 | 0,12021203 | 0,34974238 |
| 10362_at | HMG20B | 0,22649739 | 8,98282669 | 1,65075485 | 0,12023717 | 0,34976269 |
| 4066_at | LYL1 | -0,15833474 | 6,35390439 | -1,65019169 | 0,12035348 | 0,35004818 |
| 100130700_a | LOC10013070 | -0,17671604 | 4,66530025 | -1,65001297 | 0,12039041 | 0,35010274 |
| 54550_at | NECAB2 | -0,18710532 | 6,14335046 | -1,6497853 | 0,12043748 | 0,35018674 |
| 2135_at | EXTL2 | -0,13515367 | 7,80813018 | -1,64930991 | 0,1205358 | 0,35041973 |
| 25831_at | HECTD1 | 0,23778053 | 7,97894197 | 1,64907588 | 0,12058423 | 0,35050762 |
| 349075_at | ZNF713 | 0,19572242 | 3,85153946 | 1,6486684 | 0,12066859 | 0,35069992 |
| 145873_at | MESP2 | -0,19485841 | 6,20886496 | -1,64850384 | 0,12070267 | 0,35074606 |
| 4548_at | MTR | -0,14385834 | 7,85387233 | -1,64841503 | 0,12072107 | 0,35074661 |
| 10454_at | TAB1 | 0,12522488 | 6,80040246 | 1,6482585 | 0,1207535 | 0,35078792 |
| 284111_at | SLC13A5 | -0,16724025 | 4,73044225 | -1,6478641 | 0,12083526 | 0,35097249 |
| 171392_at | ZNF675 | 0,24506347 | 4,57050084 | 1,64682391 | 0,1210511 | 0,35147715 |
| 51510_at | CHMP5 | 0,21559831 | 9,41241673 | 1,64679222 | 0,12105768 | 0,35147715 |
| 27106_at | ARRDC2 | -0,14568873 | 7,34475364 | -1,64671277 | 0,12107418 | 0,35147715 |
| 246778_at | IL27 | -0,20683156 | 5,65100254 | -1,64667517 | 0,12108199 | 0,35147715 |
| 26577_at | PCOLCE2 | -0,23812564 | 3,66515965 | -1,64609832 | 0,12120186 | 0,35177209 |
| 51014_at | TMED7 | -0,25968113 | 9,28939637 | -1,64589799 | 0,12124351 | 0,35183996 |
| 1952_at | CELSR2 | 0,2433892 | 7,67162416 | 1,64561105 | 0,12130319 | 0,35196014 |
| 101929268_a | LOC10192926 | -0,23876455 | 4,46322587 | -1,64537711 | 0,12135187 | 0,35201952 |
| 8789_at | FBP2 | -0,19114384 | 4,94350898 | -1,64532531 | 0,12136265 | 0,35201952 |
| 94005_at | PIGS | 0,1766451 | 9,01891274 | 1,64524924 | 0,12137848 | 0,35201952 |
| 6297_at | SALL2 | -0,17225243 | 5,25576913 | -1,64490132 | 0,12145092 | 0,35217309 |
| 22920_at | KIFAP3 | 0,3387946 | 7,43777094 | 1,64481932 | 0,12146799 | 0,35217309 |

| | | | | | | |
|---|---|---|---|---|---|---|
| 133396_at | IL31RA | -0,11837563 | 3,32841178 | -1,64452288 | 0,12152975 | 0,35229911 |
| 5778_at | PTPN7 | -0,19900627 | 5,37296618 | -1,64425631 | 0,12158531 | 0,35240713 |
| 10084_at | PQBP1 | -0,16839836 | 9,00691797 | -1,64409843 | 0,12161822 | 0,3524495 |
| 2589_at | GALNT1 | 0,25601931 | 11,0547187 | 1,64338518 | 0,12176701 | 0,35282762 |
| 57758_at | SCUBE2 | 0,20996483 | 4,227249 | 1,64318944 | 0,12180788 | 0,35289294 |
| 606495_at | CYB5RL | -0,16079461 | 5,32613352 | -1,64281146 | 0,12188682 | 0,35306853 |
| 283373_at | ANKRD52 | 0,17400928 | 8,25247369 | 1,64226786 | 0,12200042 | 0,35334447 |
| 2200_at | FBN1 | 0,12792084 | 4,11389259 | 1,64193769 | 0,12206947 | 0,3534913 |
| 24_at | ABCA4 | -0,15765079 | 5,44639411 | -1,6417217 | 0,12211465 | 0,353569 |
| 22978_at | NT5C2 | 0,22966887 | 9,71324186 | 1,64151622 | 0,12215765 | 0,35364035 |
| 94032_at | CAMK2N2 | -0,15582462 | 6,76464564 | -1,64119152 | 0,12222563 | 0,35378397 |
| 11135_at | CDC42EP1 | -0,15866714 | 5,94705606 | -1,6410231 | 0,1222609 | 0,3538102 |
| 127281_at | FAM213B | -0,15920452 | 8,60835742 | -1,64097284 | 0,12227143 | 0,3538102 |
| 23382_at | AHCYL2 | 0,16668737 | 6,87265046 | 1,64086274 | 0,12229449 | 0,35382379 |
| 79935_at | CNTD2 | -0,22044521 | 6,72325251 | -1,64051519 | 0,12236732 | 0,35394473 |
| 1675_at | CFD | 0,60630127 | 6,93992316 | 1,64043591 | 0,12238394 | 0,35394473 |
| 116028_at | RMI2 | -0,37655936 | 9,14746264 | -1,64040026 | 0,12239142 | 0,35394473 |
| 79979_at | TRMT2B | -0,16376772 | 6,55535379 | -1,64000965 | 0,12247333 | 0,35412846 |
| 5903_at | RANBP2 | -0,15266931 | 7,91005888 | -1,63984181 | 0,12250854 | 0,35417711 |
| 8780_at | RIOK3 | 0,45951546 | 8,56662844 | 1,63945284 | 0,12259018 | 0,35435994 |
| 7385_at | UQCRC2 | -0,35940849 | 10,4210843 | -1,63933683 | 0,12261454 | 0,35437717 |
| 56521_at | DNAJC12 | 0,13940698 | 3,86375682 | 1,63912464 | 0,1226591 | 0,35437724 |
| 339231_at | ARL16 | -0,12774205 | 7,73076742 | -1,63907907 | 0,12266867 | 0,35437724 |
| 441204_at | LOC441204 | -0,12470217 | 3,91998695 | -1,63907393 | 0,12266975 | 0,35437724 |
| 145282_at | MIPOL1 | 0,13166826 | 4,87854108 | 1,63890792 | 0,12270463 | 0,35442484 |
| 3106_at | HLA-B | 0,95789151 | 10,9726196 | 1,63854455 | 0,122781 | 0,35454707 |
| 645249_at | MNX1-AS1 | -0,35599092 | 5,29862897 | -1,63853142 | 0,12278376 | 0,35454707 |
| 412_at | STS | 0,23618314 | 7,78597718 | 1,63835165 | 0,12282156 | 0,35458973 |
| 5770_at | PTPN1 | -0,10958292 | 8,99161064 | -1,63821117 | 0,1228511 | 0,35458973 |
| 51250_at | C6orf203 | 0,14352299 | 7,13029277 | 1,63815653 | 0,12286259 | 0,35458973 |
| 29950_at | SERTAD1 | 0,43730817 | 8,71624148 | 1,63811104 | 0,12287216 | 0,35458973 |
| 2049_at | EPHB3 | -0,1923875 | 6,54442413 | -1,63770199 | 0,12295823 | 0,35478497 |
| 94120_at | SYTL3 | 0,24906678 | 5,71599319 | 1,63749533 | 0,12300174 | 0,35485735 |
| 817_at | CAMK2D | 0,80248151 | 6,98311441 | 1,6370302 | 0,12309971 | 0,3550868 |
| 2719_at | GPC3 | 0,87413289 | 6,07345764 | 1,63679284 | 0,12314973 | 0,3551779 |
| 10162_at | LPCAT3 | 0,17080569 | 9,87765379 | 1,63656658 | 0,12319742 | 0,35521711 |
| 6929_at | TCF3 | -0,113439 | 8,02057972 | -1,63645504 | 0,12322094 | 0,35521711 |
| 58478_at | ENOPH1 | -0,60832972 | 9,03422901 | -1,63638989 | 0,12323468 | 0,35521711 |
| 126393_at | HSPB6 | -0,20924616 | 4,5931731 | -1,63637849 | 0,12323708 | 0,35521711 |
| 84962_at | AJUBA | 0,37330659 | 9,10053906 | 1,63600674 | 0,12331551 | 0,35530253 |
| 252884_at | ZNF396 | 0,2486624 | 3,7958106 | 1,63599675 | 0,12331762 | 0,35530253 |
| 3399_at | ID3 | 0,33220706 | 10,5654315 | 1,63597571 | 0,12332205 | 0,35530253 |
| 160365_at | CLECL1 | -0,09996838 | 2,92673274 | -1,635668 | 0,123387 | 0,35539151 |
| 54456_at | MOV10L1 | -0,12971372 | 4,81889961 | -1,63565458 | 0,12338984 | 0,35539151 |
| 4354_at | MPP1 | -0,15768124 | 5,15126579 | -1,63523159 | 0,12347917 | 0,35559129 |
| 113444_at | SMIM12 | -0,14503301 | 8,02844005 | -1,63515136 | 0,12349612 | 0,35559129 |
| 85406_at | DNAJC14 | 0,09106349 | 8,43990961 | 1,63489761 | 0,12354974 | 0,35569252 |
| 80153_at | EDC3 | 0,17010593 | 7,27679274 | 1,63435088 | 0,12366534 | 0,35597213 |
| 259173_at | ALS2CL | 0,49680084 | 7,97201693 | 1,63383678 | 0,12377413 | 0,35622772 |
| 55018_at | LINC00483 | -0,16721803 | 3,61234626 | -1,63375653 | 0,12379112 | 0,35622772 |

| | | | | | | |
|---|---|---|---|---|---|---|
| 26504_at | CNNM4 | 0,15612678 | 7,74068193 | 1,63340934 | 0,12386465 | 0,35635406 |
| 92342_at | METTL18 | 0,16771346 | 7,40975346 | 1,63337451 | 0,12387202 | 0,35635406 |
| 642273_at | FAM110C | 0,24717225 | 9,21524812 | 1,6332037 | 0,12390821 | 0,35640494 |
| 53905_at | DUOX1 | 0,39664979 | 7,16360418 | 1,63277578 | 0,12399891 | 0,35661258 |
| 91978_at | TPGS1 | 0,24285639 | 7,00153424 | 1,63244301 | 0,12406948 | 0,35671912 |
| 1382_at | CRABP2 | -0,2597739 | 6,05159627 | -1,63236938 | 0,1240851 | 0,35671912 |
| 118487_at | CHCHD1 | -0,16093958 | 9,085627 | -1,63233916 | 0,12409151 | 0,35671912 |
| 57864_at | SLC46A2 | -0,14363449 | 4,32731544 | -1,63219051 | 0,12412306 | 0,35675655 |
| 51112_at | TRAPPC12 | -0,09053457 | 6,74073223 | -1,63208414 | 0,12414563 | 0,3567682 |
| 285834_at | HCG22 | -0,24709074 | 4,30537996 | -1,63185086 | 0,12419515 | 0,35685727 |
| 4654_at | MYOD1 | -0,17408256 | 5,68207862 | -1,63160124 | 0,12424816 | 0,35695634 |
| 151056_at | PLB1 | -0,1097474 | 5,40829938 | -1,63141642 | 0,12428741 | 0,35697261 |
| 5625_at | PRODH | 0,15432785 | 5,05450454 | 1,63136677 | 0,12429796 | 0,35697261 |
| 654429_at | LRTM2 | -0,2199599 | 4,79500175 | -1,63130298 | 0,12431152 | 0,35697261 |
| 51738_at | GHRL | -0,17145943 | 4,68939668 | -1,63122567 | 0,12432795 | 0,35697261 |
| 149954_at | BPIFB4 | -0,1956496 | 4,82123632 | -1,63056396 | 0,12446863 | 0,35720832 |
| 164380_at | CST13P | -0,1387647 | 3,05926066 | -1,63042411 | 0,12449838 | 0,35720832 |
| 9487_at | PIGL | -0,65921302 | 6,03027567 | -1,63039528 | 0,12450452 | 0,35720832 |
| 51093_at | RRNAD1 | 0,15081467 | 7,42474448 | 1,6303371 | 0,1245169 | 0,35720832 |
| 25963_at | TMEM87A | 0,15843348 | 9,22614794 | 1,6303338 | 0,1245176 | 0,35720832 |
| 400566_at | C17orf97 | 0,22816667 | 6,50077587 | 1,6303164 | 0,1245213 | 0,35720832 |
| 493911_at | PHOSPHO2 | 0,24734757 | 6,53164969 | 1,63004671 | 0,1245787 | 0,35728738 |
| 9849_at | ZNF518A | 0,13166747 | 6,15456469 | 1,63001261 | 0,12458596 | 0,35728738 |
| 203427_at | SLC25A43 | -0,14848621 | 7,27239021 | -1,62988191 | 0,12461379 | 0,35730775 |
| 401207_at | C5orf63 | -0,14573211 | 4,6632722 | -1,62972535 | 0,12464713 | 0,35730775 |
| 139378_at | ADGRG4 | -0,12847465 | 3,23591997 | -1,62963693 | 0,12466596 | 0,35730775 |
| 339390_at | CLEC4G | -0,18999335 | 4,73762688 | -1,62963083 | 0,12466726 | 0,35730775 |
| 100129973_a | LOC10012997 | 0,22561941 | 4,2231582 | 1,62919697 | 0,1247597 | 0,35751951 |
| 4046_at | LSP1 | -0,16232769 | 5,1759451 | -1,62908977 | 0,12478255 | 0,35753181 |
| 165140_at | OXER1 | -0,18114203 | 6,96800297 | -1,62869793 | 0,12486611 | 0,35768662 |
| 5204_at | PFDN5 | 0,17557736 | 9,79604954 | 1,6285886 | 0,12488943 | 0,35768662 |
| 503542_at | SPRN | -0,16982499 | 5,97014978 | -1,62857519 | 0,12489229 | 0,35768662 |
| 375775_at | PNPLA7 | -0,14468492 | 5,24276368 | -1,6280461 | 0,12500521 | 0,35792987 |
| 10181_at | RBM5 | 0,17354815 | 8,83147301 | 1,62800312 | 0,12501439 | 0,35792987 |
| 30820_at | KCNIP1 | -0,13486802 | 3,90427357 | -1,62786777 | 0,12504329 | 0,35795941 |
| 54878_at | DPP8 | 0,1702279 | 8,52693628 | 1,62731919 | 0,12516049 | 0,35824168 |
| 5731_at | PTGER1 | -0,17447004 | 6,54528697 | -1,6272245 | 0,12518073 | 0,35824639 |
| 51267_at | CLEC1A | 0,19476605 | 3,6916183 | 1,62675678 | 0,12528075 | 0,35847936 |
| 8427_at | ZNF282 | 0,22355562 | 7,41546551 | 1,62659033 | 0,12531636 | 0,3584989 |
| 57053_at | CHRNA10 | -0,14156621 | 4,64150763 | -1,62649564 | 0,12533662 | 0,3584989 |
| 404220_at | C6orf201 | -0,19201812 | 3,50477667 | -1,62646391 | 0,12534341 | 0,3584989 |
| 344558_at | SH3RF3 | -0,19911293 | 5,0571302 | -1,6262625 | 0,12538652 | 0,35856896 |
| 4325_at | MMP16 | -0,09874218 | 3,7434579 | -1,62609508 | 0,12542236 | 0,35859939 |
| 54964_at | C1orf56 | 0,13435819 | 6,57070788 | 1,62603889 | 0,12543439 | 0,35859939 |
| 1947_at | EFNB1 | -0,15651933 | 6,16583766 | -1,62572379 | 0,12550188 | 0,35869411 |
| 22928_at | SEPHS2 | -0,1353071 | 10,0751403 | -1,62571032 | 0,12550477 | 0,35869411 |
| 8887_at | TAX1BP1 | 0,23256907 | 9,4056464 | 1,62553266 | 0,12554283 | 0,35874968 |
| 6445_at | SGCG | -0,13008861 | 3,58132783 | -1,62523635 | 0,12560635 | 0,35887794 |
| 101928190_a | LINC01487 | -0,10347652 | 2,98577877 | -1,62466889 | 0,12572805 | 0,35912884 |
| 246175_at | CNOT6L | 0,11291728 | 5,31861452 | 1,62465306 | 0,12573145 | 0,35912884 |

| | | | | | | |
|---|---|---|---|---|---|---|
| 3148_at | HMGB2 | -0,64207027 | 10,1685207 | -1,624351 | 0,12579628 | 0,35922045 |
| 55908_at | C19orf80 | -0,22271073 | 4,83275059 | -1,62428581 | 0,12581027 | 0,35922045 |
| 51586_at | MED15 | 0,14858646 | 8,02978171 | 1,62424298 | 0,12581947 | 0,35922045 |
| 728411_at | GUSBP1 | 0,12567784 | 3,80122887 | 1,62414581 | 0,12584033 | 0,35922678 |
| 56954_at | NIT2 | -0,35683647 | 9,78556683 | -1,62391595 | 0,1258897 | 0,35931446 |
| 101927539_a | LOC10192753 | -0,18766266 | 3,91805875 | -1,62380008 | 0,12591459 | 0,35933226 |
| 29761_at | USP25 | 0,327704 | 7,92060493 | 1,62366734 | 0,12594311 | 0,35936041 |
| 284260_at | LINC00907 | -0,09623901 | 3,31296034 | -1,62346688 | 0,12598619 | 0,35940535 |
| 23322_at | RPGRIP1L | -0,29776486 | 5,83453599 | -1,62340081 | 0,1260004 | 0,35940535 |
| 147965_at | FAM98C | 0,13386807 | 7,87634313 | 1,62330296 | 0,12602143 | 0,35940535 |
| 440275_at | EIF2AK4 | 0,22432363 | 7,73262909 | 1,62324687 | 0,12603349 | 0,35940535 |
| 2694_at | GIF | -0,12865157 | 3,51264382 | -1,62308048 | 0,12606927 | 0,35945417 |
| 23581_at | CASP14 | -0,60925745 | 6,28784803 | -1,62243437 | 0,1262083 | 0,35979731 |
| 1556_at | CYP2B7P | -0,44474722 | 5,61772261 | -1,62228779 | 0,12623985 | 0,35980807 |
| 63036_at | CELA2A | -0,13755643 | 3,98400877 | -1,62213381 | 0,12627301 | 0,35980807 |
| 6731_at | SRP72 | -0,35429748 | 8,91437591 | -1,62207325 | 0,12628606 | 0,35980807 |
| 3273_at | HRG | -0,12759953 | 4,02709112 | -1,62206986 | 0,12628679 | 0,35980807 |
| 284805_at | C20orf203 | -0,22688212 | 5,68001762 | -1,62186098 | 0,12633178 | 0,3598395 |
| 149998_at | LIPI | -0,10976943 | 2,7668363 | -1,62184522 | 0,12633518 | 0,3598395 |
| 2981_at | GUCA2B | -0,17703764 | 5,46926204 | -1,62162125 | 0,12638344 | 0,35992375 |
| 202500_at | TCTE1 | -0,17688865 | 4,84357265 | -1,62129175 | 0,12645448 | 0,36002259 |
| 101055625_a | OVCH1-AS1 | -0,18114226 | 4,11895494 | -1,62128687 | 0,12645553 | 0,36002259 |
| 222487_at | ADGRG3 | -0,15716728 | 4,4613995 | -1,62050876 | 0,12662341 | 0,36044729 |
| 8535_at | CBX4 | 0,18031849 | 7,8611596 | 1,62035454 | 0,12665671 | 0,3604888 |
| 388780_at | LOC388780 | -0,28440115 | 5,54211983 | -1,62013141 | 0,1267049 | 0,36054717 |
| 140467_at | ZNF358 | 0,11850862 | 6,97925556 | 1,62008627 | 0,12671465 | 0,36054717 |
| 100289410_a | MCF2L-AS1 | 0,15712135 | 5,15283889 | 1,61992786 | 0,12674887 | 0,36059128 |
| 285267_at | ZNF619 | -0,15694376 | 5,77014085 | -1,61972182 | 0,1267934 | 0,36062982 |
| 2250_at | FGF5 | -0,15718316 | 4,70275801 | -1,61959592 | 0,12682061 | 0,36062982 |
| 1118_at | CHIT1 | -0,14966857 | 4,46406901 | -1,61953644 | 0,12683347 | 0,36062982 |
| 101928429_a | LOC10192842 | 0,1299314 | 3,55827016 | 1,6195187 | 0,12683731 | 0,36062982 |
| 84689_at | MS4A14 | -0,13194953 | 3,32923898 | -1,61919197 | 0,12690796 | 0,36077745 |
| 6342_at | SCP2 | 0,11063466 | 9,71686041 | 1,61895364 | 0,12695952 | 0,36083682 |
| 80067_at | DCAF17 | -0,24458509 | 7,15767538 | -1,6189104 | 0,12696888 | 0,36083682 |
| 80020_at | FOXRED2 | -0,2481337 | 6,99556896 | -1,61883571 | 0,12698504 | 0,36083682 |
| 81555_at | YIPF5 | 0,17174149 | 9,22446756 | 1,61804272 | 0,12715675 | 0,36122357 |
| 8078_at | USP5 | -0,13458957 | 7,77226574 | -1,61803396 | 0,12715865 | 0,36122357 |
| 79656_at | BEND5 | 0,87734228 | 5,2046806 | 1,61782414 | 0,12720412 | 0,36126459 |
| 583_at | BBS2 | 0,38091546 | 8,09830572 | 1,61779424 | 0,1272106 | 0,36126459 |
| 8555_at | CDC14B | 0,28760969 | 6,75776428 | 1,61760466 | 0,12725169 | 0,36132803 |
| 23354_at | HAUS5 | -0,17661408 | 7,12059071 | -1,61741184 | 0,1272935 | 0,36139348 |
| 79957_at | PAQR6 | 0,27468086 | 6,20751887 | 1,61722509 | 0,12733401 | 0,3614552 |
| 5936_at | RBM4 | 0,46941699 | 7,53406161 | 1,61713717 | 0,12735308 | 0,36145608 |
| 29015_at | SLC43A3 | -0,13261005 | 4,73259812 | -1,61696857 | 0,12738966 | 0,36150664 |
| 55698_at | RADIL | -0,233113 | 5,4195937 | -1,61646281 | 0,12749946 | 0,36173113 |
| 23301_at | EHBP1 | 0,21493468 | 8,01974105 | 1,61643118 | 0,12750633 | 0,36173113 |
| 65094_at | JMJD4 | -0,28469548 | 7,30796338 | -1,61621338 | 0,12755363 | 0,36181205 |
| 90024_at | FLJ20021 | 0,26629661 | 5,88526864 | 1,6161064 | 0,12757688 | 0,3618247 |
| 28234_at | SLCO1B3 | -0,18987077 | 4,45460088 | -1,61518545 | 0,12777711 | 0,36231454 |
| 101929241_a | LOC10192924 | -0,17552635 | 3,92361143 | -1,61513906 | 0,12778721 | 0,36231454 |

| | | | | | | |
|---|---|---|---|---|---|---|
| 5498_at | PPOX | 0,16443677 | 4,94008063 | 1,61488596 | 0,12784229 | 0,36238212 |
| 730094_at | C16orf52 | 0,19034651 | 7,167641 | 1,61485666 | 0,12784867 | 0,36238212 |
| 7484_at | WNT9B | -0,17558963 | 4,69024574 | -1,61450169 | 0,12792596 | 0,36253151 |
| 79837_at | PIP4K2C | 0,12898267 | 9,58425429 | 1,61428292 | 0,12797361 | 0,36253151 |
| 149420_at | PDIK1L | 0,219411 | 7,66222946 | 1,61427658 | 0,12797499 | 0,36253151 |
| 9032_at | TM4SF5 | -0,13728926 | 5,19779808 | -1,61419329 | 0,12799314 | 0,36253151 |
| 27242_at | TNFRSF21 | 0,16653942 | 11,6354999 | 1,61418257 | 0,12799547 | 0,36253151 |
| 132241_at | RPL32P3 | 0,27336819 | 5,00527962 | 1,61407407 | 0,12801912 | 0,36254517 |
| 400745_at | SH2D5 | -0,23532281 | 5,58631629 | -1,61378825 | 0,12808142 | 0,36261198 |
| 80745_at | THUMPD2 | -0,31039343 | 6,83362293 | -1,61377661 | 0,12808396 | 0,36261198 |
| 348980_at | HCN1 | 0,0945071 | 3,156744 | 1,61367689 | 0,1281057 | 0,36261198 |
| 7980_at | TFPI2 | -0,45282701 | 7,1116552 | -1,61362045 | 0,12811801 | 0,36261198 |
| 171019_at | ADAMTS19 | 0,09867652 | 3,28082457 | 1,61336184 | 0,12817441 | 0,36265121 |
| 387914_at | SHISA2 | -0,51551423 | 6,47929283 | -1,61332508 | 0,12818243 | 0,36265121 |
| 101928694_a | LOC10192869 | -0,1181396 | 3,15019196 | -1,61329797 | 0,12818835 | 0,36265121 |
| 6927_at | HNF1A | -0,15719849 | 5,85659713 | -1,61316772 | 0,12821676 | 0,36267835 |
| 5479_at | PPIB | -0,08300051 | 11,7701884 | -1,61302239 | 0,12824848 | 0,3627148 |
| 9282_at | MED14 | -0,09191669 | 8,50023285 | -1,61288331 | 0,12827884 | 0,3627474 |
| 400_at | ARL1 | 0,14112272 | 8,89997271 | 1,61274783 | 0,12830842 | 0,36277779 |
| 91582_at | RPS19BP1 | -0,24248153 | 9,47218074 | -1,61256544 | 0,12834825 | 0,36281625 |
| 5455_at | POU3F3 | -0,18492427 | 5,21255033 | -1,61251306 | 0,12835969 | 0,36281625 |
| 9356_at | SLC22A6 | -0,17638599 | 5,14701005 | -1,61191756 | 0,12848982 | 0,36309135 |
| 6281_at | S100A10 | -0,101232 | 12,6331208 | -1,61187488 | 0,12849915 | 0,36309135 |
| 10175_at | CNIH1 | -0,19401727 | 11,0679442 | -1,61180895 | 0,12851357 | 0,36309135 |
| 257364_at | SNX33 | 0,1848983 | 7,88613386 | 1,61163214 | 0,12855223 | 0,3631058 |
| 6194_at | RPS6 | -0,11030281 | 12,553114 | -1,61149528 | 0,12858216 | 0,3631058 |
| 914_at | CD2 | -0,17385531 | 4,36757875 | -1,61145387 | 0,12859122 | 0,3631058 |
| 161145_at | TMEM229B | 0,32671831 | 6,01338555 | 1,61135929 | 0,12861191 | 0,3631058 |
| 286133_at | SCARA5 | -0,18775646 | 4,50667748 | -1,61135464 | 0,12861293 | 0,3631058 |
| 795_at | S100G | -0,1618928 | 3,35008378 | -1,61120782 | 0,12864506 | 0,36314328 |
| 146429_at | SLC22A31 | -0,16232333 | 6,21613478 | -1,61102617 | 0,12868482 | 0,36320229 |
| 9863_at | MAGI2 | 0,27907424 | 4,4259586 | 1,61091386 | 0,1287094 | 0,36321846 |
| 3632_at | INPP5A | 0,20311825 | 9,15035465 | 1,61067614 | 0,12876146 | 0,36327118 |
| 80736_at | SLC44A4 | 0,09572249 | 4,60078296 | 1,61065631 | 0,1287658 | 0,36327118 |
| 84287_at | ZDHHC16 | -0,09331532 | 9,59124839 | -1,61043803 | 0,12881361 | 0,36335286 |
| 125965_at | COX6B2 | -0,21071548 | 5,20336916 | -1,6101797 | 0,12887022 | 0,36345932 |
| 627_at | BDNF | -0,09600545 | 3,84599666 | -1,60958715 | 0,12900015 | 0,36376912 |
| 10313_at | RTN3 | 0,14086432 | 9,27725189 | 1,60950651 | 0,12901784 | 0,36376912 |
| 1327_at | COX4I1 | -0,11200056 | 11,9669618 | -1,60922298 | 0,12908005 | 0,3638489 |
| 58531_at | PRM3 | -0,17648077 | 4,66031971 | -1,6092054 | 0,12908391 | 0,3638489 |
| 84186_at | ZCCHC7 | 0,14624887 | 6,87578129 | 1,60911805 | 0,12910309 | 0,36384971 |
| 84457_at | PHYHIPL | -0,18998471 | 4,67266702 | -1,60901197 | 0,12912637 | 0,3638621 |
| 84226_at | C2orf16 | -0,15004308 | 5,63444233 | -1,60879684 | 0,12917361 | 0,36394198 |
| 286359_at | LOC286359 | -0,1509239 | 4,05332698 | -1,6086231 | 0,12921177 | 0,36399625 |
| 64218_at | SEMA4A | 0,13075331 | 7,9171055 | 1,60831781 | 0,12927885 | 0,36410714 |
| 2280_at | FKBP1A | -0,25068055 | 6,72784176 | -1,6082719 | 0,12928894 | 0,36410714 |
| 54934_at | KANSL2 | -0,18401471 | 8,04245512 | -1,60696873 | 0,12957562 | 0,36486117 |
| 92591_at | ASB16 | -0,2488887 | 6,43078233 | -1,6062973 | 0,12972355 | 0,36517933 |
| 79022_at | TMEM106C | -0,27094342 | 9,95055249 | -1,60620919 | 0,12974297 | 0,36517933 |
| 117159_at | DCD | -0,18121338 | 3,93047973 | -1,60610732 | 0,12976543 | 0,36517933 |

| | | | | | | |
|---|---|---|---|---|---|---|
| 441478_at | NRARP | -0,25765464 | 6,80872376 | -1,60603986 | 0,1297803 | 0,36517933 |
| 43847_at | KLK14 | -0,23867963 | 5,91320727 | -1,60602579 | 0,1297834 | 0,36517933 |
| 79962_at | DNAJC22 | 0,55781675 | 4,96505723 | 1,60586939 | 0,12981789 | 0,36521903 |
| 100288797_a | TMEM239 | -0,10806824 | 5,28721367 | -1,60578987 | 0,12983543 | 0,36521903 |
| 989_at | SEPT7 | -0,19930094 | 6,27108478 | -1,60527823 | 0,12994833 | 0,36546022 |
| 51201_at | ZDHHC2 | 0,74388666 | 8,32664093 | 1,60522937 | 0,12995912 | 0,36546022 |
| 83734_at | ATG10 | 0,26686061 | 5,61241297 | 1,60495372 | 0,13001998 | 0,365578 |
| 123355_at | LRRC28 | 0,18515893 | 6,7014436 | 1,60469892 | 0,13007626 | 0,36568288 |
| 5914_at | RARA | -0,13663938 | 6,11091438 | -1,60454514 | 0,13011024 | 0,36572502 |
| 145864_at | HAPLN3 | -0,16550496 | 6,56461102 | -1,60435546 | 0,13015216 | 0,36578949 |
| 51052_at | PRLH | -0,174821 | 6,05894917 | -1,60404506 | 0,13022079 | 0,36592897 |
| 253725_at | FAM21C | 0,37923908 | 6,89221022 | 1,6037189 | 0,13029293 | 0,36607829 |
| 8350_at | HIST1H3A | -0,32512028 | 5,45792558 | -1,6032585 | 0,13039483 | 0,36629254 |
| 374955_at | SPATA21 | -0,19846222 | 5,25184703 | -1,60317153 | 0,13041408 | 0,36629254 |
| 8971_at | H1FX | 0,34795973 | 8,68411654 | 1,60311666 | 0,13042623 | 0,36629254 |
| 79007_at | DBNDD1 | 0,16372998 | 5,65708797 | 1,60293262 | 0,13046699 | 0,3663536 |
| 6362_at | CCL18 | -0,109066 | 4,15337432 | -1,60274385 | 0,13050881 | 0,36641761 |
| 84293_at | FAM213A | -0,12603852 | 10,4646631 | -1,60253094 | 0,13055599 | 0,36645628 |
| 55357_at | TBC1D2 | 0,37530911 | 8,79191095 | 1,60251 | 0,13056063 | 0,36645628 |
| 4886_at | NPY1R | 1,45067236 | 5,26250004 | 1,6022904 | 0,13060931 | 0,36653951 |
| 254187_at | TSGA10IP | -0,13892421 | 6,49435846 | -1,601614 | 0,13075935 | 0,36690711 |
| 114915_at | EPB41L4A-AS1 | 0,19018425 | 8,19940388 | 1,60126774 | 0,13083621 | 0,3669661 |
| 51063_at | CALHM2 | -0,48298507 | 7,16981662 | -1,60121922 | 0,13084698 | 0,3669661 |
| 10563_at | CXCL13 | -0,10683193 | 2,92449354 | -1,60119777 | 0,13085175 | 0,3669661 |
| 55657_at | ZNF692 | -0,24238943 | 8,00453945 | -1,60117604 | 0,13085657 | 0,3669661 |
| 90060_at | CCDC120 | 0,12320562 | 6,6946829 | 1,60101865 | 0,13089153 | 0,3670107 |
| 165721_at | DNAJB8 | -0,16465895 | 4,79725791 | -1,60076539 | 0,13094779 | 0,36708719 |
| 339_at | APOBEC1 | -0,10826996 | 3,90064476 | -1,6007243 | 0,13095692 | 0,36708719 |
| 64431_at | ACTR6 | 0,1509372 | 7,40871756 | 1,60048521 | 0,13101006 | 0,36718271 |
| 6776_at | STAT5A | 0,45748403 | 6,02782025 | 1,60033952 | 0,13104245 | 0,36722006 |
| 4008_at | LMO7 | 0,18603255 | 5,99207373 | 1,6000292 | 0,13111146 | 0,36736 |
| 54947_at | LPCAT2 | -0,30768096 | 8,44711866 | -1,59941119 | 0,13124899 | 0,36769186 |
| 10053_at | AP1M2 | 0,17195993 | 8,95339792 | 1,59919383 | 0,13129739 | 0,36777397 |
| 54852_at | PAQR5 | -0,71772314 | 5,09422126 | -1,5990274 | 0,13133446 | 0,36782432 |
| 26136_at | TES | 0,15155417 | 10,5559101 | 1,59888024 | 0,13136724 | 0,36786265 |
| 100874251_a | KIRREL3-AS2 | -0,13131395 | 2,95010371 | -1,59876209 | 0,13139357 | 0,3678829 |
| 414196_at | CELF2-AS1 | -0,17970031 | 4,88242788 | -1,59860071 | 0,13142954 | 0,36793012 |
| 4215_at | MAP3K3 | 0,13046035 | 7,18780223 | 1,59803969 | 0,13155464 | 0,36820915 |
| 64090_at | GAL3ST2 | -0,19441398 | 6,03040547 | -1,59798231 | 0,13156744 | 0,36820915 |
| 388_at | RHOB | 0,20838883 | 8,71931425 | 1,59769555 | 0,13163143 | 0,36829067 |
| 79568_at | C2orf47 | -0,20389843 | 9,13209138 | -1,59768041 | 0,13163481 | 0,36829067 |
| 56062_at | KLHL4 | 0,17284624 | 4,96907196 | 1,59744209 | 0,13168801 | 0,36838601 |
| 8778_at | SIGLEC5 | -0,18361782 | 4,44695396 | -1,59731818 | 0,13171568 | 0,36840991 |
| 170954_at | PPP1R18 | -0,14793075 | 8,45696147 | -1,59687344 | 0,13181502 | 0,36863314 |
| 4000_at | LMNA | -0,10531936 | 10,0742412 | -1,59678961 | 0,13183376 | 0,36863314 |
| 10594_at | PRPF8 | -0,09507164 | 9,52779586 | -1,59668371 | 0,13185743 | 0,36864581 |
| 60313_at | GPBP1L1 | 0,16700467 | 9,0749219 | 1,59647131 | 0,13190491 | 0,36872505 |
| 100131303_a | LOC10013130 | -0,22509228 | 4,67330742 | -1,59629915 | 0,13194341 | 0,36877915 |
| 7770_at | ZNF227 | 0,1789865 | 6,79279342 | 1,59616627 | 0,13197313 | 0,3688087 |
| 84270_at | C9orf89 | -0,16323346 | 8,62745276 | -1,59563228 | 0,13209262 | 0,36908909 |

| | | | | | | |
|---|---|---|---|---|---|---|
| 27112_at | FAM155B | -0,17323965 | 5,24007136 | -1,59537333 | 0,1321506 | 0,36915884 |
| 5303_at | PIN4 | 0,14785308 | 6,39454949 | 1,59526493 | 0,13217488 | 0,36915884 |
| 503538_at | A1BG-AS1 | -0,19863735 | 4,54739587 | -1,59526405 | 0,13217508 | 0,36915884 |
| 221264_at | AK9 | 0,28939475 | 4,61682848 | 1,59512706 | 0,13220576 | 0,36917184 |
| 100131187_a | TSTD1 | 1,127789 | 7,15702373 | 1,59507217 | 0,13221806 | 0,36917184 |
| 81839_at | VANGL1 | -0,19400974 | 8,28165953 | -1,59496264 | 0,1322426 | 0,36918684 |
| 6224_at | RPS20 | -0,1599571 | 10,8169688 | -1,59474467 | 0,13229145 | 0,36926729 |
| 85021_at | REPS1 | -0,17502525 | 9,38100075 | -1,59466302 | 0,13230976 | 0,36926729 |
| 9202_at | ZMYM4 | 0,20756081 | 8,74434649 | 1,59433935 | 0,13238233 | 0,36941632 |
| 9196_at | KCNAB3 | -0,16703267 | 4,79017583 | -1,59420234 | 0,13241306 | 0,36944856 |
| 4741_at | NEFM | -0,10263099 | 3,57662886 | -1,59409485 | 0,13243718 | 0,36946232 |
| 196472_at | FAM71C | -0,17889875 | 3,7725248 | -1,59398651 | 0,13246149 | 0,36947663 |
| 9175_at | MAP3K13 | 0,16062501 | 5,64031007 | 1,59366134 | 0,13253447 | 0,36962668 |
| 3572_at | IL6ST | 0,14736884 | 6,35677348 | 1,59349095 | 0,13257273 | 0,36964416 |
| 22823_at | MTF2 | 0,19400745 | 7,58873194 | 1,5933792 | 0,13259783 | 0,36964416 |
| 23518_at | R3HDM1 | -0,48259212 | 8,61873873 | -1,59337706 | 0,13259831 | 0,36964416 |
| 81285_at | OR51E2 | -0,10749629 | 3,74597308 | -1,59324184 | 0,13262868 | 0,36967533 |
| 57705_at | WDFY4 | -0,12845111 | 3,9362026 | -1,59315518 | 0,13264815 | 0,36967611 |
| 5481_at | PPID | -0,33571708 | 8,0395811 | -1,59279052 | 0,1327301 | 0,36985098 |
| 2840_at | GPR17 | -0,15506915 | 4,25795644 | -1,59262983 | 0,13276622 | 0,36988869 |
| 126014_at | OSCAR | -0,13946553 | 4,42642595 | -1,59243003 | 0,13281115 | 0,36988869 |
| 83874_at | TBC1D10A | 0,11644448 | 7,89343024 | 1,59240022 | 0,13281785 | 0,36988869 |
| 84298_at | LLPH | -0,1954388 | 8,10789331 | -1,59238874 | 0,13282044 | 0,36988869 |
| 219557_at | C7orf62 | -0,18990958 | 4,32717559 | -1,59153454 | 0,13301268 | 0,37036506 |
| 6282_at | S100A11 | 0,1129251 | 12,8335986 | 1,59145787 | 0,13302994 | 0,37036506 |
| 7148_at | TNXB | -0,15607079 | 5,06853795 | -1,5909949 | 0,13313425 | 0,37060189 |
| 9100_at | USP10 | -0,30565591 | 8,82827694 | -1,59079694 | 0,13317887 | 0,37067253 |
| 79637_at | ARMC7 | -0,12502515 | 7,07842715 | -1,59038991 | 0,13327066 | 0,37085007 |
| 10022_at | INSL5 | -0,16369633 | 3,57326734 | -1,59034335 | 0,13328116 | 0,37085007 |
| 101928597_a | LOC10192859 | -0,15795418 | 4,26382045 | -1,59017393 | 0,13331938 | 0,37090284 |
| 285154_at | CYP1B1-AS1 | 0,08318917 | 3,19943035 | 1,58956096 | 0,13345776 | 0,37123419 |
| 199221_at | DZIP1L | -0,27731746 | 4,96618443 | -1,58930972 | 0,1335145 | 0,37133842 |
| 100506175_a | LOC10050617 | -0,19539302 | 5,03255498 | -1,58909507 | 0,13356301 | 0,37141292 |
| 51019_at | CCDC53 | 0,25215997 | 8,6901377 | 1,58902053 | 0,13357985 | 0,37141292 |
| 101926963_a | LOC10192696 | -0,15218617 | 4,55306711 | -1,58850496 | 0,13369642 | 0,37168339 |
| 572_at | BAD | -0,13821338 | 8,03337511 | -1,58814258 | 0,13377841 | 0,37185765 |
| 124152_at | IQCK | 0,14650905 | 6,86163511 | 1,58789786 | 0,1338338 | 0,37195468 |
| 55258_at | THNSL2 | 0,35306924 | 5,41201622 | 1,58781777 | 0,13385193 | 0,37195468 |
| 9980_at | DOPEY2 | 0,73103326 | 6,08726905 | 1,58736331 | 0,13395486 | 0,372187 |
| 11236_at | RNF139 | 0,21936539 | 9,49493021 | 1,58692042 | 0,13405523 | 0,37241217 |
| 3454_at | IFNAR1 | 0,2013314 | 7,58089271 | 1,58677001 | 0,13408934 | 0,37243645 |
| 3600_at | IL15 | 0,8039832 | 5,26322797 | 1,58669537 | 0,13410626 | 0,37243645 |
| 7171_at | TPM4 | -0,15836783 | 5,90149464 | -1,58662608 | 0,13412198 | 0,37243645 |
| 348235_at | SKA2 | -0,33309956 | 9,30002371 | -1,58637241 | 0,13417952 | 0,3724589 |
| 100507321_a | ERVK13-1 | 0,4670685 | 5,47089938 | 1,58634557 | 0,13418561 | 0,3724589 |
| 220_at | ALDH1A3 | 0,16468367 | 12,3233765 | 1,58633473 | 0,13418807 | 0,3724589 |
| 115123_at | MARCH3 | -0,14003813 | 5,7628616 | -1,58554746 | 0,1343668 | 0,37281024 |
| 8125_at | ANP32A | -0,26506525 | 10,0977254 | -1,5854038 | 0,13439944 | 0,37281024 |
| 643837_at | LINC01128 | -0,12522515 | 6,1307084 | -1,58529663 | 0,13442379 | 0,37281024 |
| 53345_at | TM6SF2 | -0,15182786 | 4,79285389 | -1,58518792 | 0,13444849 | 0,37281024 |

| | | | | | | |
|---|---|---|---|---|---|---|
| 80262_at | C16orf70 | -0,08704355 | 7,24042963 | -1,58514204 | 0,13445892 | 0,37281024 |
| 100128361_a | LOC10012836 | -0,1806114 | 5,60641495 | -1,58513725 | 0,13446001 | 0,37281024 |
| 1834_at | DSPP | -0,20306685 | 3,13728652 | -1,58510435 | 0,13446749 | 0,37281024 |
| 51676_at | ASB2 | -0,273659 | 4,63862634 | -1,58509559 | 0,13446948 | 0,37281024 |
| 3588_at | IL10RB | 0,45154205 | 8,8540433 | 1,58500484 | 0,13449011 | 0,37281377 |
| 729956_at | SHISA7 | -0,14322526 | 5,21257119 | -1,58488297 | 0,13451781 | 0,37283692 |
| 123099_at | DEGS2 | 0,50544382 | 6,82237485 | 1,58444261 | 0,13461797 | 0,37306085 |
| 256949_at | KANK3 | 0,18783136 | 5,34822209 | 1,58433717 | 0,13464196 | 0,37307366 |
| 114787_at | GPRIN1 | -0,16189032 | 6,40650964 | -1,5839953 | 0,13471977 | 0,37323558 |
| 26576_at | SRPK3 | -0,19830883 | 5,87680228 | -1,58377303 | 0,13477038 | 0,3733221 |
| 5578_at | PRKCA | -0,11343848 | 6,84401113 | -1,58352536 | 0,1348268 | 0,37333067 |
| 7222_at | TRPC3 | 0,10586526 | 3,27677434 | 1,58350537 | 0,13483135 | 0,37333067 |
| 25790_at | CFAP45 | -0,24732499 | 5,82182433 | -1,583474 | 0,1348385 | 0,37333067 |
| 9829_at | DNAJC6 | 0,0870817 | 3,83851906 | 1,58341914 | 0,134851 | 0,37333067 |
| 100505851_a | LOC10050585 | 0,15148409 | 3,97960749 | 1,58256182 | 0,13504646 | 0,37381808 |
| 643714_at | CASC16 | -0,1848268 | 3,83783302 | -1,58231368 | 0,13510308 | 0,37392107 |
| 101928298_a | LOC10192829 | -0,11905173 | 3,00990453 | -1,58202689 | 0,13516855 | 0,37404852 |
| 23386_at | NUDCD3 | -0,12808463 | 7,74527645 | -1,58182904 | 0,13521372 | 0,37411979 |
| 23598_at | PATZ1 | 0,33931778 | 7,37955025 | 1,58133045 | 0,13532763 | 0,37438118 |
| 126326_at | GIPC3 | -0,1737689 | 4,7615977 | -1,5811718 | 0,13536389 | 0,37442772 |
| 400958_at | LOC400958 | -0,14913096 | 3,54706087 | -1,58106053 | 0,13538933 | 0,37444432 |
| 90459_at | ERI1 | 0,19462066 | 7,85449725 | 1,5808537 | 0,13543663 | 0,37452135 |
| 286223_at | C9orf47 | -0,11787633 | 4,50571684 | -1,58069242 | 0,13547352 | 0,37456959 |
| 2914_at | GRM4 | -0,17611086 | 5,72337313 | -1,58054985 | 0,13550613 | 0,374606 |
| 56302_at | TRPV5 | -0,11956666 | 4,65965587 | -1,58043218 | 0,13553306 | 0,37462667 |
| 547_at | KIF1A | -0,15406659 | 5,56971192 | -1,58023114 | 0,13557907 | 0,37470008 |
| 343450_at | KCNT2 | 0,11377157 | 3,10869836 | 1,57978261 | 0,13568177 | 0,37493013 |
| 3101_at | HK3 | -0,15331531 | 5,67171672 | -1,57943022 | 0,13576251 | 0,37509943 |
| 9111_at | NMI | 0,52860596 | 9,12171677 | 1,57927022 | 0,13579918 | 0,37514693 |
| 5866_at | RAB3IL1 | -0,22270001 | 5,59118993 | -1,57906528 | 0,13584616 | 0,37522291 |
| 155060_at | LOC155060 | -0,16818656 | 5,66600919 | -1,57859137 | 0,13595486 | 0,37539568 |
| 23435_at | TARDBP | -0,41498946 | 7,21178808 | -1,57855562 | 0,13596306 | 0,37539568 |
| 5076_at | PAX2 | -0,13818465 | 4,70206968 | -1,57853772 | 0,13596717 | 0,37539568 |
| 54986_at | ULK4 | -0,13109945 | 4,60187785 | -1,5784037 | 0,13599793 | 0,37542678 |
| 7052_at | TGM2 | -0,10731806 | 5,34582576 | -1,57828724 | 0,13602466 | 0,37543421 |
| 9590_at | AKAP12 | 0,23150905 | 5,23040372 | 1,57822215 | 0,1360396 | 0,37543421 |
| 51720_at | UIMC1 | -0,18339398 | 7,94678594 | -1,57790807 | 0,13611172 | 0,37557943 |
| 401152_at | C4orf3 | 0,3065641 | 7,57735631 | 1,57774755 | 0,13614859 | 0,37558304 |
| 80316_at | PPP1R2P9 | -0,20223399 | 4,68483679 | -1,5777326 | 0,13615202 | 0,37558304 |
| 101926996_a | RUNDC3A-AS1 | 0,28903352 | 3,88127161 | 1,5776026 | 0,13618189 | 0,37559869 |
| 100130872_a | LOC10013087 | -0,17872515 | 5,19128868 | -1,57750372 | 0,13620461 | 0,37559869 |
| 400709_at | SIGLEC16 | -0,14105555 | 4,44608806 | -1,577399 | 0,13622868 | 0,37559869 |
| 23743_at | BHMT2 | -0,10787456 | 3,83372152 | -1,57730892 | 0,13624938 | 0,37559869 |
| 8431_at | NR0B2 | -0,13617184 | 5,40331359 | -1,57728368 | 0,13625519 | 0,37559869 |
| 9069_at | CLDN12 | 0,18914761 | 9,35228831 | 1,57704508 | 0,13631005 | 0,37568452 |
| 1774_at | DNASE1L1 | 0,20198633 | 8,1869918 | 1,57697576 | 0,13632599 | 0,37568452 |
| 10241_at | CALCOCO2 | 0,31458507 | 7,721971 | 1,57689383 | 0,13634483 | 0,37568452 |
| 286114_at | LOC286114 | -0,17327583 | 3,49944807 | -1,57666892 | 0,13639657 | 0,37577333 |
| 79934_at | ADCK4 | 0,19166276 | 6,99767511 | 1,57619145 | 0,13650646 | 0,37597459 |
| 8550_at | MAPKAPK5 | -0,13106814 | 8,09926988 | -1,5761307 | 0,13652045 | 0,37597459 |

| | | | | | | |
|---|---|---|---|---|---|---|
| 6588_at | SLN | -0,17590022 | 4,36165135 | -1,57606014 | 0,1365367 | 0,37597459 |
| 2746_at | GLUD1 | -0,42294654 | 8,69580071 | -1,57601239 | 0,13654769 | 0,37597459 |
| 100505812_a | CARD8-AS1 | 0,33422966 | 5,05740186 | 1,57562623 | 0,13663665 | 0,376114 |
| 64411_at | ARAP3 | 0,15254924 | 7,60444024 | 1,57562308 | 0,13663738 | 0,376114 |
| 6774_at | STAT3 | 0,3434624 | 9,91153911 | 1,57524412 | 0,13672472 | 0,37619035 |
| 27147_at | DENND2A | -0,1587686 | 4,32315193 | -1,57520568 | 0,13673359 | 0,37619035 |
| 283742_at | FAM98B | -0,16443024 | 8,20735703 | -1,57516738 | 0,13674242 | 0,37619035 |
| 245937_at | DEFB124 | -0,22765552 | 6,58945226 | -1,57508656 | 0,13676105 | 0,37619035 |
| 3172_at | HNF4A | -0,14830606 | 5,14483262 | -1,57507916 | 0,13676276 | 0,37619035 |
| 2115_at | ETV1 | -0,27173436 | 3,81061173 | -1,57491368 | 0,13680092 | 0,37623597 |
| 85293_at | KRTAP3-3 | -0,11997511 | 3,06909988 | -1,57483789 | 0,13681841 | 0,37623597 |
| 342926_at | ZNF677 | 0,20666483 | 3,20742089 | 1,57471524 | 0,1368467 | 0,37626007 |
| 124401_at | ANKS3 | -0,1222894 | 6,34514174 | -1,57454162 | 0,13688677 | 0,37628617 |
| 84692_at | CCDC54 | -0,13589444 | 3,39437951 | -1,57447164 | 0,13690292 | 0,37628617 |
| 523_at | ATP6V1A | -0,15970714 | 9,46195098 | -1,57429261 | 0,13694424 | 0,37628617 |
| 81797_at | OR12D3 | -0,14972439 | 4,11279702 | -1,57426057 | 0,13695164 | 0,37628617 |
| 116179_at | TGM7 | -0,15537539 | 3,81449318 | -1,57425094 | 0,13695387 | 0,37628617 |
| 10282_at | BET1 | 0,15925734 | 7,16777804 | 1,57409192 | 0,13699058 | 0,37633338 |
| 83723_at | FAM57B | -0,13210295 | 5,44495713 | -1,57395991 | 0,13702107 | 0,37636346 |
| 201158_at | TVP23C | -0,19126947 | 6,03336197 | -1,57367981 | 0,13708578 | 0,37648752 |
| 23220_at | DTX4 | 0,19749265 | 10,7502785 | 1,57262408 | 0,13732992 | 0,37710424 |
| 153579_at | BTNL9 | -0,17957223 | 5,49027226 | -1,57242664 | 0,13737562 | 0,37717596 |
| 121793_at | TEX29 | -0,19494533 | 4,36713829 | -1,57209431 | 0,13745257 | 0,37733344 |
| 54877_at | ZCCHC2 | 0,28052914 | 6,61668515 | 1,57186211 | 0,13750635 | 0,37742731 |
| 101928134_a | LOC10192813 | -0,16555062 | 4,07242205 | -1,57174537 | 0,1375334 | 0,37744777 |
| 1659_at | DHX8 | 0,07886498 | 7,84904576 | 1,57164619 | 0,13755638 | 0,37745706 |
| 26020_at | LRP10 | 0,17388267 | 8,51118687 | 1,57136868 | 0,13762071 | 0,37754864 |
| 2895_at | GRID2 | -0,18116711 | 4,4360716 | -1,57133311 | 0,13762895 | 0,37754864 |
| 51606_at | ATP6V1H | 0,13190885 | 9,08872221 | 1,5711724 | 0,13766622 | 0,3775971 |
| 285852_at | TREML4 | -0,2599955 | 3,97547626 | -1,57106314 | 0,13769156 | 0,37761283 |
| 56975_at | FAM20C | -0,22659569 | 6,12665582 | -1,5698881 | 0,13796433 | 0,37830703 |
| 134_at | ADORA1 | -0,13685748 | 5,51734056 | -1,56978343 | 0,13798865 | 0,37831987 |
| 3018_at | HIST1H2BB | -0,28257218 | 4,7365735 | -1,56960833 | 0,13802934 | 0,37837758 |
| 126119_at | JOSD2 | 0,21315018 | 7,06857419 | 1,56924566 | 0,13811366 | 0,37855484 |
| 57459_at | GATAD2B | 0,17961024 | 7,70642287 | 1,56910205 | 0,13814706 | 0,37859252 |
| 28981_at | IFT81 | 0,14066074 | 6,35268958 | 1,56885354 | 0,13820487 | 0,37860386 |
| 7006_at | TEC | -0,14711615 | 4,99623725 | -1,56883702 | 0,13820871 | 0,37860386 |
| 284415_at | VSTM1 | -0,15850528 | 3,37658654 | -1,56883081 | 0,13821016 | 0,37860386 |
| 56984_at | PSMG2 | -0,29929232 | 10,6638126 | -1,56865446 | 0,1382512 | 0,37866243 |
| 100652772_a | NNT-AS1 | 0,2841385 | 7,82873239 | 1,56828494 | 0,13833723 | 0,37880962 |
| 10002_at | NR2E3 | -0,12568027 | 4,44914385 | -1,56824653 | 0,13834617 | 0,37880962 |
| 160298_at | C11orf42 | -0,28067742 | 4,75256555 | -1,56815046 | 0,13836854 | 0,37880962 |
| 6273_at | S100A2 | -0,56718927 | 11,9402983 | -1,56808583 | 0,1383836 | 0,37880962 |
| 1945_at | EFNA4 | 0,20876109 | 8,3528762 | 1,5675783 | 0,13850186 | 0,37907948 |
| 116983_at | ACAP3 | 0,09732188 | 7,21268622 | 1,56732735 | 0,13856037 | 0,37913612 |
| 165530_at | CLEC4F | -0,17710124 | 4,77364027 | -1,5673207 | 0,13856192 | 0,37913612 |
| 7425_at | VGF | -0,18198938 | 5,18390338 | -1,56704374 | 0,13862652 | 0,37920192 |
| 26077_at | DKFZP434H16 | -0,16368441 | 4,3767636 | -1,56703726 | 0,13862803 | 0,37920192 |
| 5105_at | PCK1 | -0,14180893 | 3,72924853 | -1,56696441 | 0,13864503 | 0,37920192 |
| 10099_at | TSPAN3 | -0,16756991 | 9,28947598 | -1,56684325 | 0,1386733 | 0,3792045 |

| | | | | | | |
|---|---|---|---|---|---|---|
| 26507_at | CNNM1 | -0,1704963 | 4,76633755 | -1,56677153 | 0,13869003 | 0,3792045 |
| 100507388_a | LOC10050738 | -0,14283673 | 4,28023203 | -1,56670729 | 0,13870503 | 0,3792045 |
| 5369_at | PMCHL1 | -0,29164942 | 4,98848722 | -1,56647212 | 0,13875992 | 0,37930075 |
| 26157_at | GIMAP2 | 1,35617085 | 6,18713354 | 1,56624698 | 0,13881249 | 0,37939062 |
| 90378_at | SAMD1 | -0,31313665 | 7,07197246 | -1,56599122 | 0,13887224 | 0,37950006 |
| 133558_at | MROH2B | -0,12382182 | 3,67731166 | -1,56570105 | 0,13894005 | 0,37963025 |
| 51198_at | CDKN2A-AS1 | -0,15504094 | 4,16038779 | -1,56557951 | 0,13896846 | 0,37963025 |
| 3043_at | HBB | -0,13743783 | 5,21420395 | -1,5655344 | 0,138979 | 0,37963025 |
| 78986_at | DUSP26 | -0,14959901 | 5,1586081 | -1,56521683 | 0,13905326 | 0,37977925 |
| 329_at | BIRC2 | 0,20549924 | 9,22008089 | 1,56493165 | 0,13911998 | 0,37986017 |
| 3316_at | HSPB2 | -0,14967619 | 4,4850081 | -1,56492159 | 0,13912233 | 0,37986017 |
| 80742_at | PRR3 | -0,23451821 | 6,91853612 | -1,56458555 | 0,13920098 | 0,37998511 |
| 85397_at | RGS8 | -0,18686566 | 4,47715832 | -1,56453326 | 0,13921323 | 0,37998511 |
| 101929660_a | LOC10192966 | -0,14237505 | 3,17630422 | -1,56447326 | 0,13922727 | 0,37998511 |
| 5348_at | FXYD1 | -0,1796517 | 4,88060498 | -1,5642394 | 0,13928204 | 0,38008073 |
| 388799_at | FAM209B | -0,27365429 | 4,19747751 | -1,56411514 | 0,13931115 | 0,38008308 |
| 26658_at | OR7C2 | -0,12309303 | 5,13667854 | -1,56375172 | 0,1393963 | 0,38008308 |
| 5068_at | REG3A | -0,09884386 | 3,216005 | -1,56372251 | 0,13940315 | 0,38008308 |
| 143662_at | MUC15 | 0,99581676 | 4,40331556 | 1,5636697 | 0,13941553 | 0,38008308 |
| 197322_at | ACSF3 | -0,11184034 | 5,89928589 | -1,56366485 | 0,13941666 | 0,38008308 |
| 6717_at | SRI | 0,19814602 | 10,1136355 | 1,56365142 | 0,13941981 | 0,38008308 |
| 116285_at | ACSM1 | -0,10721538 | 3,68901815 | -1,56364628 | 0,13942102 | 0,38008308 |
| 8973_at | CHRNA6 | -0,13226523 | 4,22645766 | -1,56332879 | 0,13949546 | 0,38021101 |
| 6678_at | SPARC | -0,13877039 | 3,77100809 | -1,5632778 | 0,13950742 | 0,38021101 |
| 43849_at | KLK12 | -0,13017697 | 5,07714524 | -1,56305634 | 0,13955937 | 0,3802835 |
| 684_at | BST2 | 0,77036164 | 9,36812322 | 1,56299611 | 0,1395735 | 0,3802835 |
| 9311_at | ASIC3 | 0,15625775 | 4,84562054 | 1,56272149 | 0,13963795 | 0,3804053 |
| 1298_at | COL9A2 | 0,46130504 | 6,4133919 | 1,56240626 | 0,13971196 | 0,3805531 |
| 10474_at | TADA3 | 0,14847579 | 7,15400282 | 1,56232139 | 0,13973189 | 0,38055358 |
| 26211_at | OR2F1 | -0,13325036 | 4,55567141 | -1,56211185 | 0,13978112 | 0,38063382 |
| 84696_at | ABHD1 | 0,19561495 | 4,72378314 | 1,56202441 | 0,13980166 | 0,38063595 |
| 10643_at | IGF2BP3 | 0,41700699 | 8,74042649 | 1,56187166 | 0,13983755 | 0,38067987 |
| 4836_at | NMT1 | -0,15669058 | 7,99732207 | -1,56157205 | 0,13990798 | 0,38081484 |
| 10577_at | NPC2 | 0,13601432 | 11,428185 | 1,56149256 | 0,13992667 | 0,38081484 |
| 101927310_a | LOC10192731 | -0,11718015 | 2,89610792 | -1,56128603 | 0,13997524 | 0,38089322 |
| 4285_at | MIPEP | -0,39699702 | 6,90537792 | -1,56115456 | 0,14000617 | 0,38092357 |
| 11255_at | HRH3 | -0,11201563 | 5,97091513 | -1,5604808 | 0,14016475 | 0,38129535 |
| 100093630_a | SNHG8 | -0,3217048 | 8,31739657 | -1,5602912 | 0,14020941 | 0,38129535 |
| 51747_at | LUC7L3 | -0,18475749 | 9,09966386 | -1,56027119 | 0,14021412 | 0,38129535 |
| 5606_at | MAP2K3 | -0,34567427 | 8,63143905 | -1,56022269 | 0,14022554 | 0,38129535 |
| 100507428_a | LINC00458 | -0,12331892 | 3,01345272 | -1,55999202 | 0,14027989 | 0,38129535 |
| 647087_at | C7orf73 | 0,17533279 | 9,40150265 | 1,55998531 | 0,14028147 | 0,38129535 |
| 29110_at | TBK1 | 0,14352516 | 8,6855361 | 1,55994784 | 0,1402903 | 0,38129535 |
| 1650_at | DDOST | 0,11491488 | 11,3772431 | 1,55979992 | 0,14032516 | 0,38129535 |
| 1407_at | CRY1 | -0,25441513 | 8,08049442 | -1,55975975 | 0,14033463 | 0,38129535 |
| 84919_at | PPP1R15B | 0,15735072 | 9,80382396 | 1,55973377 | 0,14034076 | 0,38129535 |
| 57157_at | PHTF2 | -0,12738313 | 6,64323696 | -1,5595588 | 0,14038201 | 0,38135364 |
| 100129271_a | C1orf68 | -0,221483 | 5,28816739 | -1,55944201 | 0,14040955 | 0,38137467 |
| 147686_at | ZNF418 | -0,18031741 | 4,39862877 | -1,55925548 | 0,14045355 | 0,38144039 |
| 4905_at | NSF | 0,18943688 | 9,47101626 | 1,55908357 | 0,1404941 | 0,38149675 |

| | | | | | | |
|---|---|---|---|---|---|---|
| 81788_at | NUAK2 | 0,23113604 | 7,43561003 | 1,55880135 | 0,14056071 | 0,38157926 |
| 100506443_a | ITPKB-IT1 | -0,15591418 | 5,5600724 | -1,55878695 | 0,14056411 | 0,38157926 |
| 79085_at | SLC25A23 | 0,25081163 | 5,65236925 | 1,55854619 | 0,14062095 | 0,38165788 |
| 111_at | ADCY5 | -0,29459337 | 4,94422237 | -1,55849644 | 0,1406327 | 0,38165788 |
| 793_at | CALB1 | -0,15078034 | 6,25897476 | -1,55820087 | 0,14070251 | 0,38179356 |
| 114569_at | MAL2 | -0,35689719 | 11,6383805 | -1,55786779 | 0,14078122 | 0,38193511 |
| 6523_at | SLC5A1 | -0,11890943 | 4,07949063 | -1,55781232 | 0,14079433 | 0,38193511 |
| 9512_at | PMPCB | -0,12027776 | 7,82092447 | -1,55767115 | 0,1408277 | 0,38197185 |
| 8552_at | INE1 | -0,12842788 | 4,99587981 | -1,55736012 | 0,14090126 | 0,38211755 |
| 25966_at | C2CD2 | 0,26028661 | 8,72825896 | 1,55721124 | 0,14093648 | 0,38215926 |
| 3556_at | IL1RAP | -0,42504947 | 7,07314868 | -1,55684001 | 0,14102433 | 0,38231725 |
| 56923_at | NMUR2 | -0,20628706 | 4,99530654 | -1,55679729 | 0,14103444 | 0,38231725 |
| 9361_at | LONP1 | -0,33255089 | 8,27261269 | -1,55664465 | 0,14107058 | 0,38233653 |
| 160518_at | DENND5B | -0,11355447 | 3,78119698 | -1,55658399 | 0,14108494 | 0,38233653 |
| 154007_at | SNRNP48 | -0,42620913 | 7,16421936 | -1,55651577 | 0,1411011 | 0,38233653 |
| 118856_at | MMP21 | -0,15485904 | 3,65498881 | -1,55629363 | 0,14115371 | 0,3823791 |
| 56934_at | CA10 | -0,11347129 | 3,55016165 | -1,55628183 | 0,14115651 | 0,3823791 |
| 2113_at | ETS1 | 0,52686256 | 8,11149437 | 1,55597736 | 0,14122865 | 0,3824927 |
| 645323_at | LINC00461 | 0,10451122 | 4,09600311 | 1,55593726 | 0,14123816 | 0,3824927 |
| 9546_at | APBA3 | 0,08841542 | 6,53989565 | 1,55568707 | 0,14129746 | 0,38259639 |
| 55330_at | BLOC1S4 | -0,20295709 | 8,42030044 | -1,55560819 | 0,14131617 | 0,38259639 |
| 284801_at | MIR663AHG | -0,15022883 | 3,18244062 | -1,55440546 | 0,14160161 | 0,38329505 |
| 1355_at | COX15 | -0,13383289 | 7,5827275 | -1,55435322 | 0,14161402 | 0,38329505 |
| 10462_at | CLEC10A | -0,17485262 | 4,20645913 | -1,55424306 | 0,14164019 | 0,38331203 |
| 84659_at | RNASE7 | 0,55702993 | 4,94340255 | 1,55395758 | 0,14170803 | 0,38341977 |
| 5165_at | PDK3 | -0,16463012 | 7,33369879 | -1,55390802 | 0,14171981 | 0,38341977 |
| 101928378_a | PTOV1-AS2 | -0,13214967 | 6,43785452 | -1,55359366 | 0,14179455 | 0,3835681 |
| 81932_at | HDHD3 | 0,15609771 | 7,08960348 | 1,5534441 | 0,14183012 | 0,38361045 |
| 100144597_a | LINC01056 | -0,20037668 | 4,4747102 | -1,55328763 | 0,14186734 | 0,38365726 |
| 7769_at | ZNF226 | 0,26136014 | 6,47180177 | 1,55310274 | 0,14191134 | 0,38372235 |
| 10923_at | SUB1 | -0,22217433 | 9,23557937 | -1,55276371 | 0,14199204 | 0,3838144 |
| 91056_at | AP5B1 | 0,1304981 | 5,52090368 | 1,55266691 | 0,14201509 | 0,3838144 |
| 55810_at | FOXJ2 | -0,17145889 | 7,9967193 | -1,55264311 | 0,14202075 | 0,3838144 |
| 5428_at | POLG | -0,09182751 | 7,10635681 | -1,55260725 | 0,14202929 | 0,3838144 |
| 4295_at | MLN | -0,20057712 | 5,39316714 | -1,55248594 | 0,14205818 | 0,3838144 |
| 1060_at | CENPC | 0,14166947 | 7,32517992 | 1,55245762 | 0,14206493 | 0,3838144 |
| 79991_at | OBFC1 | -0,11312087 | 5,85620014 | -1,55223247 | 0,14211857 | 0,3838959 |
| 54883_at | CWC25 | -0,10266598 | 7,33107981 | -1,55214569 | 0,14213924 | 0,3838959 |
| 5090_at | PBX3 | -0,27655494 | 7,76270303 | -1,55206898 | 0,14215753 | 0,3838959 |
| 2844_at | GPR21 | -0,13146657 | 4,45517186 | -1,55199425 | 0,14217534 | 0,3838959 |
| 284366_at | KLK9 | -0,21761347 | 5,87209845 | -1,55179557 | 0,1422227 | 0,3838959 |
| 64855_at | FAM129B | -0,12774669 | 8,18904816 | -1,55174533 | 0,14223468 | 0,3838959 |
| 10589_at | DRAP1 | -0,26323924 | 9,12071107 | -1,55172057 | 0,14224058 | 0,3838959 |
| 9080_at | CLDN9 | 0,13087235 | 6,0459831 | 1,55166208 | 0,14225453 | 0,3838959 |
| 54106_at | TLR9 | -0,16114422 | 5,25396793 | -1,55155713 | 0,14227956 | 0,38390966 |
| 3557_at | IL1RN | 0,37333622 | 7,56644847 | 1,5512384 | 0,14235559 | 0,38403796 |
| 149041_at | RC3H1 | 0,09992767 | 8,21605061 | 1,55119068 | 0,14236698 | 0,38403796 |
| 93986_at | FOXP2 | 0,1829038 | 4,68284579 | 1,55101426 | 0,14240908 | 0,38406612 |
| 27031_at | NPHP3 | 0,49258984 | 6,16864148 | 1,55097984 | 0,14241729 | 0,38406612 |
| 112869_at | SGF29 | -0,13009602 | 8,32035532 | -1,5506517 | 0,14249564 | 0,38418415 |

| | | | | | | |
|---|---|---|---|---|---|---|
| 118670_at | FAM24A | -0,12772976 | 3,42800138 | -1,55059722 | 0,14250865 | 0,38418415 |
| 91937_at | TIMD4 | -0,16082045 | 3,72285235 | -1,55054596 | 0,14252089 | 0,38418415 |
| 64067_at | NPAS3 | -0,0949967 | 3,53839219 | -1,55014726 | 0,14261614 | 0,38438713 |
| 254099_at | LINC01342 | -0,15974677 | 5,69577814 | -1,54967117 | 0,14272996 | 0,38464006 |
| 100631378_a | LOC10063137 | 0,09464861 | 3,3700679 | 1,54940413 | 0,14279383 | 0,38475836 |
| 284297_at | SSC5D | -0,17198613 | 5,21857938 | -1,54831876 | 0,14305368 | 0,38540462 |
| 80267_at | EDEM3 | -0,20143791 | 7,24635737 | -1,54805908 | 0,14311591 | 0,38546919 |
| 54853_at | WDR55 | -0,09916361 | 8,19262855 | -1,54805175 | 0,14311767 | 0,38546919 |
| 339166_at | LOC339166 | -0,23125635 | 4,17278749 | -1,54796145 | 0,14313931 | 0,38547359 |
| 203228_at | C9orf72 | 0,11831306 | 4,3514387 | 1,54765713 | 0,14321228 | 0,38561618 |
| 9626_at | GUCA1C | -0,09666122 | 2,88723692 | -1,54748639 | 0,14325323 | 0,38565695 |
| 9564_at | BCAR1 | -0,23329869 | 6,67737307 | -1,54735554 | 0,14328462 | 0,38565695 |
| 3755_at | KCNG1 | 0,32642735 | 5,94794069 | 1,54734362 | 0,14328748 | 0,38565695 |
| 64699_at | TMPRSS3 | 0,51073211 | 5,15236545 | 1,5472498 | 0,14330999 | 0,38566365 |
| 158399_at | ZNF483 | -0,10836657 | 3,32965004 | -1,54712334 | 0,14334034 | 0,38569144 |
| 9603_at | NFE2L3 | -0,18298168 | 6,85797437 | -1,54697097 | 0,14337692 | 0,38573209 |
| 55633_at | TBC1D22B | 0,10977911 | 6,585678 | 1,54689357 | 0,1433955 | 0,38573209 |
| 8379_at | MAD1L1 | -0,28299875 | 7,09807103 | -1,54672833 | 0,14343518 | 0,38578495 |
| 26872_at | STEAP1 | -0,29180179 | 4,1540074 | -1,54662497 | 0,14346 | 0,38579784 |
| 196743_at | PAOX | 0,10558905 | 6,83104743 | 1,54640138 | 0,14351371 | 0,38584958 |
| 55116_at | TMEM39B | -0,29437954 | 7,70737879 | -1,54631401 | 0,1435347 | 0,38584958 |
| 90273_at | CEACAM21 | -0,14306009 | 4,12196031 | -1,54623647 | 0,14355333 | 0,38584958 |
| 8549_at | LGR5 | -0,14239016 | 3,27176805 | -1,54621138 | 0,14355936 | 0,38584958 |
| 160418_at | TMTC3 | -0,36935811 | 8,964242 | -1,54539059 | 0,14375672 | 0,38629343 |
| 118738_at | ZNF488 | -0,35664458 | 7,16005074 | -1,54535783 | 0,14376461 | 0,38629343 |
| 92922_at | CCDC102A | 0,29037034 | 6,3366553 | 1,54526459 | 0,14378704 | 0,38629983 |
| 55268_at | ECHDC2 | 0,36674651 | 7,67890655 | 1,54510475 | 0,14382551 | 0,38631932 |
| 100506769_a | C10orf71-AS1 | -0,140313 | 4,43650654 | -1,54495763 | 0,14386093 | 0,38631932 |
| 219654_at | ZCCHC24 | 0,26149239 | 7,4190169 | 1,54494784 | 0,14386328 | 0,38631932 |
| 149428_at | BNIPL | 0,39114856 | 4,32840299 | 1,54490118 | 0,14387452 | 0,38631932 |
| 220296_at | HEPACAM | -0,18588958 | 4,83326727 | -1,54464992 | 0,14393502 | 0,38642792 |
| 84314_at | TMEM107 | -0,33272016 | 6,41446472 | -1,54448525 | 0,14397469 | 0,38648055 |
| 9724_at | UTP14C | 0,13886263 | 8,29438055 | 1,54395098 | 0,14410344 | 0,38676562 |
| 80256_at | FAM214B | 0,21441397 | 6,45403191 | 1,54387799 | 0,14412104 | 0,38676562 |
| 339829_at | CCDC39 | -0,14302039 | 2,9849312 | -1,54368118 | 0,1441685 | 0,3868295 |
| 2637_at | GBX2 | -0,14959728 | 4,73386899 | -1,5434944 | 0,14421356 | 0,3868295 |
| 655_at | BMP7 | -0,40441978 | 5,00701966 | -1,54333309 | 0,14425248 | 0,3868295 |
| 22834_at | ZNF652 | 0,19980895 | 7,60623068 | 1,54330023 | 0,1442604 | 0,3868295 |
| 55311_at | ZNF444 | 0,10982718 | 7,12187219 | 1,54328055 | 0,14426515 | 0,3868295 |
| 339834_at | CCDC36 | -0,13929249 | 4,41498686 | -1,54326865 | 0,14426802 | 0,3868295 |
| 399670_at | RPL13AP17 | -0,12407665 | 5,20358083 | -1,54315376 | 0,14429575 | 0,3868295 |
| 158798_at | AKAP14 | 0,11822467 | 3,03302264 | 1,54311339 | 0,1443055 | 0,3868295 |
| 64093_at | SMOC1 | -0,1598642 | 5,04678514 | -1,54297422 | 0,14433909 | 0,38685679 |
| 5507_at | PPP1R3C | 0,75338833 | 5,99749375 | 1,54290483 | 0,14435584 | 0,38685679 |
| 100507059_a | LINC00703 | -0,14435896 | 3,48679333 | -1,54274192 | 0,14439518 | 0,3868821 |
| 283638_at | CEP170B | -0,18226546 | 9,10402751 | -1,54269938 | 0,14440546 | 0,3868821 |
| 3747_at | KCNC2 | -0,10877848 | 3,73391101 | -1,54258999 | 0,14443188 | 0,38689908 |
| 200959_at | GABRR3 | -0,10044478 | 3,04956224 | -1,54243056 | 0,14447039 | 0,38694844 |
| 84931_at | LINC01101 | -0,16662619 | 4,61475718 | -1,54213181 | 0,14454259 | 0,3870532 |
| 10184_at | LHFPL2 | 0,35397041 | 8,60167159 | 1,54210242 | 0,14454969 | 0,3870532 |

| | | | | | | |
|---|---|---|---|---|---|---|
| 2570_at | GABRR2 | -0,184786 | 5,12654462 | -1,54179795 | 0,1446233 | 0,38715738 |
| 101929315_a | LOC10192931 | -0,11544751 | 4,52293498 | -1,54174568 | 0,14463595 | 0,38715738 |
| 55216_at | C11orf57 | 0,1587523 | 7,41049046 | 1,54165703 | 0,14465739 | 0,38715738 |
| 55120_at | FANCL | -0,18334319 | 7,81186981 | -1,54158387 | 0,14467508 | 0,38715738 |
| 3758_at | KCNJ1 | -0,11948303 | 3,41218664 | -1,54152596 | 0,14468909 | 0,38715738 |
| 132671_at | SPATA18 | 0,16653292 | 4,59168564 | 1,54118715 | 0,14477108 | 0,38725817 |
| 154141_at | MBOAT1 | 0,22665521 | 9,29671749 | 1,54115482 | 0,1447789 | 0,38725817 |
| 340037_at | PRR7-AS1 | -0,14055153 | 4,49386387 | -1,54112106 | 0,14478707 | 0,38725817 |
| 84912_at | SLC35B4 | -0,1809725 | 7,3937784 | -1,54100851 | 0,14481432 | 0,38727727 |
| 23019_at | CNOT1 | -0,10165424 | 8,95850783 | -1,54085078 | 0,14485251 | 0,3872956 |
| 2157_at | F8 | 0,41553357 | 5,41685795 | 1,54081413 | 0,14486138 | 0,3872956 |
| 8303_at | SNN | 0,48074756 | 7,93996866 | 1,54018241 | 0,14501443 | 0,38765098 |
| 3627_at | CXCL10 | 1,0380049 | 5,02501177 | 1,53993729 | 0,14507385 | 0,38767253 |
| 2527_at | FUT5 | -0,18947259 | 5,07109876 | -1,53992928 | 0,14507579 | 0,38767253 |
| 9810_at | RNF40 | -0,17204035 | 7,5475112 | -1,5399001 | 0,14508287 | 0,38767253 |
| 1493_at | CTLA4 | -0,12037769 | 3,58511621 | -1,53974963 | 0,14511936 | 0,38771625 |
| 81689_at | ISCA1 | -0,12028243 | 9,26456563 | -1,53932558 | 0,14522223 | 0,3879373 |
| 2737_at | GLI3 | 0,25551343 | 7,61261001 | 1,53916884 | 0,14526027 | 0,38798511 |
| 100505758_a | PRMT5-AS1 | -0,35070858 | 4,52690629 | -1,53850616 | 0,1454212 | 0,38836109 |
| 51706_at | CYB5R1 | 0,34063632 | 9,70872983 | 1,53817605 | 0,14550142 | 0,38852146 |
| 5586_at | PKN2 | 0,19932527 | 9,57181785 | 1,53795055 | 0,14555624 | 0,38861397 |
| 3767_at | KCNJ11 | -0,17446099 | 4,57085346 | -1,53760673 | 0,14563986 | 0,38878334 |
| 80852_at | GRIP2 | -0,14975583 | 6,11044739 | -1,5368949 | 0,1458131 | 0,38915689 |
| 29988_at | SLC2A8 | -0,14971779 | 7,80566619 | -1,53686578 | 0,1458202 | 0,38915689 |
| 23061_at | TBC1D9B | 0,1288776 | 8,3730188 | 1,53662552 | 0,14587871 | 0,38925913 |
| 155054_at | ZNF425 | 0,14171204 | 5,36100543 | 1,53653469 | 0,14590084 | 0,38926425 |
| 11166_at | SOX21 | -0,19220331 | 5,09206862 | -1,53604831 | 0,14601938 | 0,38951838 |
| 64921_at | CASD1 | 0,37054545 | 6,7070757 | 1,53597796 | 0,14603653 | 0,38951838 |
| 8649_at | LAMTOR3 | 0,36226912 | 7,44751483 | 1,53575555 | 0,14609077 | 0,38960909 |
| 6689_at | SPIB | -0,15610221 | 4,61957703 | -1,53539261 | 0,14617931 | 0,38979126 |
| 6923_at | TCEB2 | -0,09497023 | 8,36343706 | -1,53530513 | 0,14620066 | 0,38979423 |
| 100506898_a | MAGOH2P | -0,22867136 | 5,33811453 | -1,53497248 | 0,14628186 | 0,38995676 |
| 3211_at | HOXB1 | -0,21761264 | 5,28630944 | -1,53473346 | 0,14634024 | 0,39005839 |
| 100129361_a | SMIM10L1 | 0,11516752 | 6,60459515 | 1,53451544 | 0,1463935 | 0,3901431 |
| 168544_at | ZNF467 | 0,31213087 | 5,69081326 | 1,53443756 | 0,14641253 | 0,3901431 |
| 10307_at | APBB3 | 0,24838549 | 6,96500869 | 1,53418515 | 0,14647421 | 0,39022973 |
| 65268_at | WNK2 | -0,18668751 | 5,79232631 | -1,53413877 | 0,14648555 | 0,39022973 |
| 113655_at | MFSD3 | -0,15562859 | 7,50689794 | -1,53387169 | 0,14655085 | 0,39034971 |
| 56300_at | IL36G | -0,16278186 | 4,19228939 | -1,53377858 | 0,14657362 | 0,39035638 |
| 10067_at | SCAMP3 | -0,12338647 | 9,34240858 | -1,53342265 | 0,1466607 | 0,390483 |
| 84524_at | ZC3H8 | -0,22972703 | 7,06123402 | -1,53341853 | 0,14666171 | 0,390483 |
| 10402_at | ST3GAL6 | -0,10075563 | 4,32918841 | -1,5332895 | 0,14669329 | 0,3905131 |
| 27300_at | ZNF544 | -0,29641487 | 7,05668155 | -1,53317254 | 0,14672191 | 0,39053534 |
| 652995_at | UCA1 | -0,32555005 | 11,3807548 | -1,53270792 | 0,14683569 | 0,39078417 |
| 7090_at | TLE3 | -0,13981194 | 7,95864863 | -1,53225623 | 0,14694636 | 0,3910247 |
| 4762_at | NEUROG1 | -0,16545958 | 5,23574359 | -1,53199683 | 0,14700996 | 0,39111782 |
| 55201_at | MAP1S | 0,22670067 | 6,76086332 | 1,53194784 | 0,14702197 | 0,39111782 |
| 4724_at | NDUFS4 | -0,23772628 | 8,1470456 | -1,53150601 | 0,14713034 | 0,39135209 |
| 100885775_a | BANCR | -0,10760392 | 3,59800962 | -1,53096264 | 0,14726372 | 0,39161496 |
| 10717_at | AP4B1 | -0,12579994 | 4,92353187 | -1,53091568 | 0,14727525 | 0,39161496 |

| | | | | | | |
|---|---|---|---|---|---|---|
| 29766_at | TMOD3 | 0,12642657 | 8,7836841 | 1,5307385 | 0,14731877 | 0,39161496 |
| 5833_at | PCYT2 | -0,09844468 | 7,34234652 | -1,53073745 | 0,14731903 | 0,39161496 |
| 101929060_a | LOC10192906 | -0,16801943 | 3,51878139 | -1,53068943 | 0,14733082 | 0,39161496 |
| 664_at | BNIP3 | -0,35068203 | 8,76958867 | -1,53055412 | 0,14736406 | 0,39164927 |
| 116173_at | CMTM5 | -0,14851808 | 5,28078826 | -1,53001964 | 0,14749543 | 0,39194433 |
| 118421_at | LINC00161 | -0,09494275 | 2,86453357 | -1,5294757 | 0,14762923 | 0,3922032 |
| 54988_at | ACSM5 | -0,15058495 | 3,97238594 | -1,52945806 | 0,14763357 | 0,3922032 |
| 10661_at | KLF1 | -0,18254246 | 5,12863424 | -1,5293599 | 0,14765772 | 0,39221329 |
| 9362_at | CPNE6 | -0,13170919 | 4,93452506 | -1,52905644 | 0,14773242 | 0,39233304 |
| 2217_at | FCGRT | 0,48427074 | 6,25558843 | 1,52883904 | 0,14778596 | 0,39233304 |
| 50846_at | DHH | -0,17199012 | 4,08587497 | -1,52881323 | 0,14779232 | 0,39233304 |
| 84941_at | HSH2D | 0,62552698 | 5,29572121 | 1,52874373 | 0,14780943 | 0,39233304 |
| 119_at | ADD2 | -0,13402401 | 5,02669556 | -1,52869607 | 0,14782117 | 0,39233304 |
| 7903_at | ST8SIA4 | 0,49023737 | 4,77319398 | 1,5286313 | 0,14783713 | 0,39233304 |
| 57511_at | COG6 | 0,19664518 | 6,848867 | 1,52859782 | 0,14784538 | 0,39233304 |
| 100292680_a | LINC00942 | -0,15979013 | 5,13228364 | -1,52766413 | 0,14807557 | 0,39283529 |
| 732_at | C8B | -0,20173735 | 4,35164031 | -1,52765623 | 0,14807752 | 0,39283529 |
| 89870_at | TRIM15 | -0,10519231 | 5,42616472 | -1,52758202 | 0,14809582 | 0,39283529 |
| 201140_at | DHRS7C | -0,11813786 | 4,04697958 | -1,52744795 | 0,14812891 | 0,39286894 |
| 29_at | ABR | 0,10829814 | 6,72108566 | 1,52724983 | 0,1481778 | 0,39291133 |
| 3667_at | IRS1 | -0,32484743 | 7,82092902 | -1,52721791 | 0,14818568 | 0,39291133 |
| 51691_at | LSM8 | 0,11160162 | 5,49042919 | 1,5269135 | 0,14826084 | 0,39302175 |
| 54497_at | HEATR5B | 0,29373362 | 7,89787031 | 1,52688398 | 0,14826813 | 0,39302175 |
| 56891_at | LGALS14 | -0,13184485 | 4,32820571 | -1,52660554 | 0,14833691 | 0,39309278 |
| 57111_at | RAB25 | 0,22479218 | 10,2304771 | 1,52658189 | 0,14834275 | 0,39309278 |
| 9640_at | ZNF592 | 0,17385733 | 8,301737 | 1,52652768 | 0,14835615 | 0,39309278 |
| 5007_at | OSBP | 0,11360657 | 8,72019174 | 1,52638329 | 0,14839183 | 0,39313325 |
| 196872_at | LINC00638 | -0,12975189 | 6,3111175 | -1,52621796 | 0,14843269 | 0,39315951 |
| 79230_at | ZNF557 | 0,16991924 | 6,12046607 | 1,52617804 | 0,14844256 | 0,39315951 |
| 51802_at | ASIC5 | -0,11496951 | 3,30775011 | -1,52577756 | 0,14854159 | 0,39333872 |
| 101927526_a | LOC10192752 | -0,14974324 | 4,85877617 | -1,52573927 | 0,14855107 | 0,39333872 |
| 11054_at | OGFR | 0,12972584 | 8,42869424 | 1,52528544 | 0,14866337 | 0,39358197 |
| 7805_at | LAPTM5 | -0,60677068 | 6,84780996 | -1,52500414 | 0,14873301 | 0,39371223 |
| 317671_at | RFESD | -0,2464217 | 5,81098955 | -1,52489915 | 0,14875901 | 0,39372116 |
| 10473_at | HMGN4 | 0,20345104 | 9,78162471 | 1,52482547 | 0,14877726 | 0,39372116 |
| 84067_at | FAM160A2 | 0,17163613 | 6,93143576 | 1,52449617 | 0,14885884 | 0,39388293 |
| 23553_at | HYAL4 | -0,22589491 | 3,87113693 | -1,52395898 | 0,148992 | 0,39412971 |
| 162_at | AP1B1 | -0,08196547 | 8,62318518 | -1,52393318 | 0,1489984 | 0,39412971 |
| 462_at | SERPINC1 | -0,12804032 | 3,6518327 | -1,52387237 | 0,14901348 | 0,39412971 |
| 883_at | CCBL1 | -0,21093957 | 6,57874476 | -1,52375249 | 0,14904321 | 0,39415423 |
| 339789_at | LINC00299 | -0,14833887 | 4,08877844 | -1,52363598 | 0,14907212 | 0,39417656 |
| 11283_at | CYP4F8 | 0,39550946 | 4,46290006 | 1,52355253 | 0,14909282 | 0,3941772 |
| 28974_at | C19orf53 | -0,19773865 | 10,4130586 | -1,52326655 | 0,1491638 | 0,39431072 |
| 390059_at | OR51M1 | -0,1425567 | 4,2254132 | -1,52301735 | 0,14922567 | 0,39442014 |
| 57452_at | GALNT16 | -0,1605424 | 4,57148388 | -1,52271906 | 0,14929975 | 0,39456182 |
| 25864_at | ABHD14A | 0,29651188 | 8,13892665 | 1,52247716 | 0,14935985 | 0,39466651 |
| 10255_at | HCG9 | -0,20371989 | 4,85869509 | -1,52218976 | 0,14943129 | 0,39479073 |
| 57409_at | MIF4GD | 0,15231831 | 7,34147953 | 1,52212312 | 0,14944785 | 0,39479073 |
| 51455_at | REV1 | 0,11680836 | 7,0452087 | 1,52173522 | 0,14954432 | 0,39490149 |
| 54778_at | RNF111 | 0,15349938 | 8,76716385 | 1,52169793 | 0,1495536 | 0,39490149 |

| | | | | | | |
|---|---|---|---|---|---|---|
| 54935_at | DUSP23 | 0,09160528 | 8,79120707 | 1,521659 | 0,14956329 | 0,39490149 |
| 128653_at | C20orf141 | -0,19685975 | 5,39945828 | -1,52158707 | 0,14958118 | 0,39490149 |
| 22824_at | HSPA4L | -0,58163116 | 6,53588024 | -1,52154244 | 0,14959228 | 0,39490149 |
| 80184_at | CEP290 | 0,15547629 | 5,77505792 | 1,52144011 | 0,14961775 | 0,39491459 |
| 27130_at | INVS | -0,23296393 | 6,9980591 | -1,52095436 | 0,14973868 | 0,39509576 |
| 4249_at | MGAT5 | -0,13806527 | 7,72642095 | -1,52092107 | 0,14974697 | 0,39509576 |
| 100379174_a | MACROD2-AS | -0,09983494 | 3,37148888 | -1,52086713 | 0,1497604 | 0,39509576 |
| 253190_at | SERHL2 | 0,28844369 | 4,53970957 | 1,5208349 | 0,14976843 | 0,39509576 |
| 400798_at | C1orf220 | 0,09464784 | 3,53070233 | 1,52054731 | 0,14984008 | 0,39523064 |
| 116442_at | RAB39B | 0,13107445 | 3,22544009 | 1,52012611 | 0,14994506 | 0,3954534 |
| 6328_at | SCN3A | 0,09402503 | 2,99680463 | 1,52002499 | 0,14997027 | 0,39546575 |
| 2294_at | FOXF1 | -0,22931716 | 4,78755627 | -1,51986214 | 0,15001088 | 0,39551723 |
| 9911_at | TMCC2 | -0,14091006 | 6,04597853 | -1,51973465 | 0,15004268 | 0,39551723 |
| 25849_at | PARM1 | 0,19620693 | 5,23509407 | 1,51969975 | 0,15005139 | 0,39551723 |
| 10316_at | NMUR1 | -0,14913396 | 4,81067566 | -1,51938025 | 0,15013112 | 0,39567324 |
| 8799_at | PEX11B | 0,31335392 | 8,37241078 | 1,51896431 | 0,15023496 | 0,39589275 |
| 23647_at | ARFIP2 | -0,1892194 | 7,49862205 | -1,51849773 | 0,15035151 | 0,3961457 |
| 57715_at | SEMA4G | -0,1263399 | 4,51963668 | -1,51810562 | 0,15044953 | 0,39630075 |
| 84264_at | HAGHL | -0,24286568 | 6,45863926 | -1,5180977 | 0,15045151 | 0,39630075 |
| 2309_at | FOXO3 | 0,26101551 | 7,79549484 | 1,51796359 | 0,15048504 | 0,39633488 |
| 2882_at | GPX7 | -0,12366517 | 3,52051382 | -1,51758997 | 0,1505785 | 0,39652682 |
| 27005_at | USP21 | 0,23692529 | 8,10544996 | 1,51747461 | 0,15060737 | 0,39654862 |
| 51289_at | RXFP3 | -0,19339957 | 4,44490404 | -1,51733733 | 0,15064172 | 0,39658487 |
| 25777_at | SUN2 | 0,15136217 | 8,7322263 | 1,51715645 | 0,150687 | 0,39658928 |
| 388581_at | FAM132A | -0,15205525 | 5,02271806 | -1,51715436 | 0,15068753 | 0,39658928 |
| 57124_at | CD248 | -0,13743905 | 5,82649674 | -1,51708393 | 0,15070516 | 0,39658928 |
| 115811_at | IQCD | 0,13457096 | 5,1965858 | 1,51698377 | 0,15073024 | 0,39659406 |
| 6755_at | SSTR5 | -0,14476444 | 5,14067242 | -1,51691223 | 0,15074816 | 0,39659406 |
| 286144_at | TRIQK | 0,2236011 | 7,82156669 | 1,5167436 | 0,15079039 | 0,39665101 |
| 22888_at | UBOX5 | -0,08714376 | 6,42196373 | -1,51662259 | 0,15082071 | 0,39667659 |
| 84779_at | NAA11 | -0,20606358 | 4,1589545 | -1,51629852 | 0,15090192 | 0,39678943 |
| 3881_at | KRT31 | -0,08842832 | 4,45569474 | -1,51621251 | 0,15092349 | 0,39678943 |
| 26716_at | OR2H1 | -0,11415573 | 3,59345847 | -1,51620484 | 0,15092541 | 0,39678943 |
| 101929123_a | LOC10192912 | -0,17711567 | 4,47497202 | -1,51606305 | 0,15096096 | 0,39682873 |
| 50514_at | DEC1 | -0,12638419 | 3,3737731 | -1,51565429 | 0,15106348 | 0,39704405 |
| 126205_at | NLRP8 | -0,13793541 | 5,06153296 | -1,51539783 | 0,15112784 | 0,39715901 |
| 9913_at | SUPT7L | 0,12764262 | 7,64250956 | 1,51489731 | 0,1512535 | 0,39743503 |
| 192134_at | B3GNT6 | -0,13090639 | 5,58149278 | -1,51479743 | 0,15127859 | 0,39744673 |
| 2870_at | GRK6 | -0,15099244 | 7,47364806 | -1,51459307 | 0,15132993 | 0,3975274 |
| 54431_at | DNAJC10 | -0,31376011 | 7,35522542 | -1,51443714 | 0,15136911 | 0,39757611 |
| 79783_at | SUGCT | 0,18478076 | 4,05208233 | 1,51409581 | 0,15145491 | 0,39773194 |
| 409_at | ARRB2 | 0,15569303 | 6,06611699 | 1,51401774 | 0,15147454 | 0,39773194 |
| 10077_at | TSPAN32 | -0,14239036 | 5,3054865 | -1,51395475 | 0,15149038 | 0,39773194 |
| 1476_at | CSTB | -0,07954627 | 12,511388 | -1,5137865 | 0,1515327 | 0,39778883 |
| 340544_at | MORF4L2-AS1 | -0,23868903 | 4,46021451 | -1,51343625 | 0,15162083 | 0,39796594 |
| 3727_at | JUND | 0,24240831 | 7,23064566 | 1,51333511 | 0,15164628 | 0,39797852 |
| 29881_at | NPC1L1 | -0,15773431 | 4,69599632 | -1,51230345 | 0,15190614 | 0,39851338 |
| 51246_at | SHISA5 | 0,28799346 | 10,2946101 | 1,51225607 | 0,15191809 | 0,39851338 |
| 200576_at | PIKFYVE | 0,17870116 | 6,43847051 | 1,51222519 | 0,15192587 | 0,39851338 |
| 129285_at | PPP1R21 | 0,17066123 | 6,96952446 | 1,51219755 | 0,15193284 | 0,39851338 |

| | | | | | | |
|---|---|---|---|---|---|---|
| 100507086_a | DLG1-AS1 | 0,2055588 | 4,77052699 | 1,5120846 | 0,15196132 | 0,39853381 |
| 83746_at | L3MBTL2 | -0,08691262 | 7,45034713 | -1,51173646 | 0,15204912 | 0,3987098 |
| 28965_at | SLC27A6 | 0,30582438 | 3,68923465 | 1,51150697 | 0,15210702 | 0,39875514 |
| 101928535_a | LOC10192853 | -0,10349644 | 3,24464481 | -1,51150383 | 0,15210781 | 0,39875514 |
| 151473_at | SLC16A14 | -0,18879479 | 4,37775239 | -1,51105724 | 0,15222054 | 0,39894415 |
| 51382_at | ATP6V1D | 0,23295075 | 9,5372576 | 1,51100107 | 0,15223473 | 0,39894415 |
| 342184_at | FMN1 | -0,13832779 | 5,30232868 | -1,51097211 | 0,15224204 | 0,39894415 |
| 55432_at | YOD1 | 0,22175118 | 5,85335153 | 1,51083292 | 0,15227719 | 0,39894942 |
| 1811_at | SLC26A3 | -0,125086 | 2,89486659 | -1,51080014 | 0,15228547 | 0,39894942 |
| 3066_at | HDAC2 | -0,16664178 | 7,41568119 | -1,51054753 | 0,15234929 | 0,39906234 |
| 6670_at | SP3 | -0,10950566 | 7,78848659 | -1,5103601 | 0,15239666 | 0,39913214 |
| 57192_at | MCOLN1 | 0,16394596 | 7,75450452 | 1,51001034 | 0,15248508 | 0,39930943 |
| 3283_at | HSD3B1 | -0,15682367 | 3,82306286 | -1,50986204 | 0,15252259 | 0,39933711 |
| 359948_at | IRF2BP2 | -0,14116021 | 8,98112912 | -1,50980461 | 0,15253712 | 0,39933711 |
| 100996492_a | LINC01314 | -0,17427758 | 4,79626222 | -1,50970048 | 0,15256346 | 0,39935179 |
| 840_at | CASP7 | 0,29047733 | 8,17692376 | 1,50958277 | 0,15259324 | 0,39937548 |
| 93082_at | NEURL3 | 0,2841361 | 6,12577355 | 1,50930964 | 0,15266236 | 0,39950211 |
| 79847_at | TMEM180 | -0,2747052 | 6,42664952 | -1,50885866 | 0,15277655 | 0,39974662 |
| 1964_at | EIF1AX | -0,50601921 | 8,71670322 | -1,50876346 | 0,15280066 | 0,39975542 |
| 619208_at | FAM229B | 0,19123607 | 6,2036604 | 1,50848258 | 0,15287183 | 0,39988729 |
| 93134_at | ZNF561 | 0,3989501 | 7,50069399 | 1,50806075 | 0,15297876 | 0,40006455 |
| 604_at | BCL6 | 0,17466819 | 7,69706115 | 1,5080514 | 0,15298113 | 0,40006455 |
| 51274_at | KLF3 | 0,12876533 | 8,72253535 | 1,50766575 | 0,15307894 | 0,40021567 |
| 25921_at | ZDHHC5 | -0,08554645 | 9,12927828 | -1,50765973 | 0,15308047 | 0,40021567 |
| 647323_at | LOC647323 | -0,17653142 | 3,60596054 | -1,506708 | 0,15332209 | 0,40070349 |
| 727800_at | RNF208 | -0,1337815 | 5,66969671 | -1,50664809 | 0,15333731 | 0,40070349 |
| 5900_at | RALGDS | 0,24962089 | 7,88233115 | 1,5066157 | 0,15334554 | 0,40070349 |
| 29780_at | PARVB | -0,106389 | 5,00056229 | -1,50659709 | 0,15335026 | 0,40070349 |
| 6332_at | SCN7A | 0,09161423 | 2,92197184 | 1,50623973 | 0,15344109 | 0,40086356 |
| 284788_at | LOC284788 | -0,09379903 | 4,03798551 | -1,5061923 | 0,15345314 | 0,40086356 |
| 3587_at | IL10RA | -0,16106278 | 4,75240108 | -1,50594556 | 0,15351588 | 0,40097307 |
| 23005_at | MAPKBP1 | 0,09651736 | 6,38660207 | 1,50565221 | 0,1535905 | 0,40108059 |
| 256051_at | ZNF549 | 0,25791307 | 4,8418487 | 1,50555657 | 0,15361483 | 0,40108059 |
| 100506668_a | NRAV | 0,15659557 | 7,71125758 | 1,50544371 | 0,15364355 | 0,40108059 |
| 137682_at | NDUFAF6 | -0,19497155 | 6,56369023 | -1,50538041 | 0,15365966 | 0,40108059 |
| 100287569_a | LINC00173 | -0,13674938 | 3,97774209 | -1,50537456 | 0,15366115 | 0,40108059 |
| 3508_at | IGHMBP2 | -0,10253527 | 6,34646741 | -1,50509286 | 0,15373286 | 0,40116301 |
| 404281_at | YY2 | -0,20644112 | 6,15864145 | -1,50500888 | 0,15375424 | 0,40116301 |
| 116039_at | OSR2 | 0,81154529 | 7,51178785 | 1,50500511 | 0,1537552 | 0,40116301 |
| 131920_at | TMEM207 | -0,12499524 | 3,81464749 | -1,50479001 | 0,15380999 | 0,40125159 |
| 116519_at | APOA5 | -0,18215016 | 5,28036726 | -1,50463942 | 0,15384835 | 0,40127135 |
| 5744_at | PTHLH | -0,42560838 | 7,35800909 | -1,50459674 | 0,15385922 | 0,40127135 |
| 100128993_a | LOC10012899 | 0,12840337 | 3,67883466 | 1,5042284 | 0,1539531 | 0,40140305 |
| 26011_at | TENM4 | -0,20019655 | 4,15246892 | -1,504228 | 0,1539532 | 0,40140305 |
| 91746_at | YTHDC1 | 0,25880408 | 7,39257737 | 1,50415334 | 0,15397223 | 0,40140305 |
| 9043_at | SPAG9 | -0,091357 | 8,03275815 | -1,50387105 | 0,15404421 | 0,40153636 |
| 57176_at | VARS2 | -0,18763016 | 7,84264684 | -1,50320458 | 0,15421427 | 0,40192526 |
| 4883_at | NPR3 | -0,1245571 | 3,93567908 | -1,50304657 | 0,15425462 | 0,40197601 |
| 101929623_a | LINC01215 | 0,11215438 | 2,79756684 | 1,50235104 | 0,1544323 | 0,4023846 |
| 8932_at | MBD2 | -0,0930773 | 5,99463572 | -1,50188516 | 0,15455141 | 0,40264049 |

| | | | | | | |
|---|---|---|---|---|---|---|
| 51163_at | DBR1 | -0,12419036 | 7,34028103 | -1,50168116 | 0,15460359 | 0,40272197 |
| 25900_at | IFFO1 | -0,14073933 | 5,66451304 | -1,50149574 | 0,15465103 | 0,40279108 |
| 60436_at | TGIF2 | 0,35740069 | 7,87145029 | 1,50104617 | 0,15476611 | 0,4030363 |
| 10879_at | SMR3B | -0,16289393 | 3,13080006 | -1,50079776 | 0,15482972 | 0,40314747 |
| 667_at | DST | 0,18340525 | 7,34185243 | 1,50067591 | 0,15486094 | 0,40315399 |
| 51380_at | CSAD | 0,34770532 | 5,41491412 | 1,50062458 | 0,15487409 | 0,40315399 |
| 100507362_a | LINC01015 | -0,16405146 | 4,51536106 | -1,50015676 | 0,15499398 | 0,40333846 |
| 285973_at | ATG9B | 0,82978827 | 5,77138687 | 1,50014783 | 0,15499627 | 0,40333846 |
| 100379224_a | LOC10037922 | 0,1121153 | 3,39921525 | 1,50010299 | 0,15500777 | 0,40333846 |
| 9559_at | VPS26A | -0,12191202 | 10,0651617 | -1,49988437 | 0,15506382 | 0,40341844 |
| 57690_at | TNRC6C | 0,15316297 | 5,73086247 | 1,49979355 | 0,15508712 | 0,40341844 |
| 23452_at | ANGPTL2 | -0,12457142 | 4,35496991 | -1,49973813 | 0,15510133 | 0,40341844 |
| 84002_at | B3GNT5 | 0,33371851 | 7,35496025 | 1,49934761 | 0,15520153 | 0,40362456 |
| 55509_at | BATF3 | -0,19970179 | 5,67666209 | -1,4991658 | 0,1552482 | 0,40369143 |
| 1299_at | COL9A3 | 0,25268322 | 6,35188428 | 1,49904154 | 0,15528011 | 0,40371423 |
| 539_at | ATP5O | -0,1815583 | 11,4379652 | -1,49895854 | 0,15530142 | 0,40371423 |
| 100874014_a | NAV2-AS2 | -0,16978411 | 4,77142258 | -1,49888679 | 0,15531984 | 0,40371423 |
| 201266_at | SLC39A11 | 0,30974842 | 7,89061728 | 1,49878815 | 0,15534518 | 0,4037256 |
| 54845_at | ESRP1 | -0,23785377 | 10,3451563 | -1,49859801 | 0,15539402 | 0,40373446 |
| 64221_at | ROBO3 | 0,30879604 | 5,44690849 | 1,49855451 | 0,1554052 | 0,40373446 |
| 779_at | CACNA1S | -0,18128184 | 4,86743392 | -1,49853013 | 0,15541146 | 0,40373446 |
| 8111_at | GPR68 | -0,13347989 | 5,51069418 | -1,49840813 | 0,15544281 | 0,40376145 |
| 164395_at | TTLL9 | -0,11261021 | 4,16967007 | -1,4983058 | 0,15546911 | 0,40377532 |
| 80352_at | RNF39 | 0,22570871 | 6,22607663 | 1,49799446 | 0,15554915 | 0,40392873 |
| 79054_at | TRPM8 | -0,10736479 | 3,76161095 | -1,49752144 | 0,15567082 | 0,40419019 |
| 3906_at | LALBA | -0,17532982 | 4,17629893 | -1,49708143 | 0,15578407 | 0,40442973 |
| 57126_at | CD177 | -0,21570208 | 6,07992237 | -1,49647654 | 0,15593988 | 0,40476094 |
| 91298_at | C12orf29 | -0,13595062 | 8,27990016 | -1,49642298 | 0,15595368 | 0,40476094 |
| 284273_at | ZADH2 | 0,10864754 | 6,82302781 | 1,49600561 | 0,15606126 | 0,40496943 |
| 4642_at | MYO1D | 0,08734987 | 6,00453287 | 1,49594825 | 0,15607605 | 0,40496943 |
| 10773_at | ZBTB6 | 0,15541257 | 6,60393561 | 1,49566685 | 0,15614863 | 0,40510318 |
| 56990_at | CDC42SE2 | 0,14516602 | 9,3608807 | 1,49543519 | 0,1562084 | 0,40520367 |
| 157807_at | CLVS1 | -0,20981699 | 4,02541605 | -1,49522049 | 0,15626381 | 0,40521952 |
| 56829_at | ZC3HAV1 | 0,17943314 | 8,81436023 | 1,49515954 | 0,15627955 | 0,40521952 |
| 388228_at | SBK1 | 0,53833152 | 7,10184567 | 1,49514415 | 0,15628352 | 0,40521952 |
| 158297_at | SAXO1 | -0,17687086 | 4,60958606 | -1,49508552 | 0,15629866 | 0,40521952 |
| 9754_at | STARD8 | -0,16712847 | 5,44781623 | -1,49493893 | 0,15633651 | 0,4052631 |
| 27125_at | AFF4 | 0,2040933 | 6,72802681 | 1,49482887 | 0,15636493 | 0,40528224 |
| 3265_at | HRAS | -0,39414572 | 8,76081779 | -1,49460208 | 0,15642351 | 0,40537944 |
| 5619_at | PRM1 | -0,12893031 | 4,73914906 | -1,49452077 | 0,15644452 | 0,40537944 |
| 80737_at | VWA7 | -0,17291039 | 6,38752196 | -1,49421243 | 0,1565242 | 0,40551287 |
| 7172_at | TPMT | 0,19068595 | 6,36960154 | 1,49411018 | 0,15655064 | 0,40551287 |
| 5616_at | PRKY | 0,30315483 | 6,12075912 | 1,49407718 | 0,15655917 | 0,40551287 |
| 22941_at | SHANK2 | 0,18807115 | 5,24203897 | 1,49344171 | 0,15672353 | 0,40588402 |
| 8310_at | ACOX3 | 0,45665362 | 7,82002265 | 1,49328553 | 0,15676395 | 0,40593412 |
| 55967_at | NDUFA12 | -0,2204027 | 10,7282729 | -1,49296601 | 0,15684666 | 0,40605138 |
| 7757_at | ZNF208 | -0,13509185 | 2,88587913 | -1,49294776 | 0,15685139 | 0,40605138 |
| 56169_at | GSDMC | -0,61400257 | 6,85288463 | -1,49246184 | 0,15697726 | 0,40631339 |
| 84899_at | TMTC4 | 0,33641755 | 6,23896561 | 1,49239418 | 0,15699479 | 0,40631339 |
| 100130691_a | LOC10013069 | 0,1253203 | 3,68787393 | 1,49211128 | 0,15706811 | 0,406446 |

| | | | | | | |
|---|---|---|---|---|---|---|
| 435_at | ASL | -0,23032285 | 8,29908076 | -1,49203129 | 0,15708885 | 0,406446 |
| 4306_at | NR3C2 | 0,13699508 | 3,8802477 | 1,49195232 | 0,15710933 | 0,406446 |
| 84839_at | RAX2 | -0,18328996 | 5,60669637 | -1,49157709 | 0,15720665 | 0,40662447 |
| 85478_at | CCDC65 | -0,11256959 | 3,29995062 | -1,49137095 | 0,15726013 | 0,40662447 |
| 90203_at | SNX21 | 0,21058938 | 6,0410794 | 1,491369 | 0,15726064 | 0,40662447 |
| 26092_at | TOR1AIP1 | 0,25482395 | 9,09728894 | 1,49136088 | 0,15726275 | 0,40662447 |
| 128674_at | PROKR2 | 0,16521798 | 5,02635185 | 1,49101788 | 0,15735178 | 0,40677887 |
| 2251_at | FGF6 | -0,11403416 | 5,57396351 | -1,49093099 | 0,15737434 | 0,40677887 |
| 2050_at | EPHB4 | -0,14841129 | 7,56241527 | -1,49088681 | 0,15738581 | 0,40677887 |
| 54856_at | GON4L | 0,13751491 | 6,43132322 | 1,49069311 | 0,15743612 | 0,40680644 |
| 259307_at | IL4I1 | 0,32219033 | 5,38341373 | 1,49068312 | 0,15743872 | 0,40680644 |
| 219699_at | UNC5B | 0,20073793 | 7,78897395 | 1,49057163 | 0,15746768 | 0,4068267 |
| 3242_at | HPD | -0,15531557 | 3,90687951 | -1,49036613 | 0,15752107 | 0,40683835 |
| 9241_at | NOG | -0,27052588 | 3,7080319 | -1,4903048 | 0,15753701 | 0,40683835 |
| 11009_at | IL24 | -0,1447603 | 3,80102579 | -1,49026775 | 0,15754664 | 0,40683835 |
| 9739_at | SETD1A | -0,10909656 | 6,68211463 | -1,49022916 | 0,15755667 | 0,40683835 |
| 53615_at | MBD3 | -0,19125473 | 8,77105363 | -1,4898613 | 0,15765231 | 0,40703074 |
| 26476_at | OR10J1 | -0,18237114 | 3,88487146 | -1,4894747 | 0,15775287 | 0,40720619 |
| 80772_at | CPTP | -0,09467727 | 7,72873111 | -1,48940672 | 0,15777056 | 0,40720619 |
| 4629_at | MYH11 | -0,11282284 | 4,73473694 | -1,48928585 | 0,15780201 | 0,40720619 |
| 4849_at | CNOT3 | 0,0919567 | 6,89512797 | 1,48927506 | 0,15780482 | 0,40720619 |
| 5538_at | PPT1 | 0,13920913 | 10,9377762 | 1,48889134 | 0,15790471 | 0,40740938 |
| 1612_at | DAPK1 | 0,39745406 | 8,30838755 | 1,48870043 | 0,15795443 | 0,40748308 |
| 3120_at | HLA-DQB2 | -0,10715049 | 5,23426558 | -1,48856113 | 0,15799071 | 0,40752211 |
| 23552_at | CDK20 | 0,15664933 | 6,57441417 | 1,48829799 | 0,15805928 | 0,40761844 |
| 8841_at | HDAC3 | -0,09674748 | 9,31197647 | -1,48822488 | 0,15807833 | 0,40761844 |
| 9363_at | RAB33A | 0,09639209 | 4,05216962 | 1,48817421 | 0,15809154 | 0,40761844 |
| 60481_at | ELOVL5 | -0,38501426 | 8,39855021 | -1,48779608 | 0,15819013 | 0,40781804 |
| 5335_at | PLCG1 | 0,13055292 | 8,10383258 | 1,48763149 | 0,15823306 | 0,40781859 |
| 440574_at | MINOS1 | -0,20877534 | 6,40676926 | -1,48756912 | 0,15824933 | 0,40781859 |
| 85416_at | ZIC5 | -0,20794465 | 4,81203573 | -1,48755176 | 0,15825385 | 0,40781859 |
| 27240_at | SIT1 | -0,16452932 | 4,83690413 | -1,487208 | 0,15834356 | 0,40796494 |
| 60482_at | SLC5A7 | 0,18043824 | 3,86058079 | 1,4871718 | 0,158353 | 0,40796494 |
| 6733_at | SRPK2 | 0,25917282 | 7,91908577 | 1,48692697 | 0,15841692 | 0,4080681 |
| 100506025_a | ISPD-AS1 | -0,1065889 | 2,92211816 | -1,48685614 | 0,15843541 | 0,4080681 |
| 2661_at | GDF9 | -0,11738678 | 4,59948208 | -1,48671258 | 0,1584729 | 0,4081101 |
| 81542_at | TMX1 | -0,17553191 | 10,5591434 | -1,48657016 | 0,15851011 | 0,40811789 |
| 169026_at | SLC30A8 | -0,15806053 | 3,93908295 | -1,48649136 | 0,15853069 | 0,40811789 |
| 92482_at | BBIP1 | 0,33598526 | 8,56901024 | 1,48645769 | 0,15853949 | 0,40811789 |
| 147699_at | PPM1N | 0,16327503 | 6,56515317 | 1,48634302 | 0,15856945 | 0,40814048 |
| 163154_at | PRR22 | -0,17109738 | 5,78163391 | -1,48608971 | 0,15863566 | 0,40825633 |
| 94122_at | SYTL5 | 0,65383694 | 5,45530098 | 1,48590787 | 0,1586832 | 0,40827082 |
| 200558_at | APLF | 0,18733464 | 3,74058036 | 1,48590604 | 0,15868367 | 0,40827082 |
| 152065_at | C3orf22 | -0,16357487 | 4,78654341 | -1,48553107 | 0,15878174 | 0,40846858 |
| 152573_at | SHISA3 | -0,13740927 | 3,61018397 | -1,48534299 | 0,15883095 | 0,40854061 |
| 26253_at | CLEC4E | -0,10573604 | 3,64529333 | -1,48516754 | 0,15887687 | 0,40860415 |
| 91624_at | NEXN | 0,95960542 | 4,56656079 | 1,48493419 | 0,15893795 | 0,40870669 |
| 732275_at | LINC00917 | -0,15568321 | 4,30906585 | -1,48484333 | 0,15896175 | 0,40871331 |
| 80127_at | CCDC176 | 0,53195138 | 4,50070543 | 1,48473782 | 0,15898937 | 0,40872979 |
| 102723701_a | LOC10272370 | -0,1861849 | 5,29131892 | -1,48462338 | 0,15901935 | 0,4087523 |

| | | | | | | |
|---|---|---|---|---|---|---|
| 222894_at | FERD3L | -0,1540677 | 5,36879916 | -1,48436777 | 0,15908631 | 0,40883887 |
| 5771_at | PTPN2 | -0,15210433 | 6,21351208 | -1,4843328 | 0,15909548 | 0,40883887 |
| 10047_at | CST8 | -0,15443238 | 3,88493816 | -1,48377263 | 0,15924232 | 0,40911279 |
| 78992_at | YIPF2 | 0,17209834 | 8,06343475 | 1,48376413 | 0,15924455 | 0,40911279 |
| 9048_at | ARTN | -0,13313962 | 5,64019621 | -1,48343806 | 0,15933008 | 0,4092346 |
| 6444_at | SGCD | -0,09759464 | 3,98003258 | -1,48339286 | 0,15934194 | 0,4092346 |
| 83933_at | HDAC10 | -0,16444026 | 6,9434507 | -1,48327877 | 0,15937187 | 0,4092346 |
| 3171_at | FOXA3 | -0,09342759 | 4,19923312 | -1,48325947 | 0,15937694 | 0,4092346 |
| 23288_at | IQCE | 0,24270414 | 6,7116115 | 1,48299205 | 0,15944713 | 0,40936026 |
| 9717_at | SEC14L5 | -0,17046313 | 4,98285265 | -1,48281522 | 0,15949356 | 0,40940009 |
| 27133_at | KCNH5 | -0,10094979 | 3,17638919 | -1,48276805 | 0,15950594 | 0,40940009 |
| 10164_at | CHST4 | -0,17489295 | 4,17717738 | -1,48269014 | 0,1595264 | 0,40940009 |
| 100506713_a | ALOX12-AS1 | 0,1243157 | 4,63006155 | 1,48258774 | 0,1595533 | 0,40941457 |
| 1463_at | NCAN | -0,14412143 | 4,82006685 | -1,48242117 | 0,15959706 | 0,40944036 |
| 8493_at | PPM1D | 0,16168037 | 6,79660158 | 1,48238766 | 0,15960586 | 0,40944036 |
| 146849_at | CCDC42 | -0,17579131 | 4,68613225 | -1,48203513 | 0,15969851 | 0,40962348 |
| 83479_at | DDX59 | 0,11459803 | 6,44691135 | 1,48177989 | 0,15976561 | 0,40974105 |
| 5264_at | PHYH | 0,32463634 | 7,90129443 | 1,48158535 | 0,15981678 | 0,40978247 |
| 1440_at | CSF3 | -0,1608258 | 6,44324847 | -1,4815567 | 0,15982431 | 0,40978247 |
| 8915_at | BCL10 | -0,12895646 | 8,81891689 | -1,48142247 | 0,15985962 | 0,40981846 |
| 347902_at | AMIGO2 | -0,1675847 | 11,3499951 | -1,48097487 | 0,15997742 | 0,41006587 |
| 10389_at | SCML2 | -0,19973646 | 4,84979459 | -1,48077247 | 0,16003071 | 0,41009989 |
| 2120_at | ETV6 | 0,42091206 | 8,03912635 | 1,48075545 | 0,16003519 | 0,41009989 |
| 84174_at | SLA2 | -0,17688856 | 4,94690826 | -1,4806819 | 0,16005456 | 0,41009989 |
| 351_at | APP | 0,13387695 | 10,8785765 | 1,48059458 | 0,16007756 | 0,41010427 |
| 143686_at | SESN3 | 0,3369014 | 5,97913324 | 1,48016191 | 0,16019155 | 0,41034174 |
| 283248_at | RCOR2 | -0,13780208 | 6,01691278 | -1,48001908 | 0,1602292 | 0,41038359 |
| 7293_at | TNFRSF4 | -0,16265834 | 5,93067384 | -1,47980156 | 0,16028654 | 0,4104759 |
| 8703_at | B4GALT3 | -0,09633202 | 6,7746147 | -1,47945206 | 0,16037872 | 0,41062129 |
| 55315_at | SLC29A3 | 0,58619052 | 8,80829328 | 1,47939153 | 0,16039469 | 0,41062129 |
| 29904_at | EEF2K | -0,26742912 | 7,43797058 | -1,47934385 | 0,16040727 | 0,41062129 |
| 90120_at | C9orf69 | 0,14546325 | 8,93317653 | 1,47919391 | 0,16044684 | 0,41063343 |
| 90113_at | VWA5B2 | -0,19818318 | 5,43514651 | -1,47913134 | 0,16046335 | 0,41063343 |
| 27120_at | DKKL1 | -0,12979998 | 5,50757094 | -1,47908354 | 0,16047596 | 0,41063343 |
| 84138_at | SLC7A6OS | -0,13869042 | 7,28212079 | -1,47893634 | 0,16051482 | 0,4106783 |
| 5820_at | PVT1 | -0,17697396 | 8,16731623 | -1,47872799 | 0,16056983 | 0,41068622 |
| 7409_at | VAV1 | -0,15256604 | 3,94199562 | -1,47868822 | 0,16058033 | 0,41068622 |
| 57643_at | ZSWIM5 | 0,18622192 | 5,55565018 | 1,47868239 | 0,16058187 | 0,41068622 |
| 50840_at | TAS2R14 | -0,16158073 | 4,10760579 | -1,47844476 | 0,16064463 | 0,4106872 |
| 3795_at | KHK | -0,15739556 | 6,02017154 | -1,47842195 | 0,16065066 | 0,4106872 |
| 158427_at | TSTD2 | 0,22362662 | 7,20012961 | 1,4784002 | 0,16065641 | 0,4106872 |
| 55929_at | DMAP1 | 0,16407284 | 8,06553106 | 1,47835808 | 0,16066753 | 0,4106872 |
| 7189_at | TRAF6 | 0,22875844 | 6,81495454 | 1,47815029 | 0,16072244 | 0,41077303 |
| 219749_at | ZNF25 | 0,24206218 | 4,66057121 | 1,4780542 | 0,16074784 | 0,41078344 |
| 7042_at | TGFB2 | 0,29706447 | 4,52187832 | 1,47795256 | 0,1607747 | 0,4107976 |
| 9075_at | CLDN2 | -0,10457772 | 5,00923382 | -1,47785959 | 0,16079928 | 0,41080591 |
| 101927257_a | LOC10192725 | -0,14291426 | 4,08059735 | -1,47765459 | 0,16085348 | 0,41085219 |
| 1010_at | CDH12 | 0,09782638 | 3,03595117 | 1,47762975 | 0,16086005 | 0,41085219 |
| 152641_at | WWC2-AS2 | 0,13965888 | 4,96701182 | 1,47747137 | 0,16090195 | 0,4109047 |
| 6660_at | SOX5 | 0,13752193 | 3,74664972 | 1,4771138 | 0,16099655 | 0,41100491 |

| | | | | | | |
|---|---|---|---|---|---|---|
| 5228_at | PGF | -0,38410925 | 5,93515593 | -1,47693622 | 0,16104356 | 0,41100491 |
| 3798_at | KIF5A | -0,21594627 | 4,82046035 | -1,47687818 | 0,16105892 | 0,41100491 |
| 64754_at | SMYD3 | -0,29849726 | 7,70697752 | -1,47687672 | 0,16105931 | 0,41100491 |
| 9421_at | HAND1 | -0,20668482 | 5,31709219 | -1,4768474 | 0,16106707 | 0,41100491 |
| 388125_at | C2CD4B | -0,18091543 | 5,56720064 | -1,47675441 | 0,16109169 | 0,41100491 |
| 59344_at | ALOXE3 | -0,17258273 | 4,30122014 | -1,47675305 | 0,16109205 | 0,41100491 |
| 4632_at | MYL1 | -0,18792361 | 3,71100172 | -1,47667819 | 0,16111188 | 0,41100491 |
| 164832_at | LONRF2 | -0,11372279 | 3,42300285 | -1,47647196 | 0,16116649 | 0,41104794 |
| 23191_at | CYFIP1 | 0,14244862 | 10,2261376 | 1,47645336 | 0,16117142 | 0,41104794 |
| 414777_at | HCG18 | -0,23636666 | 6,44124666 | -1,4763268 | 0,16120495 | 0,41107902 |
| 6827_at | SUPT4H1 | 0,14334908 | 8,85754521 | 1,4757294 | 0,16136329 | 0,41138073 |
| 103581031_a | CASC23 | -0,14282126 | 4,07407124 | -1,47567415 | 0,16137794 | 0,41138073 |
| 4990_at | SIX6 | -0,12752716 | 4,74229062 | -1,47563872 | 0,16138733 | 0,41138073 |
| 23376_at | UFL1 | 0,39833266 | 8,70392545 | 1,47538985 | 0,16145334 | 0,41141814 |
| 4057_at | LTF | -0,19854539 | 4,46404628 | -1,47535135 | 0,16146355 | 0,41141814 |
| 100506930_a | LINC00665 | -0,09660204 | 4,04969522 | -1,47534183 | 0,16146608 | 0,41141814 |
| 119395_at | CALHM3 | -0,37548458 | 4,77897106 | -1,47485218 | 0,16159603 | 0,41169478 |
| 128414_at | NKAIN4 | -0,12866262 | 4,83863713 | -1,47435111 | 0,16172909 | 0,41197931 |
| 5194_at | PEX13 | 0,21300449 | 8,21826777 | 1,47400824 | 0,1618202 | 0,41212856 |
| 64766_at | S100PBP | 0,12811853 | 7,51209778 | 1,47396091 | 0,16183278 | 0,41212856 |
| 9609_at | RAB36 | -0,2591273 | 5,28865713 | -1,47388909 | 0,16185187 | 0,41212856 |
| 90332_at | EXOC3L2 | -0,2013979 | 6,99372865 | -1,47364646 | 0,16191638 | 0,41223833 |
| 314_at | AOC2 | 0,11711669 | 5,18445265 | 1,47339825 | 0,16198239 | 0,41234059 |
| 2921_at | CXCL3 | 0,54660442 | 4,62694877 | 1,47333449 | 0,16199935 | 0,41234059 |
| 3817_at | KLK2 | -0,10941283 | 4,62604194 | -1,47314962 | 0,16204854 | 0,41241129 |
| 4683_at | NBN | -0,16417395 | 8,68168794 | -1,47302403 | 0,16208196 | 0,41244186 |
| 1902_at | LPAR1 | -0,24714453 | 6,38720534 | -1,47251982 | 0,1622162 | 0,41272892 |
| 23660_at | ZKSCAN5 | 0,17202531 | 6,85115396 | 1,47207403 | 0,16233496 | 0,41292726 |
| 100506476_a | LOC10050647 | 0,1264369 | 4,38791929 | 1,47206629 | 0,16233702 | 0,41292726 |
| 3671_at | ISLR | -0,1553737 | 4,35867129 | -1,47183661 | 0,16239824 | 0,41300525 |
| 7581_at | ZNF33A | 0,29417844 | 6,94362118 | 1,47172308 | 0,16242851 | 0,41300525 |
| 199857_at | ALG14 | -0,09505012 | 8,0602425 | -1,47169382 | 0,16243631 | 0,41300525 |
| 8908_at | GYG2 | 0,1931189 | 5,1733926 | 1,47162954 | 0,16245345 | 0,41300525 |
| 54751_at | FBLIM1 | 0,26739553 | 7,27448341 | 1,47141393 | 0,16251095 | 0,41309691 |
| 7905_at | REEP5 | 0,12031619 | 10,569087 | 1,4712282 | 0,16256049 | 0,41316833 |
| 100506526_a | LOC10050652 | -0,17305924 | 3,71016953 | -1,47106716 | 0,16260346 | 0,41322302 |
| 135656_at | DPCR1 | -0,15402985 | 4,03831888 | -1,47030242 | 0,16280764 | 0,41368733 |
| 2352_at | FOLR3 | -0,23748104 | 4,82287005 | -1,47014864 | 0,16284873 | 0,41373715 |
| 3428_at | IFI16 | 0,59976592 | 8,91177807 | 1,46980911 | 0,16293947 | 0,4139131 |
| 150577_at | LINC01104 | -0,19752394 | 4,65007204 | -1,46963982 | 0,16298473 | 0,41397348 |
| 154089_at | LINC01312 | -0,13959009 | 3,10315927 | -1,46917297 | 0,16310959 | 0,414236 |
| 25891_at | PAMR1 | -0,16832085 | 4,59891043 | -1,46907169 | 0,16313669 | 0,41424644 |
| 8358_at | HIST1H3B | -0,28142408 | 6,78970217 | -1,46899687 | 0,16315671 | 0,41424644 |
| 374291_at | NDUFS7 | -0,13091094 | 8,94927715 | -1,468878 | 0,16318853 | 0,41427261 |
| 2931_at | GSK3A | -0,13424262 | 8,51821316 | -1,46865199 | 0,16324903 | 0,4143716 |
| 221527_at | ZBTB12 | 0,29903888 | 6,72456603 | 1,46851514 | 0,16328567 | 0,41441 |
| 30012_at | TLX3 | -0,11702141 | 4,15907724 | -1,4683209 | 0,16333769 | 0,41448742 |
| 222068_at | TMED4 | 0,23246825 | 9,43607783 | 1,46812305 | 0,1633907 | 0,4145512 |
| 83887_at | TTLL2 | -0,13315331 | 4,04548546 | -1,46806643 | 0,16340587 | 0,4145512 |
| 402381_at | SOHLH1 | -0,20535606 | 6,10080478 | -1,46791317 | 0,16344694 | 0,41457117 |

| | | | | | | |
|---|---|---|---|---|---|---|
| 4544_at | MTNR1B | -0,14886757 | 5,39730488 | -1,46778973 | 0,16348002 | 0,41457117 |
| 127294_at | MYOM3 | -0,12532723 | 4,62500235 | -1,46775728 | 0,16348872 | 0,41457117 |
| 79879_at | CCDC134 | -0,32989762 | 7,16110104 | -1,46771585 | 0,16349983 | 0,41457117 |
| 80209_at | PROSER1 | -0,12980935 | 8,99713896 | -1,46738818 | 0,16358769 | 0,41469287 |
| 55576_at | STAB2 | -0,15848099 | 4,21146422 | -1,46737629 | 0,16359088 | 0,41469287 |
| 9340_at | GLP2R | 0,15057504 | 4,31447268 | 1,46728928 | 0,16361422 | 0,41469746 |
| 10385_at | BTN2A2 | 0,35781486 | 5,78899907 | 1,4669899 | 0,16369454 | 0,41481877 |
| 23081_at | KDM4C | 0,18797301 | 7,01777872 | 1,46695036 | 0,16370515 | 0,41481877 |
| 3710_at | ITPR3 | -0,13425779 | 8,12731424 | -1,46655165 | 0,16381218 | 0,41503537 |
| 55314_at | TMEM144 | 0,48797418 | 6,45348979 | 1,46638105 | 0,16385799 | 0,41504564 |
| 162993_at | ZNF846 | 0,41628063 | 4,01145272 | 1,46636802 | 0,16386149 | 0,41504564 |
| 140258_at | KRTAP13-1 | -0,15325708 | 4,05610014 | -1,46626558 | 0,163889 | 0,41504564 |
| 80856_at | KIAA1715 | -0,1500939 | 8,3138489 | -1,46621566 | 0,16390241 | 0,41504564 |
| 8972_at | MGAM | 0,52447557 | 4,95862185 | 1,46600949 | 0,1639578 | 0,41513133 |
| 26993_at | AKAP8L | 0,07827328 | 7,23806717 | 1,46569828 | 0,16404145 | 0,41528852 |
| 54820_at | NDE1 | -0,13464318 | 7,02759746 | -1,46541323 | 0,16411809 | 0,41533234 |
| 441161_at | OOEP | -0,14095011 | 3,58694617 | -1,46531998 | 0,16414317 | 0,41533234 |
| 100505676_a | RRS1-AS1 | 0,08865408 | 3,42039356 | 1,46531909 | 0,16414341 | 0,41533234 |
| 83439_at | TCF7L1 | 0,34939777 | 6,22678336 | 1,46531317 | 0,164145 | 0,41533234 |
| 114614_at | MIR155HG | 0,10231835 | 3,08898884 | 1,46505019 | 0,16421574 | 0,41545677 |
| 3000_at | GUCY2D | -0,14013588 | 5,17994132 | -1,46466982 | 0,16431811 | 0,41561612 |
| 55282_at | LRRC36 | -0,13758829 | 4,9501932 | -1,46465582 | 0,16432188 | 0,41561612 |
| 1268_at | CNR1 | 0,14789702 | 4,36460749 | 1,46390667 | 0,16452365 | 0,41607183 |
| 56848_at | SPHK2 | -0,10117014 | 6,81038354 | -1,46353735 | 0,1646232 | 0,41625231 |
| 339976_at | TRIML1 | -0,10826381 | 3,65025781 | -1,46345165 | 0,16464631 | 0,41625231 |
| 9600_at | PITPNM1 | -0,12677263 | 8,22843813 | -1,46328688 | 0,16469075 | 0,41625231 |
| 101929526_a | LOC10192952 | -0,09513841 | 4,36118989 | -1,46327339 | 0,16469438 | 0,41625231 |
| 442184_at | OR2B3 | -0,13134681 | 3,13022342 | -1,46324121 | 0,16470306 | 0,41625231 |
| 28639_at | TRBC1 | -0,36677058 | 5,19243765 | -1,46293102 | 0,16478675 | 0,41630141 |
| 80054_at | CEBPA-AS1 | -0,18124258 | 5,45351939 | -1,4629045 | 0,1647939 | 0,41630141 |
| 64374_at | SIL1 | 0,11947042 | 7,34417729 | 1,46288033 | 0,16480042 | 0,41630141 |
| 2275_at | FHL3 | -0,27897882 | 7,35692775 | -1,46284878 | 0,16480894 | 0,41630141 |
| 11309_at | SLCO2B1 | -0,14559844 | 4,54736061 | -1,46264624 | 0,1648636 | 0,41633816 |
| 200845_at | KCTD6 | -0,25810192 | 5,48289415 | -1,46263473 | 0,16486671 | 0,41633816 |
| 10273_at | STUB1 | -0,09744905 | 9,41785612 | -1,4616251 | 0,16513944 | 0,41697223 |
| 7124_at | TNF | -0,17183392 | 4,37415386 | -1,46145253 | 0,1651861 | 0,41703536 |
| 79166_at | LILRP2 | -0,12020963 | 3,86908903 | -1,46040579 | 0,16546932 | 0,41769564 |
| 160313_at | KRT19P2 | -0,15231532 | 5,26057757 | -1,46006803 | 0,16556079 | 0,41775003 |
| 101926975_a | LOC10192697 | -0,12846387 | 3,53945205 | -1,46001316 | 0,16557566 | 0,41775003 |
| 25875_at | LETMD1 | 0,25608439 | 8,55971168 | 1,45996688 | 0,16558819 | 0,41775003 |
| 220930_at | ZEB1-AS1 | -0,25769609 | 5,25806952 | -1,45996281 | 0,1655893 | 0,41775003 |
| 56135_at | PCDHAC1 | -0,1733249 | 5,1742601 | -1,4599259 | 0,1655993 | 0,41775003 |
| 100131366_a | LINC00605 | -0,19124204 | 4,09748079 | -1,4596682 | 0,16566913 | 0,41787147 |
| 6168_at | RPL37A | -0,09007128 | 12,1648282 | -1,45953142 | 0,16570621 | 0,41791027 |
| 100506527_a | LINC01007 | -0,21609426 | 4,44666649 | -1,45929455 | 0,16577043 | 0,41801751 |
| 157627_at | LINC00599 | -0,14745004 | 4,91015994 | -1,45889788 | 0,16587803 | 0,41818981 |
| 84536_at | LINC01547 | -0,11523546 | 5,29216468 | -1,45888259 | 0,16588218 | 0,41818981 |
| 63914_at | LINC01590 | 0,11529668 | 3,29176667 | 1,45863358 | 0,16594975 | 0,41830542 |
| 8140_at | SLC7A5 | -0,51616027 | 10,8169446 | -1,45831889 | 0,16603518 | 0,41846601 |
| 113878_at | DTX2 | 0,09928971 | 6,85372257 | 1,45802281 | 0,16611559 | 0,41858051 |

| | | | | | | |
|---|---|---|---|---|---|---|
| 57102_at | C12orf4 | -0,17406061 | 7,36751133 | -1,4579916 | 0,16612407 | 0,41858051 |
| 138639_at | PTPDC1 | -0,14406211 | 5,9783987 | -1,45770035 | 0,16620321 | 0,41872121 |
| 7324_at | UBE2E1 | 0,10212515 | 10,219009 | 1,45760818 | 0,16622826 | 0,41872121 |
| 3446_at | IFNA10 | -0,12686675 | 3,33372893 | -1,45754613 | 0,16624512 | 0,41872121 |
| 112703_at | FAM71E1 | 0,20702533 | 6,3110738 | 1,45735331 | 0,16629754 | 0,41876394 |
| 11187_at | PKP3 | -0,21499008 | 8,22640368 | -1,45732379 | 0,16630557 | 0,41876394 |
| 11157_at | LSM6 | -0,47854893 | 8,29502789 | -1,4571265 | 0,16635922 | 0,41884429 |
| 57480_at | PLEKHG1 | 0,43505338 | 6,13850124 | 1,45698903 | 0,16639661 | 0,41888368 |
| 100129935_a | LOC10012993 | -0,09733828 | 5,24976293 | -1,45670499 | 0,16647389 | 0,41902347 |
| 10899_at | JTB | -0,0912845 | 11,0414683 | -1,45654215 | 0,16651821 | 0,41904729 |
| 84641_at | HIATL1 | -0,12806074 | 9,75411871 | -1,45651036 | 0,16652687 | 0,41904729 |
| 28988_at | DBNL | -0,08350384 | 9,17421255 | -1,45621282 | 0,16660787 | 0,41917927 |
| 79586_at | CHPF | 0,14806342 | 7,27928728 | 1,45607963 | 0,16664415 | 0,41917927 |
| 152940_at | C4orf45 | -0,11903307 | 3,15657636 | -1,45602947 | 0,16665781 | 0,41917927 |
| 222643_at | UNC5CL | 0,1766542 | 4,83044318 | 1,45599809 | 0,16666636 | 0,41917927 |
| 79025_at | C20orf195 | -0,15424228 | 5,85586297 | -1,45580029 | 0,16672024 | 0,41926005 |
| 360205_at | PRAC2 | -0,18459195 | 6,05976291 | -1,4554037 | 0,16682833 | 0,41947545 |
| 101928940_a | LOC10192894 | 0,13239671 | 2,91139135 | 1,45532624 | 0,16684945 | 0,41947545 |
| 80007_at | C10orf88 | -0,10744684 | 6,18479787 | -1,45509505 | 0,16691249 | 0,41957918 |
| 101929464_a | LOC10192946 | -0,13233042 | 5,1988086 | -1,45451998 | 0,16706939 | 0,41989708 |
| 10042_at | HMGXB4 | -0,13949909 | 8,55300606 | -1,45447176 | 0,16708255 | 0,41989708 |
| 284100_at | YWHAEP7 | -0,22730686 | 5,26254719 | -1,4542136 | 0,16715303 | 0,42001941 |
| 28595_at | TRBV7-3 | -0,23704025 | 4,57212834 | -1,4531455 | 0,16744489 | 0,42069791 |
| 9350_at | CER1 | -0,19321824 | 4,47930321 | -1,45233011 | 0,16766798 | 0,42117488 |
| 64427_at | TTC31 | 0,17221625 | 7,82571462 | 1,4522786 | 0,16768208 | 0,42117488 |
| 5567_at | PRKACB | 0,53206864 | 7,92008452 | 1,45221197 | 0,16770032 | 0,42117488 |
| 2646_at | GCKR | -0,15652747 | 4,43911666 | -1,45196071 | 0,16776913 | 0,42129276 |
| 90_at | ACVR1 | 0,19121133 | 8,59573016 | 1,45181832 | 0,16780813 | 0,42133577 |
| 157724_at | SLC7A13 | -0,14742505 | 3,49906399 | -1,45170981 | 0,16783786 | 0,42135549 |
| 64333_at | ARHGAP9 | -0,11936598 | 4,38019745 | -1,45144218 | 0,1679112 | 0,42148468 |
| 255043_at | TMEM86B | -0,21378942 | 5,92522102 | -1,45114223 | 0,16799343 | 0,42156959 |
| 5144_at | PDE4D | -0,17409446 | 4,91386378 | -1,45111651 | 0,16800048 | 0,42156959 |
| 440836_at | ODF3B | 0,1112315 | 6,06334045 | 1,45106783 | 0,16801383 | 0,42156959 |
| 4666_at | NACA | -0,15829619 | 7,24151111 | -1,4509995 | 0,16803257 | 0,42156959 |
| 2563_at | GABRD | -0,13067644 | 5,25365114 | -1,45088781 | 0,1680632 | 0,42159154 |
| 10423_at | CDIPT | 0,16860914 | 10,720686 | 1,45051343 | 0,16816591 | 0,42176423 |
| 55146_at | ZDHHC4 | 0,12435985 | 9,22687069 | 1,45043457 | 0,16818756 | 0,42176423 |
| 84313_at | VPS25 | 0,12872278 | 9,489756 | 1,45036683 | 0,16820615 | 0,42176423 |
| 22893_at | BAHD1 | 0,13014308 | 7,30950561 | 1,45031775 | 0,16821962 | 0,42176423 |
| 257144_at | GCSAM | 0,1080712 | 3,71685222 | 1,44983387 | 0,16835249 | 0,42204242 |
| 651_at | BMP3 | 0,45533195 | 6,90814403 | 1,44959924 | 0,16841695 | 0,42214907 |
| 100507027_a | MRLN | -0,11843776 | 3,25220401 | -1,44947086 | 0,16845222 | 0,42217582 |
| 50944_at | SHANK1 | -0,09207236 | 4,69718362 | -1,4494009 | 0,16847145 | 0,42217582 |
| 9329_at | GTF3C4 | -0,22500269 | 7,55169659 | -1,44898334 | 0,16858625 | 0,4223716 |
| 4704_at | NDUFA9 | -0,18768595 | 9,83093645 | -1,44895722 | 0,16859343 | 0,4223716 |
| 170487_at | ACTL10 | 0,302709 | 6,93103671 | 1,44886964 | 0,16861751 | 0,42237701 |
| 64320_at | RNF25 | -0,13670623 | 7,49675204 | -1,4485092 | 0,16871667 | 0,42257044 |
| 10049_at | DNAJB6 | 0,24466093 | 8,79984187 | 1,44804019 | 0,16884577 | 0,42273697 |
| 6885_at | MAP3K7 | -0,17514514 | 7,74530121 | -1,44792252 | 0,16887818 | 0,42273697 |
| 218_at | ALDH3A1 | -0,25862181 | 9,83453601 | -1,44791462 | 0,16888035 | 0,42273697 |

| | | | | | | |
|---|---|---|---|---|---|---|
| 8812_at | CCNK | -0,11155297 | 8,92361418 | -1,44782599 | 0,16890476 | 0,42273697 |
| 84898_at | PLXDC2 | -0,08139183 | 3,88756858 | -1,44774681 | 0,16892657 | 0,42273697 |
| 51533_at | PHF7 | -0,21905281 | 5,53780931 | -1,44767858 | 0,16894537 | 0,42273697 |
| 54676_at | GTPBP2 | 0,1697726 | 7,51009281 | 1,44764568 | 0,16895443 | 0,42273697 |
| 340273_at | ABCB5 | -0,08889762 | 3,06681577 | -1,44763008 | 0,16895873 | 0,42273697 |
| 84327_at | ZBED3 | 0,17415529 | 6,36563733 | 1,44753878 | 0,16898388 | 0,42274376 |
| 3563_at | IL3RA | -0,14022005 | 5,01258998 | -1,44746093 | 0,16900533 | 0,42274376 |
| 6102_at | RP2 | 0,24752815 | 7,97659727 | 1,4472431 | 0,16906537 | 0,42283817 |
| 22879_at | MON1B | 0,25987459 | 7,84898913 | 1,44716471 | 0,16908698 | 0,42283817 |
| 8497_at | PPFIA4 | -0,11691489 | 4,49873354 | -1,44691671 | 0,16915536 | 0,42295426 |
| 1998_at | ELF2 | 0,29037924 | 7,818383 | 1,44658547 | 0,16924672 | 0,42312779 |
| 102724532_a | LOC10272453 | 0,34294925 | 5,70813349 | 1,44641005 | 0,16929512 | 0,42319386 |
| 7596_at | ZNF45 | 0,09522327 | 6,79630264 | 1,44576586 | 0,16947297 | 0,42346959 |
| 55243_at | KIRREL | -0,14612524 | 5,38361729 | -1,44566902 | 0,16949972 | 0,42346959 |
| 2817_at | GPC1 | -0,26843536 | 8,95402953 | -1,44566816 | 0,16949995 | 0,42346959 |
| 50674_at | NEUROG3 | -0,13232467 | 4,67050929 | -1,44563215 | 0,1695099 | 0,42346959 |
| 101928580_a | LOC10192858 | -0,09016519 | 4,37232509 | -1,44554998 | 0,1695326 | 0,42346959 |
| 29886_at | SNX8 | -0,20691204 | 6,48006525 | -1,44553289 | 0,16953732 | 0,42346959 |
| 634_at | CEACAM1 | -0,24800731 | 7,13976404 | -1,44533213 | 0,1695928 | 0,42355323 |
| 159296_at | NKX2-3 | -0,11461119 | 4,06035274 | -1,44523664 | 0,16961919 | 0,42356422 |
| 100505549_a | LOC10050554 | 0,1316304 | 4,24046296 | 1,44515327 | 0,16964223 | 0,42356686 |
| 124599_at | CD300LB | -0,18031055 | 5,62414739 | -1,4449407 | 0,169701 | 0,42365867 |
| 80834_at | TAS1R2 | -0,18024741 | 4,48721227 | -1,44443793 | 0,16984006 | 0,42393017 |
| 84910_at | TMEM87B | 0,26284287 | 8,4737755 | 1,44429577 | 0,1698794 | 0,42393017 |
| 84152_at | PPP1R1B | -0,17302607 | 5,30910913 | -1,44423993 | 0,16989486 | 0,42393017 |
| 55610_at | VPS50 | 0,15788905 | 5,50496835 | 1,44422937 | 0,16989778 | 0,42393017 |
| 440078_at | FAM66C | 0,10066421 | 3,50858489 | 1,44407008 | 0,16994187 | 0,42398527 |
| 84223_at | IQCG | 0,20694747 | 6,55787214 | 1,44389876 | 0,1699893 | 0,42404868 |
| 1791_at | DNTT | -0,10484574 | 4,143215 | -1,44344722 | 0,17011437 | 0,42430572 |
| 114790_at | STK11IP | 0,1697506 | 7,12097755 | 1,44317056 | 0,17019103 | 0,42431773 |
| 1281_at | COL3A1 | 0,23831616 | 4,72713822 | 1,4430902 | 0,17021331 | 0,42431773 |
| 125950_at | RAVER1 | -0,18683909 | 7,39409668 | -1,44304542 | 0,17022572 | 0,42431773 |
| 9516_at | LITAF | 0,27750379 | 10,24152 | 1,44290898 | 0,17026355 | 0,42431773 |
| 6789_at | STK4 | 0,0847638 | 7,52274 | 1,4428831 | 0,17027072 | 0,42431773 |
| 116987_at | AGAP1 | -0,09904349 | 7,20097549 | -1,44286933 | 0,17027454 | 0,42431773 |
| 10642_at | IGF2BP1 | -0,17073016 | 4,65690607 | -1,44278766 | 0,17029719 | 0,42431773 |
| 246_at | ALOX15 | 0,1518258 | 4,25280725 | 1,44269175 | 0,17032379 | 0,42431773 |
| 202_at | AIM1 | 0,17527477 | 9,2348853 | 1,44267958 | 0,17032716 | 0,42431773 |
| 55844_at | PPP2R2D | -0,14652696 | 7,07248638 | -1,44258669 | 0,17035292 | 0,42431773 |
| 83862_at | TMEM120A | 0,39358541 | 8,21741675 | 1,44255583 | 0,17036149 | 0,42431773 |
| 5354_at | PLP1 | -0,11332882 | 3,47791723 | -1,44236294 | 0,170415 | 0,42436501 |
| 100131827_a | ZNF717 | 0,11818053 | 6,41495778 | 1,44232859 | 0,17042453 | 0,42436501 |
| 5066_at | PAM | 0,39377156 | 8,47263904 | 1,44204858 | 0,17050224 | 0,42450364 |
| 7125_at | TNNC2 | -0,14959536 | 5,44906484 | -1,44179648 | 0,17057223 | 0,42462302 |
| 5267_at | SERPINA4 | -0,15225777 | 4,39278713 | -1,44162505 | 0,17061984 | 0,42468665 |
| 51763_at | INPP5K | 0,16304364 | 7,1984279 | 1,44141609 | 0,17067789 | 0,42477624 |
| 10763_at | NES | -0,14784002 | 4,94377164 | -1,44117021 | 0,17074621 | 0,42489138 |
| 645687_at | LINC00520 | -0,19696014 | 4,51701096 | -1,44032094 | 0,17098237 | 0,42542409 |
| 80017_at | C14orf159 | 0,33674694 | 7,15559389 | 1,44015787 | 0,17102774 | 0,42548203 |
| 10848_at | PPP1R13L | 0,23556361 | 8,54943744 | 1,4394466 | 0,17122578 | 0,4259197 |

| | | | | | | |
|---|---|---|---|---|---|---|
| 7465_at | WEE1 | 0,1268146 | 8,31198273 | 1,4392112 | 0,17129136 | 0,4259888 |
| 130617_at | ZFAND2B | 0,07397132 | 7,88424245 | 1,43918814 | 0,17129779 | 0,4259888 |
| 57580_at | PREX1 | -0,10417274 | 5,03634704 | -1,43906517 | 0,17133205 | 0,42601902 |
| 7204_at | TRIO | 0,18047093 | 7,22488738 | 1,43842622 | 0,17151021 | 0,42636046 |
| 9318_at | COPS2 | -0,21270962 | 9,73571664 | -1,43833543 | 0,17153553 | 0,42636046 |
| 84256_at | FLYWCH1 | -0,10433059 | 7,09997138 | -1,43828582 | 0,17154937 | 0,42636046 |
| 255252_at | LRRC57 | 0,18813884 | 8,14229359 | 1,43825525 | 0,1715579 | 0,42636046 |
| 23098_at | SARM1 | 0,18825646 | 5,2953324 | 1,43807514 | 0,17160816 | 0,42643034 |
| 7075_at | TIE1 | -0,12222773 | 4,23052834 | -1,43771381 | 0,17170902 | 0,4264919 |
| 9732_at | DOCK4 | 0,22375075 | 3,83565614 | 1,43770581 | 0,17171126 | 0,4264919 |
| 54344_at | DPM3 | -0,11892081 | 8,60403551 | -1,43769717 | 0,17171367 | 0,4264919 |
| 1053_at | CEBPE | -0,10851323 | 4,7146449 | -1,43761157 | 0,17173757 | 0,4264919 |
| 124989_at | EFCAB13 | 0,06965569 | 3,19172272 | 1,43758986 | 0,17174363 | 0,4264919 |
| 79896_at | THNSL1 | -0,21485895 | 5,61239777 | -1,43729768 | 0,17182524 | 0,42661071 |
| 102_at | ADAM10 | 0,19446889 | 9,78802026 | 1,43721479 | 0,1718484 | 0,42661071 |
| 63898_at | SH2D4A | 0,29789863 | 8,57142715 | 1,43713564 | 0,17187052 | 0,42661071 |
| 5290_at | PIK3CA | 0,21200753 | 7,53330447 | 1,43710147 | 0,17188007 | 0,42661071 |
| 4237_at | MFAP2 | 0,60065157 | 6,48126497 | 1,43679935 | 0,17196451 | 0,4267653 |
| 911_at | CD1C | -0,18639008 | 4,85228939 | -1,43670922 | 0,1719897 | 0,42677285 |
| 950_at | SCARB2 | 0,46955748 | 8,72992013 | 1,43624581 | 0,17211931 | 0,42703944 |
| 11332_at | ACOT7 | -0,35968873 | 8,84701908 | -1,43599275 | 0,17219012 | 0,42712188 |
| 220972_at | MARCH8 | 0,22623 | 5,85408667 | 1,43574425 | 0,17225967 | 0,42712188 |
| 80108_at | ZFP2 | 0,16015304 | 3,69982359 | 1,43568832 | 0,17227533 | 0,42712188 |
| 644890_at | MEIG1 | -0,12056651 | 4,33485751 | -1,43565389 | 0,17228497 | 0,42712188 |
| 1124_at | CHN2 | 0,30533316 | 6,49182718 | 1,43564035 | 0,17228876 | 0,42712188 |
| 59341_at | TRPV4 | 0,34510911 | 4,99741342 | 1,43552105 | 0,17232217 | 0,42712188 |
| 83541_at | FAM110A | 0,21861989 | 7,89071636 | 1,43552084 | 0,17232222 | 0,42712188 |
| 80255_at | SLC35F5 | 0,28393162 | 9,13232264 | 1,43549336 | 0,17232992 | 0,42712188 |
| 2639_at | GCDH | -0,13178304 | 7,3512523 | -1,43494267 | 0,1724842 | 0,4273833 |
| 6990_at | DYNLT3 | 0,23329995 | 10,0724928 | 1,4348954 | 0,17249744 | 0,4273833 |
| 10821_at | OR7E12P | 0,15522448 | 6,08935997 | 1,43473029 | 0,17254372 | 0,4273833 |
| 10219_at | KLRG1 | -0,1339515 | 4,69963735 | -1,43467911 | 0,17255807 | 0,4273833 |
| 54471_at | MIEF1 | -0,12578455 | 7,95666835 | -1,43459217 | 0,17258244 | 0,4273833 |
| 8676_at | STX11 | 0,14063699 | 4,50276792 | 1,43456364 | 0,17259045 | 0,4273833 |
| 128821_at | CST9L | -0,12894182 | 3,78982186 | -1,43456273 | 0,1725907 | 0,4273833 |
| 55150_at | C19orf73 | -0,1456052 | 5,64910828 | -1,43434327 | 0,17265224 | 0,42748074 |
| 140689_at | CBLN4 | -0,08676117 | 3,51491152 | -1,43402925 | 0,17274034 | 0,4276054 |
| 54544_at | CRCT1 | 0,16437439 | 3,58379966 | 1,43400556 | 0,17274699 | 0,4276054 |
| 10501_at | SEMA6B | -0,14265598 | 5,63804451 | -1,43374869 | 0,17281908 | 0,42772888 |
| 101927851_a | LOC10192785 | -0,11172077 | 3,9012664 | -1,4336207 | 0,17285501 | 0,42773473 |
| 28809_at | IGLV3-1 | -0,13160312 | 5,57806559 | -1,43353546 | 0,17287894 | 0,42773473 |
| 152007_at | GLIPR2 | 0,49854878 | 6,58828163 | 1,43350299 | 0,17288806 | 0,42773473 |
| 54438_at | GFOD1 | -0,48842116 | 6,27156023 | -1,43299257 | 0,17303143 | 0,42795 |
| 93034_at | NT5C1B | -0,12679576 | 3,15842716 | -1,43295979 | 0,17304064 | 0,42795 |
| 57179_at | KIAA1191 | -0,3845174 | 9,42895643 | -1,43272032 | 0,17310794 | 0,42795 |
| 9567_at | GTPBP1 | 0,09506568 | 6,94941101 | 1,43271061 | 0,17311067 | 0,42795 |
| 54581_at | SCAND2P | 0,08470436 | 5,32722407 | 1,43267851 | 0,17311969 | 0,42795 |
| 2045_at | EPHA7 | 0,16349216 | 3,4736013 | 1,43265308 | 0,17312684 | 0,42795 |
| 22950_at | SLC4A1AP | -0,20372287 | 9,13128466 | -1,43258616 | 0,17314565 | 0,42795 |
| 6534_at | SLC6A7 | -0,20194606 | 6,09046333 | -1,43256076 | 0,17315279 | 0,42795 |

| | | | | | | |
|---|---|---|---|---|---|---|
| 84815_at | MGC12916 | -0,10296289 | 3,83787332 | -1,43241144 | 0,17319478 | 0,42799885 |
| 64718_at | UNKL | -0,18612441 | 7,14836001 | -1,43157797 | 0,17342929 | 0,4285234 |
| 151556_at | GPR155 | 0,10653298 | 3,78091438 | 1,43145749 | 0,17346321 | 0,42854067 |
| 10422_at | UBAC1 | -0,19675774 | 8,35588254 | -1,43139511 | 0,17348077 | 0,42854067 |
| 10107_at | TRIM10 | -0,11339896 | 4,31064765 | -1,43091396 | 0,1736163 | 0,42882047 |
| 80148_at | PQLC1 | -0,29697788 | 7,47957745 | -1,43005369 | 0,17385884 | 0,42926958 |
| 23166_at | STAB1 | -0,14557842 | 5,63157678 | -1,43002418 | 0,17386717 | 0,42926958 |
| 64417_at | C5orf28 | -0,19463169 | 7,31489536 | -1,43001548 | 0,17386962 | 0,42926958 |
| 6570_at | SLC18A1 | -0,12801718 | 3,73481444 | -1,42995292 | 0,17388727 | 0,42926958 |
| 132949_at | AASDH | 0,31358016 | 6,14149921 | 1,42982018 | 0,17392472 | 0,42930702 |
| 101929549_a | LOC10192954 | -0,16476238 | 4,60250465 | -1,42966077 | 0,17396971 | 0,42935789 |
| 90423_at | ATP6V1E2 | -0,18938158 | 5,57948144 | -1,42956136 | 0,17399777 | 0,42935789 |
| 7135_at | TNNI1 | -0,15406927 | 4,48953363 | -1,42951024 | 0,1740122 | 0,42935789 |
| 10557_at | RPP38 | -0,20549047 | 7,65848613 | -1,42929757 | 0,17407225 | 0,42941007 |
| 79998_at | ANKRD53 | -0,17685754 | 5,5508942 | -1,42927743 | 0,17407793 | 0,42941007 |
| 100506826_a | MYLK-AS1 | -0,12542021 | 5,79391393 | -1,42883656 | 0,17420247 | 0,42965442 |
| 84261_at | FBXW9 | -0,23815878 | 7,37878595 | -1,42876885 | 0,1742216 | 0,42965442 |
| 9025_at | RNF8 | -0,21411012 | 6,25729888 | -1,42842496 | 0,1743188 | 0,4298391 |
| 171169_at | SPACA4 | -0,1208155 | 5,32997418 | -1,42810072 | 0,17441049 | 0,43001013 |
| 100506790_a | LOC10050679 | -0,18398437 | 3,93018344 | -1,42799579 | 0,17444017 | 0,43002827 |
| 101928937_a | LOC10192893 | -0,10746472 | 3,57663551 | -1,42783837 | 0,1744847 | 0,43003956 |
| 5251_at | PHEX | 0,29512931 | 6,55768698 | 1,42782178 | 0,1744894 | 0,43003956 |
| 56895_at | AGPAT4 | 0,34580075 | 5,28852726 | 1,42756342 | 0,17456251 | 0,43016472 |
| 8605_at | PLA2G4C | 0,10092142 | 3,41934353 | 1,42738172 | 0,17461394 | 0,43021014 |
| 2191_at | FAP | -0,16051048 | 4,57711605 | -1,42734052 | 0,17462561 | 0,43021014 |
| 3984_at | LIMK1 | -0,17759313 | 6,73702579 | -1,4270585 | 0,17470547 | 0,43035184 |
| 5009_at | OTC | -0,12206385 | 3,42860639 | -1,42672918 | 0,17479876 | 0,43050635 |
| 342897_at | NCCRP1 | -0,26781275 | 6,00784964 | -1,42664195 | 0,17482348 | 0,43050635 |
| 6650_at | CAPN15 | -0,14049498 | 6,56622999 | -1,42659801 | 0,17483593 | 0,43050635 |
| 10081_at | PDCD7 | -0,2085975 | 8,32694238 | -1,42633934 | 0,17490925 | 0,43050635 |
| 35_at | ACADS | 0,16939027 | 6,52131646 | 1,42626754 | 0,17492961 | 0,43050635 |
| 130497_at | OSR1 | 0,80939784 | 6,33485319 | 1,42625703 | 0,17493259 | 0,43050635 |
| 1587_at | ADAM3A | 0,08857073 | 2,84490826 | 1,42617394 | 0,17495615 | 0,43050635 |
| 101929125_a | LOC10192912 | -0,13031976 | 3,49007255 | -1,42616375 | 0,17495904 | 0,43050635 |
| 6853_at | SYN1 | -0,14857224 | 5,11529666 | -1,42612745 | 0,17496933 | 0,43050635 |
| 1013_at | CDH15 | -0,13109031 | 5,51089858 | -1,42509686 | 0,17526179 | 0,43117086 |
| 3512_at | JCHAIN | 0,09311035 | 2,87359519 | 1,42476437 | 0,17535623 | 0,43134811 |
| 116966_at | WDR17 | 0,10466115 | 2,87289844 | 1,42454538 | 0,17541845 | 0,43144608 |
| 11232_at | POLG2 | -0,12303254 | 5,97358458 | -1,42436708 | 0,17546913 | 0,43149213 |
| 154_at | ADRB2 | 0,57457412 | 7,78492877 | 1,42419852 | 0,17551705 | 0,43149213 |
| 1912_at | PHC2 | -0,14711857 | 6,1274744 | -1,42416792 | 0,17552575 | 0,43149213 |
| 3417_at | IDH1 | 0,23758277 | 12,3689097 | 1,42416429 | 0,17552678 | 0,43149213 |
| 571_at | BACH1 | 0,25572634 | 5,90731895 | 1,42385541 | 0,17561462 | 0,43159858 |
| 28966_at | SNX24 | 0,25368751 | 6,16044176 | 1,42385445 | 0,17561489 | 0,43159858 |
| 389906_at | LOC389906 | -0,39803993 | 7,09986643 | -1,42374587 | 0,17564578 | 0,43161943 |
| 348751_at | FTCDNL1 | -0,15051842 | 4,60197699 | -1,42354257 | 0,17570362 | 0,4317065 |
| 6710_at | SPTB | -0,113262 | 5,53165053 | -1,4234421 | 0,17573222 | 0,43172168 |
| 1135_at | CHRNA2 | -0,22294275 | 5,47862527 | -1,42303012 | 0,1758495 | 0,43195472 |
| 84134_at | TOMM40L | -0,2640178 | 7,497722 | -1,42292584 | 0,1758792 | 0,43197258 |
| 162282_at | ANKFN1 | -0,10138427 | 3,86799094 | -1,42267969 | 0,17594931 | 0,43208957 |

| | | | | | | |
|---|---|---|---|---|---|---|
| 57786_at | RBAK | 0,1287704 | 6,56027908 | 1,42260113 | 0,17597169 | 0,43208957 |
| 3675_at | ITGA3 | -0,289123 | 8,83276394 | -1,42244239 | 0,17601693 | 0,43211988 |
| 759_at | CA1 | -0,17215018 | 4,81480775 | -1,42240037 | 0,1760289 | 0,43211988 |
| 55207_at | ARL8B | 0,15732964 | 10,0308767 | 1,42215173 | 0,17609977 | 0,43223878 |
| 10299_at | MARCH6 | 0,14895368 | 8,68679705 | 1,42196253 | 0,17615372 | 0,43231611 |
| 26232_at | FBXO2 | 0,32952891 | 6,21863433 | 1,42172517 | 0,17622142 | 0,43242715 |
| 3664_at | IRF6 | 0,14981884 | 8,84494168 | 1,42159937 | 0,1762573 | 0,43244196 |
| 2683_at | B4GALT1 | 0,09520669 | 8,64336541 | 1,42154664 | 0,17627235 | 0,43244196 |
| 102723099_a | GHET1 | 0,44514101 | 5,47446683 | 1,42113994 | 0,17638842 | 0,43267161 |
| 6499_at | SKIV2L | -0,07285342 | 7,91405029 | -1,42105165 | 0,17641363 | 0,43267835 |
| 4112_at | MAGEB1 | -0,15944956 | 3,65318354 | -1,42073838 | 0,17650309 | 0,43284264 |
| 10945_at | KDELR1 | 0,1154415 | 10,3851061 | 1,42060697 | 0,17654063 | 0,43287959 |
| 730051_at | ZNF814 | 0,25169783 | 3,68279788 | 1,41998199 | 0,17671925 | 0,43326242 |
| 5447_at | POR | 0,21729801 | 9,15810155 | 1,4198874 | 0,17674629 | 0,43327358 |
| 83642_at | SELO | -0,15906125 | 8,41672492 | -1,41972345 | 0,17679318 | 0,43333337 |
| 114932_at | MRFAP1L1 | 0,16016011 | 10,1372596 | 1,41956324 | 0,17683901 | 0,43339056 |
| 57586_at | SYT13 | -0,10164153 | 4,10259886 | -1,41936383 | 0,17689606 | 0,43345446 |
| 1302_at | COL11A2 | -0,17831836 | 6,09972851 | -1,41925629 | 0,17692683 | 0,43345446 |
| 1638_at | DCT | -0,09273324 | 3,98157861 | -1,4192362 | 0,17693259 | 0,43345446 |
| 148231_at | LINC00905 | -0,11702099 | 5,18999942 | -1,4189509 | 0,17701425 | 0,43359938 |
| 55786_at | ZNF415 | 0,19205224 | 6,05691306 | 1,417947 | 0,17730187 | 0,43400709 |
| 729967_at | MORN2 | 0,16877252 | 7,96291346 | 1,4179067 | 0,17731343 | 0,43400709 |
| 51187_at | RSL24D1 | -0,20439873 | 9,18233524 | -1,41789651 | 0,17731635 | 0,43400709 |
| 5995_at | RGR | -0,15080759 | 5,00174682 | -1,41780945 | 0,17734131 | 0,43400709 |
| 91_at | ACVR1B | 0,24289017 | 6,95297428 | 1,41768596 | 0,17737672 | 0,43400709 |
| 55347_at | ABHD10 | -0,13561722 | 8,22251549 | -1,4176466 | 0,17738801 | 0,43400709 |
| 57156_at | TMEM63C | -0,17173878 | 4,18615459 | -1,41756339 | 0,17741188 | 0,43400709 |
| 84993_at | UBL7 | -0,11446741 | 8,24518806 | -1,41751325 | 0,17742626 | 0,43400709 |
| 5309_at | PITX3 | -0,13345644 | 6,04753006 | -1,41749887 | 0,17743039 | 0,43400709 |
| 5046_at | PCSK6 | -0,23126283 | 5,65318856 | -1,41742732 | 0,17745091 | 0,43400709 |
| 8712_at | PAGE1 | -0,13053589 | 3,54038943 | -1,4173674 | 0,1774681 | 0,43400709 |
| 9650_at | MTFR1 | -0,25279062 | 8,99539857 | -1,41731112 | 0,17748425 | 0,43400709 |
| 80712_at | ESX1 | -0,13633611 | 4,65334501 | -1,41723636 | 0,17750571 | 0,43400709 |
| 203547_at | VMA21 | -0,27468345 | 7,85682084 | -1,41720202 | 0,17751556 | 0,43400709 |
| 134553_at | C5orf24 | 0,13340745 | 7,61481916 | 1,41714212 | 0,17753275 | 0,43400709 |
| 6352_at | CCL5 | 0,68426899 | 5,0626413 | 1,41711272 | 0,17754119 | 0,43400709 |
| 5594_at | MAPK1 | -0,10494944 | 9,53304969 | -1,4164958 | 0,17771833 | 0,43438499 |
| 101929164_a | LOC10192916 | -0,13271607 | 4,2385821 | -1,416389 | 0,177749 | 0,43440486 |
| 23542_at | MAPK8IP2 | -0,15020509 | 5,69494904 | -1,41580062 | 0,17791811 | 0,43476297 |
| 9666_at | DZIP3 | 0,24911163 | 6,81010517 | 1,41540467 | 0,17803198 | 0,43496083 |
| 101929760_a | PIK3IP1-AS1 | 0,13549967 | 3,55816352 | 1,41536206 | 0,17804424 | 0,43496083 |
| 6276_at | S100A5 | -0,22420753 | 4,74222558 | -1,41494544 | 0,17816412 | 0,4351601 |
| 8544_at | PIR | -0,17500994 | 9,25100987 | -1,41492159 | 0,17817099 | 0,4351601 |
| 4092_at | SMAD7 | 0,45116316 | 7,07618056 | 1,41446829 | 0,17830151 | 0,43540131 |
| 9623_at | TCL1B | -0,18710582 | 4,46724694 | -1,41442161 | 0,17831495 | 0,43540131 |
| 10148_at | EBI3 | 0,17256415 | 5,16891643 | 1,41401394 | 0,17843241 | 0,4356329 |
| 6553_at | SLC9A5 | -0,17121897 | 5,07330229 | -1,41384575 | 0,17848089 | 0,43569603 |
| 79960_at | JADE1 | -0,31217028 | 6,42934091 | -1,41331256 | 0,17863464 | 0,43599222 |
| 921_at | CD5 | -0,14783303 | 5,19810379 | -1,41326802 | 0,17864749 | 0,43599222 |
| 952_at | CD38 | -0,11196125 | 5,15986529 | -1,41314987 | 0,17868158 | 0,43602017 |

| | | | | | | |
|---|---|---|---|---|---|---|
| 56005_at | MYDGF | -0,10440835 | 9,73949911 | -1,41296589 | 0,17873467 | 0,43609448 |
| 84679_at | SLC9A7 | -0,27951453 | 6,27095039 | -1,41235434 | 0,17891123 | 0,43646999 |
| 10369_at | CACNG2 | -0,13905064 | 4,10050266 | -1,41217864 | 0,17896199 | 0,43652658 |
| 661_at | POLR3D | -0,34011196 | 7,44303027 | -1,41211715 | 0,17897975 | 0,43652658 |
| 114792_at | KLHL32 | 0,10995219 | 2,97986832 | 1,4118359 | 0,17906102 | 0,43662503 |
| 1477_at | CSTF1 | -0,14423547 | 8,49407777 | -1,4118206 | 0,17906545 | 0,43662503 |
| 23217_at | ZFR2 | -0,15667723 | 5,17680551 | -1,41131452 | 0,17921177 | 0,4369153 |
| 2219_at | FCN1 | -0,13417602 | 4,83873325 | -1,41120771 | 0,17924267 | 0,4369153 |
| 10300_at | KATNB1 | -0,12450075 | 7,0751973 | -1,41111123 | 0,17927058 | 0,4369153 |
| 2662_at | GDF10 | -0,129231 | 4,05691568 | -1,41106177 | 0,17928489 | 0,4369153 |
| 57037_at | ANKMY2 | 0,24184094 | 8,36290326 | 1,4110168 | 0,1792979 | 0,4369153 |
| 1856_at | DVL2 | 0,10665513 | 7,50704195 | 1,4106744 | 0,179397 | 0,43710148 |
| 79827_at | CLMP | -0,13789843 | 3,73488742 | -1,41051001 | 0,17944459 | 0,43710169 |
| 1193_at | CLIC2 | 0,16058397 | 3,19331131 | 1,41049572 | 0,17944873 | 0,43710169 |
| 114134_at | SLC2A13 | 0,47192863 | 5,65048064 | 1,41043898 | 0,17946515 | 0,43710169 |
| 9306_at | SOCS6 | 0,27649331 | 6,39690382 | 1,40993698 | 0,17961056 | 0,43740054 |
| 549_at | AUH | 0,13498739 | 6,81487962 | 1,40969758 | 0,17967995 | 0,43743881 |
| 138311_at | FAM69B | -0,11904107 | 5,31537431 | -1,4096816 | 0,17968458 | 0,43743881 |
| 162967_at | ZNF320 | 0,13387937 | 6,95452473 | 1,40962989 | 0,17969957 | 0,43743881 |
| 284382_at | ACTL9 | -0,17543978 | 4,51014667 | -1,40956937 | 0,17971711 | 0,43743881 |
| 105376684_a | LOC10537668 | 0,12496638 | 4,86528351 | 1,40924179 | 0,1798121 | 0,4375753 |
| 19_at | ABCA1 | 0,56954399 | 4,9804551 | 1,40920434 | 0,17982296 | 0,4375753 |
| 9821_at | RB1CC1 | 0,34186222 | 8,43624087 | 1,40909891 | 0,17985354 | 0,4375753 |
| 54996_at | MARC2 | 0,2556046 | 6,42335109 | 1,40900121 | 0,17988188 | 0,4375753 |
| 27173_at | SLC39A1 | 0,16794649 | 9,1502926 | 1,40896443 | 0,17989256 | 0,4375753 |
| 11326_at | VSIG4 | -0,15723452 | 4,36133414 | -1,40875835 | 0,17995235 | 0,4375753 |
| 28395_at | IGHV4-34 | -0,16890954 | 4,79469585 | -1,40875039 | 0,17995467 | 0,4375753 |
| 256329_at | LMNTD2 | 0,18058325 | 6,00334302 | 1,4086946 | 0,17997086 | 0,4375753 |
| 28680_at | TRAV8-6 | -0,19399032 | 4,22788134 | -1,40862498 | 0,17999106 | 0,4375753 |
| 7474_at | WNT5A | 0,35504138 | 7,9393871 | 1,40854467 | 0,18001437 | 0,4375753 |
| 9540_at | TP53I3 | 0,46120801 | 8,85337455 | 1,40850084 | 0,1800271 | 0,4375753 |
| 525_at | ATP6V1B1 | 0,47964085 | 5,75249147 | 1,40830766 | 0,18008319 | 0,4375753 |
| 285753_at | CEP57L1 | -0,18355168 | 4,7320301 | -1,4082972 | 0,18008623 | 0,4375753 |
| 400410_at | ST20 | 0,22904135 | 5,22840947 | 1,40825237 | 0,18009925 | 0,4375753 |
| 284695_at | ZNF326 | -0,22139869 | 6,57236951 | -1,40814824 | 0,18012949 | 0,4375753 |
| 1592_at | CYP26A1 | 0,12178391 | 4,61170597 | 1,40812361 | 0,18013664 | 0,4375753 |
| 348_at | APOE | -0,12943631 | 5,90219127 | -1,40770806 | 0,18025737 | 0,43781337 |
| 9844_at | ELMO1 | -0,1682535 | 4,39959887 | -1,40756102 | 0,18030011 | 0,43786196 |
| 201477_at | DLGAP1-AS3 | -0,1127556 | 3,40355671 | -1,40711351 | 0,18043023 | 0,43807111 |
| 8175_at | SF3A2 | -0,15000376 | 8,6365301 | -1,4071084 | 0,18043172 | 0,43807111 |
| 146713_at | RBFOX3 | -0,13708907 | 4,34622608 | -1,40670251 | 0,1805498 | 0,43830255 |
| 84316_at | NAA38 | -0,23999084 | 8,95534674 | -1,40609969 | 0,18072529 | 0,4386733 |
| 10692_at | RRH | -0,13802995 | 3,63921456 | -1,40545226 | 0,18091393 | 0,43899383 |
| 28516_at | TRDV3 | -0,12783216 | 3,50714417 | -1,40543152 | 0,18091998 | 0,43899383 |
| 338382_at | RAB7B | 0,11782746 | 5,29837054 | 1,40540359 | 0,18092812 | 0,43899383 |
| 50717_at | DCAF8 | 0,20564877 | 6,76690617 | 1,40531487 | 0,18095398 | 0,43899383 |
| 51361_at | HOOK1 | -0,13680768 | 6,69542969 | -1,40521383 | 0,18098344 | 0,43899383 |
| 9636_at | ISG15 | 0,55545463 | 9,6520463 | 1,40517734 | 0,18099408 | 0,43899383 |
| 3137_at | HLA-J | 0,48709982 | 7,79614218 | 1,40461954 | 0,18115679 | 0,43933315 |
| 57105_at | CYSLTR2 | -0,12798717 | 3,44233155 | -1,40452014 | 0,1811858 | 0,43934819 |

| | | | | | | |
|---|---|---|---|---|---|---|
| 9348_at | NDST3 | -0,11612719 | 3,33151888 | -1,40436088 | 0,18123228 | 0,43935655 |
| 59283_at | CACNG8 | -0,16132696 | 5,53585674 | -1,40432686 | 0,18124221 | 0,43935655 |
| 1381_at | CRABP1 | -0,2046017 | 4,69774104 | -1,40426665 | 0,18125979 | 0,43935655 |
| 79742_at | CXorf36 | -0,13505124 | 4,72017108 | -1,40419579 | 0,18128048 | 0,43935655 |
| 973_at | CD79A | -0,14953766 | 5,53692985 | -1,40410752 | 0,18130625 | 0,43935932 |
| 1305_at | COL13A1 | -0,14777391 | 5,22142554 | -1,40403565 | 0,18132724 | 0,43935932 |
| 2048_at | EPHB2 | 0,1858158 | 7,74669542 | 1,40381435 | 0,18139188 | 0,43946066 |
| 6855_at | SYP | -0,11015177 | 5,20435415 | -1,4033963 | 0,18151403 | 0,43970114 |
| 283738_at | NTRK3-AS1 | -0,11183985 | 3,83103186 | -1,40331322 | 0,18153831 | 0,43970114 |
| 96597_at | TBC1D27 | -0,24458979 | 4,682229 | -1,40324035 | 0,18155961 | 0,43970114 |
| 10044_at | SH2D3C | -0,14993249 | 5,33172921 | -1,40307115 | 0,18160908 | 0,43974179 |
| 26291_at | FGF21 | -0,13050817 | 4,74384681 | -1,40299072 | 0,1816326 | 0,43974179 |
| 9622_at | KLK4 | -0,16416705 | 5,22398272 | -1,40291951 | 0,18165343 | 0,43974179 |
| 100101938_a | ANKRD26P3 | -0,07694008 | 4,34830154 | -1,40281604 | 0,18168369 | 0,43974179 |
| 155061_at | ZNF746 | 0,11301002 | 7,91520684 | 1,40278775 | 0,18169197 | 0,43974179 |
| 79156_at | PLEKHF1 | 0,30026176 | 7,53727569 | 1,40265749 | 0,18173007 | 0,43974179 |
| 10815_at | CPLX1 | -0,319207 | 4,80553514 | -1,40263655 | 0,1817362 | 0,43974179 |
| 9337_at | CNOT8 | 0,16393701 | 8,5179227 | 1,40240996 | 0,1818025 | 0,43984698 |
| 1101_at | CHAD | -0,15972474 | 5,10692973 | -1,40229981 | 0,18183474 | 0,43986973 |
| 1003_at | CDH5 | 0,15455883 | 4,81040142 | 1,40219871 | 0,18186434 | 0,43988608 |
| 84334_at | APOPT1 | -0,07803626 | 8,05521449 | -1,40178973 | 0,18198409 | 0,44002788 |
| 8204_at | NRIP1 | 0,25966611 | 9,70214864 | 1,40175349 | 0,18199471 | 0,44002788 |
| 3217_at | HOXB7 | 0,27206375 | 8,98707712 | 1,40174266 | 0,18199788 | 0,44002788 |
| 153770_at | PLAC8L1 | -0,12015791 | 4,68317715 | -1,40168649 | 0,18201434 | 0,44002788 |
| 4350_at | MPG | 0,15391188 | 8,43354555 | 1,4014388 | 0,1820869 | 0,4400716 |
| 845_at | CASQ2 | -0,17760291 | 4,06160903 | -1,4013305 | 0,18211864 | 0,4400716 |
| 81850_at | KRTAP1-3 | 0,11020555 | 4,30589443 | 1,4012848 | 0,18213204 | 0,4400716 |
| 79078_at | C1orf50 | 0,10810084 | 7,21769702 | 1,40110102 | 0,18218591 | 0,4400716 |
| 29880_at | ALG5 | -0,18457438 | 9,24824935 | -1,40094999 | 0,18223019 | 0,4400716 |
| 6272_at | SORT1 | 0,10158835 | 6,71883977 | 1,40092335 | 0,182238 | 0,4400716 |
| 11105_at | PRDM7 | -0,12123244 | 4,34277577 | -1,40087852 | 0,18225115 | 0,4400716 |
| 10347_at | ABCA7 | 0,27492449 | 6,776959 | 1,40087596 | 0,1822519 | 0,4400716 |
| 1353_at | COX11 | -0,16274821 | 7,48556516 | -1,40071393 | 0,18229942 | 0,4400716 |
| 10973_at | ASCC3 | -0,18230314 | 7,89097768 | -1,40069537 | 0,18230486 | 0,4400716 |
| 92105_at | INTS4 | -0,10429253 | 7,13967764 | -1,40069219 | 0,1823058 | 0,4400716 |
| 79943_at | ZNF696 | -0,19687021 | 5,78447609 | -1,40068957 | 0,18230657 | 0,4400716 |
| 56928_at | SPPL2B | 0,18584385 | 7,03171938 | 1,40056685 | 0,18234257 | 0,44010335 |
| 3351_at | HTR1B | -0,15864653 | 4,73802261 | -1,40014854 | 0,18246532 | 0,44031239 |
| 2786_at | GNG4 | -0,42058585 | 4,78361717 | -1,40005586 | 0,18249253 | 0,44031239 |
| 860_at | RUNX2 | -0,14163882 | 5,72278035 | -1,40003296 | 0,18249925 | 0,44031239 |
| 8798_at | DYRK4 | -0,14969316 | 5,82908998 | -1,39996022 | 0,1825206 | 0,44031239 |
| 6160_at | RPL31 | 0,1846607 | 7,345911 | 1,3996968 | 0,18259796 | 0,44044384 |
| 11129_at | CLASRP | -0,09705338 | 7,57940023 | -1,39942527 | 0,18267773 | 0,44058108 |
| 55693_at | KDM4D | -0,12799712 | 5,12353844 | -1,39898352 | 0,18280756 | 0,44083901 |
| 3560_at | IL2RB | -0,17982897 | 5,36285423 | -1,39804246 | 0,18308439 | 0,44136291 |
| 57662_at | CAMSAP3 | 0,14729583 | 6,27758789 | 1,39802455 | 0,18308966 | 0,44136291 |
| 415117_at | STX19 | 0,47973802 | 4,15610276 | 1,39801134 | 0,18309355 | 0,44136291 |
| 24140_at | FTSJ1 | -0,12449864 | 8,7938327 | -1,39751612 | 0,18323937 | 0,44165916 |
| 101928834_a | LOC10192883 | -0,1004633 | 3,34823975 | -1,39739244 | 0,1832758 | 0,4416917 |
| 166979_at | CDC20B | -0,11625805 | 4,24307915 | -1,39705583 | 0,18337499 | 0,44186982 |

| | | | | | | |
|---|---|---|---|---|---|---|
| 51228_at | GLTP | 0,16405929 | 9,74374546 | 1,39697553 | 0,18339866 | 0,44186982 |
| 84996_at | URB1-AS1 | -0,13651189 | 6,97672723 | -1,39690813 | 0,18341853 | 0,44186982 |
| 283104_at | SBF2-AS1 | 0,30860319 | 5,85335408 | 1,39648957 | 0,18354195 | 0,44211186 |
| 64231_at | MS4A6A | -0,12354295 | 3,89164466 | -1,39589336 | 0,18371787 | 0,44242809 |
| 100505624_a | GTF3C2-AS1 | -0,13142327 | 4,60323481 | -1,39582623 | 0,18373769 | 0,44242809 |
| 9118_at | INA | -0,22686792 | 7,48585801 | -1,39570041 | 0,18377483 | 0,44242809 |
| 55297_at | CCDC91 | -0,13550437 | 7,81066019 | -1,39565102 | 0,18378942 | 0,44242809 |
| 26502_at | NARF | 0,15437767 | 8,79910101 | 1,3956053 | 0,18380292 | 0,44242809 |
| 4047_at | LSS | -0,14113742 | 7,90910305 | -1,39557781 | 0,18381104 | 0,44242809 |
| 146225_at | CMTM2 | -0,1376862 | 3,83034286 | -1,3953891 | 0,18386677 | 0,44250694 |
| 439941_at | FER1L6-AS1 | -0,12871551 | 3,12473139 | -1,39514654 | 0,18393843 | 0,44252514 |
| 8477_at | GPR65 | -0,11713065 | 3,21580648 | -1,39514618 | 0,18393854 | 0,44252514 |
| 90665_at | TBL1Y | -0,11438931 | 4,27373016 | -1,39513023 | 0,18394325 | 0,44252514 |
| 28666_at | TRAV17 | 0,14901918 | 3,67336432 | 1,39503816 | 0,18397046 | 0,44253533 |
| 5420_at | PODXL | -0,47889152 | 9,78736705 | -1,3948802 | 0,18401714 | 0,44259236 |
| 101927953_a | C5orf66-AS1 | -0,09244796 | 4,63171424 | -1,39447438 | 0,18413712 | 0,44277897 |
| 5452_at | POU2F2 | -0,11429284 | 5,23345916 | -1,39440691 | 0,18415708 | 0,44277897 |
| 28663_at | TRAV20 | -0,15288017 | 3,8046782 | -1,39438457 | 0,18416369 | 0,44277897 |
| 9509_at | ADAMTS2 | -0,12871672 | 5,08748904 | -1,39413278 | 0,18423817 | 0,44290277 |
| 11311_at | VPS45 | 0,12190231 | 8,85836997 | 1,39395088 | 0,184292 | 0,44297688 |
| 57338_at | JPH3 | -0,11977236 | 4,58782981 | -1,39372406 | 0,18435913 | 0,44308296 |
| 9896_at | FIG4 | 0,35313961 | 7,70035349 | 1,39348411 | 0,18443017 | 0,44316715 |
| 22796_at | COG2 | -0,14883864 | 7,7810034 | -1,39345034 | 0,18444017 | 0,44316715 |
| 63893_at | UBE2O | -0,19996171 | 7,64874527 | -1,39320727 | 0,18451216 | 0,44323215 |
| 392459_at | CXXC1P1 | -0,14514136 | 3,9625684 | -1,3931704 | 0,18452309 | 0,44323215 |
| 3549_at | IHH | -0,12975752 | 5,56436258 | -1,39312596 | 0,18453625 | 0,44323215 |
| 101928205_a | LOC10192820 | -0,13013129 | 3,99493212 | -1,39261889 | 0,18468653 | 0,44353779 |
| 124751_at | KRBA2 | 0,10592137 | 5,78870158 | 1,39236705 | 0,1847612 | 0,4436618 |
| 64100_at | ELSPBP1 | -0,1297338 | 4,26907713 | -1,39208514 | 0,18484482 | 0,44380727 |
| 150159_at | SLC9B1 | -0,08903836 | 3,17990652 | -1,39183635 | 0,18491864 | 0,4438434 |
| 3651_at | PDX1 | 0,10347747 | 4,31232649 | 1,39181942 | 0,18492366 | 0,4438434 |
| 8289_at | ARID1A | 0,10633622 | 8,34893073 | 1,39180146 | 0,18492899 | 0,4438434 |
| 6363_at | CCL19 | -0,1568342 | 4,92176708 | -1,39166773 | 0,18496868 | 0,44384828 |
| 100289341_a | MAN1B1-AS1 | -0,32611011 | 6,54584079 | -1,39163936 | 0,1849771 | 0,44384828 |
| 3096_at | HIVEP1 | 0,08352634 | 6,16686017 | 1,39140739 | 0,18504597 | 0,44395823 |
| 285819_at | LOC285819 | -0,19564187 | 4,30151753 | -1,39129325 | 0,18507987 | 0,44398424 |
| 65997_at | RASL11B | 0,1007837 | 5,74437978 | 1,39107297 | 0,18514529 | 0,44408589 |
| 22875_at | ENPP4 | 0,34954728 | 5,77543196 | 1,3906371 | 0,18527481 | 0,44429745 |
| 101928446_a | LOC10192844 | -0,10497919 | 3,39278486 | -1,39048 | 0,18532151 | 0,44429745 |
| 8862_at | APLN | -0,1281754 | 5,73148711 | -1,39046629 | 0,18532558 | 0,44429745 |
| 7809_at | BSND | -0,13204049 | 4,80389433 | -1,39046572 | 0,18532575 | 0,44429745 |
| 128434_at | VSTM2L | -0,14394714 | 5,97995561 | -1,3903031 | 0,18537411 | 0,44435806 |
| 6616_at | SNAP25 | -0,12134778 | 4,17863552 | -1,39007994 | 0,18544048 | 0,44444049 |
| 158572_at | USP27X-AS1 | 0,18638114 | 5,30670045 | 1,39003233 | 0,18545464 | 0,44444049 |
| 338588_at | LINC00705 | -0,16663752 | 4,68192302 | -1,38986521 | 0,18550436 | 0,44448022 |
| 1757_at | SARDH | -0,11154063 | 4,91654612 | -1,38979942 | 0,18552393 | 0,44448022 |
| 1410_at | CRYAB | -0,2055185 | 4,97259952 | -1,38974395 | 0,18554044 | 0,44448022 |
| 84435_at | ADGRA1 | -0,165019 | 5,52043079 | -1,38966042 | 0,1855653 | 0,44448449 |
| 2992_at | GYG1 | -0,13550869 | 9,85567889 | -1,38944799 | 0,18562852 | 0,44458066 |
| 116085_at | SLC22A12 | -0,180295 | 5,75485225 | -1,38918679 | 0,18570629 | 0,44465857 |

| | | | | | | |
|---|---|---|---|---|---|---|
| 27095_at | TRAPPC3 | -0,07338027 | 9,22309866 | -1,38918368 | 0,18570722 | 0,44465857 |
| 4068_at | SH2D1A | -0,09783465 | 3,82332366 | -1,38908149 | 0,18573765 | 0,44467617 |
| 55832_at | CAND1 | 0,117983 | 9,69493814 | 1,38900174 | 0,18576141 | 0,44467777 |
| 100132341_a | CLUHP3 | 0,13348916 | 6,83758901 | 1,38891462 | 0,18578736 | 0,44468464 |
| 138649_at | ANKRD19P | -0,1745548 | 6,03326622 | -1,38858557 | 0,18588541 | 0,44486403 |
| 84146_at | ZNF644 | 0,17867381 | 7,84481245 | 1,38799367 | 0,18606188 | 0,44523106 |
| 10581_at | IFITM2 | 0,31904207 | 6,81789045 | 1,38769896 | 0,1861498 | 0,44535523 |
| 54557_at | SGTB | -0,11954189 | 5,06294706 | -1,38766473 | 0,18616001 | 0,44535523 |
| 100996455_a | LOC10099645 | -0,12306622 | 3,87790294 | -1,38752119 | 0,18620285 | 0,44540239 |
| 135935_at | NOBOX | -0,13805116 | 5,09332276 | -1,38733723 | 0,18625776 | 0,44547841 |
| 11334_at | TUSC2 | 0,09356417 | 7,8214179 | 1,38724514 | 0,18628525 | 0,44548885 |
| 387066_at | SNHG5 | -0,28083619 | 8,87383231 | -1,3871299 | 0,18631966 | 0,44551583 |
| #N/A | CTB-113P19,1 | -0,18217732 | 4,01472907 | -1,38681697 | 0,18641311 | 0,44568398 |
| 201475_at | RAB12 | -0,2783546 | 8,22138742 | -1,3867032 | 0,1864471 | 0,44570992 |
| 170463_at | SSBP4 | -0,16105499 | 7,63197243 | -1,38658052 | 0,18648376 | 0,44574223 |
| 132203_at | SNTN | 0,12221907 | 2,90053438 | 1,38643872 | 0,18652614 | 0,44578821 |
| 167227_at | DCP2 | 0,27287702 | 6,99058827 | 1,38604473 | 0,18664391 | 0,44598222 |
| 4192_at | MDK | 0,76335837 | 8,62989452 | 1,38601228 | 0,18665362 | 0,44598222 |
| 51270_at | TFDP3 | -0,13876327 | 3,83054612 | -1,38545303 | 0,18682092 | 0,44630344 |
| 27198_at | HCAR1 | -0,18164532 | 4,68424604 | -1,38540799 | 0,1868344 | 0,44630344 |
| 7433_at | VIPR1 | 0,25747672 | 8,95187016 | 1,38515889 | 0,18690896 | 0,44642619 |
| 100506271_a | LOC10050627 | 0,19231325 | 5,93946853 | 1,38503392 | 0,18694637 | 0,4464602 |
| 7486_at | WRN | -0,30808694 | 6,71260025 | -1,38481941 | 0,18701061 | 0,44655824 |
| 6050_at | RNH1 | -0,07264153 | 8,923121 | -1,38457232 | 0,18708462 | 0,44667961 |
| 976_at | ADGRE5 | -0,29491206 | 8,26737282 | -1,38436627 | 0,18714636 | 0,44677165 |
| 283624_at | LINC00641 | -0,23604813 | 4,4958462 | -1,3842398 | 0,18718427 | 0,44680677 |
| 81030_at | ZBP1 | 0,22718133 | 4,31095791 | 1,38409434 | 0,18722787 | 0,44683695 |
| 309_at | ANXA6 | -0,15387621 | 5,25832977 | -1,38404285 | 0,18724331 | 0,44683695 |
| 51077_at | FCF1 | -0,13642017 | 7,91049841 | -1,38392493 | 0,18727867 | 0,44686596 |
| 9882_at | TBC1D4 | 0,39985281 | 6,97736464 | 1,38352336 | 0,18739911 | 0,44709798 |
| 387104_at | SOGA3 | -0,08302003 | 3,51459226 | -1,38320414 | 0,1874949 | 0,44727111 |
| 83468_at | GLT8D2 | 0,06983914 | 3,51475269 | 1,383057 | 0,18753906 | 0,44731814 |
| 83714_at | NRIP2 | -0,1635088 | 4,52611479 | -1,38297145 | 0,18756475 | 0,44731814 |
| 140733_at | MACROD2 | 0,8673064 | 5,30263861 | 1,38285541 | 0,18759959 | 0,44731814 |
| 85004_at | RERG | -0,18506962 | 4,3198243 | -1,38282906 | 0,1876075 | 0,44731814 |
| 100289361_a | LOC10028936 | -0,1452558 | 5,77878494 | -1,38245315 | 0,18772041 | 0,44753196 |
| 28232_at | SLCO3A1 | -0,15327318 | 8,14168343 | -1,38236367 | 0,18774729 | 0,44754066 |
| 5045_at | FURIN | -0,12404304 | 7,74103518 | -1,38164698 | 0,18796274 | 0,44799878 |
| 1352_at | COX10 | -0,26919829 | 7,99275425 | -1,38138531 | 0,18804144 | 0,44811698 |
| 159963_at | SLC5A12 | 0,09783894 | 3,38329863 | 1,38131094 | 0,18806382 | 0,44811698 |
| 8612_at | PPAP2C | 0,22390653 | 9,23082956 | 1,38123132 | 0,18808778 | 0,44811698 |
| 100507584_a | LINC01016 | -0,12748719 | 4,33098515 | -1,38117282 | 0,18810538 | 0,44811698 |
| 25793_at | FBXO7 | 0,12320911 | 10,199341 | 1,3810786 | 0,18813374 | 0,44811976 |
| 57533_at | TBC1D14 | -0,16992242 | 8,56724898 | -1,38100877 | 0,18815475 | 0,44811976 |
| 965_at | CD58 | -0,17387656 | 7,25651941 | -1,38093708 | 0,18817633 | 0,44811976 |
| 64837_at | KLC2 | -0,11114548 | 7,9990325 | -1,38050666 | 0,18830594 | 0,4483538 |
| 2842_at | GPR19 | -0,34038325 | 5,21299437 | -1,3804561 | 0,18832117 | 0,4483538 |
| 3250_at | HPR | -0,12466061 | 3,90591345 | -1,38034648 | 0,18835419 | 0,44837701 |
| 9162_at | DGKI | -0,14925684 | 3,69211532 | -1,3802345 | 0,18838793 | 0,4484019 |
| 955_at | ENTPD6 | 0,07889444 | 8,26692605 | 1,38015205 | 0,18841277 | 0,44840563 |

| | | | | | | |
|---|---|---|---|---|---|---|
| 4772_at | NFATC1 | -0,24097094 | 5,29272384 | -1,37995241 | 0,18847293 | 0,44849341 |
| 260429_at | PRSS33 | -0,15765544 | 5,47805791 | -1,37970723 | 0,18854685 | 0,44861387 |
| 5883_at | RAD9A | -0,12660312 | 8,30006879 | -1,3794918 | 0,18861181 | 0,4487093 |
| 101927274_a | LOC10192727 | -0,10809489 | 4,61290644 | -1,37941975 | 0,18863354 | 0,4487093 |
| 84639_at | IL1F10 | -0,182478 | 4,5584792 | -1,37913779 | 0,1887186 | 0,4488284 |
| 100507477_a | LOC10050747 | -0,10879919 | 3,11134448 | -1,37909932 | 0,18873021 | 0,4488284 |
| 8645_at | KCNK5 | -0,40210072 | 7,06968057 | -1,3789889 | 0,18876353 | 0,44885223 |
| 339505_at | LINC01141 | -0,20406925 | 4,51169972 | -1,37880051 | 0,18882039 | 0,44893202 |
| 2081_at | ERN1 | -0,09909831 | 5,84634582 | -1,37857063 | 0,1888898 | 0,44904161 |
| 10380_at | BPNT1 | 0,21089948 | 7,19044091 | 1,37837144 | 0,18894995 | 0,44908071 |
| 497258_at | BDNF-AS | 0,0727571 | 3,18686637 | 1,37829627 | 0,18897265 | 0,44908071 |
| 140460_at | ASB7 | 0,07889343 | 6,82682293 | 1,37828458 | 0,18897619 | 0,44908071 |
| 4298_at | MLLT1 | -0,09694851 | 8,24330928 | -1,37796577 | 0,18907251 | 0,44923255 |
| 54932_at | EXD3 | 0,08541175 | 5,65996638 | 1,37791874 | 0,18908672 | 0,44923255 |
| 100131825_a | CADM3-AS1 | -0,2189594 | 4,96270197 | -1,37774097 | 0,18914045 | 0,44930479 |
| 6014_at | RIT2 | -0,10828531 | 3,30933871 | -1,37742689 | 0,18923541 | 0,44942117 |
| 79747_at | ADGB | -0,13743034 | 3,74477951 | -1,37742459 | 0,18923611 | 0,44942117 |
| 221756_at | SERPINB9P1 | -0,18883551 | 4,76843223 | -1,3771144 | 0,18932993 | 0,44955868 |
| 285889_at | LOC285889 | -0,13475906 | 3,46379347 | -1,37707886 | 0,18934068 | 0,44955868 |
| 5493_at | PPL | 0,40642854 | 9,13151956 | 1,37696697 | 0,18937454 | 0,44958364 |
| 400617_at | KCNJ2-AS1 | -0,06735084 | 3,01988714 | -1,37684031 | 0,18941287 | 0,44961923 |
| 9892_at | SNAP91 | 0,09347243 | 3,8060767 | 1,37661521 | 0,189481 | 0,44968047 |
| 1244_at | ABCC2 | -0,13960457 | 4,342207 | -1,37660082 | 0,18948535 | 0,44968047 |
| 10781_at | ZNF266 | 0,23252718 | 8,18262169 | 1,376053 | 0,18965126 | 0,44999286 |
| 140680_at | C20orf96 | -0,12779042 | 4,2086976 | -1,37593937 | 0,18968568 | 0,44999286 |
| 8853_at | ASAP2 | -0,26413633 | 9,57374974 | -1,37586875 | 0,18970708 | 0,44999286 |
| 23644_at | EDC4 | -0,11140125 | 7,87262772 | -1,3758577 | 0,18971043 | 0,44999286 |
| 146562_at | C16orf71 | -0,13378587 | 5,46149304 | -1,37488757 | 0,19000459 | 0,45063511 |
| 8856_at | NR1I2 | -0,11397189 | 4,67675483 | -1,37465632 | 0,19007476 | 0,45074605 |
| 5134_at | PDCD2 | -0,09685685 | 7,69014438 | -1,37452692 | 0,19011404 | 0,45074612 |
| 23682_at | RAB38 | 0,18562015 | 4,62264358 | 1,37450204 | 0,19012159 | 0,45074612 |
| 100129792_a | CCDC152 | -0,11923436 | 2,90060187 | -1,3744091 | 0,1901498 | 0,45075753 |
| 79848_at | CSPP1 | -0,20854608 | 6,69692445 | -1,37415127 | 0,19022809 | 0,45081183 |
| 6290_at | SAA3P | -0,11442823 | 4,19057087 | -1,37413791 | 0,19023215 | 0,45081183 |
| 613_at | BCR | -0,09406459 | 8,06318237 | -1,37397065 | 0,19028295 | 0,45081183 |
| 158158_at | RASEF | 0,13453737 | 6,23281388 | 1,37392981 | 0,19029536 | 0,45081183 |
| 101926888_a | RALY-AS1 | -0,15234389 | 6,97160841 | -1,37378624 | 0,19033898 | 0,45081183 |
| 2949_at | GSTM5 | -0,10095662 | 5,07318455 | -1,3737584 | 0,19034744 | 0,45081183 |
| 66036_at | MTMR9 | 0,21220412 | 7,7265912 | 1,37372569 | 0,19035738 | 0,45081183 |
| 10529_at | NEBL | 0,36874143 | 5,5729592 | 1,37371727 | 0,19035993 | 0,45081183 |
| 3977_at | LIFR | 0,20044376 | 3,84923823 | 1,37342275 | 0,19044945 | 0,45096838 |
| 84622_at | ZNF594 | 0,17766179 | 4,52957419 | 1,37291976 | 0,1906024 | 0,45127508 |
| 11095_at | ADAMTS8 | -0,111409 | 4,72367363 | -1,37277175 | 0,19064743 | 0,45132621 |
| 84284_at | NTPCR | -0,10493328 | 8,30596559 | -1,37234015 | 0,19077877 | 0,45154901 |
| 10747_at | MASP2 | -0,13600352 | 4,48134526 | -1,37230845 | 0,19078842 | 0,45154901 |
| 1840_at | DTX1 | -0,12647829 | 5,57400316 | -1,37203883 | 0,19087052 | 0,45161716 |
| 90987_at | ZNF251 | 0,31421776 | 6,71407227 | 1,37201537 | 0,19087766 | 0,45161716 |
| 100287387_a | LOC10028738 | 0,26682265 | 5,53459449 | 1,37195473 | 0,19089613 | 0,45161716 |
| 79873_at | NUDT18 | 0,36415223 | 6,93540636 | 1,37190592 | 0,190911 | 0,45161716 |
| 54617_at | INO80 | -0,09048498 | 7,63753842 | -1,3717189 | 0,19096797 | 0,45169646 |

| | | | | | | |
|---|---|---|---|---|---|---|
| 1896_at | EDA | -0,08152355 | 4,02031902 | -1,37138246 | 0,19107049 | 0,45186302 |
| 731789_at | LINC00202-2 | -0,14167086 | 4,72911986 | -1,37128794 | 0,1910993 | 0,45186302 |
| 4650_at | MYO9B | 0,09508125 | 7,12879919 | 1,3712484 | 0,19111135 | 0,45186302 |
| 126549_at | ANKLE1 | -0,16232615 | 4,4910742 | -1,37117996 | 0,19113222 | 0,45186302 |
| 6722_at | SRF | -0,07918652 | 7,48894873 | -1,37091025 | 0,19121446 | 0,45194647 |
| 7180_at | CRISP2 | 0,23606544 | 3,47224255 | 1,37086141 | 0,19122936 | 0,45194647 |
| 8724_at | SNX3 | 0,12687536 | 10,8179366 | 1,37083339 | 0,1912379 | 0,45194647 |
| 11133_at | KPTN | -0,12530411 | 7,34021065 | -1,37075184 | 0,19126278 | 0,45194981 |
| 125488_at | TTC39C | -0,20790685 | 6,58041896 | -1,37053046 | 0,19133032 | 0,45205395 |
| 10617_at | STAMBP | -0,16742651 | 8,11166978 | -1,37034434 | 0,19138711 | 0,45213269 |
| 4300_at | MLLT3 | 0,42502552 | 7,29045567 | 1,37012493 | 0,19145409 | 0,45220101 |
| 3682_at | ITGAE | -0,29773167 | 8,15212163 | -1,3700958 | 0,19146298 | 0,45220101 |
| 7369_at | UMOD | -0,15481963 | 4,79111559 | -1,36997694 | 0,19149927 | 0,45223127 |
| 126820_at | WDR63 | 0,17833629 | 3,29060075 | 1,36979761 | 0,19155403 | 0,45230515 |
| 4946_at | OAZ1 | -0,07554471 | 12,6188869 | -1,36956238 | 0,19162589 | 0,45241936 |
| 51060_at | TXNDC12 | -0,22217349 | 10,4685777 | -1,36937637 | 0,19168272 | 0,45249808 |
| 55905_at | RNF114 | 0,14228576 | 9,34471295 | 1,36924891 | 0,19172168 | 0,45253458 |
| 100507602_a | TRIM52-AS1 | -0,08821575 | 6,48613628 | -1,36916434 | 0,19174753 | 0,45254014 |
| 8790_at | FPGT | 0,25446488 | 5,23262443 | 1,36899881 | 0,19179813 | 0,45260411 |
| 79864_at | C11orf63 | 0,27849177 | 5,5868767 | 1,3682106 | 0,19203924 | 0,45311756 |
| 11075_at | STMN2 | -0,10016451 | 3,80413771 | -1,36804332 | 0,19209044 | 0,45318287 |
| 4917_at | NTN3 | -0,19498613 | 5,40155781 | -1,36764263 | 0,19221313 | 0,45341679 |
| 283685_at | GOLGA6L2 | -0,12527972 | 3,59990468 | -1,36747364 | 0,19226489 | 0,45348336 |
| 53632_at | PRKAG3 | -0,10811044 | 5,05105477 | -1,36736964 | 0,19229675 | 0,45350297 |
| 84894_at | LINGO1 | -0,12399354 | 4,87533608 | -1,36681025 | 0,1924682 | 0,45381772 |
| 80219_at | COQ10B | 0,14474115 | 8,54174763 | 1,36673408 | 0,19249155 | 0,45381772 |
| 339559_at | ZFP69 | -0,13432928 | 5,51924805 | -1,36652474 | 0,19255575 | 0,45381772 |
| 4669_at | NAGLU | 0,16391652 | 7,3225056 | 1,36651557 | 0,19255857 | 0,45381772 |
| 27231_at | NMRK2 | -0,15077462 | 5,5362092 | -1,36630638 | 0,19262274 | 0,45381772 |
| 252983_at | STXBP4 | 0,22271064 | 5,25683677 | 1,36629326 | 0,19262676 | 0,45381772 |
| 55808_at | ST6GALNAC1 | 0,50713458 | 5,47963294 | 1,36629053 | 0,1926276 | 0,45381772 |
| 312_at | ANXA13 | -0,15499532 | 3,59788901 | -1,36628767 | 0,19262848 | 0,45381772 |
| 389289_at | ANXA2R | -0,13200363 | 4,92313403 | -1,36624214 | 0,19264245 | 0,45381772 |
| 84717_at | HDGFRP2 | -0,09783087 | 8,04873534 | -1,36616604 | 0,1926658 | 0,45381772 |
| 123920_at | CMTM3 | 0,2086413 | 7,83764615 | 1,36594689 | 0,19273306 | 0,45392063 |
| 10993_at | SDS | -0,14191015 | 5,77220273 | -1,36544968 | 0,19288572 | 0,45422465 |
| 3717_at | JAK2 | 0,23852875 | 5,99446811 | 1,36524358 | 0,19294903 | 0,45430215 |
| 85415_at | RHPN2 | 0,435751 | 8,94778277 | 1,36518901 | 0,1929658 | 0,45430215 |
| 27102_at | EIF2AK1 | 0,15182476 | 9,23137417 | 1,36505563 | 0,19300678 | 0,45434311 |
| 972_at | CD74 | 0,19001837 | 5,75886313 | 1,36489842 | 0,1930551 | 0,45440132 |
| 7991_at | TUSC3 | 0,4149616 | 8,90659558 | 1,36453071 | 0,19316815 | 0,45458719 |
| 10590_at | SCGN | -0,12121638 | 4,27186825 | -1,36448804 | 0,19318127 | 0,45458719 |
| 79363_at | RSG1 | -0,15983832 | 5,60464624 | -1,36407406 | 0,19330861 | 0,45483129 |
| 2531_at | KDSR | 0,08910684 | 6,67847993 | 1,36384497 | 0,19337911 | 0,45491968 |
| 154197_at | PNLDC1 | -0,15796395 | 3,77960055 | -1,36374442 | 0,19341006 | 0,45491968 |
| 647288_at | CTAGE11P | -0,13119176 | 4,20343245 | -1,36367309 | 0,19343202 | 0,45491968 |
| 9617_at | MTRF1 | 0,14196081 | 5,8623758 | 1,36358575 | 0,19345891 | 0,45491968 |
| 670_at | BPHL | 0,15453627 | 7,75013618 | 1,36356837 | 0,19346426 | 0,45491968 |
| 3083_at | HGFAC | -0,14075834 | 5,00194603 | -1,36316518 | 0,19358843 | 0,4551561 |
| 646982_at | LINC00598 | -0,08455628 | 3,73148047 | -1,36307641 | 0,19361578 | 0,45516472 |

| | | | | | | |
|---|---|---|---|---|---|---|
| 374403_at | TBC1D10C | -0,16022014 | 5,83091936 | -1,36299989 | 0,19363935 | 0,45516472 |
| 1849_at | DUSP7 | -0,42365251 | 7,93079685 | -1,36265899 | 0,19374441 | 0,45535217 |
| 27289_at | RND1 | 0,20662235 | 5,5849682 | 1,36258773 | 0,19376638 | 0,45535217 |
| 79148_at | MMP28 | -0,33489235 | 7,22786761 | -1,36227449 | 0,19386297 | 0,45539553 |
| 881_at | CCIN | -0,13726203 | 4,45882593 | -1,36226232 | 0,19386672 | 0,45539553 |
| 8934_at | RAB29 | -0,15295148 | 6,95154665 | -1,36224403 | 0,19387236 | 0,45539553 |
| 286183_at | NKAIN3 | -0,14334701 | 3,80908903 | -1,36222123 | 0,19387939 | 0,45539553 |
| 4708_at | NDUFB2 | -0,08409835 | 10,4249954 | -1,3620114 | 0,19394412 | 0,45546491 |
| 2145_at | EZH1 | 0,25460271 | 6,91238608 | 1,36197218 | 0,19395622 | 0,45546491 |
| 84188_at | FAR1 | -0,34949272 | 8,59684602 | -1,36173786 | 0,19402852 | 0,45557916 |
| 2932_at | GSK3B | 0,13035617 | 7,81459867 | 1,36152469 | 0,19409432 | 0,45564035 |
| 26578_at | OSTF1 | 0,16582634 | 10,360651 | 1,36150017 | 0,19410189 | 0,45564035 |
| 57084_at | SLC17A6 | -0,11666162 | 3,5006062 | -1,3613427 | 0,19415051 | 0,45569894 |
| 266919_at | LINC00307 | -0,10630246 | 2,86254929 | -1,36103618 | 0,19424517 | 0,45586559 |
| 102724699_a | LOC10272469 | -0,24495124 | 6,84598356 | -1,36067943 | 0,1943554 | 0,45606849 |
| 144776_at | LINC00410 | -0,1372757 | 3,28671824 | -1,36060313 | 0,19437898 | 0,45606849 |
| 284837_at | AATBC | 0,8077717 | 7,1271323 | 1,36051463 | 0,19440633 | 0,45607712 |
| 25873_at | RPL36 | -0,16214003 | 11,466183 | -1,35981287 | 0,19462335 | 0,45653064 |
| 3858_at | KRT10 | -0,12697638 | 9,54267765 | -1,35963698 | 0,19467778 | 0,45655577 |
| 169792_at | GLIS3 | 0,12309274 | 4,96710699 | 1,35962505 | 0,19468147 | 0,45655577 |
| 26022_at | TMEM98 | 0,39015784 | 6,3016989 | 1,35934592 | 0,19476786 | 0,45670278 |
| 145447_at | ABHD12B | -0,11003547 | 2,99333513 | -1,35915442 | 0,19482716 | 0,45678621 |
| 83876_at | MRO | -0,09636858 | 3,49668041 | -1,35890196 | 0,19490534 | 0,45691391 |
| 3820_at | KLRB1 | -0,238932 | 3,86438481 | -1,35857595 | 0,19500634 | 0,45694301 |
| 100996485_a | C5orf66 | -0,19737996 | 5,87667931 | -1,35857188 | 0,19500761 | 0,45694301 |
| 100289678_a | ZNF783 | 0,10892523 | 7,11496222 | 1,35852418 | 0,19502239 | 0,45694301 |
| 80128_at | TRIM46 | -0,18563632 | 5,32177434 | -1,35847774 | 0,19503678 | 0,45694301 |
| 10085_at | EDIL3 | 0,27505847 | 6,7055769 | 1,3584142 | 0,19505647 | 0,45694301 |
| 4261_at | CIITA | -0,13493804 | 5,08165786 | -1,35840255 | 0,19506008 | 0,45694301 |
| 54899_at | PXK | 0,24390625 | 5,4280915 | 1,3582923 | 0,19509426 | 0,4569675 |
| 573_at | BAG1 | 0,20577899 | 8,23584521 | 1,35789835 | 0,19521641 | 0,45714088 |
| 148103_at | ZNF599 | 0,06380008 | 3,44056122 | 1,3578809 | 0,19522182 | 0,45714088 |
| 10458_at | BAIAP2 | -0,11374133 | 6,85140302 | -1,35781615 | 0,19524191 | 0,45714088 |
| 94233_at | OPN4 | -0,16367973 | 5,3328895 | -1,35774749 | 0,19526321 | 0,45714088 |
| 26164_at | MTG2 | -0,10677791 | 7,75838229 | -1,35753427 | 0,19532936 | 0,45724018 |
| 7541_at | ZBTB14 | 0,1087725 | 6,86455716 | 1,35740608 | 0,19536914 | 0,45727773 |
| 5939_at | RBMS2 | 0,22895996 | 6,9826586 | 1,35723876 | 0,19542107 | 0,45733407 |
| 64240_at | ABCG5 | -0,10652092 | 3,6705559 | -1,35717556 | 0,19544069 | 0,45733407 |
| 254158_at | CXorf58 | -0,10451277 | 3,29966376 | -1,35708026 | 0,19547028 | 0,45734774 |
| 54067_at | C21orf62-AS1 | 0,16331631 | 3,91416885 | 1,35689387 | 0,19552815 | 0,4574276 |
| 89872_at | AQP10 | 0,13904167 | 4,04082479 | 1,35643272 | 0,19567141 | 0,45756126 |
| 81572_at | PDRG1 | -0,2178507 | 8,5015482 | -1,35641169 | 0,19567794 | 0,45756126 |
| 72_at | ACTG2 | 0,25421508 | 5,00108707 | 1,35633243 | 0,19570257 | 0,45756126 |
| 118788_at | PIK3AP1 | -0,08768263 | 4,19313006 | -1,35631423 | 0,19570823 | 0,45756126 |
| 64359_at | NXN | -0,274788 | 9,02523372 | -1,35625771 | 0,19572579 | 0,45756126 |
| 27253_at | PCDH17 | 0,10118713 | 3,99028646 | 1,35609533 | 0,19577627 | 0,45756126 |
| 169611_at | OLFML2A | 0,75804695 | 7,79551696 | 1,35603891 | 0,19579381 | 0,45756126 |
| 5091_at | PC | -0,16039259 | 6,56408118 | -1,35603418 | 0,19579528 | 0,45756126 |
| 1081_at | CGA | -0,18453842 | 3,68246513 | -1,35593649 | 0,19582565 | 0,45756126 |
| 5738_at | PTGFRN | -0,18962801 | 8,78374035 | -1,35593445 | 0,19582629 | 0,45756126 |

| | | | | | | |
|---|---|---|---|---|---|---|
| 9939_at | RBM8A | -0,16777883 | 8,29432093 | -1,35583121 | 0,19585839 | 0,45756126 |
| 594855_at | CPLX3 | -0,12624326 | 4,28069532 | -1,35572084 | 0,19589272 | 0,45756126 |
| 1605_at | DAG1 | -0,14018367 | 8,01001698 | -1,35571645 | 0,19589408 | 0,45756126 |
| 924_at | CD7 | -0,12909437 | 5,36804645 | -1,35508772 | 0,19608971 | 0,45796267 |
| 90827_at | ZNF479 | 0,15974654 | 3,33669324 | 1,3548842 | 0,19615307 | 0,45805511 |
| 6898_at | TAT | -0,08562528 | 3,94729716 | -1,35478663 | 0,19618345 | 0,45807052 |
| 80313_at | LRRC27 | 0,11356392 | 5,02354776 | 1,35464473 | 0,19622764 | 0,45811818 |
| 51146_at | A4GNT | -0,14865393 | 4,21433208 | -1,3542965 | 0,19633613 | 0,4583159 |
| 4318_at | MMP9 | -0,15944952 | 5,89016274 | -1,35401001 | 0,19642541 | 0,45846876 |
| 101928460_a | LOC10192846 | -0,09755378 | 3,65623829 | -1,35383852 | 0,19647887 | 0,45851364 |
| 60343_at | FAM3A | 0,12278504 | 7,84402404 | 1,35379563 | 0,19649224 | 0,45851364 |
| 4015_at | LOX | 0,40124985 | 4,93751356 | 1,35365793 | 0,19653518 | 0,45855828 |
| 283856_at | LOC283856 | -0,16606335 | 4,55136158 | -1,35357158 | 0,19656211 | 0,45856557 |
| 4582_at | MUC1 | 0,52045663 | 7,1028642 | 1,35341937 | 0,19660959 | 0,45862079 |
| 64419_at | MTMR14 | 0,14341991 | 7,84069176 | 1,35325519 | 0,19666081 | 0,45868472 |
| 84645_at | C22orf23 | -0,11661955 | 5,40331525 | -1,35312112 | 0,19670264 | 0,45868659 |
| 26338_at | OR5L2 | -0,23232236 | 3,79321496 | -1,3531 | 0,19670923 | 0,45868659 |
| 6750_at | SST | 0,31610885 | 4,52582663 | 1,35293014 | 0,19676224 | 0,458726 |
| 26070_at | DKFZP434K02 | -0,12837579 | 5,31711813 | -1,35289326 | 0,19677376 | 0,458726 |
| 100505879_a | KIF25-AS1 | -0,11114747 | 4,55102031 | -1,35271009 | 0,19683094 | 0,45875044 |
| 254439_at | C11orf86 | -0,15266799 | 5,80052879 | -1,35263944 | 0,196853 | 0,45875044 |
| 153020_at | RASGEF1B | -0,09397283 | 3,2270327 | -1,35263083 | 0,19685569 | 0,45875044 |
| 22999_at | RIMS1 | -0,14775142 | 3,77243642 | -1,3523637 | 0,19693912 | 0,45888934 |
| 378825_at | LINC00162 | -0,15807094 | 5,10799399 | -1,35204181 | 0,19703969 | 0,45906815 |
| 5128_at | CDK17 | 0,11026544 | 7,00565125 | 1,35185491 | 0,1970981 | 0,45912073 |
| 26251_at | KCNG2 | -0,16628218 | 5,90516466 | -1,35181709 | 0,19710992 | 0,45912073 |
| 1432_at | MAPK14 | -0,16722622 | 7,74581615 | -1,35166714 | 0,1971568 | 0,45917439 |
| 4826_at | NNAT | -0,15179561 | 4,36082928 | -1,35146513 | 0,19721996 | 0,45926598 |
| 433_at | ASGR2 | -0,12177632 | 4,82511133 | -1,35132947 | 0,19726239 | 0,45930925 |
| 7781_at | SLC30A3 | -0,19427607 | 6,83341132 | -1,3511333 | 0,19732376 | 0,45939661 |
| 256957_at | HEATR9 | -0,11803954 | 3,54668991 | -1,35072822 | 0,19745052 | 0,45956727 |
| 10917_at | BTNL3 | -0,15740166 | 5,47381031 | -1,35069495 | 0,19746094 | 0,45956727 |
| 285456_at | RPL34-AS1 | 0,11434353 | 3,13807309 | 1,35060649 | 0,19748863 | 0,45956727 |
| 101928618_a | LOC10192861 | -0,11607943 | 3,31788682 | -1,35059416 | 0,19749249 | 0,45956727 |
| 117286_at | CIB3 | -0,15919858 | 4,91072259 | -1,35013725 | 0,19763558 | 0,45984469 |
| 8516_at | ITGA8 | -0,1023894 | 3,62436069 | -1,349779 | 0,19774783 | 0,45991453 |
| 50810_at | HDGFRP3 | -0,16741607 | 8,17843293 | -1,34975854 | 0,19775424 | 0,45991453 |
| 3670_at | ISL1 | -0,1313257 | 3,63248188 | -1,34972352 | 0,19776522 | 0,45991453 |
| 100506100_a | LOC10050610 | 0,14089231 | 6,9610305 | 1,34966363 | 0,19778399 | 0,45991453 |
| 1007_at | CDH9 | -0,10148936 | 3,33079504 | -1,3496605 | 0,19778497 | 0,45991453 |
| 132612_at | ADAD1 | -0,14620295 | 3,44582956 | -1,34953467 | 0,19782442 | 0,45995073 |
| 415_at | ARSE | 0,20931565 | 4,84538001 | 1,34944026 | 0,19785402 | 0,45996403 |
| 284018_at | C17orf58 | -0,17221626 | 7,86152077 | -1,34916717 | 0,19793966 | 0,46004558 |
| 170370_at | FAM170B | -0,14776524 | 4,56362076 | -1,34916436 | 0,19794054 | 0,46004558 |
| 121601_at | ANO4 | -0,10658848 | 3,62347558 | -1,34903768 | 0,19798028 | 0,46004558 |
| 200504_at | GKN2 | -0,12977536 | 3,43393821 | -1,34899081 | 0,19799498 | 0,46004558 |
| 4626_at | MYH8 | -0,09225806 | 3,08835338 | -1,34894007 | 0,1980109 | 0,46004558 |
| 2488_at | FSHB | -0,14510774 | 3,48172978 | -1,34878488 | 0,19805959 | 0,46004558 |
| 25758_at | KIAA1549L | -0,12684467 | 4,14491356 | -1,34874354 | 0,19807256 | 0,46004558 |
| 8352_at | HIST1H3C | -0,12329999 | 7,19421183 | -1,34866499 | 0,19809722 | 0,46004558 |

| | | | | | | |
|---|---|---|---|---|---|---|
| 3726_at | JUNB | 0,31463418 | 9,16239116 | 1,34864326 | 0,19810404 | 0,46004558 |
| 83937_at | RASSF4 | 0,08308599 | 4,55289529 | 1,34840697 | 0,19817821 | 0,46016235 |
| 381_at | ARF5 | 0,14391306 | 9,78928718 | 1,34809 | 0,19827774 | 0,46028539 |
| 51294_at | PCDH12 | -0,13713221 | 5,83484548 | -1,34808605 | 0,19827899 | 0,46028539 |
| 6511_at | SLC1A6 | -1,32633359 | 5,24908021 | -1,34799023 | 0,19830908 | 0,46029978 |
| 3834_at | KIF25 | -0,1303081 | 5,4790648 | -1,34782775 | 0,19836013 | 0,46031609 |
| 1133_at | CHRM5 | -0,09630441 | 3,30513625 | -1,3477467 | 0,19838559 | 0,46031609 |
| 64714_at | PDIA2 | -0,17139302 | 5,54095114 | -1,34773969 | 0,1983878 | 0,46031609 |
| 54805_at | CNNM2 | 0,08673457 | 5,58159328 | 1,34745398 | 0,19847759 | 0,46046897 |
| 54457_at | TAF7L | -0,10607746 | 3,21641505 | -1,34733004 | 0,19851655 | 0,4605039 |
| 64110_at | MAGEF1 | -0,15338546 | 9,94188451 | -1,34717026 | 0,19856679 | 0,46055738 |
| 55197_at | RPRD1A | -0,14875077 | 7,95616554 | -1,34705662 | 0,19860253 | 0,46055738 |
| 101927948_a | LOC10192794 | -0,18206368 | 3,93432056 | -1,34702861 | 0,19861134 | 0,46055738 |
| 54910_at | SEMA4C | 0,20671068 | 8,58446347 | 1,34687757 | 0,19865885 | 0,4606121 |
| 9191_at | DEDD | 0,14117285 | 7,95610846 | 1,346621 | 0,19873957 | 0,46065414 |
| 26229_at | B3GAT3 | 0,11148511 | 6,82310423 | 1,34659752 | 0,19874696 | 0,46065414 |
| 79602_at | ADIPOR2 | -0,15801683 | 9,033354 | -1,34659193 | 0,19874872 | 0,46065414 |
| 1070_at | CETN3 | -0,14864159 | 7,58642399 | -1,34632938 | 0,19883135 | 0,46079022 |
| 401262_at | CRIP3 | -0,1582708 | 5,13747009 | -1,34613754 | 0,19889175 | 0,46087474 |
| 79973_at | ZNF442 | 0,24675692 | 4,06121246 | 1,34601104 | 0,19893158 | 0,4609116 |
| 100131434_a | LINC00893 | 0,18108376 | 3,406625 | 1,34582849 | 0,19898908 | 0,46098937 |
| 64150_at | DIO3OS | -0,15115547 | 3,87207309 | -1,34537206 | 0,19913289 | 0,46123043 |
| 22827_at | PUF60 | -0,37321445 | 9,62174051 | -1,34534627 | 0,19914102 | 0,46123043 |
| 57150_at | SMIM8 | -0,11951172 | 5,60102901 | -1,34523183 | 0,19917709 | 0,46125852 |
| 100132354_a | LINC01512 | -0,18645331 | 4,77548909 | -1,34500402 | 0,19924892 | 0,46135816 |
| 3134_at | HLA-F | 0,61362072 | 8,24011476 | 1,34494343 | 0,19926802 | 0,46135816 |
| 4542_at | MYO1F | 0,19742371 | 5,75341639 | 1,34455389 | 0,19939089 | 0,46154361 |
| 340205_at | TREML1 | -0,17189607 | 5,37622638 | -1,34439812 | 0,19944005 | 0,46154361 |
| 115992_at | RNF166 | -0,1415208 | 6,77194764 | -1,34434262 | 0,19945756 | 0,46154361 |
| 644656_at | LOC644656 | -0,19079925 | 5,27566011 | -1,34432466 | 0,19946323 | 0,46154361 |
| 118442_at | GPR62 | -0,19182828 | 5,69690974 | -1,34417272 | 0,19951119 | 0,46154361 |
| 8683_at | SRSF9 | -0,13359888 | 11,2072872 | -1,34414393 | 0,19952028 | 0,46154361 |
| 9776_at | ATG13 | 0,21282006 | 7,33947444 | 1,34413723 | 0,19952239 | 0,46154361 |
| 115677_at | NOSTRIN | -0,09857394 | 3,43573998 | -1,34408208 | 0,1995398 | 0,46154361 |
| 442524_at | DPY19L2P3 | -0,17214442 | 6,13275203 | -1,34395261 | 0,19958068 | 0,46157343 |
| 100127888_a | SLCO4A1-AS1 | -0,13542292 | 5,53826844 | -1,343869 | 0,19960708 | 0,46157343 |
| 54477_at | PLEKHA5 | 0,15290333 | 6,90876374 | 1,34376379 | 0,19964031 | 0,46157343 |
| 645745_at | MT1HL1 | -0,0920002 | 6,23369851 | -1,34373771 | 0,19964854 | 0,46157343 |
| 6928_at | HNF1B | -0,10629267 | 5,74407532 | -1,34360826 | 0,19968943 | 0,46161256 |
| 4684_at | NCAM1 | -0,11682238 | 4,34513933 | -1,34330207 | 0,19978617 | 0,46178078 |
| 7988_at | ZNF212 | 0,07818823 | 7,38349393 | 1,3430469 | 0,19986682 | 0,46187897 |
| 6373_at | CXCL11 | 0,71979715 | 3,33210458 | 1,34301594 | 0,19987661 | 0,46187897 |
| 284365_at | MGC45922 | -0,14276524 | 5,34855242 | -1,34267837 | 0,19998335 | 0,46201484 |
| 2886_at | GRB7 | 0,16115916 | 7,32113281 | 1,34264516 | 0,19999385 | 0,46201484 |
| 144348_at | ZNF664 | 0,11113982 | 10,2704257 | 1,34260245 | 0,20000736 | 0,46201484 |
| 10664_at | CTCF | -0,14714187 | 9,15580084 | -1,34234434 | 0,20008901 | 0,46214803 |
| 529_at | ATP6V1E1 | 0,09639499 | 10,4513923 | 1,34218163 | 0,2001405 | 0,46221153 |
| 81693_at | AMN | -0,149077 | 5,48664971 | -1,34197294 | 0,20020655 | 0,46230864 |
| 147948_at | ZNF582 | 0,12859019 | 5,31848697 | 1,34183024 | 0,20025172 | 0,4623219 |
| 2852_at | GPER1 | -0,12773614 | 4,71795968 | -1,34180318 | 0,20026029 | 0,4623219 |

| | | | | | | |
|---|---|---|---|---|---|---|
| 8534_at | CHST1 | -0,11924613 | 5,47284364 | -1,34165563 | 0,20030701 | 0,46237435 |
| 54949_at | SDHAF2 | 0,12423372 | 8,36633453 | 1,34157712 | 0,20033188 | 0,46237634 |
| 645355_at | MIR3976HG | -0,10798689 | 3,34436106 | -1,34110684 | 0,20048086 | 0,46261175 |
| 23613_at | ZMYND8 | 0,18245179 | 9,35565419 | 1,34110355 | 0,2004819 | 0,46261175 |
| 130535_at | KCTD18 | 0,26350973 | 7,97940523 | 1,34091318 | 0,20054224 | 0,46264239 |
| 889_at | KRIT1 | 0,15843347 | 7,39507504 | 1,34091011 | 0,20054321 | 0,46264239 |
| 59067_at | IL21 | -0,15387707 | 3,75446163 | -1,34056598 | 0,20065232 | 0,4627874 |
| 23428_at | SLC7A8 | -0,29506294 | 7,86941706 | -1,34055819 | 0,20065479 | 0,4627874 |
| 54959_at | ODAM | -0,13938329 | 4,41003305 | -1,34048454 | 0,20067815 | 0,4627874 |
| 29948_at | OSGIN1 | -0,24029577 | 7,22194436 | -1,33997176 | 0,20084083 | 0,46310712 |
| 171177_at | RHOV | 0,09783469 | 7,47356981 | 1,33987242 | 0,20087236 | 0,46312438 |
| 23315_at | SLC9A8 | 0,23663164 | 7,3635499 | 1,339497 | 0,20099154 | 0,46328879 |
| 554_at | AVPR2 | -0,09788229 | 4,79842431 | -1,33935408 | 0,20103693 | 0,46328879 |
| 22992_at | KDM2A | 0,07803684 | 8,54415515 | 1,33934408 | 0,20104011 | 0,46328879 |
| 401261_at | LOC401261 | 0,40270601 | 5,31165446 | 1,33924088 | 0,20107289 | 0,46328879 |
| 27152_at | INTU | 0,26163647 | 4,76857693 | 1,33921149 | 0,20108222 | 0,46328879 |
| 2690_at | GHR | 0,12553509 | 2,96006659 | 1,3391934 | 0,20108797 | 0,46328879 |
| 6528_at | SLC5A5 | -0,15957621 | 5,70486968 | -1,33902877 | 0,20114027 | 0,46331635 |
| 57484_at | RNF150 | 0,09368355 | 4,4337891 | 1,33900434 | 0,20114803 | 0,46331635 |
| 5193_at | PEX12 | 0,17098637 | 7,85479934 | 1,33882484 | 0,20120508 | 0,46331968 |
| 729085_at | FAM198A | 0,10012477 | 3,79976125 | 1,33879724 | 0,20121385 | 0,46331968 |
| 157381_at | LINC00964 | 0,20227788 | 3,3953407 | 1,33877274 | 0,20122164 | 0,46331968 |
| 8288_at | EPX | -0,1394142 | 5,15256075 | -1,33825225 | 0,20138713 | 0,4635716 |
| 151835_at | CPNE9 | -0,11955757 | 4,73977966 | -1,33817703 | 0,20141105 | 0,4635716 |
| 101928323_a | LOC10192832 | -0,15278609 | 4,6926113 | -1,33815157 | 0,20141915 | 0,4635716 |
| 56890_at | MDM1 | -0,11737324 | 5,86712658 | -1,33812592 | 0,20142731 | 0,4635716 |
| 57231_at | SNX14 | 0,22013367 | 8,80347011 | 1,33798294 | 0,2014728 | 0,46362089 |
| 145264_at | SERPINA12 | -0,12635856 | 3,67638266 | -1,33780992 | 0,20152785 | 0,46365424 |
| 100422737_a | LOC10042273 | -0,18285806 | 4,24762192 | -1,33778611 | 0,20153543 | 0,46365424 |
| 100506710_a | EBLN3 | 0,17904253 | 8,26300125 | 1,33743786 | 0,20164628 | 0,46385387 |
| 54442_at | KCTD5 | -0,1358686 | 9,18522575 | -1,33710133 | 0,20175345 | 0,46399052 |
| 101929645_a | LOC10192964 | -0,10544304 | 2,91206092 | -1,33708982 | 0,20175712 | 0,46399052 |
| 7332_at | UBE2L3 | -0,11648715 | 8,87609889 | -1,33702442 | 0,20177795 | 0,46399052 |
| 10804_at | GJB6 | -0,49500819 | 7,87619018 | -1,33669822 | 0,20188188 | 0,46412578 |
| 4058_at | LTK | -0,11225156 | 4,63252158 | -1,33668725 | 0,20188538 | 0,46412578 |
| 54868_at | TMEM104 | -0,10561079 | 6,945848 | -1,33661297 | 0,20190905 | 0,46412578 |
| 55367_at | PIDD1 | -0,11705907 | 7,2569804 | -1,33644801 | 0,20196163 | 0,46419125 |
| 6572_at | SLC18A3 | -0,12181619 | 4,14400875 | -1,33635845 | 0,20199018 | 0,46420149 |
| 339807_at | LOC339807 | -0,13364504 | 5,3115548 | -1,33604764 | 0,2020893 | 0,46437387 |
| 51661_at | FKBP7 | 0,27667195 | 5,9130549 | 1,33588691 | 0,20214057 | 0,46443627 |
| 373156_at | GSTK1 | 0,45300203 | 10,1086234 | 1,33579703 | 0,20216924 | 0,46444676 |
| 129607_at | CMPK2 | 0,69478491 | 7,95929602 | 1,33517878 | 0,20236657 | 0,46480376 |
| 2319_at | FLOT2 | 0,37325057 | 8,46586573 | 1,33515896 | 0,2023729 | 0,46480376 |
| 124817_at | CNTD1 | -0,14155455 | 3,81067917 | -1,33493358 | 0,20244488 | 0,46483385 |
| 64109_at | CRLF2 | -0,13431404 | 4,55159923 | -1,33491367 | 0,20245123 | 0,46483385 |
| 54084_at | TSPEAR | -0,11303929 | 4,28072692 | -1,33485246 | 0,20247079 | 0,46483385 |
| 283587_at | LINC01146 | -0,07759818 | 2,74829075 | -1,33481572 | 0,20248252 | 0,46483385 |
| 6653_at | SORL1 | 0,16830455 | 9,57531043 | 1,33461741 | 0,20254588 | 0,46492389 |
| 6571_at | SLC18A2 | -0,14566136 | 3,68718785 | -1,33446042 | 0,20259605 | 0,46498364 |
| 10261_at | IGSF6 | -0,13417868 | 3,23022317 | -1,33436521 | 0,20262648 | 0,46499808 |

| | | | | | | |
|---|---|---|---|---|---|---|
| 140606_at | SELM | 0,77104188 | 5,71930283 | 1,33339955 | 0,20293532 | 0,46557349 |
| 414_at | ARSD | 0,53954239 | 6,66649782 | 1,33329431 | 0,202969 | 0,46557349 |
| 660_at | BMX | -0,12912769 | 3,65599263 | -1,33325713 | 0,2029809 | 0,46557349 |
| 55656_at | INTS8 | -0,25714684 | 8,71005412 | -1,33324788 | 0,20298387 | 0,46557349 |
| 4861_at | NPAS1 | -0,10612736 | 5,25925228 | -1,33320352 | 0,20299806 | 0,46557349 |
| 85443_at | DCLK3 | 0,10757291 | 3,60777761 | 1,33305383 | 0,20304598 | 0,46562795 |
| 340554_at | ZC3H12B | -0,10288973 | 3,44933836 | -1,33278811 | 0,20313107 | 0,46576763 |
| 8513_at | LIPF | -0,13958956 | 3,72820766 | -1,33243063 | 0,20324559 | 0,46597474 |
| 5981_at | RFC1 | -0,18320827 | 8,53420651 | -1,33211397 | 0,20334707 | 0,46615193 |
| 51127_at | TRIM17 | 0,15484779 | 4,91855926 | 1,33189105 | 0,20341854 | 0,46626027 |
| 29101_at | SSU72 | -0,08465481 | 9,97543162 | -1,33162065 | 0,20350525 | 0,46635125 |
| 65264_at | UBE2Z | 0,13014563 | 9,64698018 | 1,33161629 | 0,20350665 | 0,46635125 |
| 84439_at | HHIPL1 | -0,13904585 | 4,86427002 | -1,3313714 | 0,20358521 | 0,46641905 |
| 6758_at | SSX5 | -0,17050154 | 3,68761179 | -1,33132319 | 0,20360068 | 0,46641905 |
| 100506681_a | JARID2-AS1 | -0,13210562 | 4,41956542 | -1,33129765 | 0,20360887 | 0,46641905 |
| 7453_at | WARS | -0,24747583 | 9,36225129 | -1,33120112 | 0,20363985 | 0,46643453 |
| 2495_at | FTH1 | 0,31392032 | 9,15953062 | 1,33103791 | 0,20369223 | 0,46649905 |
| 146852_at | ODF4 | -0,09463428 | 4,61994867 | -1,33093734 | 0,20372452 | 0,46651752 |
| 54407_at | SLC38A2 | 0,16946039 | 11,6346859 | 1,33026291 | 0,20394112 | 0,46695801 |
| 60506_at | NYX | -0,12847523 | 5,65460356 | -1,33016765 | 0,20397173 | 0,46697259 |
| 55090_at | MED9 | -0,27735656 | 7,28290907 | -1,32994798 | 0,20404232 | 0,46704453 |
| 10725_at | NFAT5 | 0,31873231 | 7,35188636 | 1,32991899 | 0,20405164 | 0,46704453 |
| 117153_at | MIA2 | -0,07208866 | 3,09710603 | -1,32978363 | 0,20409516 | 0,46708863 |
| 121006_at | FAM186A | -0,11254804 | 3,02900811 | -1,32937251 | 0,20422736 | 0,46728344 |
| 57532_at | NUFIP2 | 0,1048884 | 8,97707131 | 1,32936805 | 0,2042288 | 0,46728344 |
| 125206_at | SLC5A10 | -0,12825218 | 5,66375216 | -1,3291944 | 0,20428466 | 0,46735575 |
| 51308_at | REEP2 | -0,20403617 | 6,31247942 | -1,32908448 | 0,20432003 | 0,46738115 |
| 89953_at | KLC4 | 0,1484243 | 5,82230693 | 1,32892909 | 0,20437003 | 0,46741985 |
| 56918_at | C2orf83 | -0,11979907 | 3,64712487 | -1,3287566 | 0,20442556 | 0,46741985 |
| 100507043_a | TUNAR | -0,14253901 | 4,85914983 | -1,32874112 | 0,20443054 | 0,46741985 |
| 4920_at | ROR2 | 0,1855746 | 4,06788339 | 1,32864697 | 0,20446085 | 0,46741985 |
| 23484_at | LEPROTL1 | -0,33184927 | 8,71302946 | -1,32862358 | 0,20446838 | 0,46741985 |
| 57555_at | NLGN2 | -0,14719541 | 6,89706961 | -1,32837087 | 0,20454976 | 0,46741985 |
| 26532_at | OR10H3 | -0,2368569 | 4,10281291 | -1,32832276 | 0,20456525 | 0,46741985 |
| 55374_at | TMCO6 | -0,2077006 | 7,87723864 | -1,32820335 | 0,20460371 | 0,46741985 |
| 155038_at | GIMAP8 | -0,09582507 | 3,8056159 | -1,32817023 | 0,20461438 | 0,46741985 |
| 4745_at | NELL1 | -0,09769836 | 4,38230442 | -1,32814208 | 0,20462345 | 0,46741985 |
| 100505841_a | LOC10050584 | -0,12672595 | 3,65320946 | -1,32812717 | 0,20462826 | 0,46741985 |
| 6132_at | RPL8 | -0,24867404 | 12,6730239 | -1,3280969 | 0,20463801 | 0,46741985 |
| 4800_at | NFYA | -0,11052984 | 7,7565575 | -1,32799809 | 0,20466985 | 0,46741985 |
| 100505622_a | LOC10050562 | -0,13446317 | 4,64301907 | -1,32794846 | 0,20468584 | 0,46741985 |
| 1487_at | CTBP1 | 0,0837817 | 10,1503012 | 1,32790166 | 0,20470092 | 0,46741985 |
| 5796_at | PTPRK | 0,13561445 | 10,7169435 | 1,32767476 | 0,20477405 | 0,46750653 |
| 7178_at | TPT1 | 0,12277705 | 10,0855366 | 1,32763329 | 0,20478742 | 0,46750653 |
| 2660_at | MSTN | 0,09904376 | 2,68746621 | 1,32750244 | 0,2048296 | 0,46754742 |
| 90070_at | LACRT | -0,12014164 | 4,89231259 | -1,32718782 | 0,20493107 | 0,4677236 |
| 92291_at | CAPN13 | 0,37777119 | 5,50766525 | 1,32707137 | 0,20496863 | 0,46775391 |
| 27035_at | NOX1 | -0,11001537 | 4,04555272 | -1,32614921 | 0,20526629 | 0,46834923 |
| 282763_at | OR51B5 | -0,11794479 | 3,58732481 | -1,32608692 | 0,20528641 | 0,46834923 |
| 160728_at | SLC5A8 | -0,12937351 | 4,82998975 | -1,32601799 | 0,20530868 | 0,46834923 |

| | | | | | | |
|---|---|---|---|---|---|---|
| 7007_at | TECTA | -0,12370785 | 3,8609482 | -1,32596204 | 0,20532675 | 0,46834923 |
| 9344_at | TAOK2 | 0,15933467 | 7,23755008 | 1,32576614 | 0,20539005 | 0,46843813 |
| 57467_at | HHATL | -0,15238004 | 4,42061531 | -1,32561089 | 0,20544021 | 0,46849708 |
| 10202_at | DHRS2 | 0,7324801 | 10,0738613 | 1,3251511 | 0,20558886 | 0,46878057 |
| 6424_at | SFRP4 | -0,13549208 | 3,65007805 | -1,32471349 | 0,20573042 | 0,46904782 |
| 3597_at | IL13RA1 | 0,20058428 | 8,28480754 | 1,32441892 | 0,20582575 | 0,46915592 |
| 3196_at | TLX2 | -0,13474479 | 5,21031549 | -1,32441647 | 0,20582654 | 0,46915592 |
| 11199_at | ANXA10 | -1,17354382 | 10,2122975 | -1,32427669 | 0,20587179 | 0,46920353 |
| 50831_at | TAS2R3 | -0,14439668 | 4,57175197 | -1,32416245 | 0,20590877 | 0,46923231 |
| 157638_at | FAM84B | -0,25600319 | 7,77534238 | -1,32400799 | 0,20595879 | 0,46929077 |
| 256130_at | TMEM196 | -0,10143563 | 3,3199498 | -1,32333141 | 0,206178 | 0,46970372 |
| 22907_at | DHX30 | -0,15440181 | 7,62919615 | -1,32329812 | 0,20618879 | 0,46970372 |
| 4297_at | KMT2A | 0,15821505 | 7,33108413 | 1,32315125 | 0,2062364 | 0,46975662 |
| 9479_at | MAPK8IP1 | 0,17546727 | 4,50943081 | 1,32296415 | 0,20629706 | 0,46980978 |
| 11018_at | TMED1 | 0,08860804 | 8,03005385 | 1,32292883 | 0,20630852 | 0,46980978 |
| 11025_at | LILRB3 | -0,19603731 | 6,52662778 | -1,32253918 | 0,20643491 | 0,47000118 |
| 8832_at | CD84 | -0,11211175 | 3,67614424 | -1,3224494 | 0,20646404 | 0,47000118 |
| 50854_at | C6orf48 | 0,28913426 | 9,99118946 | 1,32239819 | 0,20648066 | 0,47000118 |
| 29068_at | ZBTB44 | 0,2912188 | 8,13030896 | 1,3223689 | 0,20649016 | 0,47000118 |
| 5957_at | RCVRN | -0,09907375 | 4,26195425 | -1,32203656 | 0,20659804 | 0,47013273 |
| 26775_at | SNORA72 | -0,50199985 | 4,79498978 | -1,32199136 | 0,20661271 | 0,47013273 |
| 3218_at | HOXB8 | 0,34727265 | 6,72965759 | 1,32196529 | 0,20662118 | 0,47013273 |
| 80176_at | SPSB1 | 0,19708681 | 7,79881782 | 1,32188902 | 0,20664594 | 0,47013355 |
| 100129387_a | GABPB1-AS1 | -0,13555683 | 6,49575165 | -1,32155007 | 0,20675603 | 0,47032025 |
| 9330_at | GTF3C3 | -0,15958438 | 7,03870109 | -1,32143405 | 0,20679372 | 0,47032025 |
| 8369_at | HIST1H4G | -0,10623921 | 3,75730944 | -1,32140718 | 0,20680245 | 0,47032025 |
| 100533178_a | PRORY | -0,10536922 | 4,48061042 | -1,32130945 | 0,20683421 | 0,47032025 |
| 3762_at | KCNJ5 | -0,12694076 | 4,8049247 | -1,3212606 | 0,20685008 | 0,47032025 |
| 375686_at | SPATC1 | -0,15085249 | 5,65345272 | -1,32109828 | 0,20690284 | 0,47038467 |
| 30812_at | SOX8 | -0,15431399 | 4,75262082 | -1,32089799 | 0,20696795 | 0,47038851 |
| 55636_at | CHD7 | 0,12368584 | 7,19802833 | 1,32088225 | 0,20697306 | 0,47038851 |
| 2584_at | GALK1 | 0,17852896 | 6,71191562 | 1,32086774 | 0,20697778 | 0,47038851 |
| 9322_at | TRIP10 | 0,16646913 | 8,03071564 | 1,32076758 | 0,20701035 | 0,47040703 |
| 3601_at | IL15RA | -0,1374117 | 7,39182387 | -1,32042845 | 0,20712065 | 0,47058415 |
| 9127_at | P2RX6 | -0,18640414 | 5,14594864 | -1,32035432 | 0,20714477 | 0,47058415 |
| 51079_at | NDUFA13 | 0,14025468 | 10,5645851 | 1,32026556 | 0,20717365 | 0,47058415 |
| 11103_at | KRR1 | -0,2532161 | 8,75707135 | -1,32022756 | 0,20718601 | 0,47058415 |
| 729852_at | UMAD1 | 0,30462594 | 7,59661386 | 1,3200563 | 0,20724175 | 0,47065524 |
| 161142_at | FAM71D | -0,14594141 | 3,1077047 | -1,31987298 | 0,20730142 | 0,47069022 |
| 55065_at | SLC52A1 | 0,29212026 | 5,37630897 | 1,31985886 | 0,20730602 | 0,47069022 |
| 259240_at | WFDC9 | -0,13184455 | 4,1658787 | -1,31958979 | 0,20739363 | 0,47083364 |
| 153657_at | TTC23L | 0,11127816 | 3,55330437 | 1,3192837 | 0,20749333 | 0,47095211 |
| 229_at | ALDOB | -0,09953569 | 3,60472065 | -1,31927946 | 0,20749471 | 0,47095211 |
| 163486_at | DENND1B | 0,29682776 | 6,1746712 | 1,31912575 | 0,20754479 | 0,47096728 |
| 54522_at | ANKRD16 | -0,34175644 | 5,77614402 | -1,31910888 | 0,20755029 | 0,47096728 |
| 56925_at | LXN | 1,23930084 | 5,93751851 | 1,31872253 | 0,20767621 | 0,47114229 |
| 6000_at | RGS7 | -0,09838183 | 3,21039869 | -1,31868666 | 0,20768791 | 0,47114229 |
| 135293_at | PM20D2 | -0,41368305 | 7,64490045 | -1,31862435 | 0,20770822 | 0,47114229 |
| 120425_at | AMICA1 | -0,13964452 | 4,52097969 | -1,31856164 | 0,20772867 | 0,47114229 |
| 57211_at | ADGRG6 | 0,51218638 | 8,071304 | 1,31849713 | 0,20774971 | 0,47114229 |

| | | | | | | |
|---|---|---|---|---|---|---|
| 63941_at | NECAB3 | 0,15335034 | 7,81586705 | 1,31838482 | 0,20778634 | 0,4711649 |
| 56311_at | ANKRD7 | 0,13317696 | 3,2195722 | 1,31827667 | 0,20782161 | 0,4711649 |
| 101927249_a | PDZRN3-AS1 | -0,15214783 | 3,54671301 | -1,31824158 | 0,20783306 | 0,4711649 |
| 3702_at | ITK | -0,08525803 | 3,62979101 | -1,31769997 | 0,2080098 | 0,47151009 |
| 487_at | ATP2A1 | -0,10526745 | 4,92411095 | -1,31749085 | 0,20807807 | 0,47156529 |
| 148345_at | C1orf127 | -0,18231244 | 5,89897325 | -1,31743149 | 0,20809745 | 0,47156529 |
| 221223_at | CES5A | -0,13650153 | 4,21057849 | -1,31739393 | 0,20810972 | 0,47156529 |
| 7053_at | TGM3 | -0,12885596 | 4,15700462 | -1,31732549 | 0,20813207 | 0,47156529 |
| 3886_at | KRT35 | -0,12638342 | 3,82262586 | -1,31714972 | 0,20818948 | 0,47160386 |
| 100216545_a | KMT2E-AS1 | -0,09093269 | 6,23063624 | -1,31712347 | 0,20819806 | 0,47160386 |
| 80321_at | CEP70 | 0,1650228 | 6,68543956 | 1,31698723 | 0,20824257 | 0,47161288 |
| 6520_at | SLC3A2 | -0,36823927 | 10,5749255 | -1,31696139 | 0,20825101 | 0,47161288 |
| 51350_at | KRT76 | -0,16263193 | 4,77108867 | -1,31687197 | 0,20828023 | 0,47162361 |
| 598_at | BCL2L1 | 0,41103215 | 7,39270673 | 1,31651901 | 0,20839559 | 0,4717913 |
| 5269_at | SERPINB6 | -0,17814499 | 8,36503148 | -1,31649343 | 0,20840395 | 0,4717913 |
| 283888_at | IL21R-AS1 | -0,12197317 | 4,87296702 | -1,31642062 | 0,20842776 | 0,4717913 |
| 138474_at | TAF1L | 0,16378666 | 4,03389247 | 1,31632596 | 0,20845871 | 0,47179528 |
| 54915_at | YTHDF1 | -0,10847072 | 10,5228853 | -1,31619535 | 0,20850143 | 0,47179528 |
| 283860_at | LINC00304 | -0,1100588 | 3,66015987 | -1,31619055 | 0,20850299 | 0,47179528 |
| 6453_at | ITSN1 | 0,30894887 | 7,67745305 | 1,31584874 | 0,20861481 | 0,47199285 |
| 1769_at | DNAH8 | -0,09500772 | 3,24240631 | -1,31519628 | 0,20882838 | 0,47242056 |
| 221122_at | LOC221122 | -0,11408194 | 4,59394694 | -1,314899 | 0,20892575 | 0,47253015 |
| 10260_at | DENND4A | 0,10128696 | 6,42005739 | 1,31489365 | 0,2089275 | 0,47253015 |
| 55172_at | DNAAF2 | -0,18261776 | 8,52006003 | -1,31482371 | 0,20895041 | 0,47253015 |
| 5330_at | PLCB2 | -0,10708877 | 5,22711372 | -1,31452668 | 0,20904775 | 0,47268474 |
| 26301_at | GBGT1 | 0,50140779 | 6,0632448 | 1,31446534 | 0,20906785 | 0,47268474 |
| 3621_at | ING1 | -0,20092499 | 7,50643901 | -1,31433231 | 0,20911146 | 0,47272785 |
| 5585_at | PKN1 | 0,22768214 | 6,03506368 | 1,3139109 | 0,20924964 | 0,47292713 |
| 3697_at | ITIH1 | -0,17538329 | 4,46639975 | -1,31385474 | 0,20926806 | 0,47292713 |
| 151121_at | LOC151121 | -0,12423146 | 3,39773234 | -1,31383888 | 0,20927326 | 0,47292713 |
| 150538_at | SATB2-AS1 | -0,10460949 | 3,97557069 | -1,31371631 | 0,20931347 | 0,47293816 |
| 3814_at | KISS1 | -0,14562598 | 5,1903387 | -1,31366318 | 0,2093309 | 0,47293816 |
| 199731_at | CADM4 | 0,12708988 | 5,45947434 | 1,3135995 | 0,2093518 | 0,47293816 |
| 92196_at | DAPL1 | 0,38713148 | 5,03233743 | 1,31343025 | 0,20940734 | 0,47300815 |
| 8529_at | CYP4F2 | -0,27939087 | 5,2034641 | -1,31330681 | 0,20944785 | 0,4730442 |
| 2569_at | GABRR1 | 0,12418439 | 3,29405724 | 1,3130938 | 0,20951778 | 0,47308575 |
| 414926_at | LINC00593 | -0,13196432 | 4,31570996 | -1,31305346 | 0,20953102 | 0,47308575 |
| 51755_at | CDK12 | -0,08861302 | 7,41175165 | -1,31302635 | 0,20953993 | 0,47308575 |
| 91450_at | LOC91450 | -0,14582282 | 5,02818955 | -1,31280731 | 0,20961186 | 0,47310782 |
| 100506211_a | MIR210HG | -0,17012299 | 6,22600765 | -1,31274327 | 0,2096329 | 0,47310782 |
| 196740_at | VSTM4 | -0,10473773 | 4,40604679 | -1,31267259 | 0,20965611 | 0,47310782 |
| 7329_at | UBE2I | -0,15370415 | 8,6184402 | -1,31258528 | 0,2096848 | 0,47310782 |
| 53919_at | SLCO1C1 | -0,12105561 | 3,11777176 | -1,31255689 | 0,20969412 | 0,47310782 |
| 84669_at | USP32 | -0,11832201 | 6,17963493 | -1,31254793 | 0,20969707 | 0,47310782 |
| 128344_at | PIFO | 0,17919782 | 4,44143319 | 1,31233556 | 0,20976685 | 0,47316085 |
| 6146_at | RPL22 | -0,07832973 | 8,78082589 | -1,3123269 | 0,2097697 | 0,47316085 |
| 336_at | APOA2 | -0,10931684 | 4,08192222 | -1,31219507 | 0,20981303 | 0,47320317 |
| 400954_at | EML6 | 0,14478421 | 4,82573016 | 1,31208391 | 0,20984957 | 0,47323019 |
| 53340_at | SPA17 | -0,17498482 | 7,57119504 | -1,31169517 | 0,2099774 | 0,47342034 |
| 340168_at | DPPA5 | -0,11945031 | 3,67949553 | -1,311678 | 0,20998305 | 0,47342034 |

| | | | | | | |
|---|---|---|---|---|---|---|
| 151658_at | LINC00635 | -0,08391765 | 3,310855 | -1,31147837 | 0,21004871 | 0,47351297 |
| 5576_at | PRKAR2A | -0,10965088 | 8,29813501 | -1,31116071 | 0,21015324 | 0,47369318 |
| 92346_at | C1orf105 | -0,17361466 | 3,77830883 | -1,31050851 | 0,21036799 | 0,47412174 |
| 23268_at | DNMBP | 0,10341448 | 7,33472914 | 1,3102716 | 0,21044604 | 0,47424216 |
| 27430_at | MAT2B | 0,22298055 | 9,95740194 | 1,30994296 | 0,21055434 | 0,47439998 |
| 100506403_a | LOC10050640 | -0,13196803 | 3,83640224 | -1,30990966 | 0,21056532 | 0,47439998 |
| 124936_at | CYB5D2 | 0,31866246 | 7,47175581 | 1,30971552 | 0,21062933 | 0,47446361 |
| 4115_at | MAGEB4 | -0,10025278 | 3,36685178 | -1,30967456 | 0,21064283 | 0,47446361 |
| 100128496_a | C20orf78 | -0,18079356 | 4,29082404 | -1,30934577 | 0,21075127 | 0,47463078 |
| 101926942_a | LOC10192694 | 0,11171757 | 3,49695568 | 1,30930014 | 0,21076632 | 0,47463078 |
| 574414_at | PRR9 | -0,21800254 | 3,98335686 | -1,30907806 | 0,2108396 | 0,4747388 |
| 401612_at | SLC25A53 | 0,13992056 | 3,89649885 | 1,308991 | 0,21086833 | 0,4747388 |
| 140458_at | ASB5 | -0,08903645 | 2,93887803 | -1,3089307 | 0,21088823 | 0,4747388 |
| 5687_at | PSMA6 | -0,10220313 | 10,6659291 | -1,30827615 | 0,21110435 | 0,47505765 |
| 101928739_a | LOC10192873 | -0,08586428 | 2,81239383 | -1,30823217 | 0,21111888 | 0,47505765 |
| 9112_at | MTA1 | -0,08429441 | 8,27212751 | -1,30821401 | 0,21112487 | 0,47505765 |
| 378832_at | COL18A1-AS1 | -0,09297748 | 4,85163395 | -1,308203 | 0,21112851 | 0,47505765 |
| 79871_at | RPAP2 | -0,11262253 | 7,39035142 | -1,30759275 | 0,21133018 | 0,47545589 |
| 91304_at | TMEM259 | 0,11230456 | 8,81174426 | 1,30749256 | 0,21136331 | 0,47547488 |
| 9535_at | GMFG | -0,14353043 | 4,91792033 | -1,30729816 | 0,21142759 | 0,47555724 |
| 150368_at | FAM109B | -0,13028015 | 5,75644294 | -1,30719821 | 0,21146065 | 0,47555724 |
| 375704_at | ENHO | 0,14958326 | 5,63348049 | 1,30713311 | 0,21148219 | 0,47555724 |
| 6295_at | SAG | -0,12074992 | 4,67590046 | -1,30704666 | 0,21151078 | 0,47555724 |
| 1188_at | CLCNKB | -0,14148614 | 5,32199819 | -1,30697773 | 0,21153359 | 0,47555724 |
| 100130417_a | LOC10013041 | -0,13873184 | 5,0466981 | -1,30693404 | 0,21154804 | 0,47555724 |
| 8904_at | CPNE1 | 0,13914901 | 10,7364286 | 1,30678136 | 0,21159856 | 0,47558936 |
| 64344_at | HIF3A | -0,1116213 | 4,90761104 | -1,30674164 | 0,21161171 | 0,47558936 |
| 133308_at | SLC9B2 | -0,11652171 | 5,13458579 | -1,30664336 | 0,21164424 | 0,47559723 |
| 168374_at | ZNF92 | 0,16232803 | 5,32152004 | 1,30658186 | 0,21166459 | 0,47559723 |
| 84215_at | ZNF541 | -0,1516429 | 4,95243382 | -1,30638569 | 0,21172954 | 0,47564379 |
| 3310_at | HSPA6 | -0,18818929 | 5,07036854 | -1,3062966 | 0,21175904 | 0,47564379 |
| 81543_at | LRRC3 | -0,12923294 | 4,99598129 | -1,30629554 | 0,21175939 | 0,47564379 |
| 57132_at | CHMP1B | 0,34003731 | 7,81529809 | 1,30598604 | 0,2118619 | 0,47576471 |
| 5062_at | PAK2 | -0,13926405 | 9,42660171 | -1,30598386 | 0,21186262 | 0,47576471 |
| 5915_at | RARB | 0,15982253 | 6,43391175 | 1,30559569 | 0,21199124 | 0,4759899 |
| 8290_at | HIST3H3 | -0,13609798 | 4,79069658 | -1,30553207 | 0,21201232 | 0,4759899 |
| 51330_at | TNFRSF12A | -0,11722617 | 10,6481632 | -1,3052852 | 0,21209416 | 0,47611815 |
| 910_at | CD1B | -0,16917163 | 4,14686161 | -1,3051253 | 0,21214718 | 0,47616831 |
| 644139_at | PIRT | -0,1400592 | 4,30774251 | -1,30506873 | 0,21216594 | 0,47616831 |
| 56114_at | PCDHGA1 | -0,18023646 | 4,68987221 | -1,30483813 | 0,21224243 | 0,4762385 |
| 338322_at | NLRP10 | -0,10941165 | 3,8256921 | -1,30476455 | 0,21226684 | 0,4762385 |
| 55501_at | CHST12 | 0,16477454 | 7,32760537 | 1,30475084 | 0,21227139 | 0,4762385 |
| 5745_at | PTH1R | -0,14038376 | 4,97032004 | -1,30442321 | 0,21238012 | 0,47634074 |
| 23567_at | ZNF346 | 0,10665265 | 6,23626601 | 1,30440532 | 0,21238605 | 0,47634074 |
| 100505865_a | LINC00920 | 0,48056829 | 5,82696457 | 1,30433363 | 0,21240985 | 0,47634074 |
| 959_at | CD40LG | -0,11225729 | 4,46418715 | -1,30428503 | 0,21242598 | 0,47634074 |
| 5657_at | PRTN3 | -0,11692033 | 4,58588168 | -1,30424099 | 0,2124406 | 0,47634074 |
| 56964_at | WDR93 | -0,15883237 | 4,56010247 | -1,30393417 | 0,21254248 | 0,47651371 |
| 91316_at | GUSBP11 | 0,16447291 | 5,73658345 | 1,30347121 | 0,21269628 | 0,47678756 |
| 102723809_a | LOC10272380 | -0,12591681 | 3,18988069 | -1,30341751 | 0,21271413 | 0,47678756 |

| | | | | | | |
|---|---|---|---|---|---|---|
| 100505746_a | ITGB2-AS1 | -0,094284 | 4,15818479 | -1,30286423 | 0,21289806 | 0,47714432 |
| 9020_at | MAP3K14 | 0,41385003 | 6,48747457 | 1,30275984 | 0,21293278 | 0,47716661 |
| 7587_at | ZNF37A | -0,08856567 | 5,10736653 | -1,30251095 | 0,21301557 | 0,47729662 |
| 23229_at | ARHGEF9 | 0,13504525 | 6,38551399 | 1,3023805 | 0,21305898 | 0,47733835 |
| 79885_at | HDAC11 | 0,15462001 | 7,19555126 | 1,30194032 | 0,21320549 | 0,47755373 |
| 151300_at | LINC00608 | -0,0797433 | 3,37019142 | -1,30191216 | 0,21321487 | 0,47755373 |
| 727936_at | GXYLT2 | 0,35861725 | 5,34220968 | 1,30186825 | 0,21322949 | 0,47755373 |
| 7512_at | XPNPEP2 | -0,16174835 | 4,56104632 | -1,30168868 | 0,21328929 | 0,47763212 |
| 1117_at | CHI3L2 | -0,17160882 | 4,44093171 | -1,30160333 | 0,21331772 | 0,47764026 |
| 89884_at | LHX4 | -0,11785673 | 4,27765615 | -1,30149541 | 0,21335366 | 0,47766523 |
| 729515_at | TMEM242 | 0,10612945 | 6,1213782 | 1,30110474 | 0,21348384 | 0,47790112 |
| 23365_at | ARHGEF12 | 0,17812413 | 7,70618512 | 1,30094514 | 0,21353704 | 0,47796467 |
| 8435_at | SOAT2 | -0,15630369 | 4,97829327 | -1,30077243 | 0,21359461 | 0,478038 |
| 55435_at | AP1AR | -0,28054814 | 7,16366906 | -1,30061574 | 0,21364687 | 0,47809939 |
| 55592_at | GOLGA2P5 | 0,16655292 | 4,84107237 | 1,30049355 | 0,21368762 | 0,47813505 |
| 11061_at | LECT1 | -0,13133374 | 3,66776808 | -1,30038608 | 0,21372346 | 0,47815971 |
| 100861573_a | LINC00572 | -0,14872011 | 3,89511766 | -1,30030766 | 0,21374963 | 0,47816271 |
| 196993_at | CT62 | -0,07727334 | 3,92415719 | -1,30010687 | 0,21381662 | 0,47825704 |
| 259282_at | BOD1L1 | 0,11770161 | 7,50501174 | 1,29979772 | 0,21391981 | 0,47835118 |
| 57801_at | HES4 | 0,26147823 | 8,8416815 | 1,29978517 | 0,213924 | 0,47835118 |
| 440823_at | MIAT | -0,10958344 | 4,35863451 | -1,29975757 | 0,21393321 | 0,47835118 |
| 140834_at | LINC01620 | -0,09212139 | 3,72240497 | -1,29944354 | 0,21403807 | 0,4785301 |
| 65109_at | UPF3B | -0,14091228 | 4,94127944 | -1,29930761 | 0,21408347 | 0,47857606 |
| 1041_at | CDSN | -0,13136892 | 4,88298801 | -1,29899111 | 0,21418921 | 0,47874704 |
| 2781_at | GNAZ | -0,11188897 | 5,43305708 | -1,29888301 | 0,21422534 | 0,47874704 |
| 51657_at | STYXL1 | -0,07577405 | 8,8565357 | -1,29879756 | 0,2142539 | 0,47874704 |
| 11122_at | PTPRT | -0,13197002 | 4,78783511 | -1,29876778 | 0,21426385 | 0,47874704 |
| 26108_at | PYGO1 | 0,26839329 | 4,69870576 | 1,29870685 | 0,21428422 | 0,47874704 |
| 92359_at | CRB3 | 0,14118083 | 7,7193745 | 1,29822178 | 0,21444642 | 0,47900943 |
| 221_at | ALDH3B1 | 0,4603548 | 7,57899532 | 1,29807421 | 0,21449578 | 0,47900943 |
| 165_at | AEBP1 | -0,15199105 | 5,66215531 | -1,29797192 | 0,21453001 | 0,47900943 |
| 115861_at | NXNL1 | -0,13648748 | 7,04165356 | -1,29792787 | 0,21454475 | 0,47900943 |
| 64106_at | NPFFR1 | -0,10288176 | 5,99011189 | -1,29791742 | 0,21454824 | 0,47900943 |
| 7306_at | TYRP1 | -0,08456224 | 3,08870978 | -1,29787059 | 0,21456392 | 0,47900943 |
| 728361_at | OVOL3 | -0,15885143 | 5,21221288 | -1,2977856 | 0,21459236 | 0,47900943 |
| 84304_at | NUDT22 | 0,10551547 | 7,65480379 | 1,29776097 | 0,2146006 | 0,47900943 |
| 102724814_a | LOC10272481 | -0,11683142 | 5,5471815 | -1,2975222 | 0,21468052 | 0,4791323 |
| 22884_at | WDR37 | 0,30178797 | 7,50096042 | 1,29731478 | 0,21474997 | 0,47919107 |
| 201627_at | DENND6A | 0,16214441 | 8,76448079 | 1,29723983 | 0,21477507 | 0,47919107 |
| 57721_at | METTL14 | 0,29954665 | 7,33415507 | 1,29722068 | 0,21478148 | 0,47919107 |
| 168002_at | DACT2 | -0,15129286 | 5,89280744 | -1,29684137 | 0,21490855 | 0,47925077 |
| 101926978_a | LINC01111 | -0,13441761 | 3,44232959 | -1,29681734 | 0,2149166 | 0,47925077 |
| 115106_at | HAUS1 | -0,3217521 | 8,2258449 | -1,29681347 | 0,21491789 | 0,47925077 |
| 284252_at | KCTD1 | -0,25999924 | 6,68872682 | -1,29670254 | 0,21495507 | 0,47925077 |
| 29094_at | LGALSL | 0,15930591 | 7,66191382 | 1,29669225 | 0,21495851 | 0,47925077 |
| 4664_at | NAB1 | -0,22129856 | 6,80041789 | -1,29668446 | 0,21496112 | 0,47925077 |
| 79646_at | PANK3 | -0,13701086 | 7,73657195 | -1,29662099 | 0,2149824 | 0,47925077 |
| 28511_at | NKIRAS2 | 0,15770274 | 8,48489705 | 1,29572617 | 0,21528246 | 0,47970002 |
| 25818_at | KLK5 | -0,35540783 | 5,88863457 | -1,29572488 | 0,2152829 | 0,47970002 |
| 51199_at | NIN | -0,11622258 | 6,54738592 | -1,29568039 | 0,21529782 | 0,47970002 |

| | | | | | | |
|---|---|---|---|---|---|---|
| 259236_at | TMIE | 0,11988961 | 4,71058783 | 1,29567423 | 0,21529989 | 0,47970002 |
| 144110_at | TMEM86A | 0,15849033 | 5,34629821 | 1,29564877 | 0,21530843 | 0,47970002 |
| 6461_at | SHB | -0,17060334 | 7,74733462 | -1,2953969 | 0,21539296 | 0,47983285 |
| 128869_at | PIGU | -0,13011174 | 9,17531287 | -1,29520444 | 0,21545757 | 0,47992128 |
| 8831_at | SYNGAP1 | 0,13772643 | 5,77119266 | 1,29508041 | 0,21549922 | 0,47995854 |
| 56006_at | SMG9 | -0,16052969 | 7,92173777 | -1,29500537 | 0,21552442 | 0,47995917 |
| 7159_at | TP53BP2 | 0,09282647 | 8,61613639 | 1,29484153 | 0,21557944 | 0,48002622 |
| 5360_at | PLTP | -0,14366619 | 5,755266 | -1,29435237 | 0,2157438 | 0,48033667 |
| 347862_at | PDDC1 | -0,23063532 | 6,77720326 | -1,29411071 | 0,21582503 | 0,480462 |
| 79695_at | GALNT12 | -0,2886861 | 7,25846628 | -1,2939765 | 0,21587016 | 0,48050384 |
| 79699_at | ZYG11B | 0,20398779 | 8,5213905 | 1,29390644 | 0,21589372 | 0,48050384 |
| 344752_at | AADACL2 | 0,12558219 | 3,09099742 | 1,29356959 | 0,21600702 | 0,48066844 |
| 4258_at | MGST2 | 0,23341466 | 9,11331781 | 1,29351019 | 0,216027 | 0,48066844 |
| 9519_at | TBPL1 | 0,15853886 | 9,17925662 | 1,29346403 | 0,21604254 | 0,48066844 |
| 80739_at | C6orf25 | -0,14206849 | 5,11418961 | -1,29321891 | 0,21612502 | 0,48079643 |
| 84076_at | TKTL2 | -0,11876054 | 3,7599643 | -1,29308734 | 0,21616931 | 0,48083942 |
| 144920_at | LINC00343 | -0,10143873 | 2,90504535 | -1,2929847 | 0,21620386 | 0,48084057 |
| 66035_at | SLC2A11 | 0,0959906 | 5,97757903 | 1,29293751 | 0,21621975 | 0,48084057 |
| 29087_at | THYN1 | 0,10705741 | 7,89799174 | 1,29278493 | 0,21627113 | 0,48087844 |
| 728695_at | SPANXB1 | -0,12161152 | 3,10733569 | -1,29273867 | 0,2162867 | 0,48087844 |
| 1146_at | CHRNG | -0,15715263 | 5,88284192 | -1,29263708 | 0,21632092 | 0,480899 |
| 64403_at | CDH24 | -0,19782936 | 7,04306594 | -1,29242702 | 0,21639168 | 0,48095904 |
| 90342_at | FER1L5 | -0,1361581 | 4,1693956 | -1,29240867 | 0,21639786 | 0,48095904 |
| 1992_at | SERPINB1 | -0,11452224 | 4,40861305 | -1,29230431 | 0,21643302 | 0,48098169 |
| 100505783_a | OSER1-AS1 | 0,36831342 | 5,43702979 | 1,29216073 | 0,21648141 | 0,48102221 |
| 2905_at | GRIN2C | -0,16241298 | 4,98657421 | -1,29210201 | 0,2165012 | 0,48102221 |
| 51075_at | TMX2 | -0,19699934 | 9,64547933 | -1,2917834 | 0,2166086 | 0,48117184 |
| 9355_at | LHX2 | -0,07818093 | 4,60473505 | -1,29169151 | 0,21663959 | 0,48117184 |
| 64927_at | TTC23 | 0,08096525 | 6,15459494 | 1,29167996 | 0,21664348 | 0,48117184 |
| 118424_at | UBE2J2 | -0,11442378 | 7,6110041 | -1,29144999 | 0,21672104 | 0,48125308 |
| 2249_at | FGF4 | -0,09123809 | 4,90122251 | -1,29142335 | 0,21673003 | 0,48125308 |
| 4184_at | SMCP | -0,21747233 | 4,40811504 | -1,29115895 | 0,21681923 | 0,4813467 |
| 285613_at | RELL2 | -0,09160731 | 7,0140707 | -1,29113815 | 0,21682625 | 0,4813467 |
| 140679_at | SLC32A1 | -0,13069558 | 4,21315975 | -1,29107621 | 0,21684715 | 0,4813467 |
| 414157_at | C10orf62 | -0,13014559 | 5,60942087 | -1,29092558 | 0,21689799 | 0,48134827 |
| 4142_at | MAS1 | -0,22010562 | 4,48396642 | -1,29091414 | 0,21690185 | 0,48134827 |
| 135927_at | C7orf34 | -0,12799901 | 4,33838148 | -1,29083625 | 0,21692814 | 0,48134827 |
| 55544_at | RBM38 | -0,17394833 | 7,30761779 | -1,29077797 | 0,21694781 | 0,48134827 |
| 375748_at | ERCC6L2 | 0,07775277 | 5,36703761 | 1,29061847 | 0,21700166 | 0,48138424 |
| 164656_at | TMPRSS6 | -0,14509594 | 5,74665444 | -1,29047454 | 0,21705027 | 0,48138424 |
| 3765_at | KCNJ9 | -0,1298129 | 4,7457221 | -1,29045089 | 0,21705825 | 0,48138424 |
| 10424_at | PGRMC2 | 0,19739158 | 8,275071 | 1,29043392 | 0,21706398 | 0,48138424 |
| 6398_at | SECTM1 | 0,25780413 | 5,76371085 | 1,29025709 | 0,21712371 | 0,48141442 |
| 51384_at | WNT16 | -0,10203809 | 3,82514853 | -1,29017517 | 0,21715139 | 0,48141442 |
| 1525_at | CXADR | 0,14751602 | 9,46805137 | 1,2901448 | 0,21716165 | 0,48141442 |
| 55511_at | SAGE1 | -0,14272513 | 3,26224761 | -1,29009771 | 0,21717756 | 0,48141442 |
| 5139_at | PDE3A | -0,07421423 | 3,90489819 | -1,28993563 | 0,21723233 | 0,48148042 |
| 4688_at | NCF2 | 0,18312434 | 5,53276489 | 1,28985049 | 0,2172611 | 0,48148879 |
| 84775_at | ZNF607 | 0,10400029 | 5,62240532 | 1,28967089 | 0,21732181 | 0,48155412 |
| 253012_at | HEPACAM2 | 0,54180088 | 3,36645208 | 1,28961537 | 0,21734058 | 0,48155412 |

| | | | | | | |
|---|---|---|---|---|---|---|
| 113235_at | SLC46A1 | 0,17756616 | 6,89167332 | 1,28945256 | 0,21739563 | 0,48162069 |
| 2327_at | FMO2 | -0,12083092 | 3,52374473 | -1,28934746 | 0,21743116 | 0,48164403 |
| 3611_at | ILK | 0,15483237 | 9,15609215 | 1,28904748 | 0,21753263 | 0,4818134 |
| 3270_at | HRC | -0,15073596 | 5,12538801 | -1,28891427 | 0,2175777 | 0,48185782 |
| 10793_at | ZNF273 | 0,15210812 | 5,39227558 | 1,28879868 | 0,21761682 | 0,48186853 |
| 348094_at | ANKDD1A | 0,66202448 | 6,41787502 | 1,28875213 | 0,21763257 | 0,48186853 |
| 372_at | ARCN1 | -0,1111931 | 10,16226 | -1,28819921 | 0,21781977 | 0,482174 |
| 127247_at | ASB17 | -0,08260387 | 2,96184178 | -1,28819676 | 0,2178206 | 0,482174 |
| 146227_at | BEAN1 | 0,32890801 | 4,43412744 | 1,28780472 | 0,2179534 | 0,4823909 |
| 388610_at | TRNP1 | -0,2425297 | 7,84704422 | -1,28769718 | 0,21798984 | 0,4823909 |
| 347735_at | SERINC2 | 0,14929523 | 10,4069738 | 1,28768577 | 0,21799371 | 0,4823909 |
| 347733_at | TUBB2B | -0,11227095 | 4,06348424 | -1,28737984 | 0,2180974 | 0,48256492 |
| 79652_at | TMEM204 | -0,09600965 | 5,2082152 | -1,28727203 | 0,21813395 | 0,48259036 |
| 26233_at | FBXL6 | -0,22779079 | 7,36648382 | -1,28705493 | 0,21820757 | 0,48267045 |
| 339665_at | SLC35E4 | -0,12444214 | 5,36806181 | -1,28696616 | 0,21823768 | 0,48267045 |
| 79339_at | OR51B4 | 0,11993802 | 4,63980043 | 1,28690757 | 0,21825755 | 0,48267045 |
| 4308_at | TRPM1 | -0,09207492 | 3,67307846 | -1,28683176 | 0,21828327 | 0,48267045 |
| 2907_at | GRINA | 0,17854177 | 7,84830125 | 1,28679589 | 0,21829543 | 0,48267045 |
| 53826_at | FXYD6 | -0,12378072 | 5,2894215 | -1,28640328 | 0,21842866 | 0,48286716 |
| 55294_at | FBXW7 | 0,3064044 | 6,92355849 | 1,28638598 | 0,21843453 | 0,48286716 |
| 1388_at | ATF6B | 0,35091844 | 8,31871819 | 1,28621981 | 0,21849094 | 0,48288881 |
| 10991_at | SLC38A3 | -0,15586658 | 4,6528919 | -1,28620943 | 0,21849447 | 0,48288881 |
| 54913_at | RPP25 | -0,44984756 | 6,11582737 | -1,28593603 | 0,2185873 | 0,48302529 |
| 100505929_a | LINC01549 | 0,07713839 | 2,92342832 | 1,28575284 | 0,21864952 | 0,48302529 |
| 10890_at | RAB10 | -0,12888482 | 10,6858228 | -1,28574812 | 0,21865113 | 0,48302529 |
| 84231_at | TRAF7 | -0,09364961 | 8,03641579 | -1,28564916 | 0,21868475 | 0,48302529 |
| 56941_at | HMCES | -0,18033168 | 8,69811321 | -1,28554162 | 0,21872128 | 0,48302529 |
| 114926_at | SMIM19 | 0,29951824 | 8,77850643 | 1,28551377 | 0,21873075 | 0,48302529 |
| 1633_at | DCK | -0,5200735 | 8,28335485 | -1,28551083 | 0,21873175 | 0,48302529 |
| 9686_at | VGLL4 | 0,22380164 | 8,4158055 | 1,28518651 | 0,21884197 | 0,48311115 |
| 3665_at | IRF7 | 0,84916395 | 7,53261279 | 1,28516935 | 0,2188478 | 0,48311115 |
| 136371_at | ASB10 | -0,12364092 | 5,52936701 | -1,285108 | 0,21886866 | 0,48311115 |
| 3561_at | IL2RG | -0,15985271 | 4,21215969 | -1,28510128 | 0,21887094 | 0,48311115 |
| 811_at | CALR | 0,08644538 | 9,61985133 | 1,28470079 | 0,21900713 | 0,48334256 |
| 114131_at | UCN3 | -0,17079266 | 5,22722629 | -1,2846454 | 0,21902597 | 0,48334256 |
| 4680_at | CEACAM6 | -1,05052326 | 7,48270891 | -1,28449618 | 0,21907673 | 0,48339921 |
| 64902_at | AGXT2 | -0,1737193 | 4,23174605 | -1,28432352 | 0,21913548 | 0,48346857 |
| 9333_at | TGM5 | -0,66035365 | 5,95659827 | -1,28425628 | 0,21915836 | 0,48346857 |
| 29882_at | ANAPC2 | 0,29188081 | 7,30325151 | 1,28397668 | 0,21925353 | 0,48362313 |
| 3902_at | LAG3 | -0,22506536 | 4,83712308 | -1,283831 | 0,21930313 | 0,4836613 |
| 151112_at | ZSWIM2 | -0,10004743 | 2,75226646 | -1,28369729 | 0,21934866 | 0,4836613 |
| 5304_at | PIP | 0,23683947 | 5,17261566 | 1,28368508 | 0,21935282 | 0,4836613 |
| 26496_at | OR10A3 | -0,10860054 | 3,87363799 | -1,28360046 | 0,21938164 | 0,4836613 |
| 3203_at | HOXA6 | 0,31301709 | 6,36349304 | 1,28355719 | 0,21939638 | 0,4836613 |
| 3892_at | KRT86 | -0,13533101 | 4,9608269 | -1,28329197 | 0,21948673 | 0,48378809 |
| 136242_at | PRSS37 | -0,1623712 | 4,42188735 | -1,28324093 | 0,21950412 | 0,48378809 |
| 6942_at | TCF20 | -0,07594834 | 8,44262613 | -1,28314082 | 0,21953824 | 0,48380793 |
| 79876_at | UBA5 | -0,22329855 | 7,83677066 | -1,28275276 | 0,21967052 | 0,48402223 |
| 4622_at | MYH4 | -0,0797543 | 3,05762891 | -1,28270811 | 0,21968574 | 0,48402223 |
| 100272217_a | LOC10027221 | 0,09221846 | 3,5613917 | 1,28252875 | 0,2197469 | 0,48404866 |

| | | | | | | |
|---|---|---|---|---|---|---|
| 283383_at | ADGRD1 | 0,1153584 | 4,22597839 | 1,28245898 | 0,2197707 | 0,48404866 |
| 101928555_a | LINC01537 | -0,48227037 | 4,4859479 | -1,28245188 | 0,21977312 | 0,48404866 |
| 128826_at | MIR1-1HG | -0,15927984 | 7,38035156 | -1,28210888 | 0,21989013 | 0,48414702 |
| 2897_at | GRIK1 | -0,07983378 | 3,26840332 | -1,28206285 | 0,21990584 | 0,48414702 |
| 1960_at | EGR3 | 0,08324514 | 3,95685061 | 1,2820469 | 0,21991128 | 0,48414702 |
| 93432_at | MGAM2 | -0,13093 | 3,22373154 | -1,2820263 | 0,21991831 | 0,48414702 |
| 354_at | KLK3 | -0,09520285 | 4,70050363 | -1,28187423 | 0,21997021 | 0,48415013 |
| 51286_at | CEND1 | -0,15491078 | 6,29241386 | -1,28183217 | 0,21998457 | 0,48415013 |
| 4648_at | MYO7B | -0,12055727 | 5,4537219 | -1,28180124 | 0,21999512 | 0,48415013 |
| 2244_at | FGB | -0,09198184 | 3,42335646 | -1,28165554 | 0,22004487 | 0,48415605 |
| 815_at | CAMK2A | -0,14351827 | 5,4882857 | -1,28164612 | 0,22004808 | 0,48415605 |
| 22865_at | SLITRK3 | 0,15518453 | 4,73271963 | 1,28149453 | 0,22009984 | 0,48421462 |
| 4952_at | OCRL | 0,11462274 | 7,67777111 | 1,28126583 | 0,22017795 | 0,48433114 |
| 100038246_a | TLX1NB | -0,13040787 | 4,43335416 | -1,28116656 | 0,22021186 | 0,48435043 |
| 148423_at | C1orf52 | -0,10178606 | 7,085514 | -1,28093962 | 0,22028939 | 0,48446565 |
| 10193_at | RNF41 | 0,16351329 | 7,16959604 | 1,28086574 | 0,22031464 | 0,48446586 |
| 64860_at | ARMCX5 | 0,15781594 | 7,08950422 | 1,28074245 | 0,22035678 | 0,48450322 |
| 162968_at | ZNF497 | -0,17545391 | 4,96035503 | -1,28062743 | 0,2203961 | 0,48453435 |
| 64856_at | VWA1 | 0,17435607 | 6,55467321 | 1,28050286 | 0,22043868 | 0,48457267 |
| 26230_at | TIAM2 | 0,12129245 | 4,89293053 | 1,28035087 | 0,22049065 | 0,48463161 |
| 64236_at | PDLIM2 | -0,12969119 | 6,23016557 | -1,2802135 | 0,22053763 | 0,48467957 |
| 100128977_a | MAPT-AS1 | -0,10744011 | 4,04333492 | -1,28012296 | 0,2205686 | 0,48469232 |
| 10641_at | NPRL2 | -0,09719292 | 7,66427172 | -1,27982421 | 0,22067081 | 0,48486161 |
| 9948_at | WDR1 | -0,11685507 | 11,1394088 | -1,27953693 | 0,22076913 | 0,48499369 |
| 63035_at | BCORL1 | -0,10486648 | 5,7223478 | -1,27941286 | 0,2208116 | 0,48499369 |
| 3178_at | HNRNPA1 | -0,31376535 | 7,21554527 | -1,27939237 | 0,22081861 | 0,48499369 |
| 54621_at | VSIG10 | 0,17634332 | 8,1655269 | 1,27923384 | 0,22087289 | 0,48499369 |
| 55251_at | PCMTD2 | 0,46494148 | 9,19502201 | 1,27923383 | 0,2208729 | 0,48499369 |
| 5916_at | RARG | 0,10000915 | 7,91502076 | 1,2792073 | 0,22088198 | 0,48499369 |
| 554279_at | LINC00862 | -0,09416022 | 3,0042087 | -1,279118 | 0,22091256 | 0,48500555 |
| 28990_at | ASTE1 | -0,12839008 | 6,93171497 | -1,27901182 | 0,22094893 | 0,48501307 |
| 387923_at | SERP2 | -0,17745799 | 4,36224408 | -1,27896099 | 0,22096634 | 0,48501307 |
| 741_at | ZNHIT2 | -0,10423957 | 6,73427816 | -1,27879179 | 0,22102431 | 0,48508502 |
| 286042_at | FAM86B3P | -0,14280118 | 5,0745154 | -1,27861707 | 0,22108418 | 0,48516114 |
| 26257_at | NKX2-8 | -0,13887535 | 5,63295515 | -1,27845825 | 0,2211386 | 0,4852253 |
| 5293_at | PIK3CD | 0,12008379 | 5,60872146 | 1,27827638 | 0,22120095 | 0,48530682 |
| 286205_at | SCAI | -0,10065566 | 5,49066104 | -1,27818776 | 0,22123133 | 0,4853182 |
| 23505_at | TMEM131 | 0,10983137 | 7,34945143 | 1,27779553 | 0,22136585 | 0,485558 |
| 390940_at | PINLYP | -0,13572677 | 4,88013622 | -1,27770585 | 0,22139661 | 0,48557019 |
| 83889_at | WDR87 | -0,16713787 | 4,00816059 | -1,27751776 | 0,22146115 | 0,48565644 |
| 5034_at | P4HB | 0,07514276 | 11,6143565 | 1,27710133 | 0,22160409 | 0,48591457 |
| 94031_at | HTRA3 | -0,12864641 | 5,42516512 | -1,27684124 | 0,2216934 | 0,48596394 |
| 51291_at | GMIP | 0,10812896 | 6,81891578 | 1,27681473 | 0,2217025 | 0,48596394 |
| 84876_at | ORAI1 | -0,27098415 | 8,00277628 | -1,27677847 | 0,22171496 | 0,48596394 |
| 101927446_a | LOC10192744 | 0,09572791 | 3,16426299 | 1,27674191 | 0,22172751 | 0,48596394 |
| 55749_at | CCAR1 | -0,07699697 | 9,35000732 | -1,2764531 | 0,22182673 | 0,48608647 |
| 54785_at | BORCS6 | 0,08157915 | 6,01229659 | 1,27643227 | 0,22183389 | 0,48608647 |
| 93589_at | CACNA2D4 | -0,11190786 | 4,23341016 | -1,27614808 | 0,22193155 | 0,48624517 |
| 23300_at | ATMIN | 0,12186725 | 9,64765164 | 1,27597424 | 0,22199132 | 0,48632079 |
| 1991_at | ELANE | -0,11553923 | 4,94580751 | -1,27582438 | 0,22204284 | 0,48637836 |

| | | | | | | |
|---|---|---|---|---|---|---|
| 51780_at | KDM3B | 0,15811762 | 9,21640048 | 1,27570852 | 0,22208269 | 0,48641032 |
| 54538_at | ROBO4 | -0,12631629 | 4,97832609 | -1,27546846 | 0,22216526 | 0,48653586 |
| 197003_at | MGC15885 | -0,14789988 | 4,47286866 | -1,27533791 | 0,22221018 | 0,48656206 |
| 57617_at | VPS18 | 0,13015411 | 6,9587903 | 1,27528685 | 0,22222774 | 0,48656206 |
| 9260_at | PDLIM7 | -0,18460917 | 6,93804 | -1,27519649 | 0,22225884 | 0,48657483 |
| 9398_at | CD101 | -0,11945909 | 4,09220999 | -1,27488989 | 0,22236437 | 0,48675055 |
| 27161_at | AGO2 | -0,24993512 | 7,47505447 | -1,27472313 | 0,22242178 | 0,48677646 |
| 8658_at | TNKS | 0,14177248 | 6,52825643 | 1,27463603 | 0,22245177 | 0,48677646 |
| 8809_at | IL18R1 | 0,55533989 | 4,6484584 | 1,27448579 | 0,22250352 | 0,48677646 |
| 57763_at | ANKRA2 | 0,56921179 | 7,51290917 | 1,2744362 | 0,2225206 | 0,48677646 |
| 37_at | ACADVL | 0,19414192 | 9,56589196 | 1,27437406 | 0,222542 | 0,48677646 |
| 202374_at | STK32A | 0,16074651 | 5,04563646 | 1,2743535 | 0,22254909 | 0,48677646 |
| 340527_at | NHSL2 | -0,08928579 | 4,52013919 | -1,27434186 | 0,2225531 | 0,48677646 |
| 1131_at | CHRM3 | -0,1117007 | 4,53486147 | -1,27418273 | 0,22260792 | 0,4867892 |
| 149233_at | IL23R | -0,08164256 | 3,09544311 | -1,27417825 | 0,22260946 | 0,4867892 |
| 10298_at | PAK4 | -0,10310561 | 8,38489678 | -1,27405271 | 0,22265272 | 0,48682854 |
| 83483_at | PLVAP | -0,15167934 | 5,55646304 | -1,2737376 | 0,22276133 | 0,48701073 |
| 27165_at | GLS2 | 0,20405975 | 4,95480787 | 1,27359615 | 0,2228101 | 0,48706208 |
| 78997_at | GDAP1L1 | -0,12693595 | 5,089344 | -1,27342398 | 0,22286947 | 0,48707187 |
| 253039_at | PSMD5-AS1 | 0,46418716 | 6,96725189 | 1,27339888 | 0,22287813 | 0,48707187 |
| 3908_at | LAMA2 | -0,10050587 | 4,48733016 | -1,27336319 | 0,22289044 | 0,48707187 |
| 51725_at | FBXO40 | -0,08924392 | 4,35139717 | -1,27313893 | 0,2229678 | 0,48718566 |
| 7634_at | ZNF80 | 0,1063885 | 3,18483661 | 1,27305563 | 0,22299654 | 0,48719318 |
| 22872_at | SEC31A | 0,26439345 | 9,8769923 | 1,27291144 | 0,22304629 | 0,48724663 |
| 2109_at | ETFB | -0,23790723 | 9,21048625 | -1,27267038 | 0,22312949 | 0,48737311 |
| 57167_at | SALL4 | -0,17168029 | 5,89973018 | -1,27190402 | 0,22339417 | 0,4878959 |
| 199870_at | FAM76A | 0,09697107 | 6,01817711 | 1,2717687 | 0,22344093 | 0,48792925 |
| 148979_at | GLIS1 | -0,15217249 | 5,36647437 | -1,27171322 | 0,2234601 | 0,48792925 |
| 28662_at | TRAV21 | -0,10626899 | 3,39200404 | -1,2714663 | 0,22354545 | 0,48800504 |
| 100506497_a | LOC10050649 | -0,11299007 | 3,15146401 | -1,27140316 | 0,22356728 | 0,48800504 |
| 5549_at | PRELP | -0,16033388 | 5,21348755 | -1,27139294 | 0,22357081 | 0,48800504 |
| 2837_at | UTS2R | -0,12709128 | 7,42776399 | -1,27103233 | 0,22369551 | 0,48821745 |
| 1620_at | BRINP1 | -0,12568471 | 4,54249074 | -1,2709041 | 0,22373987 | 0,48821745 |
| 3004_at | GZMM | -0,15357393 | 4,95585367 | -1,27089172 | 0,22374415 | 0,48821745 |
| 10443_at | N4BP2L2 | 0,19140706 | 6,58242854 | 1,27068582 | 0,22381539 | 0,48831758 |
| 105372483_a | LOC10537248 | -0,13718701 | 4,88628229 | -1,27020939 | 0,2239803 | 0,48862203 |
| 200312_at | RNF215 | 0,1082381 | 6,15802297 | 1,27008602 | 0,22402302 | 0,4886546 |
| 2803_at | GOLGA4 | -0,29021643 | 9,00823276 | -1,26989669 | 0,22408859 | 0,4886546 |
| 10564_at | ARFGEF2 | 0,17598006 | 8,65087265 | 1,26989639 | 0,2240887 | 0,4886546 |
| 100506388_a | LOC10050638 | -0,11331346 | 3,47964828 | -1,26987329 | 0,2240967 | 0,4886546 |
| 284578_at | LOC284578 | -0,10446577 | 3,75390145 | -1,2697291 | 0,22414665 | 0,48869702 |
| 9283_at | GPR37L1 | -0,11025493 | 5,26618314 | -1,26950563 | 0,22422408 | 0,48869702 |
| 85460_at | ZNF518B | 0,12730729 | 7,31234568 | 1,26943449 | 0,22424873 | 0,48869702 |
| 6932_at | TCF7 | -0,09060008 | 5,02268679 | -1,26942138 | 0,22425327 | 0,48869702 |
| 57626_at | KLHL1 | 0,07134066 | 3,32858668 | 1,26930168 | 0,22429476 | 0,48869702 |
| 25763_at | HYPM | -0,16188707 | 4,33459665 | -1,26927331 | 0,22430459 | 0,48869702 |
| 148145_at | LINC00906 | -0,08197204 | 3,60479665 | -1,26924238 | 0,22431531 | 0,48869702 |
| 401014_at | TEX41 | -0,11038518 | 3,11056072 | -1,26923142 | 0,22431911 | 0,48869702 |
| 1671_at | DEFA6 | -0,13949589 | 4,74134518 | -1,2690135 | 0,22439467 | 0,48871952 |
| 4005_at | LMO2 | 0,34902991 | 5,63780061 | 1,26900292 | 0,22439834 | 0,48871952 |

| | | | | | | |
|---|---|---|---|---|---|---|
| 414236_at | C10orf55 | 0,16943871 | 3,33164124 | 1,26882873 | 0,22445874 | 0,48871952 |
| 54946_at | SLC41A3 | 0,20206437 | 8,7653679 | 1,26878851 | 0,22447269 | 0,48871952 |
| 148741_at | ANKRD35 | 0,33527222 | 5,44930257 | 1,26873092 | 0,22449267 | 0,48871952 |
| 6397_at | SEC14L1 | 0,11776082 | 7,65196135 | 1,26869265 | 0,22450594 | 0,48871952 |
| 9024_at | BRSK2 | -0,09497937 | 4,92290566 | -1,2686895 | 0,22450703 | 0,48871952 |
| 2315_at | MLANA | -0,09820947 | 3,45001236 | -1,26830882 | 0,22463912 | 0,48895178 |
| 91612_at | CHURC1 | 0,45624992 | 8,51441592 | 1,26807583 | 0,22471999 | 0,48907254 |
| 6253_at | RTN2 | 0,17587391 | 5,96510753 | 1,26799329 | 0,22474864 | 0,48907965 |
| 163778_at | SPRR4 | -0,18185356 | 6,3708956 | -1,2678806 | 0,22478777 | 0,48910954 |
| 89122_at | TRIM4 | 0,59202171 | 7,37414633 | 1,2677186 | 0,22484402 | 0,48914893 |
| 56107_at | PCDHGA9 | -0,12493185 | 5,94130152 | -1,26764419 | 0,22486986 | 0,48914893 |
| 153241_at | CEP120 | -0,19491563 | 6,64343496 | -1,2676091 | 0,22488205 | 0,48914893 |
| 81857_at | MED25 | 0,11083234 | 8,32358908 | 1,26748833 | 0,224924 | 0,48918114 |
| 56987_at | BBX | 0,18123025 | 8,41150615 | 1,26742027 | 0,22494765 | 0,48918114 |
| 55815_at | TSNAXIP1 | 0,1305085 | 5,55354222 | 1,26713374 | 0,22504721 | 0,48934241 |
| 100128731_a | OST4 | 0,18096734 | 11,6759769 | 1,26697561 | 0,22510217 | 0,48936208 |
| 114794_at | ELFN2 | -0,28181721 | 5,309523 | -1,26696154 | 0,22510706 | 0,48936208 |
| 124773_at | C17orf64 | -0,16419972 | 4,26708934 | -1,26688053 | 0,22513522 | 0,48936807 |
| 442191_at | OR14J1 | -0,15665637 | 4,14890647 | -1,26650598 | 0,22526547 | 0,48957543 |
| 146853_at | C17orf50 | -0,11429851 | 6,00059307 | -1,26646001 | 0,22528145 | 0,48957543 |
| 477_at | ATP1A2 | -0,14185504 | 4,04908102 | -1,2659134 | 0,22547164 | 0,48993348 |
| 11074_at | TRIM31 | 0,52807501 | 7,06609543 | 1,26581621 | 0,22550547 | 0,48995172 |
| 639_at | PRDM1 | 0,2083651 | 5,23293908 | 1,26566681 | 0,22555748 | 0,48996569 |
| 255411_at | LINC00935 | -0,11406779 | 3,31651184 | -1,26565162 | 0,22556277 | 0,48996569 |
| 25906_at | ANAPC15 | -0,09316976 | 8,68698815 | -1,26548847 | 0,22561958 | 0,48999061 |
| 200197_at | TMEM51-AS1 | -0,08069951 | 4,54935626 | -1,26543836 | 0,22563703 | 0,48999061 |
| 117579_at | RLN3 | -0,12853233 | 4,64833605 | -1,26539011 | 0,22565384 | 0,48999061 |
| 3201_at | HOXA4 | 0,07791602 | 6,78745696 | 1,26532649 | 0,225676 | 0,48999061 |
| 7780_at | SLC30A2 | -0,14051342 | 5,52404338 | -1,26504009 | 0,22577577 | 0,49015199 |
| 54454_at | ATAD2B | 0,12831125 | 7,29299553 | 1,26495249 | 0,22580629 | 0,49015515 |
| 5969_at | REG1P | -0,1670839 | 4,14121502 | -1,26488986 | 0,22582812 | 0,49015515 |
| 2199_at | FBLN2 | 0,18026123 | 5,5795773 | 1,26470454 | 0,22589271 | 0,49024011 |
| 474_at | ATOH1 | -0,13864691 | 5,6002029 | -1,26441485 | 0,2259937 | 0,49040404 |
| 27332_at | ZNF638 | 0,1341778 | 8,83990543 | 1,26426482 | 0,22604602 | 0,49044883 |
| 5315_at | PKM | -0,45044383 | 10,6899245 | -1,26412105 | 0,22609616 | 0,49044883 |
| 100874195_a | DIAPH3-AS1 | -0,07442675 | 3,18906324 | -1,26405921 | 0,22611774 | 0,49044883 |
| 643365_at | LINC00452 | -0,11998893 | 4,79197137 | -1,26399891 | 0,22613878 | 0,49044883 |
| 128859_at | BPIFB6 | -0,16925601 | 4,43276328 | -1,26399068 | 0,22614164 | 0,49044883 |
| 100131347_a | LOC10013134 | -0,15407618 | 3,88668857 | -1,26380747 | 0,22620557 | 0,49048638 |
| 5224_at | PGAM2 | -0,14117337 | 4,79418807 | -1,26379511 | 0,22620988 | 0,49048638 |
| 149134_at | LINC01341 | -0,16086107 | 5,0840899 | -1,26372081 | 0,22623581 | 0,49048738 |
| 100288122_a | LDLRAD4-AS1 | -0,12984359 | 4,30296833 | -1,26363309 | 0,22626642 | 0,49049855 |
| 775_at | CACNA1C | -0,11277932 | 4,00629298 | -1,2633183 | 0,22637631 | 0,49068155 |
| 1615_at | DARS | -0,24178931 | 10,1101743 | -1,26323144 | 0,22640664 | 0,49069208 |
| 913_at | CD1E | -0,09221708 | 3,60023442 | -1,26302267 | 0,22647956 | 0,49077205 |
| 10744_at | PTTG2 | -0,09913631 | 3,26192673 | -1,2629799 | 0,2264945 | 0,49077205 |
| 6459_at | SH3GL1P2 | -0,17706617 | 4,92168469 | -1,2625285 | 0,22665222 | 0,49100939 |
| 221395_at | ADGRF5 | 0,15318313 | 3,93300048 | 1,26246358 | 0,2266749 | 0,49100939 |
| 2525_at | FUT3 | 0,13658608 | 6,53850948 | 1,26244757 | 0,2266805 | 0,49100939 |
| 56916_at | SMARCAD1 | 0,29103663 | 7,79363681 | 1,26214661 | 0,22678572 | 0,49118206 |

| | | | | | | |
|---|---|---|---|---|---|---|
| 83549_at | UCK1 | 0,10014349 | 8,57678963 | 1,26196175 | 0,22685036 | 0,49126683 |
| 8659_at | ALDH4A1 | 0,19916157 | 6,92696881 | 1,26180101 | 0,22690658 | 0,49133335 |
| 440956_at | IQCF6 | -0,17405316 | 4,26743555 | -1,26164506 | 0,22696114 | 0,49139624 |
| 653390_at | RRN3P2 | 0,1668687 | 3,56987293 | 1,26122643 | 0,22710764 | 0,49165818 |
| 51604_at | PIGT | 0,20291931 | 10,414605 | 1,26069263 | 0,22729456 | 0,49200754 |
| 4059_at | BCAM | 0,1146428 | 6,83495237 | 1,26058393 | 0,22733263 | 0,49203467 |
| 83998_at | REG4 | -0,11626442 | 3,81866482 | -1,26046969 | 0,22737266 | 0,49206601 |
| 140766_at | ADAMTS14 | -0,16779817 | 5,49252558 | -1,26037372 | 0,22740629 | 0,4920835 |
| 104502416_a | FOXP1-AS1 | -0,12417362 | 3,08932183 | -1,26003422 | 0,22752528 | 0,49228568 |
| 51567_at | TDP2 | 0,25890631 | 10,5143323 | 1,25983114 | 0,22759648 | 0,49238442 |
| 85458_at | DIXDC1 | -0,10602806 | 5,06025611 | -1,25947711 | 0,22772064 | 0,49259772 |
| 101928222_a | LOC10192822 | 0,119169 | 5,31824659 | 1,25927182 | 0,22779266 | 0,4926754 |
| 54896_at | PQLC2 | 0,12414471 | 7,29012778 | 1,25914577 | 0,22783689 | 0,4926754 |
| 5207_at | PFKFB1 | -0,1284775 | 4,57100084 | -1,25910196 | 0,22785227 | 0,4926754 |
| 2239_at | GPC4 | -0,27688688 | 6,13828191 | -1,25908319 | 0,22785886 | 0,4926754 |
| 83449_at | PMFBP1 | -0,1112407 | 4,73209257 | -1,25846614 | 0,2280755 | 0,49308847 |
| 925_at | CD8A | 0,13505422 | 5,36585514 | 1,25806128 | 0,22821773 | 0,49315103 |
| 319101_at | KRT73 | -0,09655786 | 5,07410908 | -1,25799999 | 0,22823927 | 0,49315103 |
| 23466_at | CBX6 | 0,14264171 | 9,08278676 | 1,25798504 | 0,22824452 | 0,49315103 |
| 55958_at | KLHL9 | 0,30827834 | 6,81840499 | 1,25792848 | 0,2282644 | 0,49315103 |
| 117531_at | TMC1 | -0,11974981 | 3,64714848 | -1,25785274 | 0,22829102 | 0,49315103 |
| 57158_at | JPH2 | -0,11151769 | 5,95526129 | -1,25781529 | 0,22830418 | 0,49315103 |
| 3690_at | ITGB3 | -0,10816305 | 4,41536729 | -1,25780995 | 0,22830606 | 0,49315103 |
| 101929074_a | PIK3CD-AS2 | 0,11754197 | 4,15151089 | 1,25776149 | 0,22832309 | 0,49315103 |
| 441168_at | FAM26F | 0,13057819 | 4,6240831 | 1,25770456 | 0,22834311 | 0,49315103 |
| 2621_at | GAS6 | -0,2033005 | 6,86589509 | -1,25765525 | 0,22836044 | 0,49315103 |
| 84461_at | NEURL4 | -0,08666495 | 6,49284882 | -1,25748625 | 0,22841986 | 0,49322405 |
| 1621_at | DBH | -0,11236841 | 5,22666637 | -1,25729244 | 0,22848802 | 0,49331593 |
| 6935_at | ZEB1 | -0,06933073 | 3,35715257 | -1,25719903 | 0,22852088 | 0,49333158 |
| 4123_at | MAN2C1 | 0,11388531 | 6,97560703 | 1,25704968 | 0,22857342 | 0,49338971 |
| 8344_at | HIST1H2BE | -0,2320218 | 7,35451665 | -1,25670815 | 0,2286936 | 0,49359383 |
| 3181_at | HNRNPA2B1 | -0,23524903 | 10,1098387 | -1,25641382 | 0,22879722 | 0,49376213 |
| 613266_at | LOC613266 | 0,16012616 | 4,07218393 | 1,25626169 | 0,22885078 | 0,49377553 |
| 1654_at | DDX3X | -0,07788261 | 8,7434881 | -1,2562506 | 0,22885469 | 0,49377553 |
| 101927651_a | LOC10192765 | -0,12439566 | 4,03377375 | -1,25605496 | 0,2289236 | 0,49386889 |
| 8867_at | SYNJ1 | 0,17553342 | 6,63920199 | 1,25575687 | 0,22902862 | 0,49404012 |
| 55025_at | FLJ20712 | -0,13599365 | 4,5483346 | -1,25558751 | 0,22908831 | 0,49411354 |
| 3744_at | KCNA10 | -0,16193251 | 4,65448779 | -1,25531131 | 0,22918567 | 0,4942682 |
| 51337_at | THEM6 | -0,1714086 | 9,3121564 | -1,25517252 | 0,2292346 | 0,49431839 |
| 64848_at | YTHDC2 | -0,12770421 | 7,20349591 | -1,2548821 | 0,22933703 | 0,49448391 |
| 2306_at | FOXD2 | 0,24538973 | 5,21988045 | 1,25479763 | 0,22936683 | 0,49449281 |
| 374897_at | SBSN | -0,18048805 | 4,36887872 | -1,25464734 | 0,22941986 | 0,49455178 |
| 754_at | PTTG1IP | 0,33586666 | 11,101928 | 1,25453676 | 0,22945888 | 0,49458054 |
| 6530_at | SLC6A2 | -0,13868066 | 4,67833791 | -1,25437357 | 0,22951647 | 0,49463426 |
| 10935_at | PRDX3 | -0,07232949 | 11,3523395 | -1,25432064 | 0,22953515 | 0,49463426 |
| 54532_at | USP53 | 0,3919515 | 7,06275519 | 1,2540699 | 0,22962368 | 0,49465106 |
| 5049_at | PAFAH1B2 | 0,06052933 | 6,75216766 | 1,25400868 | 0,2296453 | 0,49465106 |
| 388714_at | FMO6P | -0,1370981 | 3,78534791 | -1,25396636 | 0,22966024 | 0,49465106 |
| 79805_at | VASH2 | 0,33134532 | 5,4866481 | 1,25395666 | 0,22966367 | 0,49465106 |
| 4802_at | NFYC | -0,13278133 | 7,86912767 | -1,25393493 | 0,22967134 | 0,49465106 |

| | | | | | | |
|---|---|---|---|---|---|---|
| 26173_at | INTS1 | -0,10571499 | 7,03173615 | -1,25382649 | 0,22970964 | 0,49467823 |
| 8453_at | CUL2 | -0,12082972 | 7,42677022 | -1,25366838 | 0,22976549 | 0,49470229 |
| 285343_at | TCAIM | 0,1710364 | 7,44043336 | 1,25364945 | 0,22977218 | 0,49470229 |
| 9727_at | RAB11FIP3 | 0,08893375 | 6,18419247 | 1,25354291 | 0,22980982 | 0,494713 |
| 91137_at | SLC25A46 | -0,10884891 | 9,18146296 | -1,25339865 | 0,22986079 | 0,494713 |
| 26259_at | FBXW8 | 0,28106498 | 6,22757326 | 1,25335668 | 0,22987563 | 0,494713 |
| 2785_at | GNG3 | -0,1036288 | 5,06273873 | -1,25334465 | 0,22987987 | 0,494713 |
| 85285_at | KRTAP4-1 | 0,125374 | 4,35273027 | 1,25317705 | 0,22993911 | 0,4947852 |
| 5611_at | DNAJC3 | 0,16016252 | 8,49802503 | 1,25303699 | 0,22998862 | 0,49483646 |
| 1039_at | CDR2 | -0,16054947 | 8,0395057 | -1,25284919 | 0,23005503 | 0,49492405 |
| 221191_at | PRSS54 | -0,11512581 | 4,06269755 | -1,2527637 | 0,23008526 | 0,49493382 |
| 101927657_a | LOC10192765 | -0,13981002 | 3,46523965 | -1,25257037 | 0,23015364 | 0,49496243 |
| 752_at | FMNL1 | -0,13625277 | 5,98533052 | -1,25249246 | 0,2301812 | 0,49496243 |
| 22998_at | LIMCH1 | 0,64047898 | 8,87836509 | 1,2523487 | 0,23023206 | 0,49496243 |
| 685_at | BTC | 0,48072457 | 4,77957291 | 1,25232111 | 0,23024183 | 0,49496243 |
| 8031_at | NCOA4 | 0,18943467 | 10,6708889 | 1,25230753 | 0,23024663 | 0,49496243 |
| 257397_at | TAB3 | 0,12162823 | 6,10099274 | 1,2522903 | 0,23025273 | 0,49496243 |
| 25970_at | SH2B1 | 0,07322866 | 7,19608222 | 1,25183104 | 0,23041529 | 0,49516102 |
| 100652768_a | LOC10065276 | -0,08684487 | 5,96410014 | -1,25182322 | 0,23041805 | 0,49516102 |
| 100652824_a | KIAA2012 | -0,12059683 | 3,62822259 | -1,25181143 | 0,23042223 | 0,49516102 |
| 10103_at | TSPAN1 | -0,19445981 | 9,39920047 | -1,25171023 | 0,23045806 | 0,49518279 |
| 653483_at | MLLT4-AS1 | 0,26241342 | 4,04273847 | 1,25134386 | 0,23058783 | 0,49532567 |
| 81492_at | RSPH6A | -0,15934139 | 5,62561591 | -1,25132185 | 0,23059562 | 0,49532567 |
| 10611_at | PDLIM5 | 0,21423289 | 7,5136615 | 1,25130469 | 0,2306017 | 0,49532567 |
| 23620_at | NTSR2 | -0,17668748 | 4,68745392 | -1,25120308 | 0,2306377 | 0,49534776 |
| 100861546_a | LINC00564 | 0,10909592 | 2,95717435 | 1,25090481 | 0,23074341 | 0,49551953 |
| 79609_at | VCPKMT | -0,09897593 | 7,04912131 | -1,25076814 | 0,23079185 | 0,49556833 |
| 6547_at | SLC8A3 | -0,15356174 | 4,759314 | -1,25063386 | 0,23083946 | 0,49561531 |
| 119548_at | PNLIPRP3 | -0,26076758 | 3,72968646 | -1,25052425 | 0,23087833 | 0,49562285 |
| 401588_at | ZNF674-AS1 | -0,15031848 | 5,61176437 | -1,25047884 | 0,23089443 | 0,49562285 |
| 28685_at | TRAV8-1 | -0,09707537 | 5,11771836 | -1,25024103 | 0,23097878 | 0,49569934 |
| 93624_at | TADA2B | 0,17315253 | 8,27509286 | 1,25020654 | 0,23099101 | 0,49569934 |
| 100169752_a | TEX36-AS1 | -0,09885089 | 3,09663036 | -1,250037 | 0,23105116 | 0,49569934 |
| 79831_at | KDM8 | -0,08166061 | 5,26733899 | -1,25002024 | 0,23105711 | 0,49569934 |
| 29998_at | GLTSCR1 | 0,08594714 | 7,38499609 | 1,25001566 | 0,23105873 | 0,49569934 |
| 3363_at | HTR7 | -0,15267485 | 5,29254632 | -1,24972281 | 0,23116267 | 0,49581949 |
| 130132_at | RFTN2 | 0,08659645 | 3,60776832 | 1,2497128 | 0,23116622 | 0,49581949 |
| 4856_at | NOV | -0,07920298 | 4,8199883 | -1,24923302 | 0,23133657 | 0,4960847 |
| 283489_at | CHAMP1 | -0,31054731 | 8,70791433 | -1,24921951 | 0,23134137 | 0,4960847 |
| 337878_at | KRTAP7-1 | -0,09732557 | 4,01584995 | -1,24914331 | 0,23136844 | 0,49608751 |
| 144699_at | FBXL14 | 0,16372074 | 4,64599763 | 1,24893332 | 0,23144304 | 0,49612661 |
| 57642_at | COL20A1 | -0,10580192 | 4,63822265 | -1,248879 | 0,23146234 | 0,49612661 |
| 84332_at | DYDC2 | -0,10306013 | 3,65957841 | -1,24880927 | 0,23148711 | 0,49612661 |
| 25927_at | CNRIP1 | -0,12768686 | 4,57575261 | -1,24874975 | 0,23150827 | 0,49612661 |
| 56672_at | AKIP1 | -0,15933002 | 7,47699775 | -1,2487205 | 0,23151866 | 0,49612661 |
| 283848_at | CES4A | 0,16291556 | 5,8153272 | 1,24855649 | 0,23157695 | 0,49612661 |
| 7738_at | ZNF184 | 0,19270282 | 7,15649926 | 1,2484693 | 0,23160795 | 0,49612661 |
| 253264_at | ZNF503-AS1 | 0,21352351 | 3,56820782 | 1,24844063 | 0,23161814 | 0,49612661 |
| 475_at | ATOX1 | -0,11982991 | 9,64478834 | -1,24843969 | 0,23161847 | 0,49612661 |
| 201191_at | SAMD14 | -0,1095544 | 5,16888688 | -1,24820106 | 0,23170332 | 0,49625317 |

| | | | | | | |
|---|---|---|---|---|---|---|
| 153328_at | SLC25A48 | -0,15212071 | 4,75891515 | -1,24770756 | 0,23187886 | 0,49642378 |
| 164633_at | CABP7 | -0,10679269 | 5,21115859 | -1,24766037 | 0,23189566 | 0,49642378 |
| 11237_at | RNF24 | 0,19234117 | 8,53311661 | 1,24762382 | 0,23190866 | 0,49642378 |
| 100507099_a | FRY-AS1 | 0,09702404 | 3,28462442 | 1,24761989 | 0,23191006 | 0,49642378 |
| 259234_at | DSCR10 | -0,12356907 | 5,39779151 | -1,24758694 | 0,23192179 | 0,49642378 |
| 29092_at | LINC00339 | -0,12713416 | 6,94290261 | -1,24754249 | 0,2319376 | 0,49642378 |
| 11185_at | INMT | -0,11244521 | 5,16757396 | -1,24716301 | 0,23207269 | 0,49665773 |
| 54207_at | KCNK10 | -0,083262 | 3,89701014 | -1,24697294 | 0,23214037 | 0,49674739 |
| 91133_at | L3MBTL4 | 0,09625329 | 3,55124806 | 1,24686817 | 0,23217768 | 0,49675909 |
| 8614_at | STC2 | -0,66577481 | 7,04567421 | -1,24681278 | 0,23219742 | 0,49675909 |
| 3295_at | HSD17B4 | 0,1275878 | 9,97592297 | 1,24628073 | 0,232387 | 0,49710946 |
| 417_at | ART1 | -0,12597069 | 4,76009266 | -1,24614554 | 0,23243519 | 0,49715734 |
| 255743_at | NPNT | 0,21615484 | 4,90095118 | 1,24588648 | 0,23252755 | 0,49729968 |
| 10089_at | KCNK7 | -0,11658105 | 5,52031497 | -1,24569411 | 0,23259616 | 0,49733978 |
| 9467_at | SH3BP5 | 0,20737546 | 8,16615659 | 1,24568913 | 0,23259794 | 0,49733978 |
| 85407_at | NKD1 | -0,12879668 | 4,79936508 | -1,24555586 | 0,23264547 | 0,49738529 |
| 1153_at | CIRBP | 0,31390623 | 8,34312758 | 1,24546831 | 0,23267671 | 0,49738529 |
| 91369_at | ANKRD40 | -0,17474286 | 8,44851502 | -1,24541233 | 0,23269668 | 0,49738529 |
| 101928943_a | LOC10192894 | -0,11726635 | 3,35554613 | -1,24528484 | 0,23274218 | 0,49739348 |
| 221458_at | KIF6 | -0,08724621 | 3,45251596 | -1,24523016 | 0,23276169 | 0,49739348 |
| 51324_at | SPG21 | 0,08587857 | 10,1102897 | 1,24518451 | 0,23277798 | 0,49739348 |
| 1415_at | CRYBB2 | -0,15673363 | 5,4958888 | -1,24502512 | 0,23283487 | 0,49745987 |
| 341567_at | H1FNT | -0,11813834 | 5,23865187 | -1,24483678 | 0,23290211 | 0,49754834 |
| 863_at | CBFA2T3 | -0,13189856 | 4,61844756 | -1,2447442 | 0,23293517 | 0,49756378 |
| 5669_at | PSG1 | -0,13103426 | 3,03864019 | -1,24420699 | 0,23312707 | 0,49780107 |
| 566_at | AZU1 | -0,1436311 | 4,69614493 | -1,24415712 | 0,23314489 | 0,49780107 |
| 10874_at | NMU | -0,35208239 | 5,70565246 | -1,24393633 | 0,23322379 | 0,49780107 |
| 10562_at | OLFM4 | 0,17710618 | 4,45523839 | 1,24393622 | 0,23322383 | 0,49780107 |
| 51057_at | WDPCP | 0,08475394 | 5,86427827 | 1,24383011 | 0,23326176 | 0,49780107 |
| 10400_at | PEMT | -0,26315771 | 8,63301852 | -1,24382874 | 0,23326225 | 0,49780107 |
| 643246_at | MAP1LC3B2 | 0,21053717 | 5,04774739 | 1,24380616 | 0,23327032 | 0,49780107 |
| 23312_at | DMXL2 | 0,14904624 | 7,87565369 | 1,24379031 | 0,23327599 | 0,49780107 |
| 5980_at | REV3L | 0,30229177 | 7,86135942 | 1,24378235 | 0,23327884 | 0,49780107 |
| 201255_at | LRRC45 | -0,19084129 | 7,26680238 | -1,24356176 | 0,23335771 | 0,49788896 |
| 126669_at | SHE | -0,12378677 | 4,46427191 | -1,24352259 | 0,23337172 | 0,49788896 |
| 57711_at | ZNF529 | 0,08348817 | 4,64668582 | 1,24323031 | 0,23347626 | 0,49805683 |
| 7051_at | TGM1 | -0,12012831 | 5,31336896 | -1,2430681 | 0,23353429 | 0,49807019 |
| 56920_at | SEMA3G | 0,26697082 | 5,17270495 | 1,2430313 | 0,23354746 | 0,49807019 |
| 56156_at | TEX13B | -0,154873 | 5,00985208 | -1,24296163 | 0,23357239 | 0,49807019 |
| 646214_at | LOC646214 | -0,11759173 | 6,04302862 | -1,24292376 | 0,23358595 | 0,49807019 |
| 285513_at | GPRIN3 | 0,07657689 | 3,52070731 | 1,24262864 | 0,23369158 | 0,49824027 |
| 10632_at | ATP5L | -0,11057635 | 9,74850497 | -1,2423008 | 0,23380897 | 0,49831387 |
| 125170_at | MIEF2 | 0,48119157 | 7,35653435 | 1,24228338 | 0,23381521 | 0,49831387 |
| 6285_at | S100B | -0,09543041 | 3,35971769 | -1,24224509 | 0,23382892 | 0,49831387 |
| 343035_at | RD3 | -0,10912246 | 4,73060061 | -1,24209874 | 0,23388134 | 0,49831387 |
| 22797_at | TFEC | 0,09843953 | 3,03218474 | 1,24195482 | 0,2339329 | 0,49831387 |
| 116412_at | ZNF837 | 0,08855592 | 6,24438994 | 1,24191574 | 0,23394691 | 0,49831387 |
| 55840_at | EAF2 | 0,15758962 | 6,02272399 | 1,24187505 | 0,23396149 | 0,49831387 |
| 105379045_a | LOC10537904 | -0,32798917 | 5,67273018 | -1,24186853 | 0,23396382 | 0,49831387 |
| 79839_at | CCDC102B | 0,07328287 | 2,84439809 | 1,24181334 | 0,2339836 | 0,49831387 |

| | | | | | | |
|---|---|---|---|---|---|---|
| 26339_at | OR5K1 | -0,11569511 | 3,08330301 | -1,24178358 | 0,23399426 | 0,49831387 |
| 5027_at | P2RX7 | -0,1434936 | 4,01008961 | -1,24173784 | 0,23401066 | 0,49831387 |
| 353274_at | ZNF445 | 0,08734331 | 6,80331663 | 1,24155436 | 0,23407642 | 0,49839882 |
| 101928858_a | LOC10192885 | -0,15674906 | 4,39167246 | -1,24141419 | 0,23412667 | 0,49844521 |
| 283080_at | C10orf126 | -0,08495772 | 3,04848545 | -1,24126216 | 0,23418118 | 0,49844521 |
| 29944_at | PNMA3 | -0,15323328 | 5,78058221 | -1,24123139 | 0,23419221 | 0,49844521 |
| 4799_at | NFX1 | -0,07040252 | 7,01818662 | -1,2412049 | 0,23420171 | 0,49844521 |
| 79698_at | ZMAT4 | 0,19164058 | 5,43246489 | 1,2409395 | 0,23429691 | 0,49859272 |
| 389874_at | ZCCHC13 | -0,12365056 | 4,22924933 | -1,24082402 | 0,23433833 | 0,49862579 |
| 23127_at | COLGALT2 | -0,11105594 | 4,42531345 | -1,24010974 | 0,23459471 | 0,49911618 |
| 6037_at | RNASE3 | -0,15821988 | 5,15641688 | -1,23992715 | 0,23466028 | 0,49920056 |
| 283688_at | LINC00927 | -0,099146 | 3,53970051 | -1,23979032 | 0,23470943 | 0,49924997 |
| 146_at | ADRA1D | -0,09264312 | 4,20643924 | -1,23966204 | 0,23475551 | 0,49929287 |
| 64919_at | BCL11B | 0,17238367 | 6,80193605 | 1,23941586 | 0,23484397 | 0,49942586 |
| 101928079_a | LINC01057 | 0,158484 | 5,6318674 | 1,23917687 | 0,23492987 | 0,49955339 |
| 100506457_a | LOC10050645 | -0,13888509 | 4,04803888 | -1,23893639 | 0,23501634 | 0,49968208 |
| 221711_at | SYCP2L | -0,07772457 | 3,47929844 | -1,23837105 | 0,23521969 | 0,50005926 |
| 10795_at | ZNF268 | 0,17040359 | 6,4774192 | 1,23797498 | 0,23536225 | 0,5003 |
| 9413_at | FAM189A2 | 0,15598486 | 5,77886729 | 1,23782688 | 0,23541557 | 0,5003 |
| 339483_at | MTMR9LP | -0,13066314 | 5,15757082 | -1,23777151 | 0,23543551 | 0,5003 |
| 165918_at | RNF168 | -0,08554181 | 6,00531881 | -1,2376213 | 0,2354896 | 0,5003 |
| 90850_at | ZNF598 | -0,10943802 | 7,41114328 | -1,23759267 | 0,23549991 | 0,5003 |
| 643037_at | C11orf97 | -0,0708821 | 2,86029694 | -1,23758816 | 0,23550153 | 0,5003 |
| 152024_at | LINC00691 | -0,10681407 | 3,2996057 | -1,23755151 | 0,23551474 | 0,5003 |
| 728294_at | D2HGDH | -0,14966882 | 7,80860859 | -1,23721085 | 0,23563746 | 0,50046776 |
| 8322_at | FZD4 | 0,16712023 | 5,87066153 | 1,23718806 | 0,23564567 | 0,50046776 |
| 92304_at | SCGB3A1 | -0,14027919 | 5,91187092 | -1,23661142 | 0,23585353 | 0,500854 |
| 7011_at | TEP1 | 0,11057446 | 5,95813729 | 1,23651665 | 0,23588771 | 0,50087135 |
| 3003_at | GZMK | -0,10777658 | 3,81260924 | -1,23624937 | 0,23598411 | 0,50102083 |
| 344838_at | PAQR9 | -0,11843 | 3,71694131 | -1,23615255 | 0,23601904 | 0,50103082 |
| 56157_at | TEX13A | -0,11745069 | 4,56105144 | -1,23609213 | 0,23604084 | 0,50103082 |
| 147664_at | ERVV-1 | -0,0959762 | 3,76594094 | -1,23588337 | 0,23611617 | 0,50113551 |
| 7718_at | ZNF165 | -0,27865839 | 6,12055551 | -1,23557142 | 0,23622878 | 0,5012728 |
| 80000_at | GREB1L | 0,4026893 | 5,43801813 | 1,23555869 | 0,23623337 | 0,5012728 |
| 145581_at | LRFN5 | -0,13197084 | 3,63557125 | -1,23548793 | 0,23625893 | 0,5012728 |
| 100288432_a | IL10RB-AS1 | -0,14278522 | 4,07852027 | -1,23530259 | 0,23632585 | 0,50135411 |
| 51191_at | HERC5 | 1,11944853 | 5,80007563 | 1,23523765 | 0,23634931 | 0,50135411 |
| 79730_at | NSUN7 | 0,31126389 | 4,00961286 | 1,23480222 | 0,23650662 | 0,50157981 |
| 1823_at | DSC1 | -0,10849506 | 3,31163879 | -1,23473026 | 0,23653263 | 0,50157981 |
| 3207_at | HOXA11 | -0,16771333 | 4,40749957 | -1,23459153 | 0,23658277 | 0,50157981 |
| 8631_at | SKAP1 | -0,09523361 | 4,07353963 | -1,23458854 | 0,23658385 | 0,50157981 |
| 23067_at | SETD1B | 0,14419629 | 9,42631932 | 1,23454354 | 0,23660011 | 0,50157981 |
| 160335_at | TMTC2 | 0,2604518 | 6,39800228 | 1,23451083 | 0,23661194 | 0,50157981 |
| 100507206_a | LINC00943 | -0,14524083 | 3,84948993 | -1,23428969 | 0,23669189 | 0,50162174 |
| 10052_at | GJC1 | -0,32012179 | 6,82150489 | -1,2341529 | 0,23674136 | 0,50162174 |
| 100144604_a | LINC00930 | 0,16553685 | 5,02453304 | 1,23413932 | 0,23674627 | 0,50162174 |
| 367_at | AR | -0,1196402 | 5,77260331 | -1,23409613 | 0,23676189 | 0,50162174 |
| 2593_at | GAMT | -0,10885855 | 7,22511139 | -1,23409604 | 0,23676192 | 0,50162174 |
| 83932_at | SPRTN | -0,17275689 | 6,88822268 | -1,23391072 | 0,23682896 | 0,50162804 |
| 89886_at | SLAMF9 | -0,11057719 | 4,95603886 | -1,23387855 | 0,23684059 | 0,50162804 |

| | | | | | | |
|---|---|---|---|---|---|---|
| 1020_at | CDK5 | 0,1308359 | 8,70038626 | 1,23387184 | 0,23684302 | 0,50162804 |
| 334_at | APLP2 | 0,16098839 | 11,3988215 | 1,23316502 | 0,23709883 | 0,50211465 |
| 5336_at | PLCG2 | -0,08478223 | 4,3273509 | -1,23296206 | 0,23717233 | 0,50221509 |
| 84680_at | ACCS | 0,17821804 | 5,7276505 | 1,2326856 | 0,23727247 | 0,50234514 |
| 153768_at | PRELID2 | 0,15054917 | 6,3336981 | 1,23248658 | 0,23734458 | 0,50234514 |
| 151171_at | LINC01107 | -0,17941196 | 5,2377207 | -1,23245427 | 0,23735629 | 0,50234514 |
| 146705_at | ENTHD2 | 0,18047515 | 6,95899729 | 1,23239034 | 0,23737946 | 0,50234514 |
| 51804_at | SIX4 | 0,10817389 | 6,4418605 | 1,23236081 | 0,23739016 | 0,50234514 |
| 350_at | APOH | -0,12085991 | 3,97905724 | -1,23236065 | 0,23739022 | 0,50234514 |
| 7745_at | ZKSCAN8 | 0,11515828 | 6,0125077 | 1,23227857 | 0,23741997 | 0,5023529 |
| 57154_at | SMURF1 | 0,09627588 | 7,32300626 | 1,23220273 | 0,23744746 | 0,50235589 |
| 125476_at | INO80C | -0,21353615 | 8,82763871 | -1,2319847 | 0,2375265 | 0,50245003 |
| 80342_at | TRAF3IP3 | -0,11796622 | 3,67050223 | -1,23193611 | 0,23754412 | 0,50245003 |
| 363_at | AQP6 | -0,11853595 | 6,06192605 | -1,23174153 | 0,23761469 | 0,50254411 |
| 54841_at | BIVM | 0,2803262 | 6,92598523 | 1,23161307 | 0,23766128 | 0,50258747 |
| 58525_at | WIZ | -0,08584188 | 7,11980423 | -1,23131303 | 0,23777014 | 0,50271539 |
| 10875_at | FGL2 | -0,07921488 | 3,26880992 | -1,23130248 | 0,23777397 | 0,50271539 |
| 58495_at | OVOL2 | 0,22498092 | 7,76280415 | 1,23114675 | 0,23783049 | 0,5027797 |
| 116443_at | GRIN3A | -0,09358272 | 3,58357759 | -1,23105256 | 0,23786467 | 0,50279028 |
| 26493_at | OR8B8 | -0,08518559 | 4,33250997 | -1,23098915 | 0,23788769 | 0,50279028 |
| 374387_at | DKFZp779M0( | -0,13129267 | 4,51362474 | -1,23056706 | 0,23804096 | 0,50305902 |
| 118460_at | EXOSC6 | -0,1735281 | 8,78804925 | -1,23040757 | 0,23809889 | 0,50312625 |
| 1579_at | CYP4A11 | -0,1897855 | 4,66472189 | -1,2301548 | 0,23819073 | 0,5032651 |
| 970_at | CD70 | -0,10418806 | 5,90604756 | -1,22992347 | 0,2382748 | 0,50338753 |
| 170371_at | C10orf128 | -0,09347428 | 3,64793082 | -1,22962417 | 0,23838361 | 0,50356217 |
| 283807_at | FBXL22 | -0,14154365 | 4,76971259 | -1,22939639 | 0,23846644 | 0,50368192 |
| 238_at | ALK | -0,11597025 | 4,8210233 | -1,22932352 | 0,23849295 | 0,50368267 |
| 6560_at | SLC12A4 | 0,1380608 | 6,82321712 | 1,22917621 | 0,23854653 | 0,50373972 |
| 54776_at | PPP1R12C | 0,10442359 | 7,55545547 | 1,2291055 | 0,23857226 | 0,50373972 |
| 55867_at | SLC22A11 | 0,10335298 | 4,67877701 | 1,22887556 | 0,23865593 | 0,50378602 |
| 6769_at | STAC | -0,15910864 | 5,7571397 | -1,22879668 | 0,23868464 | 0,50378602 |
| 489_at | ATP2A3 | 0,08051469 | 5,38160339 | 1,22876005 | 0,23869797 | 0,50378602 |
| 23259_at | DDHD2 | 0,14835877 | 7,53481118 | 1,22870112 | 0,23871942 | 0,50378602 |
| 223_at | ALDH9A1 | 0,14209459 | 9,89317122 | 1,22868366 | 0,23872578 | 0,50378602 |
| 167359_at | NIM1K | -0,10188042 | 4,04879579 | -1,22861408 | 0,23875111 | 0,50378602 |
| 80014_at | WWC2 | 0,24993884 | 6,64674419 | 1,22844667 | 0,23881206 | 0,50385944 |
| 5368_at | PNOC | -0,18476366 | 4,55157292 | -1,22799228 | 0,23897756 | 0,50415339 |
| 2672_at | GFI1 | -0,11439472 | 4,40739722 | -1,22721378 | 0,23926131 | 0,50469673 |
| 55193_at | PBRM1 | 0,15637564 | 7,50803735 | 1,22712805 | 0,23929257 | 0,50470741 |
| 7216_at | TRO | 0,1848243 | 4,430532 | 1,22685071 | 0,23939373 | 0,50486549 |
| 6181_at | RPLP2 | -0,10411611 | 12,6139783 | -1,22676664 | 0,2394244 | 0,5048749 |
| 5899_at | RALB | -0,12883451 | 8,5766425 | -1,22607905 | 0,23967537 | 0,5053488 |
| 2072_at | ERCC4 | 0,1031174 | 6,72476624 | 1,22597269 | 0,23971421 | 0,50537538 |
| 64850_at | ETNPPL | -0,06744628 | 3,0425425 | -1,22568557 | 0,23981908 | 0,50554114 |
| 4939_at | OAS2 | 0,90267224 | 7,77711225 | 1,22555218 | 0,23986781 | 0,50558854 |
| 255057_at | CBARP | -0,1020437 | 5,6576311 | -1,22509618 | 0,24003447 | 0,50588446 |
| 84253_at | GARNL3 | 0,16578734 | 5,16384614 | 1,22497621 | 0,24007833 | 0,50591562 |
| 2957_at | GTF2A1 | -0,18640425 | 7,3701871 | -1,22491207 | 0,24010178 | 0,50591562 |
| 221391_at | OPN5 | -0,10987219 | 3,61240669 | -1,22459327 | 0,24021837 | 0,50601875 |
| 158830_at | CXorf65 | -0,1238841 | 3,21620906 | -1,22459 | 0,24021957 | 0,50601875 |

| | | | | | | |
|---|---|---|---|---|---|---|
| 101928132_a | LOC10192813 | -0,12674499 | 3,88258782 | -1,22456276 | 0,24022953 | 0,50601875 |
| 55684_at | RABL6 | -0,20765487 | 7,52267698 | -1,22410962 | 0,24039534 | 0,50622461 |
| 128602_at | C20orf85 | -0,10517854 | 4,05013529 | -1,22407202 | 0,2404091 | 0,50622461 |
| 84886_at | C1orf198 | -0,09768385 | 9,02576813 | -1,22406197 | 0,24041278 | 0,50622461 |
| 9969_at | MED13 | 0,09735413 | 8,41291013 | 1,22400841 | 0,24043238 | 0,50622461 |
| 85290_at | KRTAP4-3 | -0,11615425 | 3,58522911 | -1,22384995 | 0,24049039 | 0,50627416 |
| 375759_at | C9orf50 | -0,11777061 | 5,36369776 | -1,22373047 | 0,24053414 | 0,50627416 |
| 4644_at | MYO5A | 0,24918799 | 6,54872082 | 1,22363947 | 0,24056746 | 0,50627416 |
| 220441_at | RNF152 | 0,12243679 | 4,1634166 | 1,22359077 | 0,24058529 | 0,50627416 |
| 57404_at | CYP20A1 | 0,11538299 | 7,00248661 | 1,22357451 | 0,24059125 | 0,50627416 |
| 987_at | LRBA | 0,23452833 | 8,68466014 | 1,22351347 | 0,24061361 | 0,50627416 |
| 93426_at | SYCE1 | -0,07815224 | 3,31350933 | -1,22318172 | 0,24073513 | 0,50642377 |
| 51530_at | ZC3HC1 | -0,17170535 | 7,72096879 | -1,22317583 | 0,24073729 | 0,50642377 |
| 347689_at | SOX2-OT | 0,07291935 | 2,95880246 | 1,22292512 | 0,24082917 | 0,50656172 |
| 1149_at | CIDEA | -0,15269415 | 4,90415637 | -1,22281305 | 0,24087025 | 0,50656926 |
| 6746_at | SSR2 | 0,08226426 | 11,377759 | 1,22276284 | 0,24088865 | 0,50656926 |
| 51122_at | COMMD2 | 0,13142024 | 8,87073982 | 1,22270012 | 0,24091165 | 0,50656926 |
| 100689074_a | POU6F2-AS2 | 0,7306797 | 3,78247335 | 1,22259781 | 0,24094915 | 0,50657767 |
| 11193_at | WBP4 | 0,15896216 | 6,58846856 | 1,22251658 | 0,24097894 | 0,50657767 |
| 9349_at | RPL23 | -0,08084147 | 12,3661754 | -1,22241196 | 0,24101731 | 0,50657767 |
| 5008_at | OSM | -0,1159054 | 3,97082744 | -1,22240233 | 0,24102084 | 0,50657767 |
| 8776_at | MTMR1 | 0,15670078 | 8,47597822 | 1,22221465 | 0,24108968 | 0,50666708 |
| 142680_at | SLC34A3 | -0,14660561 | 6,24749003 | -1,22152376 | 0,24134321 | 0,50714457 |
| 387680_at | FAM21A | 0,29743547 | 9,07059576 | 1,22127351 | 0,2414351 | 0,50728232 |
| 57669_at | EPB41L5 | 0,13245739 | 5,93715437 | 1,22111603 | 0,24149294 | 0,5073485 |
| 8928_at | FOXH1 | -0,16845763 | 5,07234375 | -1,22077064 | 0,24161982 | 0,5074234 |
| 100128139_a | LINC01233 | -0,08562343 | 2,65457668 | -1,22075142 | 0,24162689 | 0,5074234 |
| 6531_at | SLC6A3 | -0,11361092 | 6,35601147 | -1,22072749 | 0,24163568 | 0,5074234 |
| 84520_at | C14orf142 | -0,23667048 | 8,10359726 | -1,22068915 | 0,24164977 | 0,5074234 |
| 644096_at | SDHAF1 | -0,18282399 | 7,40440272 | -1,22066051 | 0,24166029 | 0,5074234 |
| 64332_at | NFKBIZ | 0,22364424 | 8,10001598 | 1,2205615 | 0,24169668 | 0,50743737 |
| 90139_at | TSPAN18 | -0,11933956 | 4,9273099 | -1,22049905 | 0,24171964 | 0,50743737 |
| 245932_at | DEFB119 | -0,08566894 | 3,356682 | -1,22040133 | 0,24175556 | 0,50744004 |
| 29070_at | CCDC113 | 0,19260606 | 5,22585942 | 1,22028387 | 0,24179874 | 0,50744004 |
| 84865_at | CCDC142 | 0,13163059 | 6,30133845 | 1,22028061 | 0,24179994 | 0,50744004 |
| 3911_at | LAMA5 | 0,28187572 | 10,0830944 | 1,22016619 | 0,24184201 | 0,50747305 |
| 8542_at | APOL1 | 0,75330767 | 9,63646847 | 1,21984449 | 0,24196032 | 0,50766602 |
| 467_at | ATF3 | 0,13408836 | 6,36662962 | 1,21971298 | 0,2420087 | 0,50770696 |
| 100628307_a | DIO2-AS1 | -0,12181387 | 3,12709105 | -1,21962951 | 0,24203941 | 0,50770696 |
| 8445_at | DYRK2 | 0,28466572 | 8,29207411 | 1,21957122 | 0,24206086 | 0,50770696 |
| 6779_at | STATH | -0,13126962 | 3,09123521 | -1,21948665 | 0,24209198 | 0,50770696 |
| 3426_at | CFI | 0,14114245 | 4,87466428 | 1,2194333 | 0,24211162 | 0,50770696 |
| 101928537_a | LINC01494 | -0,1294392 | 3,5503346 | -1,21918091 | 0,24220452 | 0,50778899 |
| 11252_at | PACSIN2 | -0,14203766 | 8,43310588 | -1,21917351 | 0,24220724 | 0,50778899 |
| 55017_at | C14orf119 | -0,08794601 | 9,24114407 | -1,21902807 | 0,24226079 | 0,50778899 |
| 23607_at | CD2AP | 0,15649377 | 7,40257394 | 1,21900137 | 0,24227062 | 0,50778899 |
| 150291_at | MORC2-AS1 | -0,16456764 | 5,60572147 | -1,21893798 | 0,24229396 | 0,50778899 |
| 6517_at | SLC2A4 | -0,12738698 | 4,38133752 | -1,21889741 | 0,2423089 | 0,50778899 |
| 55267_at | PRR34 | -0,15622268 | 5,97764908 | -1,21868964 | 0,24238543 | 0,5078941 |
| 7802_at | DNALI1 | -0,13433787 | 4,13170685 | -1,21856324 | 0,24243199 | 0,50793448 |

| | | | | | | |
|---|---|---|---|---|---|---|
| 124857_at | WFIKKN2 | -0,1459686 | 5,24193926 | -1,21849419 | 0,24245743 | 0,50793448 |
| 8079_at | MLF2 | -0,14910072 | 10,3439954 | -1,21835704 | 0,24250797 | 0,50798096 |
| 64288_at | ZSCAN31 | 0,47383621 | 7,06226769 | 1,21829085 | 0,24253236 | 0,50798096 |
| 4685_at | NCAM2 | -0,10042732 | 3,24178165 | -1,21804032 | 0,2426247 | 0,50811911 |
| 165829_at | GPR156 | -0,12926218 | 4,16965424 | -1,21718583 | 0,24293986 | 0,50868532 |
| 84167_at | C19orf44 | 0,08002035 | 5,66891216 | 1,2171641 | 0,24294788 | 0,50868532 |
| 51206_at | GP6 | 0,09987925 | 4,46206271 | 1,21675063 | 0,24310049 | 0,50894955 |
| 390616_at | ANKRD34C | 0,11689223 | 3,51857646 | 1,21621453 | 0,24329848 | 0,5093087 |
| 55256_at | ADI1 | -0,28885687 | 9,95392992 | -1,21612106 | 0,24333302 | 0,50932564 |
| 84808_at | PERM1 | -0,1241391 | 5,85442836 | -1,21591048 | 0,24341083 | 0,50941774 |
| 83902_at | KRTAP17-1 | -0,1650169 | 5,3171482 | -1,21585842 | 0,24343007 | 0,50941774 |
| 6515_at | SLC2A3 | -0,11566077 | 3,39389993 | -1,21578731 | 0,24345635 | 0,50941774 |
| 84340_at | GFM2 | -0,38680581 | 7,53209294 | -1,21537968 | 0,24360706 | 0,50963231 |
| 493_at | ATP2B4 | 0,48481484 | 7,50004446 | 1,21536683 | 0,24361181 | 0,50963231 |
| 58190_at | CTDSP1 | 0,11049607 | 8,99105618 | 1,21518391 | 0,24367946 | 0,50971847 |
| 6347_at | CCL2 | -0,10550924 | 3,24242056 | -1,21471622 | 0,2438525 | 0,51002505 |
| 6134_at | RPL10 | -0,18316452 | 6,54575881 | -1,21461395 | 0,24389035 | 0,51004883 |
| 57611_at | ISLR2 | -0,11906094 | 4,97389328 | -1,21433529 | 0,24399351 | 0,51020918 |
| 23491_at | CES3 | -0,11259707 | 5,53166742 | -1,21367099 | 0,24423956 | 0,51066826 |
| 9951_at | HS3ST4 | -0,10367104 | 4,78668189 | -1,21325819 | 0,24439256 | 0,5109327 |
| 10659_at | CELF2 | -0,07370307 | 3,50434614 | -1,21313928 | 0,24443665 | 0,51096941 |
| 101927346_a | LOC10192734 | -0,07863578 | 3,1064779 | -1,21276743 | 0,24457455 | 0,51119776 |
| 100507384_a | LOC10050738 | -0,11752129 | 4,36840042 | -1,21270163 | 0,24459896 | 0,51119776 |
| 93129_at | ORAI3 | 0,31755245 | 7,90785052 | 1,21241555 | 0,2447051 | 0,51136411 |
| 79165_at | LENG1 | -0,10177132 | 6,10826709 | -1,21222219 | 0,24477687 | 0,51145859 |
| 1346_at | COX7A1 | -0,1318321 | 4,39630484 | -1,21202603 | 0,24484968 | 0,51150339 |
| 400569_at | MED11 | 0,15194417 | 7,34239289 | 1,21171891 | 0,24496372 | 0,51150339 |
| 55037_at | PTCD3 | -0,13926413 | 7,39739444 | -1,2117069 | 0,24496818 | 0,51150339 |
| 90736_at | FAM104B | -0,08850777 | 6,4517172 | -1,21169604 | 0,24497222 | 0,51150339 |
| 554235_at | ASPDH | -0,13594719 | 5,39658409 | -1,21168809 | 0,24497517 | 0,51150339 |
| 6039_at | RNASE6 | 0,16738273 | 3,2541396 | 1,21168179 | 0,24497751 | 0,51150339 |
| 79930_at | DOK3 | -0,15600576 | 5,67219866 | -1,21166383 | 0,24498418 | 0,51150339 |
| 11170_at | FAM107A | -0,11453434 | 4,93763325 | -1,21126367 | 0,24513284 | 0,51175832 |
| 26083_at | TBC1D29 | -0,11739492 | 4,17591631 | -1,21088295 | 0,24527435 | 0,51189642 |
| 3857_at | KRT9 | -0,23618939 | 4,29902137 | -1,21076444 | 0,24531841 | 0,51189642 |
| 101927787_a | LOC10192778 | -0,14234034 | 4,47993518 | -1,21049578 | 0,24541832 | 0,51189642 |
| 5803_at | PTPRZ1 | -0,07192268 | 3,04018058 | -1,21042888 | 0,2454432 | 0,51189642 |
| 135154_at | SDHAF4 | 0,21255014 | 5,26318886 | 1,21032796 | 0,24548074 | 0,51189642 |
| 3274_at | HRH2 | -0,1241645 | 4,95178352 | -1,21032191 | 0,24548299 | 0,51189642 |
| 5250_at | SLC25A3 | -0,13935744 | 12,0633561 | -1,21030621 | 0,24548883 | 0,51189642 |
| 51141_at | INSIG2 | 0,2767987 | 8,30436894 | 1,21027808 | 0,24549929 | 0,51189642 |
| 85865_at | GTPBP10 | -0,14648499 | 5,78786131 | -1,21026566 | 0,24550392 | 0,51189642 |
| 58487_at | CREBZF | -0,07845006 | 7,31568987 | -1,21022996 | 0,2455172 | 0,51189642 |
| 100130889_a | PSORS1C3 | 0,18335081 | 3,56380076 | 1,21021695 | 0,24552204 | 0,51189642 |
| 9448_at | MAP4K4 | -0,14342747 | 7,6647056 | -1,21017292 | 0,24553842 | 0,51189642 |
| 65012_at | SLC26A10 | -0,08257177 | 4,57102181 | -1,21014243 | 0,24554976 | 0,51189642 |
| 5890_at | RAD51B | -0,08820756 | 5,27399019 | -1,20992522 | 0,2456306 | 0,51189642 |
| 50807_at | ASAP1 | -0,41326081 | 8,06985657 | -1,20988259 | 0,24564646 | 0,51189642 |
| 728283_at | LOC728283 | 0,1048423 | 4,60245557 | 1,20987271 | 0,24565014 | 0,51189642 |
| 9228_at | DLGAP2 | -0,13175539 | 4,27402077 | -1,20980399 | 0,24567572 | 0,51189642 |

| | | | | | | |
|---|---|---|---|---|---|---|
| 1464_at | CSPG4 | -0,07870644 | 4,13966913 | -1,20958739 | 0,24575635 | 0,51189642 |
| 153396_at | TMEM161B | -0,22156172 | 7,54170659 | -1,20956675 | 0,24576404 | 0,51189642 |
| 55605_at | KIF21A | 0,1387364 | 7,77580864 | 1,20951766 | 0,24578232 | 0,51189642 |
| 342977_at | NANOS3 | -0,13824423 | 6,1360296 | -1,20938003 | 0,24583357 | 0,51189642 |
| 10782_at | ZNF274 | 0,10573291 | 7,75188995 | 1,20935434 | 0,24584314 | 0,51189642 |
| 80301_at | PLEKHO2 | 0,08838383 | 7,55903996 | 1,20921341 | 0,24589563 | 0,51189642 |
| 23089_at | PEG10 | 0,33739482 | 7,3105755 | 1,20917325 | 0,24591059 | 0,51189642 |
| 339929_at | LPP-AS2 | 0,4223707 | 6,14036132 | 1,20917081 | 0,2459115 | 0,51189642 |
| 54345_at | SOX18 | -0,0867197 | 4,69657316 | -1,20914497 | 0,24592112 | 0,51189642 |
| 55504_at | TNFRSF19 | 0,10509758 | 4,61729499 | 1,20910695 | 0,24593528 | 0,51189642 |
| 10204_at | NUTF2 | -0,24188106 | 7,55886602 | -1,20908606 | 0,24594307 | 0,51189642 |
| 149773_at | APCDD1L-AS1 | -0,10086356 | 4,29128799 | -1,20898155 | 0,245982 | 0,51190667 |
| 56256_at | SERTAD4 | -0,14135787 | 6,77686191 | -1,20893018 | 0,24600115 | 0,51190667 |
| 3228_at | HOXC12 | -0,12135023 | 4,34391906 | -1,20833825 | 0,24622178 | 0,51230271 |
| 138046_at | RALYL | -0,14266689 | 3,91068377 | -1,20827691 | 0,24624466 | 0,51230271 |
| 1385_at | CREB1 | 0,21239377 | 8,41313798 | 1,20792993 | 0,24637407 | 0,5125166 |
| 100507632_a | LINC00968 | -0,11288898 | 3,01115288 | -1,20774588 | 0,24644274 | 0,51260409 |
| 101928158_a | LAMA5-AS1 | -0,13337405 | 5,82731929 | -1,20767006 | 0,24647103 | 0,51260759 |
| 284612_at | SYPL2 | -0,18301842 | 5,28005976 | -1,20735448 | 0,24658882 | 0,51264309 |
| 1116_at | CHI3L1 | -0,10418042 | 3,7577628 | -1,20732374 | 0,24660029 | 0,51264309 |
| 400941_at | LINC00487 | -0,09392518 | 3,20828097 | -1,20729921 | 0,24660945 | 0,51264309 |
| 26279_at | PLA2G2D | -0,16691754 | 5,54519617 | -1,20727954 | 0,24661679 | 0,51264309 |
| 64900_at | LPIN3 | 0,09584427 | 6,03468657 | 1,20726783 | 0,24662117 | 0,51264309 |
| 8343_at | HIST1H2BF | 0,20895481 | 8,29266562 | 1,20719168 | 0,2466496 | 0,51264687 |
| 123264_at | SLC51B | -0,09446938 | 4,15666521 | -1,2065188 | 0,24690093 | 0,51309008 |
| 1296_at | COL8A2 | -0,10064154 | 4,68223584 | -1,20647818 | 0,24691611 | 0,51309008 |
| 23398_at | PPWD1 | -0,23432443 | 7,25472248 | -1,20619751 | 0,24702101 | 0,51324882 |
| 440602_at | LOC440602 | -0,13019045 | 5,40820859 | -1,20613122 | 0,24704579 | 0,51324882 |
| 93145_at | OLFM2 | -0,13585539 | 5,5494095 | -1,20594309 | 0,24711613 | 0,51333959 |
| 27141_at | CIDEB | -0,20430733 | 5,2151135 | -1,20506236 | 0,24744562 | 0,51396863 |
| 5697_at | PYY | -0,11701456 | 5,37925537 | -1,2049683 | 0,24748083 | 0,51398634 |
| 27249_at | MMADHC | -0,14296885 | 10,7385616 | -1,20428295 | 0,2477375 | 0,51446393 |
| 60526_at | LDAH | 0,17065281 | 7,27910552 | 1,20393392 | 0,24786829 | 0,51468005 |
| 29968_at | PSAT1 | -0,58336508 | 7,81755281 | -1,20368448 | 0,24796179 | 0,51469517 |
| 57181_at | SLC39A10 | 0,13064529 | 8,35635727 | 1,20365034 | 0,24797459 | 0,51469517 |
| 117_at | ADCYAP1R1 | -0,16896268 | 5,18494183 | -1,20363061 | 0,24798199 | 0,51469517 |
| 84695_at | LOXL3 | -0,18681905 | 5,42283853 | -1,20362939 | 0,24798245 | 0,51469517 |
| 90507_at | SCRN2 | 0,1854396 | 6,86094013 | 1,20353193 | 0,24801899 | 0,51471556 |
| 93377_at | OPALIN | -0,1359031 | 4,43366337 | -1,20328971 | 0,24810983 | 0,51479978 |
| 4995_at | OR3A2 | -0,09356412 | 3,82117927 | -1,2032812 | 0,24811303 | 0,51479978 |
| 144715_at | RAD9B | 0,10533303 | 3,29246775 | 1,20314387 | 0,24816454 | 0,51480797 |
| 79633_at | FAT4 | 0,30752034 | 5,3689953 | 1,20312819 | 0,24817042 | 0,51480797 |
| 129807_at | NEU4 | -0,10233151 | 6,04029945 | -1,20278905 | 0,24829768 | 0,51497484 |
| 6098_at | ROS1 | -0,0972838 | 3,45154196 | -1,20277132 | 0,24830433 | 0,51497484 |
| 10896_at | OCLM | -0,17994175 | 4,39174536 | -1,20246854 | 0,24841799 | 0,51513427 |
| 4842_at | NOS1 | -0,11244291 | 4,32572488 | -1,20238646 | 0,24844881 | 0,51513427 |
| 79874_at | RABEP2 | 0,09148221 | 7,86199538 | 1,20232866 | 0,24847051 | 0,51513427 |
| 283987_at | HID1 | 0,20169409 | 6,40917892 | 1,20225178 | 0,24849938 | 0,51513427 |
| 9563_at | H6PD | 0,11609566 | 6,32224542 | 1,20212136 | 0,24854836 | 0,51513427 |
| 128977_at | C22orf39 | 0,18458091 | 6,96410131 | 1,20205632 | 0,24857279 | 0,51513427 |

| | | | | | | |
|---|---|---|---|---|---|---|
| 100874051_a | LZTS1-AS1 | -0,10511383 | 4,0436607 | -1,20188421 | 0,24863745 | 0,51513427 |
| 285175_at | UNC80 | -0,08680939 | 3,44875873 | -1,20188287 | 0,24863795 | 0,51513427 |
| 152756_at | FAM218A | 0,1227319 | 3,87429186 | 1,20186905 | 0,24864314 | 0,51513427 |
| 327657_at | SERPINA9 | -0,17174575 | 4,92899123 | -1,20185446 | 0,24864862 | 0,51513427 |
| 3024_at | HIST1H1A | -0,19558111 | 4,63171359 | -1,20172133 | 0,24869865 | 0,5151825 |
| 100507316_a | MINCR | -0,61723017 | 5,54014614 | -1,20143002 | 0,24880813 | 0,51529108 |
| 404672_at | GTF2H5 | 0,08201655 | 4,23133899 | 1,20142892 | 0,24880855 | 0,51529108 |
| 64895_at | PAPOLG | 0,16613239 | 6,96742895 | 1,20136835 | 0,24883132 | 0,51529108 |
| 9236_at | CCPG1 | 0,40655267 | 6,28442816 | 1,20076092 | 0,24905975 | 0,51570869 |
| 285577_at | LINC01019 | -0,1223989 | 4,15879931 | -1,20058072 | 0,24912754 | 0,51579362 |
| 63933_at | MCUR1 | -0,315316 | 8,77689978 | -1,20049094 | 0,24916133 | 0,51580692 |
| 150147_at | UMODL1-AS1 | -0,153351 | 4,48116294 | -1,20042134 | 0,24918752 | 0,51580692 |
| 131_at | ADH7 | -0,21900994 | 5,13744267 | -1,20033267 | 0,24922089 | 0,51582057 |
| 219899_at | TBCEL | 0,30544451 | 5,56526366 | 1,20008243 | 0,24931509 | 0,51595486 |
| 163175_at | LGI4 | -0,12559042 | 5,56064577 | -1,20001801 | 0,24933935 | 0,51595486 |
| 55288_at | RHOT1 | 0,11811543 | 8,54779936 | 1,19960278 | 0,24949573 | 0,51622301 |
| 85320_at | ABCC11 | 0,09464705 | 3,7880397 | 1,19945849 | 0,24955009 | 0,51623516 |
| 58484_at | NLRC4 | -0,08256506 | 3,4249562 | -1,19929774 | 0,24961066 | 0,51623516 |
| 249_at | ALPL | -0,1126318 | 4,28936876 | -1,19928719 | 0,24961463 | 0,51623516 |
| 23759_at | PPIL2 | 0,09340557 | 7,6568342 | 1,19924964 | 0,24962879 | 0,51623516 |
| 84282_at | RNF135 | 0,16495633 | 8,1359315 | 1,19920876 | 0,24964419 | 0,51623516 |
| 54619_at | CCNJ | 0,14914386 | 7,52596433 | 1,19916045 | 0,2496624 | 0,51623516 |
| 6392_at | SDHD | -0,14868327 | 8,59671719 | -1,19883714 | 0,24978428 | 0,51643174 |
| 4357_at | MPST | 0,08303973 | 8,17502018 | 1,19867971 | 0,24984364 | 0,51649904 |
| 2700_at | GJA3 | -0,10788565 | 5,06232413 | -1,19859256 | 0,24987651 | 0,51651155 |
| 79657_at | RPAP3 | -0,24435056 | 8,50848844 | -1,19835398 | 0,2499665 | 0,51662647 |
| 51398_at | WDR83OS | 0,07320176 | 9,82665006 | 1,19824392 | 0,25000802 | 0,51662647 |
| 200424_at | TET3 | 0,11247641 | 7,40279696 | 1,19823189 | 0,25001256 | 0,51662647 |
| 145407_at | C14orf37 | 0,08995712 | 3,80467876 | 1,19785701 | 0,25015404 | 0,51686337 |
| 5371_at | PML | 0,21034391 | 7,26620571 | 1,19754881 | 0,25027039 | 0,51698725 |
| 80243_at | PREX2 | -0,12084396 | 3,28940586 | -1,19738023 | 0,25033406 | 0,51698725 |
| 388284_at | C16orf86 | -0,1392041 | 6,08850174 | -1,19734679 | 0,25034669 | 0,51698725 |
| 146923_at | RUNDC1 | 0,16392117 | 7,44111674 | 1,19734016 | 0,25034919 | 0,51698725 |
| 28952_at | CCDC22 | -0,09438613 | 7,14357358 | -1,19730257 | 0,25036339 | 0,51698725 |
| 54903_at | MKS1 | -0,08671301 | 6,7629823 | -1,19727177 | 0,25037502 | 0,51698725 |
| 2843_at | GPR20 | -0,19260556 | 6,15964069 | -1,19717867 | 0,25041019 | 0,51700446 |
| 100874187_a | LINC00559 | -0,1074398 | 3,17731128 | -1,19701991 | 0,25047017 | 0,51707287 |
| 284865_at | LOC284865 | -0,19589085 | 5,16999321 | -1,19654562 | 0,25064943 | 0,51738747 |
| 129446_at | XIRP2 | -0,09951274 | 3,36708412 | -1,19629404 | 0,25074455 | 0,51752837 |
| 55503_at | TRPV6 | 0,2395401 | 6,40676113 | 1,19613281 | 0,25080553 | 0,51759877 |
| 136051_at | ZNF786 | 0,10231924 | 6,94084947 | 1,19579179 | 0,25093454 | 0,51780954 |
| 150000_at | ABCC13 | -0,14767304 | 3,78056135 | -1,19562334 | 0,25099828 | 0,51788559 |
| 286_at | ANK1 | -0,10176964 | 4,77916502 | -1,19544637 | 0,25106526 | 0,51796831 |
| 646268_at | LOC646268 | -0,10771556 | 3,54508425 | -1,19526196 | 0,25113508 | 0,51802892 |
| 26494_at | OR8G1 | -0,20100481 | 3,76686007 | -1,19522672 | 0,25114842 | 0,51802892 |
| 8087_at | FXR1 | 0,12199159 | 7,64966058 | 1,1951016 | 0,2511958 | 0,51807116 |
| 7705_at | ZNF146 | -0,11547804 | 9,21913113 | -1,19499168 | 0,25123742 | 0,51808199 |
| 10277_at | UBE4B | -0,09660478 | 7,84029856 | -1,1949457 | 0,25125484 | 0,51808199 |
| 66008_at | TRAK2 | 0,17572692 | 8,13368888 | 1,19485069 | 0,25129083 | 0,51810073 |
| 100130958_a | SYCE1L | 0,15407506 | 5,18010711 | 1,19449991 | 0,25142373 | 0,5182609 |

| | | | | | | |
|---|---|---|---|---|---|---|
| 254102_at | EHBP1L1 | -0,1196537 | 7,66027761 | -1,19446244 | 0,25143793 | 0,5182609 |
| 23113_at | CUL9 | 0,16569696 | 6,53255329 | 1,19438277 | 0,25146812 | 0,5182609 |
| 79785_at | RERGL | 0,09729432 | 3,14086604 | 1,19436165 | 0,25147613 | 0,5182609 |
| 160428_at | ALDH1L2 | -0,07049885 | 3,2836732 | -1,19420992 | 0,25153364 | 0,51830681 |
| 4147_at | MATN2 | 0,59809383 | 6,40183062 | 1,19416091 | 0,25155222 | 0,51830681 |
| 100505696_a | SH3BP5-AS1 | 0,20316868 | 4,88298516 | 1,19399996 | 0,25161324 | 0,51837709 |
| 29945_at | ANAPC4 | -0,28010923 | 8,04997169 | -1,19379947 | 0,25168927 | 0,51843077 |
| 3614_at | IMPDH1 | -0,2868416 | 8,93814462 | -1,19378932 | 0,25169312 | 0,51843077 |
| 54587_at | MXRA8 | -0,18516189 | 5,54316551 | -1,19344324 | 0,2518244 | 0,51858191 |
| 23597_at | ACOT9 | -0,13135167 | 8,53648266 | -1,19338022 | 0,25184831 | 0,51858191 |
| 1482_at | NKX2-5 | -0,09731918 | 5,88325729 | -1,19334689 | 0,25186096 | 0,51858191 |
| 79097_at | TRIM48 | -0,24144432 | 3,42769926 | -1,19331204 | 0,25187418 | 0,51858191 |
| 84083_at | ZRANB3 | -0,19978833 | 5,99132763 | -1,19281073 | 0,25206447 | 0,51874696 |
| 387644_at | LINC00202-1 | -0,137281 | 5,74641959 | -1,19277379 | 0,2520785 | 0,51874696 |
| 100506679_a | LOC10050667 | -0,16995629 | 4,3107879 | -1,19271794 | 0,2520997 | 0,51874696 |
| 5531_at | PPP4C | 0,10910514 | 9,70463595 | 1,19270531 | 0,2521045 | 0,51874696 |
| 405753_at | DUOXA2 | 0,19209634 | 5,65730708 | 1,19268109 | 0,2521137 | 0,51874696 |
| 26247_at | OR2L1P | -0,09618571 | 4,47626948 | -1,19262349 | 0,25213557 | 0,51874696 |
| 100289495_a | LOC10028949 | -0,07459576 | 5,24546013 | -1,19244834 | 0,2522021 | 0,51874696 |
| 7381_at | UQCRB | -0,07480432 | 7,22843058 | -1,19240193 | 0,25221972 | 0,51874696 |
| 57863_at | CADM3 | -0,12833722 | 5,26257976 | -1,19239873 | 0,25222094 | 0,51874696 |
| 115950_at | ZNF653 | 0,14449531 | 6,18342276 | 1,19239162 | 0,25222364 | 0,51874696 |
| 80161_at | ASMTL-AS1 | -0,13245234 | 5,92312338 | -1,19224963 | 0,25227759 | 0,51878453 |
| 1585_at | CYP11B2 | -0,17526934 | 5,48148462 | -1,19220176 | 0,25229578 | 0,51878453 |
| 8622_at | PDE8B | 0,81625145 | 4,60776328 | 1,19186691 | 0,25242303 | 0,51895478 |
| 83699_at | SH3BGRL2 | 0,49038171 | 7,67383504 | 1,19184212 | 0,25243245 | 0,51895478 |
| 29091_at | STXBP6 | 0,35110293 | 3,99354402 | 1,19164906 | 0,25250585 | 0,51905027 |
| 8643_at | PTCH2 | -0,10758492 | 5,31254428 | -1,19117034 | 0,25268791 | 0,5193691 |
| 5603_at | MAPK13 | -0,33140766 | 8,11963671 | -1,19062715 | 0,25289462 | 0,5197385 |
| 27091_at | CACNG5 | -0,13355052 | 4,26715307 | -1,19041581 | 0,25297507 | 0,51980291 |
| 8650_at | NUMB | 0,13931908 | 8,13955853 | 1,19040307 | 0,25297992 | 0,51980291 |
| 643201_at | LOC643201 | -0,13045374 | 3,59687653 | -1,19022215 | 0,25304882 | 0,51988901 |
| 375057_at | C1orf95 | -0,11643889 | 4,86486992 | -1,18995278 | 0,25315142 | 0,52004434 |
| 57605_at | PITPNM2 | -0,14278587 | 5,36875282 | -1,18979341 | 0,25321214 | 0,5201136 |
| 84191_at | FAM96A | -0,09968331 | 9,10229217 | -1,18964703 | 0,25326791 | 0,52013591 |
| 164_at | AP1G1 | 0,09502113 | 9,22247921 | 1,1896232 | 0,253277 | 0,52013591 |
| 3111_at | HLA-DOA | -0,10836859 | 5,43520872 | -1,18933517 | 0,25338678 | 0,5203059 |
| 90580_at | C19orf52 | -0,12789803 | 7,55793683 | -1,18920059 | 0,25343809 | 0,52035579 |
| 9028_at | RHBDL1 | 0,16384456 | 5,59051015 | 1,18911171 | 0,25347198 | 0,52036991 |
| 55859_at | BEX1 | -0,11603485 | 4,03943912 | -1,18903588 | 0,2535009 | 0,52037022 |
| 3206_at | HOXA10 | -0,09464807 | 5,5449238 | -1,18896964 | 0,25352616 | 0,52037022 |
| 64397_at | ZNF106 | 0,14826065 | 7,92052488 | 1,18877366 | 0,25360092 | 0,52038834 |
| 389715_at | FAM205BP | -0,10080119 | 3,74230029 | -1,18870506 | 0,25362709 | 0,52038834 |
| 3952_at | LEP | -0,16741066 | 4,80941291 | -1,18866809 | 0,25364119 | 0,52038834 |
| 7021_at | TFAP2B | -0,08430444 | 3,69088283 | -1,18866321 | 0,25364305 | 0,52038834 |
| 29896_at | TRA2A | -0,14613148 | 9,49677445 | -1,18839289 | 0,2537462 | 0,52047667 |
| 10734_at | STAG3 | -0,14021621 | 5,44887914 | -1,18834762 | 0,25376348 | 0,52047667 |
| 64241_at | ABCG8 | -0,10348857 | 5,19273549 | -1,18833797 | 0,25376716 | 0,52047667 |
| 375612_at | LHFPL3 | -0,11517036 | 3,86516459 | -1,18802448 | 0,25388684 | 0,52062579 |
| 140683_at | BPIFA2 | 0,77123347 | 5,27505511 | 1,1879941 | 0,25389843 | 0,52062579 |

| | | | | | | |
|---|---|---|---|---|---|---|
| 293_at | SLC25A6 | 0,17176159 | 11,9934348 | 1,18793513 | 0,25392095 | 0,52062579 |
| 2170_at | FABP3 | -0,12058488 | 3,88515216 | -1,18783812 | 0,253958 | 0,52064633 |
| 100507391_a | LOC10050739 | -0,13099553 | 4,78632935 | -1,18739724 | 0,25412641 | 0,52091094 |
| 374470_at | C12orf42 | -0,07181114 | 3,7438271 | -1,18735864 | 0,25414115 | 0,52091094 |
| 100861541_a | DOCK9-AS2 | -0,20289625 | 5,86760883 | -1,18718492 | 0,25420754 | 0,52099158 |
| 4007_at | PRICKLE3 | -0,125766 | 6,61289459 | -1,1869757 | 0,25428751 | 0,52106868 |
| 8774_at | NAPG | -0,22616595 | 7,30742467 | -1,18694496 | 0,25429926 | 0,52106868 |
| 10016_at | PDCD6 | 0,08893308 | 9,62528594 | 1,18667391 | 0,25440289 | 0,52122558 |
| 727897_at | MUC5B | -0,1322366 | 5,20028446 | -1,18647997 | 0,25447707 | 0,5212724 |
| 3741_at | KCNA5 | -0,15171095 | 4,89318009 | -1,18635179 | 0,2545261 | 0,5212724 |
| 646600_at | IGF2BP2-AS1 | -0,12395926 | 3,98283171 | -1,1863512 | 0,25452632 | 0,5212724 |
| 80831_at | APOL5 | -0,14942244 | 3,95974733 | -1,18620125 | 0,25458369 | 0,5212724 |
| 83938_at | C10orf11 | 0,31024598 | 4,80428669 | 1,18615089 | 0,25460296 | 0,5212724 |
| 54103_at | GSAP | 0,38313864 | 5,66998576 | 1,186141 | 0,25460674 | 0,5212724 |
| 54503_at | ZDHHC13 | -0,30440033 | 8,29824224 | -1,18611898 | 0,25461517 | 0,5212724 |
| 93556_at | EGFEM1P | 0,1252526 | 3,44864766 | 1,18565659 | 0,25479215 | 0,52148396 |
| 101927811_a | LOC10192781 | 0,206664 | 4,90225453 | 1,18565383 | 0,25479321 | 0,52148396 |
| 57596_at | BEGAIN | -0,1792974 | 4,97748302 | -1,18563681 | 0,25479972 | 0,52148396 |
| 1400_at | CRMP1 | -0,13908312 | 5,9605462 | -1,18512408 | 0,25499608 | 0,5218304 |
| 100507006_a | LOC10050700 | -0,11292999 | 5,64285173 | -1,18481223 | 0,25511557 | 0,52201947 |
| 85409_at | NKD2 | -0,1476282 | 5,54581175 | -1,18461592 | 0,25519081 | 0,52206792 |
| 51440_at | HPCAL4 | -0,11090433 | 4,77799617 | -1,18460902 | 0,25519346 | 0,52206792 |
| 160622_at | GRASP | -0,11449132 | 6,09212112 | -1,18445737 | 0,25525159 | 0,52213141 |
| 51006_at | SLC35C2 | 0,20856941 | 8,21195825 | 1,18435193 | 0,25529202 | 0,52215865 |
| 114827_at | FHAD1 | -0,08434823 | 4,16321378 | -1,18423232 | 0,25533789 | 0,52219702 |
| 80270_at | HSD3B7 | 0,10169781 | 6,66094826 | 1,18410228 | 0,25538776 | 0,52220569 |
| 80723_at | SLC35G2 | -0,42854976 | 5,63552628 | -1,18407989 | 0,25539635 | 0,52220569 |
| 541468_at | LURAP1 | -0,15474316 | 5,66226091 | -1,18388493 | 0,25547113 | 0,52223509 |
| 663_at | BNIP2 | 0,10370674 | 8,60426555 | 1,18375482 | 0,25552105 | 0,52223509 |
| 2841_at | GPR18 | -0,12291121 | 3,3510047 | -1,1837294 | 0,2555308 | 0,52223509 |
| 389043_at | LINC01120 | -0,11083489 | 3,58030952 | -1,18370122 | 0,25554162 | 0,52223509 |
| 154467_at | CCDC167 | -0,20880621 | 7,97825305 | -1,18362594 | 0,2555705 | 0,52223509 |
| 83867_at | TTTY12 | -0,13679596 | 3,68627012 | -1,18361842 | 0,25557339 | 0,52223509 |
| 284071_at | C17orf104 | 0,07634584 | 2,91996297 | 1,18329092 | 0,25569909 | 0,52243653 |
| 140862_at | ISM1 | -0,09304339 | 3,90931553 | -1,18310878 | 0,25576903 | 0,52252399 |
| 220929_at | ZNF438 | 0,11692033 | 4,60038646 | 1,18297171 | 0,25582166 | 0,5225761 |
| 2248_at | FGF3 | -0,13248367 | 5,08509699 | -1,18288613 | 0,25585453 | 0,52258782 |
| 9607_at | CARTPT | -0,12767971 | 5,04570958 | -1,18269045 | 0,25592969 | 0,52268593 |
| 3628_at | INPP1 | 0,28758092 | 8,95413103 | 1,18257756 | 0,25597307 | 0,52271909 |
| 2069_at | EREG | -0,14292349 | 4,49551071 | -1,18243765 | 0,25602683 | 0,52277345 |
| 9737_at | GPRASP1 | 0,10563191 | 4,46008204 | 1,18221257 | 0,25611333 | 0,52289465 |
| 100505938_a | LOC10050593 | 0,13831624 | 4,35178524 | 1,18201103 | 0,25619081 | 0,5229974 |
| 401_at | PHOX2A | -0,12466688 | 5,44054266 | -1,18192338 | 0,25622451 | 0,52301078 |
| 135948_at | OR2F2 | -0,0884506 | 3,59849856 | -1,18166743 | 0,25632294 | 0,52310483 |
| 105374089_a | LOC10537408 | -0,16329966 | 5,94420525 | -1,18162527 | 0,25633915 | 0,52310483 |
| 6585_at | SLIT1 | -0,10565853 | 4,68111687 | -1,18159173 | 0,25635205 | 0,52310483 |
| 8807_at | IL18RAP | -0,11481758 | 4,37272655 | -1,18146732 | 0,25639991 | 0,52313146 |
| 26060_at | APPL1 | 0,10616847 | 7,78651836 | 1,18141661 | 0,25641942 | 0,52313146 |
| 166614_at | DCLK2 | -0,11773236 | 4,46349219 | -1,18134405 | 0,25644734 | 0,52313301 |
| 7257_at | TSNAX | -0,19820045 | 9,31248233 | -1,181267 | 0,25647698 | 0,52313809 |

| | | | | | | |
|---|---|---|---|---|---|---|
| 26097_at | CHTOP | -0,10910048 | 9,17336455 | -1,18116547 | 0,25651606 | 0,52316239 |
| 283160_at | OR8D2 | -0,124589 | 3,7616541 | -1,18099494 | 0,25658169 | 0,52324085 |
| 5636_at | PRPSAP2 | -0,21678512 | 8,49826785 | -1,18082151 | 0,25664845 | 0,52332159 |
| 170425_at | LINC00858 | 0,08139877 | 3,70203176 | 1,18057199 | 0,25674453 | 0,52342961 |
| 91151_at | TIGD7 | 0,20972047 | 5,45115381 | 1,18052891 | 0,25676112 | 0,52342961 |
| 10281_at | DSCR4 | -0,11170294 | 4,87503239 | -1,18047225 | 0,25678294 | 0,52342961 |
| 10858_at | CYP46A1 | -0,15559323 | 4,85410704 | -1,18027138 | 0,25686032 | 0,52350805 |
| 55206_at | SBNO1 | -0,07648943 | 7,94094693 | -1,18022475 | 0,25687828 | 0,52350805 |
| 8483_at | CILP | -0,12656687 | 4,85624443 | -1,1801129 | 0,25692138 | 0,52350805 |
| 163081_at | ZNF567 | 0,25425261 | 5,56376714 | 1,17994076 | 0,25698771 | 0,52350805 |
| 6882_at | TAF11 | -0,10382084 | 8,24845402 | -1,1797827 | 0,25704863 | 0,52350805 |
| 255762_at | PDZD9 | -0,12572121 | 3,19295672 | -1,17977887 | 0,2570501 | 0,52350805 |
| 57556_at | SEMA6A | 0,39745335 | 7,6287201 | 1,17973112 | 0,25706851 | 0,52350805 |
| 145858_at | C15orf32 | -0,12378766 | 3,31830534 | -1,17968849 | 0,25708494 | 0,52350805 |
| 149499_at | LRRC71 | -0,14205278 | 4,43386385 | -1,17962484 | 0,25710948 | 0,52350805 |
| 8572_at | PDLIM4 | -0,09297945 | 4,40734815 | -1,17960233 | 0,25711816 | 0,52350805 |
| 101927604_a | LOC10192760 | 0,12530385 | 4,03082165 | 1,1795966 | 0,25712037 | 0,52350805 |
| 23677_at | SH3BP4 | -0,17471855 | 7,90964552 | -1,1794592 | 0,25717335 | 0,52356058 |
| 4759_at | NEU2 | -0,11178857 | 4,32893482 | -1,17938391 | 0,25720238 | 0,52356435 |
| 100132386_a | KRTAP4-9 | -0,1292762 | 4,27576774 | -1,17922399 | 0,25726406 | 0,52363456 |
| 1730_at | DIAPH2 | 0,2046231 | 6,95411574 | 1,17871952 | 0,25745869 | 0,52397535 |
| 22843_at | PPM1E | 0,08684941 | 3,6944386 | 1,17847448 | 0,25755327 | 0,52411246 |
| 100419583_a | LOC10041958 | 0,3093722 | 8,56120861 | 1,17790599 | 0,2577728 | 0,52447164 |
| 101929586_a | LOC10192958 | -0,12677286 | 4,38985631 | -1,17785046 | 0,25779425 | 0,52447164 |
| 1293_at | COL6A3 | -0,11161352 | 4,16387084 | -1,17777184 | 0,25782462 | 0,52447164 |
| 101927228_a | LOC10192722 | -0,14523065 | 3,77467077 | -1,17771922 | 0,25784495 | 0,52447164 |
| 30_at | ACAA1 | 0,26167792 | 9,12961679 | 1,17766499 | 0,2578659 | 0,52447164 |
| 286499_at | FAM133A | -0,13519435 | 3,3244078 | -1,17732784 | 0,2579962 | 0,52468126 |
| 8473_at | OGT | 0,23504309 | 8,69034713 | 1,17710965 | 0,25808055 | 0,52479739 |
| 51179_at | HAO2 | -0,09791335 | 4,04310241 | -1,17689022 | 0,25816541 | 0,52491453 |
| 100874239_a | ARHGAP26-AS | -0,14017868 | 3,97693975 | -1,17678808 | 0,25820491 | 0,52493944 |
| 54926_at | UBE2R2 | 0,09928213 | 9,65721843 | 1,17661653 | 0,25827127 | 0,52501894 |
| 10268_at | RAMP3 | -0,1461314 | 5,38871494 | -1,17618473 | 0,25843836 | 0,52530316 |
| 80303_at | EFHD1 | -0,34216245 | 5,6681364 | -1,17597612 | 0,25851911 | 0,52541186 |
| 570_at | BAAT | -0,10902955 | 4,30459071 | -1,17575845 | 0,25860339 | 0,52552771 |
| 90874_at | ZNF697 | -0,09190605 | 6,99813008 | -1,17556997 | 0,25867639 | 0,52557328 |
| 195814_at | SDR16C5 | -0,56475264 | 4,3930047 | -1,17555964 | 0,25868039 | 0,52557328 |
| 79634_at | SCRN3 | 0,15399915 | 6,13354821 | 1,17536575 | 0,25875549 | 0,52567044 |
| 27299_at | ADAMDEC1 | -0,08643481 | 3,14091514 | -1,17505971 | 0,25887408 | 0,52585589 |
| 5031_at | P2RY6 | -0,17335655 | 5,86082775 | -1,17480024 | 0,25897465 | 0,52600472 |
| 56948_at | SDR39U1 | 0,05881515 | 8,12073634 | 1,1744315 | 0,25911763 | 0,52623964 |
| 79680_at | C22orf29 | 0,17773162 | 7,42821061 | 1,17434905 | 0,25914961 | 0,5262491 |
| 283870_at | BRICD5 | -0,31299912 | 6,5731312 | -1,17411879 | 0,25923893 | 0,52629846 |
| 9312_at | KCNB2 | -0,09398468 | 3,961908 | -1,17409191 | 0,25924936 | 0,52629846 |
| 114900_at | C1QTNF4 | -0,09971843 | 4,60517712 | -1,17407511 | 0,25925588 | 0,52629846 |
| 4900_at | NRGN | -0,4548423 | 7,17335721 | -1,17364223 | 0,25942388 | 0,526584 |
| 283459_at | GATC | 0,11960001 | 8,59276327 | 1,17352233 | 0,25947042 | 0,52659138 |
| 2861_at | GPR37 | 0,20840781 | 3,8715655 | 1,17349203 | 0,25948219 | 0,52659138 |
| 27107_at | ZBTB11 | -0,13922059 | 8,85304147 | -1,17323752 | 0,25958101 | 0,52666012 |
| 115286_at | SLC25A26 | -0,08745016 | 7,23025046 | -1,1732348 | 0,25958207 | 0,52666012 |

| | | | | | | |
|---|---|---|---|---|---|---|
| 2021_at | ENDOG | -0,07779112 | 7,6668784 | -1,1731888 | 0,25959994 | 0,52666012 |
| 7169_at | TPM2 | 0,08779931 | 4,62962308 | 1,17312318 | 0,25962542 | 0,52666012 |
| 84700_at | MYO18B | -0,09961173 | 4,08930258 | -1,17302929 | 0,25966189 | 0,52667863 |
| 200150_at | PLD5 | 0,10577984 | 3,50221579 | 1,17275176 | 0,25976972 | 0,52684186 |
| 79768_at | KATNBL1 | 0,15321163 | 6,2252563 | 1,17207727 | 0,26003191 | 0,52731811 |
| 5143_at | PDE4C | -0,10616178 | 5,34270643 | -1,17200409 | 0,26006037 | 0,52732031 |
| 64801_at | ARV1 | -0,26478314 | 8,52653139 | -1,17188615 | 0,26010624 | 0,52734723 |
| 26061_at | HACL1 | -0,18721399 | 7,92004028 | -1,17182918 | 0,2601284 | 0,52734723 |
| 55611_at | OTUB1 | -0,06614706 | 9,23737838 | -1,17167328 | 0,26018905 | 0,52739345 |
| 8404_at | SPARCL1 | 0,25395571 | 4,50267345 | 1,1715702 | 0,26022916 | 0,52739345 |
| 1977_at | EIF4E | -0,29016443 | 5,4327227 | -1,17155946 | 0,26023334 | 0,52739345 |
| 2867_at | FFAR2 | -0,14832685 | 4,35276618 | -1,17146409 | 0,26027045 | 0,52741317 |
| 284358_at | MAMSTR | -0,13055182 | 6,0279785 | -1,17127398 | 0,26034444 | 0,52750762 |
| 6907_at | TBL1X | -0,08533846 | 7,24555568 | -1,17105513 | 0,26042963 | 0,52762474 |
| 146434_at | ZNF597 | 0,20185989 | 6,38766037 | 1,17095101 | 0,26047017 | 0,52765137 |
| 146723_at | C17orf77 | -0,06793386 | 3,16627101 | -1,17071375 | 0,26056257 | 0,52778304 |
| 11015_at | KDELR3 | 0,16507857 | 7,80133049 | 1,16994887 | 0,26086061 | 0,52830776 |
| 8861_at | LDB1 | 0,20496877 | 8,49204386 | 1,16990819 | 0,26087647 | 0,52830776 |
| 131578_at | LRRC15 | -0,12274383 | 4,65487571 | -1,16946409 | 0,26104964 | 0,52850761 |
| 8840_at | WISP1 | 0,06577721 | 3,63647329 | 1,16939363 | 0,26107713 | 0,52850761 |
| 100506765_a | IGSF11-AS1 | -0,09766736 | 2,94716075 | -1,16931471 | 0,26110791 | 0,52850761 |
| 57680_at | CHD8 | 0,21701234 | 7,76032397 | 1,16929007 | 0,26111753 | 0,52850761 |
| 7082_at | TJP1 | 0,19737552 | 8,69811564 | 1,16923792 | 0,26113787 | 0,52850761 |
| 1909_at | EDNRA | -0,07786536 | 3,39829092 | -1,16911959 | 0,26118404 | 0,52850761 |
| 140836_at | BANF2 | -0,11032628 | 4,25241493 | -1,16909683 | 0,26119292 | 0,52850761 |
| 1877_at | E4F1 | -0,13460562 | 7,4109864 | -1,16907746 | 0,26120048 | 0,52850761 |
| 93487_at | MAPK1IP1L | -0,10781625 | 9,94408602 | -1,16897324 | 0,26124115 | 0,52850761 |
| 400550_at | FENDRR | -0,08815991 | 4,20425044 | -1,16889961 | 0,26126988 | 0,52850761 |
| 9962_at | SLC23A2 | 0,42733661 | 6,74853524 | 1,16888149 | 0,26127696 | 0,52850761 |
| 266977_at | ADGRF1 | 0,28015973 | 8,64307743 | 1,1687558 | 0,26132602 | 0,52855135 |
| 64975_at | MRPL41 | -0,25097648 | 9,85701069 | -1,16856706 | 0,2613997 | 0,52858464 |
| 1445_at | CSK | -0,11252415 | 8,87159383 | -1,16856477 | 0,26140059 | 0,52858464 |
| 645369_at | TMEM200C | -0,11181745 | 4,67380743 | -1,16840697 | 0,26146221 | 0,52858464 |
| 253769_at | WDR27 | -0,15790386 | 5,6911609 | -1,16835691 | 0,26148176 | 0,52858464 |
| 50945_at | TBX22 | -0,09858761 | 3,02903492 | -1,16828997 | 0,26150791 | 0,52858464 |
| 1447_at | CSN2 | -0,07735488 | 3,66938189 | -1,16824866 | 0,26152404 | 0,52858464 |
| 8462_at | KLF11 | -0,10607142 | 5,84261951 | -1,16822173 | 0,26153456 | 0,52858464 |
| 85440_at | DOCK7 | -0,12488897 | 6,372748 | -1,16781037 | 0,26169527 | 0,52883037 |
| 3131_at | HLF | -0,09081315 | 3,57891103 | -1,16769874 | 0,2617389 | 0,52883037 |
| 1180_at | CLCN1 | -0,12609434 | 5,18653031 | -1,16763765 | 0,26176277 | 0,52883037 |
| 2326_at | FMO1 | -0,13097785 | 3,65645292 | -1,16762951 | 0,26176595 | 0,52883037 |
| 158055_at | C9orf163 | -0,12630985 | 5,70514074 | -1,16754474 | 0,26179909 | 0,52884186 |
| 101928419_a | LOC10192841 | -0,18625823 | 3,85134379 | -1,16710008 | 0,26197295 | 0,52913757 |
| 84668_at | FAM126A | -0,16526684 | 6,63650326 | -1,16685747 | 0,26206785 | 0,52922028 |
| 3046_at | HBE1 | -0,14619839 | 4,201686 | -1,16681005 | 0,2620864 | 0,52922028 |
| 143570_at | XRRA1 | -0,07853752 | 5,83202215 | -1,16678468 | 0,26209632 | 0,52922028 |
| 8738_at | CRADD | 0,08632438 | 6,50286393 | 1,1665094 | 0,26220404 | 0,5293823 |
| 84824_at | FCRLA | -0,06825453 | 3,4417399 | -1,16636496 | 0,26226058 | 0,52939178 |
| 100506084_a | ARL17B | 0,16348419 | 2,97897247 | 1,16634838 | 0,26226707 | 0,52939178 |
| 3897_at | L1CAM | 0,15883718 | 5,87740716 | 1,16628676 | 0,26229119 | 0,52939178 |

| | | | | | | |
|---|---|---|---|---|---|---|
| 10939_at | AFG3L2 | -0,32557041 | 8,77194746 | -1,16621045 | 0,26232106 | 0,5293966 |
| 9194_at | SLC16A7 | 0,39719442 | 8,09764185 | 1,16600625 | 0,26240101 | 0,52950249 |
| 4338_at | MOCS2 | -0,12519375 | 6,82843477 | -1,16558624 | 0,26256552 | 0,52977895 |
| 79990_at | PLEKHH3 | 0,08464021 | 6,91669322 | 1,16490355 | 0,26283308 | 0,53024299 |
| 64407_at | RGS18 | 0,07039102 | 2,71214995 | 1,16485899 | 0,26285056 | 0,53024299 |
| 344887_at | LOC344887 | -0,246981 | 7,05199823 | -1,16466632 | 0,26292611 | 0,53028188 |
| 6550_at | SLC9A3 | -0,14028274 | 5,02479204 | -1,16463607 | 0,26293797 | 0,53028188 |
| 63827_at | BCAN | -0,11485313 | 5,43832053 | -1,1645832 | 0,26295871 | 0,53028188 |
| 83959_at | SLC4A11 | 0,5212999 | 7,007991 | 1,16452904 | 0,26297995 | 0,53028188 |
| 8708_at | B3GALT1 | -0,08375653 | 3,51941003 | -1,16421534 | 0,26310302 | 0,53042706 |
| 57701_at | NCKAP5L | 0,09819542 | 6,22297057 | 1,16416078 | 0,26312443 | 0,53042706 |
| 8634_at | RTCA | -0,11639738 | 8,75762458 | -1,16413497 | 0,26313456 | 0,53042706 |
| 84106_at | PRAM1 | -0,11722617 | 4,73989164 | -1,16389773 | 0,26322767 | 0,53055924 |
| 339768_at | ESPNL | -0,19751792 | 6,2033958 | -1,16376565 | 0,26327952 | 0,53060822 |
| 3568_at | IL5RA | -0,07552345 | 3,42111736 | -1,16369233 | 0,2633083 | 0,53061072 |
| 255189_at | PLA2G4F | -0,11653973 | 5,99372046 | -1,16354643 | 0,26336559 | 0,53067065 |
| 91608_at | RASL10B | -0,09819066 | 5,75685427 | -1,1633805 | 0,26343075 | 0,53071452 |
| 9963_at | SLC23A1 | 0,20177479 | 4,6367136 | 1,16335069 | 0,26344246 | 0,53071452 |
| 7955_at | RNF217-AS1 | 0,09910293 | 3,8328276 | 1,16298655 | 0,26358552 | 0,53094718 |
| 3949_at | LDLR | -0,4151523 | 8,71175471 | -1,16275855 | 0,26367512 | 0,531028 |
| 63901_at | FAM111A | -0,13665952 | 9,17879341 | -1,16271678 | 0,26369154 | 0,531028 |
| 126259_at | TMIGD2 | 0,14978065 | 6,13238409 | 1,16262178 | 0,26372888 | 0,531028 |
| 79012_at | CAMKV | -0,14142657 | 4,6911861 | -1,16255035 | 0,26375696 | 0,531028 |
| 9147_at | NEMF | 0,1040708 | 8,36853599 | 1,16250806 | 0,26377358 | 0,531028 |
| 140462_at | ASB9 | 0,28723867 | 4,64052511 | 1,16246364 | 0,26379105 | 0,531028 |
| 84103_at | C4orf17 | -0,09957448 | 4,02004436 | -1,16238157 | 0,26382332 | 0,53103746 |
| 3824_at | KLRD1 | -0,09514935 | 3,58541155 | -1,16222997 | 0,26388293 | 0,53103804 |
| 170712_at | COX7B2 | -0,09772742 | 3,49878112 | -1,16222016 | 0,26388679 | 0,53103804 |
| 7462_at | LAT2 | -0,15188631 | 5,29542438 | -1,16217054 | 0,26390631 | 0,53103804 |
| 391_at | RHOG | -0,16082985 | 8,53798389 | -1,16205719 | 0,26395088 | 0,53107227 |
| 148696_at | LOC148696 | -0,15515685 | 4,55635312 | -1,16181702 | 0,26404537 | 0,53120689 |
| 6361_at | CCL17 | 0,23439822 | 5,58734111 | 1,16165968 | 0,26410728 | 0,53127596 |
| 6450_at | SH3BGR | 0,20755326 | 5,66095999 | 1,16156426 | 0,26414483 | 0,53129601 |
| 29066_at | ZC3H7A | 0,12440665 | 8,20676757 | 1,16147505 | 0,26417995 | 0,53131116 |
| 64430_at | PCNXL4 | 0,12288829 | 7,03371833 | 1,16135966 | 0,26422537 | 0,53131532 |
| 2040_at | STOM | 0,10895722 | 3,87862802 | 1,16127271 | 0,2642596 | 0,53131532 |
| 1809_at | DPYSL3 | -0,46679366 | 7,24554614 | -1,1612596 | 0,26426476 | 0,53131532 |
| 10895_at | PPBPP2 | -0,10457546 | 3,46706734 | -1,16084753 | 0,26442703 | 0,53158609 |
| 2487_at | FRZB | -0,07523684 | 3,09683404 | -1,16064924 | 0,26450514 | 0,53168763 |
| 3347_at | HTN3 | -0,07642882 | 2,88982024 | -1,16042927 | 0,26459182 | 0,53180637 |
| 23769_at | FLRT1 | -0,11226294 | 5,09302131 | -1,16006243 | 0,26473641 | 0,53204147 |
| 326625_at | MMAB | -0,11695708 | 7,47827401 | -1,15989512 | 0,26480238 | 0,53211853 |
| 121551_at | BTBD11 | 0,13657623 | 5,54679747 | 1,15973094 | 0,26486712 | 0,53215243 |
| 91653_at | BOC | -0,09811938 | 5,22391088 | -1,15971223 | 0,2648745 | 0,53215243 |
| 285463_at | CTBP1-AS | -0,09833988 | 4,81778468 | -1,15956902 | 0,26493099 | 0,53221041 |
| 84913_at | ATOH8 | 0,3819162 | 6,42364135 | 1,15942122 | 0,2649893 | 0,53227203 |
| 3596_at | IL13 | -0,10335797 | 5,09468315 | -1,15930706 | 0,26503434 | 0,532307 |
| 112937_at | GLB1L3 | -0,06863954 | 3,14986648 | -1,15921727 | 0,26506977 | 0,53232266 |
| 1812_at | DRD1 | 0,10433891 | 3,6514746 | 1,15867455 | 0,265284 | 0,53267347 |
| 26225_at | ARL5A | 0,1414489 | 9,52613155 | 1,15863464 | 0,26529976 | 0,53267347 |

| | | | | | | |
|---|---|---|---|---|---|---|
| 104_at | ADARB1 | 0,16521323 | 5,53902177 | 1,1583979 | 0,26539326 | 0,53272108 |
| 491_at | ATP2B2 | -0,08899954 | 4,38164711 | -1,15824272 | 0,26545456 | 0,53272108 |
| 113091_at | PTH2 | -0,15562896 | 6,55289743 | -1,15822821 | 0,26546029 | 0,53272108 |
| 51548_at | SIRT6 | -0,07124065 | 7,3200501 | -1,15822597 | 0,26546117 | 0,53272108 |
| 4850_at | CNOT4 | 0,12232167 | 6,53497489 | 1,15814606 | 0,26549275 | 0,53272108 |
| 100507431_a | LOC10050743 | -0,39026145 | 4,99586972 | -1,15811182 | 0,26550627 | 0,53272108 |
| 9905_at | SGSM2 | 0,20695714 | 7,86188601 | 1,15808452 | 0,26551706 | 0,53272108 |
| 2766_at | GMPR | 0,51293334 | 5,85776052 | 1,15792814 | 0,26557886 | 0,53272623 |
| 3141_at | HLCS | -0,19328183 | 6,57178852 | -1,15787167 | 0,26560117 | 0,53272623 |
| 171483_at | FAM9B | -0,0572556 | 3,01330808 | -1,15786253 | 0,26560478 | 0,53272623 |
| 9877_at | ZC3H11A | 0,09848502 | 9,38400834 | 1,15779811 | 0,26563025 | 0,53272623 |
| 440356_at | CDIPT-AS1 | -0,16319545 | 5,29264816 | -1,15761477 | 0,26570272 | 0,53278176 |
| 2859_at | GPR35 | -0,12386062 | 5,08637991 | -1,15758813 | 0,26571325 | 0,53278176 |
| 124460_at | SNX20 | -0,13168579 | 3,8192362 | -1,15730206 | 0,26582637 | 0,53295309 |
| 10221_at | TRIB1 | -0,11908257 | 6,86348467 | -1,15713868 | 0,26589099 | 0,53302717 |
| 101928107_a | LOC10192810 | -0,0937176 | 4,22170804 | -1,15695866 | 0,2659622 | 0,53311445 |
| 3507_at | IGHM | -0,12466956 | 5,81070363 | -1,15685597 | 0,26600283 | 0,53314042 |
| 101927081_a | LOC10192708 | -0,11778915 | 3,16342475 | -1,15651742 | 0,26613681 | 0,53335345 |
| 9446_at | GSTO1 | -0,28615992 | 9,05344593 | -1,15610542 | 0,26629993 | 0,53362484 |
| 1639_at | DCTN1 | 0,1774753 | 7,6077061 | 1,15602137 | 0,26633322 | 0,5336262 |
| 167826_at | OLIG3 | -0,08505192 | 3,68364489 | -1,1559638 | 0,26635602 | 0,5336262 |
| 6019_at | RLN2 | 0,25535913 | 4,0645766 | 1,1557469 | 0,26644194 | 0,53374283 |
| 399949_at | C11orf88 | -0,15606959 | 3,45175423 | -1,15547958 | 0,26654786 | 0,53389949 |
| 101927131_a | LOC10192713 | -0,1493494 | 4,06449216 | -1,15533549 | 0,26660497 | 0,53392906 |
| 22930_at | RAB3GAP1 | 0,17816709 | 8,38753213 | 1,15530246 | 0,26661806 | 0,53392906 |
| 8587_at | OR7E2P | 0,08881129 | 3,85713744 | 1,15521485 | 0,26665278 | 0,5339431 |
| 284898_at | LOC284898 | -0,19224572 | 4,39816334 | -1,15510192 | 0,26669756 | 0,53397724 |
| 540_at | ATP7B | 0,23589695 | 5,84804024 | 1,15494511 | 0,26675973 | 0,53404621 |
| 2779_at | GNAT1 | -0,11886241 | 4,94627444 | -1,15474747 | 0,26683811 | 0,53409611 |
| 55738_at | ARFGAP1 | 0,07524151 | 8,12693196 | 1,15469885 | 0,26685739 | 0,53409611 |
| 6886_at | TAL1 | -0,12572066 | 5,47324512 | -1,15467253 | 0,26686783 | 0,53409611 |
| 79414_at | LRFN3 | 0,24501225 | 5,99520512 | 1,15423993 | 0,26703947 | 0,53434142 |
| 126129_at | CPT1C | 0,38098259 | 5,79199782 | 1,15422377 | 0,26704588 | 0,53434142 |
| 54542_at | RC3H2 | 0,08743343 | 7,74820015 | 1,15394183 | 0,26715779 | 0,53450981 |
| 101927040_a | LOC10192704 | -0,10125319 | 3,45074412 | -1,15384164 | 0,26719757 | 0,53453388 |
| 680_at | BRS3 | -0,10129592 | 3,19543593 | -1,15365739 | 0,26727072 | 0,53457882 |
| 8481_at | OFD1 | 0,18856012 | 7,46045176 | 1,15364528 | 0,26727553 | 0,53457882 |
| 92369_at | SPSB4 | -0,10283275 | 4,93120739 | -1,15349173 | 0,26733652 | 0,53464528 |
| 7761_at | ZNF214 | 0,05947099 | 3,90030856 | 1,15338916 | 0,26737726 | 0,53467125 |
| 389458_at | RBAKDN | -0,17336611 | 5,52921377 | -1,15327793 | 0,26742144 | 0,53470409 |
| 100129460_a | DPY19L1P1 | 0,09660728 | 3,77759985 | 1,15320477 | 0,26745051 | 0,53470671 |
| 84366_at | PRAC1 | -0,07808547 | 3,51172157 | -1,15298991 | 0,26753589 | 0,53471258 |
| 8548_at | BLZF1 | 0,18952648 | 6,70564683 | 1,15296639 | 0,26754523 | 0,53471258 |
| 4023_at | LPL | -0,09741425 | 3,95482609 | -1,15292785 | 0,26756055 | 0,53471258 |
| 10120_at | ACTR1B | 0,08211987 | 8,78153116 | 1,15291796 | 0,26756448 | 0,53471258 |
| 257358_at | LINC01366 | -0,13851319 | 4,08632069 | -1,15278522 | 0,26761724 | 0,53476254 |
| 7430_at | EZR | -0,07818612 | 10,4796873 | -1,15269944 | 0,26765134 | 0,53476558 |
| 6594_at | SMARCA1 | 0,43838093 | 8,17282778 | 1,15264173 | 0,26767428 | 0,53476558 |
| 143884_at | CWF19L2 | -0,08056412 | 6,24651417 | -1,15252747 | 0,26771971 | 0,53477083 |
| 84164_at | ASCC2 | -0,12401238 | 8,60220366 | -1,15249548 | 0,26773243 | 0,53477083 |

| | | | | | | |
|---|---|---|---|---|---|---|
| 219833_at | C11orf45 | 0,17002047 | 4,49694039 | 1,15238744 | 0,26777539 | 0,53480118 |
| 5138_at | PDE2A | 0,15625337 | 5,27871011 | 1,15201429 | 0,26792382 | 0,53494852 |
| 51071_at | DERA | 0,11415201 | 8,85977857 | 1,15201188 | 0,26792478 | 0,53494852 |
| 127018_at | LYPLAL1 | 0,10212047 | 6,43972767 | 1,15199253 | 0,26793248 | 0,53494852 |
| 25842_at | ASF1A | -0,31096664 | 9,24740086 | -1,1518466 | 0,26799054 | 0,535009 |
| 1768_at | DNAH6 | -0,07621373 | 3,51834449 | -1,15161742 | 0,26808175 | 0,53513562 |
| 284361_at | EMC10 | 0,16932975 | 9,07544085 | 1,15150189 | 0,26812774 | 0,53516677 |
| 55106_at | SLFN12 | 0,7222665 | 4,93746732 | 1,1513811 | 0,26817582 | 0,53516677 |
| 10891_at | PPARGC1A | 0,14946543 | 4,72969327 | 1,15136885 | 0,2681807 | 0,53516677 |
| 10076_at | PTPRU | 0,0683479 | 8,4358051 | 1,15129749 | 0,26820911 | 0,53516803 |
| 100271722_a | LINC00899 | -0,08388436 | 4,24453324 | -1,15117881 | 0,26825637 | 0,53518787 |
| 57583_at | TMEM181 | -0,12420297 | 9,17770262 | -1,15111635 | 0,26828124 | 0,53518787 |
| 57337_at | SENP7 | 0,32980553 | 5,57843597 | 1,15106321 | 0,26830241 | 0,53518787 |
| 4756_at | NEO1 | 0,18948304 | 8,17603348 | 1,15096993 | 0,26833956 | 0,53519877 |
| 131566_at | DCBLD2 | -0,08270071 | 7,26994897 | -1,15090019 | 0,26836734 | 0,53519877 |
| 4698_at | NDUFA5 | 0,1413713 | 8,2571647 | 1,15084024 | 0,26839122 | 0,53519877 |
| 51702_at | PADI3 | -0,21538682 | 7,21152163 | -1,15068449 | 0,26845327 | 0,53522761 |
| 7455_at | ZAN | -0,10161855 | 5,4406359 | -1,15066446 | 0,26846126 | 0,53522761 |
| 8498_at | RANBP3 | -0,14484688 | 7,80008068 | -1,15054819 | 0,26850759 | 0,53526459 |
| 100507443_a | LOC10050744 | -0,12502673 | 4,07315181 | -1,15034023 | 0,26859047 | 0,53531206 |
| 124_at | ADH1A | -0,09576049 | 3,33722456 | -1,15027995 | 0,2686145 | 0,53531206 |
| 140902_at | R3HDML | -0,10889232 | 4,82435768 | -1,15027926 | 0,26861478 | 0,53531206 |
| 440131_at | LINC00544 | -0,10936568 | 4,08880073 | -1,15019529 | 0,26864825 | 0,53532339 |
| 100507540_a | LINC01504 | -0,13727633 | 4,44328406 | -1,14995975 | 0,26874216 | 0,53545514 |
| 284950_at | LOC284950 | 0,14965362 | 4,39965025 | 1,14980863 | 0,26880243 | 0,53551983 |
| 102723167_a | MIR133A1HG | -0,06717508 | 2,77246865 | -1,14959219 | 0,26888876 | 0,53563643 |
| 100507274_a | LINC01210 | -0,10782481 | 3,71477557 | -1,1493159 | 0,268999 | 0,53573718 |
| 84060_at | RBM48 | 0,11255796 | 5,66296347 | 1,14927157 | 0,26901669 | 0,53573718 |
| 55653_at | BCAS4 | 0,28756085 | 6,95208141 | 1,14917841 | 0,26905387 | 0,53573718 |
| 5651_at | TMPRSS15 | -0,10064094 | 3,40573694 | -1,14916158 | 0,26906059 | 0,53573718 |
| 136332_at | LRGUK | -0,12575594 | 3,903998 | -1,14911697 | 0,2690784 | 0,53573718 |
| 4978_at | OPCML | -0,09079584 | 3,6658315 | -1,14896906 | 0,26913744 | 0,53579935 |
| 729678_at | LINC00847 | 0,47676596 | 6,23139471 | 1,1486001 | 0,26928476 | 0,53585821 |
| 400759_at | GBP1P1 | 0,45097101 | 4,8750155 | 1,14857251 | 0,26929578 | 0,53585821 |
| 55220_at | KLHDC8A | -0,12804598 | 4,88859163 | -1,14855721 | 0,2693019 | 0,53585821 |
| 25847_at | ANAPC13 | 0,17030788 | 9,87197691 | 1,14855172 | 0,26930409 | 0,53585821 |
| 348825_at | TPRXL | -0,18863379 | 7,69572481 | -1,14854668 | 0,2693061 | 0,53585821 |
| 283948_at | NHLRC4 | -0,14112284 | 6,325649 | -1,14827125 | 0,26941613 | 0,53597126 |
| 101929631_a | LOC10192963 | -0,15298805 | 3,89855044 | -1,1481969 | 0,26944583 | 0,53597126 |
| 10866_at | HCP5 | 1,06127479 | 5,91710853 | 1,14816922 | 0,26945689 | 0,53597126 |
| 10897_at | YIF1A | -0,06658232 | 8,8565618 | -1,14812589 | 0,26947421 | 0,53597126 |
| 3240_at | HP | -0,10920836 | 3,73256308 | -1,14804798 | 0,26950534 | 0,53597784 |
| 26012_at | NSMF | -0,26462649 | 7,60656477 | -1,14763032 | 0,26967229 | 0,5362545 |
| 5327_at | PLAT | -0,45027511 | 9,8930153 | -1,1475117 | 0,26971972 | 0,53625657 |
| 51378_at | ANGPT4 | -0,12047562 | 5,20775203 | -1,14748846 | 0,26972901 | 0,53625657 |
| 57695_at | USP37 | -0,2380565 | 7,43627038 | -1,14733139 | 0,26979183 | 0,5363261 |
| 65108_at | MARCKSL1 | 0,29771349 | 10,6368059 | 1,14698327 | 0,26993108 | 0,53654756 |
| 2864_at | FFAR1 | -0,14925072 | 5,34361026 | -1,1467933 | 0,2700071 | 0,53662125 |
| 2854_at | GPR32 | 0,10126162 | 3,38191585 | 1,14673134 | 0,2700319 | 0,53662125 |
| 388630_at | TRABD2B | -0,10287169 | 4,4850288 | -1,14668179 | 0,27005173 | 0,53662125 |

| | | | | | | |
|---|---|---|---|---|---|---|
| 26000_at | TBC1D10B | -0,11596014 | 7,70004439 | -1,14631283 | 0,27019943 | 0,53685938 |
| 79582_at | SPAG16 | 0,08804482 | 4,05722347 | 1,14612755 | 0,27027363 | 0,53695141 |
| 171546_at | SPTSSA | 0,17660527 | 7,59612993 | 1,14565061 | 0,27046469 | 0,53718245 |
| 6668_at | SP2 | -0,09844865 | 7,79767354 | -1,14564225 | 0,27046804 | 0,53718245 |
| 85457_at | CIPC | 0,10709349 | 7,97551789 | 1,14562842 | 0,27047358 | 0,53718245 |
| 48_at | ACO1 | 0,190346 | 8,68809984 | 1,14538006 | 0,27057312 | 0,53732474 |
| 29906_at | ST8SIA5 | -0,12174631 | 4,75889674 | -1,14515978 | 0,27066142 | 0,53744469 |
| 768211_at | RELL1 | 0,08439787 | 7,09914324 | 1,14478846 | 0,27081033 | 0,53766972 |
| 4902_at | NRTN | 0,17352103 | 5,48081481 | 1,14473799 | 0,27083057 | 0,53766972 |
| 55700_at | MAP7D1 | 0,13443937 | 9,18822858 | 1,14443905 | 0,2709505 | 0,53771764 |
| 441178_at | LOC441178 | -0,1139357 | 4,10446941 | -1,14442965 | 0,27095427 | 0,53771764 |
| 129685_at | TAF8 | 0,07107524 | 6,7350109 | 1,14439826 | 0,27096687 | 0,53771764 |
| 641638_at | SNHG6 | -0,30658248 | 9,63326424 | -1,14434101 | 0,27098984 | 0,53771764 |
| 9447_at | AIM2 | 0,3903959 | 6,18286436 | 1,14432994 | 0,27099428 | 0,53771764 |
| 9760_at | TOX | -0,07348206 | 3,42681459 | -1,14405558 | 0,2711044 | 0,53788074 |
| 648987_at | LOC648987 | 0,12874343 | 5,58554762 | 1,14394092 | 0,27115043 | 0,53791666 |
| 653677_at | SEC1P | -0,10962035 | 4,66697426 | -1,1437143 | 0,27124143 | 0,53804177 |
| 26776_at | SNORA71B | -0,17780233 | 4,12267891 | -1,14351777 | 0,27132036 | 0,53809405 |
| 3745_at | KCNB1 | -0,11289563 | 4,02594105 | -1,14350957 | 0,27132365 | 0,53809405 |
| 5609_at | MAP2K7 | 0,10953575 | 6,53960341 | 1,14340663 | 0,271365 | 0,53812065 |
| 81609_at | SNX27 | 0,08692494 | 8,15487176 | 1,14322345 | 0,2714386 | 0,53821119 |
| 92305_at | TMEM129 | -0,08146319 | 8,87404666 | -1,14311606 | 0,27148175 | 0,53824135 |
| 51663_at | ZFR | -0,1364042 | 8,29475498 | -1,14293653 | 0,2715539 | 0,53832899 |
| 83863_at | TTTY5 | -0,1167945 | 4,34425122 | -1,14260202 | 0,27168838 | 0,53851324 |
| 10922_at | FASTK | 0,08246377 | 9,06111031 | 1,14250333 | 0,27172806 | 0,53851324 |
| 101927084_a | LINC01359 | -0,1267144 | 4,11867452 | -1,14244228 | 0,27175261 | 0,53851324 |
| 79710_at | MORC4 | -0,1215294 | 7,60278885 | -1,14236687 | 0,27178294 | 0,53851324 |
| 160762_at | CCDC63 | -0,08847431 | 4,18863996 | -1,1423577 | 0,27178662 | 0,53851324 |
| 399979_at | SNX19 | 0,21663694 | 7,55000654 | 1,14212891 | 0,27187865 | 0,53864018 |
| 101928031_a | LOC10192803 | -0,06928181 | 3,19825626 | -1,14177267 | 0,272022 | 0,53886875 |
| 54757_at | FAM20A | -0,08990143 | 3,90588806 | -1,1415356 | 0,27211742 | 0,53892915 |
| 887_at | CCKBR | -0,11351344 | 4,56623304 | -1,14146228 | 0,27214693 | 0,53892915 |
| 613212_at | CTXN3 | -0,10047846 | 3,48487286 | -1,14138583 | 0,27217771 | 0,53892915 |
| 26266_at | SLC13A4 | 0,1128616 | 5,61754057 | 1,14136216 | 0,27218725 | 0,53892915 |
| 7597_at | ZBTB25 | -0,09526888 | 6,16218808 | -1,14134151 | 0,27219556 | 0,53892915 |
| 3625_at | INHBB | 0,41755209 | 6,67320892 | 1,14127994 | 0,27222035 | 0,53892915 |
| 646168_at | LINC01330 | -0,07072617 | 2,74800633 | -1,1411172 | 0,27228589 | 0,5390035 |
| 115290_at | FBXO17 | -0,10252584 | 5,20247193 | -1,14100252 | 0,27233208 | 0,53900376 |
| 84070_at | FAM186B | -0,1505167 | 4,39984898 | -1,14087733 | 0,27238251 | 0,53900376 |
| 91614_at | DEPDC7 | -0,36154294 | 3,90972091 | -1,14083512 | 0,27239951 | 0,53900376 |
| 134288_at | TMEM174 | -0,11381861 | 3,99993827 | -1,14079905 | 0,27241405 | 0,53900376 |
| 64446_at | DNAI2 | -0,08556153 | 4,21461985 | -1,14076957 | 0,27242592 | 0,53900376 |
| 23544_at | SEZ6L | -0,11004994 | 4,89619942 | -1,14054029 | 0,27251831 | 0,53909031 |
| 6441_at | SFTPD | 0,16056073 | 4,78917052 | 1,1405221 | 0,27252564 | 0,53909031 |
| 117194_at | MRGPRX2 | -0,11045467 | 4,4425124 | -1,14033818 | 0,27259978 | 0,53918159 |
| 152992_at | TRMT44 | -0,12825755 | 6,34209929 | -1,14005801 | 0,27271273 | 0,53934962 |
| 79760_at | GEMIN7 | -0,09599791 | 7,56581283 | -1,1398129 | 0,27281158 | 0,5394509 |
| 147166_at | TRIM16L | -0,1904362 | 4,78274145 | -1,13979214 | 0,27281995 | 0,5394509 |
| 100506492_a | DSCAM-AS1 | -0,09551998 | 3,62264173 | -1,13955787 | 0,27291446 | 0,53958238 |
| 50509_at | COL5A3 | -0,12816988 | 5,91347099 | -1,13905745 | 0,27311641 | 0,53992625 |

| | | | | | | |
|---|---|---|---|---|---|---|
| 100506013_a | APELA | 0,07539777 | 2,99961246 | 1,13885978 | 0,27319621 | 0,54002859 |
| 55937_at | APOM | -0,15586266 | 5,60850524 | -1,13877411 | 0,2732308 | 0,54004155 |
| 3105_at | HLA-A | 0,63493916 | 11,3924015 | 1,13819903 | 0,2734631 | 0,54035606 |
| 10766_at | TOB2 | 0,28291975 | 7,9192865 | 1,13818678 | 0,27346805 | 0,54035606 |
| 1071_at | CETP | -0,14764005 | 4,08438733 | -1,13809117 | 0,27350668 | 0,54035606 |
| 8723_at | SNX4 | 0,16836389 | 8,6547214 | 1,13803838 | 0,27352802 | 0,54035606 |
| 400258_at | C14orf180 | -0,13077489 | 5,70237913 | -1,13803301 | 0,27353019 | 0,54035606 |
| 375323_at | LHFPL4 | -0,16001112 | 5,38102489 | -1,1377877 | 0,27362934 | 0,54046095 |
| 50861_at | STMN3 | -0,43301812 | 6,5302104 | -1,13770342 | 0,27366341 | 0,54046095 |
| 57560_at | IFT80 | 0,15222602 | 8,37262873 | 1,13761658 | 0,27369852 | 0,54046095 |
| 9455_at | HOMER2 | -0,11626771 | 4,94863498 | -1,13755394 | 0,27372385 | 0,54046095 |
| 7275_at | TUB | -0,08186289 | 5,59407929 | -1,13749643 | 0,27374711 | 0,54046095 |
| 55974_at | SLC50A1 | 0,08567068 | 8,78790041 | 1,13748527 | 0,27375162 | 0,54046095 |
| 121227_at | LRIG3 | 0,19977472 | 7,01023269 | 1,13737938 | 0,27379444 | 0,54046518 |
| 9341_at | VAMP3 | 0,13308915 | 10,0837024 | 1,13734121 | 0,27380988 | 0,54046518 |
| 28400_at | IGHV4-28 | -0,16528861 | 3,97637529 | -1,13716297 | 0,27388198 | 0,5405521 |
| 112399_at | EGLN3 | -0,22987022 | 7,69192788 | -1,13667968 | 0,27407754 | 0,54081743 |
| 57710_at | KIAA1614 | -0,16331653 | 5,83139728 | -1,13653885 | 0,27413454 | 0,54081743 |
| 4588_at | MUC6 | -0,0918852 | 4,80787561 | -1,13652596 | 0,27413976 | 0,54081743 |
| 10425_at | ARIH2 | -0,16936888 | 7,35362015 | -1,13648685 | 0,27415559 | 0,54081743 |
| 10308_at | ZNF267 | 0,14988039 | 6,77578144 | 1,1364839 | 0,27415679 | 0,54081743 |
| 112_at | ADCY6 | 0,22193869 | 7,86947029 | 1,13616174 | 0,27428723 | 0,54101935 |
| 9965_at | FGF19 | -0,12857846 | 4,77395663 | -1,13578024 | 0,27444177 | 0,54126874 |
| 4139_at | MARK1 | -0,09795718 | 6,00727179 | -1,13556279 | 0,27452988 | 0,54138709 |
| 64693_at | CTAGE1 | 0,07273935 | 3,33911191 | 1,13514744 | 0,27469824 | 0,54157343 |
| 120400_at | NXPE1 | -0,08219584 | 3,47680283 | -1,13513751 | 0,27470227 | 0,54157343 |
| 7574_at | ZNF26 | -0,14148412 | 5,48665015 | -1,13510687 | 0,27471469 | 0,54157343 |
| 254050_at | LRRC43 | -0,12065486 | 6,28285403 | -1,13505227 | 0,27473683 | 0,54157343 |
| 100861555_a | LINC00565 | -0,13648366 | 3,8684052 | -1,13493117 | 0,27478594 | 0,54161481 |
| 146556_at | C16orf89 | -0,16505068 | 4,73246875 | -1,13477826 | 0,27484796 | 0,54162532 |
| 1889_at | ECE1 | -0,08467274 | 7,91084376 | -1,13476262 | 0,2748543 | 0,54162532 |
| 7093_at | TLL2 | 0,22980794 | 4,8125654 | 1,13471006 | 0,27487562 | 0,54162532 |
| 29062_at | WDR91 | 0,05426763 | 6,16373499 | 1,13461362 | 0,27491474 | 0,54164699 |
| 100129961_a | CCNT2-AS1 | -0,12933752 | 4,76928887 | -1,13435172 | 0,27502101 | 0,54180095 |
| 317719_at | KLHL10 | -0,08009264 | 3,42859326 | -1,13415609 | 0,2751004 | 0,54180877 |
| 6010_at | RHO | -0,12743099 | 4,94155764 | -1,13415557 | 0,27510061 | 0,54180877 |
| 5268_at | SERPINB5 | -0,19830795 | 10,7096765 | -1,13413402 | 0,27510936 | 0,54180877 |
| 284541_at | CYP4A22 | -0,16758627 | 5,07635692 | -1,13402929 | 0,27515187 | 0,54182835 |
| 25896_at | INTS7 | -0,21939859 | 7,24256411 | -1,13397094 | 0,27517556 | 0,54182835 |
| 8717_at | TRADD | 0,22121783 | 8,256856 | 1,13359907 | 0,27532656 | 0,54203409 |
| 4651_at | MYO10 | -0,12645534 | 7,07793922 | -1,13357504 | 0,27533632 | 0,54203409 |
| 5154_at | PDGFA | 0,21398652 | 7,46543571 | 1,13330527 | 0,27544591 | 0,5421918 |
| 80204_at | FBXO11 | 0,21442811 | 8,58115764 | 1,13323079 | 0,27547617 | 0,5421918 |
| 27295_at | PDLIM3 | -0,0674406 | 4,50611668 | -1,13304137 | 0,27555314 | 0,5421918 |
| 158046_at | NXNL2 | -0,15805215 | 3,83817217 | -1,13301281 | 0,27556475 | 0,5421918 |
| 2669_at | GEM | -0,24557363 | 3,99188275 | -1,13299803 | 0,27557075 | 0,5421918 |
| 173_at | AFM | -0,09892382 | 3,2853781 | -1,1329622 | 0,27558531 | 0,5421918 |
| 57107_at | PDSS2 | -0,15139582 | 6,38515459 | -1,13280924 | 0,27564749 | 0,54219191 |
| 57620_at | STIM2 | -0,11806903 | 7,61281165 | -1,1327967 | 0,27565258 | 0,54219191 |
| 7049_at | TGFBR3 | 0,40561101 | 9,43687855 | 1,13270165 | 0,27569123 | 0,54219191 |

| | | | | | | |
|---|---|---|---|---|---|---|
| 1404_at | HAPLN1 | -0,07337385 | 3,25871893 | -1,1326353 | 0,2757182 | 0,54219191 |
| 26525_at | IL36RN | -0,11849781 | 4,08435855 | -1,13251217 | 0,27576827 | 0,54219191 |
| 219537_at | SMTNL1 | -0,12824442 | 5,22403691 | -1,13250222 | 0,27577231 | 0,54219191 |
| 5527_at | PPP2R5C | 0,07122168 | 8,75107646 | 1,13247742 | 0,2757824 | 0,54219191 |
| 100128511_a | NEBL-AS1 | 0,15750396 | 4,704427 | 1,13226123 | 0,27587032 | 0,54227963 |
| 90362_at | FAM110B | 0,11220333 | 4,28375114 | 1,13222928 | 0,27588332 | 0,54227963 |
| 79168_at | LILRA6 | -0,12430231 | 5,09547016 | -1,13177275 | 0,27606906 | 0,54253867 |
| 4884_at | NPTX1 | -0,12465605 | 5,42153811 | -1,13176692 | 0,27607144 | 0,54253867 |
| 23157_at | SEPT6 | 0,22949229 | 6,37177568 | 1,1316434 | 0,27612171 | 0,542552 |
| 56961_at | SHD | 0,13631781 | 4,71765802 | 1,1315882 | 0,27614418 | 0,542552 |
| 49861_at | CLDN20 | -0,12617773 | 3,40828377 | -1,13151267 | 0,27617493 | 0,542552 |
| 51438_at | MAGEC2 | -0,14729309 | 4,27554524 | -1,13131559 | 0,27625516 | 0,542552 |
| 54491_at | FAM105A | 0,11929187 | 5,69947714 | 1,13123909 | 0,27628631 | 0,542552 |
| 10143_at | CLEC3A | -0,10239488 | 4,09149956 | -1,13123374 | 0,27628849 | 0,542552 |
| 283677_at | REC114 | -0,09143921 | 4,68315519 | -1,13121528 | 0,27629601 | 0,542552 |
| 117283_at | IP6K3 | 0,12878853 | 4,00727692 | 1,13119678 | 0,27630354 | 0,542552 |
| 126668_at | TDRD10 | -0,16788952 | 4,59005805 | -1,13100509 | 0,27638161 | 0,54264665 |
| 258_at | AMBN | -0,11482965 | 4,51147768 | -1,13093335 | 0,27641083 | 0,54264665 |
| 54502_at | RBM47 | 0,19584006 | 8,13244088 | 1,13087095 | 0,27643625 | 0,54264665 |
| 83987_at | CCDC8 | -0,10442746 | 4,60657106 | -1,13055857 | 0,27656353 | 0,54275647 |
| 55122_at | AKIRIN2 | -0,15553619 | 8,57006884 | -1,13049032 | 0,27659135 | 0,54275647 |
| 647979_at | LINC00657 | 0,16114163 | 10,9644558 | 1,13046879 | 0,27660012 | 0,54275647 |
| 5739_at | PTGIR | -0,13835854 | 6,34932907 | -1,13045705 | 0,27660491 | 0,54275647 |
| 9129_at | PRPF3 | -0,13493746 | 7,72299809 | -1,13004635 | 0,27677233 | 0,54301567 |
| 79744_at | ZNF419 | 0,09036381 | 6,98455694 | 1,12999472 | 0,27679338 | 0,54301567 |
| 80343_at | SEL1L2 | -0,15630667 | 3,55853096 | -1,12975544 | 0,27689097 | 0,5431518 |
| 374920_at | C19orf68 | 0,13693377 | 6,1519829 | 1,12965744 | 0,27693094 | 0,54317491 |
| 9696_at | CROCC | 0,11680462 | 5,96628685 | 1,12944384 | 0,27701809 | 0,54327676 |
| 342371_at | ATXN1L | 0,13461018 | 8,66284451 | 1,12939191 | 0,27703928 | 0,54327676 |
| 84443_at | FRMPD3 | -0,12052253 | 5,24350162 | -1,12926907 | 0,2770894 | 0,54331974 |
| 5407_at | PNLIPRP1 | -0,11218287 | 4,79537896 | -1,12910803 | 0,27715513 | 0,54339331 |
| 3669_at | ISG20 | 0,61526519 | 6,33904327 | 1,12891624 | 0,27723342 | 0,54349149 |
| 53358_at | SHC3 | -0,1334075 | 4,94646209 | -1,12868977 | 0,27732589 | 0,54361745 |
| 148362_at | BROX | -0,08467451 | 8,44178795 | -1,12861494 | 0,27735645 | 0,54362204 |
| 163126_at | EID2 | 0,11192495 | 8,44978737 | 1,12818933 | 0,27753031 | 0,54387882 |
| 100505681_a | NAPA-AS1 | -0,13032481 | 5,96344402 | -1,12811628 | 0,27756016 | 0,54387882 |
| 9514_at | GAL3ST1 | -0,11673903 | 5,10162244 | -1,12806131 | 0,27758262 | 0,54387882 |
| 65998_at | C11orf95 | 0,15581363 | 6,21740027 | 1,1280178 | 0,27760041 | 0,54387882 |
| 117156_at | SCGB3A2 | -0,11621223 | 3,93502564 | -1,12788028 | 0,27765661 | 0,54390483 |
| 101929144_a | LOC10192914 | -0,10945212 | 3,02767259 | -1,12784715 | 0,27767015 | 0,54390483 |
| 1610_at | DAO | -0,16378368 | 4,56507725 | -1,12776217 | 0,27770489 | 0,54391756 |
| 27128_at | CYTH4 | -0,11478425 | 5,5852369 | -1,12761728 | 0,27776412 | 0,54397827 |
| 1443_at | CSH2 | -0,14519525 | 5,86046249 | -1,12721526 | 0,27792852 | 0,54424491 |
| 28773_at | IGLV9-49 | -0,08851111 | 4,77435421 | -1,12697897 | 0,27802519 | 0,54437885 |
| 1288_at | COL4A6 | 0,50797039 | 6,49544602 | 1,12688177 | 0,27806496 | 0,54440139 |
| 374860_at | ANKRD30B | -0,07559868 | 2,7267305 | -1,12644372 | 0,27824424 | 0,544668 |
| 100132596_a | XGY2 | -0,09686581 | 3,13145775 | -1,12641089 | 0,27825768 | 0,544668 |
| 100130950_a | LOC10013095 | -0,11097269 | 4,99975313 | -1,12622363 | 0,27833436 | 0,54476181 |
| 54922_at | RASIP1 | -0,12980772 | 4,60219802 | -1,1261557 | 0,27836217 | 0,54476181 |
| 114880_at | OSBPL6 | -0,17325153 | 4,4825113 | -1,12599355 | 0,27842858 | 0,54483643 |

| | | | | | | |
|---|---|---|---|---|---|---|
| 55244_at | SLC47A1 | 0,19269403 | 4,95867556 | 1,12574217 | 0,27853156 | 0,54498257 |
| 23546_at | SYNGR4 | -0,13940366 | 5,22184723 | -1,1255913 | 0,27859338 | 0,54504816 |
| 9856_at | KIAA0319 | -0,14399216 | 4,93137492 | -1,12545986 | 0,27864724 | 0,54509818 |
| 11240_at | PADI2 | 0,07172449 | 6,09659595 | 1,12527157 | 0,27872441 | 0,54519378 |
| 5589_at | PRKCSH | 0,12492543 | 8,88101008 | 1,12509186 | 0,27879809 | 0,54523766 |
| 10246_at | SLC17A2 | -0,10949588 | 3,58849185 | -1,12507877 | 0,27880345 | 0,54523766 |
| 55773_at | TBC1D23 | 0,27425137 | 8,17519451 | 1,12496626 | 0,27884958 | 0,54524208 |
| 100996263_a | LOC10099626 | -0,10612008 | 3,64481349 | -1,12493519 | 0,27886233 | 0,54524208 |
| 101927591_a | DPP10-AS3 | -0,07668083 | 3,33700901 | -1,12478217 | 0,27892508 | 0,54530943 |
| 55857_at | KIZ | 0,27754947 | 7,5544212 | 1,1247036 | 0,2789573 | 0,54531708 |
| 118663_at | BTBD16 | -0,41535519 | 8,28426144 | -1,12428973 | 0,2791271 | 0,54559364 |
| 10455_at | ECI2 | 0,61740095 | 7,05687784 | 1,12419319 | 0,27916672 | 0,54559868 |
| 254528_at | MEIOB | 0,19016219 | 3,73157647 | 1,12414542 | 0,27918633 | 0,54559868 |
| 121391_at | KRT74 | -0,10725474 | 4,89552989 | -1,12365449 | 0,27938787 | 0,54587726 |
| 7139_at | TNNT2 | -0,10399385 | 5,236541 | -1,12361056 | 0,27940591 | 0,54587726 |
| 389337_at | ARHGEF37 | 0,13443795 | 5,23872004 | 1,12359113 | 0,27941389 | 0,54587726 |
| 6344_at | SCTR | -0,14382876 | 4,54309115 | -1,12338304 | 0,27949936 | 0,5459834 |
| 3680_at | ITGA9 | -0,12901089 | 4,66694292 | -1,12328744 | 0,27953863 | 0,5459834 |
| 4665_at | NAB2 | 0,13740573 | 6,63395136 | 1,12325184 | 0,27955325 | 0,5459834 |
| 92086_at | GGTLC1 | -0,18259758 | 5,38327722 | -1,12311183 | 0,27961078 | 0,54604039 |
| 22973_at | LAMB2P1 | -0,11086905 | 4,92632074 | -1,12300586 | 0,27965432 | 0,54607007 |
| 6878_at | TAF6 | -0,17731599 | 8,53765225 | -1,12255601 | 0,27983923 | 0,54637575 |
| 85438_at | CABS1 | -0,08112657 | 2,8230034 | -1,12244287 | 0,27988575 | 0,5464062 |
| 157506_at | RDH10 | 0,17645217 | 9,00242922 | 1,12238011 | 0,27991156 | 0,5464062 |
| 79940_at | LINC00472 | -0,08412603 | 3,17775812 | -1,12211881 | 0,28001903 | 0,54653673 |
| 9885_at | OSBPL2 | 0,22761041 | 8,30070879 | 1,12204451 | 0,28004959 | 0,54653673 |
| 169436_at | STKLD1 | -0,11169564 | 4,62084014 | -1,12201059 | 0,28006355 | 0,54653673 |
| 100507240_a | VWA8-AS1 | -0,06849963 | 3,27955059 | -1,12184736 | 0,28013071 | 0,54658262 |
| 6837_at | MED22 | -0,09907283 | 7,87528099 | -1,12181551 | 0,28014381 | 0,54658262 |
| 1046_at | CDX4 | -0,11417813 | 3,784152 | -1,12169034 | 0,28019532 | 0,54662775 |
| 5730_at | PTGDS | -0,0964306 | 5,7919498 | -1,12159498 | 0,28023457 | 0,54664896 |
| 55853_at | IDI2-AS1 | -0,24534261 | 5,41254575 | -1,12142375 | 0,28030505 | 0,54673108 |
| 5650_at | KLK7 | -0,13916415 | 4,95286202 | -1,12134708 | 0,28033661 | 0,54673728 |
| 5724_at | PTAFR | 0,28649228 | 7,0279083 | 1,12110083 | 0,280438 | 0,54683531 |
| 57553_at | MICAL3 | -0,15531627 | 6,0455551 | -1,12108712 | 0,28044365 | 0,54683531 |
| 54097_at | FAM3B | 0,12214196 | 4,31653427 | 1,12052863 | 0,28067371 | 0,5472136 |
| 5635_at | PRPSAP1 | -0,09095453 | 8,97625542 | -1,12047826 | 0,28069447 | 0,5472136 |
| 10633_at | RASL10A | -0,15502744 | 5,2838141 | -1,12011968 | 0,28084227 | 0,54744633 |
| 548645_at | DNAJC25 | -0,14504093 | 6,88029809 | -1,11975332 | 0,28099333 | 0,54768537 |
| 285590_at | SH3PXD2B | -0,12167743 | 6,67634688 | -1,11959817 | 0,28105732 | 0,5477513 |
| 4846_at | NOS3 | -0,13770755 | 4,9178545 | -1,11953343 | 0,28108403 | 0,5477513 |
| 23271_at | CAMSAP2 | 0,17497194 | 8,6123294 | 1,11925006 | 0,28120094 | 0,5479237 |
| 388882_at | LOC388882 | -0,08711786 | 4,70540904 | -1,11908358 | 0,28126965 | 0,54800214 |
| 2645_at | GCK | -0,10997624 | 5,21839764 | -1,11859312 | 0,28147212 | 0,5482895 |
| 56132_at | PCDHB3 | 0,08331366 | 3,99459289 | 1,11858843 | 0,28147406 | 0,5482895 |
| 121838_at | LINC00284 | 0,14177929 | 3,61089298 | 1,11804459 | 0,28169871 | 0,54856654 |
| 132946_at | ARL9 | 0,28732795 | 5,40784575 | 1,11801084 | 0,28171265 | 0,54856654 |
| 57482_at | KIAA1211 | 0,46100332 | 4,47642841 | 1,11799988 | 0,28171718 | 0,54856654 |
| 3770_at | KCNJ14 | -0,16922163 | 5,08847142 | -1,11796838 | 0,2817302 | 0,54856654 |
| 55711_at | FAR2 | -0,4110271 | 6,10292185 | -1,11777527 | 0,28181 | 0,54866648 |

| | | | | | | |
|---|---|---|---|---|---|---|
| 718_at | C3 | 0,6847479 | 9,1966408 | 1,1176763 | 0,28185091 | 0,54869066 |
| 80207_at | OPA3 | 0,07172733 | 6,28205567 | 1,11755459 | 0,28190123 | 0,54873316 |
| 94234_at | FOXQ1 | 0,21175845 | 10,8440764 | 1,1169582 | 0,28214787 | 0,54915776 |
| 101060091_a | LOC10106009 | 0,24400757 | 4,06913055 | 1,11677213 | 0,28222485 | 0,5492521 |
| 641518_at | LEF1-AS1 | -0,10122631 | 3,41909994 | -1,11663324 | 0,28228232 | 0,54928537 |
| 3802_at | KIR2DL1 | 0,12341025 | 4,27438239 | 1,116593 | 0,28229897 | 0,54928537 |
| 5922_at | RASA2 | 0,17461463 | 6,73872906 | 1,11631873 | 0,2824125 | 0,54943813 |
| 23037_at | PDZD2 | 0,08741954 | 3,73300415 | 1,11626552 | 0,28243453 | 0,54943813 |
| 101928784_a | RORA-AS1 | 0,0958055 | 3,31132076 | 1,11594243 | 0,28256831 | 0,54957542 |
| 9217_at | VAPB | -0,06064885 | 8,7442465 | -1,11589156 | 0,28258938 | 0,54957542 |
| 29985_at | SLC39A3 | -0,08483484 | 6,63195687 | -1,11588839 | 0,2825907 | 0,54957542 |
| 57577_at | KIAA1407 | 0,41589551 | 5,57179924 | 1,11577335 | 0,28263835 | 0,5496126 |
| 154215_at | NKAIN2 | 0,46891064 | 5,22116724 | 1,11567139 | 0,28268058 | 0,54962437 |
| 7022_at | TFAP2C | 0,41001031 | 7,97830245 | 1,11561112 | 0,28270555 | 0,54962437 |
| 5119_at | CHMP1A | -0,06810002 | 9,16233942 | -1,11555213 | 0,28272999 | 0,54962437 |
| 92840_at | REEP6 | 0,22174911 | 7,70341408 | 1,11538513 | 0,28279919 | 0,5496509 |
| 8336_at | HIST1H2AM | -0,18583565 | 6,40966954 | -1,11538146 | 0,28280071 | 0,5496509 |
| 23017_at | FAIM2 | -0,12726786 | 5,26190669 | -1,11529632 | 0,282836 | 0,54965635 |
| 51167_at | CYB5R4 | -0,23922595 | 7,39457821 | -1,115193 | 0,28287882 | 0,54965635 |
| 126370_at | OR1I1 | -0,14658169 | 5,57596176 | -1,11513392 | 0,28290331 | 0,54965635 |
| 29108_at | PYCARD | 0,43681787 | 9,36378284 | 1,115038 | 0,28294307 | 0,54965635 |
| 4705_at | NDUFA10 | -0,11418664 | 5,92893925 | -1,11495963 | 0,28297556 | 0,54965635 |
| 642924_at | LINC00535 | -0,07072248 | 4,01360037 | -1,11495268 | 0,28297844 | 0,54965635 |
| 3759_at | KCNJ2 | -0,41326485 | 4,58460983 | -1,11489284 | 0,28300326 | 0,54965635 |
| 81551_at | STMN4 | -0,123051 | 4,80063682 | -1,11479591 | 0,28304345 | 0,549679 |
| 57646_at | USP28 | -0,10307716 | 7,07943465 | -1,11465336 | 0,28310256 | 0,54972244 |
| 55966_at | AJAP1 | -0,09257272 | 4,10411367 | -1,11460434 | 0,28312289 | 0,54972244 |
| 7852_at | CXCR4 | -0,07928359 | 3,62428128 | -1,11393664 | 0,28339993 | 0,55018489 |
| 265_at | AMELX | -0,10486201 | 3,30468235 | -1,11389264 | 0,28341819 | 0,55018489 |
| 57458_at | TMCC3 | 0,36154298 | 5,80051174 | 1,11376857 | 0,2834697 | 0,55022942 |
| 93621_at | MRFAP1 | -0,06034797 | 11,8171772 | -1,11345306 | 0,2836007 | 0,55042824 |
| 284688_at | LINC01142 | -0,11556163 | 3,6378322 | -1,11322952 | 0,28369354 | 0,55055297 |
| 326624_at | RAB37 | -0,15224873 | 4,0408516 | -1,11310712 | 0,28374439 | 0,55059617 |
| 3352_at | HTR1D | -0,1297324 | 4,16926449 | -1,11283194 | 0,28385873 | 0,55076257 |
| 1408_at | CRY2 | 0,17859805 | 6,61344955 | 1,11261911 | 0,28394719 | 0,55083602 |
| 1350_at | COX7C | -0,15967159 | 9,02421352 | -1,11253588 | 0,28398178 | 0,55083602 |
| 114770_at | PGLYRP2 | -0,09755817 | 4,79935497 | -1,11237228 | 0,2840498 | 0,55083602 |
| 9681_at | DEPDC5 | 0,1252217 | 6,61591204 | 1,11230336 | 0,28407846 | 0,55083602 |
| 54957_at | TXNL4B | -0,11420024 | 6,90157456 | -1,11222938 | 0,28410922 | 0,55083602 |
| 51310_at | SLC22A17 | 0,30405457 | 6,78638619 | 1,11222288 | 0,28411192 | 0,55083602 |
| 7334_at | UBE2N | -0,14450858 | 8,97615703 | -1,11222157 | 0,28411247 | 0,55083602 |
| 3225_at | HOXC9 | 0,10309298 | 4,94413158 | 1,11219058 | 0,28412535 | 0,55083602 |
| 284440_at | LINC00663 | 0,10803296 | 4,4086471 | 1,11166856 | 0,2843425 | 0,55113457 |
| 5737_at | PTGFR | 0,08384091 | 3,14805004 | 1,11166838 | 0,28434258 | 0,55113457 |
| 130560_at | SPATA3 | -0,1259692 | 5,04970073 | -1,11161406 | 0,28436518 | 0,55113457 |
| 51374_at | ATRAID | 0,10296878 | 10,7982508 | 1,11149687 | 0,28441395 | 0,55117363 |
| 4621_at | MYH3 | 0,08892571 | 3,53366486 | 1,11135456 | 0,28447318 | 0,55123296 |
| 3157_at | HMGCS1 | -0,33565821 | 8,35935551 | -1,11121949 | 0,28452941 | 0,55128645 |
| 79825_at | EFCC1 | -0,13963932 | 5,66737754 | -1,11100067 | 0,28462051 | 0,55140751 |
| 283446_at | MYO1H | -0,11161557 | 3,34907279 | -1,1106957 | 0,28474752 | 0,55151708 |

| | | | | | | |
|---|---|---|---|---|---|---|
| 140894_at | CNBD2 | -0,05992927 | 3,42754759 | -1,11057052 | 0,28479967 | 0,55151708 |
| 729614_at | FLJ37453 | 0,14976969 | 6,32522333 | 1,11046763 | 0,28484254 | 0,55151708 |
| 57418_at | WDR18 | -0,20124621 | 9,19620311 | -1,11045414 | 0,28484815 | 0,55151708 |
| 285759_at | FLJ34503 | -0,09476568 | 3,26754899 | -1,11039129 | 0,28487435 | 0,55151708 |
| 9001_at | HAP1 | -0,09704732 | 5,80208863 | -1,11035692 | 0,28488867 | 0,55151708 |
| 101927828_a | OVOL1-AS1 | -0,11141354 | 6,07870974 | -1,11034732 | 0,28489266 | 0,55151708 |
| 1119_at | CHKA | -0,18071989 | 8,11588223 | -1,1102707 | 0,2849246 | 0,55151708 |
| 9874_at | TLK1 | 0,10562299 | 7,48991265 | 1,11024634 | 0,28493474 | 0,55151708 |
| 55975_at | KLHL7 | -0,10039437 | 7,38495543 | -1,10995979 | 0,28505418 | 0,55169283 |
| 10881_at | ACTL7A | -0,09172824 | 4,8235414 | -1,10982513 | 0,28511033 | 0,55173043 |
| 11248_at | NXPH3 | -0,11968928 | 5,69589036 | -1,10977579 | 0,2851309 | 0,55173043 |
| 7525_at | YES1 | -0,17281037 | 9,45104695 | -1,10950189 | 0,28524512 | 0,55185855 |
| 11086_at | ADAM29 | -0,09427115 | 3,2408684 | -1,10943626 | 0,28527249 | 0,55185855 |
| 390927_at | ZNF793 | 0,12067124 | 5,73392432 | 1,10941095 | 0,28528305 | 0,55185855 |
| 101928009_a | LOC10192800 | -0,14627592 | 4,32983167 | -1,10912616 | 0,28540186 | 0,55203294 |
| 22954_at | TRIM32 | 0,13188112 | 6,74875961 | 1,10897555 | 0,28546471 | 0,55209907 |
| 246312_at | C21orf91-OT1 | -0,13912196 | 3,83110644 | -1,10884097 | 0,28552088 | 0,55211774 |
| 60560_at | NAA35 | -0,10189467 | 7,47328886 | -1,10881507 | 0,28553169 | 0,55211774 |
| 3955_at | LFNG | 0,16758747 | 7,11461588 | 1,10873169 | 0,28556649 | 0,55212962 |
| 8328_at | GFI1B | -0,09719142 | 5,32395748 | -1,10841811 | 0,28569741 | 0,55232731 |
| 5924_at | RASGRF2 | -0,08405927 | 4,17384844 | -1,10830466 | 0,28574479 | 0,55234987 |
| 6367_at | CCL22 | -0,09356732 | 5,57413046 | -1,1082178 | 0,28578107 | 0,55234987 |
| 348645_at | C22orf34 | -0,15154266 | 4,71651911 | -1,10818419 | 0,28579511 | 0,55234987 |
| 101928651_a | LOC10192865 | -0,08334195 | 3,15779014 | -1,10755859 | 0,2860565 | 0,5527996 |
| 5142_at | PDE4B | -0,07566325 | 3,43540182 | -1,10746909 | 0,28609391 | 0,55281643 |
| 441061_at | MARCH11 | -0,1167001 | 4,47785049 | -1,10726624 | 0,28617872 | 0,5528385 |
| 119710_at | C11orf74 | 0,1508286 | 5,03059107 | 1,10726593 | 0,28617885 | 0,5528385 |
| 221908_at | PPP1R35 | -0,10564962 | 8,39132307 | -1,10715427 | 0,28622553 | 0,5528385 |
| 286442_at | LINC01281 | 0,07201164 | 4,1705334 | 1,10711408 | 0,28624234 | 0,5528385 |
| 90485_at | ZNF835 | -0,25555602 | 5,54680102 | -1,10709856 | 0,28624883 | 0,5528385 |
| 5829_at | PXN | 0,09002406 | 8,05422734 | 1,10674351 | 0,28639734 | 0,55306986 |
| 919_at | CD247 | -0,14723414 | 4,5755243 | -1,10639182 | 0,28654449 | 0,55329856 |
| 2668_at | GDNF | -0,09212052 | 3,85565446 | -1,1062839 | 0,28658966 | 0,55333032 |
| 6943_at | TCF21 | -0,07954831 | 3,87648693 | -1,10613371 | 0,28665252 | 0,55336747 |
| 84061_at | MAGT1 | 0,13874026 | 9,39341701 | 1,10610068 | 0,28666635 | 0,55336747 |
| 728769_at | SCAMP1-AS1 | 0,2215437 | 5,89297078 | 1,10593503 | 0,28673571 | 0,55340233 |
| 80059_at | LRRTM4 | -0,08997925 | 3,16531827 | -1,1059203 | 0,28674187 | 0,55340233 |
| 102723166_a | STAM-AS1 | -0,17631146 | 3,77054821 | -1,10572354 | 0,28682427 | 0,5535059 |
| 2996_at | GYPE | -0,10624689 | 3,78162688 | -1,10559922 | 0,28687634 | 0,55352782 |
| 101410535_a | LAMTOR5-AS1 | 0,20453289 | 5,34652821 | 1,10555921 | 0,2868931 | 0,55352782 |
| 83982_at | IFI27L2 | 0,1915631 | 8,2056634 | 1,10531091 | 0,28699712 | 0,55362494 |
| 645158_at | CBX3P2 | 0,14105085 | 3,34053963 | 1,10530186 | 0,28700091 | 0,55362494 |
| 346562_at | GNAT3 | -0,07177894 | 2,94202232 | -1,10521005 | 0,28703938 | 0,5536437 |
| 23414_at | ZFPM2 | -0,09172527 | 3,24193407 | -1,10495715 | 0,28714538 | 0,55367733 |
| 80069_at | LINC00574 | -0,11527332 | 5,50854974 | -1,10495276 | 0,28714721 | 0,55367733 |
| 3866_at | KRT15 | 0,22149116 | 8,18273747 | 1,10490945 | 0,28716537 | 0,55367733 |
| 171558_at | PTCRA | -0,12416096 | 6,63764242 | -1,10489413 | 0,28717179 | 0,55367733 |
| 81537_at | SGPP1 | 0,55001529 | 9,13996929 | 1,10462528 | 0,28728451 | 0,55379379 |
| 2902_at | GRIN1 | -0,09861137 | 5,73525475 | -1,10460745 | 0,28729198 | 0,55379379 |
| 79791_at | FBXO31 | -0,09539706 | 7,04354948 | -1,10454436 | 0,28731844 | 0,55379379 |

| | | | | | | |
|---|---|---|---|---|---|---|
| 283416_at | LINC01465 | -0,12172305 | 4,38659688 | -1,1043392 | 0,28740448 | 0,55390421 |
| 23676_at | SMPX | 0,09275227 | 3,51527591 | 1,10415788 | 0,28748054 | 0,55399537 |
| 688_at | KLF5 | 0,11600057 | 10,6374498 | 1,10394703 | 0,28756901 | 0,55402456 |
| 151393_at | RMDN2 | -0,10263502 | 5,03959449 | -1,10380945 | 0,28762674 | 0,55402456 |
| 124602_at | KIF19 | 0,09358455 | 4,37914665 | 1,10379881 | 0,28763121 | 0,55402456 |
| 9912_at | ARHGAP44 | 0,24967144 | 4,86111632 | 1,10373337 | 0,28765868 | 0,55402456 |
| 23071_at | ERP44 | 0,16201174 | 7,89680544 | 1,1037292 | 0,28766043 | 0,55402456 |
| 493869_at | GPX8 | 0,28973642 | 7,74389932 | 1,10371053 | 0,28766826 | 0,55402456 |
| 51716_at | CES1P1 | -0,09072989 | 6,77809251 | -1,10358634 | 0,28772039 | 0,55406956 |
| 55344_at | PLCXD1 | -0,19247923 | 6,43325935 | -1,10307556 | 0,28793487 | 0,55436145 |
| 335_at | APOA1 | -0,12834194 | 4,92817601 | -1,10304408 | 0,28794809 | 0,55436145 |
| 26529_at | OR12D2 | -0,08302364 | 4,46224976 | -1,10298229 | 0,28797405 | 0,55436145 |
| 221241_at | LINC00305 | -0,10621171 | 3,01689226 | -1,10295127 | 0,28798708 | 0,55436145 |
| 55609_at | ZNF280C | -0,33389667 | 7,54932466 | -1,10282719 | 0,2880392 | 0,55438233 |
| 7221_at | TRPC2 | -0,11575525 | 5,52267029 | -1,10278844 | 0,28805548 | 0,55438233 |
| 63974_at | NEUROD6 | -0,13391247 | 4,33058642 | -1,10251571 | 0,28817009 | 0,55454749 |
| 374569_at | ASPG | -0,09944595 | 7,17735219 | -1,10244455 | 0,28819999 | 0,55454805 |
| 643988_at | C1orf233 | -0,11375477 | 6,84862152 | -1,10237803 | 0,28822796 | 0,55454805 |
| 124056_at | NOXO1 | -0,13909766 | 5,21606777 | -1,10221015 | 0,28829853 | 0,55462843 |
| 7477_at | WNT7B | 0,10100706 | 7,52611996 | 1,10172604 | 0,28850211 | 0,55490971 |
| 6374_at | CXCL5 | 0,07516938 | 3,74033703 | 1,10172546 | 0,28850235 | 0,55490971 |
| 148867_at | SLC30A7 | 0,12853081 | 8,75420815 | 1,10144772 | 0,2886192 | 0,55502786 |
| 9210_at | BMP15 | -0,14621498 | 3,94389921 | -1,10144248 | 0,2886214 | 0,55502786 |
| 284749_at | LINC00494 | -0,10898504 | 4,00633568 | -1,10112457 | 0,28875519 | 0,55522971 |
| 100996280_a | LINC00911 | -0,15000864 | 3,57317084 | -1,10099019 | 0,28881175 | 0,55528305 |
| 10311_at | DSCR3 | 0,14741985 | 8,18003935 | 1,10087682 | 0,28885949 | 0,55531448 |
| 285847_at | LOC285847 | -0,1220807 | 4,25374468 | -1,10081441 | 0,28888576 | 0,55531448 |
| 100506548_a | LOC10050654 | -0,11466084 | 6,38076955 | -1,10063031 | 0,28896328 | 0,55535974 |
| 57448_at | BIRC6 | 0,06628499 | 8,8057626 | 1,10061124 | 0,28897132 | 0,55535974 |
| 246176_at | GAS2L2 | -0,14887985 | 5,56885638 | -1,10041065 | 0,2890558 | 0,55535974 |
| 6280_at | S100A9 | -0,26059987 | 10,6816956 | -1,10034249 | 0,28908451 | 0,55535974 |
| 5313_at | PKLR | -0,11080483 | 4,69691607 | -1,10027635 | 0,28911238 | 0,55535974 |
| 132112_at | RTP1 | -0,08510655 | 3,96772973 | -1,10027413 | 0,28911331 | 0,55535974 |
| 8491_at | MAP4K3 | 0,20448763 | 8,46899934 | 1,10021768 | 0,2891371 | 0,55535974 |
| 126306_at | JSRP1 | -0,09582452 | 5,44704371 | -1,10021091 | 0,28913995 | 0,55535974 |
| 285093_at | RTP5 | -0,09877664 | 5,9298918 | -1,09994674 | 0,28925127 | 0,55546306 |
| 400043_at | LOC400043 | 0,16592436 | 4,95364907 | 1,0999464 | 0,28925141 | 0,55546306 |
| 255119_at | C4orf22 | -0,0726962 | 2,99873495 | -1,09973506 | 0,28934049 | 0,55557873 |
| 90624_at | LYRM7 | -0,08531081 | 7,07565169 | -1,09963923 | 0,28938089 | 0,55560093 |
| 55591_at | VEZT | 0,11646786 | 7,69903208 | 1,09938637 | 0,28948751 | 0,55575024 |
| 1781_at | DYNC1I2 | 0,16329234 | 5,80453763 | 1,09892588 | 0,28968175 | 0,55606771 |
| 57493_at | HEG1 | 0,56455662 | 8,26785441 | 1,09865699 | 0,28979521 | 0,55609475 |
| 5239_at | PGM5 | -0,07744172 | 3,91126457 | -1,09860157 | 0,2898186 | 0,55609475 |
| 54812_at | AFTPH | 0,14080046 | 8,73223242 | 1,09855768 | 0,28983713 | 0,55609475 |
| 84249_at | PSD2 | -0,09569748 | 5,26765357 | -1,09852094 | 0,28985264 | 0,55609475 |
| 353189_at | SLCO4C1 | 0,21313255 | 3,58166847 | 1,09850223 | 0,28986053 | 0,55609475 |
| 51715_at | RAB23 | -0,20461931 | 7,23898087 | -1,09848207 | 0,28986904 | 0,55609475 |
| 285386_at | TPRG1 | -0,67932862 | 5,71554139 | -1,09819162 | 0,28999167 | 0,55627459 |
| 4833_at | NME4 | -0,21220049 | 8,30921908 | -1,09805215 | 0,29005056 | 0,55628862 |
| 101928443_a | LOC10192844 | -0,15956808 | 3,98712858 | -1,09803754 | 0,29005673 | 0,55628862 |

| | | | | | | |
|---|---|---|---|---|---|---|
| 2006_at | ELN | -0,14690728 | 6,01308514 | -1,09766477 | 0,29021419 | 0,55647695 |
| 283089_at | WDR11-AS1 | -0,10157271 | 3,98321935 | -1,09758758 | 0,2902468 | 0,55647695 |
| 27134_at | TJP3 | 0,2100476 | 6,54585392 | 1,09753488 | 0,29026908 | 0,55647695 |
| 101927195_a | LINC00382 | -0,08611751 | 3,41937797 | -1,09736838 | 0,29033944 | 0,55647695 |
| 9138_at | ARHGEF1 | -0,10819453 | 7,3238666 | -1,09735132 | 0,29034665 | 0,55647695 |
| 2853_at | GPR31 | -0,17792309 | 4,68467378 | -1,09733523 | 0,29035345 | 0,55647695 |
| 577_at | ADGRB3 | 0,19055648 | 3,30576814 | 1,09732646 | 0,29035715 | 0,55647695 |
| 197259_at | MLKL | -0,39824674 | 6,5678075 | -1,09712908 | 0,29044059 | 0,55652825 |
| 60489_at | APOBEC3G | 1,01134485 | 6,43716343 | 1,09712644 | 0,2904417 | 0,55652825 |
| 8395_at | PIP5K1B | 0,37085926 | 4,43885351 | 1,09704089 | 0,29047787 | 0,55654219 |
| 259249_at | MRGPRX1 | -0,10343085 | 3,84305691 | -1,09678538 | 0,29058591 | 0,55669381 |
| 338862_at | LINC00550 | -0,09624257 | 3,0034325 | -1,09657022 | 0,29067691 | 0,55681277 |
| 150622_at | LINC01105 | -0,06856238 | 3,34449191 | -1,09643517 | 0,29073404 | 0,55686683 |
| 56105_at | PCDHGA11 | -0,10402579 | 3,60557409 | -1,09627391 | 0,29080227 | 0,55694214 |
| 27237_at | ARHGEF16 | -0,09311614 | 6,91330216 | -1,09615439 | 0,29085284 | 0,55695696 |
| 9783_at | RIMS3 | 0,14438237 | 6,07030536 | 1,09611897 | 0,29086783 | 0,55695696 |
| 338_at | APOB | -0,09887919 | 3,95820018 | -1,09591807 | 0,29095286 | 0,55706441 |
| 1760_at | DMPK | 0,0997144 | 6,61432078 | 1,0957754 | 0,29101325 | 0,55712466 |
| 101926928_a | CRAT37 | -0,09966022 | 3,00405892 | -1,09566737 | 0,29105899 | 0,55713538 |
| 9391_at | CIAO1 | -0,06081348 | 9,18935145 | -1,09560106 | 0,29108707 | 0,55713538 |
| 64838_at | FNDC4 | -0,14486234 | 5,76724073 | -1,09555229 | 0,29110772 | 0,55713538 |
| 26689_at | OR4D1 | -0,10325279 | 4,30351958 | -1,09534676 | 0,29119477 | 0,55713538 |
| 55670_at | PEX26 | -0,08861089 | 7,1780797 | -1,09531258 | 0,29120924 | 0,55713538 |
| 112936_at | VPS26B | 0,05758046 | 8,04428801 | 1,09530174 | 0,29121384 | 0,55713538 |
| 55225_at | RAVER2 | -0,10560914 | 6,99625884 | -1,09528409 | 0,29122131 | 0,55713538 |
| 5191_at | PEX7 | 0,06964155 | 6,85439484 | 1,0949217 | 0,29137484 | 0,5572898 |
| 11285_at | B4GALT7 | -0,06041185 | 7,69946741 | -1,0948312 | 0,29141319 | 0,5572898 |
| 7922_at | SLC39A7 | 0,08549553 | 8,93991412 | 1,09482213 | 0,29141704 | 0,5572898 |
| 219333_at | USP12 | -0,15886167 | 7,22364378 | -1,09482044 | 0,29141775 | 0,5572898 |
| 23120_at | ATP10B | 0,19759876 | 5,39097156 | 1,09452331 | 0,2915437 | 0,55747531 |
| 84071_at | ARMC2 | 0,14866802 | 3,53829865 | 1,09420969 | 0,29167667 | 0,55763637 |
| 104472717_a | LINC01224 | -0,09025204 | 4,1244148 | -1,09417546 | 0,29169119 | 0,55763637 |
| 23118_at | TAB2 | 0,09336812 | 8,50945217 | 1,09401952 | 0,29175733 | 0,55763637 |
| 6716_at | SRD5A2 | -0,10337755 | 3,78800099 | -1,09399239 | 0,29176884 | 0,55763637 |
| 64864_at | RFX7 | 0,28398012 | 7,90962475 | 1,09398334 | 0,29177267 | 0,55763637 |
| 3516_at | RBPJ | -0,15274206 | 8,87304308 | -1,09384511 | 0,29183131 | 0,55769311 |
| 84701_at | COX4I2 | -0,1210881 | 5,76831058 | -1,09344891 | 0,29199943 | 0,55795904 |
| 374969_at | SVBP | 0,13103008 | 7,73934064 | 1,09309892 | 0,29214801 | 0,55818757 |
| 130827_at | TMEM182 | 0,18559417 | 5,42317301 | 1,0930052 | 0,2921878 | 0,55820823 |
| 51473_at | DCDC2 | -0,09748023 | 3,4661418 | -1,09288505 | 0,29223882 | 0,55825034 |
| 9678_at | PHF14 | -0,20895109 | 6,88034489 | -1,09273253 | 0,2923036 | 0,55828563 |
| 388969_at | C2orf68 | 0,22029132 | 7,50058854 | 1,09270508 | 0,29231526 | 0,55828563 |
| 389119_at | FAM212A | -0,09795832 | 6,25845801 | -1,09260465 | 0,29235792 | 0,55831175 |
| 100287216_a | SH3RF3-AS1 | 0,09488799 | 4,10070651 | 1,09251369 | 0,29239656 | 0,5583302 |
| 6648_at | SOD2 | -0,20604177 | 7,40635042 | -1,09237795 | 0,29245424 | 0,558365 |
| 29978_at | UBQLN2 | 0,17421018 | 9,98265876 | 1,09233436 | 0,29247276 | 0,558365 |
| 28991_at | COMMD5 | -0,1803157 | 7,76137989 | -1,09193113 | 0,29264415 | 0,55858885 |
| 84662_at | GLIS2 | 0,57451965 | 7,49251591 | 1,09191476 | 0,2926511 | 0,55858885 |
| 50617_at | ATP6V0A4 | 0,24829572 | 4,62494803 | 1,0917941 | 0,2927024 | 0,55858885 |
| 51333_at | ZNF771 | -0,14018961 | 6,1541314 | -1,09176081 | 0,29271656 | 0,55858885 |

| | | | | | | |
|---|---|---|---|---|---|---|
| 478_at | ATP1A3 | -0,11332337 | 4,49902221 | -1,09170051 | 0,2927422 | 0,55858885 |
| 127396_at | ZNF684 | -0,13112681 | 4,69190472 | -1,09159194 | 0,29278837 | 0,55858885 |
| 55520_at | ELAC1 | 0,07599695 | 5,19663464 | 1,0915713 | 0,29279714 | 0,55858885 |
| 220979_at | C10orf25 | 0,23332101 | 4,45839031 | 1,09151286 | 0,292822 | 0,55858885 |
| 90121_at | TSR2 | -0,12018566 | 7,91536639 | -1,09139012 | 0,29287421 | 0,55863312 |
| 5534_at | PPP3R1 | -0,28170055 | 8,91205339 | -1,09093883 | 0,29306622 | 0,55884621 |
| 10485_at | C1orf61 | -0,15985324 | 5,99732381 | -1,09093231 | 0,29306899 | 0,55884621 |
| 442194_at | OR10C1 | -0,11889768 | 3,7410508 | -1,09091948 | 0,29307445 | 0,55884621 |
| 650669_at | GAS6-AS1 | 0,12862538 | 5,1916747 | 1,09084948 | 0,29310424 | 0,55884621 |
| 54890_at | ALKBH5 | -0,24722943 | 8,83898156 | -1,09073873 | 0,29315138 | 0,55884621 |
| 283481_at | FGF14-AS2 | -0,08311042 | 4,41871542 | -1,0906931 | 0,29317081 | 0,55884621 |
| 1310_at | COL19A1 | -0,10943489 | 4,12500778 | -1,09065035 | 0,29318901 | 0,55884621 |
| 100132911_a | DPH3P1 | -0,10471503 | 5,06204664 | -1,09053544 | 0,29323793 | 0,55888416 |
| 9887_at | SMG7 | -0,10347558 | 7,42934688 | -1,09041737 | 0,2932882 | 0,55889236 |
| 101927498_a | LINC01361 | 0,07012285 | 3,10550512 | 1,09038905 | 0,29330026 | 0,55889236 |
| 10894_at | LYVE1 | -0,08186895 | 3,3092092 | -1,09009746 | 0,29342445 | 0,5590737 |
| 101927581_a | CCDC182 | -0,08443699 | 4,81868506 | -1,08999202 | 0,29346936 | 0,55910398 |
| 728012_at | MCHR2-AS1 | -0,11502162 | 4,43305211 | -1,08967772 | 0,29360328 | 0,55915721 |
| 55296_at | TBC1D19 | 0,1809432 | 6,44245121 | 1,08965233 | 0,2936141 | 0,55915721 |
| 84900_at | RNFT2 | -0,16753848 | 5,78686666 | -1,08961751 | 0,29362894 | 0,55915721 |
| 65062_at | TMEM237 | -0,12528741 | 6,91303083 | -1,08960186 | 0,29363561 | 0,55915721 |
| 9263_at | STK17A | -0,25380377 | 6,64739994 | -1,08955822 | 0,29365421 | 0,55915721 |
| 57002_at | YAE1D1 | -0,10416579 | 7,75950559 | -1,08951772 | 0,29367147 | 0,55915721 |
| 89866_at | SEC16B | 0,12089034 | 4,12681441 | 1,08889112 | 0,29393863 | 0,55957282 |
| 7401_at | CLRN1 | -0,07209922 | 3,33629784 | -1,08886951 | 0,29394785 | 0,55957282 |
| 54360_at | CYTL1 | -0,14641751 | 4,47754649 | -1,08869801 | 0,294021 | 0,55965678 |
| 126661_at | CCDC163P | -0,06918374 | 5,95476476 | -1,08849545 | 0,29410743 | 0,55966094 |
| 440073_at | IQSEC3 | -0,08701869 | 5,18143909 | -1,0884357 | 0,29413292 | 0,55966094 |
| 346171_at | ZFP57 | -0,14375402 | 4,76488676 | -1,08839004 | 0,29415241 | 0,55966094 |
| 1026_at | CDKN1A | 0,41370198 | 9,04454697 | 1,08836442 | 0,29416334 | 0,55966094 |
| 1048_at | CEACAM5 | -0,84174621 | 5,75714876 | -1,08828282 | 0,29419816 | 0,55966094 |
| 10301_at | DLEU1 | -0,83672996 | 5,62846595 | -1,08822546 | 0,29422265 | 0,55966094 |
| 5519_at | PPP2R1B | 0,10101532 | 7,64337104 | 1,08818956 | 0,29423797 | 0,55966094 |
| 55319_at | TMA16 | -0,47530374 | 6,42135307 | -1,08814821 | 0,29425562 | 0,55966094 |
| 26223_at | FBXL21 | -0,0809273 | 3,97977056 | -1,08801847 | 0,29431101 | 0,55967729 |
| 24149_at | ZNF318 | 0,08655635 | 6,3287355 | 1,08799195 | 0,29432233 | 0,55967729 |
| 9847_at | C2CD5 | 0,12294949 | 8,77617263 | 1,08785435 | 0,29438108 | 0,55970178 |
| 65008_at | MRPL1 | -0,39602468 | 7,39367593 | -1,08782568 | 0,29439332 | 0,55970178 |
| 54944_at | LINC01521 | 0,2284064 | 5,28803313 | 1,0877358 | 0,2944317 | 0,55971951 |
| 6315_at | ATXN8OS | -0,11899849 | 3,82556998 | -1,08734182 | 0,29459998 | 0,55998416 |
| 242_at | ALOX12B | -0,11203515 | 4,38852834 | -1,08718092 | 0,29466873 | 0,56005957 |
| 9976_at | CLEC2B | -0,22652032 | 4,94854636 | -1,08704143 | 0,29472834 | 0,56011759 |
| 5961_at | PRPH2 | -0,15576059 | 4,74681169 | -1,08694311 | 0,29477036 | 0,56014219 |
| 7481_at | WNT11 | -0,44423728 | 6,37632621 | -1,08675121 | 0,29485239 | 0,5602428 |
| 55802_at | DCP1A | -0,19510425 | 7,74640619 | -1,08614617 | 0,29511112 | 0,56060919 |
| 5553_at | PRG2 | -0,08996119 | 4,43328527 | -1,0859973 | 0,29517481 | 0,56060919 |
| 100128653_a | LOC10012865 | -0,16494725 | 5,57676071 | -1,08575195 | 0,29527979 | 0,56060919 |
| 53938_at | PPIL3 | -0,14901249 | 8,99815076 | -1,08574693 | 0,29528194 | 0,56060919 |
| 6745_at | SSR1 | -0,18854568 | 9,84491615 | -1,08574113 | 0,29528443 | 0,56060919 |
| 22996_at | TTC39A | 0,09317285 | 4,91592841 | 1,08572498 | 0,29529134 | 0,56060919 |

| | | | | | | |
|---|---|---|---|---|---|---|
| 401237_at | CASC15 | 0,08217177 | 4,14839635 | 1,08570007 | 0,295302 | 0,56060919 |
| 144535_at | CFAP54 | 0,0580065 | 2,76740017 | 1,08563504 | 0,29532983 | 0,56060919 |
| 139231_at | FAM199X | -0,15672088 | 8,35366525 | -1,08563068 | 0,2953317 | 0,56060919 |
| 9397_at | NMT2 | -0,22675321 | 5,9209011 | -1,08557432 | 0,29535582 | 0,56060919 |
| 150372_at | NFAM1 | -0,09060612 | 5,86224473 | -1,08555207 | 0,29536535 | 0,56060919 |
| 8314_at | BAP1 | 0,16217038 | 7,81422648 | 1,08538643 | 0,29543625 | 0,56068853 |
| 79853_at | TM4SF20 | -0,0841141 | 3,1703859 | -1,08498163 | 0,29560959 | 0,56094089 |
| 732253_at | TDRG1 | -0,13493095 | 4,59774885 | -1,08490735 | 0,29564141 | 0,56094089 |
| 148753_at | FAM163A | -0,09092263 | 4,37001333 | -1,08487192 | 0,29565659 | 0,56094089 |
| 56967_at | C14orf132 | 0,16523495 | 5,25897027 | 1,08461221 | 0,29576785 | 0,56109673 |
| 58486_at | ZBED5 | 0,13513841 | 8,7531274 | 1,08428274 | 0,29590905 | 0,5613093 |
| 51458_at | RHCG | -0,10064464 | 4,99406522 | -1,08404015 | 0,29601304 | 0,56139992 |
| 51103_at | NDUFAF1 | -0,10234624 | 7,6697174 | -1,08403534 | 0,2960151 | 0,56139992 |
| 169522_at | KCNV2 | -0,08844744 | 3,51566806 | -1,08396025 | 0,2960473 | 0,5614057 |
| 84983_at | FAM222A-AS1 | -0,09378522 | 3,55978167 | -1,0837928 | 0,29611911 | 0,5614866 |
| 219938_at | SPATA19 | -0,13840616 | 4,3926434 | -1,08361156 | 0,29619684 | 0,56157871 |
| 10000_at | AKT3 | -0,07254067 | 3,67126841 | -1,08347493 | 0,29625545 | 0,56163456 |
| 2350_at | FOLR2 | -0,093532 | 5,43209754 | -1,08336396 | 0,29630306 | 0,56166954 |
| 5859_at | QARS | 0,10740098 | 10,8819362 | 1,08310644 | 0,29641357 | 0,56182372 |
| 59274_at | MESDC1 | -0,14009726 | 9,0332506 | -1,08262366 | 0,29662081 | 0,56206042 |
| 6665_at | SOX15 | -0,31085427 | 6,52873173 | -1,08261673 | 0,29662379 | 0,56206042 |
| 7054_at | TH | 0,13956869 | 5,50428118 | 1,08261163 | 0,29662598 | 0,56206042 |
| 8803_at | SUCLA2 | -0,14921347 | 8,98919599 | -1,0824184 | 0,29670896 | 0,56212137 |
| 1475_at | CSTA | 0,33156666 | 9,64491386 | 1,08240083 | 0,29671651 | 0,56212137 |
| 3626_at | INHBC | -0,13143866 | 4,3035459 | -1,08228544 | 0,29676607 | 0,56215998 |
| #N/A | CTC-338M12,4 | -0,1472371 | 6,83758827 | -1,081938 | 0,29691534 | 0,56235504 |
| 90313_at | TP53I13 | 0,26828094 | 7,0614083 | 1,08188575 | 0,29693779 | 0,56235504 |
| 80821_at | DDHD1 | 0,08729442 | 6,4084412 | 1,08180901 | 0,29697077 | 0,56235504 |
| 100507351_at | LOC10050735 | -0,09680611 | 4,63578546 | -1,08157655 | 0,29707069 | 0,56235504 |
| 7067_at | THRA | 0,07861741 | 6,29025439 | 1,08152606 | 0,29709239 | 0,56235504 |
| 127707_at | KLHDC7A | 0,33241606 | 6,26519533 | 1,08151463 | 0,29709731 | 0,56235504 |
| 285419_at | LINC01091 | -0,06849911 | 3,29589354 | -1,08150006 | 0,29710357 | 0,56235504 |
| 342035_at | GLDN | 0,08294343 | 4,15179098 | 1,08146765 | 0,2971175 | 0,56235504 |
| 50940_at | PDE11A | -0,06038676 | 3,44602152 | -1,08143443 | 0,29713179 | 0,56235504 |
| 9672_at | SDC3 | 0,14024132 | 6,65530275 | 1,08129262 | 0,29719276 | 0,56241518 |
| 9792_at | SERTAD2 | -0,23455043 | 7,92353491 | -1,08122378 | 0,29722236 | 0,56241595 |
| 101928684_at | LINC01209 | -0,14763986 | 3,30959494 | -1,08107839 | 0,29728489 | 0,56247902 |
| 7693_at | ZNF134 | 0,17494396 | 7,51839417 | 1,08081868 | 0,29739661 | 0,56262906 |
| 231_at | AKR1B1 | -0,44391695 | 12,9566289 | -1,08075825 | 0,29742261 | 0,56262906 |
| 9027_at | NAT8 | 0,11309639 | 3,74801171 | 1,08057833 | 0,29750003 | 0,56272025 |
| 22848_at | AAK1 | 0,08478725 | 6,33979149 | 1,08039422 | 0,29757926 | 0,56281486 |
| 51348_at | KLRF1 | 0,16896252 | 3,81985251 | 1,08019041 | 0,297667 | 0,56292552 |
| 162466_at | PHOSPHO1 | -0,10096364 | 4,31321698 | -1,07986774 | 0,29780594 | 0,56313299 |
| 8321_at | FZD1 | 0,20505213 | 6,64032162 | 1,07974978 | 0,29785674 | 0,56317377 |
| 22925_at | PLA2R1 | 0,13850226 | 4,36191953 | 1,07956724 | 0,29793537 | 0,56326249 |
| 974_at | CD79B | -0,13164003 | 4,77318191 | -1,07945521 | 0,29798363 | 0,56326249 |
| 5101_at | PCDH9 | 0,1905387 | 4,63114425 | 1,07942032 | 0,29799866 | 0,56326249 |
| 10214_at | SSX3 | -0,09957986 | 4,01777047 | -1,07936936 | 0,29802062 | 0,56326249 |
| 55619_at | DOCK10 | 0,25135303 | 3,52759744 | 1,07924711 | 0,2980733 | 0,56330678 |
| 2208_at | FCER2 | -0,08378085 | 4,52255318 | -1,07898805 | 0,29818496 | 0,56346251 |

| | | | | | | |
|---|---|---|---|---|---|---|
| 124842_at | TMEM132E | -0,11047795 | 4,77361998 | -1,07887691 | 0,29823287 | 0,56349315 |
| 101928306_a | LOC10192830 | -0,16494996 | 3,95105783 | -1,07881474 | 0,29825968 | 0,56349315 |
| 2925_at | GRPR | -0,15353087 | 4,7348947 | -1,07859601 | 0,29835399 | 0,56360356 |
| 7982_at | ST7 | -0,09840785 | 6,79456956 | -1,07854351 | 0,29837663 | 0,56360356 |
| 10612_at | TRIM3 | -0,06724647 | 4,94863409 | -1,0783076 | 0,29847839 | 0,56372474 |
| 5341_at | PLEK | -0,0933354 | 4,17339598 | -1,07815305 | 0,29854506 | 0,56372474 |
| 79642_at | ARSJ | -0,07436174 | 3,85992688 | -1,07815007 | 0,29854635 | 0,56372474 |
| 64062_at | RBM26 | -0,14716704 | 7,31133703 | -1,07812342 | 0,29855785 | 0,56372474 |
| 9358_at | ITGBL1 | 0,35030585 | 6,02628228 | 1,07803599 | 0,29859557 | 0,56374071 |
| 64123_at | ADGRL4 | 0,05975225 | 2,71996085 | 1,07782603 | 0,29868618 | 0,56378953 |
| 2296_at | FOXC1 | 0,40904584 | 10,1659978 | 1,07782046 | 0,29868858 | 0,56378953 |
| 63940_at | GPSM3 | -0,11262361 | 6,46095781 | -1,07777261 | 0,29870923 | 0,56378953 |
| 167410_at | LIX1 | -0,09290698 | 3,27552243 | -1,07748349 | 0,29883405 | 0,56396985 |
| 9869_at | SETDB1 | 0,10873357 | 6,90190869 | 1,07728791 | 0,2989185 | 0,56402343 |
| 100885778_a | NALCN-AS1 | -0,11944193 | 4,6984431 | -1,07728211 | 0,298921 | 0,56402343 |
| 8074_at | FGF23 | -0,1245697 | 4,83669777 | -1,0770167 | 0,29903564 | 0,56416064 |
| 2188_at | FANCF | 0,09638619 | 7,41060412 | 1,07697815 | 0,29905229 | 0,56416064 |
| 4063_at | LY9 | -0,14417519 | 4,67654464 | -1,07665468 | 0,29919205 | 0,56432163 |
| 2965_at | GTF2H1 | -0,13101617 | 8,2521923 | -1,07662004 | 0,29920702 | 0,56432163 |
| 51224_at | TCEB3B | -0,09869925 | 4,51066959 | -1,07657724 | 0,29922552 | 0,56432163 |
| 3484_at | IGFBP1 | -0,1321989 | 4,38941996 | -1,07608392 | 0,29943877 | 0,56465918 |
| 220416_at | LRRC63 | 0,08047586 | 2,8089081 | 1,0760276 | 0,29946312 | 0,56465918 |
| 10519_at | CIB1 | -0,16939689 | 9,80805983 | -1,07577556 | 0,29957213 | 0,56476375 |
| 10057_at | ABCC5 | 0,28761824 | 8,6984972 | 1,07571035 | 0,29960034 | 0,56476375 |
| 101929064_a | LOC10192906 | -0,0987896 | 3,71490803 | -1,07559669 | 0,29964951 | 0,56476375 |
| 2160_at | F11 | -0,07171118 | 4,79820915 | -1,07557117 | 0,29966055 | 0,56476375 |
| 284099_at | C17orf78 | -0,07638271 | 3,37118897 | -1,07556047 | 0,29966518 | 0,56476375 |
| 91734_at | IDI2 | -0,1313854 | 4,83763565 | -1,07549221 | 0,29969471 | 0,56476416 |
| 9993_at | DGCR2 | 0,07264943 | 8,24248299 | 1,07542172 | 0,29972521 | 0,56476638 |
| 388815_at | MIR99AHG | -0,07149895 | 3,07604663 | -1,07508185 | 0,2998723 | 0,56494781 |
| 7702_at | ZNF143 | -0,09881561 | 7,83551148 | -1,07502884 | 0,29989525 | 0,56494781 |
| 259197_at | NCR3 | -0,09632049 | 4,87080149 | -1,07498138 | 0,29991579 | 0,56494781 |
| 4209_at | MEF2D | 0,08777263 | 6,14039493 | 1,07492823 | 0,29993881 | 0,56494781 |
| 3305_at | HSPA1L | -0,11082143 | 5,14844536 | -1,07484667 | 0,29997411 | 0,56495907 |
| 80263_at | TRIM45 | 0,16027946 | 6,1569311 | 1,07464264 | 0,30006247 | 0,56507022 |
| 9376_at | SLC22A8 | -0,15073493 | 5,32729216 | -1,07443968 | 0,30015037 | 0,56518051 |
| 151195_at | CCNYL1 | -0,22654792 | 8,45059495 | -1,07433951 | 0,30019376 | 0,56520696 |
| 115019_at | SLC26A9 | -0,07679225 | 4,70175382 | -1,07419223 | 0,30025757 | 0,56527185 |
| 171220_at | DSTNP2 | 0,13913237 | 5,96958045 | 1,07397136 | 0,30035328 | 0,56539677 |
| 353132_at | LCE1B | -0,16352639 | 4,13442831 | -1,07364963 | 0,30049273 | 0,56549908 |
| 79748_at | LMAN1L | -0,10601267 | 5,88135094 | -1,07355188 | 0,30053511 | 0,56549908 |
| 285237_at | C3orf38 | 0,22610585 | 8,23175688 | 1,07351648 | 0,30055046 | 0,56549908 |
| 148266_at | ZNF569 | 0,20042882 | 4,69788136 | 1,07345898 | 0,30057539 | 0,56549908 |
| 54510_at | PCDH18 | 0,22688586 | 4,6967616 | 1,07344155 | 0,30058295 | 0,56549908 |
| 94039_at | ZNF101 | -0,13858559 | 6,31115363 | -1,07343967 | 0,30058376 | 0,56549908 |
| 9470_at | EIF4E2 | -0,07100279 | 8,9014206 | -1,07322488 | 0,30067691 | 0,56561908 |
| 114815_at | SORCS1 | -0,0943296 | 3,45398805 | -1,07303859 | 0,30075772 | 0,56571584 |
| 22987_at | SV2C | -0,05616272 | 3,4301738 | -1,07224461 | 0,3011023 | 0,56626545 |
| 22938_at | SNW1 | -0,06562934 | 9,29564998 | -1,07222985 | 0,30110871 | 0,56626545 |
| 642987_at | TMEM232 | 0,06728154 | 2,90930216 | 1,07195355 | 0,30122869 | 0,56643578 |

| | | | | | | |
|---|---|---|---|---|---|---|
| 6524_at | SLC5A2 | -0,10385395 | 6,19384461 | -1,07175193 | 0,30131626 | 0,56649094 |
| 51665_at | ASB1 | -0,08456769 | 6,38740525 | -1,07173897 | 0,30132189 | 0,56649094 |
| 25830_at | SULT4A1 | -0,14728116 | 5,73803921 | -1,07168291 | 0,30134624 | 0,56649094 |
| 51503_at | CWC15 | -0,1029716 | 9,00486269 | -1,07160415 | 0,30138046 | 0,56649998 |
| 4624_at | MYH6 | -0,11038685 | 3,75502296 | -1,07152045 | 0,30141683 | 0,56651306 |
| 54952_at | TRNAU1AP | 0,13010663 | 6,83808671 | 1,07137881 | 0,30147837 | 0,56657346 |
| 3049_at | HBQ1 | -0,15561468 | 5,70321315 | -1,07129401 | 0,30151523 | 0,56658744 |
| 2918_at | GRM8 | -0,08346397 | 3,51934231 | -1,07116927 | 0,30156945 | 0,56661532 |
| 55964_at | SEPT3 | -0,09425869 | 5,60728346 | -1,07106836 | 0,30161331 | 0,56661532 |
| 5024_at | P2RX3 | -0,08964188 | 3,7693381 | -1,07102998 | 0,30163 | 0,56661532 |
| 23336_at | SYNM | -0,15168217 | 6,8184874 | -1,07098921 | 0,30164772 | 0,56661532 |
| 619434_at | LINC00051 | -0,113648 | 3,57246504 | -1,07085886 | 0,3017044 | 0,5666235 |
| 9721_at | GPRIN2 | -0,09388326 | 5,3089442 | -1,0708007 | 0,30172968 | 0,5666235 |
| 647310_at | TEX22 | -0,13256044 | 5,73761982 | -1,07077622 | 0,30174033 | 0,5666235 |
| 2301_at | FOXE3 | -0,10230747 | 5,27501566 | -1,07066103 | 0,30179042 | 0,56666056 |
| 7155_at | TOP2B | 0,33510296 | 9,96829039 | 1,07052977 | 0,30184751 | 0,56666056 |
| 1524_at | CX3CR1 | 0,08904815 | 3,61316023 | 1,07051845 | 0,30185244 | 0,56666056 |
| 1677_at | DFFB | 0,104216 | 6,63064136 | 1,0704603 | 0,30187773 | 0,56666056 |
| 23788_at | MTCH2 | -0,15948505 | 8,90083053 | -1,07028188 | 0,30195535 | 0,56673281 |
| 6271_at | S100A1 | -0,1086069 | 5,24370967 | -1,07014731 | 0,3020139 | 0,56673281 |
| 56154_at | TEX15 | -0,06981142 | 3,03579714 | -1,07014 | 0,30201708 | 0,56673281 |
| 83661_at | MS4A8 | -0,10369771 | 4,53911409 | -1,07007064 | 0,30204726 | 0,56673281 |
| 83416_at | FCRL5 | -0,13109637 | 4,97806341 | -1,07003374 | 0,30206332 | 0,56673281 |
| 54108_at | CHRAC1 | -0,26288461 | 8,16206722 | -1,06994446 | 0,30210218 | 0,56675052 |
| 26050_at | SLITRK5 | -0,12667331 | 4,40276461 | -1,06984172 | 0,3021469 | 0,56677921 |
| 100506325_a | SP2-AS1 | -0,21126616 | 4,81326877 | -1,06936516 | 0,30235438 | 0,5671132 |
| 3061_at | HCRTR1 | -0,16437935 | 4,24838163 | -1,06897869 | 0,30252273 | 0,5673737 |
| 10286_at | BCAS2 | -0,15467399 | 9,27180397 | -1,06881905 | 0,30259228 | 0,56744891 |
| 6768_at | ST14 | 0,21626519 | 9,59374225 | 1,06856803 | 0,30270168 | 0,5675988 |
| 84465_at | MEGF11 | -0,08660344 | 4,69670889 | -1,06843793 | 0,3027584 | 0,56764989 |
| 84688_at | C9orf24 | -0,11810252 | 4,68109715 | -1,06820515 | 0,30285988 | 0,56772717 |
| 100128998_a | C20orf181 | -0,12703455 | 5,28059989 | -1,0681405 | 0,30288807 | 0,56772717 |
| 65979_at | PHACTR4 | 0,13655573 | 8,95853056 | 1,06811385 | 0,30289969 | 0,56772717 |
| 7322_at | UBE2D2 | -0,16530365 | 9,49298315 | -1,06807301 | 0,3029175 | 0,56772717 |
| 10129_at | FRY | 0,15842907 | 6,76465171 | 1,06781767 | 0,30302887 | 0,56785184 |
| 81669_at | CCNL2 | 0,26151368 | 8,1405768 | 1,06777017 | 0,30304959 | 0,56785184 |
| 27248_at | ERLEC1 | 0,18805991 | 9,13876725 | 1,06771776 | 0,30307246 | 0,56785184 |
| 101927588_a | LOC10192758 | -0,1021103 | 4,28274625 | -1,06761032 | 0,30311933 | 0,56788443 |
| 90993_at | CREB3L1 | -0,10421261 | 4,96249288 | -1,0674165 | 0,3032039 | 0,56798763 |
| 1785_at | DNM2 | 0,10719636 | 8,29017395 | 1,06716705 | 0,30331278 | 0,56813634 |
| 100190939_a | TPT1-AS1 | 0,09873513 | 6,53880184 | 1,06679134 | 0,30347682 | 0,56832137 |
| 3309_at | HSPA5 | -0,25231453 | 9,12799564 | -1,066739 | 0,30349967 | 0,56832137 |
| 1949_at | EFNB3 | 0,08863927 | 5,79227579 | 1,06673807 | 0,30350008 | 0,56832137 |
| 3005_at | H1F0 | 0,49467257 | 9,71185023 | 1,06654189 | 0,30358576 | 0,56842657 |
| 142685_at | ASB15 | -0,09930669 | 4,06763973 | -1,06627987 | 0,30370023 | 0,56858466 |
| 64926_at | RASAL3 | -0,13211751 | 5,93568892 | -1,0662135 | 0,30372923 | 0,56858466 |
| 83597_at | RTP3 | -0,1314121 | 4,07771122 | -1,06607423 | 0,30379009 | 0,56864333 |
| 3215_at | HOXB5 | 0,18471206 | 7,62684265 | 1,06597443 | 0,3038337 | 0,56865926 |
| 2321_at | FLT1 | -0,06565012 | 3,65642857 | -1,06588369 | 0,30387337 | 0,56865926 |
| 53347_at | UBASH3A | -0,14856839 | 3,74060953 | -1,06585213 | 0,30388716 | 0,56865926 |

| | | | | | | |
|---|---|---|---|---|---|---|
| 101927528_a | LINC01204 | -0,06294329 | 3,65746345 | -1,06570106 | 0,3039532 | 0,56871045 |
| 64582_at | GPR135 | -0,09736707 | 5,06425626 | -1,06558885 | 0,30400226 | 0,56871045 |
| 5075_at | PAX1 | -0,07742954 | 4,39890263 | -1,06558696 | 0,30400309 | 0,56871045 |
| 101928418_a | LOC10192841 | -0,0898743 | 3,29905574 | -1,0651771 | 0,30418234 | 0,56899052 |
| 219731_at | PROSER2-AS1 | -0,07756543 | 3,7016948 | -1,06505371 | 0,30423631 | 0,56903623 |
| 2863_at | GPR39 | -0,12921456 | 6,3092856 | -1,06498385 | 0,30426688 | 0,56903814 |
| 28673_at | TRAV12-2 | -0,103796 | 4,06299174 | -1,06467498 | 0,30440204 | 0,56922057 |
| 94030_at | LRRC4B | -0,09242973 | 4,61469959 | -1,0646259 | 0,30442352 | 0,56922057 |
| 339894_at | LINC00880 | -0,11666692 | 4,31167921 | -1,06429429 | 0,30456869 | 0,56934641 |
| 7091_at | TLE4 | 0,32987429 | 5,38926379 | 1,06428395 | 0,30457322 | 0,56934641 |
| 3445_at | IFNA8 | -0,08251345 | 2,97719713 | -1,06426233 | 0,30458268 | 0,56934641 |
| 158056_at | MAMDC4 | -0,20254647 | 6,36712176 | -1,06420211 | 0,30460905 | 0,56934641 |
| 6556_at | SLC11A1 | -0,09864018 | 4,90917403 | -1,06408096 | 0,30466211 | 0,56935193 |
| 1488_at | CTBP2 | -0,19243183 | 7,67809351 | -1,06399099 | 0,30470151 | 0,56935193 |
| 26692_at | OR2W1 | -0,11641424 | 3,31686803 | -1,06396787 | 0,30471163 | 0,56935193 |
| 56681_at | SAR1A | 0,08917174 | 9,40935513 | 1,06392464 | 0,30473057 | 0,56935193 |
| 50614_at | GALNT9 | -0,10436855 | 5,39340435 | -1,06385793 | 0,30475979 | 0,56935193 |
| 8739_at | HRK | -0,16235599 | 4,65920649 | -1,06360534 | 0,30487045 | 0,56950344 |
| 27077_at | B9D1 | -0,52373323 | 6,80109436 | -1,06311473 | 0,30508547 | 0,56984984 |
| 1646_at | AKR1C2 | -0,14730567 | 11,9451421 | -1,0629793 | 0,30514485 | 0,56990549 |
| 2738_at | GLI4 | 0,13245653 | 5,00427923 | 1,06273221 | 0,30525319 | 0,5700143 |
| 414260_at | LINC00619 | 0,0938572 | 3,63689039 | 1,06268018 | 0,30527601 | 0,5700143 |
| 168433_at | RNF133 | -0,06863202 | 3,41676005 | -1,062644 | 0,30529188 | 0,5700143 |
| 23746_at | AIPL1 | -0,07480236 | 5,13833167 | -1,0624687 | 0,30536878 | 0,57010261 |
| 283483_at | LINC00551 | 0,0720179 | 3,11766584 | 1,06196284 | 0,30559075 | 0,57046174 |
| 9242_at | MSC | -0,0893304 | 4,48940712 | -1,06187195 | 0,30563065 | 0,57048092 |
| 27143_at | PALD1 | -0,1102173 | 5,00505095 | -1,06158725 | 0,30575564 | 0,57065893 |
| 131540_at | ZDHHC19 | -0,11165523 | 5,50046424 | -1,06123726 | 0,30590934 | 0,57086455 |
| 254559_at | LINC00925 | -0,10928502 | 4,47061178 | -1,06120144 | 0,30592508 | 0,57086455 |
| 597_at | BCL2A1 | -0,07467808 | 3,20918971 | -1,06101973 | 0,30600491 | 0,57092683 |
| 114771_at | PGLYRP3 | -0,14027039 | 3,87769177 | -1,06099054 | 0,30601773 | 0,57092683 |
| 91461_at | PKDCC | -0,20048162 | 5,86696865 | -1,06081646 | 0,30609422 | 0,57101424 |
| 147495_at | APCDD1 | 0,1389935 | 4,96955697 | 1,06048244 | 0,30624103 | 0,57123279 |
| 63976_at | PRDM16 | -0,0920912 | 4,41581238 | -1,06013089 | 0,30639561 | 0,57146578 |
| 8086_at | AAAS | -0,07947051 | 8,03238192 | -1,05987373 | 0,30650871 | 0,57162138 |
| 84069_at | PLEKHN1 | -0,09196654 | 6,6387664 | -1,05936975 | 0,30673046 | 0,57197957 |
| 768206_at | PRCD | -0,1309852 | 5,09912203 | -1,05921481 | 0,30679866 | 0,57205051 |
| 651746_at | ANKRD33B | 0,19739922 | 5,97163418 | 1,05914839 | 0,3068279 | 0,57205051 |
| 9578_at | CDC42BPB | 0,13665777 | 8,48269996 | 1,05873706 | 0,30700901 | 0,5723072 |
| 9843_at | HEPH | -0,09513026 | 3,25498974 | -1,05870076 | 0,307025 | 0,5723072 |
| 100505984_a | LOC10050598 | -0,13161481 | 3,44732231 | -1,05861329 | 0,30706353 | 0,57232363 |
| 402_at | ARL2 | 0,09588296 | 9,23830239 | 1,05804896 | 0,30731217 | 0,57263264 |
| 56257_at | MEPCE | -0,22267705 | 8,64715519 | -1,05798675 | 0,30733959 | 0,57263264 |
| 932_at | MS4A3 | -0,07591245 | 2,92659723 | -1,05789582 | 0,30737967 | 0,57263264 |
| 55208_at | DCUN1D2 | -0,10548139 | 5,95012882 | -1,05783958 | 0,30740446 | 0,57263264 |
| 8342_at | HIST1H2BM | -0,14487698 | 6,09243641 | -1,05783407 | 0,30740689 | 0,57263264 |
| 100862679_a | NAALADL2-AS | -0,08893745 | 2,67725279 | -1,05770084 | 0,30746563 | 0,57263264 |
| 7469_at | NELFA | -0,06146799 | 8,54532414 | -1,0576977 | 0,30746701 | 0,57263264 |
| 85437_at | ZCRB1 | 0,08101032 | 9,78902895 | 1,05763905 | 0,30749287 | 0,57263264 |
| 1359_at | CPA3 | -0,19033103 | 3,50157499 | -1,05762999 | 0,30749686 | 0,57263264 |

| | | | | | | |
|---|---|---|---|---|---|---|
| 23603_at | CORO1C | -0,24813381 | 10,0859415 | -1,0573381 | 0,30762558 | 0,57278675 |
| 50863_at | NTM | -0,08944345 | 3,6817257 | -1,05730749 | 0,30763908 | 0,57278675 |
| 148756_at | OVAAL | -0,12178643 | 4,38230199 | -1,05718307 | 0,30769397 | 0,57283116 |
| 441549_at | CDNF | 0,19583481 | 3,48776483 | 1,0571186 | 0,30772241 | 0,57283116 |
| 129025_at | ZNF280A | -0,07382449 | 3,29951587 | -1,05698143 | 0,30778293 | 0,5728711 |
| 162387_at | MFSD6L | -0,09759077 | 5,21949694 | -1,05693515 | 0,30780335 | 0,5728711 |
| 90589_at | ZNF625 | 0,06957365 | 4,71582581 | 1,05657727 | 0,30796129 | 0,57310773 |
| 8123_at | PWAR5 | 0,06763375 | 3,5695074 | 1,0564424 | 0,30802083 | 0,57310773 |
| 5244_at | ABCB4 | 0,07845318 | 3,79243692 | 1,05636543 | 0,30805481 | 0,57310773 |
| 162515_at | SLC16A11 | -0,12007635 | 5,08882097 | -1,05631338 | 0,30807779 | 0,57310773 |
| 81856_at | ZNF611 | 0,11570161 | 6,42400366 | 1,05623663 | 0,30811168 | 0,57310773 |
| 10265_at | IRX5 | 0,12110264 | 6,64486082 | 1,05621849 | 0,30811969 | 0,57310773 |
| 818_at | CAMK2G | 0,15025271 | 8,44301618 | 1,05607023 | 0,30818516 | 0,57310773 |
| 5858_at | PZP | -0,10439362 | 4,02496297 | -1,05605418 | 0,30819225 | 0,57310773 |
| 414235_at | PRR26 | -0,0765553 | 4,03444897 | -1,05593344 | 0,30824558 | 0,57310773 |
| 285051_at | C2orf61 | -0,12956615 | 4,51308472 | -1,05591273 | 0,30825473 | 0,57310773 |
| 8468_at | FKBP6 | -0,07713185 | 5,00114899 | -1,05590587 | 0,30825776 | 0,57310773 |
| 10864_at | SLC22A7 | -0,09706694 | 4,89463954 | -1,05570795 | 0,30834519 | 0,57317917 |
| 147872_at | CCDC155 | -0,14218027 | 4,44091873 | -1,05563454 | 0,30837763 | 0,57317917 |
| 29065_at | ASAP1-IT1 | -0,14745494 | 4,08139699 | -1,05561684 | 0,30838545 | 0,57317917 |
| 57534_at | MIB1 | 0,20951466 | 7,75902024 | 1,05548583 | 0,30844334 | 0,5731854 |
| 387638_at | C10orf113 | 0,09437073 | 3,41449873 | 1,05547457 | 0,30844831 | 0,5731854 |
| 101930452_a | LOC10193045 | -0,12484592 | 4,69299573 | -1,0548764 | 0,30871274 | 0,57362145 |
| 124808_at | CCDC43 | -0,10809831 | 7,785043 | -1,05459689 | 0,30883636 | 0,57379579 |
| 10007_at | GNPDA1 | 0,32129319 | 9,32099633 | 1,05439233 | 0,30892685 | 0,57387101 |
| 55719_at | SLF2 | -0,09388347 | 6,83353044 | -1,05437069 | 0,30893642 | 0,57387101 |
| 8471_at | IRS4 | -0,06401469 | 3,7606606 | -1,05407081 | 0,30906913 | 0,57405739 |
| 643836_at | ZFP62 | 0,09292855 | 7,98451272 | 1,05397739 | 0,30911047 | 0,57405739 |
| 100422212_a | LOC10042221 | 0,09579729 | 3,38224865 | 1,05386147 | 0,30916178 | 0,57405739 |
| 10381_at | TUBB3 | -0,33534946 | 8,33503422 | -1,0538212 | 0,30917961 | 0,57405739 |
| 313_at | AOAH | -0,1004737 | 4,63080041 | -1,05372214 | 0,30922346 | 0,57405739 |
| 57699_at | CPNE5 | -0,13781223 | 4,58358258 | -1,05370194 | 0,3092324 | 0,57405739 |
| 441631_at | TSPAN11 | -0,09318284 | 5,43428288 | -1,05367266 | 0,30924537 | 0,57405739 |
| 6770_at | STAR | -0,10971947 | 4,32100353 | -1,05348618 | 0,30932794 | 0,57415534 |
| 3081_at | HGD | -0,07889552 | 3,5381718 | -1,05311345 | 0,30949303 | 0,57438209 |
| 9902_at | MRC2 | -0,12275849 | 6,13350968 | -1,05307574 | 0,30950974 | 0,57438209 |
| 4245_at | MGAT1 | 0,22336311 | 8,20786496 | 1,05293418 | 0,30957246 | 0,57444315 |
| 6293_at | VPS52 | 0,07756523 | 9,50103677 | 1,05263586 | 0,30970467 | 0,57458235 |
| 51741_at | WWOX | 0,07876112 | 7,14867002 | 1,05263029 | 0,30970714 | 0,57458235 |
| 7253_at | TSHR | -0,06266321 | 3,16873427 | -1,05210635 | 0,30993943 | 0,57495795 |
| 80279_at | CDK5RAP3 | 0,252034 | 8,48099056 | 1,05187583 | 0,31004167 | 0,57509223 |
| 3732_at | CD82 | 0,34884239 | 8,11538897 | 1,05163146 | 0,31015009 | 0,57520203 |
| 34_at | ACADM | 0,19376136 | 9,17829735 | 1,05155295 | 0,31018493 | 0,57520203 |
| 170692_at | ADAMTS18 | -0,09516017 | 3,75859298 | -1,05153537 | 0,31019273 | 0,57520203 |
| 4353_at | MPO | -0,07677993 | 4,48794204 | -1,0514732 | 0,31022031 | 0,57520203 |
| 5918_at | RARRES1 | -0,07384257 | 3,99412653 | -1,05130863 | 0,31029335 | 0,57528209 |
| 740_at | MRPL49 | -0,11486998 | 10,17686 | -1,05107857 | 0,31039548 | 0,57541604 |
| 27306_at | HPGDS | -0,1133696 | 3,46131871 | -1,05063894 | 0,3105907 | 0,57572254 |
| 9064_at | MAP3K6 | -0,0844164 | 6,02672004 | -1,05044869 | 0,31067521 | 0,57575859 |
| 79020_at | C7orf25 | 0,09027517 | 6,11496737 | 1,05040658 | 0,31069392 | 0,57575859 |

| | | | | | | |
|---|---|---|---|---|---|---|
| 8578_at | SCARF1 | 0,09896404 | 4,84114621 | 1,0503933 | 0,31069982 | 0,57575859 |
| 9536_at | PTGES | -0,22775515 | 10,6477067 | -1,05007365 | 0,31084186 | 0,5759664 |
| 339539_at | LOC339539 | -0,10475808 | 4,38243845 | -1,04998612 | 0,31088076 | 0,57598308 |
| 2623_at | GATA1 | -0,11609843 | 6,1615813 | -1,04950896 | 0,31109289 | 0,57629853 |
| 257106_at | ARHGAP30 | -0,08038098 | 4,9318523 | -1,04941236 | 0,31113585 | 0,57629853 |
| 92241_at | RCSD1 | -0,08355483 | 4,00405997 | -1,04940129 | 0,31114077 | 0,57629853 |
| 9502_at | XAGE2 | -0,10368855 | 3,74348427 | -1,04918772 | 0,31123577 | 0,57641905 |
| 84901_at | NFATC2IP | -0,12605886 | 8,2112508 | -1,04911778 | 0,31126688 | 0,57642125 |
| 3757_at | KCNH2 | -0,10810219 | 5,50014779 | -1,04887216 | 0,31137616 | 0,57651862 |
| 79412_at | KREMEN2 | -0,19238516 | 6,75878116 | -1,04886507 | 0,31137931 | 0,57651862 |
| 56063_at | TMEM234 | 0,06756396 | 6,3305232 | 1,0486884 | 0,31145794 | 0,57660877 |
| 114548_at | NLRP3 | -0,09524088 | 3,88573194 | -1,04851382 | 0,31153565 | 0,57669721 |
| 139065_at | SLITRK4 | -0,27338991 | 4,40170261 | -1,04844551 | 0,31156605 | 0,57669808 |
| 8733_at | GPAA1 | -0,20895011 | 9,80129751 | -1,04820266 | 0,31167418 | 0,57684279 |
| 6446_at | SGK1 | 0,3460264 | 7,81986653 | 1,04792302 | 0,31179872 | 0,57701784 |
| 404217_at | CTXN1 | -0,17440942 | 6,00325804 | -1,04783868 | 0,31183628 | 0,57702922 |
| 25803_at | SPDEF | -0,08125844 | 6,49615675 | -1,04777472 | 0,31186478 | 0,57702922 |
| 79666_at | PLEKHF2 | 0,41972603 | 8,62320687 | 1,04759251 | 0,31194596 | 0,57703624 |
| 7040_at | TGFB1 | -0,15993698 | 7,23492605 | -1,04758199 | 0,31195065 | 0,57703624 |
| 79982_at | DNAJB14 | 0,37316774 | 7,90674932 | 1,0475645 | 0,31195844 | 0,57703624 |
| 10297_at | APC2 | -0,1084013 | 5,64963367 | -1,04741013 | 0,31202723 | 0,57710807 |
| 286336_at | FAM78A | 0,39342702 | 5,81077089 | 1,04721414 | 0,31211458 | 0,57720812 |
| 375616_at | KCP | -0,10792286 | 5,45668687 | -1,0471543 | 0,31214126 | 0,57720812 |
| 155066_at | ATP6V0E2 | -0,11142264 | 7,58941949 | -1,04670501 | 0,31234159 | 0,57743145 |
| 3603_at | IL16 | -0,09419612 | 4,58261044 | -1,04669191 | 0,31234743 | 0,57743145 |
| 57091_at | CASS4 | -0,09017885 | 3,98382517 | -1,04665548 | 0,31236368 | 0,57743145 |
| 27032_at | ATP2C1 | -0,23318018 | 7,66606515 | -1,04652878 | 0,31242019 | 0,57743145 |
| 84437_at | MSANTD4 | 0,15096862 | 6,91933942 | 1,04652848 | 0,31242033 | 0,57743145 |
| 4045_at | LSAMP | -0,10071725 | 3,70000209 | -1,04648015 | 0,31244188 | 0,57743145 |
| 83877_at | TM2D2 | 0,36335847 | 8,9573296 | 1,04631688 | 0,31251473 | 0,57751067 |
| 1496_at | CTNNA2 | -0,06433367 | 3,677479 | -1,04609588 | 0,31261334 | 0,57763748 |
| 10490_at | VTI1B | 0,06009674 | 9,61710445 | 1,04590311 | 0,31269937 | 0,57774104 |
| 219681_at | ARMC3 | -0,08982482 | 3,22234266 | -1,04567225 | 0,31280243 | 0,57787602 |
| 51110_at | LACTB2 | -0,11595522 | 8,07977723 | -1,04557078 | 0,31284774 | 0,5779043 |
| 115761_at | ARL11 | -0,06317197 | 3,17158582 | -1,04543113 | 0,3129101 | 0,57796407 |
| 286076_at | BREA2 | -0,1161547 | 4,64455256 | -1,04529968 | 0,31296881 | 0,57801708 |
| 55754_at | TMEM30A | 0,2112504 | 9,30715746 | 1,04496296 | 0,31311922 | 0,57823944 |
| 100507254_a | LINC01013 | -0,0943761 | 3,57939181 | -1,044799 | 0,31319249 | 0,57829642 |
| 5332_at | PLCB4 | 0,45454907 | 3,85333284 | 1,04475954 | 0,31321012 | 0,57829642 |
| 55362_at | TMEM63B | 0,10369508 | 8,17446831 | 1,0445218 | 0,31331638 | 0,57843717 |
| 3622_at | ING2 | -0,3124268 | 7,4306696 | -1,04437922 | 0,31338011 | 0,57846255 |
| 83942_at | TSSK1B | -0,08515924 | 4,17860453 | -1,04428253 | 0,31342334 | 0,57846255 |
| 100316868_a | HOTTIP | -0,07955236 | 4,4098955 | -1,04427846 | 0,31342516 | 0,57846255 |
| 3987_at | LIMS1 | 0,24537315 | 10,2871738 | 1,04415756 | 0,31347922 | 0,57846255 |
| 6277_at | S100A6 | 0,16772263 | 11,1984787 | 1,0441552 | 0,31348028 | 0,57846255 |
| 3042_at | HBM | -0,12281827 | 4,80245179 | -1,04406923 | 0,31351872 | 0,57847406 |
| 285708_at | LINC00491 | 0,1216823 | 5,21086511 | 1,04394812 | 0,31357289 | 0,57847406 |
| 22916_at | NCBP2 | -0,088892 | 9,01715617 | -1,04392563 | 0,31358295 | 0,57847406 |
| 5030_at | P2RY4 | -0,10105373 | 4,29994084 | -1,04387267 | 0,31360663 | 0,57847406 |
| 79188_at | TMEM43 | -0,13993431 | 9,47761558 | -1,04373668 | 0,31366747 | 0,57848681 |

| | | | | | | |
|---|---|---|---|---|---|---|
| 80307_at | FER1L4 | 0,1707851 | 6,30899047 | 1,04364048 | 0,31371051 | 0,57848681 |
| 146712_at | B3GNTL1 | -0,10923784 | 6,07812912 | -1,04343824 | 0,313801 | 0,57848681 |
| 101927494_a | LOC10192749 | 0,10739192 | 3,11468405 | 1,04342925 | 0,31380502 | 0,57848681 |
| 7737_at | RNF113A | 0,08298387 | 7,40966582 | 1,04333036 | 0,31384928 | 0,57848681 |
| 64757_at | MARC1 | -0,22351534 | 6,69953113 | -1,04331426 | 0,31385649 | 0,57848681 |
| 100505515_a | LOC10050551 | -0,12676964 | 4,40076053 | -1,04316056 | 0,31392528 | 0,57848681 |
| 100506462_a | SRD5A3-AS1 | 0,12352232 | 3,15818829 | 1,04310037 | 0,31395222 | 0,57848681 |
| 170622_at | COMMD6 | 0,0988823 | 7,56162709 | 1,04308863 | 0,31395748 | 0,57848681 |
| 10586_at | MAB21L2 | -0,10284921 | 3,5194399 | -1,043063 | 0,31396895 | 0,57848681 |
| 154865_at | IQUB | 0,13179933 | 3,22632777 | 1,04303515 | 0,31398142 | 0,57848681 |
| 2847_at | MCHR1 | -0,07947415 | 4,96693866 | -1,04295012 | 0,31401949 | 0,57848681 |
| 146894_at | CD300LG | -0,11980876 | 4,64914643 | -1,04294351 | 0,31402245 | 0,57848681 |
| 2994_at | GYPB | -0,16724572 | 3,27689379 | -1,04282815 | 0,3140741 | 0,57848681 |
| 54827_at | NXPE4 | -0,11879409 | 4,00481721 | -1,0428225 | 0,31407663 | 0,57848681 |
| 6401_at | SELE | 0,11398909 | 3,48124191 | 1,04278362 | 0,31409404 | 0,57848681 |
| 55623_at | THUMPD1 | 0,09443637 | 8,59150552 | 1,0426813 | 0,31413987 | 0,57851589 |
| 2242_at | FES | -0,12112183 | 4,70325232 | -1,04257887 | 0,31418574 | 0,57854507 |
| 81872_at | KRTAP2-1 | -0,0877807 | 3,54477039 | -1,04245689 | 0,31424038 | 0,57859037 |
| 11254_at | SLC6A14 | -0,41295431 | 3,56157217 | -1,04225902 | 0,31432903 | 0,57869827 |
| 751580_at | LINC00106 | -0,12382054 | 5,43469543 | -1,04216419 | 0,31437152 | 0,57871893 |
| 145837_at | DRAIC | -0,08604445 | 3,87858973 | -1,04209988 | 0,31440033 | 0,57871893 |
| 284023_at | LOC284023 | -0,0868384 | 5,81032465 | -1,04177009 | 0,31454815 | 0,57890231 |
| 25776_at | CBY1 | 0,14612209 | 7,89503246 | 1,04157841 | 0,31463408 | 0,57890231 |
| 6787_at | NEK4 | -0,23226256 | 8,1278459 | -1,04152909 | 0,3146562 | 0,57890231 |
| 2703_at | GJA8 | -0,09821235 | 4,38144759 | -1,0415154 | 0,31466233 | 0,57890231 |
| 57101_at | ANO2 | 0,07536678 | 3,69057824 | 1,04147781 | 0,31467919 | 0,57890231 |
| 608_at | TNFRSF17 | 0,10358799 | 3,42563952 | 1,04141666 | 0,31470661 | 0,57890231 |
| 6702_at | SPRR2C | -0,10812882 | 3,95046371 | -1,04140826 | 0,31471038 | 0,57890231 |
| 131965_at | METTL6 | -0,11503756 | 5,3567332 | -1,04134136 | 0,31474038 | 0,57890231 |
| 729_at | C6 | -0,12241582 | 3,68548222 | -1,04105078 | 0,31487072 | 0,57908675 |
| 56098_at | PCDHGC4 | -0,09251381 | 3,97669854 | -1,04083892 | 0,31496577 | 0,57920626 |
| 3488_at | IGFBP5 | 0,18213932 | 4,74198795 | 1,04068338 | 0,31503557 | 0,57923241 |
| 155185_at | AMZ1 | -0,10149663 | 6,06088303 | -1,04067322 | 0,31504013 | 0,57923241 |
| 319085_at | ITPK1-AS1 | 0,10006153 | 3,8761053 | 1,04035064 | 0,31518492 | 0,57943615 |
| 1618_at | DAZL | 0,06077428 | 3,22915458 | 1,04029232 | 0,3152111 | 0,57943615 |
| 5873_at | RAB27A | -0,12165757 | 6,77824195 | -1,04014837 | 0,31527574 | 0,57946219 |
| 284661_at | LOC284661 | -0,0604708 | 3,33036992 | -1,04012679 | 0,31528543 | 0,57946219 |
| 266675_at | BEST4 | 0,07258342 | 4,06729532 | 1,03985843 | 0,31540595 | 0,57962839 |
| 94097_at | SFXN5 | -0,07559774 | 6,489224 | -1,03967461 | 0,31548852 | 0,57972483 |
| 79149_at | ZSCAN5A | -0,16840995 | 6,55346233 | -1,03959932 | 0,31552235 | 0,57973168 |
| 3737_at | KCNA2 | -0,09385813 | 4,23618384 | -1,03947189 | 0,3155796 | 0,57973246 |
| 29072_at | SETD2 | 0,10164754 | 7,62827089 | 1,03942244 | 0,31560183 | 0,57973246 |
| 645784_at | ANKRD36BP2 | -0,06653276 | 2,69769816 | -1,03939744 | 0,31561306 | 0,57973246 |
| 54798_at | DCHS2 | -0,06628215 | 3,41231263 | -1,03931801 | 0,31564875 | 0,57974275 |
| 57531_at | HACE1 | 0,13893464 | 6,93336709 | 1,03921831 | 0,31569356 | 0,57976977 |
| 7782_at | SLC30A4 | 0,06684851 | 3,95623194 | 1,03900178 | 0,31579089 | 0,57982641 |
| 165186_at | FAM179A | -0,10205629 | 4,63004548 | -1,03899946 | 0,31579194 | 0,57982641 |
| 85365_at | ALG2 | 0,07834891 | 9,09270253 | 1,03894882 | 0,31581471 | 0,57982641 |
| 5546_at | PRCC | -0,1275788 | 8,2939971 | -1,03850562 | 0,31601401 | 0,57995282 |
| 65072_at | CFLAR-AS1 | -0,08454955 | 3,40940699 | -1,03846684 | 0,31603145 | 0,57995282 |

| | | | | | | |
|---|---|---|---|---|---|---|
| 9058_at | SLC13A2 | -0,07899265 | 4,77100104 | -1,03845895 | 0,316035 | 0,57995282 |
| 8361_at | HIST1H4F | -0,13887181 | 4,25287091 | -1,03843546 | 0,31604557 | 0,57995282 |
| 143098_at | MPP7 | 0,48629805 | 6,10590288 | 1,03836731 | 0,31607622 | 0,57995282 |
| 9440_at | MED17 | 0,07555756 | 7,09086536 | 1,03822339 | 0,31614097 | 0,57995282 |
| 10371_at | SEMA3A | 0,26896215 | 5,6462974 | 1,03816454 | 0,31616745 | 0,57995282 |
| 321_at | APBA2 | -0,08294119 | 5,12117751 | -1,0380482 | 0,3162198 | 0,57995282 |
| 11045_at | UPK1A | 1,26640624 | 7,30488525 | 1,03804732 | 0,3162202 | 0,57995282 |
| 23024_at | PDZRN3 | -0,07374156 | 3,27659399 | -1,0379865 | 0,31624757 | 0,57995282 |
| 10871_at | CD300C | -0,09568411 | 5,6373741 | -1,03788295 | 0,31629417 | 0,57995282 |
| 57035_at | RSRP1 | 0,30595717 | 8,35121713 | 1,0378341 | 0,31631616 | 0,57995282 |
| 23627_at | PRND | -0,08871271 | 4,51624937 | -1,03758115 | 0,31643003 | 0,57995282 |
| 64711_at | HS3ST6 | -0,08080294 | 5,4119195 | -1,03756491 | 0,31643734 | 0,57995282 |
| 100505478_a | LOC10050547 | 0,09443141 | 5,10425469 | 1,03752622 | 0,31645476 | 0,57995282 |
| 1213_at | CLTC | -0,07593172 | 11,7345592 | -1,03748103 | 0,31647511 | 0,57995282 |
| 58499_at | ZNF462 | 0,40762107 | 6,79168816 | 1,03747559 | 0,31647755 | 0,57995282 |
| 55552_at | ZNF823 | 0,08792136 | 7,56562338 | 1,03744907 | 0,3164895 | 0,57995282 |
| 6156_at | RPL30 | -0,06482757 | 12,6079168 | -1,03744313 | 0,31649217 | 0,57995282 |
| 285500_at | LOC285500 | -0,06297607 | 3,13022268 | -1,03739569 | 0,31651354 | 0,57995282 |
| 6357_at | CCL13 | -0,09737346 | 4,87963025 | -1,03734629 | 0,31653578 | 0,57995282 |
| 390502_at | SERPINA2 | -0,10522504 | 3,55335743 | -1,03729439 | 0,31655915 | 0,57995282 |
| 5069_at | PAPPA | -0,05912357 | 3,73190568 | -1,03725693 | 0,31657602 | 0,57995282 |
| 51275_at | MAPKAPK5-AS | -0,057908 | 6,94534071 | -1,03704901 | 0,31666968 | 0,58006922 |
| 1908_at | EDN3 | -0,0975842 | 4,97816388 | -1,03696995 | 0,31670529 | 0,58007931 |
| 83655_at | LINC00208 | -0,08017864 | 3,79823997 | -1,03681646 | 0,31677444 | 0,58014927 |
| 81792_at | ADAMTS12 | -0,0940724 | 4,9164575 | -1,03675147 | 0,31680373 | 0,58014927 |
| 1995_at | ELAVL3 | -0,08295404 | 4,65627047 | -1,03647246 | 0,31692947 | 0,58032437 |
| 27074_at | LAMP3 | 0,79060482 | 7,25838565 | 1,03637518 | 0,31697332 | 0,58032631 |
| 23394_at | ADNP | 0,10054185 | 8,69809918 | 1,03633645 | 0,31699078 | 0,58032631 |
| 115752_at | DIS3L | -0,09134203 | 8,30264071 | -1,03619047 | 0,31705659 | 0,58037752 |
| 153830_at | RNF145 | -0,25779445 | 7,68033843 | -1,0361087 | 0,31709346 | 0,58037752 |
| 4647_at | MYO7A | -0,05800687 | 5,22701454 | -1,03607393 | 0,31710914 | 0,58037752 |
| 7136_at | TNNI2 | 0,16197651 | 6,32276432 | 1,03592581 | 0,31717594 | 0,58041749 |
| 81569_at | ACTL8 | -0,11964248 | 5,95068483 | -1,03582569 | 0,31722109 | 0,58041749 |
| 353500_at | BMP8A | -0,11542097 | 4,92399311 | -1,03581005 | 0,31722815 | 0,58041749 |
| 79924_at | ADM2 | -0,13323137 | 6,56139475 | -1,03575827 | 0,3172515 | 0,58041749 |
| 112464_at | PRKCDBP | 0,21604841 | 4,99797864 | 1,03535451 | 0,31743366 | 0,5806956 |
| 100505683_a | LINC01431 | 0,12516428 | 4,78719539 | 1,03486901 | 0,31765281 | 0,58104131 |
| 4179_at | CD46 | -0,15265399 | 9,92401628 | -1,03447772 | 0,3178295 | 0,58130932 |
| 100131017_a | ZNF316 | -0,08647485 | 6,97098245 | -1,0340744 | 0,31801171 | 0,58153024 |
| 25957_at | PNISR | 0,20471711 | 7,77063287 | 1,03395123 | 0,31806736 | 0,58153024 |
| 101927075_a | FGF10-AS1 | -0,13297028 | 3,57970517 | -1,03391691 | 0,31808287 | 0,58153024 |
| 51054_at | PLEKHA8P1 | 0,10410998 | 6,46166665 | 1,03391243 | 0,3180849 | 0,58153024 |
| 54847_at | SIDT1 | 0,59000451 | 6,92113902 | 1,03387629 | 0,31810123 | 0,58153024 |
| 143282_at | FGFBP3 | -0,15885329 | 5,55961537 | -1,03379138 | 0,31813961 | 0,5815452 |
| 284185_at | LINC00482 | -0,09165267 | 6,48102403 | -1,03342403 | 0,31830568 | 0,58179356 |
| 28513_at | CDH19 | -0,07180666 | 3,12696808 | -1,03321816 | 0,31839877 | 0,5819085 |
| 440829_at | SHISA8 | -0,09205387 | 5,0178915 | -1,03314411 | 0,31843226 | 0,5819145 |
| 8823_at | FGF16 | -0,11410419 | 4,69921453 | -1,03295539 | 0,31851763 | 0,58201528 |
| 774_at | CACNA1B | -0,08216932 | 4,46358443 | -1,03267086 | 0,31864636 | 0,58214771 |
| 64897_at | C12orf43 | -0,17601247 | 7,85599253 | -1,03266161 | 0,31865055 | 0,58214771 |

| | | | | | | |
|---|---|---|---|---|---|---|
| 340075_at | ARSI | -0,09412422 | 5,2382934 | -1,03256674 | 0,31869348 | 0,58217093 |
| 23541_at | SEC14L2 | -0,16827601 | 5,75998184 | -1,03234198 | 0,31879521 | 0,58230154 |
| 1393_at | CRHBP | -0,11091598 | 3,60919648 | -1,03175403 | 0,31906143 | 0,58268441 |
| 84851_at | TRIM52 | 0,22579664 | 5,99664443 | 1,03174545 | 0,31906531 | 0,58268441 |
| 1797_at | DXO | -0,12240867 | 7,36778988 | -1,03150732 | 0,31917319 | 0,58282615 |
| 83594_at | NUDT12 | 0,23507248 | 6,7732106 | 1,03140037 | 0,31922164 | 0,58285939 |
| 100302650_a | BRE-AS1 | -0,12191353 | 4,386389 | -1,03103127 | 0,31938891 | 0,58308582 |
| 3957_at | LGALS2 | -0,09324893 | 4,19377196 | -1,03099314 | 0,3194062 | 0,58308582 |
| 390205_at | LRRC10B | -0,20728546 | 5,79785344 | -1,0308302 | 0,31948006 | 0,5831654 |
| 4987_at | OPRL1 | -0,10313143 | 5,36763178 | -1,0306985 | 0,31953977 | 0,58321912 |
| 3558_at | IL2 | 0,10868868 | 3,26426284 | 1,03055301 | 0,31960575 | 0,58328427 |
| 117177_at | RAB3IP | 0,06719038 | 6,16743612 | 1,03022258 | 0,31975562 | 0,58350251 |
| 28714_at | TRAJ41 | 0,12181537 | 4,13262119 | 1,03007903 | 0,31982075 | 0,58353582 |
| 7697_at | ZNF138 | 0,13327253 | 5,42008533 | 1,03004067 | 0,31983815 | 0,58353582 |
| 92335_at | STRADA | 0,07272464 | 7,71782543 | 1,02998204 | 0,31986475 | 0,58353582 |
| 55105_at | GPATCH2 | 0,094138 | 6,39207174 | 1,02980922 | 0,31994318 | 0,58362362 |
| 63931_at | MRPS14 | -0,08228763 | 7,80271803 | -1,0297371 | 0,31997591 | 0,58362806 |
| 10345_at | TRDN | -0,05378114 | 2,81590521 | -1,02942111 | 0,32011935 | 0,58383442 |
| 4908_at | NTF3 | -0,07511018 | 4,55492219 | -1,02931506 | 0,3201675 | 0,58385822 |
| 266917_at | D21S2088E | -0,08150746 | 3,42389921 | -1,02921369 | 0,32021354 | 0,58385822 |
| 84168_at | ANTXR1 | -0,0600645 | 7,25384319 | -1,02919212 | 0,32022333 | 0,58385822 |
| 442186_at | OR2J3 | 0,08215824 | 3,90785731 | 1,02903349 | 0,32029538 | 0,58388899 |
| 9457_at | FHL5 | -0,06449734 | 2,95317272 | -1,02902148 | 0,32030083 | 0,58388899 |
| 23194_at | FBXL7 | -0,09931014 | 4,71212074 | -1,02876637 | 0,32041672 | 0,58404497 |
| 9867_at | PJA2 | 0,15285103 | 8,93104554 | 1,02851222 | 0,3205322 | 0,58413226 |
| 653121_at | ZBTB8A | 0,19318296 | 7,77197248 | 1,02850793 | 0,32053415 | 0,58413226 |
| 51412_at | ACTL6B | -0,10008526 | 4,70961484 | -1,02843921 | 0,32056538 | 0,58413226 |
| 403314_at | APOBEC4 | -0,1170921 | 3,48465518 | -1,02833657 | 0,32061203 | 0,58413226 |
| 2729_at | GCLC | 0,18705793 | 9,49186744 | 1,02832733 | 0,32061623 | 0,58413226 |
| 80011_at | FAM192A | -0,12992706 | 8,20493927 | -1,02821433 | 0,32066759 | 0,58413432 |
| 2968_at | GTF2H4 | -0,10758636 | 7,43010266 | -1,02818656 | 0,32068022 | 0,58413432 |
| 4188_at | MDFI | -0,28637613 | 7,76077611 | -1,02805607 | 0,32073954 | 0,58413432 |
| 9295_at | SRSF11 | 0,15746793 | 7,59435222 | 1,02800567 | 0,32076246 | 0,58413432 |
| 85417_at | CCNB3 | -0,08041884 | 4,63159367 | -1,02799133 | 0,32076898 | 0,58413432 |
| 164284_at | APCDD1L | -0,10043296 | 5,24860684 | -1,02777045 | 0,32086942 | 0,584262 |
| 1750_at | DLX6 | 0,20081893 | 6,23047034 | 1,02756326 | 0,32096366 | 0,58437835 |
| 221938_at | MMD2 | -0,0910177 | 5,07829396 | -1,0272739 | 0,3210953 | 0,58456278 |
| 80227_at | PAAF1 | -0,07802709 | 7,59839409 | -1,02709443 | 0,32117697 | 0,58464444 |
| 5992_at | RFX4 | -0,11995364 | 3,33368463 | -1,02702038 | 0,32121067 | 0,58464444 |
| 6314_at | ATXN7 | 0,15385821 | 7,09836631 | 1,02692229 | 0,32125532 | 0,58464444 |
| 51764_at | GNG13 | -0,17590851 | 4,39216512 | -1,02690859 | 0,32126156 | 0,58464444 |
| 7273_at | TTN | -0,07387923 | 3,34144433 | -1,02681396 | 0,32130463 | 0,5846614 |
| 5530_at | PPP3CA | 0,15555635 | 9,40444747 | 1,02675476 | 0,32133158 | 0,5846614 |
| 285627_at | LOC285627 | -0,12531329 | 3,94122395 | -1,02663563 | 0,32138582 | 0,58470487 |
| 3812_at | KIR3DL2 | -0,11389586 | 5,04222585 | -1,02631706 | 0,3215309 | 0,58490598 |
| 8558_at | CDK10 | 0,06966197 | 7,74740219 | 1,02625954 | 0,3215571 | 0,58490598 |
| 10078_at | TSSC4 | -0,09022453 | 6,76489765 | -1,02618225 | 0,3215923 | 0,58491479 |
| 6772_at | STAT1 | 0,29241739 | 9,09720569 | 1,02608591 | 0,32163619 | 0,58493785 |
| 7478_at | WNT8A | -0,08514384 | 4,04335009 | -1,0260211 | 0,32166571 | 0,58493785 |
| 54904_at | WHSC1L1 | 0,22338481 | 7,47566593 | 1,02583124 | 0,32175223 | 0,58503994 |

| | | | | | | |
|---|---|---|---|---|---|---|
| 22818_at | COPZ1 | 0,0911977 | 9,99287686 | 1,02555347 | 0,32187882 | 0,58521489 |
| 286006_at | LSMEM1 | -0,10263038 | 4,43767916 | -1,02539338 | 0,3219518 | 0,58529233 |
| 441951_at | ZFAS1 | 0,11568003 | 10,6706454 | 1,02527938 | 0,32200378 | 0,58533158 |
| 101928661_a | LOC10192866 | -0,1293669 | 4,59342595 | -1,02517821 | 0,32204991 | 0,58536019 |
| 113179_at | ADAT3 | -0,11405393 | 6,98246926 | -1,02496943 | 0,32214512 | 0,58545537 |
| 54872_at | PIGG | 0,06234453 | 6,90328391 | 1,02493011 | 0,32216305 | 0,58545537 |
| 22911_at | WDR47 | -0,08708986 | 7,99063661 | -1,02461013 | 0,32230902 | 0,58566538 |
| 1307_at | COL16A1 | 0,18364967 | 5,53337389 | 1,02450892 | 0,32235521 | 0,58569405 |
| 6861_at | SYT5 | -0,08263775 | 4,84620189 | -1,02441107 | 0,32239985 | 0,58571993 |
| 5563_at | PRKAA2 | -0,10962415 | 4,45657525 | -1,02425771 | 0,32246985 | 0,58575424 |
| 8636_at | SSNA1 | -0,16248846 | 9,19285478 | -1,02423644 | 0,32247956 | 0,58575424 |
| 219621_at | C10orf107 | 0,05992646 | 2,94590393 | 1,02377241 | 0,32269141 | 0,58608378 |
| 168537_at | GIMAP7 | 0,07309057 | 3,01984657 | 1,02353715 | 0,32279885 | 0,58622365 |
| 392617_at | ELFN1 | -0,11105115 | 5,63200967 | -1,0230603 | 0,32301671 | 0,58656399 |
| 100506776_a | TRG-AS1 | -0,09123084 | 4,28143019 | -1,022827 | 0,32312334 | 0,58670231 |
| 55421_at | NCBP3 | -0,05385847 | 6,70287562 | -1,02250122 | 0,32327227 | 0,58686618 |
| 3010_at | HIST1H1T | -0,1215807 | 4,48061317 | -1,02240804 | 0,32331488 | 0,58686618 |
| 8621_at | CDK13 | 0,07470077 | 7,9753373 | 1,02239918 | 0,32331893 | 0,58686618 |
| 8851_at | CDK5R1 | -0,12190659 | 5,70543505 | -1,02236306 | 0,32333545 | 0,58686618 |
| 56935_at | SMCO4 | -0,17171519 | 7,04705163 | -1,02214181 | 0,32343664 | 0,58693556 |
| 81539_at | SLC38A1 | -0,15744477 | 10,1749441 | -1,02213428 | 0,32344008 | 0,58693556 |
| 493812_at | HCG11 | 0,33374807 | 3,83302589 | 1,02207963 | 0,32346508 | 0,58693556 |
| 22864_at | R3HDM2 | 0,16649053 | 7,91994769 | 1,02176107 | 0,32361083 | 0,5871131 |
| 7580_at | ZNF32 | 0,21527394 | 7,12168119 | 1,02173253 | 0,32362389 | 0,5871131 |
| 6767_at | ST13 | 0,06131777 | 10,9544646 | 1,02154092 | 0,32371158 | 0,5871668 |
| 2013_at | EMP2 | 0,11667022 | 9,86080252 | 1,02153465 | 0,32371445 | 0,5871668 |
| 122509_at | IFI27L1 | -0,14575301 | 6,6559669 | -1,02133081 | 0,32380775 | 0,58728074 |
| 10272_at | FSTL3 | 0,16095785 | 7,94233215 | 1,02103508 | 0,32394316 | 0,58742346 |
| 101927636_a | LOC10192763 | -0,08257812 | 3,01332895 | -1,02102574 | 0,32394743 | 0,58742346 |
| 10230_at | NBR2 | 0,14863082 | 6,79998071 | 1,02094692 | 0,32398353 | 0,58743361 |
| 11138_at | TBC1D8 | 0,21735754 | 7,73175426 | 1,02085323 | 0,32402644 | 0,58745285 |
| 15_at | AANAT | -0,13249447 | 4,72860353 | -1,02079059 | 0,32405513 | 0,58745285 |
| 402779_at | CSMD2-AS1 | -0,14825859 | 4,62270281 | -1,02067655 | 0,32410737 | 0,58749226 |
| 54819_at | ZCCHC10 | 0,06909295 | 6,82021968 | 1,02040904 | 0,32422993 | 0,5876493 |
| 7412_at | VCAM1 | -0,07506624 | 3,72287161 | -1,02035429 | 0,32425502 | 0,5876493 |
| 57636_at | ARHGAP23 | 0,07417057 | 7,05031033 | 1,01950989 | 0,32464212 | 0,58812611 |
| 4771_at | NF2 | -0,1102775 | 6,86187319 | -1,01945976 | 0,32466511 | 0,58812611 |
| 9853_at | RUSC2 | -0,08858674 | 6,35636367 | -1,01943774 | 0,32467521 | 0,58812611 |
| 26538_at | OR10H2 | -0,11364748 | 3,36500392 | -1,01940736 | 0,32468915 | 0,58812611 |
| 55766_at | H2AFJ | 0,08873781 | 6,95909405 | 1,01935445 | 0,32471342 | 0,58812611 |
| 641467_at | TSC22D1-AS1 | 0,09504466 | 3,36248998 | 1,01927141 | 0,32475151 | 0,58812611 |
| 6900_at | CNTN2 | -0,10905179 | 4,56100352 | -1,01925873 | 0,32475733 | 0,58812611 |
| 178_at | AGL | 0,1174304 | 8,92627884 | 1,01924776 | 0,32476236 | 0,58812611 |
| 131096_at | KCNH8 | 0,07393451 | 3,05974905 | 1,01908129 | 0,32483874 | 0,5881589 |
| 7069_at | THRSP | -0,1034933 | 4,21966027 | -1,01907519 | 0,32484154 | 0,5881589 |
| 100505633_a | LINC01133 | 0,32304624 | 3,59262158 | 1,01887452 | 0,32493362 | 0,58827034 |
| 147111_at | NOTUM | -0,08919506 | 5,97446243 | -1,01877555 | 0,32497905 | 0,58829728 |
| 10140_at | TOB1 | 0,15936347 | 10,3134443 | 1,0186487 | 0,32503727 | 0,58834739 |
| 728723_at | ZBED3-AS1 | 0,10351497 | 4,33086877 | 1,01834953 | 0,32517462 | 0,58854071 |
| 124512_at | METTL23 | -0,0767656 | 9,01893068 | -1,018235 | 0,32522722 | 0,58854519 |

| | | | | | | |
|---|---|---|---|---|---|---|
| 9904_at | RBM19 | -0,07423078 | 6,90614975 | -1,01821108 | 0,3252382 | 0,58854519 |
| 101929076_a | LOC10192907 | -0,11505488 | 4,2314019 | -1,01808752 | 0,32529495 | 0,58859258 |
| 4093_at | SMAD9 | 0,11304841 | 4,90429315 | 1,01775718 | 0,3254467 | 0,58872741 |
| 100506371_a | LOC10050637 | -0,11626764 | 4,19033026 | -1,01775265 | 0,32544878 | 0,58872741 |
| 55203_at | LGI2 | -0,15105316 | 5,25992544 | -1,01772572 | 0,32546115 | 0,58872741 |
| 6494_at | SIPA1 | 0,49586326 | 6,87436726 | 1,01733273 | 0,32564176 | 0,5889988 |
| 85291_at | KRTAP4-2 | -0,10374347 | 4,94158295 | -1,01719725 | 0,32570403 | 0,5890082 |
| 26053_at | AUTS2 | -0,10356037 | 5,85565716 | -1,01714557 | 0,3257278 | 0,5890082 |
| 57818_at | G6PC2 | -0,12131812 | 4,34291444 | -1,01708199 | 0,32575703 | 0,5890082 |
| 51463_at | GPR89B | -0,16426059 | 8,78266571 | -1,01705537 | 0,32576926 | 0,5890082 |
| 51535_at | PPHLN1 | -0,07372865 | 7,14908907 | -1,0167902 | 0,3258912 | 0,58917337 |
| 3641_at | INSL4 | -0,10347057 | 3,70915049 | -1,01655637 | 0,32599875 | 0,5893125 |
| 389763_at | SPATA31D1 | -0,12413237 | 5,20460382 | -1,01633767 | 0,32609937 | 0,58943907 |
| 79192_at | IRX1 | -0,08953252 | 4,41847625 | -1,01620731 | 0,32615936 | 0,58944513 |
| 8864_at | PER2 | 0,1250298 | 7,35179504 | 1,0161233 | 0,32619802 | 0,58944513 |
| 4004_at | LMO1 | -0,08871668 | 4,49935259 | -1,0160929 | 0,32621201 | 0,58944513 |
| 28316_at | CDH20 | -0,08658923 | 3,67933298 | -1,01606441 | 0,32622512 | 0,58944513 |
| 7813_at | EVI5 | 0,11016781 | 6,12068922 | 1,01567681 | 0,32640355 | 0,58966742 |
| 54480_at | CHPF2 | 0,14496751 | 8,08169767 | 1,01566417 | 0,32640937 | 0,58966742 |
| 5157_at | PDGFRL | 0,31751745 | 5,56677107 | 1,01545351 | 0,32650637 | 0,58978735 |
| 7010_at | TEK | -0,10066886 | 3,65462017 | -1,01523497 | 0,32660703 | 0,58991385 |
| 54995_at | OXSM | -0,12415268 | 7,21874389 | -1,01511414 | 0,32666269 | 0,58995907 |
| 53371_at | NUP54 | -0,25272867 | 7,82486732 | -1,01476857 | 0,32682192 | 0,5901913 |
| 342_at | APOC1P1 | -0,15610722 | 5,62538481 | -1,01468314 | 0,32686129 | 0,59020707 |
| 63929_at | XPNPEP3 | -0,14502393 | 6,72635892 | -1,01425287 | 0,32705964 | 0,59044481 |
| 84542_at | KIAA1841 | 0,19977237 | 6,94288332 | 1,01423282 | 0,32706888 | 0,59044481 |
| 55898_at | UNC45A | 0,14983124 | 8,13460728 | 1,01419807 | 0,32708491 | 0,59044481 |
| 113510_at | HELQ | 0,17371122 | 4,93220136 | 1,01412485 | 0,32711867 | 0,59045043 |
| 55449_at | DHRS4-AS1 | 0,19401575 | 7,73013529 | 1,01392349 | 0,32721154 | 0,59052547 |
| 123872_at | DNAAF1 | -0,11377753 | 4,57158901 | -1,01384086 | 0,32724965 | 0,59052547 |
| 8936_at | WASF1 | 0,17950016 | 8,44975253 | 1,01380333 | 0,32726697 | 0,59052547 |
| 65989_at | DLK2 | 0,21037329 | 7,83612546 | 1,01364514 | 0,32733994 | 0,59052547 |
| 58509_at | CACTIN | -0,11247486 | 7,01412876 | -1,0136012 | 0,32736022 | 0,59052547 |
| 359_at | AQP2 | -0,07766842 | 5,26517288 | -1,01359586 | 0,32736268 | 0,59052547 |
| 9943_at | OXSR1 | -0,19270378 | 9,30947276 | -1,01352995 | 0,3273931 | 0,59052547 |
| 5925_at | RB1 | -0,65823531 | 5,90373539 | -1,01350308 | 0,32740549 | 0,59052547 |
| 5459_at | POU4F3 | -0,12750737 | 4,13691402 | -1,01342239 | 0,32744273 | 0,59053734 |
| 84631_at | SLITRK2 | -0,0906657 | 3,8233373 | -1,01331104 | 0,32749412 | 0,5905514 |
| 101060544_a | LINC01563 | -0,07173813 | 3,84273171 | -1,01327264 | 0,32751184 | 0,5905514 |
| 9806_at | SPOCK2 | -0,09475558 | 5,80525091 | -1,01293045 | 0,32766981 | 0,59076018 |
| 220070_at | SHANK2-AS3 | -0,08843805 | 5,19420229 | -1,01288896 | 0,32768896 | 0,59076018 |
| 9472_at | AKAP6 | -0,09136599 | 4,01242017 | -1,01273266 | 0,32776114 | 0,59078541 |
| 3880_at | KRT19 | 0,11687857 | 13,271586 | 1,01272581 | 0,3277643 | 0,59078541 |
| 101927478_a | LOC10192747 | -0,14062323 | 3,56953948 | -1,01249811 | 0,32786947 | 0,59091968 |
| 1280_at | COL2A1 | 0,07799539 | 4,57903982 | 1,01206032 | 0,32807173 | 0,59114087 |
| 79849_at | PDZD3 | -0,11289331 | 5,33864122 | -1,01205394 | 0,32807468 | 0,59114087 |
| 51727_at | CMPK1 | -0,067964 | 9,47567231 | -1,01203321 | 0,32808426 | 0,59114087 |
| 92421_at | CHMP4C | 0,11371882 | 9,07841428 | 1,01186555 | 0,32816175 | 0,59120809 |
| 83759_at | RBM4B | 0,19664898 | 8,04666829 | 1,01181968 | 0,32818295 | 0,59120809 |
| 643_at | CXCR5 | -0,09589642 | 4,66755934 | -1,01170546 | 0,32823575 | 0,59124675 |

| | | | | | | |
|---|---|---|---|---|---|---|
| 56243_at | KIAA1217 | 0,16602437 | 6,77900205 | 1,01164046 | 0,32826579 | 0,59124675 |
| 286234_at | SPATA31E1 | -0,07361823 | 4,43155779 | -1,01106974 | 0,32852972 | 0,59166578 |
| 151_at | ADRA2B | -0,11107631 | 5,68182652 | -1,01100455 | 0,32855988 | 0,59166578 |
| 100303728_a | SLC25A5-AS1 | -0,09611388 | 4,0958282 | -1,01047209 | 0,32880626 | 0,59205362 |
| 10609_at | P3H4 | 0,16604112 | 6,89266687 | 1,0104063 | 0,32883671 | 0,59205362 |
| 29958_at | DMGDH | -0,07997395 | 3,18215226 | -1,01014099 | 0,32895954 | 0,59206523 |
| 23556_at | PIGN | 0,1188654 | 7,47688776 | 1,01008581 | 0,32898509 | 0,59206523 |
| 285194_at | TUSC7 | -0,07100509 | 3,3084502 | -1,0100727 | 0,32899116 | 0,59206523 |
| 79057_at | PRRG3 | -0,1084804 | 4,50799723 | -1,01006927 | 0,32899275 | 0,59206523 |
| 101928850_a | LOC10192885 | -0,09831495 | 3,61734273 | -1,01003173 | 0,32901013 | 0,59206523 |
| 10382_at | TUBB4A | -0,17445494 | 5,07149803 | -1,00994564 | 0,32905 | 0,59206523 |
| 90196_at | SYS1 | 0,09042436 | 7,82367286 | 1,00992768 | 0,32905831 | 0,59206523 |
| 60312_at | AFAP1 | 0,30558886 | 5,52335815 | 1,00971123 | 0,32915856 | 0,59213802 |
| 255877_at | BCL6B | -0,11368198 | 5,44995023 | -1,00970759 | 0,32916025 | 0,59213802 |
| 2128_at | EVX1 | -0,09717559 | 5,05799771 | -1,00956921 | 0,32922436 | 0,59219804 |
| 65055_at | REEP1 | -0,11834919 | 4,33673206 | -1,00935645 | 0,32932293 | 0,59232004 |
| 55295_at | KLHL26 | 0,09133523 | 6,30027104 | 1,00916941 | 0,32940961 | 0,59242063 |
| 116238_at | TLCD1 | 0,22376211 | 8,07490717 | 1,00890221 | 0,32953347 | 0,59258805 |
| 644215_at | LOC644215 | -0,12860498 | 6,19269625 | -1,00867383 | 0,32963935 | 0,59272313 |
| 53841_at | CDHR5 | -0,08317301 | 4,77052597 | -1,00840633 | 0,32976341 | 0,59285268 |
| 71_at | ACTG1 | -0,06069102 | 13,2730859 | -1,00835975 | 0,32978501 | 0,59285268 |
| 9159_at | PCSK7 | -0,08222476 | 8,03988278 | -1,00831939 | 0,32980373 | 0,59285268 |
| 1366_at | CLDN7 | -0,18984528 | 8,76726594 | -1,00812425 | 0,32989426 | 0,59291375 |
| 122553_at | TRAPPC6B | 0,15935054 | 8,80579568 | 1,00790927 | 0,32999401 | 0,59291375 |
| 84142_at | FAM175A | 0,23333295 | 5,82509743 | 1,00788415 | 0,33000567 | 0,59291375 |
| 6993_at | DYNLT1 | -0,24985366 | 10,7492073 | -1,00785459 | 0,33001939 | 0,59291375 |
| 3910_at | LAMA4 | -0,114789 | 5,01301091 | -1,0078351 | 0,33002843 | 0,59291375 |
| 22_at | ABCB7 | -0,09222398 | 8,00608344 | -1,0078063 | 0,3300418 | 0,59291375 |
| 9123_at | SLC16A3 | 0,13087214 | 9,29726597 | 1,00778181 | 0,33005317 | 0,59291375 |
| 221468_at | TMEM217 | 0,07202609 | 4,42544362 | 1,00770451 | 0,33008905 | 0,59292291 |
| 91663_at | MYADM | -0,30680036 | 8,98373471 | -1,00759723 | 0,33013884 | 0,59295706 |
| 148252_at | DIRAS1 | -0,08638945 | 5,39195692 | -1,00748042 | 0,33019307 | 0,59299917 |
| 658_at | BMPR1B | -0,06525386 | 4,04265573 | -1,00724275 | 0,33030342 | 0,59306168 |
| 3301_at | DNAJA1 | -0,11734482 | 10,6419874 | -1,00721817 | 0,33031483 | 0,59306168 |
| 10290_at | SPEG | -0,10633651 | 5,24233249 | -1,00720479 | 0,33032105 | 0,59306168 |
| 9734_at | HDAC9 | 0,04742847 | 4,06206992 | 1,00708296 | 0,33037763 | 0,59306168 |
| 4824_at | NKX3-1 | 0,25008958 | 6,41264963 | 1,00707394 | 0,33038181 | 0,59306168 |
| 8425_at | LTBP4 | -0,09036633 | 5,854785 | -1,00681969 | 0,33049992 | 0,5932184 |
| 124925_at | SEZ6 | -0,10307745 | 5,17673688 | -1,00651895 | 0,33063965 | 0,59341392 |
| 79813_at | EHMT1 | 0,21890607 | 6,85383403 | 1,0062891 | 0,33074647 | 0,5934208 |
| 10332_at | CLEC4M | -0,09374174 | 3,86467852 | -1,00624283 | 0,33076798 | 0,5934208 |
| 158228_at | FAM201A | -0,160427 | 5,13314148 | -1,00620624 | 0,33078499 | 0,5934208 |
| 127_at | ADH4 | 0,07070642 | 2,96019034 | 1,00616588 | 0,33080375 | 0,5934208 |
| 81602_at | CDADC1 | -0,33916937 | 4,30954847 | -1,006126 | 0,33082229 | 0,5934208 |
| 55350_at | VNN3 | 0,06074214 | 3,67171447 | 1,00611303 | 0,33082832 | 0,5934208 |
| 79187_at | FSD1 | -0,07557828 | 6,12960936 | -1,00603664 | 0,33086384 | 0,59342924 |
| 23145_at | SSPO | -0,11018825 | 5,08711558 | -1,00596108 | 0,33089896 | 0,59343699 |
| 1558_at | CYP2C8 | 0,13331063 | 3,22654674 | 1,00582667 | 0,33096146 | 0,5934489 |
| 5136_at | PDE1A | -0,04941589 | 3,25635554 | -1,0058143 | 0,33096722 | 0,5934489 |
| 57489_at | ODF2L | 0,120793 | 6,17002996 | 1,00552853 | 0,33110013 | 0,59359713 |

| | | | | | | |
|---|---|---|---|---|---|---|
| 3514_at | IGKC | -0,11670584 | 5,35687377 | -1,00550405 | 0,33111152 | 0,59359713 |
| 644943_at | RASSF10 | -0,3689207 | 5,07120529 | -1,00538423 | 0,33116726 | 0,59361163 |
| 2987_at | GUK1 | 0,10090161 | 8,90224982 | 1,00526341 | 0,33122347 | 0,59361163 |
| 3267_at | AGFG1 | -0,08797631 | 9,16627504 | -1,00517381 | 0,33126517 | 0,59361163 |
| 158038_at | LINGO2 | -0,05546424 | 3,17883099 | -1,00511507 | 0,3312925 | 0,59361163 |
| 4314_at | MMP3 | -0,06970938 | 3,22729053 | -1,00503363 | 0,3313304 | 0,59361163 |
| 5179_at | PENK | -0,08825361 | 4,12694249 | -1,00502554 | 0,33133416 | 0,59361163 |
| 6582_at | SLC22A2 | -0,18640095 | 4,30853976 | -1,00502306 | 0,33133532 | 0,59361163 |
| 57661_at | PHRF1 | -0,07711991 | 7,05242069 | -1,00487435 | 0,33140453 | 0,59368042 |
| 285429_at | DCAF4L1 | -0,08695904 | 3,34116724 | -1,00479399 | 0,33144194 | 0,59369222 |
| 5408_at | PNLIPRP2 | -0,06352541 | 3,14365396 | -1,00466198 | 0,3315034 | 0,59374709 |
| 5967_at | REG1A | -0,10998394 | 4,54180394 | -1,00418034 | 0,33172769 | 0,59409357 |
| 116969_at | ART5 | -0,10732719 | 5,62733627 | -1,00402772 | 0,33179878 | 0,59416566 |
| 84065_at | TMEM222 | -0,0603361 | 8,62766417 | -1,00364131 | 0,33197884 | 0,59442645 |
| 23101_at | MCF2L2 | 0,065071 | 3,20225047 | 1,00358273 | 0,33200613 | 0,59442645 |
| 2857_at | GPR34 | -0,07708599 | 3,09188523 | -1,00331278 | 0,33213197 | 0,59459649 |
| 189_at | AGXT | -0,1220103 | 4,80160125 | -1,00321293 | 0,33217852 | 0,59462456 |
| 400794_at | LOC400794 | -0,12934762 | 4,37664389 | -1,00287174 | 0,33233763 | 0,5948541 |
| 56344_at | CABP5 | -0,10151836 | 4,39957338 | -1,00268284 | 0,33242575 | 0,59492479 |
| 10681_at | GNB5 | 0,14188884 | 7,65219596 | 1,00265465 | 0,33243889 | 0,59492479 |
| 101928476_a | LOC10192847 | -0,22003235 | 3,96393589 | -1,00244543 | 0,33253651 | 0,59504419 |
| 6354_at | CCL7 | -0,09239363 | 3,56736229 | -1,00234576 | 0,33258302 | 0,59506812 |
| 1087_at | CEACAM7 | -0,07959254 | 3,75184768 | -1,00228437 | 0,33261166 | 0,59506812 |
| 9940_at | DLEC1 | -0,08941378 | 4,9135915 | -1,00212779 | 0,33268475 | 0,59509386 |
| 10203_at | CALCRL | 0,05473181 | 3,2985334 | 1,00212117 | 0,33268783 | 0,59509386 |
| 101928999_a | IQCF5-AS1 | -0,08786849 | 4,01777298 | -1,00200548 | 0,33274184 | 0,59511011 |
| 339535_at | LINC01139 | 0,06865154 | 3,83976823 | 1,0019157 | 0,33278375 | 0,59511011 |
| 7424_at | VEGFC | -0,11171746 | 6,98169276 | -1,0017851 | 0,33284473 | 0,59511011 |
| 4010_at | LMX1B | -0,14070427 | 5,21817777 | -1,00169022 | 0,33288903 | 0,59511011 |
| 147_at | ADRA1B | -0,08747801 | 4,37645653 | -1,0016508 | 0,33290744 | 0,59511011 |
| 101927402_a | LOC10192740 | 0,12355575 | 3,45165334 | 1,00157614 | 0,3329423 | 0,59511011 |
| 56667_at | MUC13 | -0,06545997 | 3,6087951 | -1,00154316 | 0,33295771 | 0,59511011 |
| 100288152_a | LOC10028815 | 0,15711064 | 6,37513952 | 1,00142343 | 0,33301363 | 0,59511011 |
| 140609_at | NEK7 | -0,22124088 | 10,0173788 | -1,00139233 | 0,33302815 | 0,59511011 |
| 1396_at | CRIP1 | -0,11835079 | 4,90181955 | -1,00135465 | 0,33304575 | 0,59511011 |
| 102723927_a | LOC10272392 | -0,12047684 | 4,25506021 | -1,00133941 | 0,33305287 | 0,59511011 |
| 6954_at | TCP11 | -0,11455966 | 4,31509008 | -1,00130777 | 0,33306765 | 0,59511011 |
| 23774_at | BRD1 | 0,08956921 | 7,75351566 | 1,00106897 | 0,33317922 | 0,59525425 |
| 112939_at | NACC1 | -0,08614466 | 7,4376066 | -1,0009605 | 0,3332299 | 0,5952719 |
| 283234_at | CCDC88B | -0,10640348 | 6,4859945 | -1,00088338 | 0,33326594 | 0,5952719 |
| 930_at | CD19 | -0,16291568 | 4,98957295 | -1,00082336 | 0,33329399 | 0,5952719 |
| 340069_at | FAM170A | -0,10229925 | 5,01860973 | -1,00075602 | 0,33332546 | 0,5952719 |
| 283197_at | LINC00301 | -0,06116264 | 2,82273994 | -1,000665 | 0,33336801 | 0,5952719 |
| 256646_at | NUTM1 | -0,10169938 | 4,97369999 | -1,00061921 | 0,33338941 | 0,5952719 |
| 29106_at | SCG3 | -0,05986163 | 2,92854276 | -1,00058498 | 0,33340542 | 0,5952719 |
| 286753_at | TUSC5 | -0,12839889 | 5,34556755 | -1,00045544 | 0,33346598 | 0,59532485 |
| 3868_at | KRT16 | -0,44820707 | 6,76419769 | -1,00017079 | 0,33359908 | 0,59550728 |
| 2901_at | GRIK5 | -0,10352094 | 5,66727274 | -1,00006871 | 0,33364683 | 0,59553732 |
| 643866_at | CBLN3 | 0,13599331 | 5,56700341 | 0,99999939 | 0,33367925 | 0,59554001 |
| 54870_at | QRICH1 | -0,07856741 | 9,02358225 | -0,99977251 | 0,33378538 | 0,59563448 |

| | | | | | | |
|---|---|---|---|---|---|---|
| 57606_at | SLAIN2 | -0,11632561 | 8,15952474 | -0,99975405 | 0,33379402 | 0,59563448 |
| 101928185_a | LOC10192818 | -0,11741099 | 4,55828972 | -0,99953064 | 0,33389856 | 0,59576583 |
| 2787_at | GNG5 | 0,16951223 | 9,37359331 | 0,99933945 | 0,33398804 | 0,59587029 |
| 257203_at | DSCR9 | -0,10173514 | 4,63066844 | -0,999201 | 0,33405284 | 0,59591789 |
| 25769_at | SLC24A2 | -0,08123183 | 4,32737594 | -0,9991315 | 0,33408538 | 0,59591789 |
| 714_at | C1QC | -0,11909797 | 4,66495876 | -0,9990842 | 0,33410752 | 0,59591789 |
| 165904_at | XIRP1 | -0,10034486 | 5,51255567 | -0,99872707 | 0,33427476 | 0,59612805 |
| 26468_at | LHX6 | -0,1027767 | 5,83245711 | -0,9987004 | 0,33428725 | 0,59612805 |
| 11145_at | PLA2G16 | 0,1946725 | 7,59391342 | 0,99850656 | 0,33437804 | 0,59619263 |
| 9423_at | NTN1 | -0,13067921 | 5,60029743 | -0,99849094 | 0,33438536 | 0,59619263 |
| 4693_at | NDP | 0,09525423 | 3,63225642 | 0,99824021 | 0,33450283 | 0,59634603 |
| 55_at | ACPP | 0,20566395 | 5,15317559 | 0,99813549 | 0,3345519 | 0,59634603 |
| 100506542_a | LINC01082 | -0,10640401 | 4,13175991 | -0,99810909 | 0,33456427 | 0,59634603 |
| 4117_at | MAK | -0,0761912 | 3,81308609 | -0,99775206 | 0,33473162 | 0,59653725 |
| 55998_at | NXF5 | -0,11627829 | 4,25886503 | -0,99774808 | 0,33473349 | 0,59653725 |
| 84660_at | CCDC62 | -0,08203748 | 4,16890544 | -0,99761952 | 0,33479376 | 0,59658948 |
| 25837_at | RAB26 | 0,35563767 | 7,02500106 | 0,99745744 | 0,33486976 | 0,59666971 |
| 2581_at | GALC | 0,50134484 | 6,78259914 | 0,99726156 | 0,33496163 | 0,59675732 |
| 101929690_a | LINC01583 | 0,05572105 | 3,16425843 | 0,99722048 | 0,33498089 | 0,59675732 |
| 9091_at | PIGQ | 0,12076551 | 6,60158501 | 0,99702326 | 0,33507341 | 0,59683357 |
| 2339_at | FNTA | 0,23382365 | 9,77308379 | 0,99699714 | 0,33508566 | 0,59683357 |
| 574538_at | LOC574538 | -0,0953074 | 3,7083999 | -0,99682489 | 0,33516648 | 0,59683687 |
| 2019_at | EN1 | -0,16654965 | 5,19355698 | -0,99680968 | 0,33517362 | 0,59683687 |
| 56929_at | FEM1C | 0,10587678 | 7,85782694 | 0,99679509 | 0,33518046 | 0,59683687 |
| 6820_at | SULT2B1 | 0,22989067 | 6,39485644 | 0,99650052 | 0,33531871 | 0,5969785 |
| 2206_at | MS4A2 | -0,06232941 | 3,07920069 | -0,99647709 | 0,33532971 | 0,5969785 |
| 8443_at | GNPAT | -0,09443098 | 8,64906299 | -0,99642753 | 0,33535298 | 0,5969785 |
| 157773_at | C8orf48 | 0,39070563 | 4,59589297 | 0,99633979 | 0,33539416 | 0,59699665 |
| 2919_at | CXCL1 | 0,73150065 | 6,00139241 | 0,99621739 | 0,33545163 | 0,59700496 |
| 7035_at | TFPI | 0,33433903 | 7,41122511 | 0,99619747 | 0,33546098 | 0,59700496 |
| 387357_at | THEMIS | -0,05575618 | 2,92444991 | -0,99613183 | 0,3354918 | 0,59700496 |
| 89885_at | FATE1 | -0,15350746 | 4,92044917 | -0,99600245 | 0,33555256 | 0,59703272 |
| 9898_at | UBAP2L | -0,14694501 | 8,83523415 | -0,9959666 | 0,3355694 | 0,59703272 |
| 146439_at | CCDC64B | 0,34968079 | 7,04152412 | 0,99589568 | 0,3356027 | 0,59703684 |
| 51155_at | HN1 | -0,09208834 | 10,4195413 | -0,99558737 | 0,33574753 | 0,59723932 |
| 27443_at | CECR2 | 0,08574129 | 4,21878068 | 0,9954723 | 0,3358016 | 0,59724938 |
| 94137_at | RP1L1 | -0,1091982 | 4,53300221 | -0,99544337 | 0,33581519 | 0,59724938 |
| 2696_at | GIPR | -0,11322815 | 5,78958582 | -0,99527097 | 0,3358962 | 0,59732152 |
| 9464_at | HAND2 | 0,16702875 | 4,26653425 | 0,99522508 | 0,33591777 | 0,59732152 |
| 147746_at | HIPK4 | -0,08758674 | 6,00952891 | -0,99500905 | 0,33601931 | 0,59744693 |
| 4143_at | MAT1A | -0,10870276 | 4,86600393 | -0,99482627 | 0,33610524 | 0,59751889 |
| 101929084_a | LOC10192908 | -0,14575413 | 4,73445993 | -0,99464155 | 0,3361921 | 0,59751889 |
| 51265_at | CDKL3 | 0,0832855 | 4,37964794 | 0,99463232 | 0,33619644 | 0,59751889 |
| 8383_at | OR1A1 | -0,12641953 | 3,8100264 | -0,99460759 | 0,33620807 | 0,59751889 |
| 7776_at | ZNF236 | 0,06802015 | 6,64617768 | 0,99459311 | 0,33621488 | 0,59751889 |
| 3827_at | KNG1 | -0,08786874 | 3,29443514 | -0,99447748 | 0,33626927 | 0,59756041 |
| 91574_at | C12orf65 | -0,1508922 | 7,46314305 | -0,99432674 | 0,33634017 | 0,59759667 |
| 6521_at | SLC4A1 | -0,08473105 | 4,41437285 | -0,9942536 | 0,33637457 | 0,59759667 |
| 205251_at | LINC00116 | -0,90138273 | 4,99495974 | -0,99417531 | 0,33641141 | 0,59759667 |
| 200942_at | KLHDC8B | 0,14365239 | 5,52018826 | 0,99417031 | 0,33641376 | 0,59759667 |

| | | | | | | |
|---|---|---|---|---|---|---|
| 5096_at | PCCB | -0,2480199 | 8,69420196 | -0,99407948 | 0,33645649 | 0,5976016 |
| 27334_at | P2RY10 | -0,07705608 | 3,12901382 | -0,99403253 | 0,33647859 | 0,5976016 |
| 161931_at | ADAD2 | -0,09298803 | 6,2450918 | -0,99385159 | 0,33656373 | 0,59765676 |
| 10486_at | CAP2 | 0,28410499 | 8,09837829 | 0,99382851 | 0,3365746 | 0,59765676 |
| 343069_at | HNRNPCL1 | -0,12368518 | 5,78728359 | -0,99376875 | 0,33660272 | 0,59765676 |
| 23035_at | PHLPP2 | 0,17465423 | 7,55180244 | 0,99348868 | 0,33673456 | 0,59783575 |
| 6326_at | SCN2A | 0,08511645 | 3,17819039 | 0,99335829 | 0,33679595 | 0,59784079 |
| 27239_at | GPR162 | -0,09051782 | 5,1083958 | -0,99335081 | 0,33679947 | 0,59784079 |
| 51172_at | NAGPA | 0,16240263 | 7,15164571 | 0,99288553 | 0,33701861 | 0,59797591 |
| 5217_at | PFN2 | -0,14785164 | 11,5524937 | -0,99284173 | 0,33703924 | 0,59797591 |
| 101929172_a | LOC10192917 | -0,12204363 | 3,86233483 | -0,99275454 | 0,33708032 | 0,59797591 |
| 63894_at | VIPAS39 | 0,14232901 | 7,88814438 | 0,99272574 | 0,33709389 | 0,59797591 |
| 6199_at | RPS6KB2 | -0,10265095 | 7,88932651 | -0,99269854 | 0,3371067 | 0,59797591 |
| 91893_at | FDXACB1 | -0,08954149 | 5,62424444 | -0,99268912 | 0,33711114 | 0,59797591 |
| 63877_at | FAM204A | -0,05328896 | 6,4901796 | -0,99263363 | 0,33713729 | 0,59797591 |
| 10216_at | PRG4 | 0,171075 | 3,38913474 | 0,99260271 | 0,33715186 | 0,59797591 |
| 9797_at | TATDN2 | -0,10395719 | 7,63058282 | -0,99257151 | 0,33716656 | 0,59797591 |
| 166929_at | SGMS2 | -0,12849041 | 6,25321551 | -0,9924994 | 0,33720054 | 0,59797591 |
| 101928770_a | LINC01526 | -0,13754915 | 3,94494715 | -0,99242707 | 0,33723463 | 0,59797591 |
| 102800310_a | HAGLROS | -0,23259063 | 7,06707292 | -0,99224705 | 0,33731948 | 0,59797591 |
| 642864_at | SPATA42 | -0,09481343 | 2,91420136 | -0,99223899 | 0,33732328 | 0,59797591 |
| 23197_at | FAF2 | -0,08990474 | 9,43716387 | -0,99220865 | 0,33733758 | 0,59797591 |
| 128646_at | SIRPD | -0,09570145 | 4,49441712 | -0,9921404 | 0,33736975 | 0,59797591 |
| 3589_at | IL11 | -0,11878755 | 4,99607143 | -0,99213504 | 0,33737228 | 0,59797591 |
| 1421_at | CRYGD | -0,08137127 | 4,51564041 | -0,9919048 | 0,33748083 | 0,59809615 |
| 114991_at | ZNF618 | 0,48159008 | 6,58373056 | 0,99185946 | 0,33750221 | 0,59809615 |
| 29894_at | CPSF1 | -0,1838379 | 7,76232767 | -0,99179129 | 0,33753436 | 0,5980981 |
| 6543_at | SLC8A2 | -0,13912547 | 5,05896819 | -0,99163451 | 0,3376083 | 0,5981571 |
| 100507066_a | CLIP1-AS1 | -0,05324241 | 4,0023223 | -0,99149969 | 0,3376719 | 0,5981571 |
| 101929121_a | LINC00437 | 0,08146221 | 2,95561326 | 0,9914748 | 0,33768364 | 0,5981571 |
| 5317_at | PKP1 | -0,33565135 | 6,3087362 | -0,99145736 | 0,33769187 | 0,5981571 |
| 695_at | BTK | -0,09173483 | 4,07880391 | -0,99134256 | 0,33774603 | 0,59819802 |
| 101928017_a | UBXN10-AS1 | -0,14922887 | 5,97176891 | -0,99122887 | 0,33779967 | 0,59823803 |
| 9288_at | TAAR3 | -0,11567827 | 3,41232251 | -0,9910037 | 0,33790593 | 0,59835061 |
| 317716_at | BPIFA4P | -0,10988695 | 3,79053275 | -0,99096252 | 0,33792537 | 0,59835061 |
| 440193_at | CCDC88C | -0,12525808 | 7,07234536 | -0,99083715 | 0,33798454 | 0,59835668 |
| 91584_at | PLXNA4 | -0,06305619 | 4,29346981 | -0,99082365 | 0,33799091 | 0,59835668 |
| 2961_at | GTF2E2 | -0,24766702 | 8,05931897 | -0,99040783 | 0,33818724 | 0,5985673 |
| 51477_at | ISYNA1 | 0,31017656 | 7,79296937 | 0,99037568 | 0,33820242 | 0,5985673 |
| 54093_at | SETD4 | -0,27264131 | 6,41078147 | -0,99033329 | 0,33822245 | 0,5985673 |
| 53637_at | S1PR5 | -0,22544507 | 6,85831505 | -0,99029258 | 0,33824167 | 0,5985673 |
| 89778_at | SERPINB11 | -0,09822517 | 3,31816242 | -0,99018513 | 0,33829242 | 0,5985673 |
| 149840_at | C20orf196 | 0,08872134 | 6,08698265 | 0,99011121 | 0,33832734 | 0,5985673 |
| 400891_at | LRRC74B | -0,09366652 | 3,44971275 | -0,99000081 | 0,33837949 | 0,5985673 |
| 10536_at | P3H3 | -0,10475318 | 5,17078537 | -0,98998844 | 0,33838534 | 0,5985673 |
| 23619_at | ZIM2 | -0,0983459 | 4,67832091 | -0,98997486 | 0,33839175 | 0,5985673 |
| 162083_at | C16orf82 | -0,11097868 | 4,88649418 | -0,98991375 | 0,33842062 | 0,5985673 |
| 126231_at | ZNF573 | 0,11493331 | 6,15334254 | 0,98963996 | 0,33855 | 0,59874115 |
| 10147_at | SUGP2 | -0,06935806 | 7,43093536 | -0,98951926 | 0,33860705 | 0,59876483 |
| 57217_at | TTC7A | -0,10148275 | 6,31948555 | -0,98938978 | 0,33866825 | 0,59876483 |

| | | | | | | |
|---|---|---|---|---|---|---|
| 57332_at | CBX8 | -0,08070044 | 5,73320312 | -0,98937828 | 0,33867369 | 0,59876483 |
| 202865_at | C7orf33 | -0,08646486 | 3,56785578 | -0,98932245 | 0,33870008 | 0,59876483 |
| 338376_at | IFNE | 0,07428761 | 3,30579271 | 0,98927255 | 0,33872367 | 0,59876483 |
| 9310_at | ZNF235 | 0,10581933 | 5,41509967 | 0,98916779 | 0,3387732 | 0,59876483 |
| 286178_at | LOC286178 | 0,18014961 | 4,38608734 | 0,98915134 | 0,33878097 | 0,59876483 |
| 91947_at | ARRDC4 | 0,19277219 | 8,05483483 | 0,98856443 | 0,33905857 | 0,59920047 |
| 100507203_a | SMLR1 | -0,08633445 | 3,38668269 | -0,98839242 | 0,33913995 | 0,59926171 |
| 90835_at | C16orf93 | -0,3052869 | 5,68831989 | -0,98835969 | 0,33915544 | 0,59926171 |
| 4885_at | NPTX2 | -0,09534866 | 4,15383985 | -0,9881145 | 0,33927148 | 0,59937975 |
| 8120_at | AP3B2 | -0,08221204 | 4,18141391 | -0,98808705 | 0,33928447 | 0,59937975 |
| 101928307_a | LOC10192830 | 0,07652708 | 4,31074924 | 0,98796617 | 0,33934169 | 0,59942586 |
| 11197_at | WIF1 | -0,0829349 | 3,50498171 | -0,98777777 | 0,33943088 | 0,59952843 |
| 606553_at | C8orf49 | -0,08273963 | 3,87716234 | -0,98762195 | 0,33950467 | 0,59960378 |
| 28409_at | IGHV3-73 | 0,10345099 | 3,27815182 | 0,98737732 | 0,33962052 | 0,5997534 |
| 51302_at | CYP39A1 | 0,11416651 | 4,50324178 | 0,98721377 | 0,339698 | 0,59978181 |
| 118924_at | FRA10AC1 | -0,0590075 | 5,43610128 | -0,98721191 | 0,33969888 | 0,59978181 |
| 56834_at | GPR137 | 0,07693219 | 7,0957995 | 0,98696891 | 0,33981401 | 0,59991955 |
| 23235_at | SIK2 | -0,0672336 | 7,08139559 | -0,98663123 | 0,33997405 | 0,59991955 |
| 644353_at | ZCCHC18 | 0,16726432 | 4,73993929 | 0,98659881 | 0,33998941 | 0,59991955 |
| 85358_at | SHANK3 | -0,09989171 | 5,39883959 | -0,98659586 | 0,33999081 | 0,59991955 |
| 51429_at | SNX9 | 0,09295971 | 8,05973199 | 0,98659228 | 0,33999251 | 0,59991955 |
| 100507642_a | LOC10050764 | 0,33823364 | 5,21814581 | 0,98652894 | 0,34002253 | 0,59991955 |
| 283761_at | LINC00928 | -0,09358296 | 4,38758546 | -0,98650139 | 0,3400356 | 0,59991955 |
| 415116_at | PIM3 | -0,12398608 | 8,89106235 | -0,98637529 | 0,34009538 | 0,59991955 |
| 155_at | ADRB3 | -0,11265438 | 4,71943156 | -0,98631971 | 0,34012173 | 0,59991955 |
| 100505782_a | LOC10050578 | 0,06642344 | 3,66959598 | 0,98630553 | 0,34012846 | 0,59991955 |
| 2831_at | NPBWR1 | -0,09680418 | 5,76167071 | -0,98626898 | 0,34014579 | 0,59991955 |
| 164091_at | PAQR7 | 0,19012791 | 7,17201022 | 0,98624906 | 0,34015523 | 0,59991955 |
| 1828_at | DSG1 | 0,14511946 | 3,08323024 | 0,98612997 | 0,3402117 | 0,59991955 |
| 284385_at | ZNF561-AS1 | 0,07896445 | 5,96606399 | 0,98612744 | 0,34021291 | 0,59991955 |
| 57213_at | SPRYD7 | -0,40590741 | 4,93183458 | -0,98588664 | 0,34032712 | 0,60006602 |
| 29989_at | OBP2B | -0,10329422 | 4,66230893 | -0,98542656 | 0,34054541 | 0,60039596 |
| 100859921_a | LINC00536 | -0,16497509 | 3,74725915 | -0,98528522 | 0,34061249 | 0,60045927 |
| 146512_at | FLJ30679 | -0,10101417 | 4,42145888 | -0,98501995 | 0,34073841 | 0,60057695 |
| 57546_at | PDP2 | 0,12706771 | 6,68712729 | 0,98501324 | 0,3407416 | 0,60057695 |
| 5357_at | PLS1 | -0,13196246 | 8,39981532 | -0,98489226 | 0,34079904 | 0,60061073 |
| 100129098_a | LOC10012909 | -0,11689734 | 4,79024421 | -0,98484154 | 0,34082313 | 0,60061073 |
| 83860_at | TAF3 | -0,1125616 | 6,8181623 | -0,98465818 | 0,3409102 | 0,60068365 |
| 6618_at | SNAPC2 | -0,06536874 | 6,5534317 | -0,98462308 | 0,34092687 | 0,60068365 |
| 219525_at | OR5AK4P | -0,09724977 | 3,71407957 | -0,98444298 | 0,34101242 | 0,60077943 |
| 286077_at | FAM83H | -0,13646878 | 8,19180314 | -0,9841791 | 0,34113779 | 0,60091006 |
| 8697_at | CDC23 | -0,09522224 | 8,86488702 | -0,98415559 | 0,34114896 | 0,60091006 |
| 10522_at | DEAF1 | 0,21646446 | 6,71108084 | 0,9840209 | 0,34121297 | 0,60096785 |
| 325_at | APCS | -0,08974916 | 4,68181679 | -0,98391078 | 0,3412653 | 0,60100507 |
| 54971_at | BANP | -0,13073184 | 7,54451822 | -0,98378927 | 0,34132306 | 0,60103946 |
| 8451_at | CUL4A | -0,07032016 | 8,43582723 | -0,98373842 | 0,34134723 | 0,60103946 |
| 9572_at | NR1D1 | -0,09827157 | 7,53276687 | -0,98345619 | 0,34148141 | 0,60119223 |
| 169355_at | IDO2 | -0,06388972 | 3,30717091 | -0,98342464 | 0,34149641 | 0,60119223 |
| 23348_at | DOCK9 | -0,13589251 | 5,9659582 | -0,98316483 | 0,34161997 | 0,60135479 |
| 79861_at | TUBAL3 | 0,08974282 | 3,58754823 | 0,98269568 | 0,34184316 | 0,60163468 |

| | | | | | | |
|---|---|---|---|---|---|---|
| 4780_at | NFE2L2 | 0,15152847 | 10,3495306 | 0,98260498 | 0,34188632 | 0,60163468 |
| 1647_at | GADD45A | 0,30042514 | 9,61861779 | 0,98258087 | 0,3418978 | 0,60163468 |
| 100_at | ADA | -0,49735557 | 8,05381223 | -0,98256805 | 0,3419039 | 0,60163468 |
| 118471_at | PRAP1 | -0,10806486 | 5,12522057 | -0,98236612 | 0,34200001 | 0,60174883 |
| 286207_at | C9orf117 | -0,12770534 | 4,77091817 | -0,98216491 | 0,3420958 | 0,60186239 |
| 9320_at | TRIP12 | 0,08987837 | 9,4612778 | 0,9817836 | 0,34227737 | 0,60212685 |
| 1329_at | COX5B | -0,09372631 | 11,0268788 | -0,98171203 | 0,34231146 | 0,60213183 |
| 219347_at | TMEM254-AS | -0,06693985 | 4,99875353 | -0,9814694 | 0,34242705 | 0,6022508 |
| 10157_at | AASS | 0,4986471 | 5,63803706 | 0,9813876 | 0,34246602 | 0,6022508 |
| 4336_at | MOBP | -0,07007069 | 3,85464538 | -0,98137318 | 0,34247289 | 0,6022508 |
| 100287098_a | TONSL-AS1 | 0,11570684 | 5,1803162 | 0,98088526 | 0,34270543 | 0,60260472 |
| 10021_at | HCN4 | -0,08379759 | 4,99866831 | -0,98072078 | 0,34278384 | 0,60268758 |
| 101927770_a | LOC10192777 | -0,09370672 | 3,6508829 | -0,98051787 | 0,3428806 | 0,60280268 |
| 9886_at | RHOBTB1 | 0,14427282 | 4,4998317 | 0,9801956 | 0,3430343 | 0,60301786 |
| 83639_at | TEX101 | -0,07247363 | 4,12935786 | -0,98005324 | 0,34310222 | 0,60307288 |
| 4807_at | NHLH1 | -0,11380545 | 5,1624026 | -0,97999875 | 0,34312821 | 0,60307288 |
| 84330_at | ZNF414 | 0,10616637 | 7,54429636 | 0,97984463 | 0,34320175 | 0,6031471 |
| 55585_at | UBE2Q1 | 0,07348271 | 8,22402876 | 0,97973043 | 0,34325625 | 0,60318784 |
| 4701_at | NDUFA7 | -0,11723013 | 9,41517076 | -0,979585 | 0,34332566 | 0,60325478 |
| 5506_at | PPP1R3A | 0,05414265 | 2,87007917 | 0,97920315 | 0,34350796 | 0,60352004 |
| 50809_at | HP1BP3 | 0,09932428 | 9,59489742 | 0,97908086 | 0,34356635 | 0,60356759 |
| 339975_at | LOC339975 | -0,08128154 | 3,8792487 | -0,97893209 | 0,3436374 | 0,60360364 |
| 29082_at | CHMP4A | 0,05058514 | 8,62268895 | 0,97890668 | 0,34364954 | 0,60360364 |
| 57016_at | AKR1B10 | -0,75283477 | 9,69446414 | -0,97873523 | 0,34373143 | 0,60369244 |
| 440387_at | CTRB2 | -0,13977114 | 6,47339959 | -0,97864109 | 0,34377641 | 0,60371638 |
| 340267_at | COL28A1 | 0,07553257 | 3,95376186 | 0,97822973 | 0,34397298 | 0,60395762 |
| 256472_at | TMEM151A | -0,1255644 | 5,62046948 | -0,97815715 | 0,34400767 | 0,60395762 |
| 26952_at | SMR3A | -0,12396291 | 3,59050524 | -0,97815679 | 0,34400784 | 0,60395762 |
| 9044_at | BTAF1 | -0,17973604 | 8,039275 | -0,97802936 | 0,34406875 | 0,60398716 |
| 7289_at | TULP3 | -0,10192996 | 7,95480542 | -0,97796772 | 0,34409822 | 0,60398716 |
| 55799_at | CACNA2D3 | 0,09988901 | 3,50662949 | 0,97792482 | 0,34411873 | 0,60398716 |
| 6415_at | SEPW1 | -0,11435567 | 9,72295365 | -0,97783457 | 0,34416188 | 0,60400786 |
| 1497_at | CTNS | -0,06803146 | 6,86788945 | -0,97768532 | 0,34423325 | 0,60407808 |
| 2676_at | GFRA3 | -0,100545 | 4,74726914 | -0,97754324 | 0,34430119 | 0,60409882 |
| 50835_at | TAS2R9 | -0,10357588 | 3,4383243 | -0,97752944 | 0,34430779 | 0,60409882 |
| 84837_at | ARHGAP5-AS1 | 0,10936408 | 6,76623317 | 0,97742894 | 0,34435586 | 0,60412814 |
| 7408_at | VASP | -0,18052645 | 8,61454749 | -0,97728961 | 0,34442251 | 0,60419004 |
| 6283_at | S100A12 | 0,20187296 | 5,23318087 | 0,97706388 | 0,34453051 | 0,60432446 |
| 6868_at | ADAM17 | 0,10371799 | 8,59712768 | 0,97688429 | 0,34461645 | 0,60437469 |
| 1635_at | DCTD | -0,25719303 | 9,16808411 | -0,97687291 | 0,3446219 | 0,60437469 |
| 1830_at | DSG3 | -0,11300392 | 4,55147455 | -0,97678528 | 0,34466384 | 0,60439322 |
| 58189_at | WFDC1 | -0,11882084 | 4,57551435 | -0,97656151 | 0,34477095 | 0,60447144 |
| 7480_at | WNT10B | -0,14955855 | 6,22194148 | -0,97651 | 0,34479561 | 0,60447144 |
| 89777_at | SERPINB12 | -0,09686947 | 3,9173311 | -0,97640046 | 0,34484806 | 0,60447144 |
| 387119_at | CEP85L | -0,08937082 | 5,14509997 | -0,97639498 | 0,34485068 | 0,60447144 |
| 56979_at | PRDM9 | -0,08333905 | 3,31305041 | -0,97636435 | 0,34486535 | 0,60447144 |
| 10718_at | NRG3 | 0,39623657 | 4,36637884 | 0,97626863 | 0,34491118 | 0,60449678 |
| 646023_at | ADORA2A-AS1 | -0,10091556 | 4,62753378 | -0,97591137 | 0,34508229 | 0,60474164 |
| 56246_at | MRAP | -0,14373218 | 4,88873521 | -0,97561163 | 0,3452259 | 0,60487237 |
| 401264_at | TRAM2-AS1 | 0,17230975 | 6,44651907 | 0,97559181 | 0,34523539 | 0,60487237 |

| | | | | | | |
|---|---|---|---|---|---|---|
| 285220_at | EPHA6 | 0,05572781 | 3,16135775 | 0,97555907 | 0,34525109 | 0,60487237 |
| 9955_at | HS3ST3A1 | -0,09241832 | 3,58355984 | -0,97522428 | 0,34541154 | 0,60509845 |
| 121268_at | RHEBL1 | -0,15118324 | 5,11222015 | -0,97497867 | 0,34552929 | 0,60524968 |
| 79070_at | KDELC1 | -0,11346164 | 6,57151415 | -0,97463183 | 0,34569562 | 0,60546044 |
| 9890_at | LPPR4 | 0,08393877 | 3,29344845 | 0,97459669 | 0,34571247 | 0,60546044 |
| 55153_at | SDAD1 | -0,30105098 | 7,94567525 | -0,97449174 | 0,34576282 | 0,60549356 |
| 55250_at | ELP2 | -0,23873297 | 7,61223836 | -0,97418267 | 0,3459111 | 0,60569816 |
| 5784_at | PTPN14 | 0,24126385 | 7,77712617 | 0,97405307 | 0,34597329 | 0,60575141 |
| 2016_at | EMX1 | -0,09055776 | 4,73621463 | -0,97395475 | 0,34602048 | 0,60575141 |
| 8242_at | KDM5C | 0,16375838 | 7,23410084 | 0,97392272 | 0,34603585 | 0,60575141 |
| 55530_at | SVOP | -0,08054804 | 4,35636629 | -0,97378911 | 0,34609998 | 0,60580863 |
| 284632_at | LOC284632 | -0,09871465 | 4,12250484 | -0,97361366 | 0,34618421 | 0,60590101 |
| 10566_at | AKAP3 | 0,06470959 | 4,48430219 | 0,97341111 | 0,34628147 | 0,60601616 |
| 100130155_a | MIR124-2HG | -0,07213993 | 3,20841208 | -0,97318105 | 0,34639196 | 0,60615446 |
| 60437_at | CDH26 | 0,13618733 | 4,55708644 | 0,973093 | 0,34643425 | 0,6061734 |
| 1380_at | CR2 | -0,06468538 | 3,34650357 | -0,97293668 | 0,34650935 | 0,60624973 |
| 4720_at | NDUFS2 | -0,08754245 | 9,59999099 | -0,97245232 | 0,3467421 | 0,60643113 |
| 7041_at | TGFB1I1 | 0,35352933 | 7,83947255 | 0,97239819 | 0,34676812 | 0,60643113 |
| 27020_at | NPTN | 0,0618011 | 11,7303492 | 0,9723971 | 0,34676864 | 0,60643113 |
| 63904_at | DUSP21 | -0,0894472 | 4,18335446 | -0,97239515 | 0,34676958 | 0,60643113 |
| 54508_at | EPB41L4A-AS1 | 0,22123962 | 5,52730011 | 0,97239337 | 0,34677044 | 0,60643113 |
| 2149_at | F2R | 0,18596373 | 5,1826277 | 0,9720769 | 0,34692258 | 0,60664213 |
| 23515_at | MORC3 | 0,3631726 | 8,06997016 | 0,97191219 | 0,34700178 | 0,60668303 |
| 55671_at | PPP4R3A | -0,11079697 | 8,6536115 | -0,97189727 | 0,34700896 | 0,60668303 |
| 9107_at | MTMR6 | -0,09010048 | 6,49961439 | -0,97166503 | 0,34712066 | 0,60682324 |
| 284339_at | TMEM145 | -0,09180032 | 4,60958655 | -0,97145584 | 0,3472213 | 0,60694409 |
| 101928809_a | LOC10192880 | -0,06184079 | 3,2508832 | -0,9713512 | 0,34727165 | 0,60697702 |
| 101928710_a | LOC10192871 | -0,18262881 | 4,18220495 | -0,97127974 | 0,34730603 | 0,60698205 |
| 80863_at | PRRT1 | 0,23574995 | 5,49013177 | 0,9712064 | 0,34734132 | 0,60698695 |
| 5379_at | PMS2P1 | 0,05671096 | 8,13090794 | 0,97104703 | 0,34741802 | 0,60698695 |
| 948_at | CD36 | 0,58438378 | 4,92251377 | 0,97104284 | 0,34742004 | 0,60698695 |
| 113622_at | ADPRHL1 | -0,08114058 | 5,34301267 | -0,97098324 | 0,34744873 | 0,60698695 |
| 7033_at | TFF3 | -0,08772253 | 5,1266954 | -0,97094655 | 0,34746639 | 0,60698695 |
| 55616_at | ASAP3 | 0,24546195 | 6,45886423 | 0,97086528 | 0,34750551 | 0,60700025 |
| 4337_at | MOCS1 | 0,14293456 | 6,06497284 | 0,97068717 | 0,34759126 | 0,60709498 |
| 51634_at | RBMX2 | -0,11228048 | 8,10912477 | -0,97049565 | 0,34768348 | 0,60720099 |
| 56341_at | PRMT8 | -0,06692072 | 4,02744907 | -0,97019156 | 0,34782995 | 0,60740171 |
| 79948_at | LPPR3 | -0,10381449 | 5,59733994 | -0,97010118 | 0,34787349 | 0,60742268 |
| 10368_at | CACNG3 | -0,09402455 | 4,0916467 | -0,9697444 | 0,3480454 | 0,60766777 |
| 137735_at | ABRA | -0,06238828 | 3,26476134 | -0,96916151 | 0,34832639 | 0,60808733 |
| 2318_at | FLNC | -0,07514341 | 5,72696606 | -0,96911495 | 0,34834884 | 0,60808733 |
| 20_at | ABCA2 | -0,10224731 | 6,001052 | -0,96886718 | 0,34846833 | 0,60821674 |
| 10886_at | NPFFR2 | -0,0699138 | 3,19074765 | -0,96883031 | 0,34848612 | 0,60821674 |
| 3981_at | LIG4 | 0,19552092 | 6,2387858 | 0,96851904 | 0,34863629 | 0,60838 |
| 219623_at | TMEM26 | -0,09436783 | 3,22208099 | -0,96850547 | 0,34864283 | 0,60838 |
| 6721_at | SREBF2 | 0,11888098 | 7,86291467 | 0,96835291 | 0,34871645 | 0,60840732 |
| 140564_at | APOBEC3D | -0,06210387 | 6,10809422 | -0,96834212 | 0,34872166 | 0,60840732 |
| 5624_at | PROC | 0,31845092 | 7,28701774 | 0,96807502 | 0,34885057 | 0,60857712 |
| 284756_at | C20orf197 | -0,07891297 | 3,33475223 | -0,96777994 | 0,34899303 | 0,60876622 |
| 246269_at | LACE1 | 0,10059136 | 4,48662045 | 0,96771958 | 0,34902218 | 0,60876622 |

| | | | | | | |
|---|---|---|---|---|---|---|
| 100506649_a | PXN-AS1 | -0,12978005 | 6,66199336 | -0,96759897 | 0,34908042 | 0,60879951 |
| 8439_at | NSMAF | -0,14472235 | 7,96232794 | -0,96753569 | 0,34911098 | 0,60879951 |
| 284294_at | LOC284294 | -0,1341394 | 3,64619715 | -0,96745983 | 0,34914762 | 0,60879951 |
| 55930_at | MYO5C | 0,43272783 | 5,44549609 | 0,96734943 | 0,34920095 | 0,60879951 |
| 150696_at | PROM2 | 0,14329068 | 9,31651484 | 0,96727642 | 0,34923621 | 0,60879951 |
| 4852_at | NPY | -0,09322031 | 4,77827027 | -0,96725319 | 0,34924744 | 0,60879951 |
| 4055_at | LTBR | -0,11088556 | 9,37630454 | -0,96722202 | 0,34926249 | 0,60879951 |
| 83896_at | KRTAP3-1 | -0,14355374 | 5,51854328 | -0,96715048 | 0,34929706 | 0,60880467 |
| #N/A | GS1-259H13,2 | 0,13481253 | 5,21555366 | 0,96702853 | 0,34935598 | 0,60885228 |
| 65059_at | RAPH1 | -0,09470736 | 6,96176793 | -0,96686675 | 0,34943416 | 0,60893344 |
| 4267_at | CD99 | 0,09774808 | 9,91419082 | 0,96667172 | 0,34952842 | 0,6090426 |
| 22915_at | MMRN1 | 0,05168758 | 2,9395143 | 0,96626287 | 0,34972609 | 0,60933191 |
| 140775_at | SMCR8 | 0,17509215 | 7,33434535 | 0,9658773 | 0,34991257 | 0,60960168 |
| 55273_at | TMEM100 | -0,09820037 | 2,88994837 | -0,9656822 | 0,35000695 | 0,60971098 |
| 51363_at | CHST15 | 0,10254646 | 6,28475721 | 0,9654106 | 0,35013838 | 0,60988477 |
| 26153_at | KIF26A | -0,11463066 | 4,96524201 | -0,96507913 | 0,35029883 | 0,61010907 |
| 642597_at | C18orf42 | -0,08130511 | 3,89447231 | -0,96494793 | 0,35036234 | 0,61012281 |
| 51149_at | C5orf45 | -0,115315 | 6,49813581 | -0,96482337 | 0,35042266 | 0,61012281 |
| 6991_at | TCTE3 | -0,07942505 | 3,53031685 | -0,96480917 | 0,35042954 | 0,61012281 |
| 255783_at | INAFM1 | 0,13508671 | 7,60866558 | 0,96477222 | 0,35044743 | 0,61012281 |
| 5260_at | PHKG1 | -0,1029142 | 4,85702866 | -0,96473576 | 0,35046508 | 0,61012281 |
| 2308_at | FOXO1 | 0,2378498 | 7,29072881 | 0,96463806 | 0,3505124 | 0,61015004 |
| 353324_at | SPATA12 | -0,09536928 | 4,25014841 | -0,96452233 | 0,35056845 | 0,61019247 |
| 100505877_a | UBE2E2-AS1 | -0,12355636 | 3,37657354 | -0,96395959 | 0,3508411 | 0,61056479 |
| 2695_at | GIP | -0,10933441 | 3,89076829 | -0,96386715 | 0,35088591 | 0,61056479 |
| 54476_at | RNF216 | 0,06966789 | 6,55739728 | 0,96386076 | 0,350889 | 0,61056479 |
| 36_at | ACADSB | 0,15136395 | 6,28001984 | 0,96381432 | 0,35091151 | 0,61056479 |
| 5047_at | PAEP | -0,08568636 | 4,30229941 | -0,96375383 | 0,35094084 | 0,61056479 |
| 219970_at | GLYATL2 | -0,10424373 | 3,73417056 | -0,96368797 | 0,35097276 | 0,61056519 |
| 85352_at | KIAA1644 | -0,08176844 | 3,8886563 | -0,96359698 | 0,35101687 | 0,61058678 |
| 10439_at | OLFM1 | -0,09097359 | 4,54661187 | -0,96288737 | 0,35136102 | 0,61111866 |
| 79969_at | ATAT1 | 0,1085018 | 6,24009676 | 0,96269006 | 0,35145675 | 0,61111866 |
| 9019_at | MPZL1 | -0,08778637 | 8,70222453 | -0,96268725 | 0,35145812 | 0,61111866 |
| 2811_at | GP1BA | -0,07769435 | 5,28630985 | -0,96253416 | 0,35153241 | 0,61111866 |
| 81849_at | ST6GALNAC5 | 0,14355109 | 3,8317926 | 0,96251702 | 0,35154072 | 0,61111866 |
| 6239_at | RREB1 | 0,11257611 | 6,86544941 | 0,96246483 | 0,35156605 | 0,61111866 |
| 100996694_a | LOC10099669 | -0,11815068 | 3,86169765 | -0,96244525 | 0,35157556 | 0,61111866 |
| 245711_at | SPDYA | 0,07829779 | 3,13467707 | 0,96244342 | 0,35157644 | 0,61111866 |
| 199699_at | DAND5 | -0,16567547 | 4,88022438 | -0,96224121 | 0,3516746 | 0,61123412 |
| 6051_at | RNPEP | -0,23261918 | 9,75825118 | -0,96188566 | 0,35184723 | 0,61147899 |
| 147463_at | ANKRD29 | 0,12810482 | 3,38839384 | 0,9612224 | 0,35216942 | 0,61198371 |
| 5453_at | POU3F1 | -0,14406599 | 5,81483058 | -0,96108696 | 0,35223524 | 0,61204288 |
| 56288_at | PARD3 | 0,15858072 | 7,04063233 | 0,96101096 | 0,35227217 | 0,61205185 |
| 196415_at | C12orf77 | -0,09661205 | 3,19691024 | -0,96069928 | 0,35242368 | 0,61225986 |
| 92521_at | SPECC1 | 0,10148852 | 6,43863721 | 0,9605653 | 0,35248882 | 0,61231781 |
| 869_at | CBLN1 | -0,12210654 | 4,13022124 | -0,96048665 | 0,35252707 | 0,61232903 |
| 391356_at | PTRHD1 | 0,15602985 | 8,81137354 | 0,96029231 | 0,35262158 | 0,61243796 |
| 100506779_a | BZRAP1-AS1 | -0,10435056 | 3,75953491 | -0,96013699 | 0,35269712 | 0,61249176 |
| 2904_at | GRIN2B | -0,08468534 | 3,77123152 | -0,96007823 | 0,35272571 | 0,61249176 |
| 85371_at | KIAA1656 | -0,09037884 | 4,37216462 | -0,96003253 | 0,35274794 | 0,61249176 |

| | | | | | | |
|---|---|---|---|---|---|---|
| 255022_at | CALHM1 | -0,13245717 | 4,72914901 | -0,95987906 | 0,3528226 | 0,61250724 |
| 56659_at | KCNK13 | -0,0948319 | 4,53248578 | -0,95982966 | 0,35284664 | 0,61250724 |
| 222029_at | DKFZp434L19 | 0,09896458 | 3,14067279 | 0,95978386 | 0,35286893 | 0,61250724 |
| 101929265_a | LINC01428 | -0,08869291 | 3,35190071 | -0,9597528 | 0,35288404 | 0,61250724 |
| 84514_at | GHDC | 0,31710837 | 6,07034919 | 0,95953429 | 0,35299039 | 0,61263662 |
| 257313_at | UTS2B | 0,09305202 | 3,25635584 | 0,95932938 | 0,35309013 | 0,61275452 |
| 6150_at | MRPL23 | -0,24658168 | 8,42994725 | -0,95926106 | 0,35312339 | 0,61275704 |
| 83547_at | RILP | 0,11489504 | 6,89788771 | 0,95917709 | 0,35316427 | 0,61277278 |
| 269_at | AMHR2 | -0,09261297 | 5,0170112 | -0,95903701 | 0,35323248 | 0,61283592 |
| 60485_at | SAV1 | 0,13167992 | 7,77346904 | 0,9587362 | 0,35337898 | 0,61298187 |
| 57624_at | NYAP2 | 0,08215285 | 3,49581303 | 0,9587336 | 0,35338024 | 0,61298187 |
| 101926987_a | DLGAP4-AS1 | -0,09320853 | 4,37302563 | -0,9585289 | 0,35347996 | 0,61307978 |
| 4843_at | NOS2 | -0,14840056 | 5,30677553 | -0,95848707 | 0,35350034 | 0,61307978 |
| 953_at | ENTPD1 | -0,04670148 | 3,51629723 | -0,95839373 | 0,35354582 | 0,61310345 |
| 285555_at | STPG2 | -0,11666476 | 3,8529255 | -0,95812647 | 0,35367606 | 0,61323311 |
| 51531_at | C9orf156 | 0,12428712 | 6,72075781 | 0,95806707 | 0,35370501 | 0,61323311 |
| 3786_at | KCNQ3 | -0,10925449 | 4,52157669 | -0,95804433 | 0,35371609 | 0,61323311 |
| 5799_at | PTPRN2 | -0,09664649 | 4,56708573 | -0,95795249 | 0,35376086 | 0,61324437 |
| 80146_at | UXS1 | -0,16137394 | 9,00010478 | -0,95790039 | 0,35378626 | 0,61324437 |
| 117532_at | TMC2 | -0,08979673 | 5,00720406 | -0,9577785 | 0,35384568 | 0,61329219 |
| 1607_at | DGKB | -0,0593318 | 3,03032622 | -0,95764562 | 0,35391047 | 0,61332989 |
| 10842_at | PPP1R17 | -0,10704458 | 4,31012602 | -0,95760328 | 0,35393112 | 0,61332989 |
| 112812_at | FDX1L | -0,18933456 | 8,10439642 | -0,95753217 | 0,35396579 | 0,61333481 |
| 4157_at | MC1R | 0,22122548 | 5,91487506 | 0,95742712 | 0,35401703 | 0,61333518 |
| 10138_at | YAF2 | -0,10517614 | 5,74166144 | -0,95739042 | 0,35403492 | 0,61333518 |
| 342574_at | KRT27 | -0,11800714 | 3,72585818 | -0,95732464 | 0,35406701 | 0,61333518 |
| 285605_at | DTWD2 | -0,0938999 | 6,6646888 | -0,9572706 | 0,35409337 | 0,61333518 |
| 440900_at | LINC01191 | -0,10933403 | 3,95214681 | -0,95685059 | 0,35429828 | 0,61358014 |
| 55824_at | PAG1 | -0,44366472 | 7,12491856 | -0,95685016 | 0,35429849 | 0,61358014 |
| 5564_at | PRKAB1 | 0,13425247 | 7,91060402 | 0,95650845 | 0,35446527 | 0,61381377 |
| 57540_at | PTCHD2 | -0,0939382 | 5,83403336 | -0,95641958 | 0,35450865 | 0,61383372 |
| 116362_at | RBP7 | -0,13389576 | 4,23341328 | -0,95603652 | 0,35469568 | 0,61408667 |
| 23563_at | CHST5 | -0,06704378 | 5,65828678 | -0,95596884 | 0,35472874 | 0,61408667 |
| 83700_at | JAM3 | -0,0678046 | 3,91928615 | -0,95583893 | 0,35479219 | 0,61408667 |
| 101928616_a | LINC00354 | -0,08694787 | 3,25659449 | -0,95582489 | 0,35479905 | 0,61408667 |
| 84734_at | FAM167B | -0,122037 | 4,97966024 | -0,95579401 | 0,35481413 | 0,61408667 |
| 4012_at | LNPEP | -0,18989178 | 7,35046329 | -0,95568667 | 0,35486657 | 0,61412224 |
| 50832_at | TAS2R4 | -0,10308966 | 3,56808916 | -0,95556408 | 0,35492646 | 0,61413614 |
| 10791_at | VAMP5 | 0,07559097 | 6,27808469 | 0,95552104 | 0,35494749 | 0,61413614 |
| 5523_at | PPP2R3A | 0,5631013 | 6,89353411 | 0,95547447 | 0,35497024 | 0,61413614 |
| 23066_at | CAND2 | -0,0905516 | 5,054021 | -0,95536116 | 0,35502561 | 0,61417677 |
| 114783_at | LMTK3 | 0,09821404 | 4,82983536 | 0,95507111 | 0,35516737 | 0,61436684 |
| 11264_at | PXMP4 | 0,16325618 | 6,71867784 | 0,95495181 | 0,35522569 | 0,61441254 |
| 101929372_a | LOC10192937 | -0,10914107 | 4,87688856 | -0,95482033 | 0,35528997 | 0,61446855 |
| 114987_at | WDR31 | 0,12611636 | 5,28654546 | 0,95416593 | 0,35561003 | 0,61493816 |
| 23632_at | CA14 | -0,11300262 | 4,69068254 | -0,95413461 | 0,35562535 | 0,61493816 |
| 51114_at | ZDHHC9 | -0,08404504 | 8,31025444 | -0,95404303 | 0,35567016 | 0,61495051 |
| 100507468_a | LOC10050746 | -0,11570739 | 3,54756549 | -0,95398953 | 0,35569634 | 0,61495051 |
| 374659_at | HDDC3 | -0,09014101 | 8,15549254 | -0,95390729 | 0,35573658 | 0,61496201 |
| 89790_at | SIGLEC10 | -0,08886459 | 4,10698265 | -0,95378336 | 0,35579723 | 0,61496201 |

| | | | | | | |
|---|---|---|---|---|---|---|
| 144501_at | KRT80 | -0,13146958 | 8,23646614 | -0,95378022 | 0,35579877 | 0,61496201 |
| 26001_at | RNF167 | 0,08997781 | 8,16033761 | 0,9534958 | 0,35593799 | 0,61514377 |
| 997_at | CDC34 | -0,04914949 | 8,23891057 | -0,95343492 | 0,35596779 | 0,61514377 |
| 283480_at | FGF14-IT1 | -0,08471514 | 3,63821421 | -0,95310279 | 0,35613042 | 0,61532855 |
| 752014_at | CEMP1 | -0,16809724 | 4,49778825 | -0,95308608 | 0,35613861 | 0,61532855 |
| 283471_at | TMPRSS12 | -0,0821592 | 3,30009839 | -0,95298235 | 0,35618941 | 0,61536113 |
| 54898_at | ELOVL2 | -0,05757786 | 3,53958902 | -0,95285575 | 0,35625142 | 0,61541307 |
| 8854_at | ALDH1A2 | -0,06461877 | 4,49580272 | -0,95249261 | 0,35642934 | 0,61556006 |
| 94081_at | SFXN1 | -0,17620576 | 8,6249545 | -0,95248691 | 0,35643213 | 0,61556006 |
| 101928513_a | CATIP-AS1 | -0,06854597 | 5,75797974 | -0,9524864 | 0,35643238 | 0,61556006 |
| 101928376_a | IL12A-AS1 | -0,06944558 | 3,73183985 | -0,95236619 | 0,35649129 | 0,61560661 |
| 55303_at | GIMAP4 | -0,13616361 | 3,39777253 | -0,95219246 | 0,35657644 | 0,61569846 |
| 84148_at | KAT8 | -0,05761244 | 6,70998776 | -0,95210223 | 0,35662067 | 0,61571964 |
| 7531_at | YWHAE | -0,11228088 | 10,3738123 | -0,9518975 | 0,35672104 | 0,61581768 |
| 5271_at | SERPINB8 | -0,08892976 | 6,9750822 | -0,95185598 | 0,3567414 | 0,61581768 |
| 114825_at | PWWP2A | 0,04855777 | 6,21668194 | 0,95165823 | 0,35683837 | 0,61592988 |
| 160777_at | CCDC60 | -0,14427753 | 5,46306508 | -0,95149768 | 0,35691711 | 0,61598187 |
| 6614_at | SIGLEC1 | -0,09859697 | 5,24363026 | -0,95146643 | 0,35693244 | 0,61598187 |
| 55094_at | GPATCH1 | -0,06580368 | 7,4654321 | -0,95130775 | 0,35701028 | 0,616061 |
| 81027_at | TUBB1 | -0,17879591 | 4,48702912 | -0,95116206 | 0,35708175 | 0,61612915 |
| 101929582_a | LINC01241 | -0,06123141 | 3,26101332 | -0,95087581 | 0,35722222 | 0,61631632 |
| 6545_at | SLC7A4 | -0,10082877 | 5,21910273 | -0,95077396 | 0,35727221 | 0,61634736 |
| 6781_at | STC1 | -0,06829718 | 3,94976761 | -0,95035389 | 0,35747844 | 0,61663735 |
| 283518_at | KCNRG | -0,23976715 | 3,94351905 | -0,95027847 | 0,35751547 | 0,61663735 |
| 29929_at | ALG6 | -0,07879601 | 8,12412642 | -0,95023597 | 0,35753634 | 0,61663735 |
| 2762_at | GMDS | 0,10338411 | 8,03741602 | 0,94995355 | 0,35767505 | 0,61681679 |
| 5169_at | ENPP3 | -0,07808851 | 3,25077669 | -0,949876 | 0,35771315 | 0,61681679 |
| 8464_at | SUPT3H | -0,06820877 | 6,47782261 | -0,94982858 | 0,35773645 | 0,61681679 |
| 388381_at | C17orf98 | -0,1219414 | 3,94850239 | -0,94941689 | 0,35793874 | 0,61701746 |
| 23013_at | SPEN | 0,12374034 | 8,75633811 | 0,94941547 | 0,35793943 | 0,61701746 |
| 5825_at | ABCD3 | -0,12942 | 8,59678454 | -0,94937413 | 0,35795975 | 0,61701746 |
| 64084_at | CLSTN2 | -0,11357234 | 3,93015952 | -0,94928072 | 0,35800566 | 0,61701746 |
| 284451_at | ODF3L2 | -0,13962872 | 5,8445307 | -0,94920588 | 0,35804245 | 0,61701746 |
| 100270804_a | LOC10027080 | 0,22836851 | 4,5864346 | 0,94917509 | 0,35805759 | 0,61701746 |
| 202018_at | TAPT1 | -0,17700168 | 7,31975609 | -0,94909836 | 0,35809531 | 0,61701746 |
| 6375_at | XCL1 | 0,09894087 | 3,90735275 | 0,94907035 | 0,35810908 | 0,61701746 |
| 283045_at | LOC283045 | -0,08065734 | 4,8248721 | -0,9488009 | 0,35824157 | 0,61714408 |
| 1718_at | DHCR24 | -0,08102254 | 11,1910313 | -0,94879058 | 0,35824664 | 0,61714408 |
| 54987_at | C1orf123 | -0,06393595 | 9,00073935 | -0,9486091 | 0,3583359 | 0,61720557 |
| 4858_at | NOVA2 | -0,08227116 | 4,63390107 | -0,9485877 | 0,35834642 | 0,61720557 |
| 100128788_a | SRRM2-AS1 | 0,09676712 | 5,35170162 | 0,94850195 | 0,3583886 | 0,61722303 |
| 55778_at | ZNF839 | 0,10309594 | 6,19718662 | 0,9482762 | 0,35849966 | 0,6173591 |
| 54535_at | CCHCR1 | -0,08288338 | 6,89017569 | -0,9479555 | 0,35865747 | 0,61757565 |
| 64219_at | PJA1 | 0,12623808 | 8,30314598 | 0,94773193 | 0,35876751 | 0,61770992 |
| 10653_at | SPINT2 | 0,13781735 | 11,0582036 | 0,94753389 | 0,35886501 | 0,6178085 |
| 222389_at | BEND7 | 0,14624937 | 5,58994786 | 0,9474154 | 0,35892335 | 0,6178085 |
| 283131_at | NEAT1 | 0,49903714 | 8,96668953 | 0,94739075 | 0,35893549 | 0,6178085 |
| 84623_at | KIRREL3 | -0,09175548 | 4,51484325 | -0,94729183 | 0,3589842 | 0,6178085 |
| 79058_at | ASPSCR1 | 0,09083383 | 7,64499543 | 0,94728617 | 0,35898698 | 0,6178085 |
| 7016_at | TESK1 | -0,07469047 | 8,39449437 | -0,94722481 | 0,3590172 | 0,6178085 |

| | | | | | | |
|---|---|---|---|---|---|---|
| 10235_at | RASGRP2 | -0,09209826 | 4,44097912 | -0,94696414 | 0,3591456 | 0,61797425 |
| 200403_at | VWA3B | -0,06034877 | 3,43624973 | -0,94680973 | 0,35922167 | 0,61804993 |
| 83473_at | KATNAL2 | 0,09321739 | 4,31582995 | 0,94673113 | 0,3592604 | 0,61806136 |
| 100309464_a | OTX2-AS1 | -0,09961243 | 3,23255369 | -0,9466028 | 0,35932363 | 0,61809902 |
| 143501_at | C11orf40 | -0,1006078 | 3,29368349 | -0,94650216 | 0,35937323 | 0,61809902 |
| 54439_at | RBM27 | -0,19309382 | 8,53661436 | -0,94649136 | 0,35937855 | 0,61809902 |
| 142910_at | LIPJ | -0,04587335 | 2,91519539 | -0,9462954 | 0,35947514 | 0,61820995 |
| 2550_at | GABBR1 | 0,1061613 | 6,40860747 | 0,94607241 | 0,35958507 | 0,6182498 |
| 104355146_a | LINC00396 | 0,08048606 | 3,88934846 | 0,94604807 | 0,35959707 | 0,6182498 |
| 165324_at | UBXN2A | -0,16864646 | 7,2784231 | -0,9459727 | 0,35963424 | 0,6182498 |
| 3950_at | LECT2 | -0,07824375 | 3,46302929 | -0,94591278 | 0,35966378 | 0,6182498 |
| 3964_at | LGALS8 | 0,08293992 | 9,16725341 | 0,94584266 | 0,35969836 | 0,6182498 |
| 55916_at | NXT2 | 0,12221901 | 7,7768545 | 0,94580537 | 0,35971675 | 0,6182498 |
| 144571_at | A2M-AS1 | 0,28401662 | 4,34642372 | 0,94579274 | 0,35972298 | 0,6182498 |
| 440104_at | TMEM198B | 0,12341264 | 6,46904331 | 0,94567271 | 0,35978217 | 0,61829637 |
| 100129195_a | ZSCAN16-AS1 | 0,05421465 | 3,78431203 | 0,94542918 | 0,35990231 | 0,61830736 |
| 100507290_a | ZNF865 | -0,08442515 | 6,73116818 | -0,94538619 | 0,35992351 | 0,61830736 |
| 203_at | AK1 | -0,20973103 | 7,83734616 | -0,94535118 | 0,35994079 | 0,61830736 |
| 8313_at | AXIN2 | -0,07231749 | 4,6537434 | -0,94520663 | 0,36001211 | 0,61830736 |
| 58500_at | ZNF250 | 0,25175481 | 6,01222457 | 0,94518107 | 0,36002472 | 0,61830736 |
| 51686_at | OAZ3 | -0,1190032 | 5,57102557 | -0,94516268 | 0,3600338 | 0,61830736 |
| 2078_at | ERG | -0,05194581 | 3,57455692 | -0,94515778 | 0,36003621 | 0,61830736 |
| 4214_at | MAP3K1 | 0,283726 | 7,41626548 | 0,94513926 | 0,36004536 | 0,61830736 |
| 345557_at | PLCXD3 | 0,21920568 | 5,6937323 | 0,945046 | 0,36009138 | 0,61833127 |
| 355_at | FAS | -0,34724443 | 6,7266842 | -0,94496584 | 0,36013094 | 0,61834409 |
| 54143_at | LINC00308 | -0,07123479 | 3,49521029 | -0,94488968 | 0,36016853 | 0,61835351 |
| 157753_at | TMEM74 | 0,06372104 | 3,37559459 | 0,94437452 | 0,36042287 | 0,61871082 |
| 22926_at | ATF6 | 0,13662866 | 8,30853253 | 0,94431522 | 0,36045216 | 0,61871082 |
| 255167_at | LINC01018 | -0,14466093 | 3,85114308 | -0,9442167 | 0,36050082 | 0,61871082 |
| 8815_at | BANF1 | -0,25832372 | 9,21255292 | -0,94417027 | 0,36052375 | 0,61871082 |
| 1301_at | COL11A1 | 0,06827803 | 3,50930421 | 0,94414294 | 0,36053725 | 0,61871082 |
| 80185_at | TTI2 | -0,27884203 | 7,06873887 | -0,94399713 | 0,36060928 | 0,61877931 |
| 6542_at | SLC7A2 | -0,12242671 | 4,54974435 | -0,94375766 | 0,36072759 | 0,61892719 |
| 4798_at | NFRKB | 0,12104945 | 7,37660212 | 0,94362674 | 0,36079229 | 0,61896205 |
| 5159_at | PDGFRB | -0,12606864 | 5,20993893 | -0,9435865 | 0,36081217 | 0,61896205 |
| 1491_at | CTH | -0,09212651 | 4,47602856 | -0,94347059 | 0,36086946 | 0,6190052 |
| 170589_at | GPHA2 | -0,07640533 | 5,38379312 | -0,94338441 | 0,36091206 | 0,61902315 |
| 63978_at | PRDM14 | -0,10678999 | 4,67093195 | -0,94316222 | 0,3610219 | 0,61915641 |
| 246330_at | PELI3 | -0,0907099 | 7,01300467 | -0,94286816 | 0,3611673 | 0,61935064 |
| 126567_at | C2CD4C | -0,12528614 | 6,18983152 | -0,94264846 | 0,36127596 | 0,61946356 |
| 8735_at | MYH13 | -0,10446279 | 4,17714601 | -0,942605 | 0,36129747 | 0,61946356 |
| 154288_at | KHDC3L | -0,10676902 | 5,23065749 | -0,94234676 | 0,36142522 | 0,61957823 |
| 388533_at | KRTDAP | -0,07474436 | 3,81318899 | -0,94233978 | 0,36142868 | 0,61957823 |
| 132228_at | LSMEM2 | -0,13650655 | 5,81349759 | -0,94225911 | 0,36146859 | 0,61958507 |
| 51237_at | MZB1 | -0,09632143 | 4,85907435 | -0,94214922 | 0,36152297 | 0,61958507 |
| 101928273_a | LOC10192827 | -0,1141993 | 3,61411385 | -0,94205428 | 0,36156996 | 0,61958507 |
| 28584_at | TRBV10-2 | -0,12589214 | 4,12484685 | -0,94204006 | 0,361577 | 0,61958507 |
| 7287_at | TULP1 | -0,06902952 | 4,65616302 | -0,94196537 | 0,36161396 | 0,61958507 |
| 8748_at | ADAM20 | -0,06324426 | 4,16701596 | -0,94194176 | 0,36162565 | 0,61958507 |
| 158326_at | FREM1 | 0,07444667 | 4,15866019 | 0,94181151 | 0,36169012 | 0,61964042 |

| | | | | | | |
|---|---|---|---|---|---|---|
| 3199_at | HOXA2 | 0,10416142 | 5,31551589 | 0,94173344 | 0,36172877 | 0,61965152 |
| 285045_at | LINC00486 | -0,07193882 | 3,27623617 | -0,94094559 | 0,36211896 | 0,62026477 |
| 159989_at | CCDC67 | -0,04803779 | 2,82681973 | -0,94076562 | 0,36220814 | 0,62036236 |
| 51338_at | MS4A4A | -0,07713046 | 2,97997174 | -0,94055031 | 0,36231484 | 0,62048994 |
| 9370_at | ADIPOQ | -0,12742055 | 3,51541935 | -0,94017275 | 0,36250201 | 0,62075529 |
| 275_at | AMT | 0,16872578 | 6,42629762 | 0,93999548 | 0,36258991 | 0,62085061 |
| 2060_at | EPS15 | 0,09901372 | 8,51324403 | 0,93988605 | 0,36264418 | 0,62088834 |
| 27124_at | INPP5J | 0,06814812 | 6,05953711 | 0,93970556 | 0,36273369 | 0,62098642 |
| 9211_at | LGI1 | -0,05210054 | 3,10990351 | -0,93959067 | 0,36279069 | 0,62100445 |
| 54991_at | C1orf159 | -0,09682781 | 6,55443536 | -0,93955436 | 0,3628087 | 0,62100445 |
| 1038_at | CDR1 | -0,05683925 | 3,2355617 | -0,93939672 | 0,36288691 | 0,62108313 |
| 140823_at | ROMO1 | -0,12354566 | 10,3898957 | -0,93881898 | 0,36317365 | 0,62148568 |
| 139628_at | FOXR2 | -0,06090342 | 3,12279197 | -0,93877043 | 0,36319775 | 0,62148568 |
| 3092_at | HIP1 | -0,24344242 | 5,63048702 | -0,93865533 | 0,3632549 | 0,62148568 |
| 26523_at | AGO1 | 0,12326844 | 7,1409085 | 0,93849893 | 0,36333256 | 0,62148568 |
| 57622_at | LRFN1 | -0,10819001 | 6,10676316 | -0,93848022 | 0,36334186 | 0,62148568 |
| 5274_at | SERPINI1 | 0,64494864 | 5,13178473 | 0,93842917 | 0,36336721 | 0,62148568 |
| 5126_at | PCSK2 | -0,06543114 | 3,89123482 | -0,93841233 | 0,36337557 | 0,62148568 |
| 801_at | CALM1 | -0,09884152 | 10,5276247 | -0,93840296 | 0,36338022 | 0,62148568 |
| 2074_at | ERCC6 | 0,15288007 | 6,36057611 | 0,93827556 | 0,3634435 | 0,62153872 |
| 125150_at | ZSWIM7 | 0,31548319 | 6,78505357 | 0,93798856 | 0,36358607 | 0,62172734 |
| 29796_at | UQCR10 | -0,05298612 | 9,70759501 | -0,93783841 | 0,36366068 | 0,62173184 |
| 51248_at | PDZD11 | -0,12365747 | 8,63781397 | -0,93780053 | 0,3636795 | 0,62173184 |
| 57408_at | LRTM1 | -0,07131175 | 3,89331553 | -0,93778839 | 0,36368553 | 0,62173184 |
| 2917_at | GRM7 | -0,0645073 | 3,42860965 | -0,93764893 | 0,36375484 | 0,62179514 |
| 9459_at | ARHGEF6 | 0,12118612 | 5,72914252 | 0,93755481 | 0,36380162 | 0,62181993 |
| 54716_at | SLC6A20 | -0,09930933 | 4,50965727 | -0,93745929 | 0,3638491 | 0,6218459 |
| 101928276_a | LINC01227 | 0,06202715 | 3,23935831 | 0,93729366 | 0,36393144 | 0,62193145 |
| 51573_at | GDE1 | 0,07485496 | 10,1340094 | 0,93709336 | 0,36403103 | 0,6219723 |
| 285800_at | PRR18 | -0,13471549 | 5,49383655 | -0,93707859 | 0,36403837 | 0,6219723 |
| 55051_at | NRDE2 | 0,06296846 | 6,1176409 | 0,93701577 | 0,36406961 | 0,6219723 |
| 5288_at | PIK3C2G | -0,10687558 | 4,10572792 | -0,93694875 | 0,36410294 | 0,6219723 |
| 148213_at | ZNF681 | 0,07486669 | 5,30899505 | 0,93692091 | 0,36411678 | 0,6219723 |
| 201294_at | UNC13D | 0,23840637 | 7,15492175 | 0,93671586 | 0,36421877 | 0,62199847 |
| 60684_at | TRAPPC11 | 0,12504928 | 7,39106663 | 0,9366853 | 0,36423398 | 0,62199847 |
| 79145_at | CHCHD7 | -0,06700725 | 7,84441328 | -0,93667435 | 0,36423942 | 0,62199847 |
| 441072_at | FLJ31104 | -0,09935023 | 3,9416815 | -0,93663044 | 0,36426127 | 0,62199847 |
| 253152_at | EPHX4 | -0,22114615 | 4,68124674 | -0,93618465 | 0,36448308 | 0,62232207 |
| 255027_at | MPV17L | -0,15741864 | 7,09483116 | -0,93596009 | 0,36459485 | 0,62240124 |
| 1821_at | DRP2 | -0,07416963 | 4,29177463 | -0,93591765 | 0,36461598 | 0,62240124 |
| 3746_at | KCNC1 | -0,07912319 | 4,14150957 | -0,93589676 | 0,36462638 | 0,62240124 |
| 150365_at | MEI1 | -0,09498419 | 4,68944661 | -0,93559455 | 0,36477685 | 0,62249066 |
| 100507634_a | LOC10050763 | -0,06522532 | 4,60210119 | -0,93554627 | 0,36480089 | 0,62249066 |
| 8531_at | YBX3 | -0,09257782 | 10,9297857 | -0,9355396 | 0,36480421 | 0,62249066 |
| 101928201_a | LOC10192820 | -0,07145221 | 3,72792499 | -0,9354695 | 0,36483912 | 0,62249066 |
| 23042_at | PDXDC1 | 0,10109354 | 9,78201784 | 0,93546704 | 0,36484035 | 0,62249066 |
| 3891_at | KRT85 | -0,08956232 | 5,33581984 | -0,9352495 | 0,3649487 | 0,62262038 |
| 375056_at | MIA3 | 0,17253304 | 6,77486036 | 0,93469546 | 0,36522475 | 0,62303616 |
| 338758_at | LINC00936 | 0,11589305 | 6,3529123 | 0,9343407 | 0,36540158 | 0,62328262 |
| 285224_at | DNAJB8-AS1 | 0,08171481 | 3,65310507 | 0,93418646 | 0,36547848 | 0,6233586 |

| | | | | | | |
|---|---|---|---|---|---|---|
| 170825_at | GSX2 | -0,10911982 | 5,76639121 | -0,93400219 | 0,36557037 | 0,62342178 |
| 221393_at | ADGRF4 | -0,31003869 | 6,86094561 | -0,93396943 | 0,36558671 | 0,62342178 |
| 138009_at | DCAF4L2 | 0,05427368 | 3,51039932 | 0,93391748 | 0,36561262 | 0,62342178 |
| 51678_at | MPP6 | -0,3143639 | 5,37712183 | -0,93355856 | 0,36579166 | 0,62362665 |
| 2515_at | ADAM2 | -0,10056042 | 3,03049749 | -0,93354683 | 0,36579751 | 0,62362665 |
| 24145_at | PANX1 | 0,09502987 | 8,00936816 | 0,93339142 | 0,36587506 | 0,62370365 |
| 101929512_a | LOC10192951 | -0,08923664 | 3,26254149 | -0,93329071 | 0,36592531 | 0,62373412 |
| 5190_at | PEX6 | 0,07321093 | 7,25111497 | 0,93302254 | 0,36605916 | 0,62389869 |
| 100533106_a | ZHX1-C8orf76 | -0,08222757 | 5,40778163 | -0,93293382 | 0,36610345 | 0,62389869 |
| 2926_at | GRSF1 | -0,1957484 | 8,58990193 | -0,93287747 | 0,36613158 | 0,62389869 |
| 93953_at | ACRC | 0,13298428 | 4,75464896 | 0,93283774 | 0,36615141 | 0,62389869 |
| 11316_at | COPE | 0,09967226 | 9,91418577 | 0,93266788 | 0,36623623 | 0,62398801 |
| 89795_at | NAV3 | -0,04913589 | 3,44054612 | -0,9324852 | 0,36632745 | 0,62408823 |
| 286097_at | MICU3 | 0,06984486 | 2,64310697 | 0,9323708 | 0,36638459 | 0,62413038 |
| 56943_at | ENY2 | 0,10396551 | 7,68577735 | 0,93213954 | 0,36650011 | 0,62422887 |
| 197196_at | LINC00311 | -0,10948329 | 4,16038766 | -0,93212531 | 0,36650722 | 0,62422887 |
| 169044_at | COL22A1 | -0,09841529 | 4,65716452 | -0,93203247 | 0,3665536 | 0,62425268 |
| 286053_at | NSMCE2 | -0,09790983 | 6,96604333 | -0,931825 | 0,36665727 | 0,62437402 |
| 221303_at | FAM162B | -0,08826414 | 4,63511906 | -0,93167058 | 0,36673444 | 0,6244199 |
| 92949_at | ADAMTSL1 | -0,07417025 | 3,94798391 | -0,93160102 | 0,36676921 | 0,6244199 |
| 2542_at | SLC37A4 | -0,05000818 | 7,9445524 | -0,93152243 | 0,36680849 | 0,6244199 |
| 283692_at | CPEB1-AS1 | -0,09380351 | 4,05547261 | -0,93151167 | 0,36681387 | 0,6244199 |
| 83694_at | RPS6KL1 | -0,11745947 | 5,32323144 | -0,93141116 | 0,36686412 | 0,62445026 |
| 81888_at | HYI | 0,14564311 | 7,03608932 | 0,93133391 | 0,36690274 | 0,6244539 |
| 101927653_a | LOC10192765 | -0,06143462 | 3,78281294 | -0,9312772 | 0,36693109 | 0,6244539 |
| 257101_at | ZNF683 | -0,10047821 | 5,89971358 | -0,93115583 | 0,36699178 | 0,624502 |
| 84569_at | LYZL1 | -0,07496 | 3,11318748 | -0,93091983 | 0,3671098 | 0,62460041 |
| 55023_at | PHIP | -0,12504371 | 8,35245356 | -0,93091052 | 0,36711446 | 0,62460041 |
| 55672_at | NBPF1 | 0,18103084 | 8,13798075 | 0,93060581 | 0,36726689 | 0,62478348 |
| 51160_at | VPS28 | 0,27289935 | 9,88587998 | 0,93054722 | 0,3672962 | 0,62478348 |
| 80763_at | SPX | -0,12516066 | 3,85189804 | -0,93050093 | 0,36731936 | 0,62478348 |
| 116372_at | LYPD1 | 0,77835745 | 7,34573745 | 0,93015057 | 0,36749469 | 0,62497633 |
| 941_at | CD80 | -0,08607817 | 3,70224098 | -0,93009261 | 0,3675237 | 0,62497633 |
| 26471_at | NUPR1 | 0,09992945 | 4,76088302 | 0,93007988 | 0,36753008 | 0,62497633 |
| 10168_at | ZNF197 | 0,09865221 | 5,26356599 | 0,93000806 | 0,36756603 | 0,62498229 |
| 1240_at | CMKLR1 | -0,08064255 | 4,36076909 | -0,92988148 | 0,36762939 | 0,62503486 |
| 11131_at | CAPN11 | -0,09153735 | 4,39104176 | -0,92961611 | 0,36776226 | 0,62513708 |
| 84978_at | FRMD5 | 0,07513346 | 5,24692124 | 0,92954406 | 0,36779834 | 0,62513708 |
| 9437_at | NCR1 | -0,10004738 | 4,91255542 | -0,92953955 | 0,3678006 | 0,62513708 |
| 100506571_a | LOC10050657 | -0,10372434 | 4,83878582 | -0,92950217 | 0,36781932 | 0,62513708 |
| 101927934_a | LOC10192793 | -0,15185617 | 4,80795707 | -0,92942571 | 0,36785762 | 0,625147 |
| 100507436_a | MICA | -0,38837643 | 8,65512619 | -0,92919496 | 0,36797321 | 0,62528828 |
| 3036_at | HAS1 | -0,08736562 | 5,13455033 | -0,92897929 | 0,36808126 | 0,62541671 |
| 340061_at | TMEM173 | 0,10718077 | 7,49209092 | 0,9288887 | 0,36812665 | 0,62543867 |
| 64326_at | RFWD2 | 0,11301532 | 8,09374597 | 0,92870885 | 0,36821678 | 0,62553664 |
| 221079_at | ARL5B | 0,12643556 | 8,32546931 | 0,92788788 | 0,36862841 | 0,62611791 |
| 84233_at | TMEM126A | -0,13504891 | 9,11396473 | -0,92788504 | 0,36862983 | 0,62611791 |
| 59336_at | PRDM13 | -0,08811336 | 4,37863362 | -0,92780742 | 0,36866877 | 0,62611791 |
| 285016_at | FAM150B | 0,08219907 | 4,60768083 | 0,92776717 | 0,36868896 | 0,62611791 |
| 152002_at | XXYLT1 | -0,17519242 | 7,52759015 | -0,9275044 | 0,36882079 | 0,62628657 |

| | | | | | | |
|---|---|---|---|---|---|---|
| 140690_at | CTCFL | -0,06826317 | 3,23149507 | -0,92736401 | 0,36889124 | 0,62630577 |
| 54520_at | CCDC93 | -0,07918726 | 7,15211623 | -0,92735228 | 0,36889712 | 0,62630577 |
| 7389_at | UROD | 0,05905114 | 8,68872864 | 0,9272622 | 0,36894233 | 0,62630724 |
| 813_at | CALU | -0,16078732 | 9,2472501 | -0,92722098 | 0,36896302 | 0,62630724 |
| 10086_at | HHLA1 | -0,08505266 | 5,08080521 | -0,9269163 | 0,36911596 | 0,62633926 |
| 26575_at | RGS17 | -0,5074568 | 4,95323812 | -0,92691448 | 0,36911687 | 0,62633926 |
| 64210_at | MMS19 | -0,07482999 | 8,26020128 | -0,9268471 | 0,3691507 | 0,62633926 |
| 3638_at | INSIG1 | -0,24088267 | 9,25596032 | -0,92671393 | 0,36921756 | 0,62633926 |
| 22898_at | DENND3 | 0,17390668 | 6,42318409 | 0,92668972 | 0,36922972 | 0,62633926 |
| 55741_at | EDEM2 | 0,19062425 | 7,87920792 | 0,92660318 | 0,36927318 | 0,62633926 |
| 55231_at | CCDC87 | 0,07246223 | 5,47372049 | 0,92653121 | 0,36930932 | 0,62633926 |
| 54536_at | EXOC6 | -0,1070793 | 6,87102536 | -0,92651187 | 0,36931903 | 0,62633926 |
| 6591_at | SNAI2 | 0,09970282 | 10,8624234 | 0,92648372 | 0,36933317 | 0,62633926 |
| 55022_at | PID1 | -0,08764345 | 3,90687127 | -0,92645534 | 0,36934742 | 0,62633926 |
| 79621_at | RNASEH2B | -0,31085502 | 4,79235099 | -0,9264136 | 0,36936839 | 0,62633926 |
| 25876_at | SPEF1 | 0,09626974 | 5,10920359 | 0,92640629 | 0,36937206 | 0,62633926 |
| 338773_at | TMEM119 | -0,07791749 | 4,14945015 | -0,92632445 | 0,36941317 | 0,62634018 |
| 22829_at | NLGN4Y | 0,26241803 | 4,42950924 | 0,92627574 | 0,36943763 | 0,62634018 |
| 129790_at | C7orf13 | 0,18050494 | 5,97525194 | 0,92591497 | 0,36961889 | 0,62659234 |
| 9478_at | CABP1 | -0,07084806 | 4,54568341 | -0,92573041 | 0,36971165 | 0,6266309 |
| 51316_at | PLAC8 | 0,27367354 | 8,41097689 | 0,92572515 | 0,36971429 | 0,6266309 |
| 130574_at | LYPD6 | 0,38967354 | 6,46007623 | 0,92567553 | 0,36973923 | 0,6266309 |
| 27190_at | IL17B | -0,11989123 | 4,54938524 | -0,92543808 | 0,36985859 | 0,62677804 |
| 6045_at | RNF2 | 0,10158288 | 7,64656673 | 0,92489114 | 0,37013362 | 0,62718894 |
| 9834_at | KIAA0125 | -0,09527197 | 4,44126169 | -0,92480659 | 0,37017615 | 0,62720354 |
| 84915_at | FAM222A | -0,19729062 | 6,514299 | -0,92464026 | 0,37025982 | 0,62720354 |
| 83696_at | TRAPPC9 | -0,16983582 | 6,25836352 | -0,92461778 | 0,37027113 | 0,62720354 |
| 199786_at | FAM129C | -0,08815802 | 5,171157 | -0,92461512 | 0,37027247 | 0,62720354 |
| 9293_at | GPR52 | -0,12216299 | 4,16389751 | -0,92440324 | 0,37037909 | 0,62727999 |
| 79446_at | WDR25 | 0,07294789 | 6,30270161 | 0,92439599 | 0,37038273 | 0,62727999 |
| 84648_at | LCE3D | -0,11634938 | 4,43657251 | -0,92421845 | 0,37047208 | 0,62735051 |
| 729658_at | LOC729658 | -0,11526092 | 4,16270267 | -0,92418383 | 0,37048951 | 0,62735051 |
| 219348_at | PLAC9 | -0,12306531 | 4,28212041 | -0,92401401 | 0,37057498 | 0,62742809 |
| 255101_at | CCDC108 | -0,09538204 | 4,41878248 | -0,92391535 | 0,37062465 | 0,62742809 |
| 83394_at | PITPNM3 | -0,14269134 | 5,18449358 | -0,92385671 | 0,37065418 | 0,62742809 |
| 100131816_a | UBE2DNL | -0,11116645 | 4,75570145 | -0,92383401 | 0,37066561 | 0,62742809 |
| 51339_at | DACT1 | -0,0716776 | 4,00990026 | -0,92375835 | 0,3707037 | 0,62743743 |
| 50937_at | CDON | -0,11602692 | 5,73372404 | -0,92348213 | 0,3708428 | 0,62761613 |
| 55704_at | CCDC88A | -0,41247053 | 6,39410253 | -0,92341931 | 0,37087444 | 0,62761613 |
| 101928068_a | LOC10192806 | 0,10387598 | 4,28327205 | 0,92323266 | 0,37096846 | 0,62768244 |
| 338328_at | GPIHBP1 | -0,11168896 | 5,4189394 | -0,92321214 | 0,3709788 | 0,62768244 |
| 79792_at | GSDMD | 0,22334272 | 8,12111829 | 0,92293035 | 0,37112077 | 0,62776931 |
| 2178_at | FANCE | -0,08255002 | 7,61158232 | -0,92292054 | 0,37112572 | 0,62776931 |
| 102724511_a | LOC10272451 | -0,10778132 | 3,78908004 | -0,9228291 | 0,3711718 | 0,62776931 |
| 100507300_a | ALKBH3-AS1 | -0,07509824 | 3,6874971 | -0,92281858 | 0,3711771 | 0,62776931 |
| 1636_at | ACE | -0,07718618 | 4,45037752 | -0,92278685 | 0,37119309 | 0,62776931 |
| 9203_at | ZMYM3 | 0,12539448 | 6,86983078 | 0,9227117 | 0,37123097 | 0,62777826 |
| 57215_at | THAP11 | -0,11659506 | 8,83450248 | -0,92257894 | 0,37129788 | 0,62779907 |
| 100128782_a | LINC00476 | 0,10478821 | 4,94918126 | 0,92251592 | 0,37132965 | 0,62779907 |
| 2969_at | GTF2I | 0,12700902 | 8,71251372 | 0,92245382 | 0,37136096 | 0,62779907 |

| | | | | | | |
|---|---|---|---|---|---|---|
| 9201_at | DCLK1 | -0,06557474 | 3,26815367 | -0,92241083 | 0,37138263 | 0,62779907 |
| 100505702_a | LINC01094 | -0,06931435 | 3,22434435 | -0,92231369 | 0,37143161 | 0,62779907 |
| 3736_at | KCNA1 | -0,07187945 | 3,80718423 | -0,92229937 | 0,37143882 | 0,62779907 |
| 23392_at | KIAA0368 | -0,07340588 | 8,36864919 | -0,92206964 | 0,37155467 | 0,6278987 |
| 26330_at | GAPDHS | 0,07796098 | 4,38901119 | 0,9220532 | 0,37156296 | 0,6278987 |
| 374973_at | TEX38 | -0,12327086 | 4,33571247 | -0,92183679 | 0,37167211 | 0,62798458 |
| 392307_at | FAM221B | -0,10284004 | 4,5065893 | -0,92182317 | 0,37167898 | 0,62798458 |
| 339779_at | PRR30 | -0,08329687 | 5,53485988 | -0,92172116 | 0,37173044 | 0,6279917 |
| 85479_at | DNAJC5B | -0,10930935 | 4,01138625 | -0,92167071 | 0,37175589 | 0,6279917 |
| 1002_at | CDH4 | -0,08877626 | 4,65884274 | -0,92162094 | 0,371781 | 0,6279917 |
| 84530_at | SRRM4 | -0,10668526 | 4,44733791 | -0,92151738 | 0,37183325 | 0,62799961 |
| 22806_at | IKZF3 | -0,06031822 | 4,33377862 | -0,92148243 | 0,37185088 | 0,62799961 |
| 3495_at | IGHD | -0,12042329 | 5,88443448 | -0,92125307 | 0,37196663 | 0,62814002 |
| 57381_at | RHOJ | -0,05322009 | 3,50771002 | -0,92108718 | 0,37205036 | 0,62817644 |
| 130367_at | SGPP2 | 0,08081818 | 3,76104345 | 0,92108112 | 0,37205342 | 0,62817644 |
| 54437_at | SEMA5B | -0,09611581 | 6,19989815 | -0,92083992 | 0,37217518 | 0,62832696 |
| 442028_at | LOC442028 | 0,0968479 | 3,79053396 | 0,92070975 | 0,37224091 | 0,62834095 |
| 6492_at | SIM1 | -0,07335077 | 3,69200141 | -0,92060281 | 0,37229491 | 0,62834095 |
| 2833_at | CXCR3 | -0,08728419 | 5,6692218 | -0,92051521 | 0,37233915 | 0,62834095 |
| 4694_at | NDUFA1 | 0,05580842 | 10,3719434 | 0,92050576 | 0,37234392 | 0,62834095 |
| 7130_at | TNFAIP6 | -0,05824393 | 3,48045838 | -0,92050052 | 0,37234657 | 0,62834095 |
| 132243_at | H1FOO | -0,10875041 | 5,62695384 | -0,92008586 | 0,37255604 | 0,62863936 |
| 7464_at | CORO2A | 0,10163788 | 8,22108628 | 0,91993465 | 0,37263244 | 0,62870837 |
| 101928697_a | LOC10192869 | -0,07328667 | 3,25357731 | -0,91987162 | 0,37266429 | 0,62870837 |
| 125336_at | LOXHD1 | -0,09183206 | 4,26655508 | -0,91981115 | 0,37269485 | 0,62870837 |
| 54328_at | GPR173 | -0,09387783 | 4,84333943 | -0,91949422 | 0,37285504 | 0,62892352 |
| 23464_at | GCAT | -0,12213796 | 7,42798565 | -0,91937579 | 0,37291491 | 0,62896943 |
| 201176_at | ARHGAP27 | 0,08275341 | 6,6126982 | 0,91920337 | 0,37300209 | 0,6290614 |
| 140828_at | LINC00261 | -0,09044733 | 3,64565964 | -0,91907878 | 0,37306509 | 0,62911257 |
| 494115_at | RBMXL1 | 0,2403999 | 7,49145502 | 0,91887634 | 0,37316748 | 0,62923015 |
| 1289_at | COL5A1 | 0,38370188 | 7,03288663 | 0,91874057 | 0,37323616 | 0,62928991 |
| 51252_at | FAM178B | -0,14941148 | 6,19048619 | -0,91867711 | 0,37326826 | 0,62928991 |
| 55089_at | SLC38A4 | -0,05540114 | 2,90722805 | -0,9183192 | 0,37344936 | 0,62954013 |
| 206338_at | LVRN | -0,07687861 | 3,05014876 | -0,91816483 | 0,37352749 | 0,6295865 |
| 27113_at | BBC3 | 0,07621682 | 6,66616465 | 0,91813569 | 0,37354224 | 0,6295865 |
| 54738_at | FEV | -0,12272419 | 5,14876345 | -0,91789391 | 0,37366463 | 0,62969343 |
| 100874433_a | IQCJ-SCHIP1-A | -0,12663992 | 4,24353858 | -0,91786677 | 0,37367837 | 0,62969343 |
| 6906_at | SERPINA7 | -0,07497883 | 3,31597234 | -0,91781665 | 0,37370374 | 0,62969343 |
| 4880_at | NPPC | -0,12996974 | 5,4717079 | -0,91774448 | 0,37374029 | 0,62969991 |
| 10335_at | MRVI1 | -0,08348236 | 4,88361495 | -0,91764635 | 0,37378998 | 0,62972856 |
| 6549_at | SLC9A2 | 0,15408152 | 4,84168985 | 0,91756166 | 0,37383287 | 0,62973486 |
| 339168_at | TMEM95 | -0,12049607 | 5,63741349 | -0,91750986 | 0,3738591 | 0,62973486 |
| 92312_at | MEX3A | 0,13657784 | 6,65480299 | 0,9174107 | 0,37390932 | 0,62975018 |
| 11243_at | PMF1 | -0,06263364 | 8,82154884 | -0,9173223 | 0,3739541 | 0,62975018 |
| 101927159_a | LOC10192715 | -0,08557156 | 3,78476703 | -0,91725671 | 0,37398732 | 0,62975018 |
| 92270_at | ATP6AP1L | 0,21688702 | 4,95557834 | 0,91719928 | 0,37401642 | 0,62975018 |
| 8419_at | BFSP2 | -0,09941637 | 3,9173756 | -0,91709076 | 0,3740714 | 0,62975018 |
| 140710_at | SOGA1 | -0,09443263 | 6,18055875 | -0,91705616 | 0,37408892 | 0,62975018 |
| 441355_at | MIR2052HG | -0,11524086 | 3,75364936 | -0,91701392 | 0,37411033 | 0,62975018 |
| 114908_at | TMEM123 | 0,12229935 | 11,364028 | 0,91697078 | 0,37413218 | 0,62975018 |

| | | | | | | |
|---|---|---|---|---|---|---|
| 8835_at | SOCS2 | 0,07455111 | 4,01485791 | 0,9169111 | 0,37416243 | 0,62975018 |
| 151011_at | SEPT10 | 0,25509777 | 8,22914373 | 0,91684126 | 0,37419782 | 0,62975472 |
| 283876_at | LINC00921 | -0,0892203 | 4,75363106 | -0,91655864 | 0,37434106 | 0,62989073 |
| 80036_at | TRPM3 | -0,05868616 | 3,90818766 | -0,91651056 | 0,37436543 | 0,62989073 |
| 653240_at | KRTAP4-11 | -0,1127983 | 4,64175312 | -0,91648827 | 0,37437673 | 0,62989073 |
| 101927244_a | LOC10192724 | -0,06080639 | 3,0614358 | -0,91628153 | 0,37448155 | 0,62998492 |
| 54463_at | FAM134B | 0,13086859 | 4,88657661 | 0,91624884 | 0,37449812 | 0,62998492 |
| 50632_at | CALY | -0,08716525 | 5,56333564 | -0,91612825 | 0,37455927 | 0,63003276 |
| 54210_at | TREM1 | -0,1024748 | 3,94036685 | -0,91559321 | 0,37483065 | 0,63039018 |
| 148581_at | UBE2U | -0,06272989 | 2,91558852 | -0,91555315 | 0,37485098 | 0,63039018 |
| 51562_at | MBIP | 0,21431438 | 8,06469684 | 0,91544208 | 0,37490733 | 0,63039018 |
| 30819_at | KCNIP2 | -0,07570482 | 4,32500022 | -0,91539685 | 0,37493029 | 0,63039018 |
| 56751_at | BARHL1 | -0,07885316 | 6,57151536 | -0,91538681 | 0,37493538 | 0,63039018 |
| 84896_at | ATAD1 | -0,05761523 | 7,87603497 | -0,91519244 | 0,37503402 | 0,63045385 |
| 124221_at | PRSS30P | -0,0856747 | 5,14859085 | -0,91516004 | 0,37505047 | 0,63045385 |
| 101752400_a | CAPN10-AS1 | -0,16491045 | 4,44000624 | -0,91509404 | 0,37508397 | 0,63045385 |
| 5257_at | PHKB | 0,12491857 | 8,46435901 | 0,91502612 | 0,37511844 | 0,63045385 |
| 6258_at | RXRG | 0,09707262 | 4,08290103 | 0,91498977 | 0,3751369 | 0,63045385 |
| 8819_at | SAP30 | -0,26223962 | 7,16236478 | -0,91470182 | 0,37528309 | 0,63062598 |
| 54880_at | BCOR | 0,07407446 | 6,72782025 | 0,9145494 | 0,37536049 | 0,63062598 |
| 1390_at | CREM | -0,07093322 | 5,42589086 | -0,91450322 | 0,37538394 | 0,63062598 |
| 57529_at | RGAG1 | -0,0959631 | 4,2856514 | -0,91447939 | 0,37539604 | 0,63062598 |
| 8675_at | STX16 | 0,14293248 | 8,27743451 | 0,91446568 | 0,37540301 | 0,63062598 |
| 4324_at | MMP15 | -0,08158571 | 6,34023945 | -0,91431396 | 0,37548007 | 0,63070043 |
| 5192_at | PEX10 | -0,13918019 | 7,98073472 | -0,91416413 | 0,37555618 | 0,63077327 |
| 55349_at | CHDH | -0,06164737 | 4,7851176 | -0,91405682 | 0,3756107 | 0,63077877 |
| 1588_at | CYP19A1 | -0,09044958 | 3,52802304 | -0,9140229 | 0,37562793 | 0,63077877 |
| 101929181_a | LOC10192918 | -0,11172398 | 3,41380226 | -0,91396433 | 0,37565769 | 0,63077877 |
| 6656_at | SOX1 | -0,08651802 | 4,86869427 | -0,91386293 | 0,37570922 | 0,63081031 |
| 79776_at | ZFHX4 | -0,09978183 | 3,75410254 | -0,91359811 | 0,37584381 | 0,63098128 |
| 128338_at | DRAM2 | 0,12643933 | 8,75717554 | 0,91322325 | 0,37603438 | 0,63116882 |
| 10786_at | SLC17A3 | -0,10080287 | 3,74640162 | -0,91321429 | 0,37603893 | 0,63116882 |
| 342667_at | STAC2 | -0,08292685 | 5,99203083 | -0,91318503 | 0,37605381 | 0,63116882 |
| 10984_at | KCNQ1OT1 | -0,0466199 | 4,71777809 | -0,91263085 | 0,37633567 | 0,63147868 |
| 55015_at | PRPF39 | -0,09859218 | 7,48950261 | -0,91262882 | 0,3763367 | 0,63147868 |
| 112840_at | WDR89 | -0,12502188 | 7,14961316 | -0,91257093 | 0,37636616 | 0,63147868 |
| 9652_at | TTC37 | 0,14746883 | 8,19629604 | 0,912486 | 0,37640937 | 0,63147868 |
| 10652_at | YKT6 | -0,12510135 | 8,97912104 | -0,91247977 | 0,37641254 | 0,63147868 |
| 2848_at | GPR25 | -0,08929289 | 4,82983566 | -0,9124354 | 0,37643512 | 0,63147868 |
| 10869_at | USP19 | 0,0477621 | 7,09996527 | 0,91227381 | 0,37651734 | 0,63156161 |
| 134187_at | POU5F2 | -0,0943481 | 5,19972381 | -0,91205392 | 0,37662926 | 0,63169433 |
| 79002_at | C19orf43 | -0,09809097 | 10,3862264 | -0,9119436 | 0,37668542 | 0,63173352 |
| 101928823_a | LOC10192882 | -0,07379294 | 3,2545996 | -0,91187878 | 0,37671841 | 0,63173386 |
| 101927049_a | PLCE1-AS2 | -0,08566718 | 3,33551355 | -0,91175243 | 0,37678275 | 0,63178674 |
| 23513_at | SCRIB | -0,27975685 | 8,63718603 | -0,9113494 | 0,37698799 | 0,63207587 |
| 7466_at | WFS1 | 0,20869674 | 8,45628168 | 0,91120889 | 0,37705956 | 0,63208886 |
| 81494_at | CFHR5 | -0,08913413 | 2,88793055 | -0,91120534 | 0,37706137 | 0,63208886 |
| 22924_at | MAPRE3 | 0,08186572 | 6,33335435 | 0,91103635 | 0,37714746 | 0,63213649 |
| 56252_at | YLPM1 | -0,04717573 | 8,566678 | -0,91101923 | 0,37715618 | 0,63213649 |
| 101927668_a | LOC10192766 | -0,05818708 | 3,62550183 | -0,91094927 | 0,37719183 | 0,63213649 |

| | | | | | | |
|---|---|---|---|---|---|---|
| 153_at | ADRB1 | 0,12147807 | 5,13033321 | 0,91083597 | 0,37724956 | 0,63213649 |
| 158381_at | ATP8B5P | -0,04831496 | 3,00932145 | -0,91080331 | 0,37726621 | 0,63213649 |
| 120066_at | OR5P3 | -0,08000109 | 3,93536896 | -0,91072318 | 0,37730704 | 0,63213649 |
| 25771_at | TBC1D22A | 0,12202671 | 6,99122435 | 0,91069875 | 0,37731949 | 0,63213649 |
| 8792_at | TNFRSF11A | 0,24069788 | 5,88058375 | 0,91001385 | 0,37766866 | 0,63266644 |
| 148413_at | LOC148413 | 0,09869004 | 7,66445616 | 0,90980564 | 0,37777485 | 0,63273485 |
| 2890_at | GRIA1 | -0,06690701 | 3,39978672 | -0,90974066 | 0,377808 | 0,63273485 |
| 5093_at | PCBP1 | -0,12220594 | 10,8417835 | -0,9097267 | 0,37781512 | 0,63273485 |
| 25996_at | REXO2 | -0,13207612 | 9,07498616 | -0,90967618 | 0,37784089 | 0,63273485 |
| 9725_at | TMEM63A | 0,09379419 | 7,8351066 | 0,90955618 | 0,37790211 | 0,63273856 |
| 101928123_at | LOC10192812 | 0,13675658 | 5,47027565 | 0,90954307 | 0,3779088 | 0,63273856 |
| 129530_at | LYG1 | 0,08615422 | 5,46978538 | 0,90928394 | 0,37804102 | 0,63290249 |
| 100506142_at | LOC10050614 | -0,0916837 | 4,33401474 | -0,90920199 | 0,37808284 | 0,63290249 |
| 114819_at | CROCCP3 | -0,0687138 | 4,92549998 | -0,90915804 | 0,37810527 | 0,63290249 |
| 129049_at | SGSM1 | -0,11119118 | 4,68333397 | -0,9090673 | 0,37815158 | 0,63292501 |
| 4731_at | NDUFV3 | -0,20369699 | 8,45079705 | -0,90893116 | 0,37822108 | 0,63292963 |
| 57451_at | TENM2 | -0,16458102 | 5,16160022 | -0,90884639 | 0,37826435 | 0,63292963 |
| 6857_at | SYT1 | 0,09257833 | 4,86616291 | 0,90880945 | 0,37828321 | 0,63292963 |
| 154043_at | CNKSR3 | 0,21303502 | 6,28051153 | 0,90877116 | 0,37830276 | 0,63292963 |
| 91828_at | EXOC3L4 | -0,07516711 | 5,19919231 | -0,90874007 | 0,37831863 | 0,63292963 |
| 3859_at | KRT12 | -0,07429498 | 3,35212571 | -0,90863151 | 0,37837406 | 0,6329674 |
| 79026_at | AHNAK | -0,0860778 | 7,8482623 | -0,90855722 | 0,37841199 | 0,63297588 |
| 23219_at | FBXO28 | -0,1087371 | 8,01417831 | -0,90848657 | 0,37844808 | 0,63298127 |
| 623_at | BDKRB1 | -0,13893358 | 5,47448755 | -0,90816549 | 0,37861207 | 0,63310691 |
| 164592_at | CCDC116 | -0,11701617 | 5,44424509 | -0,90815699 | 0,37861641 | 0,63310691 |
| 153163_at | MGC32805 | 0,08146929 | 3,0426537 | 0,90810938 | 0,37864073 | 0,63310691 |
| 55884_at | WSB2 | -0,10518152 | 11,484725 | -0,90808212 | 0,37865466 | 0,63310691 |
| 90826_at | PRMT9 | 0,18092889 | 6,85204965 | 0,90786882 | 0,37876364 | 0,63323416 |
| 84277_at | DNAJC30 | 0,08499823 | 7,68991164 | 0,90755979 | 0,37892157 | 0,63344323 |
| 10578_at | GNLY | -0,08138681 | 4,74313054 | -0,90714713 | 0,37913253 | 0,63362785 |
| 10847_at | SRCAP | 0,05969999 | 6,89193483 | 0,90713484 | 0,37913881 | 0,63362785 |
| 58491_at | ZNF71 | 0,11584179 | 6,3898477 | 0,90711878 | 0,37914703 | 0,63362785 |
| 345_at | APOC3 | -0,10831118 | 4,77163535 | -0,90707881 | 0,37916746 | 0,63362785 |
| 54953_at | C1orf27 | -0,10621951 | 6,87332366 | -0,90702206 | 0,37919649 | 0,63362785 |
| 84699_at | CREB3L3 | -0,07537666 | 3,86951404 | -0,90669509 | 0,37936371 | 0,63381832 |
| 10551_at | AGR2 | 0,26527351 | 9,4543166 | 0,90663781 | 0,37939301 | 0,63381832 |
| 56899_at | ANKS1B | -0,0553556 | 3,36333869 | -0,90658559 | 0,37941973 | 0,63381832 |
| 57614_at | KIAA1468 | 0,06880361 | 7,31455995 | 0,90652294 | 0,37945178 | 0,63381832 |
| 348807_at | CCDC37 | -0,07706021 | 4,72319042 | -0,90647758 | 0,37947499 | 0,63381832 |
| 140465_at | MYL6B | 0,21320217 | 8,73785689 | 0,90632472 | 0,3795532 | 0,63389399 |
| 283999_at | TMEM235 | -0,08751987 | 4,62683638 | -0,90604971 | 0,37969394 | 0,63407407 |
| 414201_at | ZNF32-AS3 | -0,06966738 | 3,46268417 | -0,90558947 | 0,37992956 | 0,63441254 |
| 4783_at | NFIL3 | -0,20685396 | 7,30477889 | -0,90501797 | 0,38022227 | 0,63482307 |
| 50515_at | CHST11 | -0,29637695 | 7,16707836 | -0,90492902 | 0,38026784 | 0,63482307 |
| 22801_at | ITGA11 | -0,08599967 | 4,24045825 | -0,9048852 | 0,38029029 | 0,63482307 |
| 84929_at | FIBCD1 | -0,20968737 | 4,87273796 | -0,90485214 | 0,38030723 | 0,63482307 |
| 3001_at | GZMA | -0,07456714 | 3,63508741 | -0,90477759 | 0,38034543 | 0,63483182 |
| 26246_at | OR2L2 | -0,09619625 | 3,58769567 | -0,90469614 | 0,38038717 | 0,63484648 |
| 55248_at | TMEM206 | -0,1317675 | 7,4787298 | -0,90453543 | 0,38046954 | 0,63488467 |
| 56656_at | OR2S2 | -0,10436443 | 4,80698496 | -0,90452288 | 0,38047597 | 0,63488467 |

| | | | | | | |
|---|---|---|---|---|---|---|
| 388456_at | LOC388456 | -0,09184787 | 3,82989335 | -0,90440391 | 0,38053695 | 0,63493142 |
| 5333_at | PLCD1 | 0,30484268 | 7,60621487 | 0,90397238 | 0,3807582 | 0,63519117 |
| 3885_at | KRT34 | -0,07987906 | 3,77422826 | -0,90397164 | 0,38075858 | 0,63519117 |
| 151903_at | CCDC12 | -0,07952854 | 8,40873067 | -0,90342989 | 0,38103646 | 0,63548562 |
| 55914_at | ERBB2IP | 0,19656326 | 8,68724618 | 0,90340022 | 0,38105168 | 0,63548562 |
| 347744_at | C6orf52 | -0,07914172 | 4,38969095 | -0,9033755 | 0,38106437 | 0,63548562 |
| 57010_at | CABP4 | -0,10983883 | 6,32310681 | -0,90334747 | 0,38107874 | 0,63548562 |
| 100129726_a | LINC01126 | -0,09552123 | 4,2668002 | -0,90326215 | 0,38112253 | 0,63548562 |
| 51015_at | ISOC1 | -0,25934856 | 9,60810665 | -0,9032417 | 0,38113302 | 0,63548562 |
| 51527_at | GSKIP | 0,10774137 | 9,26838827 | 0,90305723 | 0,38122769 | 0,63558845 |
| 100874068_a | GRTP1-AS1 | -0,08594894 | 3,86220616 | -0,9029537 | 0,38128083 | 0,63560777 |
| 339456_at | TMEM52 | -0,12755299 | 5,43483356 | -0,90287644 | 0,38132048 | 0,63560777 |
| 2920_at | CXCL2 | 0,67591996 | 5,22469311 | 0,90265071 | 0,38143637 | 0,63560777 |
| 9924_at | PAN2 | 0,30699486 | 7,84184842 | 0,90260625 | 0,38145919 | 0,63560777 |
| 5913_at | RAPSN | 0,08825888 | 4,48790431 | 0,90254417 | 0,38149107 | 0,63560777 |
| 339568_at | LOC339568 | -0,08309393 | 3,40425457 | -0,90251893 | 0,38150403 | 0,63560777 |
| 50511_at | SYCP3 | -0,14373653 | 3,16529536 | -0,90249115 | 0,38151829 | 0,63560777 |
| 4982_at | TNFRSF11B | -0,07232231 | 3,64389299 | -0,90244279 | 0,38154313 | 0,63560777 |
| 9715_at | FAM131B | -0,08479218 | 4,24295152 | -0,90243887 | 0,38154514 | 0,63560777 |
| 245934_at | DEFB121 | -0,06682021 | 3,39367282 | -0,90238801 | 0,38157126 | 0,63560777 |
| 10392_at | NOD1 | 0,08309751 | 5,9123391 | 0,9023277 | 0,38160223 | 0,63560777 |
| 27004_at | TCL6 | -0,09979221 | 3,73927651 | -0,90220632 | 0,38166458 | 0,63561632 |
| 51059_at | FAM135B | -0,07420364 | 4,0318883 | -0,90218921 | 0,38167336 | 0,63561632 |
| 55325_at | UFSP2 | -0,14295848 | 7,50137271 | -0,9015565 | 0,38199844 | 0,63609188 |
| 117854_at | TRIM6 | 0,36431824 | 6,15140636 | 0,90145281 | 0,38205173 | 0,63609188 |
| 7706_at | TRIM25 | 0,20085128 | 8,65919196 | 0,90132461 | 0,38211763 | 0,63609188 |
| 4818_at | NKG7 | -0,10320744 | 5,23861683 | -0,90127503 | 0,38214312 | 0,63609188 |
| 79739_at | TTLL7 | 0,2361611 | 5,7781362 | 0,90126698 | 0,38214726 | 0,63609188 |
| 2052_at | EPHX1 | 0,32384544 | 9,42690846 | 0,90118926 | 0,38218721 | 0,63609188 |
| 163115_at | ZNF781 | 0,06767077 | 3,17850971 | 0,9011679 | 0,38219819 | 0,63609188 |
| 3791_at | KDR | -0,06333448 | 3,6582815 | -0,90110491 | 0,38223058 | 0,63609188 |
| 27296_at | TP53TG5 | -0,10567127 | 4,84687506 | -0,90102964 | 0,38226928 | 0,63609188 |
| 8822_at | FGF17 | -0,10629109 | 5,30955859 | -0,90099102 | 0,38228914 | 0,63609188 |
| 26249_at | KLHL3 | 0,19494753 | 6,4893546 | 0,9007904 | 0,38239231 | 0,63615745 |
| 25827_at | FBXL2 | 0,33680981 | 6,30219795 | 0,90068125 | 0,38244845 | 0,63615745 |
| 55556_at | ENOSF1 | -0,33797066 | 6,97651598 | -0,90066365 | 0,3824575 | 0,63615745 |
| 150709_at | ANKAR | -0,09578031 | 3,26626938 | -0,90065753 | 0,38246065 | 0,63615745 |
| 10320_at | IKZF1 | -0,07384036 | 3,97899547 | -0,90055011 | 0,3825159 | 0,63617066 |
| 3700_at | ITIH4 | -0,09289524 | 5,20991947 | -0,90051368 | 0,38253464 | 0,63617066 |
| 10758_at | TRAF3IP2 | -0,05381398 | 7,44211202 | -0,9003425 | 0,38262271 | 0,63626219 |
| 340602_at | CXorf67 | -0,07038284 | 4,04767981 | -0,90017936 | 0,38270665 | 0,63634685 |
| 5318_at | PKP2 | 0,08951374 | 7,55382683 | 0,90008294 | 0,38275627 | 0,63637442 |
| 63951_at | DMRTA1 | -0,09140379 | 4,64163664 | -0,89990764 | 0,38284649 | 0,63644645 |
| 9308_at | CD83 | -0,28697751 | 7,01570252 | -0,89986292 | 0,38286951 | 0,63644645 |
| 101929504_a | LOC10192950 | -0,10962919 | 3,83808916 | -0,89980619 | 0,38289871 | 0,63644645 |
| 3699_at | ITIH3 | -0,11217523 | 4,34762341 | -0,89972268 | 0,3829417 | 0,63646299 |
| 134829_at | CLVS2 | -0,08192273 | 3,34306269 | -0,89958315 | 0,38301353 | 0,63652745 |
| 10587_at | TXNRD2 | 0,09129955 | 7,6574705 | 0,8992387 | 0,38319091 | 0,63671769 |
| 5322_at | PLA2G5 | -0,07637403 | 3,69402394 | -0,89923248 | 0,38319411 | 0,63671769 |
| 3995_at | FADS3 | 0,22411839 | 7,62498108 | 0,89912475 | 0,3832496 | 0,63674403 |

| | | | | | | |
|---|---|---|---|---|---|---|
| 387707_at | CC2D2B | -0,09397231 | 3,84200201 | -0,89907087 | 0,38327735 | 0,63674403 |
| 711_at | ERC2-IT1 | -0,07436299 | 3,81696345 | -0,89900918 | 0,38330913 | 0,63674403 |
| 10724_at | MGEA5 | 0,16378471 | 8,53706535 | 0,89883694 | 0,38339786 | 0,63682006 |
| 4674_at | NAP1L2 | -0,08672388 | 4,95457351 | -0,89873177 | 0,38345205 | 0,63682006 |
| 79171_at | RBM42 | -0,12951664 | 8,73537483 | -0,89872784 | 0,38345408 | 0,63682006 |
| 51552_at | RAB14 | 0,05651126 | 9,55051527 | 0,89830016 | 0,38367449 | 0,6370492 |
| 101929765_a | LINC01585 | -0,07332416 | 4,46323182 | -0,89827442 | 0,38368776 | 0,6370492 |
| 8301_at | PICALM | -0,1658976 | 9,30573785 | -0,89815632 | 0,38374864 | 0,6370492 |
| 1016_at | CDH18 | -0,09298973 | 4,06386117 | -0,89808008 | 0,38378794 | 0,6370492 |
| 206358_at | SLC36A1 | 0,05921929 | 6,8465308 | 0,89802512 | 0,38381628 | 0,6370492 |
| 730112_at | FAM166B | -0,10993689 | 4,76391062 | -0,89801083 | 0,38382365 | 0,6370492 |
| 116_at | ADCYAP1 | -0,06086348 | 3,95231861 | -0,89796407 | 0,38384776 | 0,6370492 |
| 9722_at | NOS1AP | -0,0756119 | 6,02419949 | -0,89792224 | 0,38386932 | 0,6370492 |
| 90957_at | DHX57 | -0,06702977 | 7,11846372 | -0,89787388 | 0,38389426 | 0,6370492 |
| 5878_at | RAB5C | 0,13719425 | 10,2624486 | 0,89781861 | 0,38392276 | 0,6370492 |
| 7267_at | TTC3 | 0,18152513 | 9,30965449 | 0,89760876 | 0,38403099 | 0,63713103 |
| 730_at | C7 | -0,05958767 | 3,43109549 | -0,89756609 | 0,384053 | 0,63713103 |
| 81688_at | C6orf62 | 0,14070025 | 9,25260598 | 0,89753061 | 0,3840713 | 0,63713103 |
| 28999_at | KLF15 | -0,10475324 | 5,67720261 | -0,89740421 | 0,38413651 | 0,63718432 |
| 57178_at | ZMIZ1 | 0,08445988 | 10,0313517 | 0,8972819 | 0,38419961 | 0,63723412 |
| 713_at | C1QB | -0,09300108 | 4,33573158 | -0,89709587 | 0,3842956 | 0,63733845 |
| 101929657_a | LOC10192965 | 0,11594427 | 4,51496843 | 0,89672584 | 0,38448658 | 0,63760029 |
| 3234_at | HOXD8 | 0,16998373 | 6,30364887 | 0,89625207 | 0,38473119 | 0,63793631 |
| 196074_at | METTL15 | -0,12656475 | 5,91174477 | -0,89615807 | 0,38477974 | 0,63793631 |
| 1136_at | CHRNA3 | -0,06890665 | 4,67750337 | -0,89614099 | 0,38478856 | 0,63793631 |
| 414300_at | LOC414300 | -0,06608868 | 3,29739167 | -0,89559928 | 0,38506841 | 0,63831897 |
| 10595_at | ERN2 | -0,09712448 | 5,89893871 | -0,89549937 | 0,38512004 | 0,63831897 |
| 9313_at | MMP20 | -0,06161818 | 3,18909853 | -0,89545488 | 0,38514303 | 0,63831897 |
| 283693_at | ACTG1P17 | -0,05611571 | 4,31898222 | -0,89543768 | 0,38515192 | 0,63831897 |
| 51082_at | POLR1D | 0,072667 | 9,34770942 | 0,89514776 | 0,38530177 | 0,63847198 |
| 100271832_a | LOC10027183 | -0,0759407 | 3,91312046 | -0,89513082 | 0,38531053 | 0,63847198 |
| 55384_at | MEG3 | -0,07527064 | 4,63651187 | -0,89495132 | 0,38540333 | 0,63851864 |
| 7837_at | PXDN | 0,27357224 | 5,45443781 | 0,89486087 | 0,3854501 | 0,63851864 |
| 7223_at | TRPC4 | -0,04962738 | 3,15692709 | -0,89485632 | 0,38545245 | 0,63851864 |
| 163882_at | CNST | -0,16687615 | 6,25791144 | -0,8948199 | 0,38547128 | 0,63851864 |
| 51545_at | ZNF581 | 0,1174018 | 8,33714064 | 0,8945904 | 0,38558997 | 0,63866033 |
| 80298_at | MTERF2 | 0,15807462 | 6,12340722 | 0,8944231 | 0,3856765 | 0,63874873 |
| 25994_at | HIGD1A | -0,20002085 | 7,72512285 | -0,89397729 | 0,38590716 | 0,63907579 |
| 317648_at | NOP14-AS1 | 0,08608508 | 6,52871579 | 0,89376984 | 0,38601452 | 0,63919864 |
| 101929247_a | LINC01231 | -0,10171574 | 4,86386052 | -0,89344147 | 0,38618451 | 0,639371 |
| 26238_at | LINC01558 | -0,10620512 | 4,055122 | -0,89339491 | 0,38620862 | 0,639371 |
| 497189_at | TIFAB | -0,1037114 | 5,15910755 | -0,8933471 | 0,38623337 | 0,639371 |
| 337876_at | CHSY3 | -0,35097262 | 4,9885655 | -0,89331232 | 0,38625138 | 0,639371 |
| 100506733_a | FAM170B-AS1 | -0,04521406 | 3,12517135 | -0,8932156 | 0,38630147 | 0,63939896 |
| 255330_at | NUP210P1 | -0,06963437 | 4,55368294 | -0,89309163 | 0,38636567 | 0,63945028 |
| 10309_at | CCNO | 0,23329123 | 6,32907262 | 0,89296352 | 0,38643202 | 0,63946779 |
| 166348_at | KBTBD12 | -0,06421408 | 3,28505504 | -0,89294081 | 0,38644378 | 0,63946779 |
| 2195_at | FAT1 | -0,08270451 | 10,3271541 | -0,89287892 | 0,38647584 | 0,63946779 |
| 482_at | ATP1B2 | -0,11237221 | 5,48437528 | -0,89225129 | 0,38680105 | 0,63991487 |
| 646627_at | LYPD8 | -0,08000734 | 4,40987144 | -0,89219194 | 0,38683181 | 0,63991487 |

| | | | | | | |
|---|---|---|---|---|---|---|
| 4925_at | NUCB2 | 0,17391872 | 6,09553095 | 0,89216255 | 0,38684704 | 0,63991487 |
| 4753_at | NELL2 | 0,49195083 | 5,68144705 | 0,89204746 | 0,3869067 | 0,63991487 |
| 80023_at | NRSN2 | -0,06554217 | 5,61102001 | -0,89203699 | 0,38691213 | 0,63991487 |
| 220980_at | TMEM72-AS1 | -0,10871107 | 4,86809122 | -0,89174706 | 0,38706245 | 0,64010851 |
| 25972_at | UNC50 | 0,09294167 | 8,62923708 | 0,89149101 | 0,38719524 | 0,64026242 |
| 333926_at | PPM1J | -0,07940574 | 5,55867721 | -0,89143943 | 0,38722199 | 0,64026242 |
| 283232_at | TMEM80 | 0,16909481 | 7,07399393 | 0,89115485 | 0,38736962 | 0,64043858 |
| 64786_at | TBC1D15 | 0,10707174 | 6,47957206 | 0,89110589 | 0,38739502 | 0,64043858 |
| 23094_at | SIPA1L3 | 0,101506 | 7,23266111 | 0,89102764 | 0,38743562 | 0,64045073 |
| 10087_at | COL4A3BP | 0,10626705 | 7,1930833 | 0,89083321 | 0,38753651 | 0,64056254 |
| 283897_at | C16orf54 | -0,68773839 | 5,48294303 | -0,89024967 | 0,38783943 | 0,64100823 |
| 165631_at | PARP15 | -0,06262436 | 3,03887254 | -0,89017561 | 0,38787788 | 0,64101679 |
| 10846_at | PDE10A | -0,0466772 | 3,62363906 | -0,88993142 | 0,3880047 | 0,64117136 |
| 374786_at | EFCAB5 | -0,05659697 | 3,12382273 | -0,88977344 | 0,38808676 | 0,64121414 |
| 196385_at | DNAH10 | -0,06771067 | 3,97825809 | -0,88975341 | 0,38809717 | 0,64121414 |
| 84634_at | KISS1R | -0,10139784 | 4,80816497 | -0,88962599 | 0,38816336 | 0,64124851 |
| 8786_at | RGS11 | -0,08838155 | 4,4136466 | -0,88958522 | 0,38818454 | 0,64124851 |
| 201456_at | FBXO15 | -0,05739575 | 3,77121943 | -0,88938165 | 0,38829032 | 0,64128385 |
| 905_at | CCNT2 | 0,14791072 | 6,69082611 | 0,88936627 | 0,38829831 | 0,64128385 |
| 5618_at | PRLR | -0,06077511 | 3,19417109 | -0,88935183 | 0,38830581 | 0,64128385 |
| 51129_at | ANGPTL4 | -0,21265686 | 6,04131891 | -0,88893456 | 0,38852269 | 0,64151805 |
| 7018_at | TF | -0,08554732 | 4,03107243 | -0,88891423 | 0,38853326 | 0,64151805 |
| 92689_at | FAM114A1 | 0,24365545 | 8,44267603 | 0,88888678 | 0,38854753 | 0,64151805 |
| 56660_at | KCNK12 | -0,08601954 | 5,09195177 | -0,88877929 | 0,38860341 | 0,64155532 |
| 90865_at | IL33 | -0,07641858 | 3,55026789 | -0,8885096 | 0,38874365 | 0,64173185 |
| 29941_at | PKN3 | 0,1932613 | 5,92275081 | 0,88840383 | 0,38879866 | 0,6417674 |
| 116437_at | LINC01257 | -0,07303076 | 3,37260641 | -0,88834007 | 0,38883182 | 0,6417674 |
| 117285_at | DEFB118 | -0,09868577 | 4,42651717 | -0,88815186 | 0,38892973 | 0,641874 |
| 80168_at | MOGAT2 | 0,10764807 | 5,94889575 | 0,88807355 | 0,38897047 | 0,64188624 |
| 30832_at | ZNF354C | 0,04732643 | 5,06206192 | 0,88798886 | 0,38901453 | 0,64190396 |
| 101927193_a | CHL1-AS1 | -0,07519921 | 3,2725661 | -0,88784515 | 0,3890893 | 0,64197236 |
| 5686_at | PSMA5 | 0,22916619 | 6,9249746 | 0,88776872 | 0,38912908 | 0,64198299 |
| 22905_at | EPN2 | -0,10466509 | 7,56795266 | -0,88749443 | 0,38927183 | 0,64216351 |
| 56033_at | BARX1 | 0,18969433 | 5,11329092 | 0,88727813 | 0,38938444 | 0,6422789 |
| 28641_at | TRAV40 | 0,06141711 | 3,24617668 | 0,88723198 | 0,38940846 | 0,6422789 |
| 94121_at | SYTL4 | -0,20860918 | 7,33121513 | -0,88694412 | 0,38955836 | 0,64244779 |
| 51135_at | IRAK4 | 0,10987014 | 6,60006809 | 0,88684879 | 0,38960801 | 0,64244779 |
| 55254_at | TMEM39A | -0,12794162 | 8,07172969 | -0,8868432 | 0,38961092 | 0,64244779 |
| 1582_at | CYP8B1 | -0,09222237 | 4,31125039 | -0,88676172 | 0,38965336 | 0,64246278 |
| 219738_at | C10orf35 | -0,2480174 | 7,25982196 | -0,88666643 | 0,38970299 | 0,64248962 |
| 2241_at | FER | 0,06381032 | 6,48534604 | 0,88646598 | 0,38980742 | 0,6425815 |
| 3371_at | TNC | -0,15497463 | 4,55629352 | -0,8864314 | 0,38982544 | 0,6425815 |
| 60676_at | PAPPA2 | -0,0937614 | 4,67696687 | -0,88623603 | 0,38992724 | 0,64266679 |
| 9351_at | SLC9A3R2 | 0,10804366 | 5,49303358 | 0,88620405 | 0,38994391 | 0,64266679 |
| 11215_at | AKAP11 | -0,08659508 | 8,55987769 | -0,88610789 | 0,38999402 | 0,64267183 |
| 10020_at | GNE | -0,22270992 | 8,8837078 | -0,88607015 | 0,39001369 | 0,64267183 |
| 56963_at | RGMA | -0,09895104 | 6,29740681 | -0,88593623 | 0,39008349 | 0,64273187 |
| 6948_at | TCN2 | 0,15871818 | 5,72064591 | 0,88574218 | 0,39018465 | 0,64284357 |
| 729993_at | SHISA9 | -0,05759765 | 3,73503526 | -0,8856631 | 0,39022588 | 0,64285651 |
| 8997_at | KALRN | -0,05766411 | 3,90633724 | -0,88535469 | 0,3903867 | 0,64306645 |

| | | | | | | |
|---|---|---|---|---|---|---|
| 79024_at | SMIM2 | -0,05197844 | 3,68715947 | -0,88485297 | 0,39064842 | 0,64344255 |
| 23369_at | PUM2 | 0,10868557 | 9,6854626 | 0,88475615 | 0,39069894 | 0,64347074 |
| 256021_at | LINC01619 | 0,08709496 | 3,11046543 | 0,88395486 | 0,39111722 | 0,64409391 |
| 9154_at | SLC28A1 | -0,1030459 | 4,29491528 | -0,88390321 | 0,39114419 | 0,64409391 |
| 55825_at | PECR | 0,16620259 | 4,67498549 | 0,88359458 | 0,39130538 | 0,64430427 |
| 148254_at | ZNF555 | 0,05071663 | 4,19190864 | 0,88340199 | 0,39140599 | 0,64439169 |
| 91977_at | MYOZ3 | -0,10328371 | 5,20431418 | -0,88333354 | 0,39144175 | 0,64439169 |
| 283933_at | ZNF843 | -0,11807672 | 5,73802887 | -0,88319092 | 0,39151628 | 0,64439169 |
| 6921_at | TCEB1 | -0,12564967 | 8,61535355 | -0,88318576 | 0,39151897 | 0,64439169 |
| 7473_at | WNT3 | 0,08447735 | 5,56970119 | 0,88311831 | 0,39155422 | 0,64439169 |
| 29982_at | NRBF2 | 0,13367645 | 7,52367844 | 0,8831088 | 0,39155919 | 0,64439169 |
| 337970_at | KRTAP19-3 | -0,08298143 | 3,64499897 | -0,88301006 | 0,39161079 | 0,64442155 |
| 283847_at | CCDC79 | -0,0648094 | 3,37727864 | -0,88291227 | 0,3916619 | 0,64445061 |
| 23527_at | ACAP2 | -0,10573203 | 7,84536671 | -0,88274624 | 0,39174869 | 0,64445821 |
| 730249_at | IRG1 | -0,0907341 | 3,17834684 | -0,88271041 | 0,39176742 | 0,64445821 |
| 92745_at | SLC38A5 | 0,09494231 | 5,10906222 | 0,88269362 | 0,3917762 | 0,64445821 |
| 266747_at | RGL4 | -0,10177867 | 4,10097128 | -0,88264742 | 0,39180035 | 0,64445821 |
| 2155_at | F7 | -0,08442115 | 5,75700985 | -0,88249257 | 0,39188131 | 0,64448677 |
| 645191_at | LINGO3 | -0,14864907 | 7,16921201 | -0,88248622 | 0,39188463 | 0,64448677 |
| 339260_at | LOC339260 | -0,11789135 | 4,26532033 | -0,88198747 | 0,39214547 | 0,64480825 |
| 644660_at | RAD21-AS1 | 0,07628477 | 3,92649158 | 0,88198445 | 0,39214705 | 0,64480825 |
| 9379_at | NRXN2 | -0,0687579 | 4,94581454 | -0,88165493 | 0,39231945 | 0,64503667 |
| 643236_at | TMEM72 | -0,06287959 | 4,50268286 | -0,88143269 | 0,39243575 | 0,64512468 |
| 387509_at | GPR153 | -0,14724868 | 7,33072508 | -0,88142465 | 0,39243996 | 0,64512468 |
| 4103_at | MAGEA4 | 0,09593983 | 3,47512323 | 0,88131159 | 0,39249914 | 0,6451669 |
| 9923_at | ZBTB40 | 0,05996673 | 6,97273028 | 0,88120222 | 0,39255638 | 0,64517706 |
| 100506622_a | LINC00540 | -0,08789695 | 3,64755455 | -0,88111266 | 0,39260327 | 0,64517706 |
| 284083_at | C17orf47 | -0,11253893 | 3,86361545 | -0,88108323 | 0,39261868 | 0,64517706 |
| 1158_at | CKM | -0,07664025 | 4,23391903 | -0,88104385 | 0,39263929 | 0,64517706 |
| 9334_at | B4GALT5 | -0,13601148 | 9,71575704 | -0,88071849 | 0,39280966 | 0,64539035 |
| 3915_at | LAMC1 | 0,1938247 | 9,93016367 | 0,880668 | 0,3928361 | 0,64539035 |
| 5997_at | RGS2 | 0,23363171 | 7,51568684 | 0,88050852 | 0,39291963 | 0,64547253 |
| 3635_at | INPP5D | 0,21766858 | 7,43931916 | 0,88034725 | 0,39300411 | 0,64555625 |
| 256126_at | SYCE2 | -0,10068913 | 4,48399717 | -0,88027348 | 0,39304276 | 0,64556469 |
| 128077_at | LIX1L | 0,10884855 | 6,73501707 | 0,88015948 | 0,39310249 | 0,64557677 |
| 2643_at | GCH1 | -0,24533247 | 8,69229962 | -0,88013151 | 0,39311714 | 0,64557677 |
| 81035_at | COLEC12 | -0,07396256 | 3,8775155 | -0,87985378 | 0,39326269 | 0,64564646 |
| 100533184_a | ARHGAP19-SL | -0,15043716 | 3,33371203 | -0,87983563 | 0,3932722 | 0,64564646 |
| 286411_at | LINC00632 | -0,07661375 | 3,23525155 | -0,87982341 | 0,3932786 | 0,64564646 |
| 8600_at | TNFSF11 | -0,14827608 | 3,61784135 | -0,8797947 | 0,39329365 | 0,64564646 |
| 1810_at | DR1 | -0,16635116 | 8,83443838 | -0,879604 | 0,39339361 | 0,64575553 |
| 84570_at | COL25A1 | -0,06547046 | 3,57679458 | -0,87947185 | 0,39346289 | 0,64576621 |
| 440603_at | BCL2L15 | -0,07094676 | 3,77518526 | -0,8794138 | 0,39349333 | 0,64576621 |
| 136853_at | SSC4D | -0,10115713 | 6,05355304 | -0,87936165 | 0,39352067 | 0,64576621 |
| 57130_at | ATP13A1 | -0,0726191 | 8,53905024 | -0,87933583 | 0,39353421 | 0,64576621 |
| 25895_at | METTL21B | 0,12386932 | 7,10440686 | 0,8791876 | 0,39361194 | 0,64581802 |
| 2850_at | GPR27 | -0,08616353 | 6,47341006 | -0,87913318 | 0,39364048 | 0,64581802 |
| 140873_at | C20orf173 | -0,09441424 | 4,15345615 | -0,87905494 | 0,39368151 | 0,64581802 |
| 54414_at | SIAE | 0,24052651 | 7,31291267 | 0,8790199 | 0,39369989 | 0,64581802 |
| 441025_at | LOC441025 | -0,08275663 | 3,78206498 | -0,87894523 | 0,39373906 | 0,64582727 |

| | | | | | | |
|---|---|---|---|---|---|---|
| 155051_at | CRYGN | -0,08951733 | 4,7177649 | -0,87857731 | 0,39393208 | 0,64603548 |
| 100507331_a | ZSWIM8-AS1 | -0,12384443 | 5,80510486 | -0,87857542 | 0,39393307 | 0,64603548 |
| 6487_at | ST3GAL3 | -0,06590193 | 5,52765506 | -0,87838595 | 0,39403249 | 0,64614351 |
| 57585_at | CRAMP1 | 0,16064385 | 6,95153253 | 0,87813146 | 0,39416606 | 0,6462931 |
| 219790_at | RTKN2 | -0,11746224 | 4,78447047 | -0,8780843 | 0,39419081 | 0,6462931 |
| 9653_at | HS2ST1 | 0,09863967 | 8,21623031 | 0,87781528 | 0,39433205 | 0,64642463 |
| 121273_at | C12orf54 | 0,14352916 | 4,12011306 | 0,87780366 | 0,39433815 | 0,64642463 |
| 101928420_a | LINC01490 | 0,05803594 | 3,27318226 | 0,87766077 | 0,39441318 | 0,64649261 |
| 100652856_a | LINC00408 | 0,07638616 | 3,71201084 | 0,87729641 | 0,39460455 | 0,64666444 |
| 25984_at | KRT23 | -0,46887414 | 5,62334776 | -0,87725257 | 0,39462758 | 0,64666444 |
| 2011_at | MARK2 | 0,155856 | 7,40194805 | 0,87717681 | 0,39466738 | 0,64666444 |
| 1297_at | COL9A1 | -0,09114006 | 4,88399236 | -0,87710445 | 0,3947054 | 0,64666444 |
| 58985_at | IL22RA1 | 0,09890169 | 5,89494904 | 0,87709842 | 0,39470857 | 0,64666444 |
| 308_at | ANXA5 | -0,35916909 | 10,7827235 | -0,87707774 | 0,39471943 | 0,64666444 |
| 23154_at | NCDN | -0,07743044 | 7,11804566 | -0,87692856 | 0,39479782 | 0,64673785 |
| 90668_at | LRRC16B | -0,07316773 | 5,08884026 | -0,87679172 | 0,39486973 | 0,64680064 |
| 10150_at | MBNL2 | 0,12356301 | 8,49848199 | 0,8765818 | 0,39498006 | 0,64692134 |
| 126374_at | WTIP | 0,10342784 | 6,69624795 | 0,87652374 | 0,39501058 | 0,64692134 |
| 8528_at | DDO | 0,24990583 | 5,4601707 | 0,87629685 | 0,39512986 | 0,64702796 |
| 2691_at | GHRH | -0,09109009 | 4,33530575 | -0,87627212 | 0,39514286 | 0,64702796 |
| 102724927_a | LOC10272492 | 0,06154724 | 2,9351549 | 0,87614139 | 0,3952116 | 0,64708551 |
| 26508_at | HEYL | -0,07847974 | 5,29806454 | -0,87594131 | 0,39531682 | 0,64720278 |
| 79674_at | VEPH1 | 0,15909988 | 4,96740464 | 0,8750532 | 0,3957841 | 0,64781279 |
| 253044_at | LINGO1-AS1 | -0,08770705 | 4,15458892 | -0,87504983 | 0,39578587 | 0,64781279 |
| 84501_at | SPIRE2 | 0,17581772 | 5,93416765 | 0,8750019 | 0,3958111 | 0,64781279 |
| 3069_at | HDLBP | -0,08691204 | 9,04617052 | -0,87497751 | 0,39582394 | 0,64781279 |
| 6590_at | SLPI | 0,07527974 | 11,8200345 | 0,87480418 | 0,39591518 | 0,64788199 |
| 55007_at | FAM118A | 0,25790648 | 7,42311419 | 0,8747694 | 0,39593349 | 0,64788199 |
| 1414_at | CRYBB1 | -0,10610996 | 5,55345313 | -0,8745791 | 0,39603369 | 0,64799091 |
| 79812_at | MMRN2 | 0,38041472 | 5,22748345 | 0,87447651 | 0,39608772 | 0,64802426 |
| 6360_at | CCL16 | -0,09693188 | 4,01898957 | -0,87440851 | 0,39612352 | 0,64802781 |
| 5620_at | PRM2 | -0,08214266 | 4,31993292 | -0,87428401 | 0,3961891 | 0,64808004 |
| 114788_at | CSMD3 | -0,07238662 | 2,97663227 | -0,87419585 | 0,39623553 | 0,64810096 |
| 90293_at | KLHL13 | -0,11916354 | 6,03118804 | -0,87409596 | 0,39628815 | 0,648132 |
| 3695_at | ITGB7 | -0,17945806 | 5,86234089 | -0,87402407 | 0,39632603 | 0,64813892 |
| 259286_at | TAS2R40 | -0,10239975 | 3,40486888 | -0,87382218 | 0,3964324 | 0,64823512 |
| 340351_at | AGBL3 | -0,07289738 | 2,8759946 | -0,87378468 | 0,39645216 | 0,64823512 |
| 440307_at | TTLL13P | -0,0839118 | 4,21066055 | -0,87334077 | 0,39668612 | 0,64856122 |
| 6122_at | RPL3 | -0,07380578 | 12,2536009 | -0,87325386 | 0,39673194 | 0,64856122 |
| 4864_at | NPC1 | 0,11653747 | 6,07560416 | 0,87319999 | 0,39676034 | 0,64856122 |
| 101929260_a | LOC10192926 | -0,06408012 | 3,67057366 | -0,8731508 | 0,39678628 | 0,64856122 |
| 746_at | TMEM258 | -0,1121483 | 11,0295188 | -0,87303234 | 0,39684873 | 0,64860827 |
| 8846_at | ALKBH1 | -0,09294082 | 7,28287008 | -0,87296646 | 0,39688347 | 0,64861002 |
| 6707_at | SPRR3 | -0,71409312 | 6,71768445 | -0,87284394 | 0,39694809 | 0,64866059 |
| 84938_at | ATG4C | -0,07888058 | 6,60185094 | -0,8727283 | 0,39700908 | 0,64867574 |
| 100506433_a | LINC00648 | 0,08507045 | 3,86032778 | 0,8726515 | 0,39704958 | 0,64867574 |
| 958_at | CD40 | -0,06500007 | 4,05367959 | -0,87259304 | 0,39708042 | 0,64867574 |
| 441009_at | CPEB2-AS1 | -0,05426858 | 3,23428851 | -0,87257098 | 0,39709206 | 0,64867574 |
| 10284_at | SAP18 | 0,04222311 | 8,49233116 | 0,87245629 | 0,39715256 | 0,64871956 |
| 145820_at | LINC00924 | -0,05306131 | 3,12351548 | -0,87220589 | 0,39728468 | 0,64888034 |

| | | | | | | |
|---|---|---|---|---|---|---|
| 1584_at | CYP11B1 | -0,08478454 | 4,24315017 | -0,87209317 | 0,39734416 | 0,64892247 |
| 161835_at | FSIP1 | 0,05335117 | 3,16729647 | 0,87200651 | 0,3973899 | 0,64894215 |
| 1184_at | CLCN5 | -0,08780907 | 4,99766365 | -0,87169156 | 0,39755614 | 0,64905253 |
| 10466_at | COG5 | 0,09463039 | 8,1352409 | 0,87165927 | 0,39757319 | 0,64905253 |
| 135892_at | TRIM50 | -0,09511468 | 4,69005872 | -0,87164778 | 0,39757926 | 0,64905253 |
| 8820_at | HESX1 | 0,06628529 | 3,8383302 | 0,87162314 | 0,39759227 | 0,64905253 |
| 10778_at | ZNF271P | 0,10180085 | 5,15453229 | 0,87148663 | 0,39766434 | 0,64911517 |
| 10205_at | MPZL2 | -0,15285818 | 9,88071873 | -0,8709357 | 0,39795532 | 0,6495351 |
| 5989_at | RFX1 | 0,06748178 | 6,65803973 | 0,87082369 | 0,39801449 | 0,64953735 |
| 9832_at | JAKMIP2 | -0,07075115 | 3,22544984 | -0,87077447 | 0,39804049 | 0,64953735 |
| 28968_at | SLC6A16 | -0,06603427 | 3,96103196 | -0,87074161 | 0,39805785 | 0,64953735 |
| 9780_at | PIEZO1 | 0,08766939 | 9,34624393 | 0,87066589 | 0,39809787 | 0,64954762 |
| 51397_at | COMMD10 | -0,06117387 | 6,58278089 | -0,87054246 | 0,39816309 | 0,64959902 |
| 4782_at | NFIC | 0,07723279 | 7,44415851 | 0,87024244 | 0,39832166 | 0,64980269 |
| 94134_at | ARHGAP12 | 0,22502623 | 7,11580045 | 0,8700479 | 0,3984245 | 0,64991542 |
| 100507392_a | SENCR | -0,12989827 | 4,48697465 | -0,86973491 | 0,39859 | 0,65013033 |
| 6356_at | CCL11 | -0,07700507 | 4,11004135 | -0,86958178 | 0,39867099 | 0,65020737 |
| 1044_at | CDX1 | -0,08818893 | 6,21700441 | -0,86949974 | 0,39871438 | 0,65021432 |
| 8284_at | KDM5D | 0,55030265 | 5,97248284 | 0,86944609 | 0,39874276 | 0,65021432 |
| 80774_at | LIMD2 | 0,11005836 | 5,60802808 | 0,86912997 | 0,39891 | 0,65043197 |
| 305_at | ANXA2P3 | -0,13173834 | 4,69791509 | -0,86896635 | 0,39899658 | 0,6505056 |
| 652972_at | LRRC37A5P | -0,0983621 | 3,89968139 | -0,86891701 | 0,39902269 | 0,6505056 |
| 23051_at | ZHX3 | -0,068806 | 5,92731515 | -0,86877322 | 0,39909879 | 0,6505746 |
| 84911_at | ZNF382 | 0,14097009 | 3,41432769 | 0,86848151 | 0,39925321 | 0,65072486 |
| 6431_at | SRSF6 | -0,2044248 | 9,74644049 | -0,86847147 | 0,39925853 | 0,65072486 |
| 2561_at | GABRB2 | -0,06300926 | 3,24482127 | -0,86828615 | 0,39935665 | 0,65082971 |
| 57619_at | SHROOM3 | -0,09489115 | 5,76864555 | -0,86809665 | 0,399457 | 0,65088862 |
| 9153_at | SLC28A2 | -0,05844852 | 3,66922903 | -0,86806223 | 0,39947523 | 0,65088862 |
| 6185_at | RPN2 | -0,09434952 | 12,0870039 | -0,86797575 | 0,39952103 | 0,65088862 |
| 10637_at | LEFTY1 | -0,07581919 | 4,61625371 | -0,86796268 | 0,39952796 | 0,65088862 |
| 3681_at | ITGAD | -0,11368231 | 4,28234688 | -0,86787065 | 0,39957671 | 0,65091299 |
| 7760_at | ZNF213 | 0,0569462 | 6,90592932 | 0,8676493 | 0,39969397 | 0,65104895 |
| 4897_at | NRCAM | -0,07807375 | 4,15109514 | -0,8675029 | 0,39977154 | 0,65112025 |
| 158801_at | NKAPP1 | 0,09842313 | 5,48594292 | 0,86734274 | 0,39985641 | 0,65117548 |
| 51312_at | SLC25A37 | -0,31246908 | 7,02985217 | -0,86731132 | 0,39987306 | 0,65117548 |
| 200879_at | LIPH | 0,27910885 | 6,61250001 | 0,86721941 | 0,39992177 | 0,65119975 |
| 115352_at | FCRL3 | -0,08029184 | 3,589193 | -0,8671404 | 0,39996365 | 0,6512129 |
| 284611_at | FAM102B | -0,36700135 | 8,13044931 | -0,86692188 | 0,40007949 | 0,65134645 |
| 11149_at | BVES | 0,16026622 | 3,94280724 | 0,86667881 | 0,40020837 | 0,65150121 |
| 196475_at | RMST | -0,06623138 | 3,3513248 | -0,86642172 | 0,40034471 | 0,65166808 |
| 26123_at | TCTN3 | -0,07197391 | 8,71365034 | -0,86634187 | 0,40038706 | 0,65168196 |
| 7472_at | WNT2 | -0,09470868 | 4,26651175 | -0,86621388 | 0,40045495 | 0,65173739 |
| 101927167_a | GATA2-AS1 | 0,14223296 | 7,5322308 | 0,86606471 | 0,40053409 | 0,65181112 |
| 23629_at | BRD7P3 | -0,06649256 | 3,19092374 | -0,86599599 | 0,40057055 | 0,65181538 |
| 79037_at | PVRIG | 0,14713329 | 5,51509218 | 0,86591865 | 0,40061159 | 0,6518271 |
| 11184_at | MAP4K1 | -0,07736913 | 5,41873286 | -0,86576661 | 0,40069227 | 0,65185539 |
| 10972_at | TMED10 | 0,09286554 | 11,5449343 | 0,86570362 | 0,4007257 | 0,65185539 |
| 56947_at | MFF | 0,09197369 | 9,23329345 | 0,86565256 | 0,40075279 | 0,65185539 |
| 101059948_a | LOC10105994 | -0,09849513 | 4,24432002 | -0,86563082 | 0,40076433 | 0,65185539 |
| 3565_at | IL4 | -0,0627328 | 3,4159531 | -0,86542227 | 0,40087503 | 0,65198039 |

| | | | | | | |
|---|---|---|---|---|---|---|
| 101929693_a | RARA-AS1 | 0,1332875 | 5,91428948 | 0,86535122 | 0,40091276 | 0,65198669 |
| 6451_at | SH3BGRL | 0,95513845 | 8,12935326 | 0,86489522 | 0,40115489 | 0,6522797 |
| 101927616_a | LOC10192761 | 0,06519405 | 2,8836946 | 0,86488437 | 0,40116065 | 0,6522797 |
| 400752_at | LINC01144 | -0,1402371 | 5,49068115 | -0,86481763 | 0,40119609 | 0,65228227 |
| 55246_at | CCDC25 | -0,0728834 | 7,74772582 | -0,8647089 | 0,40125385 | 0,65232111 |
| 5671_at | PSG3 | 0,08334335 | 3,4214999 | 0,86464437 | 0,40128813 | 0,65232179 |
| 84288_at | EFCAB2 | 0,1481193 | 5,29740823 | 0,86445754 | 0,40138738 | 0,65242808 |
| 9751_at | SNPH | 0,13561284 | 6,4061856 | 0,86431286 | 0,40146425 | 0,65244909 |
| 10630_at | PDPN | 0,08887582 | 5,87064662 | 0,86426071 | 0,40149197 | 0,65244909 |
| 124930_at | ANKRD13B | -0,10596233 | 6,28249053 | -0,86424198 | 0,40150192 | 0,65244909 |
| 256369_at | LINC00521 | -0,0909857 | 4,35707568 | -0,86406691 | 0,40159496 | 0,65254523 |
| 55508_at | SLC35E3 | -0,13551447 | 8,08828293 | -0,86382234 | 0,40172496 | 0,65270141 |
| 10914_at | PAPOLA | -0,07869707 | 9,64863253 | -0,86373114 | 0,40177345 | 0,65272513 |
| 113277_at | TMEM106A | 0,08817781 | 4,71715634 | 0,86309254 | 0,40211305 | 0,65314626 |
| 54921_at | CHTF8 | -0,08022962 | 10,0393122 | -0,86307718 | 0,40212122 | 0,65314626 |
| 54_at | ACP5 | -0,09570894 | 4,96167841 | -0,86305242 | 0,40213439 | 0,65314626 |
| 9956_at | HS3ST2 | -0,1305795 | 5,42284238 | -0,86298031 | 0,40217275 | 0,6531535 |
| 100506585_a | LOC10050658 | -0,08357067 | 4,88732642 | -0,86288729 | 0,40222224 | 0,65317488 |
| 100506495_a | LIFR-AS1 | -0,06243907 | 2,88336608 | -0,86282809 | 0,40225373 | 0,65317488 |
| 794_at | CALB2 | -0,10499831 | 4,3170262 | -0,86274687 | 0,40229695 | 0,65319 |
| 957_at | ENTPD5 | 0,1257176 | 6,61160219 | 0,86243388 | 0,40246352 | 0,65340537 |
| 51115_at | RMDN1 | -0,171696 | 6,44190698 | -0,86232396 | 0,40252202 | 0,65343962 |
| 10001_at | MED6 | 0,10076135 | 8,56753312 | 0,86226679 | 0,40255245 | 0,65343962 |
| 6814_at | STXBP3 | 0,14283915 | 8,4238116 | 0,86215692 | 0,40261095 | 0,65347949 |
| 84694_at | GJA10 | -0,1231429 | 3,95848964 | -0,86207958 | 0,40265212 | 0,65349127 |
| 3912_at | LAMB1 | 0,17874864 | 9,37692193 | 0,8619268 | 0,40273347 | 0,65355271 |
| 4887_at | NPY2R | -0,08699048 | 4,07657776 | -0,86183729 | 0,40278113 | 0,65355271 |
| 2765_at | GML | -0,1016147 | 3,93209403 | -0,86177098 | 0,40281645 | 0,65355271 |
| 342892_at | ZNF850 | -0,28085962 | 6,07714875 | -0,86173943 | 0,40283325 | 0,65355271 |
| 1339_at | COX6A2 | -0,11450329 | 5,4287748 | -0,86168991 | 0,40285962 | 0,65355271 |
| 8029_at | CUBN | -0,08892166 | 3,59509004 | -0,86159152 | 0,40291203 | 0,65358268 |
| 2303_at | FOXC2 | -0,08109423 | 5,15181908 | -0,86148661 | 0,40296791 | 0,65359571 |
| 11055_at | ZPBP | 0,0810673 | 3,50550211 | 0,86137044 | 0,40302979 | 0,65359571 |
| 81796_at | SLCO5A1 | -0,05180824 | 3,5070282 | -0,86134382 | 0,40304398 | 0,65359571 |
| 401312_at | LOC401312 | -0,07688229 | 3,31730037 | -0,86132167 | 0,40305577 | 0,65359571 |
| 1237_at | CCR8 | -0,0987958 | 3,85958409 | -0,86118815 | 0,40312692 | 0,65365604 |
| 10979_at | FERMT2 | 0,32117687 | 7,29855896 | 0,86105207 | 0,40319943 | 0,65371179 |
| 84658_at | ADGRE3 | -0,09936574 | 3,93848375 | -0,86099626 | 0,40322917 | 0,65371179 |
| 9758_at | FRMPD4 | -0,06854917 | 3,9034876 | -0,86078247 | 0,40334311 | 0,65384149 |
| 29927_at | SEC61A1 | -0,07180987 | 10,2055116 | -0,86069061 | 0,40339208 | 0,65386583 |
| 10388_at | SYCP2 | 0,29420222 | 4,13348695 | 0,86059548 | 0,40344279 | 0,65386772 |
| 133418_at | EMB | 0,12793378 | 6,20582433 | 0,8605353 | 0,40347487 | 0,65386772 |
| 116835_at | HSPA12B | -0,07871799 | 4,97754759 | -0,86048507 | 0,40350165 | 0,65386772 |
| 3990_at | LIPC | 0,39450398 | 4,97918677 | 0,86043375 | 0,40352902 | 0,65386772 |
| 10978_at | CLP1 | -0,05824799 | 7,71805913 | -0,86034431 | 0,4035767 | 0,65387137 |
| 339479_at | BRINP3 | -0,05398265 | 3,15538699 | -0,8603022 | 0,40359916 | 0,65387137 |
| 190_at | NR0B1 | -0,07009947 | 4,66493445 | -0,86012723 | 0,40369247 | 0,65396754 |
| 79953_at | SYNDIG1 | -0,48011863 | 5,64930572 | -0,85998009 | 0,40377095 | 0,65403967 |
| 79705_at | LRRK1 | 0,13558625 | 5,52305616 | 0,85991188 | 0,40380733 | 0,65404361 |
| 1069_at | CETN2 | -0,12845635 | 9,67769638 | -0,85984063 | 0,40384534 | 0,65405017 |

| | | | | | | |
|---|---|---|---|---|---|---|
| 3788_at | KCNS2 | -0,11853481 | 5,12691837 | -0,8596385 | 0,40395318 | 0,65416982 |
| 6870_at | TACR3 | 0,06850484 | 3,82074621 | 0,85936336 | 0,4041 | 0,65435257 |
| 22904_at | SBNO2 | 0,09429326 | 6,39803565 | 0,85928038 | 0,40414428 | 0,65436927 |
| 2692_at | GHRHR | -0,10434186 | 5,00871497 | -0,85917043 | 0,40420296 | 0,65438985 |
| 100133205_a | LINC00240 | 0,10811315 | 3,53613834 | 0,85912928 | 0,40422493 | 0,65438985 |
| 115560_at | ZNF501 | -0,08647106 | 3,8811057 | -0,85891786 | 0,40433779 | 0,65451755 |
| 83546_at | RTBDN | -0,09715863 | 5,54268875 | -0,85846762 | 0,40457822 | 0,65485171 |
| 203523_at | ZNF449 | 0,08594358 | 5,54888604 | 0,85807369 | 0,40478865 | 0,65513728 |
| 100507050_a | TPBGL | -0,08445663 | 4,73419699 | -0,85766403 | 0,40500757 | 0,65527436 |
| 286530_at | P2RY8 | -0,0993805 | 4,51195431 | -0,85765538 | 0,40501219 | 0,65527436 |
| 1419_at | CRYGB | -0,09691898 | 4,00203108 | -0,85764596 | 0,40501723 | 0,65527436 |
| 23331_at | TTC28 | 0,05031192 | 4,82773903 | 0,85763033 | 0,40502558 | 0,65527436 |
| 401884_at | MGC57346 | 0,08117453 | 7,18004537 | 0,85748473 | 0,40510341 | 0,65527436 |
| 203259_at | FAM219A | -0,11493877 | 6,6690748 | -0,85743599 | 0,40512946 | 0,65527436 |
| 83742_at | MARVELD1 | -0,27168543 | 8,66359887 | -0,85737602 | 0,40516152 | 0,65527436 |
| 80254_at | CEP63 | -0,25322419 | 6,58889493 | -0,85736023 | 0,40516996 | 0,65527436 |
| 830_at | CAPZA2 | 0,09117261 | 9,72027161 | 0,85734237 | 0,40517951 | 0,65527436 |
| 4881_at | NPR1 | -0,07621254 | 5,64594102 | -0,85714656 | 0,4052842 | 0,65534304 |
| 4992_at | OR1F1 | -0,10964201 | 5,05007788 | -0,85713569 | 0,40529002 | 0,65534304 |
| 3090_at | HIC1 | -0,06876724 | 4,95633157 | -0,85693844 | 0,4053955 | 0,65545858 |
| 54737_at | MPHOSPH8 | -0,07892845 | 7,32112394 | -0,85683842 | 0,405449 | 0,65546808 |
| 728116_at | ZBTB8B | 0,07002024 | 4,55386983 | 0,8568002 | 0,40546943 | 0,65546808 |
| 339416_at | ANKRD45 | -0,05888433 | 3,12722662 | -0,85669766 | 0,40552428 | 0,65550174 |
| 8360_at | HIST1H4D | -0,18647669 | 4,99185851 | -0,85652651 | 0,40561585 | 0,65559473 |
| 161291_at | TMEM30B | 0,09066184 | 8,66779066 | 0,85639698 | 0,40568514 | 0,65565173 |
| 29999_at | FSCN3 | -0,12581227 | 4,75267511 | -0,85631087 | 0,40573122 | 0,65566241 |
| 460_at | ASTN1 | -0,08585319 | 3,58868119 | -0,85623134 | 0,40577378 | 0,65566241 |
| 30818_at | KCNIP3 | -0,08761033 | 5,7368972 | -0,85619381 | 0,40579387 | 0,65566241 |
| 9366_at | RAB9BP1 | 0,05517619 | 2,75114127 | 0,85610083 | 0,40584362 | 0,65568781 |
| 1429_at | CRYZ | -0,40130151 | 7,9002164 | -0,85603601 | 0,40587832 | 0,65568887 |
| 401024_at | FSIP2 | 0,07520251 | 3,11548033 | 0,85569138 | 0,40606281 | 0,65588583 |
| 26043_at | UBXN7 | 0,09868739 | 8,19032217 | 0,85566444 | 0,40607723 | 0,65588583 |
| 3209_at | HOXA13 | 0,06597144 | 6,4964855 | 0,85555882 | 0,40613378 | 0,65588583 |
| 641649_at | TMEM91 | 0,15170828 | 5,60827523 | 0,85539816 | 0,40621981 | 0,65588583 |
| 55626_at | AMBRA1 | 0,10659771 | 7,04760058 | 0,8553712 | 0,40623425 | 0,65588583 |
| 2770_at | GNAI1 | 0,33775158 | 8,77910108 | 0,85535357 | 0,40624369 | 0,65588583 |
| 26267_at | FBXO10 | -0,0636805 | 4,92083712 | -0,85533288 | 0,40625477 | 0,65588583 |
| 2220_at | FCN2 | -0,11138333 | 3,91298884 | -0,85529953 | 0,40627263 | 0,65588583 |
| 4118_at | MAL | 0,08855707 | 5,26050877 | 0,85511034 | 0,40637396 | 0,65597201 |
| 51101_at | ZC2HC1A | 0,13910727 | 7,15906766 | 0,85507272 | 0,40639412 | 0,65597201 |
| 29800_at | ZDHHC1 | 0,08530601 | 5,06552657 | 0,85499955 | 0,40643331 | 0,6559803 |
| 23399_at | CTDNEP1 | -0,15498734 | 8,05557559 | -0,85491407 | 0,40647911 | 0,65599925 |
| 256933_at | NPB | 0,0823605 | 7,00500469 | 0,85479818 | 0,4065412 | 0,6560445 |
| 57591_at | MKL1 | 0,10343393 | 6,36202404 | 0,85454497 | 0,40667689 | 0,65620849 |
| 199223_at | TTC21A | 0,17738714 | 5,07233019 | 0,85422086 | 0,40685062 | 0,65643382 |
| 285033_at | STARD7-AS1 | -0,0818507 | 5,84011635 | -0,85411713 | 0,40690622 | 0,65646856 |
| 56_at | ACRV1 | -0,08338027 | 4,13630007 | -0,85351096 | 0,4072313 | 0,6568644 |
| 6522_at | SLC4A2 | 0,15945243 | 7,46313404 | 0,85349343 | 0,4072407 | 0,6568644 |
| 1398_at | CRK | 0,08823154 | 8,88593205 | 0,85346886 | 0,40725388 | 0,6568644 |
| 65258_at | MPPE1 | 0,20009208 | 7,19641258 | 0,85333336 | 0,40732657 | 0,65688857 |

| | | | | | | |
|---|---|---|---|---|---|---|
| 200844_at | C3orf67 | 0,06631278 | 3,38528721 | 0,85331378 | 0,40733707 | 0,65688857 |
| 151258_at | SLC38A11 | 0,08504292 | 3,30837871 | 0,85295255 | 0,4075309 | 0,65714614 |
| 80111_at | C3orf36 | -0,1006657 | 4,38768276 | -0,85248205 | 0,40778346 | 0,65747468 |
| 29924_at | EPN1 | -0,06543388 | 7,15031326 | -0,85244582 | 0,40780291 | 0,65747468 |
| 4695_at | NDUFA2 | 0,05751137 | 9,41120133 | 0,85210266 | 0,40798718 | 0,65771671 |
| 641700_at | ECSCR | -0,09400151 | 5,35337969 | -0,85192484 | 0,40808269 | 0,6578133 |
| 23232_at | TBC1D12 | 0,06332842 | 6,18006537 | 0,85180707 | 0,40814595 | 0,6578133 |
| 6447_at | SCG5 | -0,07367495 | 4,04608961 | -0,85179775 | 0,40815096 | 0,6578133 |
| 128861_at | BPIFA3 | -0,08547314 | 5,31727374 | -0,85170994 | 0,40819813 | 0,6578133 |
| 2705_at | GJB1 | -0,08617759 | 5,32106857 | -0,85167326 | 0,40821784 | 0,6578133 |
| 285547_at | LINC01097 | -0,08622339 | 4,08516651 | -0,85126445 | 0,40843752 | 0,65804464 |
| 50604_at | IL20 | -0,11201234 | 3,82052722 | -0,85108202 | 0,40853557 | 0,65804464 |
| 8392_at | OR3A3 | -0,12710224 | 5,22604781 | -0,85107492 | 0,40853939 | 0,65804464 |
| 10741_at | RBBP9 | 0,07311577 | 7,30148228 | 0,85106805 | 0,40854308 | 0,65804464 |
| 100506742_a | CASP12 | -0,06448038 | 3,4700535 | -0,85103093 | 0,40856304 | 0,65804464 |
| 100506907_a | CCDC37-AS1 | -0,07023826 | 4,01730735 | -0,8509877 | 0,40858628 | 0,65804464 |
| 339442_at | LINC01343 | -0,06684492 | 4,42894569 | -0,85092251 | 0,40862132 | 0,65804464 |
| 83943_at | IMMP2L | 0,07155531 | 6,70775521 | 0,85087834 | 0,40864507 | 0,65804464 |
| 80230_at | RUFY1 | -0,05105493 | 8,18799692 | -0,85079053 | 0,40869228 | 0,65804464 |
| 25999_at | CLIP3 | -0,10184876 | 5,42060858 | -0,85075519 | 0,40871128 | 0,65804464 |
| 5550_at | PREP | 0,06853041 | 7,72091699 | 0,85066395 | 0,40876034 | 0,65804464 |
| 5350_at | PLN | -0,06163104 | 2,87091363 | -0,85064352 | 0,40877133 | 0,65804464 |
| 83539_at | CHST9 | -0,05443635 | 2,9596276 | -0,85035342 | 0,40892735 | 0,65824079 |
| 2207_at | FCER1G | -0,11130838 | 4,83389673 | -0,8502483 | 0,40898389 | 0,6582768 |
| 2626_at | GATA4 | -0,08010271 | 5,0901969 | -0,85013859 | 0,40904291 | 0,65830028 |
| 127733_at | UBXN10 | -0,07307335 | 4,63139902 | -0,85009414 | 0,40906683 | 0,65830028 |
| 1015_at | CDH17 | -0,08615195 | 3,98000751 | -0,85001364 | 0,40911014 | 0,65831285 |
| 5587_at | PRKD1 | 0,14317394 | 4,36841694 | 0,84995258 | 0,40914299 | 0,65831285 |
| 51526_at | OSER1 | 0,15278906 | 8,14755168 | 0,84967304 | 0,40929342 | 0,65849989 |
| 57698_at | SHTN1 | 0,25328163 | 7,76423614 | 0,84913261 | 0,40958435 | 0,65891291 |
| 5562_at | PRKAA1 | 0,16582557 | 7,71627803 | 0,84892826 | 0,40969439 | 0,6590349 |
| 5525_at | PPP2R5A | -0,10601226 | 7,32749452 | -0,84878302 | 0,40977261 | 0,65905936 |
| 6195_at | RPS6KA1 | -0,1465196 | 8,09559877 | -0,8487363 | 0,40979778 | 0,65905936 |
| 4794_at | NFKBIE | 0,20763674 | 6,9361262 | 0,84867723 | 0,40982959 | 0,65905936 |
| 132720_at | C4orf32 | -0,10955062 | 9,00544743 | -0,84864595 | 0,40984644 | 0,65905936 |
| 5569_at | PKIA | -0,14251858 | 8,26902058 | -0,84839396 | 0,4099822 | 0,65922263 |
| 57616_at | TSHZ3 | -0,18630934 | 5,03047549 | -0,84817301 | 0,41010125 | 0,65935902 |
| 147646_at | C19orf84 | -0,1195559 | 6,18059908 | -0,8480148 | 0,41018652 | 0,65944107 |
| 55249_at | YY1AP1 | 0,05128434 | 9,02307992 | 0,84770733 | 0,41035226 | 0,65965248 |
| 6311_at | ATXN2 | 0,05759683 | 8,22571442 | 0,84763849 | 0,41038937 | 0,65965709 |
| 4776_at | NFATC4 | -0,08616612 | 4,93124761 | -0,84756009 | 0,41043164 | 0,65966998 |
| 9580_at | SOX13 | -0,0538308 | 7,65823056 | -0,84748288 | 0,41047327 | 0,65968185 |
| 440508_at | CLEC4GP1 | -0,07159457 | 4,8935746 | -0,84735475 | 0,41054237 | 0,65973786 |
| 6323_at | SCN1A | -0,07196045 | 3,06780946 | -0,84713001 | 0,41066358 | 0,65987759 |
| 124912_at | SPACA3 | -0,08562427 | 4,32228982 | -0,8469134 | 0,41078042 | 0,66001028 |
| 284086_at | NEK8 | 0,12631729 | 6,63190029 | 0,84680084 | 0,41084115 | 0,66002684 |
| 84883_at | AIFM2 | -0,29353957 | 7,83825524 | -0,84676729 | 0,41085925 | 0,66002684 |
| 4109_at | MAGEA10 | -0,09850753 | 4,19315774 | -0,84646119 | 0,41102443 | 0,66022797 |
| 51449_at | PCYOX1 | 0,10107096 | 9,23066714 | 0,84640825 | 0,411053 | 0,66022797 |
| 26707_at | OR2J2 | -0,05239125 | 2,7223393 | -0,84631833 | 0,41110153 | 0,66025086 |

| | | | | | | |
|---|---|---|---|---|---|---|
| 8784_at | TNFRSF18 | -0,07723953 | 5,64664359 | -0,84592204 | 0,41131547 | 0,66053939 |
| 9499_at | MYOT | 0,07200243 | 3,05555317 | 0,84578408 | 0,41138997 | 0,66060395 |
| 53833_at | IL20RB | -0,1739109 | 6,2160266 | -0,84566351 | 0,41145508 | 0,66063982 |
| 3158_at | HMGCS2 | 0,13629611 | 4,25851033 | 0,84561571 | 0,4114809 | 0,66063982 |
| 7881_at | KCNAB1 | 0,08360863 | 4,86065947 | 0,84531206 | 0,41164492 | 0,66084807 |
| 8505_at | PARG | -0,13644155 | 7,53995441 | -0,84523885 | 0,41168447 | 0,66085649 |
| 286514_at | MAGEB18 | 0,07366422 | 3,03262239 | 0,84508811 | 0,41176591 | 0,66088681 |
| 285668_at | C5orf64 | -0,06128527 | 3,43016881 | -0,84507689 | 0,41177197 | 0,66088681 |
| 56145_at | PCDHA3 | -0,12201111 | 4,69325904 | -0,8448277 | 0,41190663 | 0,66104786 |
| 58_at | ACTA1 | 0,08461037 | 5,3537323 | 0,84470026 | 0,41197551 | 0,66110332 |
| 91703_at | ACY3 | 0,06365725 | 5,43967664 | 0,84452191 | 0,41207192 | 0,66120295 |
| 81575_at | APOLD1 | -0,07022662 | 6,17434349 | -0,84430619 | 0,41218855 | 0,66129342 |
| 6738_at | TROVE2 | 0,16306965 | 7,70387639 | 0,84429063 | 0,41219696 | 0,66129342 |
| 59349_at | KLHL12 | 0,12636438 | 8,39052588 | 0,84397404 | 0,41236816 | 0,66142874 |
| 57120_at | GOPC | -0,11960598 | 7,85501299 | -0,84397083 | 0,4123699 | 0,66142874 |
| 200172_at | SLFNL1 | -0,08799861 | 5,95922955 | -0,84394417 | 0,41238432 | 0,66142874 |
| 83893_at | SPATA16 | -0,07928011 | 3,69327785 | -0,84376806 | 0,41247958 | 0,66152644 |
| 167153_at | PAPD4 | 0,23175847 | 8,10176762 | 0,84358475 | 0,41257875 | 0,66154026 |
| 2701_at | GJA4 | -0,08893942 | 5,21704404 | -0,84352 | 0,41261378 | 0,66154026 |
| 4811_at | NID1 | 0,31422347 | 6,28767334 | 0,84346994 | 0,41264087 | 0,66154026 |
| 1780_at | DYNC1I1 | 0,41735304 | 5,32737826 | 0,84337981 | 0,41268964 | 0,66154026 |
| 2815_at | GP9 | -0,07964233 | 6,04739073 | -0,84334675 | 0,41270753 | 0,66154026 |
| 55195_at | C14orf105 | -0,05373237 | 2,92799062 | -0,84334226 | 0,41270995 | 0,66154026 |
| 140876_at | FAM65C | -0,06482278 | 4,34989892 | -0,84329129 | 0,41273754 | 0,66154026 |
| 6279_at | S100A8 | 0,15431516 | 8,52732129 | 0,84324436 | 0,41276293 | 0,66154026 |
| 100128098_a | ST8SIA6-AS1 | -0,0837885 | 4,5141781 | -0,84306836 | 0,41285819 | 0,66162259 |
| 3689_at | ITGB2 | -0,07497782 | 4,69268327 | -0,84302254 | 0,412883 | 0,66162259 |
| 152195_at | NUDT16P1 | 0,1275921 | 5,10946675 | 0,84249619 | 0,41316797 | 0,66200753 |
| 285521_at | COX18 | -0,18902072 | 7,92773007 | -0,84241626 | 0,41321126 | 0,66200753 |
| 387082_at | SUMO4 | -0,08810233 | 8,41376225 | -0,84238844 | 0,41322632 | 0,66200753 |
| 84109_at | QRFPR | -0,06152681 | 3,10375101 | -0,841885 | 0,41349904 | 0,66230776 |
| 51347_at | TAOK3 | 0,14649693 | 8,17967843 | 0,84187235 | 0,41350589 | 0,66230776 |
| 26003_at | GORASP2 | -0,06112358 | 10,2859901 | -0,84180201 | 0,413544 | 0,66230776 |
| 9946_at | CRYZL1 | -0,26204518 | 7,45493529 | -0,84178864 | 0,41355125 | 0,66230776 |
| 284904_at | SEC14L4 | -0,07945307 | 4,2475798 | -0,84172515 | 0,41358565 | 0,66230779 |
| 859_at | CAV3 | -0,1099636 | 5,31663957 | -0,8416014 | 0,41365272 | 0,66236012 |
| 6847_at | SYCP1 | -0,04591482 | 2,72690127 | -0,84139718 | 0,4137634 | 0,66248229 |
| 400761_at | FLJ27354 | -0,1046998 | 3,95885118 | -0,84131653 | 0,41380712 | 0,66249722 |
| 25948_at | KBTBD2 | -0,10062073 | 7,41582295 | -0,84124873 | 0,41384388 | 0,662501 |
| 140901_at | STK35 | 0,05491754 | 8,35602984 | 0,84107438 | 0,4139384 | 0,66255525 |
| 4107_at | MAGEA8 | -0,07695726 | 4,15723452 | -0,84105266 | 0,41395017 | 0,66255525 |
| 55130_at | ARMC4 | -0,06336691 | 3,21528841 | -0,84099591 | 0,41398095 | 0,66255525 |
| 7707_at | ZNF148 | 0,13649827 | 7,6826061 | 0,84090227 | 0,41403172 | 0,66258147 |
| 221946_at | LOC221946 | -0,07743183 | 4,4077545 | -0,84083134 | 0,41407019 | 0,66258797 |
| 286002_at | SLC26A4-AS1 | -0,07602086 | 2,90182602 | -0,84069088 | 0,41414637 | 0,66263621 |
| 90499_at | FAM95A | -0,0897974 | 3,91603867 | -0,84064891 | 0,41416913 | 0,66263621 |
| 4689_at | NCF4 | 0,31480539 | 5,52617895 | 0,84049309 | 0,41425365 | 0,66266567 |
| 8355_at | HIST1H3G | -0,12025178 | 7,83772853 | -0,8404594 | 0,41427193 | 0,66266567 |
| 28964_at | GIT1 | -0,07159462 | 8,54189994 | -0,84042471 | 0,41429075 | 0,66266567 |
| 50508_at | NOX3 | -0,09053003 | 3,31830637 | -0,84030862 | 0,41435373 | 0,66271138 |

| | | | | | | |
|---|---|---|---|---|---|---|
| 100129662_a | FGF13-AS1 | -0,07684135 | 3,49745495 | -0,84023486 | 0,41439375 | 0,66272037 |
| 100507487_a | LOC10050748 | 0,2467087 | 4,55709734 | 0,83998487 | 0,4145294 | 0,66286614 |
| 116931_at | MED12L | 0,08533941 | 4,83741889 | 0,83994006 | 0,41455372 | 0,66286614 |
| 7200_at | TRH | -0,11362217 | 5,03853213 | -0,83985092 | 0,41460211 | 0,66288834 |
| 50486_at | G0S2 | -0,10036524 | 4,73652556 | -0,83976465 | 0,41464893 | 0,66288834 |
| 2554_at | GABRA1 | 0,05670469 | 3,2025588 | 0,83972427 | 0,41467085 | 0,66288834 |
| 23412_at | COMMD3 | 0,23430143 | 8,20724874 | 0,83948575 | 0,41480033 | 0,66296162 |
| 23531_at | MMD | 0,36404281 | 8,96830987 | 0,83945908 | 0,41481482 | 0,66296162 |
| 100874143_a | LINC00358 | -0,0778385 | 3,30922355 | -0,83933579 | 0,41488176 | 0,66296162 |
| 100289230_a | LOC10028923 | -0,17772101 | 5,24440027 | -0,83930342 | 0,41489934 | 0,66296162 |
| 51765_at | STK26 | -0,10594748 | 8,41689329 | -0,83928753 | 0,41490797 | 0,66296162 |
| 5733_at | PTGER3 | -0,05149997 | 3,24778316 | -0,83925951 | 0,41492318 | 0,66296162 |
| 404550_at | C16orf74 | -0,16294972 | 8,35203109 | -0,83911903 | 0,41499947 | 0,66302852 |
| 684959_at | SNORA25 | -0,24067551 | 5,98580948 | -0,83893554 | 0,41509914 | 0,66313275 |
| 796_at | CALCA | -0,0566753 | 4,14905864 | -0,8388312 | 0,41515581 | 0,66316829 |
| 348932_at | SLC6A18 | -0,09494785 | 5,1715972 | -0,83873466 | 0,41520826 | 0,66319708 |
| 11092_at | C9orf9 | 0,17722177 | 5,27030468 | 0,8382702 | 0,41546065 | 0,66354518 |
| 1174_at | AP1S1 | -0,07033285 | 9,15385536 | -0,83801578 | 0,41559894 | 0,66371103 |
| 283521_at | LINC00282 | -0,09147545 | 3,87475683 | -0,83790893 | 0,41565703 | 0,66374876 |
| 114_at | ADCY8 | -0,07889223 | 3,88734198 | -0,83769027 | 0,41577591 | 0,66386777 |
| 203238_at | CCDC171 | -0,06617038 | 3,65291839 | -0,83763291 | 0,41580711 | 0,66386777 |
| 23274_at | CLEC16A | 0,08677986 | 6,36659 | 0,83758172 | 0,41583494 | 0,66386777 |
| 132989_at | C4orf36 | 0,15562961 | 4,40479571 | 0,83742532 | 0,41592 | 0,66394854 |
| 79931_at | TNIP3 | 0,06626188 | 3,09380791 | 0,83731119 | 0,41598208 | 0,66399261 |
| 10893_at | MMP24 | -0,09761864 | 4,9081203 | -0,83697889 | 0,41616286 | 0,66414576 |
| 51299_at | NRN1 | -0,09427792 | 4,22143798 | -0,83696317 | 0,41617141 | 0,66414576 |
| 2297_at | FOXD1 | -0,4287746 | 6,3272167 | -0,83694471 | 0,41618146 | 0,66414576 |
| 57050_at | UTP3 | -0,22566263 | 8,26256606 | -0,83652855 | 0,41640794 | 0,66438266 |
| 9542_at | NRG2 | -0,07182065 | 4,52273265 | -0,83652064 | 0,41641224 | 0,66438266 |
| 5137_at | PDE1C | -0,05702831 | 3,65120399 | -0,83648181 | 0,41643338 | 0,66438266 |
| 53616_at | ADAM22 | 0,04101044 | 3,91742679 | 0,8363823 | 0,41648755 | 0,66441405 |
| 100316904_a | SAP25 | 0,09846689 | 7,39971062 | 0,83631374 | 0,41652487 | 0,66441857 |
| 4636_at | MYL5 | 0,10974213 | 6,65272942 | 0,83607483 | 0,41665495 | 0,66456661 |
| 54039_at | PCBP3 | -0,0848887 | 4,96784779 | -0,83601656 | 0,41668668 | 0,66456661 |
| 149647_at | FAM71A | -0,08840464 | 4,73149251 | -0,8357785 | 0,41681633 | 0,66471835 |
| 6725_at | SRMS | -0,10986249 | 4,50433742 | -0,83541457 | 0,41701457 | 0,66497945 |
| 10406_at | WFDC2 | 0,35131076 | 6,62361909 | 0,83497036 | 0,41725664 | 0,66526611 |
| 57689_at | LRRC4C | -0,05246613 | 3,28666522 | -0,83494432 | 0,41727083 | 0,66526611 |
| 11234_at | HPS5 | 0,17379455 | 7,11545917 | 0,83489457 | 0,41729795 | 0,66526611 |
| 79661_at | NEIL1 | 0,12638258 | 5,98438062 | 0,83457897 | 0,41746999 | 0,66548531 |
| 353322_at | ANKRD37 | -0,35310765 | 6,42823391 | -0,8343712 | 0,41758328 | 0,66552476 |
| 80164_at | PRR36 | 0,12101905 | 5,88337772 | 0,83436981 | 0,41758404 | 0,66552476 |
| 731157_at | LOC731157 | -0,0781916 | 4,19686677 | -0,83416199 | 0,41769737 | 0,66552476 |
| 8756_at | ADAM7 | -0,05165573 | 3,09802898 | -0,83411016 | 0,41772565 | 0,66552476 |
| 4242_at | MFNG | -0,15174678 | 5,12467654 | -0,83408151 | 0,41774127 | 0,66552476 |
| 4114_at | MAGEB3 | 0,06851346 | 3,6718581 | 0,83401687 | 0,41777653 | 0,66552476 |
| 29119_at | CTNNA3 | -0,0493698 | 3,07400026 | -0,83398063 | 0,4177963 | 0,66552476 |
| 158763_at | ARHGAP36 | -0,05754006 | 3,69122567 | -0,83397756 | 0,41779797 | 0,66552476 |
| 93550_at | ZFAND4 | 0,0790946 | 3,66008937 | 0,83396343 | 0,41780568 | 0,66552476 |
| 138199_at | CARNMT1 | -0,07765829 | 6,90324868 | -0,83389473 | 0,41784316 | 0,66552942 |

| | | | | | | |
|---|---|---|---|---|---|---|
| 136991_at | ASZ1 | 0,05596524 | 2,76630757 | 0,83372169 | 0,41793757 | 0,66556771 |
| 261726_at | TIPRL | -0,07689362 | 8,49223672 | -0,83371123 | 0,41794327 | 0,66556771 |
| 5740_at | PTGIS | -0,11342173 | 4,36311575 | -0,83362235 | 0,41799177 | 0,66556771 |
| 5670_at | PSG2 | -0,10901208 | 3,4278662 | -0,83359736 | 0,41800541 | 0,66556771 |
| 5500_at | PPP1CB | 0,13463525 | 9,70846184 | 0,83343192 | 0,41809569 | 0,66565329 |
| 9926_at | LPGAT1 | -0,06401014 | 7,7598484 | -0,83334176 | 0,41814489 | 0,66565329 |
| 65056_at | GPBP1 | 0,11175727 | 8,72258463 | 0,83323417 | 0,41820362 | 0,66565329 |
| 2558_at | GABRA5 | -0,0563297 | 3,70242597 | -0,83319771 | 0,41822352 | 0,66565329 |
| 11330_at | CTRC | -0,08049705 | 4,97015314 | -0,83313377 | 0,41825842 | 0,66565329 |
| 6597_at | SMARCA4 | -0,05357846 | 8,87281265 | -0,83311898 | 0,41826649 | 0,66565329 |
| 23387_at | SIK3 | 0,19004753 | 6,47960426 | 0,83298508 | 0,4183396 | 0,66571463 |
| 55125_at | CEP192 | -0,21067432 | 7,11037069 | -0,83269077 | 0,41850029 | 0,66591535 |
| 284930_at | LOC284930 | -0,07661155 | 3,97721474 | -0,83261742 | 0,41854035 | 0,66592408 |
| 2824_at | GPM6B | -0,08999984 | 3,98788094 | -0,83251445 | 0,41859659 | 0,66595855 |
| 3062_at | HCRTR2 | 0,0907471 | 3,87677098 | 0,83244492 | 0,41863457 | 0,66596396 |
| 115701_at | ALPK2 | -0,06507341 | 3,56796078 | -0,8321501 | 0,41879562 | 0,66616515 |
| 56851_at | EMC7 | -0,11595804 | 9,67181254 | -0,83196626 | 0,41889606 | 0,6662264 |
| 23431_at | AP4E1 | 0,06439033 | 6,3184939 | 0,83195302 | 0,4189033 | 0,6662264 |
| 221937_at | FOXK1 | -0,10120875 | 7,34252772 | -0,83180781 | 0,41898265 | 0,66628788 |
| 284521_at | OR2L13 | 0,06181669 | 3,28549889 | 0,83175571 | 0,41901113 | 0,66628788 |
| 170482_at | CLEC4C | -0,0592585 | 3,26801492 | -0,83143145 | 0,41918836 | 0,66638147 |
| 6696_at | SPP1 | 0,17820558 | 8,319645 | 0,83142875 | 0,41918984 | 0,66638147 |
| 137196_at | CCDC26 | -0,07466222 | 3,08600534 | -0,83142199 | 0,41919354 | 0,66638147 |
| 150248_at | C22orf15 | -0,07814713 | 4,93473988 | -0,83139487 | 0,41920836 | 0,66638147 |
| 100505912_a | LOC10050591 | 0,07897887 | 4,06181335 | 0,83110205 | 0,41936846 | 0,66649479 |
| 161003_at | STOML3 | -0,07391797 | 3,98799478 | -0,83107318 | 0,41938425 | 0,66649479 |
| 158248_at | TTC16 | -0,08835292 | 4,97202831 | -0,83096733 | 0,41944214 | 0,66649479 |
| 27254_at | CSDC2 | -0,09211806 | 6,18162032 | -0,83091831 | 0,41946894 | 0,66649479 |
| 23341_at | DNAJC16 | 0,13484849 | 7,70061873 | 0,83088812 | 0,41948546 | 0,66649479 |
| 147011_at | PROCA1 | -0,07687603 | 5,33661339 | -0,83086634 | 0,41949737 | 0,66649479 |
| 1340_at | COX6B1 | -0,06245532 | 11,8272824 | -0,83082159 | 0,41952185 | 0,66649479 |
| 27109_at | ATP5S | 0,0720081 | 6,7788589 | 0,83070773 | 0,41958413 | 0,66653876 |
| 55764_at | IFT122 | -0,20000704 | 6,68429469 | -0,83029782 | 0,41980839 | 0,66684002 |
| 2130_at | EWSR1 | -0,06124675 | 7,15357914 | -0,83021436 | 0,41985406 | 0,66685758 |
| 388327_at | C17orf100 | 0,21534259 | 5,62942454 | 0,8301446 | 0,41989224 | 0,66686324 |
| 143941_at | TTC36 | -0,07924108 | 5,78192735 | -0,82982781 | 0,42006563 | 0,66693651 |
| 54715_at | RBFOX1 | -0,07085651 | 4,63326681 | -0,82982247 | 0,42006855 | 0,66693651 |
| 8842_at | PROM1 | 0,07579671 | 3,10809988 | 0,82980552 | 0,42007783 | 0,66693651 |
| 9321_at | TRIP11 | 0,12794024 | 7,08304019 | 0,8297356 | 0,42011611 | 0,66693651 |
| 339298_at | LOC339298 | -0,08406687 | 3,89070888 | -0,82967469 | 0,42014946 | 0,66693651 |
| 926_at | CD8B | -0,06380213 | 3,49262247 | -0,82966751 | 0,42015338 | 0,66693651 |
| 9331_at | B4GALT6 | 0,27440884 | 5,92675384 | 0,82961756 | 0,42018073 | 0,66693651 |
| 57519_at | STARD9 | -0,06931667 | 4,0390434 | -0,82948993 | 0,42025062 | 0,66698382 |
| 83605_at | CCM2 | -0,05291583 | 8,14010146 | -0,82943666 | 0,42027979 | 0,66698382 |
| 5972_at | REN | -0,18955718 | 5,53105043 | -0,8293621 | 0,42032062 | 0,66699366 |
| 6452_at | SH3BP2 | 0,06720886 | 7,30110378 | 0,82920657 | 0,42040579 | 0,66707239 |
| 145173_at | B3GLCT | -0,11896765 | 6,63873401 | -0,82914506 | 0,42043949 | 0,66707239 |
| 101927599_a | ZNF529-AS1 | 0,09435044 | 4,51110693 | 0,82904629 | 0,42049359 | 0,66709255 |
| 6497_at | SKI | -0,08890192 | 8,34221598 | -0,82899542 | 0,42052146 | 0,66709255 |
| 84140_at | FAM161A | -0,14160808 | 5,4031582 | -0,82887966 | 0,42058487 | 0,667094 |

| | | | | | | |
|---|---|---|---|---|---|---|
| 11020_at | IFT27 | -0,05382251 | 6,34743144 | -0,82886732 | 0,42059163 | 0,667094 |
| 23504_at | RIMBP2 | -0,06911236 | 4,04083717 | -0,82875911 | 0,42065092 | 0,6671331 |
| 57468_at | SLC12A5 | -0,07847515 | 3,68480811 | -0,82864444 | 0,42071375 | 0,66717782 |
| 822_at | CAPG | 0,37976191 | 9,06681283 | 0,82830255 | 0,42090112 | 0,66738994 |
| 51260_at | PBDC1 | 0,11129397 | 8,29536551 | 0,82827394 | 0,42091681 | 0,66738994 |
| 92737_at | DNER | 0,07186652 | 3,82167812 | 0,82809788 | 0,42101332 | 0,66748725 |
| 23705_at | CADM1 | 0,96093261 | 7,71521128 | 0,82803557 | 0,42104748 | 0,66748725 |
| 8941_at | CDK5R2 | 0,0781123 | 5,16410896 | 0,82790866 | 0,42111706 | 0,66754262 |
| 440590_at | ZYG11A | -0,09936482 | 4,25695556 | -0,82764467 | 0,42126182 | 0,66767266 |
| 2740_at | GLP1R | -0,08137518 | 5,0589686 | -0,82763265 | 0,42126841 | 0,66767266 |
| 101928773_a | LINC01449 | 0,06619697 | 3,0667633 | 0,82726698 | 0,42146899 | 0,66783997 |
| 25941_at | TPGS2 | -0,14078015 | 7,96274559 | -0,82723398 | 0,4214871 | 0,66783997 |
| 387694_at | SH2D4B | -0,08396508 | 4,1202334 | -0,82721701 | 0,42149641 | 0,66783997 |
| 127841_at | LINC00628 | -0,09238946 | 3,57318803 | -0,82718705 | 0,42151284 | 0,66783997 |
| 197131_at | UBR1 | 0,07061261 | 7,4259995 | 0,82712419 | 0,42154733 | 0,66783997 |
| 9454_at | HOMER3 | 0,23305064 | 7,76768738 | 0,8266056 | 0,42183193 | 0,66819864 |
| 3845_at | KRAS | 0,07094719 | 7,66173705 | 0,82658525 | 0,4218431 | 0,66819864 |
| 28652_at | TRAV30 | -0,13741575 | 5,44283281 | -0,8263942 | 0,42194798 | 0,66820266 |
| 102724009_a | LOC10272400 | -0,08954868 | 5,68110999 | -0,82629488 | 0,42200251 | 0,66820266 |
| 5800_at | PTPRO | -0,06040196 | 3,28778342 | -0,82623766 | 0,42203393 | 0,66820266 |
| 9958_at | USP15 | 0,11909062 | 6,9565034 | 0,82623685 | 0,42203437 | 0,66820266 |
| 146664_at | MGAT5B | -0,08756211 | 5,68134183 | -0,82623359 | 0,42203616 | 0,66820266 |
| 144742_at | LINC00934 | -0,06777168 | 3,71606666 | -0,82620153 | 0,42205377 | 0,66820266 |
| 5877_at | RABIF | -0,09702423 | 7,53028535 | -0,82612537 | 0,42209559 | 0,66821395 |
| 84281_at | C2orf88 | -0,05950397 | 3,90277034 | -0,82594085 | 0,42219692 | 0,66831671 |
| 254263_at | CNIH2 | -0,08442646 | 6,55168731 | -0,82588083 | 0,42222989 | 0,66831671 |
| 158521_at | FMR1NB | -0,08309635 | 3,7684109 | -0,82578301 | 0,42228361 | 0,66834683 |
| 55600_at | ITLN1 | 0,36617915 | 4,84246021 | 0,82552768 | 0,42242388 | 0,6685139 |
| 2619_at | GAS1 | 0,0946567 | 4,75705001 | 0,82545854 | 0,42246187 | 0,6685191 |
| 64769_at | MEAF6 | -0,04523212 | 6,30229583 | -0,82515214 | 0,42263024 | 0,6687306 |
| 105378592_a | LOC10537859 | -0,11646343 | 5,15041997 | -0,82503237 | 0,42269607 | 0,66877982 |
| 201651_at | AADACP1 | 0,22617224 | 6,85578615 | 0,82468229 | 0,42288852 | 0,66898696 |
| 9736_at | USP34 | 0,12007382 | 7,29575543 | 0,82461572 | 0,42292512 | 0,66898696 |
| 10352_at | WARS2 | -0,11212863 | 6,7287603 | -0,8246047 | 0,42293117 | 0,66898696 |
| 28908_at | IGKV4-1 | -0,11438905 | 5,15533018 | -0,82451532 | 0,42298032 | 0,66900976 |
| 7779_at | SLC30A1 | 0,07945277 | 7,94673586 | 0,82430695 | 0,42309491 | 0,66908986 |
| 3418_at | IDH2 | 0,13416119 | 8,92054727 | 0,82429691 | 0,42310043 | 0,66908986 |
| 51130_at | ASB3 | 0,15718167 | 7,63846338 | 0,82417518 | 0,42316738 | 0,66914081 |
| 2631_at | GBAS | 0,09179149 | 10,0449573 | 0,82391007 | 0,42331322 | 0,66931647 |
| 23040_at | MYT1L | -0,05968662 | 3,52007947 | -0,82381422 | 0,42336596 | 0,66934491 |
| 9784_at | SNX17 | -0,0413825 | 10,130264 | -0,82373758 | 0,42340812 | 0,66935664 |
| 9562_at | MINPP1 | 0,13583888 | 8,35205703 | 0,82357975 | 0,42349497 | 0,669439 |
| 84632_at | AFAP1L2 | -0,30060789 | 4,9200805 | -0,8233702 | 0,4236103 | 0,66955444 |
| 127391_at | TMCO2 | -0,06352014 | 3,76671103 | -0,82327417 | 0,42366316 | 0,66955444 |
| 116236_at | ABHD15 | 0,18965373 | 7,24175768 | 0,82325761 | 0,42367228 | 0,66955444 |
| 6370_at | CCL25 | -0,14610305 | 4,40889472 | -0,82314289 | 0,42373543 | 0,6695993 |
| 140701_at | ABHD16B | -0,10011074 | 4,80641944 | -0,82293068 | 0,42385226 | 0,66968579 |
| 256471_at | MFSD8 | 0,22609305 | 6,99081731 | 0,8229172 | 0,42385969 | 0,66968579 |
| 81559_at | TRIM11 | -0,04490738 | 7,81220688 | -0,82278376 | 0,42393316 | 0,66974695 |
| 81576_at | CCDC130 | 0,08526809 | 7,38023104 | 0,82259147 | 0,42403906 | 0,66985406 |

| | | | | | | |
|---|---|---|---|---|---|---|
| 90990_at | KIFC2 | 0,28235059 | 7,02258495 | 0,82253437 | 0,42407051 | 0,66985406 |
| 10636_at | RGS14 | -0,0719367 | 5,7585295 | -0,82229051 | 0,42420484 | 0,6700113 |
| 283849_at | EXOC3L1 | -0,07395977 | 5,7929699 | -0,82220236 | 0,4242534 | 0,67003307 |
| 23190_at | UBXN4 | 0,20152723 | 9,6594925 | 0,82207946 | 0,42432112 | 0,67008507 |
| 57521_at | RPTOR | -0,06023377 | 7,33953172 | -0,82200276 | 0,42436339 | 0,67009689 |
| 353116_at | RILPL1 | 0,09237629 | 7,61098258 | 0,82180887 | 0,42447024 | 0,67021067 |
| 131583_at | FAM43A | -0,10887587 | 5,98296344 | -0,82163691 | 0,42456502 | 0,6702889 |
| 7080_at | NKX2-1 | 0,08687343 | 5,5461733 | 0,82159272 | 0,42458938 | 0,6702889 |
| 7710_at | ZNF154 | -0,08701022 | 3,66618454 | -0,8213936 | 0,42469914 | 0,67040724 |
| 51608_at | GET4 | 0,04779527 | 8,09035815 | 0,82122453 | 0,42479236 | 0,67049945 |
| 125972_at | CALR3 | -0,0494133 | 3,79587502 | -0,82114427 | 0,42483662 | 0,67051436 |
| 51084_at | CRYL1 | 0,09377536 | 6,31792776 | 0,82084994 | 0,42499894 | 0,6707156 |
| 84964_at | ALKBH6 | -0,06532502 | 7,73527396 | -0,82065975 | 0,42510386 | 0,67082622 |
| 255480_at | TBX5-AS1 | -0,09111579 | 3,25894489 | -0,82043117 | 0,42522997 | 0,67097025 |
| 10857_at | PGRMC1 | 0,07985914 | 11,1074135 | 0,82030624 | 0,42529891 | 0,67102407 |
| 729238_at | SFTPA2 | -0,09441398 | 3,59151235 | -0,82020178 | 0,42535655 | 0,67104218 |
| 148398_at | SAMD11 | -0,07910305 | 6,58192282 | -0,82014995 | 0,42538516 | 0,67104218 |
| 3012_at | HIST1H2AE | -0,12331613 | 5,55802603 | -0,82009606 | 0,4254149 | 0,67104218 |
| 11245_at | GPR176 | 0,10631858 | 5,59561265 | 0,81991809 | 0,42551313 | 0,67107786 |
| 5145_at | PDE6A | 0,08564662 | 4,06433222 | 0,81986729 | 0,42554117 | 0,67107786 |
| 1825_at | DSC3 | -0,32899136 | 4,0017919 | -0,81982373 | 0,42556522 | 0,67107786 |
| 10179_at | RBM7 | 0,11097797 | 7,0471434 | 0,81959845 | 0,4256896 | 0,67107786 |
| 222642_at | TSPO2 | -0,09334805 | 4,57946891 | -0,81957123 | 0,42570462 | 0,67107786 |
| 10437_at | IFI30 | 0,37935882 | 9,58318898 | 0,81949416 | 0,42574718 | 0,67107786 |
| 116092_at | DNTTIP1 | -0,08069809 | 9,29205822 | -0,81946181 | 0,42576505 | 0,67107786 |
| 439921_at | MXRA7 | -0,07231885 | 4,68286594 | -0,81946063 | 0,4257657 | 0,67107786 |
| 3679_at | ITGA7 | -0,06200249 | 5,03647787 | -0,81945136 | 0,42577082 | 0,67107786 |
| 23305_at | ACSL6 | -0,06866812 | 3,58250547 | -0,81942407 | 0,42578589 | 0,67107786 |
| 3166_at | HMX1 | -0,10801667 | 6,74215383 | -0,81899428 | 0,42602328 | 0,67136728 |
| 163227_at | ZNF100 | -0,10286864 | 7,30768229 | -0,81891951 | 0,42606458 | 0,67136728 |
| 11085_at | ADAM30 | -0,05894585 | 3,16786073 | -0,81890232 | 0,42607408 | 0,67136728 |
| 8898_at | MTMR2 | -0,24655223 | 9,22045276 | -0,81879689 | 0,42613233 | 0,67140414 |
| 116447_at | TOP1MT | -0,23389414 | 8,10096493 | -0,8186081 | 0,42623665 | 0,6714657 |
| 202299_at | LINC01554 | 0,05383476 | 4,20556801 | 0,81860002 | 0,42624111 | 0,6714657 |
| 100128239_a | LOC10012823 | -0,08558264 | 4,57677999 | -0,81845156 | 0,42632316 | 0,67148551 |
| 85019_at | TMEM241 | -0,15994293 | 7,23487751 | -0,81844998 | 0,42632403 | 0,67148551 |
| 1406_at | CRX | -0,0742149 | 4,21759596 | -0,81838283 | 0,42636115 | 0,67148551 |
| 80215_at | RUNX1-IT1 | -0,14635799 | 4,62520301 | -0,81832497 | 0,42639313 | 0,67148551 |
| 283298_at | OLFML1 | -0,06269406 | 3,11283167 | -0,81810376 | 0,42651541 | 0,67162319 |
| 9625_at | AATK | -0,10699039 | 6,05419734 | -0,81754168 | 0,42682623 | 0,67199497 |
| 116843_at | SLC18B1 | 0,26341539 | 8,40078723 | 0,81752832 | 0,42683361 | 0,67199497 |
| 50813_at | COPS7A | 0,07458265 | 9,40889121 | 0,81747609 | 0,42686251 | 0,67199497 |
| 84280_at | BTBD10 | -0,12748159 | 8,94299118 | -0,81742447 | 0,42689106 | 0,67199497 |
| 101927021_a | LOC10192702 | -0,08368739 | 5,15740862 | -0,81719391 | 0,42701861 | 0,67214083 |
| 219855_at | SLC37A2 | 0,31188549 | 7,5596349 | 0,81703644 | 0,42710574 | 0,67222304 |
| 1361_at | CPB2 | -0,07340638 | 3,09003545 | -0,81689968 | 0,42718141 | 0,67228722 |
| 5875_at | RABGGTA | 0,07882387 | 7,86963596 | 0,81681385 | 0,42722892 | 0,67230557 |
| 439931_at | THAP7-AS1 | -0,08465656 | 5,5992002 | -0,8166322 | 0,42732946 | 0,67230557 |
| 23111_at | SPG20 | 0,48518658 | 6,37648995 | 0,81656835 | 0,4273648 | 0,67230557 |
| 51385_at | ZNF589 | -0,04723141 | 5,14053132 | -0,81650511 | 0,42739981 | 0,67230557 |

| | | | | | | |
|---|---|---|---|---|---|---|
| 124995_at | MRPL10 | -0,08638107 | 7,93086364 | -0,81650376 | 0,42740055 | 0,67230557 |
| 80110_at | ZNF614 | -0,05310909 | 6,47788975 | -0,81650027 | 0,42740249 | 0,67230557 |
| 130162_at | CLHC1 | 0,33175564 | 5,04731779 | 0,81630963 | 0,42750803 | 0,67239379 |
| 79869_at | CPSF7 | -0,04965971 | 9,08886232 | -0,81625273 | 0,42753954 | 0,67239379 |
| 4702_at | NDUFA8 | -0,07634416 | 9,62908166 | -0,81615043 | 0,42759618 | 0,67239379 |
| 256714_at | MAP7D2 | 0,08072478 | 4,04524967 | 0,8161468 | 0,42759819 | 0,67239379 |
| 4725_at | NDUFS5 | -0,1428657 | 11,2590315 | -0,81588193 | 0,42774488 | 0,67256955 |
| 5990_at | RFX2 | -0,08006978 | 6,23316705 | -0,81570447 | 0,42784318 | 0,6726692 |
| 79899_at | PRR5L | -0,08365562 | 5,25802847 | -0,81553844 | 0,42793515 | 0,67275453 |
| 23070_at | CMTR1 | 0,15398256 | 8,24937653 | 0,8154337 | 0,42799319 | 0,67275453 |
| 255193_at | CSNK1G2-AS1 | -0,11048605 | 5,71501054 | -0,8154138 | 0,42800421 | 0,67275453 |
| 57644_at | MYH7B | -0,07520884 | 4,71182753 | -0,81535436 | 0,42803715 | 0,67275453 |
| 126208_at | ZNF787 | -0,16601742 | 7,23853805 | -0,81521945 | 0,42811191 | 0,67281714 |
| 57597_at | BAHCC1 | 0,06064045 | 6,48240682 | 0,81502425 | 0,42822009 | 0,67293226 |
| 5994_at | RFXAP | -0,05280295 | 5,64927189 | -0,81478733 | 0,42835142 | 0,67308373 |
| 729121_at | RGPD4-AS1 | -0,06634215 | 2,91905019 | -0,81464825 | 0,42842853 | 0,67310477 |
| 55266_at | TMEM19 | -0,06120341 | 8,86155314 | -0,81463713 | 0,42843469 | 0,67310477 |
| 100129528_a | MUC8 | -0,15004133 | 6,900595 | -0,8144797 | 0,42852198 | 0,673187 |
| 50964_at | SOST | -0,08494715 | 4,14915125 | -0,81431722 | 0,42861209 | 0,67319792 |
| 254128_at | NIFK-AS1 | 0,05097941 | 5,96443714 | 0,81423184 | 0,42865945 | 0,67319792 |
| 26222_at | DGCR10 | -0,05913688 | 2,80593313 | -0,81422131 | 0,42866529 | 0,67319792 |
| 164312_at | LRRN4 | -0,08799381 | 4,5645011 | -0,81416014 | 0,42869921 | 0,67319792 |
| 2185_at | PTK2B | 0,09669999 | 6,71113864 | 0,81415209 | 0,42870368 | 0,67319792 |
| 4593_at | MUSK | -0,04567213 | 3,21960759 | -0,81402302 | 0,42877527 | 0,67325547 |
| 147184_at | TMEM99 | -0,1594536 | 7,02106401 | -0,81384135 | 0,42887606 | 0,67335883 |
| 55069_at | TMEM248 | -0,06309906 | 9,56702795 | -0,81349266 | 0,42906954 | 0,67360572 |
| 80731_at | THSD7B | 0,45840613 | 5,19801628 | 0,81343194 | 0,42910324 | 0,67360572 |
| 951_at | CD37 | -0,08127247 | 4,35811252 | -0,81320607 | 0,42922861 | 0,67371474 |
| 84836_at | ABHD14B | 0,13987016 | 6,72468821 | 0,81317497 | 0,42924588 | 0,67371474 |
| 6728_at | SRP19 | -0,06891798 | 5,86921959 | -0,81303147 | 0,42932554 | 0,67371474 |
| 6866_at | TAC3 | -0,08775407 | 5,56476203 | -0,81301647 | 0,42933387 | 0,67371474 |
| 9986_at | RCE1 | -0,08082671 | 8,17474538 | -0,8129918 | 0,42934757 | 0,67371474 |
| 56244_at | BTNL2 | -0,07462106 | 4,88291628 | -0,81271021 | 0,42950392 | 0,67390519 |
| 57727_at | NCOA5 | 0,06086231 | 8,08961684 | 0,81262531 | 0,42955107 | 0,67392428 |
| 2334_at | AFF2 | -0,07709823 | 4,04541853 | -0,81255293 | 0,42959127 | 0,67393246 |
| 1901_at | S1PR1 | 0,05929016 | 3,38440289 | 0,81219541 | 0,42978987 | 0,674186 |
| 254170_at | FBXO33 | 0,12347482 | 7,3018343 | 0,81213598 | 0,42982289 | 0,674186 |
| 23264_at | ZC3H7B | 0,07490831 | 8,09988949 | 0,81185627 | 0,42997832 | 0,67437488 |
| 100124700_a | HOTAIR | 0,08752838 | 3,95867961 | 0,81167824 | 0,43007726 | 0,67446197 |
| 7185_at | TRAF1 | 0,06274752 | 4,36753236 | 0,81156814 | 0,43013846 | 0,67446197 |
| 5789_at | PTPRD | 0,05612906 | 3,5107478 | 0,81156738 | 0,43013888 | 0,67446197 |
| 100507034_a | PITRM1-AS1 | -0,06382019 | 4,05781464 | -0,81147105 | 0,43019243 | 0,67449103 |
| 282966_at | C10orf53 | -0,05808657 | 4,90941681 | -0,8109686 | 0,4304718 | 0,67473647 |
| 114793_at | FMNL2 | -0,28931909 | 7,09551914 | -0,81093812 | 0,43048875 | 0,67473647 |
| 5743_at | PTGS2 | -0,54600458 | 6,63879645 | -0,81092171 | 0,43049788 | 0,67473647 |
| 338809_at | C12orf74 | -0,06340953 | 3,17828582 | -0,81091843 | 0,43049971 | 0,67473647 |
| 10668_at | CGRRF1 | -0,1099249 | 6,79526526 | -0,81087455 | 0,43052411 | 0,67473647 |
| 1789_at | DNMT3B | 0,07418269 | 7,56542627 | 0,81074534 | 0,43059598 | 0,6747942 |
| 79570_at | NKAIN1 | 0,07976557 | 5,38844463 | 0,81061007 | 0,43067122 | 0,67485722 |
| 222255_at | ATXN7L1 | 0,06507591 | 5,20197883 | 0,81021523 | 0,43089091 | 0,67507521 |

| | | | | | | |
|---|---|---|---|---|---|---|
| 79729_at | SH3D21 | -0,0768466 | 5,78152532 | -0,81019594 | 0,43090165 | 0,67507521 |
| 114299_at | PALM2 | -0,08482013 | 3,87590474 | -0,81017109 | 0,43091547 | 0,67507521 |
| 28803_at | IGLV3-10 | -0,12463098 | 5,18056584 | -0,81007019 | 0,43097163 | 0,67510828 |
| 63982_at | ANO3 | 0,3679603 | 4,98056537 | 0,80994371 | 0,43104203 | 0,67513303 |
| 55604_at | LRRC16A | 0,20121296 | 7,58638417 | 0,80991586 | 0,43105752 | 0,67513303 |
| 152831_at | KLB | 0,14410828 | 4,44932554 | 0,80979352 | 0,43112563 | 0,6751848 |
| 729506_at | LOC729506 | -0,077293 | 4,1388554 | -0,80961468 | 0,43122519 | 0,67528582 |
| 10048_at | RANBP9 | 0,08691755 | 9,23394249 | 0,80945465 | 0,43131429 | 0,67536893 |
| 121340_at | SP7 | -0,0905962 | 5,17161456 | -0,80936674 | 0,43136325 | 0,67536893 |
| 4049_at | LTA | -0,10713574 | 4,43706802 | -0,80933048 | 0,43138344 | 0,67536893 |
| 2002_at | ELK1 | -0,06187684 | 7,19595647 | -0,80924538 | 0,43143084 | 0,67538823 |
| 185_at | AGTR1 | -0,04957751 | 3,43232444 | -0,80905558 | 0,43153655 | 0,67549883 |
| 55897_at | MESP1 | -0,07019099 | 5,70834877 | -0,80879376 | 0,4316824 | 0,67563982 |
| 116729_at | PPP1R27 | -0,09783807 | 5,49332244 | -0,80873656 | 0,43171427 | 0,67563982 |
| 101928211_a | LOC10192821 | 0,04765591 | 2,78616537 | 0,80870503 | 0,43173184 | 0,67563982 |
| 256364_at | EML3 | -0,04222557 | 7,18383189 | -0,80862278 | 0,43177767 | 0,67565665 |
| 283432_at | LINC00485 | 0,04542966 | 2,87616673 | 0,80850795 | 0,43184166 | 0,67570188 |
| 91156_at | IGFN1 | -0,06352283 | 4,43051957 | -0,80840854 | 0,43189706 | 0,67571559 |
| 27146_at | FAM184B | 0,08657265 | 4,36188727 | 0,80832104 | 0,43194582 | 0,67571559 |
| 55301_at | OLAH | -0,0629596 | 3,10447444 | -0,80830341 | 0,43195565 | 0,67571559 |
| 23708_at | GSPT2 | -0,06994059 | 3,53580064 | -0,80816762 | 0,43203134 | 0,67573125 |
| 4633_at | MYL2 | -0,0952821 | 3,85602495 | -0,80815366 | 0,43203912 | 0,67573125 |
| 79987_at | SVEP1 | -0,057385 | 4,4273176 | -0,80809665 | 0,4320709 | 0,67573125 |
| 65217_at | PCDH15 | 0,0431845 | 2,82720523 | 0,80801906 | 0,43211416 | 0,67574404 |
| 10838_at | ZNF275 | -0,07557745 | 7,0912373 | -0,80791387 | 0,43217281 | 0,67575563 |
| 225689_at | MAPK15 | -0,08923128 | 5,43230715 | -0,80787993 | 0,43219173 | 0,67575563 |
| 105373132_a | LOC10537313 | -0,11687364 | 4,14375387 | -0,80771675 | 0,43228272 | 0,67584303 |
| 155382_at | VPS37D | 0,08449284 | 6,30955947 | 0,80752809 | 0,43238793 | 0,67587604 |
| 84866_at | TMEM25 | 0,27562159 | 6,14729305 | 0,80750558 | 0,43240048 | 0,67587604 |
| 4620_at | MYH2 | -0,05288138 | 3,08010524 | -0,80742695 | 0,43244434 | 0,67587604 |
| 100533182_a | SEC24B-AS1 | 0,06545246 | 5,58119131 | 0,80737523 | 0,43247319 | 0,67587604 |
| 148281_at | SYT6 | -0,07930261 | 4,45252409 | -0,80736434 | 0,43247926 | 0,67587604 |
| 114327_at | EFHC1 | 0,18830314 | 6,00370022 | 0,80701743 | 0,4326728 | 0,67607947 |
| 729723_at | DNAJC27-AS1 | -0,07466261 | 4,46732813 | -0,80695395 | 0,43270822 | 0,67607947 |
| 168451_at | THAP5 | 0,1327464 | 5,63803536 | 0,80688388 | 0,43274732 | 0,67607947 |
| 101928731_a | LOC10192873 | -0,05980679 | 3,40206535 | -0,80687939 | 0,43274982 | 0,67607947 |
| 23020_at | SNRNP200 | -0,07185805 | 10,4672871 | -0,80666239 | 0,43287092 | 0,67621382 |
| 3394_at | IRF8 | -0,08525322 | 4,64728161 | -0,80649939 | 0,4329619 | 0,6763011 |
| 23022_at | PALLD | 0,355367 | 9,8190659 | 0,80633919 | 0,43305133 | 0,67638594 |
| 57709_at | SLC7A14 | -0,07711587 | 3,91434935 | -0,80625009 | 0,43310108 | 0,67640879 |
| 147741_at | ZNF560 | 0,05081362 | 3,01566786 | 0,80605151 | 0,43321196 | 0,67652711 |
| 10324_at | KLHL41 | 0,05948115 | 3,2807328 | 0,80591174 | 0,43329001 | 0,67658818 |
| 10552_at | ARPC1A | -0,08435146 | 10,8662307 | -0,80582729 | 0,43333718 | 0,67658818 |
| 284751_at | LINC01270 | -0,06603078 | 4,72098136 | -0,80579281 | 0,43335643 | 0,67658818 |
| 101929212_a | SMIM2-AS1 | 0,06570977 | 4,02545669 | 0,80535246 | 0,43360242 | 0,67691737 |
| 338094_at | FAM151A | -0,09986139 | 5,01748932 | -0,80523783 | 0,43366647 | 0,6769625 |
| 105369187_a | LOC10536918 | -0,08933256 | 3,69457876 | -0,80506014 | 0,43376577 | 0,67703768 |
| 339965_at | CCDC158 | -0,09556734 | 3,11260898 | -0,80493947 | 0,43383321 | 0,67703768 |
| 103689912_a | TBX2-AS1 | -0,25635459 | 6,48639386 | -0,80490974 | 0,43384983 | 0,67703768 |
| 1770_at | DNAH9 | -0,0651485 | 4,43333956 | -0,80484231 | 0,43388751 | 0,67703768 |

| | | | | | | |
|---|---|---|---|---|---|---|
| 4760_at | NEUROD1 | 0,05539921 | 3,42778006 | 0,80483721 | 0,43389037 | 0,67703768 |
| 3077_at | HFE | 0,08126683 | 6,5245234 | 0,80468829 | 0,43397361 | 0,67707206 |
| 144132_at | DNHD1 | -0,07214283 | 4,24890261 | -0,80467204 | 0,4339827 | 0,67707206 |
| 55869_at | HDAC8 | 0,03828754 | 7,12856897 | 0,8044233 | 0,43412177 | 0,67723418 |
| 101929120_a | LINC01351 | 0,05393475 | 3,26107888 | 0,80434744 | 0,43416418 | 0,6772451 |
| 7319_at | UBE2A | -0,09697199 | 10,4058103 | -0,80423334 | 0,43422799 | 0,6772451 |
| 199713_at | NLRP7 | 0,47849153 | 5,9532898 | 0,80422216 | 0,43423424 | 0,6772451 |
| 283208_at | P4HA3 | -0,06225788 | 4,82852197 | -0,80399787 | 0,43435969 | 0,6773859 |
| 6675_at | UAP1 | -0,09151892 | 8,71001961 | -0,80384047 | 0,43444774 | 0,67746837 |
| 29843_at | SENP1 | -0,15196479 | 7,70891457 | -0,80373358 | 0,43450753 | 0,67750677 |
| 55182_at | RNF220 | -0,0974884 | 8,19910732 | -0,80362849 | 0,43456633 | 0,6775436 |
| 84248_at | FYTTD1 | 0,073955 | 9,53722161 | 0,80345703 | 0,43466228 | 0,67763835 |
| 101930085_a | LOC10193008 | 0,2028507 | 6,86435987 | 0,80311577 | 0,43485327 | 0,67784949 |
| 725_at | C4BPB | -0,07939572 | 4,15219041 | -0,8030893 | 0,43486809 | 0,67784949 |
| 56171_at | DNAH7 | 0,05876109 | 3,74922577 | 0,80293708 | 0,4349533 | 0,67792745 |
| 9023_at | CH25H | 0,05689449 | 3,91354451 | 0,80283279 | 0,43501169 | 0,67794081 |
| 345611_at | IRGM | 0,06529387 | 2,96236043 | 0,80274261 | 0,43506218 | 0,67794081 |
| 11181_at | TREH | -0,0913908 | 3,79344901 | -0,80270997 | 0,43508046 | 0,67794081 |
| 6812_at | STXBP1 | 0,26767696 | 7,42018861 | 0,80267034 | 0,43510265 | 0,67794081 |
| 22894_at | DIS3 | 0,09268054 | 7,0626792 | 0,80255053 | 0,43516974 | 0,67794962 |
| 22919_at | MAPRE1 | -0,0788109 | 10,4216323 | -0,80253456 | 0,43517869 | 0,67794962 |
| 971_at | CD72 | 0,07304763 | 4,40349157 | 0,80241217 | 0,43524723 | 0,67800157 |
| 54851_at | ANKRD49 | 0,13294681 | 7,91605953 | 0,80216524 | 0,43538554 | 0,67816218 |
| 151126_at | ZNF385B | 0,06037061 | 3,54042541 | 0,80198631 | 0,43548579 | 0,6781886 |
| 10330_at | CNPY2 | -0,05345212 | 9,26823732 | -0,80196577 | 0,43549729 | 0,6781886 |
| 1448_at | CSN3 | -0,06914729 | 3,48359765 | -0,80190816 | 0,43552957 | 0,6781886 |
| 1837_at | DTNA | -0,05865903 | 3,58200711 | -0,80179792 | 0,43559134 | 0,6781886 |
| 64168_at | NECAB1 | -0,05348527 | 3,12251646 | -0,80170454 | 0,43564367 | 0,6781886 |
| 6693_at | SPN | -0,07533708 | 4,74936513 | -0,8016889 | 0,43565244 | 0,6781886 |
| 5930_at | RBBP6 | 0,04404154 | 8,23079037 | 0,80168402 | 0,43565517 | 0,6781886 |
| 10912_at | GADD45G | -0,11729927 | 5,92052194 | -0,80162736 | 0,43568693 | 0,6781886 |
| 11281_at | POU6F2 | -0,06521126 | 3,91215827 | -0,80156949 | 0,43571936 | 0,6781886 |
| 51156_at | SERPINA10 | -0,08023845 | 3,93413349 | -0,80149544 | 0,43576087 | 0,6781984 |
| 820_at | CAMP | 0,07243545 | 5,72869557 | 0,80133145 | 0,43585279 | 0,67828665 |
| 9765_at | ZFYVE16 | 0,12180642 | 6,37791082 | 0,80083101 | 0,43613338 | 0,67862915 |
| 284276_at | LINC00908 | -0,08333052 | 4,93581657 | -0,80081328 | 0,43614332 | 0,67862915 |
| 101927150_a | L3MBTL4-AS1 | -0,08821342 | 3,97180416 | -0,80065297 | 0,43623323 | 0,67871422 |
| 64116_at | SLC39A8 | -0,20021086 | 6,51950646 | -0,80048435 | 0,43632782 | 0,67873863 |
| 6640_at | SNTA1 | 0,09754566 | 7,1082348 | 0,8004706 | 0,43633553 | 0,67873863 |
| 146880_at | LOC146880 | 0,09521278 | 5,64725652 | 0,80043656 | 0,43635462 | 0,67873863 |
| 101927715_a | LINC01029 | -0,07616998 | 3,70805028 | -0,80002473 | 0,4365857 | 0,67894245 |
| 342538_at | NACA2 | -0,0653988 | 3,013744 | -0,7999553 | 0,43662466 | 0,67894245 |
| 284369_at | SIGLECL1 | -0,08044452 | 3,5923458 | -0,79976171 | 0,43673331 | 0,67894245 |
| 105373764_a | LOC10537376 | 0,06703013 | 3,11547369 | 0,79975871 | 0,436735 | 0,67894245 |
| 6799_at | SULT1A2 | 0,20804062 | 6,64268665 | 0,79975433 | 0,43673746 | 0,67894245 |
| 125144_at | LRRC75A-AS1 | -0,12021026 | 7,11536965 | -0,79974072 | 0,43674509 | 0,67894245 |
| 55224_at | ETNK2 | 0,14856475 | 5,82833075 | 0,79967452 | 0,43678225 | 0,67894245 |
| 94_at | ACVRL1 | -0,09826225 | 4,85950463 | -0,79967079 | 0,43678435 | 0,67894245 |
| 7448_at | VTN | -0,06916103 | 4,67391 | -0,79963779 | 0,43680287 | 0,67894245 |
| 5584_at | PRKCI | 0,16270265 | 10,1981473 | 0,79941713 | 0,43692675 | 0,6790802 |

| | | | | | | |
|---|---|---|---|---|---|---|
| 81832_at | NETO1 | -0,10770679 | 3,4753168 | -0,79927916 | 0,43700422 | 0,67913152 |
| 55607_at | PPP1R9A | 0,04490095 | 3,84778676 | 0,79920746 | 0,43704448 | 0,67913152 |
| 1622_at | DBI | 0,07365686 | 11,1596532 | 0,79916997 | 0,43706554 | 0,67913152 |
| 10072_at | DPP3 | 0,05239076 | 9,78754599 | 0,79878045 | 0,43728431 | 0,67938907 |
| 25829_at | TMEM184B | -0,05805038 | 9,35207611 | -0,7987426 | 0,43730557 | 0,67938907 |
| 64816_at | CYP3A43 | 0,07497236 | 3,58757744 | 0,79868649 | 0,43733709 | 0,67938907 |
| 128989_at | TANGO2 | -0,09102149 | 8,01904889 | -0,79841842 | 0,43748771 | 0,67956825 |
| 285955_at | SPDYE1 | 0,12296623 | 4,80705664 | 0,79828411 | 0,43756319 | 0,67963068 |
| 101927565_a | LINC01267 | -0,08100142 | 3,98161441 | -0,79809516 | 0,43766938 | 0,67963561 |
| 83957_at | FRMD8P1 | -0,07222591 | 6,11007403 | -0,79807652 | 0,43767986 | 0,67963561 |
| 81491_at | GPR63 | 0,23170056 | 5,66252211 | 0,79804563 | 0,43769722 | 0,67963561 |
| 374864_at | CCDC178 | -0,09109483 | 4,26754218 | -0,79800905 | 0,43771778 | 0,67963561 |
| 54665_at | RSBN1 | 0,06505956 | 7,3901671 | 0,79796076 | 0,43774493 | 0,67963561 |
| 100506795_a | LINC01354 | 0,05915616 | 3,41115615 | 0,79790183 | 0,43777806 | 0,67963561 |
| 23538_at | OR52A1 | -0,06942027 | 3,52736464 | -0,79769986 | 0,4378916 | 0,67973029 |
| 6718_at | AKR1D1 | -0,05646664 | 3,16294586 | -0,79766782 | 0,43790962 | 0,67973029 |
| 6925_at | TCF4 | 0,04222579 | 3,90762947 | 0,79747161 | 0,43801995 | 0,67978804 |
| 100287015_a | LOC10028701 | -0,17513284 | 4,90773365 | -0,79740123 | 0,43805953 | 0,67978804 |
| 100505619_a | LINC00919 | -0,05040921 | 3,69793372 | -0,79729239 | 0,43812074 | 0,67978804 |
| 4868_at | NPHS1 | -0,07498408 | 4,14821794 | -0,79729216 | 0,43812086 | 0,67978804 |
| 157310_at | PEBP4 | -0,10409282 | 4,39847431 | -0,79723272 | 0,4381543 | 0,67978804 |
| 23500_at | DAAM2 | -0,08975091 | 5,02837197 | -0,79722515 | 0,43815856 | 0,67978804 |
| 100505659_a | LINC00967 | -0,08630163 | 3,85209199 | -0,7971593 | 0,4381956 | 0,67979075 |
| 1740_at | DLG2 | 0,05940807 | 3,60682321 | 0,79691228 | 0,43833456 | 0,67979379 |
| 23119_at | HIC2 | 0,10815204 | 6,69192184 | 0,79680039 | 0,43839751 | 0,67979379 |
| 1084_at | CEACAM3 | -0,09362925 | 5,58230801 | -0,79679567 | 0,43840017 | 0,67979379 |
| 338963_at | LOC338963 | -0,09751902 | 5,51452156 | -0,79675355 | 0,43842386 | 0,67979379 |
| 677779_at | LINC00488 | 0,09025452 | 3,49691543 | 0,79675189 | 0,4384248 | 0,67979379 |
| 3658_at | IREB2 | 0,13372273 | 8,46985753 | 0,79671739 | 0,43844421 | 0,67979379 |
| 101928708_a | LOC10192870 | -0,0817791 | 4,6036192 | -0,79671673 | 0,43844458 | 0,67979379 |
| 148930_at | KNCN | -0,09549404 | 4,88935741 | -0,79664363 | 0,43848572 | 0,67980285 |
| 26526_at | TSPAN16 | -0,07043265 | 3,95534038 | -0,79645772 | 0,43859034 | 0,67983163 |
| 283899_at | INO80E | -0,26128887 | 7,2881363 | -0,79639582 | 0,43862518 | 0,67983163 |
| 84654_at | SPZ1 | 0,04394841 | 2,86247759 | 0,79631138 | 0,43867271 | 0,67983163 |
| 29113_at | C6orf15 | -0,14202925 | 5,97595995 | -0,7963089 | 0,43867411 | 0,67983163 |
| 255631_at | COL24A1 | 0,0529617 | 3,67617962 | 0,7962971 | 0,43868075 | 0,67983163 |
| 10818_at | FRS2 | 0,07239467 | 6,49701147 | 0,79607956 | 0,43880321 | 0,67996671 |
| 4485_at | MST1 | 0,06683801 | 6,65406661 | 0,79577561 | 0,43897435 | 0,6801772 |
| 5910_at | RAP1GDS1 | -0,37435214 | 7,9800782 | -0,79568313 | 0,43902643 | 0,68020318 |
| 80757_at | TMEM121 | -0,06612562 | 5,24155192 | -0,79545354 | 0,43915574 | 0,68030973 |
| 57708_at | MIER1 | 0,0829962 | 7,35393474 | 0,79543562 | 0,43916584 | 0,68030973 |
| 147929_at | ZNF565 | 0,06831473 | 5,95267286 | 0,79520905 | 0,43929347 | 0,68045273 |
| 57536_at | KIAA1328 | -0,08204864 | 4,91440798 | -0,79506788 | 0,43937301 | 0,68052121 |
| 389023_at | DPP10-AS1 | -0,08115463 | 3,97112192 | -0,79493605 | 0,43944729 | 0,68053111 |
| 7358_at | UGDH | -0,17970146 | 10,5457601 | -0,79482678 | 0,43950887 | 0,68053111 |
| 158219_at | TTC39B | 0,12282418 | 5,54734798 | 0,79481706 | 0,43951435 | 0,68053111 |
| 57794_at | SUGP1 | 0,06256812 | 8,04092701 | 0,79480576 | 0,43952072 | 0,68053111 |
| 92283_at | ZNF461 | 0,1097953 | 4,40720527 | 0,79455531 | 0,43966188 | 0,68069497 |
| 26275_at | HIBCH | -0,13262824 | 7,77603193 | -0,79425382 | 0,43983185 | 0,68090339 |
| 23432_at | GPR161 | 0,09762652 | 6,0532902 | 0,79398374 | 0,43998414 | 0,68103412 |

| | | | | | | |
|---|---|---|---|---|---|---|
| 51761_at | ATP8A2 | -0,07167277 | 4,00013202 | -0,79397866 | 0,43998701 | 0,68103412 |
| 145946_at | SPATA8 | -0,08541407 | 4,68979503 | -0,79379104 | 0,44009282 | 0,68114318 |
| 4656_at | MYOG | -0,09643341 | 5,13492321 | -0,79361301 | 0,44019324 | 0,68123788 |
| 101928012_a | LOC10192801 | 0,04505435 | 2,96770703 | 0,79355719 | 0,44022474 | 0,68123788 |
| 114990_at | VASN | -0,12706642 | 5,62727093 | -0,79347918 | 0,44026875 | 0,68125126 |
| 747_at | DAGLA | -0,11663757 | 5,85429917 | -0,79307435 | 0,44049718 | 0,68154998 |
| 253143_at | PRR14L | 0,06120954 | 7,6644906 | 0,79293619 | 0,44057516 | 0,68159861 |
| 157922_at | CAMSAP1 | -0,04653264 | 6,52901241 | -0,79289327 | 0,44059938 | 0,68159861 |
| 55472_at | RBM12B-AS1 | -0,07734537 | 3,9214382 | -0,79282258 | 0,44063929 | 0,6816056 |
| 101927701_a | LOC10192770 | -0,09317589 | 3,78587727 | -0,79229847 | 0,44093521 | 0,68193889 |
| 84221_at | SPATC1L | -0,11053753 | 7,59601939 | -0,79229534 | 0,44093697 | 0,68193889 |
| 85480_at | TSLP | -0,07262527 | 2,96285274 | -0,79223538 | 0,44097083 | 0,68193889 |
| 400027_at | LINC00938 | 0,07896689 | 6,29721228 | 0,79219019 | 0,44099635 | 0,68193889 |
| 10449_at | ACAA2 | 0,1612769 | 5,6671068 | 0,79209341 | 0,44105102 | 0,68194958 |
| 342898_at | SYCN | -0,12163594 | 4,88031918 | -0,79202905 | 0,44108736 | 0,68194958 |
| 205564_at | SENP5 | 0,09019578 | 7,20096566 | 0,79193622 | 0,4411398 | 0,68194958 |
| 56994_at | CHPT1 | -0,17368739 | 6,82510704 | -0,79188849 | 0,44116676 | 0,68194958 |
| 280636_at | C11orf31 | 0,16987119 | 9,03824613 | 0,79186457 | 0,44118028 | 0,68194958 |
| 56623_at | INPP5E | 0,07074727 | 7,71164373 | 0,79168713 | 0,44128053 | 0,68204981 |
| 1141_at | CHRNB2 | -0,07977102 | 5,88381759 | -0,7912266 | 0,44154078 | 0,6823535 |
| 5836_at | PYGL | 0,30230681 | 10,4203987 | 0,79121408 | 0,44154785 | 0,6823535 |
| 8339_at | HIST1H2BG | -0,11091308 | 6,18663131 | -0,79101455 | 0,44166064 | 0,68247304 |
| 23349_at | PHF24 | -0,07993734 | 4,61131733 | -0,79081292 | 0,44177464 | 0,68259443 |
| 283165_at | KIRREL3-AS3 | -0,10159717 | 4,74006935 | -0,79066536 | 0,44185807 | 0,68266859 |
| 101928961_a | LOC10192896 | -0,0502505 | 2,93853075 | -0,79038227 | 0,44201817 | 0,68277618 |
| 203074_at | PRSS55 | -0,07460668 | 4,09153331 | -0,7903428 | 0,44204049 | 0,68277618 |
| 84307_at | ZNF397 | 0,13643621 | 5,27862595 | 0,79026429 | 0,4420849 | 0,68277618 |
| 101927358_a | LOC10192735 | -0,04594627 | 2,86878807 | -0,79017363 | 0,44213619 | 0,68277618 |
| 101929180_a | LOC10192918 | -0,11716953 | 4,67703334 | -0,79010978 | 0,44217231 | 0,68277618 |
| 149708_at | WFDC5 | 0,06001219 | 4,23847591 | 0,7901067 | 0,44217405 | 0,68277618 |
| 29916_at | SNX11 | 0,06261962 | 7,83376839 | 0,79010356 | 0,44217582 | 0,68277618 |
| 79822_at | ARHGAP28 | 0,05785014 | 4,15597276 | 0,78982916 | 0,44233108 | 0,68296117 |
| 81565_at | NDEL1 | 0,05333301 | 6,88975474 | 0,78971637 | 0,44239491 | 0,68298112 |
| 23287_at | AGTPBP1 | 0,08183202 | 3,93007701 | 0,78968102 | 0,44241491 | 0,68298112 |
| 100506311_a | HOTAIRM1 | 0,2440804 | 8,07917571 | 0,78945611 | 0,4425422 | 0,68307118 |
| 284325_at | C19orf54 | 0,06662598 | 7,97181248 | 0,78942492 | 0,44255986 | 0,68307118 |
| 54807_at | ZNF586 | 0,06887966 | 7,02737911 | 0,78938999 | 0,44257963 | 0,68307118 |
| 8324_at | FZD7 | -0,18656641 | 7,63999641 | -0,78920129 | 0,44268645 | 0,68318131 |
| 2743_at | GLRB | 0,13607866 | 4,05138378 | 0,78896113 | 0,44282243 | 0,68328448 |
| 114897_at | C1QTNF1 | -0,08201309 | 4,63340484 | -0,78895792 | 0,44282425 | 0,68328448 |
| 5331_at | PLCB3 | -0,09825313 | 7,6066374 | -0,788727 | 0,44295502 | 0,68341412 |
| 114786_at | XKR4 | -0,05801168 | 4,14614642 | -0,78868426 | 0,44297922 | 0,68341412 |
| 54102_at | CLIC6 | -0,05059256 | 3,80116971 | -0,78847464 | 0,44309796 | 0,68348585 |
| 133619_at | PRRC1 | 0,11407975 | 9,77039789 | 0,78837863 | 0,44315235 | 0,68348585 |
| 440132_at | LINC00545 | -0,09805849 | 3,93995995 | -0,78829046 | 0,4432023 | 0,68348585 |
| 387978_at | LINC01551 | -0,0784167 | 3,88133953 | -0,78827215 | 0,44321267 | 0,68348585 |
| 220963_at | SLC16A9 | -0,16246954 | 4,75786075 | -0,78826128 | 0,44321883 | 0,68348585 |
| 4087_at | SMAD2 | -0,03637173 | 7,83902092 | -0,78822638 | 0,44323861 | 0,68348585 |
| 102723335_a | LOC10272333 | 0,08250762 | 4,45893417 | 0,78812163 | 0,44329796 | 0,68352266 |
| 100270679_a | LINC01432 | 0,07734993 | 3,63693068 | 0,78798361 | 0,44337617 | 0,68358854 |

| | | | | | | |
|---|---|---|---|---|---|---|
| 100131067_a | CKMT2-AS1 | 0,05166123 | 5,06352058 | 0,78766695 | 0,44355564 | 0,68381051 |
| 9612_at | NCOR2 | -0,06377665 | 6,74800841 | -0,78746985 | 0,44366738 | 0,68392803 |
| 101927292_a | LOC10192729 | -0,06116329 | 3,08704524 | -0,78730912 | 0,44375851 | 0,68401086 |
| 353174_at | ZACN | -0,15470296 | 5,57506143 | -0,78724983 | 0,44379213 | 0,68401086 |
| 6754_at | SSTR4 | -0,12492705 | 5,21230289 | -0,78710054 | 0,44387678 | 0,68406469 |
| 80724_at | ACAD10 | 0,04396695 | 6,12153914 | 0,78706299 | 0,44389808 | 0,68406469 |
| 133874_at | C5orf58 | 0,05264832 | 3,1940037 | 0,78677201 | 0,44406312 | 0,68416731 |
| 101928571_a | ZNF649-AS1 | -0,10001894 | 4,49466639 | -0,78673166 | 0,44408601 | 0,68416731 |
| 253943_at | YTHDF3 | 0,09174276 | 8,71187772 | 0,78671957 | 0,44409287 | 0,68416731 |
| 9486_at | CHST10 | -0,17698419 | 6,59344096 | -0,78669512 | 0,44410674 | 0,68416731 |
| 84224_at | NBPF3 | 0,07074086 | 4,6493629 | 0,78657358 | 0,44417569 | 0,68417909 |
| 101927814_a | LOC10192781 | -0,065665 | 3,95709428 | -0,78653212 | 0,44419921 | 0,68417909 |
| 388272_at | C16orf87 | -0,14784366 | 6,86476209 | -0,78640618 | 0,44427067 | 0,68417909 |
| 146850_at | PIK3R6 | -0,07361716 | 4,73180713 | -0,78638366 | 0,44428345 | 0,68417909 |
| 63920_at | ZBED8 | 0,12255941 | 6,03850936 | 0,78635467 | 0,4442999 | 0,68417909 |
| 11247_at | NXPH4 | -0,09040573 | 6,62563058 | -0,78630605 | 0,44432749 | 0,68417909 |
| 115201_at | ATG4A | -0,07008644 | 7,00167881 | -0,7859672 | 0,4445198 | 0,6844205 |
| 3772_at | KCNJ15 | 0,16325701 | 8,77330821 | 0,78583553 | 0,44459454 | 0,68448087 |
| 219990_at | OOSP2 | -0,07096671 | 3,05326904 | -0,78553031 | 0,44476783 | 0,68469294 |
| 57085_at | AGTRAP | -0,20915818 | 7,84883353 | -0,78522881 | 0,44493905 | 0,68490178 |
| 9181_at | ARHGEF2 | -0,04089544 | 6,64275897 | -0,78514599 | 0,44498609 | 0,68491947 |
| 25782_at | RAB3GAP2 | 0,10312115 | 8,31038398 | 0,78480878 | 0,44517765 | 0,68515957 |
| 101929681_a | LOC10192968 | 0,13282029 | 4,28640397 | 0,78459987 | 0,44529636 | 0,6852875 |
| 170680_at | PSORS1C2 | -0,11557449 | 5,45595496 | -0,78448045 | 0,44536422 | 0,68533719 |
| 100505854_a | APTR | 0,08835053 | 6,81606181 | 0,78427556 | 0,44548067 | 0,68537947 |
| 64386_at | MMP25 | -0,09760837 | 5,72281586 | -0,78424721 | 0,44549678 | 0,68537947 |
| 54972_at | TMEM132A | 0,09854054 | 6,64446111 | 0,7842443 | 0,44549844 | 0,68537947 |
| 3840_at | KPNA4 | 0,10039419 | 8,57860228 | 0,78402446 | 0,44562341 | 0,68544243 |
| 22908_at | SACM1L | -0,18730116 | 9,21406326 | -0,78400711 | 0,44563327 | 0,68544243 |
| 57648_at | KIAA1522 | 0,10456456 | 9,95775515 | 0,78398453 | 0,44564611 | 0,68544243 |
| 283547_at | LINC00639 | -0,05990821 | 3,15881133 | -0,78361428 | 0,44585664 | 0,6856738 |
| 414919_at | C8orf82 | 0,13912444 | 8,06149476 | 0,78354593 | 0,44589551 | 0,6856738 |
| 7748_at | ZNF195 | 0,12225209 | 7,09215068 | 0,7835322 | 0,44590332 | 0,6856738 |
| 28986_at | MAGEH1 | 0,35900824 | 6,48475598 | 0,78322655 | 0,44607718 | 0,68585666 |
| 374768_at | SPEM1 | -0,08749997 | 5,10148766 | -0,78315778 | 0,4461163 | 0,68585666 |
| 5818_at | PVRL1 | -0,05352692 | 7,40021715 | -0,78313536 | 0,44612905 | 0,68585666 |
| 4958_at | OMD | -0,04928565 | 2,92702915 | -0,78294903 | 0,44623507 | 0,68591643 |
| 81626_at | SHCBP1L | -0,05451476 | 3,05168232 | -0,78294186 | 0,44623915 | 0,68591643 |
| 154822_at | LINC00689 | 0,06858033 | 4,22722471 | 0,78268345 | 0,4463862 | 0,68608684 |
| 5197_at | PF4V1 | 0,06639619 | 3,70124945 | 0,78260579 | 0,4464304 | 0,68608684 |
| 80319_at | CXXC4 | 0,08569656 | 4,98468228 | 0,78252432 | 0,44647677 | 0,68608684 |
| 79074_at | C2orf49 | 0,11031738 | 6,89766093 | 0,78244064 | 0,4465244 | 0,68608684 |
| 1420_at | CRYGC | -0,07600428 | 4,87598491 | -0,78243415 | 0,4465281 | 0,68608684 |
| 3652_at | IPP | 0,07645552 | 6,02158443 | 0,78218189 | 0,44667171 | 0,68623747 |
| 101928401_a | LOC10192840 | -0,09500257 | 3,55678305 | -0,7821368 | 0,44669738 | 0,68623747 |
| 122402_at | TDRD9 | 0,06851673 | 3,18743622 | 0,7820025 | 0,44677385 | 0,68630021 |
| 101927994_a | LINC01087 | 0,081505 | 3,05653801 | 0,7818551 | 0,44685779 | 0,68637441 |
| 54839_at | LRRC49 | -0,12083087 | 5,94278432 | -0,7817534 | 0,44691571 | 0,68640865 |
| 144809_at | FAM216B | -0,06902644 | 3,08920307 | -0,7815136 | 0,4470523 | 0,68641203 |
| 54887_at | UHRF1BP1 | 0,19351869 | 7,44001186 | 0,78148155 | 0,44707056 | 0,68641203 |

| | | | | | | |
|---|---|---|---|---|---|---|
| 79400_at | NOX5 | -0,08938403 | 5,18204395 | -0,78144371 | 0,44709211 | 0,68641203 |
| 64211_at | LHX5 | -0,10493581 | 4,99658115 | -0,78135028 | 0,44714534 | 0,68641203 |
| 79173_at | C19orf57 | -0,06326739 | 6,43728901 | -0,78134322 | 0,44714937 | 0,68641203 |
| 9497_at | SLC4A7 | -0,15627194 | 6,12120647 | -0,78129769 | 0,4471753 | 0,68641203 |
| 200081_at | TXLNA | -0,07503151 | 8,13416716 | -0,78121453 | 0,44722269 | 0,68641203 |
| 79140_at | CCDC28B | 0,06008699 | 5,7459862 | 0,78118682 | 0,44723848 | 0,68641203 |
| 84193_at | SETD3 | -0,05141715 | 7,57854579 | -0,78118658 | 0,44723861 | 0,68641203 |
| 501_at | ALDH7A1 | 0,38136106 | 4,52427524 | 0,78111667 | 0,44727845 | 0,68641848 |
| 3889_at | KRT83 | -0,06730269 | 5,41363582 | -0,78099096 | 0,44735009 | 0,68643827 |
| 133688_at | UGT3A1 | -0,06983929 | 3,78069959 | -0,78095357 | 0,4473714 | 0,68643827 |
| 104472718_a | LINC00508 | -0,13761684 | 4,07879216 | -0,7808592 | 0,44742518 | 0,68643827 |
| 54540_at | FAM193B | 0,13138583 | 7,397575 | 0,78084393 | 0,44743389 | 0,68643827 |
| 57152_at | SLURP1 | -0,05332698 | 4,77560675 | -0,78073871 | 0,44749386 | 0,68644357 |
| 23379_at | ICE1 | 0,06948775 | 8,61758647 | 0,78071283 | 0,44750861 | 0,68644357 |
| 27159_at | CHIA | -0,06315541 | 4,48180698 | -0,78041637 | 0,44767762 | 0,6864499 |
| 488_at | ATP2A2 | -0,06723258 | 9,41056283 | -0,78041625 | 0,44767769 | 0,6864499 |
| 3815_at | KIT | 0,28484981 | 4,73740126 | 0,78040495 | 0,44768413 | 0,6864499 |
| 219844_at | HYLS1 | -0,22223213 | 6,81371608 | -0,78039322 | 0,44769082 | 0,6864499 |
| 56127_at | PCDHB9 | 0,06123463 | 5,03979373 | 0,78037992 | 0,4476984 | 0,6864499 |
| 100996570_a | NFIA-AS2 | 0,05208097 | 3,62015437 | 0,78032448 | 0,44773002 | 0,6864499 |
| 285758_at | LINC01268 | -0,09486765 | 3,2438325 | -0,78026806 | 0,44776219 | 0,6864499 |
| 57149_at | LYRM1 | -0,05303202 | 8,29309365 | -0,77997318 | 0,44793035 | 0,68665306 |
| 389376_at | SFTA2 | -0,09574562 | 4,20135399 | -0,77976738 | 0,44804775 | 0,68674657 |
| 100506475_a | EGFLAM-AS2 | -0,0538307 | 3,42091983 | -0,77974124 | 0,44806265 | 0,68674657 |
| 9921_at | RNF10 | 0,06070169 | 8,75044588 | 0,77946825 | 0,44821841 | 0,68681488 |
| 100505576_a | LINC00672 | -0,10163323 | 4,3667986 | -0,77943729 | 0,44823607 | 0,68681488 |
| 64689_at | GORASP1 | 0,12619752 | 7,67145955 | 0,77941883 | 0,44824661 | 0,68681488 |
| 576_at | ADGRB2 | -0,11563407 | 5,80731811 | -0,77936192 | 0,44827908 | 0,68681488 |
| 57732_at | ZFYVE28 | -0,08859527 | 5,43104026 | -0,77930981 | 0,44830882 | 0,68681488 |
| 115509_at | ZNF689 | -0,10368419 | 7,78910328 | -0,7792882 | 0,44832115 | 0,68681488 |
| 8638_at | OASL | 0,70230946 | 7,45090085 | 0,77856106 | 0,44873623 | 0,68737225 |
| 22866_at | CNKSR2 | -0,05088075 | 3,20872499 | -0,77852583 | 0,44875634 | 0,68737225 |
| 317762_at | CCDC85C | -0,06767634 | 7,82688218 | -0,77833362 | 0,44886611 | 0,68745922 |
| 338653_at | KCNQ1-AS1 | -0,09085884 | 4,49284921 | -0,77826758 | 0,44890383 | 0,68745922 |
| 724087_at | LINC01545 | -0,0508631 | 3,85395911 | -0,77811126 | 0,44899311 | 0,68745922 |
| 79956_at | ERMP1 | 0,0943089 | 10,702924 | 0,77810241 | 0,44899817 | 0,68745922 |
| 259296_at | TAS2R50 | 0,10860128 | 3,35977812 | 0,77802444 | 0,44904271 | 0,68745922 |
| 400500_at | BCAR4 | 0,08796214 | 3,78887671 | 0,77801546 | 0,44904784 | 0,68745922 |
| 998_at | CDC42 | 0,04098705 | 5,38832378 | 0,77793322 | 0,44909482 | 0,68745922 |
| 55084_at | SOBP | -0,11960866 | 5,1203152 | -0,77792655 | 0,44909863 | 0,68745922 |
| 414245_at | DNAJC9-AS1 | -0,0952176 | 4,45500167 | -0,77783011 | 0,44915373 | 0,68748894 |
| 7450_at | VWF | -0,06924548 | 4,65130475 | -0,7777022 | 0,44922681 | 0,68754617 |
| 149465_at | CFAP57 | -0,1140794 | 4,35831411 | -0,77750769 | 0,44933796 | 0,6876572 |
| 285550_at | FAM200B | -0,05929713 | 6,15503319 | -0,77744403 | 0,44937434 | 0,6876572 |
| 157695_at | TDRP | 0,25217864 | 5,02945544 | 0,77738785 | 0,44940645 | 0,6876572 |
| 5629_at | PROX1 | 0,05460849 | 3,78515667 | 0,77731569 | 0,44944769 | 0,68766568 |
| 9369_at | NRXN3 | 0,5322942 | 5,79045325 | 0,77698759 | 0,44963525 | 0,68788583 |
| 8436_at | SDPR | 0,05981429 | 3,80443945 | 0,77693905 | 0,449663 | 0,68788583 |
| 2331_at | FMOD | -0,06829122 | 4,735581 | -0,776854 | 0,44971163 | 0,68790559 |
| 283659_at | PRTG | -0,05358509 | 3,36845844 | -0,77666864 | 0,44981763 | 0,6879546 |

| | | | | | | |
|---|---|---|---|---|---|---|
| 79762_at | C1orf115 | 0,43370634 | 5,74537239 | 0,77661998 | 0,44984545 | 0,6879546 |
| 2302_at | FOXJ1 | -0,10639619 | 4,69212018 | -0,77654257 | 0,44988973 | 0,6879546 |
| 2980_at | GUCA2A | -0,15670998 | 5,048242 | -0,77649322 | 0,44991795 | 0,6879546 |
| 63910_at | SLC17A9 | -0,06878713 | 5,54833555 | -0,776455 | 0,44993981 | 0,6879546 |
| 10983_at | CCNI | 0,1837199 | 10,8972822 | 0,77641854 | 0,44996067 | 0,6879546 |
| 83856_at | FSD1L | 0,03656444 | 3,0631624 | 0,77636085 | 0,44999366 | 0,6879546 |
| 51168_at | MYO15A | 0,06163296 | 4,59053041 | 0,77616804 | 0,45010396 | 0,68804752 |
| 283554_at | GPR137C | 0,25474571 | 4,50482548 | 0,77612973 | 0,45012588 | 0,68804752 |
| 150726_at | FBXO41 | 0,14404621 | 6,16118116 | 0,7760566 | 0,45016772 | 0,68805687 |
| #N/A | CTB-12O2,1 | 0,08741308 | 3,40630991 | 0,77598165 | 0,4502106 | 0,68806782 |
| 26284_at | ERAL1 | -0,15453319 | 8,56898392 | -0,77585092 | 0,45028541 | 0,68812756 |
| 1536_at | CYBB | -0,06205963 | 3,94462507 | -0,77574404 | 0,45034657 | 0,68816643 |
| 55191_at | NADSYN1 | 0,17433329 | 8,63105447 | 0,77557124 | 0,45044547 | 0,68826296 |
| 4775_at | NFATC3 | -0,07589211 | 8,04160051 | -0,77545659 | 0,4505111 | 0,68830863 |
| 55731_at | FAM222B | 0,05727507 | 7,49277061 | 0,77530433 | 0,45059826 | 0,68836545 |
| 56999_at | ADAMTS9 | -0,07524887 | 3,69544358 | -0,77520075 | 0,45065756 | 0,68836545 |
| 326_at | AIRE | -0,08694164 | 5,90180644 | -0,77515983 | 0,45068099 | 0,68836545 |
| 101928495_a | LOC10192849 | 0,07842051 | 3,13805573 | 0,77514195 | 0,45069122 | 0,68836545 |
| 917_at | CD3G | -0,10906226 | 3,86609442 | -0,77480706 | 0,450883 | 0,6885541 |
| 84066_at | TEX35 | -0,06362126 | 3,91698004 | -0,77480142 | 0,45088623 | 0,6885541 |
| 151254_at | ALS2CR11 | -0,04674544 | 2,79539831 | -0,77462445 | 0,45098759 | 0,68865429 |
| 84618_at | NT5C1A | -0,0897878 | 5,46655574 | -0,77444922 | 0,45108797 | 0,68875297 |
| 9248_at | GPR50 | -0,06446848 | 3,84270469 | -0,77418858 | 0,4512373 | 0,68892637 |
| 101929261_a | LOC10192926 | -0,04839745 | 3,13599847 | -0,77409926 | 0,45128848 | 0,68894991 |
| 200373_at | CFAP221 | -0,11014879 | 3,98124264 | -0,77396612 | 0,45136478 | 0,68901178 |
| 256309_at | CCDC110 | 0,04744841 | 2,7951877 | 0,77387974 | 0,45141428 | 0,68903275 |
| 157869_at | SBSPON | -0,05282702 | 3,43632632 | -0,77381231 | 0,45145293 | 0,68903715 |
| 23630_at | KCNE5 | -0,11998111 | 5,07077672 | -0,77374356 | 0,45149234 | 0,6890427 |
| 101927774_a | LINC00824 | -0,09510498 | 4,44470679 | -0,77336556 | 0,45170904 | 0,6893188 |
| 5238_at | PGM3 | 0,12443219 | 7,32378341 | 0,77324003 | 0,45178102 | 0,68932325 |
| 199777_at | ZNF626 | -0,0372979 | 3,26450086 | -0,77323336 | 0,45178484 | 0,68932325 |
| 11182_at | SLC2A6 | -0,20128708 | 7,7359332 | -0,77313315 | 0,45184231 | 0,68932325 |
| 8787_at | RGS9 | -0,05222983 | 3,88184147 | -0,77302703 | 0,45190317 | 0,68932325 |
| 5892_at | RAD51D | -0,09025871 | 6,09898125 | -0,77302 | 0,4519072 | 0,68932325 |
| 100130519_a | TMEM221 | -0,0630069 | 6,34052051 | -0,77298607 | 0,45192666 | 0,68932325 |
| 4250_at | SCGB2A2 | 0,04625184 | 2,96359998 | 0,77251912 | 0,45219453 | 0,68960205 |
| 64478_at | CSMD1 | -0,05213601 | 3,78012891 | -0,7724109 | 0,45225662 | 0,68960205 |
| 100506834_a | LOC10050683 | 0,04277693 | 2,90887962 | 0,77240199 | 0,45226174 | 0,68960205 |
| 92017_at | SNX29 | 0,05938087 | 4,31775685 | 0,77239723 | 0,45226447 | 0,68960205 |
| 1804_at | DPP6 | -0,06893412 | 4,56423433 | -0,77235545 | 0,45228844 | 0,68960205 |
| 54849_at | DEF8 | -0,20983161 | 7,99619177 | -0,771999 | 0,45249301 | 0,68985935 |
| 57558_at | USP35 | 0,08819199 | 6,57763324 | 0,77183396 | 0,45258775 | 0,68994919 |
| 79858_at | NEK11 | 0,16009598 | 5,36819864 | 0,77166727 | 0,45268345 | 0,69004046 |
| 2055_at | CLN8 | -0,07419594 | 6,4004423 | -0,77145449 | 0,45280563 | 0,69017208 |
| 3778_at | KCNMA1 | -0,05492474 | 3,45170536 | -0,77137313 | 0,45285235 | 0,69018869 |
| 51655_at | RASD1 | -0,09593783 | 6,01027474 | -0,77126792 | 0,45291277 | 0,69021789 |
| 2650_at | GCNT1 | -0,15756192 | 6,80007651 | -0,77109875 | 0,45300994 | 0,69021789 |
| 10570_at | DPYSL4 | -0,12514109 | 4,91669442 | -0,7710948 | 0,45301221 | 0,69021789 |
| 84867_at | PTPN5 | -0,0707439 | 4,48237241 | -0,77109022 | 0,45301484 | 0,69021789 |
| 5806_at | PTX3 | -1,03252468 | 6,41274957 | -0,77066279 | 0,4532604 | 0,69051302 |

| | | | | | | |
|---|---|---|---|---|---|---|
| 3887_at | KRT81 | -0,10172859 | 5,52583181 | -0,77062827 | 0,45328024 | 0,69051302 |
| 101927586_a | LINC01271 | -0,08262734 | 3,8386856 | -0,77046839 | 0,45337211 | 0,69052783 |
| 2978_at | GUCA1A | -0,06048446 | 4,68488455 | -0,77046239 | 0,45337556 | 0,69052783 |
| 100506428_a | CBR3-AS1 | 0,25386908 | 7,742433 | 0,77038453 | 0,4534203 | 0,69052783 |
| 64080_at | RBKS | 0,17800891 | 5,94275348 | 0,77036184 | 0,45343334 | 0,69052783 |
| 253017_at | TECRL | 0,04264497 | 2,6377393 | 0,77016016 | 0,45354927 | 0,69061315 |
| 6554_at | SLC10A1 | -0,09918322 | 4,11831834 | -0,77011441 | 0,45357557 | 0,69061315 |
| 80094_at | BIN3-IT1 | -0,06279724 | 4,86946833 | -0,77007725 | 0,45359693 | 0,69061315 |
| 143678_at | C11orf94 | 0,06992759 | 5,2511529 | 0,76994317 | 0,45367401 | 0,69067592 |
| 79788_at | ZNF665 | 0,10384144 | 5,22899445 | 0,76974634 | 0,45378718 | 0,69075185 |
| 378_at | ARF4 | 0,11547487 | 11,2682819 | 0,7697317 | 0,4537956 | 0,69075185 |
| 57828_at | CATSPERG | -0,06721428 | 4,39963374 | -0,76962972 | 0,45385424 | 0,69078653 |
| 56978_at | PRDM8 | -0,08328383 | 5,14979113 | -0,76938128 | 0,45399713 | 0,69094939 |
| 170384_at | FUT11 | -0,2633045 | 6,71459339 | -0,76931894 | 0,45403299 | 0,69094939 |
| 408263_at | FNDC9 | -0,0523574 | 3,70735333 | -0,76922336 | 0,45408797 | 0,69095527 |
| 54867_at | TMEM214 | -0,04580686 | 8,53911549 | -0,76913354 | 0,45413964 | 0,69095527 |
| 101927125_a | ELFN1-AS1 | -0,11926525 | 5,05241109 | -0,76909768 | 0,45416027 | 0,69095527 |
| 54829_at | ASPN | -0,0432947 | 3,18564142 | -0,7689566 | 0,45424144 | 0,69095527 |
| 4322_at | MMP13 | 0,31204668 | 6,45812977 | 0,76892512 | 0,45425956 | 0,69095527 |
| 7173_at | TPO | -0,08977512 | 4,21524213 | -0,76886975 | 0,45429141 | 0,69095527 |
| 100996671_a | LOC10099667 | -0,08096606 | 3,49597963 | -0,76884748 | 0,45430423 | 0,69095527 |
| 1824_at | DSC2 | -0,1801591 | 5,60045032 | -0,76881347 | 0,4543238 | 0,69095527 |
| 8527_at | DGKD | -0,05816528 | 7,93293382 | -0,7686613 | 0,45441137 | 0,69103389 |
| 11314_at | CD300A | -0,0720461 | 4,24246675 | -0,7685101 | 0,45449839 | 0,69106415 |
| 1024_at | CDK8 | 0,09390154 | 8,0081839 | 0,76850063 | 0,45450384 | 0,69106415 |
| 6691_at | SPINK2 | 0,16050137 | 4,8148919 | 0,76838781 | 0,45456878 | 0,69106415 |
| 27284_at | SULT1B1 | -0,10358274 | 3,77093715 | -0,76833281 | 0,45460044 | 0,69106415 |
| 116449_at | CLNK | -0,06606047 | 3,79815654 | -0,76831508 | 0,45461065 | 0,69106415 |
| 101928370_a | LOC10192837 | -0,08211123 | 3,83379319 | -0,76816597 | 0,4546965 | 0,69114011 |
| 132204_at | SYNPR | -0,08940462 | 3,69984386 | -0,76758065 | 0,45503356 | 0,69156314 |
| 23338_at | JADE2 | 0,05911787 | 8,07183128 | 0,76755801 | 0,45504661 | 0,69156314 |
| 1850_at | DUSP8 | -0,09283022 | 6,80758064 | -0,76738645 | 0,45514543 | 0,69165876 |
| 23209_at | MLC1 | -0,05940067 | 4,79142095 | -0,76729866 | 0,45519601 | 0,69168105 |
| 6439_at | SFTPB | -0,08568912 | 5,23593715 | -0,76706582 | 0,45533017 | 0,69183034 |
| 50649_at | ARHGEF4 | -0,13714853 | 5,50397133 | -0,76699445 | 0,45537129 | 0,69183826 |
| 101929407_a | PTPRD-AS1 | 0,08030228 | 2,94026658 | 0,7665772 | 0,45561178 | 0,69214903 |
| 342527_at | SMTNL2 | -0,07769053 | 4,7511183 | -0,76609839 | 0,45588785 | 0,69246475 |
| 8200_at | GDF5 | -0,0799495 | 5,22163138 | -0,76608343 | 0,45589648 | 0,69246475 |
| 5297_at | PI4KA | 0,27958055 | 8,37340089 | 0,76602972 | 0,45592745 | 0,69246475 |
| 55515_at | ASIC4 | -0,09666043 | 5,45717114 | -0,76562063 | 0,45616342 | 0,69272441 |
| 140771_at | SMCR5 | -0,06674234 | 3,83295959 | -0,76560863 | 0,45617034 | 0,69272441 |
| 401067_at | IQCF3 | -0,06831737 | 4,1352709 | -0,76544252 | 0,45626617 | 0,69281533 |
| 23241_at | PACS2 | 0,04526449 | 8,115341 | 0,76525566 | 0,45637399 | 0,69289293 |
| 57587_at | CFAP97 | -0,0600938 | 4,81531841 | -0,76522928 | 0,45638922 | 0,69289293 |
| 24142_at | NAT6 | -0,05332227 | 6,32443742 | -0,76512829 | 0,45644749 | 0,69291687 |
| 79895_at | ATP8B4 | 0,05167909 | 3,09381532 | 0,76507729 | 0,45647693 | 0,69291687 |
| 54734_at | RAB39A | -0,08350562 | 4,1316598 | -0,76443937 | 0,45684519 | 0,69340411 |
| 57496_at | MKL2 | 0,04332839 | 7,76094142 | 0,7643504 | 0,45689657 | 0,69340411 |
| 386678_at | KRTAP10-11 | -0,09320568 | 5,16329301 | -0,76429199 | 0,4569303 | 0,69340411 |
| 406988_at | MIR205 | -0,36071832 | 6,81704199 | -0,7642719 | 0,4569419 | 0,69340411 |

| | | | | | | |
|---|---|---|---|---|---|---|
| 23332_at | CLASP1 | 0,07120064 | 8,59999095 | 0,76411072 | 0,45703499 | 0,69346227 |
| 55366_at | LGR4 | 0,07419848 | 6,86641886 | 0,76408088 | 0,45705222 | 0,69346227 |
| 100506207_a | LOC10050620 | 0,05327905 | 3,58779815 | 0,76401328 | 0,45709127 | 0,69346689 |
| 84517_at | ACTRT3 | 0,29320474 | 5,46658104 | 0,76394226 | 0,45713229 | 0,69347451 |
| 92806_at | CENPBD1 | 0,14176703 | 6,85039496 | 0,76372063 | 0,45726033 | 0,69361412 |
| 79777_at | ACBD4 | 0,1214674 | 5,75900929 | 0,76338777 | 0,45745266 | 0,69385123 |
| 642976_at | GRIK1-AS1 | 0,07184145 | 3,29277224 | 0,76325003 | 0,45753227 | 0,69391734 |
| 7341_at | SUMO1 | 0,15483008 | 7,3547695 | 0,76301239 | 0,45766964 | 0,69407103 |
| 23302_at | WSCD1 | -0,0806712 | 4,70228653 | -0,76293207 | 0,45771607 | 0,6940868 |
| 439927_at | LINC01555 | 0,15382193 | 4,24497661 | 0,76261353 | 0,45790024 | 0,6942587 |
| #N/A | GS1-124K5,4 | -0,10699053 | 6,22966353 | -0,76260857 | 0,45790311 | 0,6942587 |
| 64834_at | ELOVL1 | -0,07755969 | 10,6508499 | -0,762549 | 0,45793755 | 0,6942587 |
| 2134_at | EXTL1 | -0,07611846 | 5,31736492 | -0,7624369 | 0,45800238 | 0,69430234 |
| 320_at | APBA1 | 0,10208221 | 5,10215896 | 0,76233815 | 0,4580595 | 0,69433428 |
| 3816_at | KLK1 | -0,07357014 | 5,85795842 | -0,7621185 | 0,45818655 | 0,69433809 |
| 85569_at | GALP | 0,06949436 | 3,79599568 | 0,7621095 | 0,45819176 | 0,69433809 |
| 80198_at | MUS81 | -0,08657739 | 8,78342159 | -0,76208204 | 0,45820764 | 0,69433809 |
| 10742_at | RAI2 | 0,25612284 | 5,47572408 | 0,76206691 | 0,4582164 | 0,69433809 |
| 5359_at | PLSCR1 | 0,28852219 | 9,69038429 | 0,76202225 | 0,45824223 | 0,69433809 |
| 30814_at | PLA2G2E | -0,09794413 | 5,40847427 | -0,76172662 | 0,45841328 | 0,69445689 |
| 220359_at | TIGD3 | -0,08020352 | 6,07550464 | -0,76170603 | 0,4584252 | 0,69445689 |
| 133482_at | SLCO6A1 | 0,13161693 | 3,39098635 | 0,76169981 | 0,45842879 | 0,69445689 |
| 55245_at | UQCC1 | -0,11661819 | 7,43489843 | -0,76148793 | 0,45855141 | 0,6945455 |
| 115795_at | FANCD2OS | -0,09710951 | 4,99065217 | -0,76138154 | 0,45861299 | 0,6945455 |
| 9702_at | CEP57 | -0,05666431 | 7,98352992 | -0,76135862 | 0,45862626 | 0,6945455 |
| 4437_at | MSH3 | -0,15103906 | 7,0480224 | -0,76134955 | 0,45863151 | 0,6945455 |
| #N/A | HGC6,3 | -0,07462528 | 5,29831494 | -0,76128403 | 0,45866944 | 0,69454833 |
| 9172_at | MYOM2 | 0,10432537 | 4,9426178 | 0,76114609 | 0,45874929 | 0,69457149 |
| 27344_at | PCSK1N | 0,18545816 | 7,32014757 | 0,76113304 | 0,45875684 | 0,69457149 |
| 283571_at | PROX2 | 0,06346373 | 3,42380717 | 0,76106002 | 0,45879912 | 0,6945809 |
| 3653_at | IPW | 0,27812887 | 5,95666418 | 0,76096702 | 0,45885296 | 0,69460783 |
| 2562_at | GABRB3 | 0,1592284 | 4,29454737 | 0,76082859 | 0,45893312 | 0,69466177 |
| 80759_at | KHDC1 | -0,09712519 | 5,39278663 | -0,76078093 | 0,45896072 | 0,69466177 |
| 100507537_a | LOC10050753 | -0,067001 | 3,04337377 | -0,76048317 | 0,45913317 | 0,69482802 |
| 27019_at | DNAI1 | -0,08519575 | 4,70702837 | -0,7604667 | 0,4591427 | 0,69482802 |
| 4130_at | MAP1A | -0,08344832 | 4,42955094 | -0,76032636 | 0,459224 | 0,69489172 |
| 401124_at | DTHD1 | -0,05282209 | 3,02728897 | -0,76026949 | 0,45925694 | 0,69489172 |
| 83698_at | CALN1 | -0,07646839 | 3,86539538 | -0,76014396 | 0,45932967 | 0,69494717 |
| 128408_at | BHLHE23 | -0,0829275 | 4,8762232 | -0,76004164 | 0,45938895 | 0,69498228 |
| 284370_at | ZNF615 | 0,09691516 | 5,90270944 | 0,75978084 | 0,45954008 | 0,69511688 |
| 100505621_a | C11orf72 | -0,11598428 | 6,03215924 | -0,75976356 | 0,45955009 | 0,69511688 |
| 23141_at | ANKLE2 | -0,05651786 | 8,20510066 | -0,75969299 | 0,459591 | 0,69512416 |
| 2813_at | GP2 | -0,08507776 | 4,22953775 | -0,75959643 | 0,45964696 | 0,69515423 |
| 23495_at | TNFRSF13B | -0,08855862 | 6,31618114 | -0,75934378 | 0,45979342 | 0,69530174 |
| 100506302_a | LOC10050630 | -0,14934333 | 5,72157116 | -0,75918382 | 0,45988616 | 0,69530174 |
| 151888_at | BTLA | -0,06757449 | 3,53464636 | -0,75916202 | 0,4598988 | 0,69530174 |
| 91074_at | ANKRD30A | -0,07633736 | 3,12627708 | -0,75911227 | 0,45992764 | 0,69530174 |
| 100996583_a | LOC10099658 | -0,07635142 | 4,32470682 | -0,75907027 | 0,459952 | 0,69530174 |
| 149371_at | EXOC8 | 0,08987589 | 7,54613888 | 0,75905462 | 0,45996107 | 0,69530174 |
| 57128_at | LYRM4 | -0,21913072 | 6,41494339 | -0,75895199 | 0,46002059 | 0,69533715 |

| | | | | | | |
|---|---|---|---|---|---|---|
| 113730_at | KLHDC7B | 0,1966229 | 5,51651646 | 0,75870312 | 0,46016493 | 0,69546995 |
| 4625_at | MYH7 | -0,11361026 | 4,43098605 | -0,758676 | 0,46018066 | 0,69546995 |
| 55363_at | HEMGN | -0,0606005 | 2,84372253 | -0,75841189 | 0,46033387 | 0,69564692 |
| 80019_at | UBTD1 | 0,07579696 | 6,902714 | 0,75819529 | 0,46045954 | 0,69573873 |
| 7593_at | MZF1 | 0,20098957 | 6,88075063 | 0,75818268 | 0,46046686 | 0,69573873 |
| 8224_at | SYN3 | -0,0803923 | 5,10556224 | -0,75790201 | 0,46062974 | 0,69593025 |
| 83607_at | AMMECR1L | -0,09352202 | 8,11216152 | -0,75774133 | 0,460723 | 0,69598333 |
| 137872_at | ADHFE1 | -0,12618229 | 6,23946301 | -0,75771698 | 0,46073713 | 0,69598333 |
| 196792_at | FAM24B | -0,11221711 | 5,82639794 | -0,75761622 | 0,46079562 | 0,6960171 |
| 10512_at | SEMA3C | 0,21044302 | 9,26688829 | 0,75745376 | 0,46088994 | 0,69610498 |
| 157927_at | C9orf62 | -0,07087305 | 5,5606052 | -0,75720602 | 0,46103379 | 0,69626481 |
| 286151_at | FBXO43 | 0,06764623 | 4,083276 | 0,75713573 | 0,46107461 | 0,69626481 |
| 81833_at | SPACA1 | 0,04871914 | 3,23286965 | 0,75700643 | 0,4611497 | 0,69626481 |
| 26515_at | TIMM10B | 0,07844502 | 7,45093495 | 0,75697446 | 0,46116827 | 0,69626481 |
| 26094_at | DCAF4 | 0,08781479 | 7,3074021 | 0,75696031 | 0,46117649 | 0,69626481 |
| 83707_at | TRPT1 | 0,07296913 | 8,07514097 | 0,75675001 | 0,46129865 | 0,69630912 |
| 78990_at | OTUB2 | 0,09296617 | 5,94193211 | 0,75673292 | 0,46130857 | 0,69630912 |
| 5768_at | QSOX1 | 0,11248011 | 9,54143559 | 0,7567231 | 0,46131428 | 0,69630912 |
| 23173_at | METAP1 | -0,16492459 | 9,77842208 | -0,75664504 | 0,46135963 | 0,696323 |
| 92749_at | DRC1 | -0,07479645 | 4,10924445 | -0,75654737 | 0,46141637 | 0,69635409 |
| 222194_at | RSBN1L | 0,10614106 | 6,77405423 | 0,75645698 | 0,46146889 | 0,69637879 |
| 90333_at | ZNF468 | -0,07210945 | 7,99667378 | -0,75583189 | 0,46183218 | 0,69677567 |
| 284338_at | PRR19 | -0,07388296 | 5,44981995 | -0,75551718 | 0,46201516 | 0,69677567 |
| 51227_at | PIGP | 0,16421378 | 9,12601453 | 0,75546369 | 0,46204626 | 0,69677567 |
| 26038_at | CHD5 | -0,08598919 | 4,59126477 | -0,75540316 | 0,46208145 | 0,69677567 |
| 5449_at | POU1F1 | 0,04530244 | 2,7786171 | 0,75532431 | 0,46212731 | 0,69677567 |
| 23532_at | PRAME | -0,07218094 | 4,25119256 | -0,75532326 | 0,46212792 | 0,69677567 |
| 7150_at | TOP1 | -0,1622023 | 9,8363952 | -0,75529839 | 0,46214238 | 0,69677567 |
| 57574_at | MARCH4 | -0,0806154 | 5,07218812 | -0,7552951 | 0,4621443 | 0,69677567 |
| 28955_at | DEXI | -0,1111442 | 8,49301361 | -0,7552587 | 0,46216546 | 0,69677567 |
| 3383_at | ICAM1 | 0,05590236 | 5,60800228 | 0,75525395 | 0,46216823 | 0,69677567 |
| 25854_at | FAM149A | 0,43750002 | 5,4467357 | 0,75512153 | 0,46224525 | 0,69677567 |
| 138428_at | PTRH1 | -0,13194871 | 6,81017194 | -0,75511338 | 0,46224998 | 0,69677567 |
| 341640_at | FREM2 | 0,06314792 | 3,21908834 | 0,75503877 | 0,46229339 | 0,69677567 |
| 728978_at | UNC5B-AS1 | 0,18773677 | 5,59951504 | 0,75502061 | 0,46230395 | 0,69677567 |
| 83882_at | TSPAN10 | -0,10502147 | 5,70266647 | -0,75501134 | 0,46230934 | 0,69677567 |
| 23132_at | RAD54L2 | -0,04418284 | 7,15616267 | -0,7549007 | 0,4623737 | 0,69677567 |
| 124044_at | SPATA2L | 0,06673938 | 7,41649894 | 0,75488863 | 0,46238073 | 0,69677567 |
| 4747_at | NEFL | -0,05052664 | 3,62553425 | -0,75488475 | 0,46238298 | 0,69677567 |
| 8895_at | CPNE3 | -0,13784447 | 10,207285 | -0,75464488 | 0,46252254 | 0,69680266 |
| 645188_at | LOC645188 | -0,05842466 | 3,05956041 | -0,75462881 | 0,46253189 | 0,69680266 |
| 1392_at | CRH | -0,05029571 | 3,55806646 | -0,75460863 | 0,46254363 | 0,69680266 |
| 643355_at | LOC643355 | -0,08642752 | 3,80880341 | -0,75460527 | 0,46254559 | 0,69680266 |
| 121457_at | IKBIP | -0,19778165 | 6,49463566 | -0,7544097 | 0,4626594 | 0,69680378 |
| 7024_at | TFCP2 | -0,0489828 | 7,57763033 | -0,7543982 | 0,46266609 | 0,69680378 |
| 55356_at | SLC22A15 | 0,22574392 | 5,88812286 | 0,754362 | 0,46268716 | 0,69680378 |
| 101927849_a | LOC10192784 | -0,06853862 | 4,20849574 | -0,75435536 | 0,46269102 | 0,69680378 |
| 54862_at | CC2D1A | 0,04758934 | 6,86759291 | 0,75413127 | 0,46282145 | 0,69694572 |
| 4804_at | NGFR | -0,0640891 | 4,65577411 | -0,75371557 | 0,46306346 | 0,69725565 |
| 26065_at | LSM14A | 0,06943646 | 9,52605652 | 0,75347977 | 0,46320078 | 0,69729611 |

| | | | | | | |
|---|---|---|---|---|---|---|
| 100130733_a | LRRC70 | -0,06304002 | 3,14223958 | -0,75345007 | 0,46321808 | 0,69729611 |
| 57495_at | NWD2 | -0,07594821 | 3,41001075 | -0,75341019 | 0,4632413 | 0,69729611 |
| 1948_at | EFNB2 | -0,19914326 | 8,55766784 | -0,75336515 | 0,46326754 | 0,69729611 |
| 7350_at | UCP1 | 0,0634791 | 3,7524446 | 0,75324455 | 0,46333778 | 0,69729611 |
| 54516_at | MTRF1L | -0,06030873 | 3,50737432 | -0,75321838 | 0,46335303 | 0,69729611 |
| 102724156_a | LOC10272415 | 0,1000378 | 4,96148161 | 0,75321453 | 0,46335527 | 0,69729611 |
| 814_at | CAMK4 | -0,07306261 | 4,00269821 | -0,7531722 | 0,46337993 | 0,69729611 |
| 8929_at | PHOX2B | -0,05975274 | 4,43171919 | -0,75278509 | 0,46360546 | 0,697581 |
| 4914_at | NTRK1 | -0,15398395 | 4,51085668 | -0,75269284 | 0,46365922 | 0,6976074 |
| 79152_at | FA2H | 0,04754595 | 6,11663263 | 0,75255488 | 0,46373962 | 0,69766812 |
| 8852_at | AKAP4 | -0,06513895 | 3,77283624 | -0,75249326 | 0,46377553 | 0,69766812 |
| 50615_at | IL21R | -0,06843734 | 4,16986771 | -0,75243716 | 0,46380823 | 0,69766812 |
| 1819_at | DRG2 | -0,24539094 | 8,62234904 | -0,75237217 | 0,46384611 | 0,69767062 |
| 84691_at | FAM71F1 | -0,06169358 | 3,73144942 | -0,75202221 | 0,46405013 | 0,69786645 |
| 401585_at | LOC401585 | -0,05699871 | 5,3168542 | -0,75200962 | 0,46405747 | 0,69786645 |
| 150379_at | PNPLA5 | -0,08243361 | 5,02676022 | -0,75196241 | 0,46408499 | 0,69786645 |
| 100616668_a | TPTE2P5 | 0,06255556 | 3,76441985 | 0,75160873 | 0,46429124 | 0,69802827 |
| 340348_at | TSPAN33 | 0,07403167 | 6,75199732 | 0,75154093 | 0,46433078 | 0,69802827 |
| 100294362_a | LOC10029436 | -0,07624022 | 3,84713568 | -0,75150645 | 0,4643509 | 0,69802827 |
| 390_at | RND3 | 0,16045168 | 10,1429893 | 0,7514809 | 0,4643658 | 0,69802827 |
| 7439_at | BEST1 | -0,08282381 | 5,14116624 | -0,7514672 | 0,46437379 | 0,69802827 |
| 10418_at | SPON1 | -0,04637812 | 3,62335919 | -0,75133574 | 0,46445048 | 0,69806836 |
| 10663_at | CXCR6 | -0,06389648 | 4,21213082 | -0,75129724 | 0,46447293 | 0,69806836 |
| 79740_at | ZBBX | -0,05290463 | 3,05226472 | -0,75076169 | 0,46478542 | 0,69848351 |
| 55274_at | PHF10 | -0,11516384 | 8,58635169 | -0,74993705 | 0,46526685 | 0,69913766 |
| 55228_at | PNMAL1 | -0,07433447 | 4,15698272 | -0,74984386 | 0,46532127 | 0,69913766 |
| 90639_at | COX19 | 0,04748113 | 5,82918919 | 0,74982962 | 0,46532959 | 0,69913766 |
| 389203_at | SMIM20 | -0,10480526 | 8,18071763 | -0,74976598 | 0,46536676 | 0,69913897 |
| 25937_at | WWTR1 | 0,08850372 | 8,29092402 | 0,74965195 | 0,46543336 | 0,69918233 |
| 2771_at | GNAI2 | 0,25231312 | 9,58691509 | 0,74959228 | 0,46546822 | 0,69918233 |
| 7224_at | TRPC5 | -0,05392761 | 3,17201187 | -0,74911465 | 0,46574727 | 0,69953497 |
| 26268_at | FBXO9 | 0,08485982 | 8,26137812 | 0,74902604 | 0,46579905 | 0,69953497 |
| 80728_at | ARHGAP39 | -0,14358447 | 5,27323483 | -0,74895244 | 0,46584207 | 0,69953497 |
| 80006_at | TRAPPC13 | -0,15813767 | 5,87530384 | -0,74894188 | 0,46584824 | 0,69953497 |
| 54567_at | DLL4 | -0,06543314 | 5,84051346 | -0,74879843 | 0,46593208 | 0,69956983 |
| 339855_at | KY | -0,05371473 | 5,00052806 | -0,74877323 | 0,46594681 | 0,69956983 |
| 163702_at | IFNLR1 | 0,04826427 | 6,27412294 | 0,74871576 | 0,4659804 | 0,69956983 |
| 6351_at | CCL4 | -0,10826226 | 4,57543379 | -0,74851857 | 0,46609567 | 0,69966854 |
| 6252_at | RTN1 | 0,05538517 | 3,64074957 | 0,74841155 | 0,46615824 | 0,69966854 |
| 23129_at | PLXND1 | -0,20408898 | 6,39111014 | -0,74836596 | 0,4661849 | 0,69966854 |
| 400456_at | LINC01197 | 0,04382446 | 3,17887471 | 0,74835477 | 0,46619144 | 0,69966854 |
| 131377_at | KLHL40 | -0,07110793 | 4,19135882 | -0,74827946 | 0,46623548 | 0,69968012 |
| 84561_at | SLC12A8 | 0,12440129 | 6,46374472 | 0,74810051 | 0,46634013 | 0,6997196 |
| 4584_at | MUC3A | -0,07210174 | 5,65098101 | -0,74806061 | 0,46636346 | 0,6997196 |
| 285501_at | LINC01098 | -0,040625 | 3,26881242 | -0,74804812 | 0,46637076 | 0,6997196 |
| 8667_at | EIF3H | -0,05958284 | 11,4104542 | -0,74777738 | 0,46652912 | 0,69985216 |
| 125115_at | KRT40 | -0,04817748 | 3,22091182 | -0,74777283 | 0,46653178 | 0,69985216 |
| 26148_at | C10orf12 | -0,10972481 | 5,21250735 | -0,74765544 | 0,46660045 | 0,69990068 |
| 8874_at | ARHGEF7 | 0,05028168 | 6,52740856 | 0,74730376 | 0,46680621 | 0,70014883 |
| 135644_at | TRIM40 | -0,07720713 | 3,95837688 | -0,74724845 | 0,46683858 | 0,70014883 |

| | | | | | | |
|---|---|---|---|---|---|---|
| 2845_at | GPR22 | -0,08186608 | 2,90494371 | -0,74689434 | 0,46704584 | 0,70040513 |
| 80131_at | LRRC8E | 0,09868454 | 6,42103578 | 0,74672256 | 0,46714639 | 0,70049137 |
| 54329_at | GPR85 | 0,18805097 | 4,78772683 | 0,74667187 | 0,46717607 | 0,70049137 |
| 101927020_a | LOC10192702 | -0,06153655 | 3,51858571 | -0,74658929 | 0,46722442 | 0,70050934 |
| 11212_at | PROSC | -0,15245803 | 9,19395623 | -0,74643148 | 0,46731682 | 0,70058977 |
| 51050_at | PI15 | -0,04349616 | 3,14678415 | -0,74634695 | 0,46736632 | 0,70058977 |
| 56936_at | CCDC177 | -0,10184224 | 5,32888954 | -0,74631134 | 0,46738717 | 0,70058977 |
| 100129175_a | LOC10012917 | -0,05945696 | 4,52059652 | -0,74622078 | 0,46744021 | 0,70061475 |
| 59340_at | HRH4 | -0,04913136 | 3,14803585 | -0,74600482 | 0,4675667 | 0,70074981 |
| 5610_at | EIF2AK2 | 0,16920778 | 9,03136991 | 0,74567989 | 0,46775705 | 0,70098056 |
| 152503_at | SH3D19 | 0,12227705 | 6,77921479 | 0,74553472 | 0,46784211 | 0,70102685 |
| 100293516_a | ZNF587B | -0,32691955 | 6,40495775 | -0,74544941 | 0,4678921 | 0,70102685 |
| 196463_at | PLBD2 | 0,09913456 | 6,9667509 | 0,74541782 | 0,46791062 | 0,70102685 |
| 9811_at | CTIF | 0,0809008 | 6,15206128 | 0,74537875 | 0,46793351 | 0,70102685 |
| 138307_at | LCN8 | -0,06298124 | 4,4977687 | -0,74521406 | 0,46803003 | 0,70111692 |
| 171389_at | NLRP6 | -0,0851022 | 4,46727161 | -0,7450087 | 0,46815041 | 0,70115633 |
| 100506655_a | LCMT1-AS2 | -0,10530369 | 4,22856676 | -0,74500155 | 0,46815459 | 0,70115633 |
| 100652931_a | LOC10065293 | -0,12319432 | 4,08513725 | -0,74498289 | 0,46816554 | 0,70115633 |
| 5793_at | PTPRG | -0,30201604 | 5,87242857 | -0,74486765 | 0,4682331 | 0,70120299 |
| 27309_at | ZNF330 | -0,28482723 | 7,29055204 | -0,74453017 | 0,46843097 | 0,7014448 |
| 101927592_a | LOC10192759 | -0,06436339 | 3,87297791 | -0,74436876 | 0,46852563 | 0,70150002 |
| 147138_at | TMC8 | -0,07340971 | 5,23396708 | -0,74434309 | 0,46854069 | 0,70150002 |
| 100874074_a | DLEU7-AS1 | -0,10061777 | 3,85555889 | -0,74404796 | 0,4687138 | 0,70170467 |
| 116254_at | GINM1 | 0,1543234 | 9,40945676 | 0,74393327 | 0,46878109 | 0,70171636 |
| 83873_at | GPR61 | -0,07748716 | 4,24442264 | -0,74390056 | 0,46880028 | 0,70171636 |
| 10438_at | C1D | -0,14688505 | 7,21691037 | -0,74384837 | 0,4688309 | 0,70171636 |
| 25859_at | PART1 | -0,09791165 | 5,23952667 | -0,74373328 | 0,46889843 | 0,70176291 |
| 10600_at | USP16 | -0,14693981 | 9,26686476 | -0,74362663 | 0,46896101 | 0,7017919 |
| 690_at | BTF3P11 | -0,12471364 | 4,16188861 | -0,74347261 | 0,4690514 | 0,7017919 |
| 6422_at | SFRP1 | 0,16630516 | 5,01069634 | 0,74341861 | 0,46908309 | 0,7017919 |
| 225_at | ABCD2 | 0,0489253 | 2,67478896 | 0,74335416 | 0,46912092 | 0,7017919 |
| 6251_at | RSU1 | 0,12891679 | 8,94400951 | 0,74332472 | 0,4691382 | 0,7017919 |
| 101927543_a | LOC10192754 | 0,05825667 | 3,62796846 | 0,74330315 | 0,46915086 | 0,7017919 |
| 57687_at | VAT1L | -0,08349636 | 3,92391895 | -0,74326573 | 0,46917283 | 0,7017919 |
| 101929229_a | LOC10192922 | -0,08910258 | 3,57266338 | -0,74318652 | 0,46921932 | 0,70180696 |
| 7314_at | UBB | 0,07844337 | 12,2062528 | 0,74300296 | 0,46932708 | 0,70191364 |
| 2960_at | GTF2E1 | -0,08815579 | 8,2594519 | -0,7429368 | 0,46936593 | 0,70191723 |
| 57214_at | CEMIP | -0,06097722 | 4,27656689 | -0,74269113 | 0,46951018 | 0,70203855 |
| 285954_at | INHBA-AS1 | -0,07448459 | 3,56323786 | -0,74263289 | 0,46954438 | 0,70203855 |
| 284835_at | LINC00323 | 0,05003184 | 3,17807935 | 0,74261245 | 0,46955639 | 0,70203855 |
| 8428_at | STK24 | -0,10021864 | 10,3242743 | -0,74246145 | 0,46964507 | 0,70207545 |
| 100303755_a | PET117 | -0,16610926 | 7,02961346 | -0,74244632 | 0,46965396 | 0,70207545 |
| 10393_at | ANAPC10 | -0,24086111 | 5,89983909 | -0,74217966 | 0,4698106 | 0,70225511 |
| 55809_at | TRERF1 | -0,06806222 | 7,22125319 | -0,74194704 | 0,46994727 | 0,70240489 |
| 3764_at | KCNJ8 | -0,08296646 | 4,46776368 | -0,7417847 | 0,47004267 | 0,70249297 |
| 145226_at | RDH12 | -0,11926884 | 5,12439609 | -0,74166204 | 0,47011475 | 0,70253259 |
| 3752_at | KCND3 | -0,06571867 | 4,99789768 | -0,74161548 | 0,47014211 | 0,70253259 |
| 283254_at | HARBI1 | 0,13074434 | 5,35257525 | 0,74132529 | 0,47031268 | 0,70273295 |
| 146212_at | KCTD19 | -0,08241499 | 4,75586727 | -0,74116738 | 0,47040552 | 0,70278654 |
| 6566_at | SLC16A1 | -0,47461589 | 6,15024351 | -0,74114018 | 0,47042151 | 0,70278654 |

| | | | | | | |
|---|---|---|---|---|---|---|
| 51043_at | ZBTB7B | -0,05852645 | 6,51513662 | -0,74085859 | 0,47058709 | 0,70297938 |
| 10960_at | LMAN2 | -0,0674848 | 10,2285804 | -0,74051732 | 0,4707878 | 0,70315823 |
| 100128554_a | LOC10012855 | -0,05057469 | 3,08144801 | -0,74047511 | 0,47081263 | 0,70315823 |
| 56901_at | NDUFA4L2 | -0,08566804 | 5,60822805 | -0,74043687 | 0,47083513 | 0,70315823 |
| 56158_at | TEX12 | 0,04404181 | 2,86877605 | 0,74028037 | 0,4709272 | 0,70315823 |
| 3142_at | HLX | 0,07121987 | 5,66708681 | 0,7402411 | 0,4709503 | 0,70315823 |
| 92129_at | RIPPLY1 | -0,07103356 | 4,04399709 | -0,74023551 | 0,47095359 | 0,70315823 |
| 3861_at | KRT14 | -0,26859492 | 6,41756186 | -0,74021968 | 0,4709629 | 0,70315823 |
| 100131655_a | LOC10013165 | -0,06786009 | 5,20910227 | -0,74008723 | 0,47104084 | 0,70315823 |
| 837_at | CASP4 | 0,09592084 | 8,81580483 | 0,74003841 | 0,47106956 | 0,70315823 |
| 105370941_a | LOC10537094 | 0,11325183 | 4,39190077 | 0,74003453 | 0,47107184 | 0,70315823 |
| 26272_at | FBXO4 | -0,07672434 | 4,33570569 | -0,73973072 | 0,47125064 | 0,7033706 |
| 100505942_a | LOC10050594 | -0,05326177 | 4,65403298 | -0,73959756 | 0,47132902 | 0,70342979 |
| 646736_at | LOC646736 | -0,0523515 | 3,29017151 | -0,7395304 | 0,47136855 | 0,70342979 |
| 101927839_a | LOC10192783 | -0,08431405 | 4,71635444 | -0,73947725 | 0,47139984 | 0,70342979 |
| 734_at | OSGIN2 | 0,16918491 | 7,63079704 | 0,73933553 | 0,47148327 | 0,70349979 |
| 196047_at | EMX2OS | 0,1482423 | 4,92309339 | 0,73914844 | 0,47159343 | 0,70359495 |
| 50833_at | TAS2R16 | -0,07807937 | 3,26873788 | -0,73910315 | 0,4716201 | 0,70359495 |
| 83660_at | TLN2 | 0,04636645 | 5,00719579 | 0,73880279 | 0,47179698 | 0,70380433 |
| 221092_at | HNRNPUL2 | -0,075745 | 9,24374584 | -0,73867371 | 0,47187301 | 0,70386324 |
| 100129858_a | SCOC-AS1 | 0,05881757 | 4,1045557 | 0,73860525 | 0,47191335 | 0,7038689 |
| 145376_at | PPP1R36 | -0,06650806 | 4,86613307 | -0,73849007 | 0,4719812 | 0,70389324 |
| 121643_at | FOXN4 | -0,07396807 | 5,6254659 | -0,73845349 | 0,47200275 | 0,70389324 |
| 162962_at | ZNF836 | 0,09134127 | 4,0082447 | 0,73837378 | 0,47204971 | 0,70390877 |
| 81889_at | FAHD1 | -0,13305441 | 8,21727818 | -0,73830896 | 0,4720879 | 0,70391123 |
| 389170_at | LEKR1 | 0,09523897 | 4,03637226 | 0,73786426 | 0,47234997 | 0,70424748 |
| 344018_at | FIGLA | -0,07475799 | 3,59534156 | -0,73773697 | 0,472425 | 0,70430483 |
| 29085_at | PHPT1 | 0,05508541 | 9,06575661 | 0,73756474 | 0,47252653 | 0,70440168 |
| 1816_at | DRD5 | -0,08957007 | 5,08510788 | -0,73741808 | 0,472613 | 0,70443317 |
| 56776_at | FMN2 | -0,05314834 | 3,31918696 | -0,73733618 | 0,47266129 | 0,70443317 |
| 7360_at | UGP2 | -0,07098504 | 9,56173539 | -0,73732421 | 0,47266834 | 0,70443317 |
| 3170_at | FOXA2 | -0,13708983 | 4,9305302 | -0,73723616 | 0,47272027 | 0,70443317 |
| 170691_at | ADAMTS17 | -0,07751168 | 4,70413165 | -0,7372188 | 0,4727305 | 0,70443317 |
| 57159_at | TRIM54 | -0,06650811 | 4,26677742 | -0,7370714 | 0,47281743 | 0,70450821 |
| 283078_at | MKX | 0,04970814 | 3,68741629 | 0,73694335 | 0,47289295 | 0,70456624 |
| 140700_at | SAMD10 | -0,06244607 | 6,4747578 | -0,73675199 | 0,47300583 | 0,70467992 |
| 80032_at | ZNF556 | 0,07021654 | 5,30160196 | 0,73652901 | 0,47313738 | 0,70480192 |
| 84812_at | PLCD4 | 0,10748683 | 4,33341273 | 0,73648915 | 0,4731609 | 0,70480192 |
| 5196_at | PF4 | -0,11835445 | 4,45184761 | -0,7364166 | 0,47320371 | 0,70481118 |
| 493861_at | EID3 | 0,05514038 | 4,42893869 | 0,73616243 | 0,47335369 | 0,70493638 |
| 7046_at | TGFBR1 | 0,07906983 | 6,07819304 | 0,73611943 | 0,47337907 | 0,70493638 |
| 166647_at | ADGRA3 | -0,1533854 | 6,41788977 | -0,73608812 | 0,47339755 | 0,70493638 |
| 286262_at | TPRN | -0,09091997 | 7,42787808 | -0,73599858 | 0,4734504 | 0,70496058 |
| 163479_at | FNDC7 | -0,07770737 | 4,51358971 | -0,73581599 | 0,47355818 | 0,70500391 |
| 85360_at | SYDE1 | -0,08968341 | 6,60284186 | -0,73579291 | 0,47357181 | 0,70500391 |
| 56911_at | MAP3K7CL | 0,09631077 | 4,14229813 | 0,73576329 | 0,47358929 | 0,70500391 |
| 128272_at | ARHGEF19 | 0,2270622 | 7,16232756 | 0,73558078 | 0,47369705 | 0,70510982 |
| 2693_at | GHSR | -0,06735526 | 4,34140578 | -0,73544484 | 0,47377731 | 0,70512442 |
| 79671_at | NLRX1 | 0,11217619 | 6,88876034 | 0,73544018 | 0,47378006 | 0,70512442 |
| 147906_at | DACT3 | -0,05658305 | 6,52629781 | -0,7352216 | 0,47390915 | 0,70526204 |

| | | | | | | |
|---|---|---|---|---|---|---|
| 6911_at | TBX6 | -0,05738563 | 5,709529 | -0,73486307 | 0,47412092 | 0,70552269 |
| 32_at | ACACB | 0,04356844 | 4,82714168 | 0,7346711 | 0,47423433 | 0,70563695 |
| 57191_at | VN1R1 | 0,08571885 | 4,11688701 | 0,73445165 | 0,47436401 | 0,70577538 |
| 100874107_a | MTUS2-AS1 | -0,09666664 | 4,31651304 | -0,73427741 | 0,47446698 | 0,70587407 |
| 23566_at | LPAR3 | 0,14611264 | 5,00141985 | 0,73417047 | 0,47453018 | 0,70591358 |
| 6910_at | TBX5 | -0,0545851 | 4,38472872 | -0,73390065 | 0,47468968 | 0,70609632 |
| 283463_at | MUC19 | -0,11635517 | 3,96717406 | -0,73371871 | 0,47479725 | 0,70613795 |
| 4735_at | SEPT2 | -0,04964763 | 10,5691394 | -0,73371812 | 0,4747976 | 0,70613795 |
| 30835_at | CD209 | -0,08516862 | 4,87160583 | -0,73366731 | 0,47482764 | 0,70613795 |
| 366_at | AQP9 | -0,06730716 | 4,24396214 | -0,73329776 | 0,47504618 | 0,70640841 |
| 25959_at | KANK2 | 0,21814952 | 8,11117299 | 0,73320476 | 0,47510119 | 0,70643568 |
| 56099_at | PCDHGB7 | 0,05155121 | 3,67700565 | 0,7329554 | 0,4752487 | 0,70658666 |
| 5079_at | PAX5 | -0,06832749 | 5,20736636 | -0,73287211 | 0,47529798 | 0,70658666 |
| 55016_at | MARCH1 | -0,04570613 | 3,21925065 | -0,73275498 | 0,47536728 | 0,70658666 |
| 100287765_a | LINC00630 | -0,08032139 | 5,13508309 | -0,73269451 | 0,47540306 | 0,70658666 |
| 50943_at | FOXP3 | -0,05840014 | 4,92527793 | -0,73269404 | 0,47540334 | 0,70658666 |
| 29942_at | PURG | -0,06169841 | 3,86629202 | -0,73266112 | 0,47542282 | 0,70658666 |
| 4851_at | NOTCH1 | 0,08255527 | 7,36333211 | 0,73255714 | 0,47548435 | 0,70662359 |
| 100132078_a | LOC10013207 | -0,12491568 | 4,27907162 | -0,73249348 | 0,47552202 | 0,70662506 |
| 26256_at | CABYR | 0,23090338 | 6,7037984 | 0,73206123 | 0,47577788 | 0,70691655 |
| 105_at | ADARB2 | -0,06240801 | 4,34786265 | -0,73203809 | 0,47579158 | 0,70691655 |
| 339366_at | ADAMTSL5 | -0,08133703 | 5,72571929 | -0,73185831 | 0,47589802 | 0,70702017 |
| 392465_at | GLOD5 | -0,06917547 | 4,81068862 | -0,73164668 | 0,47602333 | 0,70714689 |
| 636_at | BICD1 | 0,0599145 | 4,47963655 | 0,73159028 | 0,47605673 | 0,70714689 |
| 342357_at | ZKSCAN2 | 0,060583 | 6,10928564 | 0,73148692 | 0,47611794 | 0,70718328 |
| 26740_at | OR1J2 | -0,07834423 | 4,62499326 | -0,73142331 | 0,47615562 | 0,70718471 |
| 55214_at | P3H2 | -0,07806952 | 7,32634285 | -0,73124402 | 0,47626182 | 0,70721172 |
| 1051_at | CEBPB | -0,1031209 | 10,8085412 | -0,73119842 | 0,47628883 | 0,70721172 |
| 56146_at | PCDHA2 | -0,0500274 | 3,11160096 | -0,73119064 | 0,47629344 | 0,70721172 |
| 28677_at | TRAV9-2 | 0,08576253 | 3,64528681 | 0,73114469 | 0,47632066 | 0,70721172 |
| 101929413_a | LOC10192941 | -0,05294465 | 3,51629672 | -0,73101144 | 0,4763996 | 0,70723638 |
| 80740_at | LY6G6C | -0,10905736 | 5,55619377 | -0,73097659 | 0,47642025 | 0,70723638 |
| 168090_at | C6orf118 | -0,04917626 | 3,56216576 | -0,7308871 | 0,47647328 | 0,70723638 |
| 57413_at | TMIGD3 | -0,08766308 | 3,69743082 | -0,73084996 | 0,47649529 | 0,70723638 |
| 11172_at | INSL6 | -0,06211827 | 3,91139816 | -0,73075953 | 0,47654887 | 0,70723638 |
| 85452_at | CFAP74 | -0,11913383 | 5,42969994 | -0,73074487 | 0,47655756 | 0,70723638 |
| 3683_at | ITGAL | 0,06883121 | 4,35254809 | 0,73063982 | 0,47661981 | 0,70727156 |
| 57565_at | KLHL14 | -0,07690556 | 3,48170114 | -0,73052954 | 0,47668517 | 0,70727156 |
| 150771_at | ITPRIPL1 | -0,10108038 | 4,45942198 | -0,73051901 | 0,47669141 | 0,70727156 |
| 26248_at | OR2K2 | -0,10016094 | 3,47581155 | -0,73003545 | 0,47697807 | 0,70762345 |
| 4317_at | MMP8 | -0,05588399 | 3,20235776 | -0,72999501 | 0,47700205 | 0,70762345 |
| 610_at | HCN2 | -0,06810124 | 4,81516039 | -0,72979571 | 0,47712023 | 0,70774426 |
| 494558_at | C9orf135-AS1 | -0,09463294 | 3,49055958 | -0,72964228 | 0,47721122 | 0,70779722 |
| 11013_at | TMSB15A | 0,45770929 | 4,97360312 | 0,72961159 | 0,47722942 | 0,70779722 |
| 7434_at | VIPR2 | -0,06021611 | 5,12768046 | -0,72929856 | 0,4774151 | 0,70797097 |
| 283867_at | LINC00922 | -0,04886216 | 4,2052745 | -0,7291216 | 0,47752009 | 0,70797097 |
| 256236_at | NAPSB | -0,073115 | 4,28107974 | -0,72911016 | 0,47752687 | 0,70797097 |
| 162979_at | ZNF296 | -0,09349914 | 6,5671861 | -0,72911001 | 0,47752697 | 0,70797097 |
| 100505832_a | PROX1-AS1 | -0,08065248 | 3,26919742 | -0,72910433 | 0,47753033 | 0,70797097 |
| 3839_at | KPNA3 | -0,77711215 | 7,16440649 | -0,72891372 | 0,47764344 | 0,70808416 |

| | | | | | | |
|---|---|---|---|---|---|---|
| 10732_at | TCFL5 | -0,11297005 | 8,49978946 | -0,72875876 | 0,4777354 | 0,70812582 |
| 8698_at | S1PR4 | -0,06389161 | 6,09196939 | -0,72874247 | 0,47774506 | 0,70812582 |
| 81928_at | CABLES2 | -0,05769028 | 8,00025994 | -0,72862426 | 0,47781522 | 0,70817532 |
| 2158_at | F9 | -0,0737968 | 3,17005367 | -0,72847811 | 0,47790198 | 0,70824941 |
| 3640_at | INSL3 | -0,08278319 | 4,96417896 | -0,72834203 | 0,47798276 | 0,70827637 |
| 100188953_a | LINC00092 | -0,0840738 | 5,68632857 | -0,72822989 | 0,47804933 | 0,70827637 |
| 84629_at | TNRC18 | 0,07011448 | 6,85813589 | 0,72812741 | 0,47811018 | 0,70827637 |
| 29890_at | RBM15B | 0,07770651 | 8,00896007 | 0,72812443 | 0,47811195 | 0,70827637 |
| 83875_at | BCO2 | -0,09626743 | 4,05216497 | -0,72806388 | 0,4781479 | 0,70827637 |
| 1501_at | CTNND2 | -0,05492756 | 4,04651554 | -0,72799871 | 0,4781866 | 0,70827637 |
| 49860_at | CRNN | -0,08800108 | 5,08386855 | -0,72799363 | 0,47818962 | 0,70827637 |
| 54979_at | HRASLS2 | 0,20380362 | 5,25173174 | 0,72792329 | 0,47823139 | 0,70827637 |
| 2519_at | FUCA2 | -0,18321689 | 8,5769318 | -0,72789012 | 0,47825109 | 0,70827637 |
| 9858_at | PPP1R26 | -0,05270389 | 7,0891003 | -0,72773303 | 0,47834439 | 0,70836008 |
| 79147_at | FKRP | 0,06371239 | 6,61776892 | 0,72718876 | 0,47866771 | 0,70867667 |
| 9657_at | IQCB1 | -0,17446472 | 7,49791907 | -0,72717161 | 0,4786779 | 0,70867667 |
| 100132057_a | LOC10013205 | -0,10914679 | 4,74562326 | -0,72712368 | 0,47870638 | 0,70867667 |
| 1306_at | COL15A1 | 0,05052661 | 3,65880677 | 0,72709169 | 0,47872539 | 0,70867667 |
| 80003_at | PCNXL2 | 0,09712288 | 6,01980775 | 0,72706354 | 0,47874212 | 0,70867667 |
| 144486_at | CEP83-AS1 | -0,07550922 | 5,03278738 | -0,72680612 | 0,4788951 | 0,70873875 |
| 401474_at | SAMD12 | 0,09396672 | 3,63446633 | 0,72679164 | 0,47890371 | 0,70873875 |
| 101929093_a | LINC01350 | -0,07297204 | 3,56138996 | -0,72674651 | 0,47893053 | 0,70873875 |
| 57728_at | WDR19 | 0,27094553 | 6,37295737 | 0,72674534 | 0,47893123 | 0,70873875 |
| 338739_at | CSTF3-AS1 | -0,05464857 | 2,95311961 | -0,72650997 | 0,47907114 | 0,70885042 |
| 79689_at | STEAP4 | 0,13608078 | 3,74914887 | 0,7264234 | 0,47912261 | 0,70885042 |
| 399_at | RHOH | -0,09765285 | 4,17779638 | -0,72639138 | 0,47914165 | 0,70885042 |
| 961_at | CD47 | 0,07229717 | 8,34469487 | 0,72633496 | 0,47917519 | 0,70885042 |
| 9778_at | KIAA0232 | 0,18389568 | 7,04551355 | 0,726276 | 0,47921025 | 0,70885042 |
| 6620_at | SNCB | -0,06459511 | 5,65069185 | -0,72624701 | 0,47922748 | 0,70885042 |
| 3792_at | KEL | -0,10696494 | 5,53927319 | -0,72618089 | 0,4792668 | 0,70885414 |
| 1444_at | CSHL1 | -0,09823371 | 5,93863149 | -0,72583987 | 0,47946961 | 0,70909966 |
| 55791_at | LRIF1 | 0,14041765 | 8,41660399 | 0,72547473 | 0,47968681 | 0,70922556 |
| 100506172_a | LINC01568 | -0,06836801 | 3,0091234 | -0,72541771 | 0,47972073 | 0,70922556 |
| 3357_at | HTR2B | -0,11101418 | 4,28555934 | -0,72541502 | 0,47972234 | 0,70922556 |
| 60559_at | SPCS3 | -0,25289545 | 8,93283739 | -0,72540479 | 0,47972842 | 0,70922556 |
| 6004_at | RGS16 | -0,2500089 | 7,65105899 | -0,7253873 | 0,47973883 | 0,70922556 |
| 57212_at | TP73-AS1 | -0,07657667 | 5,24156151 | -0,72503609 | 0,47994782 | 0,70942342 |
| 644873_at | LINC01184 | 0,10173041 | 5,45882002 | 0,72494672 | 0,48000101 | 0,70942342 |
| 117144_at | CATSPER1 | 0,08108612 | 6,22914729 | 0,72491234 | 0,48002147 | 0,70942342 |
| 284021_at | MILR1 | 0,15735504 | 4,23148985 | 0,72489415 | 0,4800323 | 0,70942342 |
| 3490_at | IGFBP7 | 0,22856146 | 6,75117953 | 0,72484593 | 0,480061 | 0,70942342 |
| 101929384_a | LOC10192938 | -0,094692 | 4,13799262 | -0,72479111 | 0,48009363 | 0,70942342 |
| 7482_at | WNT2B | 0,07227027 | 4,34563365 | 0,72471892 | 0,4801366 | 0,70943249 |
| 100652740_a | PYCARD-AS1 | 0,0905101 | 5,22665415 | 0,72465272 | 0,48017601 | 0,7094363 |
| 54546_at | RNF186 | -0,10011248 | 5,13023871 | -0,72452172 | 0,480254 | 0,70946051 |
| 11021_at | RAB35 | -0,08068379 | 8,34589379 | -0,72450148 | 0,48026605 | 0,70946051 |
| 245_at | ALOX12P2 | -0,0732595 | 6,158041 | -0,72430862 | 0,48038089 | 0,70953483 |
| 113763_at | ZBED6CL | 0,34413459 | 7,52907213 | 0,72429325 | 0,48039004 | 0,70953483 |
| 79890_at | RIN3 | -0,06865837 | 6,59656778 | -0,72414367 | 0,48047911 | 0,70961199 |
| 57552_at | NCEH1 | 0,17380774 | 8,49368463 | 0,72382247 | 0,48067042 | 0,70981929 |

| | | | | | | |
|---|---|---|---|---|---|---|
| 101927314_a | LOC10192731 | -0,04992308 | 2,93757053 | -0,72378427 | 0,48069318 | 0,70981929 |
| 168975_at | CNBD1 | -0,03850983 | 2,97698216 | -0,72359153 | 0,480808 | 0,70993336 |
| 4155_at | MBP | -0,06656187 | 6,70496252 | -0,72353089 | 0,48084413 | 0,70993336 |
| 2937_at | GSS | -0,06882767 | 9,58565995 | -0,72345809 | 0,48088751 | 0,70994298 |
| 10651_at | MTX2 | -0,14730639 | 9,02887816 | -0,72306624 | 0,48112102 | 0,71023329 |
| 9573_at | GDF3 | -0,07326814 | 4,5015345 | -0,72296995 | 0,48117841 | 0,71026358 |
| 222662_at | LHFPL5 | -0,0941656 | 4,72322964 | -0,72266795 | 0,48135844 | 0,71047488 |
| 5429_at | POLH | -0,12111375 | 7,00480826 | -0,72236222 | 0,48154074 | 0,71061603 |
| 2780_at | GNAT2 | 0,05944946 | 3,79095893 | 0,72233991 | 0,48155405 | 0,71061603 |
| 8447_at | DOC2B | -0,0694935 | 4,77807525 | -0,72232196 | 0,48156474 | 0,71061603 |
| 7432_at | VIP | 0,06034007 | 3,07625369 | 0,72222763 | 0,481621 | 0,71064461 |
| 51744_at | CD244 | -0,07225884 | 4,12851568 | -0,72215886 | 0,48166202 | 0,71065069 |
| 112752_at | IFT43 | 0,0696662 | 7,24393993 | 0,72194463 | 0,4817898 | 0,7106602 |
| 257407_at | C2orf72 | -0,06482939 | 5,58498499 | -0,72185528 | 0,4818431 | 0,7106602 |
| 79585_at | CORO7 | -0,12615909 | 6,21619428 | -0,72184269 | 0,48185061 | 0,7106602 |
| 25834_at | MGAT4C | -0,04051546 | 2,75272174 | -0,72179966 | 0,48187628 | 0,7106602 |
| 282969_at | FUOM | -0,09595446 | 4,88272501 | -0,7216972 | 0,48193741 | 0,7106602 |
| 1418_at | CRYGA | -0,08126342 | 3,80891455 | -0,72167613 | 0,48194998 | 0,7106602 |
| 2067_at | ERCC1 | 0,08597232 | 8,11517248 | 0,72165641 | 0,48196175 | 0,7106602 |
| 81029_at | WNT5B | 0,08741683 | 4,6563019 | 0,7216533 | 0,4819636 | 0,7106602 |
| 6013_at | RLN1 | 0,07073265 | 3,79265363 | 0,72155995 | 0,4820193 | 0,71066752 |
| 57453_at | DSCAML1 | -0,10700388 | 4,65424633 | -0,72152132 | 0,48204235 | 0,71066752 |
| 1778_at | DYNC1H1 | 0,05812961 | 8,07216254 | 0,72129562 | 0,48217703 | 0,71080559 |
| 54499_at | TMCO1 | 0,07916395 | 9,26870659 | 0,72123025 | 0,48221605 | 0,71080559 |
| 387758_at | FIBIN | -0,05384966 | 3,48759821 | -0,72114707 | 0,4822657 | 0,71080559 |
| 6519_at | SLC3A1 | -0,09657103 | 3,62819498 | -0,72111707 | 0,4822836 | 0,71080559 |
| 401494_at | HACD4 | 0,51115573 | 5,68975131 | 0,72099001 | 0,48235945 | 0,71085603 |
| 153769_at | SH3RF2 | -0,10729938 | 6,5235189 | -0,72087169 | 0,48243008 | 0,71085603 |
| 1272_at | CNTN1 | -0,13869143 | 4,32606741 | -0,72077388 | 0,48248847 | 0,71085603 |
| 8999_at | CDKL2 | 0,05401929 | 3,69709277 | 0,72075509 | 0,48249969 | 0,71085603 |
| 4062_at | LY6H | 0,09764726 | 4,82622501 | 0,72075065 | 0,48250234 | 0,71085603 |
| 51046_at | ST8SIA3 | -0,05249647 | 3,50170221 | -0,72050458 | 0,48264927 | 0,71090943 |
| 85012_at | TCEAL3 | 0,10962024 | 8,43946081 | 0,72041972 | 0,48269995 | 0,71090943 |
| 7355_at | SLC35A2 | -0,07446972 | 7,82910278 | -0,72041542 | 0,48270251 | 0,71090943 |
| 8907_at | AP1M1 | -0,10176028 | 8,47113923 | -0,72031145 | 0,4827646 | 0,71090943 |
| 6694_at | SPP2 | -0,1323307 | 4,5586481 | -0,72030668 | 0,48276745 | 0,71090943 |
| 79088_at | ZNF426 | 0,11519176 | 7,13699278 | 0,72028016 | 0,48278329 | 0,71090943 |
| 640_at | BLK | -0,06048768 | 4,5348386 | -0,72020704 | 0,48282697 | 0,71090943 |
| 56971_at | CEACAM19 | -0,0718137 | 6,42189561 | -0,72019555 | 0,48283383 | 0,71090943 |
| 6478_at | SIAH2 | 0,09572366 | 8,92487201 | 0,71970739 | 0,48312546 | 0,71122424 |
| 146722_at | CD300LF | -0,09220767 | 4,56067881 | -0,71966585 | 0,48315028 | 0,71122424 |
| 56261_at | GPCPD1 | -0,17554629 | 6,25231081 | -0,71965224 | 0,48315841 | 0,71122424 |
| 4825_at | NKX6-1 | -0,10865986 | 4,94233102 | -0,71935376 | 0,48333678 | 0,71143245 |
| 105371184_a | LOC10537118 | -0,08986896 | 6,02186624 | -0,71922283 | 0,48341504 | 0,71145991 |
| 91120_at | ZNF682 | 0,0736488 | 4,60432756 | 0,71919896 | 0,48342931 | 0,71145991 |
| 55252_at | ASXL2 | 0,06082052 | 7,26998302 | 0,71908511 | 0,48349736 | 0,71148373 |
| 101927176_a | LINC01069 | -0,07131219 | 3,9492038 | -0,7190483 | 0,48351937 | 0,71148373 |
| 100506215_a | PAXBP1-AS1 | 0,07355542 | 3,91643854 | 0,71881608 | 0,4836582 | 0,71163366 |
| 8148_at | TAF15 | -0,12557647 | 8,64177229 | -0,71854004 | 0,48382326 | 0,71182215 |
| 9145_at | SYNGR1 | 0,17455601 | 6,79326456 | 0,71840245 | 0,48390555 | 0,71188885 |

| | | | | | | |
|---|---|---|---|---|---|---|
| 101928386_a | LOC10192838 | -0,05534695 | 3,01277983 | -0,71826765 | 0,48398618 | 0,71195309 |
| 91875_at | TTC5 | -0,12357126 | 6,96243785 | -0,71815264 | 0,48405497 | 0,71199992 |
| 92092_at | ZC3HAV1L | -0,10199627 | 4,56417841 | -0,7179193 | 0,48419457 | 0,71215087 |
| 22803_at | XRN2 | 0,0514059 | 8,96284678 | 0,71770966 | 0,48432 | 0,71228098 |
| 4948_at | OCA2 | -0,06931207 | 4,23927919 | -0,71753245 | 0,48442605 | 0,71238255 |
| 259232_at | NALCN | -0,23763924 | 5,05618183 | -0,71735253 | 0,48453374 | 0,71248652 |
| 8644_at | AKR1C3 | 0,08026982 | 10,6829328 | 0,71725702 | 0,4845909 | 0,71251619 |
| 729359_at | PLIN4 | -0,12287131 | 5,63696648 | -0,71713753 | 0,48466244 | 0,71256698 |
| 101928417_a | LOC10192841 | 0,04776051 | 2,90954321 | 0,71705154 | 0,48471391 | 0,71258827 |
| 114905_at | C1QTNF7 | -0,05557109 | 3,30868864 | -0,71670827 | 0,48491945 | 0,71283603 |
| 101927248_a | LOC10192724 | -0,07925428 | 3,81016064 | -0,71648958 | 0,48505042 | 0,71297415 |
| 205717_at | KIAA2018 | 0,06008945 | 6,05181658 | 0,71641013 | 0,48509801 | 0,71298969 |
| 8938_at | BAIAP3 | 0,04853574 | 6,43420926 | 0,71591773 | 0,485393 | 0,71334788 |
| 7739_at | ZNF185 | 0,21974031 | 8,91662134 | 0,71587972 | 0,48541577 | 0,71334788 |
| 80162_at | ATHL1 | -0,05002885 | 6,01029315 | -0,71568004 | 0,48553543 | 0,7134693 |
| 23192_at | ATG4B | 0,0345668 | 7,91513685 | 0,7155967 | 0,48558537 | 0,71348826 |
| 6404_at | SELPLG | -0,06731525 | 5,4505497 | -0,71537497 | 0,48571828 | 0,7136291 |
| 79891_at | ZNF671 | -0,05635383 | 4,94103417 | -0,71521994 | 0,48581121 | 0,71371121 |
| 100996286_a | LOC10099628 | -0,08674005 | 4,90989393 | -0,71496805 | 0,48596223 | 0,71387863 |
| 161436_at | EML5 | 0,04297576 | 3,75049595 | 0,71490469 | 0,48600023 | 0,71388 |
| 101929467_a | LURAP1L-AS1 | -0,04863306 | 2,92449586 | -0,71480679 | 0,48605893 | 0,71388404 |
| 246744_at | STH | -0,10626758 | 5,38557919 | -0,7147765 | 0,4860771 | 0,71388404 |
| 22944_at | KIN | 0,1179859 | 4,98269298 | 0,71469686 | 0,48612486 | 0,71389976 |
| 100861544_a | LINC00557 | -0,06764958 | 3,7703744 | -0,71426103 | 0,48638628 | 0,71422922 |
| 128497_at | SPATA25 | -0,04768616 | 4,575492 | -0,71409992 | 0,48648294 | 0,71430333 |
| 3386_at | ICAM4 | -0,0670364 | 3,86063901 | -0,7140533 | 0,48651091 | 0,71430333 |
| 4718_at | NDUFC2 | 0,10361301 | 7,67953397 | 0,71395558 | 0,48656954 | 0,71431921 |
| 51195_at | RAPGEFL1 | -0,08633403 | 7,90858804 | -0,71383181 | 0,48664382 | 0,71431921 |
| 2906_at | GRIN2D | -0,15783422 | 5,16056165 | -0,71381941 | 0,48665126 | 0,71431921 |
| 100499194_a | LOC10049919 | -0,05642633 | 4,3517177 | -0,71378562 | 0,48667154 | 0,71431921 |
| 51460_at | SFMBT1 | 0,06636392 | 6,8151143 | 0,71372272 | 0,48670929 | 0,71431921 |
| 4435_at | CITED1 | 0,0692022 | 5,29516148 | 0,71366451 | 0,48674422 | 0,71431921 |
| 84203_at | TXNDC2 | -0,06243927 | 5,05737919 | -0,71337132 | 0,48692021 | 0,71452304 |
| 112802_at | KRT71 | -0,08757746 | 4,9826113 | -0,71327274 | 0,48697939 | 0,71455545 |
| 158401_at | C9orf84 | 0,05769835 | 3,39742307 | 0,71291968 | 0,48719138 | 0,71468838 |
| 25820_at | ARIH1 | 0,07756942 | 8,77762111 | 0,71280261 | 0,48726169 | 0,71468838 |
| 9645_at | MICAL2 | -0,28055219 | 7,81180739 | -0,71278561 | 0,4872719 | 0,71468838 |
| 23144_at | ZC3H3 | -0,07554929 | 5,61944305 | -0,71277728 | 0,4872769 | 0,71468838 |
| 51733_at | UPB1 | -0,05819771 | 4,54922313 | -0,71277278 | 0,48727961 | 0,71468838 |
| 101929312_a | LOC10192931 | -0,06119252 | 3,13299395 | -0,71275115 | 0,48729259 | 0,71468838 |
| 3350_at | HTR1A | -0,08666458 | 4,62582836 | -0,71268288 | 0,48733359 | 0,7146941 |
| 9473_at | THEMIS2 | -0,07331289 | 4,46175499 | -0,71234735 | 0,48753515 | 0,71488724 |
| 30811_at | HUNK | -0,05741886 | 4,44628578 | -0,71234008 | 0,48753951 | 0,71488724 |
| 79600_at | TCTN1 | 0,16402846 | 7,37333817 | 0,71218822 | 0,48763075 | 0,7149666 |
| 137392_at | FAM92A1 | 0,08644376 | 5,23819184 | 0,71208652 | 0,48769186 | 0,71497495 |
| 23008_at | KLHDC10 | 0,05513329 | 7,8992055 | 0,7120552 | 0,48771068 | 0,71497495 |
| 90249_at | UNC5A | -0,07912784 | 5,36446236 | -0,71188939 | 0,48781032 | 0,7150666 |
| 134111_at | UBE2QL1 | -0,05395937 | 3,30147394 | -0,71172564 | 0,48790874 | 0,71512754 |
| 221476_at | PI16 | -0,07384536 | 5,88339413 | -0,71169668 | 0,48792614 | 0,71512754 |
| 8913_at | CACNA1G | -0,04873671 | 4,71382928 | -0,71154842 | 0,48801526 | 0,71516411 |

| | | | | | | |
|---|---|---|---|---|---|---|
| 126375_at | ZNF792 | -0,0893341 | 5,92804344 | -0,71147973 | 0,48805655 | 0,71516411 |
| 26099_at | SZRD1 | -0,11856196 | 8,98064538 | -0,71140869 | 0,48809926 | 0,71516411 |
| 5998_at | RGS3 | -0,05469654 | 5,63063564 | -0,71140813 | 0,4880996 | 0,71516411 |
| 253970_at | SFTA3 | 0,0565124 | 3,28197862 | 0,71119295 | 0,48822897 | 0,71525126 |
| 23091_at | ZC3H13 | -0,04160424 | 7,32590984 | -0,71109781 | 0,48828617 | 0,71525126 |
| 4208_at | MEF2C | 0,19297614 | 6,44129398 | 0,71107984 | 0,48829698 | 0,71525126 |
| 150244_at | ZDHHC8P1 | 0,14738117 | 6,42342691 | 0,71105557 | 0,48831158 | 0,71525126 |
| 5021_at | OXTR | -0,08171139 | 5,13111899 | -0,71100043 | 0,48834473 | 0,71525126 |
| 9638_at | FEZ1 | 0,20344392 | 5,08274585 | 0,71086916 | 0,48842368 | 0,71526811 |
| 29984_at | RHOD | -0,11995125 | 9,62696535 | -0,71081462 | 0,48845648 | 0,71526811 |
| 100288198_a | HMGN3-AS1 | 0,06314625 | 4,22522469 | 0,71079608 | 0,48846763 | 0,71526811 |
| 131118_at | DNAJC19 | -0,06464028 | 7,90344321 | -0,71054889 | 0,48861632 | 0,7154172 |
| 348110_at | ARPIN | -0,05647118 | 5,23373229 | -0,71050333 | 0,48864373 | 0,7154172 |
| 50700_at | RDH8 | -0,08048926 | 4,36874116 | -0,71014279 | 0,48886065 | 0,71566156 |
| 4071_at | TM4SF1 | -0,84925119 | 7,85759358 | -0,71010244 | 0,48888493 | 0,71566156 |
| 27036_at | SIGLEC7 | -0,06065907 | 4,69538144 | -0,70980023 | 0,48906681 | 0,71586174 |
| 2745_at | GLRX | 0,2196732 | 7,04064969 | 0,70972849 | 0,48910999 | 0,71586174 |
| 54578_at | UGT1A6 | 0,1038707 | 9,3962655 | 0,70968999 | 0,48913317 | 0,71586174 |
| 54112_at | GPR88 | -0,05686956 | 4,27651313 | -0,70958317 | 0,48919746 | 0,71590145 |
| 286101_at | ZNF252P | 0,15919544 | 6,60627218 | 0,70942934 | 0,48929008 | 0,71595074 |
| 79100_at | LINC00626 | -0,09644399 | 3,5038609 | -0,70940376 | 0,48930548 | 0,71595074 |
| 130749_at | CPO | -0,05108234 | 3,5315427 | -0,70934013 | 0,48934378 | 0,7159524 |
| 121260_at | SLC15A4 | -0,0950965 | 8,51034898 | -0,7092046 | 0,48942539 | 0,71600251 |
| 4090_at | SMAD5 | -0,07699356 | 7,9906175 | -0,70915798 | 0,48945346 | 0,71600251 |
| 4316_at | MMP7 | 0,09778418 | 4,23920894 | 0,70909808 | 0,48948954 | 0,71600251 |
| 558_at | AXL | 0,15807791 | 6,6014304 | 0,70896376 | 0,48957043 | 0,71603775 |
| 168850_at | ZNF800 | -0,06498326 | 6,24243393 | -0,70893462 | 0,48958798 | 0,71603775 |
| 79184_at | BRCC3 | -0,10832233 | 7,95826975 | -0,70870982 | 0,48972338 | 0,71606177 |
| 163589_at | TDRD5 | 0,39010154 | 4,93229818 | 0,70867641 | 0,48974351 | 0,71606177 |
| 224_at | ALDH3A2 | -0,19805014 | 9,2899324 | -0,70865946 | 0,48975372 | 0,71606177 |
| 91807_at | MYLK3 | -0,04845712 | 3,79043403 | -0,70855829 | 0,48981466 | 0,71606177 |
| 5013_at | OTX1 | -0,05423055 | 5,96798357 | -0,70855334 | 0,48981765 | 0,71606177 |
| 114795_at | TMEM132B | -0,04708781 | 3,23917582 | -0,70853708 | 0,48982744 | 0,71606177 |
| 7101_at | NR2E1 | -0,05910671 | 3,5813753 | -0,70771791 | 0,4903211 | 0,71670257 |
| 319103_at | SNORD8 | -0,11043745 | 4,37813548 | -0,70767014 | 0,4903499 | 0,71670257 |
| 102724508_a | THCAT158 | -0,05123391 | 4,67845411 | -0,70762451 | 0,4903774 | 0,71670257 |
| 93973_at | ACTR8 | 0,05196255 | 7,10049885 | 0,70744877 | 0,49048335 | 0,7167994 |
| 2776_at | GNAQ | 0,15141921 | 8,72115221 | 0,70739119 | 0,49051807 | 0,7167994 |
| 53904_at | MYO3A | -0,04143805 | 3,55319502 | -0,70701579 | 0,49074445 | 0,71707581 |
| 196500_at | PIANP | -0,07464846 | 5,1877504 | -0,70690275 | 0,49081263 | 0,71712104 |
| 5243_at | ABCB1 | 0,62288503 | 5,55547912 | 0,70677777 | 0,49088802 | 0,71712624 |
| 7485_at | WRB | -0,10845444 | 8,5091701 | -0,70677342 | 0,49089065 | 0,71712624 |
| 395_at | ARHGAP6 | -0,06494191 | 3,71743661 | -0,70669903 | 0,49093552 | 0,7171374 |
| 100506385_a | LINC01426 | -0,05700268 | 3,04291169 | -0,70645175 | 0,49108471 | 0,71729546 |
| 100137047_a | JMJD7 | 0,17565348 | 7,5904901 | 0,70639625 | 0,49111819 | 0,71729546 |
| 80346_at | REEP4 | 0,09233332 | 9,24822119 | 0,70607525 | 0,49131191 | 0,71752398 |
| 284825_at | LOC284825 | -0,08301694 | 3,95780219 | -0,70583304 | 0,49145811 | 0,71754907 |
| 64595_at | TTTY15 | 0,23867092 | 5,22213607 | 0,70583043 | 0,49145968 | 0,71754907 |
| 79411_at | GLB1L | 0,15064315 | 5,7410831 | 0,70580765 | 0,49147343 | 0,71754907 |
| 257169_at | C9orf43 | -0,09474325 | 4,17082314 | -0,70575954 | 0,49150247 | 0,71754907 |

| | | | | | | |
|---|---|---|---|---|---|---|
| 101927237_a | LOC10192723 | 0,04880466 | 2,94787781 | 0,70573822 | 0,49151534 | 0,71754907 |
| 9496_at | TBX4 | -0,05882411 | 4,96936213 | -0,70562613 | 0,49158302 | 0,71759348 |
| 1953_at | MEGF6 | 0,20513584 | 7,09385877 | 0,70548223 | 0,4916699 | 0,71766593 |
| 9143_at | SYNGR3 | 0,13616781 | 5,216204 | 0,7053446 | 0,491753 | 0,71771442 |
| 140469_at | MYO3B | 0,05138634 | 4,26097845 | 0,70525727 | 0,49180573 | 0,71771442 |
| 101927433_a | LINC01255 | -0,05018699 | 3,32557995 | -0,7052421 | 0,4918149 | 0,71771442 |
| 4352_at | MPL | -0,06316827 | 4,5055791 | -0,705126 | 0,49188501 | 0,71776237 |
| 55568_at | GALNT10 | -0,16292223 | 7,73204402 | -0,70500353 | 0,49195898 | 0,71781593 |
| 30813_at | VSX1 | -0,0647415 | 4,28011452 | -0,70480534 | 0,4920787 | 0,71791617 |
| 2172_at | FABP6 | 0,22443048 | 6,42938982 | 0,7047664 | 0,49210222 | 0,71791617 |
| 100862680_a | LINC00507 | -0,05532342 | 3,0058853 | -0,70466588 | 0,49216295 | 0,71795039 |
| 84464_at | SLX4 | -0,04991002 | 6,57564727 | -0,7045828 | 0,49221315 | 0,71795864 |
| 55971_at | BAIAP2L1 | -0,04438666 | 8,33432521 | -0,70453314 | 0,49224315 | 0,71795864 |
| 388335_at | TMEM220 | -0,06601103 | 3,47508567 | -0,70424221 | 0,49241895 | 0,71813904 |
| 6843_at | VAMP1 | 0,05639817 | 6,17773037 | 0,70420507 | 0,4924414 | 0,71813904 |
| 125893_at | ZNF816 | 0,07960099 | 6,59840358 | 0,70413466 | 0,49248395 | 0,71814673 |
| 1142_at | CHRNB3 | 0,064538 | 3,5229872 | 0,70399127 | 0,49257062 | 0,71818852 |
| 8269_at | TMEM187 | 0,25268421 | 6,95038841 | 0,70396388 | 0,49258718 | 0,71818852 |
| 84842_at | HPDL | -0,35485651 | 5,52648791 | -0,70386212 | 0,49264869 | 0,71822385 |
| 26130_at | GAPVD1 | 0,05391995 | 8,11283475 | 0,7037896 | 0,49269253 | 0,7182334 |
| 4158_at | MC2R | -0,07358699 | 3,8837927 | -0,70339025 | 0,49293399 | 0,71853102 |
| 100507389_a | LOC10050738 | -0,07445968 | 3,60973311 | -0,70313991 | 0,49308539 | 0,71867216 |
| 374907_at | B3GNT8 | 0,09993238 | 5,53040248 | 0,70310677 | 0,49310543 | 0,71867216 |
| 131408_at | FAM131A | 0,26442781 | 5,80993922 | 0,70302711 | 0,49315361 | 0,718688 |
| 100885798_a | LINC00210 | -0,08701146 | 3,59234369 | -0,70276604 | 0,49331155 | 0,71886378 |
| 1185_at | CLCN6 | 0,06930163 | 7,35301063 | 0,70254832 | 0,49344327 | 0,71896829 |
| 5058_at | PAK1 | 0,22503127 | 6,88825053 | 0,70252413 | 0,49345791 | 0,71896829 |
| 10876_at | EDDM3A | 0,039646 | 3,1260736 | 0,70233939 | 0,4935697 | 0,71907678 |
| 94235_at | GNG8 | -0,06006209 | 4,53570176 | -0,70216276 | 0,4936766 | 0,71917813 |
| 7733_at | ZNF180 | 0,04321919 | 6,21814614 | 0,70199011 | 0,4937811 | 0,71927597 |
| 1643_at | DDB2 | -0,12678079 | 8,38059343 | -0,70181217 | 0,49388882 | 0,71937848 |
| 56106_at | PCDHGA10 | -0,07316933 | 5,77094496 | -0,70167116 | 0,4939742 | 0,71944843 |
| 4355_at | MPP2 | -0,05650179 | 5,99335426 | -0,70155175 | 0,4940465 | 0,7194949 |
| 7918_at | GPANK1 | 0,08402251 | 7,32612683 | 0,7014951 | 0,4940808 | 0,7194949 |
| 3002_at | GZMB | -0,0576521 | 4,40567007 | -0,70133479 | 0,49417789 | 0,71958188 |
| 79846_at | CFAP69 | 0,13748426 | 4,08368755 | 0,70088412 | 0,49445086 | 0,71979227 |
| 8742_at | TNFSF12 | -0,09433882 | 4,81751412 | -0,7008804 | 0,49445312 | 0,71979227 |
| 7534_at | YWHAZ | -0,07318005 | 11,4374949 | -0,70086179 | 0,49446439 | 0,71979227 |
| 101060019_a | LOC10106001 | -0,06686615 | 3,03721172 | -0,70079513 | 0,49450478 | 0,71979227 |
| 23313_at | KIAA0930 | 0,07026088 | 8,32989511 | 0,70078782 | 0,49450921 | 0,71979227 |
| 347365_at | ITIH6 | -0,0717116 | 4,19132086 | -0,70071579 | 0,49455285 | 0,7198014 |
| 219854_at | TMEM218 | 0,11141295 | 5,43278767 | 0,70035603 | 0,49477085 | 0,72006429 |
| 100170229_a | SRRM5 | -0,08544575 | 4,25462105 | -0,70024899 | 0,49483573 | 0,7201043 |
| 375484_at | SIMC1 | 0,14850152 | 5,71535671 | 0,70008904 | 0,49493268 | 0,72019097 |
| 8630_at | HSD17B6 | -0,24655676 | 4,84569312 | -0,69992668 | 0,4950311 | 0,72023799 |
| 100505890_a | SORCS3-AS1 | -0,06222619 | 4,12938795 | -0,69991239 | 0,49503977 | 0,72023799 |
| 10540_at | DCTN2 | 0,05051716 | 10,0559746 | 0,69977321 | 0,49512414 | 0,72028075 |
| 137994_at | LETM2 | -0,09628212 | 4,34332315 | -0,6996821 | 0,49517939 | 0,72028075 |
| 101929116_a | LOC10192911 | -0,07121297 | 4,00424132 | -0,6996456 | 0,49520152 | 0,72028075 |
| 58530_at | LY6G6D | -0,08794768 | 4,76746982 | -0,69958281 | 0,4952396 | 0,72028075 |

| | | | | | | |
|---|---|---|---|---|---|---|
| 101928371_a | LOC10192837 | -0,09756909 | 4,24882196 | -0,69955556 | 0,49525612 | 0,72028075 |
| 55049_at | C19orf60 | -0,05665568 | 8,49111208 | -0,69903124 | 0,49557413 | 0,72068885 |
| 6005_at | RHAG | -0,06019091 | 4,00473206 | -0,69885296 | 0,49568228 | 0,72079171 |
| 6355_at | CCL8 | -0,06805234 | 3,17003399 | -0,69858276 | 0,49584623 | 0,72097569 |
| 8672_at | EIF4G3 | -0,06940238 | 7,13030518 | -0,69850474 | 0,49589358 | 0,72099011 |
| 23741_at | EID1 | 0,10411317 | 9,29879679 | 0,69838783 | 0,49596453 | 0,72102206 |
| 221416_at | C6orf223 | -0,08098675 | 4,66484904 | -0,69834518 | 0,49599041 | 0,72102206 |
| 84765_at | ZNF577 | 0,05094007 | 4,60357295 | 0,69813469 | 0,49611818 | 0,72115337 |
| 84217_at | ZMYND12 | 0,08151563 | 3,84613984 | 0,69792764 | 0,49624387 | 0,72124282 |
| 1008_at | CDH10 | 0,09406223 | 4,42779065 | 0,69787139 | 0,49627802 | 0,72124282 |
| 5516_at | PPP2CB | 0,17910055 | 6,81215813 | 0,69776646 | 0,49634173 | 0,72124282 |
| 161502_at | C15orf26 | -0,04702821 | 3,12013557 | -0,69775742 | 0,49634722 | 0,72124282 |
| 9823_at | ARMCX2 | 0,38632517 | 4,76777521 | 0,69767581 | 0,49639677 | 0,72124282 |
| 101929718_a | LOC10192971 | -0,14251585 | 4,89921667 | -0,69766331 | 0,49640436 | 0,72124282 |
| 9745_at | ZNF536 | -0,03593357 | 3,52997568 | -0,69749029 | 0,49650944 | 0,72134107 |
| 54879_at | ST7L | 0,06011891 | 5,51187555 | 0,69736966 | 0,4965827 | 0,72136145 |
| 6734_at | SRPR | 0,05366773 | 9,55289446 | 0,69734388 | 0,49659836 | 0,72136145 |
| 339327_at | ZNF546 | 0,03645548 | 3,12472 | 0,69726021 | 0,49664918 | 0,72138087 |
| 5623_at | PSPN | -0,09840056 | 5,10331927 | -0,69701425 | 0,49679859 | 0,72149217 |
| 25807_at | RHBDD3 | -0,05563788 | 7,77717856 | -0,69701075 | 0,49680071 | 0,72149217 |
| 12_at | SERPINA3 | -0,09426993 | 5,55358138 | -0,69691723 | 0,49685753 | 0,72152028 |
| 9839_at | ZEB2 | -0,04339474 | 3,31432062 | -0,69674375 | 0,49696293 | 0,72161895 |
| 56952_at | PRTFDC1 | 0,06447694 | 3,15516097 | 0,69661533 | 0,49704097 | 0,72167786 |
| 51090_at | PLLP | 0,43353621 | 6,90608715 | 0,69654118 | 0,49708603 | 0,72168889 |
| 283358_at | B4GALNT3 | -0,06566677 | 5,76137223 | -0,696309 | 0,49722715 | 0,72181518 |
| 286103_at | ZNF252P-AS1 | 0,12391291 | 4,23793399 | 0,69624747 | 0,49726455 | 0,72181518 |
| 2322_at | FLT3 | -0,08866863 | 4,07745032 | -0,69621311 | 0,49728543 | 0,72181518 |
| 154075_at | SAMD3 | 0,04892578 | 3,07778953 | 0,69612324 | 0,49734006 | 0,72184009 |
| 8174_at | MADCAM1 | -0,10694773 | 5,61312865 | -0,69598535 | 0,49742389 | 0,72190736 |
| 84080_at | ENKD1 | -0,1155647 | 6,47940252 | -0,69566289 | 0,49761996 | 0,72200148 |
| 56980_at | PRDM10 | -0,03959996 | 7,6130736 | -0,69559778 | 0,49765955 | 0,72200148 |
| 85280_at | KRTAP9-4 | -0,12351945 | 4,42802595 | -0,6955605 | 0,49768223 | 0,72200148 |
| 642280_at | ZNF876P | 0,05311321 | 3,06521522 | 0,69548408 | 0,4977287 | 0,72200148 |
| 51272_at | BET1L | -0,18974294 | 6,7093898 | -0,69547908 | 0,49773174 | 0,72200148 |
| 91179_at | SCARF2 | -0,06378523 | 5,32853224 | -0,69545084 | 0,49774892 | 0,72200148 |
| 27233_at | SULT1C4 | -0,07356352 | 3,29006301 | -0,69544722 | 0,49775111 | 0,72200148 |
| 23211_at | ZC3H4 | -0,05285782 | 9,12715244 | -0,69529727 | 0,49784232 | 0,72203342 |
| 7296_at | TXNRD1 | -0,08563162 | 10,5355388 | -0,69528777 | 0,4978481 | 0,72203342 |
| 10172_at | ZNF256 | 0,10206486 | 5,57680044 | 0,69511869 | 0,49795095 | 0,72212688 |
| 55353_at | LAPTM4B | -0,05788745 | 11,364913 | -0,69505859 | 0,49798751 | 0,72212688 |
| 26240_at | FAM50B | -0,10522763 | 6,97427065 | -0,69483696 | 0,49812236 | 0,72226805 |
| 105370059_a | LOC10537005 | -0,06315299 | 3,59514798 | -0,69472663 | 0,49818949 | 0,72227819 |
| 23657_at | SLC7A11 | -0,50587378 | 9,5160964 | -0,69470223 | 0,49820434 | 0,72227819 |
| 64097_at | EPB41L4A | 0,3782763 | 6,6116745 | 0,69463187 | 0,49824716 | 0,72228589 |
| 54973_at | CPSF3L | -0,04781769 | 8,69984621 | -0,69456286 | 0,49828915 | 0,72229242 |
| 2880_at | GPX5 | -0,06073239 | 3,84132298 | -0,69450121 | 0,49832667 | 0,72229246 |
| 79645_at | EFCAB1 | -0,05390686 | 3,55448286 | -0,69430484 | 0,4984462 | 0,72241134 |
| 55612_at | FERMT1 | -0,12210145 | 9,4961144 | -0,69386527 | 0,4987138 | 0,72274481 |
| 58986_at | TMEM8A | 0,10527487 | 8,63570925 | 0,69359969 | 0,49887554 | 0,72292481 |
| 221692_at | PHACTR1 | 0,04678269 | 3,98614028 | 0,69335594 | 0,49902399 | 0,72308554 |

| | | | | | | |
|---|---|---|---|---|---|---|
| 222545_at | GPRC6A | -0,08940454 | 4,31958884 | -0,69326356 | 0,49908026 | 0,72311268 |
| 10004_at | NAALADL1 | -0,04183041 | 5,09252819 | -0,69307858 | 0,49919295 | 0,72319747 |
| 9865_at | TRIL | -0,06166791 | 4,51492678 | -0,69303268 | 0,49922091 | 0,72319747 |
| 285533_at | RNF175 | -0,07279576 | 4,33589152 | -0,69298263 | 0,49925141 | 0,72319747 |
| 284798_at | LOC284798 | -0,0571334 | 4,47878073 | -0,69285525 | 0,49932903 | 0,72320286 |
| 200205_at | IBA57 | 0,04598086 | 6,00685918 | 0,69281662 | 0,49935257 | 0,72320286 |
| 29124_at | LGALS13 | -0,10622253 | 3,50222721 | -0,69279167 | 0,49936777 | 0,72320286 |
| 221749_at | PXDC1 | 0,18488242 | 7,86389731 | 0,6926871 | 0,49943149 | 0,72320933 |
| 27236_at | ARFIP1 | 0,17179736 | 7,80989051 | 0,69264543 | 0,49945688 | 0,72320933 |
| 6137_at | RPL13 | -0,08702731 | 6,9241577 | -0,69259952 | 0,49948486 | 0,72320933 |
| 90134_at | KCNH7 | -0,07352642 | 3,61849931 | -0,69244577 | 0,49957857 | 0,72329064 |
| 163404_at | LPPR5 | 0,07865421 | 3,38322565 | 0,69226697 | 0,49968756 | 0,72339407 |
| 729296_at | LOC729296 | -0,05561432 | 3,25876303 | -0,69210951 | 0,49978355 | 0,72347865 |
| 2997_at | GYS1 | -0,06609009 | 8,34502402 | -0,69199928 | 0,49985075 | 0,72352157 |
| 116135_at | LRRC3B | 0,04168739 | 3,83239744 | 0,6915587 | 0,50011942 | 0,72382298 |
| 2587_at | GALR1 | 0,05870379 | 4,00426239 | 0,69153457 | 0,50013414 | 0,72382298 |
| 100130452_a | LOC10013045 | -0,08840403 | 3,38684034 | -0,69135294 | 0,50024493 | 0,72389878 |
| 6830_at | SUPT6H | 0,06611067 | 7,51012282 | 0,69132548 | 0,50026167 | 0,72389878 |
| 861_at | RUNX1 | -0,06100848 | 6,26817865 | -0,69117454 | 0,50035375 | 0,72395874 |
| 84000_at | TMPRSS13 | 0,07957535 | 5,66931731 | 0,69113435 | 0,50037828 | 0,72395874 |
| 23218_at | NBEAL2 | 0,11551416 | 6,64525712 | 0,69105986 | 0,50042372 | 0,72397012 |
| 23460_at | ABCA6 | 0,04221518 | 3,153663 | 0,69096961 | 0,50047878 | 0,7239954 |
| 10929_at | SRSF8 | -0,05192821 | 8,46734365 | -0,69084843 | 0,50055273 | 0,72402898 |
| 1232_at | CCR3 | 0,07170826 | 3,80198402 | 0,69068997 | 0,50064942 | 0,72402898 |
| 138162_at | C9orf116 | 0,08794216 | 6,70547016 | 0,69068696 | 0,50065126 | 0,72402898 |
| 3992_at | FADS1 | -0,55552326 | 9,2438657 | -0,69068519 | 0,50065234 | 0,72402898 |
| 1608_at | DGKG | -0,04890219 | 4,32323243 | -0,69053833 | 0,50074197 | 0,72410424 |
| 3860_at | KRT13 | -0,18974485 | 12,8981031 | -0,69031105 | 0,5008807 | 0,72424528 |
| 100144603_a | CHKB-AS1 | 0,15185258 | 6,92900851 | 0,69023645 | 0,50092624 | 0,72424528 |
| 101929529_a | LOC10192952 | -0,07277432 | 3,57917098 | -0,69019377 | 0,5009523 | 0,72424528 |
| 128387_at | TATDN3 | 0,08592343 | 6,35594906 | 0,68944189 | 0,50141144 | 0,72478376 |
| 4363_at | ABCC1 | -0,05812751 | 9,19874462 | -0,68940507 | 0,50143393 | 0,72478376 |
| 8050_at | PDHX | -0,1432367 | 8,84653521 | -0,68931856 | 0,50148678 | 0,72478376 |
| 100505920_a | LOC10050592 | -0,06292038 | 4,43027607 | -0,68919297 | 0,5015635 | 0,72478376 |
| 10083_at | USH1C | -0,07733012 | 4,94033429 | -0,68917332 | 0,5015755 | 0,72478376 |
| 11284_at | PNKP | 0,16345673 | 8,43065149 | 0,68917254 | 0,50157598 | 0,72478376 |
| 5680_at | PSG11 | -0,14776141 | 3,81305283 | -0,68915264 | 0,50158814 | 0,72478376 |
| 59_at | ACTA2 | 0,23610075 | 7,20413762 | 0,68865688 | 0,50189108 | 0,7251121 |
| 8340_at | HIST1H2BL | -0,11916032 | 5,57974476 | -0,68858148 | 0,50193716 | 0,7251121 |
| 8844_at | KSR1 | -0,06125681 | 5,96482005 | -0,68856593 | 0,50194667 | 0,7251121 |
| 78989_at | COLEC11 | 0,08175366 | 4,81677308 | 0,68853094 | 0,50196805 | 0,7251121 |
| 153745_at | FAM71B | -0,06554429 | 3,90094306 | -0,68845757 | 0,5020129 | 0,7251121 |
| 79664_at | ICE2 | -0,06772124 | 6,82489363 | -0,68838691 | 0,50205609 | 0,7251121 |
| 127003_at | C1orf194 | -0,08945489 | 5,40947122 | -0,68834965 | 0,50207887 | 0,7251121 |
| 100130964_a | LOC10013096 | -0,06384103 | 2,99874677 | -0,68822749 | 0,50215354 | 0,72513898 |
| 57731_at | SPTBN4 | -0,04566826 | 5,13824852 | -0,68819606 | 0,50217276 | 0,72513898 |
| 149699_at | GTSF1L | -0,05936989 | 3,71936656 | -0,68806748 | 0,50225138 | 0,72519813 |
| 6756_at | SSX1 | -0,05745116 | 3,12704758 | -0,68778884 | 0,50242176 | 0,72538977 |
| 399972_at | ST3GAL4-AS1 | 0,16042589 | 6,76557082 | 0,68752789 | 0,50258135 | 0,72556581 |
| 6891_at | TAP2 | 0,14668438 | 7,65288877 | 0,68736231 | 0,50268264 | 0,72565765 |

| | | | | | | |
|---|---|---|---|---|---|---|
| 29777_at | ABT1 | -0,11570577 | 8,48354659 | -0,68727736 | 0,50273461 | 0,72567829 |
| 3973_at | LHCGR | -0,05094109 | 3,11691821 | -0,68701626 | 0,50289435 | 0,72574809 |
| 113146_at | AHNAK2 | -0,08114136 | 5,85669198 | -0,68694977 | 0,50293504 | 0,72574809 |
| 80025_at | PANK2 | 0,0920272 | 9,12177528 | 0,68694585 | 0,50293744 | 0,72574809 |
| 84274_at | COQ5 | 0,07820713 | 8,4203887 | 0,68689061 | 0,50297124 | 0,72574809 |
| 102724201_a | LOC10272420 | -0,05275743 | 2,90491854 | -0,68689044 | 0,50297135 | 0,72574809 |
| 55665_at | URGCP | -0,0632258 | 7,7958536 | -0,68673083 | 0,50306902 | 0,72583466 |
| 785_at | CACNB4 | -0,05409678 | 3,82614796 | -0,68660635 | 0,5031452 | 0,72586591 |
| 368_at | ABCC6 | -0,0571423 | 4,42543976 | -0,6865723 | 0,50316604 | 0,72586591 |
| 115727_at | RASGRP4 | -0,07273829 | 4,76384855 | -0,68648582 | 0,50321898 | 0,72588792 |
| 55048_at | VPS37C | 0,10414641 | 8,25913983 | 0,68638905 | 0,50327821 | 0,725919 |
| 79701_at | OGFOD3 | -0,04148663 | 7,86146569 | -0,68613027 | 0,50343665 | 0,72603907 |
| 9866_at | TRIM66 | -0,10537591 | 5,02980478 | -0,68607667 | 0,50346946 | 0,72603907 |
| 54802_at | TRIT1 | 0,05640662 | 8,48220627 | 0,68606839 | 0,50347453 | 0,72603907 |
| 100996654_a | LOC10099665 | -0,08277285 | 4,42961005 | -0,68590361 | 0,50357543 | 0,72613021 |
| 2304_at | FOXE1 | 0,31772505 | 5,26721248 | 0,68581897 | 0,50362726 | 0,7261506 |
| 83737_at | ITCH | 0,04710733 | 7,79661835 | 0,68574783 | 0,50367083 | 0,72615906 |
| 57104_at | PNPLA2 | 0,08031107 | 7,87894935 | 0,68551861 | 0,50381123 | 0,72627881 |
| 4863_at | NPAT | -0,12902608 | 7,23798941 | -0,68543856 | 0,50386026 | 0,72627881 |
| 2332_at | FMR1 | 0,17177948 | 9,64561309 | 0,68530465 | 0,50394229 | 0,72627881 |
| 4835_at | NQO2 | -0,24215183 | 8,99865318 | -0,68527755 | 0,50395889 | 0,72627881 |
| 83482_at | SCRT1 | -0,08191406 | 4,86792303 | -0,68527309 | 0,50396163 | 0,72627881 |
| 4281_at | MID1 | 0,05514721 | 9,61180327 | 0,68523363 | 0,5039858 | 0,72627881 |
| 53947_at | A4GALT | 0,08004033 | 5,8494892 | 0,68518137 | 0,50401781 | 0,72627881 |
| 27252_at | KLHL20 | 0,09025061 | 7,550783 | 0,68503697 | 0,50410629 | 0,72635197 |
| 389073_at | C2orf80 | 0,06561532 | 3,20508116 | 0,68451581 | 0,50442569 | 0,72675782 |
| 51226_at | COPZ2 | -0,09257007 | 6,21073704 | -0,68440609 | 0,50449294 | 0,72680035 |
| 81545_at | FBXO38 | 0,08414583 | 7,48438933 | 0,68425442 | 0,50458592 | 0,72687995 |
| 100506406_a | LOC10050640 | 0,07076787 | 3,71630792 | 0,68407702 | 0,50469469 | 0,72690356 |
| 101929122_a | LOC10192912 | -0,06053435 | 4,25869904 | -0,68404847 | 0,5047122 | 0,72690356 |
| 847_at | CAT | 0,24318653 | 9,82802995 | 0,68399636 | 0,50474415 | 0,72690356 |
| 221481_at | ARMC12 | -0,05845814 | 5,11879597 | -0,68391755 | 0,50479248 | 0,72690356 |
| 84525_at | HOPX | 0,32944616 | 5,1657224 | 0,68384531 | 0,50483678 | 0,72690356 |
| 23593_at | HEBP2 | -0,08126275 | 10,6490795 | -0,68377562 | 0,50487952 | 0,72690356 |
| 132014_at | IL17RE | -0,07099853 | 6,13585982 | -0,68372226 | 0,50491224 | 0,72690356 |
| 2798_at | GNRHR | -0,05474984 | 3,17252498 | -0,68370134 | 0,50492507 | 0,72690356 |
| 5626_at | PROP1 | -0,08619399 | 5,13227316 | -0,68367384 | 0,50494194 | 0,72690356 |
| 2832_at | NPBWR2 | -0,0775897 | 4,1275533 | -0,68331937 | 0,50515938 | 0,72711917 |
| 30827_at | CXXC1 | -0,03843184 | 8,18927396 | -0,68330661 | 0,5051672 | 0,72711917 |
| 23193_at | GANAB | -0,05951832 | 10,2826756 | -0,68318073 | 0,50524443 | 0,72717599 |
| 9375_at | TM9SF2 | 0,16795943 | 11,4476593 | 0,68305104 | 0,50532401 | 0,72721694 |
| 9026_at | HIP1R | 0,06778167 | 6,64739056 | 0,68299698 | 0,50535718 | 0,72721694 |
| 55779_at | CFAP44 | 0,21964856 | 5,63684146 | 0,6829257 | 0,50540092 | 0,72721694 |
| 10771_at | ZMYND11 | -0,09489059 | 7,17245381 | -0,68283026 | 0,50545949 | 0,72721694 |
| 57575_at | PCDH10 | 0,04179 | 3,93744931 | 0,68282675 | 0,50546164 | 0,72721694 |
| 11341_at | SCRG1 | -0,08809433 | 3,51869673 | -0,68262348 | 0,5055864 | 0,72731534 |
| 163688_at | CALML6 | -0,05761628 | 5,13533631 | -0,6825198 | 0,50565004 | 0,72731534 |
| 93474_at | ZNF670 | -0,25543291 | 5,82151344 | -0,68247738 | 0,50567608 | 0,72731534 |
| 8745_at | ADAM23 | -0,15546131 | 5,68593085 | -0,68246925 | 0,50568107 | 0,72731534 |
| 114026_at | ZIM3 | 0,03997446 | 2,85406571 | 0,68220496 | 0,50584332 | 0,72749439 |

| | | | | | | |
|---|---|---|---|---|---|---|
| 5275_at | SERPINB13 | 0,09502988 | 6,06557842 | 0,68211344 | 0,50589951 | 0,72752089 |
| 2247_at | FGF2 | -0,0635614 | 3,99087975 | -0,68203541 | 0,50594742 | 0,72753548 |
| 641364_at | SLC7A11-AS1 | -0,07124786 | 3,71366328 | -0,68191396 | 0,50602201 | 0,72757577 |
| 92126_at | DSEL | 0,06359684 | 4,40613961 | 0,68183444 | 0,50607084 | 0,72757577 |
| 204474_at | PDILT | 0,05898677 | 3,54427471 | 0,68180526 | 0,50608876 | 0,72757577 |
| 56965_at | PARP6 | 0,2932061 | 7,13504225 | 0,68163765 | 0,5061917 | 0,72766946 |
| 115749_at | C12orf56 | 0,44403722 | 4,85969704 | 0,68142095 | 0,50632481 | 0,72776429 |
| 341405_at | ANKRD33 | -0,06632154 | 4,39913459 | -0,68126882 | 0,50641827 | 0,72776429 |
| 6305_at | SBF1 | 0,05913768 | 7,83749708 | 0,68126101 | 0,50642307 | 0,72776429 |
| 5081_at | PAX7 | -0,0657929 | 4,3276031 | -0,68123653 | 0,50643811 | 0,72776429 |
| 1767_at | DNAH5 | -0,06219409 | 4,27586362 | -0,68122276 | 0,50644657 | 0,72776429 |
| 161882_at | ZFPM1 | -0,04772945 | 7,06699925 | -0,68106409 | 0,50654406 | 0,72785008 |
| 64789_at | EXO5 | -0,12795106 | 6,46610369 | -0,68080146 | 0,50670546 | 0,72802769 |
| 9218_at | VAPA | -0,18901399 | 7,2054108 | -0,68068717 | 0,5067757 | 0,72807431 |
| 23149_at | FCHO1 | -0,07670558 | 5,24809107 | -0,68050663 | 0,50688667 | 0,72813338 |
| 341947_at | COX8C | 0,0599655 | 3,5878283 | 0,68049728 | 0,50689242 | 0,72813338 |
| 7404_at | UTY | 0,18332845 | 4,28647895 | 0,68034245 | 0,5069876 | 0,7282158 |
| 10488_at | CREB3 | -0,04710746 | 9,40112664 | -0,680011 | 0,50719139 | 0,7284542 |
| 124535_at | HSF5 | 0,05136336 | 3,61272614 | 0,6798992 | 0,50726014 | 0,72849862 |
| 5535_at | PPP3R2 | -0,0446751 | 4,09811462 | -0,67969889 | 0,50738333 | 0,7284998 |
| 101928303_a | LOC10192830 | -0,06115757 | 3,30975298 | -0,67963572 | 0,50742219 | 0,7284998 |
| 27230_at | SERP1 | -0,07492358 | 10,5778758 | -0,67955326 | 0,50747291 | 0,7284998 |
| 79992_at | AGPAT4-IT1 | -0,05412339 | 5,4633578 | -0,67950824 | 0,5075006 | 0,7284998 |
| 6581_at | SLC22A3 | 0,04809458 | 3,15554248 | 0,67949192 | 0,50751064 | 0,7284998 |
| 23609_at | MKRN2 | -0,03788228 | 8,10209026 | -0,67946752 | 0,50752565 | 0,7284998 |
| 1155_at | TBCB | 0,05051354 | 9,8767325 | 0,67946746 | 0,50752569 | 0,7284998 |
| 93183_at | PIGM | -0,19936931 | 7,42700393 | -0,67932747 | 0,50761181 | 0,7285618 |
| 100128164_a | LOC10012816 | -0,06071421 | 4,1800139 | -0,67926333 | 0,50765127 | 0,7285618 |
| 388394_at | RPRML | -0,07316515 | 5,01161847 | -0,67910001 | 0,50775176 | 0,7285618 |
| 100507533_a | SOX21-AS1 | -0,49682285 | 5,41241887 | -0,67907592 | 0,50776658 | 0,7285618 |
| 23520_at | ANP32C | -0,04192766 | 3,70848374 | -0,67897694 | 0,50782748 | 0,7285618 |
| 3927_at | LASP1 | 0,06359694 | 10,9414543 | 0,67894106 | 0,50784956 | 0,7285618 |
| 169693_at | TMEM252 | -0,05986092 | 3,48435562 | -0,67887881 | 0,50788787 | 0,7285618 |
| 64137_at | ABCG4 | -0,06525072 | 5,55860887 | -0,67885999 | 0,50789945 | 0,7285618 |
| 79867_at | TCTN2 | -0,15227136 | 7,14561774 | -0,67884403 | 0,50790928 | 0,7285618 |
| 149461_at | CLDN19 | -0,08824468 | 6,01427353 | -0,67865806 | 0,50802373 | 0,7286289 |
| 1004_at | CDH6 | 0,47271425 | 7,32981843 | 0,67864511 | 0,50803171 | 0,7286289 |
| 6664_at | SOX11 | -0,04481071 | 4,70905885 | -0,67850002 | 0,50812102 | 0,72870274 |
| 5782_at | PTPN12 | -0,14741076 | 7,84259025 | -0,67836296 | 0,50820539 | 0,72876548 |
| 430_at | ASCL2 | 0,05753637 | 4,54932261 | 0,67822389 | 0,50829101 | 0,72876548 |
| 23269_at | MGA | 0,11126568 | 7,42104471 | 0,67822149 | 0,50829249 | 0,72876548 |
| 23429_at | RYBP | 0,17597967 | 8,73908264 | 0,67816259 | 0,50832876 | 0,72876548 |
| 10276_at | NET1 | -0,13737601 | 9,8967164 | -0,67812171 | 0,50835393 | 0,72876548 |
| 2862_at | MLNR | -0,09147969 | 4,41672689 | -0,6780587 | 0,50839273 | 0,72876686 |
| 23787_at | MTCH1 | -0,05278036 | 11,0903435 | -0,67796905 | 0,50844793 | 0,72879176 |
| 5319_at | PLA2G1B | 0,04958749 | 3,9221845 | 0,67776641 | 0,50857272 | 0,72888994 |
| 92170_at | MTG1 | -0,07608588 | 8,45547898 | -0,67773494 | 0,5085921 | 0,72888994 |
| 2168_at | FABP1 | -0,0552413 | 4,27543594 | -0,67758504 | 0,50868444 | 0,7289503 |
| 939_at | CD27 | -0,08116381 | 5,31277737 | -0,67754369 | 0,50870991 | 0,7289503 |
| 7547_at | ZIC3 | -0,07789516 | 3,53534941 | -0,67736168 | 0,50882203 | 0,72905674 |

| | | | | | | |
|---|---|---|---|---|---|---|
| 1499_at | CTNNB1 | -0,12253561 | 9,87963112 | -0,67707269 | 0,50900008 | 0,72925761 |
| 63935_at | PCIF1 | -0,04727369 | 7,4326104 | -0,67675246 | 0,50919743 | 0,72946802 |
| 100506660_a | DDX11-AS1 | -0,07078281 | 3,4861449 | -0,67671149 | 0,50922268 | 0,72946802 |
| 280664_at | WFDC10B | -0,05641152 | 3,55597751 | -0,67658627 | 0,50929987 | 0,72952434 |
| 28465_at | IGHV1-46 | -0,09500392 | 6,11709583 | -0,6764811 | 0,50936469 | 0,72956295 |
| 5356_at | PLRG1 | -0,16902525 | 8,17247353 | -0,67632766 | 0,50945929 | 0,72959126 |
| 26261_at | FBXO24 | -0,0504777 | 4,59237496 | -0,67632616 | 0,50946021 | 0,72959126 |
| 60592_at | SCOC | 0,17365656 | 9,15648505 | 0,67599705 | 0,50966314 | 0,72982761 |
| 7621_at | ZNF70 | 0,05440138 | 6,2089787 | 0,67580298 | 0,50978282 | 0,72994473 |
| 27232_at | GNMT | -0,08778897 | 5,21960299 | -0,67566051 | 0,50987069 | 0,72995134 |
| 79084_at | WDR77 | -0,1534387 | 9,20371173 | -0,67555834 | 0,50993371 | 0,72995134 |
| 643332_at | ECRP | -0,09405186 | 3,60818331 | -0,675524 | 0,5099549 | 0,72995134 |
| 84856_at | LINC00839 | -0,06575537 | 5,12822191 | -0,67533847 | 0,51006935 | 0,72995134 |
| 4134_at | MAP4 | -0,08136147 | 7,03205057 | -0,67525162 | 0,51012293 | 0,72995134 |
| 5365_at | PLXNB3 | -0,08411533 | 5,96837648 | -0,67524065 | 0,5101297 | 0,72995134 |
| 645638_at | WFDC21P | 0,2548396 | 5,89188275 | 0,67521732 | 0,5101441 | 0,72995134 |
| 9611_at | NCOR1 | 0,15931822 | 8,18560177 | 0,67521669 | 0,51014448 | 0,72995134 |
| 221472_at | FGD2 | 0,06218059 | 5,78896599 | 0,67519113 | 0,51016025 | 0,72995134 |
| 349565_at | NMNAT3 | -0,04930165 | 5,30563022 | -0,67518121 | 0,51016637 | 0,72995134 |
| 131873_at | COL6A6 | -0,06566436 | 3,42032772 | -0,67500693 | 0,51027392 | 0,73001505 |
| 254428_at | SLC41A1 | -0,10357471 | 8,35802874 | -0,67498268 | 0,51028888 | 0,73001505 |
| 79947_at | DHDDS | -0,07527085 | 8,01769011 | -0,6749248 | 0,51032459 | 0,73001505 |
| 66002_at | CYP4F12 | 0,29820628 | 6,61876781 | 0,67469565 | 0,51046602 | 0,73008145 |
| 285761_at | DCBLD1 | 0,08133923 | 7,78010409 | 0,6746616 | 0,51048704 | 0,73008145 |
| 56477_at | CCL28 | 0,06193832 | 4,27107405 | 0,67451245 | 0,5105791 | 0,73008145 |
| 84343_at | HPS3 | 0,0827437 | 8,63126977 | 0,67450326 | 0,51058478 | 0,73008145 |
| 8547_at | FCN3 | -0,09233826 | 4,92162661 | -0,67445317 | 0,51061569 | 0,73008145 |
| 3782_at | KCNN3 | -0,05444699 | 3,85882834 | -0,67443005 | 0,51062997 | 0,73008145 |
| 440173_at | LOC440173 | 0,48873699 | 5,49995878 | 0,67435112 | 0,5106787 | 0,73008145 |
| 2900_at | GRIK4 | -0,06545025 | 4,29837121 | -0,6743036 | 0,51070803 | 0,73008145 |
| 199974_at | CYP4Z1 | 0,10231905 | 3,86324824 | 0,67429698 | 0,51071212 | 0,73008145 |
| 1465_at | CSRP1 | -0,12756066 | 10,1977292 | -0,67400729 | 0,51089099 | 0,73024518 |
| 57535_at | KIAA1324 | -0,05155883 | 4,49113228 | -0,67398871 | 0,51090247 | 0,73024518 |
| 440730_at | TRIM67 | -0,06381782 | 5,56038763 | -0,67382813 | 0,51100163 | 0,73033272 |
| 7375_at | USP4 | 0,09087822 | 8,077554 | 0,67364592 | 0,51111417 | 0,73035735 |
| 1589_at | CYP21A2 | 0,0824979 | 5,39552698 | 0,67361603 | 0,51113263 | 0,73035735 |
| 116159_at | CYYR1 | -0,05158822 | 3,5473873 | -0,67357189 | 0,5111599 | 0,73035735 |
| 439949_at | PRKCQ-AS1 | -0,1383184 | 4,21970387 | -0,67355469 | 0,51117052 | 0,73035735 |
| 222611_at | ADGRF2 | 0,06868107 | 3,41230464 | 0,67342778 | 0,51124892 | 0,73041518 |
| 54860_at | MS4A12 | -0,07421018 | 3,35511856 | -0,67334759 | 0,51129846 | 0,73043179 |
| 101927027_a | LOC10192702 | 0,13746677 | 5,85843749 | 0,67326756 | 0,5113479 | 0,73044825 |
| 8876_at | VNN1 | 0,04071861 | 3,08429073 | 0,67290841 | 0,51156983 | 0,73071108 |
| 55602_at | CDKN2AIP | 0,22361046 | 7,23167631 | 0,67281725 | 0,51162617 | 0,73073736 |
| 10558_at | SPTLC1 | 0,05023245 | 8,01145852 | 0,67272515 | 0,51168309 | 0,73076448 |
| 101928052_a | LOC10192805 | 0,05154994 | 2,75003294 | 0,67251662 | 0,51181198 | 0,73089437 |
| 163747_at | LEXM | -0,08986827 | 4,89450103 | -0,67239828 | 0,51188514 | 0,73094465 |
| 100126791_a | EGOT | -0,08642859 | 3,45303993 | -0,67230432 | 0,51194323 | 0,73097342 |
| 6641_at | SNTB1 | -0,04470895 | 4,07952507 | -0,67222696 | 0,51199106 | 0,73097979 |
| 5376_at | PMP22 | -0,15127272 | 5,22583184 | -0,67217435 | 0,51202359 | 0,73097979 |
| 255220_at | TXNDC8 | -0,07080771 | 3,91646383 | -0,67210248 | 0,51206802 | 0,73098906 |

| | | | | | | |
|---|---|---|---|---|---|---|
| 25817_at | FAM19A5 | -0,05421833 | 4,27711736 | -0,67182033 | 0,5122425 | 0,73113053 |
| 8022_at | LHX3 | -0,08171912 | 5,03348264 | -0,67181947 | 0,51224304 | 0,73113053 |
| 81579_at | PLA2G12A | -0,1829209 | 8,16182101 | -0,67154565 | 0,5124124 | 0,73125218 |
| 54876_at | DCAF16 | -0,10850599 | 7,39019042 | -0,67151616 | 0,51243064 | 0,73125218 |
| 100048912_a | CDKN2B-AS1 | -0,05520524 | 3,44510569 | -0,67139724 | 0,5125042 | 0,73125218 |
| 1308_at | COL17A1 | 0,15108015 | 6,94565296 | 0,67136329 | 0,51252521 | 0,73125218 |
| 3707_at | ITPKB | -0,08432632 | 5,70807253 | -0,67130371 | 0,51256207 | 0,73125218 |
| 3579_at | CXCR2 | 0,30067106 | 4,61638316 | 0,67125595 | 0,51259162 | 0,73125218 |
| 719_at | C3AR1 | -0,06420887 | 3,7530266 | -0,67122006 | 0,51261382 | 0,73125218 |
| 53373_at | TPCN1 | -0,08960055 | 7,00850276 | -0,67119075 | 0,51263196 | 0,73125218 |
| 7380_at | UPK3A | 0,08468528 | 4,87013122 | 0,67092903 | 0,51279391 | 0,7314163 |
| 4438_at | MSH4 | -0,06258272 | 3,02409819 | -0,67082304 | 0,5128595 | 0,7314163 |
| 5431_at | POLR2B | -0,13798164 | 9,92718556 | -0,67082075 | 0,51286092 | 0,7314163 |
| 10421_at | CD2BP2 | -0,05062141 | 8,11601095 | -0,67073508 | 0,51291394 | 0,73143776 |
| 975_at | CD81 | 0,11112528 | 11,0653846 | 0,67054147 | 0,51303378 | 0,73155449 |
| 65220_at | NADK | -0,05048576 | 8,04516851 | -0,6703593 | 0,51314655 | 0,73162073 |
| 9403_at | SEP15 | -0,09364709 | 11,1171889 | -0,67029127 | 0,51318867 | 0,73162073 |
| 150568_at | LINC01102 | -0,06957877 | 3,28487037 | -0,67028239 | 0,51319417 | 0,73162073 |
| 1182_at | CLCN3 | 0,19824769 | 8,45902425 | 0,66991773 | 0,51341996 | 0,73187633 |
| 1943_at | EFNA2 | -0,10748416 | 5,07012352 | -0,66982306 | 0,51347859 | 0,73187633 |
| 4009_at | LMX1A | -0,10882825 | 4,44670831 | -0,66980877 | 0,51348744 | 0,73187633 |
| 5314_at | PKHD1 | -0,06361555 | 3,86803346 | -0,66929603 | 0,51380505 | 0,73219333 |
| 1641_at | DCX | 0,04658695 | 3,29227001 | 0,6692801 | 0,51381492 | 0,73219333 |
| 91012_at | CERS5 | 0,05241897 | 6,72942747 | 0,66921696 | 0,51385404 | 0,73219333 |
| 6403_at | SELP | -0,05508866 | 4,60314058 | -0,6691576 | 0,51389082 | 0,73219333 |
| 7391_at | USF1 | 0,10815042 | 7,43236705 | 0,66914294 | 0,5138999 | 0,73219333 |
| 60681_at | FKBP10 | -0,06251786 | 5,5553246 | -0,66906157 | 0,51395032 | 0,73221101 |
| 10794_at | ZNF460 | 0,09163351 | 5,59216413 | 0,66863208 | 0,51421649 | 0,73245148 |
| 8728_at | ADAM19 | 0,06795475 | 4,97389574 | 0,66860817 | 0,51423131 | 0,73245148 |
| 10317_at | B3GALT5 | -0,09433109 | 3,80419736 | -0,66860084 | 0,51423586 | 0,73245148 |
| 339123_at | JMJD8 | -0,03649877 | 8,76618118 | -0,66854382 | 0,5142712 | 0,73245148 |
| 3623_at | INHA | 0,06017402 | 4,51341371 | 0,66826814 | 0,5144421 | 0,73264071 |
| 3831_at | KLC1 | -0,08664241 | 8,47959442 | -0,66814528 | 0,51451828 | 0,73267204 |
| 50487_at | PLA2G3 | -0,09941614 | 5,21656679 | -0,66810997 | 0,51454017 | 0,73267204 |
| 8635_at | RNASET2 | 0,12660098 | 8,9005338 | 0,66791894 | 0,51465863 | 0,73278654 |
| 5212_at | VIT | -0,05015296 | 4,15691211 | -0,6676801 | 0,51480675 | 0,73294327 |
| 6405_at | SEMA3F | 0,10547613 | 8,46587309 | 0,6675355 | 0,51489645 | 0,73301679 |
| 2230_at | FDX1 | -0,08330213 | 7,98264131 | -0,66744199 | 0,51495445 | 0,7330452 |
| 93273_at | LEMD1 | -0,0743866 | 3,47517273 | -0,66717706 | 0,51511882 | 0,73318113 |
| 100130717_a | CECR5-AS1 | -0,07734376 | 3,85562403 | -0,66714778 | 0,51513698 | 0,73318113 |
| 113189_at | CHST14 | -0,14949062 | 8,43716674 | -0,66710403 | 0,51516413 | 0,73318113 |
| 6331_at | SCN5A | 0,05469455 | 4,85735076 | 0,66704064 | 0,51520346 | 0,73318294 |
| 148156_at | ZNF558 | 0,14509462 | 7,35365822 | 0,66688239 | 0,51530166 | 0,73326851 |
| 259215_at | LY6G6F | -0,09649667 | 4,7237213 | -0,66678503 | 0,51536208 | 0,73330032 |
| 92595_at | ZNF764 | -0,04819092 | 6,90217766 | -0,66668314 | 0,51542532 | 0,73333614 |
| 64746_at | ACBD3 | 0,0621415 | 9,61986345 | 0,66659471 | 0,51548021 | 0,73336006 |
| 64784_at | CRTC3 | 0,0667871 | 8,86433088 | 0,66594088 | 0,51588613 | 0,73388336 |
| 200035_at | NUDT17 | 0,08142682 | 6,26752164 | 0,66574265 | 0,51600925 | 0,73400429 |
| 4025_at | LPO | -0,08259122 | 4,5873781 | -0,66531889 | 0,51627246 | 0,73432448 |
| 84539_at | MCHR2 | -0,07982217 | 4,30911977 | -0,66511371 | 0,51639995 | 0,73445158 |

| | | | | | | |
|---|---|---|---|---|---|---|
| 101928851_a | LOC10192885 | -0,06058301 | 3,58399242 | -0,66501342 | 0,51646226 | 0,73448598 |
| 495_at | ATP4A | -0,06919486 | 4,43364988 | -0,66492861 | 0,51651496 | 0,73450669 |
| 9655_at | SOCS5 | 0,12582391 | 8,10672274 | 0,66472395 | 0,51664215 | 0,73456458 |
| 339822_at | LINC01115 | -0,06330967 | 3,78257325 | -0,66469966 | 0,51665724 | 0,73456458 |
| 53822_at | FXYD7 | -0,07167783 | 5,0607403 | -0,66465102 | 0,51668748 | 0,73456458 |
| 55643_at | BTBD2 | 0,11181813 | 7,96724359 | 0,66460268 | 0,51671752 | 0,73456458 |
| 115572_at | FAM46B | 0,19162083 | 5,60331366 | 0,66450062 | 0,51678096 | 0,73456458 |
| 64420_at | SUSD1 | 0,21844767 | 5,69684863 | 0,66449499 | 0,51678446 | 0,73456458 |
| 202243_at | CCDC125 | 0,12984716 | 5,6332534 | 0,66440561 | 0,51684002 | 0,73458934 |
| 79977_at | GRHL2 | 0,08778247 | 9,73638638 | 0,6642382 | 0,5169441 | 0,73462377 |
| 728392_at | LOC728392 | -0,07971386 | 5,01947117 | -0,66415138 | 0,51699808 | 0,73462377 |
| 342918_at | C19orf81 | 0,10521916 | 5,42689038 | 0,66407476 | 0,51704572 | 0,73462377 |
| 8794_at | TNFRSF10C | -0,04218526 | 3,46532541 | -0,66401597 | 0,51708227 | 0,73462377 |
| 55668_at | GPATCH2L | 0,03684174 | 6,37865131 | 0,6639697 | 0,51711104 | 0,73462377 |
| 100128176_a | LOC10012817 | -0,07529055 | 4,17728617 | -0,6639628 | 0,51711534 | 0,73462377 |
| 105369214_a | LOC10536921 | 0,09975607 | 3,92259976 | 0,6639196 | 0,5171422 | 0,73462377 |
| 285367_at | RPUSD3 | -0,14324471 | 8,50372984 | -0,66387596 | 0,51716934 | 0,73462377 |
| 220108_at | FAM124A | -0,08179207 | 4,54209967 | -0,66377358 | 0,51723301 | 0,73466004 |
| 3768_at | KCNJ12 | -0,06965267 | 4,83630676 | -0,6634765 | 0,51741778 | 0,7348683 |
| 117248_at | GALNT15 | -0,05861729 | 3,85851391 | -0,66322316 | 0,51757539 | 0,7350161 |
| 60496_at | AASDHPPT | -0,0753483 | 9,29187869 | -0,66318655 | 0,51759816 | 0,7350161 |
| 10526_at | IPO8 | -0,08369513 | 6,77560466 | -0,66311819 | 0,5176407 | 0,73502232 |
| 54732_at | TMED9 | 0,05483914 | 10,4782148 | 0,66289509 | 0,51777952 | 0,73516524 |
| 23034_at | SAMD4A | -0,11813988 | 6,02838906 | -0,66274381 | 0,51787366 | 0,73524472 |
| 63946_at | DMRTC2 | -0,05387888 | 4,23536343 | -0,66267589 | 0,51791593 | 0,73525054 |
| 29909_at | GPR171 | -0,05596331 | 3,28214172 | -0,66259828 | 0,51796424 | 0,73526493 |
| 3934_at | LCN2 | 0,38412545 | 9,57871311 | 0,6623727 | 0,51810465 | 0,73537729 |
| 3562_at | IL3 | -0,06901755 | 3,6749863 | -0,66234846 | 0,51811974 | 0,73537729 |
| 4601_at | MXI1 | 0,06143323 | 7,97359418 | 0,6619933 | 0,51834087 | 0,73563694 |
| 91464_at | ISX | -0,1034024 | 4,06343818 | -0,66184708 | 0,51843192 | 0,73571195 |
| 10257_at | ABCC4 | 0,30262135 | 7,53325229 | 0,66177395 | 0,51847746 | 0,73572238 |
| 246119_at | TTTY10 | -0,05788588 | 3,76792792 | -0,66125245 | 0,51880229 | 0,73612909 |
| 57657_at | HCN3 | -0,1009732 | 5,41898676 | -0,66116561 | 0,51885639 | 0,73615163 |
| 101928327_a | LOC10192832 | 0,06582687 | 3,45330297 | 0,66099706 | 0,51896141 | 0,73617272 |
| 26289_at | AK5 | -0,05641722 | 4,12184223 | -0,6609648 | 0,51898151 | 0,73617272 |
| 11280_at | SCN11A | 0,04919565 | 2,98499243 | 0,66095775 | 0,51898591 | 0,73617272 |
| 388503_at | C3P1 | -0,07501208 | 4,47315673 | -0,66089274 | 0,51902642 | 0,73617598 |
| 6909_at | TBX2 | -0,08164422 | 6,79526506 | -0,66062198 | 0,51919516 | 0,73632875 |
| 84292_at | WDR83 | -0,07100322 | 6,72313291 | -0,66059723 | 0,51921058 | 0,73632875 |
| 1602_at | DACH1 | -0,04139883 | 3,07263891 | -0,66025368 | 0,51942474 | 0,73650617 |
| 56547_at | MMP26 | -0,05936078 | 3,80250315 | -0,66024301 | 0,51943139 | 0,73650617 |
| 101927017_a | LINC01180 | -0,08038498 | 4,18579222 | -0,66021253 | 0,51945039 | 0,73650617 |
| 101928539_a | LOC10192853 | -0,03786232 | 2,97273474 | -0,66007174 | 0,51953816 | 0,73657641 |
| 8858_at | PROZ | -0,05364283 | 5,28583706 | -0,66000531 | 0,51957959 | 0,73658093 |
| 3456_at | IFNB1 | 0,05181282 | 3,48443665 | 0,65986212 | 0,51966888 | 0,73665329 |
| 8525_at | DGKZ | -0,17801821 | 9,43390043 | -0,65976076 | 0,51973208 | 0,73668391 |
| 252995_at | FNDC5 | -0,08401779 | 5,46090563 | -0,65970482 | 0,51976697 | 0,73668391 |
| 390999_at | PRAMEF12 | -0,09125675 | 4,03314492 | -0,65947774 | 0,5199086 | 0,73677802 |
| 1577_at | CYP3A5 | 0,49318944 | 6,06443878 | 0,65947571 | 0,51990986 | 0,73677802 |
| 25923_at | ATL3 | -0,08087798 | 7,42426736 | -0,65921494 | 0,52007253 | 0,73695434 |

| | | | | | | |
|---|---|---|---|---|---|---|
| 57396_at | CLK4 | 0,18634828 | 6,89424205 | 0,65914634 | 0,52011533 | 0,73696077 |
| 100506125_a | LOC10050612 | 0,08383527 | 4,90991526 | 0,65902706 | 0,52018976 | 0,73701201 |
| 170689_at | ADAMTS15 | 0,11327459 | 6,79789443 | 0,65884976 | 0,52030039 | 0,73706997 |
| 131375_at | LYZL4 | -0,09778658 | 4,69373364 | -0,65872872 | 0,52037592 | 0,73706997 |
| 8028_at | MLLT10 | -0,10028342 | 6,398297 | -0,65866593 | 0,52041511 | 0,73706997 |
| 23275_at | POFUT2 | -0,04830089 | 6,26626734 | -0,65859237 | 0,52046101 | 0,73706997 |
| 93611_at | FBXO44 | -0,05978957 | 5,47508465 | -0,65854071 | 0,52049326 | 0,73706997 |
| 635_at | BHMT | -0,13169398 | 4,51775803 | -0,65843844 | 0,52055709 | 0,73706997 |
| 8189_at | SYMPK | 0,07711397 | 7,49199879 | 0,65842672 | 0,52056441 | 0,73706997 |
| 55751_at | TMEM184C | 0,15549203 | 9,52791541 | 0,65842079 | 0,52056811 | 0,73706997 |
| 4618_at | MYF6 | -0,06903198 | 3,87838851 | -0,6584097 | 0,52057503 | 0,73706997 |
| 56548_at | CHST7 | -0,08384142 | 5,80306848 | -0,65834548 | 0,52061512 | 0,73707254 |
| 1266_at | CNN3 | 0,34305169 | 8,90482873 | 0,658248 | 0,52067597 | 0,73710452 |
| 221184_at | CPNE2 | 0,11146956 | 8,63392995 | 0,6580374 | 0,52080745 | 0,73719709 |
| 157739_at | TDH | -0,07039395 | 4,990621 | -0,65798013 | 0,52084322 | 0,73719709 |
| 9633_at | MTL5 | 0,06896161 | 4,61023076 | 0,65795938 | 0,52085617 | 0,73719709 |
| 9705_at | ST18 | -0,04787596 | 3,06477807 | -0,65764717 | 0,52105113 | 0,7374182 |
| 4501_at | MT1X | -0,1158117 | 10,0527486 | -0,65758661 | 0,52108896 | 0,7374182 |
| 401442_at | LOC401442 | -0,07717099 | 3,65922878 | -0,65747793 | 0,52115683 | 0,73746008 |
| 83900_at | KRTAP9-3 | -0,0637765 | 4,1282189 | -0,65697943 | 0,52146825 | 0,73784655 |
| 404744_at | NPSR1-AS1 | -0,07165509 | 3,80657925 | -0,65681984 | 0,52156797 | 0,73793345 |
| 138716_at | RPP25L | -0,06537657 | 8,96108222 | -0,65672826 | 0,5216252 | 0,73796021 |
| 58155_at | PTBP2 | 0,0833515 | 6,30872386 | 0,65655126 | 0,52173582 | 0,73805085 |
| 93100_at | NAPRT | 0,12188676 | 9,60294007 | 0,65646259 | 0,52179124 | 0,73805085 |
| 80381_at | CD276 | -0,07807957 | 8,36283406 | -0,65639146 | 0,5218357 | 0,73805085 |
| 150142_at | ZNF295-AS1 | -0,09719721 | 4,10932158 | -0,65638054 | 0,52184253 | 0,73805085 |
| 745_at | MYRF | -0,05783484 | 5,27369548 | -0,65627377 | 0,52190927 | 0,73809106 |
| 102723165_a | LINC01169 | -0,05363328 | 3,89954404 | -0,65612896 | 0,5219998 | 0,73809963 |
| 10186_at | LHFP | 0,23730543 | 8,22437768 | 0,65612803 | 0,52200038 | 0,73809963 |
| 29923_at | HILPDA | 0,11724116 | 8,46660973 | 0,65607597 | 0,52203293 | 0,73809963 |
| 10657_at | KHDRBS1 | -0,06905803 | 10,0897085 | -0,65601891 | 0,5220686 | 0,73809963 |
| 64518_at | TEKT3 | -0,05164917 | 3,39886423 | -0,65584701 | 0,52217609 | 0,73819741 |
| 284485_at | RIIAD1 | -0,08482415 | 4,68424885 | -0,65569606 | 0,52227048 | 0,73827667 |
| 352909_at | DNAAF3 | -0,08280926 | 5,84607835 | -0,65540147 | 0,52245472 | 0,73840133 |
| 54788_at | DNAJB12 | -0,04149647 | 7,41003352 | -0,65538799 | 0,52246315 | 0,73840133 |
| 285735_at | LINC00326 | -0,06549139 | 3,05796609 | -0,65537119 | 0,52247366 | 0,73840133 |
| 8001_at | GLRA3 | -0,04147157 | 2,86212574 | -0,65529058 | 0,52252408 | 0,73841841 |
| 6683_at | SPAST | -0,08914522 | 7,27160767 | -0,65520549 | 0,52257731 | 0,73843205 |
| 84826_at | SFT2D3 | -0,04095066 | 7,69971373 | -0,65515259 | 0,5226104 | 0,73843205 |
| 490_at | ATP2B1 | -0,08292386 | 7,92962568 | -0,65504326 | 0,5226788 | 0,73846514 |
| 4801_at | NFYB | -0,05048724 | 7,2251287 | -0,65499262 | 0,52271049 | 0,73846514 |
| 8620_at | NPFF | -0,06082336 | 5,24199974 | -0,65487284 | 0,52278543 | 0,73851685 |
| 2027_at | ENO3 | -0,10089569 | 4,81103926 | -0,65451995 | 0,52300627 | 0,73877464 |
| 91750_at | LIN52 | -0,09976961 | 6,79842722 | -0,65439979 | 0,52308147 | 0,73882669 |
| 87_at | ACTN1 | -0,19738288 | 10,7950962 | -0,65417027 | 0,52322515 | 0,73897544 |
| 81793_at | TLR10 | -0,04255264 | 3,54857271 | -0,65410775 | 0,52326428 | 0,73897653 |
| 100499489_a | LOC10049948 | -0,05773712 | 4,92759076 | -0,6539775 | 0,52334583 | 0,73903751 |
| 170392_at | OIT3 | -0,0513726 | 3,58390802 | -0,65379992 | 0,52345702 | 0,73914034 |
| 101928894_a | LOC10192889 | -0,05317886 | 3,98978132 | -0,65345335 | 0,52367406 | 0,73939261 |
| 101929450_a | LOC10192945 | 0,04526958 | 3,4041488 | 0,65289177 | 0,52402585 | 0,73983243 |

| | | | | | | |
|---|---|---|---|---|---|---|
| 84616_at | KRTAP4-4 | 0,06721361 | 4,0694608 | 0,65283348 | 0,52406237 | 0,73983243 |
| 51151_at | SLC45A2 | -0,06605352 | 4,2326591 | -0,65271389 | 0,52413731 | 0,73988399 |
| 54900_at | LAX1 | 0,05881329 | 3,97949196 | 0,65257179 | 0,52422636 | 0,73992886 |
| 57666_at | FBRSL1 | -0,05316172 | 7,19481788 | -0,65253091 | 0,52425198 | 0,73992886 |
| 79906_at | MORN1 | -0,07423759 | 5,30152134 | -0,65247929 | 0,52428433 | 0,73992886 |
| 196883_at | ADCY4 | 0,05632201 | 5,89057189 | 0,65227311 | 0,52441356 | 0,74001303 |
| 4988_at | OPRM1 | -0,05019825 | 3,38445938 | -0,65226156 | 0,5244208 | 0,74001303 |
| 10888_at | GPR83 | -0,0633583 | 3,90893985 | -0,65214676 | 0,52449276 | 0,74003133 |
| 1113_at | CHGA | -0,05005405 | 4,72865467 | -0,6521183 | 0,5245106 | 0,74003133 |
| 55081_at | IFT57 | -0,09677499 | 7,39540459 | -0,65200888 | 0,5245792 | 0,7400739 |
| 2252_at | FGF7 | -0,05000376 | 3,09505044 | -0,65179307 | 0,52471451 | 0,74017134 |
| 286135_at | FAM183CP | -0,04218002 | 4,1291283 | -0,65177615 | 0,52472511 | 0,74017134 |
| 441024_at | MTHFD2L | -0,06509664 | 4,02712448 | -0,65138072 | 0,52497309 | 0,74046692 |
| 100506791_a | LINC01511 | -0,08712595 | 5,15217088 | -0,65109948 | 0,5251495 | 0,7406615 |
| 122945_at | NOXRED1 | -0,0778448 | 4,04351692 | -0,65096124 | 0,52523622 | 0,74072958 |
| 100506343_a | FAM212B-AS1 | -0,04687929 | 5,44097796 | -0,65075258 | 0,52536714 | 0,74085998 |
| 60529_at | ALX4 | -0,08006804 | 6,03950437 | -0,65021541 | 0,52570426 | 0,7412811 |
| 133584_at | EGFLAM | -0,06412099 | 4,86598737 | -0,6501468 | 0,52574733 | 0,74128757 |
| 3586_at | IL10 | -0,04129342 | 3,73094616 | -0,64999777 | 0,52584088 | 0,74136521 |
| 2580_at | GAK | 0,07218883 | 9,0145986 | 0,64949333 | 0,52615762 | 0,74175748 |
| 8574_at | AKR7A2 | -0,07517127 | 9,82676382 | -0,64927049 | 0,52629757 | 0,74189975 |
| 7841_at | MOGS | -0,07108828 | 8,26384374 | -0,64920999 | 0,52633556 | 0,74189975 |
| 80129_at | CCDC170 | -0,04856372 | 3,42463711 | -0,64877403 | 0,52660944 | 0,74223148 |
| 115703_at | ARHGAP33 | -0,07602667 | 6,24483133 | -0,64864843 | 0,52668836 | 0,74227051 |
| 120376_at | COLCA2 | 0,27557587 | 6,10910449 | 0,6486073 | 0,5267142 | 0,74227051 |
| 60675_at | PROK2 | 0,06554419 | 3,80904305 | 0,64852082 | 0,52676855 | 0,7422928 |
| 63892_at | THADA | -0,09097106 | 6,68631916 | -0,64844414 | 0,52681674 | 0,7423064 |
| 147650_at | SPACA6P | 0,19281921 | 6,1894298 | 0,64833241 | 0,52688695 | 0,74235104 |
| 100506070_a | RBFADN | -0,04038673 | 3,71985402 | -0,6482237 | 0,52695528 | 0,74237955 |
| 8345_at | HIST1H2BH | -0,07592987 | 7,41455839 | -0,6481071 | 0,52702857 | 0,74237955 |
| 201181_at | ZNF385C | 0,07195731 | 4,83220819 | 0,64798947 | 0,52710251 | 0,74237955 |
| 131034_at | CPNE4 | -0,04589391 | 3,12255629 | -0,64797533 | 0,5271114 | 0,74237955 |
| 7252_at | TSHB | -0,04671926 | 3,6458314 | -0,64793432 | 0,52713718 | 0,74237955 |
| 8320_at | EOMES | -0,05554601 | 4,80489395 | -0,64773845 | 0,52726033 | 0,74237955 |
| 388789_at | LINC00493 | -0,05302207 | 9,05502277 | -0,64771331 | 0,52727613 | 0,74237955 |
| 1725_at | DHPS | -0,0483326 | 8,87155116 | -0,64771043 | 0,52727795 | 0,74237955 |
| 143187_at | VTI1A | 0,03162348 | 5,72822294 | 0,64769692 | 0,52728644 | 0,74237955 |
| 388336_at | SHISA6 | -0,06806197 | 3,76029923 | -0,64766691 | 0,5273053 | 0,74237955 |
| 64771_at | C6orf106 | -0,08615284 | 8,06338711 | -0,64757527 | 0,52736293 | 0,74237955 |
| 9482_at | STX8 | 0,04399874 | 6,85892864 | 0,64749338 | 0,52741443 | 0,74237955 |
| 253868_at | C20orf166-AS | -0,06924697 | 6,09676105 | -0,64744193 | 0,52744678 | 0,74237955 |
| 55697_at | VAC14 | 0,10847134 | 7,07285837 | 0,64738228 | 0,52748429 | 0,74237955 |
| 4145_at | MATK | -0,07965271 | 5,30930118 | -0,64725996 | 0,52756122 | 0,74237955 |
| 94015_at | TTYH2 | -0,05289363 | 5,58524656 | -0,64721995 | 0,52758638 | 0,74237955 |
| 54576_at | UGT1A8 | 0,58384484 | 7,11243978 | 0,64721281 | 0,52759088 | 0,74237955 |
| 101927553_a | LOC10192755 | -0,06638282 | 3,99448215 | -0,64713712 | 0,52763848 | 0,74237955 |
| 144193_at | AMDHD1 | -0,07196613 | 3,9701859 | -0,64702106 | 0,52771148 | 0,74237955 |
| 166012_at | CHST13 | -0,0644576 | 5,10623774 | -0,64701316 | 0,52771646 | 0,74237955 |
| 5544_at | PRB3 | -0,05892253 | 5,79765782 | -0,64700697 | 0,52772035 | 0,74237955 |
| 285877_at | POM121L12 | -0,10839394 | 5,83635397 | -0,64690757 | 0,52778288 | 0,74237955 |

| | | | | | | |
|---|---|---|---|---|---|---|
| 56731_at | SLC2A4RG | -0,06503946 | 7,88471609 | -0,64689054 | 0,52779359 | 0,74237955 |
| 23478_at | SEC11A | -0,03899694 | 10,4512325 | -0,64675971 | 0,5278759 | 0,74244111 |
| 29118_at | DDX25 | -0,06410203 | 3,42130085 | -0,64662595 | 0,52796006 | 0,74250527 |
| 91694_at | LONRF1 | 0,16680617 | 7,4860616 | 0,64654899 | 0,52800849 | 0,74251917 |
| 128876_at | FAM83C | -0,0564135 | 4,30199213 | -0,64626357 | 0,5281881 | 0,74271753 |
| 23250_at | ATP11A | -0,10778649 | 6,37280419 | -0,64619369 | 0,52823208 | 0,74272515 |
| 131177_at | FAM3D | -0,06103018 | 5,32758199 | -0,64595209 | 0,52838416 | 0,74288476 |
| 730102_at | LOC730102 | -0,04905334 | 2,97733585 | -0,64586313 | 0,52844016 | 0,74290763 |
| 79852_at | EPHX3 | -0,0802881 | 4,76237581 | -0,64577517 | 0,52849553 | 0,74290763 |
| 57194_at | ATP10A | -0,0762135 | 4,60810684 | -0,64574246 | 0,52851613 | 0,74290763 |
| 5868_at | RAB5A | 0,076865 | 9,71881734 | 0,64562291 | 0,5285914 | 0,74295922 |
| 653160_at | LOC653160 | -0,07476807 | 6,19511699 | -0,64543008 | 0,52871281 | 0,74307566 |
| 54882_at | ANKHD1 | 0,05801336 | 7,68277995 | 0,64514796 | 0,52889049 | 0,74326159 |
| 2273_at | FHL1 | -0,05468413 | 4,33672966 | -0,6450675 | 0,52894117 | 0,74326159 |
| 26051_at | PPP1R16B | -0,05567013 | 4,04880232 | -0,64503622 | 0,52896087 | 0,74326159 |
| 222826_at | FAM217A | -0,05603142 | 3,40092915 | -0,64476211 | 0,52913354 | 0,74341748 |
| 5351_at | PLOD1 | -0,06946575 | 9,10126654 | -0,64473759 | 0,52914899 | 0,74341748 |
| 23171_at | GPD1L | 0,28812684 | 8,92506509 | 0,64452631 | 0,52928211 | 0,74350358 |
| 146177_at | VWA3A | -0,08102468 | 4,86375361 | -0,6445178 | 0,52928747 | 0,74350358 |
| 5802_at | PTPRS | -0,10712508 | 7,60398013 | -0,64427798 | 0,5294386 | 0,74364011 |
| 6559_at | SLC12A3 | -0,06019076 | 4,71037647 | -0,64417079 | 0,52950615 | 0,74364011 |
| 1627_at | DBN1 | 0,12459025 | 7,4539322 | 0,64407119 | 0,52956892 | 0,74364011 |
| 254394_at | MCM9 | 0,07756723 | 5,56241705 | 0,64403972 | 0,52958876 | 0,74364011 |
| 101928748_a | LOC10192874 | -0,05501707 | 3,61317965 | -0,64399984 | 0,52961389 | 0,74364011 |
| 80829_at | ZFP91 | -0,15766183 | 9,30516372 | -0,64399191 | 0,52961889 | 0,74364011 |
| 22854_at | NTNG1 | -0,05411836 | 3,82478441 | -0,64393479 | 0,5296549 | 0,74364011 |
| 3742_at | KCNA6 | -0,07473801 | 4,49939962 | -0,643817 | 0,52972915 | 0,74369016 |
| 5149_at | PDE6H | -0,05606419 | 3,33459519 | -0,64371287 | 0,52979479 | 0,74372811 |
| 55571_at | CNOT11 | -0,06369283 | 9,71340939 | -0,6435792 | 0,52987907 | 0,74379221 |
| 122616_at | C14orf79 | 0,07004312 | 6,39456365 | 0,64331605 | 0,530045 | 0,74394194 |
| 291_at | SLC25A4 | -0,0830037 | 7,26372724 | -0,64328754 | 0,53006298 | 0,74394194 |
| 3604_at | TNFRSF9 | -0,04620637 | 3,7551871 | -0,64315384 | 0,53014729 | 0,74400607 |
| 55512_at | SMPD3 | 0,09813026 | 5,12381857 | 0,64297383 | 0,53026083 | 0,7441112 |
| 152189_at | CMTM8 | -0,14005183 | 7,46246693 | -0,64275083 | 0,5304015 | 0,74425437 |
| 85016_at | C11orf70 | 0,25356348 | 4,1992556 | 0,64250843 | 0,53055443 | 0,74441474 |
| 9920_at | KBTBD11 | 0,24437275 | 6,21213806 | 0,64236064 | 0,53064768 | 0,74449136 |
| 846_at | CASR | -0,05277193 | 4,26730355 | -0,64185843 | 0,53096464 | 0,74472833 |
| 140_at | ADORA3 | -0,08560099 | 4,02584167 | -0,64184109 | 0,53097558 | 0,74472833 |
| 83884_at | SLC25A2 | -0,05953774 | 4,33010637 | -0,64184017 | 0,53097616 | 0,74472833 |
| 22878_at | TRAPPC8 | 0,12904577 | 8,26363137 | 0,6418317 | 0,53098151 | 0,74472833 |
| 100287482_a | SMKR1 | -0,11672081 | 5,28058762 | -0,64178673 | 0,5310099 | 0,74472833 |
| 150383_at | CDPF1 | -0,1273193 | 6,52206284 | -0,64161099 | 0,53112084 | 0,74474703 |
| 4124_at | MAN2A1 | 0,10180277 | 8,90223135 | 0,64155638 | 0,53115532 | 0,74474703 |
| 5291_at | PIK3CB | -0,10820241 | 7,1668906 | -0,64153973 | 0,53116583 | 0,74474703 |
| 6329_at | SCN4A | -0,08096464 | 4,17120227 | -0,64150389 | 0,53118845 | 0,74474703 |
| 728279_at | KRTAP2-2 | -0,06378901 | 3,41785273 | -0,64145941 | 0,53121654 | 0,74474703 |
| 134265_at | AFAP1L1 | -0,16033689 | 5,22792458 | -0,64138871 | 0,53126118 | 0,74475541 |
| 1728_at | NQO1 | 0,17025738 | 12,0147165 | 0,6412054 | 0,53137693 | 0,7448467 |
| 254312_at | LINC00710 | -0,06162641 | 3,40336197 | -0,64110682 | 0,53143918 | 0,7448467 |
| 651430_at | FLJ38576 | 0,07418074 | 3,27005843 | 0,64110189 | 0,5314423 | 0,7448467 |

| | | | | | | |
|---|---|---|---|---|---|---|
| 27065_at | NSG1 | -0,09409604 | 5,379199 | -0,64093828 | 0,53154563 | 0,74493733 |
| 100289019_a | SLC25A25-AS1 | -0,19438279 | 6,43242651 | -0,64077252 | 0,53165033 | 0,74502986 |
| 2512_at | FTL | -0,10341217 | 13,1120647 | -0,64070155 | 0,53169517 | 0,74503848 |
| 136259_at | KLF14 | -0,05312649 | 4,96002213 | -0,64053241 | 0,53180202 | 0,74513401 |
| 90768_at | LOC90768 | -0,05612661 | 4,40612305 | -0,64046258 | 0,53184614 | 0,74514162 |
| 3444_at | IFNA7 | 0,07807633 | 2,73765945 | 0,64032676 | 0,53193196 | 0,74520766 |
| 6815_at | STYX | 0,05878659 | 7,06324221 | 0,64011012 | 0,53206885 | 0,74534523 |
| 55364_at | IMPACT | -0,15532394 | 7,39842367 | -0,63997239 | 0,5321559 | 0,74537669 |
| 10215_at | OLIG2 | -0,04330136 | 4,18940237 | -0,63993634 | 0,53217868 | 0,74537669 |
| 105274304_a | LOC10527430 | 0,06084126 | 5,3649018 | 0,63983112 | 0,53224518 | 0,74537669 |
| 9213_at | XPR1 | 0,14564955 | 7,90486856 | 0,63980695 | 0,53226046 | 0,74537669 |
| 9498_at | SLC4A8 | -0,06886464 | 3,93081779 | -0,63976847 | 0,53228479 | 0,74537669 |
| 4100_at | MAGEA1 | -0,1886031 | 4,14317878 | -0,63965464 | 0,53235674 | 0,74542326 |
| 102723704_a | LOC10272370 | 0,05313454 | 3,22364466 | 0,63944759 | 0,53248763 | 0,74555235 |
| 8038_at | ADAM12 | -0,08269686 | 6,90838077 | -0,63935625 | 0,53254538 | 0,74557902 |
| 5794_at | PTPRH | -0,08020541 | 5,09588669 | -0,63919446 | 0,53264768 | 0,74561416 |
| 728081_at | LINC00290 | 0,04255478 | 3,09229982 | 0,63913804 | 0,53268336 | 0,74561416 |
| 3787_at | KCNS1 | -0,08581099 | 5,05974887 | -0,63913291 | 0,53268661 | 0,74561416 |
| 117196_at | MRGPRX4 | -0,04493994 | 3,7892102 | -0,63864801 | 0,53299329 | 0,74595644 |
| 55660_at | PRPF40A | -0,10396282 | 8,92580004 | -0,63856174 | 0,53304787 | 0,74595644 |
| 151174_at | LOC151174 | -0,07643272 | 5,24888702 | -0,63848989 | 0,53309332 | 0,74595644 |
| 65990_at | FAM173A | -0,05701701 | 8,09494032 | -0,63847309 | 0,53310395 | 0,74595644 |
| 6663_at | SOX10 | -0,07527754 | 5,37189908 | -0,63833807 | 0,53318937 | 0,74595644 |
| 51024_at | FIS1 | 0,10062714 | 9,5126475 | 0,63828043 | 0,53322584 | 0,74595644 |
| 101927769_a | LOC10192776 | -0,05131228 | 3,20727592 | -0,63827169 | 0,53323137 | 0,74595644 |
| 197187_at | SNAI3-AS1 | 0,06072744 | 5,31260036 | 0,63825658 | 0,53324094 | 0,74595644 |
| 51411_at | BIN2 | 0,07596694 | 3,84684972 | 0,6381659 | 0,53329831 | 0,74598253 |
| 29803_at | REPIN1 | 0,09012594 | 6,4696228 | 0,63804074 | 0,53337751 | 0,74601884 |
| 84667_at | HES7 | 0,05989854 | 5,52896694 | 0,63791893 | 0,5334546 | 0,74601884 |
| 50618_at | ITSN2 | 0,09631074 | 6,74594105 | 0,63791016 | 0,53346015 | 0,74601884 |
| 100507201_a | LOC10050720 | -0,03967242 | 2,98371179 | -0,63788009 | 0,53347918 | 0,74601884 |
| 728743_at | LOC728743 | 0,04887758 | 5,72356835 | 0,63778451 | 0,53353967 | 0,74604927 |
| 79001_at | VKORC1 | -0,0614898 | 9,81731905 | -0,63758066 | 0,53366871 | 0,74617554 |
| 255031_at | LINC00957 | 0,08016252 | 6,31412429 | 0,63739016 | 0,53378931 | 0,74628998 |
| 10816_at | SPINT3 | -0,07387714 | 3,60751564 | -0,63706429 | 0,53399564 | 0,74648922 |
| 22979_at | EFR3B | -0,0935397 | 4,96625277 | -0,63704269 | 0,53400931 | 0,74648922 |
| 7201_at | TRHR | -0,08608613 | 3,7152057 | -0,63683164 | 0,53414297 | 0,74662188 |
| 245935_at | DEFB122 | 0,06080989 | 3,43124689 | 0,63658746 | 0,53429763 | 0,74678387 |
| 26048_at | ZNF500 | 0,0399584 | 6,5572225 | 0,63651374 | 0,53434433 | 0,74679495 |
| 2814_at | GP5 | -0,05153781 | 4,16055718 | -0,63642953 | 0,53439767 | 0,74681532 |
| 133491_at | C5orf47 | 0,0391501 | 3,25020448 | 0,6363048 | 0,53447669 | 0,74687157 |
| 1236_at | CCR7 | 0,05907898 | 4,24117024 | 0,63606498 | 0,53462865 | 0,74702971 |
| 2983_at | GUCY1B3 | -0,04364204 | 3,5184441 | -0,63593684 | 0,53470985 | 0,74708898 |
| 1088_at | CEACAM8 | 0,05194901 | 3,36354659 | 0,63573369 | 0,5348386 | 0,74721467 |
| 114132_at | SIGLEC11 | -0,05885256 | 4,07536223 | -0,63546306 | 0,53501014 | 0,74733179 |
| 388963_at | C2orf81 | -0,05322679 | 6,53753329 | -0,63538868 | 0,53505729 | 0,74733179 |
| 338785_at | KRT79 | -0,05360581 | 4,6749813 | -0,6353524 | 0,53508028 | 0,74733179 |
| 9991_at | PTBP3 | -0,13160374 | 8,79450374 | -0,63527006 | 0,53513249 | 0,74733179 |
| 730227_at | LINC01136 | -0,06907666 | 4,54476962 | -0,63520371 | 0,53517455 | 0,74733179 |
| 3982_at | LIM2 | -0,06200097 | 4,5230072 | -0,63517912 | 0,53519014 | 0,74733179 |

| | | | | | | |
|---|---|---|---|---|---|---|
| 284307_at | ZIK1 | -0,20933261 | 6,57806967 | -0,63517303 | 0,535194 | 0,74733179 |
| 27012_at | KCNV1 | -0,03890573 | 3,44718919 | -0,6349376 | 0,53534329 | 0,74748607 |
| 100506504_a | LOC10050650 | -0,06343693 | 6,03063606 | -0,63486211 | 0,53539116 | 0,74749872 |
| 2916_at | GRM6 | 0,07118219 | 5,29848221 | 0,63469043 | 0,53550004 | 0,74750059 |
| 139728_at | PNCK | -0,08163409 | 5,95235872 | -0,63468311 | 0,53550468 | 0,74750059 |
| 403_at | ARL3 | 0,0420807 | 6,10078579 | 0,63467643 | 0,53550891 | 0,74750059 |
| 654841_at | LOC654841 | 0,06108894 | 4,4263418 | 0,63446738 | 0,53564151 | 0,74763151 |
| 9620_at | CELSR1 | -0,11280123 | 6,57344274 | -0,63418861 | 0,53581836 | 0,74782416 |
| 9826_at | ARHGEF11 | 0,07590148 | 6,79819205 | 0,63408412 | 0,53588465 | 0,7478625 |
| 643677_at | CCDC168 | -0,03645731 | 3,03828946 | -0,63383207 | 0,53604459 | 0,74803151 |
| 79628_at | SH3TC2 | -0,19459624 | 5,30438496 | -0,63371167 | 0,53612099 | 0,74806078 |
| 64924_at | SLC30A5 | -0,17259384 | 7,89180762 | -0,63367663 | 0,53614323 | 0,74806078 |
| 10127_at | ZNF263 | 0,04938769 | 7,00222373 | 0,63352996 | 0,53623632 | 0,74812517 |
| 23384_at | SPECC1L | -0,03326141 | 6,63843799 | -0,63348154 | 0,53626706 | 0,74812517 |
| 101926962_a | LOC10192696 | -0,06940437 | 3,64114411 | -0,63336072 | 0,53634375 | 0,74815567 |
| 85315_at | PAQR8 | -0,08786721 | 6,0925449 | -0,63327908 | 0,53639558 | 0,74815567 |
| 9762_at | LZTS3 | 0,04794993 | 5,45175339 | 0,63323872 | 0,5364212 | 0,74815567 |
| 55506_at | H2AFY2 | 0,11018309 | 8,33823145 | 0,63320236 | 0,53644428 | 0,74815567 |
| 284408_at | ZNF790-AS1 | 0,23075125 | 4,31632278 | 0,63287863 | 0,53664982 | 0,74833093 |
| 137970_at | UNC5D | 0,070067 | 3,65457684 | 0,63284314 | 0,53667236 | 0,74833093 |
| 374887_at | YJEFN3 | -0,06919857 | 5,48107083 | -0,63274309 | 0,53673589 | 0,74833093 |
| 2995_at | GYPC | -0,07650389 | 4,99350966 | -0,63271937 | 0,53675096 | 0,74833093 |
| 8506_at | CNTNAP1 | -0,05836885 | 4,86902431 | -0,63267582 | 0,53677861 | 0,74833093 |
| 22899_at | ARHGEF15 | -0,0942751 | 5,17462234 | -0,63263738 | 0,53680303 | 0,74833093 |
| 57604_at | KIAA1456 | 0,05278057 | 3,98308168 | 0,63257473 | 0,53684281 | 0,74833223 |
| 388531_at | RGS9BP | -0,06694617 | 3,88303677 | -0,63239853 | 0,53695473 | 0,74836665 |
| 83606_at | GUCD1 | 0,09796228 | 9,1297199 | 0,63235924 | 0,53697969 | 0,74836665 |
| 126433_at | FBXO27 | -0,10673236 | 5,9901402 | -0,63231813 | 0,5370058 | 0,74836665 |
| 23266_at | ADGRL2 | 0,06464077 | 5,38697243 | 0,63224348 | 0,53705322 | 0,74836665 |
| 10139_at | ARFRP1 | -0,10091209 | 7,7231464 | -0,63223005 | 0,53706176 | 0,74836665 |
| 100507470_a | GRID1-AS1 | -0,07034351 | 3,37072342 | -0,63199131 | 0,53721343 | 0,74852385 |
| 6092_at | ROBO2 | 0,05276045 | 3,86316574 | 0,63188898 | 0,53727845 | 0,7485603 |
| 1047_at | CLGN | 0,45897606 | 4,38310563 | 0,63163497 | 0,53743986 | 0,74873104 |
| 93517_at | SDR42E1 | -0,19173379 | 6,39259581 | -0,63157239 | 0,53747963 | 0,74873229 |
| 4159_at | MC3R | -0,06093038 | 4,29532369 | -0,63132702 | 0,53763559 | 0,74887788 |
| 84795_at | PYROXD2 | 0,31154248 | 6,79980167 | 0,63128563 | 0,5376619 | 0,74887788 |
| 94160_at | ABCC12 | -0,05120568 | 3,57979064 | -0,63113997 | 0,53775448 | 0,74895269 |
| 54566_at | EPB41L4B | -0,04871888 | 5,25313608 | -0,63085628 | 0,53793485 | 0,74914972 |
| 116986_at | AGAP2 | -0,0524231 | 4,77794624 | -0,63043322 | 0,53820388 | 0,74947021 |
| 51027_at | BOLA1 | -0,03941007 | 6,97582747 | -0,63030116 | 0,53828788 | 0,74953299 |
| 131450_at | CD200R1 | -0,04622578 | 2,87177458 | -0,63005646 | 0,53844354 | 0,74969555 |
| 9481_at | SLC25A27 | 0,12816829 | 3,75425032 | 0,62984464 | 0,5385783 | 0,74982898 |
| 201780_at | SLC10A4 | 0,09888315 | 4,35037119 | 0,62968993 | 0,53867674 | 0,74986814 |
| 83706_at | FERMT3 | -0,10169733 | 5,69333927 | -0,62967808 | 0,53868428 | 0,74986814 |
| 167838_at | TXLNB | -0,04559665 | 3,08233772 | -0,62940004 | 0,53886122 | 0,75006024 |
| 64167_at | ERAP2 | 0,14352329 | 8,48505344 | 0,62932673 | 0,53890788 | 0,75007099 |
| 4034_at | LRCH4 | 0,03737912 | 7,19521295 | 0,62921265 | 0,53898049 | 0,75011785 |
| 6688_at | SPI1 | -0,05414043 | 3,84970414 | -0,62906705 | 0,53907317 | 0,75019264 |
| 25797_at | QPCT | 0,52261877 | 4,62628635 | 0,62889582 | 0,53918218 | 0,75029013 |
| 146691_at | TOM1L2 | 0,18657237 | 7,68256813 | 0,62863193 | 0,5393502 | 0,7504626 |

| | | | | | | |
|---|---|---|---|---|---|---|
| 6018_at | RLF | -0,08205261 | 8,17452853 | -0,62857879 | 0,53938404 | 0,7504626 |
| 2038_at | EPB42 | -0,0658133 | 4,47596076 | -0,62842224 | 0,53948373 | 0,7505471 |
| 1826_at | DSCAM | -0,04012195 | 3,69546941 | -0,62815799 | 0,53965204 | 0,75072703 |
| 387755_at | INSC | -0,07224705 | 3,83110222 | -0,62804742 | 0,53972246 | 0,75077079 |
| 79803_at | HPS6 | -0,07909517 | 7,62410505 | -0,62777207 | 0,53989788 | 0,75090925 |
| 23140_at | ZZEF1 | 0,07112973 | 6,92791789 | 0,62772672 | 0,53992677 | 0,75090925 |
| 5596_at | MAPK4 | -0,05125539 | 4,60449246 | -0,62770761 | 0,53993895 | 0,75090925 |
| 8715_at | NOL4 | -0,04055012 | 3,41667974 | -0,62758216 | 0,54001888 | 0,75096619 |
| 23093_at | TTLL5 | -0,08655609 | 8,31025389 | -0,6274663 | 0,54009271 | 0,75098637 |
| 140803_at | TRPM6 | -0,06701843 | 3,66259349 | -0,62743703 | 0,54011136 | 0,75098637 |
| 8838_at | WISP3 | 0,46970939 | 4,9443884 | 0,62730499 | 0,54019551 | 0,75101208 |
| 101927360_a | LINC01103 | -0,07298858 | 3,28247619 | -0,62725368 | 0,54022821 | 0,75101208 |
| 100147773_a | LOC10014777 | -0,05308522 | 3,97714866 | -0,62722449 | 0,54024682 | 0,75101208 |
| 641365_at | LINC00616 | -0,05171395 | 3,03816654 | -0,62712703 | 0,54030894 | 0,75101647 |
| 5414_at | SEPT4 | -0,04699215 | 4,43545266 | -0,6270972 | 0,54032795 | 0,75101647 |
| 6775_at | STAT4 | -0,1001374 | 6,15124175 | -0,62692678 | 0,54043658 | 0,75111326 |
| 8287_at | USP9Y | 0,52920614 | 4,62728198 | 0,62685931 | 0,54047959 | 0,75111885 |
| 256356_at | GK5 | 0,08830609 | 4,88880591 | 0,62679016 | 0,54052367 | 0,75112592 |
| 51585_at | PCF11 | -0,07569767 | 7,74963979 | -0,62661941 | 0,54063254 | 0,75122301 |
| 259308_at | FAM205A | -0,04381735 | 4,25849792 | -0,62650915 | 0,54070285 | 0,75123276 |
| 57547_at | ZNF624 | 0,32638525 | 5,4755314 | 0,62647946 | 0,54072178 | 0,75123276 |
| 388115_at | C15orf52 | 0,13747844 | 7,67353692 | 0,62642492 | 0,54075655 | 0,75123276 |
| 79933_at | SYNPO2L | 0,04601942 | 4,3992207 | 0,62614398 | 0,54093572 | 0,75137884 |
| 101928098_a | LOC10192809 | -0,06002875 | 3,5911531 | -0,62613772 | 0,54093971 | 0,75137884 |
| 51666_at | ASB4 | -0,04748461 | 3,67735139 | -0,62605381 | 0,54099323 | 0,751399 |
| 57834_at | CYP4F11 | -0,12041515 | 6,4350584 | -0,62562269 | 0,54126826 | 0,75172013 |
| 85464_at | SSH2 | 0,07906286 | 6,27416587 | 0,62556902 | 0,5413025 | 0,75172013 |
| 54088_at | LINC00113 | -0,0964818 | 3,76481899 | -0,62536542 | 0,54143241 | 0,75182406 |
| 7133_at | TNFRSF1B | 0,22507281 | 5,67372581 | 0,62532941 | 0,54145539 | 0,75182406 |
| 3208_at | HPCA | 0,07333639 | 4,72511384 | 0,62502217 | 0,54165147 | 0,75204211 |
| 54602_at | NDFIP2 | 0,20252961 | 9,50848817 | 0,62481075 | 0,54178642 | 0,75214981 |
| 23029_at | RBM34 | -0,0557554 | 6,95746016 | -0,62477831 | 0,54180713 | 0,75214981 |
| 26985_at | AP3M1 | 0,07257346 | 8,41473162 | 0,6246684 | 0,54187729 | 0,752193 |
| 11102_at | RPP14 | -0,11264973 | 7,57770483 | -0,62451329 | 0,54197632 | 0,75227626 |
| 266812_at | NAP1L5 | 0,31171499 | 5,5723907 | 0,6244311 | 0,5420288 | 0,75229489 |
| 706_at | TSPO | 0,04872339 | 10,5670106 | 0,62421387 | 0,54216751 | 0,75238016 |
| 285972_at | LINC00996 | -0,05791552 | 3,35580809 | -0,62415961 | 0,54220216 | 0,75238016 |
| 2020_at | EN2 | -0,0844931 | 5,40216774 | -0,6241514 | 0,54220741 | 0,75238016 |
| 79850_at | FAM57A | -0,184373 | 8,64594039 | -0,6239223 | 0,54235373 | 0,75247487 |
| 4037_at | LRP3 | -0,13616504 | 6,70358179 | -0,6239222 | 0,54235379 | 0,75247487 |
| 142686_at | ASB14 | -0,06941743 | 3,47519988 | -0,62372937 | 0,54247696 | 0,75256412 |
| 140578_at | CHODL | 0,07510834 | 4,16547238 | 0,62366199 | 0,54252001 | 0,75256412 |
| 118490_at | MSS51 | 0,12010509 | 4,45143498 | 0,62363801 | 0,54253532 | 0,75256412 |
| 2056_at | EPO | -0,05201865 | 4,87742545 | -0,62315607 | 0,54284325 | 0,75293704 |
| 247_at | ALOX15B | -0,05553621 | 4,85993799 | -0,62308646 | 0,54288774 | 0,75294453 |
| 145978_at | LINC00052 | 0,05461696 | 3,68271362 | 0,62291786 | 0,54299549 | 0,75303975 |
| 9274_at | BCL7C | -0,069922 | 9,00988027 | -0,62280099 | 0,54307019 | 0,7530673 |
| 10631_at | POSTN | -0,04257305 | 3,63075047 | -0,62271511 | 0,54312509 | 0,7530673 |
| 139735_at | ZFP92 | -0,06933947 | 5,23342861 | -0,6227033 | 0,54313264 | 0,7530673 |
| 51714_at | SELT | -0,05028723 | 11,0802908 | -0,62247015 | 0,54328168 | 0,7532047 |

| | | | | | | |
|---|---|---|---|---|---|---|
| 55066_at | PDPR | 0,16138373 | 7,31185317 | 0,62242596 | 0,54330993 | 0,7532047 |
| 94107_at | TMEM203 | -0,13402359 | 9,18333003 | -0,62233255 | 0,54336966 | 0,75323328 |
| 1581_at | CYP7A1 | -0,04726621 | 2,99586739 | -0,62225204 | 0,54342114 | 0,75325044 |
| 101929592_at | LOC10192959 | -0,0497459 | 2,92050948 | -0,62208074 | 0,54353067 | 0,75333331 |
| 50939_at | IMPG2 | 0,04079256 | 3,21946756 | 0,62199938 | 0,5435827 | 0,75333331 |
| 30850_at | CDR2L | 0,05581907 | 7,92278862 | 0,62197507 | 0,54359825 | 0,75333331 |
| 57713_at | SFMBT2 | -0,05402147 | 5,02052717 | -0,62167031 | 0,54379317 | 0,75354923 |
| 50855_at | PARD6A | 0,10376071 | 6,49489894 | 0,62159699 | 0,54384007 | 0,75356001 |
| 5328_at | PLAU | 0,29671525 | 7,93904816 | 0,62143337 | 0,54394474 | 0,75360709 |
| 1813_at | DRD2 | -0,06891334 | 5,34161639 | -0,62140051 | 0,54396576 | 0,75360709 |
| 7549_at | ZNF2 | 0,03698393 | 6,15950731 | 0,62136042 | 0,54399141 | 0,75360709 |
| 25973_at | PARS2 | 0,04396625 | 7,03301474 | 0,6212498 | 0,54406218 | 0,75365093 |
| 64122_at | FN3K | -0,06098135 | 4,35892949 | -0,62084206 | 0,54432311 | 0,75393215 |
| 55283_at | MCOLN3 | -0,1054817 | 3,65325489 | -0,62074654 | 0,54438424 | 0,75393215 |
| 54505_at | DHX29 | -0,2296514 | 8,54163955 | -0,62068326 | 0,54442474 | 0,75393215 |
| 101928778_at | LOC10192877 | -0,08482612 | 4,55882755 | -0,62065809 | 0,54444085 | 0,75393215 |
| 3400_at | ID4 | 0,47832598 | 7,22338451 | 0,62048462 | 0,54455189 | 0,75393215 |
| 79686_at | LINC00341 | 0,05426739 | 4,21810874 | 0,62044965 | 0,54457428 | 0,75393215 |
| 1756_at | DMD | -0,1042402 | 3,80030338 | -0,62042256 | 0,54459162 | 0,75393215 |
| 399693_at | CCDC187 | -0,06285542 | 5,67169744 | -0,6204181 | 0,54459447 | 0,75393215 |
| 9852_at | EPM2AIP1 | -0,27261347 | 5,56779522 | -0,62037889 | 0,54461958 | 0,75393215 |
| 1950_at | EGF | -0,12257576 | 4,00345015 | -0,62032108 | 0,54465658 | 0,75393215 |
| 388135_at | C15orf59 | -0,05803334 | 4,55792393 | -0,61998657 | 0,54487077 | 0,75411556 |
| 1311_at | COMP | -0,06667826 | 4,61880781 | -0,61993174 | 0,54490587 | 0,75411556 |
| 345222_at | MSANTD1 | -0,07348982 | 5,55885175 | -0,61993072 | 0,54490653 | 0,75411556 |
| 85461_at | TANC1 | 0,10213217 | 8,66130483 | 0,61977338 | 0,54500729 | 0,75420082 |
| 85403_at | EAF1 | -0,04681266 | 8,96073546 | -0,61956808 | 0,54513878 | 0,75429349 |
| 2664_at | GDI1 | -0,08955664 | 8,93096998 | -0,61951863 | 0,54517046 | 0,75429349 |
| 389084_at | C2orf82 | 0,12525024 | 6,15993497 | 0,61944252 | 0,54521921 | 0,75429349 |
| 79177_at | ZNF576 | -0,10092492 | 7,20545222 | -0,61942429 | 0,54523089 | 0,75429349 |
| 338667_at | LOC338667 | -0,078891 | 4,22790245 | -0,61929574 | 0,54531324 | 0,75435324 |
| 151534_at | LBX2-AS1 | -0,05962414 | 5,40037562 | -0,61914054 | 0,54541267 | 0,75440391 |
| 81697_at | OR2B2 | -0,06051669 | 3,04202787 | -0,61911631 | 0,54542819 | 0,75440391 |
| 152586_at | MGAT4D | -0,05023722 | 2,78684005 | -0,6189545 | 0,54553187 | 0,75449314 |
| 100506413_at | DOCK4-AS1 | -0,05640535 | 3,2831386 | -0,6187416 | 0,5456683 | 0,75460334 |
| 10417_at | SPON2 | -0,0752097 | 5,57054738 | -0,6187079 | 0,5456899 | 0,75460334 |
| 7352_at | UCP3 | 0,04624821 | 4,98020693 | 0,6186008 | 0,54575854 | 0,75464409 |
| 285381_at | DPH3 | -0,069077 | 8,03714002 | -0,61836271 | 0,54591115 | 0,75480093 |
| 9616_at | RNF7 | -0,07077174 | 8,71366334 | -0,61825413 | 0,54598075 | 0,75484298 |
| 23041_at | MON2 | 0,1060415 | 6,92986449 | 0,61813725 | 0,54605568 | 0,75489239 |
| 192683_at | SCAMP5 | -0,06047402 | 5,4885772 | -0,61798291 | 0,54615464 | 0,75497502 |
| 123624_at | AGBL1 | -0,05436393 | 3,56568822 | -0,617906 | 0,54620395 | 0,754989 |
| 150160_at | CCT8L2 | -0,05835079 | 5,47893241 | -0,617658 | 0,54636299 | 0,75515464 |
| 84418_at | CYSTM1 | 0,08544653 | 9,38659796 | 0,6172903 | 0,54659883 | 0,75537458 |
| 8905_at | AP1S2 | -0,11284909 | 5,89042099 | -0,61728762 | 0,54660054 | 0,75537458 |
| 286256_at | LCN12 | 0,0567882 | 5,40137298 | 0,61722243 | 0,54664236 | 0,75537818 |
| 100506606_at | LOC10050660 | 0,05237465 | 4,42463047 | 0,61704825 | 0,5467541 | 0,75547839 |
| 57501_at | KIAA1257 | 0,04586092 | 4,22060522 | 0,61654253 | 0,5470786 | 0,75585111 |
| 7871_at | SLMAP | 0,09980902 | 6,97874596 | 0,61635584 | 0,54719843 | 0,75585111 |
| 27136_at | MORC1 | 0,03647586 | 2,87774098 | 0,61634593 | 0,54720478 | 0,75585111 |

| | | | | | | |
|---|---|---|---|---|---|---|
| 100862662_a | ALDH1L1-AS2 | -0,06483922 | 5,66476002 | -0,61633969 | 0,54720879 | 0,75585111 |
| 23167_at | EFR3A | 0,23163216 | 9,51592972 | 0,61626246 | 0,54725836 | 0,75585111 |
| 2688_at | GH1 | -0,05360832 | 5,17825745 | -0,61620242 | 0,5472969 | 0,75585111 |
| 339345_at | NANOS2 | -0,06109129 | 4,32769863 | -0,61619991 | 0,54729851 | 0,75585111 |
| 126248_at | WDR88 | -0,05334025 | 3,53113408 | -0,61601304 | 0,54741847 | 0,75594601 |
| 348093_at | RBPMS2 | -0,05528745 | 4,9636736 | -0,61586646 | 0,54751258 | 0,75594601 |
| 29965_at | CDIP1 | 0,1525486 | 6,01825002 | 0,61582551 | 0,54753888 | 0,75594601 |
| 1209_at | CLPTM1 | 0,05856307 | 9,43416866 | 0,61580564 | 0,54755163 | 0,75594601 |
| 2153_at | F5 | 0,29617628 | 4,41969291 | 0,61578724 | 0,54756344 | 0,75594601 |
| 246705_at | LINC00314 | -0,04983147 | 2,85417165 | -0,61558798 | 0,54769139 | 0,75601849 |
| 79908_at | BTNL8 | -0,0488051 | 3,62249069 | -0,6155262 | 0,54773107 | 0,75601849 |
| 136263_at | SSMEM1 | 0,0631796 | 3,29602673 | 0,61552212 | 0,54773369 | 0,75601849 |
| 101927972_a | LOC10192797 | -0,0648834 | 3,55085598 | -0,6153917 | 0,54781745 | 0,75603399 |
| 101927152_a | LINC01212 | 0,05029929 | 3,23147541 | 0,61538241 | 0,54782342 | 0,75603399 |
| 397_at | ARHGDIB | -0,19937118 | 9,21884901 | -0,61530082 | 0,54787582 | 0,75605214 |
| 139341_at | FUNDC1 | 0,15124572 | 7,89769346 | 0,61517605 | 0,54795596 | 0,75610857 |
| 80741_at | LY6G5C | 0,17090783 | 5,77051115 | 0,61507566 | 0,54802044 | 0,75614338 |
| 221120_at | ALKBH3 | -0,10858208 | 8,06622361 | -0,61484543 | 0,54816835 | 0,75626913 |
| 51123_at | ZNF706 | -0,10518857 | 7,11806372 | -0,61481158 | 0,5481901 | 0,75626913 |
| 2875_at | GPT | -0,06203749 | 6,30206103 | -0,61464 | 0,54830034 | 0,75636517 |
| 9569_at | GTF2IRD1 | 0,05806713 | 8,45230032 | 0,61457854 | 0,54833983 | 0,75636517 |
| 100652999_a | LOC10065299 | -0,089166 | 5,41521211 | -0,6145199 | 0,54837751 | 0,75636517 |
| 2264_at | FGFR4 | -0,04997435 | 5,43079528 | -0,61381872 | 0,5488282 | 0,75693259 |
| 2702_at | GJA5 | 0,06736333 | 3,6959027 | 0,61361964 | 0,54895619 | 0,75705492 |
| 23_at | ABCF1 | 0,0413439 | 9,51049325 | 0,61355686 | 0,54899656 | 0,75705639 |
| 162517_at | FBXO39 | -0,06028279 | 3,94481806 | -0,61347696 | 0,54904793 | 0,75707304 |
| 9077_at | DIRAS3 | -0,06313374 | 3,66782707 | -0,61317994 | 0,54923895 | 0,75714368 |
| 390063_at | OR51I1 | -0,06429007 | 4,03980769 | -0,61317612 | 0,5492414 | 0,75714368 |
| 285696_at | LOC285696 | 0,05489337 | 4,27914155 | 0,61312883 | 0,54927182 | 0,75714368 |
| 644809_at | C15orf56 | -0,0818257 | 4,7283533 | -0,61305877 | 0,54931688 | 0,75714368 |
| 100507062_a | PSMD6-AS2 | 0,0547234 | 4,4623732 | 0,61305798 | 0,54931739 | 0,75714368 |
| 64111_at | NPVF | -0,05573054 | 3,08213792 | -0,6130306 | 0,549335 | 0,75714368 |
| 84707_at | BEX2 | 0,2655483 | 6,18126728 | 0,61283479 | 0,54946095 | 0,7572631 |
| 978_at | CDA | 0,19466472 | 6,87134089 | 0,61263629 | 0,54958866 | 0,75738491 |
| 91522_at | COL23A1 | -0,08702456 | 5,75824909 | -0,61246199 | 0,5497008 | 0,75748527 |
| 2039_at | DMTN | 0,05677287 | 6,2347938 | 0,61215148 | 0,54990062 | 0,75761368 |
| 135228_at | CD109 | 0,08523956 | 6,90769673 | 0,61206083 | 0,54995896 | 0,75761368 |
| 9463_at | PICK1 | -0,05429416 | 7,08049936 | -0,6120529 | 0,54996407 | 0,75761368 |
| 10093_at | ARPC4 | -0,03876685 | 9,86587377 | -0,61200714 | 0,54999352 | 0,75761368 |
| 91181_at | NUP210L | -0,05058732 | 3,12035684 | -0,61198039 | 0,55001074 | 0,75761368 |
| 646522_at | LOC646522 | -0,068352 | 3,77047888 | -0,61195051 | 0,55002997 | 0,75761368 |
| 9543_at | IGDCC3 | 0,13399954 | 5,20363944 | 0,61161055 | 0,55024881 | 0,75782264 |
| 1944_at | EFNA3 | -0,07137321 | 6,56509918 | -0,61159262 | 0,55026036 | 0,75782264 |
| 728175_at | LOC728175 | -0,06371431 | 4,18679122 | -0,61128014 | 0,55046155 | 0,75804553 |
| 25936_at | NSL1 | -0,09003602 | 6,11657416 | -0,61111326 | 0,55056902 | 0,75813932 |
| 22943_at | DKK1 | 0,13927084 | 7,87237936 | 0,61098918 | 0,55064893 | 0,75819516 |
| 8974_at | P4HA2 | 0,15376117 | 9,22049765 | 0,61040084 | 0,55102792 | 0,75863856 |
| 64753_at | CCDC136 | 0,06658982 | 4,3883376 | 0,61036701 | 0,55104972 | 0,75863856 |
| 222553_at | SLC35F1 | -0,07022703 | 5,37484759 | -0,610208 | 0,55115218 | 0,75872539 |
| 4284_at | MIP | -0,05157015 | 3,80234116 | -0,61002063 | 0,55127292 | 0,75878817 |

| | | | | | | |
|---|---|---|---|---|---|---|
| 123591_at | C15orf27 | -0,0524497 | 4,23665043 | -0,60996086 | 0,55131144 | 0,75878817 |
| 23181_at | DIP2A | -0,09083342 | 6,23393305 | -0,60995386 | 0,55131595 | 0,75878817 |
| 84924_at | ZNF566 | 0,05473672 | 4,86214063 | 0,60987029 | 0,55136981 | 0,75880808 |
| 55370_at | PPP4R1L | -0,03332253 | 4,49042161 | -0,609776 | 0,55143058 | 0,7588375 |
| 43_at | ACHE | -0,05038745 | 4,6685066 | -0,60942778 | 0,55165506 | 0,75905501 |
| 710_at | SERPING1 | -0,05126089 | 4,00781562 | -0,60940855 | 0,55166745 | 0,75905501 |
| 3455_at | IFNAR2 | -0,06713706 | 7,6487003 | -0,60933192 | 0,55171686 | 0,75906877 |
| 57007_at | ACKR3 | 0,18978895 | 9,63383188 | 0,60920483 | 0,5517988 | 0,7591033 |
| 11155_at | LDB3 | 0,12011842 | 5,1311319 | 0,60913869 | 0,55184144 | 0,7591033 |
| 411_at | ARSB | -0,06558714 | 5,55399983 | -0,60907604 | 0,55188185 | 0,7591033 |
| 6093_at | ROCK1 | -0,05826305 | 6,83202097 | -0,60904853 | 0,55189958 | 0,7591033 |
| 4360_at | MRC1 | 0,05544541 | 3,14163486 | 0,6089191 | 0,55198305 | 0,75913142 |
| 151613_at | TTC14 | 0,14578628 | 6,84586172 | 0,60889461 | 0,55199885 | 0,75913142 |
| 79647_at | AKIRIN1 | -0,06021617 | 8,9302185 | -0,60873793 | 0,5520999 | 0,75921619 |
| 89857_at | KLHL6 | -0,05087277 | 3,72251582 | -0,6085731 | 0,55220621 | 0,75930818 |
| 120892_at | LRRK2 | -0,03560153 | 3,20007372 | -0,60819399 | 0,55245078 | 0,75953813 |
| 29844_at | TFPT | 0,16040835 | 7,90325978 | 0,60815652 | 0,55247496 | 0,75953813 |
| 26226_at | FBXW4P1 | 0,05946705 | 6,01498868 | 0,60812535 | 0,55249507 | 0,75953813 |
| 283214_at | LOC283214 | -0,03668759 | 3,46811765 | -0,60806016 | 0,55253713 | 0,75953813 |
| 8161_at | COIL | 0,08563388 | 8,12240617 | 0,60800831 | 0,55257059 | 0,75953813 |
| 5780_at | PTPN9 | -0,02776424 | 7,69376987 | -0,60787524 | 0,55265646 | 0,75960196 |
| 153918_at | ZC2HC1B | 0,05640205 | 3,66077872 | 0,60753137 | 0,5528784 | 0,75985278 |
| 643160_at | CYMP | -0,0542768 | 4,50083365 | -0,60711266 | 0,5531487 | 0,76017003 |
| 643749_at | TRAF3IP2-AS1 | -0,0451972 | 4,60039453 | -0,60685514 | 0,55331498 | 0,76030577 |
| 57722_at | IGDCC4 | -0,05978021 | 4,40964418 | -0,60683743 | 0,55332641 | 0,76030577 |
| 285796_at | PACRG-AS1 | -0,06357847 | 4,06036177 | -0,60661978 | 0,55346697 | 0,76044467 |
| 2591_at | GALNT3 | 0,15333816 | 10,0099228 | 0,60649312 | 0,55354877 | 0,76050282 |
| 8515_at | ITGA10 | -0,1070094 | 4,97146184 | -0,60642803 | 0,55359082 | 0,76050634 |
| 6706_at | SPRR2G | -0,08866775 | 5,79766467 | -0,60635639 | 0,55363709 | 0,76051568 |
| 399806_at | LBX1-AS1 | -0,04607479 | 5,27538291 | -0,60619143 | 0,55374365 | 0,76060781 |
| 64779_at | MTHFSD | 0,03063287 | 5,46598114 | 0,60612611 | 0,55378585 | 0,76061154 |
| 6440_at | SFTPC | -0,0624017 | 5,81620169 | -0,60597106 | 0,55388603 | 0,76069489 |
| 2235_at | FECH | -0,09231142 | 6,95044989 | -0,60572006 | 0,55404821 | 0,76086339 |
| 1521_at | CTSW | -0,05614003 | 4,98317893 | -0,60550644 | 0,55418627 | 0,76099872 |
| 54831_at | BEST2 | -0,05955106 | 4,50992496 | -0,6052322 | 0,55436352 | 0,76118786 |
| 127833_at | SYT2 | -0,06136441 | 5,88264181 | -0,60498073 | 0,55452608 | 0,7613568 |
| 10495_at | ENOX2 | -0,04704033 | 6,96186464 | -0,60489784 | 0,55457967 | 0,76137611 |
| 57568_at | SIPA1L2 | 0,25501709 | 6,86253169 | 0,60452614 | 0,55482003 | 0,76161296 |
| 80154_at | GOLGA2P10 | 0,1769433 | 7,49072434 | 0,60450876 | 0,55483126 | 0,76161296 |
| 92255_at | LMBRD2 | -0,07459875 | 5,73403524 | -0,60441735 | 0,55489038 | 0,76163983 |
| 81851_at | KRTAP1-1 | -0,04132044 | 4,83018564 | -0,60426504 | 0,55498889 | 0,76167603 |
| 1739_at | DLG1 | 0,07592257 | 8,36920484 | 0,60423708 | 0,55500697 | 0,76167603 |
| 140886_at | PABPC5 | 0,04560031 | 2,97987772 | 0,60419318 | 0,55503537 | 0,76167603 |
| 64129_at | TINAGL1 | -0,10109562 | 9,59945103 | -0,60412625 | 0,55507867 | 0,76168118 |
| 51409_at | HEMK1 | 0,05934593 | 6,51026129 | 0,60400882 | 0,55515463 | 0,76173115 |
| 164045_at | HFM1 | -0,04380017 | 2,79978862 | -0,60359726 | 0,5554209 | 0,7619768 |
| 55647_at | RAB20 | 0,03855444 | 8,85971076 | 0,60352144 | 0,55546997 | 0,7619768 |
| 7123_at | CLEC3B | -0,07610575 | 5,16856679 | -0,60351453 | 0,55547444 | 0,7619768 |
| 59339_at | PLEKHA2 | 0,10209712 | 7,97972239 | 0,60348756 | 0,55549189 | 0,7619768 |
| 340371_at | NRBP2 | 0,3630708 | 6,88885417 | 0,60339918 | 0,55554909 | 0,762001 |

| | | | | | | |
|---|---|---|---|---|---|---|
| 374739_at | TEPP | -0,07048847 | 5,11275445 | -0,60306573 | 0,5557649 | 0,76224274 |
| 202559_at | KHDRBS2 | -0,08641632 | 3,95579463 | -0,60290041 | 0,55587192 | 0,76233524 |
| 7025_at | NR2F1 | -0,30900736 | 6,30556636 | -0,60270002 | 0,55600165 | 0,76238869 |
| 102724434_a | LOC10272443 | -0,06986118 | 4,24534091 | -0,60267273 | 0,55601932 | 0,76238869 |
| 11011_at | TLK2 | -0,03794137 | 6,65554442 | -0,6026568 | 0,55602963 | 0,76238869 |
| 5189_at | PEX1 | 0,12000993 | 7,03670982 | 0,60245348 | 0,55616128 | 0,76251493 |
| 79413_at | ZBED2 | 0,52500523 | 6,49513745 | 0,60230437 | 0,55625784 | 0,76254662 |
| 55230_at | USP40 | 0,25499033 | 8,3292421 | 0,60223645 | 0,55630183 | 0,76254662 |
| 6484_at | ST3GAL4 | 0,06351055 | 7,26185241 | 0,60223441 | 0,55630315 | 0,76254662 |
| 147323_at | STARD6 | 0,04440409 | 3,22169981 | 0,60213507 | 0,55636748 | 0,76257561 |
| 256987_at | SERINC5 | 0,15016304 | 10,1479333 | 0,6020795 | 0,55640348 | 0,76257561 |
| 54954_at | FAM120C | -0,04616199 | 6,31619337 | -0,60201554 | 0,5564449 | 0,76257813 |
| 8085_at | KMT2D | 0,04295836 | 7,27338672 | 0,60194914 | 0,55648791 | 0,76258281 |
| 80350_at | LPAL2 | -0,06563455 | 3,82733073 | -0,60159758 | 0,55671566 | 0,76278289 |
| 6596_at | HLTF | -0,11249622 | 8,88134531 | -0,60149149 | 0,5567844 | 0,76278289 |
| 6103_at | RPGR | -0,10840416 | 4,92556984 | -0,60145955 | 0,55680509 | 0,76278289 |
| 55831_at | EMC3 | -0,0724692 | 8,41834 | -0,60144659 | 0,55681349 | 0,76278289 |
| 25814_at | ATXN10 | 0,03419755 | 9,26182383 | 0,60141816 | 0,55683191 | 0,76278289 |
| 319_at | APOF | 0,03788014 | 3,38263118 | 0,60120495 | 0,55697007 | 0,7629179 |
| 6496_at | SIX3 | -0,04421153 | 4,09525641 | -0,60096416 | 0,55712613 | 0,76304121 |
| 10123_at | ARL4C | -0,14968233 | 7,68172223 | -0,60094382 | 0,55713931 | 0,76304121 |
| 101929684_a | LINC01520 | -0,07466128 | 4,11835407 | -0,60087736 | 0,55718239 | 0,76304595 |
| 22889_at | KIAA0907 | -0,13352019 | 9,26331796 | -0,60080727 | 0,55722782 | 0,76305392 |
| 81556_at | VWA9 | 0,04053709 | 8,88533859 | 0,6002941 | 0,55756051 | 0,76342171 |
| 2124_at | EVI2B | -0,05845074 | 3,03980783 | -0,60027074 | 0,55757566 | 0,76342171 |
| 348761_at | SPATA3-AS1 | 0,06376788 | 4,18377023 | 0,60017188 | 0,55763977 | 0,76345521 |
| 7690_at | ZNF131 | -0,09958567 | 7,61537154 | -0,59978248 | 0,55789231 | 0,76374669 |
| 28683_at | TRAV8-3 | 0,09780695 | 5,12112752 | 0,59925852 | 0,55823222 | 0,7639699 |
| 2290_at | FOXG1 | -0,05154695 | 5,02239567 | -0,59924477 | 0,55824114 | 0,7639699 |
| 100506696_a | PCAT6 | 0,14728537 | 6,46975577 | 0,59918927 | 0,55827715 | 0,7639699 |
| 4222_at | MEOX1 | -0,05367712 | 4,00366535 | -0,59917807 | 0,55828442 | 0,7639699 |
| 2538_at | G6PC | -0,04070859 | 3,53682977 | -0,59913225 | 0,55831415 | 0,7639699 |
| 64002_at | PCGEM1 | 0,05609862 | 3,26840145 | 0,59912603 | 0,55831819 | 0,7639699 |
| 100129482_a | ZNF37BP | -0,08347529 | 4,71320714 | -0,59905882 | 0,5583618 | 0,7639699 |
| 149775_at | GNAS-AS1 | -0,19703295 | 4,10705532 | -0,59904211 | 0,55837264 | 0,7639699 |
| 84791_at | LINC00467 | -0,14309541 | 6,96765233 | -0,5987893 | 0,55853671 | 0,76409906 |
| 5893_at | RAD52 | -0,04812991 | 5,88081467 | -0,59877442 | 0,55854637 | 0,76409906 |
| 64061_at | TSPYL2 | 0,10055293 | 7,53871624 | 0,59868826 | 0,55860229 | 0,76412129 |
| 27177_at | IL36B | -0,04379121 | 3,30480731 | -0,59855141 | 0,55869112 | 0,76418853 |
| 284119_at | PTRF | 0,11431322 | 6,80789734 | 0,5982064 | 0,5589151 | 0,76441233 |
| 54936_at | ADPRHL2 | -0,05035133 | 8,91121057 | -0,59817713 | 0,5589341 | 0,76441233 |
| 53343_at | NUDT9 | -0,09976011 | 7,88603843 | -0,59801469 | 0,55903958 | 0,76446811 |
| 79643_at | CHMP6 | 0,05181499 | 7,6914513 | 0,59785884 | 0,55914078 | 0,76446811 |
| 84522_at | JAGN1 | -0,04049561 | 9,38796561 | -0,59781979 | 0,55916614 | 0,76446811 |
| 266553_at | OFCC1 | -0,04712535 | 3,84441856 | -0,59779222 | 0,55918405 | 0,76446811 |
| 203076_at | C8orf74 | -0,04358643 | 4,70212917 | -0,59778742 | 0,55918717 | 0,76446811 |
| 146433_at | IL34 | -0,04685934 | 6,07406317 | -0,59761179 | 0,55930123 | 0,76446811 |
| 81623_at | DEFB126 | 0,06501896 | 3,38553371 | 0,59748984 | 0,55938044 | 0,76446811 |
| 1233_at | CCR4 | -0,07586428 | 4,95641934 | -0,5973085 | 0,55949824 | 0,76446811 |
| 219995_at | MS4A15 | -0,06028959 | 4,35041098 | -0,59729886 | 0,5595045 | 0,76446811 |

| | | | | | | |
|---|---|---|---|---|---|---|
| 9425_at | CDYL | 0,04077092 | 8,27644039 | 0,59727938 | 0,55951716 | 0,76446811 |
| 4773_at | NFATC2 | 0,10042912 | 5,19984014 | 0,5972629 | 0,55952786 | 0,76446811 |
| 223082_at | ZNRF2 | 0,13653622 | 7,81586423 | 0,59725501 | 0,55953299 | 0,76446811 |
| 541565_at | C8orf58 | 0,09758639 | 6,8222324 | 0,59720635 | 0,5595646 | 0,76446811 |
| 152815_at | THAP6 | 0,13241123 | 5,82478772 | 0,59719268 | 0,55957348 | 0,76446811 |
| 403312_at | LOC403312 | -0,0476117 | 3,49517731 | -0,59711235 | 0,55962567 | 0,76446811 |
| 6576_at | SLC25A1 | -0,16499069 | 10,0776855 | -0,59705123 | 0,55966538 | 0,76446811 |
| 5158_at | PDE6B | 0,08308539 | 5,26652207 | 0,59703575 | 0,55967544 | 0,76446811 |
| 2911_at | GRM1 | -0,03858316 | 3,96183872 | -0,59696729 | 0,55971992 | 0,76446811 |
| 496_at | ATP4B | -0,05669434 | 4,46425658 | -0,59695344 | 0,55972893 | 0,76446811 |
| 146310_at | RNF151 | -0,05042942 | 5,07823833 | -0,59679261 | 0,55983343 | 0,76455664 |
| 100528062_a | ARMCX5-GPR. | 0,13038268 | 6,21512425 | 0,59669479 | 0,559897 | 0,76458925 |
| 23526_at | HMHA1 | 0,0566325 | 5,59992709 | 0,5965348 | 0,56000098 | 0,76463535 |
| 151649_at | PP2D1 | -0,04446987 | 2,74374228 | -0,59652069 | 0,56001015 | 0,76463535 |
| 222865_at | TMEM130 | 0,05559044 | 4,15754687 | 0,59635199 | 0,5601198 | 0,76469488 |
| 5613_at | PRKX | -0,08249233 | 6,97093978 | -0,59627313 | 0,56017106 | 0,76469488 |
| 100874110_a | GLYCTK-AS1 | -0,06876327 | 4,64877814 | -0,5962704 | 0,56017284 | 0,76469488 |
| 151477_at | LINC00471 | -0,05175315 | 3,61725144 | -0,59609156 | 0,5602891 | 0,76479938 |
| 100506275_a | KRBOX1-AS1 | 0,06436904 | 4,69745167 | 0,59582179 | 0,5604645 | 0,76496299 |
| 83401_at | ELOVL3 | -0,05163424 | 3,99221845 | -0,59578506 | 0,56048838 | 0,76496299 |
| 387763_at | C11orf96 | 0,05400326 | 4,48526709 | 0,59566746 | 0,56056485 | 0,76501316 |
| 8677_at | STX10 | -0,03544964 | 8,4024419 | -0,5955142 | 0,56066452 | 0,76507023 |
| 146059_at | CDAN1 | -0,05078144 | 7,53199022 | -0,59540815 | 0,56073349 | 0,76507023 |
| 285692_at | LOC285692 | -0,04472991 | 3,08904318 | -0,59537438 | 0,56075546 | 0,76507023 |
| 11141_at | IL1RAPL1 | -0,06644972 | 3,86727115 | -0,59535887 | 0,56076554 | 0,76507023 |
| 55005_at | RMND1 | -0,06231233 | 6,43931641 | -0,59518673 | 0,56087752 | 0,76516881 |
| 283731_at | LOC283731 | 0,04035744 | 3,42811542 | 0,59504623 | 0,56096891 | 0,7652393 |
| 79796_at | ALG9 | 0,06000897 | 7,93901671 | 0,59452966 | 0,56130501 | 0,76564357 |
| 79577_at | CDC73 | -0,06854657 | 8,02354961 | -0,59446262 | 0,56134864 | 0,76564887 |
| 346389_at | MACC1 | -0,31101453 | 6,02734936 | -0,59429373 | 0,56145856 | 0,76574457 |
| 134466_at | ZNF300P1 | -0,05048449 | 3,08447922 | -0,59419165 | 0,561525 | 0,76578096 |
| 28526_at | TRDC | -0,04773408 | 3,40278674 | -0,59407556 | 0,56160056 | 0,76580217 |
| 4985_at | OPRD1 | -0,04906785 | 5,70876398 | -0,59404561 | 0,56162006 | 0,76580217 |
| 64579_at | NDST4 | -0,06584842 | 3,96867585 | -0,59377954 | 0,56179327 | 0,76598413 |
| 9597_at | SMAD5-AS1 | -0,05912046 | 4,05498207 | -0,59345499 | 0,56200459 | 0,76621803 |
| 55534_at | MAML3 | 0,11313024 | 6,56151877 | 0,59337502 | 0,56205667 | 0,76623479 |
| 100141515_a | C17orf99 | -0,11999198 | 5,34230414 | -0,59296654 | 0,56232271 | 0,76654323 |
| 156_at | ADRBK1 | -0,07081272 | 7,63732864 | -0,5928564 | 0,56239446 | 0,76658678 |
| 373861_at | HILS1 | 0,0594968 | 5,02613997 | 0,59248649 | 0,56263545 | 0,76681642 |
| 23243_at | ANKRD28 | -0,08868488 | 7,59022512 | -0,59247561 | 0,56264254 | 0,76681642 |
| 84687_at | PPP1R9B | 0,0370409 | 5,95782575 | 0,59219329 | 0,56282651 | 0,76701289 |
| 5896_at | RAG1 | 0,04818834 | 3,38785205 | 0,59210365 | 0,56288494 | 0,76703824 |
| 2555_at | GABRA2 | 0,03204902 | 2,90398734 | 0,59193603 | 0,56299418 | 0,76713284 |
| 81794_at | ADAMTS10 | -0,05601972 | 5,41325651 | -0,59149065 | 0,56328452 | 0,76747416 |
| 4752_at | NEK3 | 0,11263735 | 4,77441762 | 0,59128122 | 0,56342107 | 0,76748786 |
| 55540_at | IL17RB | -0,09418365 | 5,0803141 | -0,59122325 | 0,56345887 | 0,76748786 |
| 5306_at | PITPNA | 0,05953706 | 8,43664574 | 0,59119522 | 0,56347715 | 0,76748786 |
| 6817_at | SULT1A1 | 0,04737093 | 6,57934889 | 0,59115768 | 0,56350163 | 0,76748786 |
| 58512_at | DLGAP3 | -0,05690713 | 5,96017509 | -0,59109293 | 0,56354385 | 0,76748786 |
| 90592_at | ZNF700 | 0,08390762 | 6,0591899 | 0,59107524 | 0,56355539 | 0,76748786 |

| | | | | | | |
|---|---|---|---|---|---|---|
| 154442_at | BVES-AS1 | 0,03662033 | 3,22441664 | 0,59100902 | 0,56359858 | 0,76748786 |
| 6320_at | CLEC11A | -0,06145233 | 6,01128691 | -0,59097397 | 0,56362143 | 0,76748786 |
| 5355_at | PLP2 | -0,20665457 | 10,4253218 | -0,59092533 | 0,56365316 | 0,76748786 |
| 1559_at | CYP2C9 | -0,04099952 | 3,94531052 | -0,59083583 | 0,56371154 | 0,7675131 |
| 8924_at | HERC2 | -0,04867029 | 7,34864083 | -0,59069636 | 0,5638025 | 0,76751437 |
| 84074_at | QRICH2 | -0,07986875 | 4,81085656 | -0,59069585 | 0,56380284 | 0,76751437 |
| 6007_at | RHD | -0,06326806 | 5,38047309 | -0,59064451 | 0,56383633 | 0,76751437 |
| 107_at | ADCY1 | 0,03650069 | 6,28453904 | 0,59059006 | 0,56387184 | 0,76751437 |
| 285556_at | LOC285556 | 0,05032793 | 3,87268953 | 0,59033706 | 0,5640369 | 0,76768314 |
| 283687_at | ST20-AS1 | 0,18902671 | 6,1010146 | 0,59027784 | 0,56407554 | 0,76768314 |
| 116511_at | MAS1L | -0,08945844 | 3,74666866 | -0,58995633 | 0,56428533 | 0,76787295 |
| 1473_at | CST5 | -0,09163105 | 4,78038819 | -0,58982575 | 0,56437055 | 0,76787295 |
| 9290_at | GPR55 | -0,04376015 | 4,15653035 | -0,58981712 | 0,56437619 | 0,76787295 |
| 7020_at | TFAP2A | 0,13741495 | 7,4661174 | 0,58967908 | 0,56446628 | 0,76787295 |
| 643923_at | LOC643923 | -0,06789961 | 4,27551049 | -0,5896648 | 0,5644756 | 0,76787295 |
| 23373_at | CRTC1 | -0,06067516 | 6,9467653 | -0,58959426 | 0,56452164 | 0,76787295 |
| 4707_at | NDUFB1 | 0,03754375 | 9,92995914 | 0,58953741 | 0,56455875 | 0,76787295 |
| 53829_at | P2RY13 | -0,03911442 | 3,28902069 | -0,58953213 | 0,5645622 | 0,76787295 |
| 101926951_a | LOC10192695 | 0,08805954 | 3,31354301 | 0,58942452 | 0,56463245 | 0,76787295 |
| 2099_at | ESR1 | 0,04193248 | 5,35426317 | 0,58942153 | 0,5646344 | 0,76787295 |
| 285352_at | KIF9-AS1 | -0,05434324 | 5,17327581 | -0,58934257 | 0,56468594 | 0,76787295 |
| 6801_at | STRN | 0,03899426 | 7,05853558 | 0,58932964 | 0,56469439 | 0,76787295 |
| 387856_at | CCDC184 | -0,06254566 | 4,86665873 | -0,5892313 | 0,56475859 | 0,76787295 |
| 196477_at | CCER1 | 0,05334688 | 3,42073482 | 0,58919691 | 0,56478104 | 0,76787295 |
| 65988_at | ZNF747 | -0,08306264 | 7,0884235 | -0,58911714 | 0,56483312 | 0,76787295 |
| 5597_at | MAPK6 | 0,08151559 | 10,2494829 | 0,58908699 | 0,56485281 | 0,76787295 |
| 134121_at | C5orf49 | -0,06129698 | 3,79982522 | -0,58882856 | 0,56502156 | 0,76801736 |
| 4194_at | MDM4 | -0,08549605 | 6,15672263 | -0,58880168 | 0,56503912 | 0,76801736 |
| 728448_at | PPIEL | -0,09221422 | 4,48222541 | -0,58872228 | 0,56509097 | 0,76801736 |
| 338324_at | S100A7A | -0,06788463 | 3,64956922 | -0,58864952 | 0,56513849 | 0,76801736 |
| 9167_at | COX7A2L | 0,04354552 | 9,75018446 | 0,58861906 | 0,56515839 | 0,76801736 |
| 101927003_a | PKIA-AS1 | 0,03762557 | 3,00966098 | 0,588551 | 0,56520284 | 0,76802358 |
| 1137_at | CHRNA4 | -0,05859602 | 4,64696268 | -0,58840369 | 0,56529906 | 0,76810015 |
| 283404_at | LINC00592 | 0,05988828 | 4,58013744 | 0,58822826 | 0,56541366 | 0,76810364 |
| 389058_at | SP5 | -0,07200699 | 5,06385851 | -0,58821817 | 0,56542025 | 0,76810364 |
| 51207_at | DUSP13 | -0,06244204 | 5,14050804 | -0,58821664 | 0,56542125 | 0,76810364 |
| 7704_at | ZBTB16 | -0,05369466 | 4,4715887 | -0,58770584 | 0,565755 | 0,76850283 |
| 27148_at | STK36 | 0,12734162 | 7,29981199 | 0,58757399 | 0,56584118 | 0,76856569 |
| 9584_at | RBM39 | 0,07682726 | 6,97081077 | 0,58744759 | 0,56592379 | 0,76858313 |
| 8566_at | PDXK | 0,12685282 | 9,88151291 | 0,58743224 | 0,56593382 | 0,76858313 |
| 157574_at | FBXO16 | 0,13244575 | 5,81969345 | 0,58734544 | 0,56599055 | 0,768606 |
| 388743_at | CAPN8 | -0,41024271 | 5,60674078 | -0,58721134 | 0,56607821 | 0,76867084 |
| 140801_at | RPL10L | -0,04066813 | 5,71266583 | -0,58713276 | 0,56612958 | 0,7686864 |
| 27092_at | CACNG4 | 0,10036707 | 7,44913046 | 0,5870554 | 0,56618015 | 0,76868694 |
| 286023_at | FLJ40288 | -0,06549966 | 5,30085347 | -0,58701008 | 0,56620978 | 0,76868694 |
| 7539_at | ZFP37 | -0,06225816 | 3,4803645 | -0,58672519 | 0,56639605 | 0,76888563 |
| 26091_at | HERC4 | 0,0710195 | 7,46974577 | 0,58626099 | 0,56669963 | 0,76924354 |
| 6777_at | STAT5B | 0,07970329 | 7,11283581 | 0,58615773 | 0,56676717 | 0,76928101 |
| 126070_at | ZNF440 | -0,10253165 | 6,92336741 | -0,58588035 | 0,56694862 | 0,76942192 |
| 57148_at | RALGAPB | 0,0691475 | 8,55733526 | 0,58587691 | 0,56695088 | 0,76942192 |

| | | | | | | |
|---|---|---|---|---|---|---|
| 10468_at | FST | -0,07599565 | 4,67627623 | -0,58563434 | 0,56710959 | 0,76953892 |
| 220082_at | SPERT | -0,04210966 | 4,39997176 | -0,58562303 | 0,56711699 | 0,76953892 |
| 51582_at | AZIN1 | -0,08108126 | 9,9668131 | -0,58544886 | 0,56723096 | 0,76963936 |
| 101929607_a | LOC10192960 | 0,05509203 | 3,14547604 | 0,58514112 | 0,56743236 | 0,7698584 |
| 100505687_a | LINC00888 | 0,20981789 | 7,22637742 | 0,58498147 | 0,56753686 | 0,76986992 |
| 79136_at | LY6G6E | -0,08537284 | 4,95598367 | -0,58490576 | 0,56758642 | 0,76986992 |
| 671_at | BPI | -0,04857541 | 3,57935701 | -0,58488852 | 0,56759771 | 0,76986992 |
| 25874_at | MPC2 | 0,0796351 | 7,79973034 | 0,58488392 | 0,56760072 | 0,76986992 |
| 100505894_a | TMEM161B-A | -0,08729916 | 5,28830949 | -0,58469722 | 0,56772294 | 0,76992813 |
| 285195_at | SLC9A9 | -0,06911559 | 5,2046701 | -0,58457328 | 0,56780409 | 0,76992813 |
| 340419_at | RSPO2 | -0,03719186 | 3,35768869 | -0,58455794 | 0,56781413 | 0,76992813 |
| 5272_at | SERPINB9 | 0,04755315 | 3,91916072 | 0,58453361 | 0,56783007 | 0,76992813 |
| 399821_at | FLJ37035 | 0,0615659 | 3,72259557 | 0,58446726 | 0,56787351 | 0,76992813 |
| 5599_at | MAPK8 | -0,08305518 | 7,02214196 | -0,58445209 | 0,56788345 | 0,76992813 |
| 81577_at | GFOD2 | -0,02670814 | 6,54598533 | -0,58431473 | 0,56797339 | 0,76999588 |
| 2674_at | GFRA1 | -0,0411023 | 3,79123143 | -0,58424839 | 0,56801684 | 0,77000058 |
| 79727_at | LIN28A | -0,09682467 | 3,50702582 | -0,58392679 | 0,56822747 | 0,77023191 |
| 56884_at | FSTL5 | 0,03167703 | 2,95912896 | 0,58379789 | 0,5683119 | 0,77029216 |
| 10516_at | FBLN5 | -0,06417063 | 4,29013741 | -0,58368955 | 0,56838287 | 0,77033414 |
| 283025_at | LINC01553 | -0,06395707 | 3,30022965 | -0,58347679 | 0,56852226 | 0,77046885 |
| 54969_at | C4orf27 | -0,14314253 | 7,64516234 | -0,58324411 | 0,56867472 | 0,77062125 |
| 10006_at | ABI1 | 0,13300815 | 8,4662969 | 0,58303712 | 0,56881037 | 0,77075085 |
| 84970_at | C1orf94 | -0,06297871 | 4,86804866 | -0,58281852 | 0,56895363 | 0,77085495 |
| 50_at | ACO2 | -0,06263029 | 9,39253308 | -0,58272735 | 0,5690134 | 0,77085495 |
| 84897_at | TBRG1 | -0,06187503 | 7,19855344 | -0,58254701 | 0,56913161 | 0,77085495 |
| 117289_at | TAGAP | 0,04543323 | 3,66407895 | 0,58249386 | 0,56916646 | 0,77085495 |
| 79917_at | MAGIX | 0,05073684 | 5,13059479 | 0,58245043 | 0,56919493 | 0,77085495 |
| 11031_at | RAB31 | -0,12108242 | 6,53191349 | -0,58236834 | 0,56924875 | 0,77085495 |
| 57504_at | MTA3 | 0,06972527 | 6,54220172 | 0,58231621 | 0,56928293 | 0,77085495 |
| 286827_at | TRIM59 | -0,19197921 | 6,74022293 | -0,58228793 | 0,56930147 | 0,77085495 |
| 8740_at | TNFSF14 | 0,04226797 | 4,51071268 | 0,58228396 | 0,56930407 | 0,77085495 |
| 677766_at | SCARNA2 | -0,05018422 | 4,83855702 | -0,58226016 | 0,56931968 | 0,77085495 |
| 7349_at | UCN | 0,0532651 | 4,98862688 | 0,58218708 | 0,5693676 | 0,77085495 |
| 80762_at | NDFIP1 | -0,05629936 | 9,30439596 | -0,58207568 | 0,56944065 | 0,77085495 |
| 148709_at | LOC148709 | -0,0587119 | 4,32398735 | -0,58206349 | 0,56944864 | 0,77085495 |
| 100874243_a | PRICKLE2-AS3 | -0,04221198 | 3,14817228 | -0,58205898 | 0,56945159 | 0,77085495 |
| 101928238_a | LOC10192823 | -0,03123993 | 3,48985691 | -0,5820043 | 0,56948745 | 0,77085495 |
| 343637_at | RSPO4 | -0,0518576 | 4,96299482 | -0,58189537 | 0,56955889 | 0,77089748 |
| 101929705_a | LOC10192970 | -0,05926749 | 4,32844871 | -0,58176179 | 0,56964649 | 0,77096188 |
| 3811_at | KIR3DL1 | -0,07890474 | 4,3796799 | -0,58158322 | 0,56976362 | 0,77106622 |
| 26290_at | GALNT8 | -0,04753126 | 4,27829404 | -0,58145983 | 0,56984456 | 0,77112158 |
| 100506470_a | LOC10050647 | -0,07549862 | 5,16950146 | -0,58119747 | 0,57001668 | 0,77130032 |
| 114823_at | LENG8 | 0,09125221 | 7,4427798 | 0,58098377 | 0,5701569 | 0,77139419 |
| 8942_at | KYNU | 0,21881982 | 5,87526656 | 0,58080657 | 0,57027318 | 0,77139419 |
| 389320_at | TEX43 | -0,07137307 | 3,18106222 | -0,58079937 | 0,57027791 | 0,77139419 |
| 2004_at | ELK3 | -0,06780836 | 6,84941038 | -0,58078799 | 0,57028537 | 0,77139419 |
| 79368_at | FCRL2 | -0,04873416 | 3,98286656 | -0,58078661 | 0,57028628 | 0,77139419 |
| 26189_at | OR1A2 | 0,06699093 | 3,34678376 | 0,5807124 | 0,57033498 | 0,77140589 |
| 390058_at | OR51B6 | -0,04291305 | 3,75209383 | -0,5804085 | 0,57053445 | 0,77158631 |
| 4153_at | MBL2 | 0,05469907 | 3,38298508 | 0,58038712 | 0,57054848 | 0,77158631 |

| | | | | | | |
|---|---|---|---|---|---|---|
| 3725_at | JUN | -0,12668902 | 8,33021742 | -0,58026088 | 0,57063135 | 0,77164421 |
| 100129794_a | SLC25A21-AS1 | 0,09581165 | 5,00685131 | 0,58015885 | 0,57069834 | 0,77168062 |
| 55237_at | VRTN | -0,0460128 | 5,41664291 | -0,58003037 | 0,57078269 | 0,77171687 |
| 55233_at | MOB1A | -0,08861545 | 9,24544595 | -0,57996716 | 0,57082418 | 0,77171687 |
| 54496_at | PRMT7 | 0,04460139 | 7,45348467 | 0,57993496 | 0,57084533 | 0,77171687 |
| 158833_at | AWAT1 | -0,05876073 | 4,16416378 | -0,57977018 | 0,57095353 | 0,77171787 |
| 10135_at | NAMPT | -0,10207912 | 9,6783171 | -0,57975833 | 0,57096131 | 0,77171787 |
| 79845_at | RNF122 | -0,03849869 | 6,10271248 | -0,5797508 | 0,57096626 | 0,77171787 |
| 1534_at | CYB561 | -0,03954101 | 8,70353275 | -0,57955616 | 0,57109408 | 0,77178707 |
| 285286_at | LINC00877 | -0,06370691 | 3,78211079 | -0,57955081 | 0,57109759 | 0,77178707 |
| 29995_at | LMCD1 | 0,05490308 | 5,47870781 | 0,57908851 | 0,57140125 | 0,77203248 |
| 54977_at | SLC25A38 | -0,05942474 | 8,24378034 | -0,57908299 | 0,57140488 | 0,77203248 |
| 54866_at | PPP1R14D | -0,07580905 | 5,03280028 | -0,57894736 | 0,57149398 | 0,77203248 |
| 5475_at | PPEF1 | -0,03874515 | 3,11748012 | -0,57893943 | 0,57149919 | 0,77203248 |
| 105370939_a | LOC10537093 | -0,0545163 | 4,50158952 | -0,57890151 | 0,57152411 | 0,77203248 |
| 1788_at | DNMT3A | 0,05240288 | 6,34866803 | 0,57887296 | 0,57154286 | 0,77203248 |
| 619207_at | SCART1 | -0,05347055 | 5,40288442 | -0,57884728 | 0,57155973 | 0,77203248 |
| 23704_at | KCNE4 | -0,05207314 | 3,88784611 | -0,57866449 | 0,57167984 | 0,77214057 |
| 26095_at | PTPN20 | 0,09619519 | 2,97785387 | 0,5785796 | 0,57173563 | 0,77216177 |
| 3082_at | HGF | -0,03115049 | 3,13586655 | -0,57829995 | 0,57191941 | 0,77232259 |
| 25774_at | GSTTP1 | -0,10042482 | 5,04778383 | -0,57813309 | 0,57202908 | 0,77232259 |
| 161247_at | FITM1 | -0,07454289 | 4,83594081 | -0,57813063 | 0,5720307 | 0,77232259 |
| 1808_at | DPYSL2 | -0,30293189 | 9,89240426 | -0,5780596 | 0,57207739 | 0,77232259 |
| 55784_at | MCTP2 | -0,08688844 | 6,13073569 | -0,57797902 | 0,57213036 | 0,77232259 |
| 91149_at | RAPGEF4-AS1 | 0,06709657 | 3,6292587 | 0,57795419 | 0,57214668 | 0,77232259 |
| 27179_at | IL36A | -0,0532174 | 5,26197858 | -0,57785896 | 0,57220929 | 0,77232259 |
| 192668_at | CYS1 | 0,06733843 | 4,90916161 | 0,57780535 | 0,57224454 | 0,77232259 |
| 79925_at | SPEF2 | -0,0448792 | 3,16792418 | -0,57775249 | 0,57227929 | 0,77232259 |
| 254122_at | SNX32 | -0,04010686 | 4,86210522 | -0,57768203 | 0,57232562 | 0,77232259 |
| 123096_at | SLC25A29 | 0,12404354 | 6,08225247 | 0,5776717 | 0,57233241 | 0,77232259 |
| 119587_at | CPXM2 | -0,04761124 | 5,10562322 | -0,57763576 | 0,57235604 | 0,77232259 |
| 352999_at | C6orf58 | -0,05470654 | 3,70591521 | -0,57760552 | 0,57237592 | 0,77232259 |
| 387723_at | LINC00959 | 0,20129266 | 5,03477861 | 0,57752126 | 0,57243132 | 0,77234325 |
| 83716_at | CRISPLD2 | -0,26061278 | 6,88153541 | -0,57734383 | 0,572548 | 0,77243571 |
| 7932_at | OR2H2 | 0,05887324 | 3,6286505 | 0,5772951 | 0,57258005 | 0,77243571 |
| 63876_at | PKNOX2 | -0,05694058 | 4,74356871 | -0,57709345 | 0,57271267 | 0,77252112 |
| 11072_at | DUSP14 | 0,0743811 | 7,77896869 | 0,57707688 | 0,57272357 | 0,77252112 |
| 285268_at | ZNF621 | 0,04742791 | 6,03446547 | 0,57697862 | 0,5727882 | 0,7725542 |
| 26074_at | CFAP61 | -0,03973643 | 3,64630359 | -0,57686596 | 0,57286231 | 0,77260006 |
| 7343_at | UBTF | -0,07445017 | 7,53859054 | -0,5765453 | 0,57307327 | 0,77281448 |
| 4829_at | NMBR | -0,03857079 | 3,57904997 | -0,57639842 | 0,57316991 | 0,77281448 |
| 400955_at | LINC01122 | 0,0345103 | 3,10143102 | 0,57631754 | 0,57322313 | 0,77281448 |
| 134510_at | UBLCP1 | 0,07248888 | 8,22812057 | 0,57631391 | 0,57322552 | 0,77281448 |
| 81853_at | TMEM14B | -0,10303545 | 10,6313968 | -0,57628677 | 0,57324338 | 0,77281448 |
| 10045_at | SH2D3A | -0,05469032 | 7,98159967 | -0,57625466 | 0,57326451 | 0,77281448 |
| 256764_at | WDR72 | 0,09726424 | 4,94819428 | 0,57619749 | 0,57330213 | 0,77281448 |
| 832_at | CAPZB | 0,03707108 | 10,7189215 | 0,57606518 | 0,5733892 | 0,77285341 |
| 91975_at | ZNF300 | -0,05696599 | 5,83103185 | -0,57593822 | 0,57347277 | 0,77285341 |
| 54440_at | SASH3 | -0,07669532 | 4,54065174 | -0,57593348 | 0,57347589 | 0,77285341 |
| 138724_at | C9orf131 | -0,07916731 | 5,03849309 | -0,57590976 | 0,5734915 | 0,77285341 |

| | | | | | | |
|---|---|---|---|---|---|---|
| 4808_at | NHLH2 | -0,03447178 | 3,71202753 | -0,57572053 | 0,57361606 | 0,77293989 |
| 23109_at | DDN | -0,0813384 | 5,80198154 | -0,57558767 | 0,57370352 | 0,77293989 |
| 1576_at | CYP3A4 | -0,06206992 | 5,12222849 | -0,57557226 | 0,57371366 | 0,77293989 |
| 54884_at | RETSAT | 0,14487698 | 8,88651245 | 0,57556845 | 0,57371617 | 0,77293989 |
| 100505540_a | SLIT1-AS1 | -0,07172826 | 3,66130506 | -0,57540662 | 0,57382271 | 0,77302937 |
| 197021_at | LCTL | -0,04004676 | 3,28451179 | -0,57523747 | 0,57393409 | 0,77309311 |
| 339622_at | LOC339622 | 0,03099406 | 2,70051987 | 0,57521285 | 0,5739503 | 0,77309311 |
| 5018_at | OXA1L | 0,05439015 | 10,4892491 | 0,5750866 | 0,57403343 | 0,77315103 |
| 3937_at | LCP2 | 0,05562282 | 3,66383861 | 0,57478221 | 0,5742339 | 0,77336696 |
| 80217_at | CFAP43 | -0,03359634 | 2,98562445 | -0,57471544 | 0,57427788 | 0,77337212 |
| 133690_at | CAPSL | -0,04163598 | 3,26875507 | -0,57443979 | 0,57445946 | 0,77356257 |
| 84432_at | PROK1 | 0,06825737 | 4,8827923 | 0,57432193 | 0,57453711 | 0,77361305 |
| 56253_at | CRTAM | -0,0443771 | 3,21371635 | -0,57426078 | 0,5745774 | 0,77361322 |
| 9577_at | BRE | -0,08831283 | 8,35507457 | -0,57415207 | 0,57464902 | 0,77365558 |
| 1993_at | ELAVL2 | -0,22592091 | 4,37702461 | -0,57366314 | 0,57497122 | 0,77398168 |
| 1198_at | CLK3 | 0,04633808 | 5,96625957 | 0,57366256 | 0,5749716 | 0,77398168 |
| 55181_at | SMG8 | 0,06107812 | 7,80857739 | 0,57306503 | 0,5753655 | 0,7744578 |
| 1258_at | CNGB1 | -0,05364229 | 4,92891443 | -0,57299798 | 0,5754097 | 0,77446319 |
| 54518_at | APBB1IP | -0,03213275 | 3,87965781 | -0,57292369 | 0,57545869 | 0,77447501 |
| 148022_at | TICAM1 | -0,05168287 | 6,80548316 | -0,57259945 | 0,57567251 | 0,77470864 |
| 378805_at | LINC-PINT | -0,05485289 | 4,54831445 | -0,57238794 | 0,57581201 | 0,77479856 |
| 222256_at | CDHR3 | -0,02925939 | 3,71412853 | -0,57237618 | 0,57581977 | 0,77479856 |
| 158376_at | LINC00961 | -0,08989774 | 4,7418491 | -0,57228964 | 0,57587685 | 0,77481885 |
| 10060_at | ABCC9 | 0,02809416 | 3,16737516 | 0,57223136 | 0,57591529 | 0,77481885 |
| 64388_at | GREM2 | -0,03918214 | 3,66631698 | -0,57168214 | 0,57627764 | 0,7752522 |
| 8448_at | DOC2A | 0,11468336 | 5,68541208 | 0,57143051 | 0,5764437 | 0,7753796 |
| 54989_at | ZNF770 | -0,12025995 | 6,92077137 | -0,57136759 | 0,57648523 | 0,7753796 |
| 57226_at | LYRM2 | -0,05233388 | 7,187867 | -0,57135566 | 0,5764931 | 0,7753796 |
| 11128_at | POLR3A | -0,0278498 | 7,47437966 | -0,57111546 | 0,57665164 | 0,77549264 |
| 375607_at | NAT16 | -0,08385579 | 5,14895996 | -0,57110634 | 0,57665767 | 0,77549264 |
| 64412_at | GZF1 | -0,04929363 | 6,93184092 | -0,57102091 | 0,57671406 | 0,77551433 |
| 402160_at | CFAP99 | -0,08270825 | 5,62351217 | -0,57085805 | 0,57682157 | 0,77560477 |
| 9726_at | ZNF646 | 0,06124394 | 5,93148624 | 0,57078033 | 0,57687289 | 0,77561963 |
| 2913_at | GRM3 | 0,0579999 | 3,70522995 | 0,5706064 | 0,57698772 | 0,77571988 |
| 4938_at | OAS1 | -0,16863611 | 9,36791688 | -0,57049104 | 0,5770639 | 0,77576815 |
| 91283_at | MSANTD3 | -0,06399582 | 7,53791627 | -0,56994225 | 0,57742634 | 0,77613875 |
| 25894_at | PLEKHG4 | 0,1013878 | 6,00882415 | 0,56986804 | 0,57747536 | 0,77613875 |
| 55026_at | TMEM255A | 0,07387184 | 3,91014795 | 0,5698347 | 0,57749739 | 0,77613875 |
| 83637_at | ZMIZ2 | -0,03563509 | 7,60820696 | -0,56982963 | 0,57750073 | 0,77613875 |
| 2037_at | EPB41L2 | -0,37461997 | 6,69954725 | -0,56945928 | 0,57774541 | 0,7763685 |
| 7264_at | TSTA3 | -0,23229963 | 9,61157629 | -0,56939077 | 0,57779069 | 0,7763685 |
| 23410_at | SIRT3 | 0,06939038 | 6,45731619 | 0,56938788 | 0,5777926 | 0,7763685 |
| 653_at | BMP5 | -0,05057314 | 3,74023601 | -0,5689807 | 0,57806168 | 0,77667588 |
| 4036_at | LRP2 | -0,0452718 | 3,54334898 | -0,56879443 | 0,5781848 | 0,77678712 |
| 100507605_a | LINC00837 | 0,03844989 | 3,49035111 | 0,56851849 | 0,57836721 | 0,77693725 |
| 145258_at | GSC | -0,05418112 | 5,38259389 | -0,56850337 | 0,57837721 | 0,77693725 |
| 133923_at | ZNF474 | 0,0363261 | 3,45382018 | 0,56836902 | 0,57846603 | 0,77700238 |
| 83741_at | TFAP2D | -0,045044 | 3,78817738 | -0,56818442 | 0,57858809 | 0,77711215 |
| 4713_at | NDUFB7 | -0,06829021 | 9,83297176 | -0,56810863 | 0,57863821 | 0,77712528 |
| 93661_at | CAPZA3 | -0,05323761 | 3,0906451 | -0,56798973 | 0,57871684 | 0,77717267 |

| | | | | | | |
|---|---|---|---|---|---|---|
| 124404_at | SEPT12 | -0,046404 | 6,05393052 | -0,56793326 | 0,57875419 | 0,77717267 |
| 64405_at | CDH22 | -0,05839024 | 5,34569852 | -0,56773356 | 0,57888627 | 0,77728617 |
| 8539_at | API5 | -0,08285319 | 7,33467305 | -0,56768344 | 0,57891941 | 0,77728617 |
| 147645_at | VSIG10L | -0,13036195 | 6,4342808 | -0,56757821 | 0,57898902 | 0,77732544 |
| 101929730_a | LINC01352 | -0,03865518 | 3,02717867 | -0,56744768 | 0,57907536 | 0,77735805 |
| 9988_at | DMTF1 | 0,11263815 | 8,23935534 | 0,56740891 | 0,57910101 | 0,77735805 |
| 8452_at | CUL3 | -0,03982109 | 8,53310472 | -0,56735849 | 0,57913437 | 0,77735805 |
| 8847_at | DLEU2 | -0,04435833 | 3,82453573 | -0,56719003 | 0,57924582 | 0,77745348 |
| 155006_at | TMEM213 | -0,0548797 | 3,93953638 | -0,56661754 | 0,57962467 | 0,77788371 |
| 55102_at | ATG2B | 0,04995887 | 6,69897396 | 0,5665836 | 0,57964713 | 0,77788371 |
| 8899_at | PRPF4B | -0,045419 | 8,24001366 | -0,56643907 | 0,5797428 | 0,77788967 |
| 8536_at | CAMK1 | 0,12328834 | 5,45454833 | 0,56643554 | 0,57974514 | 0,77788967 |
| 574407_at | C1orf145 | 0,03437933 | 4,01808462 | 0,56639386 | 0,57977273 | 0,77788967 |
| 1_at | A1BG | -0,0621143 | 4,61178381 | -0,56633141 | 0,57981406 | 0,77789095 |
| 5080_at | PAX6 | 0,03269093 | 3,55461551 | 0,56604899 | 0,58000104 | 0,77808761 |
| 51450_at | PRRX2 | -0,07086589 | 6,65789677 | -0,56590039 | 0,58009942 | 0,7781654 |
| 100506714_a | NUP50-AS1 | -0,05149701 | 6,09389859 | -0,56543871 | 0,58040516 | 0,77852132 |
| 84068_at | SLC10A7 | 0,08232567 | 4,93291061 | 0,56514613 | 0,58059896 | 0,77869519 |
| 24141_at | LAMP5 | 0,0447812 | 4,44129173 | 0,56511883 | 0,58061705 | 0,77869519 |
| 84417_at | C2orf40 | -0,05288543 | 3,49033962 | -0,56505994 | 0,58065606 | 0,77869519 |
| 100287177_a | EML2-AS1 | -0,07893616 | 4,88049579 | -0,56481171 | 0,58082052 | 0,77885515 |
| 162540_at | SPPL2C | -0,06854449 | 5,06261172 | -0,56475785 | 0,58085621 | 0,77885515 |
| 25791_at | NGEF | -0,04955562 | 5,33652817 | -0,5645163 | 0,58101627 | 0,77901555 |
| 6533_at | SLC6A6 | -0,06153653 | 7,50353501 | -0,56395087 | 0,58139103 | 0,7793017 |
| 55180_at | LINS1 | -0,08403331 | 5,86991479 | -0,56384991 | 0,58145796 | 0,7793017 |
| 4122_at | MAN2A2 | 0,094738 | 6,75156832 | 0,56382729 | 0,58147295 | 0,7793017 |
| 100859930_a | HEIH | 0,1692731 | 8,23628167 | 0,56375604 | 0,58152019 | 0,7793017 |
| 55118_at | CRTAC1 | -0,05395303 | 4,96509853 | -0,56374249 | 0,58152917 | 0,7793017 |
| 83550_at | GPR101 | -0,05366144 | 4,76781346 | -0,56372852 | 0,58153843 | 0,7793017 |
| 79718_at | TBL1XR1 | 0,07071484 | 9,60529456 | 0,56372478 | 0,58154091 | 0,7793017 |
| 255130_at | IGFBP7-AS1 | -0,06751564 | 3,21139804 | -0,5636643 | 0,58158101 | 0,7793017 |
| 57103_at | TIGAR | -0,11324899 | 7,92603476 | -0,56364502 | 0,58159379 | 0,7793017 |
| 91754_at | NEK9 | 0,05351518 | 7,63206052 | 0,5634489 | 0,58172383 | 0,77937315 |
| 255061_at | TAC4 | -0,08797755 | 4,82916469 | -0,56344256 | 0,58172804 | 0,77937315 |
| 101929406_a | LOC10192940 | -0,06769815 | 3,95633178 | -0,56313681 | 0,5819308 | 0,77954773 |
| 4643_at | MYO1E | 0,19980985 | 7,44578629 | 0,56312403 | 0,58193928 | 0,77954773 |
| 80210_at | ARMC9 | -0,04332851 | 5,74531742 | -0,56304453 | 0,581992 | 0,77956415 |
| 374946_at | DRAXIN | -0,06732167 | 4,32654203 | -0,562839 | 0,58212833 | 0,77966669 |
| 130888_at | FBXO36 | 0,15128095 | 6,31065722 | 0,56280707 | 0,58214951 | 0,77966669 |
| 1800_at | DPEP1 | -0,06376606 | 5,8890045 | -0,5626538 | 0,58225119 | 0,77974865 |
| 55339_at | WDR33 | -0,05281303 | 6,33257644 | -0,56229206 | 0,5824912 | 0,77996408 |
| 101929284_a | LOC10192928 | -0,09500963 | 4,30860095 | -0,56223774 | 0,58252724 | 0,77996408 |
| 5518_at | PPP2R1A | 0,0784003 | 10,0187457 | 0,56222828 | 0,58253352 | 0,77996408 |
| 8100_at | IFT88 | 0,07816609 | 6,17034588 | 0,56204851 | 0,58265282 | 0,78006959 |
| 440028_at | LOC440028 | 0,05357957 | 4,76155893 | 0,56186952 | 0,58277161 | 0,78007353 |
| 841_at | CASP8 | -0,06868185 | 7,64923888 | -0,56180693 | 0,58281316 | 0,78007353 |
| 51282_at | SCAND1 | 0,06432501 | 9,27837756 | 0,56177752 | 0,58283267 | 0,78007353 |
| 343_at | AQP8 | -0,04617188 | 5,33928355 | -0,56175473 | 0,5828478 | 0,78007353 |
| 3384_at | ICAM2 | -0,25813351 | 5,69858173 | -0,561739 | 0,58285824 | 0,78007353 |
| 101929517_a | LOC10192951 | -0,05429784 | 3,80165011 | -0,5615759 | 0,58296651 | 0,78008357 |

| | | | | | | |
|---|---|---|---|---|---|---|
| 51066_at | SSUH2 | 0,07808904 | 5,14449216 | 0,56151831 | 0,58300474 | 0,78008357 |
| 55805_at | LRP2BP | -0,049405 | 3,71499676 | -0,56148751 | 0,58302519 | 0,78008357 |
| 157697_at | ERICH1 | 0,04627732 | 5,94534085 | 0,56142428 | 0,58306717 | 0,78008357 |
| 126917_at | IFFO2 | -0,11846947 | 8,0998101 | -0,56142268 | 0,58306823 | 0,78008357 |
| 7786_at | MAP3K12 | -0,05514396 | 6,26151838 | -0,56128968 | 0,58315653 | 0,78014752 |
| 91768_at | CABLES1 | 0,20888444 | 6,72840642 | 0,56110437 | 0,58327957 | 0,78025794 |
| 2628_at | GATM | 0,54874849 | 5,77701782 | 0,56092105 | 0,5834013 | 0,78036659 |
| 129293_at | TRABD2A | -0,0733019 | 5,46649229 | -0,56072888 | 0,58352893 | 0,78043446 |
| 9208_at | LRRFIP1 | -0,04702065 | 8,00616461 | -0,56068553 | 0,58355772 | 0,78043446 |
| 9909_at | DENND4B | -0,05796381 | 8,42682202 | -0,56064007 | 0,58358792 | 0,78043446 |
| 9750_at | FAM65B | 0,03793162 | 3,9297165 | 0,56060065 | 0,5836141 | 0,78043446 |
| 55328_at | RNLS | -0,06104381 | 4,77272785 | -0,56050258 | 0,58367924 | 0,78046739 |
| 66005_at | CHID1 | -0,08504865 | 7,92695627 | -0,56038903 | 0,58375467 | 0,78051407 |
| 387695_at | C10orf99 | 0,32619666 | 5,31544606 | 0,56015672 | 0,583909 | 0,78065854 |
| 81876_at | RAB1B | 0,0883714 | 9,68779944 | 0,56010439 | 0,58394377 | 0,78065854 |
| 9053_at | MAP7 | -0,03843414 | 8,49376693 | -0,5599287 | 0,5840605 | 0,78076041 |
| 85379_at | KIAA1671 | -0,08195104 | 7,10535983 | -0,55961501 | 0,58426896 | 0,78096686 |
| 152206_at | CCDC13 | -0,05985488 | 5,03873297 | -0,55952315 | 0,58433002 | 0,78096686 |
| 6006_at | RHCE | -0,07838864 | 4,67047373 | -0,55951328 | 0,58433657 | 0,78096686 |
| 9244_at | CRLF1 | -0,19328435 | 5,9454528 | -0,55942604 | 0,58439456 | 0,78099017 |
| 54626_at | HES2 | -0,06926919 | 5,16028065 | -0,55900739 | 0,58467286 | 0,78130789 |
| 129563_at | DIS3L2 | 0,03500978 | 6,57894233 | 0,55889484 | 0,58474768 | 0,78135368 |
| 64857_at | PLEKHG2 | 0,05870139 | 6,5209354 | 0,55873645 | 0,584853 | 0,7814402 |
| 23633_at | KPNA6 | 0,03958582 | 8,76521182 | 0,55857547 | 0,58496005 | 0,78152902 |
| 101929497_a | LOC10192949 | -0,04846996 | 3,00596718 | -0,55804572 | 0,58531238 | 0,78194552 |
| 23400_at | ATP13A2 | 0,0733821 | 7,83218761 | 0,55793126 | 0,58538853 | 0,78199301 |
| 64072_at | CDH23 | 0,07362623 | 5,39179459 | 0,55780379 | 0,58547333 | 0,78205206 |
| 100506046_a | GAS5-AS1 | 0,06790933 | 4,08773552 | 0,55765705 | 0,58557096 | 0,782072 |
| 26628_at | OR7E47P | -0,08055438 | 5,71891407 | -0,55760181 | 0,58560772 | 0,782072 |
| 100289509_a | KCNIP2-AS1 | -0,05998257 | 4,57288254 | -0,55759829 | 0,58561006 | 0,782072 |
| 245802_at | MS4A6E | -0,06211244 | 2,89484735 | -0,55734739 | 0,58577702 | 0,78224074 |
| 220136_at | CFAP53 | 0,06080041 | 3,51645394 | 0,5572696 | 0,58582879 | 0,78225564 |
| 125058_at | TBC1D16 | -0,0461784 | 6,777128 | -0,55705513 | 0,58597152 | 0,78233701 |
| 25799_at | ZNF324 | -0,0538024 | 7,04221814 | -0,55697842 | 0,58602258 | 0,78233701 |
| 113746_at | ODF3 | -0,06075531 | 4,5947823 | -0,55697647 | 0,58602388 | 0,78233701 |
| 1674_at | DES | -0,04871671 | 6,074075 | -0,55693396 | 0,58605218 | 0,78233701 |
| 221301_at | FAM26D | -0,04943134 | 3,40448092 | -0,55677021 | 0,58616118 | 0,7824283 |
| 115207_at | KCTD12 | -0,3464291 | 6,39534417 | -0,55657526 | 0,58629096 | 0,78251313 |
| 285095_at | LOC285095 | -0,06962371 | 4,46912283 | -0,55655271 | 0,58630598 | 0,78251313 |
| 130340_at | AP1S3 | 0,07470232 | 8,15200873 | 0,55644434 | 0,58637813 | 0,78255521 |
| 339788_at | LINC00298 | 0,0559334 | 3,11407248 | 0,55628336 | 0,58648533 | 0,78261143 |
| 362_at | AQP5 | -0,05590261 | 5,33978884 | -0,55623353 | 0,58651851 | 0,78261143 |
| 56650_at | CLDND1 | -0,1281068 | 9,9578794 | -0,55619804 | 0,58654214 | 0,78261143 |
| 5286_at | PIK3C2A | 0,05199468 | 7,4390527 | 0,5559965 | 0,58667635 | 0,78273629 |
| 339883_at | C3orf35 | -0,08779607 | 3,69010331 | -0,55573496 | 0,58685056 | 0,78291449 |
| 683_at | BST1 | -0,05729551 | 4,05419683 | -0,55566901 | 0,58689449 | 0,78291887 |
| 79797_at | ZNF408 | -0,05620335 | 6,96661137 | -0,55536513 | 0,58709693 | 0,78310769 |
| 728606_at | PCAT18 | -0,05074239 | 3,47028122 | -0,5553345 | 0,58711734 | 0,78310769 |
| 23416_at | KCNH3 | -0,05416843 | 5,77933667 | -0,55520854 | 0,58720126 | 0,78314588 |
| 126075_at | CCDC159 | 0,08791709 | 6,25553943 | 0,55516949 | 0,58722728 | 0,78314588 |

| | | | | | | |
|---|---|---|---|---|---|---|
| 100506071_a | LOC10050607 | -0,043333 | 3,1691462 | -0,55492367 | 0,58739109 | 0,78327302 |
| 4627_at | MYH9 | 0,07919093 | 9,52574759 | 0,55490439 | 0,58740394 | 0,78327302 |
| 26052_at | DNM3 | 0,0262986 | 3,51500609 | 0,55470497 | 0,58753684 | 0,78339601 |
| 8463_at | TEAD2 | -0,04688202 | 5,16863763 | -0,55449567 | 0,58767635 | 0,78342387 |
| 54149_at | C21orf91 | 0,0985078 | 7,35889016 | 0,5544453 | 0,58770992 | 0,78342387 |
| 117245_at | HRASLS5 | -0,04540812 | 4,44332806 | -0,55444222 | 0,58771198 | 0,78342387 |
| 4232_at | MEST | -0,12494813 | 10,4262244 | -0,55442956 | 0,58772041 | 0,78342387 |
| 56606_at | SLC2A9 | -0,21970429 | 7,65112154 | -0,55436238 | 0,5877652 | 0,78342935 |
| 163131_at | ZNF780B | 0,03677558 | 4,40702691 | 0,55419201 | 0,58787878 | 0,78351466 |
| 7515_at | XRCC1 | -0,06123876 | 8,57732863 | -0,55412702 | 0,58792211 | 0,78351466 |
| 79919_at | C2orf54 | 0,13058586 | 5,31430988 | 0,55408335 | 0,58795123 | 0,78351466 |
| 11059_at | WWP1 | -0,15086443 | 8,57222874 | -0,55385719 | 0,58810203 | 0,78362055 |
| 5499_at | PPP1CA | -0,02907688 | 11,2847311 | -0,55384217 | 0,58811204 | 0,78362055 |
| 4520_at | MTF1 | 0,06586236 | 7,80647717 | 0,55342443 | 0,58839064 | 0,78380138 |
| 11194_at | ABCB8 | -0,05650627 | 6,62697977 | -0,5533919 | 0,58841234 | 0,78380138 |
| 79576_at | NKAP | -0,05866918 | 7,40623726 | -0,55338342 | 0,58841799 | 0,78380138 |
| 716_at | C1S | 0,14206281 | 4,48861006 | 0,55333188 | 0,58845237 | 0,78380138 |
| 55041_at | PLEKHB2 | 0,09761392 | 9,23996627 | 0,55328776 | 0,5884818 | 0,78380138 |
| 56155_at | TEX14 | -0,05399283 | 3,84178027 | -0,55317492 | 0,58855707 | 0,78380138 |
| 51233_at | DRICH1 | -0,05831954 | 4,26499263 | -0,55312816 | 0,58858827 | 0,78380138 |
| 64795_at | RMND5A | 0,10073169 | 7,69538271 | 0,55311057 | 0,58860001 | 0,78380138 |
| 54165_at | DCUN1D1 | 0,03648651 | 6,14377175 | 0,55308012 | 0,58862032 | 0,78380138 |
| 170062_at | FAM47B | -0,04834804 | 3,66113622 | -0,55302865 | 0,58865466 | 0,78380138 |
| 2869_at | GRK5 | -0,20658805 | 6,10834447 | -0,55291738 | 0,58872889 | 0,78384605 |
| 11057_at | ABHD2 | 0,05937026 | 8,09799583 | 0,55283902 | 0,58878118 | 0,78386148 |
| 116448_at | OLIG1 | -0,04881807 | 4,09675644 | -0,55266602 | 0,58889661 | 0,78395737 |
| 7535_at | ZAP70 | -0,04958533 | 4,87344302 | -0,55259289 | 0,58894541 | 0,78395737 |
| 51300_at | TIMMDC1 | -0,09202577 | 10,6483163 | -0,55254811 | 0,58897529 | 0,78395737 |
| 2922_at | GRP | -0,05553254 | 3,70085411 | -0,55193763 | 0,58938276 | 0,78444553 |
| 909_at | CD1A | -0,07970396 | 4,17848146 | -0,55182519 | 0,58945782 | 0,78448316 |
| 5370_at | PMCHL2 | 0,08078715 | 3,96600213 | 0,55173062 | 0,58952096 | 0,78448316 |
| 84547_at | PGBD1 | -0,17556836 | 5,09686611 | -0,55171227 | 0,58953321 | 0,78448316 |
| 51332_at | SPTBN5 | 0,06000502 | 5,74038075 | 0,5515763 | 0,589624 | 0,78454977 |
| 65981_at | CAPRIN2 | 0,07538967 | 6,91752095 | 0,55134431 | 0,58977891 | 0,78466179 |
| 25962_at | KIAA1429 | -0,05186288 | 7,16951773 | -0,55132823 | 0,58978965 | 0,78466179 |
| 162514_at | TRPV3 | 0,08210947 | 4,92302335 | 0,55107445 | 0,58995914 | 0,78483307 |
| 23180_at | RFTN1 | -0,40126654 | 6,90162519 | -0,55090491 | 0,59007238 | 0,78488498 |
| 339210_at | C17orf67 | -0,04341168 | 4,65943228 | -0,55089403 | 0,59007965 | 0,78488498 |
| 619455_at | SPANXA2-OT1 | -0,07181521 | 3,40011439 | -0,55067242 | 0,59022769 | 0,78502768 |
| 552_at | AVPR1A | -0,03678027 | 3,50736006 | -0,55050575 | 0,59033904 | 0,78512157 |
| 84100_at | ARL6 | -0,07555939 | 5,60665703 | -0,55023122 | 0,59052247 | 0,78531131 |
| 51161_at | C3orf18 | 0,0352032 | 5,88510246 | 0,5500773 | 0,59062533 | 0,78536765 |
| 3112_at | HLA-DOB | 0,07329601 | 5,04481535 | 0,55001363 | 0,59066788 | 0,78536765 |
| 338321_at | NLRP9 | -0,07681745 | 4,61822995 | -0,54996192 | 0,59070244 | 0,78536765 |
| 8110_at | DPF3 | -0,0440599 | 4,21265192 | -0,54984001 | 0,59078392 | 0,78536765 |
| 1787_at | TRDMT1 | -0,11648938 | 3,87603731 | -0,5497948 | 0,59081414 | 0,78536765 |
| 284067_at | C17orf105 | 0,04385933 | 3,19954549 | 0,54978498 | 0,5908207 | 0,78536765 |
| 80055_at | PGAP1 | 0,13941708 | 6,62457105 | 0,5497408 | 0,59085023 | 0,78536765 |
| 101927286_a | LOC10192728 | -0,04687256 | 3,19909754 | -0,54943825 | 0,59105248 | 0,78555118 |
| 4889_at | NPY5R | 0,05224683 | 3,84797346 | 0,54941224 | 0,59106987 | 0,78555118 |

| | | | | | | |
|---|---|---|---|---|---|---|
| 9744_at | ACAP1 | 0,05400696 | 5,32190871 | 0,54916532 | 0,59123496 | 0,78571638 |
| 65996_at | CENPBD1P1 | -0,08107446 | 7,3195491 | -0,54909436 | 0,5912824 | 0,78572523 |
| 50507_at | NOX4 | 0,06056647 | 3,17858747 | 0,54890565 | 0,5914086 | 0,78583871 |
| 284069_at | FAM171A2 | -0,04871661 | 5,30607454 | -0,5487663 | 0,59150178 | 0,78590832 |
| 89869_at | PLCZ1 | -0,02998578 | 2,94565733 | -0,54858325 | 0,59162421 | 0,78601678 |
| 1318_at | SLC31A2 | 0,19720682 | 7,75152254 | 0,54819875 | 0,59188142 | 0,78625206 |
| 7044_at | LEFTY2 | -0,06792617 | 4,90325915 | -0,54819647 | 0,59188294 | 0,78625206 |
| 27315_at | PGAP2 | -0,03230318 | 7,82426088 | -0,54781189 | 0,59214027 | 0,78653964 |
| 130951_at | M1AP | -0,04046141 | 3,76559324 | -0,54760719 | 0,59227725 | 0,78666735 |
| 79696_at | ZC2HC1C | 0,15705393 | 4,92326658 | 0,54750041 | 0,59234871 | 0,78670802 |
| 136288_at | C7orf57 | -0,06172996 | 3,33746365 | -0,54736766 | 0,59243756 | 0,78677179 |
| 100192379_a | PP12613 | -0,10851612 | 4,10843951 | -0,54726039 | 0,59250936 | 0,78678709 |
| 10073_at | SNUPN | 0,09332836 | 7,41425656 | 0,54721674 | 0,59253858 | 0,78678709 |
| 5229_at | PGGT1B | 0,07311143 | 6,71632345 | 0,54715276 | 0,59258141 | 0,78678709 |
| 5814_at | PURB | -0,06964034 | 8,52942171 | -0,54710637 | 0,59261246 | 0,78678709 |
| 51571_at | FAM49B | -0,20584383 | 7,541373 | -0,54703777 | 0,59265839 | 0,78679383 |
| 59343_at | SENP2 | 0,07110748 | 8,34769238 | 0,5467036 | 0,59288212 | 0,7869736 |
| 223075_at | CCDC129 | -0,05489215 | 4,02971075 | -0,54664951 | 0,59291834 | 0,7869736 |
| 375757_at | SWI5 | 0,07978665 | 7,93430911 | 0,54657856 | 0,59296585 | 0,7869736 |
| 79820_at | CATSPERB | 0,03835542 | 3,52349878 | 0,54656213 | 0,59297685 | 0,7869736 |
| 8076_at | MFAP5 | -0,03683234 | 3,21322095 | -0,54649551 | 0,59302146 | 0,7869736 |
| 6001_at | RGS10 | 0,1825238 | 8,53490335 | 0,54646943 | 0,59303893 | 0,7869736 |
| 93643_at | TJAP1 | 0,02962366 | 7,60595113 | 0,54637676 | 0,59310099 | 0,78700174 |
| 55973_at | BCAP29 | -0,08783976 | 7,41411363 | -0,54620266 | 0,59321759 | 0,78710225 |
| 85450_at | ITPRIP | 0,14654497 | 6,53685721 | 0,54610527 | 0,59328282 | 0,78713459 |
| 386597_at | RNF144A-AS1 | -0,07023021 | 4,1274112 | -0,54596528 | 0,59337659 | 0,78718032 |
| 7323_at | UBE2D3 | 0,092095 | 8,35954103 | 0,54593179 | 0,59339903 | 0,78718032 |
| 100287114_a | LINC00421 | -0,07152424 | 5,20217213 | -0,54580696 | 0,59348265 | 0,78723704 |
| 100144596_a | LINC00029 | -0,04198904 | 5,48624117 | -0,54553714 | 0,59366342 | 0,78742261 |
| 114881_at | OSBPL7 | 0,08249886 | 6,10578247 | 0,54523537 | 0,59386564 | 0,78761639 |
| 51808_at | PHAX | -0,11573894 | 7,80471482 | -0,54519709 | 0,59389129 | 0,78761639 |
| 7791_at | ZYX | 0,07579986 | 8,29819048 | 0,54494276 | 0,59406174 | 0,7877882 |
| 221188_at | ADGRG5 | 0,04630968 | 5,44301327 | 0,54486003 | 0,5941172 | 0,7878075 |
| 1173_at | AP2M1 | -0,06504281 | 11,2259255 | -0,54471552 | 0,59421406 | 0,78788171 |
| 9411_at | ARHGAP29 | -0,28486807 | 7,69176371 | -0,54456938 | 0,59431203 | 0,78795738 |
| 80169_at | CTC1 | 0,03590333 | 5,58762523 | 0,54441146 | 0,59441791 | 0,78804351 |
| 90806_at | ANGEL2 | 0,05407106 | 7,09935898 | 0,54422264 | 0,59454451 | 0,78814959 |
| 84690_at | SPATA22 | -0,03664423 | 2,79543323 | -0,54408625 | 0,59463597 | 0,78814959 |
| 6792_at | CDKL5 | -0,05970265 | 4,92935763 | -0,54407643 | 0,59464256 | 0,78814959 |
| 404037_at | HAPLN4 | -0,09616737 | 4,74496284 | -0,54403956 | 0,59466728 | 0,78814959 |
| 9223_at | MAGI1 | 0,02416299 | 4,76334363 | 0,54398704 | 0,5947025 | 0,78814959 |
| 6938_at | TCF12 | 0,08786889 | 8,50065852 | 0,5437903 | 0,59483444 | 0,78827022 |
| 912_at | CD1D | -0,04114876 | 3,96216125 | -0,54371504 | 0,59488492 | 0,78828288 |
| 23589_at | CARHSP1 | 0,10055163 | 8,06080898 | 0,54360483 | 0,59495884 | 0,78832661 |
| 80031_at | SEMA6D | 0,06796656 | 3,98598267 | 0,54334877 | 0,59513062 | 0,78849997 |
| 100132116_a | ACTA2-AS1 | 0,04180834 | 4,19839401 | 0,54327084 | 0,5951829 | 0,788515 |
| 79142_at | PHF23 | -0,05641161 | 7,90575934 | -0,54312768 | 0,59527895 | 0,78858802 |
| 79782_at | LRRC31 | 0,03348428 | 2,98827297 | 0,54301504 | 0,59535453 | 0,78863391 |
| 94101_at | ORMDL1 | 0,06803875 | 7,55494062 | 0,54288781 | 0,5954399 | 0,78867681 |
| 26039_at | SS18L1 | -0,14051017 | 8,25040501 | -0,54284474 | 0,5954688 | 0,78867681 |

| | | | | | | |
|---|---|---|---|---|---|---|
| 57582_at | KCNT1 | -0,05734312 | 4,40806653 | -0,54277387 | 0,59551636 | 0,78868557 |
| 1152_at | CKB | 0,15500884 | 7,57311414 | 0,54253591 | 0,59567607 | 0,78884285 |
| 398_at | ARHGDIG | -0,05292412 | 5,8100965 | -0,54241419 | 0,59575776 | 0,78887993 |
| 283284_at | IGSF22 | 0,04511801 | 5,28047517 | 0,54237216 | 0,59578598 | 0,78887993 |
| 101927034_a | ROR1-AS1 | -0,03097148 | 2,93021511 | -0,54204774 | 0,59600376 | 0,78911405 |
| 100506677_a | AA06 | -0,05718304 | 3,94851951 | -0,54185166 | 0,59613541 | 0,78923411 |
| 441094_at | NR2F1-AS1 | -0,10884209 | 4,21592958 | -0,54172248 | 0,59622215 | 0,7892947 |
| 10670_at | RRAGA | 0,05750916 | 9,7786684 | 0,54151517 | 0,59636136 | 0,78934911 |
| 933_at | CD22 | -0,05516276 | 4,74253457 | -0,54151476 | 0,59636164 | 0,78934911 |
| 57513_at | CASKIN2 | 0,05007828 | 7,14575091 | 0,54143891 | 0,59641257 | 0,78934911 |
| 5999_at | RGS4 | -0,08076622 | 4,20033372 | -0,54140691 | 0,59643407 | 0,78934911 |
| 122970_at | ACOT4 | 0,08647205 | 4,38491313 | 0,54133465 | 0,5964826 | 0,78934911 |
| 100133612_a | LINC01134 | -0,07009018 | 6,28455077 | -0,54129517 | 0,59650911 | 0,78934911 |
| 283875_at | LINC00514 | -0,04446332 | 4,45314563 | -0,54115291 | 0,59660467 | 0,78940103 |
| 2565_at | GABRG1 | 0,0268465 | 3,03364562 | 0,54108985 | 0,59664702 | 0,78940103 |
| 222546_at | RFX6 | -0,04207797 | 3,05401898 | -0,54099826 | 0,59670855 | 0,78940103 |
| 126147_at | NTN5 | 0,0423215 | 4,91568434 | 0,54095434 | 0,59673806 | 0,78940103 |
| 101928526_a | LOC10192852 | -0,06417244 | 4,44804959 | -0,54093173 | 0,59675325 | 0,78940103 |
| 79624_at | ARMT1 | 0,09892649 | 9,04229777 | 0,54067703 | 0,59692436 | 0,7894991 |
| 221178_at | SPATA13 | -0,05180617 | 5,68315251 | -0,54064611 | 0,59694514 | 0,7894991 |
| 115825_at | WDFY2 | -0,18197691 | 5,91012536 | -0,54059722 | 0,59697799 | 0,7894991 |
| 11017_at | SNRNP27 | -0,11203783 | 8,67872103 | -0,54057736 | 0,59699133 | 0,7894991 |
| 56969_at | RPL23AP32 | -0,08590077 | 5,59770085 | -0,54044507 | 0,59708023 | 0,78956246 |
| 55790_at | CSGALNACT1 | -0,04793697 | 3,54649446 | -0,54032407 | 0,59716153 | 0,78961578 |
| 146223_at | CMTM4 | 0,13791069 | 8,11927392 | 0,5400077 | 0,59737416 | 0,78980448 |
| 4481_at | MSR1 | -0,03120685 | 3,29488596 | -0,53998972 | 0,59738625 | 0,78980448 |
| 5732_at | PTGER2 | 0,05642168 | 3,74770163 | 0,53989224 | 0,59745177 | 0,7898369 |
| 6138_at | RPL15 | 0,0923603 | 11,0435923 | 0,53957627 | 0,59766418 | 0,78994705 |
| 10850_at | CCL27 | -0,05764969 | 4,37762032 | -0,53955516 | 0,59767837 | 0,78994705 |
| 91752_at | ZNF804A | 0,05535663 | 3,99740785 | 0,53954343 | 0,59768626 | 0,78994705 |
| 64756_at | ATPAF1 | -0,05208816 | 8,69971658 | -0,53952428 | 0,59769913 | 0,78994705 |
| 728637_at | MEIKIN | -0,0574722 | 3,05376219 | -0,53934821 | 0,59781751 | 0,79001717 |
| 50837_at | TAS2R7 | -0,03571962 | 3,08014371 | -0,53932339 | 0,5978342 | 0,79001717 |
| 101927907_a | LOC10192790 | -0,06366961 | 3,13739732 | -0,53924151 | 0,59788926 | 0,79003572 |
| 51266_at | CLEC1B | -0,06733978 | 4,01918108 | -0,53911324 | 0,59797552 | 0,79009551 |
| 3630_at | INS | 0,07289828 | 4,22131322 | 0,53869622 | 0,59825598 | 0,79041187 |
| 26167_at | PCDHB5 | 0,13474035 | 5,46676013 | 0,53856901 | 0,59834155 | 0,79047071 |
| 284739_at | LINC00176 | 0,07120531 | 5,02734188 | 0,5382413 | 0,59856202 | 0,79070774 |
| 153733_at | CCDC112 | -0,09001024 | 5,4248281 | -0,53773929 | 0,59889983 | 0,79109973 |
| 57470_at | LRRC47 | -0,0385297 | 10,2760365 | -0,53760398 | 0,5989909 | 0,79112403 |
| 56128_at | PCDHB8 | -0,09566318 | 4,33674907 | -0,53756654 | 0,59901609 | 0,79112403 |
| 1316_at | KLF6 | -0,07720026 | 7,84618791 | -0,53752889 | 0,59904144 | 0,79112403 |
| 11343_at | MGLL | -0,18469908 | 6,72418869 | -0,53744756 | 0,59909618 | 0,7911421 |
| 85509_at | MBD3L1 | -0,04755894 | 3,39379541 | -0,53732768 | 0,59917688 | 0,79115368 |
| 6337_at | SCNN1A | -0,05846192 | 7,67635323 | -0,53728984 | 0,59920235 | 0,79115368 |
| 84376_at | HOOK3 | 0,10829047 | 7,12198801 | 0,53722824 | 0,59924382 | 0,79115368 |
| 56654_at | NPDC1 | 0,19186063 | 8,0213407 | 0,53719048 | 0,59926924 | 0,79115368 |
| 9989_at | PPP4R1 | -0,13968171 | 9,9680386 | -0,53677595 | 0,59954834 | 0,79142593 |
| 168391_at | GALNTL5 | -0,04087528 | 3,20003735 | -0,53676215 | 0,59955763 | 0,79142593 |
| 3231_at | HOXD1 | 0,19784495 | 6,84159096 | 0,53665125 | 0,59963231 | 0,79147028 |

| | | | | | | |
|---|---|---|---|---|---|---|
| 57472_at | CNOT6 | 0,04072479 | 8,88247124 | 0,53645042 | 0,59976756 | 0,79159455 |
| 57415_at | C3orf14 | 0,10064699 | 4,26224197 | 0,53633331 | 0,59984643 | 0,79164096 |
| 91683_at | SYT12 | -0,05911454 | 4,96815665 | -0,53624414 | 0,59990649 | 0,79164096 |
| 256979_at | SUN3 | -0,03618829 | 3,25243677 | -0,53621517 | 0,59992601 | 0,79164096 |
| 55107_at | ANO1 | 0,29385122 | 5,71265801 | 0,53571572 | 0,60026247 | 0,79203069 |
| 7507_at | XPA | 0,11958741 | 7,08274746 | 0,53561239 | 0,60033209 | 0,79206829 |
| 22934_at | RPIA | 0,06625446 | 8,37704767 | 0,53507127 | 0,60069676 | 0,79249515 |
| 6121_at | RPE65 | -0,0387828 | 2,93790577 | -0,53490512 | 0,60080875 | 0,79254306 |
| 101927482_a | LOC10192748 | 0,04688666 | 2,80738707 | 0,53489532 | 0,60081536 | 0,79254306 |
| 116369_at | SLC26A8 | -0,04969612 | 3,73340039 | -0,5345709 | 0,60103407 | 0,79272412 |
| 338694_at | LOC338694 | -0,04856322 | 4,622931 | -0,53453481 | 0,6010584 | 0,79272412 |
| 2710_at | GK | 0,11479114 | 5,52422533 | 0,53448366 | 0,60109288 | 0,79272412 |
| 90522_at | YIF1B | 0,02885055 | 6,83938374 | 0,53444755 | 0,60111723 | 0,79272412 |
| 25989_at | ULK3 | 0,07199356 | 7,02499297 | 0,53421458 | 0,60127432 | 0,7927662 |
| 147525_at | LINC00526 | 0,0659957 | 5,56471313 | 0,53408824 | 0,60135952 | 0,7927662 |
| 728215_at | FAM155A | -0,03303657 | 3,50037914 | -0,53406753 | 0,60137349 | 0,7927662 |
| 22880_at | MORC2 | -0,06441501 | 7,35482471 | -0,53406494 | 0,60137523 | 0,7927662 |
| 101927043_a | LOC10192704 | 0,04611032 | 3,49396833 | 0,53396185 | 0,60144476 | 0,7927662 |
| 63977_at | PRDM15 | -0,04722185 | 6,18257053 | -0,5339478 | 0,60145423 | 0,7927662 |
| 100873930_a | DLG3-AS1 | -0,04883221 | 5,56310784 | -0,53383911 | 0,60152754 | 0,7927662 |
| 642475_at | MROH6 | -0,09575548 | 7,16108327 | -0,53381109 | 0,60154644 | 0,7927662 |
| 10658_at | CELF1 | 0,05715536 | 8,48175594 | 0,53380311 | 0,60155182 | 0,7927662 |
| 101928663_a | LOC10192866 | -0,04419875 | 3,33980802 | -0,53378958 | 0,60156095 | 0,7927662 |
| 57060_at | PCBP4 | 0,07958549 | 7,07767154 | 0,53372895 | 0,60160184 | 0,7927662 |
| 25980_at | AAR2 | 0,03109482 | 8,5301734 | 0,53363123 | 0,60166776 | 0,79279882 |
| 84085_at | FBXO30 | -0,07397612 | 8,55951691 | -0,53335575 | 0,6018536 | 0,79298946 |
| 9436_at | NCR2 | -0,05264378 | 5,53521313 | -0,53325824 | 0,60191939 | 0,7930219 |
| 100128025_a | WWTR1-AS1 | -0,05231287 | 3,88249861 | -0,53317273 | 0,60197709 | 0,79304367 |
| 8604_at | SLC25A12 | -0,06752466 | 7,69917622 | -0,53309832 | 0,60202729 | 0,79305558 |
| 100505933_a | ADD3-AS1 | -0,05343682 | 3,98576341 | -0,53295266 | 0,60212558 | 0,79313082 |
| 285315_at | C3orf33 | 0,09876983 | 4,41992033 | 0,53253872 | 0,60240495 | 0,79340148 |
| 201633_at | TIGIT | 0,05846898 | 4,14869836 | 0,53252614 | 0,60241344 | 0,79340148 |
| 54951_at | COMMD8 | -0,05921627 | 9,15863623 | -0,53234956 | 0,60253263 | 0,79346931 |
| 4359_at | MPZ | -0,06912539 | 5,19251177 | -0,53230191 | 0,60256479 | 0,79346931 |
| 57020_at | C16orf62 | 0,21775594 | 7,55156032 | 0,53226678 | 0,60258851 | 0,79346931 |
| 114960_at | TSGA13 | -0,07138432 | 3,56281878 | -0,53207678 | 0,60271678 | 0,79352834 |
| 5005_at | ORM2 | -0,06047308 | 3,54090273 | -0,53196665 | 0,60279114 | 0,79352834 |
| 64940_at | STAG3L4 | -0,08658992 | 6,78365031 | -0,53189185 | 0,60284164 | 0,79352834 |
| 8412_at | BCAR3 | -0,0950399 | 8,27798111 | -0,53186833 | 0,60285752 | 0,79352834 |
| 55527_at | FEM1A | -0,04524051 | 7,86322824 | -0,53186571 | 0,60285929 | 0,79352834 |
| 284367_at | SIGLEC17P | -0,07340427 | 3,61864673 | -0,53180557 | 0,6028999 | 0,79352834 |
| 100126784_a | LOC10012678 | -0,05544031 | 4,84534492 | -0,53177328 | 0,6029217 | 0,79352834 |
| 60626_at | RIC8A | -0,44011058 | 7,60740302 | -0,53164609 | 0,60300759 | 0,79354877 |
| 139285_at | AMER1 | 0,05124029 | 5,10950821 | 0,53162828 | 0,60301962 | 0,79354877 |
| 101927682_a | CD81-AS1 | -0,0292132 | 4,99329203 | -0,53131268 | 0,60323276 | 0,79377503 |
| 53339_at | BTBD1 | -0,03772172 | 10,272779 | -0,53123154 | 0,60328757 | 0,79379293 |
| 255512_at | TOLLIP-AS1 | -0,0842661 | 5,1434215 | -0,53104606 | 0,60341286 | 0,79390356 |
| 3760_at | KCNJ3 | -0,03809802 | 3,45212715 | -0,53097945 | 0,60345786 | 0,79390853 |
| 2947_at | GSTM3 | 0,33080447 | 7,85562997 | 0,53053874 | 0,60375562 | 0,79424603 |
| 81573_at | ANKRD13C | 0,06366024 | 7,16981651 | 0,53030196 | 0,60391563 | 0,79431592 |

| | | | | | | |
|---|---|---|---|---|---|---|
| 57549_at | IGSF9 | -0,10282629 | 7,33699921 | -0,53027986 | 0,60393057 | 0,79431592 |
| 213_at | ALB | -0,03300282 | 2,95676689 | -0,53019778 | 0,60398604 | 0,79431592 |
| 3269_at | HRH1 | -0,11653439 | 6,86955919 | -0,53017638 | 0,6040005 | 0,79431592 |
| 57379_at | AICDA | -0,04283134 | 3,8427309 | -0,53015503 | 0,60401493 | 0,79431592 |
| 115004_at | MB21D1 | -0,35635542 | 6,24503356 | -0,52994775 | 0,60415503 | 0,79441374 |
| 339778_at | C2orf70 | -0,06800248 | 5,37812234 | -0,52992295 | 0,60417179 | 0,79441374 |
| 55365_at | TMEM176A | -0,09262522 | 5,68874941 | -0,52982871 | 0,6042355 | 0,79444327 |
| 100506122_a | LOC10050612 | -0,04544887 | 3,74880448 | -0,5296782 | 0,60433725 | 0,79452282 |
| 8626_at | TP63 | -0,14126964 | 7,71342774 | -0,52959855 | 0,60439109 | 0,79453939 |
| 101060691_a | NUTM2B-AS1 | 0,12143521 | 5,39533561 | 0,5294634 | 0,60448247 | 0,79457073 |
| 22918_at | CD93 | -0,05545437 | 4,35567854 | -0,52944127 | 0,60449743 | 0,79457073 |
| 23554_at | TSPAN12 | -0,13755321 | 7,24069868 | -0,52920415 | 0,60465776 | 0,79472725 |
| 101928519_a | LOC10192851 | -0,03161783 | 2,72297955 | -0,52899412 | 0,6047998 | 0,79484004 |
| 26036_at | ZNF451 | -0,02972158 | 6,97654741 | -0,52895522 | 0,60482611 | 0,79484004 |
| 1974_at | EIF4A2 | 0,07861315 | 11,6610624 | 0,52886783 | 0,60488521 | 0,79486349 |
| 199964_at | TMEM61 | -0,04071761 | 5,0792531 | -0,52872872 | 0,60497931 | 0,7949329 |
| 1979_at | EIF4EBP2 | 0,08861471 | 8,80167945 | 0,52861372 | 0,60505709 | 0,79498089 |
| 29969_at | MDFIC | -0,12268984 | 7,50648425 | -0,52846754 | 0,60515598 | 0,79505659 |
| 149684_at | LOC149684 | -0,04272289 | 3,95219028 | -0,52839726 | 0,60520353 | 0,79506483 |
| 5037_at | PEBP1 | -0,04389028 | 9,98026239 | -0,52829013 | 0,60527601 | 0,79510582 |
| 318_at | NUDT2 | -0,06838459 | 6,79806899 | -0,52816144 | 0,60536307 | 0,79512254 |
| 116113_at | FOXP4 | 0,02517418 | 6,98140097 | 0,5281493 | 0,60537129 | 0,79512254 |
| 51285_at | RASL12 | -0,04951174 | 5,26492705 | -0,52796111 | 0,60549863 | 0,79519639 |
| 102723665_a | LOC10272366 | 0,10922747 | 5,27343756 | 0,52794418 | 0,60551008 | 0,79519639 |
| 29123_at | ANKRD11 | 0,10843715 | 8,29242611 | 0,52782044 | 0,60559382 | 0,79520052 |
| 7592_at | ZNF41 | 0,04358939 | 5,79743538 | 0,52778112 | 0,60562043 | 0,79520052 |
| 339105_at | PRSS53 | -0,08905256 | 6,06419671 | -0,52772234 | 0,60566021 | 0,79520052 |
| 3569_at | IL6 | 0,04811048 | 4,30219258 | 0,52769553 | 0,60567835 | 0,79520052 |
| 57579_at | FAM135A | 0,12131818 | 7,30588892 | 0,52758516 | 0,60575305 | 0,7952444 |
| 6336_at | SCN10A | -0,05149724 | 4,24557691 | -0,52750252 | 0,60580899 | 0,79526363 |
| 347732_at | CATSPER3 | -0,06739745 | 3,98101473 | -0,52722473 | 0,60599703 | 0,79545627 |
| 1939_at | EIF2D | -0,11806365 | 8,00296536 | -0,52690037 | 0,60621663 | 0,7956903 |
| 114884_at | OSBPL10 | -0,05223578 | 6,36469548 | -0,52681313 | 0,6062757 | 0,79571362 |
| 23423_at | TMED3 | 0,04871291 | 7,06086782 | 0,52657709 | 0,60643554 | 0,79577387 |
| 4283_at | CXCL9 | -0,06953276 | 3,86541058 | -0,52657039 | 0,60644007 | 0,79577387 |
| 2915_at | GRM5 | -0,04773969 | 4,56758068 | -0,52656232 | 0,60644554 | 0,79577387 |
| 5834_at | PYGB | 0,07364532 | 8,77789035 | 0,52624893 | 0,60665779 | 0,79599816 |
| 25932_at | CLIC4 | 0,15980443 | 8,43339555 | 0,52611211 | 0,60675047 | 0,79606554 |
| 9328_at | GTF3C5 | -0,10445626 | 8,92225913 | -0,52520839 | 0,60736278 | 0,79676999 |
| 8456_at | FOXN1 | 0,06349002 | 4,79650709 | 0,52515457 | 0,60739926 | 0,79676999 |
| 132200_at | C3orf49 | -0,07869943 | 4,47102177 | -0,52509897 | 0,60743694 | 0,79676999 |
| 400831_at | C20orf202 | -0,04519598 | 5,40668063 | -0,5250636 | 0,60746091 | 0,79676999 |
| 100131564_a | LOC10013156 | 0,23249957 | 6,33367853 | 0,52490755 | 0,60756668 | 0,79676999 |
| 10618_at | TGOLN2 | 0,04740601 | 9,02770827 | 0,52486705 | 0,60759413 | 0,79676999 |
| 26018_at | LRIG1 | -0,08079799 | 4,16015137 | -0,52484137 | 0,60761154 | 0,79676999 |
| 56301_at | SLC7A10 | -0,05824358 | 5,65406497 | -0,52483141 | 0,60761829 | 0,79676999 |
| 222166_at | MTURN | 0,06728368 | 5,62708887 | 0,52471298 | 0,60769857 | 0,79680323 |
| 27132_at | CPNE7 | -0,05175458 | 5,95167707 | -0,52461576 | 0,60776448 | 0,79680323 |
| 1456_at | CSNK1G3 | -0,06842455 | 8,30368153 | -0,52457554 | 0,60779175 | 0,79680323 |
| 554202_at | MIR31HG | -0,04214405 | 4,33767444 | -0,52454994 | 0,6078091 | 0,79680323 |

| | | | | | | |
|---|---|---|---|---|---|---|
| 79949_at | PLEKHS1 | 0,24621837 | 3,75226032 | 0,52436872 | 0,60793197 | 0,79691008 |
| 186_at | AGTR2 | 0,0406205 | 3,61331756 | 0,52427542 | 0,60799523 | 0,79693877 |
| 6869_at | TACR1 | 0,04773363 | 3,88876675 | 0,52410448 | 0,60811115 | 0,79703647 |
| 10464_at | PIBF1 | 0,06179516 | 6,96060721 | 0,52376632 | 0,60834049 | 0,79722771 |
| 78994_at | PRR14 | 0,03548402 | 7,87623819 | 0,52374293 | 0,60835635 | 0,79722771 |
| 160065_at | PATE1 | -0,03357249 | 3,16319106 | -0,52370627 | 0,60838121 | 0,79722771 |
| 284348_at | LYPD5 | 0,13757255 | 6,36240872 | 0,52361933 | 0,60844018 | 0,79724684 |
| 79857_at | FLJ13224 | 0,0357697 | 4,36569341 | 0,52356271 | 0,60847859 | 0,79724684 |
| 79139_at | DERL1 | -0,08201695 | 9,88231961 | -0,52331568 | 0,60864617 | 0,79737161 |
| 22874_at | PLEKHA6 | 0,0883336 | 6,94930147 | 0,5233003 | 0,6086566 | 0,79737161 |
| 114294_at | LACTB | 0,0461927 | 7,19050112 | 0,52318486 | 0,60873492 | 0,79741998 |
| 2222_at | FDFT1 | -0,15580378 | 11,6469881 | -0,52300711 | 0,60885553 | 0,79752374 |
| 219287_at | AMER2 | 0,04738869 | 4,15631643 | 0,52272404 | 0,60904762 | 0,79770154 |
| 3547_at | IGSF1 | -0,05868738 | 3,88707591 | -0,52263644 | 0,60910707 | 0,79770154 |
| 8409_at | UXT | 0,0356036 | 10,0380859 | 0,52262401 | 0,6091155 | 0,79770154 |
| 101928333_a | LOC10192833 | -0,04470736 | 3,54191085 | -0,5224999 | 0,60919974 | 0,79775762 |
| 886_at | CCKAR | -0,05989729 | 4,7714003 | -0,52194094 | 0,60957918 | 0,79813442 |
| 9807_at | IP6K1 | 0,049953 | 7,66395268 | 0,52190509 | 0,60960352 | 0,79813442 |
| 3574_at | IL7 | -0,13845155 | 4,56385552 | -0,52189292 | 0,60961178 | 0,79813442 |
| 23519_at | ANP32D | 0,05296777 | 3,20935268 | 0,52183135 | 0,60965359 | 0,79813491 |
| 79939_at | SLC35E1 | 0,03790757 | 8,92917498 | 0,52176901 | 0,60969592 | 0,79813608 |
| 101928002_a | LOC10192800 | -0,03695323 | 2,91049716 | -0,52164189 | 0,60978223 | 0,79819483 |
| 148645_at | LINC00337 | -0,06642648 | 4,45063068 | -0,52135615 | 0,60997628 | 0,79839458 |
| 100129597_a | LINC00462 | -0,04324104 | 3,95156009 | -0,52124932 | 0,61004883 | 0,79843529 |
| 401562_at | LCNL1 | -0,06065587 | 5,75736585 | -0,52117742 | 0,61009767 | 0,79844497 |
| 101927275_a | NDUFV2-AS1 | -0,06825925 | 3,53068479 | -0,52093399 | 0,61026302 | 0,79860711 |
| 83697_at | SLC4A9 | -0,04086813 | 4,04833771 | -0,52082182 | 0,61033922 | 0,79865257 |
| 285848_at | PNPLA1 | 0,0605851 | 4,20159114 | 0,52061644 | 0,61047876 | 0,79874497 |
| 5168_at | ENPP2 | -0,03245821 | 3,2433181 | -0,52059582 | 0,61049277 | 0,79874497 |
| 1747_at | DLX3 | 0,07486162 | 4,8029951 | 0,52040077 | 0,6106253 | 0,7988641 |
| 1072_at | CFL1 | -0,10740485 | 11,7776043 | -0,52026988 | 0,61071424 | 0,79887346 |
| 151651_at | EFHB | 0,0322666 | 2,98661944 | 0,52026819 | 0,61071539 | 0,79887346 |
| 122416_at | ANKRD9 | 0,15686333 | 8,29887344 | 0,52012185 | 0,61081484 | 0,79887794 |
| 1956_at | EGFR | -0,02785938 | 7,50244834 | -0,51996165 | 0,61092372 | 0,79887794 |
| 28659_at | TRAV24 | -0,05334562 | 3,90269779 | -0,5199365 | 0,61094081 | 0,79887794 |
| 4110_at | MAGEA11 | -0,04465233 | 3,34450699 | -0,51993323 | 0,61094303 | 0,79887794 |
| 29933_at | GPR132 | -0,03968342 | 4,78570136 | -0,51993125 | 0,61094438 | 0,79887794 |
| 84516_at | DCTN5 | 0,07089786 | 7,53396443 | 0,51988582 | 0,61097526 | 0,79887794 |
| 4301_at | MLLT4 | 0,03566187 | 5,58988045 | 0,51981205 | 0,6110254 | 0,79887794 |
| 1363_at | CPE | 0,59430059 | 7,05783807 | 0,51971142 | 0,6110938 | 0,79887794 |
| 55188_at | RIC8B | -0,03366089 | 6,2785014 | -0,51970193 | 0,61110025 | 0,79887794 |
| 29935_at | RPA4 | 0,04305858 | 3,91057496 | 0,51962361 | 0,61115349 | 0,79887794 |
| 54596_at | L1TD1 | -0,03304068 | 3,48852525 | -0,51954653 | 0,61120589 | 0,79887794 |
| 55260_at | TMEM143 | 0,11125864 | 7,02278762 | 0,51953096 | 0,61121648 | 0,79887794 |
| 79161_at | TMEM243 | 0,06749909 | 8,90436833 | 0,51937135 | 0,61132499 | 0,79893458 |
| 340206_at | TREML3P | -0,05047635 | 4,58659287 | -0,51933284 | 0,61135118 | 0,79893458 |
| 100499177_a | THAP9-AS1 | -0,14910532 | 6,94608508 | -0,5192842 | 0,61138424 | 0,79893458 |
| 55322_at | C5orf22 | -0,10148565 | 8,03042636 | -0,51915704 | 0,61147071 | 0,79896983 |
| 5726_at | TAS2R38 | -0,06707566 | 4,50280581 | -0,51911936 | 0,61149633 | 0,79896983 |
| 84229_at | DRC7 | -0,06680026 | 4,77786299 | -0,51903436 | 0,61155413 | 0,79896983 |

| | | | | | | |
|---|---|---|---|---|---|---|
| 84720_at | PIGO | -0,03985524 | 7,73071302 | -0,51900054 | 0,61157713 | 0,79896983 |
| 90523_at | MLIP | -0,05440818 | 3,30738393 | -0,51887218 | 0,61166442 | 0,79902969 |
| 84309_at | NUDT16L1 | -0,03964359 | 7,486501 | -0,51875865 | 0,61174163 | 0,79906758 |
| 101926913_a | LOC10192691 | 0,03374857 | 3,04763592 | 0,51870754 | 0,61177639 | 0,79906758 |
| 51555_at | PEX5L | -0,02990785 | 3,29894347 | -0,51858889 | 0,61185709 | 0,79911832 |
| 441394_at | SUGT1P1 | 0,0527595 | 3,8722433 | 0,51841897 | 0,61197267 | 0,79911832 |
| 92558_at | CCDC64 | -0,05733014 | 5,91615421 | -0,51837864 | 0,61200011 | 0,79911832 |
| 9697_at | TRAM2 | -0,1025512 | 7,81266172 | -0,51834703 | 0,61202161 | 0,79911832 |
| 103352670_a | LINC01419 | 0,05532959 | 3,05721463 | 0,51834548 | 0,61202267 | 0,79911832 |
| 4515_at | MTCP1 | -0,14164962 | 5,24375242 | -0,51815201 | 0,61215428 | 0,799236 |
| 6506_at | SLC1A2 | 0,03138709 | 3,54255953 | 0,51796149 | 0,61228391 | 0,7993378 |
| 11034_at | DSTN | 0,03790004 | 11,4130666 | 0,51791543 | 0,61231525 | 0,7993378 |
| 100507506_a | LOC10050750 | -0,04593347 | 3,84547032 | -0,51769089 | 0,61246804 | 0,79941298 |
| 23057_at | NMNAT2 | -0,14211894 | 5,18738725 | -0,51766018 | 0,61248894 | 0,79941298 |
| 10019_at | SH2B3 | 0,09564364 | 7,30422914 | 0,51752006 | 0,6125843 | 0,79941298 |
| 169200_at | TMEM64 | -0,0856773 | 7,64495187 | -0,51747699 | 0,61261361 | 0,79941298 |
| 3766_at | KCNJ10 | 0,04293931 | 4,03381189 | 0,5174718 | 0,61261714 | 0,79941298 |
| 5077_at | PAX3 | -0,04162642 | 3,86542187 | -0,51742213 | 0,61265095 | 0,79941298 |
| 90853_at | SPOCD1 | -0,20632466 | 7,43414025 | -0,51737322 | 0,61268424 | 0,79941298 |
| 1670_at | DEFA5 | -0,05654812 | 4,46877031 | -0,51734296 | 0,61270483 | 0,79941298 |
| 66000_at | TMEM108 | -0,05249092 | 3,89562316 | -0,51708905 | 0,61287767 | 0,79958432 |
| 503519_at | LINC00929 | -0,03902164 | 4,4247772 | -0,51694417 | 0,6129763 | 0,79965884 |
| 203100_at | HTRA4 | -0,05595225 | 3,70097884 | -0,51643951 | 0,61331991 | 0,79991439 |
| 58526_at | MID1IP1 | -0,16303677 | 7,87023341 | -0,51643017 | 0,61332627 | 0,79991439 |
| 1441_at | CSF3R | -0,0529675 | 4,82211017 | -0,51640959 | 0,61334028 | 0,79991439 |
| 100996307_a | LIPE-AS1 | -0,04837029 | 4,32272714 | -0,51637291 | 0,61336526 | 0,79991439 |
| 101929423_a | MEF2C-AS1 | -0,05011601 | 4,4754938 | -0,51629171 | 0,61342056 | 0,79991439 |
| 338864_at | LINC00347 | -0,03502307 | 3,07080175 | -0,51628025 | 0,61342836 | 0,79991439 |
| 5106_at | PCK2 | -0,06341281 | 7,74628582 | -0,51622958 | 0,61346287 | 0,79991439 |
| 60598_at | KCNK15 | 0,07817573 | 4,59830452 | 0,51612898 | 0,61353139 | 0,79992018 |
| 51292_at | GMPR2 | 0,09107163 | 8,89344848 | 0,51604781 | 0,61358667 | 0,79992018 |
| 763_at | CA5A | -0,05417793 | 4,56310218 | -0,51604015 | 0,61359189 | 0,79992018 |
| 10518_at | CIB2 | 0,10332695 | 5,68457913 | 0,51577474 | 0,61377268 | 0,8000883 |
| 316_at | AOX1 | -0,05123401 | 3,7580158 | -0,51572888 | 0,61380392 | 0,8000883 |
| 5099_at | PCDH7 | -0,06291924 | 7,81369579 | -0,51562502 | 0,61387468 | 0,80012639 |
| 117195_at | MRGPRX3 | -0,05328259 | 3,63645408 | -0,51549891 | 0,61396059 | 0,80018423 |
| 3784_at | KCNQ1 | -0,07016024 | 6,50757478 | -0,51522624 | 0,61414638 | 0,80037222 |
| 28674_at | TRAV12-1 | -0,04990697 | 3,3812773 | -0,51513649 | 0,61420754 | 0,80039659 |
| 256435_at | ST6GALNAC3 | 0,03985463 | 3,62759855 | 0,51507686 | 0,61424818 | 0,80039659 |
| 388722_at | C1orf53 | -0,09515464 | 7,20710716 | -0,51501224 | 0,61429222 | 0,80039983 |
| 6642_at | SNX1 | 0,03863127 | 8,53043738 | 0,51482979 | 0,61441657 | 0,8005077 |
| 55760_at | DHX32 | -0,05884184 | 8,91720696 | -0,51472166 | 0,61449026 | 0,80054957 |
| 285671_at | RNF180 | 0,02624835 | 2,93505336 | 0,51463019 | 0,61455261 | 0,80057665 |
| 646113_at | LINC00643 | 0,03474778 | 3,11622212 | 0,51395503 | 0,61501291 | 0,80112211 |
| 2524_at | FUT2 | 0,07435167 | 5,03836013 | 0,51369895 | 0,61518754 | 0,80127083 |
| 79840_at | NHEJ1 | 0,05267386 | 7,42631776 | 0,51364897 | 0,61522163 | 0,80127083 |
| 51321_at | AMZ2 | -0,0409313 | 9,6752056 | -0,5135931 | 0,61525973 | 0,80127083 |
| 26231_at | LRRC29 | 0,08203252 | 5,5315328 | 0,51344557 | 0,61536036 | 0,80127083 |
| 643711_at | LOC643711 | -0,02941453 | 3,02005732 | -0,51343925 | 0,61536467 | 0,80127083 |
| 147007_at | TMEM199 | 0,03322725 | 7,93674076 | 0,51342166 | 0,61537666 | 0,80127083 |

| | | | | | | |
|---|---|---|---|---|---|---|
| 122664_at | TPPP2 | -0,03820787 | 3,70071809 | -0,51331284 | 0,61545089 | 0,80131332 |
| 100130331_a | LOC10013033 | -0,05153095 | 3,96204634 | -0,51302911 | 0,61564444 | 0,80151115 |
| 79750_at | ZNF385D | -0,03828145 | 3,29121463 | -0,51276945 | 0,61582161 | 0,80168761 |
| 10542_at | LAMTOR5 | -0,02904923 | 9,4606884 | -0,51239818 | 0,61607496 | 0,80188357 |
| 4969_at | OGN | 0,03652819 | 2,87610258 | 0,51236838 | 0,6160953 | 0,80188357 |
| 93233_at | CCDC114 | -0,04649981 | 4,60363344 | -0,51236586 | 0,61609702 | 0,80188357 |
| 6940_at | ZNF354A | -0,15911266 | 6,8430713 | -0,51222428 | 0,61619364 | 0,80192757 |
| 100128071_a | FAM229A | -0,05296058 | 6,35679118 | -0,51219433 | 0,61621409 | 0,80192757 |
| 54020_at | SLC37A1 | 0,14687319 | 8,46520697 | 0,51207098 | 0,61629828 | 0,80198296 |
| 28763_at | IGLVI-70 | -0,05618057 | 4,59946279 | -0,51198805 | 0,61635489 | 0,80200244 |
| 149018_at | LELP1 | 0,06101771 | 4,15541645 | 0,51183982 | 0,61645607 | 0,80206264 |
| 56674_at | TMEM9B | 0,05701361 | 9,0457656 | 0,51172517 | 0,61653435 | 0,80206264 |
| 140290_at | TCP10L | -0,04241539 | 4,25179776 | -0,51165168 | 0,61658452 | 0,80206264 |
| 283403_at | C12orf80 | -0,05356766 | 4,34249474 | -0,51165035 | 0,61658543 | 0,80206264 |
| 3054_at | HCFC1 | -0,03636358 | 7,69458968 | -0,51161532 | 0,61660934 | 0,80206264 |
| 6564_at | SLC15A1 | -0,10779617 | 4,80787593 | -0,51155064 | 0,6166535 | 0,80206593 |
| 101927066_a | LOC10192706 | -0,03764003 | 3,01831723 | -0,5114101 | 0,61674946 | 0,80213658 |
| 3550_at | IK | 0,04961026 | 9,70284084 | 0,51113232 | 0,61693915 | 0,80232911 |
| 406_at | ARNTL | -0,13251976 | 6,36587674 | -0,51084363 | 0,61713632 | 0,8025129 |
| 105377135_a | LOC10537713 | -0,14048115 | 5,73126917 | -0,51080068 | 0,61716566 | 0,8025129 |
| 401546_at | C9orf152 | 0,05277361 | 3,28902453 | 0,51069814 | 0,6172357 | 0,8025129 |
| 101928259_a | LOC10192825 | -0,05490288 | 3,13862585 | -0,51068143 | 0,61724711 | 0,8025129 |
| 65078_at | RTN4R | 0,11648762 | 6,61453193 | 0,51057673 | 0,61731863 | 0,80255172 |
| 84312_at | BRMS1L | 0,04860348 | 6,45462012 | 0,5103841 | 0,61745023 | 0,80266863 |
| 253128_at | LINC00612 | -0,05871907 | 3,38460464 | -0,51016654 | 0,61759888 | 0,80280769 |
| 84614_at | ZBTB37 | -0,03655195 | 4,55879168 | -0,50993555 | 0,61775672 | 0,80295868 |
| 10627_at | MYL12A | -0,05532252 | 8,48378722 | -0,5098682 | 0,61780274 | 0,80296432 |
| 56914_at | OTOR | -0,04522146 | 4,171426 | -0,50978711 | 0,61785816 | 0,80298217 |
| 11056_at | DDX52 | 0,04230772 | 7,88013379 | 0,50965366 | 0,61794936 | 0,80304652 |
| 727957_at | MROH1 | -0,13193642 | 6,77660461 | -0,50958189 | 0,61799842 | 0,80305609 |
| 389541_at | LAMTOR4 | -0,03676937 | 8,23801327 | -0,50912527 | 0,61831055 | 0,8034075 |
| 11136_at | SLC7A9 | 0,04197083 | 4,98896592 | 0,50897453 | 0,61841361 | 0,80344245 |
| 22977_at | AKR7A3 | -0,05346047 | 5,99589853 | -0,50896391 | 0,61842087 | 0,80344245 |
| 414243_at | LINC00595 | -0,05125788 | 4,20713131 | -0,50881871 | 0,61852015 | 0,80347235 |
| 10634_at | GAS2L1 | -0,03555325 | 7,85161755 | -0,50880826 | 0,6185273 | 0,80347235 |
| 7097_at | TLR2 | 0,2657948 | 6,55336801 | 0,50870528 | 0,61859772 | 0,80350963 |
| 84649_at | DGAT2 | -0,06062866 | 8,34470711 | -0,50854502 | 0,61870731 | 0,8035978 |
| 130_at | ADH6 | -0,04819097 | 3,48003503 | -0,50845895 | 0,61876617 | 0,80362006 |
| 5063_at | PAK3 | 0,0287676 | 3,41919822 | 0,50832474 | 0,61885796 | 0,80368509 |
| 140545_at | RNF32 | 0,02957222 | 4,4973836 | 0,50825289 | 0,61890711 | 0,80369473 |
| 57335_at | ZNF286A | 0,14276956 | 6,43371 | 0,50803521 | 0,619056 | 0,80379126 |
| 137362_at | GOT1L1 | -0,03321628 | 3,30538639 | -0,5080222 | 0,6190649 | 0,80379126 |
| 7163_at | TPD52 | 0,10943749 | 8,37054385 | 0,50787445 | 0,61916598 | 0,80386832 |
| 7483_at | WNT9A | 0,04121416 | 5,44417053 | 0,50767227 | 0,6193043 | 0,80399371 |
| 54714_at | CNGB3 | 0,06478832 | 3,41559 | 0,50757623 | 0,61937001 | 0,80402483 |
| 401320_at | LOC401320 | 0,08997124 | 5,68415137 | 0,5074553 | 0,61945276 | 0,80407806 |
| 9068_at | ANGPTL1 | -0,02773702 | 3,04623078 | -0,5072231 | 0,61961165 | 0,80423013 |
| 6658_at | SOX3 | -0,0505638 | 5,05678562 | -0,50715902 | 0,61965551 | 0,80423286 |
| 84868_at | HAVCR2 | -0,04797302 | 4,34800517 | -0,50699964 | 0,61976459 | 0,80432024 |
| 256380_at | SCML4 | -0,03448234 | 3,55437089 | -0,50688997 | 0,61983965 | 0,80436346 |

| | | | | | | |
|---|---|---|---|---|---|---|
| 904_at | CCNT1 | -0,03938575 | 8,17895408 | -0,50679198 | 0,61990672 | 0,80439631 |
| 26007_at | TKFC | -0,0819571 | 6,74957924 | -0,50659185 | 0,62004372 | 0,80448972 |
| 81931_at | ZNF93 | -0,16428452 | 6,42070257 | -0,50654164 | 0,6200781 | 0,80448972 |
| 101926915_a | LOC10192691 | -0,0702885 | 4,01160239 | -0,5065038 | 0,620104 | 0,80448972 |
| 219793_at | TBATA | -0,06427322 | 3,88413533 | -0,50632708 | 0,62022499 | 0,8045925 |
| 401190_at | RGS7BP | -0,04208863 | 3,38400586 | -0,50614619 | 0,62034885 | 0,80466036 |
| 2638_at | GC | -0,03841537 | 2,92914204 | -0,50612867 | 0,62036085 | 0,80466036 |
| 2196_at | FAT2 | 0,101769 | 7,22673426 | 0,50586845 | 0,62053905 | 0,8047845 |
| 57047_at | PLSCR2 | 0,03184607 | 3,31088098 | 0,5058669 | 0,62054011 | 0,8047845 |
| 3756_at | KCNH1 | -0,0450913 | 3,87878389 | -0,50572377 | 0,62063813 | 0,80485744 |
| 54863_at | TOR4A | -0,0617791 | 7,62558752 | -0,50561283 | 0,62071412 | 0,80490179 |
| 10975_at | UQCR11 | 0,03431645 | 8,53871236 | 0,50545645 | 0,62082124 | 0,8049865 |
| 283377_at | SPRYD4 | -0,03861287 | 7,87097054 | -0,50515209 | 0,62102974 | 0,8051776 |
| 101927417_a | LOC10192741 | -0,03752527 | 3,16442386 | -0,50511928 | 0,62105222 | 0,8051776 |
| 245936_at | DEFB123 | -0,06559555 | 4,21535826 | -0,50498722 | 0,6211427 | 0,80524072 |
| 28785_at | IGLV4-60 | 0,06397621 | 4,2984387 | 0,50468136 | 0,62135228 | 0,80545821 |
| 3128_at | HLA-DRB6 | 0,03980778 | 5,3215165 | 0,50443847 | 0,62151874 | 0,80557418 |
| 2796_at | GNRH1 | 0,04984456 | 3,75647568 | 0,50437016 | 0,62156556 | 0,80557418 |
| 9917_at | FAM20B | 0,05011196 | 8,8066858 | 0,50432812 | 0,62159437 | 0,80557418 |
| 24148_at | PRPF6 | 0,0734332 | 9,06615236 | 0,50415549 | 0,6217127 | 0,80557418 |
| 399947_at | C11orf87 | 0,02584286 | 3,2921749 | 0,50405826 | 0,62177935 | 0,80557418 |
| 56000_at | NXF3 | -0,04392836 | 3,42445168 | -0,50401651 | 0,62180797 | 0,80557418 |
| 5151_at | PDE8A | 0,06961572 | 8,60272383 | 0,50399001 | 0,62182613 | 0,80557418 |
| 23039_at | XPO7 | -0,13530409 | 8,46582538 | -0,50397526 | 0,62183625 | 0,80557418 |
| 57061_at | HYMAI | -0,05007705 | 3,66747734 | -0,50387665 | 0,62190385 | 0,80557418 |
| 3851_at | KRT4 | -0,76238488 | 8,11613725 | -0,50379931 | 0,62195688 | 0,80557418 |
| 23601_at | CLEC5A | 0,04957653 | 2,96429547 | 0,50379197 | 0,62196191 | 0,80557418 |
| 1742_at | DLG4 | 0,03848545 | 4,34540867 | 0,50371835 | 0,62201238 | 0,80557418 |
| 3743_at | KCNA7 | 0,03464738 | 4,06773187 | 0,50368977 | 0,62203197 | 0,80557418 |
| 57110_at | HRASLS | -0,03991421 | 3,53923173 | -0,50368899 | 0,62203251 | 0,80557418 |
| 388585_at | HES5 | -0,0420861 | 5,07238907 | -0,50363571 | 0,62206904 | 0,80557418 |
| 123970_at | C16orf78 | -0,05794036 | 3,73438576 | -0,50339324 | 0,62223531 | 0,80573533 |
| 101927884_a | LOC10192788 | -0,09767297 | 4,51069362 | -0,50329565 | 0,62230223 | 0,80576781 |
| 55669_at | MFN1 | -0,07803329 | 8,63363482 | -0,50307975 | 0,6224503 | 0,80585893 |
| 8496_at | PPFIBP1 | -0,04539535 | 6,43826694 | -0,50307104 | 0,62245627 | 0,80585893 |
| 219771_at | CCNY | 0,09927729 | 7,39177346 | 0,50298522 | 0,62251513 | 0,80587618 |
| 28776_at | IGLV7-43 | -0,062746 | 3,99191722 | -0,50292963 | 0,62255326 | 0,80587618 |
| 440699_at | LRRC52 | -0,05585771 | 4,89045175 | -0,50283347 | 0,62261922 | 0,80589765 |
| 83931_at | STK40 | -0,03773931 | 7,21242875 | -0,50260214 | 0,62277791 | 0,80589765 |
| 10361_at | NPM2 | -0,05059574 | 6,59165494 | -0,50259903 | 0,62278005 | 0,80589765 |
| 388951_at | TSPYL6 | -0,05076217 | 3,46123741 | -0,50244534 | 0,62288549 | 0,80589765 |
| 440556_at | LINC00982 | -0,04563748 | 4,43828609 | -0,50243467 | 0,62289281 | 0,80589765 |
| 132158_at | GLYCTK | -0,07097309 | 6,0968018 | -0,50241363 | 0,62290724 | 0,80589765 |
| 4248_at | MGAT3 | -0,09312012 | 5,51720977 | -0,5023681 | 0,62293848 | 0,80589765 |
| 6535_at | SLC6A8 | -0,18383459 | 9,13418859 | -0,50234111 | 0,622957 | 0,80589765 |
| 10401_at | PIAS3 | 0,10860803 | 8,37338686 | 0,50230757 | 0,62298001 | 0,80589765 |
| 53335_at | BCL11A | 0,15734858 | 4,38189532 | 0,50229562 | 0,62298821 | 0,80589765 |
| 55954_at | ZMAT5 | 0,03546431 | 7,09710058 | 0,50223289 | 0,62303126 | 0,80589922 |
| 7546_at | ZIC2 | -0,24456204 | 8,61327768 | -0,50211244 | 0,62311391 | 0,80595201 |
| 862_at | RUNX1T1 | -0,02375799 | 3,29668543 | -0,5016212 | 0,62345105 | 0,80630892 |

| | | | | | | |
|---|---|---|---|---|---|---|
| 1439_at | CSF2RB | 0,05340402 | 4,50450397 | 0,50158838 | 0,62347357 | 0,80630892 |
| 4703_at | NEB | -0,03875325 | 4,03741738 | -0,50125088 | 0,62370525 | 0,80655439 |
| 440560_at | PRAMEF11 | -0,0873697 | 3,45421969 | -0,50117143 | 0,6237598 | 0,80656055 |
| 8994_at | LIMD1 | -0,04432781 | 7,32205355 | -0,50101038 | 0,62387037 | 0,80656055 |
| 56052_at | ALG1 | 0,06273325 | 6,15432286 | 0,50096764 | 0,62389971 | 0,80656055 |
| 9908_at | G3BP2 | -0,15283199 | 8,71038295 | -0,50090805 | 0,62394063 | 0,80656055 |
| 100505776_a | LINC01541 | -0,03233054 | 2,91042412 | -0,50085283 | 0,62397855 | 0,80656055 |
| 2123_at | EVI2A | -0,03934146 | 2,93543973 | -0,50083284 | 0,62399228 | 0,80656055 |
| 161253_at | REM2 | -0,04124416 | 5,18908832 | -0,50081707 | 0,62400311 | 0,80656055 |
| 57127_at | RHBG | 0,06580252 | 7,31216005 | 0,50064648 | 0,62412026 | 0,80661652 |
| 9929_at | JOSD1 | 0,02961441 | 9,63030914 | 0,50063206 | 0,62413016 | 0,80661652 |
| 7325_at | UBE2E2 | -0,1588978 | 4,84441993 | -0,50037831 | 0,62430443 | 0,80677078 |
| 9900_at | SV2A | 0,27342907 | 5,26761686 | 0,50033631 | 0,62433328 | 0,80677078 |
| 117155_at | CATSPER2 | 0,12339769 | 4,33799142 | 0,50020848 | 0,62442108 | 0,80680596 |
| 10645_at | CAMKK2 | -0,03061332 | 7,14981923 | -0,50014462 | 0,62446494 | 0,80680596 |
| 102800317_a | LOC400927-C! | 0,09668153 | 4,43601468 | 0,50007857 | 0,62451032 | 0,80680596 |
| 254173_at | TTLL10 | -0,06322035 | 5,48279741 | -0,50005277 | 0,62452804 | 0,80680596 |
| 2000_at | ELF4 | 0,06416337 | 9,20894548 | 0,49968509 | 0,62478065 | 0,80696467 |
| 200539_at | ANKRD23 | -0,06667114 | 5,91927934 | -0,49965666 | 0,62480018 | 0,80696467 |
| 2658_at | GDF2 | 0,03750224 | 5,53950666 | 0,49963638 | 0,62481412 | 0,80696467 |
| 2986_at | GUCY2F | -0,04046623 | 3,88786111 | -0,49951683 | 0,62489626 | 0,80696467 |
| 2632_at | GBE1 | -0,11001977 | 9,38110478 | -0,49948605 | 0,62491741 | 0,80696467 |
| 57497_at | LRFN2 | 0,04441031 | 4,49462673 | 0,49948081 | 0,62492101 | 0,80696467 |
| 101927798_a | LOC10192779 | -0,03791807 | 3,16473115 | -0,49944716 | 0,62494414 | 0,80696467 |
| 1364_at | CLDN4 | -0,0505279 | 10,0476036 | -0,49930459 | 0,62504211 | 0,80703708 |
| 158724_at | FAM47A | -0,06272088 | 3,57184686 | -0,49899403 | 0,62525555 | 0,80723805 |
| 339451_at | KLHL17 | 0,03950754 | 6,22953871 | 0,49895617 | 0,62528158 | 0,80723805 |
| 91289_at | LMF2 | 0,04935223 | 8,80867379 | 0,49889346 | 0,62532468 | 0,8072396 |
| 23184_at | MESDC2 | -0,05469256 | 7,84474922 | -0,49873309 | 0,62543492 | 0,80729608 |
| 283629_at | TSSK4 | 0,05699486 | 3,88178443 | 0,49865715 | 0,62548712 | 0,80729608 |
| 285593_at | LOC285593 | -0,03971397 | 3,65705469 | -0,49864691 | 0,62549416 | 0,80729608 |
| 9353_at | SLIT2 | -0,03411955 | 3,40752922 | -0,49840992 | 0,62565709 | 0,80738961 |
| 6913_at | TBX15 | -0,04087524 | 4,06835628 | -0,49840516 | 0,62566036 | 0,80738961 |
| 84630_at | TTBK1 | -0,05918344 | 5,30434513 | -0,49835862 | 0,62569237 | 0,80738961 |
| 4223_at | MEOX2 | 0,0475154 | 3,51940425 | 0,49829435 | 0,62573656 | 0,80739255 |
| 80868_at | HCG4B | -0,04859978 | 4,31730281 | -0,49802559 | 0,62592136 | 0,80757691 |
| 22862_at | FNDC3A | 0,16678606 | 5,11780624 | 0,49779126 | 0,62608252 | 0,80773073 |
| 100506368_a | LOC10050636 | -0,05905873 | 4,61530483 | -0,49772164 | 0,6261304 | 0,80773841 |
| 56949_at | XAB2 | -0,08448905 | 6,7224457 | -0,49764645 | 0,62618212 | 0,80774679 |
| 283585_at | LOC283585 | -0,05523812 | 3,00660203 | -0,49759027 | 0,62622076 | 0,80774679 |
| 92104_at | TTC30A | -0,07546433 | 6,12119926 | -0,4973496 | 0,62638632 | 0,80790623 |
| 257194_at | NEGR1 | -0,0269492 | 3,03009629 | -0,49714349 | 0,62652811 | 0,80803502 |
| 3290_at | HSD11B1 | 0,03866625 | 3,60858271 | 0,49703177 | 0,62660498 | 0,80808005 |
| 384_at | ARG2 | -0,24569448 | 6,83947131 | -0,49681421 | 0,62675468 | 0,80816457 |
| 55843_at | ARHGAP15 | 0,03820212 | 3,52006216 | 0,49676474 | 0,62678872 | 0,80816457 |
| 25992_at | SNED1 | -0,04776934 | 4,82449587 | -0,49675361 | 0,62679638 | 0,80816457 |
| 284114_at | TMEM102 | 0,04575784 | 6,31381573 | 0,4963933 | 0,62704435 | 0,80843018 |
| 55170_at | PRMT6 | -0,12989084 | 7,80291344 | -0,49625688 | 0,62713824 | 0,80846462 |
| 57406_at | ABHD6 | 0,08848681 | 6,50173242 | 0,49623254 | 0,627155 | 0,80846462 |
| 7447_at | VSNL1 | -0,12381424 | 5,81202818 | -0,49599536 | 0,62731827 | 0,80857629 |

| | | | | | | |
|---|---|---|---|---|---|---|
| 163033_at | ZNF579 | -0,04243864 | 7,37620016 | -0,49597741 | 0,62733062 | 0,80857629 |
| 53828_at | FXYD4 | -0,08832432 | 4,15819283 | -0,49592376 | 0,62736756 | 0,80857629 |
| 138065_at | RNF183 | -0,07543503 | 4,57947296 | -0,49546243 | 0,6276852 | 0,80893155 |
| 89876_at | MAATS1 | -0,02659717 | 3,51401735 | -0,49517526 | 0,62788296 | 0,80913228 |
| 5672_at | PSG4 | 0,24440239 | 5,96590006 | 0,49492632 | 0,62805442 | 0,8092455 |
| 729582_at | DIRC3 | 0,04006609 | 4,10217559 | 0,49491227 | 0,6280641 | 0,8092455 |
| 100861552_a | LINC00558 | 0,03737453 | 3,09780994 | 0,49486473 | 0,62809684 | 0,8092455 |
| 79823_at | CAMKMT | -0,06267213 | 5,95908798 | -0,49469589 | 0,62821315 | 0,80934121 |
| 84838_at | ZNF496 | -0,04750724 | 6,34422668 | -0,49455505 | 0,62831017 | 0,80941208 |
| 4832_at | NME3 | 0,24380445 | 7,65976938 | 0,49423708 | 0,62852925 | 0,80958859 |
| 84503_at | ZNF527 | -0,03584367 | 6,03404015 | -0,49418229 | 0,628567 | 0,80958859 |
| 5232_at | PGK2 | -0,05787922 | 4,18920181 | -0,49417319 | 0,62857327 | 0,80958859 |
| 100506599_a | PPP1R26-AS1 | -0,08254291 | 4,75622934 | -0,4941087 | 0,62861771 | 0,8095917 |
| 816_at | CAMK2B | -0,03800016 | 5,42417893 | -0,4937381 | 0,62887311 | 0,80982592 |
| 112609_at | MRAP2 | -0,09119708 | 3,97100575 | -0,4937228 | 0,62888366 | 0,80982592 |
| 51693_at | TRAPPC2L | -0,08783712 | 6,94066424 | -0,49319725 | 0,62924593 | 0,81023826 |
| 187_at | APLNR | -0,06821119 | 4,49273171 | -0,49298198 | 0,62939435 | 0,81037244 |
| 2899_at | GRIK3 | -0,03571123 | 4,1151926 | -0,49292409 | 0,62943427 | 0,81037244 |
| 7138_at | TNNT1 | 0,10876812 | 5,51833305 | 0,49271573 | 0,62957794 | 0,81050323 |
| 100505881_a | MAGI2-AS3 | -0,03680905 | 3,43120154 | -0,49258811 | 0,62966595 | 0,81056236 |
| 7087_at | ICAM5 | -0,06670674 | 5,37218254 | -0,4924656 | 0,62975044 | 0,81061696 |
| 101927876_a | LOC10192787 | 0,06058912 | 3,82473031 | 0,49235855 | 0,62982427 | 0,81065782 |
| 55275_at | VPS53 | -0,02749294 | 6,1255494 | -0,49223272 | 0,62991107 | 0,81068437 |
| 91749_at | KIAA1919 | -0,07113844 | 6,31656804 | -0,49218632 | 0,62994307 | 0,81068437 |
| 4710_at | NDUFB4 | 0,06802431 | 7,39182824 | 0,4921456 | 0,62997116 | 0,81068437 |
| 57410_at | SCYL1 | -0,03996304 | 8,85034766 | -0,49200782 | 0,6300662 | 0,81075252 |
| 7251_at | TSG101 | 0,13835466 | 9,44437053 | 0,49180855 | 0,63020368 | 0,81087525 |
| 10788_at | IQGAP2 | 0,05162681 | 3,21931557 | 0,49171075 | 0,63027115 | 0,81087864 |
| 161829_at | EXD1 | 0,02790269 | 3,00416216 | 0,49168271 | 0,6302905 | 0,81087864 |
| 211_at | ALAS1 | -0,05489893 | 8,70426649 | -0,4913225 | 0,63053905 | 0,81104982 |
| 2671_at | GFER | -0,02776328 | 6,75206078 | -0,49131503 | 0,6305442 | 0,81104982 |
| 6862_at | T | -0,07291361 | 5,57424145 | -0,49130682 | 0,63054987 | 0,81104982 |
| 79738_at | BBS10 | -0,06285278 | 6,89821867 | -0,4908038 | 0,63089705 | 0,81131336 |
| 55745_at | AP5M1 | -0,05705863 | 7,40595905 | -0,4907435 | 0,63093867 | 0,81131336 |
| 87769_at | GGACT | 0,03445567 | 4,90773928 | 0,49071656 | 0,63095727 | 0,81131336 |
| 284346_at | ZNF575 | 0,06054876 | 6,13337455 | 0,49067785 | 0,63098399 | 0,81131336 |
| 114898_at | C1QTNF2 | 0,0637133 | 4,97520584 | 0,49067426 | 0,63098647 | 0,81131336 |
| 161198_at | CLEC14A | -0,06244081 | 4,60620323 | -0,49064385 | 0,63100746 | 0,81131336 |
| 131870_at | NUDT16 | 0,05883646 | 6,50165827 | 0,4905152 | 0,63109627 | 0,8113271 |
| 84709_at | MGARP | 0,07027752 | 4,91797415 | 0,49047064 | 0,63112704 | 0,8113271 |
| 337879_at | KRTAP8-1 | -0,04591336 | 4,61927798 | -0,49044066 | 0,63114774 | 0,8113271 |
| 148203_at | ZNF738 | 0,05209543 | 5,47726077 | 0,49038234 | 0,631188 | 0,8113271 |
| 760_at | CA2 | -0,22007263 | 5,92059663 | -0,49021631 | 0,63130264 | 0,8113271 |
| 64232_at | MS4A5 | -0,05848806 | 3,50018768 | -0,49015991 | 0,63134158 | 0,8113271 |
| 207063_at | DHRSX | 0,05028559 | 7,64626365 | 0,49014204 | 0,63135392 | 0,8113271 |
| 101927123_a | LOC10192712 | 0,04359335 | 3,73060044 | 0,49014034 | 0,63135509 | 0,8113271 |
| 152225_at | LOC152225 | -0,0395904 | 2,98357383 | -0,49006253 | 0,63140882 | 0,81134042 |
| 3823_at | KLRC3 | 0,0605539 | 3,15651386 | 0,49000333 | 0,6314497 | 0,81134042 |
| 100128893_a | GATA6-AS1 | -0,07344567 | 4,5995346 | -0,48979049 | 0,63159669 | 0,81147516 |
| 5273_at | SERPINB10 | -0,04369113 | 3,24241872 | -0,48963752 | 0,63170234 | 0,81151864 |

| | | | | | | |
|---|---|---|---|---|---|---|
| 112487_at | DTD2 | 0,04615163 | 7,55628822 | 0,48951714 | 0,63178548 | 0,81151864 |
| 6197_at | RPS6KA3 | 0,05602012 | 8,66762999 | 0,48950018 | 0,6317972 | 0,81151864 |
| 51138_at | COPS4 | -0,13086216 | 8,95191118 | -0,48949751 | 0,63179905 | 0,81151864 |
| 11278_at | KLF12 | 0,03324688 | 4,96638188 | 0,48938228 | 0,63187864 | 0,81156676 |
| 7132_at | TNFRSF1A | -0,05863863 | 10,3113088 | -0,48930202 | 0,63193408 | 0,81158385 |
| 6677_at | SPAM1 | -0,0464273 | 3,29772841 | -0,48923599 | 0,6319797 | 0,81158833 |
| 28661_at | TRAV22 | -0,04278534 | 3,51494553 | -0,48902055 | 0,63212854 | 0,81172535 |
| 9146_at | HGS | 0,03540229 | 8,82873461 | 0,48893359 | 0,63218862 | 0,8117484 |
| 56655_at | POLE4 | -0,16189901 | 8,24614146 | -0,48886693 | 0,63223468 | 0,81175342 |
| 58158_at | NEUROD4 | -0,05566862 | 4,29671495 | -0,4885756 | 0,63243599 | 0,81195778 |
| 10394_at | PRG3 | -0,05252302 | 5,24300384 | -0,48837972 | 0,63257137 | 0,81200831 |
| 11070_at | TMEM115 | 0,04715938 | 7,99262541 | 0,48835001 | 0,6325919 | 0,81200831 |
| 56603_at | CYP26B1 | -0,0807069 | 5,74695012 | -0,48833567 | 0,63260181 | 0,81200831 |
| 9796_at | PHYHIP | -0,04202723 | 4,99751524 | -0,4882469 | 0,63266317 | 0,81201354 |
| 92235_at | DUSP27 | 0,03268948 | 4,1416751 | 0,4882078 | 0,63269019 | 0,81201354 |
| 83894_at | TTC29 | -0,03178489 | 2,98665833 | -0,48808593 | 0,63277443 | 0,81206754 |
| 6489_at | ST8SIA1 | -0,031504 | 3,22424453 | -0,4877696 | 0,63299311 | 0,81228849 |
| 282618_at | IFNL1 | 0,06202526 | 4,66927465 | 0,48770351 | 0,6330388 | 0,81228849 |
| 2962_at | GTF2F1 | -0,03510585 | 9,00876705 | -0,48763862 | 0,63308366 | 0,81228849 |
| 29851_at | ICOS | -0,0407973 | 3,33562354 | -0,48759291 | 0,63311527 | 0,81228849 |
| 125981_at | ACER1 | -0,04904777 | 4,62634321 | -0,48710994 | 0,63344924 | 0,81265845 |
| 132332_at | TMEM155 | -0,0465786 | 3,57815909 | -0,4870539 | 0,63348799 | 0,81265845 |
| 54212_at | SNTG1 | -0,02706219 | 3,83677742 | -0,48662398 | 0,63378537 | 0,81298578 |
| 2203_at | FBP1 | -0,05109796 | 4,89481923 | -0,48653407 | 0,63384756 | 0,81301143 |
| 90525_at | SHF | -0,07835035 | 5,46507492 | -0,48624916 | 0,63404467 | 0,8132101 |
| 83983_at | TSSK6 | -0,03293999 | 5,30192198 | -0,48616766 | 0,63410107 | 0,81322828 |
| 339344_at | MYPOP | -0,04012683 | 6,55883092 | -0,48607982 | 0,63416184 | 0,81325208 |
| 440944_at | THUMPD3-AS | -0,1287152 | 6,6059657 | -0,48581084 | 0,63434798 | 0,81338678 |
| 1936_at | EEF1D | -0,10574301 | 10,2665386 | -0,485806 | 0,63435132 | 0,81338678 |
| 64343_at | AZI2 | 0,05520296 | 7,15596201 | 0,48509089 | 0,63484631 | 0,81396728 |
| 284551_at | LINC01226 | -0,03890618 | 4,65212952 | -0,48485983 | 0,63500628 | 0,8141182 |
| 9468_at | PCYT1B | -0,03622923 | 3,63525575 | -0,48473278 | 0,63509425 | 0,81417679 |
| 100506054_a | RNASEH1-AS1 | -0,13936525 | 6,80662215 | -0,4846636 | 0,63514215 | 0,81418402 |
| 148824_at | GCSAML-AS1 | 0,03085391 | 3,04939746 | 0,48449693 | 0,63525757 | 0,81427778 |
| 101929147_a | LOC10192914 | -0,05119989 | 3,63932853 | -0,48420639 | 0,63545878 | 0,8144815 |
| 375346_at | TMEM110 | 0,06226453 | 6,23927228 | 0,48402305 | 0,63558577 | 0,81448626 |
| 2047_at | EPHB1 | -0,03423795 | 4,17465644 | -0,48396985 | 0,63562262 | 0,81448626 |
| 23436_at | CELA3B | -0,05006737 | 5,0878962 | -0,48396574 | 0,63562547 | 0,81448626 |
| 5199_at | CFP | -0,05059446 | 4,2442762 | -0,48384808 | 0,63570697 | 0,81448626 |
| 9782_at | MATR3 | 0,03765772 | 8,88571149 | 0,48384273 | 0,63571068 | 0,81448626 |
| 1553_at | CYP2A13 | -0,04797711 | 5,07279165 | -0,48379069 | 0,63574673 | 0,81448626 |
| 2286_at | FKBP2 | 0,05114798 | 8,90557969 | 0,48377374 | 0,63575847 | 0,81448626 |
| 10691_at | GMEB1 | -0,06320243 | 7,27604793 | -0,48345767 | 0,63597745 | 0,81463722 |
| 1815_at | DRD4 | -0,06548794 | 5,51291042 | -0,48341418 | 0,63600759 | 0,81463722 |
| 101927342_a | LOC10192734 | -0,04499249 | 3,15812969 | -0,48339609 | 0,63602012 | 0,81463722 |
| 100507244_a | MAMDC2-AS1 | 0,03693499 | 3,61943215 | 0,48328481 | 0,63609723 | 0,81463722 |
| 5155_at | PDGFB | 0,08927069 | 5,68630123 | 0,48324086 | 0,63612768 | 0,81463722 |
| 10655_at | DMRT2 | 0,05549801 | 5,06308897 | 0,48321102 | 0,63614836 | 0,81463722 |
| 253639_at | ZNF620 | -0,08385873 | 4,08674014 | -0,48317642 | 0,63617234 | 0,81463722 |
| 8273_at | SLC10A3 | -0,08090569 | 8,64616875 | -0,48310639 | 0,63622087 | 0,81463984 |

| | | | | | | |
|---|---|---|---|---|---|---|
| 8559_at | PRPF18 | -0,07541368 | 7,13149245 | -0,48296366 | 0,63631978 | 0,81463984 |
| 51639_at | SF3B6 | -0,05381717 | 10,5084028 | -0,48293599 | 0,63633896 | 0,81463984 |
| 8433_at | UTF1 | -0,04973791 | 5,94050598 | -0,48292937 | 0,63634354 | 0,81463984 |
| 163_at | AP2B1 | -0,03708779 | 9,59522603 | -0,48276203 | 0,63645953 | 0,81473417 |
| 5064_at | PALM | 0,09000734 | 6,45787478 | 0,48260283 | 0,63656988 | 0,81478057 |
| 27160_at | INGX | -0,04627047 | 4,72010189 | -0,48254317 | 0,63661123 | 0,81478057 |
| 65125_at | WNK1 | 0,04232022 | 8,80638078 | 0,48247805 | 0,63665638 | 0,81478057 |
| 80201_at | HKDC1 | -0,09307418 | 5,82448169 | -0,48246566 | 0,63666496 | 0,81478057 |
| 286128_at | ZFP41 | -0,06752729 | 6,72865283 | -0,48198328 | 0,6369994 | 0,81509257 |
| 338339_at | CLEC4D | -0,04808017 | 3,75956327 | -0,48194351 | 0,63702698 | 0,81509257 |
| 387718_at | TEX36 | -0,03295843 | 3,16264487 | -0,48189965 | 0,63705739 | 0,81509257 |
| 84268_at | RPAIN | 0,02492675 | 6,63136452 | 0,48186991 | 0,63707801 | 0,81509257 |
| 29018_at | FOXN3-AS2 | -0,05948134 | 4,393464 | -0,48167976 | 0,63720988 | 0,81517255 |
| 113402_at | SFT2D1 | -0,06896665 | 9,90851913 | -0,48162397 | 0,63724857 | 0,81517255 |
| 5430_at | POLR2A | 0,03835219 | 6,75314006 | 0,4815967 | 0,63726748 | 0,81517255 |
| 84895_at | FAM73B | 0,05593076 | 7,09736783 | 0,48146015 | 0,63736219 | 0,81523956 |
| 170394_at | PWWP2B | 0,0437815 | 6,52061863 | 0,48132291 | 0,63745738 | 0,81530718 |
| 3983_at | ABLIM1 | -0,09963331 | 8,92372142 | -0,48102704 | 0,63766261 | 0,81551553 |
| 399473_at | SPRED3 | -0,05492791 | 6,43212898 | -0,4809062 | 0,63774645 | 0,8155602 |
| 9674_at | KIAA0040 | -0,04595184 | 7,34333232 | -0,48085464 | 0,63778222 | 0,8155602 |
| 6368_at | CCL23 | -0,03114449 | 3,16967602 | -0,48059451 | 0,63796271 | 0,81573685 |
| 3846_at | KRTAP5-9 | -0,05618086 | 5,36402322 | -0,4803046 | 0,63816388 | 0,81593992 |
| 10075_at | HUWE1 | -0,03548688 | 6,99169638 | -0,48019046 | 0,6382431 | 0,81598704 |
| 153469_at | JAKMIP2-AS1 | -0,06987486 | 3,7613518 | -0,48012873 | 0,63828594 | 0,81598766 |
| 64399_at | HHIP | -0,03078886 | 3,23255857 | -0,47999069 | 0,63838175 | 0,81605598 |
| 197_at | AHSG | -0,03866688 | 3,85117941 | -0,47989643 | 0,63844717 | 0,81608546 |
| 79068_at | FTO | 0,10642143 | 9,17560448 | 0,47972754 | 0,63856441 | 0,81618116 |
| 4956_at | ODF1 | -0,06349302 | 4,47173267 | -0,47939488 | 0,63879536 | 0,81642218 |
| 347730_at | LRRTM1 | -0,0397687 | 4,33821586 | -0,47886301 | 0,63916469 | 0,81684 |
| 57143_at | ADCK1 | 0,04988196 | 6,83244661 | 0,47864986 | 0,63931273 | 0,81688669 |
| 5649_at | RELN | -0,04572994 | 4,54162253 | -0,47848561 | 0,63942681 | 0,81688669 |
| 6100_at | RP9 | -0,10747435 | 7,25757151 | -0,47842467 | 0,63946914 | 0,81688669 |
| 100506195_a | LARGE-AS1 | -0,03027026 | 3,00160865 | -0,47837973 | 0,63950036 | 0,81688669 |
| 7047_at | TGM4 | -0,04679183 | 4,39223316 | -0,47837821 | 0,63950142 | 0,81688669 |
| 158800_at | RHOXF1 | -0,05613247 | 4,70089304 | -0,4783396 | 0,63952823 | 0,81688669 |
| 2764_at | GMFB | 0,12740418 | 8,87979556 | 0,47831873 | 0,63954273 | 0,81688669 |
| 388507_at | ZNF788 | 0,31816521 | 4,97959292 | 0,47831692 | 0,63954399 | 0,81688669 |
| 22936_at | ELL2 | -0,19880642 | 8,52697996 | -0,47826093 | 0,63958289 | 0,81688669 |
| 3292_at | HSD17B1 | -0,04886582 | 5,88995945 | -0,4780909 | 0,63970101 | 0,8169834 |
| 2152_at | F3 | -0,11689829 | 8,928334 | -0,47794149 | 0,63980482 | 0,81703299 |
| 90411_at | MCFD2 | -0,04751681 | 9,2935914 | -0,47785263 | 0,63986656 | 0,81703299 |
| 55554_at | KLK15 | 0,04288028 | 4,38875722 | 0,47785188 | 0,63986709 | 0,81703299 |
| 653308_at | ASAH2B | -0,0559329 | 5,78248035 | -0,47729066 | 0,6402571 | 0,81743239 |
| 101927759_a | LOC10192775 | -0,0525881 | 4,4366448 | -0,47727965 | 0,64026475 | 0,81743239 |
| 56270_at | WDR45B | -0,02987965 | 9,73615102 | -0,47711813 | 0,64037702 | 0,81752154 |
| 84448_at | ABLIM2 | -0,05030383 | 5,61110161 | -0,47691991 | 0,64051481 | 0,81764326 |
| 85294_at | KRTAP2-4 | -0,06567088 | 3,2627443 | -0,47662843 | 0,64071745 | 0,81784774 |
| 84735_at | CNDP1 | -0,05377483 | 3,93456578 | -0,47644035 | 0,64084822 | 0,81796046 |
| 2057_at | EPOR | -0,04706128 | 5,80590648 | -0,47636182 | 0,64090283 | 0,81797596 |
| 64806_at | IL25 | -0,05155091 | 4,69084631 | -0,47618106 | 0,64102852 | 0,81808218 |

| | | | | | | |
|---|---|---|---|---|---|---|
| 220388_at | CCDC89 | 0,13843424 | 4,08760249 | 0,47609178 | 0,64109062 | 0,81808532 |
| 6546_at | SLC8A1 | 0,04351735 | 3,63452727 | 0,4760554 | 0,64111592 | 0,81808532 |
| 54089_at | LINC00112 | -0,06155827 | 4,26117852 | -0,47567864 | 0,64137797 | 0,81836549 |
| 100506289_a | LOC10050628 | 0,03447343 | 3,35804878 | 0,47544508 | 0,64154044 | 0,81839809 |
| 80099_at | C7orf69 | 0,05922319 | 4,02232645 | 0,47543819 | 0,64154523 | 0,81839809 |
| 8459_at | TPST2 | 0,04178941 | 8,62448777 | 0,47540916 | 0,64156543 | 0,81839809 |
| 3781_at | KCNN2 | -0,03611183 | 4,00967276 | -0,47536102 | 0,64159892 | 0,81839809 |
| 101928167_a | LOC10192816 | 0,04291633 | 3,20279273 | 0,4752805 | 0,64165494 | 0,81839809 |
| 11156_at | PTP4A3 | -0,03709783 | 5,86231487 | -0,47527548 | 0,64165843 | 0,81839809 |
| 4440_at | MSI1 | -0,05210878 | 4,42061647 | -0,47468964 | 0,64206608 | 0,81886381 |
| 100144748_a | KLLN | -0,04546956 | 5,58640881 | -0,4744535 | 0,64223044 | 0,81896329 |
| 6532_at | SLC6A4 | -0,03739901 | 3,64766738 | -0,47443813 | 0,64224113 | 0,81896329 |
| 151306_at | GPBAR1 | -0,06534373 | 5,70990288 | -0,47439431 | 0,64227163 | 0,81896329 |
| 8674_at | VAMP4 | -0,11181527 | 6,42555434 | -0,47430987 | 0,64233041 | 0,81898403 |
| 10870_at | HCST | -0,05805754 | 5,37660065 | -0,47402071 | 0,6425317 | 0,81911973 |
| 79888_at | LPCAT1 | 0,07392646 | 9,36209922 | 0,47398083 | 0,64255946 | 0,81911973 |
| 91355_at | LRP5L | -0,0396852 | 6,30227747 | -0,47397372 | 0,64256441 | 0,81911973 |
| 268_at | AMH | -0,08466824 | 6,66388745 | -0,47388011 | 0,64262958 | 0,8191486 |
| 286467_at | FIRRE | -0,04475314 | 4,46074825 | -0,47378911 | 0,64269295 | 0,81917516 |
| 202151_at | RANBP3L | 0,03226047 | 3,07547105 | 0,47359997 | 0,64282464 | 0,8192401 |
| 6097_at | RORC | -0,05177687 | 4,94042551 | -0,47359377 | 0,64282896 | 0,8192401 |
| 150946_at | GAREML | -0,05752502 | 5,94001723 | -0,47333104 | 0,64301192 | 0,81941906 |
| 101927901_a | LOC10192790 | -0,03419887 | 3,46668312 | -0,4732602 | 0,64306125 | 0,81942772 |
| 100288123_a | LOC10028812 | -0,03515057 | 4,68569749 | -0,47319743 | 0,64310496 | 0,81942922 |
| 205_at | AK4 | -0,34274823 | 7,63996608 | -0,4730778 | 0,64318829 | 0,81945131 |
| 23499_at | MACF1 | 0,11571098 | 6,8771783 | 0,47304349 | 0,64321218 | 0,81945131 |
| 55336_at | FBXL8 | -0,05864804 | 5,07681392 | -0,4729893 | 0,64324993 | 0,81945131 |
| 2137_at | EXTL3 | -0,10153541 | 7,24202685 | -0,47260685 | 0,64351634 | 0,8197365 |
| 23381_at | SMG5 | -0,04663271 | 7,65196661 | -0,47253343 | 0,64356749 | 0,81973803 |
| 51705_at | EMCN | -0,03532183 | 3,0673543 | -0,47243158 | 0,64363845 | 0,81973803 |
| 54413_at | NLGN3 | -0,0302964 | 4,4670006 | -0,47242188 | 0,64364521 | 0,81973803 |
| 284697_at | BTBD8 | -0,04651937 | 3,58075628 | -0,47217475 | 0,64381741 | 0,81986049 |
| 100507257_a | MEG9 | -0,0459515 | 4,15981797 | -0,472111 | 0,64386183 | 0,81986049 |
| 2204_at | FCAR | -0,02951965 | 3,80252576 | -0,47209604 | 0,64387226 | 0,81986049 |
| 10654_at | PMVK | 0,04169521 | 7,86966189 | 0,47203956 | 0,64391161 | 0,81986049 |
| 340591_at | CA5BP1 | 0,04375219 | 7,84807033 | 0,47191971 | 0,64399513 | 0,81991264 |
| 101927051_a | LOC10192705 | -0,02341407 | 5,17067485 | -0,47164705 | 0,64418516 | 0,82010038 |
| 83857_at | TMTC1 | 0,15431213 | 4,99726892 | 0,47146478 | 0,64431221 | 0,82020792 |
| 9197_at | SLC33A1 | 0,05292837 | 8,53784356 | 0,47127044 | 0,64444769 | 0,82026455 |
| 54205_at | CYCS | -0,0978851 | 6,24540235 | -0,47116176 | 0,64452345 | 0,82026455 |
| 116841_at | SNAP47 | -0,07432743 | 8,42864883 | -0,47114376 | 0,644536 | 0,82026455 |
| 84628_at | NTNG2 | -0,04410201 | 4,54576153 | -0,47108844 | 0,64457457 | 0,82026455 |
| 80199_at | FUZ | 0,07124268 | 6,70232921 | 0,47104236 | 0,6446067 | 0,82026455 |
| 100874021_a | MAGI2-AS2 | -0,05494963 | 3,19896511 | -0,47094033 | 0,64467783 | 0,82026455 |
| 127002_at | ATXN7L2 | 0,04793013 | 4,91097007 | 0,47083843 | 0,64474888 | 0,82026455 |
| 23108_at | RAP1GAP2 | -0,07703084 | 5,98759124 | -0,47083656 | 0,64475019 | 0,82026455 |
| 649446_at | DLGAP1-AS1 | -0,12182707 | 6,60346246 | -0,47076535 | 0,64479984 | 0,82026455 |
| 57693_at | ZNF317 | -0,05607182 | 7,85094339 | -0,47051622 | 0,64497357 | 0,82026455 |
| 55906_at | ZC4H2 | 0,09888381 | 5,46357802 | 0,47048905 | 0,64499252 | 0,82026455 |
| 141_at | ADPRH | -0,02503378 | 3,94758853 | -0,47045865 | 0,64501372 | 0,82026455 |

| | | | | | | |
|---|---|---|---|---|---|---|
| 23768_at | FLRT2 | 0,21883599 | 5,78132005 | 0,47039681 | 0,64505685 | 0,82026455 |
| 26258_at | BLOC1S6 | 0,08407152 | 8,26775039 | 0,47027994 | 0,64513836 | 0,82026455 |
| 4744_at | NEFH | 0,03487029 | 4,01728602 | 0,47024526 | 0,64516255 | 0,82026455 |
| 83878_at | USHBP1 | 0,03719877 | 5,26642961 | 0,47018197 | 0,64520669 | 0,82026455 |
| 8896_at | BUD31 | -0,07176718 | 9,34036531 | -0,47016523 | 0,64521837 | 0,82026455 |
| 3362_at | HTR6 | -0,06336791 | 4,62095954 | -0,47015276 | 0,64522707 | 0,82026455 |
| 10408_at | MYCNOS | -0,03680834 | 4,27949619 | -0,47012828 | 0,64524415 | 0,82026455 |
| 55124_at | PIWIL2 | -0,03658249 | 3,74550909 | -0,46999896 | 0,64533435 | 0,82026455 |
| 3340_at | NDST1 | 0,03375694 | 7,80667193 | 0,46996906 | 0,64535521 | 0,82026455 |
| 2359_at | FPR3 | -0,04498941 | 3,80169919 | -0,46996472 | 0,64535824 | 0,82026455 |
| 28642_at | TRAV39 | 0,03560699 | 3,44050632 | 0,46989587 | 0,64540627 | 0,82026455 |
| 5443_at | POMC | -0,04817893 | 5,12747847 | -0,4698471 | 0,64544029 | 0,82026455 |
| 80714_at | PBX4 | -0,21450365 | 6,1833995 | -0,46981892 | 0,64545995 | 0,82026455 |
| 440117_at | LOC440117 | -0,04151575 | 3,280341 | -0,4697797 | 0,64548731 | 0,82026455 |
| 253573_at | LOC253573 | 0,04423683 | 3,06088023 | 0,46975229 | 0,64550643 | 0,82026455 |
| 203430_at | ZCCHC5 | -0,04161098 | 4,1273476 | -0,46959108 | 0,6456189 | 0,82030963 |
| 374654_at | KIF7 | 0,06107132 | 6,74465829 | 0,46949512 | 0,64568586 | 0,82030963 |
| 6863_at | TAC1 | 0,06636125 | 3,29331874 | 0,46943867 | 0,64572525 | 0,82030963 |
| 80021_at | TMEM62 | -0,09051177 | 6,25365695 | -0,46942568 | 0,64573431 | 0,82030963 |
| 286451_at | YIPF6 | 0,05411741 | 7,65171958 | 0,46931664 | 0,64581039 | 0,82030963 |
| 858_at | CAV2 | 0,29279833 | 9,90606424 | 0,46928624 | 0,64583161 | 0,82030963 |
| 466_at | ATF1 | 0,03176451 | 8,45909098 | 0,46927422 | 0,64584 | 0,82030963 |
| 3044_at | HBBP1 | -0,0330061 | 3,910006 | -0,46890433 | 0,64609814 | 0,8205834 |
| 56267_at | CCBL2 | 0,08159485 | 8,31173183 | 0,46853058 | 0,64635902 | 0,82082128 |
| 7363_at | UGT2B4 | 0,03315338 | 2,67812321 | 0,46851391 | 0,64637066 | 0,82082128 |
| 10278_at | EFS | 0,31828992 | 7,08445718 | 0,46842339 | 0,64643385 | 0,82084741 |
| 120065_at | OR5P2 | -0,07957498 | 3,96571377 | -0,46814923 | 0,64662526 | 0,82095179 |
| 137209_at | ZNF572 | -0,05196557 | 3,91511486 | -0,46812724 | 0,64664061 | 0,82095179 |
| 5678_at | PSG9 | 0,05381325 | 5,24273178 | 0,46810231 | 0,64665801 | 0,82095179 |
| 80145_at | THOC7 | -0,09038272 | 9,13155781 | -0,46806149 | 0,64668652 | 0,82095179 |
| 23615_at | PYY2 | 0,05289129 | 5,12152244 | 0,46794778 | 0,64676592 | 0,82096089 |
| 10111_at | RAD50 | -0,10267348 | 7,92506257 | -0,46792915 | 0,64677893 | 0,82096089 |
| 23671_at | TMEFF2 | 0,03397385 | 3,22690292 | 0,46740079 | 0,64714792 | 0,82137513 |
| 145942_at | TMCO5A | 0,02421717 | 3,15513836 | 0,46733007 | 0,64719732 | 0,82138371 |
| 11063_at | SOX30 | 0,03977437 | 3,38699728 | 0,467098 | 0,64735942 | 0,82153532 |
| 79603_at | CERS4 | 0,05136584 | 5,30429404 | 0,4670237 | 0,64741132 | 0,82154706 |
| 51322_at | WAC | -0,05102139 | 8,2897496 | -0,46685624 | 0,64752832 | 0,82163227 |
| 221656_at | KDM1B | 0,07746861 | 6,42066399 | 0,46677084 | 0,64758798 | 0,82163227 |
| 54857_at | GDPD2 | -0,06821264 | 5,58139354 | -0,46665997 | 0,64766545 | 0,82163227 |
| 3552_at | IL1A | -0,5119796 | 7,56934358 | -0,46658048 | 0,64772099 | 0,82163227 |
| 54331_at | GNG2 | -0,02983678 | 3,98490281 | -0,46641558 | 0,64783622 | 0,82163227 |
| 10695_at | CNPY3 | 0,07537079 | 8,14426195 | 0,46639003 | 0,64785408 | 0,82163227 |
| 102724312_a | LOC10272431 | -0,05721611 | 5,45270641 | -0,4663453 | 0,64788533 | 0,82163227 |
| 7048_at | TGFBR2 | 0,15133078 | 8,98641024 | 0,46628827 | 0,64792519 | 0,82163227 |
| 85313_at | PPIL4 | -0,07791375 | 7,66325044 | -0,46627345 | 0,64793554 | 0,82163227 |
| 51454_at | GULP1 | 0,16975338 | 4,12202927 | 0,46626033 | 0,64794471 | 0,82163227 |
| 100134229_a | JHDM1D-AS1 | 0,04466366 | 5,88419146 | 0,46622409 | 0,64797004 | 0,82163227 |
| 11154_at | AP4S1 | 0,026504 | 5,46003972 | 0,46615397 | 0,64801904 | 0,82163227 |
| 130355_at | C2orf76 | -0,05248282 | 6,68353473 | -0,46613405 | 0,64803297 | 0,82163227 |
| 440704_at | LOC440704 | -0,03527878 | 3,23583703 | -0,46591154 | 0,64818848 | 0,82173334 |

| | | | | | | |
|---|---|---|---|---|---|---|
| 121952_at | METTL21EP | -0,04736767 | 3,58338303 | -0,46589793 | 0,648198 | 0,82173334 |
| 23262_at | PPIP5K2 | 0,10139088 | 8,44883652 | 0,46566815 | 0,64835862 | 0,82187866 |
| 254427_at | PROSER2 | -0,09583054 | 7,69997997 | -0,46560308 | 0,64840411 | 0,82187866 |
| 84272_at | YIPF4 | 0,09867568 | 6,58020473 | 0,46555084 | 0,64844063 | 0,82187866 |
| 160419_at | C12orf50 | -0,0420892 | 3,0175334 | -0,46546258 | 0,64850233 | 0,82189421 |
| 2520_at | GAST | -0,04789585 | 5,16213581 | -0,46541123 | 0,64853823 | 0,82189421 |
| 1068_at | CETN1 | -0,05596082 | 4,65155886 | -0,46526507 | 0,64864042 | 0,82196964 |
| 90506_at | LRRC46 | -0,0420574 | 4,71514827 | -0,46507953 | 0,64877016 | 0,82207996 |
| 230_at | ALDOC | -0,27491691 | 8,03067606 | -0,46467252 | 0,6490548 | 0,82238654 |
| 132160_at | PPM1M | 0,04420865 | 5,44093757 | 0,46435539 | 0,64927661 | 0,82261348 |
| 161753_at | ODF3L1 | -0,06036406 | 4,38389554 | -0,46426044 | 0,64934303 | 0,8226235 |
| 146857_at | SLFN13 | 0,20860025 | 4,07181858 | 0,46416458 | 0,64941009 | 0,8226235 |
| 644538_at | SMIM10 | -0,05181908 | 3,9167924 | -0,46416095 | 0,64941263 | 0,8226235 |
| 23283_at | CSTF2T | 0,03972567 | 7,74788601 | 0,46380288 | 0,64966315 | 0,82288672 |
| 100505811_a | HNCAT21 | 0,03303869 | 3,38724429 | 0,46373102 | 0,64971343 | 0,8228963 |
| 1129_at | CHRM2 | -0,05914099 | 3,75568693 | -0,46361238 | 0,64979645 | 0,82294734 |
| 29974_at | A1CF | -0,02736465 | 3,42282668 | -0,46348919 | 0,64988266 | 0,82300241 |
| 246213_at | SLC17A8 | 0,03514132 | 3,0467502 | 0,46334008 | 0,649987 | 0,82306515 |
| 1438_at | CSF2RA | -0,04456637 | 4,24922428 | -0,46329628 | 0,65001766 | 0,82306515 |
| 27023_at | FOXB1 | -0,04870166 | 4,49224916 | -0,46318163 | 0,6500979 | 0,82309099 |
| 84870_at | RSPO3 | 0,03669444 | 3,53993117 | 0,46310332 | 0,65015272 | 0,82309099 |
| 3097_at | HIVEP2 | 0,07609197 | 7,55121042 | 0,46308398 | 0,65016625 | 0,82309099 |
| 4922_at | NTS | -0,04785852 | 3,54331353 | -0,4630179 | 0,65021251 | 0,82309545 |
| 56882_at | CDC42SE1 | 0,04424973 | 9,18604632 | 0,4628254 | 0,65034725 | 0,82313446 |
| 23333_at | DPY19L1 | -0,06716204 | 8,51729089 | -0,46282267 | 0,65034917 | 0,82313446 |
| 102724386_a | LINC01393 | 0,03585479 | 3,41516574 | 0,46279074 | 0,65037152 | 0,82313446 |
| 51177_at | PLEKHO1 | 0,18350491 | 5,88129547 | 0,46242195 | 0,65062971 | 0,82340714 |
| 84878_at | ZBTB45 | 0,02672096 | 5,96387188 | 0,46226419 | 0,65074017 | 0,82349283 |
| 2169_at | FABP2 | -0,03497024 | 3,2699837 | -0,46220108 | 0,65078436 | 0,82349466 |
| 5545_at | PRB4 | -0,04179456 | 4,3971364 | -0,46202349 | 0,65090873 | 0,8235577 |
| 2689_at | GH2 | -0,05345261 | 3,65941973 | -0,46200783 | 0,65091969 | 0,8235577 |
| 55179_at | FAIM | -0,19588909 | 7,28587606 | -0,46179119 | 0,65107142 | 0,82369557 |
| 2849_at | GPR26 | -0,03630881 | 3,62744393 | -0,46171284 | 0,65112629 | 0,82371089 |
| 55316_at | RSAD1 | 0,06869332 | 7,56512126 | 0,46160276 | 0,65120339 | 0,82372317 |
| 346606_at | MOGAT3 | -0,05808911 | 4,16364638 | -0,46156068 | 0,65123287 | 0,82372317 |
| 27199_at | OXGR1 | -0,07555633 | 5,76740622 | -0,46151584 | 0,65126428 | 0,82372317 |
| 147183_at | KRT25 | 0,05057919 | 3,49396501 | 0,46142744 | 0,6513262 | 0,8237474 |
| 101928120_a | LOC10192812 | -0,04884318 | 5,88902658 | -0,46128108 | 0,65142873 | 0,82382298 |
| 6345_at | SRL | 0,04399203 | 4,86119658 | 0,46107397 | 0,65157383 | 0,82395239 |
| 253724_at | TTC41P | -0,03944236 | 3,61982739 | -0,46082356 | 0,65174929 | 0,82408974 |
| 63948_at | DMRTB1 | -0,05742849 | 5,4279371 | -0,46079683 | 0,65176801 | 0,82408974 |
| 27439_at | CECR6 | -0,06654808 | 6,10786866 | -0,4606465 | 0,65187336 | 0,82416884 |
| 7652_at | ZNF99 | -0,0234225 | 2,71165305 | -0,46053535 | 0,65195125 | 0,82417404 |
| 80157_at | CWH43 | -0,0293539 | 2,94245009 | -0,46051854 | 0,65196304 | 0,82417404 |
| 9016_at | SLC25A14 | -0,02979775 | 6,72950866 | -0,46036452 | 0,65207098 | 0,82421952 |
| 84935_at | MEDAG | -0,04534669 | 4,23758155 | -0,46027107 | 0,65213648 | 0,82421952 |
| 761_at | CA3 | 0,03314265 | 3,70785425 | 0,46015003 | 0,65222132 | 0,82421952 |
| 8348_at | HIST1H2BO | -0,06180866 | 4,61744774 | -0,46011453 | 0,6522462 | 0,82421952 |
| 84141_at | EVA1A | 0,10287548 | 6,07242686 | 0,46010729 | 0,65225127 | 0,82421952 |
| 80335_at | WDR82 | -0,08897719 | 9,80304052 | -0,46006717 | 0,6522794 | 0,82421952 |

| | | | | | | |
|---|---|---|---|---|---|---|
| 5153_at | PDE1B | -0,0442328 | 5,26733066 | -0,46003988 | 0,65229853 | 0,82421952 |
| 114822_at | RHPN1 | -0,0600021 | 5,97544156 | -0,45989568 | 0,65239961 | 0,82423715 |
| 6376_at | CX3CL1 | -0,0452917 | 5,19095089 | -0,45978725 | 0,65247562 | 0,82423715 |
| 148738_at | HFE2 | -0,03600894 | 3,60575189 | -0,45977445 | 0,6524846 | 0,82423715 |
| 23293_at | SMG6 | 0,03195641 | 6,21431173 | 0,45975257 | 0,65249994 | 0,82423715 |
| 10254_at | STAM2 | 0,05971321 | 7,9116225 | 0,45965846 | 0,65256592 | 0,82423715 |
| 339541_at | C1orf228 | 0,04007397 | 3,89148146 | 0,45965376 | 0,65256921 | 0,82423715 |
| 2792_at | GNGT1 | 0,1324432 | 3,42185497 | 0,45927414 | 0,6528354 | 0,82449551 |
| 51327_at | AHSP | -0,04131176 | 4,39869115 | -0,45918385 | 0,65289871 | 0,82449551 |
| 55100_at | WDR70 | 0,05983739 | 7,59393706 | 0,45914937 | 0,65292289 | 0,82449551 |
| 58515_at | SELK | -0,11708771 | 9,13470722 | -0,45911788 | 0,65294497 | 0,82449551 |
| 100507534_a | LOC10050753 | -0,04342296 | 3,37910666 | -0,45896104 | 0,65305497 | 0,82458036 |
| 123879_at | DCUN1D3 | -0,05632905 | 7,22407862 | -0,45876298 | 0,65319389 | 0,82463179 |
| 23380_at | SRGAP2 | 0,03246359 | 7,42459342 | 0,45872597 | 0,65321985 | 0,82463179 |
| 285326_at | LINC00692 | -0,02834659 | 2,80694409 | -0,45871985 | 0,65322414 | 0,82463179 |
| 474344_at | GIMAP6 | -0,03638697 | 3,67305017 | -0,45825003 | 0,65355372 | 0,82494862 |
| 6714_at | SRC | 0,02329606 | 7,10584667 | 0,45823999 | 0,65356076 | 0,82494862 |
| 29991_at | OBP2A | 0,047747 | 4,3478112 | 0,45816371 | 0,65361428 | 0,82496213 |
| 943_at | TNFRSF8 | -0,04373509 | 5,66157653 | -0,45760121 | 0,65400899 | 0,82540623 |
| 79094_at | CHAC1 | 0,0755904 | 6,44845797 | 0,45749565 | 0,65408308 | 0,82544564 |
| 11213_at | IRAK3 | 0,24176285 | 4,25211149 | 0,45713473 | 0,65433641 | 0,82564254 |
| 613126_at | CYP51A1-AS1 | -0,0375009 | 3,08379575 | -0,45709446 | 0,65436468 | 0,82564254 |
| 6622_at | SNCA | 0,04038986 | 5,20910642 | 0,45709018 | 0,65436768 | 0,82564254 |
| 200186_at | CRTC2 | -0,03092848 | 7,63806978 | -0,45697679 | 0,65444728 | 0,82568889 |
| 89796_at | NAV1 | 0,08623326 | 6,76679961 | 0,45671999 | 0,65462757 | 0,82586226 |
| 101927460_a | LOC10192746 | -0,05654885 | 3,48471236 | -0,45663555 | 0,65468686 | 0,82588297 |
| 2098_at | ESD | -0,10979468 | 8,6018073 | -0,45651743 | 0,65476979 | 0,82588449 |
| 9897_at | KIAA0196 | 0,07897587 | 9,04065406 | 0,45651171 | 0,65477381 | 0,82588449 |
| 83953_at | FCAMR | -0,03835238 | 3,63129989 | -0,45642535 | 0,65483445 | 0,8259069 |
| 5216_at | PFN1 | -0,13099414 | 10,0115019 | -0,45610524 | 0,65505925 | 0,82613633 |
| 786_at | CACNG1 | -0,05315649 | 5,43092159 | -0,45603285 | 0,65511009 | 0,82614636 |
| 78999_at | LRFN4 | -0,04897799 | 6,99280853 | -0,45561825 | 0,65540131 | 0,8264419 |
| 8721_at | EDF1 | 0,02981308 | 10,4474377 | 0,45549044 | 0,6554911 | 0,8264419 |
| 126206_at | NLRP5 | 0,03502131 | 4,07359625 | 0,45527411 | 0,65564308 | 0,8264419 |
| 83590_at | TMUB1 | 0,03212463 | 8,17093018 | 0,45526212 | 0,65565151 | 0,8264419 |
| 170679_at | PSORS1C1 | -0,05186524 | 4,56601887 | -0,45525757 | 0,6556547 | 0,8264419 |
| 85329_at | LGALS12 | -0,04834353 | 4,45085568 | -0,45523054 | 0,6556737 | 0,8264419 |
| 3687_at | ITGAX | -0,05359904 | 4,86328028 | -0,45521332 | 0,65568579 | 0,8264419 |
| 55278_at | QRSL1 | -0,10025271 | 6,68903331 | -0,45521065 | 0,65568767 | 0,8264419 |
| 22847_at | ZNF507 | -0,03883392 | 7,06579433 | -0,45502845 | 0,65581569 | 0,82652321 |
| 100268168_a | LOC10026816 | -0,04291401 | 4,23190908 | -0,45499671 | 0,655838 | 0,82652321 |
| 10058_at | ABCB6 | 0,22963432 | 7,71726379 | 0,4547957 | 0,65597925 | 0,82664714 |
| 55064_at | SPATA6L | -0,03086853 | 3,57953897 | -0,45406764 | 0,65649098 | 0,82709914 |
| 1807_at | DPYS | -0,04395449 | 3,52459125 | -0,45402507 | 0,65652091 | 0,82709914 |
| 285000_at | LOC285000 | -0,05193759 | 4,02294015 | -0,45398142 | 0,6565516 | 0,82709914 |
| 79720_at | VPS37B | -0,04708133 | 7,75298609 | -0,45394029 | 0,65658051 | 0,82709914 |
| 1961_at | EGR4 | -0,04499951 | 5,36261061 | -0,45386872 | 0,65663083 | 0,82709914 |
| 439933_at | LOC439933 | 0,03253455 | 3,04843075 | 0,45384648 | 0,65664647 | 0,82709914 |
| 10180_at | RBM6 | 0,04442092 | 7,6084803 | 0,45381822 | 0,65666633 | 0,82709914 |
| 282991_at | BLOC1S2 | -0,0569179 | 9,7230563 | -0,45377518 | 0,6566966 | 0,82709914 |

| | | | | | | |
|---|---|---|---|---|---|---|
| 101927821_a | LINC01425 | -0,04338694 | 4,26369387 | -0,4537357 | 0,65672436 | 0,82709914 |
| 51561_at | IL23A | 0,08504292 | 5,42511526 | 0,45363463 | 0,65679542 | 0,82713456 |
| 127670_at | TEDDM1 | -0,03654434 | 3,44204573 | -0,45341659 | 0,65694874 | 0,82727356 |
| 1347_at | COX7A2 | 0,03029627 | 10,2798411 | 0,45322357 | 0,65708448 | 0,8273904 |
| 954_at | ENTPD2 | -0,05315963 | 5,76774341 | -0,45314791 | 0,6571377 | 0,82740333 |
| 347475_at | CCDC160 | 0,03438837 | 3,32034771 | 0,45306742 | 0,65719431 | 0,82742052 |
| 5308_at | PITX2 | 0,02883504 | 3,88991623 | 0,4528742 | 0,65733021 | 0,8275168 |
| 2647_at | BLOC1S1 | 0,11398488 | 8,65428931 | 0,4527712 | 0,65740266 | 0,8275168 |
| 6786_at | STIM1 | 0,02679226 | 6,82203424 | 0,45277011 | 0,65740343 | 0,8275168 |
| 64096_at | GFRA4 | -0,05162892 | 4,81652301 | -0,45268944 | 0,65746018 | 0,8275168 |
| 399726_at | CASC10 | 0,06650171 | 5,91907337 | 0,45259684 | 0,65752532 | 0,8275168 |
| 387640_at | SKIDA1 | 0,08664349 | 4,36342197 | 0,45259227 | 0,65752853 | 0,8275168 |
| 26046_at | LTN1 | -0,05782318 | 7,15590745 | -0,45213841 | 0,65784786 | 0,82783387 |
| 100506757_a | LINC00629 | 0,04077839 | 2,88502168 | 0,45211203 | 0,65786642 | 0,82783387 |
| 9414_at | TJP2 | -0,03998025 | 7,13862007 | -0,45180893 | 0,65807971 | 0,82804817 |
| 41_at | ASIC1 | -0,06831659 | 5,52131331 | -0,45123431 | 0,65848416 | 0,82850296 |
| 3239_at | HOXD13 | 0,03970495 | 4,80075575 | 0,45103286 | 0,65862598 | 0,82862727 |
| 117157_at | SH2D1B | -0,03618164 | 3,58913731 | -0,45082189 | 0,65877451 | 0,82876002 |
| 63905_at | MANBAL | -0,03857733 | 9,80532423 | -0,45052615 | 0,65898275 | 0,82896479 |
| 56889_at | TM9SF3 | 0,05420005 | 10,3092182 | 0,4504685 | 0,65902335 | 0,82896479 |
| 101927709_a | LOC10192770 | 0,04403699 | 3,16229245 | 0,45034962 | 0,65910707 | 0,82901596 |
| 101929153_a | LOC10192915 | -0,03251906 | 3,29479892 | -0,4501459 | 0,65925055 | 0,82908966 |
| 114836_at | SLAMF6 | -0,06358099 | 4,10793039 | -0,4501442 | 0,65925174 | 0,82908966 |
| 25981_at | DNAH1 | -0,04391221 | 4,92291811 | -0,44997358 | 0,65937192 | 0,82915844 |
| 339761_at | CYP27C1 | -0,35622338 | 5,64089091 | -0,44994433 | 0,65939252 | 0,82915844 |
| 1762_at | DMWD | -0,0416201 | 7,07902747 | -0,4495601 | 0,65966319 | 0,82944465 |
| 386758_at | ZNF582-AS1 | 0,11760903 | 5,27106872 | 0,44942355 | 0,6597594 | 0,82951147 |
| 7166_at | TPH1 | 0,04865829 | 3,03108932 | 0,44931286 | 0,65983738 | 0,82955538 |
| 101929762_a | LOC10192976 | -0,03865661 | 3,23983487 | -0,44919094 | 0,65992329 | 0,82960923 |
| 653140_at | FAM228A | 0,0608636 | 3,71762106 | 0,44889045 | 0,66013504 | 0,82979015 |
| 9253_at | NUMBL | 0,08417315 | 6,72365514 | 0,44875451 | 0,66023084 | 0,82979015 |
| 23639_at | LRRC6 | 0,0467194 | 3,93524198 | 0,4487507 | 0,66023353 | 0,82979015 |
| 3899_at | AFF3 | -0,03156619 | 3,89827019 | -0,44874222 | 0,6602395 | 0,82979015 |
| 101929694_a | LOC10192969 | -0,04042658 | 3,08770293 | -0,4485804 | 0,66035356 | 0,82983655 |
| 257240_at | KLHL34 | 0,0376495 | 4,45906681 | 0,44856759 | 0,66036259 | 0,82983655 |
| 101929717_a | LOC10192971 | -0,04679291 | 4,18327486 | -0,44842117 | 0,66046579 | 0,82987565 |
| 55083_at | KIF26B | -0,03835051 | 5,35967699 | -0,44834166 | 0,66052184 | 0,82987565 |
| 3006_at | HIST1H1C | -0,13554769 | 8,30855095 | -0,4481913 | 0,66062783 | 0,82987565 |
| 102800447_a | LOC10280044 | -0,03715466 | 3,65039743 | -0,44816929 | 0,66064335 | 0,82987565 |
| 101927396_a | LINC01514 | 0,04045737 | 3,88748861 | 0,44816718 | 0,66064484 | 0,82987565 |
| 55147_at | RBM23 | 0,07205128 | 9,75816429 | 0,44815676 | 0,66065218 | 0,82987565 |
| 6788_at | STK3 | 0,0763992 | 5,8859198 | 0,44804885 | 0,66072826 | 0,829887 |
| 29117_at | BRD7 | -0,03752368 | 8,58798161 | -0,44795843 | 0,66079201 | 0,829887 |
| 4330_at | MN1 | -0,04978976 | 4,61090183 | -0,44793911 | 0,66080563 | 0,829887 |
| 100507117_a | ASAP1-IT2 | -0,07218489 | 3,69436388 | -0,44789951 | 0,66083355 | 0,829887 |
| 23207_at | PLEKHM2 | -0,04675893 | 8,00560735 | -0,4478153 | 0,66089292 | 0,82990746 |
| 5605_at | MAP2K2 | -0,05796244 | 9,46056979 | -0,44741689 | 0,66117387 | 0,83020498 |
| 7849_at | PAX8 | 0,17628949 | 6,73078256 | 0,44728566 | 0,66126642 | 0,83020498 |
| 23732_at | FRRS1L | -0,04998584 | 4,22096796 | -0,44725462 | 0,66128831 | 0,83020498 |
| 51117_at | COQ4 | 0,12966202 | 8,28737782 | 0,44722029 | 0,66131252 | 0,83020498 |

| | | | | | | |
|---|---|---|---|---|---|---|
| 83854_at | ANGPTL6 | 0,049108 | 5,57455556 | 0,44717375 | 0,66134534 | 0,83020498 |
| 144453_at | BEST3 | 0,02301042 | 3,29824089 | 0,44699811 | 0,66146923 | 0,83030639 |
| 283310_at | OTOGL | -0,11874695 | 4,21527797 | -0,44652544 | 0,66180267 | 0,83066925 |
| 7263_at | TST | 0,03314592 | 9,69012103 | 0,44646608 | 0,66184455 | 0,83066925 |
| 55696_at | RBM22 | -0,03613865 | 8,79887647 | -0,44619896 | 0,66203303 | 0,83085166 |
| 2697_at | GJA1 | -0,10395527 | 5,9228902 | -0,44601884 | 0,66216013 | 0,83095704 |
| 22883_at | CLSTN1 | -0,04245509 | 9,19947215 | -0,44588434 | 0,66225505 | 0,83098098 |
| 123016_at | TTC8 | 0,055566 | 8,44398683 | 0,44586955 | 0,66226548 | 0,83098098 |
| 285908_at | LINC00174 | -0,04631994 | 6,14952058 | -0,44578721 | 0,6623236 | 0,83098514 |
| 10294_at | DNAJA2 | -0,08061331 | 8,46334004 | -0,4457067 | 0,66238042 | 0,83098514 |
| 9685_at | CLINT1 | -0,06183245 | 9,44092474 | -0,44568149 | 0,66239821 | 0,83098514 |
| 116441_at | TM4SF18 | -0,03577421 | 3,14503247 | -0,44559322 | 0,66246051 | 0,83100917 |
| 127845_at | GOLT1A | -0,07869596 | 5,13850009 | -0,44549019 | 0,66253324 | 0,83104629 |
| 347736_at | NME9 | -0,05710764 | 3,65511056 | -0,44522553 | 0,66272007 | 0,8312265 |
| 23256_at | SCFD1 | -0,05189087 | 7,90121073 | -0,444759 | 0,66304944 | 0,83152672 |
| 285704_at | RGMB | -0,0668884 | 6,15767403 | -0,44467812 | 0,66310656 | 0,83152672 |
| 55608_at | ANKRD10 | -0,0926652 | 7,79631537 | -0,44465162 | 0,66312527 | 0,83152672 |
| 100652929_a | HP09025 | -0,06313939 | 4,6466097 | -0,44464197 | 0,66313209 | 0,83152672 |
| 101928496_a | LINC01492 | 0,03026156 | 2,92465208 | 0,44436719 | 0,66332613 | 0,8317159 |
| 83992_at | CTTNBP2 | 0,03173247 | 3,19824243 | 0,44427827 | 0,66338893 | 0,8317405 |
| 285_at | ANGPT2 | 0,02580587 | 3,54504985 | 0,44419477 | 0,66344791 | 0,8317603 |
| 10827_at | FAM114A2 | 0,07605611 | 6,37709949 | 0,44398057 | 0,6635992 | 0,83189584 |
| 4358_at | MPV17 | 0,03758595 | 9,12904728 | 0,44372452 | 0,66378008 | 0,83206844 |
| 153218_at | SPINK13 | 0,03134538 | 2,79828414 | 0,44364043 | 0,66383949 | 0,83208876 |
| 389114_at | ZNF662 | -0,05478525 | 4,06083179 | -0,44339682 | 0,66401159 | 0,83219551 |
| 158067_at | AK8 | -0,06343769 | 5,81631081 | -0,44335981 | 0,66403775 | 0,83219551 |
| 79734_at | KCTD17 | -0,04540858 | 6,33884624 | -0,44333644 | 0,66405426 | 0,83219551 |
| 91442_at | FAAP24 | -0,07784392 | 6,5101278 | -0,44312658 | 0,66420255 | 0,83226926 |
| 55757_at | UGGT2 | -0,04007577 | 6,825535 | -0,44309178 | 0,66422714 | 0,83226926 |
| 4166_at | CHST6 | -0,04766177 | 4,89907803 | -0,44306972 | 0,66424273 | 0,83226926 |
| 51028_at | VPS36 | 0,1246556 | 8,26357415 | 0,44291584 | 0,66435148 | 0,83227983 |
| 140832_at | WFDC10A | -0,03859886 | 3,78762959 | -0,44289567 | 0,66436573 | 0,83227983 |
| 8927_at | BSN | -0,05962729 | 4,46238621 | -0,44287437 | 0,66438078 | 0,83227983 |
| 57018_at | CCNL1 | 0,10437815 | 8,6190026 | 0,44267018 | 0,6645251 | 0,83240649 |
| 101927640_a | LOC10192764 | -0,04985091 | 4,0808108 | -0,44232432 | 0,66476957 | 0,83265857 |
| 56097_at | PCDHGC5 | -0,03994942 | 4,12995834 | -0,4421665 | 0,66488114 | 0,83271792 |
| 221294_at | NT5DC1 | 0,05854132 | 8,08681507 | 0,442135 | 0,66490341 | 0,83271792 |
| 2794_at | GNL1 | 0,0254547 | 6,30926742 | 0,44171667 | 0,66519919 | 0,8329944 |
| 10859_at | LILRB1 | 0,03126485 | 3,79004892 | 0,44165352 | 0,66524384 | 0,8329944 |
| 728927_at | ZNF736 | -0,02433012 | 4,18946934 | -0,44163929 | 0,6652539 | 0,8329944 |
| 51162_at | EGFL7 | 0,08768603 | 6,81843579 | 0,44152409 | 0,66533537 | 0,83304226 |
| 6455_at | SH3GL1 | 0,03918542 | 9,38436744 | 0,44135467 | 0,66545518 | 0,83313812 |
| 8829_at | NRP1 | -0,11139829 | 5,99890957 | -0,44120681 | 0,66555976 | 0,83317856 |
| 10434_at | LYPLA1 | -0,04473575 | 10,011423 | -0,44118669 | 0,66557399 | 0,83317856 |
| 90141_at | EFCAB11 | -0,07561961 | 7,90797003 | -0,44101667 | 0,66569424 | 0,83324698 |
| 255374_at | MBLAC1 | 0,0463415 | 6,50132315 | 0,4409871 | 0,66571516 | 0,83324698 |
| 284427_at | SLC25A41 | -0,05016441 | 4,83574092 | -0,44088426 | 0,6657879 | 0,83328389 |
| 6854_at | SYN2 | -0,03063602 | 4,45016333 | -0,44082179 | 0,66583209 | 0,83328506 |
| 8418_at | CMAHP | 0,02687976 | 3,17892152 | 0,44056769 | 0,66601186 | 0,83335118 |
| 101927363_a | LOC10192736 | -0,05484092 | 3,26502264 | -0,44048006 | 0,66607386 | 0,83335118 |

| | | | | | | |
|---|---|---|---|---|---|---|
| 3553_at | IL1B | -0,18714787 | 7,86568574 | -0,4404584 | 0,66608918 | 0,83335118 |
| 344595_at | DUBR | 0,04827017 | 4,28495311 | 0,44039955 | 0,66613082 | 0,83335118 |
| 788_at | SLC25A20 | 0,1545235 | 6,6181773 | 0,44038528 | 0,66614092 | 0,83335118 |
| 257160_at | RNF214 | -0,03354399 | 7,09431793 | -0,44038021 | 0,6661445 | 0,83335118 |
| 23612_at | PHLDA3 | 0,06475149 | 8,629546 | 0,44021967 | 0,6662581 | 0,83341064 |
| 10568_at | SLC34A2 | -0,0461559 | 4,18322885 | -0,44019076 | 0,66627855 | 0,83341064 |
| 25800_at | SLC39A6 | 0,07361603 | 9,06524573 | 0,44011808 | 0,66632999 | 0,83342085 |
| 1036_at | CDO1 | -0,04217813 | 3,74676298 | -0,43988196 | 0,66649708 | 0,83351958 |
| 6598_at | SMARCB1 | 0,05605271 | 8,89066942 | 0,43978791 | 0,66656365 | 0,83351958 |
| 1880_at | GPR183 | -0,02686353 | 2,96640451 | -0,43971544 | 0,66661493 | 0,83351958 |
| 101928491_a | LOC10192849 | 0,04464896 | 3,70954914 | 0,43971007 | 0,66661873 | 0,83351958 |
| 140628_at | GATA5 | -0,05186909 | 4,85502271 | -0,43966253 | 0,66665238 | 0,83351958 |
| 977_at | CD151 | 0,09442904 | 9,37792718 | 0,43963226 | 0,66667381 | 0,83351958 |
| 7572_at | ZNF24 | 0,06588815 | 7,49223582 | 0,43957857 | 0,66671181 | 0,83351958 |
| 83715_at | ESPN | 0,04709933 | 5,48159564 | 0,43941975 | 0,66682423 | 0,83360602 |
| 64220_at | STRA6 | 0,05327541 | 5,52601949 | 0,4392268 | 0,66696082 | 0,83372267 |
| 5004_at | ORM1 | 0,0496702 | 3,14219787 | 0,43911002 | 0,66704349 | 0,8337719 |
| 100216546_a | LINC01004 | 0,05668056 | 4,29533345 | 0,43871225 | 0,66732513 | 0,83406981 |
| 84251_at | SGIP1 | -0,02820794 | 3,06667789 | -0,43840056 | 0,66754585 | 0,83421423 |
| 6872_at | TAF1 | -0,02539572 | 6,24467946 | -0,43838902 | 0,66755402 | 0,83421423 |
| 284_at | ANGPT1 | 0,03969532 | 4,21492024 | 0,43836562 | 0,66757059 | 0,83421423 |
| 83866_at | TTTY11 | -0,06070276 | 4,3065264 | -0,43814325 | 0,66772809 | 0,83432003 |
| 54627_at | MAP10 | -0,0298546 | 3,37619245 | -0,43808705 | 0,6677679 | 0,83432003 |
| 84636_at | GPR174 | -0,02616809 | 2,83238268 | -0,43806262 | 0,6677852 | 0,83432003 |
| 11258_at | DCTN3 | 0,06215443 | 9,80445181 | 0,43785682 | 0,66793098 | 0,83438168 |
| 22953_at | P2RX2 | 0,06271017 | 5,41487722 | 0,43783152 | 0,6679489 | 0,83438168 |
| 11275_at | KLHL2 | -0,17530062 | 7,99753228 | -0,43780952 | 0,66796448 | 0,83438168 |
| 51307_at | FAM53C | 0,07711373 | 8,48046092 | 0,4377376 | 0,66801543 | 0,83439121 |
| 1114_at | CHGB | -0,03628347 | 3,78541634 | -0,4376568 | 0,66807267 | 0,83440861 |
| 9684_at | LRRC14 | -0,12641015 | 7,01109483 | -0,43747153 | 0,66820393 | 0,83447398 |
| 30845_at | EHD3 | -0,09473441 | 6,88039661 | -0,43743189 | 0,66823202 | 0,83447398 |
| 10178_at | TENM1 | -0,04053475 | 3,8598467 | -0,43732482 | 0,66830788 | 0,83447398 |
| 3773_at | KCNJ16 | 0,03061027 | 3,40845418 | 0,4373212 | 0,66831044 | 0,83447398 |
| 5178_at | PEG3 | -0,03247454 | 3,20639134 | -0,4372599 | 0,66835388 | 0,83447398 |
| 23359_at | FAM189A1 | -0,03438763 | 5,12203777 | -0,43721607 | 0,66838494 | 0,83447398 |
| 84173_at | ELMOD3 | -0,03488566 | 6,43833564 | -0,4370683 | 0,66848964 | 0,83451983 |
| 440503_at | PLIN5 | 0,051183 | 6,33379058 | 0,43704197 | 0,66850831 | 0,83451983 |
| 4482_at | MSRA | -0,09651592 | 7,20241162 | -0,43693868 | 0,6685815 | 0,83452056 |
| 27013_at | CNPPD1 | 0,02794035 | 7,28853918 | 0,43691887 | 0,66859554 | 0,83452056 |
| 84552_at | PARD6G | -0,03621053 | 6,02146357 | -0,4367326 | 0,66872756 | 0,83455006 |
| 101927117_a | PLCG1-AS1 | 0,05315404 | 3,78133435 | 0,43671827 | 0,66873771 | 0,83455006 |
| 9919_at | SEC16A | 0,04935969 | 8,92858518 | 0,43668022 | 0,66876468 | 0,83455006 |
| 152816_at | C4orf26 | 0,05177564 | 3,4762467 | 0,43664101 | 0,66879247 | 0,83455006 |
| 200931_at | SLC51A | 0,05652972 | 3,8749727 | 0,43655544 | 0,66885312 | 0,83457168 |
| 4974_at | OMG | -0,06294122 | 4,51962522 | -0,4363961 | 0,66896606 | 0,83459033 |
| 29071_at | C1GALT1C1 | 0,04047366 | 7,69271491 | 0,43629445 | 0,66903812 | 0,83459033 |
| 162494_at | RHBDL3 | -0,03912555 | 4,97203196 | -0,43627374 | 0,6690528 | 0,83459033 |
| 3598_at | IL13RA2 | -0,06839583 | 3,49292261 | -0,43623504 | 0,66908024 | 0,83459033 |
| 9096_at | TBX18 | 0,02451925 | 3,59131657 | 0,43622875 | 0,66908469 | 0,83459033 |
| 51778_at | MYOZ2 | -0,02652794 | 2,94753687 | -0,43592623 | 0,66929917 | 0,83475157 |

| | | | | | | |
|---|---|---|---|---|---|---|
| 100126793_a | GHRLOS | 0,05606307 | 4,90147112 | 0,43592417 | 0,66930063 | 0,83475157 |
| 57488_at | ESYT2 | 0,03596211 | 9,01099356 | 0,43573831 | 0,66943242 | 0,83482116 |
| 60401_at | EDA2R | -0,06149105 | 5,40774413 | -0,43565365 | 0,66949245 | 0,83482116 |
| 11068_at | CYB561D2 | 0,03628533 | 6,84612104 | 0,43551409 | 0,66959141 | 0,83482116 |
| 90673_at | PPP1R3E | -0,03765511 | 6,25617535 | -0,43550728 | 0,66959624 | 0,83482116 |
| 11024_at | LILRA1 | -0,03532766 | 3,62889725 | -0,43548855 | 0,66960952 | 0,83482116 |
| 101927808_a | CLSTN2-AS1 | -0,03343522 | 3,97391069 | -0,43547878 | 0,66961645 | 0,83482116 |
| 84914_at | ZNF587 | 0,08007649 | 6,97678106 | 0,43479125 | 0,67010409 | 0,83537127 |
| 9949_at | AMMECR1 | -0,47279943 | 6,22753851 | -0,43473438 | 0,67014443 | 0,83537127 |
| 54556_at | ING3 | 0,04017667 | 6,91149194 | 0,4345375 | 0,67028411 | 0,83549131 |
| 9935_at | MAFB | 0,38568252 | 6,6604572 | 0,43427008 | 0,67047385 | 0,83565773 |
| 9034_at | CCRL2 | -0,04997599 | 4,93743022 | -0,43422705 | 0,67050438 | 0,83565773 |
| 23213_at | SULF1 | -0,04621399 | 4,57037035 | -0,43401894 | 0,67065205 | 0,83578769 |
| 163183_at | SYNE4 | -0,14915585 | 6,70188872 | -0,43383361 | 0,67078357 | 0,83589753 |
| 100507650_a | RNF212B | 0,04555678 | 3,89803906 | 0,43367689 | 0,67089479 | 0,83597131 |
| 55190_at | NUDT11 | 0,05492848 | 4,19171459 | 0,43362789 | 0,67092957 | 0,83597131 |
| 57113_at | TRPC7 | -0,0296826 | 3,59563706 | -0,43356533 | 0,67097397 | 0,83597255 |
| 1040_at | CDS1 | -0,07679472 | 8,11770152 | -0,43335653 | 0,67112219 | 0,83602845 |
| 23462_at | HEY1 | -0,30135494 | 6,73020022 | -0,43330908 | 0,67115587 | 0,83602845 |
| 375387_at | NRROS | -0,04564239 | 4,8188048 | -0,43330029 | 0,67116211 | 0,83602845 |
| 92014_at | SLC25A51 | 0,03612783 | 7,49398302 | 0,43320486 | 0,67122985 | 0,83602845 |
| 80194_at | TMEM134 | 0,03274181 | 7,60950953 | 0,43311931 | 0,67129059 | 0,83602845 |
| 390595_at | UBAP1L | -0,05114481 | 5,83966056 | -0,43311158 | 0,67129607 | 0,83602845 |
| 6752_at | SSTR2 | -0,04384647 | 4,47755266 | -0,43307416 | 0,67132264 | 0,83602845 |
| 401265_at | KLHL31 | 0,02885145 | 3,90806617 | 0,43281563 | 0,67150619 | 0,83618963 |
| 55850_at | USE1 | -0,0243789 | 8,51823361 | -0,43276958 | 0,67153888 | 0,83618963 |
| 246329_at | STAC3 | 0,05217091 | 5,19684339 | 0,43256223 | 0,67168612 | 0,8363189 |
| 201305_at | SPNS3 | 0,04347803 | 5,44365772 | 0,43234947 | 0,67183721 | 0,83640701 |
| 116228_at | COX20 | -0,07308723 | 8,27086748 | -0,43231976 | 0,67185831 | 0,83640701 |
| 10295_at | BCKDK | -0,04888485 | 7,9800254 | -0,43223241 | 0,67192034 | 0,83640701 |
| 255928_at | SYT14 | 0,03862483 | 3,24485408 | 0,43221801 | 0,67193057 | 0,83640701 |
| 80726_at | KIAA1683 | 0,06973245 | 4,96109155 | 0,43171546 | 0,67228753 | 0,83679728 |
| 79703_at | C11orf80 | 0,11372585 | 6,68218243 | 0,43163568 | 0,6723442 | 0,83681374 |
| 358_at | AQP1 | -0,03780211 | 4,50899467 | -0,43140017 | 0,67251152 | 0,83687992 |
| 4162_at | MCAM | -0,03314813 | 4,60200966 | -0,43139303 | 0,67251659 | 0,83687992 |
| 29883_at | CNOT7 | -0,14789221 | 7,73038963 | -0,43137738 | 0,67252771 | 0,83687992 |
| 340156_at | MYLK4 | -0,08764939 | 5,40303324 | -0,43103616 | 0,67277016 | 0,83710702 |
| 4236_at | MFAP1 | -0,04389685 | 8,83635513 | -0,43098822 | 0,67280422 | 0,83710702 |
| 10481_at | HOXB13 | -0,06831035 | 4,88458726 | -0,43089514 | 0,67287037 | 0,83710702 |
| 21_at | ABCA3 | -0,03790841 | 5,19984462 | -0,43087592 | 0,67288403 | 0,83710702 |
| 221955_at | DAGLB | 0,03869499 | 7,4212141 | 0,43055016 | 0,67311555 | 0,83716024 |
| 22798_at | LAMB4 | -0,02671737 | 3,36330102 | -0,43054499 | 0,67311922 | 0,83716024 |
| 60680_at | CELF5 | 0,04025993 | 4,21586765 | 0,43053715 | 0,6731248 | 0,83716024 |
| 60484_at | HAPLN2 | -0,04482048 | 5,00396671 | -0,43052128 | 0,67313607 | 0,83716024 |
| 137492_at | VPS37A | -0,11535448 | 7,42000715 | -0,43050998 | 0,67314411 | 0,83716024 |
| 643224_at | TUBBP5 | 0,3209946 | 6,15786444 | 0,43044564 | 0,67318984 | 0,83716306 |
| 4986_at | OPRK1 | -0,03831131 | 3,55858887 | -0,42992888 | 0,67355719 | 0,83756582 |
| 6920_at | TCEA3 | 0,02929553 | 5,27706197 | 0,42974127 | 0,67369058 | 0,83763421 |
| 84787_at | SUV420H2 | -0,05299883 | 6,50940179 | -0,42968222 | 0,67373256 | 0,83763421 |
| 85462_at | FHDC1 | -0,06026354 | 6,42558665 | -0,42966804 | 0,67374264 | 0,83763421 |

| | | | | | | |
|---|---|---|---|---|---|---|
| 27163_at | NAAA | -0,02872511 | 6,41957288 | -0,42931821 | 0,67399141 | 0,83788941 |
| 1265_at | CNN2 | 0,04775475 | 9,96100801 | 0,42871237 | 0,67442231 | 0,83837099 |
| 4113_at | MAGEB2 | 0,04449855 | 3,70648006 | 0,42840998 | 0,67463743 | 0,83853639 |
| 84885_at | ZDHHC12 | 0,04178588 | 8,961253 | 0,42840296 | 0,67464243 | 0,83853639 |
| 131368_at | ZPLD1 | 0,02551669 | 3,46513656 | 0,42832887 | 0,67469514 | 0,8385478 |
| 2568_at | GABRP | 0,43784366 | 5,12835886 | 0,42808918 | 0,67486567 | 0,83870564 |
| 1879_at | EBF1 | -0,02764661 | 3,61406663 | -0,42784292 | 0,67504091 | 0,83880628 |
| 1374_at | CPT1A | -0,04212172 | 5,94816759 | -0,42778025 | 0,67508551 | 0,83880628 |
| 339488_at | TFAP2E | -0,04841232 | 5,95366923 | -0,42776572 | 0,67509585 | 0,83880628 |
| 101559451_a | LOC10155945 | -0,03445574 | 4,00356134 | -0,42770626 | 0,67513816 | 0,83880628 |
| 378925_at | RNF148 | 0,03779906 | 3,25672587 | 0,42766943 | 0,67516437 | 0,83880628 |
| 11096_at | ADAMTS5 | -0,03670928 | 3,94828089 | -0,42734257 | 0,67539701 | 0,83892771 |
| 4076_at | CAPRIN1 | -0,07198934 | 8,92776429 | -0,42734205 | 0,67539739 | 0,83892771 |
| 644794_at | LOC644794 | 0,06870773 | 5,53530411 | 0,42730212 | 0,6754258 | 0,83892771 |
| 222698_at | NKAPL | -0,03085384 | 3,44354122 | -0,42728735 | 0,67543632 | 0,83892771 |
| 4342_at | MOS | -0,04815348 | 4,32611396 | -0,42715737 | 0,67552884 | 0,83894318 |
| 5127_at | CDK16 | -0,0418191 | 8,62720603 | -0,42714748 | 0,67553588 | 0,83894318 |
| 55584_at | CHRNA9 | 0,0501806 | 3,45199051 | 0,42704992 | 0,67560533 | 0,8389617 |
| 101929025_a | LINC01516 | -0,04257763 | 3,34986407 | -0,42700417 | 0,6756379 | 0,8389617 |
| 5715_at | PSMD9 | -0,03354058 | 7,3375176 | -0,42686645 | 0,67573594 | 0,83902936 |
| 6456_at | SH3GL2 | 0,02563388 | 4,06484491 | 0,42669958 | 0,67585475 | 0,83912279 |
| 221002_at | RASGEF1A | -0,08042412 | 4,4641416 | -0,42646254 | 0,67602353 | 0,83925083 |
| 102659288_a | KCCAT333 | 0,59512379 | 6,29399604 | 0,42643236 | 0,67604502 | 0,83925083 |
| 205860_at | TRIML2 | -0,03357147 | 4,214312 | -0,42629345 | 0,67614394 | 0,83931955 |
| 60468_at | BACH2 | -0,03851862 | 3,86832015 | -0,42612577 | 0,67626335 | 0,83941368 |
| 388407_at | C17orf82 | -0,05068881 | 6,63932822 | -0,42576713 | 0,67651879 | 0,83967664 |
| 280_at | AMY2B | 0,20769584 | 6,52814103 | 0,42564061 | 0,67660891 | 0,83973211 |
| 51427_at | ZNF107 | 0,02692876 | 5,74672598 | 0,42558199 | 0,67665067 | 0,83973211 |
| 23591_at | FAM215A | -0,05890155 | 5,50054717 | -0,4255117 | 0,67670074 | 0,83974014 |
| 55768_at | NGLY1 | -0,09242196 | 8,42980961 | -0,42522374 | 0,67690589 | 0,83989789 |
| 57600_at | FNIP2 | 0,11780642 | 7,12656289 | 0,42521087 | 0,67691506 | 0,83989789 |
| 10800_at | CYSLTR1 | -0,02226952 | 3,29202526 | -0,42484123 | 0,67717843 | 0,84011342 |
| 140881_at | DEFB129 | -0,04071487 | 3,8024505 | -0,42480651 | 0,67720317 | 0,84011342 |
| 101928865_a | LOC10192886 | -0,02875508 | 3,39253423 | -0,42475274 | 0,6772415 | 0,84011342 |
| 23358_at | USP24 | 0,03293932 | 7,97040753 | 0,42468495 | 0,6772898 | 0,84011342 |
| 8326_at | FZD9 | -0,03960384 | 5,6148614 | -0,42466106 | 0,67730683 | 0,84011342 |
| 55811_at | ADCY10 | -0,02555533 | 3,27287026 | -0,42440485 | 0,67748943 | 0,84018275 |
| 23125_at | CAMTA2 | 0,02449048 | 7,06201572 | 0,42426602 | 0,67758837 | 0,84018275 |
| 9889_at | ZBED4 | 0,03810825 | 7,68159805 | 0,4242314 | 0,67761305 | 0,84018275 |
| 9851_at | KIAA0753 | 0,03347548 | 6,99382984 | 0,42412055 | 0,67769207 | 0,84018275 |
| 6708_at | SPTA1 | 0,03853784 | 3,86463587 | 0,42411286 | 0,67769754 | 0,84018275 |
| 101929198_a | ATP13A4-AS1 | 0,02062174 | 2,77966252 | 0,42409457 | 0,67771059 | 0,84018275 |
| 100873985_a | MED14OS | -0,03876976 | 4,97919309 | -0,42405819 | 0,67773652 | 0,84018275 |
| 90187_at | EMILIN3 | 0,05355581 | 6,15305117 | 0,42404304 | 0,67774732 | 0,84018275 |
| 10736_at | SIX2 | 0,14398593 | 6,22036817 | 0,42397334 | 0,677797 | 0,84018275 |
| 84225_at | ZMYND15 | 0,04797738 | 4,76582307 | 0,4239707 | 0,67779888 | 0,84018275 |
| 140685_at | ZBTB46 | 0,08457712 | 5,9991536 | 0,42385906 | 0,67787847 | 0,84022004 |
| 4543_at | MTNR1A | -0,04495153 | 4,62902154 | -0,42380613 | 0,67791621 | 0,84022004 |
| 256586_at | LYSMD2 | -0,25190363 | 7,65792245 | -0,42370697 | 0,6779869 | 0,8402536 |
| 283385_at | MORN3 | -0,03842563 | 5,69387818 | -0,42344103 | 0,67817652 | 0,84035457 |

| | | | | | | |
|---|---|---|---|---|---|---|
| 286046_at | XKR6 | -0,02834146 | 4,59093809 | -0,42342142 | 0,67819049 | 0,84035457 |
| 100852409_a | NEGR1-IT1 | 0,04586008 | 3,4172515 | 0,42334565 | 0,67824453 | 0,84035457 |
| 51335_at | NGRN | 0,03936103 | 9,8838441 | 0,42330848 | 0,67827103 | 0,84035457 |
| 3176_at | HNMT | 0,17956489 | 6,28866186 | 0,42327268 | 0,67829655 | 0,84035457 |
| 6829_at | SUPT5H | 0,02406123 | 8,2488656 | 0,42322561 | 0,67833012 | 0,84035457 |
| 259287_at | TAS2R41 | -0,03988819 | 4,85767815 | -0,42297946 | 0,67850566 | 0,84051798 |
| 101928388_a | PLA2G4E-AS1 | 0,03058958 | 3,18860058 | 0,42279113 | 0,67863998 | 0,84063031 |
| 282706_at | DAOA-AS1 | -0,03466751 | 3,03136284 | -0,42250207 | 0,67884617 | 0,84083164 |
| 10141_at | LINC01587 | -0,04412897 | 3,23585416 | -0,42230492 | 0,6789868 | 0,84095177 |
| 56269_at | IRGC | -0,04979457 | 4,40471151 | -0,42212288 | 0,67911667 | 0,84105854 |
| 103910_at | MYL12B | 0,06321221 | 9,99097358 | 0,42204851 | 0,67916974 | 0,84107018 |
| 338692_at | ANKRD13D | 0,04192915 | 6,46322913 | 0,42175314 | 0,67938049 | 0,84127709 |
| 10427_at | SEC24B | -0,09092782 | 9,196646 | -0,42158592 | 0,67949982 | 0,84133212 |
| 4052_at | LTBP1 | -0,12180318 | 6,57992865 | -0,42156845 | 0,67951228 | 0,84133212 |
| 55801_at | IL26 | 0,02637486 | 2,71417366 | 0,4212296 | 0,67975412 | 0,84157745 |
| 282996_at | RBM20 | -0,04748882 | 3,51691856 | -0,42112972 | 0,6798254 | 0,84159834 |
| 101927944_a | FAM53B-AS1 | -0,04205901 | 3,69166613 | -0,42108354 | 0,67985837 | 0,84159834 |
| 3294_at | HSD17B2 | -0,28373735 | 5,34598447 | -0,42099239 | 0,67992343 | 0,84162479 |
| 201617_at | LINC00870 | -0,05079552 | 3,49471305 | -0,42069658 | 0,6801346 | 0,84178384 |
| 374378_at | GALNT18 | -0,13644367 | 6,82356744 | -0,42068997 | 0,68013932 | 0,84178384 |
| 50626_at | CYHR1 | 0,10362259 | 7,48522377 | 0,42021474 | 0,68047863 | 0,84205305 |
| 100130418_a | CECR7 | 0,03221983 | 4,49278331 | 0,42015872 | 0,68051863 | 0,84205305 |
| 55586_at | MIOX | -0,05843139 | 4,49833648 | -0,42014852 | 0,68052591 | 0,84205305 |
| 84255_at | SLC37A3 | -0,07956571 | 8,29635546 | -0,42012061 | 0,68054584 | 0,84205305 |
| 1045_at | CDX2 | -0,05713774 | 4,62820074 | -0,41996507 | 0,68065691 | 0,84205305 |
| 56413_at | LTB4R2 | 0,05119626 | 4,4987271 | 0,41991972 | 0,6806893 | 0,84205305 |
| 9441_at | MED26 | -0,02846176 | 6,52050197 | -0,41990763 | 0,68069794 | 0,84205305 |
| 7772_at | ZNF229 | 0,06276923 | 4,01790831 | 0,41989559 | 0,68070654 | 0,84205305 |
| 3918_at | LAMC2 | -0,06347606 | 6,99047124 | -0,41970113 | 0,68084542 | 0,84216941 |
| 219409_at | GSX1 | -0,03175027 | 5,05877914 | -0,41964145 | 0,68088804 | 0,84216941 |
| 23318_at | ZCCHC11 | 0,04316288 | 7,34432846 | 0,41954615 | 0,68095611 | 0,84219857 |
| 6286_at | S100P | -0,04463137 | 12,9499956 | -0,41930813 | 0,68112613 | 0,84219857 |
| 162239_at | ZFP1 | 0,04666792 | 6,81037285 | 0,41930183 | 0,68113063 | 0,84219857 |
| 27197_at | GPR82 | 0,02012489 | 2,91952172 | 0,41929413 | 0,68113613 | 0,84219857 |
| 100874018_a | EPN2-AS1 | -0,06002544 | 3,81703356 | -0,41923964 | 0,68117505 | 0,84219857 |
| 54901_at | CDKAL1 | 0,12740666 | 6,65532384 | 0,41923878 | 0,68117567 | 0,84219857 |
| 2357_at | FPR1 | -0,05900607 | 4,96674267 | -0,41915286 | 0,68123705 | 0,84219857 |
| 4135_at | MAP6 | -0,04166982 | 4,61904031 | -0,41900363 | 0,68134366 | 0,84219857 |
| 9515_at | STXBP5L | 0,02889824 | 3,00022875 | 0,41895212 | 0,68138046 | 0,84219857 |
| 283461_at | C12orf40 | -0,03315501 | 2,86157255 | -0,4189477 | 0,68138362 | 0,84219857 |
| 3739_at | KCNA4 | -0,03871985 | 4,82898004 | -0,41887386 | 0,68143638 | 0,84219857 |
| 84923_at | FAM104A | 0,03890297 | 8,93096589 | 0,41879607 | 0,68149196 | 0,84219857 |
| 85445_at | CNTNAP4 | -0,03604081 | 4,24839183 | -0,41873459 | 0,68153589 | 0,84219857 |
| 132851_at | SPATA4 | -0,03970994 | 4,90337403 | -0,41865551 | 0,6815924 | 0,84219857 |
| 25998_at | IBTK | 0,05389482 | 8,58137298 | 0,41861001 | 0,68162491 | 0,84219857 |
| 26010_at | SPATS2L | 0,04528103 | 8,7969476 | 0,41858056 | 0,68164595 | 0,84219857 |
| 441666_at | LOC441666 | 0,04049828 | 2,84776882 | 0,41853906 | 0,68167561 | 0,84219857 |
| 134218_at | DNAJC21 | -0,02890892 | 7,11767069 | -0,41837602 | 0,68179212 | 0,84219857 |
| 56946_at | C11orf30 | -0,03913004 | 7,23534937 | -0,41836806 | 0,6817978 | 0,84219857 |
| 54784_at | ALKBH4 | -0,03787186 | 6,53096609 | -0,41836172 | 0,68180234 | 0,84219857 |

| | | | | | | |
|---|---|---|---|---|---|---|
| 6856_at | SYPL1 | -0,04184926 | 10,9591511 | -0,41832335 | 0,68182976 | 0,84219857 |
| 1059_at | CENPB | -0,09105212 | 6,93671925 | -0,41823739 | 0,68189119 | 0,84222044 |
| 781_at | CACNA2D1 | 0,16152917 | 4,45561209 | 0,41815297 | 0,68195153 | 0,84224096 |
| 920_at | CD4 | -0,04843849 | 5,65131722 | -0,41784437 | 0,6821721 | 0,84245937 |
| 4854_at | NOTCH3 | -0,04452911 | 7,25607279 | -0,41776022 | 0,68223225 | 0,84247964 |
| 286148_at | DPY19L4 | 0,05457879 | 9,04798978 | 0,41769082 | 0,68228186 | 0,84248689 |
| 56670_at | SUCNR1 | 0,03909605 | 3,63794586 | 0,41760104 | 0,68234604 | 0,84251214 |
| 2201_at | FBN2 | -0,32536332 | 7,22133571 | -0,41710175 | 0,68270302 | 0,84273453 |
| 7317_at | UBA1 | 0,04933492 | 10,1270209 | 0,41696087 | 0,68280375 | 0,84273453 |
| 100286844_a | BCDIN3D-AS1 | -0,04719996 | 4,42084807 | -0,41688217 | 0,68286003 | 0,84273453 |
| 51703_at | ACSL5 | 0,10134979 | 9,10609572 | 0,4168769 | 0,6828638 | 0,84273453 |
| 3676_at | ITGA4 | -0,02995831 | 3,05664911 | -0,41679072 | 0,68292543 | 0,84273453 |
| 9477_at | MED20 | 0,04710475 | 7,9571859 | 0,41675918 | 0,68294799 | 0,84273453 |
| 4137_at | MAPT | -0,03425593 | 5,24921741 | -0,41672526 | 0,68297225 | 0,84273453 |
| 167_at | CRISP1 | 0,03907737 | 3,22890198 | 0,41672213 | 0,68297449 | 0,84273453 |
| 124997_at | WDR81 | 0,04244043 | 7,50405631 | 0,41671777 | 0,6829776 | 0,84273453 |
| 100125288_a | ZGLP1 | -0,06229619 | 5,9399215 | -0,41669004 | 0,68299743 | 0,84273453 |
| 284576_at | LEMD1-AS1 | 0,02795723 | 3,37870876 | 0,41667611 | 0,6830074 | 0,84273453 |
| 55253_at | TYW1 | -0,04891786 | 7,91031321 | -0,41649627 | 0,68313602 | 0,84283924 |
| 90326_at | THAP3 | -0,04211754 | 6,10908396 | -0,41619305 | 0,68335291 | 0,84301134 |
| 4185_at | ADAM11 | -0,02632251 | 5,42497128 | -0,41613977 | 0,68339102 | 0,84301134 |
| 6645_at | SNTB2 | 0,08277228 | 8,15688891 | 0,41611772 | 0,6834068 | 0,84301134 |
| 25898_at | RCHY1 | 0,11067385 | 6,84748317 | 0,41601372 | 0,6834812 | 0,84304913 |
| #N/A | LL22NC03-75I | -0,04041473 | 4,6147966 | -0,41558832 | 0,68378554 | 0,84336022 |
| 90632_at | LINC00473 | 0,04590114 | 4,09000572 | 0,41544172 | 0,68389044 | 0,84336022 |
| 10919_at | EHMT2 | -0,02392828 | 7,32365188 | -0,41539145 | 0,68392642 | 0,84336022 |
| 2649_at | NR6A1 | 0,02352224 | 5,84698824 | 0,41538697 | 0,68392962 | 0,84336022 |
| 23187_at | PHLDB1 | 0,05063514 | 6,83083271 | 0,41535527 | 0,68395231 | 0,84336022 |
| 11098_at | PRSS23 | -0,22253519 | 6,68759906 | -0,41528192 | 0,6840048 | 0,84337095 |
| 284395_at | LOC284395 | -0,03953871 | 5,30306155 | -0,41510609 | 0,68413063 | 0,84342515 |
| 51209_at | RAB9B | 0,12852403 | 5,86866125 | 0,41508322 | 0,68414699 | 0,84342515 |
| 374354_at | NHLRC2 | -0,10928833 | 5,76929639 | -0,41502448 | 0,68418904 | 0,84342515 |
| 29953_at | TRHDE | 0,02781931 | 3,10711944 | 0,41497578 | 0,68422389 | 0,84342515 |
| 79034_at | C7orf26 | -0,02822264 | 7,94544793 | -0,41481255 | 0,68434072 | 0,84351519 |
| 27348_at | TOR1B | 0,03283766 | 8,24235891 | 0,41451191 | 0,68455592 | 0,84372645 |
| 5775_at | PTPN4 | -0,06312974 | 6,29501817 | -0,4141786 | 0,68479454 | 0,84396655 |
| 219972_at | MPEG1 | 0,04588599 | 3,59559304 | 0,41405025 | 0,68488643 | 0,84402581 |
| 10855_at | HPSE | 0,20601815 | 5,43800419 | 0,41393864 | 0,68496635 | 0,84407029 |
| 6865_at | TACR2 | -0,04872984 | 5,04322471 | -0,41377948 | 0,68508032 | 0,84415673 |
| 3233_at | HOXD4 | -0,10046512 | 6,75074517 | -0,4136577 | 0,68516752 | 0,84421018 |
| 7337_at | UBE3A | -0,0612339 | 8,13457666 | -0,41351666 | 0,68526853 | 0,84424298 |
| 284649_at | SMG7-AS1 | 0,04854484 | 3,85652888 | 0,41349814 | 0,6852818 | 0,84424298 |
| 10068_at | IL18BP | 0,03842556 | 5,84187433 | 0,41326663 | 0,68544761 | 0,84439325 |
| 140459_at | ASB6 | -0,02933526 | 7,44054783 | -0,41315502 | 0,68552755 | 0,84442601 |
| 124783_at | SPATA32 | -0,03771063 | 4,05039891 | -0,41307859 | 0,68558229 | 0,84442601 |
| 101928646_a | ARNTL2-AS1 | -0,03431051 | 2,85685981 | -0,4130459 | 0,68560571 | 0,84442601 |
| 2159_at | F10 | -0,03909995 | 4,90056765 | -0,4128672 | 0,68573372 | 0,84447756 |
| 3801_at | KIFC3 | -0,03330628 | 6,21008702 | -0,41283637 | 0,68575581 | 0,84447756 |
| 199720_at | GGN | -0,04184643 | 5,47825199 | -0,41279103 | 0,68578829 | 0,84447756 |
| 57821_at | CCDC181 | 0,03667011 | 3,64711683 | 0,41274269 | 0,68582292 | 0,84447756 |

| | | | | | | |
|---|---|---|---|---|---|---|
| 100506675_a | LOC10050667 | -0,03666315 | 3,65219279 | -0,41252562 | 0,68597844 | 0,84449148 |
| 84033_at | OBSCN | -0,03266914 | 4,76426307 | -0,41249011 | 0,68600388 | 0,84449148 |
| 2984_at | GUCY2C | -0,03670711 | 3,36853096 | -0,41241101 | 0,68606056 | 0,84449148 |
| 101928551_a | LOC10192855 | -0,02781022 | 3,12541436 | -0,41240806 | 0,68606267 | 0,84449148 |
| 114112_at | TXNRD3 | -0,12022794 | 6,79524675 | -0,41240572 | 0,68606435 | 0,84449148 |
| 150082_at | LCA5L | 0,05345103 | 4,19313451 | 0,41235977 | 0,68609727 | 0,84449148 |
| 6561_at | SLC13A1 | 0,0239296 | 2,82767819 | 0,41209442 | 0,68628742 | 0,84467155 |
| 29919_at | C18orf8 | -0,11413407 | 7,73123737 | -0,41195534 | 0,68638708 | 0,84474025 |
| 4253_at | CTAGE5 | 0,05282489 | 5,36799752 | 0,41181208 | 0,68648976 | 0,84477335 |
| 10445_at | MCRS1 | 0,03686734 | 8,9083508 | 0,41179543 | 0,68650169 | 0,84477335 |
| 10863_at | ADAM28 | 0,0949751 | 5,51474263 | 0,41150893 | 0,68670704 | 0,84493797 |
| 90649_at | ZNF486 | -0,03483645 | 3,83844844 | -0,41148432 | 0,68672468 | 0,84493797 |
| 11317_at | RBPJL | -0,0409118 | 4,92861431 | -0,41142519 | 0,68676706 | 0,84493797 |
| 90011_at | KIR3DX1 | -0,06020032 | 3,96872152 | -0,41116925 | 0,68695054 | 0,84510252 |
| 389932_at | AKR1C6P | -0,02654667 | 3,91375447 | -0,41107284 | 0,68701965 | 0,84510252 |
| 9275_at | BCL7B | -0,07866866 | 8,70676605 | -0,41105503 | 0,68703242 | 0,84510252 |
| 90231_at | KIAA2013 | -0,03476135 | 7,87095052 | -0,41092384 | 0,68712648 | 0,84512127 |
| 3993_at | LLGL2 | 0,06003152 | 7,88817865 | 0,41091139 | 0,6871354 | 0,84512127 |
| 6391_at | SDHC | 0,06011414 | 6,61173655 | 0,41047913 | 0,68744536 | 0,84542363 |
| 55068_at | ENOX1 | 0,13440806 | 4,61347582 | 0,41044613 | 0,68746902 | 0,84542363 |
| 780776_at | TVP23A | -0,03120218 | 4,14306199 | -0,41022734 | 0,68762593 | 0,84543167 |
| 219743_at | TYSND1 | -0,03431769 | 7,32611778 | -0,41020579 | 0,68764139 | 0,84543167 |
| 6540_at | SLC6A13 | -0,04552507 | 5,09934782 | -0,41019876 | 0,68764643 | 0,84543167 |
| 56124_at | PCDHB12 | 0,04047322 | 4,89412703 | 0,41019223 | 0,68765111 | 0,84543167 |
| 6595_at | SMARCA2 | 0,07941176 | 7,38823706 | 0,40979927 | 0,68793297 | 0,8457041 |
| 7302_at | TYRO3P | -0,0357905 | 4,15625422 | -0,40961244 | 0,68806699 | 0,8457041 |
| 133121_at | ENPP6 | -0,03498418 | 3,26362204 | -0,40956397 | 0,68810177 | 0,8457041 |
| 85449_at | KIAA1755 | -0,03798625 | 4,80697234 | -0,40951037 | 0,68814022 | 0,8457041 |
| 93035_at | PKHD1L1 | 0,02528834 | 2,91445034 | 0,40935755 | 0,68824986 | 0,8457041 |
| 100506866_a | TTN-AS1 | 0,03992857 | 4,54970147 | 0,40930081 | 0,68829057 | 0,8457041 |
| 2324_at | FLT4 | -0,03783075 | 4,59914038 | -0,40928804 | 0,68829973 | 0,8457041 |
| 201164_at | PLD6 | -0,07857498 | 5,73638449 | -0,40925307 | 0,68832482 | 0,8457041 |
| 134526_at | ACOT12 | 0,04478052 | 3,2753094 | 0,40911813 | 0,68842164 | 0,8457041 |
| 148898_at | ZNF436-AS1 | -0,10116724 | 6,17606339 | -0,40908441 | 0,68844584 | 0,8457041 |
| 160287_at | LDHAL6A | 0,02770974 | 3,34019393 | 0,40907068 | 0,68845569 | 0,8457041 |
| 84945_at | ABHD13 | 0,0720658 | 6,90658452 | 0,40905291 | 0,68846844 | 0,8457041 |
| 340024_at | SLC6A19 | -0,04160254 | 5,1942603 | -0,4090496 | 0,68847081 | 0,8457041 |
| 64375_at | IKZF4 | 0,04177476 | 6,16918034 | 0,40902656 | 0,68848735 | 0,8457041 |
| 158434_at | LOC158434 | -0,03200605 | 3,62128857 | -0,40861155 | 0,68878517 | 0,84592611 |
| 10648_at | SCGB1D1 | -0,03877614 | 3,35638619 | -0,40860412 | 0,68879051 | 0,84592611 |
| 2797_at | GNRH2 | -0,05343087 | 5,25789653 | -0,40856502 | 0,68881857 | 0,84592611 |
| 755_at | C21orf2 | 0,06674777 | 6,25043198 | 0,40852995 | 0,68884374 | 0,84592611 |
| 9840_at | TESPA1 | 0,02546177 | 3,78142588 | 0,40824989 | 0,68904475 | 0,84611902 |
| 150678_at | MYEOV2 | -0,03363211 | 8,70692051 | -0,40815875 | 0,68911018 | 0,84614542 |
| 3298_at | HSF2 | 0,05443877 | 6,78759036 | 0,40793402 | 0,6892715 | 0,84628956 |
| 6407_at | SEMG2 | -0,02988679 | 3,27787746 | -0,40757602 | 0,68952853 | 0,8465145 |
| 84727_at | SPSB2 | 0,07655368 | 7,2768779 | 0,40755644 | 0,68954259 | 0,8465145 |
| 7368_at | UGT8 | 0,03426169 | 3,69955049 | 0,40747142 | 0,68960364 | 0,8465335 |
| #N/A | CTD-2194D22 | -0,03743882 | 3,36893201 | -0,40741247 | 0,68964597 | 0,8465335 |
| 80122_at | MAP3K19 | 0,02403905 | 3,5484055 | 0,40726973 | 0,68974847 | 0,84655253 |

| | | | | | | |
|---|---|---|---|---|---|---|
| 2281_at | FKBP1B | -0,03837668 | 4,67607898 | -0,40722913 | 0,68977763 | 0,84655253 |
| 55361_at | PI4K2A | 0,05855267 | 8,4058188 | 0,40717606 | 0,68981573 | 0,84655253 |
| 128817_at | CSTL1 | 0,03425351 | 3,40894614 | 0,40706379 | 0,68989636 | 0,84655253 |
| 23046_at | KIF21B | 0,05898943 | 5,60017679 | 0,40703299 | 0,68991848 | 0,84655253 |
| 9582_at | APOBEC3B | 0,08093081 | 8,28878266 | 0,40698479 | 0,6899531 | 0,84655253 |
| 7057_at | THBS1 | 0,05494754 | 8,65559006 | 0,40689841 | 0,69001514 | 0,84655253 |
| 183_at | AGT | -0,03899884 | 4,45265285 | -0,40687756 | 0,69003011 | 0,84655253 |
| 10343_at | PKDREJ | -0,02297147 | 4,46402489 | -0,40684014 | 0,69005699 | 0,84655253 |
| 6386_at | SDCBP | 0,08337644 | 11,1897499 | 0,40671012 | 0,69015038 | 0,84661318 |
| 10110_at | SGK2 | -0,04144551 | 4,59570377 | -0,40652819 | 0,69028107 | 0,84671957 |
| 84726_at | PRRC2B | 0,0381876 | 8,16667738 | 0,40646516 | 0,69032634 | 0,84672119 |
| 254896_at | LOC254896 | -0,04816234 | 4,87599347 | -0,40616821 | 0,69053967 | 0,84689241 |
| 5023_at | P2RX1 | -0,0318932 | 5,22393052 | -0,40611158 | 0,69058035 | 0,84689241 |
| 349408_at | TLR8-AS1 | 0,04992248 | 4,13996501 | 0,40608726 | 0,69059783 | 0,84689241 |
| 55112_at | WDR60 | 0,05842276 | 6,97172307 | 0,40595905 | 0,69068995 | 0,84694195 |
| 78998_at | RHPN1-AS1 | -0,06544681 | 5,76848615 | -0,40590866 | 0,69072616 | 0,84694195 |
| 333929_at | SNAI3 | 0,03805866 | 5,72010705 | 0,40575625 | 0,69083567 | 0,84695816 |
| 83844_at | USP26 | -0,03825714 | 3,44576374 | -0,40574366 | 0,69084472 | 0,84695816 |
| 56123_at | PCDHB13 | 0,04380153 | 4,19914956 | 0,40570669 | 0,69087128 | 0,84695816 |
| 9180_at | OSMR | -0,14671897 | 7,486212 | -0,40528878 | 0,69117161 | 0,84726081 |
| 124402_at | UBALD1 | -0,03482917 | 7,33931981 | -0,40524076 | 0,69120613 | 0,84726081 |
| 157777_at | MCMDC2 | 0,03945615 | 3,15651736 | 0,40505358 | 0,69134067 | 0,84737181 |
| 91433_at | RCCD1 | -0,08124952 | 7,57831828 | -0,40493489 | 0,69142599 | 0,84742246 |
| 1009_at | CDH11 | -0,02699756 | 3,48964597 | -0,40478108 | 0,69153655 | 0,84748009 |
| 134147_at | CMBL | -0,13077704 | 6,23230734 | -0,40474707 | 0,691561 | 0,84748009 |
| 6423_at | SFRP2 | 0,0329102 | 3,93024726 | 0,40466546 | 0,69161967 | 0,84749808 |
| 207_at | AKT1 | -0,08584874 | 9,7766603 | -0,40451744 | 0,69172609 | 0,84755267 |
| 7528_at | YY1 | -0,03646172 | 9,88146255 | -0,40448109 | 0,69175222 | 0,84755267 |
| 255025_at | LINC00879 | -0,04008717 | 3,42482345 | -0,40425656 | 0,69191366 | 0,84765639 |
| 6555_at | SLC10A2 | -0,04181386 | 3,59188501 | -0,40419749 | 0,69195613 | 0,84765639 |
| 255082_at | CASC2 | 0,03143282 | 3,48181015 | 0,40417975 | 0,69196889 | 0,84765639 |
| 10188_at | TNK2 | -0,02245199 | 6,22983776 | -0,40410259 | 0,69202438 | 0,84767045 |
| 29966_at | STRN3 | -0,05518456 | 7,96716667 | -0,40364906 | 0,69235053 | 0,84784498 |
| 153364_at | MBLAC2 | -0,0965685 | 4,24712694 | -0,40362212 | 0,69236991 | 0,84784498 |
| 56254_at | RNF20 | -0,04356768 | 8,27254074 | -0,40361399 | 0,69237576 | 0,84784498 |
| 22852_at | ANKRD26 | -0,07031383 | 5,04327845 | -0,40357902 | 0,69240091 | 0,84784498 |
| 128488_at | WFDC12 | -0,07084209 | 5,22422064 | -0,40353883 | 0,69242982 | 0,84784498 |
| 91603_at | ZNF830 | 0,06033143 | 7,04471139 | 0,40353727 | 0,69243094 | 0,84784498 |
| 54836_at | BSPRY | -0,07381913 | 6,01576033 | -0,40343047 | 0,69250776 | 0,84788514 |
| 79894_at | ZNF672 | 0,04635945 | 7,69806802 | 0,40332529 | 0,69258341 | 0,84791362 |
| 4585_at | MUC4 | -0,06221828 | 4,24108572 | -0,40315128 | 0,69270859 | 0,84791362 |
| 100507057_a | MFI2-AS1 | -0,05028311 | 6,23983481 | -0,40312604 | 0,69272675 | 0,84791362 |
| 84072_at | HORMAD1 | 0,26191896 | 4,9186936 | 0,40295626 | 0,69284889 | 0,84791362 |
| 5028_at | P2RY1 | -0,11637567 | 5,42792507 | -0,40289479 | 0,69289311 | 0,84791362 |
| 100129034_a | LOC10012903 | -0,05606506 | 6,17022865 | -0,40284718 | 0,69292737 | 0,84791362 |
| 6840_at | SVIL | 0,14472174 | 7,82783715 | 0,40280143 | 0,69296028 | 0,84791362 |
| 4878_at | NPPA | -0,04307821 | 5,34676921 | -0,40279935 | 0,69296178 | 0,84791362 |
| 2103_at | ESRRB | -0,03810939 | 4,18209506 | -0,40279725 | 0,69296329 | 0,84791362 |
| 54492_at | NEURL1B | 0,13828557 | 7,34415128 | 0,40278627 | 0,69297119 | 0,84791362 |
| 100127983_a | C8orf88 | -0,02803615 | 3,56380591 | -0,40261136 | 0,69309704 | 0,84791688 |

| | | | | | | |
|---|---|---|---|---|---|---|
| 64641_at | EBF2 | -0,03442307 | 3,53347026 | -0,40256947 | 0,69312718 | 0,84791688 |
| 285492_at | LINC00955 | -0,03596774 | 3,46917641 | -0,40254473 | 0,69314498 | 0,84791688 |
| 139411_at | PTCHD1 | -0,03440869 | 3,56209958 | -0,40253786 | 0,69314993 | 0,84791688 |
| 55970_at | GNG12 | 0,07115251 | 8,11536528 | 0,40239299 | 0,69325418 | 0,8479389 |
| 101929570_a | LOC10192957 | 0,0318405 | 3,0749124 | 0,4023905 | 0,69325597 | 0,8479389 |
| 9132_at | KCNQ4 | -0,02811251 | 4,52969502 | -0,40222804 | 0,69337288 | 0,84802805 |
| 6780_at | STAU1 | -0,03978278 | 10,842871 | -0,4020466 | 0,69350346 | 0,8481339 |
| 58524_at | DMRT3 | -0,03477862 | 3,81703759 | -0,40192204 | 0,69359311 | 0,8481897 |
| 4774_at | NFIA | -0,06158075 | 8,36178 | -0,40183564 | 0,6936553 | 0,84820277 |
| 26005_at | C2CD3 | 0,03833506 | 7,4920298 | 0,40178484 | 0,69369186 | 0,84820277 |
| 100506237_a | NKX2-1-AS1 | -0,02840753 | 3,8215002 | -0,40169254 | 0,6937583 | 0,84823016 |
| 55129_at | ANO10 | 0,07502689 | 7,93995392 | 0,40157003 | 0,69384649 | 0,8482327 |
| 400236_at | FOXN3-AS1 | 0,06849417 | 7,18909925 | 0,40156732 | 0,69384844 | 0,8482327 |
| 1123_at | CHN1 | 0,10088505 | 5,91116901 | 0,4012817 | 0,69405406 | 0,8482833 |
| 51208_at | CLDN18 | -0,02701188 | 4,32409246 | -0,40118429 | 0,69412419 | 0,8482833 |
| 581_at | BAX | -0,08467603 | 8,45154354 | -0,40115242 | 0,69414714 | 0,8482833 |
| 55028_at | C17orf80 | -0,02791771 | 6,78605815 | -0,40108661 | 0,69419452 | 0,8482833 |
| 55628_at | ZNF407 | 0,01992299 | 4,03732634 | 0,40108624 | 0,69419479 | 0,8482833 |
| 101928505_a | LOC10192850 | 0,09440638 | 3,84152073 | 0,40105983 | 0,6942138 | 0,8482833 |
| 439_at | ASNA1 | 0,0520683 | 9,46970719 | 0,40099832 | 0,69425809 | 0,8482833 |
| 90379_at | DCAF15 | -0,05999828 | 6,29376369 | -0,4009605 | 0,69428532 | 0,8482833 |
| 783_at | CACNB2 | -0,03101105 | 3,95606303 | -0,40095933 | 0,69428617 | 0,8482833 |
| 5948_at | RBP2 | -0,04922441 | 4,11836829 | -0,40073204 | 0,69444984 | 0,84842945 |
| 100130264_a | LOC10013026 | -0,04233883 | 3,42982968 | -0,40054643 | 0,6945835 | 0,84850474 |
| 3268_at | AGFG2 | 0,03298558 | 6,86590101 | 0,40052414 | 0,69459956 | 0,84850474 |
| 89822_at | KCNK17 | -0,05763948 | 4,38593483 | -0,40034001 | 0,69473217 | 0,84861292 |
| 285527_at | FRYL | 0,06788117 | 7,15263213 | 0,40010421 | 0,69490201 | 0,84874559 |
| 118461_at | C10orf71 | 0,03206832 | 4,00254688 | 0,40001135 | 0,6949689 | 0,84874559 |
| 56034_at | PDGFC | 0,15491596 | 6,36265182 | 0,40000571 | 0,69497296 | 0,84874559 |
| 5881_at | RAC3 | 0,04130684 | 5,9950923 | 0,39994295 | 0,69501817 | 0,84874699 |
| 6385_at | SDC4 | -0,06799055 | 10,8894081 | -0,39975223 | 0,69515556 | 0,84886096 |
| 79754_at | ASB13 | -0,02838351 | 7,91048894 | -0,39946209 | 0,69536459 | 0,84906239 |
| 6317_at | SERPINB3 | 0,06902952 | 5,28018078 | 0,39932103 | 0,69546623 | 0,84913267 |
| 84267_at | C9orf64 | 0,30995319 | 6,89822746 | 0,39914351 | 0,69559415 | 0,84923502 |
| 1903_at | S1PR3 | -0,08159526 | 6,77054353 | -0,3989542 | 0,69573058 | 0,84934775 |
| 57578_at | UNC79 | -0,03955107 | 4,39516215 | -0,39881454 | 0,69583123 | 0,84941679 |
| 54893_at | MTMR10 | 0,02454109 | 6,33026378 | 0,39827157 | 0,69622259 | 0,84984068 |
| 64849_at | SLC13A3 | -0,03467811 | 4,97060554 | -0,39799327 | 0,69642322 | 0,84998635 |
| 10302_at | SNAPC5 | -0,03330105 | 5,20992075 | -0,39798362 | 0,69643018 | 0,84998635 |
| 115209_at | OMA1 | 0,08701821 | 7,99801562 | 0,39785595 | 0,69652222 | 0,85004483 |
| 83795_at | KCNK16 | -0,05008468 | 5,90573779 | -0,39765849 | 0,6966646 | 0,85006895 |
| 51619_at | UBE2D4 | 0,02490703 | 7,39065131 | 0,39759662 | 0,69670921 | 0,85006895 |
| 134353_at | LSM11 | 0,02616977 | 5,76088911 | 0,39758968 | 0,69671421 | 0,85006895 |
| 93190_at | C1orf158 | 0,04513898 | 3,83363857 | 0,39756782 | 0,69672998 | 0,85006895 |
| 7994_at | KAT6A | 0,12143606 | 8,13722789 | 0,39752252 | 0,69676264 | 0,85006895 |
| 199990_at | FAAP20 | 0,02636933 | 5,57251062 | 0,39743087 | 0,69682873 | 0,85009575 |
| 64184_at | EDDM3B | 0,02545912 | 2,94722679 | 0,39732964 | 0,69690173 | 0,85013096 |
| 93058_at | COQ10A | 0,03897487 | 6,83310401 | 0,3970563 | 0,69709886 | 0,85031759 |
| 90135_at | BTBD6 | -0,03178149 | 9,57210566 | -0,39665379 | 0,69738919 | 0,8505716 |
| 339201_at | ASB16-AS1 | 0,09580161 | 6,85362663 | 0,39664516 | 0,69739542 | 0,8505716 |

| | | | | | | |
|---|---|---|---|---|---|---|
| 100287227_a | TIPARP-AS1 | -0,05111462 | 4,56594717 | -0,39654555 | 0,69746727 | 0,85060494 |
| 3459_at | IFNGR1 | 0,07375407 | 9,24597528 | 0,39648485 | 0,69751106 | 0,85060494 |
| 116349_at | EXOC3-AS1 | -0,08802439 | 5,85458655 | -0,39636664 | 0,69759634 | 0,85065508 |
| 1741_at | DLG3 | -0,0436039 | 6,90272574 | -0,39617273 | 0,69773624 | 0,85077182 |
| 10768_at | AHCYL1 | 0,04115503 | 10,2571919 | 0,39604236 | 0,69783031 | 0,85083266 |
| 11189_at | CELF3 | -0,05323598 | 6,17153889 | -0,39597329 | 0,69788014 | 0,85083957 |
| 100505555_a | LOC10050555 | -0,02842198 | 4,48061311 | -0,39575756 | 0,69803582 | 0,85092922 |
| 55686_at | MREG | -0,05695449 | 7,11417376 | -0,39574896 | 0,69804202 | 0,85092922 |
| 4478_at | MSN | -0,04930887 | 6,42691123 | -0,39533061 | 0,69834394 | 0,8512434 |
| 57537_at | SORCS2 | 0,18027265 | 5,81898618 | 0,39511901 | 0,69849668 | 0,85130001 |
| 4711_at | NDUFB5 | -0,03973652 | 10,7705311 | -0,39511842 | 0,6984971 | 0,85130001 |
| 203069_at | R3HCC1 | -0,10518148 | 8,33820061 | -0,39494288 | 0,69862381 | 0,85130001 |
| 284340_at | CXCL17 | 0,11371312 | 6,33780598 | 0,39494206 | 0,6986244 | 0,85130001 |
| 166863_at | RBM46 | 0,02201757 | 3,1861868 | 0,39490045 | 0,69865444 | 0,85130001 |
| 92483_at | LDHAL6B | -0,02761897 | 3,70896312 | -0,39484642 | 0,69869345 | 0,85130001 |
| 101927641_a | LOC10192764 | -0,04228962 | 3,46450498 | -0,3948377 | 0,69869974 | 0,85130001 |
| 51426_at | POLK | 0,11519598 | 6,52428756 | 0,39455986 | 0,69890033 | 0,85142465 |
| 79018_at | GID4 | -0,15610297 | 6,42630983 | -0,39448698 | 0,69895294 | 0,85142465 |
| 10208_at | USPL1 | 0,03471443 | 7,69077827 | 0,39438939 | 0,69902341 | 0,85142465 |
| 54586_at | EQTN | -0,02399228 | 2,87743796 | -0,3943857 | 0,69902607 | 0,85142465 |
| 79719_at | AAGAB | -0,03540765 | 7,85492537 | -0,39434739 | 0,69905374 | 0,85142465 |
| 100144602_a | EPHA5-AS1 | -0,04403082 | 4,36238102 | -0,39423597 | 0,69913419 | 0,85142465 |
| 100505658_a | LOC10050565 | -0,02595555 | 3,58650943 | -0,39421192 | 0,69915155 | 0,85142465 |
| 200523_at | TEX37 | -0,05224909 | 4,82174784 | -0,39420626 | 0,69915564 | 0,85142465 |
| 63906_at | GPATCH3 | 0,03840296 | 6,8729023 | 0,39401381 | 0,69929462 | 0,85154006 |
| 221400_at | TDRD6 | 0,03843642 | 3,14476197 | 0,3938459 | 0,69941588 | 0,85163388 |
| 120534_at | ARL14EP | 0,05703777 | 7,16252035 | 0,3937441 | 0,6994894 | 0,85166957 |
| 55138_at | FAM90A1 | -0,05410826 | 4,94959011 | -0,39356208 | 0,69962087 | 0,8517758 |
| 145482_at | PTGR2 | -0,12309826 | 4,36567121 | -0,39343722 | 0,69971105 | 0,85178486 |
| 346007_at | EYS | 0,0199332 | 2,94886805 | 0,3934063 | 0,69973338 | 0,85178486 |
| 4994_at | OR3A1 | -0,05308821 | 3,72349987 | -0,39336812 | 0,69976096 | 0,85178486 |
| 23551_at | RASD2 | -0,05062221 | 5,76191853 | -0,39328041 | 0,69982432 | 0,85180816 |
| 4761_at | NEUROD2 | -0,03853468 | 5,73922864 | -0,39304678 | 0,6999931 | 0,85195976 |
| 84446_at | BRSK1 | -0,05151903 | 5,17262963 | -0,39297694 | 0,70004355 | 0,85196734 |
| 8888_at | MCM3AP | -0,0536294 | 7,33053191 | -0,3926695 | 0,70026568 | 0,85218383 |
| 126006_at | PCP2 | 0,06366099 | 4,76681983 | 0,39254747 | 0,70035386 | 0,8522373 |
| 476_at | ATP1A1 | 0,02414627 | 10,8966495 | 0,39243298 | 0,7004366 | 0,85228414 |
| 5187_at | PER1 | 0,03661196 | 6,15938062 | 0,39217803 | 0,70062084 | 0,8524538 |
| 8476_at | CDC42BPA | 0,07864881 | 7,78197126 | 0,39207949 | 0,70069205 | 0,8524538 |
| 100129427_a | DDC-AS1 | -0,02894536 | 4,68033546 | -0,39191801 | 0,70080876 | 0,8524538 |
| 121536_at | AEBP2 | -0,02985835 | 6,72538969 | -0,39186135 | 0,70084972 | 0,8524538 |
| 167465_at | ZNF366 | 0,02523528 | 3,39243145 | 0,39178796 | 0,70090277 | 0,8524538 |
| 341880_at | SLC35F4 | 0,02437107 | 3,06487934 | 0,39174474 | 0,70093401 | 0,8524538 |
| 7179_at | TPTE | -0,03454203 | 2,97726216 | -0,39171938 | 0,70095234 | 0,8524538 |
| 9947_at | MAGEC1 | 0,02731042 | 3,56240675 | 0,39163179 | 0,70101566 | 0,8524538 |
| 83989_at | FAM172A | 0,14907908 | 6,4344965 | 0,39159822 | 0,70103992 | 0,8524538 |
| 267012_at | DAOA | -0,02384467 | 3,21556599 | -0,39159674 | 0,70104099 | 0,8524538 |
| 285423_at | LOC285423 | 0,04567031 | 3,48413726 | 0,39156549 | 0,70106358 | 0,8524538 |
| 7066_at | THPO | -0,02800805 | 3,99074213 | -0,3914546 | 0,70114375 | 0,8524538 |
| 642757_at | TBC1D22A-AS | 0,03713453 | 3,72405442 | 0,39142439 | 0,70116558 | 0,8524538 |

| | | | | | | |
|---|---|---|---|---|---|---|
| 26027_at | ACOT11 | 0,04813666 | 7,42813207 | 0,3913829 | 0,70119558 | 0,8524538 |
| 3119_at | HLA-DQB1 | -0,02262861 | 3,55026164 | -0,39113851 | 0,70137226 | 0,85259344 |
| 389206_at | BEND4 | -0,03029686 | 3,57921987 | -0,39110159 | 0,70139896 | 0,85259344 |
| 100506963_a | TCEB3-AS1 | 0,06721733 | 6,4325818 | 0,39098071 | 0,70148636 | 0,85264362 |
| 158131_at | OR1Q1 | -0,04045298 | 4,2524638 | -0,39092205 | 0,70152877 | 0,85264362 |
| 10614_at | HEXIM1 | 0,029906 | 8,8669609 | 0,3908396 | 0,7015884 | 0,85266229 |
| 257062_at | CATSPERD | 0,03084642 | 3,36521617 | 0,39077071 | 0,70163821 | 0,85266904 |
| 26873_at | OPLAH | 0,12292862 | 5,92155987 | 0,39063787 | 0,70173428 | 0,85273198 |
| 8660_at | IRS2 | 0,06021164 | 7,41449565 | 0,39017828 | 0,70206667 | 0,85304324 |
| 3656_at | IRAK2 | -0,08576074 | 6,86092655 | -0,39016124 | 0,70207899 | 0,85304324 |
| 6242_at | RTKN | 0,03706775 | 4,69376862 | 0,38995618 | 0,70222732 | 0,85316965 |
| 574432_at | IBA57-AS1 | -0,03534516 | 4,45480555 | -0,38980996 | 0,7023331 | 0,85324435 |
| 145645_at | C15orf43 | 0,02487357 | 2,82294654 | 0,3897152 | 0,70240165 | 0,85327382 |
| 51256_at | TBC1D7 | -0,08267859 | 8,20611734 | -0,38943127 | 0,70260707 | 0,85346954 |
| 2885_at | GRB2 | -0,05085468 | 9,08898596 | -0,389249 | 0,70273895 | 0,85357591 |
| 26205_at | GMEB2 | -0,0562436 | 8,45815863 | -0,38910823 | 0,70284081 | 0,85362241 |
| 286464_at | CFAP47 | -0,02102833 | 3,13818994 | -0,38906103 | 0,70287496 | 0,85362241 |
| 285203_at | EOGT | 0,243199 | 7,57021638 | 0,38901238 | 0,70291017 | 0,85362241 |
| 2044_at | EPHA5 | -0,02632603 | 3,228555 | -0,38870845 | 0,70313012 | 0,85383569 |
| 1043_at | CD52 | 0,05966768 | 4,69870293 | 0,3885631 | 0,70323532 | 0,85390961 |
| 390066_at | OR52D1 | -0,04764493 | 4,46230972 | -0,38826699 | 0,70344966 | 0,8540135 |
| 59345_at | GNB4 | -0,17402368 | 6,87345814 | -0,38824082 | 0,7034686 | 0,8540135 |
| 101928271_a | LOC10192827 | -0,03124593 | 2,66049391 | -0,38822292 | 0,70348156 | 0,8540135 |
| 101927058_a | LOC10192705 | -0,03571531 | 3,10919236 | -0,38817593 | 0,70351557 | 0,8540135 |
| 90167_at | FRMD7 | 0,02552509 | 3,6388163 | 0,38813866 | 0,70354255 | 0,8540135 |
| 2829_at | XCR1 | 0,04913874 | 4,84289255 | 0,38770915 | 0,70385351 | 0,85433712 |
| 5015_at | OTX2 | 0,02550023 | 3,47148569 | 0,38755594 | 0,70396444 | 0,85441793 |
| 119391_at | GSTO2 | 0,17750025 | 6,84867034 | 0,38740184 | 0,70407602 | 0,85443424 |
| 57824_at | HMHB1 | -0,03052063 | 4,32272054 | -0,38734268 | 0,70411887 | 0,85443424 |
| 728537_at | LINC01125 | -0,04260165 | 4,06535002 | -0,38723978 | 0,70419338 | 0,85443424 |
| 27022_at | FOXD3 | 0,03194648 | 4,46628691 | 0,38721062 | 0,7042145 | 0,85443424 |
| 339524_at | LINC01140 | 0,03373143 | 4,23712414 | 0,38718253 | 0,70423484 | 0,85443424 |
| 100506465_a | LINC01234 | -0,11308086 | 3,5598294 | -0,38716987 | 0,70424401 | 0,85443424 |
| 51390_at | AIG1 | 0,04570031 | 5,21913751 | 0,38697787 | 0,70438307 | 0,85449611 |
| 132789_at | GNPDA2 | -0,05468694 | 6,99005199 | -0,38697695 | 0,70438373 | 0,85449611 |
| 64763_at | ZNF574 | -0,03800324 | 7,40278168 | -0,38678233 | 0,70452469 | 0,85461329 |
| 142684_at | RAB40A | -0,02986921 | 3,05810962 | -0,38667125 | 0,70460515 | 0,85465707 |
| 8139_at | GAN | 0,06886119 | 7,33155225 | 0,38655287 | 0,7046909 | 0,85470727 |
| 125113_at | KRT222 | 0,0222422 | 2,75516954 | 0,38644731 | 0,70476737 | 0,85474619 |
| 8714_at | ABCC3 | 0,0703975 | 8,36053133 | 0,38628046 | 0,70488824 | 0,85483896 |
| 51068_at | NMD3 | -0,06652626 | 9,94876034 | -0,38612181 | 0,70500318 | 0,85492453 |
| 22844_at | FRMPD1 | -0,0446977 | 4,22525269 | -0,3859234 | 0,70514693 | 0,85504503 |
| 81608_at | FIP1L1 | -0,07571361 | 7,25730231 | -0,38570978 | 0,70530173 | 0,85512817 |
| 4682_at | NUBP1 | -0,04355831 | 8,55118677 | -0,38570625 | 0,70530428 | 0,85512817 |
| 91775_at | NXPE3 | 0,05650671 | 6,07100262 | 0,38551494 | 0,70544291 | 0,85524242 |
| 4255_at | MGMT | 0,08532066 | 6,99462467 | 0,38500728 | 0,70581085 | 0,8555472 |
| 22917_at | ZP1 | -0,03792218 | 4,85326205 | -0,38492634 | 0,70586952 | 0,8555472 |
| 401081_at | FLJ22763 | -0,03002044 | 3,03191515 | -0,38492061 | 0,70587367 | 0,8555472 |
| 129852_at | C2orf73 | 0,02988453 | 3,38940763 | 0,38488246 | 0,70590132 | 0,8555472 |
| 23439_at | ATP1B4 | -0,02528331 | 3,38391553 | -0,3848617 | 0,70591638 | 0,8555472 |

| | | | | | | |
|---|---|---|---|---|---|---|
| 29774_at | POM121L9P | -0,05525061 | 4,16709311 | -0,38468647 | 0,7060434 | 0,85563177 |
| 146378_at | C16orf92 | -0,0363244 | 5,02739021 | -0,38461125 | 0,70609793 | 0,85563177 |
| 157657_at | C8orf37 | -0,07811735 | 6,44649733 | -0,38458162 | 0,70611941 | 0,85563177 |
| 2492_at | FSHR | -0,03759994 | 3,83901591 | -0,3844631 | 0,70620533 | 0,85564828 |
| 7356_at | SCGB1A1 | -0,06601424 | 4,28845771 | -0,38444028 | 0,70622188 | 0,85564828 |
| 101927523_a | LOC10192752 | -0,03075723 | 3,77912146 | -0,38426426 | 0,7063495 | 0,85571961 |
| 79694_at | MANEA | -0,07197871 | 6,8566013 | -0,38422318 | 0,70637928 | 0,85571961 |
| 83892_at | KCTD10 | 0,02586592 | 8,21307465 | 0,38417528 | 0,70641402 | 0,85571961 |
| 53354_at | PANK1 | -0,05356255 | 7,00460382 | -0,38401749 | 0,70652843 | 0,85575136 |
| 124245_at | ZC3H18 | -0,06300515 | 7,02866943 | -0,38397366 | 0,70656022 | 0,85575136 |
| 8685_at | MARCO | -0,05589534 | 5,13927114 | -0,38395534 | 0,7065735 | 0,85575136 |
| 30846_at | EHD2 | 0,11159056 | 7,85577639 | 0,38374426 | 0,70672657 | 0,85588294 |
| 100874063_a | CLYBL-AS2 | 0,03005002 | 2,94912716 | 0,38361901 | 0,70681741 | 0,85593913 |
| 92906_at | HNRNPLL | 0,0497124 | 7,63511431 | 0,38327312 | 0,70706828 | 0,85618911 |
| 162461_at | TMEM92 | -0,06367021 | 6,11408635 | -0,38298929 | 0,70727417 | 0,85630287 |
| 101928022_a | LINC01214 | -0,04521206 | 5,09850762 | -0,38295031 | 0,70730245 | 0,85630287 |
| 27145_at | FILIP1 | 0,02571147 | 3,33358365 | 0,38290137 | 0,70733795 | 0,85630287 |
| 8796_at | SCEL | -0,36498741 | 5,46880081 | -0,38289849 | 0,70734004 | 0,85630287 |
| 3354_at | HTR1E | 0,04398824 | 4,64673798 | 0,38272703 | 0,70746443 | 0,85633801 |
| 55132_at | LARP1B | 0,04585359 | 6,02335222 | 0,3826541 | 0,70751734 | 0,85633801 |
| 1720_at | LOC1720 | 0,03110615 | 3,11990097 | 0,38259089 | 0,7075632 | 0,85633801 |
| 27285_at | TEKT2 | -0,04886918 | 4,95389386 | -0,38258779 | 0,70756545 | 0,85633801 |
| 5473_at | PPBP | 0,02728447 | 3,28239197 | 0,38252539 | 0,70761073 | 0,85633801 |
| 113452_at | TMEM54 | -0,08250509 | 9,23040734 | -0,38249084 | 0,7076358 | 0,85633801 |
| 7095_at | SEC62 | 0,0391559 | 7,94268094 | 0,38237813 | 0,70771758 | 0,85635352 |
| 393_at | ARHGAP4 | 0,02800131 | 5,84012086 | 0,38235063 | 0,70773753 | 0,85635352 |
| 51522_at | TMEM14C | 0,06963385 | 9,86954946 | 0,38215466 | 0,70787974 | 0,85645692 |
| 100128126_a | STAU2-AS1 | -0,05654304 | 4,51318035 | -0,38207468 | 0,70793778 | 0,85645692 |
| 102723831_a | LOC10272383 | -0,02616016 | 4,84343027 | -0,38204906 | 0,70795637 | 0,85645692 |
| 26063_at | DECR2 | 0,03254647 | 7,70755189 | 0,38198204 | 0,70800501 | 0,85646198 |
| 9110_at | MTMR4 | -0,02585182 | 8,46374979 | -0,38178373 | 0,70814893 | 0,85658229 |
| 152742_at | LINC01085 | 0,03714374 | 3,44965545 | 0,38169361 | 0,70821435 | 0,85660762 |
| 55879_at | GABRQ | 0,04357446 | 5,80878333 | 0,3816017 | 0,70828106 | 0,85663452 |
| 79571_at | GCC1 | 0,02953063 | 7,00111162 | 0,3814602 | 0,70838376 | 0,85670495 |
| 100507501_a | LINC01569 | 0,04861141 | 6,02327653 | 0,38121142 | 0,70856436 | 0,85686248 |
| 2624_at | GATA2 | -0,04194005 | 8,44670234 | -0,38109146 | 0,70865145 | 0,85686248 |
| 8839_at | WISP2 | -0,07879545 | 5,74074766 | -0,38104757 | 0,70868331 | 0,85686248 |
| 28670_at | TRAV13-2 | 0,03775174 | 3,138083 | 0,38098169 | 0,70873114 | 0,85686248 |
| 10096_at | ACTR3 | 0,05572473 | 8,73219303 | 0,3809744 | 0,70873644 | 0,85686248 |
| 3458_at | IFNG | -0,02154913 | 2,99191442 | -0,3807895 | 0,70887068 | 0,856971 |
| 340390_at | KIAA1875 | 0,04011207 | 5,25324953 | 0,38064535 | 0,70897535 | 0,85704031 |
| 57829_at | ZP4 | -0,05328443 | 3,66210092 | -0,380588 | 0,70901699 | 0,85704031 |
| 55613_at | MTMR8 | -0,03259291 | 3,45256635 | -0,38018996 | 0,70930605 | 0,85730017 |
| 101929441_a | LOC10192944 | -0,05665103 | 4,56437904 | -0,3801694 | 0,70932098 | 0,85730017 |
| 101927438_a | LOC10192743 | -0,0365582 | 3,69073279 | -0,38003147 | 0,70942116 | 0,85736745 |
| 6247_at | RS1 | -0,02667232 | 3,76422198 | -0,37986195 | 0,70954428 | 0,85746245 |
| 10817_at | FRS3 | -0,02915335 | 5,4401306 | -0,37944734 | 0,70984546 | 0,85777142 |
| 130612_at | TMEM198 | -0,05033923 | 5,76087671 | -0,37927657 | 0,70996952 | 0,85777142 |
| 10806_at | SDCCAG8 | 0,06184243 | 5,34011438 | 0,37924377 | 0,70999335 | 0,85777142 |
| 6364_at | CCL20 | 0,27904199 | 5,34629277 | 0,37923957 | 0,70999641 | 0,85777142 |

| | | | | | | |
|---|---|---|---|---|---|---|
| 84081_at | NSRP1 | -0,07358688 | 7,61631263 | -0,37920352 | 0,7100226 | 0,85777142 |
| 80380_at | PDCD1LG2 | -0,0478072 | 4,41338287 | -0,37877903 | 0,71033103 | 0,85807545 |
| 283212_at | KLHL35 | 0,10538582 | 5,47685119 | 0,37872021 | 0,71037378 | 0,85807545 |
| 57683_at | ZDBF2 | 0,12737989 | 4,59775357 | 0,37867325 | 0,7104079 | 0,85807545 |
| 22850_at | ADNP2 | -0,05518843 | 8,03823755 | -0,37855128 | 0,71049654 | 0,85810485 |
| 3607_at | FOXK2 | -0,0425358 | 8,31312652 | -0,37844487 | 0,71057387 | 0,85810485 |
| 128864_at | C20orf144 | -0,05477084 | 5,09776387 | -0,3783683 | 0,71062952 | 0,85810485 |
| 23761_at | PISD | 0,06340922 | 7,80843541 | 0,37836305 | 0,71063334 | 0,85810485 |
| 57139_at | RGL3 | 0,0335095 | 5,67586066 | 0,37833328 | 0,71065497 | 0,85810485 |
| 29802_at | VPREB3 | -0,03460207 | 5,00295899 | -0,3780296 | 0,7108757 | 0,85822993 |
| 5236_at | PGM1 | -0,2224122 | 8,35528122 | -0,37796967 | 0,71091926 | 0,85822993 |
| 64776_at | C11orf1 | 0,02644787 | 8,01762917 | 0,3779598 | 0,71092644 | 0,85822993 |
| 128025_at | WDR64 | 0,03003042 | 3,44617349 | 0,37790619 | 0,7109654 | 0,85822993 |
| 51100_at | SH3GLB1 | 0,04311384 | 9,61628334 | 0,3778843 | 0,71098132 | 0,85822993 |
| 9997_at | SCO2 | -0,08828795 | 8,03082614 | -0,37773109 | 0,7110927 | 0,85826998 |
| 124590_at | USH1G | -0,03643608 | 6,53520805 | -0,37761396 | 0,71117785 | 0,85826998 |
| 7529_at | YWHAB | 0,02429946 | 12,0495677 | 0,37756385 | 0,71121428 | 0,85826998 |
| 161357_at | MDGA2 | 0,02210882 | 2,94044138 | 0,37740786 | 0,71132769 | 0,85826998 |
| 79442_at | LRRC2 | 0,02057538 | 2,90834175 | 0,37738743 | 0,71134255 | 0,85826998 |
| 132724_at | TMPRSS11B | -0,03306376 | 2,99749284 | -0,37732082 | 0,71139098 | 0,85826998 |
| 590_at | BCHE | -0,0294224 | 3,02199306 | -0,37729474 | 0,71140994 | 0,85826998 |
| 383_at | ARG1 | -0,02344792 | 3,53956875 | -0,37729115 | 0,71141255 | 0,85826998 |
| 283237_at | TTC9C | 0,03740233 | 7,84948813 | 0,37728709 | 0,7114155 | 0,85826998 |
| 24144_at | TFIP11 | -0,02671918 | 8,11413042 | -0,37661535 | 0,71190398 | 0,85877711 |
| 757_at | TMEM50B | 0,06862235 | 8,06634664 | 0,37657272 | 0,71193499 | 0,85877711 |
| 255626_at | HIST1H2BA | -0,03048929 | 2,71077112 | -0,37648619 | 0,71199793 | 0,85877711 |
| 84624_at | FNDC1 | -0,03194058 | 3,50444785 | -0,37646384 | 0,71201418 | 0,85877711 |
| 6508_at | SLC4A3 | 0,07554573 | 6,01026761 | 0,37639319 | 0,71206557 | 0,85878531 |
| 100507466_a | DPH6-AS1 | 0,02310155 | 3,38012811 | 0,37624198 | 0,71217555 | 0,85886052 |
| 7732_at | RNF112 | -0,03635971 | 4,88652164 | -0,37618488 | 0,71221709 | 0,85886052 |
| 5469_at | MED1 | 0,04689231 | 8,49884003 | 0,37590037 | 0,71242406 | 0,85894618 |
| 4241_at | MFI2 | 0,06100689 | 6,17422362 | 0,3757754 | 0,71251498 | 0,85894618 |
| 54811_at | ZNF562 | -0,05527812 | 6,83990167 | -0,37570989 | 0,71256264 | 0,85894618 |
| 3996_at | LLGL1 | 0,06984143 | 7,53662033 | 0,3756977 | 0,71257151 | 0,85894618 |
| 101929620_a | LINC01243 | -0,02361688 | 3,26176419 | -0,37567184 | 0,71259033 | 0,85894618 |
| 2651_at | GCNT2 | 0,23074167 | 6,14943901 | 0,37564067 | 0,712613 | 0,85894618 |
| 1844_at | DUSP2 | 0,25061107 | 6,97576302 | 0,37559545 | 0,71264591 | 0,85894618 |
| 285382_at | C3orf70 | 0,18775143 | 5,0796528 | 0,37552023 | 0,71270064 | 0,85894618 |
| 8065_at | CUL5 | 0,03730048 | 7,21466215 | 0,3754368 | 0,71276134 | 0,85894618 |
| 100507462_a | GVQW2 | 0,04945291 | 3,6900496 | 0,37541768 | 0,71277525 | 0,85894618 |
| 9619_at | ABCG1 | -0,04433196 | 5,54819601 | -0,37540084 | 0,7127875 | 0,85894618 |
| 4660_at | PPP1R12B | 0,04994227 | 5,73058953 | 0,37535176 | 0,71282322 | 0,85894618 |
| 2307_at | FOXS1 | -0,04000431 | 4,60938539 | -0,37520018 | 0,71293352 | 0,85902536 |
| 3712_at | IVD | -0,03391778 | 7,8022029 | -0,37501894 | 0,71306541 | 0,8591171 |
| 286190_at | LACTB2-AS1 | -0,02322259 | 2,95117059 | -0,37490654 | 0,71314721 | 0,8591171 |
| 192666_at | KRT24 | -0,0352854 | 4,54387288 | -0,37489926 | 0,71315252 | 0,8591171 |
| 1294_at | COL7A1 | 0,19232036 | 7,44198169 | 0,37475019 | 0,71326101 | 0,8591171 |
| 158314_at | LINC00475 | 0,04276202 | 3,81401373 | 0,37466538 | 0,71332274 | 0,8591171 |
| 148870_at | CCDC27 | -0,04151145 | 4,65815896 | -0,37461691 | 0,71335802 | 0,8591171 |
| 53942_at | CNTN5 | -0,02234522 | 3,36617569 | -0,37460134 | 0,71336935 | 0,8591171 |

| | | | | | | |
|---|---|---|---|---|---|---|
| 63939_at | FAM217B | 0,10515502 | 8,05371029 | 0,37444576 | 0,7134826 | 0,8591171 |
| 100130522_a | PARD6G-AS1 | -0,02860887 | 5,66045319 | -0,37434108 | 0,7135588 | 0,8591171 |
| 5935_at | RBM3 | -0,06344554 | 8,55873858 | -0,37427636 | 0,71360592 | 0,8591171 |
| 221074_at | SLC39A12 | -0,03865396 | 2,90604607 | -0,37427552 | 0,71360652 | 0,8591171 |
| 6934_at | TCF7L2 | -0,03367482 | 7,8461966 | -0,37422819 | 0,71364098 | 0,8591171 |
| 284257_at | BOD1L2 | -0,03415254 | 4,83275059 | -0,37421412 | 0,71365122 | 0,8591171 |
| 5747_at | PTK2 | 0,09819711 | 9,36538771 | 0,37409194 | 0,71374017 | 0,8591171 |
| 203522_at | DDX26B | 0,19819212 | 5,49222203 | 0,3740667 | 0,71375855 | 0,8591171 |
| 5346_at | PLIN1 | -0,05110473 | 6,03404723 | -0,37402904 | 0,71378596 | 0,8591171 |
| 101927950_a | LOC10192795 | -0,04714436 | 5,02312376 | -0,3738138 | 0,71394267 | 0,8591171 |
| 10699_at | CORIN | 0,02173095 | 3,86395757 | 0,37378697 | 0,71396221 | 0,8591171 |
| 11148_at | HHLA2 | -0,02478014 | 3,06614652 | -0,37364114 | 0,71406839 | 0,8591171 |
| 101929486_a | LOC10192948 | -0,03723906 | 3,26926895 | -0,37362658 | 0,71407899 | 0,8591171 |
| 59269_at | HIVEP3 | -0,02856612 | 5,35826484 | -0,37361908 | 0,71408446 | 0,8591171 |
| 2891_at | GRIA2 | 0,01863125 | 2,92218007 | 0,37361357 | 0,71408847 | 0,8591171 |
| 201798_at | TIGD4 | 0,02351659 | 3,45555223 | 0,3734709 | 0,71419236 | 0,8591171 |
| 653808_at | ZG16 | 0,02904265 | 4,62969713 | 0,37346358 | 0,71419769 | 0,8591171 |
| 4212_at | MEIS2 | 0,05848758 | 8,82955233 | 0,37336698 | 0,71426803 | 0,8591171 |
| 23523_at | CABIN1 | 0,02392338 | 6,75047139 | 0,37332609 | 0,71429781 | 0,8591171 |
| 126859_at | AXDND1 | 0,07131419 | 4,50289533 | 0,3733187 | 0,71430319 | 0,8591171 |
| 420_at | ART4 | -0,04388874 | 3,1053772 | -0,37330253 | 0,71431497 | 0,8591171 |
| 50651_at | SLC45A1 | -0,0355543 | 5,0064189 | -0,37327656 | 0,71433388 | 0,8591171 |
| 57121_at | LPAR5 | 0,05177326 | 6,98309309 | 0,37325767 | 0,71434764 | 0,8591171 |
| 353514_at | LILRA5 | -0,02632722 | 3,50694092 | -0,37296769 | 0,71455883 | 0,85931744 |
| 400620_at | LOC400620 | -0,02088668 | 3,74197735 | -0,37279011 | 0,71468818 | 0,85941934 |
| 400710_at | FLJ26850 | 0,02881888 | 4,01804473 | 0,37259968 | 0,71482689 | 0,85953248 |
| 222235_at | FBXL13 | 0,04136607 | 3,23028978 | 0,37229986 | 0,71504531 | 0,85974145 |
| 150572_at | SMYD1 | -0,03748001 | 4,29779612 | -0,37211167 | 0,71518242 | 0,85983512 |
| 191585_at | PLAC4 | -0,03135789 | 4,52453202 | -0,37206536 | 0,71521616 | 0,85983512 |
| 3487_at | IGFBP4 | 0,31735116 | 8,49416498 | 0,37200914 | 0,71525712 | 0,85983512 |
| 4923_at | NTSR1 | -0,050259 | 5,25249616 | -0,37187969 | 0,71535145 | 0,8598483 |
| 85015_at | USP45 | -0,04768569 | 5,96118763 | -0,37187157 | 0,71535736 | 0,8598483 |
| 9639_at | ARHGEF10 | 0,05975374 | 8,17918325 | 0,37175531 | 0,71544208 | 0,85985829 |
| 5082_at | PDCL | 0,02853747 | 7,38365954 | 0,37173149 | 0,71545944 | 0,85985829 |
| 10954_at | PDIA5 | -0,08178551 | 8,03013794 | -0,37167639 | 0,71549959 | 0,85985829 |
| 101928781_a | TMEM26-AS1 | 0,02530863 | 2,87274387 | 0,37148922 | 0,71563599 | 0,85995919 |
| 28658_at | TRAV25 | 0,02893626 | 2,94954117 | 0,37143867 | 0,71567283 | 0,85995919 |
| 101928893_a | LOC10192889 | 0,02949929 | 2,84003893 | 0,37108271 | 0,71593226 | 0,86016396 |
| 168400_at | DDX53 | 0,02572251 | 3,03698118 | 0,37108231 | 0,71593256 | 0,86016396 |
| 6500_at | SKP1 | -0,03586295 | 8,61959488 | -0,37089249 | 0,71607092 | 0,8601972 |
| 5579_at | PRKCB | 0,02147641 | 4,66309463 | 0,37082099 | 0,71612304 | 0,8601972 |
| 11083_at | DIDO1 | 0,04102682 | 7,50740679 | 0,3707647 | 0,71616408 | 0,8601972 |
| 79981_at | FRMD1 | -0,04105872 | 5,23570828 | -0,37075502 | 0,71617113 | 0,8601972 |
| 201853_at | LINC00504 | -0,02342369 | 3,49589217 | -0,37073805 | 0,7161835 | 0,8601972 |
| 347344_at | ZNF81 | -0,02280816 | 5,03941683 | -0,37040866 | 0,71642364 | 0,86027047 |
| 23114_at | NFASC | 0,02120202 | 4,68782683 | 0,3703859 | 0,71644023 | 0,86027047 |
| 931_at | MS4A1 | -0,02472751 | 3,4599526 | -0,37034819 | 0,71646772 | 0,86027047 |
| 55167_at | MSL2 | -0,03691331 | 9,44856752 | -0,37029172 | 0,7165089 | 0,86027047 |
| 6748_at | SSR4 | 0,05079109 | 10,0240724 | 0,37021331 | 0,71656607 | 0,86027047 |
| 57188_at | ADAMTSL3 | -0,01968402 | 3,75219998 | -0,37020953 | 0,71656882 | 0,86027047 |

| | | | | | | |
|---|---|---|---|---|---|---|
| 10666_at | CD226 | -0,02266976 | 2,96472048 | -0,3700657 | 0,7166737 | 0,86027047 |
| 150967_at | PKI55 | 0,05102827 | 6,03517592 | 0,37005531 | 0,71668128 | 0,86027047 |
| 115827_at | RAB3C | -0,02599973 | 3,19892023 | -0,36997788 | 0,71673774 | 0,86027047 |
| 130507_at | UBR3 | -0,04274659 | 8,9477068 | -0,36995579 | 0,71675385 | 0,86027047 |
| 6751_at | SSTR1 | -0,02827326 | 4,08665044 | -0,36992294 | 0,7167778 | 0,86027047 |
| 5456_at | POU3F4 | 0,03709953 | 4,16856032 | 0,36991936 | 0,71678041 | 0,86027047 |
| 27040_at | LAT | 0,0325682 | 5,87260619 | 0,36971803 | 0,71692724 | 0,86034268 |
| 2017_at | CTTN | -0,02888252 | 9,52679322 | -0,36954324 | 0,71705471 | 0,86034268 |
| 1144_at | CHRND | -0,03834369 | 4,42635677 | -0,36952864 | 0,71706536 | 0,86034268 |
| 29765_at | TMOD4 | -0,04739173 | 4,41357031 | -0,3694859 | 0,71709653 | 0,86034268 |
| 91523_at | PCED1B | -0,22144429 | 7,24731653 | -0,36947236 | 0,7171064 | 0,86034268 |
| 25893_at | TRIM58 | 0,02981138 | 3,76438054 | 0,36940077 | 0,71715862 | 0,86034268 |
| 100128822_a | LINC01003 | 0,06241488 | 7,41400731 | 0,36939855 | 0,71716024 | 0,86034268 |
| 100505964_a | LINC01304 | -0,02505696 | 3,84896749 | -0,369343 | 0,71720075 | 0,86034268 |
| 144811_at | LACC1 | 0,12996671 | 4,67310826 | 0,36928571 | 0,71724254 | 0,86034268 |
| 9443_at | MED7 | 0,03290598 | 7,7449568 | 0,36911666 | 0,71736585 | 0,86041015 |
| 23493_at | HEY2 | -0,0735356 | 4,96732072 | -0,3690767 | 0,717395 | 0,86041015 |
| 7058_at | THBS2 | -0,04916968 | 4,4677118 | -0,3690037 | 0,71744825 | 0,86041015 |
| 51133_at | KCTD3 | 0,07590922 | 8,69567532 | 0,36896367 | 0,71747745 | 0,86041015 |
| 400680_at | LINC00664 | -0,02756276 | 3,1767692 | -0,36887647 | 0,71754106 | 0,86043286 |
| 100507410_a | C1QTNF1-AS1 | 0,03335712 | 4,15390599 | 0,36837703 | 0,71790545 | 0,86076001 |
| 29855_at | UBN1 | 0,03443462 | 7,75102314 | 0,36831329 | 0,71795196 | 0,86076001 |
| 140890_at | SREK1 | -0,06886192 | 6,55831257 | -0,36830976 | 0,71795454 | 0,86076001 |
| 9860_at | LRIG2 | 0,0397269 | 6,34263192 | 0,36825758 | 0,71799262 | 0,86076001 |
| 126069_at | ZNF491 | -0,05665046 | 4,90920966 | -0,36778147 | 0,71834006 | 0,86112295 |
| 259283_at | MDS2 | 0,03062229 | 4,24393188 | 0,36771315 | 0,71838992 | 0,86112913 |
| 10745_at | PHTF1 | 0,07110083 | 7,12462123 | 0,36760513 | 0,71846876 | 0,86117004 |
| 79630_at | C1orf54 | -0,03285694 | 4,69279178 | -0,36749041 | 0,7185525 | 0,86121682 |
| 101928035_a | LOC10192803 | -0,0268442 | 3,14679789 | -0,36726315 | 0,71871838 | 0,86131461 |
| 54797_at | MED18 | 0,0359544 | 6,21330967 | 0,36725134 | 0,71872701 | 0,86131461 |
| 101927502_a | LOC10192750 | -0,03727859 | 4,0596581 | -0,36719487 | 0,71876823 | 0,86131461 |
| 251_at | ALPPL2 | 0,14658053 | 6,27026484 | 0,36708041 | 0,71885179 | 0,86136116 |
| 284615_at | ANKRD34A | 0,04079406 | 5,04103233 | 0,36679562 | 0,7190597 | 0,86155669 |
| 23208_at | SYT11 | 0,02639746 | 3,7290369 | 0,36667808 | 0,71914552 | 0,86160593 |
| 203414_at | LINC01560 | 0,07839288 | 5,68439582 | 0,36647531 | 0,71929358 | 0,86172972 |
| 9476_at | NAPSA | -0,05686994 | 5,59196888 | -0,36622535 | 0,71947611 | 0,86188345 |
| 3232_at | HOXD3 | -0,03456849 | 4,60358408 | -0,36609274 | 0,71957295 | 0,86188345 |
| 26002_at | MOXD1 | 0,04643127 | 4,40490396 | 0,36608912 | 0,71957559 | 0,86188345 |
| 377_at | ARF3 | 0,03213613 | 9,62112589 | 0,36605299 | 0,71960198 | 0,86188345 |
| 23524_at | SRRM2 | 0,03419482 | 8,75606367 | 0,36599325 | 0,71964561 | 0,86188345 |
| 599_at | BCL2L2 | 0,02485754 | 6,39267007 | 0,36544309 | 0,72004746 | 0,86220871 |
| 163933_at | FAM43B | 0,03190301 | 5,85935656 | 0,3653497 | 0,72011568 | 0,86220871 |
| 3346_at | HTN1 | 0,03508291 | 2,68939349 | 0,36531388 | 0,72014185 | 0,86220871 |
| 201134_at | CEP112 | 0,02766353 | 3,85673182 | 0,36530753 | 0,72014648 | 0,86220871 |
| 25861_at | DFNB31 | 0,02423768 | 5,74255102 | 0,36525661 | 0,72018369 | 0,86220871 |
| 4750_at | NEK1 | 0,15710985 | 6,26953401 | 0,36522594 | 0,72020609 | 0,86220871 |
| 3321_at | IGSF3 | 0,0706036 | 7,16602629 | 0,36519252 | 0,72023051 | 0,86220871 |
| 657_at | BMPR1A | -0,05122205 | 7,33537066 | -0,36494081 | 0,72041441 | 0,86233841 |
| 51128_at | SAR1B | -0,04549865 | 8,37938857 | -0,36492168 | 0,72042839 | 0,86233841 |
| 9808_at | KIAA0087 | 0,03504974 | 3,38908581 | 0,36458911 | 0,72067139 | 0,86257567 |

| | | | | | | |
|---|---|---|---|---|---|---|
| 162998_at | OR7D2 | 0,14100064 | 4,00076488 | 0,36443403 | 0,72078472 | 0,86262262 |
| 100169989_a | DBIL5P2 | -0,03625555 | 3,46138679 | -0,36441288 | 0,72080017 | 0,86262262 |
| 10873_at | ME3 | 0,13600624 | 5,61797238 | 0,36429807 | 0,72088407 | 0,8626336 |
| 221833_at | SP8 | 0,02897611 | 3,87304917 | 0,36427778 | 0,72089891 | 0,8626336 |
| 6462_at | SHBG | 0,03334771 | 5,69719097 | 0,3640833 | 0,72104104 | 0,86271175 |
| 9150_at | CTDP1 | 0,02706221 | 7,00994263 | 0,36406585 | 0,72105379 | 0,86271175 |
| 123775_at | C16orf46 | 0,04122363 | 5,50581584 | 0,36395808 | 0,72113256 | 0,86275241 |
| 4168_at | MCF2 | -0,0196041 | 2,96480926 | -0,36381348 | 0,72123826 | 0,86282527 |
| 132430_at | PABPC4L | 0,02624184 | 3,26283318 | 0,36373804 | 0,72129339 | 0,86283765 |
| 643733_at | LOC643733 | -0,0296372 | 3,91448827 | -0,36364251 | 0,72136323 | 0,8628676 |
| 23086_at | EXPH5 | -0,18751334 | 6,51413349 | -0,36310693 | 0,72175478 | 0,86328235 |
| 348808_at | NPHP3-AS1 | -0,05631866 | 4,56920654 | -0,36288641 | 0,72191602 | 0,8634187 |
| 101927211_a | LOC10192721 | 0,02342147 | 3,03123313 | 0,36274309 | 0,72202082 | 0,8634187 |
| 153090_at | DAB2IP | 0,07957812 | 7,02766897 | 0,36268687 | 0,72206193 | 0,8634187 |
| 135250_at | RAET1E | 0,08881395 | 6,65942341 | 0,36267985 | 0,72206707 | 0,8634187 |
| 10082_at | GPC6 | -0,02960505 | 4,00854166 | -0,36264454 | 0,72209288 | 0,8634187 |
| 285311_at | C3orf56 | 0,03527091 | 3,53570811 | 0,36248661 | 0,72220838 | 0,86345312 |
| 56255_at | TMX4 | -0,02671081 | 8,45437645 | -0,36248259 | 0,72221132 | 0,86345312 |
| 283314_at | C1RL-AS1 | 0,02935793 | 4,85087387 | 0,36240538 | 0,72226779 | 0,86346704 |
| 126272_at | EID2B | 0,07710909 | 5,24661957 | 0,36221011 | 0,7224106 | 0,86356466 |
| 121256_at | TMEM132D | -0,03024148 | 4,54769572 | -0,36214825 | 0,72245585 | 0,86356466 |
| 29098_at | RANGRF | -0,11613374 | 6,89582251 | -0,36206038 | 0,72252012 | 0,86356466 |
| 5741_at | PTH | 0,0278797 | 2,81023852 | 0,36202826 | 0,72254362 | 0,86356466 |
| 80817_at | CEP44 | -0,05088198 | 5,26257389 | -0,36194275 | 0,72260616 | 0,86356466 |
| 5463_at | POU6F1 | 0,02671477 | 4,51569397 | 0,36190232 | 0,72263573 | 0,86356466 |
| 100506365_a | OTUD6B-AS1 | -0,03962342 | 7,66723562 | -0,3618647 | 0,72266326 | 0,86356466 |
| 51496_at | CTDSPL2 | -0,05306515 | 8,28046621 | -0,361575 | 0,72287519 | 0,86376433 |
| 80237_at | ELL3 | -0,02526993 | 5,40869039 | -0,3612279 | 0,72312914 | 0,86386262 |
| 56896_at | DPYSL5 | 0,03185092 | 4,51672255 | 0,36117206 | 0,72316999 | 0,86386262 |
| 10015_at | PDCD6IP | 0,09122903 | 9,59871402 | 0,3611482 | 0,72318745 | 0,86386262 |
| 5073_at | PARN | -0,0557122 | 8,46200966 | -0,36114676 | 0,7231885 | 0,86386262 |
| 4056_at | LTC4S | -0,04387066 | 6,2315419 | -0,36112816 | 0,72320211 | 0,86386262 |
| 753_at | LDLRAD4 | 0,02658956 | 4,61992355 | 0,36105299 | 0,72325711 | 0,86386262 |
| 115330_at | GPR146 | 0,02947377 | 4,60907786 | 0,3609938 | 0,72330043 | 0,86386262 |
| 65975_at | STK33 | -0,06063693 | 4,46759706 | -0,36093207 | 0,7233456 | 0,86386262 |
| 6729_at | SRP54 | 0,03795314 | 9,08496816 | 0,36089005 | 0,72337635 | 0,86386262 |
| 148170_at | CDC42EP5 | -0,21180145 | 8,28773106 | -0,36082128 | 0,72342667 | 0,86386262 |
| 3355_at | HTR1F | 0,02516677 | 3,54648025 | 0,36078838 | 0,72345075 | 0,86386262 |
| 57089_at | ENTPD7 | 0,09386022 | 8,49034257 | 0,36070714 | 0,7235102 | 0,86388006 |
| 9354_at | UBE4A | -0,03752719 | 10,2539527 | -0,36063128 | 0,72356572 | 0,8638928 |
| 221154_at | MICU2 | 0,06560886 | 8,45003112 | 0,36051549 | 0,72365046 | 0,86394043 |
| 6557_at | SLC12A1 | -0,0201976 | 2,82700258 | -0,36021201 | 0,72387258 | 0,86415206 |
| 10802_at | SEC24A | 0,04746606 | 7,58814178 | 0,36009867 | 0,72395555 | 0,86419754 |
| 100169890_a | PEG3-AS1 | 0,03894684 | 3,36930928 | 0,35999811 | 0,72402916 | 0,86423186 |
| 100996295_a | DNAH17-AS1 | -0,03786102 | 4,39594387 | -0,35985192 | 0,72413617 | 0,86424799 |
| 653583_at | PHLDB3 | 0,04594868 | 6,48811332 | 0,35982131 | 0,72415859 | 0,86424799 |
| 140706_at | CCM2L | 0,03526238 | 5,48031027 | 0,35979578 | 0,72417727 | 0,86424799 |
| 9469_at | CHST3 | -0,02376764 | 5,74754435 | -0,35961344 | 0,72431077 | 0,86435376 |
| 60495_at | HPSE2 | 0,04166423 | 4,30724287 | 0,35940816 | 0,72446106 | 0,86435988 |
| 286075_at | ZNF707 | -0,05992004 | 5,86975462 | -0,35937942 | 0,72448211 | 0,86435988 |

| | | | | | | |
|---|---|---|---|---|---|---|
| 5294_at | PIK3CG | -0,02131426 | 3,22500832 | -0,35936291 | 0,7244942 | 0,86435988 |
| 10109_at | ARPC2 | -0,03222277 | 12,1761343 | -0,35936129 | 0,72449538 | 0,86435988 |
| 56241_at | SUSD2 | -0,03845152 | 5,44745651 | -0,35925539 | 0,72457292 | 0,86439885 |
| 84989_at | JMJD1C-AS1 | 0,06081148 | 6,53053175 | 0,3591098 | 0,72467953 | 0,86447249 |
| 7957_at | EPM2A | 0,04239461 | 5,88814073 | 0,35891525 | 0,724822 | 0,8645889 |
| 81544_at | GDPD5 | -0,03396735 | 7,16364667 | -0,35876386 | 0,72493287 | 0,86466761 |
| 8115_at | TCL1A | -0,04243128 | 4,40574449 | -0,35861114 | 0,72504472 | 0,86474748 |
| 101927069_a | LOC10192706 | -0,03205323 | 2,96425545 | -0,35851192 | 0,72511739 | 0,86475518 |
| 6836_at | SURF4 | 0,0301297 | 10,0809855 | 0,35847973 | 0,72514097 | 0,86475518 |
| 139599_at | MAGEE2 | 0,02496 | 3,2445378 | 0,35829874 | 0,72527354 | 0,86481379 |
| 8437_at | RASAL1 | -0,0255182 | 5,90292511 | -0,35829005 | 0,72527991 | 0,86481379 |
| 101928700_a | LOC10192870 | 0,03323701 | 3,31291889 | 0,35800003 | 0,72549235 | 0,86499649 |
| 101927166_a | LOC10192716 | -0,04890126 | 4,61309509 | -0,3578066 | 0,72563406 | 0,86499649 |
| 55109_at | AGGF1 | -0,07688278 | 8,27384446 | -0,35780476 | 0,72563542 | 0,86499649 |
| 10017_at | BCL2L10 | -0,03550555 | 5,09472428 | -0,35777645 | 0,72565615 | 0,86499649 |
| 284349_at | ZNF283 | 0,05001391 | 4,84809799 | 0,35777441 | 0,72565765 | 0,86499649 |
| 51364_at | ZMYND10 | -0,04344003 | 4,95826184 | -0,35742423 | 0,72591422 | 0,86521481 |
| 440_at | ASNS | -0,10594979 | 8,25820495 | -0,35734063 | 0,72597548 | 0,86521481 |
| 10814_at | CPLX2 | -0,03684405 | 4,35680633 | -0,35729256 | 0,72601071 | 0,86521481 |
| 101929473_a | LOC10192947 | -0,03652791 | 3,58129267 | -0,35719273 | 0,72608386 | 0,86521481 |
| 64772_at | ENGASE | 0,08210293 | 7,3870624 | 0,35713133 | 0,72612886 | 0,86521481 |
| 22897_at | CEP164 | -0,02911236 | 6,28168896 | -0,35713073 | 0,72612929 | 0,86521481 |
| 3890_at | KRT84 | -0,03854215 | 4,07351511 | -0,35709536 | 0,72615521 | 0,86521481 |
| 84940_at | CORO6 | -0,03302344 | 4,99747723 | -0,3568762 | 0,72631583 | 0,8653064 |
| 55076_at | TMEM45A | -0,17975373 | 7,50668368 | -0,35686789 | 0,72632192 | 0,8653064 |
| 642345_at | MIR4500HG | 0,03175012 | 3,56390173 | 0,35648448 | 0,72660293 | 0,86546425 |
| 113220_at | KIF12 | -0,03578984 | 5,87556128 | -0,35647999 | 0,72660623 | 0,86546425 |
| 101927416_a | LOC10192741 | -0,03198668 | 4,55936515 | -0,35645037 | 0,72662794 | 0,86546425 |
| 220965_at | FAM13C | 0,02461918 | 4,16488962 | 0,35644192 | 0,72663414 | 0,86546425 |
| 57507_at | ZNF608 | 0,17423987 | 7,0212304 | 0,35621462 | 0,72680076 | 0,86559158 |
| 260425_at | MAGI3 | 0,03291788 | 6,32712086 | 0,356092 | 0,72689065 | 0,86559158 |
| 55679_at | LIMS2 | -0,03541376 | 5,72793882 | -0,35605277 | 0,72691941 | 0,86559158 |
| 57706_at | DENND1A | 0,01881996 | 6,69586734 | 0,3560509 | 0,72692078 | 0,86559158 |
| 257068_at | PLCXD2 | 0,0448765 | 5,23408027 | 0,3559687 | 0,72698104 | 0,86560982 |
| 1557_at | CYP2C19 | -0,0365644 | 3,98882257 | -0,35574733 | 0,72714335 | 0,86574957 |
| 9144_at | SYNGR2 | 0,04791007 | 9,65343239 | 0,3553691 | 0,72742068 | 0,8659489 |
| 101929239_a | LOC10192923 | -0,01947768 | 3,20686921 | -0,35526505 | 0,72749699 | 0,8659489 |
| 221710_at | SMIM13 | -0,08219918 | 7,51424336 | -0,3552617 | 0,72749944 | 0,8659489 |
| 56131_at | PCDHB4 | 0,02131066 | 3,68634254 | 0,35517454 | 0,72756336 | 0,8659489 |
| 26343_at | OR5E1P | -0,05096638 | 3,77634503 | -0,35516597 | 0,72756965 | 0,8659489 |
| 2894_at | GRID1 | 0,0470965 | 5,48462789 | 0,35515119 | 0,72758049 | 0,8659489 |
| 10174_at | SORBS3 | -0,04246721 | 6,14607954 | -0,35491784 | 0,72775162 | 0,86606094 |
| 22949_at | PTGR1 | 0,03134288 | 4,96313488 | 0,35490022 | 0,72776455 | 0,86606094 |
| 200298_at | LINC00528 | 0,03447858 | 3,6507891 | 0,35479309 | 0,72784313 | 0,86610095 |
| 101928043_a | LOC10192804 | -0,03458442 | 4,03476392 | -0,35459227 | 0,72799042 | 0,86622271 |
| 2266_at | FGG | -0,03457405 | 3,45447456 | -0,35438705 | 0,72814096 | 0,86632446 |
| 93664_at | CADPS2 | 0,32491762 | 6,24974353 | 0,35435309 | 0,72816588 | 0,86632446 |
| 57562_at | CEP126 | 0,03352063 | 4,49357275 | 0,35417474 | 0,72829672 | 0,86642661 |
| 100133308_a | RSU1P2 | -0,02527261 | 3,31037165 | -0,35393295 | 0,72847411 | 0,86656845 |
| 2534_at | FYN | 0,04795441 | 4,73597758 | 0,3538896 | 0,72850592 | 0,86656845 |

| | | | | | | |
|---|---|---|---|---|---|---|
| 5071_at | PARK2 | -0,02095037 | 3,48232324 | -0,35379221 | 0,72857738 | 0,86659994 |
| 100170765_a | ERICH4 | -0,03365381 | 5,31753467 | -0,35355073 | 0,72875457 | 0,86672103 |
| 4855_at | NOTCH4 | -0,03176562 | 5,04798262 | -0,35353083 | 0,72876917 | 0,86672103 |
| 9383_at | TSIX | -0,16220815 | 4,33683736 | -0,35345809 | 0,72882255 | 0,866731 |
| 6538_at | SLC6A11 | 0,04673312 | 5,40307338 | 0,35327548 | 0,72895656 | 0,86675474 |
| 3714_at | JAG2 | 0,11445099 | 8,06501863 | 0,35324736 | 0,7289772 | 0,86675474 |
| 170850_at | KCNG3 | -0,03522999 | 3,96052352 | -0,35317272 | 0,72903197 | 0,86675474 |
| 8653_at | DDX3Y | 0,3087662 | 5,53520221 | 0,35316126 | 0,72904038 | 0,86675474 |
| 23457_at | ABCB9 | -0,03315056 | 5,60785952 | -0,35312432 | 0,72906749 | 0,86675474 |
| 22802_at | CLCA4 | -0,09345921 | 5,89765158 | -0,35299914 | 0,72915937 | 0,86681048 |
| 50805_at | IRX4 | 0,03960133 | 5,0638275 | 0,35254256 | 0,72949452 | 0,86715538 |
| 10665_at | C6orf10 | -0,02562912 | 2,88432783 | -0,35236384 | 0,72962571 | 0,86725781 |
| 374407_at | DNAJB13 | -0,0354894 | 5,18790007 | -0,35229888 | 0,7296734 | 0,86726098 |
| 5870_at | RAB6A | 0,02988417 | 8,38802568 | 0,35210763 | 0,72981381 | 0,86732312 |
| 27288_at | RBMXL2 | 0,0348567 | 4,62343222 | 0,35210502 | 0,72981573 | 0,86732312 |
| 135295_at | SRSF12 | 0,02617131 | 3,97834483 | 0,35161795 | 0,73017337 | 0,86769461 |
| 101929082_a | LINC01333 | 0,03135024 | 3,32966627 | 0,35147488 | 0,73027844 | 0,86771196 |
| 387_at | RHOA | -0,0541221 | 11,4418395 | -0,35144345 | 0,73030152 | 0,86771196 |
| 113278_at | SLC52A3 | -0,0443464 | 6,68897384 | -0,35131954 | 0,73039252 | 0,86771196 |
| 10981_at | RAB32 | 0,15281189 | 8,27539152 | 0,35131351 | 0,73039695 | 0,86771196 |
| 84848_at | MIR503HG | -0,04817796 | 5,07657457 | -0,35129138 | 0,7304132 | 0,86771196 |
| 64428_at | NARFL | -0,0194375 | 7,22631094 | -0,3511623 | 0,730508 | 0,86777107 |
| 9716_at | AQR | -0,02714575 | 8,57629078 | -0,35105159 | 0,73058932 | 0,86781415 |
| 245812_at | CNPY4 | -0,02494099 | 6,57360359 | -0,35072799 | 0,73082702 | 0,86803027 |
| 833_at | CARS | 0,04025149 | 7,82803607 | 0,35063933 | 0,73089214 | 0,86803027 |
| 51259_at | TMEM216 | 0,05071089 | 8,25810053 | 0,35061985 | 0,73090645 | 0,86803027 |
| 143888_at | KDELC2 | -0,16930811 | 7,0205867 | -0,35045407 | 0,73102825 | 0,86812139 |
| 7755_at | ZNF205 | -0,03904366 | 6,44825131 | -0,35016463 | 0,73124089 | 0,86830819 |
| 765_at | CA6 | -0,02986967 | 3,76623579 | -0,35011726 | 0,7312757 | 0,86830819 |
| 4086_at | SMAD1 | 0,07748231 | 8,04404172 | 0,34966424 | 0,73160858 | 0,86850494 |
| 85289_at | KRTAP4-5 | 0,03324852 | 4,31116049 | 0,34957846 | 0,73167162 | 0,86850494 |
| 2982_at | GUCY1A3 | -0,02150602 | 3,5114851 | -0,34950225 | 0,73172763 | 0,86850494 |
| 10152_at | ABI2 | 0,03088226 | 8,05284473 | 0,34948474 | 0,7317405 | 0,86850494 |
| 5891_at | MOK | 0,06639597 | 6,19990881 | 0,34944696 | 0,73176827 | 0,86850494 |
| 8702_at | B4GALT4 | 0,03983539 | 7,92948183 | 0,34941263 | 0,73179349 | 0,86850494 |
| 10671_at | DCTN6 | -0,08179775 | 9,05453694 | -0,34930946 | 0,73186932 | 0,86850494 |
| 4430_at | MYO1B | -0,11526732 | 10,1177845 | -0,34930893 | 0,73186971 | 0,86850494 |
| 203447_at | NRK | 0,02560389 | 3,12746481 | 0,34919504 | 0,73195341 | 0,86850494 |
| 283345_at | RPL13P5 | -0,02323679 | 6,42067453 | -0,3491567 | 0,7319816 | 0,86850494 |
| 54660_at | PCDHB18P | 0,05162116 | 4,55150842 | 0,3491167 | 0,732011 | 0,86850494 |
| 11190_at | CEP250 | -0,01856453 | 6,00640844 | -0,34910493 | 0,73201964 | 0,86850494 |
| 253650_at | ANKRD18A | -0,05024628 | 3,34808517 | -0,3490106 | 0,73208899 | 0,86850494 |
| 10397_at | NDRG1 | -0,05612743 | 9,98327341 | -0,3489826 | 0,73210957 | 0,86850494 |
| 51087_at | YBX2 | -0,04912735 | 6,12050172 | -0,34879533 | 0,73224722 | 0,86850494 |
| 83897_at | KRTAP3-2 | 0,02227002 | 3,42956301 | 0,34878152 | 0,73225737 | 0,86850494 |
| 2712_at | GK2 | -0,03439199 | 3,82518146 | -0,34874893 | 0,73228133 | 0,86850494 |
| 22820_at | COPG1 | 0,11156119 | 10,0173554 | 0,3487382 | 0,73228922 | 0,86850494 |
| 252839_at | TMEM9 | 0,03912126 | 8,76118174 | 0,34872619 | 0,73229805 | 0,86850494 |
| 553103_at | LOC553103 | 0,07501097 | 5,97703833 | 0,34842346 | 0,73252061 | 0,86870198 |
| 25945_at | PVRL3 | 0,3195728 | 7,25654349 | 0,34837753 | 0,73255438 | 0,86870198 |

| | | | | | | |
|---|---|---|---|---|---|---|
| 115350_at | FCRL1 | 0,02584127 | 3,53359097 | 0,34818485 | 0,73269605 | 0,86879656 |
| 8992_at | ATP6V0E1 | -0,06128606 | 11,7079627 | -0,348124 | 0,73274079 | 0,86879656 |
| 4224_at | MEP1A | 0,03811495 | 3,34753802 | 0,34802591 | 0,73281291 | 0,86879656 |
| 11014_at | KDELR2 | -0,03426218 | 10,8829543 | -0,3480237 | 0,73281454 | 0,86879656 |
| 9415_at | FADS2 | -0,04069321 | 7,59380046 | -0,34788491 | 0,73291659 | 0,86886408 |
| 79843_at | FAM124B | -0,03586042 | 3,98842947 | -0,34780283 | 0,73297696 | 0,86888216 |
| 135886_at | WBSCR28 | 0,05566389 | 4,3057446 | 0,34763728 | 0,7330987 | 0,86896341 |
| 151484_at | LOC151484 | -0,03941093 | 4,53394526 | -0,34758694 | 0,73313572 | 0,86896341 |
| 100505716_a | LOC10050571 | -0,04219717 | 3,5357477 | -0,34741708 | 0,73326065 | 0,86905801 |
| 26512_at | INTS6 | -0,32763241 | 5,56856664 | -0,34707975 | 0,73350876 | 0,86929859 |
| 9513_at | FXR2 | 0,02294483 | 7,61576179 | 0,34693182 | 0,73361758 | 0,86936831 |
| 56616_at | DIABLO | -0,03578694 | 9,46512878 | -0,34687707 | 0,73365786 | 0,86936831 |
| 9719_at | ADAMTSL2 | 0,03020328 | 4,67861121 | 0,34633579 | 0,73405608 | 0,86975709 |
| 84850_at | GLIS3-AS1 | -0,0416263 | 4,91973359 | -0,34619309 | 0,73416108 | 0,86975709 |
| 53831_at | GPR84 | -0,05000391 | 4,13233911 | -0,34616608 | 0,73418095 | 0,86975709 |
| 11040_at | PIM2 | -0,05944159 | 7,38330874 | -0,34614395 | 0,73419724 | 0,86975709 |
| 1130_at | LYST | -0,06011067 | 5,71344613 | -0,34612428 | 0,73421171 | 0,86975709 |
| 7157_at | TP53 | 0,12623585 | 8,27355396 | 0,34595871 | 0,73433355 | 0,86984793 |
| 3804_at | KIR2DL3 | -0,05651459 | 6,72434012 | -0,3458362 | 0,7344237 | 0,86985431 |
| 8682_at | PEA15 | 0,0487574 | 9,66966406 | 0,34582866 | 0,73442925 | 0,86985431 |
| 369_at | ARAF | -0,03253497 | 7,53715075 | -0,34559081 | 0,73460429 | 0,87000815 |
| 5241_at | PGR | -0,02339197 | 3,24298483 | -0,34551406 | 0,73466078 | 0,87002155 |
| 119369_at | NUDT9P1 | 0,03227782 | 3,26150258 | 0,34528499 | 0,73482938 | 0,87014766 |
| 11006_at | LILRB4 | -0,04025127 | 4,1235763 | -0,34524664 | 0,73485761 | 0,87014766 |
| 642559_at | POU5F1P3 | -0,06562856 | 4,94635314 | -0,34513701 | 0,73493831 | 0,87018972 |
| 51085_at | MLXIPL | -0,03999612 | 6,16769807 | -0,34492239 | 0,73509629 | 0,87032329 |
| 1178_at | CLC | -0,03939925 | 3,84243957 | -0,34485719 | 0,73514429 | 0,87032662 |
| 9435_at | CHST2 | -0,03824736 | 5,40410693 | -0,34462559 | 0,7353148 | 0,87040211 |
| 8567_at | MADD | 0,02209094 | 8,42738763 | 0,34462552 | 0,73531485 | 0,87040211 |
| 159162_at | RBMY2FP | -0,02570928 | 3,60613862 | -0,34457485 | 0,73535216 | 0,87040211 |
| 6442_at | SGCA | -0,03972774 | 5,06975041 | -0,34452509 | 0,73538879 | 0,87040211 |
| 85495_at | RPPH1 | -0,08446593 | 6,96981405 | -0,34429221 | 0,73556026 | 0,87048478 |
| 5631_at | PRPS1 | -0,0821436 | 9,10750562 | -0,34420181 | 0,73562682 | 0,87048478 |
| 100506405_a | LOC10050640 | -0,03196483 | 3,87476885 | -0,34407105 | 0,73572311 | 0,87048478 |
| 26049_at | FAM169A | -0,05349397 | 6,87124374 | -0,34405725 | 0,73573327 | 0,87048478 |
| 55702_at | CCDC94 | -0,02273925 | 7,85257839 | -0,34405563 | 0,73573447 | 0,87048478 |
| 120379_at | PIH1D2 | 0,03931058 | 4,26013028 | 0,34401898 | 0,73576146 | 0,87048478 |
| 7441_at | VPREB1 | -0,02857811 | 3,95447618 | -0,34400063 | 0,73577497 | 0,87048478 |
| 150135_at | LINC00479 | -0,0694041 | 5,05226799 | -0,34386821 | 0,73587249 | 0,87048501 |
| 8302_at | KLRC4 | -0,01906234 | 2,63534038 | -0,34379198 | 0,73592862 | 0,87048501 |
| 10660_at | LBX1 | -0,03654909 | 4,98905606 | -0,34374379 | 0,73596411 | 0,87048501 |
| 64174_at | DPEP2 | -0,03598474 | 4,56186606 | -0,34364172 | 0,73603929 | 0,87048501 |
| 5255_at | PHKA1 | -0,05751031 | 6,92214818 | -0,34363321 | 0,73604556 | 0,87048501 |
| 81627_at | TRMT1L | -0,04625559 | 7,33035377 | -0,3436322 | 0,7360463 | 0,87048501 |
| 29801_at | ZDHHC8 | -0,03442562 | 6,97835275 | -0,3435412 | 0,73611333 | 0,87050565 |
| 1836_at | SLC26A2 | 0,10450645 | 8,3672197 | 0,3434858 | 0,73615413 | 0,87050565 |
| 10880_at | ACTL7B | -0,02413223 | 5,09487879 | -0,34274129 | 0,73670256 | 0,87106729 |
| 284958_at | NT5DC4 | -0,03988478 | 3,62542576 | -0,34264268 | 0,73677522 | 0,87106729 |
| 51056_at | LAP3 | 0,08358721 | 9,22991936 | 0,34259626 | 0,73680942 | 0,87106729 |
| 441461_at | STX17-AS1 | 0,03481811 | 5,19427186 | 0,34255091 | 0,73684283 | 0,87106729 |

| | | | | | | |
|---|---|---|---|---|---|---|
| 161635_at | CSNK1A1P1 | 0,02524922 | 4,15020839 | 0,34253414 | 0,73685519 | 0,87106729 |
| 101927411_a | LOC10192741 | -0,0319311 | 3,97942661 | -0,34238917 | 0,73696201 | 0,8711103 |
| 91039_at | DPP9 | 0,04873353 | 7,17928894 | 0,34236202 | 0,73698201 | 0,8711103 |
| 8013_at | NR4A3 | -0,02228717 | 3,94120239 | -0,34221068 | 0,73709353 | 0,87114934 |
| 729522_at | AACSP1 | 0,04880341 | 4,4773713 | 0,34216021 | 0,73713071 | 0,87114934 |
| 5276_at | SERPINI2 | -0,03880718 | 3,46010909 | -0,34209917 | 0,7371757 | 0,87114934 |
| 8633_at | UNC5C | 0,02782504 | 4,10786009 | 0,34205269 | 0,73720995 | 0,87114934 |
| 126003_at | TRAPPC5 | 0,04526782 | 9,47927113 | 0,34201033 | 0,73724117 | 0,87114934 |
| 9380_at | GRHPR | -0,06296528 | 9,18044414 | -0,34182535 | 0,73737749 | 0,87125698 |
| 23272_at | FAM208A | -0,0506494 | 7,79360914 | -0,34149131 | 0,73762369 | 0,87149442 |
| 353219_at | KAAG1 | -0,0319539 | 4,3750217 | -0,34137746 | 0,73770761 | 0,87154012 |
| 101927843_a | LOC10192784 | 0,02931939 | 3,72515297 | 0,34120508 | 0,73783468 | 0,87158403 |
| 57190_at | SEPN1 | 0,038344 | 7,75954485 | 0,34117001 | 0,73786052 | 0,87158403 |
| 414899_at | BLID | -0,03289961 | 3,37322494 | -0,34110655 | 0,73790731 | 0,87158403 |
| 80778_at | ZNF34 | 0,10022675 | 5,69484433 | 0,34108151 | 0,73792576 | 0,87158403 |
| 91607_at | SLFN11 | 0,11438979 | 4,23197342 | 0,34084701 | 0,73809864 | 0,87173478 |
| 2528_at | FUT6 | -0,02556512 | 6,07117235 | -0,34069797 | 0,73820852 | 0,8718111 |
| 944_at | TNFSF8 | 0,02699463 | 3,66939075 | 0,34060147 | 0,73827967 | 0,87182383 |
| 130916_at | MTERF4 | -0,02626555 | 6,16518352 | -0,34056058 | 0,73830983 | 0,87182383 |
| 3104_at | ZBTB48 | -0,0549771 | 7,03070092 | -0,34048125 | 0,73836832 | 0,87183946 |
| 57571_at | CARNS1 | -0,04715341 | 5,35355442 | -0,34038663 | 0,73843809 | 0,8718684 |
| 10882_at | C1QL1 | -0,03401477 | 6,6608776 | -0,34024798 | 0,73854032 | 0,87190338 |
| 10738_at | RFPL3 | -0,0541421 | 4,48159404 | -0,3402086 | 0,73856936 | 0,87190338 |
| 115399_at | LRRC56 | 0,04445092 | 6,19804542 | 0,34011414 | 0,73863902 | 0,87190338 |
| 100130557_a | NFYC-AS1 | -0,04008259 | 5,44930819 | -0,34010092 | 0,73864877 | 0,87190338 |
| 23514_at | SPIDR | 0,02770708 | 6,75177361 | 0,33953983 | 0,73906259 | 0,87233839 |
| 5423_at | POLB | -0,10811106 | 6,71252237 | -0,3392354 | 0,73928715 | 0,87252531 |
| 137868_at | SGCZ | 0,03396508 | 4,34545366 | 0,3391928 | 0,73931857 | 0,87252531 |
| 4495_at | MT1G | 0,030672 | 6,42697063 | 0,33909781 | 0,73938865 | 0,87252531 |
| 220323_at | OAF | 0,06691607 | 7,76014846 | 0,33907953 | 0,73940213 | 0,87252531 |
| 5607_at | MAP2K5 | 0,02584835 | 6,67929022 | 0,3387588 | 0,73963875 | 0,87268374 |
| 23418_at | CRB1 | -0,02401277 | 3,4136065 | -0,33867469 | 0,73970081 | 0,87268374 |
| 124976_at | SPNS2 | -0,07228625 | 6,23276932 | -0,33865876 | 0,73971256 | 0,87268374 |
| 54873_at | PALMD | -0,29420041 | 6,80150573 | -0,33852564 | 0,73981079 | 0,87268374 |
| 3678_at | ITGA5 | 0,03758764 | 7,25561536 | 0,33851089 | 0,73982167 | 0,87268374 |
| 9276_at | COPB2 | 0,07387214 | 10,2477368 | 0,33847981 | 0,73984461 | 0,87268374 |
| 25878_at | MXRA5 | 0,32677762 | 6,27233393 | 0,33837526 | 0,73992175 | 0,87268374 |
| 4619_at | MYH1 | -0,02312588 | 2,90551833 | -0,33835037 | 0,73994012 | 0,87268374 |
| 286319_at | TUSC1 | -0,02113484 | 9,3031762 | -0,33834495 | 0,73994412 | 0,87268374 |
| 22931_at | RAB18 | 0,06229648 | 8,0864331 | 0,33821129 | 0,74004275 | 0,87273547 |
| 81858_at | SHARPIN | 0,08500472 | 8,3115081 | 0,33816273 | 0,74007859 | 0,87273547 |
| 144360_at | LINC00477 | -0,03836178 | 3,55056024 | -0,3379284 | 0,74025152 | 0,87288595 |
| 145567_at | TTC7B | 0,05082721 | 7,5312637 | 0,33784902 | 0,7403101 | 0,8729016 |
| 7923_at | HSD17B8 | 0,05855314 | 7,70532704 | 0,33765777 | 0,74045126 | 0,87301461 |
| 100505878_a | LOC10050587 | 0,0275613 | 2,88544262 | 0,33751066 | 0,74055984 | 0,87307733 |
| 101928516_a | LOC10192851 | 0,02508746 | 3,30963518 | 0,3374442 | 0,7406089 | 0,87307733 |
| 100506680_a | SACS-AS1 | 0,02198261 | 3,35986667 | 0,33728331 | 0,74072767 | 0,87307733 |
| 283869_at | NPW | -0,02849099 | 4,35215687 | -0,3372789 | 0,74073092 | 0,87307733 |
| 7454_at | WAS | -0,04781488 | 5,50274447 | -0,33727868 | 0,74073108 | 0,87307733 |
| 100506994_a | PTPRG-AS1 | 0,05362339 | 3,25907915 | 0,33697741 | 0,74095349 | 0,87321958 |

| | | | | | | |
|---|---|---|---|---|---|---|
| 401097_at | C3orf80 | -0,01868084 | 3,57411849 | -0,33695672 | 0,74096876 | 0,87321958 |
| 127731_at | VWA5B1 | -0,03139153 | 4,50413275 | -0,3369226 | 0,74099396 | 0,87321958 |
| 375033_at | PEAR1 | -0,04733529 | 5,40300839 | -0,33683828 | 0,74105621 | 0,87321958 |
| 220202_at | ATOH7 | 0,0223271 | 3,47855993 | 0,33680819 | 0,74107843 | 0,87321958 |
| 79364_at | ZXDC | 0,04027368 | 7,4020821 | 0,3366012 | 0,74123126 | 0,87327658 |
| 1196_at | CLK2 | 0,0422819 | 8,60717791 | 0,33658152 | 0,74124578 | 0,87327658 |
| 440348_at | NPIPB15 | -0,1738638 | 5,46390184 | -0,33650787 | 0,74130017 | 0,87327658 |
| 682_at | BSG | 0,09969666 | 10,4028303 | 0,33649708 | 0,74130814 | 0,87327658 |
| 28_at | ABO | -0,02953152 | 5,75598986 | -0,33626163 | 0,741482 | 0,87342798 |
| 84678_at | KDM2B | -0,04243853 | 8,48898512 | -0,33607427 | 0,74162037 | 0,87353755 |
| 729440_at | CCDC61 | 0,0340583 | 6,21411485 | 0,33587856 | 0,74176491 | 0,87363393 |
| 27068_at | PPA2 | -0,07970949 | 9,01429818 | -0,33584066 | 0,74179289 | 0,87363393 |
| 163071_at | ZNF114 | -0,08291404 | 4,25274292 | -0,33562 | 0,74195588 | 0,87371454 |
| 286059_at | LOC286059 | 0,02143069 | 3,84962235 | 0,33560668 | 0,74196571 | 0,87371454 |
| 100130894_a | VAC14-AS1 | -0,03135417 | 4,12807754 | -0,33551901 | 0,74203047 | 0,87371454 |
| 101929591_a | KIZ-AS1 | -0,04361588 | 3,44221829 | -0,33550237 | 0,74204277 | 0,87371454 |
| 1875_at | E2F5 | 0,06358939 | 6,86799594 | 0,33521621 | 0,74225415 | 0,87391001 |
| 7494_at | XBP1 | -0,05100586 | 9,22243042 | -0,33504113 | 0,74238349 | 0,87400766 |
| 147968_at | CAPN12 | 0,02564641 | 4,87765154 | 0,33485708 | 0,74251947 | 0,87400766 |
| 5921_at | RASA1 | -0,09040436 | 8,60331108 | -0,33482791 | 0,74254102 | 0,87400766 |
| 2775_at | GNAO1 | -0,02819965 | 4,21908135 | -0,33481376 | 0,74255148 | 0,87400766 |
| 26019_at | UPF2 | -0,08060857 | 7,67392256 | -0,33479688 | 0,74256395 | 0,87400766 |
| 554201_at | CCDC148-AS1 | 0,02876593 | 3,19232146 | 0,33461101 | 0,74270129 | 0,8741159 |
| 1822_at | ATN1 | -0,02866344 | 7,34053471 | -0,33442864 | 0,74283604 | 0,87421728 |
| 25942_at | SIN3A | 0,0257559 | 7,93015753 | 0,33437159 | 0,7428782 | 0,87421728 |
| 132625_at | ZFP42 | -0,02097088 | 3,28509785 | -0,33411875 | 0,74306505 | 0,87434783 |
| 149643_at | SPATA45 | -0,02699576 | 3,63406232 | -0,33409863 | 0,74307991 | 0,87434783 |
| 102724612_a | LOC10272461 | 0,02788609 | 3,51763768 | 0,33382113 | 0,743285 | 0,87452227 |
| 116535_at | MRGPRF | -0,03962428 | 5,17803536 | -0,333716 | 0,74336271 | 0,87452227 |
| 3284_at | HSD3B2 | 0,02606838 | 3,31320009 | 0,33371377 | 0,74336436 | 0,87452227 |
| 400643_at | LINC00668 | -0,04174733 | 5,740485 | -0,33339449 | 0,74360036 | 0,87469885 |
| 8911_at | CACNA1I | -0,03186037 | 5,65865381 | -0,33330825 | 0,74366411 | 0,87469885 |
| 771_at | CA12 | -0,37384396 | 8,30635831 | -0,33326748 | 0,74369425 | 0,87469885 |
| 147409_at | DSG4 | -0,0799694 | 3,5179757 | -0,33322975 | 0,74372214 | 0,87469885 |
| 118429_at | ANTXR2 | -0,08350828 | 5,13033114 | -0,33320356 | 0,7437415 | 0,87469885 |
| 11081_at | KERA | -0,03898504 | 2,96431192 | -0,3331344 | 0,74379263 | 0,87470559 |
| 284402_at | SCGB2B2 | -0,03204694 | 3,97824095 | -0,33303232 | 0,7438681 | 0,87474093 |
| 143471_at | PSMA8 | -0,01832038 | 2,72466036 | -0,33269907 | 0,74411449 | 0,87491894 |
| 57526_at | PCDH19 | 0,04448513 | 5,11394042 | 0,33267565 | 0,74413181 | 0,87491894 |
| 83259_at | PCDH11Y | -0,02622383 | 3,12020836 | -0,33264329 | 0,74415573 | 0,87491894 |
| 5195_at | PEX14 | 0,03721979 | 7,88398736 | 0,33238674 | 0,74434544 | 0,87508857 |
| 7414_at | VCL | -0,03634217 | 8,56976164 | -0,3319919 | 0,74463744 | 0,87532705 |
| 8227_at | AKAP17A | -0,06373136 | 7,35293731 | -0,33198955 | 0,74463917 | 0,87532705 |
| 10342_at | TFG | -0,05879418 | 10,8960076 | -0,33191119 | 0,74469713 | 0,87534176 |
| 27442_at | CECR3 | -0,03101754 | 3,80901878 | -0,33128036 | 0,74516374 | 0,87582019 |
| 201931_at | TMEM192 | -0,10354791 | 7,52797066 | -0,33123799 | 0,74519509 | 0,87582019 |
| 57403_at | RAB22A | 0,03436851 | 9,07035446 | 0,33116244 | 0,74525098 | 0,87583244 |
| 134429_at | STARD4 | -0,21057113 | 6,32158285 | -0,33093361 | 0,74542027 | 0,87597796 |
| 100505545_a | LINC01095 | -0,04050013 | 3,40847217 | -0,33068799 | 0,745602 | 0,87612047 |
| 389895_at | LOC389895 | -0,03731473 | 4,29794303 | -0,33064677 | 0,74563251 | 0,87612047 |

| | | | | | | |
|---|---|---|---|---|---|---|
| 159195_at | USP54 | 0,04817593 | 6,99217473 | 0,33039211 | 0,74582094 | 0,87628843 |
| 1397_at | CRIP2 | -0,33274106 | 7,42707092 | -0,33025898 | 0,74591946 | 0,87635073 |
| 91300_at | R3HDM4 | 0,01998989 | 8,58087403 | 0,33013139 | 0,74601388 | 0,87640821 |
| 101928162_a | LOC10192816 | 0,02529784 | 3,35494799 | 0,33004298 | 0,74607932 | 0,87643163 |
| 100505797_a | LOC10050579 | -0,05452484 | 3,53413903 | -0,32990555 | 0,74618103 | 0,87644915 |
| 57528_at | KCTD16 | 0,07883151 | 4,42197635 | 0,32989987 | 0,74618523 | 0,87644915 |
| 3932_at | LCK | -0,09567662 | 5,80674117 | -0,32965288 | 0,74636804 | 0,87660377 |
| 9953_at | HS3ST3B1 | 0,02801406 | 3,6607736 | 0,32956812 | 0,74643079 | 0,87660377 |
| 4170_at | MCL1 | -0,06051267 | 9,82717322 | -0,32953758 | 0,74645339 | 0,87660377 |
| 93166_at | PRDM6 | -0,0453477 | 5,18338711 | -0,32917269 | 0,74672351 | 0,87686753 |
| 8424_at | BBOX1 | -0,18690211 | 4,54652605 | -0,32901023 | 0,74684378 | 0,87689859 |
| 4589_at | MUC7 | 0,02418533 | 3,1024315 | 0,32896138 | 0,74687994 | 0,87689859 |
| 10249_at | GLYAT | -0,03142735 | 3,67064849 | -0,32895249 | 0,74688653 | 0,87689859 |
| 9711_at | KIAA0226 | 0,03301354 | 6,87415429 | 0,32829991 | 0,74736973 | 0,87741243 |
| 10867_at | TSPAN9 | 0,02080563 | 6,67496144 | 0,32808394 | 0,74752967 | 0,87754672 |
| 57545_at | CC2D2A | 0,04275127 | 5,10705582 | 0,32797345 | 0,7476115 | 0,8775893 |
| 11144_at | DMC1 | 0,01998862 | 3,29605635 | 0,32773051 | 0,74779144 | 0,87774634 |
| 8749_at | ADAM18 | -0,02057799 | 3,12901207 | -0,32766978 | 0,74783642 | 0,87774634 |
| 83786_at | FRMD8 | -0,02738705 | 7,6729309 | -0,32754096 | 0,74793184 | 0,87780243 |
| 285598_at | ARL10 | -0,03681333 | 5,11355382 | -0,32748223 | 0,74797534 | 0,87780243 |
| 80816_at | ASXL3 | -0,02415882 | 3,64872516 | -0,32725903 | 0,74814069 | 0,87781877 |
| 55888_at | ZKSCAN7 | 0,01893817 | 3,64834608 | 0,32723563 | 0,74815802 | 0,87781877 |
| 152118_at | C3orf79 | -0,02399004 | 2,97619281 | -0,32722357 | 0,74816695 | 0,87781877 |
| 8710_at | SERPINB7 | -0,02484889 | 4,07375295 | -0,3271931 | 0,74818953 | 0,87781877 |
| 535_at | ATP6V0A1 | -0,04298966 | 6,97670139 | -0,32715586 | 0,74821712 | 0,87781877 |
| 6695_at | SPOCK1 | -0,12916234 | 6,89102499 | -0,32688792 | 0,74841562 | 0,87799818 |
| 121214_at | SDR9C7 | 0,03313546 | 3,96281994 | 0,32670965 | 0,74854771 | 0,87809966 |
| 283352_at | LOC283352 | -0,03233321 | 3,25682933 | -0,32664073 | 0,74859878 | 0,87810609 |
| 340017_at | LOC340017 | 0,02147306 | 3,08428322 | 0,32650143 | 0,748702 | 0,87816624 |
| 6284_at | S100A13 | 0,04805322 | 8,98187036 | 0,32644849 | 0,74874123 | 0,87816624 |
| 2354_at | FOSB | 0,0335796 | 5,75043519 | 0,32629517 | 0,74885484 | 0,87824601 |
| 221409_at | SPATS1 | -0,02690309 | 4,48125657 | -0,32610879 | 0,74899297 | 0,87835453 |
| 26015_at | RPAP1 | -0,02855728 | 7,33684799 | -0,325836 | 0,74919513 | 0,87853813 |
| 7694_at | ZNF135 | 0,03452373 | 4,13261082 | 0,32536656 | 0,7495431 | 0,87887627 |
| 100507564_a | LOC10050756 | -0,04628183 | 5,21879281 | -0,32532387 | 0,74957474 | 0,87887627 |
| 8940_at | TOP3B | 0,03836603 | 6,6439908 | 0,32519055 | 0,74967358 | 0,87889982 |
| 23096_at | IQSEC2 | -0,03757059 | 5,59847392 | -0,32515354 | 0,74970102 | 0,87889982 |
| 64127_at | NOD2 | 0,03452738 | 4,00617116 | 0,32511214 | 0,74973171 | 0,87889982 |
| 284757_at | MIR646HG | -0,08110555 | 3,90957007 | -0,32497052 | 0,7498367 | 0,87894473 |
| 55756_at | INTS9 | -0,07245829 | 7,20740546 | -0,3249224 | 0,74987238 | 0,87894473 |
| 23002_at | DAAM1 | 0,02765935 | 9,06433006 | 0,32482196 | 0,74994685 | 0,87894473 |
| 647946_at | LINC00669 | -0,10208526 | 3,87921727 | -0,32481429 | 0,74995253 | 0,87894473 |
| 255275_at | MYADML2 | 0,03707161 | 5,32496191 | 0,32468447 | 0,75004879 | 0,87900122 |
| 26_at | AOC1 | 0,28890538 | 5,88807785 | 0,32457903 | 0,75012697 | 0,87900122 |
| 51070_at | NOSIP | 0,03826666 | 8,87338144 | 0,32456467 | 0,75013762 | 0,87900122 |
| 100506049_a | LRRC72 | -0,03596685 | 3,85167183 | -0,32424933 | 0,75037145 | 0,87915716 |
| 2523_at | FUT1 | -0,03704865 | 7,14325865 | -0,32423195 | 0,75038435 | 0,87915716 |
| 1805_at | DPT | -0,02626316 | 3,85513324 | -0,32420056 | 0,75040762 | 0,87915716 |
| 400242_at | DICER1-AS1 | 0,04595354 | 4,73770261 | 0,32390901 | 0,75062384 | 0,87929009 |
| 55293_at | UEVLD | 0,05900519 | 7,7471493 | 0,32390298 | 0,75062831 | 0,87929009 |

| | | | | | | |
|---|---|---|---|---|---|---|
| 5795_at | PTPRJ | 0,0392999 | 6,58913268 | 0,32386293 | 0,75065802 | 0,87929009 |
| 54332_at | GDAP1 | 0,03707123 | 6,95796936 | 0,32362516 | 0,75083438 | 0,87944318 |
| 54843_at | SYTL2 | 0,19121642 | 6,45616083 | 0,32348055 | 0,75094164 | 0,87951535 |
| 6096_at | RORB | 0,03132593 | 3,63843585 | 0,3233734 | 0,75102112 | 0,87955495 |
| 1365_at | CLDN3 | 0,05641529 | 5,72219503 | 0,32314173 | 0,75119298 | 0,87967639 |
| 7503_at | XIST | -0,38320904 | 5,23396942 | -0,32311051 | 0,75121615 | 0,87967639 |
| 51061_at | TXNDC11 | 0,03363641 | 7,6293422 | 0,32303558 | 0,75127173 | 0,87968801 |
| 140735_at | DYNLL2 | -0,0699261 | 8,81863065 | -0,32241382 | 0,75173306 | 0,88015025 |
| 101928304_a | LOC10192830 | -0,04502088 | 4,70530754 | -0,32234275 | 0,7517858 | 0,88015025 |
| 3661_at | IRF3 | -0,03643545 | 8,49054312 | -0,32231879 | 0,75180357 | 0,88015025 |
| 1642_at | DDB1 | -0,01868489 | 10,4617028 | -0,32169959 | 0,75226312 | 0,88063473 |
| 54886_at | LPPR1 | 0,02171439 | 3,77212729 | 0,3216251 | 0,7523184 | 0,88064593 |
| 100133991_a | MAP3K14-AS1 | 0,03408959 | 5,00247959 | 0,32154689 | 0,75237645 | 0,88066036 |
| 29115_at | SAP30BP | -0,02213293 | 7,60375601 | -0,32143742 | 0,75245771 | 0,88066318 |
| 7639_at | ZNF85 | -0,06088095 | 5,90076578 | -0,32142047 | 0,7524703 | 0,88066318 |
| 64901_at | RANBP17 | 0,03790245 | 4,64665258 | 0,32115009 | 0,752671 | 0,88076674 |
| 171482_at | FAM9A | -0,03055681 | 3,09437967 | -0,32113576 | 0,75268165 | 0,88076674 |
| 8460_at | TPST1 | -0,03890625 | 7,07609601 | -0,32111143 | 0,75269971 | 0,88076674 |
| 2317_at | FLNB | 0,04298414 | 9,79872613 | 0,32102211 | 0,75276602 | 0,88076674 |
| 55020_at | TTC38 | -0,02280101 | 7,13817276 | -0,3209872 | 0,75279194 | 0,88076674 |
| 23770_at | FKBP8 | -0,02938341 | 7,23412143 | -0,32093172 | 0,75283312 | 0,88076674 |
| 3479_at | IGF1 | -0,02651392 | 3,80029998 | -0,32063123 | 0,75305623 | 0,88097426 |
| 112724_at | RDH13 | 0,08144165 | 8,32799704 | 0,32019139 | 0,75338283 | 0,88128634 |
| 1472_at | CST4 | -0,11769697 | 7,47379886 | -0,32009838 | 0,7534519 | 0,88128634 |
| 26239_at | LCE2B | -0,03877445 | 3,83698497 | -0,32008714 | 0,75346025 | 0,88128634 |
| 286083_at | LOC286083 | -0,0352344 | 3,83763283 | -0,31990231 | 0,75359751 | 0,88138961 |
| 28825_at | IGLV1-40 | -0,02583668 | 4,5596595 | -0,31977226 | 0,7536941 | 0,88138961 |
| 56651_at | LINC00470 | 0,04712226 | 3,09301363 | 0,31976844 | 0,75369694 | 0,88138961 |
| 8048_at | CSRP3 | -0,02746218 | 3,87374225 | -0,3196612 | 0,75377658 | 0,88138961 |
| 10913_at | EDAR | -0,04589049 | 6,73881832 | -0,31957153 | 0,75384319 | 0,88138961 |
| 58527_at | ABRACL | -0,08634614 | 9,29526119 | -0,31955824 | 0,75385306 | 0,88138961 |
| 2975_at | GTF3C1 | 0,01901441 | 8,31655695 | 0,31942214 | 0,75395416 | 0,88138961 |
| 3067_at | HDC | 0,0344168 | 4,85167706 | 0,31934841 | 0,75400893 | 0,88138961 |
| 28978_at | TMEM14A | 0,042685 | 10,4611775 | 0,31928972 | 0,75405252 | 0,88138961 |
| 283229_at | CRACR2B | 0,06340017 | 5,66857094 | 0,31928489 | 0,75405611 | 0,88138961 |
| 2257_at | FGF12 | -0,02074172 | 4,04314221 | -0,31926442 | 0,75407131 | 0,88138961 |
| 4635_at | MYL4 | -0,03527557 | 5,43754452 | -0,31916953 | 0,7541418 | 0,88138961 |
| 101926892_a | LOC10192689 | -0,02239244 | 2,9270008 | -0,31916743 | 0,75414336 | 0,88138961 |
| 101927112_a | LINC01581 | 0,02837966 | 3,22704944 | 0,31907007 | 0,75421569 | 0,88142067 |
| 83758_at | RBP5 | 0,02152879 | 4,81742253 | 0,31898737 | 0,75427713 | 0,88143899 |
| 6525_at | SMTN | -0,08537017 | 6,73595349 | -0,31882825 | 0,75439534 | 0,88148006 |
| 114784_at | CSMD2 | 0,03750389 | 4,02704291 | 0,3187755 | 0,75443454 | 0,88148006 |
| 1345_at | COX6C | -0,03201164 | 10,6825213 | -0,31867949 | 0,75450587 | 0,88148006 |
| 84747_at | UNC119B | -0,03246629 | 9,1159516 | -0,31849449 | 0,75464332 | 0,88148006 |
| 83544_at | DNAL1 | 0,04444583 | 5,23144093 | 0,31848835 | 0,75464788 | 0,88148006 |
| 125111_at | GJD3 | -0,03397785 | 6,33986068 | -0,31838344 | 0,75472584 | 0,88148006 |
| 4246_at | SCGB2A1 | 0,03127244 | 3,30647878 | 0,31837152 | 0,7547347 | 0,88148006 |
| 2670_at | GFAP | -0,037728 | 4,55547269 | -0,31834279 | 0,75475605 | 0,88148006 |
| 283663_at | LINC00926 | -0,02951099 | 5,13771991 | -0,31832971 | 0,75476577 | 0,88148006 |
| 1108_at | CHD4 | 0,02782964 | 8,93101203 | 0,31832418 | 0,75476987 | 0,88148006 |

| | | | | | | |
|---|---|---|---|---|---|---|
| 100996301_a | FOXD3-AS1 | -0,03759641 | 4,71954752 | -0,31823538 | 0,75483586 | 0,88150368 |
| 51176_at | LEF1 | 0,02232927 | 3,75075772 | 0,31813898 | 0,7549075 | 0,8815339 |
| 100506229_a | LINC01093 | 0,03941583 | 3,60515363 | 0,31789309 | 0,75509023 | 0,88169383 |
| 25979_at | DHRS7B | -0,09915418 | 8,14883107 | -0,31780074 | 0,75515887 | 0,88172053 |
| 23762_at | OSBP2 | -0,04085321 | 6,0511843 | -0,31757605 | 0,75532587 | 0,88182593 |
| 101927124_a | LOC10192712 | 0,02143353 | 3,29377896 | 0,3175561 | 0,75534069 | 0,88182593 |
| 26054_at | SENP6 | -0,03614661 | 7,39169666 | -0,31739056 | 0,75546374 | 0,88184577 |
| 122830_at | NAA30 | -0,03210208 | 8,82027227 | -0,3172984 | 0,75553224 | 0,88184577 |
| 80183_at | KIAA0226L | 0,02157672 | 3,24390442 | 0,31727393 | 0,75555043 | 0,88184577 |
| 253559_at | CADM2 | -0,01603651 | 3,26490825 | -0,31716084 | 0,7556345 | 0,88184577 |
| 5978_at | REST | 0,05853724 | 6,9442978 | 0,3171356 | 0,75565326 | 0,88184577 |
| 4604_at | MYBPC1 | 0,03090833 | 3,26896483 | 0,31709202 | 0,75568566 | 0,88184577 |
| 144124_at | OR10A5 | 0,037847 | 3,46963304 | 0,31704076 | 0,75572376 | 0,88184577 |
| 79006_at | METRN | -0,09709861 | 7,18754821 | -0,31704054 | 0,75572392 | 0,88184577 |
| 26088_at | GGA1 | 0,03027763 | 8,976275 | 0,31653927 | 0,7560966 | 0,88211836 |
| 7554_at | ZNF8 | -0,03679998 | 6,56901335 | -0,31653786 | 0,75609765 | 0,88211836 |
| 284933_at | LOC284933 | -0,02974831 | 4,48386223 | -0,3165347 | 0,7561 | 0,88211836 |
| 9168_at | TMSB10 | 0,07568629 | 12,6611641 | 0,31644561 | 0,75616624 | 0,88211836 |
| 9054_at | NFS1 | -0,01823583 | 8,28734621 | -0,31641837 | 0,7561865 | 0,88211836 |
| 259285_at | TAS2R39 | 0,05251777 | 4,16575205 | 0,31629417 | 0,75627885 | 0,88217112 |
| 56142_at | PCDHA6 | -0,03351135 | 4,07591812 | -0,31623436 | 0,75632332 | 0,88217112 |
| 85463_at | ZC3H12C | 0,07747408 | 7,68327027 | 0,31608897 | 0,75643143 | 0,88224381 |
| 54777_at | CFAP46 | 0,02826333 | 4,35843048 | 0,31582877 | 0,75662494 | 0,88240356 |
| 57151_at | LYZL6 | -0,02456295 | 4,66456463 | -0,31573889 | 0,75669178 | 0,88240356 |
| 2936_at | GSR | 0,08000896 | 9,94232445 | 0,31572 | 0,75670583 | 0,88240356 |
| 23474_at | ETHE1 | 0,035025 | 9,58748243 | 0,31549372 | 0,75687413 | 0,88254639 |
| 100506243_a | KRBOX1 | 0,02820979 | 3,99075755 | 0,31541588 | 0,75693203 | 0,88256048 |
| 51528_at | JKAMP | -0,0273563 | 9,31813463 | -0,31531628 | 0,75700611 | 0,88259344 |
| 100507171_a | BOLA3-AS1 | -0,02176323 | 4,27851597 | -0,31524435 | 0,75705961 | 0,88260239 |
| 9158_at | FIBP | 0,0218039 | 10,3064089 | 0,31497619 | 0,75725909 | 0,88273171 |
| 3662_at | IRF4 | -0,02625141 | 4,50686376 | -0,31491613 | 0,75730377 | 0,88273171 |
| 152789_at | JAKMIP1 | -0,02676746 | 3,95839914 | -0,31491044 | 0,75730801 | 0,88273171 |
| 7542_at | ZFPL1 | 0,02815506 | 7,85312952 | 0,31483156 | 0,75736669 | 0,88274669 |
| 80264_at | ZNF430 | 0,03267624 | 6,09462969 | 0,31475868 | 0,75742091 | 0,88275647 |
| 51574_at | LARP7 | 0,08345495 | 7,68719418 | 0,31465206 | 0,75750023 | 0,88276728 |
| 92162_at | TMEM88 | 0,02829413 | 6,06625329 | 0,31434398 | 0,75772946 | 0,88276728 |
| 51474_at | LIMA1 | 0,09215353 | 9,75637314 | 0,31431968 | 0,75774754 | 0,88276728 |
| 10739_at | RFPL2 | -0,03728406 | 4,76023419 | -0,314222 | 0,75782022 | 0,88276728 |
| 170960_at | ZNF721 | -0,04205512 | 8,04662833 | -0,3141153 | 0,75789961 | 0,88276728 |
| 101927829_a | LOC10192782 | -0,03053022 | 3,09983629 | -0,31411208 | 0,75790201 | 0,88276728 |
| 11107_at | PRDM5 | 0,02102912 | 3,26624532 | 0,31410249 | 0,75790915 | 0,88276728 |
| 5147_at | PDE6D | -0,02001719 | 6,9655206 | -0,31408622 | 0,75792125 | 0,88276728 |
| 4221_at | MEN1 | 0,03648589 | 8,49630802 | 0,31401076 | 0,75797741 | 0,88276728 |
| 3059_at | HCLS1 | 0,20899943 | 5,17682704 | 0,31398267 | 0,75799831 | 0,88276728 |
| 11162_at | NUDT6 | -0,06033058 | 6,1495522 | -0,31395807 | 0,75801662 | 0,88276728 |
| 79092_at | CARD14 | 0,02287308 | 5,19898728 | 0,31395103 | 0,75802185 | 0,88276728 |
| 387882_at | C12orf75 | -0,29057225 | 7,15724779 | -0,31394555 | 0,75802593 | 0,88276728 |
| 57462_at | KIAA1161 | -0,03714584 | 6,23376863 | -0,31375038 | 0,75817118 | 0,88286904 |
| 641515_at | MGAT4EP | 0,03165148 | 3,49942927 | 0,31370496 | 0,75820498 | 0,88286904 |
| 2826_at | CCR10 | -0,02695448 | 4,37561623 | -0,31344539 | 0,75839817 | 0,88304061 |

| | | | | | | |
|---|---|---|---|---|---|---|
| 3483_at | IGFALS | 0,0272771 | 5,70117579 | 0,31318085 | 0,75859507 | 0,88321649 |
| 257177_at | CFAP126 | 0,03620132 | 3,98403179 | 0,31311832 | 0,75864162 | 0,8832173 |
| 10351_at | ABCA8 | -0,02349263 | 2,86721167 | -0,31302645 | 0,75871001 | 0,88324354 |
| 118425_at | PCAT4 | 0,04099147 | 3,34805759 | 0,31294484 | 0,75877076 | 0,88326089 |
| 85377_at | MICALL1 | -0,03858711 | 9,74909244 | -0,31266874 | 0,7589763 | 0,88344677 |
| 5172_at | SLC26A4 | -0,01814921 | 3,16735136 | -0,31249592 | 0,75910497 | 0,88354314 |
| 221061_at | FAM171A1 | -0,02896011 | 4,58433273 | -0,31233565 | 0,7592243 | 0,8835827 |
| 57134_at | MAN1C1 | 0,06669889 | 5,40737463 | 0,31231653 | 0,75923853 | 0,8835827 |
| 399948_at | COLCA1 | 0,08525717 | 5,97207612 | 0,31226545 | 0,75927657 | 0,8835827 |
| 2731_at | GLDC | -0,11735181 | 5,62225855 | -0,31219435 | 0,75932951 | 0,88359094 |
| 2834_at | PRLHR | -0,02463495 | 4,34045448 | -0,31208535 | 0,75941067 | 0,883632 |
| 26035_at | GLCE | 0,0770698 | 7,36648828 | 0,3117966 | 0,75962569 | 0,88375895 |
| 89890_at | KBTBD6 | -0,05526729 | 7,49098469 | -0,31174426 | 0,75966467 | 0,88375895 |
| 1538_at | CYLC1 | -0,02730954 | 3,3122277 | -0,31170695 | 0,75969245 | 0,88375895 |
| 64180_at | DPEP3 | -0,03908771 | 4,90500207 | -0,31153849 | 0,75981792 | 0,88375895 |
| 1746_at | DLX2 | -0,05854505 | 5,14864782 | -0,31153481 | 0,75982065 | 0,88375895 |
| 112885_at | PHF21B | -0,02340719 | 4,7571367 | -0,3115093 | 0,75983966 | 0,88375895 |
| 84953_at | MICALCL | 0,0671776 | 5,22720771 | 0,31148206 | 0,75985994 | 0,88375895 |
| 11082_at | ESM1 | -0,02041262 | 3,15727935 | -0,31144599 | 0,7598868 | 0,88375895 |
| 4613_at | MYCN | 0,02438605 | 5,15409217 | 0,31134534 | 0,75996177 | 0,88379278 |
| 8881_at | CDC16 | -0,05307463 | 8,93841898 | -0,31116841 | 0,76009356 | 0,88389268 |
| 2110_at | ETFDH | 0,06242753 | 6,95919177 | 0,31108102 | 0,76015865 | 0,88391501 |
| 199_at | AIF1 | -0,02988572 | 4,96200784 | -0,31094828 | 0,76025753 | 0,88397663 |
| 650655_at | ABCA17P | -0,03605171 | 4,15933598 | -0,3108757 | 0,7603116 | 0,88398613 |
| 1649_at | DDIT3 | -0,05889629 | 6,81819087 | -0,31080346 | 0,76036541 | 0,88399535 |
| 57554_at | LRRC7 | -0,02134752 | 2,9583366 | -0,31051664 | 0,76057908 | 0,88414486 |
| 54997_at | TESC | 0,03426293 | 5,06636875 | 0,31049844 | 0,76059264 | 0,88414486 |
| 101928583_a | LOC10192858 | -0,01602131 | 3,26042996 | -0,310446 | 0,76063171 | 0,88414486 |
| 349136_at | WDR86 | -0,02585067 | 4,50900274 | -0,31033905 | 0,76071139 | 0,88415757 |
| 84034_at | EMILIN2 | -0,05844707 | 5,60156579 | -0,31030811 | 0,76073445 | 0,88415757 |
| 133957_at | CCDC127 | 0,02459099 | 8,50957444 | 0,30998315 | 0,76097658 | 0,88438563 |
| 5055_at | SERPINB2 | -0,19790065 | 6,2711551 | -0,30986068 | 0,76106784 | 0,88439781 |
| 522_at | ATP5J | -0,0320028 | 10,606397 | -0,30984586 | 0,76107888 | 0,88439781 |
| 81550_at | TDRD3 | 0,01819747 | 7,27829461 | 0,30976932 | 0,76113592 | 0,88441073 |
| 55856_at | ACOT13 | -0,09311477 | 9,37134183 | -0,30960891 | 0,76125546 | 0,88443999 |
| 65110_at | UPF3A | 0,03296408 | 6,24354333 | 0,30956356 | 0,76128925 | 0,88443999 |
| 23408_at | SIRT5 | 0,01553984 | 6,58853987 | 0,30955071 | 0,76129883 | 0,88443999 |
| 139716_at | GAB3 | 0,02504294 | 4,49761636 | 0,30940776 | 0,76140537 | 0,88449326 |
| 791114_at | PWRN1 | -0,02603624 | 3,16202238 | -0,30932128 | 0,76146983 | 0,88449326 |
| 128710_at | SLX4IP | -0,02551113 | 6,7591444 | -0,30930435 | 0,76148244 | 0,88449326 |
| 101928886_a | LOC10192888 | -0,02794027 | 3,78004958 | -0,30922099 | 0,76154457 | 0,88451209 |
| 170591_at | S100Z | -0,02870548 | 3,75429999 | -0,30906527 | 0,76166063 | 0,88457797 |
| 114799_at | ESCO1 | 0,04695744 | 6,48243801 | 0,30901007 | 0,76170179 | 0,88457797 |
| 101928140_a | LOC10192814 | 0,02031607 | 3,20726564 | 0,30896007 | 0,76173905 | 0,88457797 |
| 11000_at | SLC27A3 | 0,06087728 | 7,81068152 | 0,30881764 | 0,76184522 | 0,88464793 |
| 221662_at | RBM24 | 0,01930328 | 3,14633802 | 0,30865225 | 0,76196851 | 0,88470387 |
| 122786_at | FRMD6 | 0,0677943 | 8,91153346 | 0,3086298 | 0,76198525 | 0,88470387 |
| 101929162_a | LOC10192916 | -0,03798647 | 3,39841036 | -0,30847353 | 0,76210174 | 0,88474365 |
| 122618_at | PLD4 | -0,0337892 | 4,81305572 | -0,30843035 | 0,76213394 | 0,88474365 |
| 100133985_a | LOC10013398 | -0,030156 | 4,17100678 | -0,30835699 | 0,76218863 | 0,88474365 |

| | | | | | | |
|---|---|---|---|---|---|---|
| 57449_at | PLEKHG5 | -0,02328393 | 5,95970413 | -0,30833741 | 0,76220323 | 0,88474365 |
| 2_at | A2M | 0,0315025 | 3,73154963 | 0,30807328 | 0,76240016 | 0,88487237 |
| 145438_at | FRMD6-AS1 | -0,04201157 | 5,03641141 | -0,30806545 | 0,762406 | 0,88487237 |
| 1109_at | AKR1C4 | 0,02402515 | 3,29954853 | 0,30793147 | 0,7625059 | 0,88492729 |
| 3605_at | IL17A | -0,02754486 | 3,71536689 | -0,30783868 | 0,76257509 | 0,88492729 |
| 11223_at | MST1L | -0,04128015 | 5,09921322 | -0,30776382 | 0,76263091 | 0,88492729 |
| 100129827_a | MRVI1-AS1 | -0,03968446 | 5,13196061 | -0,30775468 | 0,76263773 | 0,88492729 |
| 8218_at | CLTCL1 | -0,02637693 | 5,09645684 | -0,30769395 | 0,76268301 | 0,88492729 |
| 84671_at | ZNF347 | 0,04600262 | 4,97213758 | 0,30746815 | 0,7628514 | 0,88506936 |
| 6335_at | SCN9A | 0,01734199 | 3,02600237 | 0,30739473 | 0,76290615 | 0,88507957 |
| 140711_at | TLDC2 | -0,03242208 | 4,13024182 | -0,30728548 | 0,76298763 | 0,88512079 |
| 63934_at | ZNF667 | -0,02710492 | 3,51981653 | -0,3070264 | 0,76318086 | 0,8852633 |
| 196527_at | ANO6 | -0,03379428 | 9,70444843 | -0,30699754 | 0,76320239 | 0,8852633 |
| 53940_at | FTHL17 | 0,03572941 | 4,20068659 | 0,30652567 | 0,76355437 | 0,88561825 |
| 23360_at | FNBP4 | -0,09983619 | 6,72876803 | -0,30645926 | 0,76360392 | 0,88562239 |
| 666_at | BOK | 0,03156152 | 7,80817551 | 0,30628518 | 0,76373378 | 0,88571968 |
| 6895_at | TARBP2 | -0,02761703 | 8,11629929 | -0,30611572 | 0,76386022 | 0,88581298 |
| 2559_at | GABRA6 | -0,02628063 | 3,22349951 | -0,30574531 | 0,76413659 | 0,88603501 |
| 4696_at | NDUFA3 | 0,04624281 | 10,2828776 | 0,30573107 | 0,76414721 | 0,88603501 |
| 9452_at | ITM2A | -0,02011414 | 3,28772713 | -0,30559678 | 0,76424742 | 0,88603501 |
| 94059_at | LENG9 | 0,09128188 | 6,6587093 | 0,30558381 | 0,7642571 | 0,88603501 |
| 65124_at | SOWAHC | 0,15161922 | 9,37509838 | 0,30550104 | 0,76431886 | 0,88603501 |
| 54923_at | LIME1 | -0,0452621 | 6,7084507 | -0,30543981 | 0,76436456 | 0,88603501 |
| 10994_at | ILVBL | 0,0636275 | 8,42830073 | 0,30532218 | 0,76445234 | 0,88603501 |
| 56913_at | C1GALT1 | -0,05275509 | 7,34131142 | -0,30530805 | 0,76446288 | 0,88603501 |
| 4256_at | MGP | -0,09004868 | 4,03921503 | -0,30530435 | 0,76446565 | 0,88603501 |
| 344905_at | ATP13A5 | 0,03109872 | 3,38411855 | 0,30522817 | 0,7645225 | 0,88604759 |
| 51588_at | PIAS4 | 0,02632155 | 7,60218467 | 0,30479827 | 0,76484336 | 0,88636612 |
| 26273_at | FBXO3 | 0,08244007 | 6,85205261 | 0,3046324 | 0,76496717 | 0,88645627 |
| 55285_at | RBM41 | 0,02246521 | 6,37869734 | 0,30444812 | 0,76510474 | 0,88648825 |
| 92070_at | CTBP1-AS2 | 0,08006415 | 7,78055291 | 0,30440365 | 0,76513793 | 0,88648825 |
| 7225_at | TRPC6 | -0,02311353 | 3,29084958 | -0,30435702 | 0,76517274 | 0,88648825 |
| 5266_at | PI3 | -0,15453672 | 6,39500445 | -0,30434884 | 0,76517885 | 0,88648825 |
| 200132_at | TCTEX1D1 | 0,01789594 | 3,31389897 | 0,30410756 | 0,76535897 | 0,88664361 |
| 55734_at | ZFP64 | 0,01360389 | 6,95906446 | 0,30398877 | 0,76544766 | 0,88669302 |
| 2173_at | FABP7 | -0,01964733 | 3,51965211 | -0,30381009 | 0,76558106 | 0,88679423 |
| 101927574_a | LINC01186 | -0,03959081 | 4,47915108 | -0,30349943 | 0,76581303 | 0,88696888 |
| 88_at | ACTN2 | -0,01976995 | 4,02244526 | -0,30348484 | 0,76582393 | 0,88696888 |
| 5226_at | PGD | -0,02366007 | 11,417533 | -0,30335829 | 0,76591843 | 0,88698627 |
| 100132891_a | MSC-AS1 | 0,02889959 | 4,65618623 | 0,30330915 | 0,76595512 | 0,88698627 |
| 23254_at | KAZN | 0,03047204 | 6,08202278 | 0,30327974 | 0,76597709 | 0,88698627 |
| 1427_at | CRYGS | -0,05745269 | 4,92409879 | -0,3031625 | 0,76606464 | 0,88703434 |
| 729059_at | C1orf234 | 0,03806071 | 3,45097823 | 0,30302539 | 0,76616704 | 0,88709958 |
| 11177_at | BAZ1A | -0,05865219 | 9,17530555 | -0,30266341 | 0,76643739 | 0,88731193 |
| 340542_at | BEX5 | 0,09644155 | 5,14090661 | 0,30265648 | 0,76644257 | 0,88731193 |
| 51105_at | PHF20L1 | -0,07001921 | 6,99103613 | -0,30240387 | 0,76663126 | 0,88737776 |
| 84971_at | ATG4D | 0,0253279 | 7,50558402 | 0,30239872 | 0,76663511 | 0,88737776 |
| 26287_at | ANKRD2 | 0,04714384 | 6,13564575 | 0,30231711 | 0,76669606 | 0,88737776 |
| 11127_at | KIF3A | 0,03946985 | 6,83392367 | 0,30230078 | 0,76670827 | 0,88737776 |
| 101928847_a | LOC10192884 | -0,02546136 | 3,73578272 | -0,30221784 | 0,76677022 | 0,88737776 |

| | | | | | | |
|---|---|---|---|---|---|---|
| 100506274_a | LOC10050627 | -0,02779247 | 3,07637849 | -0,30220995 | 0,76677611 | 0,88737776 |
| 253018_at | HCG27 | -0,04068032 | 4,89833662 | -0,30213006 | 0,7668358 | 0,88737776 |
| 100506272_a | LOC10050627 | -0,02447443 | 3,67136961 | -0,30208302 | 0,76687094 | 0,88737776 |
| 151987_at | PPP4R2 | -0,11438881 | 7,61737942 | -0,30202534 | 0,76691403 | 0,88737776 |
| 124626_at | ZPBP2 | -0,03206565 | 3,21597249 | -0,30189789 | 0,76700925 | 0,88741548 |
| 56907_at | SPIRE1 | 0,09228705 | 6,6991614 | 0,30185838 | 0,76703876 | 0,88741548 |
| 479_at | ATP12A | 0,05393345 | 4,25878216 | 0,30174901 | 0,76712048 | 0,88745672 |
| 5763_at | PTMS | 0,23976653 | 7,44349181 | 0,30156065 | 0,76726121 | 0,88756622 |
| 11202_at | KLK8 | -0,20266817 | 6,25206623 | -0,30143244 | 0,76735702 | 0,88762375 |
| 55663_at | ZNF446 | 0,04521547 | 6,81870212 | 0,30133141 | 0,76743251 | 0,88765777 |
| 10944_at | C11orf58 | -0,05831226 | 6,85016143 | -0,30125278 | 0,76749127 | 0,88767243 |
| 57144_at | PAK7 | 0,02279408 | 3,65735801 | 0,30112972 | 0,76758322 | 0,88768803 |
| 361_at | AQP4 | 0,01594081 | 3,03922586 | 0,30111139 | 0,76759693 | 0,88768803 |
| 27246_at | RNF115 | -0,04760456 | 8,24107055 | -0,30104579 | 0,76764595 | 0,88769143 |
| 101927450_a | LOC10192745 | -0,01989293 | 3,03582937 | -0,30079131 | 0,76783613 | 0,88785806 |
| 1528_at | CYB5A | -0,04007492 | 9,36539066 | -0,30070123 | 0,76790345 | 0,88788261 |
| 100507297_a | TMEM44-AS1 | 0,0723775 | 5,57541985 | 0,30060453 | 0,76797573 | 0,88790787 |
| 56986_at | DTWD1 | -0,02571795 | 5,66831147 | -0,30052486 | 0,76803528 | 0,88790787 |
| 827_at | CAPN6 | 0,02941687 | 3,75193934 | 0,30048698 | 0,76806359 | 0,88790787 |
| 222008_at | VSTM2A | -0,02085156 | 3,459433 | -0,29986666 | 0,76852729 | 0,88839062 |
| 8365_at | HIST1H4H | -0,06638405 | 6,31511885 | -0,29927321 | 0,76897099 | 0,88881023 |
| 56287_at | GKN1 | 0,16975639 | 4,85024855 | 0,29925772 | 0,76898257 | 0,88881023 |
| 9545_at | RAB3D | 0,0803797 | 6,15147678 | 0,29876541 | 0,76935072 | 0,88918239 |
| 100192386_a | FLJ16779 | -0,03141869 | 3,9537443 | -0,29853512 | 0,76952295 | 0,88932809 |
| 5919_at | RARRES2 | -0,03217025 | 4,93062922 | -0,29836689 | 0,76964878 | 0,8893898 |
| 23326_at | USP22 | -0,06306694 | 10,0503665 | -0,29834027 | 0,76966869 | 0,8893898 |
| 126520_at | PLK5 | -0,02995793 | 5,63026222 | -0,29816099 | 0,76980278 | 0,88940917 |
| 619351_at | LINC00589 | -0,04130995 | 4,54917916 | -0,29808723 | 0,76985796 | 0,88940917 |
| 100507064_a | TEX26-AS1 | -0,02347859 | 3,3132061 | -0,29808466 | 0,76985988 | 0,88940917 |
| 4061_at | LY6E | -0,12187179 | 9,59993274 | -0,29807094 | 0,76987014 | 0,88940917 |
| 9922_at | IQSEC1 | -0,05757156 | 7,44590001 | -0,29798351 | 0,76993554 | 0,88943139 |
| 139420_at | PPP4R3CP | 0,0195558 | 2,87444305 | 0,29768599 | 0,77015811 | 0,88957499 |
| 579_at | NKX3-2 | 0,03509175 | 5,16976011 | 0,29764389 | 0,77018961 | 0,88957499 |
| 79066_at | METTL16 | -0,04941989 | 6,32381413 | -0,29763214 | 0,77019839 | 0,88957499 |
| 9630_at | GNA14 | -0,03136342 | 4,02703832 | -0,29754897 | 0,77026062 | 0,88959352 |
| 79178_at | THTPA | 0,0792646 | 7,15355421 | 0,29741242 | 0,77036278 | 0,88964293 |
| 200261_at | LINC00656 | -0,0359554 | 5,4836942 | -0,29736832 | 0,77039577 | 0,88964293 |
| 101927900_a | LINC01477 | -0,02844029 | 3,65789615 | -0,29719901 | 0,77052245 | 0,88973588 |
| 11117_at | EMILIN1 | 0,03137313 | 5,07059393 | 0,29687604 | 0,77076412 | 0,88991747 |
| 388323_at | GLTPD2 | -0,03302971 | 5,44649362 | -0,29686537 | 0,7707721 | 0,88991747 |
| 10327_at | AKR1A1 | -0,05081167 | 10,0085568 | -0,29670001 | 0,77089585 | 0,89000699 |
| 29099_at | COMMD9 | -0,04029709 | 7,37306983 | -0,29637022 | 0,77114266 | 0,89021355 |
| 10_at | NAT2 | -0,03595197 | 4,35603252 | -0,2963046 | 0,77119176 | 0,89021355 |
| 3960_at | LGALS4 | -0,0529686 | 5,16252277 | -0,2962757 | 0,7712134 | 0,89021355 |
| 57165_at | GJC2 | -0,03224304 | 6,37585087 | -0,29619995 | 0,7712701 | 0,89022565 |
| 4084_at | MXD1 | 0,09540167 | 6,75301974 | 0,2958194 | 0,77155494 | 0,89045173 |
| 10350_at | ABCA9 | 0,02136932 | 3,19919231 | 0,29581475 | 0,77155842 | 0,89045173 |
| 28904_at | IGKV1D-8 | -0,0421441 | 3,77297061 | -0,29549179 | 0,77180019 | 0,8905593 |
| 5458_at | POU4F2 | -0,0293311 | 3,97082654 | -0,29548843 | 0,7718027 | 0,8905593 |
| 1241_at | LTB4R | -0,05040333 | 5,34329589 | -0,29546696 | 0,77181877 | 0,8905593 |

| | | | | | | |
|---|---|---|---|---|---|---|
| 101927420_a | LOC10192742 | -0,05851918 | 5,92279678 | -0,29542082 | 0,77185331 | 0,8905593 |
| 5524_at | PPP2R4 | -0,03052689 | 8,3658621 | -0,29532304 | 0,77192652 | 0,8905593 |
| 161394_at | SAMD15 | 0,03563097 | 4,67621782 | 0,29527959 | 0,77195905 | 0,8905593 |
| 81870_at | KRTAP9-9 | -0,02936347 | 3,21224918 | -0,29525795 | 0,77197525 | 0,8905593 |
| 161725_at | OTUD7A | -0,02470967 | 4,82725029 | -0,29512518 | 0,77207465 | 0,89062064 |
| 65061_at | CDK15 | -0,02702668 | 4,30790367 | -0,29491896 | 0,77222906 | 0,89063878 |
| 26276_at | VPS33B | -0,03677786 | 7,07943097 | -0,29489658 | 0,77224581 | 0,89063878 |
| 248_at | ALPI | 0,03078699 | 5,05336451 | 0,29484809 | 0,77228212 | 0,89063878 |
| 57569_at | ARHGAP20 | -0,08747811 | 5,07877509 | -0,29475682 | 0,77235047 | 0,89063878 |
| 11120_at | BTN2A1 | 0,04986036 | 7,64245074 | 0,2947562 | 0,77235093 | 0,89063878 |
| 6884_at | TAF13 | -0,1047685 | 5,89356563 | -0,29467711 | 0,77241015 | 0,89063878 |
| 149483_at | CCDC17 | -0,06947727 | 5,09842832 | -0,29467194 | 0,77241403 | 0,89063878 |
| 80258_at | EFHC2 | 0,01656708 | 3,14982016 | 0,29457973 | 0,77248307 | 0,8906522 |
| 388387_at | LINC00671 | -0,03937064 | 4,56331495 | -0,29453291 | 0,77251814 | 0,8906522 |
| 90527_at | DUOXA1 | -0,06326189 | 6,30254291 | -0,29427331 | 0,77271255 | 0,89079211 |
| 386724_at | AMIGO3 | 0,06505704 | 7,09918369 | 0,2941232 | 0,77282497 | 0,89079211 |
| 123_at | PLIN2 | -0,25496081 | 7,6366585 | -0,2940015 | 0,77291611 | 0,89079211 |
| 1738_at | DLD | -0,0414322 | 8,2242542 | -0,29397118 | 0,77293883 | 0,89079211 |
| 151647_at | FAM19A4 | 0,02354716 | 3,28989301 | 0,29394377 | 0,77295935 | 0,89079211 |
| 206412_at | C6orf163 | 0,01541294 | 3,67051258 | 0,29391556 | 0,77298048 | 0,89079211 |
| 222663_at | SCUBE3 | 0,02129541 | 5,5245493 | 0,2938858 | 0,77300277 | 0,89079211 |
| 65999_at | LRRC61 | 0,10780398 | 7,77656699 | 0,2938414 | 0,77303603 | 0,89079211 |
| 51284_at | TLR7 | -0,01884439 | 3,82339311 | -0,29381516 | 0,77305569 | 0,89079211 |
| 116115_at | ZNF526 | 0,03282672 | 6,80108131 | 0,29373592 | 0,77311504 | 0,89080721 |
| 100505826_a | STK4-AS1 | -0,04119121 | 5,62755277 | -0,29342795 | 0,77334573 | 0,89096822 |
| 7274_at | TTPA | -0,03685948 | 3,92911049 | -0,29342589 | 0,77334728 | 0,89096822 |
| 64745_at | METTL17 | -0,05188171 | 9,18932173 | -0,29335494 | 0,77340043 | 0,89097616 |
| 4991_at | OR1D2 | -0,05553624 | 5,29353222 | -0,29301939 | 0,77365181 | 0,89118233 |
| 64843_at | ISL2 | -0,06016886 | 5,80102645 | -0,2929754 | 0,77368476 | 0,89118233 |
| 23361_at | ZNF629 | 0,01932539 | 7,48004837 | 0,29289721 | 0,77374335 | 0,89118233 |
| 25925_at | ZNF521 | -0,01845723 | 3,01582716 | -0,29286906 | 0,77376444 | 0,89118233 |
| 5746_at | PTH2R | 0,02678089 | 3,62356996 | 0,29258159 | 0,77397983 | 0,89135209 |
| 55521_at | TRIM36 | -0,03723341 | 4,04803616 | -0,29253962 | 0,77401128 | 0,89135209 |
| 5502_at | PPP1R1A | -0,02094037 | 4,73102393 | -0,29248708 | 0,77405065 | 0,89135209 |
| 90288_at | EFCAB12 | -0,03906244 | 4,96578923 | -0,2922067 | 0,77426076 | 0,89145683 |
| 100302691_a | LINC00184 | 0,02842278 | 3,87099531 | 0,29220068 | 0,77426527 | 0,89145683 |
| 158135_at | TTLL11 | -0,02126794 | 5,56115207 | -0,29214595 | 0,77430629 | 0,89145683 |
| 100506115_a | LINC01272 | 0,03903325 | 4,31371786 | 0,29211867 | 0,77432673 | 0,89145683 |
| 123036_at | TC2N | 0,05532823 | 7,66478416 | 0,29184177 | 0,77453425 | 0,89164246 |
| 80152_at | CENPT | 0,03068156 | 6,58360404 | 0,29137027 | 0,77488766 | 0,89192452 |
| 6785_at | ELOVL4 | 0,11246112 | 4,6181786 | 0,2913421 | 0,77490878 | 0,89192452 |
| 54059_at | YBEY | 0,10698753 | 7,21223561 | 0,29132956 | 0,77491818 | 0,89192452 |
| 26998_at | FETUB | -0,0205201 | 3,30447128 | -0,29088111 | 0,77525436 | 0,89225815 |
| 7553_at | ZNF7 | 0,0621574 | 6,85971304 | 0,29062274 | 0,77544806 | 0,89242777 |
| 1748_at | DLX4 | 0,03351245 | 4,69211069 | 0,29041513 | 0,77560373 | 0,89255359 |
| 3221_at | HOXC4 | -0,06664151 | 6,39171415 | -0,29019335 | 0,77577002 | 0,89265514 |
| 57817_at | HAMP | -0,02872978 | 4,9223512 | -0,29015176 | 0,77580122 | 0,89265514 |
| 65992_at | DDRGK1 | 0,04314687 | 9,14629482 | 0,29011205 | 0,77583099 | 0,89265514 |
| 254863_at | TMEM256 | 0,05661889 | 8,51099748 | 0,2899416 | 0,77595881 | 0,89274889 |
| 819_at | CAMLG | 0,02047479 | 8,6910981 | 0,28957163 | 0,77623628 | 0,89301478 |

| | | | | | | |
|---|---|---|---|---|---|---|
| 22985_at | ACIN1 | -0,03525064 | 7,48748527 | -0,28940603 | 0,77636048 | 0,89307972 |
| 284454_at | LOC284454 | -0,05992272 | 7,11283647 | -0,28937274 | 0,77638545 | 0,89307972 |
| 55012_at | PPP2R3C | 0,03444976 | 8,00253052 | 0,28897354 | 0,77668489 | 0,89337081 |
| 22836_at | RHOBTB3 | 0,02502122 | 11,0886406 | 0,28888358 | 0,77675238 | 0,89337488 |
| 29887_at | SNX10 | -0,04353928 | 7,28941384 | -0,28884518 | 0,77678118 | 0,89337488 |
| 340074_at | LOC340074 | -0,03637055 | 4,1321205 | -0,28828211 | 0,77720362 | 0,89380736 |
| 83895_at | KRTAP1-5 | 0,02972733 | 4,71895133 | 0,28766352 | 0,77766779 | 0,89415408 |
| 63925_at | ZNF335 | -0,03680059 | 6,3216751 | -0,28765115 | 0,77767708 | 0,89415408 |
| 11163_at | NUDT4 | 0,05391759 | 6,70637729 | 0,28763335 | 0,77769043 | 0,89415408 |
| 654433_at | PAX8-AS1 | 0,02054516 | 4,55272042 | 0,28757915 | 0,77773111 | 0,89415408 |
| 100131138_a | LINC01405 | -0,02261197 | 3,03600304 | -0,28757104 | 0,77773719 | 0,89415408 |
| 57419_at | SLC24A3 | 0,13732228 | 5,08325224 | 0,28745219 | 0,77782639 | 0,89416011 |
| 389_at | RHOC | -0,02733332 | 10,7295579 | -0,28741367 | 0,7778553 | 0,89416011 |
| 84196_at | USP48 | -0,02077205 | 7,3301773 | -0,28733214 | 0,77791649 | 0,89416011 |
| 100996579_a | LOC10099657 | -0,02426559 | 5,78326765 | -0,28728292 | 0,77795343 | 0,89416011 |
| 57115_at | PGLYRP4 | -0,07819117 | 5,79300214 | -0,2872548 | 0,77797454 | 0,89416011 |
| 55423_at | SIRPG | -0,04404529 | 3,9339083 | -0,28718508 | 0,77802686 | 0,89416376 |
| 402573_at | C7orf61 | -0,04551722 | 5,81540967 | -0,28712687 | 0,77807055 | 0,89416376 |
| 1459_at | CSNK2A2 | 0,02948287 | 8,67844802 | 0,28700593 | 0,77816133 | 0,89421474 |
| 1859_at | DYRK1A | 0,01908974 | 6,32027581 | 0,28682549 | 0,77829677 | 0,89431703 |
| 3226_at | HOXC10 | -0,06008423 | 4,66263144 | -0,28674666 | 0,77835594 | 0,89433167 |
| 100506082_a | LOC10050608 | -0,01911118 | 3,99456116 | -0,28655955 | 0,7784964 | 0,89436556 |
| 57455_at | REXO1 | -0,01988372 | 6,42252851 | -0,28649106 | 0,77854782 | 0,89436556 |
| 6483_at | ST3GAL2 | 0,02186492 | 6,1522515 | 0,28647036 | 0,77856336 | 0,89436556 |
| 10217_at | CTDSPL | -0,08013066 | 8,62286945 | -0,28645998 | 0,77857115 | 0,89436556 |
| 203245_at | NAIF1 | 0,02318654 | 7,10220095 | 0,2862734 | 0,77871123 | 0,89447313 |
| 10206_at | TRIM13 | -0,23377052 | 5,90163481 | -0,28608069 | 0,77885591 | 0,89453352 |
| 3216_at | HOXB6 | 0,14176369 | 7,47066575 | 0,28594392 | 0,7789586 | 0,89453352 |
| 127435_at | PODN | -0,0331063 | 4,95253401 | -0,28586137 | 0,77902058 | 0,89453352 |
| 221785_at | ZSCAN25 | -0,01839219 | 6,80073274 | -0,28585867 | 0,77902261 | 0,89453352 |
| 4867_at | NPHP1 | -0,04771062 | 5,00595209 | -0,28584351 | 0,77903399 | 0,89453352 |
| 389432_at | SAMD5 | 0,13917121 | 6,13858955 | 0,28578682 | 0,77907656 | 0,89453352 |
| 5648_at | MASP1 | -0,02459563 | 4,4404925 | -0,28577043 | 0,77908887 | 0,89453352 |
| 388942_at | LOC388942 | -0,07501225 | 4,79059475 | -0,2855174 | 0,77927887 | 0,89463721 |
| 9647_at | PPM1F | 0,01791555 | 7,20963971 | 0,28541777 | 0,77935368 | 0,89463721 |
| 337_at | APOA4 | -0,02844255 | 5,30771886 | -0,28540077 | 0,77936645 | 0,89463721 |
| 84059_at | ADGRV1 | 0,040331 | 3,7259781 | 0,28534309 | 0,77940977 | 0,89463721 |
| 8785_at | MATN4 | -0,03020857 | 5,48534648 | -0,28534093 | 0,77941139 | 0,89463721 |
| 100507600_a | LOC10050760 | -0,03148503 | 3,84795233 | -0,28525012 | 0,77947959 | 0,89466178 |
| 138948_at | DBH-AS1 | 0,03976251 | 4,59956572 | 0,28518873 | 0,77952569 | 0,89466178 |
| 9378_at | NRXN1 | 0,02250708 | 3,48150817 | 0,28502178 | 0,77965108 | 0,89475238 |
| 284001_at | CCDC57 | -0,03033645 | 5,65149632 | -0,28495409 | 0,77970192 | 0,89475742 |
| 23562_at | CLDN14 | 0,019243 | 4,47274813 | 0,28474716 | 0,77985734 | 0,89488246 |
| 56143_at | PCDHA5 | 0,03026181 | 3,8093915 | 0,28445922 | 0,78007362 | 0,89504072 |
| 1395_at | CRHR2 | -0,02964979 | 4,38988269 | -0,28443983 | 0,78008818 | 0,89504072 |
| 3071_at | NCKAP1L | -0,03035547 | 4,80657791 | -0,28434839 | 0,78015688 | 0,89505735 |
| 100506603_a | LOC10050660 | -0,02174558 | 5,81783413 | -0,28427935 | 0,78020874 | 0,89505735 |
| 389677_at | RBM12B | 0,02118184 | 6,19230911 | 0,28422508 | 0,78024951 | 0,89505735 |
| 9752_at | PCDHA9 | -0,02750747 | 3,85855102 | -0,28416711 | 0,78029306 | 0,89505735 |
| 146845_at | CFAP52 | -0,02689231 | 3,96801426 | -0,28406219 | 0,78037188 | 0,89505735 |

| | | | | | | |
|---|---|---|---|---|---|---|
| 63908_at | NAPB | 0,04015562 | 7,11797165 | 0,28401954 | 0,78040392 | 0,89505735 |
| 643314_at | KIAA0754 | -0,03394609 | 3,66029475 | -0,28398758 | 0,78042793 | 0,89505735 |
| 50613_at | UBQLN3 | -0,02965096 | 4,05027686 | -0,28390026 | 0,78049353 | 0,89507634 |
| 100507421_a | TMEM178B | -0,02181766 | 4,93326962 | -0,28384185 | 0,78053742 | 0,89507634 |
| 22933_at | SIRT2 | 0,04377251 | 7,78849365 | 0,28373434 | 0,7806182 | 0,89511568 |
| 8569_at | MKNK1 | -0,0471906 | 8,5767686 | -0,28353641 | 0,78076691 | 0,89523292 |
| 148479_at | PHF13 | -0,02721441 | 8,2826423 | -0,28345361 | 0,78082913 | 0,89525096 |
| 101954204_a | LINC00644 | 0,01466786 | 2,7523779 | 0,28317055 | 0,78104183 | 0,89541346 |
| 9701_at | PPP6R2 | 0,0386949 | 7,03413937 | 0,28312853 | 0,78107341 | 0,89541346 |
| 100652770_a | DSG2-AS1 | -0,0975108 | 4,69525284 | -0,28307943 | 0,78111031 | 0,89541346 |
| 339400_at | FLG-AS1 | 0,04796074 | 4,21733739 | 0,28291029 | 0,78123741 | 0,89550588 |
| 79154_at | DHRS11 | -0,13199743 | 7,5896652 | -0,28268521 | 0,78140658 | 0,89564649 |
| 63895_at | PIEZO2 | -0,01513199 | 3,83891673 | -0,28259927 | 0,78147117 | 0,89566723 |
| 57038_at | RARS2 | -0,01870882 | 7,59312976 | -0,28218422 | 0,78178314 | 0,89587572 |
| 123688_at | HYKK | -0,06952984 | 5,98281434 | -0,28217989 | 0,78178639 | 0,89587572 |
| 23547_at | LILRA4 | -0,0901767 | 5,47336339 | -0,28216388 | 0,78179843 | 0,89587572 |
| 9108_at | MTMR7 | -0,02455446 | 3,26659854 | -0,28210977 | 0,78183911 | 0,89587572 |
| 254956_at | MORN5 | -0,02387596 | 3,375863 | -0,28179115 | 0,78207862 | 0,89602531 |
| 1776_at | DNASE1L3 | -0,0185474 | 3,48837258 | -0,28178517 | 0,78208312 | 0,89602531 |
| 120227_at | CYP2R1 | -0,05491503 | 6,19339627 | -0,28175047 | 0,7821092 | 0,89602531 |
| 64792_at | IFT22 | -0,01678021 | 7,99337462 | -0,28165905 | 0,78217794 | 0,89605077 |
| 192286_at | HIGD2A | -0,03464564 | 10,0705168 | -0,28152863 | 0,78227599 | 0,89605728 |
| 9389_at | SLC22A14 | -0,0393078 | 6,47568547 | -0,28147776 | 0,78231424 | 0,89605728 |
| 155368_at | WBSCR27 | -0,0350399 | 6,56174173 | -0,28146587 | 0,78232318 | 0,89605728 |
| 100129196_a | MATN1-AS1 | 0,03616801 | 4,2355588 | 0,28134081 | 0,7824172 | 0,89611169 |
| 134864_at | TAAR1 | -0,04092972 | 2,99663287 | -0,28125169 | 0,78248421 | 0,89611702 |
| 50804_at | MYEF2 | -0,02314076 | 3,278408 | -0,28121088 | 0,7825149 | 0,89611702 |
| 10059_at | DNM1L | 0,03575298 | 8,74837712 | 0,28092807 | 0,78272755 | 0,89624649 |
| 9070_at | ASH2L | -0,05835341 | 8,56547433 | -0,2809224 | 0,78273182 | 0,89624649 |
| 283514_at | SIAH3 | 0,01557493 | 3,79879271 | 0,28080904 | 0,78281706 | 0,89624649 |
| 285600_at | KIAA0825 | -0,02361515 | 2,86556648 | -0,28078643 | 0,78283407 | 0,89624649 |
| #N/A | LL22NC03-63E | -0,03695576 | 4,17611603 | -0,28068944 | 0,78290701 | 0,89624649 |
| 340533_at | KIAA2022 | 0,0173032 | 3,32196341 | 0,2806893 | 0,78290711 | 0,89624649 |
| 4017_at | LOXL2 | -0,03848334 | 5,97906877 | -0,28057662 | 0,78299185 | 0,89624715 |
| 221718_at | LINC00518 | 0,02033467 | 3,06094897 | 0,28052478 | 0,78303084 | 0,89624715 |
| 150356_at | CHADL | 0,02214312 | 5,89586378 | 0,2804865 | 0,78305963 | 0,89624715 |
| 168741_at | PER4 | 0,03698883 | 3,79800651 | 0,28038111 | 0,78313889 | 0,89624715 |
| 84626_at | KRBA1 | 0,02920221 | 7,0036507 | 0,28037921 | 0,78314032 | 0,89624715 |
| 116988_at | AGAP3 | 0,02796853 | 7,90443871 | 0,28027748 | 0,78321683 | 0,89625583 |
| 1362_at | CPD | 0,083422 | 8,65696619 | 0,28024539 | 0,78324096 | 0,89625583 |
| 23221_at | RHOBTB2 | -0,0650672 | 7,33998657 | -0,28013411 | 0,78332466 | 0,89629837 |
| 1603_at | DAD1 | -0,02041672 | 11,6072215 | -0,28002373 | 0,78340769 | 0,89634012 |
| 143903_at | LAYN | 0,01749272 | 3,88517804 | 0,279857 | 0,7835331 | 0,89642531 |
| 139105_at | BEND2 | -0,01907049 | 3,2283973 | -0,27980101 | 0,78357522 | 0,89642531 |
| 27039_at | PKD2L2 | 0,01450681 | 2,93157623 | 0,27968612 | 0,78366164 | 0,89643598 |
| 6046_at | BRD2 | 0,05881581 | 9,56264324 | 0,27964144 | 0,78369526 | 0,89643598 |
| 3708_at | ITPR1 | 0,04470561 | 5,69948215 | 0,27957263 | 0,78374701 | 0,89643598 |
| 81621_at | KAZALD1 | -0,03494031 | 6,06746799 | -0,2794728 | 0,78382212 | 0,89643598 |
| 374618_at | TEX9 | -0,0751322 | 4,95080192 | -0,27944608 | 0,78384222 | 0,89643598 |
| 717_at | C2 | 0,02231242 | 3,94947605 | 0,27941744 | 0,78386377 | 0,89643598 |

| | | | | | | |
|---|---|---|---|---|---|---|
| 338069_at | ST7-OT4 | 0,02280448 | 3,38705293 | 0,27925443 | 0,7839864 | 0,89652269 |
| 4943_at | TBC1D25 | 0,02879829 | 6,95476151 | 0,27919294 | 0,78403267 | 0,89652269 |
| 2622_at | GAS8 | -0,03650173 | 7,17088648 | -0,27885657 | 0,78428575 | 0,89675885 |
| 4968_at | OGG1 | 0,01975648 | 6,26037508 | 0,27868788 | 0,78441268 | 0,89685075 |
| 199899_at | LINC00466 | -0,0382491 | 4,47896404 | -0,27854624 | 0,78451927 | 0,89691938 |
| 4634_at | MYL3 | -0,0372313 | 5,76437814 | -0,27846145 | 0,78458308 | 0,89693894 |
| 9456_at | HOMER1 | -0,03469609 | 8,39182454 | -0,27839976 | 0,7846295 | 0,89693894 |
| 137695_at | TMEM68 | 0,05020068 | 7,85939696 | 0,27817663 | 0,78479742 | 0,89707765 |
| 10500_at | SEMA6C | -0,02620034 | 5,96563473 | -0,27777649 | 0,78509858 | 0,89736866 |
| 643977_at | FLJ32255 | 0,08418537 | 6,7092965 | 0,27753129 | 0,78528314 | 0,89745523 |
| 1773_at | DNASE1 | -0,02383333 | 5,49590348 | -0,27750263 | 0,78530472 | 0,89745523 |
| 10669_at | CGREF1 | -0,03130133 | 5,1969542 | -0,27749017 | 0,78531409 | 0,89745523 |
| 147719_at | LYPD4 | -0,02244281 | 3,67586349 | -0,27739639 | 0,78538468 | 0,89746621 |
| 947_at | CD34 | 0,03384832 | 4,93228464 | 0,27735362 | 0,78541688 | 0,89746621 |
| 102724965_a | LOC10272496 | 0,05872174 | 4,83544099 | 0,27709588 | 0,78561091 | 0,89763467 |
| 7752_at | ZNF200 | 0,02040673 | 7,35888496 | 0,27700507 | 0,78567927 | 0,89765954 |
| 432_at | ASGR1 | -0,03151454 | 4,61362157 | -0,27689378 | 0,78576306 | 0,89770202 |
| 647219_at | ASCL5 | -0,01956991 | 4,73841046 | -0,27679383 | 0,78583831 | 0,89773475 |
| 6583_at | SLC22A4 | -0,12318482 | 5,96281229 | -0,27645364 | 0,78609445 | 0,89797411 |
| 54753_at | ZNF853 | -0,02157589 | 4,74388135 | -0,27623804 | 0,7862568 | 0,8981063 |
| 123041_at | SLC24A4 | -0,03306803 | 3,78153068 | -0,27582927 | 0,78656463 | 0,89838615 |
| 9755_at | TBKBP1 | 0,01917673 | 5,78908484 | 0,27573431 | 0,78663615 | 0,89838615 |
| 202309_at | GAPT | -0,01922244 | 2,98502839 | -0,27562865 | 0,78671572 | 0,89838615 |
| 8731_at | RNMT | -0,05440689 | 5,66123675 | -0,27552755 | 0,78679187 | 0,89838615 |
| 2539_at | G6PD | 0,0768874 | 10,3389814 | 0,27549037 | 0,78681987 | 0,89838615 |
| 10267_at | RAMP1 | 0,02583507 | 6,27114731 | 0,27544606 | 0,78685324 | 0,89838615 |
| 115557_at | ARHGEF25 | -0,02861555 | 5,57562273 | -0,27542093 | 0,78687217 | 0,89838615 |
| 101928604_a | ZBTB46-AS1 | -0,0350395 | 4,63837513 | -0,27541731 | 0,7868749 | 0,89838615 |
| 283129_at | C11orf85 | -0,03494778 | 3,97502894 | -0,27520757 | 0,78703288 | 0,89851327 |
| 4602_at | MYB | -0,06583883 | 5,88243734 | -0,27493221 | 0,7872403 | 0,89869681 |
| 3559_at | IL2RA | 0,01948113 | 4,26857338 | 0,27482198 | 0,78732334 | 0,89873834 |
| 57542_at | KLHL42 | -0,01731978 | 7,75714752 | -0,27463781 | 0,78746209 | 0,89884346 |
| 55740_at | ENAH | 0,02179591 | 8,14965436 | 0,27438056 | 0,7876559 | 0,89898476 |
| 11064_at | CNTRL | -0,02530065 | 5,34918634 | -0,27434961 | 0,78767922 | 0,89898476 |
| 56130_at | PCDHB6 | 0,02565509 | 3,48732795 | 0,27405333 | 0,78790246 | 0,89918627 |
| 6793_at | STK10 | -0,02112115 | 7,10028715 | -0,27382569 | 0,78807399 | 0,89923904 |
| 30816_at | ERVW-1 | 0,02078305 | 3,30766431 | 0,27381135 | 0,78808479 | 0,89923904 |
| 999_at | CDH1 | -0,0433929 | 9,91471763 | -0,27380611 | 0,78808874 | 0,89923904 |
| 10798_at | OR5I1 | -0,04590858 | 3,6096854 | -0,27365247 | 0,78820453 | 0,89931788 |
| 6223_at | RPS19 | -0,01561417 | 11,9446182 | -0,27347477 | 0,78833844 | 0,89934346 |
| 387895_at | LINC00944 | 0,03592468 | 4,80271399 | 0,27344458 | 0,78836119 | 0,89934346 |
| 126133_at | ALDH16A1 | 0,02086734 | 8,02309439 | 0,27337253 | 0,78841549 | 0,89934346 |
| 374877_at | C19orf45 | -0,02101702 | 5,10841337 | -0,27332815 | 0,78844894 | 0,89934346 |
| 100128590_a | SLC8A1-AS1 | -0,02001202 | 3,2221361 | -0,27331297 | 0,78846038 | 0,89934346 |
| 57692_at | MAGEE1 | 0,03566794 | 5,90778448 | 0,2731881 | 0,78855449 | 0,89939755 |
| 11037_at | STON1 | 0,03661347 | 5,18727135 | 0,27258503 | 0,78900906 | 0,89986274 |
| 55553_at | SOX6 | -0,02963945 | 4,46727327 | -0,27235744 | 0,78918063 | 0,90000512 |
| 8382_at | NME5 | 0,02548787 | 3,40417409 | 0,27198457 | 0,78946174 | 0,90027241 |
| 100240728_a | LOC10024072 | -0,02059905 | 4,26350972 | -0,2717647 | 0,78962752 | 0,90040815 |
| 1435_at | CSF1 | -0,0245818 | 6,35993431 | -0,27115569 | 0,79008675 | 0,90087849 |

| | | | | | | |
|---|---|---|---|---|---|---|
| 64410_at | KLHL25 | -0,01993668 | 6,56386491 | -0,27080237 | 0,79035322 | 0,90108849 |
| 101927704_a | MIR3663HG | 0,02160337 | 3,20410701 | 0,27078743 | 0,79036448 | 0,90108849 |
| 1506_at | CTRL | 0,03639312 | 5,32206998 | 0,27052491 | 0,79056249 | 0,90121459 |
| 6558_at | SLC12A2 | -0,06887676 | 8,99213902 | -0,27051673 | 0,79056866 | 0,90121459 |
| 260434_at | PYDC1 | -0,03051453 | 3,85004923 | -0,27042391 | 0,79063867 | 0,90124106 |
| 1655_at | DDX5 | 0,02887262 | 10,0923974 | 0,2701653 | 0,79083375 | 0,90133612 |
| 84619_at | ZGPAT | -0,02293506 | 7,94335771 | -0,27013341 | 0,79085781 | 0,90133612 |
| 55192_at | DNAJC17 | -0,02228367 | 6,94662503 | -0,27012728 | 0,79086243 | 0,90133612 |
| 1379_at | CR1L | 0,02912483 | 3,42667148 | 0,27001074 | 0,79095035 | 0,90138298 |
| 284648_at | LOC284648 | -0,02369932 | 4,68839523 | -0,269923 | 0,79101653 | 0,90139855 |
| 55001_at | TTC22 | 0,05438614 | 6,67634771 | 0,26986857 | 0,79105759 | 0,90139855 |
| 148223_at | C19orf25 | 0,02099679 | 7,60257487 | 0,26963738 | 0,79123202 | 0,90154397 |
| 1471_at | CST3 | 0,05673263 | 10,1937343 | 0,26917271 | 0,79158262 | 0,90185725 |
| 492307_at | C8orf22 | -0,02388139 | 3,31777208 | -0,26912373 | 0,79161958 | 0,90185725 |
| 472_at | ATM | -0,07180494 | 5,87146507 | -0,26902186 | 0,79169644 | 0,90185725 |
| 388650_at | FAM69A | 0,02935837 | 8,27578142 | 0,26895119 | 0,79174977 | 0,90185725 |
| 55502_at | HES6 | -0,06299959 | 6,72171161 | -0,26890611 | 0,79178379 | 0,90185725 |
| 528_at | ATP6V1C1 | 0,03872451 | 8,02730694 | 0,2689007 | 0,79178787 | 0,90185725 |
| 55829_at | VIMP | -0,04259334 | 9,32402445 | -0,26853407 | 0,79206456 | 0,90211905 |
| 79741_at | CCDC7 | 0,01707567 | 3,11194242 | 0,2683399 | 0,7922111 | 0,90223261 |
| 3281_at | HSBP1 | 0,01757431 | 9,70294954 | 0,26818721 | 0,79232635 | 0,90231052 |
| 5729_at | PTGDR | -0,01989667 | 3,41360946 | -0,26798392 | 0,79247978 | 0,9024259 |
| 23242_at | COBL | -0,05175597 | 4,62634628 | -0,26782562 | 0,79259928 | 0,9024259 |
| 10687_at | PNMA2 | 0,02255505 | 3,68159543 | 0,26776925 | 0,79264183 | 0,9024259 |
| 84944_at | MAEL | 0,0244365 | 3,18828194 | 0,26774135 | 0,79266289 | 0,9024259 |
| 339745_at | SPOPL | 0,05266056 | 7,26986344 | 0,26773024 | 0,79267127 | 0,9024259 |
| 92691_at | TMEM169 | -0,0258581 | 4,24884425 | -0,2676806 | 0,79270875 | 0,9024259 |
| 101927606_a | LOC10192760 | -0,02946825 | 4,33727661 | -0,26752095 | 0,79282927 | 0,90250976 |
| 23284_at | ADGRL3 | 0,01398253 | 3,21344687 | 0,26698419 | 0,79323451 | 0,90287846 |
| 348378_at | FAM159A | 0,03725992 | 4,9227689 | 0,26696778 | 0,7932469 | 0,90287846 |
| 100861554_a | LINC00563 | -0,03271055 | 4,43971437 | -0,26664562 | 0,79349016 | 0,90308377 |
| 140699_at | MROH8 | 0,02265882 | 4,26572321 | 0,26659965 | 0,79352487 | 0,90308377 |
| 6536_at | SLC6A9 | 0,05076999 | 6,27148524 | 0,26654263 | 0,79356793 | 0,90308377 |
| 60509_at | AGBL5 | 0,03010683 | 7,60369654 | 0,26647686 | 0,79361759 | 0,90308694 |
| 3229_at | HOXC13 | 0,04147714 | 6,63684851 | 0,26608102 | 0,79391652 | 0,90330734 |
| 54875_at | CNTLN | 0,07758331 | 4,36533551 | 0,26596519 | 0,79400401 | 0,90330734 |
| 22794_at | CASC3 | 0,02002693 | 8,84846869 | 0,26590312 | 0,79405089 | 0,90330734 |
| 202051_at | SPATA24 | -0,04014204 | 7,39937408 | -0,26579843 | 0,79412995 | 0,90330734 |
| 85413_at | SLC22A16 | -0,01965531 | 3,78808007 | -0,26578512 | 0,79414 | 0,90330734 |
| 2974_at | GUCY1B2 | -0,03986252 | 4,10591695 | -0,26570404 | 0,79420125 | 0,90330734 |
| 101927406_a | LOC10192740 | -0,02693241 | 3,71341721 | -0,26568986 | 0,79421195 | 0,90330734 |
| 79778_at | MICALL2 | -0,02629594 | 7,0491453 | -0,26567874 | 0,79422036 | 0,90330734 |
| 5568_at | PRKACG | -0,02596044 | 5,33611123 | -0,26558462 | 0,79429145 | 0,90330734 |
| 79912_at | PYROXD1 | 0,06472886 | 7,10827982 | 0,26557 | 0,79430249 | 0,90330734 |
| 284656_at | EPHA10 | -0,02464551 | 4,90393649 | -0,26553741 | 0,79432711 | 0,90330734 |
| 945_at | CD33 | -0,0281264 | 5,03748952 | -0,26537244 | 0,79445172 | 0,90335777 |
| 4099_at | MAG | -0,03336602 | 4,1858985 | -0,26535454 | 0,79446524 | 0,90335777 |
| 114876_at | OSBPL1A | 0,08913764 | 8,12072214 | 0,26516919 | 0,79460525 | 0,90346364 |
| 28444_at | IGHV3-21 | 0,02115106 | 3,45167706 | 0,26478093 | 0,79489858 | 0,90370513 |
| 93463_at | LOC93463 | 0,01899869 | 3,04289912 | 0,26476386 | 0,79491147 | 0,90370513 |

| | | | | | | |
|---|---|---|---|---|---|---|
| 124935_at | SLC43A2 | -0,02558899 | 6,27421037 | -0,26457913 | 0,79505104 | 0,90381045 |
| 101928283_a | LOC10192828 | -0,02510042 | 3,00711412 | -0,26446299 | 0,79513879 | 0,90384952 |
| 84295_at | PHF6 | -0,026019 | 7,24254006 | -0,26440945 | 0,79517924 | 0,90384952 |
| 27063_at | ANKRD1 | 0,07055142 | 4,21198822 | 0,26432358 | 0,79524413 | 0,90386993 |
| 56666_at | PANX2 | 0,02230355 | 6,94461748 | 0,26413047 | 0,79539004 | 0,90396006 |
| 257_at | ALX3 | -0,03372197 | 5,24099813 | -0,26409443 | 0,79541728 | 0,90396006 |
| 84922_at | FIZ1 | 0,01838592 | 6,57123337 | 0,26383776 | 0,79561124 | 0,90409109 |
| 56172_at | ANKH | 0,08568422 | 6,09103756 | 0,26380328 | 0,79563729 | 0,90409109 |
| 85009_at | MGC16025 | -0,01977645 | 3,54982966 | -0,26375554 | 0,79567338 | 0,90409109 |
| 116154_at | PHACTR3 | -0,04753411 | 4,6144 | -0,2636361 | 0,79576364 | 0,90414032 |
| 387316_at | VN1R10P | -0,03568066 | 3,62400857 | -0,26300796 | 0,7962384 | 0,90456514 |
| 54700_at | RRN3 | -0,03547459 | 8,94921606 | -0,26298073 | 0,79625898 | 0,90456514 |
| 728485_at | LOC728485 | -0,05877874 | 5,16788652 | -0,26295502 | 0,79627841 | 0,90456514 |
| 27255_at | CNTN6 | -0,01588563 | 3,34268023 | -0,26281226 | 0,79638632 | 0,90463438 |
| 3421_at | IDH3G | 0,01565218 | 9,68379406 | 0,26271406 | 0,79646056 | 0,90466024 |
| 10344_at | CCL26 | -0,02398627 | 3,48831196 | -0,26261967 | 0,79653191 | 0,90466024 |
| 84179_at | MFSD7 | -0,0322611 | 6,33318761 | -0,26259577 | 0,79654998 | 0,90466024 |
| 5996_at | RGS1 | 0,01739389 | 2,85450735 | 0,2625299 | 0,79659978 | 0,90466345 |
| 9592_at | IER2 | -0,02230839 | 9,37396439 | -0,26239988 | 0,79669808 | 0,90467589 |
| 3219_at | HOXB9 | 0,2012916 | 6,86919052 | 0,26239117 | 0,79670466 | 0,90467589 |
| 55780_at | ERMARD | 0,05562458 | 7,33537446 | 0,2622549 | 0,79680768 | 0,9046864 |
| 79096_at | C11orf49 | 0,05911601 | 8,08425439 | 0,26225469 | 0,79680784 | 0,9046864 |
| 246126_at | TXLNGY | 0,11826795 | 4,57447769 | 0,26200511 | 0,79699654 | 0,90484731 |
| 6690_at | SPINK1 | 0,01768994 | 3,27074991 | 0,26192697 | 0,79705562 | 0,90485879 |
| 9095_at | TBX19 | 0,02422357 | 5,38596646 | 0,26186749 | 0,7971006 | 0,90485879 |
| 23164_at | MPRIP | -0,09246866 | 7,45650386 | -0,26164394 | 0,79726963 | 0,9049454 |
| 79684_at | MSANTD2 | -0,03010458 | 6,47645606 | -0,26164233 | 0,79727085 | 0,9049454 |
| 51592_at | TRIM33 | 0,01941687 | 9,04946268 | 0,26149898 | 0,79737925 | 0,90498386 |
| 2121_at | EVC | -0,0172954 | 5,63534364 | -0,26143876 | 0,79742479 | 0,90498386 |
| 2977_at | GUCY1A2 | -0,01944011 | 3,54442182 | -0,26141114 | 0,79744568 | 0,90498386 |
| 80336_at | PABPC1L | -0,07835197 | 6,9450359 | -0,2612899 | 0,79753736 | 0,90498408 |
| 1758_at | DMP1 | 0,0132421 | 2,96357161 | 0,26127614 | 0,79754777 | 0,90498408 |
| 55848_at | PLGRKT | 0,02264914 | 7,55681025 | 0,26122452 | 0,79758681 | 0,90498408 |
| 1235_at | CCR6 | 0,01822654 | 3,1629466 | 0,2611144 | 0,79767008 | 0,90502526 |
| 728084_at | LOC728084 | -0,01596173 | 3,63827697 | -0,26097647 | 0,7977744 | 0,90505785 |
| 401282_at | DKFZp451B08 | -0,02525163 | 4,12413233 | -0,26081384 | 0,7978974 | 0,90505785 |
| 7336_at | UBE2V2 | -0,02262962 | 7,31240391 | -0,26075173 | 0,79794438 | 0,90505785 |
| 3074_at | HEXB | -0,06987827 | 10,0517478 | -0,26072585 | 0,79796395 | 0,90505785 |
| 283050_at | ZMIZ1-AS1 | -0,02384957 | 4,67118067 | -0,26065061 | 0,79802087 | 0,90505785 |
| 102724323_a | LOC10272432 | -0,02687069 | 3,89481276 | -0,26064873 | 0,79802229 | 0,90505785 |
| 339318_at | ZNF181 | 0,02116851 | 4,17934579 | 0,26063418 | 0,79803329 | 0,90505785 |
| 5146_at | PDE6C | 0,01722741 | 2,64031659 | 0,26057945 | 0,79807468 | 0,90505785 |
| 50512_at | PODXL2 | -0,0323169 | 5,98940728 | -0,26042625 | 0,79819057 | 0,90513598 |
| 2046_at | EPHA8 | -0,0341196 | 5,95954267 | -0,26027634 | 0,79830397 | 0,90521129 |
| 8687_at | KRT38 | -0,0300141 | 4,17658075 | -0,25993143 | 0,79856489 | 0,905405 |
| 23549_at | DNPEP | -0,02299705 | 9,55165558 | -0,25992626 | 0,7985688 | 0,905405 |
| 26056_at | RAB11FIP5 | 0,05054926 | 7,06280283 | 0,25974045 | 0,79870938 | 0,90551108 |
| 101927272_a | LOC10192727 | 0,0176412 | 2,88602607 | 0,25963093 | 0,79879224 | 0,90555172 |
| 735_at | C9 | 0,02598603 | 3,06196064 | 0,25940142 | 0,79896589 | 0,9056484 |
| 3956_at | LGALS1 | -0,10361667 | 9,04975276 | -0,25930916 | 0,79903569 | 0,9056484 |

| | | | | | | |
|---|---|---|---|---|---|---|
| 25976_at | TIPARP | 0,03113254 | 9,03973839 | 0,25927877 | 0,79905869 | 0,9056484 |
| 441601_at | LOC441601 | -0,03332001 | 4,68640872 | -0,25926967 | 0,79906558 | 0,9056484 |
| 3888_at | KRT82 | -0,03207586 | 4,94969095 | -0,25909035 | 0,79920126 | 0,90574889 |
| 83417_at | FCRL4 | -0,01807367 | 3,37090935 | -0,25876556 | 0,79944705 | 0,90596753 |
| 63899_at | NSUN3 | -0,07324545 | 7,1429726 | -0,25867702 | 0,79951405 | 0,90596753 |
| 283777_at | FAM169B | -0,02630347 | 4,1852001 | -0,25863813 | 0,79954349 | 0,90596753 |
| 83551_at | TAAR8 | 0,01655213 | 4,09812434 | 0,25851184 | 0,79963907 | 0,90596753 |
| 114984_at | FLYWCH2 | -0,05502832 | 7,82651887 | -0,25850041 | 0,79964771 | 0,90596753 |
| 83888_at | FGFBP2 | -0,02085347 | 3,10189284 | -0,25837514 | 0,79974252 | 0,90596753 |
| 7373_at | COL14A1 | -0,01852519 | 3,40168157 | -0,25836206 | 0,79975242 | 0,90596753 |
| 80776_at | B9D2 | 0,02661368 | 6,44076871 | 0,25824184 | 0,79984341 | 0,90596753 |
| 652_at | BMP4 | -0,04535511 | 6,30110294 | -0,25823354 | 0,7998497 | 0,90596753 |
| 10242_at | KCNMB2 | 0,02139954 | 3,26524592 | 0,25821399 | 0,7998645 | 0,90596753 |
| 9846_at | GAB2 | -0,03014441 | 5,46174309 | -0,25801195 | 0,80001742 | 0,90599255 |
| 51124_at | IER3IP1 | 0,02608552 | 9,67616253 | 0,25800924 | 0,80001947 | 0,90599255 |
| 23026_at | MYO16 | 0,02230189 | 4,05450131 | 0,25794233 | 0,80007012 | 0,90599255 |
| 287_at | ANK2 | 0,01955638 | 3,43928856 | 0,2578462 | 0,80014289 | 0,90599255 |
| 8220_at | DGCR14 | -0,0160314 | 6,87113914 | -0,25781315 | 0,8001679 | 0,90599255 |
| 100507098_a | ADAMTS9-AS2 | -0,026918 | 3,4626983 | -0,25781198 | 0,80016878 | 0,90599255 |
| 2165_at | F13B | 0,02131336 | 3,01043088 | 0,25761547 | 0,80031754 | 0,90603956 |
| 2529_at | FUT7 | -0,01957284 | 4,84720872 | -0,25758434 | 0,80034111 | 0,90603956 |
| 23528_at | ZNF281 | 0,05845055 | 7,94633456 | 0,25757074 | 0,80035141 | 0,90603956 |
| 22913_at | RALY | 0,03551764 | 9,49597904 | 0,25740865 | 0,80047411 | 0,90612522 |
| 200772_at | LOC200772 | 0,0242758 | 5,06464002 | 0,25668081 | 0,80102518 | 0,90669574 |
| 55713_at | ZNF334 | 0,0124257 | 3,23880323 | 0,25621076 | 0,80138113 | 0,90689769 |
| 101927659_a | SMAD1-AS2 | -0,02505994 | 3,04084454 | -0,25620781 | 0,80138336 | 0,90689769 |
| 55800_at | SCN3B | 0,02598282 | 3,98862843 | 0,25620294 | 0,80138705 | 0,90689769 |
| 6158_at | RPL28 | 0,0293646 | 8,68083484 | 0,25615361 | 0,80142441 | 0,90689769 |
| 7638_at | ZNF221 | -0,02098809 | 4,60602113 | -0,25613435 | 0,801439 | 0,90689769 |
| 64321_at | SOX17 | -0,02187708 | 4,40689104 | -0,25579536 | 0,80169573 | 0,90703277 |
| 2788_at | GNG7 | -0,0237311 | 4,8790121 | -0,255785 | 0,80170358 | 0,90703277 |
| 11178_at | LZTS1 | -0,01921174 | 4,92159943 | -0,25575789 | 0,80172411 | 0,90703277 |
| 2138_at | EYA1 | -0,02494909 | 5,7571474 | -0,25572805 | 0,80174672 | 0,90703277 |
| 8795_at | TNFRSF10B | 0,08197266 | 8,46499602 | 0,25550713 | 0,80191405 | 0,9071688 |
| 799_at | CALCR | 0,02235259 | 3,97137247 | 0,25541341 | 0,80198503 | 0,90719582 |
| 440982_at | LOC440982 | 0,01605009 | 3,07420467 | 0,25534609 | 0,80203603 | 0,90719846 |
| 5108_at | PCM1 | 0,05199523 | 7,32986299 | 0,25523872 | 0,80211736 | 0,90719846 |
| 29926_at | GMPPA | -0,03002934 | 7,80222035 | -0,25522382 | 0,80212865 | 0,90719846 |
| 57733_at | GBA3 | 0,01551974 | 3,71072108 | 0,25505018 | 0,80226018 | 0,90729395 |
| 100506068_a | DACT3-AS1 | -0,02016245 | 3,47121705 | -0,25495009 | 0,802336 | 0,90731134 |
| 126364_at | LRRC25 | -0,03052049 | 4,78509075 | -0,25490151 | 0,80237281 | 0,90731134 |
| 10712_at | FAM189B | 0,01966015 | 8,66348908 | 0,25479899 | 0,80245048 | 0,90731134 |
| 6608_at | SMO | -0,02455643 | 6,36612351 | -0,25473106 | 0,80250194 | 0,90731134 |
| 374443_at | LOC374443 | -0,03377799 | 6,76245409 | -0,25471902 | 0,80251106 | 0,90731134 |
| 101927587_a | LOC10192758 | 0,01587985 | 3,32047471 | 0,2545343 | 0,80265101 | 0,90739369 |
| 730755_at | KRTAP2-3 | -0,03085468 | 4,12602397 | -0,25449852 | 0,80267812 | 0,90739369 |
| 11060_at | WWP2 | 0,02612227 | 6,65329816 | 0,25443266 | 0,80272802 | 0,90739686 |
| 84062_at | DTNBP1 | 0,02483688 | 7,01158832 | 0,25433079 | 0,8028052 | 0,90741084 |
| 64922_at | LRRC19 | -0,01576606 | 2,85359824 | -0,25424726 | 0,8028685 | 0,90741084 |
| 63924_at | CIDEC | -0,03824559 | 5,55988975 | -0,25422982 | 0,80288171 | 0,90741084 |

| | | | | | | |
|---|---|---|---|---|---|---|
| 8817_at | FGF18 | -0,01939205 | 5,14657338 | -0,25391956 | 0,8031168 | 0,90760386 |
| 64645_at | HIAT1 | 0,04639674 | 9,95308652 | 0,25388007 | 0,80314673 | 0,90760386 |
| 3991_at | LIPE | -0,02122886 | 5,60620463 | -0,25375163 | 0,80324406 | 0,9076606 |
| 81696_at | OR5V1 | -0,01734331 | 3,54706578 | -0,25347186 | 0,80345608 | 0,90784693 |
| 64805_at | P2RY12 | -0,01527087 | 3,09959652 | -0,25325221 | 0,80362255 | 0,90798177 |
| 167681_at | PRSS35 | -0,01610121 | 3,37627081 | -0,25314848 | 0,80370117 | 0,90801734 |
| 113246_at | C12orf57 | -0,04617592 | 9,5054909 | -0,2530168 | 0,80380097 | 0,90807684 |
| 11151_at | CORO1A | -0,14261733 | 6,75983115 | -0,25285336 | 0,80392486 | 0,90816353 |
| 7538_at | ZFP36 | 0,01820966 | 8,46997844 | 0,25278272 | 0,80397841 | 0,90817077 |
| 4591_at | TRIM37 | -0,02635727 | 7,40900746 | -0,25251405 | 0,80418207 | 0,90834755 |
| 5529_at | PPP2R5E | -0,02065307 | 8,99897308 | -0,25236911 | 0,80429194 | 0,90838046 |
| 150275_at | CCDC117 | -0,01396242 | 7,30348128 | -0,25231932 | 0,80432968 | 0,90838046 |
| 284217_at | LAMA1 | -0,02392708 | 4,29206561 | -0,25222573 | 0,80440063 | 0,90838046 |
| 58511_at | DNASE2B | -0,02316918 | 3,06158108 | -0,25217314 | 0,80444051 | 0,90838046 |
| 8909_at | ENDOU | 0,0208766 | 3,73736207 | 0,2521646 | 0,80444698 | 0,90838046 |
| 54963_at | UCKL1 | 0,02055573 | 8,0789637 | 0,25193942 | 0,80461771 | 0,90847746 |
| 101929488_a | LOC10192948 | -0,02281614 | 4,75843492 | -0,25192688 | 0,80462721 | 0,90847746 |
| 731424_at | LOC731424 | -0,02950328 | 3,74851148 | -0,25186274 | 0,80467584 | 0,90847912 |
| 5520_at | PPP2R2A | 0,0454495 | 8,53661231 | 0,25174074 | 0,80476834 | 0,90848308 |
| 8398_at | PLA2G6 | -0,03471528 | 6,62377321 | -0,2517337 | 0,80477368 | 0,90848308 |
| 4828_at | NMB | 0,08880632 | 9,1404346 | 0,251635 | 0,80484852 | 0,90849001 |
| 349667_at | RTN4RL2 | 0,0355462 | 5,4890462 | 0,25149018 | 0,80495833 | 0,90849001 |
| 9420_at | CYP7B1 | 0,019395 | 3,47517565 | 0,2514748 | 0,80496999 | 0,90849001 |
| 105374693_a | LOC10537469 | -0,03644448 | 3,65533846 | -0,25139864 | 0,80502774 | 0,90849001 |
| 91782_at | CHMP7 | -0,07801689 | 7,87583403 | -0,25138913 | 0,80503495 | 0,90849001 |
| 83447_at | SLC25A31 | 0,01388347 | 2,8929079 | 0,25135243 | 0,80506278 | 0,90849001 |
| 100505967_a | LINC00645 | -0,01236238 | 2,86119573 | -0,25128981 | 0,80511027 | 0,90849037 |
| 100130987_a | LOC10013098 | -0,04963367 | 6,41765551 | -0,25112767 | 0,80523322 | 0,90855378 |
| 835_at | CASP2 | 0,0544872 | 8,67956325 | 0,25109131 | 0,80526079 | 0,90855378 |
| 126695_at | KDF1 | 0,02424474 | 7,40884038 | 0,25084809 | 0,80544524 | 0,90870866 |
| 9703_at | KIAA0100 | 0,02904238 | 8,13290637 | 0,25076521 | 0,80550811 | 0,90872636 |
| 84063_at | KIRREL2 | -0,03072731 | 4,50309772 | -0,250405 | 0,80578131 | 0,90898134 |
| 57835_at | SLC4A5 | -0,01981317 | 4,34056713 | -0,25020069 | 0,80593628 | 0,9090736 |
| 100128292_a | DLG5-AS1 | -0,02049188 | 4,39485626 | -0,25014498 | 0,80597853 | 0,9090736 |
| 5988_at | RFPL1 | 0,01393266 | 2,93717297 | 0,25011052 | 0,80600467 | 0,9090736 |
| 2042_at | EPHA3 | -0,0161102 | 3,13890284 | -0,24993528 | 0,80613761 | 0,90914453 |
| 10861_at | SLC26A1 | -0,01902308 | 4,91389484 | -0,24989793 | 0,80616594 | 0,90914453 |
| 56160_at | NDNL2 | 0,03351141 | 7,17237377 | 0,24984097 | 0,80620915 | 0,90914453 |
| 116224_at | FAM122A | 0,0491581 | 7,22787103 | 0,24973338 | 0,80629077 | 0,90915742 |
| 5617_at | PRL | -0,02792308 | 3,83924532 | -0,24970147 | 0,80631498 | 0,90915742 |
| 100131755_a | ARMCX4 | -0,01348672 | 3,6316755 | -0,24959466 | 0,80639601 | 0,90919557 |
| 255967_at | PAN3 | 0,03299814 | 8,9133619 | 0,24931931 | 0,80660491 | 0,90937787 |
| 55054_at | ATG16L1 | -0,01808852 | 6,66152354 | -0,24923153 | 0,80667151 | 0,90939974 |
| 152405_at | C3orf30 | -0,02211587 | 3,14340311 | -0,24884368 | 0,8069658 | 0,90965946 |
| 2868_at | GRK4 | -0,01646439 | 3,66722243 | -0,24873485 | 0,80704838 | 0,90965946 |
| 9287_at | TAAR2 | 0,02871724 | 3,74928083 | 0,2486501 | 0,80711269 | 0,90965946 |
| 57481_at | KIAA1210 | 0,02009025 | 3,32387323 | 0,24862122 | 0,80713461 | 0,90965946 |
| 79887_at | PLBD1 | 0,14525747 | 6,51419963 | 0,24861673 | 0,80713801 | 0,90965946 |
| 29911_at | HOOK2 | 0,05598062 | 8,36905065 | 0,24810939 | 0,80752303 | 0,91004015 |
| 100505835_a | LINC01532 | -0,02091456 | 3,8952925 | -0,24767049 | 0,80785616 | 0,9103623 |

| | | | | | | |
|---|---|---|---|---|---|---|
| 6400_at | SEL1L | 0,02038139 | 8,05834556 | 0,24741625 | 0,80804914 | 0,91048406 |
| 158431_at | ZNF782 | 0,02508253 | 4,75986057 | 0,24740361 | 0,80805874 | 0,91048406 |
| 29922_at | NME7 | 0,03031602 | 7,20439454 | 0,24715979 | 0,80824382 | 0,91063934 |
| 57119_at | EPPIN | -0,02578388 | 3,17416407 | -0,24679705 | 0,80851921 | 0,91085804 |
| 200010_at | SLC5A9 | 0,02372167 | 4,71827017 | 0,24675331 | 0,80855241 | 0,91085804 |
| 55596_at | ZCCHC8 | -0,02278568 | 6,74986419 | -0,24671726 | 0,80857979 | 0,91085804 |
| 158787_at | RIBC1 | -0,02057278 | 4,97265158 | -0,24654893 | 0,80870759 | 0,91094873 |
| 79153_at | GDPD3 | 0,07920374 | 6,75783804 | 0,2463401 | 0,80886615 | 0,91101599 |
| 56917_at | MEIS3 | 0,03303669 | 5,53425094 | 0,2462489 | 0,8089354 | 0,91101599 |
| 79087_at | ALG12 | -0,02301435 | 7,60223998 | -0,24621375 | 0,80896209 | 0,91101599 |
| 83871_at | RAB34 | 0,29700694 | 8,58122855 | 0,2461431 | 0,80901574 | 0,91101599 |
| 80331_at | DNAJC5 | 0,01436574 | 8,3105619 | 0,24609497 | 0,80905228 | 0,91101599 |
| 50619_at | DEF6 | -0,03175548 | 6,86520188 | -0,24603703 | 0,80909629 | 0,91101599 |
| 102467079_a | LOC10246707 | -0,03006952 | 3,35124043 | -0,24603431 | 0,80909835 | 0,91101599 |
| 4204_at | MECP2 | 0,03068703 | 7,38233391 | 0,245797 | 0,80927856 | 0,91104094 |
| 127933_at | UHMK1 | -0,06367872 | 8,77729928 | -0,24577455 | 0,80929561 | 0,91104094 |
| 84814_at | PPAPDC3 | -0,02458729 | 5,40869714 | -0,24576224 | 0,80930496 | 0,91104094 |
| 149830_at | PRNT | 0,02341789 | 3,87843612 | 0,2456457 | 0,80939346 | 0,91104094 |
| 84789_at | MGC2889 | -0,01712715 | 3,48497047 | -0,24562342 | 0,80941038 | 0,91104094 |
| 79838_at | TMC5 | -0,01438664 | 3,76269982 | -0,24547373 | 0,80952407 | 0,91104094 |
| 79884_at | MAP9 | 0,11294757 | 4,79793407 | 0,24546225 | 0,80953279 | 0,91104094 |
| 6036_at | RNASE2 | -0,0229923 | 3,07847262 | -0,24541481 | 0,80956882 | 0,91104094 |
| 5968_at | REG1B | -0,02156297 | 3,75728044 | -0,24531081 | 0,8096478 | 0,91104094 |
| 729420_at | LMO7DN | -0,01789906 | 4,16612838 | -0,24530976 | 0,8096486 | 0,91104094 |
| 54854_at | FAM83E | -0,02455024 | 5,18477637 | -0,24529164 | 0,80966236 | 0,91104094 |
| 7625_at | ZNF74 | -0,01851684 | 6,52938698 | -0,24525782 | 0,80968805 | 0,91104094 |
| 28958_at | COA3 | 0,02480112 | 9,70860819 | 0,24516057 | 0,80976191 | 0,91107083 |
| 4697_at | NDUFA4 | -0,02971797 | 8,84218845 | -0,24499701 | 0,80988615 | 0,91109353 |
| 59350_at | RXFP1 | 0,01182026 | 2,98655442 | 0,24489343 | 0,80996482 | 0,91109353 |
| 56937_at | PMEPA1 | 0,067491 | 9,1911075 | 0,24486669 | 0,80998513 | 0,91109353 |
| 26057_at | ANKRD17 | 0,06617759 | 8,50459035 | 0,24484643 | 0,81000053 | 0,91109353 |
| 10716_at | TBR1 | -0,02913976 | 3,81884797 | -0,24482267 | 0,81001857 | 0,91109353 |
| 92797_at | HELB | 0,01656149 | 4,71411409 | 0,24471741 | 0,81009853 | 0,91113026 |
| 57142_at | RTN4 | 0,02208312 | 10,7879658 | 0,244562 | 0,81021658 | 0,91116269 |
| 1716_at | DGUOK | -0,0239693 | 8,92542674 | -0,24447137 | 0,81028544 | 0,91116269 |
| 63826_at | SRR | 0,07573582 | 6,25428625 | 0,2444469 | 0,81030402 | 0,91116269 |
| 9586_at | CREB5 | -0,0189754 | 3,9783819 | -0,24435134 | 0,81037662 | 0,91116269 |
| 131544_at | CRYBG3 | 0,04249213 | 7,80809987 | 0,24433551 | 0,81038864 | 0,91116269 |
| 55567_at | DNAH3 | 0,01543034 | 4,56565424 | 0,24424422 | 0,810458 | 0,91116269 |
| 388567_at | ZNF749 | 0,01839404 | 4,28686068 | 0,2442436 | 0,81045847 | 0,91116269 |
| 126638_at | RPTN | -0,0314654 | 3,32949937 | -0,24417185 | 0,81051298 | 0,9111708 |
| 641977_at | SEPT7P2 | -0,0406276 | 4,89400828 | -0,24402881 | 0,81062165 | 0,91123978 |
| 258010_at | SVIP | -0,03868041 | 7,64078522 | -0,24378612 | 0,81080605 | 0,91137293 |
| 283600_at | SLC25A47 | 0,01907611 | 5,04750907 | 0,24363368 | 0,81092188 | 0,91137293 |
| 65977_at | PLEKHA3 | 0,0191244 | 6,93589158 | 0,24360844 | 0,81094106 | 0,91137293 |
| 11267_at | SNF8 | 0,04489222 | 9,01781968 | 0,24350759 | 0,81101769 | 0,91137293 |
| 10962_at | MLLT11 | 0,23368545 | 9,81048194 | 0,24350134 | 0,81102243 | 0,91137293 |
| #N/A | GS1-24F4,2 | 0,01744556 | 2,95300754 | 0,24349932 | 0,81102397 | 0,91137293 |
| 171017_at | ZNF384 | 0,01351096 | 8,56537114 | 0,24325998 | 0,81120584 | 0,91144538 |
| 100507629_a | LINC00658 | -0,02182045 | 4,43749736 | -0,24315997 | 0,81128184 | 0,91144538 |

| | | | | | | |
|---|---|---|---|---|---|---|
| 101927735_a | LOC10192773 | -0,01993691 | 3,65598653 | -0,2431516 | 0,8112882 | 0,91144538 |
| 2018_at | EMX2 | 0,19056759 | 5,60890523 | 0,24303689 | 0,81137538 | 0,91144538 |
| 2675_at | GFRA2 | -0,02140112 | 4,52544913 | -0,24301412 | 0,81139268 | 0,91144538 |
| 113828_at | FAM83F | 0,05652509 | 6,92144708 | 0,24299169 | 0,81140973 | 0,91144538 |
| 127579_at | DCST2 | 0,0261815 | 4,32460534 | 0,24297862 | 0,81141966 | 0,91144538 |
| 2256_at | FGF11 | 0,021083 | 6,09479702 | 0,242854 | 0,81151437 | 0,91149862 |
| 10117_at | ENAM | -0,03406198 | 4,05209717 | -0,24238452 | 0,81187119 | 0,9117862 |
| 150209_at | AIFM3 | 0,0204255 | 5,59404987 | 0,24237001 | 0,81188222 | 0,9117862 |
| 8292_at | COLQ | -0,0292187 | 4,90426362 | -0,24230965 | 0,8119281 | 0,9117862 |
| 5450_at | POU2AF1 | -0,14076467 | 5,22167776 | -0,24226748 | 0,81196016 | 0,9117862 |
| 2014_at | EMP3 | 0,02915611 | 4,04623248 | 0,24220576 | 0,81200707 | 0,9117862 |
| 197358_at | NLRC3 | -0,02983045 | 3,95472042 | -0,24173391 | 0,81236576 | 0,91208449 |
| 4060_at | LUM | -0,48502268 | 7,14675561 | -0,24172441 | 0,81237298 | 0,91208449 |
| 284274_at | SMIM21 | -0,01837036 | 3,34447824 | -0,24166943 | 0,81241477 | 0,91208449 |
| 10497_at | UNC13B | 0,03114655 | 6,8486651 | 0,24158392 | 0,81247978 | 0,91210432 |
| 148418_at | SAMD13 | 0,12579923 | 5,17446988 | 0,24146327 | 0,81257151 | 0,91214626 |
| 28962_at | OSTM1 | 0,06358227 | 6,58865934 | 0,24141022 | 0,81261184 | 0,91214626 |
| 1005_at | CDH7 | 0,02406326 | 5,12754026 | 0,24129218 | 0,81270158 | 0,91219383 |
| 11033_at | ADAP1 | -0,03061557 | 7,52644101 | -0,24116972 | 0,81279469 | 0,91224519 |
| 100129128_a | KHDC1L | -0,02668872 | 3,81392465 | -0,24082301 | 0,81305832 | 0,91244685 |
| 129787_at | TMEM18 | 0,01974294 | 8,18268288 | 0,24080882 | 0,8130691 | 0,91244685 |
| 56912_at | IFT46 | -0,02969714 | 7,87910534 | -0,24073237 | 0,81312724 | 0,91245893 |
| 55139_at | ANKZF1 | -0,05584868 | 6,96561362 | -0,24056171 | 0,81325701 | 0,91250903 |
| 767_at | CA8 | -0,01440876 | 3,09565153 | -0,24054907 | 0,81326662 | 0,91250903 |
| 5897_at | RAG2 | -0,01601162 | 2,83605332 | -0,24037617 | 0,81339811 | 0,9126034 |
| 6579_at | SLCO1A2 | 0,02100064 | 3,1623832 | 0,24007932 | 0,81362387 | 0,91280353 |
| 6299_at | SALL1 | -0,01871878 | 3,60950713 | -0,23992943 | 0,81373786 | 0,91284432 |
| 50636_at | ANO7 | 0,02479451 | 6,49990647 | 0,2399069 | 0,813755 | 0,91284432 |
| 50484_at | RRM2B | 0,06282744 | 8,52636448 | 0,23983858 | 0,81380696 | 0,91284944 |
| 100506840_a | IQCF4 | -0,01310378 | 3,15594285 | -0,23944274 | 0,81410804 | 0,91309087 |
| 6514_at | SLC2A2 | -0,02064122 | 3,44916803 | -0,23937716 | 0,81415793 | 0,91309087 |
| 193629_at | LINC00189 | -0,018832 | 3,35638364 | -0,23936866 | 0,8141644 | 0,91309087 |
| 5965_at | RECQL | -0,0211641 | 7,97259028 | -0,23926952 | 0,81423981 | 0,91309402 |
| 92822_at | ZNF276 | 0,01533181 | 6,29620427 | 0,23924034 | 0,81426201 | 0,91309402 |
| 168507_at | PKD1L1 | -0,02081421 | 3,94642127 | -0,23874598 | 0,81463809 | 0,91346257 |
| 27189_at | IL17C | -0,03378249 | 6,27509432 | -0,23853958 | 0,81479512 | 0,91358547 |
| 101927129_a | LINC01580 | -0,02614366 | 3,9978191 | -0,23842455 | 0,81488265 | 0,91363043 |
| 100130476_a | LOC10013047 | 0,01514162 | 3,63376466 | 0,23831634 | 0,81496498 | 0,91366956 |
| 9814_at | SFI1 | -0,03270005 | 6,45805243 | -0,23807059 | 0,81515197 | 0,91381022 |
| 946_at | SIGLEC6 | -0,02929842 | 5,6820283 | -0,23802676 | 0,81518532 | 0,91381022 |
| 79759_at | ZNF668 | -0,03914984 | 6,1378014 | -0,23789571 | 0,81528504 | 0,91383232 |
| 1373_at | CPS1 | 0,02166185 | 3,76741344 | 0,23787617 | 0,81529991 | 0,91383232 |
| 9673_at | SLC25A44 | 0,02879301 | 8,05916223 | 0,23774002 | 0,81540352 | 0,91389527 |
| 100852408_a | EGFLAM-AS4 | -0,02042754 | 3,3940797 | -0,23756722 | 0,81553502 | 0,91398947 |
| 101929052_a | LINC00836 | 0,01309133 | 3,46800502 | 0,23750358 | 0,81558345 | 0,91399058 |
| 100996425_a | PRCAT47 | 0,02466326 | 5,1205226 | 0,23742468 | 0,8156435 | 0,91400469 |
| 101926906_a | SLC16A12-AS1 | 0,02743613 | 3,61824911 | 0,23705718 | 0,8159232 | 0,91421344 |
| 56849_at | TCEAL7 | 0,01590682 | 3,20497262 | 0,23705521 | 0,8159247 | 0,91421344 |
| 126204_at | NLRP13 | -0,03486918 | 4,31166425 | -0,23695524 | 0,81600079 | 0,91422996 |
| 138240_at | C9orf57 | -0,02020166 | 3,22115951 | -0,23691113 | 0,81603436 | 0,91422996 |

| | | | | | | |
|---|---|---|---|---|---|---|
| 84455_at | EFCAB7 | 0,03722749 | 6,67496945 | 0,23675477 | 0,81615338 | 0,91431012 |
| 9495_at | AKAP5 | -0,01799612 | 3,5830172 | -0,23656375 | 0,81629878 | 0,91441983 |
| 8694_at | DGAT1 | 0,07489692 | 8,55988172 | 0,23621603 | 0,81656348 | 0,9146061 |
| 162417_at | NAGS | 0,0354556 | 5,9776092 | 0,2361939 | 0,81658033 | 0,9146061 |
| 7857_at | SCG2 | -0,01869833 | 3,52322803 | -0,2361582 | 0,8166075 | 0,9146061 |
| 117143_at | TADA1 | -0,02199603 | 7,95169143 | -0,2360915 | 0,81665828 | 0,91460979 |
| 8233_at | ZRSR2 | -0,03007633 | 6,81213161 | -0,23599804 | 0,81672944 | 0,91463631 |
| 5454_at | POU3F2 | 0,02105642 | 4,50978636 | 0,23568834 | 0,81696523 | 0,91482901 |
| 100288730_a | PAN3-AS1 | -0,02012156 | 5,28344666 | -0,23560408 | 0,81702938 | 0,91482901 |
| 1656_at | DDX6 | 0,02193526 | 8,21278267 | 0,23553839 | 0,8170794 | 0,91482901 |
| 56203_at | LMOD3 | -0,01635833 | 3,09623355 | -0,23547345 | 0,81712885 | 0,91482901 |
| 65267_at | WNK3 | -0,01266898 | 3,31411757 | -0,23538619 | 0,81719529 | 0,91482901 |
| 57000_at | GSN-AS1 | 0,03055869 | 4,49010331 | 0,23536614 | 0,81721056 | 0,91482901 |
| 339229_at | OXLD1 | 0,01346653 | 7,71217655 | 0,23533541 | 0,81723395 | 0,91482901 |
| 1634_at | DCN | -0,02066767 | 3,897415 | -0,23525018 | 0,81729886 | 0,9148485 |
| 645591_at | LHFPL3-AS1 | 0,01949272 | 3,40492243 | 0,23495889 | 0,81752067 | 0,91497853 |
| 169841_at | ZNF169 | -0,01218211 | 4,73275628 | -0,23495126 | 0,81752648 | 0,91497853 |
| 414319_at | LINC00587 | 0,01476681 | 3,08488727 | 0,23489349 | 0,81757047 | 0,91497853 |
| 9098_at | USP6 | 0,02273077 | 4,7203299 | 0,23484813 | 0,81760502 | 0,91497853 |
| 3356_at | HTR2A | -0,01718955 | 3,42945849 | -0,23460595 | 0,81778946 | 0,91509365 |
| 643911_at | CRNDE | -0,051115 | 7,7640008 | -0,2345883 | 0,8178029 | 0,91509365 |
| 203102_at | ADAM32 | -0,01848563 | 2,97457615 | -0,23448603 | 0,81788078 | 0,91510227 |
| 65266_at | WNK4 | 0,01606536 | 4,2110128 | 0,23445343 | 0,81790561 | 0,91510227 |
| 29122_at | PRSS50 | -0,02803976 | 5,46929161 | -0,23434635 | 0,81798717 | 0,91511802 |
| 55111_at | PLEKHJ1 | 0,03050851 | 8,55588704 | 0,23426496 | 0,81804916 | 0,91511802 |
| 26659_at | OR7A5 | 0,02914019 | 3,87143638 | 0,23421655 | 0,81808603 | 0,91511802 |
| 126123_at | IZUMO2 | 0,02828329 | 4,10147768 | 0,23417138 | 0,81812043 | 0,91511802 |
| 9382_at | COG1 | -0,03774588 | 8,02259417 | -0,23412309 | 0,81815722 | 0,91511802 |
| 304_at | ANXA2P2 | 0,02654444 | 9,35909646 | 0,233874 | 0,81834695 | 0,91520028 |
| 729683_at | LOC729683 | -0,01909734 | 5,93539688 | -0,23386064 | 0,81835713 | 0,91520028 |
| 7975_at | MAFK | -0,02218174 | 7,01554636 | -0,2337894 | 0,81841139 | 0,91520028 |
| 8576_at | STK16 | 0,02750032 | 6,59699112 | 0,23377704 | 0,8184208 | 0,91520028 |
| 150992_at | LINC00309 | 0,02687586 | 3,28110916 | 0,23352488 | 0,8186129 | 0,91522952 |
| 2802_at | GOLGA3 | -0,02226503 | 8,06429454 | -0,23352417 | 0,81861344 | 0,91522952 |
| 79955_at | PDZD7 | -0,03086874 | 5,21900838 | -0,23344036 | 0,81867729 | 0,91522952 |
| 6469_at | SHH | -0,07399803 | 6,75440192 | -0,23342563 | 0,8186885 | 0,91522952 |
| 407738_at | FAM19A1 | -0,01681401 | 3,34098046 | -0,23341645 | 0,8186955 | 0,91522952 |
| 148113_at | CILP2 | -0,0178791 | 4,93277676 | -0,23328448 | 0,81879603 | 0,91522952 |
| 140807_at | KRT72 | -0,02060306 | 4,68248639 | -0,23328007 | 0,81879939 | 0,91522952 |
| 57333_at | RCN3 | 0,03390622 | 5,64648292 | 0,23324377 | 0,81882705 | 0,91522952 |
| 1031_at | CDKN2C | -0,05221849 | 7,37998465 | -0,23293154 | 0,81906493 | 0,91540068 |
| 101927282_a | LOC10192728 | 0,01931958 | 3,51516322 | 0,23291802 | 0,81907523 | 0,91540068 |
| 1775_at | DNASE1L2 | -0,05078205 | 5,6299012 | -0,23284097 | 0,81913394 | 0,91541318 |
| 93010_at | B3GNT7 | -0,02994632 | 4,55069383 | -0,23270368 | 0,81923855 | 0,91547698 |
| 128_at | ADH5 | 0,08135882 | 9,32952419 | 0,23257586 | 0,81933594 | 0,91548946 |
| 54514_at | DDX4 | 0,01346927 | 2,98952981 | 0,23253772 | 0,819365 | 0,91548946 |
| 134549_at | SHROOM1 | -0,05451515 | 6,84614048 | -0,23246319 | 0,81942179 | 0,91548946 |
| 27283_at | TINAG | 0,02438036 | 4,09324178 | 0,23243953 | 0,81943982 | 0,91548946 |
| 728613_at | LOC728613 | 0,0408398 | 5,10881101 | 0,23214364 | 0,8196653 | 0,91568826 |
| 55040_at | EPN3 | -0,0256955 | 8,06039355 | -0,23180664 | 0,81992212 | 0,91592205 |

| | | | | | | |
|---|---|---|---|---|---|---|
| 64087_at | MCCC2 | -0,08516964 | 8,39521311 | -0,23156004 | 0,82011007 | 0,91607887 |
| 100128285_a | DNM1P35 | -0,02776328 | 5,1174484 | -0,23123942 | 0,82035444 | 0,91625982 |
| 132_at | ADK | -0,04335957 | 9,47738621 | -0,2312227 | 0,82036719 | 0,91625982 |
| 144245_at | ALG10B | -0,04006045 | 5,27915765 | -0,23101519 | 0,82052536 | 0,91636861 |
| 170961_at | ANKRD24 | -0,02090976 | 5,43413202 | -0,23097009 | 0,82055974 | 0,91636861 |
| 619279_at | ZNF704 | 0,07364744 | 6,91406266 | 0,23087366 | 0,82063325 | 0,91639758 |
| 100507291_a | LOC10050729 | 0,05966396 | 6,150405 | 0,23078166 | 0,82070337 | 0,91642276 |
| 128209_at | KLF17 | -0,03392447 | 3,97976201 | -0,23071366 | 0,82075521 | 0,91642752 |
| 3452_at | IFNA21 | 0,02213339 | 3,91852371 | 0,23047538 | 0,82093687 | 0,91649237 |
| 6783_at | SULT1E1 | -0,03094238 | 4,30598031 | -0,23045546 | 0,82095205 | 0,91649237 |
| 57608_at | KIAA1462 | 0,08455062 | 4,52549826 | 0,23045025 | 0,82095602 | 0,91649237 |
| 5078_at | PAX4 | 0,0174288 | 4,7305309 | 0,23018494 | 0,82115829 | 0,91666505 |
| 55902_at | ACSS2 | -0,02027506 | 7,89973907 | -0,22978312 | 0,82146467 | 0,9169021 |
| 26059_at | ERC2 | -0,02051911 | 3,37986981 | -0,22978158 | 0,82146584 | 0,9169021 |
| 80173_at | IFT74 | 0,04701136 | 4,10705044 | 0,22960623 | 0,82159955 | 0,91695335 |
| 10494_at | STK25 | 0,02142736 | 8,45841849 | 0,22959651 | 0,82160696 | 0,91695335 |
| 130271_at | PLEKHH2 | 0,03070858 | 3,92254068 | 0,22934936 | 0,82179543 | 0,91711056 |
| 151295_at | SLC23A3 | 0,06895211 | 5,62115618 | 0,22919118 | 0,82191605 | 0,91719204 |
| 26184_at | OR1F2P | -0,02517272 | 3,96750567 | -0,22908129 | 0,82199986 | 0,91723242 |
| 147947_at | ZNF542P | 0,0508868 | 5,98561997 | 0,22872959 | 0,82226808 | 0,91741661 |
| 9731_at | CEP104 | 0,03093217 | 7,74464562 | 0,22869607 | 0,82229365 | 0,91741661 |
| 101978719_a | LINC00970 | 0,01784642 | 4,92852679 | 0,22865217 | 0,82232713 | 0,91741661 |
| 91647_at | ATPAF2 | -0,06718082 | 6,41074567 | -0,22856864 | 0,82239084 | 0,91741661 |
| 27123_at | DKK2 | 0,02259421 | 3,40111556 | 0,22855263 | 0,82240306 | 0,91741661 |
| 253738_at | EBF3 | -0,0236114 | 3,81580855 | -0,22839092 | 0,8225264 | 0,91750107 |
| 144097_at | C11orf84 | -0,03924275 | 6,81375347 | -0,22805946 | 0,82277923 | 0,91772996 |
| 4763_at | NF1 | -0,035128 | 7,39768836 | -0,2277093 | 0,82304635 | 0,91792768 |
| 3214_at | HOXB4 | -0,03461884 | 7,36572344 | -0,22770215 | 0,82305181 | 0,91792768 |
| 5316_at | PKNOX1 | 0,01203448 | 6,41810571 | 0,22745991 | 0,82323661 | 0,91808063 |
| 11336_at | EXOC3 | 0,01808036 | 7,5923522 | 0,22730979 | 0,82335115 | 0,91814162 |
| 29992_at | PILRA | -0,01919867 | 4,74440809 | -0,22726329 | 0,82338662 | 0,91814162 |
| 51098_at | IFT52 | 0,02868742 | 8,41019981 | 0,2270502 | 0,82354921 | 0,91824238 |
| 2846_at | LPAR4 | -0,02071478 | 2,83405763 | -0,22701991 | 0,82357232 | 0,91824238 |
| 11344_at | TWF2 | 0,01664108 | 8,33854503 | 0,22678515 | 0,82375145 | 0,91835659 |
| 5532_at | PPP3CB | 0,02674162 | 7,94690988 | 0,22662583 | 0,82387302 | 0,91835659 |
| 1510_at | CTSE | -0,14644677 | 6,7101248 | -0,22661134 | 0,82388408 | 0,91835659 |
| 2346_at | FOLH1 | 0,01608106 | 3,31162734 | 0,2265828 | 0,82390585 | 0,91835659 |
| 646762_at | LOC646762 | 0,02234697 | 6,18083311 | 0,22657327 | 0,82391313 | 0,91835659 |
| 7292_at | TNFSF4 | -0,0201205 | 3,48085622 | -0,22644622 | 0,82401008 | 0,91839804 |
| 10647_at | SCGB1D2 | 0,02602962 | 3,2392008 | 0,22639958 | 0,82404567 | 0,91839804 |
| 165100_at | C2orf57 | -0,03852767 | 5,51247684 | -0,22626705 | 0,82414681 | 0,91845763 |
| 53820_at | RIPPLY3 | 0,03129469 | 5,00633916 | 0,22616176 | 0,82422716 | 0,91849403 |
| 80028_at | FBXL18 | -0,02449737 | 6,87492283 | -0,22601696 | 0,82433767 | 0,91851716 |
| 1515_at | CTSV | 0,0471119 | 7,42095774 | 0,22600961 | 0,82434328 | 0,91851716 |
| 50839_at | TAS2R10 | -0,01644079 | 3,16185833 | -0,22585176 | 0,82446376 | 0,91855376 |
| 6883_at | TAF12 | -0,04131006 | 7,94953123 | -0,22580062 | 0,82450279 | 0,91855376 |
| 90861_at | HN1L | 0,02495112 | 9,71264893 | 0,22577913 | 0,82451919 | 0,91855376 |
| 8073_at | PTP4A2 | -0,02489252 | 10,9908044 | -0,22557338 | 0,82467623 | 0,91863969 |
| 55324_at | ABCF3 | 0,01499243 | 8,0612844 | 0,22545109 | 0,82476957 | 0,91863969 |
| 7544_at | ZFY | 0,01368033 | 4,25468726 | 0,22533394 | 0,82485899 | 0,91863969 |

| | | | | | | |
|---|---|---|---|---|---|---|
| 4893_at | NRAS | 0,05553743 | 9,16620842 | 0,22533121 | 0,82486108 | 0,91863969 |
| 196410_at | METTL7B | -0,05583707 | 5,70534907 | -0,22527663 | 0,82490274 | 0,91863969 |
| 283316_at | CD163L1 | 0,10541938 | 5,16412093 | 0,22520792 | 0,82495519 | 0,91863969 |
| 64426_at | SUDS3 | 0,04182675 | 8,9239277 | 0,22518491 | 0,82497276 | 0,91863969 |
| 8346_at | HIST1H2BI | 0,03895462 | 5,27019633 | 0,22509519 | 0,82504124 | 0,91863969 |
| 1571_at | CYP2E1 | -0,01747375 | 4,14779942 | -0,22508839 | 0,82504643 | 0,91863969 |
| 147339_at | C18orf25 | -0,02042189 | 5,78996018 | -0,22505331 | 0,82507321 | 0,91863969 |
| 9039_at | UBA3 | -0,03698072 | 9,13064288 | -0,22488579 | 0,8252011 | 0,91866593 |
| 284418_at | FAM71E2 | -0,01603013 | 5,29319822 | -0,22484906 | 0,82522914 | 0,91866593 |
| 283038_at | LOC283038 | -0,01500326 | 3,58515202 | -0,22478668 | 0,82527675 | 0,91866593 |
| 7812_at | CSDE1 | -0,01779421 | 11,2426218 | -0,22471063 | 0,82533481 | 0,91866593 |
| 101929335_a | ADAMTS9-AS1 | 0,03350419 | 3,31513293 | 0,22471008 | 0,82533523 | 0,91866593 |
| 2494_at | NR5A2 | -0,01493776 | 3,91256604 | -0,22460205 | 0,82541771 | 0,91868172 |
| 442213_at | PTCHD4 | 0,03476072 | 4,23850344 | 0,22456656 | 0,8254448 | 0,91868172 |
| 7161_at | TP73 | -0,01695399 | 5,86055791 | -0,22443044 | 0,82554873 | 0,9187443 |
| 162972_at | ZNF550 | 0,02025576 | 6,52011835 | 0,22436385 | 0,82559957 | 0,9187478 |
| 646658_at | SYNDIG1L | -0,01911346 | 5,44027373 | -0,22419724 | 0,82572678 | 0,91883629 |
| 105376204_a | LOC10537620 | -0,0207542 | 4,08732583 | -0,22411562 | 0,8257891 | 0,91885256 |
| 101927380_a | C9orf41-AS1 | -0,0277719 | 3,93931749 | -0,22352917 | 0,82623691 | 0,91929773 |
| 80760_at | ITIH5 | -0,01492381 | 4,18554573 | -0,22334747 | 0,82637567 | 0,91939901 |
| 100505547_a | LINC01208 | 0,01637657 | 4,23383112 | 0,22318687 | 0,82649832 | 0,91948236 |
| 348938_at | NIPAL4 | -0,06536371 | 7,90790438 | -0,22308163 | 0,82657869 | 0,91951868 |
| 246777_at | SPESP1 | -0,0173729 | 2,84606986 | -0,22277652 | 0,82681172 | 0,91964708 |
| 79465_at | ULBP3 | -0,0477155 | 6,29266743 | -0,22276543 | 0,82682019 | 0,91964708 |
| 8618_at | CADPS | 0,0175858 | 3,55322905 | 0,22274297 | 0,82683734 | 0,91964708 |
| 11250_at | GPR45 | 0,01529957 | 5,29659742 | 0,22260991 | 0,82693898 | 0,91970702 |
| 8707_at | B3GALT2 | -0,01565203 | 3,00175919 | -0,22238646 | 0,82710966 | 0,9198118 |
| 138255_at | C9orf135 | -0,01916576 | 3,16427851 | -0,22236154 | 0,82712869 | 0,9198118 |
| 9038_at | TAAR5 | -0,0344286 | 4,45125868 | -0,22227698 | 0,82719329 | 0,91983053 |
| 93429_at | DKFZp434J02 | 0,0344712 | 5,24900446 | 0,22219281 | 0,82725758 | 0,91984893 |
| 54674_at | LRRN3 | -0,02006335 | 3,67101335 | -0,22212695 | 0,82730789 | 0,91985177 |
| 102606465_a | LOC10260646 | -0,08976626 | 5,9880307 | -0,22204118 | 0,82737341 | 0,91985358 |
| 257236_at | CCDC96 | 0,03483594 | 5,32282566 | 0,22195325 | 0,82744059 | 0,91985358 |
| 105376910_a | LOC10537691 | 0,03278598 | 3,87772601 | 0,2219373 | 0,82745277 | 0,91985358 |
| 5931_at | RBBP7 | -0,01832905 | 10,7588668 | -0,22172517 | 0,82761483 | 0,91998064 |
| 84620_at | ST6GAL2 | -0,01364582 | 3,68337347 | -0,22160395 | 0,82770744 | 0,92001583 |
| 153572_at | IRX2 | -0,02258165 | 4,7663865 | -0,2215587 | 0,82774201 | 0,92001583 |
| 6651_at | SON | 0,01581084 | 9,45761352 | 0,22142531 | 0,82784393 | 0,92007602 |
| 441150_at | C6orf226 | 0,02510159 | 6,94166857 | 0,22130489 | 0,82793593 | 0,92012519 |
| 2893_at | GRIA4 | -0,01153448 | 3,16338701 | -0,22108769 | 0,82810188 | 0,92025652 |
| 119467_at | CLRN3 | -0,02133223 | 3,28705069 | -0,22081721 | 0,82830857 | 0,92039456 |
| 10640_at | EXOC5 | 0,02645288 | 8,79005864 | 0,22074557 | 0,8283633 | 0,92039456 |
| 83850_at | ESYT3 | 0,01474808 | 3,50550303 | 0,22073754 | 0,82836944 | 0,92039456 |
| 26245_at | OR2M4 | 0,02638346 | 3,60323057 | 0,22055423 | 0,82850952 | 0,92047478 |
| 126823_at | KLHDC9 | 0,0421265 | 5,87063804 | 0,22047484 | 0,82857019 | 0,92047478 |
| 92370_at | PXYLP1 | -0,02638411 | 7,78010878 | -0,22040365 | 0,82862459 | 0,92047478 |
| 64844_at | MARCH7 | -0,02393021 | 9,25213876 | -0,22039294 | 0,82863278 | 0,92047478 |
| 9628_at | RGS6 | 0,0135 | 4,76914583 | 0,22001752 | 0,8289197 | 0,92074041 |
| 81033_at | KCNH6 | -0,02077363 | 4,39684657 | -0,21957917 | 0,82925474 | 0,92098585 |
| 23443_at | SLC35A3 | -0,04199741 | 8,02182579 | -0,21955763 | 0,8292712 | 0,92098585 |

| | | | | | | |
|---|---|---|---|---|---|---|
| 8270_at | LAGE3 | -0,02302978 | 7,87333929 | -0,21951634 | 0,82930276 | 0,92098585 |
| 8030_at | CCDC6 | -0,02696591 | 9,84343507 | -0,21947821 | 0,82933191 | 0,92098585 |
| 641339_at | ZNF674 | 0,01413153 | 3,06925194 | 0,21930975 | 0,82946068 | 0,92106906 |
| 25978_at | CHMP2B | -0,02877444 | 9,49027219 | -0,21925508 | 0,82950247 | 0,92106906 |
| 345757_at | FAM174A | 0,04695545 | 8,22127159 | 0,21908959 | 0,82962898 | 0,92115644 |
| 220074_at | LRTOMT | 0,02986612 | 6,25961849 | 0,21893186 | 0,82974956 | 0,92123722 |
| 10391_at | CORO2B | -0,02880197 | 4,2380636 | -0,2187359 | 0,82989937 | 0,92124471 |
| 100506409_a | ELOVL2-AS1 | 0,01478615 | 3,64724979 | 0,21869268 | 0,82993241 | 0,92124471 |
| 83463_at | MXD3 | -0,02919026 | 7,44816288 | -0,21868785 | 0,8299361 | 0,92124471 |
| 360_at | AQP3 | -0,17679521 | 10,9928708 | -0,2185869 | 0,83001328 | 0,92124471 |
| 5208_at | PFKFB2 | -0,02000932 | 5,9533536 | -0,21855562 | 0,8300372 | 0,92124471 |
| 1080_at | CFTR | 0,01467373 | 3,2842564 | 0,21849762 | 0,83008154 | 0,92124471 |
| 51205_at | ACP6 | -0,01635721 | 7,0917234 | -0,2184268 | 0,83013568 | 0,92124471 |
| 56853_at | CELF4 | -0,01844272 | 4,4251563 | -0,21842258 | 0,83013891 | 0,92124471 |
| 55846_at | ITFG2 | -0,01714964 | 7,25069901 | -0,21831662 | 0,83021992 | 0,9212523 |
| 84271_at | POLDIP3 | 0,03169273 | 8,79930782 | 0,21821575 | 0,83029705 | 0,9212523 |
| 23099_at | ZBTB43 | 0,01880653 | 6,62852482 | 0,21818598 | 0,83031981 | 0,9212523 |
| 100507140_a | LOC10050714 | -0,02096721 | 3,27086252 | -0,21816345 | 0,83033704 | 0,9212523 |
| 57463_at | AMIGO1 | -0,02332499 | 5,97974476 | -0,21791245 | 0,83052896 | 0,92141216 |
| 154872_at | C7orf77 | -0,02169382 | 3,47861755 | -0,2177137 | 0,83068094 | 0,92147988 |
| 3929_at | LBP | 0,02306489 | 4,46243544 | 0,21770751 | 0,83068567 | 0,92147988 |
| 100505536_a | ISM1-AS1 | -0,0174394 | 3,52772248 | -0,21753196 | 0,83081992 | 0,92151175 |
| 55096_at | EBLN2 | 0,02845159 | 5,28444306 | 0,21750119 | 0,83084345 | 0,92151175 |
| 101928995_a | LOC10192899 | 0,02136513 | 3,45464171 | 0,21743039 | 0,8308976 | 0,92151175 |
| 3975_at | LHX1 | -0,06254358 | 5,00488564 | -0,21741972 | 0,83090575 | 0,92151175 |
| 4013_at | VWA5A | 0,01817761 | 4,34357391 | 0,21735209 | 0,83095747 | 0,92151605 |
| 25960_at | ADGRA2 | -0,02224236 | 4,76790889 | -0,21702815 | 0,83120522 | 0,92171923 |
| 6879_at | TAF7 | 0,02895304 | 9,38370526 | 0,21698741 | 0,83123638 | 0,92171923 |
| 8228_at | PNPLA4 | -0,05301572 | 7,73940706 | -0,21689551 | 0,83130666 | 0,9217441 |
| 122622_at | ADSSL1 | 0,03500709 | 7,50975721 | 0,21678716 | 0,83138954 | 0,92178293 |
| 10407_at | SPAG11B | -0,02475824 | 3,53907745 | -0,21655302 | 0,83156862 | 0,92190151 |
| 285830_at | HLA-F-AS1 | 0,04009055 | 7,66054598 | 0,21652219 | 0,83159221 | 0,92190151 |
| 8295_at | TRRAP | 0,0196134 | 7,41718717 | 0,21626993 | 0,83178517 | 0,92202155 |
| 55117_at | SLC6A15 | -0,01050117 | 3,194828 | -0,21625549 | 0,83179621 | 0,92202155 |
| 414194_at | CCNYL2 | 0,01665098 | 3,49283579 | 0,2161481 | 0,83187836 | 0,92205955 |
| 5976_at | UPF1 | -0,02989629 | 8,23273563 | -0,21604974 | 0,83195361 | 0,92208989 |
| 66037_at | BOLL | 0,01689725 | 4,01971835 | 0,21546808 | 0,8323986 | 0,92253002 |
| 22873_at | DZIP1 | 0,16849765 | 5,84106434 | 0,21521908 | 0,83258912 | 0,92268501 |
| 7571_at | ZNF23 | -0,01547604 | 5,88663247 | -0,21512982 | 0,83265741 | 0,92268501 |
| 54905_at | CYP2W1 | -0,02386574 | 5,25263123 | -0,21509748 | 0,83268215 | 0,92268501 |
| 83733_at | SLC25A18 | 0,01882513 | 4,40212251 | 0,21462202 | 0,83304598 | 0,92303507 |
| 79567_at | FAM65A | -0,03753593 | 7,80921143 | -0,21449389 | 0,83314403 | 0,92309062 |
| 145748_at | LYSMD4 | 0,03611311 | 6,45802836 | 0,21433339 | 0,83326686 | 0,92314073 |
| 1644_at | DDC | 0,01660562 | 4,40314775 | 0,21430954 | 0,83328511 | 0,92314073 |
| 1687_at | DFNA5 | 0,14126673 | 7,1114337 | 0,21412519 | 0,83342619 | 0,92324393 |
| 339512_at | C1orf110 | -0,01645883 | 3,48729348 | -0,21372788 | 0,83373028 | 0,92352768 |
| 148823_at | GCSAML | 0,01092488 | 3,32649854 | 0,2136233 | 0,83381032 | 0,92356323 |
| 7399_at | USH2A | 0,01140456 | 3,01735945 | 0,21350155 | 0,83390351 | 0,92358994 |
| 1893_at | ECM1 | -0,02230918 | 5,30734301 | -0,21345765 | 0,83393712 | 0,92358994 |
| 11100_at | HNRNPUL1 | 0,02103853 | 9,07829913 | 0,21337866 | 0,83399758 | 0,92358994 |

| | | | | | | |
|---|---|---|---|---|---|---|
| 10163_at | WASF2 | 0,01833288 | 9,2177073 | 0,21330887 | 0,834051 | 0,92358994 |
| 80133_at | MROH9 | -0,01832776 | 2,85944054 | -0,21321745 | 0,83412098 | 0,92358994 |
| 9177_at | HTR3B | -0,0262963 | 3,60032215 | -0,21321597 | 0,83412211 | 0,92358994 |
| 9915_at | ARNT2 | -0,08046439 | 5,72536512 | -0,21313273 | 0,83418584 | 0,92360741 |
| 9785_at | DHX38 | -0,03092785 | 7,5912036 | -0,21271759 | 0,83450364 | 0,92386488 |
| 356_at | FASLG | -0,01763987 | 3,82931398 | -0,21269682 | 0,83451954 | 0,92386488 |
| 9021_at | SOCS3 | -0,02007396 | 5,50818247 | -0,21264102 | 0,83456225 | 0,92386488 |
| 4628_at | MYH10 | -0,0287179 | 7,83367282 | -0,21251053 | 0,83466216 | 0,92392238 |
| 5083_at | PAX9 | 0,0199695 | 6,09644798 | 0,21240213 | 0,83474516 | 0,92394066 |
| 6568_at | SLC17A1 | 0,01361103 | 3,00906824 | 0,21227273 | 0,83484423 | 0,92394066 |
| 100507173_a | LINC01012 | 0,02229178 | 3,4024746 | 0,21226534 | 0,83484988 | 0,92394066 |
| 3257_at | HPS1 | -0,02322802 | 7,0068539 | -0,21219337 | 0,83490499 | 0,92394066 |
| 85453_at | TSPYL5 | -0,29789704 | 5,59452655 | -0,21217572 | 0,8349185 | 0,92394066 |
| 404266_at | HOXB-AS3 | 0,03458116 | 5,24373415 | 0,21163538 | 0,83533225 | 0,92434542 |
| 25819_at | NOCT | -0,02248816 | 6,10528655 | -0,21150032 | 0,83543567 | 0,92440676 |
| 5542_at | PRB1 | -0,037736 | 5,62688815 | -0,21143243 | 0,83548767 | 0,92441119 |
| 8711_at | TNK1 | 0,01712621 | 7,01322696 | 0,2107546 | 0,83600678 | 0,92483989 |
| 84073_at | MYCBPAP | -0,02397052 | 3,91406839 | -0,21074973 | 0,83601051 | 0,92483989 |
| 642938_at | FAM196A | 0,0136109 | 3,11050076 | 0,21073843 | 0,83601916 | 0,92483989 |
| 387751_at | GVINP1 | -0,01975513 | 3,7430693 | -0,21057665 | 0,83614308 | 0,92490062 |
| 63928_at | CHP2 | 0,01846836 | 5,17531201 | 0,21054139 | 0,83617008 | 0,92490062 |
| 643783_at | USP46-AS1 | 0,05352612 | 5,15427426 | 0,2103513 | 0,83631568 | 0,92495763 |
| 9113_at | LATS1 | -0,02162453 | 5,71275539 | -0,21030733 | 0,83634936 | 0,92495763 |
| 84767_at | TRIM51 | 0,03324465 | 4,32364865 | 0,21028603 | 0,83636568 | 0,92495763 |
| 7227_at | TRPS1 | 0,01409531 | 3,53973219 | 0,2098135 | 0,83672765 | 0,92530483 |
| 80329_at | ULBP1 | 0,03837721 | 5,4269404 | 0,20963201 | 0,83686669 | 0,92536644 |
| 114805_at | GALNT13 | -0,01042201 | 3,19249346 | -0,20961537 | 0,83687944 | 0,92536644 |
| 286370_at | LOC286370 | -0,01383054 | 3,51881812 | -0,20945461 | 0,8370026 | 0,92544949 |
| 101928335_a | LOC10192833 | -0,02696564 | 3,524967 | -0,20934125 | 0,83708945 | 0,9254924 |
| 401074_at | LINC00960 | -0,0327907 | 7,31279586 | -0,20903858 | 0,83732135 | 0,92569566 |
| 150_at | ADRA2A | 0,02335018 | 4,07857377 | 0,20877337 | 0,83752456 | 0,92582856 |
| 140767_at | NRSN1 | -0,02232064 | 4,10087197 | -0,20869557 | 0,83758418 | 0,92582856 |
| 1759_at | DNM1 | -0,025904 | 4,94538398 | -0,20861425 | 0,83764649 | 0,92582856 |
| 64748_at | LPPR2 | 0,03096467 | 6,41813999 | 0,20856967 | 0,83768065 | 0,92582856 |
| 84232_at | MAF1 | -0,04939272 | 8,62439907 | -0,20852101 | 0,83771794 | 0,92582856 |
| 201595_at | STT3B | 0,02600603 | 11,0598481 | 0,20850535 | 0,83772993 | 0,92582856 |
| 51200_at | CPA4 | 0,06289476 | 7,33528153 | 0,20824579 | 0,83792884 | 0,92593829 |
| 51032_at | CELA2B | 0,02069576 | 4,09207534 | 0,20822923 | 0,83794154 | 0,92593829 |
| 116540_at | MRPL53 | -0,04942946 | 10,3516015 | -0,2081876 | 0,83797344 | 0,92593829 |
| 83755_at | KRTAP4-12 | -0,0201236 | 3,19199122 | -0,20801895 | 0,83810269 | 0,92602799 |
| 283238_at | SLC22A24 | -0,02160463 | 3,60929213 | -0,20781766 | 0,83825695 | 0,92614532 |
| 5571_at | PRKAG1 | 0,02204564 | 9,33440285 | 0,20744329 | 0,83854388 | 0,92638724 |
| 55082_at | ARGLU1 | 0,03078422 | 8,58857336 | 0,20740648 | 0,8385721 | 0,92638724 |
| 55911_at | APOBR | -0,01973415 | 5,35557142 | -0,20728415 | 0,83866587 | 0,92640505 |
| 7678_at | ZNF124 | 0,05857267 | 5,69652812 | 0,207233 | 0,83870507 | 0,92640505 |
| 5537_at | PPP6C | 0,01975818 | 9,41306691 | 0,20719722 | 0,8387325 | 0,92640505 |
| 26045_at | LRRTM2 | -0,02315185 | 3,32214578 | -0,20705935 | 0,83883818 | 0,92646866 |
| 91392_at | ZNF502 | -0,06529011 | 4,0606504 | -0,2068968 | 0,83896278 | 0,92655315 |
| 1462_at | VCAN | -0,13342408 | 4,8536031 | -0,20674175 | 0,83908164 | 0,92663129 |
| 28972_at | SPCS1 | -0,02114291 | 11,0846548 | -0,20664193 | 0,83915816 | 0,92666268 |

| | | | | | | |
|---|---|---|---|---|---|---|
| 9530_at | BAG4 | -0,06974238 | 7,1358616 | -0,20635382 | 0,83937904 | 0,92683285 |
| 10050_at | SLC17A4 | -0,02801071 | 4,19930422 | -0,20631539 | 0,8394085 | 0,92683285 |
| 4129_at | MAOB | -0,02103349 | 4,88843494 | -0,20620487 | 0,83949323 | 0,92687328 |
| 9400_at | RECQL5 | 0,0171126 | 6,48517976 | 0,20590356 | 0,83972425 | 0,92706138 |
| 10952_at | SEC61B | -0,01371733 | 11,2935311 | -0,20580858 | 0,83979707 | 0,92706138 |
| 55526_at | DHTKD1 | -0,08530318 | 7,51705886 | -0,20573288 | 0,83985512 | 0,92706138 |
| 11183_at | MAP4K5 | 0,02524928 | 8,48302456 | 0,20573159 | 0,83985611 | 0,92706138 |
| 345630_at | FBLL1 | 0,0389583 | 4,77599626 | 0,20556811 | 0,83998146 | 0,92709989 |
| 7855_at | FZD5 | 0,01886235 | 7,97352188 | 0,20556056 | 0,83998725 | 0,92709989 |
| 2819_at | GPD1 | -0,04002179 | 4,86669863 | -0,20539157 | 0,84011683 | 0,92710066 |
| 3674_at | ITGA2B | -0,01512975 | 4,94488484 | -0,2053556 | 0,84014442 | 0,92710066 |
| 150274_at | HSCB | 0,03586596 | 6,23509075 | 0,20535084 | 0,84014807 | 0,92710066 |
| 1832_at | DSP | 0,0828824 | 10,1799676 | 0,20530859 | 0,84018046 | 0,92710066 |
| 79155_at | TNIP2 | -0,01218752 | 8,83434669 | -0,20516301 | 0,8402921 | 0,92717074 |
| 5552_at | SRGN | 0,35488561 | 8,517407 | 0,20491611 | 0,84048144 | 0,92726564 |
| 129684_at | CNTNAP5 | -0,01487526 | 3,53530319 | -0,20488303 | 0,84050682 | 0,92726564 |
| 290_at | ANPEP | -0,02065024 | 4,14222012 | -0,20486255 | 0,84052252 | 0,92726564 |
| 63950_at | DMRTA2 | -0,01714354 | 4,71757915 | -0,20463299 | 0,84069858 | 0,92730395 |
| 65080_at | MRPL44 | -0,03125046 | 7,98957115 | -0,2045278 | 0,84077926 | 0,92730395 |
| 55291_at | PPP6R3 | -0,00998652 | 9,85687909 | -0,20449587 | 0,84080374 | 0,92730395 |
| 80005_at | DOCK5 | -0,02034495 | 7,51062684 | -0,20448949 | 0,84080864 | 0,92730395 |
| 5406_at | PNLIP | -0,01914888 | 3,54088789 | -0,20440384 | 0,84087433 | 0,92730395 |
| 284723_at | SLC25A34 | 0,02800086 | 5,6087357 | 0,20440005 | 0,84087723 | 0,92730395 |
| 729178_at | STXBP5-AS1 | 0,03673335 | 4,6731904 | 0,2043779 | 0,84089422 | 0,92730395 |
| 624_at | BDKRB2 | -0,07735764 | 6,83436651 | -0,20387329 | 0,84128127 | 0,92767061 |
| 5305_at | PIP4K2A | 0,0617364 | 7,1292616 | 0,20381885 | 0,84132303 | 0,92767061 |
| 93587_at | TRMT10A | -0,02375326 | 5,42221343 | -0,20354865 | 0,84153031 | 0,92776012 |
| 10331_at | B3GNT3 | 0,02810886 | 8,08812689 | 0,20346588 | 0,8415938 | 0,92776012 |
| 51255_at | RNF181 | 0,01394057 | 10,1831349 | 0,2034305 | 0,84162094 | 0,92776012 |
| 79844_at | ZDHHC11 | 0,03735641 | 5,66795326 | 0,20337467 | 0,84166378 | 0,92776012 |
| 101927410_a | LOC10192741 | -0,01902066 | 4,33153857 | -0,20332096 | 0,84170498 | 0,92776012 |
| 100996511_a | LINC01355 | 0,02933106 | 4,30478001 | 0,20328518 | 0,84173243 | 0,92776012 |
| 28781_at | IGLV5-45 | -0,01473848 | 3,70848811 | -0,20327355 | 0,84174135 | 0,92776012 |
| 1134_at | CHRNA1 | -0,02077205 | 3,83817761 | -0,20307191 | 0,84189604 | 0,92787753 |
| 63932_at | CXorf56 | 0,01572999 | 4,24775116 | 0,20289703 | 0,84203022 | 0,92797231 |
| 6580_at | SLC22A1 | -0,02593215 | 4,238304 | -0,20283222 | 0,84207994 | 0,92797402 |
| 100505609_a | LOC10050560 | -0,01346552 | 3,16367525 | -0,20258839 | 0,84226703 | 0,9281271 |
| 55031_at | USP47 | 0,07813596 | 7,69240655 | 0,20229985 | 0,84248843 | 0,92831796 |
| 55425_at | GPALPP1 | 0,01675583 | 7,25547264 | 0,20219785 | 0,84256669 | 0,9283511 |
| 1143_at | CHRNB4 | -0,02160401 | 4,72885742 | -0,20206292 | 0,84267024 | 0,92841208 |
| 400128_at | TUSC8 | -0,01349926 | 3,16197262 | -0,20192297 | 0,84277764 | 0,9284773 |
| 387597_at | ILDR2 | 0,01859044 | 3,59396287 | 0,20182048 | 0,84285628 | 0,92848414 |
| 5310_at | PKD1 | 0,01074642 | 5,99849255 | 0,20178926 | 0,84288024 | 0,92848414 |
| 643338_at | C15orf62 | -0,02103966 | 5,51876599 | -0,20168097 | 0,84296335 | 0,92849284 |
| 286177_at | LOC286177 | -0,01968117 | 3,29496394 | -0,20156486 | 0,84305246 | 0,92849284 |
| 2132_at | EXT2 | -0,02879418 | 9,01135282 | -0,20150273 | 0,84310014 | 0,92849284 |
| 338651_at | KRTAP5-AS1 | 0,02963766 | 4,47206436 | 0,20147731 | 0,84311965 | 0,92849284 |
| 100505718_a | LINC01592 | -0,02008923 | 3,07581092 | -0,20146494 | 0,84312914 | 0,92849284 |
| 4504_at | MT3 | 0,02175778 | 5,79167737 | 0,20121424 | 0,84332155 | 0,92865163 |
| 64593_at | RBMY3AP | 0,02184123 | 3,05497777 | 0,20104137 | 0,84345424 | 0,92874465 |

| | | | | | | |
|---|---|---|---|---|---|---|
| 10769_at | PLK2 | 0,07974369 | 8,7094796 | 0,20062454 | 0,84377419 | 0,92904385 |
| 4065_at | LY75 | 0,21595315 | 5,8508088 | 0,20038666 | 0,84395679 | 0,9291918 |
| 253962_at | CACNA1G-AS1 | -0,02220374 | 4,53060115 | -0,20006798 | 0,84420143 | 0,92932851 |
| 118881_at | COMTD1 | -0,04355146 | 7,89117872 | -0,20005731 | 0,84420962 | 0,92932851 |
| 8045_at | RASSF7 | -0,01559962 | 8,34919421 | -0,20002474 | 0,84423463 | 0,92932851 |
| 84627_at | ZNF469 | 0,02531717 | 5,11104085 | 0,19993961 | 0,84429998 | 0,92932851 |
| 51150_at | SDF4 | -0,01202112 | 9,4154465 | -0,19986923 | 0,84435402 | 0,92932851 |
| 165082_at | ADGRF3 | 0,01669263 | 4,29319643 | 0,19984785 | 0,84437043 | 0,92932851 |
| 4053_at | LTBP2 | 0,03769465 | 7,12837457 | 0,19976669 | 0,84443274 | 0,929344 |
| 130026_at | ICA1L | 0,01240704 | 4,35133215 | 0,19963834 | 0,84453129 | 0,92937534 |
| 441046_at | GUSBP5 | 0,07547805 | 5,62210347 | 0,19959971 | 0,84456094 | 0,92937534 |
| 6307_at | MSMO1 | 0,07814718 | 10,7679248 | 0,19945513 | 0,84467196 | 0,92937534 |
| 58505_at | OSTC | -0,03071443 | 10,909593 | -0,19942837 | 0,8446925 | 0,92937534 |
| 6676_at | SPAG4 | 0,01826107 | 5,83965962 | 0,1994154 | 0,84470245 | 0,92937534 |
| 84684_at | INSM2 | -0,03008759 | 4,75576355 | -0,19924418 | 0,84483392 | 0,92944725 |
| 23049_at | SMG1 | 0,01349467 | 8,00323781 | 0,19915094 | 0,84490552 | 0,92944725 |
| 6871_at | TADA2A | 0,01389849 | 6,22499785 | 0,19914176 | 0,84491257 | 0,92944725 |
| 92999_at | ZBTB47 | -0,01274534 | 6,25875294 | -0,19899195 | 0,8450276 | 0,92946426 |
| 56944_at | OLFML3 | 0,0267131 | 4,33964287 | 0,19899177 | 0,84502774 | 0,92946426 |
| 26166_at | RGS22 | -0,0200716 | 3,79636325 | -0,19893311 | 0,84507278 | 0,92946426 |
| 85442_at | KNDC1 | -0,02541117 | 5,22986569 | -0,1988246 | 0,84515611 | 0,92950283 |
| 1938_at | EEF2 | -0,01277287 | 12,3519748 | -0,19862802 | 0,84530707 | 0,92961395 |
| 100499171_at | LINC00276 | 0,01585846 | 3,05743332 | 0,19856735 | 0,84535366 | 0,92961395 |
| 387486_at | LINC00320 | -0,01446652 | 3,06677324 | -0,19808475 | 0,84572429 | 0,92996843 |
| 220002_at | CYB561A3 | -0,02425227 | 8,79729357 | -0,19786013 | 0,84589681 | 0,93002714 |
| 130120_at | REG3G | -0,01613953 | 3,44149791 | -0,197856 | 0,84589998 | 0,93002714 |
| 27129_at | HSPB7 | -0,01931661 | 4,54741454 | -0,19782665 | 0,84592252 | 0,93002714 |
| 26047_at | CNTNAP2 | -0,02249195 | 3,91075609 | -0,19767292 | 0,8460406 | 0,93005132 |
| 5470_at | PPEF2 | 0,01889301 | 3,20651878 | 0,1976723 | 0,84604108 | 0,93005132 |
| 142689_at | ASB12 | 0,01878562 | 3,92335785 | 0,19746705 | 0,84619873 | 0,93017155 |
| 11132_at | CAPN10 | 0,01443943 | 6,50907475 | 0,19731048 | 0,846319 | 0,9302385 |
| 27030_at | MLH3 | 0,04501884 | 6,59213284 | 0,19722594 | 0,84638394 | 0,9302385 |
| 150280_at | HORMAD2 | 0,01955554 | 3,81961483 | 0,19715857 | 0,84643569 | 0,9302385 |
| 55891_at | LENEP | -0,02592858 | 4,84366549 | -0,19710777 | 0,84647472 | 0,9302385 |
| 6493_at | SIM2 | -0,05131458 | 7,10348972 | -0,19707343 | 0,8465011 | 0,9302385 |
| 283579_at | C14orf178 | 0,01452025 | 5,26979202 | 0,19689094 | 0,84664128 | 0,93033947 |
| 51547_at | SIRT7 | 0,01996619 | 8,59735287 | 0,19682646 | 0,84669082 | 0,93034084 |
| 284424_at | MIR7-3HG | -0,01961996 | 4,07727146 | -0,19650434 | 0,84693829 | 0,93055403 |
| 867_at | CBL | 0,03139525 | 7,20388999 | 0,19644815 | 0,84698145 | 0,93055403 |
| 729307_at | LOC729307 | -0,01934246 | 3,24134632 | -0,19615265 | 0,8472085 | 0,9307132 |
| 100189589_at | DCTN1-AS1 | -0,02160063 | 3,40373128 | -0,19609333 | 0,84725407 | 0,9307132 |
| 6845_at | VAMP7 | 0,02271027 | 9,92199876 | 0,19607093 | 0,84727128 | 0,9307132 |
| 64506_at | CPEB1 | -0,01965905 | 3,83663052 | -0,19580549 | 0,84747524 | 0,93087216 |
| 53405_at | CLIC5 | -0,02490411 | 4,26135526 | -0,19571493 | 0,84754483 | 0,93087216 |
| 341032_at | C11orf53 | 0,12950758 | 7,11483564 | 0,19569393 | 0,84756096 | 0,93087216 |
| 51386_at | EIF3L | -0,02085968 | 11,4129108 | -0,19535718 | 0,84781974 | 0,93109277 |
| 26062_at | HYALP1 | -0,01610829 | 3,17768238 | -0,19530674 | 0,8478585 | 0,93109277 |
| 3855_at | KRT7 | -0,02698552 | 12,610323 | -0,19490989 | 0,84816348 | 0,93135695 |
| 100873979_at | RBMS3-AS3 | -0,02030048 | 3,34805563 | -0,19486788 | 0,84819577 | 0,93135695 |
| 2571_at | GAD1 | -0,06546603 | 5,81849921 | -0,19476332 | 0,84827613 | 0,9313921 |

| | | | | | | |
|---|---|---|---|---|---|---|
| 2972_at | BRF1 | 0,01170469 | 6,52531496 | 0,1946233 | 0,84838375 | 0,93142543 |
| 101805491_a | LOC10180549 | 0,0141009 | 3,05682662 | 0,19449776 | 0,84848023 | 0,93142543 |
| 100874323_a | HOXA10-AS | -0,10263989 | 5,65999289 | -0,19447227 | 0,84849982 | 0,93142543 |
| 929_at | CD14 | -0,03156466 | 4,46797452 | -0,19444063 | 0,84852414 | 0,93142543 |
| 10245_at | TIMM17B | -0,0257682 | 8,48401084 | -0,19435286 | 0,84859161 | 0,93142543 |
| 54545_at | MTMR12 | 0,01219793 | 7,04548464 | 0,19426269 | 0,84866091 | 0,93142543 |
| 51214_at | IGF2-AS | -0,01700263 | 6,02510918 | -0,19422809 | 0,8486875 | 0,93142543 |
| 29895_at | MYLPF | 0,02875324 | 5,95238848 | 0,19422055 | 0,84869331 | 0,93142543 |
| 84553_at | FAXC | -0,01317634 | 4,19857035 | -0,19398784 | 0,84887218 | 0,93152474 |
| 654790_at | PCP4L1 | -0,02191731 | 4,55958527 | -0,19387885 | 0,84895596 | 0,93152474 |
| 58506_at | SCAF1 | 0,01837381 | 5,97554791 | 0,19382682 | 0,84899595 | 0,93152474 |
| 2312_at | FLG | 0,01406585 | 3,90640646 | 0,19376787 | 0,84904127 | 0,93152474 |
| 115948_at | CCDC151 | -0,01672613 | 4,78178476 | -0,19369239 | 0,84909929 | 0,93152474 |
| 26953_at | RANBP6 | 0,0368998 | 8,66224579 | 0,19366892 | 0,84911733 | 0,93152474 |
| 285755_at | PPIL6 | 0,02104007 | 4,01432808 | 0,19366246 | 0,8491223 | 0,93152474 |
| 4690_at | NCK1 | -0,04218869 | 6,8595269 | -0,19335749 | 0,84935675 | 0,93172888 |
| 56676_at | ASCL3 | -0,02066333 | 3,87866017 | -0,19308543 | 0,84956592 | 0,93190526 |
| 10650_at | PRELID3A | -0,02392893 | 4,42744375 | -0,19272662 | 0,84984179 | 0,93215479 |
| 10161_at | LPAR6 | -0,01316985 | 3,31957567 | -0,19255587 | 0,84997308 | 0,93224571 |
| 7367_at | UGT2B17 | 0,01573896 | 2,7688126 | 0,19241944 | 0,85007798 | 0,93230768 |
| 200185_at | KRTCAP2 | 0,01177986 | 10,6983567 | 0,19210508 | 0,85031971 | 0,93251971 |
| 113835_at | ZNF257 | 0,03047191 | 3,33781552 | 0,19186916 | 0,85050113 | 0,93266556 |
| 221336_at | BEND6 | -0,03897876 | 5,07167014 | -0,19166915 | 0,85065494 | 0,93273517 |
| 940_at | CD28 | 0,00973459 | 3,67317304 | 0,1916607 | 0,85066145 | 0,93273517 |
| 8976_at | WASL | 0,03052547 | 8,94847457 | 0,19158603 | 0,85071887 | 0,93274504 |
| 9051_at | PSTPIP1 | -0,02413634 | 5,48012555 | -0,19149052 | 0,85079233 | 0,93274786 |
| 79798_at | ARMC5 | 0,02985522 | 6,18267144 | 0,19139998 | 0,85086196 | 0,93274786 |
| 83442_at | SH3BGRL3 | -0,05705039 | 10,190417 | -0,19139381 | 0,85086671 | 0,93274786 |
| 101929536_a | LOC10192953 | 0,03825017 | 4,72783639 | 0,19086212 | 0,85127564 | 0,93301875 |
| 1278_at | COL1A2 | 0,0193498 | 4,58595084 | 0,1908456 | 0,85128835 | 0,93301875 |
| 22989_at | MYH15 | -0,01316014 | 3,31638316 | -0,19080765 | 0,85131755 | 0,93301875 |
| 208_at | AKT2 | 0,0123559 | 7,85985275 | 0,19079315 | 0,8513287 | 0,93301875 |
| 57477_at | SHROOM4 | 0,03983664 | 4,74617619 | 0,19075766 | 0,85135599 | 0,93301875 |
| 29903_at | CCDC106 | 0,08396727 | 5,97230999 | 0,19062188 | 0,85146043 | 0,93308012 |
| 101927059_a | LOC10192705 | 0,01472178 | 3,08603401 | 0,19041526 | 0,85161938 | 0,93319904 |
| 222950_at | NYAP1 | -0,02168007 | 5,70712776 | -0,19035486 | 0,85166584 | 0,93319904 |
| 6495_at | SIX1 | -0,05815999 | 7,98367489 | -0,19002557 | 0,85191915 | 0,93335039 |
| 23624_at | CBLC | -0,14511403 | 5,89816963 | -0,19001471 | 0,85192751 | 0,93335039 |
| 1632_at | ECI1 | 0,02067474 | 8,73886367 | 0,18998635 | 0,85194933 | 0,93335039 |
| 101927151_a | LOC10192715 | -0,01601507 | 6,91024829 | -0,18985291 | 0,85205198 | 0,93336803 |
| 81490_at | PTDSS2 | 0,02006543 | 7,22085677 | 0,18979137 | 0,85209934 | 0,93336803 |
| 92340_at | PRR29 | -0,01772567 | 5,91475393 | -0,18977648 | 0,85211079 | 0,93336803 |
| 23095_at | KIF1B | -0,01362044 | 8,05833864 | -0,1895475 | 0,85228696 | 0,93341639 |
| 58475_at | MS4A7 | 0,0110083 | 3,35881816 | 0,1895082 | 0,8523172 | 0,93341639 |
| 26219_at | OR1J4 | -0,01455964 | 3,25293165 | -0,18949218 | 0,85232952 | 0,93341639 |
| 79724_at | ZNF768 | 0,02872397 | 7,32452339 | 0,18940813 | 0,85239419 | 0,93341639 |
| 80790_at | CMIP | -0,0232759 | 8,08815411 | -0,18940419 | 0,85239722 | 0,93341639 |
| 5837_at | PYGM | -0,01757025 | 5,27763795 | -0,18924771 | 0,85251762 | 0,93344275 |
| 5588_at | PRKCQ | 0,04255749 | 5,28160203 | 0,18924694 | 0,85251821 | 0,93344275 |
| 7181_at | NR2C1 | -0,02734231 | 6,03091492 | -0,18912248 | 0,85261398 | 0,93349455 |

| | | | | | | |
|---|---|---|---|---|---|---|
| 80332_at | ADAM33 | 0,01608337 | 5,16712186 | 0,18905752 | 0,85266396 | 0,93349621 |
| 10288_at | LILRB2 | -0,01526727 | 4,02466307 | -0,18878577 | 0,85287306 | 0,93367207 |
| 59351_at | PBOV1 | 0,01989466 | 3,63149127 | 0,188661 | 0,85296908 | 0,93372412 |
| 4901_at | NRL | -0,02283255 | 5,27942259 | -0,18848974 | 0,85310087 | 0,93381532 |
| 221424_at | LRRC73 | -0,01925236 | 4,82854966 | -0,18833358 | 0,85322104 | 0,9338938 |
| 122961_at | ISCA2 | 0,01535379 | 8,48659085 | 0,18825136 | 0,85328432 | 0,93390877 |
| 836_at | CASP3 | 0,05511567 | 8,39963719 | 0,1881898 | 0,85333169 | 0,93390877 |
| 29931_at | LINC00312 | -0,01522434 | 4,15118338 | -0,18810561 | 0,85339648 | 0,93392495 |
| 58538_at | MPP4 | -0,01191121 | 3,22166513 | -0,1880353 | 0,8534506 | 0,93392495 |
| 27333_at | GOLIM4 | -0,03947788 | 7,59669727 | -0,18798161 | 0,85349192 | 0,93392495 |
| #N/A | CTD-3080P12, | -0,01992733 | 3,78365847 | -0,18789251 | 0,85356049 | 0,9339411 |
| 84926_at | SPRYD3 | 0,0202841 | 7,77429855 | 0,18783277 | 0,85360647 | 0,9339411 |
| 169834_at | ZNF883 | -0,12086143 | 4,14411559 | -0,18777344 | 0,85365213 | 0,9339411 |
| 481_at | ATP1B1 | -0,07159497 | 10,8806424 | -0,1876411 | 0,85375399 | 0,9339995 |
| 54511_at | HMGCLL1 | 0,01141847 | 3,1673535 | 0,18755871 | 0,85381741 | 0,93401583 |
| 344405_at | PRORSD1P | -0,02332812 | 4,43254455 | -0,18735052 | 0,85397765 | 0,93413807 |
| 101927295_a | LINC01360 | 0,01420183 | 2,88148552 | 0,18724732 | 0,85405709 | 0,93417192 |
| 81615_at | TMEM163 | -0,01762878 | 4,03650373 | -0,18704253 | 0,85421472 | 0,93422925 |
| 729652_at | LOC729652 | -0,02591434 | 3,52664223 | -0,18699672 | 0,85424999 | 0,93422925 |
| 100506725_a | LOC10050672 | -0,02302952 | 3,75998415 | -0,18693493 | 0,85429756 | 0,93422925 |
| 84329_at | HVCN1 | 0,01648826 | 6,05474708 | 0,18688711 | 0,85433436 | 0,93422925 |
| 100507194_a | LOC10050719 | -0,01783302 | 3,45676375 | -0,18684996 | 0,85436296 | 0,93422925 |
| 5566_at | PRKACA | -0,02282175 | 7,06035178 | -0,18680121 | 0,85440049 | 0,93422925 |
| 196051_at | PPAPDC1A | -0,0207485 | 3,99709069 | -0,18671322 | 0,85446823 | 0,93425028 |
| 64777_at | RMND5B | -0,01554002 | 8,1475301 | -0,18652138 | 0,85461591 | 0,93435872 |
| 153562_at | MARVELD2 | 0,02106652 | 7,11055167 | 0,18644939 | 0,85467133 | 0,93436628 |
| 25828_at | TXN2 | -0,01377608 | 9,31611333 | -0,18628758 | 0,8547959 | 0,93444944 |
| 168667_at | BMPER | 0,01416186 | 3,82145585 | 0,18602932 | 0,85499475 | 0,93461377 |
| 2940_at | GSTA3 | -0,01953022 | 3,90176393 | -0,18595042 | 0,85505549 | 0,93462713 |
| 26281_at | FGF20 | 0,01336665 | 4,29990342 | 0,18583183 | 0,8551468 | 0,9346739 |
| 169966_at | FAM46D | 0,01519373 | 3,01469547 | 0,18563088 | 0,85530152 | 0,93477792 |
| 55321_at | TMEM74B | -0,03919146 | 5,83357664 | -0,18558217 | 0,85533903 | 0,93477792 |
| 282775_at | OR5J2 | -0,01908876 | 3,3115293 | -0,18538734 | 0,85548905 | 0,93488202 |
| 116337_at | PANX3 | -0,01423162 | 3,29125592 | -0,18520062 | 0,85563283 | 0,93488202 |
| 166968_at | MIER3 | 0,04356465 | 6,56231223 | 0,18513767 | 0,85568131 | 0,93488202 |
| 8530_at | CST7 | -0,02367605 | 5,77450941 | -0,18512365 | 0,8556921 | 0,93488202 |
| 26664_at | OR7C1 | -0,01660806 | 3,50499834 | -0,18506577 | 0,85573667 | 0,93488202 |
| 100506286_a | LOC10050628 | -0,02170649 | 3,57276005 | -0,18498662 | 0,85579762 | 0,93488202 |
| 3903_at | LAIR1 | 0,01598686 | 5,13557037 | 0,18496006 | 0,85581808 | 0,93488202 |
| 575_at | ADGRB1 | -0,01536181 | 5,52700335 | -0,18491153 | 0,85585545 | 0,93488202 |
| 27293_at | SMPDL3B | 0,03998575 | 5,55979185 | 0,18489125 | 0,85587107 | 0,93488202 |
| 105377621_a | LOC10537762 | -0,0313039 | 5,06580554 | -0,18473173 | 0,85599392 | 0,93496319 |
| 80781_at | COL18A1 | 0,09427115 | 6,85276159 | 0,18453978 | 0,85614174 | 0,93507163 |
| 91748_at | ELMSAN1 | -0,01719289 | 6,83532098 | -0,18431489 | 0,85631495 | 0,93515969 |
| 51734_at | MSRB1 | 0,02312312 | 9,40327973 | 0,18419324 | 0,85640864 | 0,93515969 |
| 203260_at | CCDC107 | 0,0167792 | 6,57882858 | 0,18418865 | 0,85641218 | 0,93515969 |
| 8639_at | AOC3 | 0,02418445 | 4,20581674 | 0,18412092 | 0,85646435 | 0,93515969 |
| 285782_at | CAGE1 | 0,01207571 | 2,7733292 | 0,18411994 | 0,8564651 | 0,93515969 |
| 51314_at | NME8 | -0,0159792 | 3,1634443 | -0,18404651 | 0,85652166 | 0,93516844 |
| 101927111_a | SUCLG2-AS1 | 0,01012991 | 3,98296373 | 0,18377722 | 0,85672908 | 0,93534189 |

| | | | | | | |
|---|---|---|---|---|---|---|
| 55599_at | RNPC3 | -0,02798118 | 7,02450886 | -0,18343756 | 0,85699072 | 0,93551928 |
| 5352_at | PLOD2 | 0,0787287 | 9,74656347 | 0,18339111 | 0,8570265 | 0,93551928 |
| 56905_at | C15orf39 | -0,01552691 | 7,35622722 | -0,18337714 | 0,85703726 | 0,93551928 |
| 83752_at | LONP2 | 0,01326778 | 8,82368955 | 0,18326369 | 0,85712465 | 0,93554052 |
| 153478_at | PLEKHG4B | -0,02222154 | 4,56052619 | -0,18322579 | 0,85715385 | 0,93554052 |
| 26152_at | ZNF337 | -0,02492375 | 6,89440101 | -0,18304883 | 0,85729018 | 0,93563533 |
| 7348_at | UPK1B | 0,02490341 | 9,58303416 | 0,18298693 | 0,85733786 | 0,93563533 |
| 26280_at | IL1RAPL2 | -0,01838416 | 3,70453594 | -0,18283321 | 0,85745628 | 0,9356962 |
| 4140_at | MARK3 | 0,01740849 | 8,59855366 | 0,18271201 | 0,85754966 | 0,9356962 |
| 1814_at | DRD3 | 0,01546401 | 3,75441245 | 0,18268613 | 0,8575696 | 0,9356962 |
| 2328_at | FMO3 | 0,0102171 | 3,15436372 | 0,18266232 | 0,85758794 | 0,9356962 |
| 284406_at | ZFP82 | -0,02789737 | 5,3134391 | -0,1825746 | 0,85765553 | 0,93570784 |
| 101929083_a | CASC6 | 0,01770588 | 3,06399237 | 0,18252238 | 0,85769576 | 0,93570784 |
| 4340_at | MOG | 0,01391586 | 3,72006832 | 0,18227931 | 0,85788303 | 0,93580113 |
| 143425_at | SYT9 | -0,01923423 | 3,65673338 | -0,18219499 | 0,857948 | 0,93580113 |
| 29095_at | ORMDL2 | -0,04661659 | 10,3501293 | -0,18215192 | 0,85798119 | 0,93580113 |
| 1029_at | CDKN2A | 0,01178918 | 4,22873639 | 0,18209638 | 0,85802399 | 0,93580113 |
| 2878_at | GPX3 | 0,11265924 | 7,45783526 | 0,18203844 | 0,85806863 | 0,93580113 |
| 89792_at | GAL3ST3 | 0,01749385 | 5,58282906 | 0,1820331 | 0,85807274 | 0,93580113 |
| 84942_at | WDR73 | 0,00939845 | 8,12261881 | 0,18195929 | 0,85812962 | 0,93581017 |
| 9817_at | KEAP1 | -0,02205588 | 9,64728324 | -0,18181142 | 0,85824356 | 0,93588144 |
| #N/A | CTD-2350J17, | -0,01368111 | 3,17511685 | -0,18169683 | 0,85833186 | 0,93588866 |
| 253430_at | IPMK | 0,01483945 | 5,49706562 | 0,18167673 | 0,85834735 | 0,93588866 |
| 5252_at | PHF1 | -0,0096706 | 7,71075114 | -0,18146454 | 0,85851086 | 0,93596277 |
| 55211_at | DPPA4 | 0,01269347 | 3,240431 | 0,18142439 | 0,8585418 | 0,93596277 |
| 3750_at | KCND1 | -0,02402529 | 4,82820229 | -0,18139217 | 0,85856663 | 0,93596277 |
| 129804_at | FBLN7 | -0,0195547 | 4,7215545 | -0,18133631 | 0,85860968 | 0,93596277 |
| 55532_at | SLC30A10 | 0,01592376 | 3,55995957 | 0,1812027 | 0,85871265 | 0,93599269 |
| 11126_at | CD160 | 0,02588111 | 3,43322514 | 0,1811429 | 0,85875873 | 0,93599269 |
| 28457_at | IGHV2-5 | -0,01787755 | 5,43566129 | -0,18111155 | 0,85878289 | 0,93599269 |
| 3606_at | IL18 | -0,10912997 | 7,33069608 | -0,1809962 | 0,85887179 | 0,93603662 |
| 132884_at | EVC2 | 0,04732586 | 4,53203905 | 0,18091615 | 0,85893348 | 0,9360509 |
| 222183_at | SRRM3 | -0,01440724 | 6,65055423 | -0,18077208 | 0,85904452 | 0,93611894 |
| 254773_at | LYG2 | 0,0181783 | 4,48552869 | 0,18052391 | 0,85923579 | 0,93627441 |
| 388182_at | SPATA41 | 0,0129913 | 3,95454755 | 0,18032744 | 0,85938722 | 0,93633623 |
| 101060391_a | LOC10106039 | 0,01421825 | 3,38756088 | 0,18032417 | 0,85938974 | 0,93633623 |
| 101669762_a | BLACAT1 | 0,01839948 | 3,64312451 | 0,18016631 | 0,85951142 | 0,93641584 |
| 2791_at | GNG11 | -0,10485274 | 5,00430418 | -0,18005946 | 0,85959378 | 0,93645261 |
| 221496_at | LEMD2 | 0,00996331 | 7,41327486 | 0,17977982 | 0,85980934 | 0,93658349 |
| 114798_at | SLITRK1 | 0,01789492 | 3,45946772 | 0,17973093 | 0,85984703 | 0,93658349 |
| 256710_at | GLIPR1L1 | 0,01284608 | 3,93944315 | 0,17971438 | 0,85985979 | 0,93658349 |
| 148304_at | C1orf74 | -0,03781645 | 6,33802808 | -0,17953669 | 0,85999676 | 0,93665519 |
| 57125_at | PLXDC1 | 0,01001842 | 3,99695396 | 0,17950285 | 0,86002286 | 0,93665519 |
| 26033_at | ATRNL1 | 0,010225 | 3,15006597 | 0,1794107 | 0,86009389 | 0,93667959 |
| 401134_at | LOC401134 | -0,00851137 | 2,81051982 | -0,1792891 | 0,86018764 | 0,93672873 |
| 10218_at | ANGPTL7 | -0,01496807 | 4,42781125 | -0,1791213 | 0,860317 | 0,93674478 |
| 8623_at | ASMTL | -0,06994804 | 5,94743513 | -0,17898106 | 0,86042513 | 0,93674478 |
| 144402_at | CPNE8 | -0,05096427 | 4,23380368 | -0,1789574 | 0,86044337 | 0,93674478 |
| 219578_at | ZNF804B | -0,01471327 | 3,45621941 | -0,17893781 | 0,86045847 | 0,93674478 |
| 177_at | AGER | -0,01445142 | 5,10168849 | -0,17889182 | 0,86049393 | 0,93674478 |

| | | | | | | |
|---|---|---|---|---|---|---|
| 219539_at | YPEL4 | -0,01853517 | 4,28623402 | -0,17889153 | 0,86049415 | 0,93674478 |
| 8857_at | FCGBP | 0,01729892 | 4,93808979 | 0,17871127 | 0,86063313 | 0,93684313 |
| 23248_at | RPRD2 | 0,01423997 | 7,81177188 | 0,17854431 | 0,86076187 | 0,93690546 |
| 646903_at | LOC646903 | -0,07662934 | 5,21138546 | -0,17844524 | 0,86083826 | 0,93690546 |
| 10777_at | ARPP21 | -0,00880419 | 3,20662718 | -0,17834689 | 0,86091409 | 0,93690546 |
| 219770_at | GJD4 | -0,01552023 | 4,72562248 | -0,17833246 | 0,86092521 | 0,93690546 |
| 100861523_a | RPS6KA2-AS1 | -0,01788454 | 4,49748457 | -0,17829685 | 0,86095267 | 0,93690546 |
| 100132356_a | LOC10013235 | 0,03063241 | 4,10952639 | 0,17825854 | 0,86098222 | 0,93690546 |
| 151871_at | DPPA2 | -0,02076736 | 3,29638531 | -0,17813338 | 0,86107872 | 0,93692513 |
| 8128_at | ST8SIA2 | 0,00944593 | 3,89425366 | 0,17810894 | 0,86109757 | 0,93692513 |
| 8590_at | OR6A2 | -0,02745759 | 3,94328304 | -0,17790576 | 0,86125424 | 0,93704267 |
| 5097_at | PCDH1 | -0,02263067 | 6,99029648 | -0,17778628 | 0,86134638 | 0,93708944 |
| 132299_at | OCIAD2 | 0,01478019 | 9,84055377 | 0,17772386 | 0,86139452 | 0,93708944 |
| 144717_at | FAM109A | 0,0154128 | 6,84028222 | 0,17755667 | 0,86152346 | 0,93717678 |
| 84666_at | RETNLB | -0,01611944 | 4,12230869 | -0,17745889 | 0,86159886 | 0,93720588 |
| 8556_at | CDC14A | -0,00947346 | 4,68168069 | -0,17711771 | 0,86186199 | 0,93743564 |
| 100996597_a | LINC00690 | -0,01696219 | 2,89985052 | -0,17702846 | 0,86193082 | 0,93743564 |
| 26505_at | CNNM3 | 0,01030088 | 6,79891391 | 0,17699571 | 0,86195608 | 0,93743564 |
| 374655_at | ZNF710 | -0,010312 | 6,04669394 | -0,17668653 | 0,86219456 | 0,93764206 |
| 9416_at | DDX23 | 0,01090021 | 9,44252043 | 0,17660497 | 0,86225746 | 0,93765753 |
| 147657_at | ZNF480 | -0,04088423 | 7,19067817 | -0,17646469 | 0,86236567 | 0,93772226 |
| 5162_at | PDHB | -0,02237449 | 8,97080048 | -0,17638746 | 0,86242524 | 0,9377341 |
| 28959_at | TMEM176B | 0,02293751 | 4,66662945 | 0,17619964 | 0,86257011 | 0,93783869 |
| 55806_at | HR | 0,02740007 | 6,73240716 | 0,1759755 | 0,86274302 | 0,9379356 |
| 7188_at | TRAF5 | -0,02559761 | 5,47421085 | -0,17595786 | 0,86275663 | 0,9379356 |
| 2557_at | GABRA4 | -0,01518626 | 3,91410068 | -0,17584515 | 0,86284358 | 0,93797719 |
| 25786_at | DGCR11 | -0,02565148 | 6,46391309 | -0,17545119 | 0,86314751 | 0,93825464 |
| 9104_at | RGN | 0,01331751 | 3,40216121 | 0,17534782 | 0,86322725 | 0,93828837 |
| 5325_at | PLAGL1 | 0,14201731 | 6,14473873 | 0,17481378 | 0,86363929 | 0,93866172 |
| 645682_at | POU5F1P4 | -0,0278246 | 4,23953767 | -0,17474256 | 0,86369425 | 0,93866172 |
| 101928449_a | LOC10192844 | 0,01571808 | 3,33497924 | 0,17471318 | 0,86371692 | 0,93866172 |
| 339669_at | TEX33 | -0,01442578 | 3,38070841 | -0,17454532 | 0,86384644 | 0,93869737 |
| 285401_at | LINC00698 | -0,01311843 | 3,78237679 | -0,17448356 | 0,86389409 | 0,93869737 |
| 5104_at | SERPINA5 | -0,01287899 | 4,36805386 | -0,1744812 | 0,86389592 | 0,93869737 |
| 1690_at | COCH | -0,04553873 | 5,8834536 | -0,17440527 | 0,8639545 | 0,93870808 |
| 91661_at | ZNF765 | -0,01524915 | 6,96546016 | -0,17428059 | 0,86405072 | 0,93872083 |
| 124872_at | B4GALNT2 | 0,02042139 | 4,49888031 | 0,17425471 | 0,86407069 | 0,93872083 |
| 7879_at | RAB7A | -0,01946898 | 10,6860954 | -0,1741437 | 0,86415635 | 0,93872083 |
| 7290_at | HIRA | -0,02726783 | 8,87111493 | -0,17413747 | 0,86416116 | 0,93872083 |
| 80196_at | RNF34 | 0,01716269 | 9,12295646 | 0,17388365 | 0,86435703 | 0,93881641 |
| 5894_at | RAF1 | -0,01968633 | 9,50631092 | -0,17386769 | 0,86436935 | 0,93881641 |
| 54760_at | PCSK4 | -0,0185954 | 6,53379851 | -0,17383398 | 0,86439537 | 0,93881641 |
| 346653_at | FAM71F2 | -0,02439303 | 4,55919588 | -0,1737217 | 0,86448201 | 0,93885759 |
| 668_at | FOXL2 | -0,03033899 | 4,67397399 | -0,17355802 | 0,86460833 | 0,93894184 |
| 146771_at | TCAM1P | 0,03402328 | 3,37973203 | 0,17349142 | 0,86465973 | 0,93894472 |
| 101927597_a | NCK1-AS1 | 0,02139679 | 6,67014653 | 0,17304047 | 0,86500779 | 0,93926973 |
| 50652_at | PCA3 | -0,01427776 | 3,55214218 | -0,17287388 | 0,86513636 | 0,93935639 |
| 7062_at | TCHH | -0,08120704 | 4,6509268 | -0,17275093 | 0,86523127 | 0,93938195 |
| 347734_at | SLC35B2 | -0,02748403 | 9,53079953 | -0,17265859 | 0,86530255 | 0,93938195 |
| 441054_at | C4orf47 | 0,01115286 | 3,59761227 | 0,17265385 | 0,8653062 | 0,93938195 |

| | | | | | | |
|---|---|---|---|---|---|---|
| 7570_at | ZNF22 | 0,13488947 | 6,37458842 | 0,17237929 | 0,86551814 | 0,93955909 |
| 10463_at | SLC30A9 | 0,03100426 | 9,30583595 | 0,1722601 | 0,86561015 | 0,9395709 |
| 378828_at | LINC00317 | -0,01639543 | 3,63544413 | -0,17216306 | 0,86568506 | 0,9395709 |
| 112574_at | SNX18 | -0,01343966 | 6,80457032 | -0,17213112 | 0,86570972 | 0,9395709 |
| 8512_at | MBL1P | -0,02300127 | 4,76692098 | -0,1720668 | 0,86575937 | 0,9395709 |
| 79648_at | MCPH1 | -0,0573107 | 5,60846571 | -0,17204927 | 0,8657729 | 0,9395709 |
| 338872_at | C1QTNF9 | 0,01869109 | 3,02066366 | 0,17192807 | 0,86586647 | 0,9396195 |
| 101929584_a | LOC10192958 | -0,01867091 | 4,35002061 | -0,17184821 | 0,86592812 | 0,93963347 |
| 594_at | BCKDHB | -0,02906602 | 6,43060829 | -0,17144085 | 0,86624262 | 0,93980207 |
| 81501_at | DCSTAMP | -0,01633313 | 3,83401049 | -0,17139544 | 0,86627768 | 0,93980207 |
| 5281_at | PIGF | -0,01877615 | 7,6704263 | -0,1713694 | 0,86629778 | 0,93980207 |
| 27164_at | SALL3 | 0,01250979 | 4,40585492 | 0,17133641 | 0,86632325 | 0,93980207 |
| 374899_at | ZNF829 | 0,03468867 | 4,69258837 | 0,17132472 | 0,86633228 | 0,93980207 |
| 10677_at | AVIL | 0,02853676 | 4,77873317 | 0,17126512 | 0,8663783 | 0,93980207 |
| 101929110_a | CASC18 | 0,01187076 | 3,52526299 | 0,17120462 | 0,86642501 | 0,93980207 |
| 400238_at | LINC00642 | 0,01265701 | 3,92413 | 0,17100725 | 0,86657741 | 0,93988159 |
| 78991_at | PCYOX1L | 0,03624262 | 7,97821733 | 0,17098328 | 0,86659591 | 0,93988159 |
| 51441_at | YTHDF2 | -0,02556572 | 9,97973973 | -0,17068114 | 0,86682921 | 0,94008169 |
| 4041_at | LRP5 | 0,02406588 | 8,19668553 | 0,17056488 | 0,86691898 | 0,94012613 |
| 113178_at | SCAMP4 | 0,01452309 | 7,13309739 | 0,17041808 | 0,86703235 | 0,94019613 |
| 100506724_a | LINC00901 | 0,01449512 | 2,85832576 | 0,1702923 | 0,86712948 | 0,94024853 |
| 51768_at | TM7SF3 | 0,05634882 | 8,84796698 | 0,17000395 | 0,86735216 | 0,94038431 |
| 7564_at | ZNF16 | 0,04531083 | 6,67016018 | 0,17000372 | 0,86735233 | 0,94038431 |
| 64092_at | SAMSN1 | 0,00845864 | 3,13425959 | 0,16973594 | 0,86755913 | 0,94046736 |
| 56956_at | LHX9 | -0,01104584 | 4,03356862 | -0,16968745 | 0,86759658 | 0,94046736 |
| 51311_at | TLR8 | -0,01149135 | 3,4651668 | -0,16963776 | 0,86763496 | 0,94046736 |
| 154664_at | ABCA13 | 0,00795623 | 3,04027139 | 0,16962855 | 0,86764207 | 0,94046736 |
| 387338_at | NSUN4 | -0,03167334 | 7,52582397 | -0,16958845 | 0,86767304 | 0,94046736 |
| 10461_at | MERTK | 0,04675565 | 4,61004862 | 0,16944118 | 0,86778678 | 0,94049296 |
| 170506_at | DHX36 | -0,03498085 | 8,73890036 | -0,16939557 | 0,86782202 | 0,94049296 |
| 100506126_a | LINC00867 | 0,01696278 | 3,65877303 | 0,16936822 | 0,86784313 | 0,94049296 |
| 7475_at | WNT6 | 0,02310147 | 5,45999773 | 0,16923633 | 0,86794501 | 0,94055044 |
| 56163_at | RNF17 | 0,00915932 | 3,37875277 | 0,16863677 | 0,86840812 | 0,94099936 |
| 90355_at | C5orf30 | -0,0214264 | 7,49779947 | -0,16848478 | 0,86852553 | 0,94107364 |
| 10584_at | COLEC10 | 0,010378 | 3,71949583 | 0,16827176 | 0,86869008 | 0,94118097 |
| 8608_at | RDH16 | 0,03752911 | 5,35734094 | 0,16823005 | 0,8687223 | 0,94118097 |
| 101928896_a | LOC10192889 | -0,0157135 | 3,92527098 | -0,16814278 | 0,86878972 | 0,94120108 |
| 7392_at | USF2 | 0,01034379 | 8,79965846 | 0,16800086 | 0,86889936 | 0,94126692 |
| 148137_at | PROSER3 | 0,01180514 | 5,89490589 | 0,16763951 | 0,86917853 | 0,94151639 |
| 5923_at | RASGRF1 | 0,01076533 | 4,75435248 | 0,16733791 | 0,86941155 | 0,94171585 |
| 1539_at | CYLC2 | -0,01595096 | 3,04515427 | -0,16706537 | 0,86962213 | 0,94172806 |
| 23742_at | NPAP1 | -0,02138021 | 4,89067714 | -0,16705687 | 0,8696287 | 0,94172806 |
| 57117_at | INTS12 | -0,04170238 | 7,01961237 | -0,16703425 | 0,86964618 | 0,94172806 |
| 50616_at | IL22 | -0,00915657 | 3,71576091 | -0,16695217 | 0,86970961 | 0,94172806 |
| 146279_at | TEKT5 | 0,02456102 | 4,88779545 | 0,16694611 | 0,86971429 | 0,94172806 |
| 57716_at | PRX | 0,01855077 | 5,9723855 | 0,16694369 | 0,86971615 | 0,94172806 |
| 54529_at | ASNSD1 | 0,01741603 | 8,83364138 | 0,16684282 | 0,8697941 | 0,94175952 |
| 26032_at | SUSD5 | 0,02049031 | 4,16415845 | 0,16672817 | 0,86988269 | 0,94180251 |
| 163259_at | DENND2C | 0,02503796 | 5,17876205 | 0,16653778 | 0,87002981 | 0,94190886 |
| 8490_at | RGS5 | -0,0360613 | 5,44728302 | -0,16643273 | 0,87011099 | 0,9419438 |

| | | | | | | |
|---|---|---|---|---|---|---|
| 55845_at | BRK1 | -0,02203203 | 10,8141127 | -0,16632436 | 0,87019474 | 0,94195646 |
| 5042_at | PABPC3 | -0,01820478 | 9,88525484 | -0,16628261 | 0,870227 | 0,94195646 |
| 389124_at | IQCF5 | -0,01901382 | 4,23607179 | -0,16617243 | 0,87031215 | 0,94195646 |
| 1838_at | DTNB | 0,03506441 | 6,62823374 | 0,16615194 | 0,87032798 | 0,94195646 |
| 6324_at | SCN1B | 0,01976971 | 5,59293627 | 0,16610122 | 0,87036718 | 0,94195646 |
| 400360_at | C15orf54 | 0,01206352 | 3,1380611 | 0,16600902 | 0,87043843 | 0,94198064 |
| 10810_at | WASF3 | 0,03669609 | 6,62586326 | 0,16593517 | 0,8704955 | 0,94198949 |
| 100505875_a | LINC01088 | 0,17208429 | 6,05087401 | 0,1657467 | 0,87064116 | 0,94201512 |
| 10629_at | TAF6L | -0,01904458 | 6,92040726 | -0,16567286 | 0,87069823 | 0,94201512 |
| 1731_at | SEPT1 | -0,01299523 | 4,71471574 | -0,16564472 | 0,87071998 | 0,94201512 |
| 116444_at | GRIN3B | 0,02136764 | 5,37584459 | 0,16562274 | 0,87073697 | 0,94201512 |
| 6619_at | SNAPC3 | -0,03101704 | 8,09602685 | -0,16558815 | 0,8707637 | 0,94201512 |
| 51375_at | SNX7 | 0,03725758 | 8,39062145 | 0,1653659 | 0,87093548 | 0,94214804 |
| 101928751_a | LOC10192875 | 0,0193577 | 4,02323171 | 0,16524416 | 0,87102958 | 0,94217915 |
| 284677_at | C1orf204 | 0,02408423 | 5,00021046 | 0,16520213 | 0,87106206 | 0,94217915 |
| 400940_at | LOC400940 | -0,02109903 | 3,51447832 | -0,16498104 | 0,87123295 | 0,94231108 |
| 23576_at | DDAH1 | -0,01688735 | 7,50128972 | -0,16473841 | 0,8714205 | 0,94244565 |
| 286184_at | LINC01289 | 0,01358713 | 3,71560246 | 0,16465197 | 0,87148732 | 0,94244565 |
| 56165_at | TDRD1 | 0,12339359 | 4,42377882 | 0,1646302 | 0,87150414 | 0,94244565 |
| 1600_at | DAB1 | -0,02085939 | 4,23686853 | -0,16453471 | 0,87157797 | 0,94247257 |
| 283143_at | LINC00900 | 0,01540553 | 3,99065544 | 0,1642656 | 0,87178599 | 0,94260979 |
| 8153_at | RND2 | 0,00983815 | 4,0731899 | 0,16424396 | 0,87180273 | 0,94260979 |
| 2923_at | PDIA3 | -0,0165377 | 8,08492289 | -0,16405057 | 0,87195223 | 0,94266584 |
| 55338_at | LOC55338 | 0,01277153 | 3,18579152 | 0,1640503 | 0,87195244 | 0,94266584 |
| 26585_at | GREM1 | 0,01262208 | 3,55829 | 0,16387127 | 0,87209085 | 0,94272221 |
| 4152_at | MBD1 | -0,01951218 | 7,06547805 | -0,16369072 | 0,87223044 | 0,94272221 |
| 101929019_a | LOC10192901 | -0,0144088 | 3,48141353 | -0,16360783 | 0,87229453 | 0,94272221 |
| 100506783_a | HOXD-AS2 | -0,01782717 | 6,2121424 | -0,16359476 | 0,87230463 | 0,94272221 |
| 103091865_a | BRWD1-IT2 | 0,01918382 | 5,38075536 | 0,16355106 | 0,87233841 | 0,94272221 |
| 51290_at | ERGIC2 | 0,02925904 | 9,94058918 | 0,16354361 | 0,87234417 | 0,94272221 |
| 375189_at | PFN4 | 0,0189835 | 4,39991817 | 0,16348848 | 0,8723868 | 0,94272221 |
| 101928559_a | LINC01467 | 0,01181194 | 3,0862933 | 0,16344465 | 0,87242069 | 0,94272221 |
| 64101_at | LRRC4 | 0,02291266 | 4,39920662 | 0,16336042 | 0,87248581 | 0,94272221 |
| 10787_at | NCKAP1 | -0,02284365 | 10,829564 | -0,16328506 | 0,87254408 | 0,94272221 |
| 7122_at | CLDN5 | -0,01203884 | 5,22582291 | -0,1632209 | 0,87259369 | 0,94272221 |
| 100288413_a | ERVMER34-1 | -0,08188739 | 5,32419812 | -0,16311448 | 0,87267598 | 0,94272221 |
| 2254_at | FGF9 | -0,01088804 | 3,02007457 | -0,1631135 | 0,87267673 | 0,94272221 |
| 1455_at | CSNK1G2 | 0,01737829 | 8,62601213 | 0,1630498 | 0,87272598 | 0,94272221 |
| 103752554_a | MYCNUT | -0,01449897 | 4,06190774 | -0,16303339 | 0,87273868 | 0,94272221 |
| 100506119_a | LINC01503 | -0,05455036 | 9,20014162 | -0,16270407 | 0,87299332 | 0,94292065 |
| 55781_at | RIOK2 | -0,02690744 | 7,00985164 | -0,16266919 | 0,87302029 | 0,94292065 |
| 151531_at | UPP2 | -0,01228989 | 3,54984243 | -0,16255973 | 0,87310493 | 0,9429592 |
| 102546226_a | LOC10254622 | -0,01002062 | 2,89328545 | -0,1624596 | 0,87318236 | 0,94298996 |
| 337867_at | UBAC2 | 0,02903882 | 8,46302447 | 0,16206385 | 0,87348842 | 0,94326759 |
| 440072_at | LINC00167 | 0,01611301 | 4,4592477 | 0,16189937 | 0,87361562 | 0,94330744 |
| 126868_at | MAB21L3 | 0,07108607 | 6,4156609 | 0,1618895 | 0,87362325 | 0,94330744 |
| 30817_at | ADGRE2 | -0,02057129 | 4,68631344 | -0,16164198 | 0,87381468 | 0,94346125 |
| 2596_at | GAP43 | -0,01350029 | 3,45346117 | -0,16145493 | 0,87395935 | 0,9435429 |
| 28727_at | TRAJ28 | -0,02018679 | 6,10259362 | -0,16141755 | 0,87398826 | 0,9435429 |
| 5048_at | PAFAH1B1 | 0,01330945 | 8,13949429 | 0,16130936 | 0,87407194 | 0,94358036 |

| | | | | | | |
|---|---|---|---|---|---|---|
| 5270_at | SERPINE2 | -0,16431715 | 6,38737064 | -0,16119047 | 0,8741639 | 0,94360925 |
| 9121_at | SLC16A5 | 0,01448525 | 8,52560826 | 0,16114809 | 0,87419668 | 0,94360925 |
| 103695435_a | BBOX1-AS1 | -0,09070346 | 5,52378536 | -0,16085209 | 0,87442564 | 0,94367114 |
| 4821_at | NKX2-2 | 0,01353357 | 3,87488093 | 0,16084202 | 0,87443342 | 0,94367114 |
| 1672_at | DEFB1 | -0,11819306 | 5,0658502 | -0,16083399 | 0,87443964 | 0,94367114 |
| 6275_at | S100A4 | 0,04994624 | 9,58310019 | 0,16080085 | 0,87446527 | 0,94367114 |
| 5092_at | PCBD1 | 0,0155332 | 8,02278511 | 0,16066906 | 0,87456722 | 0,94367114 |
| 25924_at | MYRIP | -0,01156783 | 3,40629539 | -0,16063876 | 0,87459065 | 0,94367114 |
| 388011_at | LINC01550 | 0,01763281 | 3,7194144 | 0,16063063 | 0,87459694 | 0,94367114 |
| 4351_at | MPI | 0,01582932 | 7,65336111 | 0,16035251 | 0,87481208 | 0,94385041 |
| 563_at | AZGP1 | 0,01423892 | 3,40092378 | 0,16027291 | 0,87487366 | 0,94385146 |
| 10491_at | CRTAP | -0,01828343 | 8,95854394 | -0,16007771 | 0,87502467 | 0,94385146 |
| 4608_at | MYBPH | -0,02077036 | 6,0153482 | -0,16005578 | 0,87504164 | 0,94385146 |
| 282616_at | IFNL2 | 0,02456745 | 4,84024839 | 0,1600386 | 0,87505493 | 0,94385146 |
| 221883_at | HOXA11-AS | 0,01388599 | 4,13223468 | 0,16000183 | 0,87508338 | 0,94385146 |
| 100507424_a | LOC10050742 | -0,03052319 | 4,80945436 | -0,15997124 | 0,87510704 | 0,94385146 |
| 79789_at | CLMN | -0,08497769 | 5,22808553 | -0,15986689 | 0,87518777 | 0,94388568 |
| 3645_at | INSRR | -0,0145055 | 4,93214314 | -0,159587 | 0,87540431 | 0,94403466 |
| 55466_at | DNAJA4 | 0,0249578 | 9,14547133 | 0,15956164 | 0,87542393 | 0,94403466 |
| 85451_at | UNK | 0,01196732 | 7,49888114 | 0,15933794 | 0,87559702 | 0,94414649 |
| 642968_at | FAM163B | -0,01336707 | 5,25556104 | -0,15930092 | 0,87562566 | 0,94414649 |
| 3800_at | KIF5C | -0,02074833 | 5,45071061 | -0,15920747 | 0,87569796 | 0,9441716 |
| 6736_at | SRY | 0,02212162 | 3,68101776 | 0,15894467 | 0,8759013 | 0,94433798 |
| 220004_at | PPP1R32 | -0,02164497 | 6,32230705 | -0,1586721 | 0,87611221 | 0,94451251 |
| 3175_at | ONECUT1 | -0,01773412 | 5,1917851 | -0,1585672 | 0,87619338 | 0,94454716 |
| 91373_at | UAP1L1 | -0,02056759 | 5,53021042 | -0,15826569 | 0,87642671 | 0,94474582 |
| 100996624_a | LOC10099662 | 0,01270663 | 3,24502411 | 0,15813676 | 0,87652648 | 0,94480049 |
| 147081_at | CRHR1-IT1 | 0,01858707 | 4,86373084 | 0,1580037 | 0,87662946 | 0,94485862 |
| 54429_at | TAS2R5 | -0,01316507 | 4,36936127 | -0,15775101 | 0,87682502 | 0,9450099 |
| 27153_at | ZNF777 | -0,01414335 | 6,44434972 | -0,15761976 | 0,8769266 | 0,9450099 |
| 100131496_a | LOC10013149 | 0,01677384 | 4,16141738 | 0,15757258 | 0,87696311 | 0,9450099 |
| 81472_at | OR2C3 | -0,01424228 | 3,55774086 | -0,15754282 | 0,87698614 | 0,9450099 |
| 79963_at | ABCA11P | 0,03371783 | 6,04319331 | 0,15750145 | 0,87701816 | 0,9450099 |
| 115908_at | CTHRC1 | 0,02059809 | 3,91992425 | 0,15744201 | 0,87706417 | 0,9450099 |
| 5653_at | KLK6 | 0,10004838 | 6,75485688 | 0,15710316 | 0,87732643 | 0,94523962 |
| 89765_at | RSPH1 | -0,08982461 | 5,72757962 | -0,15682017 | 0,87754548 | 0,94533541 |
| 59353_at | TMEM35 | 0,01740416 | 3,82741047 | 0,15679929 | 0,87756164 | 0,94533541 |
| 3361_at | HTR5A | -0,01576803 | 4,65041587 | -0,15679808 | 0,87756257 | 0,94533541 |
| 100130581_a | LINC00910 | 0,01144358 | 4,43822904 | 0,15672888 | 0,87761614 | 0,94534025 |
| 10902_at | BRD8 | 0,01539815 | 7,37023203 | 0,15658088 | 0,8777307 | 0,94541078 |
| 6529_at | SLC6A1 | -0,0148058 | 3,67422854 | -0,15640373 | 0,87786783 | 0,94550562 |
| 396_at | ARHGDIA | -0,04076238 | 8,68429848 | -0,156307 | 0,87794271 | 0,94553341 |
| 7429_at | VIL1 | 0,02403717 | 4,24457264 | 0,15583126 | 0,878311 | 0,94577212 |
| 23371_at | TNS2 | 0,01582074 | 5,93275801 | 0,15581061 | 0,87832699 | 0,94577212 |
| 55213_at | RCBTB1 | -0,15331995 | 5,44608018 | -0,15578022 | 0,87835051 | 0,94577212 |
| 22809_at | ATF5 | -0,03699829 | 6,5013622 | -0,155767 | 0,87836075 | 0,94577212 |
| 100506489_a | LOC10050648 | -0,01469369 | 3,57757897 | -0,15568974 | 0,87842056 | 0,94578365 |
| 1160_at | CKMT2 | -0,01635806 | 4,33701418 | -0,15559767 | 0,87849185 | 0,94580754 |
| 54752_at | FNDC8 | -0,01504521 | 4,77257977 | -0,15545435 | 0,87860281 | 0,94587414 |
| 92703_at | TMEM183A | -0,03452694 | 6,53937808 | -0,15524868 | 0,87876204 | 0,94599269 |

| | | | | | | |
|---|---|---|---|---|---|---|
| 5125_at | PCSK5 | -0,02490322 | 4,62825876 | -0,15484285 | 0,87907627 | 0,94627808 |
| 129450_at | TYW5 | 0,01345232 | 6,00703744 | 0,15462951 | 0,87924147 | 0,94640301 |
| 4238_at | MFAP3 | 0,01576253 | 8,07581562 | 0,15425503 | 0,87953144 | 0,94656105 |
| 57198_at | ATP8B2 | 0,01910766 | 5,60835331 | 0,15424994 | 0,87953538 | 0,94656105 |
| 9945_at | GFPT2 | -0,0119156 | 4,53394629 | -0,15422345 | 0,8795559 | 0,94656105 |
| 526_at | ATP6V1B2 | 0,04129234 | 8,91049923 | 0,15417853 | 0,87959068 | 0,94656105 |
| 6844_at | VAMP2 | -0,01308164 | 5,74635921 | -0,15412263 | 0,87963398 | 0,94656105 |
| 100379661_a | GRIK1-AS2 | -0,01013897 | 3,47319726 | -0,15400015 | 0,87972882 | 0,94658427 |
| 26492_at | OR8G2 | -0,02531749 | 4,01890021 | -0,15396785 | 0,87975384 | 0,94658427 |
| 64418_at | TMEM168 | 0,04292744 | 7,23745831 | 0,1536045 | 0,88003522 | 0,94678009 |
| 101927870_a | LOC10192787 | 0,00897593 | 2,89188531 | 0,15349846 | 0,88011735 | 0,94678009 |
| 9214_at | FCMR | -0,02491901 | 5,18073945 | -0,15347953 | 0,88013201 | 0,94678009 |
| 101929279_a | LOC10192927 | -0,02281526 | 4,34043793 | -0,15347899 | 0,88013243 | 0,94678009 |
| 3303_at | HSPA1A | 0,27998945 | 9,00540341 | 0,1533886 | 0,88020243 | 0,94680252 |
| 91408_at | BTF3L4 | 0,02229014 | 7,53089586 | 0,15327442 | 0,88029087 | 0,94684477 |
| 23476_at | BRD4 | -0,01791027 | 8,3846159 | -0,15301338 | 0,88049305 | 0,94697332 |
| 9086_at | EIF1AY | 0,18345711 | 5,08445314 | 0,15299317 | 0,8805087 | 0,94697332 |
| 152078_at | PQLC2L | -0,03109801 | 3,73801146 | -0,15281206 | 0,88064898 | 0,94704719 |
| 1829_at | DSG2 | 0,0653458 | 9,69262127 | 0,15274821 | 0,88069843 | 0,94704719 |
| 100874253_a | LINC00853 | 0,01593981 | 3,96211086 | 0,15267188 | 0,88075756 | 0,94704719 |
| 51009_at | DERL2 | 0,01514986 | 7,687347 | 0,1526506 | 0,88077405 | 0,94704719 |
| 4247_at | MGAT2 | -0,02250024 | 8,42091063 | -0,15257143 | 0,88083537 | 0,94706026 |
| 100506083_a | LOC10050608 | -0,01842048 | 3,22081677 | -0,15244794 | 0,88093102 | 0,94706175 |
| 401551_at | WDR38 | -0,01613707 | 5,37280027 | -0,15244271 | 0,88093508 | 0,94706175 |
| 6712_at | SPTBN2 | 0,04273336 | 6,12936806 | 0,15227209 | 0,88106724 | 0,94715097 |
| 57827_at | C6orf47 | 0,01220656 | 7,84887056 | 0,15206208 | 0,88122992 | 0,9472211 |
| 3718_at | JAK3 | -0,01961394 | 4,78401258 | -0,15201072 | 0,88126971 | 0,9472211 |
| 115362_at | GBP5 | -0,0095534 | 3,49265853 | -0,15199744 | 0,88127999 | 0,9472211 |
| 9595_at | CYTIP | 0,0113643 | 3,57510389 | 0,15178959 | 0,88144101 | 0,9473413 |
| 10251_at | SPRY3 | -0,01502339 | 4,54715491 | -0,15144209 | 0,88171023 | 0,94749933 |
| 196996_at | GRAMD2 | -0,06166401 | 5,37185538 | -0,15142611 | 0,88172261 | 0,94749933 |
| 79015_at | LINC01260 | -0,01820099 | 5,68529693 | -0,1513246 | 0,88180125 | 0,94749933 |
| 100129503_a | PP13 | -0,02386198 | 5,73968798 | -0,15128607 | 0,8818311 | 0,94749933 |
| 64800_at | EFCAB6 | -0,00972148 | 3,63259216 | -0,15122869 | 0,88187555 | 0,94749933 |
| 129080_at | EMID1 | -0,01154834 | 6,01433756 | -0,15116981 | 0,88192117 | 0,94749933 |
| 100506108_a | SIX3-AS1 | 0,01940558 | 4,82878738 | 0,15115537 | 0,88193236 | 0,94749933 |
| 23370_at | ARHGEF18 | 0,0239243 | 9,29585206 | 0,15106416 | 0,88200303 | 0,94752241 |
| 23062_at | GGA2 | -0,01367326 | 7,52579957 | -0,1507916 | 0,88221421 | 0,94756977 |
| 6430_at | SRSF5 | 0,03159025 | 10,3174507 | 0,15076897 | 0,88223174 | 0,94756977 |
| 51596_at | CUTA | 0,02111381 | 10,4922044 | 0,15076498 | 0,88223483 | 0,94756977 |
| 1745_at | DLX1 | -0,07191769 | 6,28045725 | -0,15075331 | 0,88224387 | 0,94756977 |
| 100130992_a | LOC10013099 | 0,01219759 | 3,41991343 | 0,15068271 | 0,88229858 | 0,94757569 |
| 55796_at | MBNL3 | 0,02413385 | 6,15041123 | 0,15046757 | 0,88246528 | 0,94770188 |
| 22901_at | ARSG | 0,01000494 | 4,02499143 | 0,1502866 | 0,8826055 | 0,94779963 |
| 58494_at | JAM2 | -0,01176294 | 3,65558093 | -0,15020616 | 0,88266784 | 0,94780027 |
| 57634_at | EP400 | -0,01335146 | 7,42898374 | -0,15015884 | 0,8827045 | 0,94780027 |
| 101060004_a | LINC01523 | 0,0168883 | 3,2874338 | 0,14999823 | 0,88282896 | 0,94788107 |
| 128439_at | SNHG11 | -0,02341797 | 7,94682335 | -0,14981338 | 0,8829722 | 0,94798203 |
| 4810_at | NHS | -0,03156805 | 6,22956908 | -0,14961968 | 0,88312231 | 0,94808188 |
| 23534_at | TNPO3 | -0,01744126 | 9,24633348 | -0,14954968 | 0,88317656 | 0,94808188 |

| | | | | | | |
|---|---|---|---|---|---|---|
| 6599_at | SMARCC1 | 0,02628865 | 8,53668251 | 0,14950284 | 0,88321286 | 0,94808188 |
| 91948_at | LINC00923 | -0,01630742 | 4,39127804 | -0,14938817 | 0,88330173 | 0,94808842 |
| 90956_at | ADCK2 | 0,01538997 | 7,53415081 | 0,14932041 | 0,88335424 | 0,94808842 |
| 1234_at | CCR5 | 0,01575165 | 5,0987856 | 0,14929193 | 0,88337631 | 0,94808842 |
| 9570_at | GOSR2 | -0,00993472 | 6,86709493 | -0,14924095 | 0,88341582 | 0,94808842 |
| 2898_at | GRIK2 | -0,01527129 | 4,48205533 | -0,14910122 | 0,88352411 | 0,94815181 |
| 644997_at | PIK3CD-AS1 | -0,01051502 | 6,09052077 | -0,1490181 | 0,88358853 | 0,94816812 |
| 117166_at | WFIKKN1 | 0,01189378 | 6,7692692 | 0,1488817 | 0,88369425 | 0,94822084 |
| 11277_at | TREX1 | 0,01804902 | 8,03661043 | 0,14882769 | 0,88373611 | 0,94822084 |
| 100506874_a | LINC00933 | -0,02016093 | 3,98876662 | -0,14870621 | 0,88383027 | 0,94826905 |
| 5010_at | CLDN11 | 0,01424458 | 4,96986342 | 0,14836491 | 0,88409481 | 0,94845369 |
| 23251_at | KIAA1024 | -0,0168736 | 4,35180598 | -0,14835713 | 0,88410084 | 0,94845369 |
| 2567_at | GABRG3 | 0,00954832 | 3,42918515 | 0,14818827 | 0,88423173 | 0,94854128 |
| 122042_at | RXFP2 | 0,01363718 | 3,04150232 | 0,1480512 | 0,88433797 | 0,94860243 |
| 25903_at | OLFML2B | -0,01231075 | 3,94976516 | -0,14797187 | 0,88439947 | 0,94861557 |
| 79598_at | CEP97 | -0,01638787 | 5,93349326 | -0,147733 | 0,88458464 | 0,94876135 |
| 9076_at | CLDN1 | 0,06619848 | 11,0064032 | 0,14761767 | 0,88467404 | 0,94880441 |
| 84263_at | HSDL2 | 0,03269104 | 7,83631874 | 0,14714655 | 0,88503927 | 0,94909869 |
| 23510_at | KCTD2 | -0,00973641 | 8,02406529 | -0,14713661 | 0,88504697 | 0,94909869 |
| 84858_at | ZNF503 | 0,01481048 | 5,79679119 | 0,14703587 | 0,88512507 | 0,94912961 |
| 8411_at | EEA1 | 0,01251201 | 7,57259355 | 0,14658991 | 0,88547083 | 0,94938021 |
| 9399_at | STOML1 | -0,01258038 | 7,00684027 | -0,14650781 | 0,88553448 | 0,94938021 |
| 121599_at | SPIC | 0,01057606 | 2,78824407 | 0,14647798 | 0,88555761 | 0,94938021 |
| 586_at | BCAT1 | 0,18446158 | 7,59107031 | 0,14643597 | 0,88559018 | 0,94938021 |
| 5265_at | SERPINA1 | -0,01099484 | 4,28060772 | -0,1464166 | 0,8856052 | 0,94938021 |
| 9934_at | P2RY14 | -0,00941059 | 2,90690322 | -0,14626333 | 0,88572404 | 0,94940782 |
| 84719_at | LINC00260 | -0,02182756 | 4,05700611 | -0,14620065 | 0,88577264 | 0,94940782 |
| 4507_at | MTAP | -0,00856289 | 3,38460259 | -0,14609021 | 0,88585828 | 0,94940782 |
| 115265_at | DDIT4L | -0,02200998 | 4,66287862 | -0,14597843 | 0,88594495 | 0,94940782 |
| 1238_at | ACKR2 | -0,01868973 | 4,59806157 | -0,14597149 | 0,88595033 | 0,94940782 |
| 57223_at | PPP4R3B | -0,01436564 | 8,794733 | -0,14592205 | 0,88598867 | 0,94940782 |
| 10943_at | MSL3 | 0,00923106 | 6,80496365 | 0,14589439 | 0,88601012 | 0,94940782 |
| 55892_at | MYNN | 0,01577914 | 7,35573977 | 0,14587487 | 0,88602525 | 0,94940782 |
| 55638_at | SYBU | 0,11604835 | 5,75880002 | 0,14576266 | 0,88611227 | 0,94940817 |
| 57679_at | ALS2 | -0,00992552 | 6,46626748 | -0,14574734 | 0,88612415 | 0,94940817 |
| 5764_at | PTN | -0,01161496 | 3,76404794 | -0,14543932 | 0,886363 | 0,94955286 |
| 2782_at | GNB1 | -0,02098445 | 10,8601595 | -0,1454122 | 0,88638403 | 0,94955286 |
| 55959_at | SULF2 | -0,06448444 | 6,55160514 | -0,14538248 | 0,88640707 | 0,94955286 |
| 147685_at | C19orf18 | -0,02091702 | 4,34834226 | -0,14509559 | 0,88662956 | 0,949728 |
| 23200_at | ATP11B | -0,01915555 | 8,18712732 | -0,14495544 | 0,88673825 | 0,949728 |
| 29114_at | TAGLN3 | -0,01145861 | 4,34769321 | -0,14492502 | 0,88676184 | 0,949728 |
| 145270_at | PRIMA1 | -0,03194326 | 5,47675552 | -0,14491736 | 0,88676778 | 0,949728 |
| 54749_at | EPDR1 | -0,14048109 | 8,06139912 | -0,14474104 | 0,88690452 | 0,94973089 |
| 1852_at | DUSP9 | 0,01696386 | 7,66165996 | 0,14472603 | 0,88691617 | 0,94973089 |
| 2888_at | GRB14 | 0,04274886 | 4,015037 | 0,14472315 | 0,8869184 | 0,94973089 |
| 148_at | ADRA1A | -0,00869188 | 3,34478813 | -0,14441555 | 0,88715697 | 0,94993355 |
| 1012_at | CDH13 | 0,02281127 | 3,76189694 | 0,14432992 | 0,88722338 | 0,94995187 |
| 642852_at | LOC642852 | -0,01475448 | 7,02656257 | -0,1442181 | 0,88731011 | 0,94995452 |
| 6899_at | TBX1 | 0,02389726 | 4,9001391 | 0,14419956 | 0,88732449 | 0,94995452 |
| 116541_at | MRPL54 | -0,00929657 | 8,35896838 | -0,14412896 | 0,88737925 | 0,94996035 |

| | | | | | | |
|---|---|---|---|---|---|---|
| 64145_at | RBSN | 0,01146557 | 6,49295291 | 0,14405742 | 0,88743473 | 0,94996695 |
| 2835_at | GPR12 | -0,01453203 | 4,42113003 | -0,14384813 | 0,88759707 | 0,95008793 |
| 777_at | CACNA1E | -0,01140986 | 3,60121128 | -0,14365605 | 0,88774606 | 0,95019461 |
| 5798_at | PTPRN | 0,02006255 | 5,13215304 | 0,14357777 | 0,88780678 | 0,9502068 |
| 339977_at | LRRC66 | -0,00866065 | 3,10920104 | -0,1434346 | 0,88791784 | 0,95027286 |
| 79970_at | ZNF767P | 0,03653915 | 6,51066991 | 0,14325899 | 0,88805406 | 0,95031579 |
| 4921_at | DDR2 | 0,01175929 | 4,2733704 | 0,14321231 | 0,88809027 | 0,95031579 |
| 4081_at | MAB21L1 | 0,04121735 | 3,94924684 | 0,14315386 | 0,88813561 | 0,95031579 |
| 195827_at | AAED1 | -0,04723311 | 7,36141279 | -0,14312851 | 0,88815528 | 0,95031579 |
| 79770_at | TXNDC15 | -0,01227103 | 8,62066896 | -0,14279853 | 0,88841127 | 0,95053689 |
| 643210_at | EHMT1-IT1 | 0,01398746 | 5,02730172 | 0,14271991 | 0,88847226 | 0,95054248 |
| 57664_at | PLEKHA4 | -0,01222878 | 5,02331793 | -0,1426642 | 0,88851548 | 0,95054248 |
| 284379_at | LOC284379 | 0,01440853 | 4,28970981 | 0,14260099 | 0,88856452 | 0,95054248 |
| 92610_at | TIFA | 0,05835108 | 7,67585687 | 0,14219463 | 0,88887979 | 0,95074964 |
| 137902_at | PXDNL | 0,01003972 | 3,54390857 | 0,14216293 | 0,88890438 | 0,95074964 |
| 282973_at | JAKMIP3 | -0,02225052 | 5,39150967 | -0,14216053 | 0,88890624 | 0,95074964 |
| 7579_at | ZSCAN20 | 0,0111999 | 4,65816355 | 0,14172281 | 0,88924586 | 0,95106008 |
| 27202_at | C5AR2 | 0,01056398 | 3,84085123 | 0,14147495 | 0,88943819 | 0,95121296 |
| 4233_at | MET | -0,02406898 | 8,31244756 | -0,14136996 | 0,88951965 | 0,95124727 |
| 254887_at | ZDHHC23 | -0,03131028 | 6,36716046 | -0,14128135 | 0,88958842 | 0,95126799 |
| 338657_at | CCDC84 | 0,03833181 | 7,52119536 | 0,1410898 | 0,88973705 | 0,95137412 |
| 79958_at | DENND1C | -0,02331152 | 4,51615679 | -0,14091525 | 0,8898725 | 0,95143533 |
| 3460_at | IFNGR2 | -0,0188367 | 9,28902333 | -0,14088375 | 0,88989695 | 0,95143533 |
| 127795_at | C1orf87 | -0,01063817 | 3,40350406 | -0,14082509 | 0,88994247 | 0,95143533 |
| 53834_at | FGFRL1 | 0,00980356 | 6,42294049 | 0,14073963 | 0,89000878 | 0,95143557 |
| 340481_at | ZDHHC21 | -0,01433184 | 5,71855786 | -0,1406452 | 0,89008207 | 0,95143557 |
| 1154_at | CISH | -0,01179324 | 6,48344082 | -0,14063386 | 0,89009087 | 0,95143557 |
| 9757_at | KMT2B | -0,01060951 | 6,38977613 | -0,14041735 | 0,89025889 | 0,95156236 |
| 28562_at | TRBV25-1 | 0,01237818 | 2,84969908 | 0,14020595 | 0,89042295 | 0,95168142 |
| 11279_at | KLF8 | 0,00793789 | 4,8631168 | 0,14010118 | 0,89050426 | 0,95168142 |
| 101928622_a | LOC10192862 | -0,01371708 | 3,02760969 | -0,14008285 | 0,89051849 | 0,95168142 |
| 85301_at | COL27A1 | -0,02446343 | 6,41161361 | -0,13973177 | 0,89079097 | 0,95191981 |
| 440119_at | FZD10-AS1 | 0,01572971 | 5,18371936 | 0,13961458 | 0,89088193 | 0,95195189 |
| 2274_at | FHL2 | -0,02567394 | 9,10130959 | -0,13945445 | 0,89100621 | 0,95195189 |
| 79444_at | BIRC7 | 0,01288843 | 5,2312252 | 0,13937047 | 0,8910714 | 0,95195189 |
| 100505666_a | LOC10050566 | -0,02048385 | 4,83506415 | -0,13934687 | 0,89108972 | 0,95195189 |
| 130540_at | ALS2CR12 | -0,00966452 | 3,57433936 | -0,13920848 | 0,89119714 | 0,95195189 |
| 6236_at | RRAD | 0,0082687 | 4,17087424 | 0,13917024 | 0,89122681 | 0,95195189 |
| 51108_at | METTL9 | -0,00898623 | 8,32067996 | -0,13912137 | 0,89126475 | 0,95195189 |
| 54941_at | RNF125 | -0,01122487 | 3,44545502 | -0,13904306 | 0,89132554 | 0,95195189 |
| 492_at | ATP2B3 | -0,01114382 | 4,53594944 | -0,13892002 | 0,89142104 | 0,95195189 |
| 145474_at | LOC145474 | 0,01195969 | 3,37010228 | 0,13890163 | 0,89143532 | 0,95195189 |
| 200634_at | KRTCAP3 | 0,08210743 | 7,49575121 | 0,13889704 | 0,89143889 | 0,95195189 |
| 55255_at | WDR41 | 0,04761012 | 7,67120094 | 0,13888656 | 0,89144701 | 0,95195189 |
| 153643_at | FAM81B | 0,01379969 | 3,16780935 | 0,13880137 | 0,89151314 | 0,95195189 |
| 158880_at | USP51 | 0,01876709 | 4,64793103 | 0,13869107 | 0,89159877 | 0,95195189 |
| 5937_at | RBMS1 | 0,02226715 | 9,84816947 | 0,13862462 | 0,89165035 | 0,95195189 |
| 285672_at | SREK1IP1 | -0,03531655 | 7,13333232 | -0,13862191 | 0,89165245 | 0,95195189 |
| 29993_at | PACSIN1 | -0,01173505 | 5,39776495 | -0,13861076 | 0,89166111 | 0,95195189 |
| 285941_at | C7orf71 | 0,01695008 | 3,44360639 | 0,13841754 | 0,89181111 | 0,95201218 |

| | | | | | | |
|---|---|---|---|---|---|---|
| 57720_at | GPR107 | -0,009781 | 8,07109673 | -0,13835289 | 0,89186129 | 0,95201218 |
| 3485_at | IGFBP2 | -0,03244777 | 7,64548245 | -0,13834702 | 0,89186585 | 0,95201218 |
| 143684_at | FAM76B | -0,01562084 | 6,41759262 | -0,13814467 | 0,89202293 | 0,9521271 |
| 101927067_a | LINC01607 | -0,00955366 | 3,38520293 | -0,13783942 | 0,89225991 | 0,95232728 |
| 57168_at | ASPHD2 | 0,03273743 | 5,37815801 | 0,13766772 | 0,89239322 | 0,95239684 |
| 79626_at | TNFAIP8L2 | 0,01307037 | 4,79002423 | 0,1376281 | 0,89242398 | 0,95239684 |
| 285349_at | ZNF660 | 0,00805023 | 3,04725905 | 0,1371616 | 0,89278618 | 0,9527306 |
| 51343_at | FZR1 | 0,01831315 | 6,9822278 | 0,13663145 | 0,89319782 | 0,95311709 |
| 864_at | RUNX3 | -0,01086339 | 5,10914906 | -0,13648274 | 0,8933133 | 0,95318751 |
| 5641_at | LGMN | 0,0343171 | 10,693439 | 0,13639406 | 0,89338215 | 0,95320818 |
| 8755_at | ADAM6 | -0,01410873 | 3,98578339 | -0,13613666 | 0,89358204 | 0,95336864 |
| 83999_at | KREMEN1 | -0,01481173 | 6,86137593 | -0,13606627 | 0,8936367 | 0,95337415 |
| 126536_at | LINC00661 | 0,01330994 | 4,23241654 | 0,13589894 | 0,89376664 | 0,95345998 |
| 84236_at | RHBDD1 | 0,01264264 | 7,27726716 | 0,13582034 | 0,89382768 | 0,95347229 |
| 10970_at | CKAP4 | -0,03765817 | 9,38339664 | -0,1356995 | 0,89392153 | 0,9534934 |
| 116093_at | DIRC1 | -0,01459552 | 3,06294958 | -0,13566738 | 0,89394647 | 0,9534934 |
| 8521_at | GCM1 | -0,01542806 | 3,55279557 | -0,13559084 | 0,89400591 | 0,95350401 |
| 339593_at | LOC339593 | -0,0124915 | 3,58395967 | -0,13549295 | 0,89408194 | 0,9535323 |
| 158866_at | ZDHHC15 | 0,00826303 | 2,8520994 | 0,13531756 | 0,89421815 | 0,95354534 |
| 11176_at | BAZ2A | -0,02440231 | 7,33839695 | -0,13529378 | 0,89423662 | 0,95354534 |
| 80114_at | BICC1 | 0,00874657 | 3,40205859 | 0,13528599 | 0,89424267 | 0,95354534 |
| 105371932_a | LOC10537193 | 0,02444269 | 4,59657224 | 0,13521961 | 0,89429423 | 0,95354753 |
| 728039_at | SSR4P1 | 0,01552969 | 4,97850359 | 0,13509416 | 0,89439166 | 0,95358706 |
| 81578_at | COL21A1 | 0,01064588 | 3,36749425 | 0,13497186 | 0,89448665 | 0,95358706 |
| 80309_at | SPHKAP | -0,01233487 | 3,54840644 | -0,13496793 | 0,8944897 | 0,95358706 |
| 26297_at | SERGEF | -0,01466947 | 6,61497679 | -0,13491693 | 0,89452931 | 0,95358706 |
| 339804_at | C2orf74 | 0,01481564 | 4,01607712 | 0,13483046 | 0,89459647 | 0,95360588 |
| 100507419_a | MMP25-AS1 | 0,01663322 | 5,75094024 | 0,13470046 | 0,89469744 | 0,95366073 |
| 157273_at | LOC157273 | 0,01077289 | 3,13468928 | 0,13437193 | 0,89495262 | 0,95387995 |
| 54436_at | SH3TC1 | 0,01729724 | 6,7903304 | 0,1341399 | 0,89513285 | 0,95401926 |
| 2245_at | FGD1 | 0,0196162 | 6,3589283 | 0,13406358 | 0,89519214 | 0,95402967 |
| 7145_at | TNS1 | 0,02150036 | 5,96776457 | 0,13382854 | 0,89537472 | 0,95417146 |
| 84239_at | ATP13A4 | -0,00843251 | 3,57258945 | -0,13367305 | 0,89549551 | 0,95417317 |
| 4291_at | MLF1 | -0,03106441 | 6,36750417 | -0,13365689 | 0,89550806 | 0,95417317 |
| 3439_at | IFNA1 | 0,00946509 | 2,61240663 | 0,13363518 | 0,89552493 | 0,95417317 |
| 83988_at | NCALD | 0,01088766 | 3,73184026 | 0,13324681 | 0,89582663 | 0,95444184 |
| 55234_at | SMU1 | 0,02088207 | 6,43611082 | 0,13312779 | 0,8959191 | 0,95448756 |
| 149297_at | FAM78B | 0,01395919 | 4,14759131 | 0,13303247 | 0,89599315 | 0,95451366 |
| 130013_at | ACMSD | 0,01326069 | 3,42506633 | 0,13289313 | 0,89610141 | 0,9545762 |
| 79016_at | DDA1 | 0,00987309 | 7,71425179 | 0,13278054 | 0,89618888 | 0,95461659 |
| 94026_at | POM121L2 | -0,01069181 | 4,94425466 | -0,13271071 | 0,89624313 | 0,95462159 |
| 7423_at | VEGFB | -0,02158326 | 7,20292604 | -0,13263714 | 0,8963003 | 0,95462969 |
| 10134_at | BCAP31 | -0,02064591 | 11,4093962 | -0,13239263 | 0,89649027 | 0,95477924 |
| 100130776_a | AGAP2-AS1 | 0,01937707 | 5,87346974 | 0,13209615 | 0,89672063 | 0,95497177 |
| 1179_at | CLCA1 | -0,0094809 | 3,82262697 | -0,13203185 | 0,89677059 | 0,95497219 |
| 129642_at | MBOAT2 | -0,03580478 | 7,93600705 | -0,13190852 | 0,89686642 | 0,95500871 |
| 64421_at | DCLRE1C | -0,02319021 | 6,09725496 | -0,13179225 | 0,89695677 | 0,95500871 |
| 10367_at | MICU1 | 0,0200036 | 7,82097755 | 0,13178414 | 0,89696307 | 0,95500871 |
| 55924_at | FAM212B | 0,0117093 | 6,2287437 | 0,13172342 | 0,89701025 | 0,95500871 |
| 83598_at | LRRC2-AS1 | -0,01687263 | 3,92665543 | -0,13165345 | 0,89706462 | 0,95500871 |

| | | | | | | |
|---|---|---|---|---|---|---|
| 148523_at | CIART | -0,00940095 | 4,57132218 | -0,13155266 | 0,89714294 | 0,95500871 |
| 5716_at | PSMD10 | 0,01103389 | 10,1280444 | 0,13148386 | 0,8971964 | 0,95500871 |
| 101241892_a | NPTN-IT1 | -0,01854397 | 6,61466036 | -0,13147729 | 0,89720151 | 0,95500871 |
| 4018_at | LPA | -0,0109709 | 3,41892093 | -0,13138623 | 0,89727227 | 0,95503125 |
| 8291_at | DYSF | -0,01726771 | 6,1649758 | -0,13095646 | 0,89760625 | 0,95533395 |
| 3497_at | IGHE | -0,08317626 | 5,61846778 | -0,13066394 | 0,89783358 | 0,95547537 |
| 2313_at | FLI1 | 0,00842289 | 3,42020297 | 0,13065783 | 0,89783833 | 0,95547537 |
| 79804_at | HAND2-AS1 | 0,01142895 | 3,2066742 | 0,13045587 | 0,89799529 | 0,95554422 |
| 26064_at | RAI14 | -0,02528034 | 9,44714967 | -0,13044693 | 0,89800223 | 0,95554422 |
| 26145_at | IRF2BP1 | 0,00738744 | 6,97406147 | 0,13022026 | 0,89817839 | 0,95567888 |
| 5774_at | PTPN3 | 0,01520473 | 7,18117363 | 0,12993502 | 0,8984001 | 0,95575226 |
| 8690_at | JRKL | -0,01094589 | 7,58583968 | -0,12993494 | 0,89840016 | 0,95575226 |
| 84675_at | TRIM55 | 0,00959964 | 3,53589401 | 0,12990289 | 0,89842506 | 0,95575226 |
| 642587_at | MIR205HG | -0,04770969 | 10,961764 | -0,12984161 | 0,89847269 | 0,95575226 |
| 28442_at | IGHV3-23 | 0,01688607 | 5,10963896 | 0,12981234 | 0,89849544 | 0,95575226 |
| 389792_at | IER5L | -0,02715932 | 7,40570214 | -0,1297086 | 0,89857608 | 0,95578526 |
| 255798_at | SMCO1 | 0,01185858 | 3,40974183 | 0,12963393 | 0,89863411 | 0,95579421 |
| 2783_at | GNB2 | -0,01759597 | 10,3634912 | -0,12936819 | 0,89884067 | 0,95596112 |
| 7499_at | XG | -0,02297023 | 5,80054518 | -0,12922573 | 0,89895141 | 0,95602611 |
| 11249_at | NXPH2 | -0,00814851 | 3,70343146 | -0,12888587 | 0,89921559 | 0,95625428 |
| 4091_at | SMAD6 | -0,02777355 | 6,89079853 | -0,12865266 | 0,89939688 | 0,95631779 |
| 55652_at | SLC48A1 | 0,02772662 | 7,56083276 | 0,12853065 | 0,89949173 | 0,95631779 |
| 283981_at | LINC00685 | 0,02771694 | 6,1469708 | 0,12848894 | 0,89952415 | 0,95631779 |
| 6539_at | SLC6A12 | 0,01897409 | 4,72144739 | 0,12842792 | 0,8995716 | 0,95631779 |
| 80320_at | SP6 | -0,04303392 | 6,92809368 | -0,12842377 | 0,89957482 | 0,95631779 |
| 51011_at | FAHD2A | 0,01673592 | 5,91480534 | 0,12841474 | 0,89958184 | 0,95631779 |
| 79722_at | ANKRD55 | -0,00806927 | 3,57486339 | -0,12834692 | 0,89963456 | 0,95631779 |
| 1469_at | CST1 | 0,13481437 | 5,96197126 | 0,1282638 | 0,89969919 | 0,95631779 |
| 10201_at | NME6 | -0,01286508 | 7,20791287 | -0,12822675 | 0,89972799 | 0,95631779 |
| 57576_at | KIF17 | -0,01417216 | 5,82986972 | -0,12817042 | 0,89977177 | 0,95631779 |
| 283507_at | SUGT1P3 | 0,00833234 | 3,01292722 | 0,12790173 | 0,89998067 | 0,95646636 |
| 79170_at | PRR15L | 0,04549735 | 5,09992882 | 0,12785826 | 0,90001447 | 0,95646636 |
| 4217_at | MAP3K5 | 0,03085505 | 8,98017455 | 0,12779902 | 0,90006052 | 0,95646636 |
| 440026_at | TMEM41B | 0,02963417 | 8,61399623 | 0,12772082 | 0,90012132 | 0,95647821 |
| 100996634_a | LOC10099663 | -0,01194488 | 3,94927298 | -0,12750052 | 0,9002926 | 0,95660745 |
| 647107_at | LINC01192 | 0,01324294 | 3,38347757 | 0,12739812 | 0,90037222 | 0,95661463 |
| 6881_at | TAF10 | 0,02281076 | 10,2962987 | 0,12736408 | 0,90039868 | 0,95661463 |
| 642946_at | FLVCR1-AS1 | -0,06359412 | 6,63939861 | -0,12713615 | 0,9005759 | 0,95675014 |
| 101929224_a | LOC10192922 | -0,02551445 | 4,52175031 | -0,12702348 | 0,90066351 | 0,9567556 |
| 646576_at | HHIP-AS1 | -0,01014289 | 4,95046566 | -0,12700179 | 0,90068038 | 0,9567556 |
| 257218_at | SHPRH | 0,0149569 | 7,18096169 | 0,12673266 | 0,90088964 | 0,95692513 |
| 54937_at | SOHLH2 | 0,01122746 | 4,22527168 | 0,12663653 | 0,9009644 | 0,9569432 |
| 11228_at | RASSF8 | -0,00964988 | 5,60482645 | -0,12656214 | 0,90102224 | 0,9569432 |
| 23473_at | CAPN7 | 0,02308689 | 8,37989729 | 0,12651704 | 0,9010573 | 0,9569432 |
| 9532_at | BAG2 | -0,13115306 | 6,29153507 | -0,12644428 | 0,90111389 | 0,9569432 |
| 94086_at | HSPB9 | -0,01794776 | 4,17777684 | -0,12639019 | 0,90115595 | 0,9569432 |
| 100131814_a | LINC00271 | -0,01645616 | 3,79211863 | -0,12632746 | 0,90120473 | 0,9569432 |
| 90204_at | ZSWIM1 | -0,00730088 | 8,40651752 | -0,12620308 | 0,90130145 | 0,95699315 |
| 127124_at | ATP6V1G3 | -0,01468605 | 4,10599866 | -0,1260767 | 0,90139972 | 0,95704475 |
| 2101_at | ESRRA | 0,01032561 | 7,38215276 | 0,12592559 | 0,90151724 | 0,95711676 |

| | | | | | | |
|---|---|---|---|---|---|---|
| 120224_at | TMEM45B | -0,00962504 | 4,31030749 | -0,12581083 | 0,90160649 | 0,95715876 |
| 342945_at | ZSCAN22 | -0,01976078 | 6,49740347 | -0,12572126 | 0,90167614 | 0,95717996 |
| 3790_at | KCNS3 | 0,01618244 | 7,93888571 | 0,12542025 | 0,90191024 | 0,9573757 |
| 9553_at | MRPL33 | 0,01366784 | 9,02254319 | 0,12511471 | 0,90214787 | 0,95757518 |
| 124565_at | SLC38A10 | 0,00827462 | 7,56689805 | 0,12504414 | 0,90220275 | 0,95758067 |
| 9950_at | GOLGA5 | -0,01743635 | 9,36230108 | -0,12478068 | 0,90240767 | 0,95774539 |
| 3805_at | KIR2DL4 | -0,00834624 | 4,57390467 | -0,12455061 | 0,90258661 | 0,95776404 |
| 101929319_a | LOC10192931 | -0,01486196 | 3,40730286 | -0,12453323 | 0,90260013 | 0,95776404 |
| 9748_at | SLK | -0,02852622 | 9,04521551 | -0,12452169 | 0,90260911 | 0,95776404 |
| 11168_at | PSIP1 | 0,13982473 | 6,98917612 | 0,12450239 | 0,90262412 | 0,95776404 |
| 8501_at | SLC43A1 | 0,01216926 | 4,79079956 | 0,124401 | 0,90270298 | 0,95779496 |
| 9099_at | USP2 | -0,00927433 | 4,8231815 | -0,12419248 | 0,90286517 | 0,95784738 |
| 196528_at | ARID2 | -0,00873378 | 7,94193011 | -0,12417269 | 0,90288057 | 0,95784738 |
| 8406_at | SRPX | 0,06015107 | 5,77740774 | 0,1241457 | 0,90290156 | 0,95784738 |
| #N/A | CTD-2297D10 | -0,01583348 | 4,19822963 | -0,12393908 | 0,90306228 | 0,95790984 |
| 676_at | BRDT | 0,01120937 | 3,32788893 | 0,1239288 | 0,90307028 | 0,95790984 |
| 25961_at | NUDT13 | 0,04905307 | 4,43440054 | 0,12385628 | 0,9031267 | 0,95790984 |
| 338579_at | ANKRD30BP3 | -0,01175849 | 3,30278204 | -0,12379716 | 0,90317268 | 0,95790984 |
| 23015_at | GOLGA8A | -0,07222847 | 7,61720926 | -0,12375037 | 0,90320908 | 0,95790984 |
| 871_at | SERPINH1 | 0,09453982 | 9,829232 | 0,12353591 | 0,90337591 | 0,95803403 |
| 220164_at | DOK6 | 0,00789953 | 3,36662134 | 0,12346287 | 0,90343273 | 0,95804154 |
| 100506827_a | LVCAT1 | 0,01194601 | 3,11403409 | 0,12330052 | 0,90355902 | 0,95812273 |
| 64778_at | FNDC3B | 0,01661082 | 10,0680582 | 0,12313304 | 0,90368931 | 0,95820814 |
| 5600_at | MAPK11 | -0,02630151 | 6,05851589 | -0,12306681 | 0,90374084 | 0,95821003 |
| 83853_at | ROPN1L | 0,00937354 | 4,53527521 | 0,1229717 | 0,90381483 | 0,95823574 |
| 100132735_a | LOC10013273 | -0,01185264 | 3,03429349 | -0,12282694 | 0,90392745 | 0,95824293 |
| 9388_at | LIPG | -0,1031769 | 7,33909083 | -0,12269771 | 0,90402799 | 0,95824293 |
| 116328_at | C8orf34 | 0,0084379 | 3,30730922 | 0,12269681 | 0,90402869 | 0,95824293 |
| 9073_at | CLDN8 | 0,13364331 | 5,1119006 | 0,12268452 | 0,90403825 | 0,95824293 |
| 139170_at | DCAF12L1 | 0,01132034 | 4,23487797 | 0,12264328 | 0,90407034 | 0,95824293 |
| 2903_at | GRIN2A | -0,0099445 | 3,51573628 | -0,12227466 | 0,90435713 | 0,95849416 |
| 51807_at | TUBA8 | -0,01184196 | 5,15716628 | -0,1222004 | 0,90441491 | 0,95850266 |
| 57325_at | CSRP2BP | -0,01242433 | 6,01962351 | -0,12151252 | 0,90495014 | 0,95895601 |
| 79574_at | EPS8L3 | 0,01539825 | 5,57351268 | 0,12143013 | 0,90501425 | 0,95895601 |
| 2272_at | FHIT | 0,01063474 | 4,04628362 | 0,12141331 | 0,90502733 | 0,95895601 |
| 140686_at | WFDC3 | -0,00950946 | 4,89346079 | -0,12139472 | 0,9050418 | 0,95895601 |
| 646_at | BNC1 | -0,00876233 | 3,22451117 | -0,12079302 | 0,90551002 | 0,95938052 |
| 25806_at | VAX2 | 0,01171673 | 5,17952386 | 0,12073986 | 0,90555139 | 0,95938052 |
| 100287902_a | LGALS8-AS1 | -0,0154187 | 5,3870399 | -0,12068785 | 0,90559186 | 0,95938052 |
| 56833_at | SLAMF8 | -0,01272624 | 4,13439375 | -0,1203556 | 0,90585042 | 0,95957563 |
| 25794_at | FSCN2 | 0,01099826 | 5,07859219 | 0,12032318 | 0,90587566 | 0,95957563 |
| 26138_at | LINC00588 | 0,00909583 | 3,49137159 | 0,12015248 | 0,9060085 | 0,95959287 |
| 56997_at | ADCK3 | 0,00943139 | 6,82625282 | 0,12010751 | 0,9060435 | 0,95959287 |
| 84651_at | SPINK7 | -0,01486658 | 3,91728576 | -0,1200712 | 0,90607176 | 0,95959287 |
| 79868_at | ALG13 | -0,01209397 | 6,85255497 | -0,12004623 | 0,90609119 | 0,95959287 |
| 149111_at | CNIH3 | -0,00882913 | 4,03818433 | -0,11987683 | 0,90622303 | 0,95967973 |
| 26086_at | GPSM1 | 0,01275173 | 6,80419252 | 0,11973717 | 0,90633173 | 0,95974208 |
| 5050_at | PAFAH1B3 | 0,01323576 | 9,64097122 | 0,11946493 | 0,90654362 | 0,95991368 |
| 25956_at | SEC31B | 0,02463605 | 5,23099147 | 0,11935086 | 0,9066324 | 0,95995492 |
| 27342_at | RABGEF1 | -0,01426089 | 7,70717529 | -0,11902902 | 0,90688291 | 0,96016739 |

| | | | | | | |
|---|---|---|---|---|---|---|
| 79650_at | USB1 | -0,01848203 | 7,63230446 | -0,11895421 | 0,90694114 | 0,96017626 |
| 84750_at | FUT10 | 0,01013986 | 6,28966168 | 0,1187967 | 0,90706374 | 0,96025329 |
| 6425_at | SFRP5 | -0,01116153 | 4,73278375 | -0,11865038 | 0,90717764 | 0,96032109 |
| 83481_at | EPPK1 | 0,04046819 | 8,71419636 | 0,1184775 | 0,90731221 | 0,96039909 |
| 10507_at | SEMA4D | -0,01048562 | 5,64957857 | -0,11842762 | 0,90735103 | 0,96039909 |
| 150381_at | PRR34-AS1 | 0,06700462 | 5,98441816 | 0,11825931 | 0,90748205 | 0,96048499 |
| 6049_at | RNF6 | -0,01195468 | 6,23087179 | -0,11816084 | 0,9075587 | 0,96051334 |
| 100628315_a | DNM3OS | -0,00905961 | 3,37246527 | -0,11805354 | 0,90764223 | 0,96054897 |
| 9229_at | DLGAP1 | 0,00589599 | 3,67910066 | 0,11797969 | 0,90769972 | 0,96055704 |
| 254778_at | C8orf46 | 0,00672273 | 3,60104436 | 0,1176834 | 0,90793038 | 0,96074835 |
| 113829_at | SLC35A4 | 0,01493497 | 9,29573833 | 0,11752504 | 0,90805367 | 0,96082603 |
| 101929745_a | LOC10192974 | -0,01016859 | 3,117886 | -0,1174239 | 0,9081324 | 0,96082725 |
| 730101_at | LOC730101 | 0,05221961 | 5,49865817 | 0,11739541 | 0,90815459 | 0,96082725 |
| 2041_at | EPHA1 | 0,00764172 | 8,33547549 | 0,11725976 | 0,90826019 | 0,96088621 |
| 100129461_a | LOC10012946 | 0,01031193 | 5,00141503 | 0,11701505 | 0,90845071 | 0,96103288 |
| 55140_at | ELP3 | 0,01692945 | 7,97989642 | 0,11695354 | 0,90849861 | 0,96103288 |
| 3748_at | KCNC3 | -0,01020805 | 5,37912525 | -0,11688548 | 0,90855159 | 0,96103615 |
| 196403_at | DTX3 | -0,01088618 | 5,04368695 | -0,11676671 | 0,90864406 | 0,96105168 |
| 150353_at | DNAJB7 | 0,00996247 | 3,11695861 | 0,11673847 | 0,90866605 | 0,96105168 |
| 8808_at | IL1RL2 | -0,01219313 | 4,13706671 | -0,11635002 | 0,9089685 | 0,96131878 |
| 84869_at | CBR4 | 0,01358977 | 7,43744409 | 0,11597373 | 0,9092615 | 0,96155096 |
| 11235_at | PDCD10 | -0,01577397 | 9,93692333 | -0,11593987 | 0,90928787 | 0,96155096 |
| 9087_at | TMSB4Y | 0,02058359 | 6,08716399 | 0,11576006 | 0,90942788 | 0,96164623 |
| 100996251_a | LOC10099625 | 0,01208333 | 3,4145176 | 0,11558963 | 0,90956059 | 0,96167608 |
| 3443_at | IFNA6 | -0,01328698 | 3,5382185 | -0,11556073 | 0,9095831 | 0,96167608 |
| 57209_at | ZNF248 | 0,02869656 | 5,41494741 | 0,11553147 | 0,90960588 | 0,96167608 |
| 286554_at | BCORP1 | -0,00993617 | 3,30951006 | -0,11527803 | 0,90980324 | 0,96183194 |
| 114885_at | OSBPL11 | 0,01014226 | 8,02174015 | 0,11517458 | 0,90988381 | 0,96186336 |
| 9699_at | RIMS2 | 0,02426435 | 4,59369383 | 0,11504408 | 0,90998543 | 0,96186336 |
| 80177_at | MYCT1 | 0,01686173 | 3,87345395 | 0,11492707 | 0,91007655 | 0,96186336 |
| 79607_at | FAM118B | 0,0239142 | 7,67262707 | 0,11487746 | 0,91011519 | 0,96186336 |
| 53336_at | CPXCR1 | 0,00588834 | 2,93968395 | 0,11485505 | 0,91013264 | 0,96186336 |
| 150350_at | ENTHD1 | 0,00708703 | 2,9704903 | 0,11481893 | 0,91016077 | 0,96186336 |
| 3567_at | IL5 | -0,00790423 | 2,92255896 | -0,11471678 | 0,91024032 | 0,96186336 |
| 84759_at | PCGF1 | 0,03382524 | 7,45584347 | 0,11470425 | 0,91025008 | 0,96186336 |
| 83852_at | SETDB2 | -0,04085455 | 4,02684337 | -0,1146628 | 0,91028236 | 0,96186336 |
| 7140_at | TNNT3 | 0,02280143 | 5,87711599 | 0,11449276 | 0,91041478 | 0,96195052 |
| 54918_at | CMTM6 | 0,01279771 | 10,3579925 | 0,11434049 | 0,91053338 | 0,96196003 |
| 126328_at | NDUFA11 | -0,01539491 | 9,68233136 | -0,11428253 | 0,91057852 | 0,96196003 |
| 5906_at | RAP1A | 0,04103825 | 6,19576902 | 0,11425447 | 0,91060037 | 0,96196003 |
| 11012_at | KLK11 | -0,01326629 | 5,04921776 | -0,1141693 | 0,9106667 | 0,96196003 |
| 400120_at | SERTM1 | 0,00758232 | 3,74396005 | 0,11416062 | 0,91067347 | 0,96196003 |
| 400548_at | LOC400548 | 0,01145768 | 4,97991194 | 0,11404103 | 0,91076661 | 0,96196921 |
| 10959_at | TMED2 | 0,00920569 | 11,7933051 | 0,11402122 | 0,91078204 | 0,96196921 |
| 57631_at | LRCH2 | -0,08030794 | 5,05341346 | -0,11371447 | 0,91102096 | 0,9621688 |
| 101928728_a | LOC10192872 | 0,01509561 | 4,30088577 | 0,1135849 | 0,91112187 | 0,96218457 |
| 101929528_a | LOC10192952 | -0,00911924 | 3,45350722 | -0,11356704 | 0,91113579 | 0,96218457 |
| 140883_at | ZNF280B | 0,01499876 | 4,6665509 | 0,11343892 | 0,91123558 | 0,9622177 |
| 9537_at | TP53I11 | 0,01508236 | 5,4747094 | 0,11339849 | 0,91126707 | 0,9622177 |
| 84079_at | ANKRD27 | -0,01260021 | 8,89725067 | -0,113234 | 0,91139519 | 0,96230024 |

| | | | | | | |
|---|---|---|---|---|---|---|
| 3713_at | IVL | 0,05523318 | 8,64464292 | 0,11302642 | 0,91155689 | 0,96241822 |
| 79872_at | CBLL1 | -0,01549817 | 7,57877576 | -0,11286244 | 0,91168462 | 0,96246641 |
| 80274_at | SCUBE1 | -0,01031108 | 5,07509601 | -0,11283952 | 0,91170247 | 0,96246641 |
| 5862_at | RAB2A | 0,01875911 | 9,09636027 | 0,11250556 | 0,91196262 | 0,9626526 |
| 6457_at | SH3GL3 | -0,00981501 | 4,35086993 | -0,11247952 | 0,91198291 | 0,9626526 |
| 81854_at | ZNF205-AS1 | -0,01411013 | 4,63992493 | -0,11242067 | 0,91202875 | 0,9626526 |
| 339874_at | LOC339874 | -0,00872124 | 3,43449466 | -0,11213948 | 0,9122478 | 0,96280321 |
| 27181_at | SIGLEC8 | -0,0084164 | 4,60116456 | -0,11209621 | 0,91228151 | 0,96280321 |
| 6602_at | SMARCD1 | 0,00750773 | 7,35159936 | 0,11200292 | 0,91235418 | 0,96280321 |
| 56262_at | LRRC8A | -0,01020329 | 8,97819325 | -0,11198085 | 0,91237138 | 0,96280321 |
| 51601_at | LIPT1 | -0,01643828 | 6,12707084 | -0,11172125 | 0,91257362 | 0,96294513 |
| 89797_at | NAV2 | 0,01999984 | 6,97836184 | 0,11165499 | 0,91262524 | 0,96294513 |
| 100009676_a | ZBTB11-AS1 | -0,01047795 | 6,09961951 | -0,11161573 | 0,91265583 | 0,96294513 |
| 10905_at | MAN1A2 | -0,01174198 | 7,55318975 | -0,11119737 | 0,91298177 | 0,9632181 |
| 23415_at | KCNH4 | -0,01061289 | 5,01708728 | -0,11114639 | 0,9130215 | 0,9632181 |
| 26212_at | OR2B6 | 0,00897347 | 3,53202913 | 0,11109113 | 0,91306455 | 0,9632181 |
| 55145_at | THAP1 | -0,02645813 | 7,76185179 | -0,11093868 | 0,91318333 | 0,96329064 |
| 10194_at | TSHZ1 | 0,00735079 | 6,42784621 | 0,11058842 | 0,91345623 | 0,96341546 |
| 132954_at | PDCL2 | -0,00776991 | 2,77547712 | -0,11053737 | 0,91349601 | 0,96341546 |
| 101929154_a | LOC10192915 | 0,01067844 | 3,01040169 | 0,11051888 | 0,91351042 | 0,96341546 |
| 27238_at | GPKOW | -0,01585322 | 8,45509151 | -0,11047938 | 0,91354119 | 0,96341546 |
| 338817_at | LINC01252 | 0,01556292 | 3,59781341 | 0,11046587 | 0,91355172 | 0,96341546 |
| 145783_at | LOC145783 | -0,00989882 | 4,51240048 | -0,11039729 | 0,91360516 | 0,96341907 |
| 81871_at | KRTAP4-6 | 0,01323769 | 5,76104757 | 0,10999741 | 0,91391675 | 0,96357094 |
| 285346_at | ZNF852 | 0,01296266 | 3,84324076 | 0,10999659 | 0,91391739 | 0,96357094 |
| 54942_at | FAM206A | 0,00883008 | 8,13958853 | 0,1098722 | 0,91401432 | 0,96357094 |
| 24146_at | CLDN15 | 0,01027596 | 7,61085408 | 0,10981514 | 0,91405878 | 0,96357094 |
| 57663_at | USP29 | 0,01554976 | 3,46225942 | 0,10975124 | 0,91410858 | 0,96357094 |
| 22983_at | MAST1 | 0,01330946 | 5,03943117 | 0,10974638 | 0,91411236 | 0,96357094 |
| 8826_at | IQGAP1 | 0,01168705 | 10,1692966 | 0,10960323 | 0,91422391 | 0,96357094 |
| 5884_at | RAD17 | -0,02965721 | 7,2230202 | -0,10942703 | 0,91436122 | 0,96357094 |
| 4074_at | M6PR | 0,01175306 | 10,0359692 | 0,10936785 | 0,91440734 | 0,96357094 |
| 257396_at | LOC257396 | -0,02014233 | 4,61930267 | -0,10929894 | 0,91446104 | 0,96357094 |
| 84933_at | C8orf76 | 0,0146688 | 7,93531743 | 0,10928777 | 0,91446975 | 0,96357094 |
| 9629_at | CLCA3P | -0,00688638 | 2,78212094 | -0,10927789 | 0,91447744 | 0,96357094 |
| 2551_at | GABPA | -0,0218518 | 7,46974325 | -0,10925164 | 0,9144979 | 0,96357094 |
| 84460_at | ZMAT1 | -0,00661208 | 2,81084599 | -0,10918472 | 0,91455005 | 0,96357094 |
| 3617_at | IMPG1 | 0,00827057 | 2,98782372 | 0,10913675 | 0,91458743 | 0,96357094 |
| 79817_at | MOB3B | -0,03749154 | 7,07348028 | -0,10908285 | 0,91462944 | 0,96357094 |
| 2887_at | GRB10 | 0,01933003 | 5,83609419 | 0,10904665 | 0,91465765 | 0,96357094 |
| 23604_at | DAPK2 | 0,04528223 | 5,39145232 | 0,10901733 | 0,9146805 | 0,96357094 |
| 79865_at | TREML2 | 0,00863257 | 5,25265474 | 0,10891318 | 0,91476167 | 0,96357094 |
| 152578_at | LINC01618 | 0,00907305 | 3,15272335 | 0,10889665 | 0,91477455 | 0,96357094 |
| 23468_at | CBX5 | 0,04118037 | 8,78790174 | 0,10885433 | 0,91480753 | 0,96357094 |
| 84056_at | KATNAL1 | 0,01575089 | 5,56942537 | 0,10872663 | 0,91490705 | 0,96357094 |
| 4784_at | NFIX | 0,00857223 | 6,56651853 | 0,10863528 | 0,91497824 | 0,96357094 |
| 3339_at | HSPG2 | 0,0186678 | 8,2483209 | 0,10856253 | 0,91503494 | 0,96357094 |
| 10095_at | ARPC1B | -0,00674819 | 10,7531039 | -0,10855113 | 0,91504383 | 0,96357094 |
| 283576_at | ZDHHC22 | 0,00982788 | 4,66699735 | 0,10854354 | 0,91504974 | 0,96357094 |
| 100506022_a | LOC10050602 | 0,01407958 | 4,94957271 | 0,1084237 | 0,91514314 | 0,96359521 |

| | | | | | | |
|---|---|---|---|---|---|---|
| 885_at | CCK | 0,04289517 | 6,25798441 | 0,10838559 | 0,91517284 | 0,96359521 |
| 7691_at | ZNF132 | -0,04904593 | 4,48868417 | -0,10817118 | 0,91533995 | 0,96369367 |
| 3475_at | IFRD1 | -0,02416664 | 7,39478769 | -0,10813723 | 0,91536641 | 0,96369367 |
| 11076_at | TPPP | -0,02412924 | 5,24552563 | -0,10803384 | 0,915447 | 0,96372584 |
| 5132_at | PDC | 0,00680366 | 2,83706257 | 0,10792059 | 0,91553526 | 0,96373973 |
| 25792_at | CIZ1 | -0,01148153 | 8,4659793 | -0,10788853 | 0,91556025 | 0,96373973 |
| 100505876_a | CEBPZOS | 0,01088866 | 7,02909157 | 0,10750218 | 0,91586138 | 0,96400403 |
| 221421_at | RSPH9 | 0,01163717 | 3,7175666 | 0,10733922 | 0,9159884 | 0,96404712 |
| 1470_at | CST2 | -0,05081326 | 5,95496347 | -0,10732125 | 0,91600241 | 0,96404712 |
| 4320_at | MMP11 | 0,00987401 | 5,55303981 | 0,10717063 | 0,91611982 | 0,96406593 |
| 8615_at | USO1 | 0,02628049 | 9,07800681 | 0,10716991 | 0,91612038 | 0,96406593 |
| 10498_at | CARM1 | 0,01334924 | 8,71630339 | 0,10703905 | 0,91622238 | 0,9641206 |
| 100507398_a | INTS6-AS1 | -0,03357077 | 4,40992688 | -0,10689393 | 0,9163355 | 0,96418696 |
| 10533_at | ATG7 | -0,01328636 | 6,89906371 | -0,1067716 | 0,91643086 | 0,96419209 |
| 57524_at | CASKIN1 | -0,00814285 | 6,15193815 | -0,10672774 | 0,91646505 | 0,96419209 |
| 84617_at | TUBB6 | 0,07453905 | 7,05429682 | 0,10665118 | 0,91652473 | 0,96419209 |
| 5150_at | PDE7A | -0,02344488 | 6,42199265 | -0,10663084 | 0,91654058 | 0,96419209 |
| 5161_at | PDHA2 | 0,00995756 | 4,70135385 | 0,10652766 | 0,91662102 | 0,96421513 |
| 30848_at | CTAG2 | -0,00979117 | 5,17156796 | -0,10647431 | 0,9166626 | 0,96421513 |
| 84939_at | MUM1 | 0,02308832 | 7,2275387 | 0,10634042 | 0,91676698 | 0,96424908 |
| 55079_at | FEZF2 | 0,00740448 | 4,08343992 | 0,10630449 | 0,91679499 | 0,96424908 |
| 551_at | AVP | -0,01240959 | 5,4699027 | -0,10623461 | 0,91684947 | 0,96425373 |
| 23155_at | CLCC1 | 0,00936065 | 7,80361879 | 0,10610632 | 0,91694948 | 0,96429871 |
| 54543_at | TOMM7 | 0,01121613 | 10,2493563 | 0,10605131 | 0,91699236 | 0,96429871 |
| 27287_at | VENTX | -0,0148773 | 5,49633925 | -0,10575174 | 0,91722591 | 0,96447227 |
| 100506555_a | PVRL3-AS1 | 0,01286914 | 3,53597673 | 0,10569307 | 0,91727164 | 0,96447227 |
| 9510_at | ADAMTS1 | -0,09598987 | 7,00092499 | -0,10564694 | 0,91730761 | 0,96447227 |
| 81616_at | ACSBG2 | 0,01149501 | 4,12063623 | 0,1054498 | 0,9174613 | 0,96450603 |
| 1292_at | COL6A2 | 0,00908533 | 5,69515827 | 0,10539672 | 0,91750269 | 0,96450603 |
| 163351_at | GBP6 | -0,03697343 | 5,06723069 | -0,10535341 | 0,91753645 | 0,96450603 |
| 152519_at | NIPAL1 | -0,03117054 | 7,31732781 | -0,10527968 | 0,91759393 | 0,96450603 |
| 91862_at | MARVELD3 | -0,01245222 | 7,13754175 | -0,10524707 | 0,91761936 | 0,96450603 |
| 375061_at | FAM89A | 0,00846823 | 6,71556381 | 0,10522042 | 0,91764014 | 0,96450603 |
| 11231_at | SEC63 | 0,00961388 | 7,84722415 | 0,1050557 | 0,91776857 | 0,96458839 |
| 286122_at | C8orf31 | 0,03026959 | 5,65361915 | 0,10498779 | 0,91782151 | 0,9645914 |
| 768096_at | HAR1A | -0,02151947 | 4,48587632 | -0,10477189 | 0,91798985 | 0,96471568 |
| 731779_at | LINC01300 | -0,0093382 | 3,22997181 | -0,10469784 | 0,91804759 | 0,96472373 |
| 388566_at | ZNF470 | -0,01289978 | 4,20411652 | -0,10462147 | 0,91810713 | 0,96473367 |
| 101927074_a | LINC00682 | 0,00821267 | 4,16495424 | 0,10441875 | 0,9182652 | 0,96482507 |
| 431707_at | LHX8 | -0,00699431 | 3,02487658 | -0,10438144 | 0,91829429 | 0,96482507 |
| 23347_at | SMCHD1 | -0,02625078 | 7,07932346 | -0,10410401 | 0,91851061 | 0,96499972 |
| 51004_at | COQ6 | -0,01510965 | 7,55882976 | -0,10376384 | 0,91877586 | 0,96517235 |
| 101929495_a | LINC01339 | 0,00808822 | 4,06219936 | 0,10371462 | 0,91881424 | 0,96517235 |
| 22895_at | RPH3A | 0,00795011 | 4,59103054 | 0,10357725 | 0,91892136 | 0,96517235 |
| 101927196_a | LOC10192719 | 0,00763042 | 3,05804597 | 0,10354072 | 0,91894985 | 0,96517235 |
| 317761_at | C14orf39 | -0,00602322 | 2,80464766 | -0,10351297 | 0,91897149 | 0,96517235 |
| 220047_at | CCDC83 | -0,0068223 | 2,92870199 | -0,10349616 | 0,9189846 | 0,96517235 |
| 401497_at | LINC01242 | 0,00757524 | 2,80551836 | 0,1034435 | 0,91902566 | 0,96517235 |
| 207107_at | SFTA1P | 0,01329667 | 3,77600705 | 0,10332971 | 0,9191144 | 0,96521292 |
| 101927990_a | HRAT13 | -0,00716177 | 3,28372908 | -0,1031925 | 0,91922139 | 0,96527266 |

| | | | | | | |
|---|---|---|---|---|---|---|
| 84677_at | DSCR8 | 0,00920495 | 3,27930961 | 0,10304663 | 0,91933515 | 0,96533949 |
| 3440_at | IFNA2 | -0,00561929 | 3,12035415 | -0,10293411 | 0,9194229 | 0,965345 |
| 10228_at | STX6 | 0,01400198 | 7,91615569 | 0,10282627 | 0,919507 | 0,965345 |
| 54507_at | ADAMTSL4 | -0,01569498 | 4,96549467 | -0,10278472 | 0,9195394 | 0,965345 |
| 197320_at | ZNF778 | -0,01099932 | 5,22295906 | -0,10278287 | 0,91954085 | 0,965345 |
| 56998_at | CTNNBIP1 | 0,02136158 | 7,69602154 | 0,1027043 | 0,91960212 | 0,96535146 |
| 147429_at | AQP4-AS1 | -0,00886975 | 3,07528668 | -0,10261371 | 0,91967277 | 0,96535146 |
| 89887_at | ZNF628 | -0,00785665 | 5,91070084 | -0,1025822 | 0,91969735 | 0,96535146 |
| 4239_at | MFAP4 | -0,01172741 | 4,14302248 | -0,10239869 | 0,91984047 | 0,96544908 |
| 56996_at | SLC12A9 | -0,00728695 | 6,75887444 | -0,10221863 | 0,91998089 | 0,96549747 |
| 4043_at | LRPAP1 | -0,01385876 | 8,85827739 | -0,10221103 | 0,91998682 | 0,96549747 |
| 637_at | BID | 0,01556443 | 9,60459194 | 0,10198267 | 0,92016493 | 0,96563178 |
| 6017_at | RLBP1 | 0,01015211 | 4,7495099 | 0,10182247 | 0,92028987 | 0,96571029 |
| 117581_at | TWIST2 | 0,01772062 | 5,51576131 | 0,10165052 | 0,92042398 | 0,96579841 |
| 5788_at | PTPRC | 0,00540272 | 2,88157653 | 0,10156402 | 0,92049145 | 0,96581659 |
| 28671_at | TRAV13-1 | -0,00853366 | 2,78511562 | -0,10135864 | 0,92065164 | 0,96593206 |
| 25915_at | NDUFAF3 | -0,03215631 | 9,28710362 | -0,10119892 | 0,92077622 | 0,96595997 |
| 80704_at | SLC19A3 | 0,00742805 | 3,64070819 | 0,10118092 | 0,92079026 | 0,96595997 |
| 101669764_a | GPR1-AS | -0,0078706 | 3,35369271 | -0,10107533 | 0,92087262 | 0,96595997 |
| 64131_at | XYLT1 | 0,02741131 | 5,11790306 | 0,10103904 | 0,92090093 | 0,96595997 |
| 375295_at | LINC01116 | 0,00983867 | 5,06684086 | 0,10088695 | 0,92101957 | 0,96595997 |
| 25913_at | POT1 | -0,01406625 | 7,96697563 | -0,10080818 | 0,921081 | 0,96595997 |
| 100507433_a | ZNF571-AS1 | -0,00803519 | 3,84416315 | -0,10078154 | 0,92110179 | 0,96595997 |
| 6419_at | SETMAR | -0,00659638 | 5,83824701 | -0,10074565 | 0,92112978 | 0,96595997 |
| 64788_at | LMF1 | -0,01497011 | 5,80632687 | -0,10071847 | 0,92115098 | 0,96595997 |
| 163255_at | ZNF540 | -0,00666753 | 3,34871898 | -0,10068165 | 0,92117971 | 0,96595997 |
| 57471_at | ERMN | -0,01017999 | 3,10776496 | -0,10026452 | 0,92150509 | 0,96624857 |
| 60370_at | AVPI1 | -0,01634437 | 9,27163302 | -0,10017086 | 0,92157816 | 0,96626261 |
| 93654_at | ST7-AS2 | 0,00718527 | 3,05838375 | 0,10011876 | 0,9216188 | 0,96626261 |
| 55589_at | BMP2K | -0,0313832 | 6,01093275 | -0,09997863 | 0,92172811 | 0,96630869 |
| 646962_at | HRCT1 | 0,01219243 | 5,69454208 | 0,09993381 | 0,92176308 | 0,96630869 |
| 1982_at | EIF4G2 | -0,00630827 | 11,8245686 | -0,0998041 | 0,92186427 | 0,96636218 |
| 3595_at | IL12RB2 | 0,00762291 | 3,57560745 | 0,09952666 | 0,9220807 | 0,96649349 |
| 729013_at | ZBED5-AS1 | -0,0446955 | 8,20931821 | -0,0995149 | 0,92208988 | 0,96649349 |
| 55529_at | TMEM55A | -0,03596695 | 5,08342585 | -0,0993834 | 0,92219247 | 0,96654842 |
| 5213_at | PFKM | -0,01885373 | 8,19772178 | -0,09918476 | 0,92234744 | 0,96663454 |
| 51005_at | AMDHD2 | 0,00900455 | 6,30953862 | 0,09906514 | 0,92244076 | 0,96663454 |
| 10220_at | GDF11 | 0,02911783 | 6,32037951 | 0,09902722 | 0,92247035 | 0,96663454 |
| 1796_at | DOK1 | 0,03438749 | 6,47966707 | 0,0990208 | 0,92247536 | 0,96663454 |
| 4332_at | MNDA | 0,00688401 | 2,9168314 | 0,09877059 | 0,92267057 | 0,96665478 |
| 2585_at | GALK2 | -0,00900904 | 5,95480057 | -0,09876014 | 0,92267872 | 0,96665478 |
| 733_at | C8G | -0,01210674 | 5,42841961 | -0,09875381 | 0,92268366 | 0,96665478 |
| 1030_at | CDKN2B | -0,00909451 | 4,27108729 | -0,09867941 | 0,92274171 | 0,96665478 |
| 101928741_a | LOC10192874 | -0,01049135 | 4,01835669 | -0,09867445 | 0,92274558 | 0,96665478 |
| 9254_at | CACNA2D2 | -0,00660595 | 5,21818574 | -0,0981446 | 0,92315898 | 0,96700645 |
| 100505583_a | LINC00838 | -0,01151729 | 3,86784108 | -0,09811552 | 0,92318167 | 0,96700645 |
| 1436_at | CSF1R | -0,01256926 | 4,83585646 | -0,09796582 | 0,92329848 | 0,96707621 |
| 79961_at | DENND2D | -0,05518157 | 6,42093285 | -0,0978931 | 0,92335522 | 0,96708306 |
| 374882_at | TMEM205 | 0,03387007 | 9,24965636 | 0,09771383 | 0,9234951 | 0,96717698 |
| 55803_at | ADAP2 | 0,00731596 | 5,17814259 | 0,09754735 | 0,923625 | 0,96726044 |

| | | | | | | |
|---|---|---|---|---|---|---|
| 28656_at | TRAV26-2 | 0,00734808 | 3,63254974 | 0,09746201 | 0,9236916 | 0,96727759 |
| 11016_at | ATF7 | -0,01153808 | 6,92000747 | -0,09732641 | 0,92379741 | 0,96733581 |
| 3691_at | ITGB4 | 0,00979458 | 8,22759185 | 0,09712942 | 0,92395113 | 0,96744418 |
| 129531_at | MITD1 | 0,00716508 | 7,31992235 | 0,09682951 | 0,92418516 | 0,96760747 |
| 200895_at | DHFRL1 | 0,03308744 | 6,20708126 | 0,09680084 | 0,92420754 | 0,96760747 |
| 4502_at | MT2A | 0,03495071 | 12,0018131 | 0,09655396 | 0,9244002 | 0,96775243 |
| 102723742_a | LINC01500 | -0,00594336 | 3,20263682 | -0,09647205 | 0,92446412 | 0,96775243 |
| 140850_at | DEFB127 | 0,00736031 | 2,94216027 | 0,09643027 | 0,92449672 | 0,96775243 |
| 100294720_a | NHEG1 | 0,01054187 | 4,01232257 | 0,09611915 | 0,92473951 | 0,96795399 |
| 339500_at | ZNF678 | -0,01045928 | 5,23827616 | -0,09581673 | 0,92497554 | 0,96804411 |
| 401463_at | LOC401463 | 0,00844683 | 3,34495006 | 0,09581403 | 0,92497764 | 0,96804411 |
| 25825_at | BACE2 | -0,0278237 | 7,72257704 | -0,09575998 | 0,92501983 | 0,96804411 |
| 90799_at | CEP95 | 0,01531946 | 6,82465488 | 0,0956458 | 0,92510894 | 0,96804411 |
| 83656_at | FAM167A-AS1 | -0,00833038 | 3,85504103 | -0,09563437 | 0,92511786 | 0,96804411 |
| 81617_at | CAB39L | -0,08781811 | 6,42314544 | -0,09556931 | 0,92516863 | 0,96804411 |
| 122651_at | RNASE11 | -0,0064615 | 3,00267718 | -0,0955581 | 0,92517739 | 0,96804411 |
| 128611_at | ZNF831 | 0,00709752 | 3,88227527 | 0,09541559 | 0,92528861 | 0,96805708 |
| 4038_at | LRP4 | -0,02904696 | 7,83736658 | -0,09539073 | 0,92530801 | 0,96805708 |
| 4040_at | LRP6 | 0,00730032 | 6,68847157 | 0,09534903 | 0,92534055 | 0,96805708 |
| 283140_at | LOC283140 | 0,01368195 | 5,27659495 | 0,09517748 | 0,92547445 | 0,96812207 |
| 51171_at | HSD17B14 | 0,01026822 | 5,6240575 | 0,09514066 | 0,92550319 | 0,96812207 |
| 102724094_a | LOC10272409 | 0,01247393 | 5,47137629 | 0,09501606 | 0,92560044 | 0,96816757 |
| 400654_at | LINC01538 | -0,00968898 | 3,07296738 | -0,09493094 | 0,92566688 | 0,96816757 |
| 339862_at | LOC339862 | -0,00673457 | 3,08050944 | -0,09489175 | 0,92569746 | 0,96816757 |
| 8689_at | KRT36 | 0,01253989 | 3,87449965 | 0,09481059 | 0,92576081 | 0,96818126 |
| 3593_at | IL12B | 0,00665505 | 3,64738282 | 0,09469982 | 0,92584727 | 0,96821911 |
| 9219_at | MTA2 | -0,02660927 | 7,9758987 | -0,09450565 | 0,92599882 | 0,96827841 |
| 113230_at | LOC113230 | -0,02134968 | 5,42184531 | -0,09449837 | 0,92600451 | 0,96827841 |
| 79625_at | NDNF | -0,00944025 | 4,44352852 | -0,09419764 | 0,92623924 | 0,96847129 |
| 55212_at | BBS7 | 0,0290376 | 5,93882702 | 0,09399519 | 0,92639728 | 0,96858395 |
| 149620_at | CHIAP2 | 0,00747747 | 3,38161219 | 0,09380903 | 0,92654259 | 0,96868331 |
| 26127_at | FGFR1OP2 | -0,01678374 | 7,4541194 | -0,09370629 | 0,92662279 | 0,96871458 |
| 145741_at | C2CD4A | 0,00851657 | 4,82697685 | 0,09354091 | 0,92675189 | 0,96872809 |
| 166752_at | FREM3 | 0,0064745 | 3,14830878 | 0,09339392 | 0,92686663 | 0,96872809 |
| 401022_at | HAGLR | -0,01931944 | 5,813158 | -0,09338791 | 0,92687132 | 0,96872809 |
| 101927495_a | LOC10192749 | -0,00844045 | 3,51083791 | -0,09319571 | 0,92702136 | 0,96872809 |
| 57187_at | THOC2 | 0,0111968 | 8,06004079 | 0,09314362 | 0,92706203 | 0,96872809 |
| 256158_at | HMCN2 | 0,00821581 | 5,15975804 | 0,09312552 | 0,92707615 | 0,96872809 |
| 285780_at | LY86-AS1 | -0,00764851 | 3,25196365 | -0,09310919 | 0,92708891 | 0,96872809 |
| 9466_at | IL27RA | -0,01284543 | 5,75268245 | -0,09296545 | 0,92720112 | 0,96872809 |
| 10409_at | BASP1 | 0,07576276 | 9,67898815 | 0,09289439 | 0,92725659 | 0,96872809 |
| 4857_at | NOVA1 | 0,02297257 | 4,55983067 | 0,09279734 | 0,92733235 | 0,96872809 |
| 7060_at | THBS4 | -0,01072259 | 5,31712469 | -0,09279256 | 0,92733609 | 0,96872809 |
| 57563_at | KLHL8 | 0,02756118 | 7,09791847 | 0,09278775 | 0,92733985 | 0,96872809 |
| 117583_at | PARD3B | -0,00935913 | 4,14458023 | -0,09257917 | 0,92750268 | 0,96872809 |
| 164684_at | WBP2NL | -0,00743093 | 3,07821403 | -0,09251252 | 0,92755472 | 0,96872809 |
| 100505768_a | LINC01364 | 0,00662183 | 2,87714735 | 0,09246933 | 0,92758843 | 0,96872809 |
| 58503_at | PROL1 | -0,00874127 | 3,82921045 | -0,09244753 | 0,92760546 | 0,96872809 |
| 2877_at | GPX2 | -0,01676548 | 12,0731571 | -0,0923301 | 0,92769713 | 0,96872809 |
| 220001_at | VWCE | -0,00694129 | 5,68233249 | -0,09225452 | 0,92775614 | 0,96872809 |

| | | | | | | |
|---|---|---|---|---|---|---|
| 23199_at | GSE1 | -0,01308148 | 7,62257782 | -0,09224523 | 0,92776339 | 0,96872809 |
| 9753_at | ZSCAN12 | 0,00463372 | 3,67276456 | 0,09223631 | 0,92777036 | 0,96872809 |
| 51094_at | ADIPOR1 | -0,00888636 | 9,96166498 | -0,09218943 | 0,92780696 | 0,96872809 |
| 50836_at | TAS2R8 | -0,00764819 | 2,87464829 | -0,092178 | 0,92781588 | 0,96872809 |
| 55742_at | PARVA | -0,00908323 | 7,62981854 | -0,09214813 | 0,9278392 | 0,96872809 |
| 100507507_a | LOC10050750 | 0,01294616 | 6,83541182 | 0,09214369 | 0,92784267 | 0,96872809 |
| 5911_at | RAP2A | -0,00909917 | 8,24284489 | -0,09202987 | 0,92793153 | 0,96875545 |
| 23521_at | RPL13A | -0,01042392 | 11,1624874 | -0,09198129 | 0,92796946 | 0,96875545 |
| 51131_at | PHF11 | -0,10165244 | 5,94870775 | -0,09190993 | 0,92802517 | 0,96876112 |
| 8323_at | FZD6 | -0,0150084 | 10,460796 | -0,09154006 | 0,92831395 | 0,96901006 |
| 100874241_a | STARD13-AS | -0,00542143 | 2,91826008 | -0,09112957 | 0,92863446 | 0,96921344 |
| 55714_at | TENM3 | -0,01196154 | 5,24276096 | -0,09103952 | 0,92870477 | 0,96921344 |
| 894_at | CCND2 | 0,01150014 | 5,43325134 | 0,09100271 | 0,92873352 | 0,96921344 |
| 145497_at | LRRC74A | -0,00766141 | 4,15331498 | -0,09097289 | 0,92875679 | 0,96921344 |
| 6904_at | TBCD | 0,00793828 | 7,7858284 | 0,09094678 | 0,92877719 | 0,96921344 |
| 135112_at | NCOA7 | -0,01943475 | 9,86967599 | -0,09086296 | 0,92884263 | 0,96921344 |
| 23683_at | PRKD3 | 0,02538836 | 7,42215572 | 0,0907776 | 0,92890929 | 0,96921344 |
| 140625_at | ACTRT2 | -0,0108363 | 4,8668343 | -0,090775 | 0,92891131 | 0,96921344 |
| 2070_at | EYA4 | 0,02828917 | 4,12956671 | 0,09040153 | 0,92920294 | 0,96946521 |
| 29103_at | DNAJC15 | -0,02529735 | 8,26461744 | -0,09026917 | 0,92930629 | 0,96952053 |
| 84107_at | ZIC4 | 0,00903381 | 4,06143203 | 0,08997127 | 0,92953892 | 0,96971071 |
| 644844_at | PHGR1 | -0,01571638 | 5,22850466 | -0,08952512 | 0,92988733 | 0,97002164 |
| 28648_at | TRAV34 | 0,01016738 | 3,08348752 | 0,08933921 | 0,93003251 | 0,97009596 |
| 51023_at | MRPS18C | 0,01318517 | 3,8363772 | 0,08923201 | 0,93011623 | 0,97009596 |
| 101929748_a | LOC10192974 | -0,00881409 | 5,15325579 | -0,08918665 | 0,93015165 | 0,97009596 |
| 9404_at | LPXN | -0,026627 | 6,77646901 | -0,08917594 | 0,93016002 | 0,97009596 |
| 728095_at | LOC728095 | -0,00990312 | 4,79605301 | -0,08910106 | 0,93021849 | 0,97010442 |
| 8351_at | HIST1H3D | -0,02125819 | 6,86619767 | -0,08887375 | 0,93039601 | 0,97023703 |
| 143503_at | OR51E1 | -0,00911198 | 3,99338702 | -0,08875126 | 0,93049168 | 0,97028426 |
| 151278_at | CCDC140 | 0,00739873 | 4,04759428 | 0,08856059 | 0,93064059 | 0,97035899 |
| 204851_at | HIPK1 | -0,0069254 | 6,9635679 | -0,08852571 | 0,93066783 | 0,97035899 |
| 55855_at | FAM45B | 0,01651022 | 6,75829115 | 0,088466 | 0,93071446 | 0,97035899 |
| 57539_at | WDR35 | 0,00853103 | 6,07796962 | 0,088266 | 0,93087066 | 0,97038516 |
| 90019_at | SYT8 | 0,01045201 | 7,37137164 | 0,08823824 | 0,93089235 | 0,97038516 |
| 11010_at | GLIPR1 | 0,03684141 | 7,58343698 | 0,0881844 | 0,9309344 | 0,97038516 |
| 23614_at | PPY2P | 0,01124036 | 4,05398488 | 0,08812605 | 0,93097997 | 0,97038516 |
| 6700_at | SPRR2A | 0,01266113 | 3,94831277 | 0,08808704 | 0,93101044 | 0,97038516 |
| 29100_at | TMEM208 | 0,00477784 | 8,52496093 | 0,08804687 | 0,93104182 | 0,97038516 |
| 221091_at | LRRN4CL | -0,00929292 | 4,93156231 | -0,08777673 | 0,93125281 | 0,97051871 |
| 283635_at | FAM177A1 | 0,01148659 | 7,39339755 | 0,0877538 | 0,93127072 | 0,97051871 |
| 85025_at | TMEM60 | 0,00854677 | 8,12211472 | 0,08762559 | 0,93137086 | 0,97056046 |
| 25870_at | SUMF2 | 0,02338397 | 9,32970695 | 0,08752139 | 0,93145225 | 0,97056046 |
| 284998_at | LINC01114 | -0,00933496 | 4,42218946 | -0,08749461 | 0,93147316 | 0,97056046 |
| 3190_at | HNRNPK | 0,00432206 | 11,6068811 | 0,08744448 | 0,93151232 | 0,97056046 |
| 100506127_a | LOC10050612 | -0,0051283 | 3,55607756 | -0,08724094 | 0,9316713 | 0,97067361 |
| 4820_at | NKTR | -0,01323085 | 6,91834459 | -0,08717579 | 0,93172219 | 0,97067412 |
| 7291_at | TWIST1 | 0,00761493 | 4,0682889 | 0,08709587 | 0,93178462 | 0,97068666 |
| 7917_at | BAG6 | 0,00573952 | 9,73087273 | 0,08678972 | 0,93202376 | 0,97088328 |
| 287015_at | TRIM42 | -0,00726032 | 3,56749423 | -0,08669343 | 0,93209897 | 0,97090912 |
| 404201_at | WDFY3-AS2 | -0,00857882 | 4,71003874 | -0,08645682 | 0,9322838 | 0,97097738 |

| | | | | | | |
|---|---|---|---|---|---|---|
| 25839_at | COG4 | -0,00887431 | 8,78918302 | -0,08643606 | 0,93230002 | 0,97097738 |
| 340094_at | LINC01020 | -0,0050994 | 2,84235916 | -0,08641596 | 0,93231571 | 0,97097738 |
| 145501_at | ISM2 | -0,0096912 | 4,79622802 | -0,0860796 | 0,93257846 | 0,97119852 |
| 285888_at | CNPY1 | 0,00828764 | 3,2261185 | 0,08597696 | 0,93265865 | 0,97122951 |
| 100289255_a | LINC00675 | -0,01370581 | 4,95434692 | -0,08588663 | 0,93272922 | 0,97125049 |
| 79905_at | TMC7 | 0,0259556 | 5,88124316 | 0,08576681 | 0,93282282 | 0,97129546 |
| 534_at | ATP6V1G2 | 0,00753876 | 4,27039452 | 0,08554321 | 0,93299749 | 0,97140636 |
| 50999_at | TMED5 | 0,0083041 | 8,40123613 | 0,08550136 | 0,93303019 | 0,97140636 |
| 333_at | APLP1 | -0,0133141 | 6,29315568 | -0,08534959 | 0,93314876 | 0,9714773 |
| 284131_at | ENDOV | 0,01148467 | 6,38956377 | 0,08526178 | 0,93321736 | 0,97149622 |
| 4888_at | NPY6R | 0,00712661 | 3,86782047 | 0,08481607 | 0,93356557 | 0,97168208 |
| 388886_at | LRRC75B | -0,01306503 | 6,15490679 | -0,08480337 | 0,93357549 | 0,97168208 |
| 29930_at | PCDHB1 | 0,00800587 | 3,97859335 | 0,08478498 | 0,93358986 | 0,97168208 |
| 284076_at | TTLL6 | 0,00630113 | 4,28694837 | 0,08477498 | 0,93359767 | 0,97168208 |
| 10326_at | SIRPB1 | 0,00925952 | 3,94092086 | 0,08462347 | 0,93371605 | 0,97169374 |
| 100132987_a | LINC00856 | -0,00966932 | 4,16425461 | -0,08457477 | 0,93375409 | 0,97169374 |
| #N/A | GS1-279B7,1 | -0,01101837 | 4,09083027 | -0,08447428 | 0,93383261 | 0,97169374 |
| 65009_at | NDRG4 | -0,0187636 | 6,83328054 | -0,08435889 | 0,93392276 | 0,97169374 |
| 8706_at | B3GALNT1 | -0,11010712 | 6,6918441 | -0,08433178 | 0,93394395 | 0,97169374 |
| 10735_at | STAG2 | 0,04602085 | 7,58810847 | 0,08427646 | 0,93398716 | 0,97169374 |
| 7175_at | TPR | 0,00924916 | 7,6582703 | 0,08424297 | 0,93401333 | 0,97169374 |
| 139135_at | PASD1 | -0,00978214 | 3,63197866 | -0,08421162 | 0,93403783 | 0,97169374 |
| 93653_at | ST7-AS1 | -0,00796178 | 3,85904001 | -0,08417957 | 0,93406286 | 0,97169374 |
| 100507250_a | LOC10050725 | 0,01269328 | 4,28870838 | 0,08400894 | 0,93419618 | 0,97177994 |
| 147945_at | NLRP4 | -0,00679219 | 3,36508494 | -0,08369054 | 0,93444496 | 0,97191087 |
| 1466_at | CSRP2 | -0,0563874 | 9,87057584 | -0,08368177 | 0,93445181 | 0,97191087 |
| 9235_at | IL32 | 0,01424943 | 5,21917753 | 0,08365414 | 0,9344734 | 0,97191087 |
| 28563_at | TRBV24-1 | -0,01079421 | 4,11591846 | -0,0835787 | 0,93453234 | 0,97191969 |
| 963_at | CD53 | 0,00714023 | 5,12769769 | 0,08346212 | 0,93462344 | 0,97196196 |
| 5053_at | PAH | 0,00437584 | 2,96261094 | 0,08317349 | 0,93484897 | 0,97214401 |
| 696_at | BTN1A1 | -0,00952585 | 4,31567749 | -0,08287735 | 0,93508037 | 0,97225602 |
| 442721_at | LMOD2 | 0,01052464 | 3,79279544 | 0,0828269 | 0,9351198 | 0,97225602 |
| 10749_at | KIF1C | -0,00770193 | 7,31154589 | -0,08276683 | 0,93516674 | 0,97225602 |
| 116068_at | LYSMD3 | 0,02368368 | 7,90625177 | 0,08271762 | 0,93520519 | 0,97225602 |
| 57623_at | ZFAT | -0,01143847 | 6,33057282 | -0,08271269 | 0,93520904 | 0,97225602 |
| 55298_at | RNF121 | -0,00635324 | 7,62147986 | -0,0824279 | 0,93543158 | 0,97243489 |
| 79893_at | GGNBP2 | 0,00728691 | 8,73935273 | 0,08222172 | 0,9355927 | 0,97249545 |
| 6999_at | TDO2 | 0,01137702 | 3,65756309 | 0,0822027 | 0,93560756 | 0,97249545 |
| 7342_at | UBP1 | -0,00568262 | 9,51013281 | -0,08215748 | 0,9356429 | 0,97249545 |
| 101927219_a | HNF4A-AS1 | 0,00541857 | 3,34978066 | 0,08202902 | 0,93574328 | 0,97249545 |
| 105371397_a | LOC10537139 | -0,0095143 | 3,86734388 | -0,08200035 | 0,93576569 | 0,97249545 |
| 101927007_a | LINC01365 | -0,00806788 | 3,79526047 | -0,08194249 | 0,93581091 | 0,97249545 |
| 4225_at | MEP1B | 0,0076382 | 3,72959602 | 0,08190113 | 0,93584323 | 0,97249545 |
| 3576_at | CXCL8 | 0,08044815 | 6,36555215 | 0,08182241 | 0,93590475 | 0,97250691 |
| 23092_at | ARHGAP26 | 0,02039162 | 6,92093886 | 0,08163381 | 0,93605214 | 0,97256785 |
| 100134259_a | LINC01119 | -0,0114134 | 4,36753564 | -0,08152754 | 0,93613519 | 0,97256785 |
| 25759_at | SHC2 | -0,01202039 | 5,61038891 | -0,08149295 | 0,93616222 | 0,97256785 |
| 9315_at | NREP | 0,00652326 | 3,36137771 | 0,08148894 | 0,93616535 | 0,97256785 |
| 2316_at | FLNA | 0,02572036 | 9,71519524 | 0,08113172 | 0,93644452 | 0,97280541 |
| 388588_at | SMIM1 | 0,00730008 | 6,44070516 | 0,08101276 | 0,93653749 | 0,97284953 |

| | | | | | | |
|---|---|---|---|---|---|---|
| 6916_at | TBXAS1 | -0,01497207 | 4,6919508 | -0,08062099 | 0,93684368 | 0,97307933 |
| 2681_at | GGTA1P | -0,00684408 | 3,78974941 | -0,08060043 | 0,93685975 | 0,97307933 |
| 831_at | CAST | -0,01396118 | 10,6814626 | -0,08037822 | 0,93703343 | 0,97318526 |
| 100329135_a | TRPC5OS | 0,00757185 | 3,00022869 | 0,08034066 | 0,93706277 | 0,97318526 |
| 11216_at | AKAP10 | -0,01492645 | 6,52826927 | -0,07994107 | 0,93737509 | 0,97345714 |
| 163782_at | KANK4 | 0,01315207 | 4,55213242 | 0,0798337 | 0,93745902 | 0,97349181 |
| 285740_at | LOC285740 | 0,00835343 | 3,25508394 | 0,07955325 | 0,93767823 | 0,97364518 |
| 5592_at | PRKG1 | -0,00499078 | 3,58610883 | -0,07951541 | 0,9377078 | 0,97364518 |
| 56134_at | PCDHAC2 | -0,00643953 | 4,83281419 | -0,07934204 | 0,93784332 | 0,9737334 |
| 766_at | CA7 | 0,00661098 | 5,01757177 | 0,07913287 | 0,93800681 | 0,97385067 |
| 101929454_a | LOC10192945 | -0,00641913 | 3,23706536 | -0,07898487 | 0,9381225 | 0,97390718 |
| 374987_at | NEXN-AS1 | 0,00683212 | 3,47877179 | 0,07893386 | 0,93816237 | 0,97390718 |
| 1803_at | DPP4 | 0,00408773 | 3,60173472 | 0,07871285 | 0,93833513 | 0,97403404 |
| 6474_at | SHOX2 | -0,00587331 | 4,46650046 | -0,07844279 | 0,93854623 | 0,97420067 |
| 84273_at | NOA1 | 0,01322992 | 8,93475626 | 0,07820101 | 0,93873524 | 0,97430489 |
| 54708_at | MARCH5 | -0,01007998 | 8,27704875 | -0,07816583 | 0,93876274 | 0,97430489 |
| 284525_at | SLC9C2 | -0,00532217 | 2,76447366 | -0,0780897 | 0,93882225 | 0,97430489 |
| 51704_at | GPRC5B | -0,01570204 | 6,24020399 | -0,07805555 | 0,93884895 | 0,97430489 |
| 5655_at | KLK10 | -0,02024397 | 5,84527666 | -0,07773276 | 0,93910129 | 0,9744538 |
| 49854_at | ZBTB21 | -0,01146179 | 7,83593942 | -0,07766734 | 0,93915243 | 0,9744538 |
| 145389_at | SLC38A6 | -0,01649559 | 7,74479413 | -0,07761715 | 0,93919167 | 0,9744538 |
| 116832_at | RPL39L | -0,05103073 | 5,0885752 | -0,07752146 | 0,93926647 | 0,9744538 |
| 54994_at | GID8 | 0,01075056 | 10,0234775 | 0,07751583 | 0,93927087 | 0,9744538 |
| 28778_at | IGLV6-57 | 0,00888535 | 4,77313306 | 0,07743735 | 0,93933223 | 0,9744538 |
| 440184_at | LINC00238 | 0,00653628 | 2,91972553 | 0,07741903 | 0,93934655 | 0,9744538 |
| 51065_at | RPS27L | 0,01049732 | 6,45236445 | 0,07702104 | 0,93965769 | 0,97469164 |
| 4345_at | CD200 | 0,00460754 | 3,18864861 | 0,07676941 | 0,93985442 | 0,97469164 |
| 79583_at | TMEM231 | 0,0101214 | 5,81186625 | 0,0766203 | 0,939971 | 0,97469164 |
| 26058_at | GIGYF2 | -0,0059057 | 7,72261041 | -0,07660228 | 0,93998508 | 0,97469164 |
| 100128640_a | ACVR2B-AS1 | 0,01166975 | 4,53922231 | 0,07657931 | 0,94000304 | 0,97469164 |
| 768239_at | PSAPL1 | -0,00937444 | 5,84058709 | -0,0765692 | 0,94001094 | 0,97469164 |
| 23617_at | TSSK2 | -0,0060654 | 4,58180535 | -0,0764756 | 0,94008412 | 0,97469164 |
| 5753_at | PTK6 | 0,01571538 | 7,95449734 | 0,07644801 | 0,94010569 | 0,97469164 |
| 100996342_a | LOC10099634 | 0,00630813 | 5,2474874 | 0,07634378 | 0,94018719 | 0,97469164 |
| 255926_at | ADAM5 | 0,00820259 | 3,37594595 | 0,07631867 | 0,94020682 | 0,97469164 |
| 54742_at | LY6K | 0,046608 | 5,78552343 | 0,07631147 | 0,94021245 | 0,97469164 |
| 7013_at | TERF1 | 0,00666979 | 7,87675145 | 0,07630829 | 0,94021493 | 0,97469164 |
| 115548_at | FCHO2 | -0,02754021 | 7,30167206 | -0,0762844 | 0,94023362 | 0,97469164 |
| 4915_at | NTRK2 | -0,00434827 | 3,86813384 | -0,07621333 | 0,94028918 | 0,97469679 |
| 4148_at | MATN3 | 0,03715634 | 4,46004659 | 0,07614762 | 0,94034055 | 0,97469759 |
| 122060_at | SLAIN1 | 0,00952639 | 3,53626432 | 0,07600767 | 0,94044998 | 0,97475857 |
| 322_at | APBB1 | 0,00801454 | 5,21371315 | 0,07586521 | 0,94056136 | 0,97479075 |
| 5345_at | SERPINF2 | 0,01011917 | 5,03706048 | 0,07583851 | 0,94058224 | 0,97479075 |
| 4150_at | MAZ | -0,0231556 | 7,89008535 | -0,07569444 | 0,94069488 | 0,97483164 |
| 4327_at | MMP19 | -0,00668855 | 5,12138557 | -0,07565859 | 0,94072291 | 0,97483164 |
| 118672_at | PSTK | 0,00502868 | 5,19287218 | 0,0755862 | 0,94077951 | 0,97483786 |
| 22982_at | DIP2C | -0,01469503 | 6,68826717 | -0,0754805 | 0,94086216 | 0,97486479 |
| 339666_at | LOC339666 | 0,00876528 | 4,99861315 | 0,0753623 | 0,94095458 | 0,97486479 |
| 10114_at | HIPK3 | -0,0108902 | 6,69782178 | -0,07535879 | 0,94095733 | 0,97486479 |
| 153642_at | ARSK | -0,02408805 | 6,3701343 | -0,07511432 | 0,94114848 | 0,97500342 |

| | | | | | | |
|---|---|---|---|---|---|---|
| 146330_at | FBXL16 | -0,01301751 | 6,1919956 | -0,07502874 | 0,9412154 | 0,97500342 |
| 6346_at | CCL1 | -0,00966143 | 4,01257579 | -0,07498589 | 0,9412489 | 0,97500342 |
| 56924_at | PAK6 | 0,00896692 | 7,17270315 | 0,07492872 | 0,9412936 | 0,97500342 |
| 2268_at | FGR | 0,00545033 | 5,73186809 | 0,07482257 | 0,9413766 | 0,97503697 |
| 54465_at | ETAA1 | -0,00828608 | 7,38801237 | -0,07468523 | 0,941484 | 0,97509577 |
| 23355_at | VPS8 | -0,01252672 | 7,88112441 | -0,07456894 | 0,94157493 | 0,97513752 |
| 28688_at | TRAV5 | 0,00808169 | 3,4817762 | 0,07405512 | 0,94197672 | 0,975482 |
| 1102_at | RCBTB2 | 0,03342908 | 4,82496234 | 0,07401406 | 0,94200883 | 0,975482 |
| 9294_at | S1PR2 | 0,00544178 | 6,32707732 | 0,07394013 | 0,94206664 | 0,97548942 |
| 4050_at | LTB | 0,00747472 | 6,21348722 | 0,07371675 | 0,94224132 | 0,97561786 |
| 123283_at | TARSL2 | -0,00508188 | 5,46908955 | -0,07363694 | 0,94230374 | 0,97563004 |
| 81894_at | SLC25A28 | -0,00978876 | 6,99517709 | -0,07350878 | 0,94240396 | 0,97568136 |
| 63970_at | TP53AIP1 | -0,00475074 | 4,1164164 | -0,07338669 | 0,94249944 | 0,97572777 |
| 5173_at | PDYN | 0,00576387 | 3,81565294 | 0,07329457 | 0,94257148 | 0,97574991 |
| 84269_at | CHCHD5 | -0,01584855 | 7,73300682 | -0,07300621 | 0,94279699 | 0,9759201 |
| 100359394_a | LINC00102 | 0,00482788 | 3,07435279 | 0,07295478 | 0,94283721 | 0,9759201 |
| 440279_at | UNC13C | -0,00358989 | 2,91179678 | -0,07281269 | 0,94294833 | 0,97598269 |
| 127495_at | LRRC39 | 0,00980208 | 3,66531753 | 0,07274582 | 0,94300063 | 0,97598437 |
| 3381_at | IBSP | -0,00457623 | 3,5452427 | -0,07266823 | 0,94306131 | 0,9759853 |
| 147700_at | KLC3 | 0,00697091 | 6,66134514 | 0,07261511 | 0,94310286 | 0,9759853 |
| 152_at | ADRA2C | -0,01862946 | 6,3685327 | -0,0723888 | 0,94327984 | 0,97611601 |
| 4311_at | MME | 0,0076913 | 4,59826927 | 0,07221129 | 0,94341867 | 0,97614841 |
| 80013_at | FAM188A | -0,01250133 | 6,28898236 | -0,0721304 | 0,94348194 | 0,97614841 |
| 51132_at | RLIM | 0,00696801 | 7,68585365 | 0,07210602 | 0,94350101 | 0,97614841 |
| 778_at | CACNA1F | -0,00968681 | 4,8872639 | -0,0720896 | 0,94351385 | 0,97614841 |
| 2314_at | FLII | -0,03109278 | 9,44277918 | -0,07184662 | 0,94370388 | 0,97629258 |
| 3612_at | IMPA1 | 0,01251922 | 8,79383823 | 0,07168716 | 0,94382861 | 0,97629627 |
| 54940_at | OCIAD1 | -0,00988073 | 10,1519518 | -0,07164688 | 0,94386011 | 0,97629627 |
| 54494_at | C11orf71 | 0,01022517 | 6,46458686 | 0,07160738 | 0,943891 | 0,97629627 |
| 9712_at | USP6NL | -0,01005046 | 6,23750104 | -0,07158286 | 0,94391018 | 0,97629627 |
| 341_at | APOC1 | 0,03376025 | 6,07655187 | 0,07139417 | 0,94405776 | 0,9763965 |
| 51268_at | PIPOX | -0,00632904 | 3,80388808 | -0,07126724 | 0,94415704 | 0,97641117 |
| 343990_at | KIAA1211L | -0,01580864 | 5,28643075 | -0,07124642 | 0,94417333 | 0,97641117 |
| 3543_at | IGLL1 | -0,00623977 | 4,10505569 | -0,07104413 | 0,94433155 | 0,97651707 |
| 3204_at | HOXA7 | 0,05251457 | 6,93399175 | 0,07092081 | 0,94442801 | 0,97651707 |
| 392_at | ARHGAP1 | 0,01230269 | 9,09931651 | 0,07086532 | 0,94447141 | 0,97651707 |
| 182_at | JAG1 | -0,00975422 | 8,3850632 | -0,07085118 | 0,94448247 | 0,97651707 |
| 10487_at | CAP1 | -0,00511442 | 11,4824249 | -0,07077808 | 0,94453965 | 0,97651707 |
| 159013_at | CXorf38 | 0,01175339 | 7,60345629 | 0,07072663 | 0,94457989 | 0,97651707 |
| 128854_at | TSPY26P | 0,00809328 | 5,36199675 | 0,07063862 | 0,94464873 | 0,97653582 |
| 101927359_a | LOC10192735 | 0,00416657 | 2,85265449 | 0,07048019 | 0,94477265 | 0,97661152 |
| 64130_at | LIN7B | 0,00552465 | 6,82359688 | 0,07040113 | 0,9448345 | 0,97662304 |
| 100506922_a | LOC10050692 | 0,01664976 | 5,87764821 | 0,07000023 | 0,94514809 | 0,97689476 |
| 84676_at | TRIM63 | 0,00862991 | 3,89518719 | 0,06989315 | 0,94523185 | 0,97692891 |
| 51222_at | ZNF219 | -0,00775976 | 7,04599227 | -0,06976524 | 0,9453319 | 0,9769799 |
| 7326_at | UBE2G1 | -0,00655275 | 8,85639951 | -0,06956538 | 0,94548825 | 0,97708906 |
| 101928020_a | LOC10192802 | 0,00511612 | 2,80552338 | 0,06944025 | 0,94558613 | 0,97713779 |
| 125_at | ADH1B | 0,00675932 | 3,18829987 | 0,06933323 | 0,94566985 | 0,97717189 |
| 797_at | CALCB | 0,00650687 | 3,47414123 | 0,06917998 | 0,94578973 | 0,97719389 |
| 135138_at | PACRG | 0,00463543 | 3,80407909 | 0,06917631 | 0,9457926 | 0,97719389 |

| | | | | | | |
|---|---|---|---|---|---|---|
| 9148_at | NEURL1 | -0,00520282 | 5,44672551 | -0,06901316 | 0,94592023 | 0,97727334 |
| 54585_at | LZTFL1 | -0,01738131 | 5,91047852 | -0,06863625 | 0,94621509 | 0,97752554 |
| 7840_at | ALMS1 | 0,00493993 | 7,00854583 | 0,06854697 | 0,94628493 | 0,97754527 |
| 55222_at | LRRC20 | -0,01034436 | 6,96945741 | -0,06818843 | 0,94656543 | 0,9777826 |
| 100507135_a | SPG20-AS1 | 0,00598567 | 3,28710573 | 0,06809959 | 0,94663493 | 0,97780195 |
| 79366_at | HMGN5 | 0,0292647 | 4,36412557 | 0,06798066 | 0,94672797 | 0,97784562 |
| 79683_at | ZDHHC14 | -0,00695867 | 5,71303968 | -0,06789289 | 0,94679663 | 0,97784665 |
| 3624_at | INHBA | -0,00998616 | 5,1020251 | -0,06784421 | 0,94683472 | 0,97784665 |
| 81532_at | MOB2 | -0,00376405 | 6,59056438 | -0,06778151 | 0,94688378 | 0,97784665 |
| 9326_at | ZNHIT3 | -0,0078072 | 8,54973337 | -0,06770068 | 0,94694701 | 0,97784665 |
| 146513_at | LOC146513 | -0,00616873 | 3,78646767 | -0,06765497 | 0,94698278 | 0,97784665 |
| 389421_at | LIN28B | -0,00833982 | 3,25375675 | -0,06732122 | 0,94724389 | 0,97794219 |
| 10819_at | OR7E14P | 0,01154841 | 7,38938778 | 0,0673138 | 0,9472497 | 0,97794219 |
| 1452_at | CSNK1A1 | 0,01136303 | 10,4153536 | 0,06730447 | 0,947257 | 0,97794219 |
| 3064_at | HTT | 0,00512865 | 8,01608969 | 0,06727714 | 0,94727838 | 0,97794219 |
| 57407_at | NMRAL1 | 0,00624117 | 8,25933061 | 0,06702947 | 0,94747216 | 0,97808983 |
| 401288_at | LINC00242 | 0,01035101 | 5,14391892 | 0,06693381 | 0,947547 | 0,97809082 |
| 55323_at | LARP6 | 0,00698609 | 7,72662777 | 0,06689844 | 0,94757467 | 0,97809082 |
| 7703_at | PCGF2 | 0,00876275 | 7,78332968 | 0,06674251 | 0,94769667 | 0,97816151 |
| 100134713_a | NDUFB2-AS1 | -0,0068044 | 5,84463145 | -0,06664461 | 0,94777328 | 0,97816151 |
| 30833_at | NT5C | -0,0055817 | 7,4599204 | -0,06661622 | 0,94779549 | 0,97816151 |
| 100885781_a | LINC00348 | -0,00458611 | 2,96225951 | -0,06645623 | 0,94792067 | 0,97820067 |
| 114757_at | CYGB | -0,00700221 | 5,19472372 | -0,06635304 | 0,94800141 | 0,97820067 |
| 57514_at | ARHGAP31 | 0,00592899 | 4,07226221 | 0,0663365 | 0,94801435 | 0,97820067 |
| 101_at | ADAM8 | 0,00826593 | 5,9328667 | 0,06617752 | 0,94813874 | 0,97820067 |
| 1514_at | CTSL | -0,01269457 | 8,1121739 | -0,06614266 | 0,94816602 | 0,97820067 |
| 144608_at | C12orf60 | 0,0084467 | 4,53830095 | 0,06612323 | 0,94818122 | 0,97820067 |
| 2928_at | GSC2 | -0,00609073 | 4,55050918 | -0,06611341 | 0,9481889 | 0,97820067 |
| 23291_at | FBXW11 | 0,00709228 | 7,50192117 | 0,06575716 | 0,94846765 | 0,97843584 |
| 5261_at | PHKG2 | -0,00468157 | 6,23309062 | -0,06561684 | 0,94857746 | 0,97846308 |
| 123803_at | NTAN1 | 0,01665226 | 7,94824216 | 0,06557456 | 0,94861054 | 0,97846308 |
| 57805_at | CCAR2 | -0,02687623 | 8,79801467 | -0,06547724 | 0,94868669 | 0,97846308 |
| 101928988_a | LOC10192898 | 0,00932793 | 3,08287511 | 0,0654442 | 0,94871254 | 0,97846308 |
| 283120_at | H19 | 0,06253143 | 7,56330463 | 0,06539884 | 0,94874804 | 0,97846308 |
| 56675_at | NRIP3 | -0,02113597 | 5,74497301 | -0,06524453 | 0,94886878 | 0,9784845 |
| 643763_at | UG0898H09 | -0,0055649 | 3,1604455 | -0,06521428 | 0,94889245 | 0,9784845 |
| 84725_at | PLEKHA8 | 0,00538987 | 5,24871171 | 0,06517756 | 0,94892119 | 0,9784845 |
| 1842_at | ECM2 | 0,01013778 | 3,31241504 | 0,06495835 | 0,94909272 | 0,97860898 |
| 10411_at | RAPGEF3 | 0,01258339 | 6,41170977 | 0,06480063 | 0,94921614 | 0,97868385 |
| 9694_at | EMC2 | 0,01599989 | 8,79087594 | 0,06471653 | 0,94928194 | 0,97869932 |
| 26013_at | L3MBTL1 | -0,00894481 | 4,77473708 | -0,06463983 | 0,94934197 | 0,97870882 |
| 219931_at | TPCN2 | -0,00788355 | 6,57340492 | -0,06444956 | 0,94949086 | 0,9787232 |
| 55343_at | SLC35C1 | 0,01079491 | 7,36512035 | 0,06431489 | 0,94959625 | 0,9787232 |
| 146456_at | TMED6 | -0,00584793 | 3,7411662 | -0,06430552 | 0,94960358 | 0,9787232 |
| 1300_at | COL10A1 | -0,00536651 | 3,72373796 | -0,06425283 | 0,94964481 | 0,9787232 |
| 261729_at | STEAP2 | -0,0092893 | 4,43388023 | -0,06422752 | 0,94966462 | 0,9787232 |
| 101929524_a | LOC10192952 | 0,00549533 | 3,75361788 | 0,06421859 | 0,94967161 | 0,9787232 |
| 100507299_a | SMC5-AS1 | -0,00463081 | 4,53652298 | -0,06412239 | 0,94974689 | 0,9787232 |
| 11101_at | ATE1 | -0,00819828 | 7,32519968 | -0,06410259 | 0,94976239 | 0,9787232 |
| 100505641_a | FGD5-AS1 | 0,0051096 | 10,5076015 | 0,06371104 | 0,9500688 | 0,97898659 |

| | | | | | | |
|---|---|---|---|---|---|---|
| 101928795_a | LINC01448 | 0,00609666 | 3,2329449 | 0,06354466 | 0,950199 | 0,97901663 |
| 7038_at | TG | 0,00574287 | 4,9614356 | 0,06349188 | 0,95024031 | 0,97901663 |
| 254028_at | LOC254028 | -0,00389093 | 4,39506154 | -0,06347622 | 0,95025257 | 0,97901663 |
| 51557_at | LGSN | -0,00468274 | 2,98486289 | -0,06341402 | 0,95030125 | 0,97901663 |
| 100885848_a | PTGES3L | 0,0075542 | 3,50591233 | 0,06314216 | 0,950514 | 0,97914025 |
| 1399_at | CRKL | 0,00490577 | 8,00187938 | 0,06312615 | 0,95052653 | 0,97914025 |
| 5865_at | RAB3B | -0,00893634 | 4,23679754 | -0,06306584 | 0,95057373 | 0,97914025 |
| 101927348_a | LOC10192734 | -0,00613041 | 3,23618287 | -0,06270932 | 0,95085275 | 0,97937528 |
| 23152_at | CIC | -0,00496281 | 7,41670668 | -0,06258022 | 0,95095379 | 0,97942698 |
| 3694_at | ITGB6 | 0,01561359 | 8,81873027 | 0,06233828 | 0,95114315 | 0,97954385 |
| 84873_at | ADGRG7 | -0,00420237 | 3,08809786 | -0,06230529 | 0,95116896 | 0,97954385 |
| 9812_at | KIAA0141 | -0,00971229 | 7,20058887 | -0,06219237 | 0,95125734 | 0,9795825 |
| 124923_at | SGK494 | -0,01171885 | 5,76832245 | -0,06208683 | 0,95133995 | 0,97961519 |
| 340146_at | SLC35D3 | 0,00529073 | 4,22566307 | 0,0617933 | 0,95156968 | 0,97979938 |
| 13_at | AADAC | 0,02380316 | 6,9814437 | 0,06170848 | 0,95163607 | 0,97980819 |
| 4967_at | OGDH | -0,0045913 | 7,60510089 | -0,06165239 | 0,95167997 | 0,97980819 |
| 144108_at | SPTY2D1 | 0,00990137 | 6,42524278 | 0,06157527 | 0,95174033 | 0,97981797 |
| 285636_at | C5orf51 | 0,00880177 | 8,72477275 | 0,06144332 | 0,95184361 | 0,97987192 |
| 57228_at | SMAGP | 0,01994062 | 8,73986992 | 0,06128861 | 0,9519647 | 0,97992284 |
| 101928001_a | PRKCA-AS1 | 0,00545418 | 2,95819129 | 0,06125014 | 0,95199481 | 0,97992284 |
| 374355_at | CCDC172 | 0,00369102 | 3,30921008 | 0,06114393 | 0,95207794 | 0,97995604 |
| 26469_at | PTPN18 | 0,00778249 | 7,59141331 | 0,06107847 | 0,95212918 | 0,97995642 |
| 113691_at | TUBA3FP | -0,00810683 | 5,44768544 | -0,06084661 | 0,95231066 | 0,9800774 |
| 89874_at | SLC25A21 | -0,01586067 | 4,24180851 | -0,06069316 | 0,95243077 | 0,9800774 |
| 255426_at | RASGEF1C | 0,00519469 | 4,87132243 | 0,06065339 | 0,9524619 | 0,9800774 |
| 8170_at | SLC14A2 | -0,00666416 | 3,90459588 | -0,06047612 | 0,95260066 | 0,9800774 |
| 1915_at | EEF1A1 | 0,02114319 | 7,73383954 | 0,06042898 | 0,95263756 | 0,9800774 |
| 5813_at | PURA | -0,00852101 | 9,18172874 | -0,06038152 | 0,95267471 | 0,9800774 |
| 644246_at | KANSL1-AS1 | 0,01340557 | 6,08774454 | 0,06036406 | 0,95268838 | 0,9800774 |
| 57670_at | KIAA1549 | 0,00409942 | 6,24161883 | 0,06036066 | 0,95269103 | 0,9800774 |
| 7405_at | UVRAG | 0,00447272 | 7,46690072 | 0,06034329 | 0,95270463 | 0,9800774 |
| 84893_at | FBXO18 | 0,00979763 | 7,67470724 | 0,06016937 | 0,95284077 | 0,98016511 |
| 6262_at | RYR2 | 0,00383597 | 3,4362976 | 0,06003637 | 0,95294488 | 0,98021986 |
| 9412_at | MED21 | 0,01119081 | 8,87611859 | 0,05987949 | 0,95306768 | 0,98025926 |
| 65244_at | SPATS2 | 0,01392203 | 7,81890421 | 0,05985741 | 0,95308497 | 0,98025926 |
| 118427_at | OLFM3 | -0,00421208 | 3,07973669 | -0,05967766 | 0,95322567 | 0,98035163 |
| 203859_at | ANO5 | 0,00842973 | 4,17248444 | 0,05954497 | 0,95332953 | 0,98039505 |
| 79676_at | OGFOD2 | -0,00709195 | 7,60259897 | -0,05949369 | 0,95336968 | 0,98039505 |
| 219988_at | PATL1 | 0,01537567 | 8,64118864 | 0,05941743 | 0,95342938 | 0,9804041 |
| 50865_at | HEBP1 | 0,01305186 | 9,42850215 | 0,05917095 | 0,95362233 | 0,98050034 |
| 27098_at | CLUL1 | -0,00583124 | 3,59305747 | -0,05916782 | 0,95362477 | 0,98050034 |
| 51560_at | RAB6B | -0,012903 | 5,30371293 | -0,0589006 | 0,95383396 | 0,98066143 |
| 857_at | CAV1 | -0,03184731 | 10,3327704 | -0,05883762 | 0,95388326 | 0,98066143 |
| 7410_at | VAV2 | -0,00583745 | 6,88292982 | -0,05857231 | 0,95409095 | 0,98071029 |
| 89766_at | UMODL1 | 0,0052963 | 3,72471819 | 0,05846929 | 0,95417161 | 0,98071029 |
| 6334_at | SCN8A | -0,00335276 | 3,78813124 | -0,05846613 | 0,95417408 | 0,98071029 |
| 388341_at | LRRC75A | 0,01020192 | 4,96714296 | 0,05841882 | 0,95421111 | 0,98071029 |
| 728024_at | LOC728024 | -0,0098797 | 4,00177616 | -0,05838753 | 0,95423561 | 0,98071029 |
| 400618_at | SOX9-AS1 | -0,00399202 | 3,23866967 | -0,0583867 | 0,95423626 | 0,98071029 |
| 100129617_a | LOC10012961 | 0,00450535 | 6,18831762 | 0,05816332 | 0,95441114 | 0,98083769 |

| | | | | | | |
|---|---|---|---|---|---|---|
| 23142_at | DCUN1D4 | 0,01336098 | 6,79638466 | 0,05800428 | 0,95453564 | 0,98091331 |
| 123606_at | NIPA1 | 0,00415287 | 6,60003643 | 0,05791056 | 0,95460901 | 0,98093638 |
| 56479_at | KCNQ5 | -0,00356689 | 3,2585638 | -0,05767021 | 0,95479718 | 0,9810774 |
| 94025_at | MUC16 | 0,01533276 | 4,65752913 | 0,05746028 | 0,95496153 | 0,98119394 |
| 203062_at | TSNARE1 | -0,0073342 | 5,90174783 | -0,05734776 | 0,95504962 | 0,98119727 |
| 338004_at | LINC00226 | 0,00499621 | 3,27355267 | 0,05725228 | 0,95512437 | 0,98119727 |
| 139081_at | MAGEC3 | -0,00531303 | 3,83589194 | -0,05725059 | 0,9551257 | 0,98119727 |
| 30837_at | SOCS7 | 0,00346245 | 7,69632948 | 0,05719589 | 0,95516852 | 0,98119727 |
| 5977_at | DPF2 | -0,00359516 | 9,67315618 | -0,05704698 | 0,9552851 | 0,9812647 |
| 64324_at | NSD1 | 0,00439058 | 6,89384575 | 0,05671606 | 0,95554419 | 0,9814785 |
| 50856_at | CLEC4A | 0,00350759 | 3,68925608 | 0,05657591 | 0,95565392 | 0,98153222 |
| 100874194_a | CLDN10-AS1 | -0,00711098 | 3,40574669 | -0,0565191 | 0,9556984 | 0,98153222 |
| 348013_at | TMEM255B | -0,00632134 | 4,43866556 | -0,05634888 | 0,95583167 | 0,98157198 |
| 54902_at | TTC19 | 0,02372801 | 9,07576573 | 0,05633949 | 0,95583903 | 0,98157198 |
| 1401_at | CRP | 0,00368456 | 3,56260147 | 0,05612477 | 0,95600714 | 0,98169229 |
| 7504_at | XK | 0,01583059 | 4,25432784 | 0,05578767 | 0,95627108 | 0,98183295 |
| 10840_at | ALDH1L1 | -0,01786827 | 8,18788452 | -0,05573243 | 0,95631433 | 0,98183295 |
| 399694_at | SHC4 | 0,00305363 | 3,48910107 | 0,0557149 | 0,95632806 | 0,98183295 |
| 65123_at | INTS3 | 0,00594634 | 8,62986877 | 0,05563374 | 0,9563916 | 0,98183295 |
| 9970_at | NR1I3 | -0,00581861 | 4,10273192 | -0,05562432 | 0,95639898 | 0,98183295 |
| 148534_at | TMEM56 | -0,01169653 | 6,19154425 | -0,0555284 | 0,95647408 | 0,98185773 |
| 3236_at | HOXD10 | 0,00534338 | 4,97246278 | 0,05538314 | 0,95658782 | 0,98192216 |
| 101929460_a | LOC10192946 | 0,0045156 | 4,39708083 | 0,05526242 | 0,95668234 | 0,98196686 |
| 54914_at | FOCAD | 0,01708941 | 6,99392286 | 0,05506038 | 0,95684054 | 0,98201522 |
| 100874012_a | NAV2-AS5 | -0,00570234 | 3,24248358 | -0,05501673 | 0,95687472 | 0,98201522 |
| 23150_at | FRMD4B | 0,02591646 | 5,68514609 | 0,05500692 | 0,9568824 | 0,98201522 |
| 80072_at | HEXA-AS1 | 0,00484121 | 3,72453 | 0,05493815 | 0,95693625 | 0,98201817 |
| 101927157_a | LOC10192715 | -0,00797665 | 4,72293465 | -0,05472789 | 0,95710089 | 0,9821348 |
| 115817_at | DHRS1 | -0,02075438 | 8,0528743 | -0,0545142 | 0,95726822 | 0,98222806 |
| 3642_at | INSM1 | 0,0045331 | 4,53878872 | 0,05448158 | 0,95729376 | 0,98222806 |
| 6794_at | STK11 | 0,00346031 | 7,30000303 | 0,05411626 | 0,95757982 | 0,98246924 |
| 29951_at | PDZRN4 | 0,00432699 | 3,42313423 | 0,05388879 | 0,95775794 | 0,98251097 |
| 3299_at | HSF4 | 0,00807723 | 6,64504958 | 0,05387562 | 0,95776826 | 0,98251097 |
| 2560_at | GABRB1 | 0,00410922 | 3,22490705 | 0,05376153 | 0,95785759 | 0,98251097 |
| 4717_at | NDUFC1 | -0,01454588 | 7,36940314 | -0,05373936 | 0,95787496 | 0,98251097 |
| 245938_at | DEFB125 | -0,00542853 | 3,23002272 | -0,05373864 | 0,95787552 | 0,98251097 |
| 65243_at | ZFP69B | -0,00852616 | 4,82749742 | -0,05341814 | 0,9581265 | 0,98267674 |
| 57654_at | UVSSA | 0,00707014 | 6,76538372 | 0,05334985 | 0,95817997 | 0,98267674 |
| 29125_at | C11orf21 | 0,00618006 | 4,92041172 | 0,05333683 | 0,95819017 | 0,98267674 |
| 51534_at | VTA1 | -0,00479868 | 9,42505899 | -0,05296913 | 0,95847811 | 0,98290426 |
| 2139_at | EYA2 | -0,05605387 | 6,57200064 | -0,05292322 | 0,95851407 | 0,98290426 |
| 114609_at | TIRAP | -0,00263429 | 6,41098974 | -0,05256193 | 0,958797 | 0,98309609 |
| 7320_at | UBE2B | -0,00726013 | 8,24682642 | -0,052554 | 0,95880321 | 0,98309609 |
| 100507547_a | LOC10050754 | -0,01094882 | 5,21151183 | -0,05244322 | 0,95888996 | 0,98313271 |
| 283130_at | SLC25A45 | 0,0097728 | 5,31631071 | 0,0521883 | 0,9590896 | 0,98314716 |
| 6263_at | RYR3 | 0,00452979 | 3,6130202 | 0,05214546 | 0,95912315 | 0,98314716 |
| 100506540_a | SPTY2D1-AS1 | 0,00722271 | 3,86767737 | 0,05211659 | 0,95914576 | 0,98314716 |
| 26146_at | TRAF3IP1 | -0,0058557 | 6,64860423 | -0,05208786 | 0,95916826 | 0,98314716 |
| 100873942_a | PHEX-AS1 | 0,00494693 | 3,87500235 | 0,05205065 | 0,9591974 | 0,98314716 |
| 2998_at | GYS2 | -0,00339803 | 3,0477538 | -0,05203419 | 0,95921029 | 0,98314716 |

| | | | | | | |
|---|---|---|---|---|---|---|
| 171023_at | ASXL1 | 0,00403064 | 7,92319035 | 0,05178612 | 0,95940457 | 0,98328395 |
| 750_at | GAS8-AS1 | -0,00369064 | 3,76830034 | -0,05173343 | 0,95944583 | 0,98328395 |
| 51309_at | ARMCX1 | 0,00491025 | 3,56153084 | 0,05153389 | 0,9596021 | 0,98339178 |
| 54553_at | DKFZP434I07: | -0,00985778 | 6,59280895 | -0,05129015 | 0,95979299 | 0,98343831 |
| 6992_at | PPP1R11 | 0,00703993 | 9,53461456 | 0,05125384 | 0,95982143 | 0,98343831 |
| 1446_at | CSN1S1 | 0,00344903 | 3,19462822 | 0,05123335 | 0,95983748 | 0,98343831 |
| 474354_at | LRRC18 | -0,00524145 | 3,63378969 | -0,05115745 | 0,95989692 | 0,98343831 |
| 79814_at | AGMAT | 0,00487992 | 4,59219652 | 0,05110423 | 0,9599386 | 0,98343831 |
| 4653_at | MYOC | 0,00499045 | 4,41140108 | 0,05108479 | 0,95995383 | 0,98343831 |
| 5396_at | PRRX1 | -0,0042113 | 4,01815282 | -0,05086469 | 0,96012621 | 0,9835626 |
| 5598_at | MAPK7 | 0,0138164 | 7,11464194 | 0,05073664 | 0,9602265 | 0,98356288 |
| 9518_at | GDF15 | -0,03816232 | 8,97411763 | -0,05073395 | 0,9602286 | 0,98356288 |
| 199920_at | C1orf168 | 0,00339593 | 3,27924235 | 0,05065044 | 0,96029401 | 0,98357758 |
| 10752_at | CHL1 | -0,0027084 | 3,0164928 | -0,05055841 | 0,96036608 | 0,9835991 |
| 5871_at | MAP4K2 | 0,00760174 | 5,89977182 | 0,05042768 | 0,96046848 | 0,98364582 |
| 3988_at | LIPA | 0,01271446 | 10,7500459 | 0,05036978 | 0,96051382 | 0,98364582 |
| 7305_at | TYROBP | -0,00537393 | 4,55648094 | -0,05019343 | 0,96065195 | 0,98373496 |
| 142679_at | DUSP19 | 0,00649076 | 4,24964899 | 0,04991068 | 0,96087341 | 0,98383236 |
| 22800_at | RRAS2 | -0,00627104 | 8,68292513 | -0,04987538 | 0,96090106 | 0,98383236 |
| 5824_at | PEX19 | -0,00547999 | 8,56326964 | -0,04986815 | 0,96090672 | 0,98383236 |
| 1378_at | CR1 | -0,00309875 | 3,11504403 | -0,04977252 | 0,96098162 | 0,98383236 |
| 144423_at | GLT1D1 | -0,00570541 | 4,37803122 | -0,04974595 | 0,96100243 | 0,98383236 |
| 11325_at | DDX42 | -0,00427957 | 8,37054433 | -0,04951452 | 0,9611837 | 0,98392214 |
| 155435_at | RBM33 | -0,00434141 | 6,42404215 | -0,04950357 | 0,96119228 | 0,98392214 |
| 3177_at | SLC29A2 | -0,00825976 | 6,33598663 | -0,0494321 | 0,96124825 | 0,98392715 |
| 9314_at | KLF4 | -0,01922995 | 5,84784378 | -0,04922222 | 0,96141264 | 0,98402771 |
| 23281_at | MTUS2 | -0,02611827 | 5,31898142 | -0,04917623 | 0,96144866 | 0,98402771 |
| 6095_at | RORA | -0,00276439 | 3,42361416 | -0,04899708 | 0,96158899 | 0,98411904 |
| 400931_at | MIRLET7BHG | 0,00722096 | 7,05733229 | 0,04879793 | 0,96174498 | 0,98419146 |
| 10953_at | TOMM34 | 0,01094556 | 9,78622716 | 0,04871104 | 0,96181303 | 0,98419146 |
| 5973_at | RENBP | -0,00942286 | 5,14424414 | -0,0485685 | 0,96192469 | 0,98419146 |
| 3021_at | H3F3B | 0,0067984 | 8,49426461 | 0,04856707 | 0,96192581 | 0,98419146 |
| 338799_at | LINC01089 | -0,00851426 | 7,02885733 | -0,04855658 | 0,96193402 | 0,98419146 |
| 339290_at | LINC00667 | -0,008903 | 6,33279775 | -0,04851538 | 0,96196629 | 0,98419146 |
| 92597_at | MOB1B | 0,01669706 | 7,95352883 | 0,04813135 | 0,96226711 | 0,9844312 |
| 6476_at | SI | -0,00251982 | 2,79362961 | -0,04808574 | 0,96230283 | 0,9844312 |
| 56648_at | EIF5A2 | -0,02264658 | 5,23056524 | -0,04777764 | 0,96254418 | 0,9846258 |
| 51281_at | ANKMY1 | -0,00480925 | 5,34869627 | -0,04727883 | 0,96293491 | 0,9849732 |
| 26191_at | PTPN22 | -0,00256202 | 2,81315451 | -0,04709413 | 0,9630796 | 0,98505765 |
| 54539_at | NDUFB11 | 0,0062725 | 11,1323195 | 0,04699239 | 0,96315929 | 0,98505765 |
| 57512_at | GPR158 | -0,00651303 | 4,18193552 | -0,04697759 | 0,96317089 | 0,98505765 |
| 80233_at | FAAP100 | -0,00546303 | 7,11242859 | -0,04612184 | 0,96384126 | 0,98568602 |
| 79345_at | OR51B2 | -0,0045043 | 4,65667163 | -0,04606264 | 0,96388764 | 0,98568602 |
| 9575_at | CLOCK | 0,01430314 | 7,81705304 | 0,04572761 | 0,9641501 | 0,98587478 |
| 51362_at | CDC40 | -0,01026777 | 7,19688373 | -0,04569635 | 0,96417458 | 0,98587478 |
| 401577_at | CD99P1 | -0,00902987 | 6,66149121 | -0,04554595 | 0,96429241 | 0,98593641 |
| 219736_at | STOX1 | 0,0070225 | 4,26898168 | 0,04548499 | 0,96434017 | 0,98593641 |
| 55236_at | UBA6 | 0,010521 | 8,44525707 | 0,04538847 | 0,96441578 | 0,98593641 |
| 100506530_a | LOC10050653 | 0,00453733 | 3,38649818 | 0,04535808 | 0,96443959 | 0,98593641 |
| 3397_at | ID1 | 0,00910246 | 12,465947 | 0,0451487 | 0,96460363 | 0,98601747 |

| | | | | | | |
|---|---|---|---|---|---|---|
| 100192378_a | ZFHX4-AS1 | -0,00309217 | 3,20590139 | -0,04508804 | 0,96465115 | 0,98601747 |
| 1261_at | CNGA3 | -0,00447437 | 3,76476032 | -0,04502771 | 0,96469841 | 0,98601747 |
| 100129716_a | ARRDC3-AS1 | 0,00477125 | 4,6543426 | 0,04497147 | 0,96474247 | 0,98601747 |
| 4641_at | MYO1C | -0,00696221 | 8,31588238 | -0,04493018 | 0,96477482 | 0,98601747 |
| 79862_at | ZNF669 | -0,00818193 | 6,58877553 | -0,04469006 | 0,96496293 | 0,98611359 |
| 83891_at | SNX25 | 0,00752774 | 6,06464264 | 0,04463332 | 0,96500739 | 0,98611359 |
| 5000_at | ORC4 | -0,01617816 | 7,60365485 | -0,04461411 | 0,96502244 | 0,98611359 |
| 170261_at | ZCCHC12 | 0,00397666 | 3,66092439 | 0,04446299 | 0,96514083 | 0,98614915 |
| 3212_at | HOXB2 | 0,01860491 | 8,74903346 | 0,044439 | 0,96515963 | 0,98614915 |
| 340107_at | LOC340107 | -0,00444691 | 3,30564521 | -0,04422838 | 0,96532464 | 0,98623569 |
| 84988_at | PPP1R16A | -0,01372526 | 7,46075897 | -0,04416937 | 0,96537088 | 0,98623569 |
| 11161_at | C14orf1 | 0,00509853 | 10,4282679 | 0,04409525 | 0,96542895 | 0,98623569 |
| 143241_at | DYDC1 | 0,00381984 | 2,87121542 | 0,0440695 | 0,96544912 | 0,98623569 |
| 256691_at | MAMDC2 | -0,00380226 | 3,67013352 | -0,04381049 | 0,96565205 | 0,98639068 |
| 7727_at | ZNF174 | 0,00281842 | 6,06865836 | 0,04358132 | 0,96583159 | 0,98650898 |
| 118430_at | MUCL1 | -0,00687275 | 5,55000735 | -0,04353193 | 0,96587029 | 0,98650898 |
| 55290_at | BRF2 | 0,01242298 | 7,57308346 | 0,04329532 | 0,96605567 | 0,98657673 |
| 5283_at | PIGH | 0,00459571 | 7,36847652 | 0,04325941 | 0,96608381 | 0,98657673 |
| 256076_at | COL6A5 | 0,00279076 | 3,35035523 | 0,04325117 | 0,96609027 | 0,98657673 |
| 503569_at | RGMB-AS1 | -0,00317462 | 4,23527906 | -0,04311893 | 0,96619388 | 0,98663023 |
| 9517_at | SPTLC2 | -0,00347578 | 7,95840092 | -0,04281257 | 0,96643391 | 0,98674116 |
| 6635_at | SNRPE | 0,00278062 | 4,94195516 | 0,04276431 | 0,96647172 | 0,98674116 |
| 241_at | ALOX5AP | 0,00905084 | 4,87648515 | 0,04274374 | 0,96648784 | 0,98674116 |
| 56938_at | ARNTL2 | 0,00497888 | 7,07044602 | 0,04270624 | 0,96651722 | 0,98674116 |
| 79746_at | ECHDC3 | 0,0042035 | 4,62394849 | 0,04265339 | 0,96655863 | 0,98674116 |
| 55317_at | AP5S1 | 0,00348524 | 7,84996522 | 0,0425178 | 0,96666487 | 0,98679731 |
| 23451_at | SF3B1 | -0,0038802 | 10,0754546 | -0,0423942 | 0,96676171 | 0,98680141 |
| 89882_at | TPD52L3 | -0,00329114 | 3,72444021 | -0,04236556 | 0,96678415 | 0,98680141 |
| 55909_at | BIN3 | -0,00655696 | 7,40580825 | -0,04227093 | 0,9668583 | 0,98680141 |
| 54549_at | SDK2 | -0,00256532 | 4,7793719 | -0,04225116 | 0,96687379 | 0,98680141 |
| 388325_at | SCIMP | -0,00599938 | 4,31017141 | -0,04211155 | 0,96698318 | 0,98686077 |
| 89781_at | HPS4 | 0,0026363 | 6,77021464 | 0,04188607 | 0,96715985 | 0,98698878 |
| 5787_at | PTPRB | -0,00286831 | 4,85563502 | -0,04169156 | 0,96731225 | 0,98709201 |
| 100506436_a | SPAG5-AS1 | -0,00553797 | 4,64256959 | -0,04161725 | 0,96737048 | 0,98709914 |
| 7142_at | TNP2 | -0,00498877 | 3,57037268 | -0,04152142 | 0,96744557 | 0,98712347 |
| 222658_at | KCTD20 | 0,00931671 | 7,78466326 | 0,04133989 | 0,96758781 | 0,98720555 |
| 100506374_a | LOC10050637 | -0,00416626 | 4,98619868 | -0,04118266 | 0,96771101 | 0,98720555 |
| 10949_at | HNRNPA0 | -0,00540378 | 10,2637535 | -0,04117733 | 0,96771519 | 0,98720555 |
| 79191_at | IRX3 | 0,01002905 | 9,68876455 | 0,04111676 | 0,96776265 | 0,98720555 |
| 2990_at | GUSB | -0,00513566 | 9,12950611 | -0,04094316 | 0,96789868 | 0,98720555 |
| 29956_at | CERS2 | 0,00333375 | 10,4858563 | 0,04089779 | 0,96793423 | 0,98720555 |
| 728450_at | LINC00687 | 0,00548438 | 4,08610331 | 0,04089247 | 0,9679384 | 0,98720555 |
| 10743_at | RAI1 | 0,00805961 | 7,83762641 | 0,04076393 | 0,96803912 | 0,98720555 |
| 23087_at | TRIM35 | 0,01029436 | 7,03768244 | 0,04069537 | 0,96809284 | 0,98720555 |
| 8943_at | AP3D1 | 0,00302557 | 9,83902399 | 0,04066962 | 0,96811302 | 0,98720555 |
| 101928435_a | LOC10192843 | -0,00498935 | 4,42904871 | -0,04059727 | 0,96816971 | 0,98720555 |
| 57446_at | NDRG3 | -0,00934226 | 8,33472002 | -0,04059688 | 0,96817002 | 0,98720555 |
| 100133319_a | PRO1804 | -0,00384234 | 3,45284276 | -0,04056851 | 0,96819225 | 0,98720555 |
| 79091_at | METTL22 | -0,00419563 | 6,88695135 | -0,04042468 | 0,96830495 | 0,98722124 |
| 10928_at | RALBP1 | 0,00869812 | 7,88944869 | 0,04033347 | 0,96837643 | 0,98722124 |

| | | | | | | |
|---|---|---|---|---|---|---|
| 84985_at | FAM83A | -0,00607462 | 9,70862947 | -0,04027199 | 0,9684246 | 0,98722124 |
| 22808_at | MRAS | -0,00351515 | 5,09673416 | -0,04026012 | 0,9684339 | 0,98722124 |
| 3489_at | IGFBP6 | -0,02377312 | 10,5728311 | -0,04018239 | 0,96849481 | 0,98722124 |
| 6003_at | RGS13 | -0,00242646 | 3,03253341 | -0,04015645 | 0,96851513 | 0,98722124 |
| 4313_at | MMP2 | 0,01387758 | 9,73339207 | 0,03986625 | 0,96874254 | 0,98723527 |
| 25871_at | C3orf17 | -0,006523 | 7,50717225 | -0,03978695 | 0,96880468 | 0,98723527 |
| 22846_at | VASH1 | 0,00462964 | 5,22298955 | 0,03962891 | 0,96892852 | 0,98723527 |
| 57172_at | CAMK1G | -0,00386481 | 4,23568619 | -0,03942626 | 0,96908732 | 0,98723527 |
| 100505918_a | LOC10050591 | 0,00554827 | 4,40816041 | 0,0394253 | 0,96908808 | 0,98723527 |
| 286676_at | ILDR1 | 0,00397918 | 5,10135902 | 0,03931215 | 0,96917675 | 0,98723527 |
| 23014_at | FBXO21 | 0,00540191 | 9,20674553 | 0,03928367 | 0,96919906 | 0,98723527 |
| 9788_at | MTSS1 | -0,00461565 | 5,49804396 | -0,03927103 | 0,96920897 | 0,98723527 |
| 56888_at | KCMF1 | 0,0031235 | 7,82496554 | 0,03925305 | 0,96922306 | 0,98723527 |
| 100507156_a | LOC10050715 | -0,00638571 | 5,39576924 | -0,03921208 | 0,96925517 | 0,98723527 |
| 51617_at | HMP19 | -0,00464274 | 3,9189544 | -0,03917649 | 0,96928305 | 0,98723527 |
| 155465_at | AGR3 | 0,03622561 | 4,18177965 | 0,0391611 | 0,96929512 | 0,98723527 |
| 25920_at | NELFB | 0,00809412 | 9,18204644 | 0,03914231 | 0,96930984 | 0,98723527 |
| 81698_at | LINC00597 | 0,00425868 | 3,41096206 | 0,0390511 | 0,96938131 | 0,98723527 |
| 54970_at | TTC12 | 0,00420093 | 6,12385737 | 0,03900508 | 0,96941737 | 0,98723527 |
| 641372_at | ACOT6 | 0,00366024 | 3,64441883 | 0,03888866 | 0,96950861 | 0,98723527 |
| 80150_at | ASRGL1 | 0,01311662 | 6,00888976 | 0,03874908 | 0,96961799 | 0,98723527 |
| 100130855_a | USP3-AS1 | -0,00922197 | 4,0257097 | -0,03874155 | 0,96962389 | 0,98723527 |
| 83930_at | STARD3NL | 0,00937689 | 9,17957893 | 0,03874105 | 0,96962429 | 0,98723527 |
| 50852_at | TRAT1 | 0,00398535 | 2,96073549 | 0,03869046 | 0,96966393 | 0,98723527 |
| 8034_at | SLC25A16 | 0,00468609 | 5,97085427 | 0,03864191 | 0,96970198 | 0,98723527 |
| 10100_at | TSPAN2 | 0,01128894 | 6,21988387 | 0,03861963 | 0,96971944 | 0,98723527 |
| 642636_at | RAD21L1 | 0,00211207 | 2,78037086 | 0,03861426 | 0,96972364 | 0,98723527 |
| 100652857_a | TMCO5B | 0,00367851 | 2,90197669 | 0,0385176 | 0,96979939 | 0,98723527 |
| 9247_at | GCM2 | -0,00429361 | 3,81145259 | -0,03850386 | 0,96981016 | 0,98723527 |
| 25928_at | SOSTDC1 | -0,00320259 | 3,43444869 | -0,03834567 | 0,96993412 | 0,98730929 |
| 57057_at | TBX20 | -0,00229509 | 3,9207348 | -0,03825344 | 0,97000641 | 0,98733069 |
| 128853_at | DUSP15 | -0,004208 | 4,72589427 | -0,03803441 | 0,97017805 | 0,98743275 |
| 9261_at | MAPKAPK2 | -0,00357877 | 7,2341963 | -0,03799467 | 0,9702092 | 0,98743275 |
| 5170_at | PDPK1 | -0,00206347 | 8,76110006 | -0,03774499 | 0,97040487 | 0,98757972 |
| 129880_at | BBS5 | -0,00324459 | 3,56626112 | -0,03767046 | 0,97046327 | 0,98758698 |
| 65985_at | AACS | -0,00635957 | 8,63119976 | -0,03737318 | 0,97069625 | 0,98775022 |
| 4640_at | MYO1A | 0,00417677 | 4,53640321 | 0,03733492 | 0,97072623 | 0,98775022 |
| 114803_at | MYSM1 | -0,00535333 | 6,09163134 | -0,03709501 | 0,97091425 | 0,98786559 |
| 55689_at | YEATS2 | 0,00359009 | 7,73688412 | 0,03691824 | 0,97105278 | 0,98786559 |
| 4321_at | MMP12 | 0,00491519 | 4,27186414 | 0,03688902 | 0,97107568 | 0,98786559 |
| 64114_at | TMBIM1 | 0,00372662 | 11,4709424 | 0,03682536 | 0,97112558 | 0,98786559 |
| 56478_at | EIF4ENIF1 | -0,00327221 | 8,11698383 | -0,03678268 | 0,97115902 | 0,98786559 |
| 158809_at | MAGEB6 | -0,00321958 | 3,51540877 | -0,03674564 | 0,97118805 | 0,98786559 |
| 92292_at | GLYATL1 | -0,00316225 | 3,48174762 | -0,03673219 | 0,9711986 | 0,98786559 |
| 149603_at | RNF187 | 0,00454459 | 8,52978727 | 0,0364936 | 0,97138558 | 0,98796617 |
| 317701_at | VN1R2 | 0,00449737 | 3,53084421 | 0,03647513 | 0,97140006 | 0,98796617 |
| 54472_at | TOLLIP | -0,00714858 | 7,50175377 | -0,03633049 | 0,97151341 | 0,9880293 |
| 23007_at | PLCH1 | 0,00196882 | 3,46037342 | 0,03602498 | 0,97175285 | 0,98817128 |
| 55539_at | KCNQ1DN | -0,00308697 | 5,37209609 | -0,03597453 | 0,97179239 | 0,98817128 |
| 23148_at | NACAD | 0,00347828 | 5,43163174 | 0,03595599 | 0,97180692 | 0,98817128 |

| | | | | | | |
|---|---|---|---|---|---|---|
| #N/A | GS1-600G8,3 | 0,00260045 | 3,3641189 | 0,03581109 | 0,97192049 | 0,98822882 |
| 85001_at | MGC16275 | 0,00377492 | 4,50829483 | 0,03575287 | 0,97196612 | 0,98822882 |
| 101928989_a | LOC10192898 | -0,0024696 | 4,34882787 | -0,03547736 | 0,97218204 | 0,9883962 |
| 399474_at | TMEM200B | 0,00617108 | 6,3236008 | 0,03529886 | 0,97232194 | 0,98848626 |
| 419_at | ART3 | -0,00864469 | 3,49661795 | -0,0350615 | 0,97250798 | 0,98858295 |
| 824_at | CAPN2 | -0,01537946 | 11,1644766 | -0,03504656 | 0,97251969 | 0,98858295 |
| 22845_at | DOLK | 0,00423114 | 8,03892119 | 0,03497164 | 0,97257841 | 0,98858544 |
| 6919_at | TCEA2 | -0,00709828 | 7,47876631 | -0,03491247 | 0,97262479 | 0,98858544 |
| 93185_at | IGSF8 | 0,00307259 | 7,13875392 | 0,0347503 | 0,97275189 | 0,98866247 |
| 100272147_a | CMC4 | 0,00702993 | 7,88487924 | 0,03453103 | 0,97292375 | 0,98878496 |
| 102724316_a | SVIL-AS1 | 0,00371378 | 5,37410904 | 0,03430907 | 0,97309772 | 0,98889241 |
| 114780_at | PKD1L2 | -0,00621117 | 4,74513028 | -0,03426514 | 0,97313215 | 0,98889241 |
| 3055_at | HCK | 0,004379 | 4,87867867 | 0,03385975 | 0,97344989 | 0,98916312 |
| 151790_at | WDR49 | -0,00177323 | 2,89462036 | -0,03378761 | 0,97350644 | 0,98916692 |
| 23317_at | DNAJC13 | 0,00493836 | 7,21541601 | 0,03372395 | 0,97355633 | 0,98916692 |
| 7345_at | UCHL1 | 0,02932422 | 7,06144061 | 0,03364441 | 0,97361867 | 0,98917809 |
| 7643_at | ZNF90 | 0,00291794 | 3,02761414 | 0,03353229 | 0,97370656 | 0,9892152 |
| 100506380_a | LINC01006 | 0,00404444 | 4,33984451 | 0,0332688 | 0,97391308 | 0,98937283 |
| 79083_at | MLPH | -0,01231435 | 8,77377394 | -0,03312662 | 0,97402452 | 0,98942082 |
| 5947_at | RBP1 | 0,0032867 | 4,63279838 | 0,03306938 | 0,97406939 | 0,98942082 |
| 7205_at | TRIP6 | 0,01333488 | 9,24950453 | 0,03301195 | 0,97411441 | 0,98942082 |
| 116379_at | IL22RA2 | -0,0030526 | 3,03784999 | -0,03275454 | 0,97431618 | 0,98942585 |
| 101929557_a | LINC01395 | 0,00289669 | 3,73078288 | 0,03266045 | 0,97438992 | 0,98942585 |
| 151827_at | LRRC34 | 0,00227824 | 2,91222565 | 0,03264952 | 0,97439849 | 0,98942585 |
| 23231_at | SEL1L3 | 0,0087377 | 7,57992524 | 0,03264068 | 0,97440542 | 0,98942585 |
| 101928973_a | LOC10192897 | 0,00351519 | 5,13922852 | 0,03259469 | 0,97444147 | 0,98942585 |
| 80323_at | CCDC68 | -0,01627222 | 4,787841 | -0,03245468 | 0,97455121 | 0,98942585 |
| 80167_at | C4orf29 | 0,00679174 | 5,92679154 | 0,0324072 | 0,97458843 | 0,98942585 |
| 153339_at | TMEM167A | 0,00851365 | 9,60971069 | 0,0323959 | 0,97459729 | 0,98942585 |
| 3360_at | HTR4 | -0,0022564 | 4,06041345 | -0,03235106 | 0,97463244 | 0,98942585 |
| 56832_at | IFNK | 0,00216175 | 2,84956334 | 0,03231772 | 0,97465857 | 0,98942585 |
| 29957_at | SLC25A24 | -0,00308892 | 9,60039355 | -0,03228481 | 0,97468436 | 0,98942585 |
| 5141_at | PDE4A | 0,00241482 | 5,51999127 | 0,03217894 | 0,97476735 | 0,98945794 |
| 100129250_a | TOPORS-AS1 | 0,00658619 | 6,26201319 | 0,03202799 | 0,97488567 | 0,98951617 |
| 157769_at | FAM91A1 | -0,00609533 | 9,58805775 | -0,03193203 | 0,97496089 | 0,98951617 |
| 7036_at | TFR2 | -0,00369012 | 5,6232555 | -0,03190916 | 0,97497882 | 0,98951617 |
| 200232_at | FAM209A | 0,00328346 | 3,54349742 | 0,03184082 | 0,97503239 | 0,98951841 |
| 146802_at | SLC47A2 | 0,00486413 | 4,84760877 | 0,03170192 | 0,97514126 | 0,98957676 |
| 283932_at | FBXL19-AS1 | -0,00281496 | 4,58166655 | -0,03132345 | 0,97543794 | 0,98982568 |
| 5100_at | PCDH8 | 0,00266582 | 3,83016258 | 0,03123548 | 0,97550689 | 0,98983629 |
| 60677_at | CELF6 | 0,00307985 | 4,2265634 | 0,03111544 | 0,97560098 | 0,98983629 |
| 101927766_a | LOC10192776 | -0,00262833 | 4,38522648 | -0,03111045 | 0,9756049 | 0,98983629 |
| 90324_at | CCDC97 | -0,00207914 | 7,17013698 | -0,0310479 | 0,97565393 | 0,98983629 |
| 118812_at | MORN4 | 0,00278118 | 6,18328622 | 0,03091609 | 0,97575725 | 0,98988898 |
| 118_at | ADD1 | -0,0042818 | 9,16820484 | -0,03076441 | 0,97587615 | 0,98995747 |
| 239_at | ALOX12 | 0,01878078 | 5,94578398 | 0,03063541 | 0,97597727 | 0,98996636 |
| 79608_at | RIC3 | -0,00284742 | 3,88750914 | -0,0306221 | 0,9759877 | 0,98996636 |
| 100507267_a | LINC01187 | -0,00249387 | 3,16991931 | -0,03055466 | 0,97604057 | 0,98996786 |
| 203197_at | C9orf91 | 0,00497749 | 6,23679546 | 0,0301517 | 0,97635645 | 0,99020974 |
| 3633_at | INPP5B | -0,00285613 | 6,66310777 | -0,03010755 | 0,97639106 | 0,99020974 |

| | | | | | | |
|---|---|---|---|---|---|---|
| 203054_at | ADCK5 | -0,00606104 | 6,36977241 | -0,03003769 | 0,97644582 | 0,99020974 |
| 1611_at | DAP | 0,00254891 | 9,57045224 | 0,02998773 | 0,97648499 | 0,99020974 |
| 2774_at | GNAL | 0,00167641 | 3,87999446 | 0,02991615 | 0,9765411 | 0,99020974 |
| 161823_at | ADAL | -0,0032892 | 6,38685584 | -0,02981896 | 0,97661728 | 0,99020974 |
| 79758_at | DHRS12 | -0,00290313 | 5,51783655 | -0,02964993 | 0,97674979 | 0,99020974 |
| 23215_at | PRRC2C | 0,00366638 | 9,11220156 | 0,02964899 | 0,97675053 | 0,99020974 |
| 838_at | CASP5 | 0,00245745 | 4,57785935 | 0,02964674 | 0,97675229 | 0,99020974 |
| 54834_at | GDAP2 | -0,00369785 | 6,83086118 | -0,02943968 | 0,97691461 | 0,99020974 |
| 5514_at | PPP1R10 | 0,00487429 | 8,2093523 | 0,02939957 | 0,97694605 | 0,99020974 |
| 6833_at | ABCC8 | 0,0024513 | 5,63785299 | 0,02931373 | 0,97701334 | 0,99020974 |
| 51365_at | PLA1A | 0,00218338 | 3,60792338 | 0,02929242 | 0,97703005 | 0,99020974 |
| 55655_at | NLRP2 | -0,0282046 | 7,28379262 | -0,02927738 | 0,97704183 | 0,99020974 |
| 30062_at | RAX | -0,0030245 | 6,1029587 | -0,02918626 | 0,97711327 | 0,99020974 |
| 285989_at | ZNF789 | -0,00444996 | 5,58046909 | -0,02918174 | 0,97711681 | 0,99020974 |
| 22858_at | ICK | -0,00824338 | 6,71851074 | -0,02913566 | 0,97715293 | 0,99020974 |
| 6649_at | SOD3 | 0,00502122 | 5,82105439 | 0,02900719 | 0,97725364 | 0,99021946 |
| 3029_at | HAGH | 0,00594984 | 8,25688974 | 0,02899228 | 0,97726534 | 0,99021946 |
| 140947_at | DCANP1 | -0,00277194 | 4,03635252 | -0,02892316 | 0,97731952 | 0,99022228 |
| 51166_at | AADAT | 0,00751755 | 6,6301152 | 0,02884495 | 0,97738083 | 0,99022945 |
| 79816_at | TLE6 | 0,00708504 | 5,67160214 | 0,02878298 | 0,97742941 | 0,99022945 |
| 164237_at | WFDC13 | -0,00378274 | 3,53756498 | -0,02855337 | 0,97760941 | 0,99030563 |
| 2068_at | ERCC2 | 0,00272902 | 7,15330321 | 0,02849287 | 0,97765684 | 0,99030563 |
| 133686_at | NADK2 | 0,00648953 | 7,10474576 | 0,02846845 | 0,97767598 | 0,99030563 |
| 9830_at | TRIM14 | 0,00561737 | 8,4301312 | 0,02842474 | 0,97771025 | 0,99030563 |
| 100506858_a | LOC10050685 | -0,00318617 | 3,95596086 | -0,02832675 | 0,97778707 | 0,99033137 |
| 79365_at | BHLHE41 | 0,00946284 | 8,11566707 | 0,0279834 | 0,97805623 | 0,99051956 |
| 50853_at | VILL | -0,02228913 | 7,19049805 | -0,02795854 | 0,97807572 | 0,99051956 |
| 256536_at | TCERG1L | -0,00275178 | 4,35600491 | -0,02777391 | 0,97822046 | 0,99061407 |
| 2213_at | FCGR2B | -0,0043294 | 3,79772246 | -0,02756116 | 0,97838725 | 0,9906925 |
| 9450_at | LY86 | 0,003402 | 4,73809089 | 0,02754391 | 0,97840077 | 0,9906925 |
| 84859_at | LRCH3 | 0,00174003 | 6,45167077 | 0,02725087 | 0,97863051 | 0,99084757 |
| 115111_at | SLC26A7 | 0,00153474 | 2,99740007 | 0,02720015 | 0,97867027 | 0,99084757 |
| 23072_at | HECW1 | -0,00261137 | 4,62101915 | -0,02715173 | 0,97870823 | 0,99084757 |
| 84176_at | MYH16 | -0,00260908 | 3,90423376 | -0,0269121 | 0,9788961 | 0,9909333 |
| 115273_at | RAB42 | 0,00492453 | 3,70861566 | 0,02686806 | 0,97893062 | 0,9909333 |
| 64881_at | PCDH20 | 0,00199145 | 3,08576021 | 0,02684687 | 0,97894724 | 0,9909333 |
| 28660_at | TRAV23DV6 | -0,00207624 | 2,92350853 | -0,02666623 | 0,97908885 | 0,99098563 |
| 147837_at | ZNF563 | 0,00213579 | 4,06812739 | 0,02661663 | 0,97912774 | 0,99098563 |
| 400960_at | PCBP1-AS1 | 0,00222206 | 4,80831305 | 0,02658406 | 0,97915328 | 0,99098563 |
| 162966_at | ZNF600 | 0,00475359 | 6,560002 | 0,02624105 | 0,97942219 | 0,99116891 |
| 59352_at | LGR6 | 0,00368544 | 5,24843552 | 0,0262218 | 0,97943728 | 0,99116891 |
| 100862704_a | LINC00441 | -0,00228075 | 3,62977318 | -0,02613181 | 0,97950783 | 0,99117531 |
| 5366_at | PMAIP1 | -0,0097929 | 7,95481694 | -0,02599204 | 0,97961741 | 0,99117531 |
| 23434_at | LINC01565 | 0,00251309 | 5,40937504 | 0,02598111 | 0,97962598 | 0,99117531 |
| 134701_at | RIPPLY2 | -0,00227028 | 4,48620483 | -0,02593925 | 0,9796588 | 0,99117531 |
| 7162_at | TPBG | 0,00271128 | 9,35121471 | 0,02588559 | 0,97970087 | 0,99117531 |
| 284186_at | TMEM105 | 0,00273465 | 6,49588661 | 0,02575971 | 0,97979955 | 0,99118643 |
| 2999_at | GZMH | 0,00373367 | 4,60572702 | 0,02563451 | 0,97989771 | 0,99118643 |
| 55033_at | FKBP14 | 0,00478731 | 7,94323128 | 0,02556278 | 0,97995395 | 0,99118643 |
| 6944_at | VPS72 | -0,00222063 | 9,15268763 | -0,02555878 | 0,97995709 | 0,99118643 |

| | | | | | | |
|---|---|---|---|---|---|---|
| 27257_at | LSM1 | -0,00855659 | 8,68256118 | -0,0255434 | 0,97996914 | 0,99118643 |
| 8629_at | JRK | -0,00412359 | 6,43760342 | -0,02535537 | 0,98011656 | 0,99128349 |
| 1052_at | CEBPD | -0,00634491 | 9,40336061 | -0,02512819 | 0,98029467 | 0,99141158 |
| 728591_at | CCDC169 | 0,00159888 | 3,20067127 | 0,02505283 | 0,98035376 | 0,99141928 |
| 148229_at | ATP8B3 | 0,00250442 | 4,63286726 | 0,02490502 | 0,98046964 | 0,99148442 |
| 1000_at | CDH2 | -0,00273124 | 5,82864714 | -0,02465272 | 0,98066745 | 0,99163239 |
| 388698_at | FLG2 | 0,00210423 | 3,35061242 | 0,02449277 | 0,98079286 | 0,99170714 |
| 729799_at | SEC14L1P1 | -0,00453255 | 4,58538068 | -0,02437489 | 0,98088528 | 0,99174854 |
| 6330_at | SCN4B | -0,00230104 | 4,16589982 | -0,02414431 | 0,98106606 | 0,99184632 |
| 379_at | ARL4D | -0,00217754 | 7,7990832 | -0,02407628 | 0,9811194 | 0,99184632 |
| 91147_at | TMEM67 | 0,00338057 | 4,74920815 | 0,02402093 | 0,9811628 | 0,99184632 |
| 80035_at | ANP32A-IT1 | 0,00173577 | 3,84283138 | 0,02398884 | 0,98118795 | 0,99184632 |
| 1315_at | COPB1 | 0,0060571 | 10,2718751 | 0,02390823 | 0,98125116 | 0,99185816 |
| 2615_at | LRRC32 | -0,00229389 | 5,10842892 | -0,02367427 | 0,9814346 | 0,99199153 |
| 29850_at | TRPM5 | -0,00288255 | 4,58132177 | -0,02346967 | 0,98159501 | 0,99207349 |
| 84519_at | ACRBP | -0,00340358 | 4,8480237 | -0,02334201 | 0,9816951 | 0,99207349 |
| 399512_at | SLC25A35 | -0,00223961 | 6,2581818 | -0,02326002 | 0,98175939 | 0,99207349 |
| 6569_at | SLC34A1 | 0,00206143 | 5,20847834 | 0,02325015 | 0,98176712 | 0,99207349 |
| 10740_at | RFPL1S | 0,00251003 | 3,36233834 | 0,02324241 | 0,98177319 | 0,99207349 |
| 101929225_a | MCM8-AS1 | -0,00200112 | 3,09355996 | -0,02303447 | 0,98193623 | 0,99218619 |
| 9604_at | RNF14 | 0,00339602 | 7,75445282 | 0,02277186 | 0,98214213 | 0,99234219 |
| 7076_at | TIMP1 | 0,00358518 | 10,3471485 | 0,02267204 | 0,98222039 | 0,99236922 |
| 3033_at | HADH | -0,00832096 | 9,13632554 | -0,02251351 | 0,98234469 | 0,99243898 |
| 101928846_a | HIPK1-AS1 | -0,00145112 | 3,08369137 | -0,02245255 | 0,98239249 | 0,99243898 |
| 55346_at | TCP11L1 | -0,00465378 | 5,91465643 | -0,02231616 | 0,98249943 | 0,99244411 |
| 148641_at | SLC35F3 | 0,00142314 | 4,31487399 | 0,02231466 | 0,98250061 | 0,99244411 |
| 1428_at | CRYM | -0,00233993 | 5,57312859 | -0,02212183 | 0,9826518 | 0,9924916 |
| 6369_at | CCL24 | -0,00210519 | 4,94760864 | -0,02209644 | 0,9826717 | 0,9924916 |
| 6850_at | SYK | -0,00728692 | 7,25382571 | -0,02205539 | 0,98270389 | 0,9924916 |
| 3060_at | HCRT | 0,00227111 | 4,81466741 | 0,02199185 | 0,98275371 | 0,9924916 |
| 154796_at | AMOT | -0,00317084 | 6,38552409 | -0,02155699 | 0,98309468 | 0,99278389 |
| 8736_at | MYOM1 | 0,00249876 | 4,05923056 | 0,02121917 | 0,98335956 | 0,99293085 |
| 23316_at | CUX2 | -0,0022971 | 4,92122996 | -0,02120223 | 0,98337285 | 0,99293085 |
| 345895_at | RSPH4A | 0,00228414 | 3,79606632 | 0,02116126 | 0,98340497 | 0,99293085 |
| 100506730_a | LOC10050673 | -0,00247937 | 6,37531529 | -0,02110844 | 0,98344638 | 0,99293085 |
| 201163_at | FLCN | 0,00561921 | 6,75730502 | 0,02094472 | 0,98357475 | 0,99300796 |
| 201243_at | C17orf74 | -0,00173789 | 3,96240815 | -0,02087954 | 0,98362586 | 0,99300796 |
| 3904_at | LAIR2 | -0,00180303 | 4,2850615 | -0,02074126 | 0,98373429 | 0,99306538 |
| 5140_at | PDE3B | 0,00346824 | 5,48525058 | 0,02055465 | 0,98388061 | 0,99308329 |
| 400830_at | DEFB132 | 0,00217088 | 3,9760333 | 0,02054111 | 0,98389123 | 0,99308329 |
| 302_at | ANXA2 | -0,00161738 | 11,5125505 | -0,02052138 | 0,98390669 | 0,99308329 |
| 1733_at | DIO1 | -0,00335665 | 3,62706894 | -0,02027169 | 0,98410248 | 0,99321686 |
| 2205_at | FCER1A | 0,00136027 | 3,46340516 | 0,0202211 | 0,98414214 | 0,99321686 |
| 4617_at | MYF5 | -0,00188132 | 3,85800748 | -0,02004311 | 0,98428171 | 0,99326568 |
| 4151_at | MB | -0,01362343 | 8,49453222 | -0,02002789 | 0,98429364 | 0,99326568 |
| 6443_at | SGCB | 0,00769634 | 7,37438936 | 0,0197925 | 0,98447822 | 0,99339989 |
| 2892_at | GRIA3 | 0,00118452 | 3,07820566 | 0,019707 | 0,98454526 | 0,9934155 |
| 11318_at | GPR182 | -0,00138317 | 4,65721839 | -0,01960633 | 0,98462419 | 0,99344311 |
| 54763_at | ROPN1 | -0,00207126 | 3,64256364 | -0,01952644 | 0,98468684 | 0,99345428 |
| 140893_at | RBBP8NL | 0,001697 | 5,84013162 | 0,01924512 | 0,98490743 | 0,99353659 |

| | | | | | | |
|---|---|---|---|---|---|---|
| 57182_at | ANKRD50 | -0,00682949 | 8,29590496 | -0,01924299 | 0,9849091 | 0,99353659 |
| 154790_at | CLEC2L | 0,00189738 | 5,93161325 | 0,01922506 | 0,98492316 | 0,99353659 |
| 2262_at | GPC5 | -0,00105307 | 3,10684396 | -0,01910492 | 0,98501737 | 0,99357959 |
| 7092_at | TLL1 | 0,0014839 | 3,64370656 | 0,01827329 | 0,98566947 | 0,99413281 |
| 170302_at | ARX | -0,00204505 | 5,13327449 | -0,01825482 | 0,98568396 | 0,99413281 |
| 8573_at | CASK | -0,00176968 | 8,48603942 | -0,01816415 | 0,98575505 | 0,99413281 |
| 388552_at | BLOC1S3 | 0,00121908 | 7,45769241 | 0,01813174 | 0,98578047 | 0,99413281 |
| 4160_at | MC4R | -0,00244951 | 3,96684616 | -0,0180764 | 0,98582386 | 0,99413281 |
| 10845_at | CLPX | 0,00186742 | 8,03770089 | 0,01792088 | 0,98594581 | 0,99420375 |
| 285370_at | LINC00606 | -0,0012968 | 3,82890339 | -0,01784869 | 0,98600242 | 0,99420879 |
| 399909_at | PCNXL3 | 0,00115152 | 8,5020998 | 0,01764382 | 0,98616307 | 0,99431539 |
| 158696_at | LINC00889 | 0,00183036 | 3,28956942 | 0,01755575 | 0,98623213 | 0,99431539 |
| 8446_at | DUSP11 | -0,00302591 | 9,27785232 | -0,01742191 | 0,98633708 | 0,99431539 |
| 9409_at | PEX16 | -0,00338345 | 7,41167009 | -0,01738012 | 0,98636985 | 0,99431539 |
| 1630_at | DCC | -0,00105453 | 3,28253117 | -0,01731972 | 0,98641721 | 0,99431539 |
| 3936_at | LCP1 | -0,01158801 | 9,04643672 | -0,0173189 | 0,98641785 | 0,99431539 |
| 100506895_a | LOC10050689 | 0,00143295 | 3,83006304 | 0,01713647 | 0,98656091 | 0,99440756 |
| 51008_at | ASCC1 | 0,00128422 | 8,08635339 | 0,01670845 | 0,98689655 | 0,99469381 |
| 340596_at | LHFPL1 | -0,00161812 | 3,6162796 | -0,01648122 | 0,98707473 | 0,99473046 |
| 7137_at | TNNI3 | 0,00174273 | 4,92085013 | 0,01644551 | 0,98710273 | 0,99473046 |
| 83604_at | TMEM47 | -0,02128991 | 9,57161852 | -0,01637228 | 0,98716016 | 0,99473046 |
| 414332_at | LCN10 | 0,00196369 | 5,42785538 | 0,01625255 | 0,98725405 | 0,99473046 |
| 284013_at | VMO1 | -0,00366882 | 4,94816606 | -0,01623463 | 0,9872681 | 0,99473046 |
| 9079_at | LDB2 | 0,00113012 | 3,39449055 | 0,01622333 | 0,98727696 | 0,99473046 |
| 2299_at | FOXI1 | 0,00159292 | 4,60237723 | 0,01620112 | 0,98729438 | 0,99473046 |
| 10990_at | LILRB5 | -0,00183895 | 4,07801341 | -0,01598602 | 0,98746305 | 0,99484837 |
| 5211_at | PFKL | 0,00306947 | 8,86897224 | 0,01580499 | 0,98760501 | 0,99488249 |
| 79605_at | PGBD5 | 0,00202001 | 5,09721892 | 0,01565696 | 0,98772109 | 0,99488249 |
| 100505538_a | RBM26-AS1 | -0,00346889 | 4,80114097 | -0,01561073 | 0,98775735 | 0,99488249 |
| 839_at | CASP6 | -0,00404156 | 8,0612899 | -0,01560407 | 0,98776257 | 0,99488249 |
| 8995_at | TNFSF18 | -0,0016279 | 4,12691387 | -0,01554898 | 0,98780576 | 0,99488249 |
| 122046_at | TEX26 | -0,00123149 | 2,96979401 | -0,01554766 | 0,9878068 | 0,99488249 |
| 215_at | ABCD1 | -0,002309 | 6,95676443 | -0,01539081 | 0,9879298 | 0,99489188 |
| 101928314_a | LOC10192831 | -0,00100495 | 3,13771854 | -0,01536198 | 0,98795241 | 0,99489188 |
| 4790_at | NFKB1 | 0,00510889 | 8,44264687 | 0,01533181 | 0,98797607 | 0,99489188 |
| 196441_at | ZFC3H1 | 0,00397693 | 8,14297868 | 0,01527232 | 0,98802272 | 0,99489188 |
| 6372_at | CXCL6 | -0,00150381 | 3,7047121 | -0,01510954 | 0,98815037 | 0,99496841 |
| 390916_at | NUDT19 | 0,00246746 | 9,57754389 | 0,01503302 | 0,98821038 | 0,99497682 |
| 30011_at | SH3KBP1 | 0,00402197 | 7,71573222 | 0,01474062 | 0,98843967 | 0,99514188 |
| 201799_at | TMEM154 | 0,00372856 | 7,81867069 | 0,01469221 | 0,98847763 | 0,99514188 |
| 54541_at | DDIT4 | 0,00382507 | 10,9144554 | 0,01444378 | 0,98867245 | 0,99528599 |
| 54768_at | HYDIN | 0,00153042 | 4,21052615 | 0,0141322 | 0,98891679 | 0,99547994 |
| 8467_at | SMARCA5 | 0,00431199 | 9,10278519 | 0,01403674 | 0,98899165 | 0,99550328 |
| 170572_at | HTR3C | 0,00211389 | 4,6673947 | 0,01392164 | 0,98908191 | 0,9955421 |
| 100049716_a | LOC10004971 | 0,00232358 | 5,07160011 | 0,01383738 | 0,98914799 | 0,9955566 |
| 112950_at | MED8 | 0,00247393 | 7,99822736 | 0,01349692 | 0,98941498 | 0,99577329 |
| 152559_at | PAQR3 | -0,00414614 | 5,54339861 | -0,01339216 | 0,98949713 | 0,99579971 |
| 8403_at | SOX14 | 0,00131179 | 4,31709996 | 0,01325337 | 0,98960597 | 0,99579971 |
| 2993_at | GYPA | -0,00069212 | 3,34532108 | -0,01323977 | 0,98961664 | 0,99579971 |
| 6318_at | SERPINB4 | -0,00230553 | 4,91000846 | -0,01313555 | 0,98969837 | 0,99579971 |

| | | | | | | |
|---|---|---|---|---|---|---|
| 5928_at | RBBP4 | 0,00123765 | 7,51333515 | 0,01308865 | 0,98973514 | 0,99579971 |
| 140612_at | ZFP28 | 0,00067517 | 3,87212625 | 0,01297179 | 0,98982678 | 0,99579971 |
| 50838_at | TAS2R13 | 0,00121706 | 3,31105925 | 0,01295193 | 0,98984236 | 0,99579971 |
| 100132476_a | KRTAP4-7 | -0,00185797 | 4,6321404 | -0,01284906 | 0,98992303 | 0,99579971 |
| 1411_at | CRYBA1 | -0,0007424 | 3,27209071 | -0,01280229 | 0,98995971 | 0,99579971 |
| 29888_at | STRN4 | 0,00160498 | 7,39434044 | 0,01277158 | 0,98998379 | 0,99579971 |
| 9687_at | GREB1 | 0,00077921 | 3,85277828 | 0,01273832 | 0,99000988 | 0,99579971 |
| 137835_at | TMEM71 | 0,00102456 | 3,50152889 | 0,01260882 | 0,99011143 | 0,99584986 |
| 23138_at | N4BP3 | -0,00225062 | 5,01806931 | -0,01210613 | 0,99050565 | 0,99619435 |
| 2232_at | FDXR | -0,00133363 | 7,9428633 | -0,01202262 | 0,99057114 | 0,9962082 |
| 221078_at | NSUN6 | -0,00268215 | 5,46526381 | -0,01178713 | 0,99075582 | 0,99623974 |
| 8507_at | ENC1 | 0,0032554 | 7,5594528 | 0,01176775 | 0,99077101 | 0,99623974 |
| 146325_at | PRR35 | -0,00096496 | 6,69679469 | -0,01176738 | 0,99077131 | 0,99623974 |
| 146547_at | PRSS36 | -0,0013919 | 5,90016214 | -0,01171884 | 0,99080937 | 0,99623974 |
| 84171_at | LOXL4 | -0,00226682 | 6,10320543 | -0,01149205 | 0,99098722 | 0,99630536 |
| 100131691_a | MZF1-AS1 | -0,00116141 | 4,57415511 | -0,01144795 | 0,99102181 | 0,99630536 |
| 134860_at | TAAR9 | 0,00173356 | 3,98405622 | 0,01143777 | 0,99102979 | 0,99630536 |
| 101927547_a | ANKRD44-IT1 | 0,00094228 | 2,94576681 | 0,01108685 | 0,991305 | 0,99653001 |
| 58504_at | ARHGAP22 | -0,00101889 | 4,45139281 | -0,01078979 | 0,99153796 | 0,99654171 |
| 84529_at | C15orf41 | -0,00073907 | 6,86123929 | -0,01064293 | 0,99165313 | 0,99654171 |
| 4216_at | MAP3K4 | 0,00073168 | 8,43289117 | 0,01054069 | 0,99173331 | 0,99654171 |
| 5783_at | PTPN13 | 0,00333623 | 8,57078295 | 0,01045515 | 0,99180039 | 0,99654171 |
| 574036_at | SERTAD4-AS1 | -0,00160723 | 5,78501218 | -0,01036163 | 0,99187373 | 0,99654171 |
| 54435_at | HCG4 | 0,00079467 | 4,21690034 | 0,01023877 | 0,99197008 | 0,99654171 |
| 127254_at | ERICH3 | -0,00060599 | 3,02728744 | -0,01017043 | 0,99202368 | 0,99654171 |
| 656_at | BMP8B | 0,00131417 | 5,77002588 | 0,01016456 | 0,99202828 | 0,99654171 |
| 26154_at | ABCA12 | 0,00426719 | 5,72107092 | 0,01013339 | 0,99205273 | 0,99654171 |
| 84302_at | TMEM246 | 0,00566272 | 6,54431382 | 0,01011539 | 0,99206684 | 0,99654171 |
| 162333_at | MARCH10 | -0,00072367 | 4,15133454 | -0,01009319 | 0,99208426 | 0,99654171 |
| 29079_at | MED4 | -0,0016683 | 7,19837871 | -0,01005125 | 0,99211715 | 0,99654171 |
| 130872_at | AHSA2 | 0,0032258 | 6,57422886 | 0,01001754 | 0,99214358 | 0,99654171 |
| 10266_at | RAMP2 | 0,00131366 | 4,93119911 | 0,0100062 | 0,99215247 | 0,99654171 |
| 79802_at | HHIPL2 | 0,00076011 | 4,6832227 | 0,00992779 | 0,99221396 | 0,99654171 |
| 2950_at | GSTP1 | -0,00125246 | 12,1218773 | -0,00988096 | 0,99225069 | 0,99654171 |
| 80022_at | MYO15B | 0,00092935 | 5,812189 | 0,00987979 | 0,99225161 | 0,99654171 |
| 5468_at | PPARG | -0,000837 | 10,1562668 | -0,0098428 | 0,99228062 | 0,99654171 |
| 203328_at | SUSD3 | 0,00082284 | 5,08084228 | 0,00976903 | 0,99233847 | 0,99654171 |
| 113451_at | AZIN2 | 0,00071898 | 5,46777938 | 0,00975267 | 0,9923513 | 0,99654171 |
| 441733_at | PRKXP1 | 0,00067471 | 4,92549367 | 0,00942271 | 0,99261007 | 0,99674962 |
| 10008_at | KCNE3 | -0,00079486 | 3,82878559 | -0,00926867 | 0,99273088 | 0,99681896 |
| 83608_at | C18orf21 | 0,00113877 | 7,73145189 | 0,00884352 | 0,9930643 | 0,9970598 |
| 101926890_a | DAB1-AS1 | -0,00085245 | 3,88501977 | -0,00883083 | 0,99307425 | 0,9970598 |
| 54726_at | OTUD4 | -0,0021928 | 5,40501027 | -0,00875953 | 0,99313017 | 0,99706398 |
| 2246_at | FGF1 | 0,00060065 | 3,73007048 | 0,00853198 | 0,99330862 | 0,99719116 |
| 79640_at | C22orf46 | -0,00055842 | 5,9700078 | -0,00835158 | 0,9934501 | 0,99726315 |
| 527_at | ATP6V0C | 0,00100152 | 11,4555243 | 0,00830852 | 0,99348387 | 0,99726315 |
| 10580_at | SORBS1 | -0,00128706 | 5,09521929 | -0,00818862 | 0,99357791 | 0,99730557 |
| 105373991_a | LOC10537399 | 0,00137164 | 4,80642519 | 0,00797852 | 0,99374267 | 0,99737377 |
| 5339_at | PLEC | 0,00084584 | 7,22503751 | 0,00794108 | 0,99377204 | 0,99737377 |
| 147920_at | IGFL2 | 0,0019125 | 4,1505649 | 0,00790392 | 0,99380118 | 0,99737377 |

| | | | | | | |
|---|---|---|---|---|---|---|
| 135946_at | OR6B1 | 0,00112206 | 3,91513384 | 0,00767804 | 0,99397833 | 0,99749958 |
| 1370_at | CPN2 | -0,00095496 | 3,81701066 | -0,00754472 | 0,99408289 | 0,99750769 |
| 6956_at | TRAV6 | -0,00098211 | 3,59503437 | -0,00753568 | 0,99408997 | 0,99750769 |
| 810_at | CALML3 | -0,00057073 | 5,77614636 | -0,00734717 | 0,99423781 | 0,99760407 |
| 23626_at | SPO11 | 0,00049304 | 2,76407143 | 0,00725328 | 0,99431145 | 0,99761318 |
| 117145_at | THEM4 | -0,00499561 | 6,10570558 | -0,00720352 | 0,99435047 | 0,99761318 |
| 100131213_a | ZNF503-AS2 | -0,00062135 | 5,71385507 | -0,00694923 | 0,9945499 | 0,9977613 |
| 5861_at | RAB1A | -0,00074302 | 9,82865501 | -0,00688319 | 0,9946017 | 0,9977613 |
| 283596_at | SNHG10 | 0,00189277 | 5,92678089 | 0,00650071 | 0,99490166 | 0,99800486 |
| 375196_at | LOC375196 | -0,00055909 | 3,65275249 | -0,00644149 | 0,9949481 | 0,99800486 |
| 25930_at | PTPN23 | 0,00030086 | 6,63842158 | 0,00633925 | 0,99502828 | 0,9980078 |
| 5122_at | PCSK1 | -0,00063901 | 3,26241538 | -0,00630563 | 0,99505466 | 0,9980078 |
| 101929520_a | LINC00993 | -0,00071771 | 2,98097165 | -0,00622195 | 0,99512028 | 0,99802166 |
| 7984_at | ARHGEF5 | -0,00064325 | 8,07537806 | -0,00572379 | 0,99551097 | 0,99830938 |
| 260436_at | FDCSP | 0,00074526 | 3,8321736 | 0,00566205 | 0,99555939 | 0,99830938 |
| 84891_at | ZSCAN10 | -0,00059303 | 5,37255552 | -0,00562396 | 0,99558926 | 0,99830938 |
| 922_at | CD5L | 0,00047605 | 3,83445858 | 0,00552891 | 0,99566381 | 0,99830938 |
| 7288_at | TULP2 | 0,0004359 | 4,62378677 | 0,00552574 | 0,99566629 | 0,99830938 |
| 9475_at | ROCK2 | 0,00097136 | 8,23146318 | 0,0054107 | 0,99575652 | 0,99834788 |
| 149628_at | PYHIN1 | -0,00035783 | 3,40077808 | -0,00520985 | 0,99591404 | 0,99840148 |
| 283673_at | EWSAT1 | 0,00061399 | 4,93213483 | 0,0051545 | 0,99595745 | 0,99840148 |
| 79811_at | SLTM | -0,00038784 | 9,0868627 | -0,00504255 | 0,99604525 | 0,99840148 |
| 8546_at | AP3B1 | 0,00114654 | 8,06862075 | 0,00488708 | 0,99616718 | 0,99840148 |
| 2255_at | FGF10 | -0,00056154 | 3,85401692 | -0,00486874 | 0,99618156 | 0,99840148 |
| 30010_at | NXPH1 | -0,00046284 | 3,71467214 | -0,00484009 | 0,99620403 | 0,99840148 |
| 5879_at | RAC1 | 0,00043162 | 11,5087717 | 0,00481769 | 0,9962216 | 0,99840148 |
| 729083_at | LOC729083 | 0,0003021 | 3,36083095 | 0,00481384 | 0,99622462 | 0,99840148 |
| 29034_at | CPS1-IT1 | 0,00040468 | 3,76570388 | 0,00428284 | 0,99664107 | 0,99865921 |
| 101927815_a | LOC10192781 | -0,0003723 | 3,22451715 | -0,00428199 | 0,99664173 | 0,99865921 |
| 54796_at | BNC2 | -0,00032577 | 3,7678157 | -0,00424958 | 0,99666715 | 0,99865921 |
| 84966_at | IGSF21 | -0,00044387 | 4,46988377 | -0,00415311 | 0,99674281 | 0,99865921 |
| 89941_at | RHOT2 | 0,00028001 | 8,34141918 | 0,00410082 | 0,99678382 | 0,99865921 |
| 254268_at | AKNAD1 | -0,00025721 | 3,0872526 | -0,00403804 | 0,99683305 | 0,99865921 |
| 7102_at | TSPAN7 | 0,00235972 | 7,89877596 | 0,0040232 | 0,99684469 | 0,99865921 |
| 1910_at | EDNRB | -0,00028973 | 3,12814801 | -0,0038344 | 0,99699276 | 0,99875561 |
| 55086_at | CXorf57 | -0,00225561 | 8,07917691 | -0,00353274 | 0,99722935 | 0,99888226 |
| 153527_at | ZMAT2 | -0,00022095 | 9,02748173 | -0,0034885 | 0,99726404 | 0,99888226 |
| 23151_at | GRAMD4 | 0,00024286 | 6,53893681 | 0,00347484 | 0,99727476 | 0,99888226 |
| 80143_at | SIKE1 | 0,00091239 | 7,27787129 | 0,00325719 | 0,99744545 | 0,99900129 |
| 147687_at | ZNF417 | 0,00022536 | 4,03406953 | 0,00314965 | 0,9975298 | 0,99900318 |
| 92154_at | MTSS1L | -0,00035145 | 6,7049402 | -0,00312253 | 0,99755107 | 0,99900318 |
| 64781_at | CERK | 0,00154182 | 8,23556254 | 0,00304494 | 0,99761192 | 0,99900398 |
| 84640_at | USP38 | 0,00095539 | 7,6478177 | 0,00298925 | 0,99765559 | 0,99900398 |
| 28512_at | NKIRAS1 | 0,00059223 | 7,19610514 | 0,00276012 | 0,9978353 | 0,99913199 |
| 968_at | CD68 | 0,000641 | 6,61981359 | 0,0024766 | 0,99805765 | 0,9993027 |
| 100506627_a | DCDC5 | 0,00015101 | 2,82880564 | 0,00226021 | 0,99822736 | 0,99942067 |
| 8939_at | FUBP3 | -0,00021628 | 9,08261825 | -0,00204545 | 0,9983958 | 0,9994782 |
| 51026_at | GOLT1B | 0,00015088 | 8,14120584 | 0,0019373 | 0,99848061 | 0,9994782 |
| 7545_at | ZIC1 | 0,00010389 | 3,28463936 | 0,00184258 | 0,9985549 | 0,9994782 |
| 283592_at | FAM181A-AS1 | -0,00018944 | 3,90917805 | -0,00183451 | 0,99856123 | 0,9994782 |

| | | | | | | |
|---|---|---|---|---|---|---|
| 7978_at | MTERF1 | 0,00022165 | 6,57720516 | 0,00182488 | 0,99856879 | 0,9994782 |
| 51507_at | RTFDC1 | -0,0002115 | 9,13374663 | -0,00176888 | 0,9986127 | 0,9994782 |
| 101927224_a | LINC01049 | 0,00021099 | 4,22738481 | 0,00172384 | 0,99864802 | 0,9994782 |
| 222962_at | SLC29A4 | -0,00016873 | 5,65306673 | -0,00144119 | 0,9988697 | 0,99964812 |
| 55510_at | DDX43 | 6,96E-05 | 2,84109757 | 0,00106357 | 0,99916586 | 0,99982247 |
| 55816_at | DOK5 | -8,52E-05 | 3,83273668 | -0,00081551 | 0,99936041 | 0,99982247 |
| 100874375_a | FAM155A-IT1 | -8,14E-05 | 3,32153788 | -0,00078827 | 0,99938177 | 0,99982247 |
| 84657_at | LINC00852 | 7,53E-05 | 4,61352555 | 0,00072505 | 0,99943136 | 0,99982247 |
| 10911_at | UTS2 | -5,38E-05 | 3,10612371 | -0,00072482 | 0,99943154 | 0,99982247 |
| 6433_at | SFSWAP | -7,77E-05 | 8,17549444 | -0,00071337 | 0,99944052 | 0,99982247 |
| 728819_at | C1GALT1C1L | 0,00034898 | 4,33417729 | 0,00068086 | 0,99946601 | 0,99982247 |
| 415056_at | CNTFR-AS1 | -5,14E-05 | 4,14718842 | -0,0006398 | 0,99949822 | 0,99982247 |
| 196383_at | RILPL2 | -0,00014961 | 8,24063455 | -0,00060094 | 0,99952869 | 0,99982247 |
| 83450_at | LRRC48 | 4,64E-05 | 4,52306806 | 0,00051843 | 0,9995934 | 0,99982247 |
| 149047_at | MGC27382 | 4,86E-05 | 2,86595069 | 0,00049108 | 0,99961486 | 0,99982247 |
| 100190938_a | RAMP2-AS1 | -3,41E-05 | 4,05953408 | -0,00029807 | 0,99976623 | 0,99992195 |
| 1121_at | CHM | -1,66E-05 | 5,58877304 | -0,00019643 | 0,99984595 | 0,99994977 |
| 255403_at | ZNF718 | -1,58E-05 | 4,55523787 | -7,24E-05 | 0,99994325 | 0,9999636 |
| 114818_at | KLHL29 | 1,46E-05 | 7,05519754 | 4,64E-05 | 0,9999636 | 0,9999636 |

Supplementary Table 3. AR regulon gene set in NMIBC (UROMOL cohort)

| Regulon | Gene | Mode |
|---|---|---|
| AR | A4GALT | -1 |
| AR | ABCD3 | -1 |
| AR | ABHD12 | -1 |
| AR | ACKR2 | -1 |
| AR | ACOT7 | -1 |
| AR | ACSF2 | -1 |
| AR | ACSM3 | -1 |
| AR | ADAM8 | -1 |
| AR | ADAP1 | -1 |
| AR | ADCY1 | -1 |
| AR | AK2 | -1 |
| AR | AKR1B1 | -1 |
| AR | ALAS1 | -1 |
| AR | ALDH3B2 | -1 |
| AR | AMIGO2 | -1 |
| AR | AMOTL2 | -1 |
| AR | ANKDD1A | -1 |
| AR | ANKRD22 | -1 |
| AR | ANKRD35 | -1 |
| AR | ANKRD65 | -1 |
| AR | ANXA9 | -1 |
| AR | AP1B1 | -1 |
| AR | APLP2 | -1 |
| AR | APOBEC3G | -1 |
| AR | APOBEC3H | -1 |
| AR | APOBR | -1 |
| AR | ARHGAP10 | -1 |
| AR | ARHGAP40 | -1 |
| AR | ARID3A | -1 |
| AR | ARL4D | -1 |
| AR | ARRB1 | -1 |
| AR | ATG9B | -1 |
| AR | ATP2C2 | -1 |
| AR | AUNIP | -1 |
| AR | BAIAP2L2 | -1 |
| AR | BAIAP3 | -1 |
| AR | BCAR3 | -1 |
| AR | BCAS4 | -1 |
| AR | BCAT2 | -1 |
| AR | BCL2L1 | -1 |
| AR | BCL7A | -1 |
| AR | BLM | -1 |
| AR | BLNK | -1 |
| AR | BNIPL | -1 |
| AR | BPGM | -1 |
| AR | BTBD3 | -1 |

| | | |
|---|---|---|
| AR | C16orf87 | -1 |
| AR | C1QTNF6 | -1 |
| AR | CABP4 | -1 |
| AR | CACNA1D | -1 |
| AR | CACNG4 | -1 |
| AR | CAMK2G | -1 |
| AR | CAPG | -1 |
| AR | CATSPER1 | -1 |
| AR | CBX6 | -1 |
| AR | CCNE1 | -1 |
| AR | CCNJL | -1 |
| AR | CCR7 | -1 |
| AR | CDC42EP3 | -1 |
| AR | CDK2 | -1 |
| AR | CDR2L | -1 |
| AR | CDS1 | -1 |
| AR | CEBPA | -1 |
| AR | CECR2 | -1 |
| AR | CEP250 | -1 |
| AR | CEP85 | -1 |
| AR | CERS3 | -1 |
| AR | CHI3L2 | -1 |
| AR | CHST3 | -1 |
| AR | CITED4 | -1 |
| AR | CKLF | -1 |
| AR | CLDN4 | -1 |
| AR | CLDN9 | -1 |
| AR | CLIP2 | -1 |
| AR | CLN3 | -1 |
| AR | CNGB1 | -1 |
| AR | CPZ | -1 |
| AR | CRYBG2 | -1 |
| AR | CST1 | -1 |
| AR | CST4 | -1 |
| AR | CST6 | -1 |
| AR | CXorf40A | -1 |
| AR | CXXC5 | -1 |
| AR | CYP2J2 | -1 |
| AR | CYP3A7 | -1 |
| AR | CYSRT1 | -1 |
| AR | CYSTM1 | -1 |
| AR | DENND2C | -1 |
| AR | DENND2D | -1 |
| AR | DEPP1 | -1 |
| AR | DIPK1A | -1 |
| AR | DMXL2 | -1 |
| AR | DNMT3B | -1 |
| AR | DOCK3 | -1 |
| AR | DRD1 | -1 |
| AR | DUSP5 | -1 |

| | | |
|---|---|---|
| AR | DUSP9 | -1 |
| AR | E2F1 | -1 |
| AR | EMP2 | -1 |
| AR | ENDOD1 | -1 |
| AR | ENGASE | -1 |
| AR | ENTPD5 | -1 |
| AR | EPB41L1 | -1 |
| AR | EPHB2 | -1 |
| AR | EPHB3 | -1 |
| AR | EPS8L3 | -1 |
| AR | ERBB3 | -1 |
| AR | ERP27 | -1 |
| AR | ERRFI1 | -1 |
| AR | EVX1 | -1 |
| AR | FAM107B | -1 |
| AR | FAM118A | -1 |
| AR | FAM198B-AS1 | -1 |
| AR | FAM25A | -1 |
| AR | FAM3C | -1 |
| AR | FAM49A | -1 |
| AR | FBXO27 | -1 |
| AR | FCRLB | -1 |
| AR | FER1L4 | -1 |
| AR | FLVCR2 | -1 |
| AR | FMO9P | -1 |
| AR | FOLR1 | -1 |
| AR | FREM2 | -1 |
| AR | FUT1 | -1 |
| AR | GALE | -1 |
| AR | GCNT4 | -1 |
| AR | GDA | -1 |
| AR | GDF15 | -1 |
| AR | GDPD3 | -1 |
| AR | GNE | -1 |
| AR | GPR78 | -1 |
| AR | GPT2 | -1 |
| AR | GSDMA | -1 |
| AR | HABP2 | -1 |
| AR | HAGH | -1 |
| AR | HAVCR1 | -1 |
| AR | HCAR1 | -1 |
| AR | HCAR2 | -1 |
| AR | HCAR3 | -1 |
| AR | HCLS1 | -1 |
| AR | HID1 | -1 |
| AR | HIP1R | -1 |
| AR | HIST3H2A | -1 |
| AR | HMGCS2 | -1 |
| AR | HSD17B14 | -1 |
| AR | HSD3B1 | -1 |

| | | |
|---|---|---|
| AR | HSPB8 | -1 |
| AR | HTRA3 | -1 |
| AR | HYAL4 | -1 |
| AR | IER5 | -1 |
| AR | IFFO2 | -1 |
| AR | IGDCC3 | -1 |
| AR | IGFL2-AS1 | -1 |
| AR | IGSF3 | -1 |
| AR | IGSF9 | -1 |
| AR | IL1A | -1 |
| AR | IL1B | -1 |
| AR | IL22RA1 | -1 |
| AR | IL36RN | -1 |
| AR | IL6R | -1 |
| AR | INPP4A | -1 |
| AR | INPP4B | -1 |
| AR | INPP5A | -1 |
| AR | IQCD | -1 |
| AR | IRAK2 | -1 |
| AR | IRF5 | -1 |
| AR | ITSN2 | -1 |
| AR | KCNG1 | -1 |
| AR | KCNN4 | -1 |
| AR | KCTD17 | -1 |
| AR | KIFC3 | -1 |
| AR | KLHDC7A | -1 |
| AR | KLHDC8B | -1 |
| AR | KLHL6 | -1 |
| AR | KRT18 | -1 |
| AR | KRT23 | -1 |
| AR | KRT33B | -1 |
| AR | LACTB2 | -1 |
| AR | LCE1E | -1 |
| AR | LINC00456 | -1 |
| AR | LINC00475 | -1 |
| AR | LINC00482 | -1 |
| AR | LINC00840 | -1 |
| AR | LINC01443 | -1 |
| AR | LINC01541 | -1 |
| AR | LINC01764 | -1 |
| AR | LPAR3 | -1 |
| AR | LPAR5 | -1 |
| AR | LYST | -1 |
| AR | MAFF | -1 |
| AR | MAN1C1 | -1 |
| AR | MAP2K1 | -1 |
| AR | MAP3K9 | -1 |
| AR | MCRIP1 | -1 |
| AR | MDM2 | -1 |
| AR | MMP19 | -1 |

| | | |
|---|---|---|
| AR | MMRN2 | -1 |
| AR | MOCS1 | -1 |
| AR | MOSPD1 | -1 |
| AR | MPHOSPH6 | -1 |
| AR | MTCP1 | -1 |
| AR | MTSS1L | -1 |
| AR | NAB2 | -1 |
| AR | NAGK | -1 |
| AR | NCEH1 | -1 |
| AR | NDRG2 | -1 |
| AR | NEDD9 | -1 |
| AR | NGEF | -1 |
| AR | NOCT | -1 |
| AR | OTOF | -1 |
| AR | OVGP1 | -1 |
| AR | P2RY2 | -1 |
| AR | PALM3 | -1 |
| AR | PAQR5 | -1 |
| AR | PAQR7 | -1 |
| AR | PARD6B | -1 |
| AR | PCSK6 | -1 |
| AR | PDZD3 | -1 |
| AR | PELI1 | -1 |
| AR | PHLDB3 | -1 |
| AR | PI4K2A | -1 |
| AR | PKIA | -1 |
| AR | PLA2G15 | -1 |
| AR | PLA2G2F | -1 |
| AR | PLAU | -1 |
| AR | PLCXD1 | -1 |
| AR | PLEKHH1 | -1 |
| AR | PLPP1 | -1 |
| AR | PNPLA1 | -1 |
| AR | PNPLA8 | -1 |
| AR | POU3F1 | -1 |
| AR | PPCDC | -1 |
| AR | PPP1R14C | -1 |
| AR | PPP1R14D | -1 |
| AR | PPP1R3C | -1 |
| AR | PPP2R2D | -1 |
| AR | PPP4R1 | -1 |
| AR | PQLC2 | -1 |
| AR | PRAG1 | -1 |
| AR | PTK2B | -1 |
| AR | PTPRE | -1 |
| AR | PVALB | -1 |
| AR | QPRT | -1 |
| AR | RAB11FIP4 | -1 |
| AR | RAB20 | -1 |
| AR | RAI14 | -1 |

| | | |
|---|---|---|
| AR | RBBP8NL | -1 |
| AR | RCC1 | -1 |
| AR | REG4 | -1 |
| AR | RFC5 | -1 |
| AR | RFX2 | -1 |
| AR | RGS16 | -1 |
| AR | RHBG | -1 |
| AR | RNASE7 | -1 |
| AR | RNF186 | -1 |
| AR | RNF223 | -1 |
| AR | RPIA | -1 |
| AR | RRP1 | -1 |
| AR | S100A10 | -1 |
| AR | S100A4 | -1 |
| AR | SCARA3 | -1 |
| AR | SCPEP1 | -1 |
| AR | SEMA3B | -1 |
| AR | SEMA7A | -1 |
| AR | SGK2 | -1 |
| AR | SGPL1 | -1 |
| AR | SGSM3 | -1 |
| AR | SH3BP5 | -1 |
| AR | SH3TC2 | -1 |
| AR | SH3TC2-DT | -1 |
| AR | SIGLEC5 | -1 |
| AR | SLC11A2 | -1 |
| AR | SLC1A6 | -1 |
| AR | SLC1A7 | -1 |
| AR | SLC20A1 | -1 |
| AR | SLC27A2 | -1 |
| AR | SLC29A1 | -1 |
| AR | SLC2A6 | -1 |
| AR | SLC30A2 | -1 |
| AR | SLC31A2 | -1 |
| AR | SLC35E4 | -1 |
| AR | SLC39A2 | -1 |
| AR | SLC46A1 | -1 |
| AR | SLC4A3 | -1 |
| AR | SLC6A2 | -1 |
| AR | SLC6A6 | -1 |
| AR | SLC9A7 | -1 |
| AR | SLCO4A1 | -1 |
| AR | SMOC1 | -1 |
| AR | SNCG | -1 |
| AR | SOWAHB | -1 |
| AR | SPAG9 | -1 |
| AR | SPATA13 | -1 |
| AR | SRC | -1 |
| AR | ST3GAL4 | -1 |
| AR | STAMBPL1 | -1 |

| | | |
|---|---|---|
| AR | STK32C | -1 |
| AR | STX3 | -1 |
| AR | SULT2A1 | -1 |
| AR | SUN2 | -1 |
| AR | SVOPL | -1 |
| AR | SYNJ2 | -1 |
| AR | SYT7 | -1 |
| AR | TAGLN2 | -1 |
| AR | TBC1D30 | -1 |
| AR | TBC1D8 | -1 |
| AR | TDO2 | -1 |
| AR | TDRKH | -1 |
| AR | TEPP | -1 |
| AR | TESK2 | -1 |
| AR | TH | -1 |
| AR | TIGD2 | -1 |
| AR | TMEM184B | -1 |
| AR | TMEM265 | -1 |
| AR | TMEM40 | -1 |
| AR | TMEM62 | -1 |
| AR | TMEM97 | -1 |
| AR | TMPRSS13 | -1 |
| AR | TMPRSS6 | -1 |
| AR | TNFRSF12A | -1 |
| AR | TOMM34 | -1 |
| AR | TPST2 | -1 |
| AR | TRAFD1 | -1 |
| AR | TRIB3 | -1 |
| AR | TRIM17 | -1 |
| AR | TRNP1 | -1 |
| AR | TRPV3 | -1 |
| AR | TSPAN1 | -1 |
| AR | TTC39A | -1 |
| AR | TTC7B | -1 |
| AR | TUFT1 | -1 |
| AR | UBASH3B | -1 |
| AR | UBXN10 | -1 |
| AR | UCA1 | -1 |
| AR | UNC13D | -1 |
| AR | UPK3A | -1 |
| AR | USH1G | -1 |
| AR | UXS1 | -1 |
| AR | VCL | -1 |
| AR | VGLL1 | -1 |
| AR | VGLL3 | -1 |
| AR | VRK1 | -1 |
| AR | VSIG10 | -1 |
| AR | VWA1 | -1 |
| AR | VWA5B1 | -1 |
| AR | WDR77 | -1 |

| | | |
|---|---|---|
| AR | WDR91 | -1 |
| AR | WNT7A | -1 |
| AR | WWC1 | -1 |
| AR | ZBTB8B | -1 |
| AR | ZC3H12A | -1 |
| AR | ZDHHC8P1 | -1 |
| AR | ZNF175 | -1 |
| AR | ZNF319 | -1 |
| AR | ZNF350-AS1 | -1 |
| AR | ZNF432 | -1 |
| AR | ZNF488 | -1 |
| AR | ZNF823 | -1 |
| AR | ZNF841 | -1 |
| AR | ZNRF1 | -1 |
| AR | AASS | 1 |
| AR | ACSS3 | 1 |
| AR | ACTRT3 | 1 |
| AR | ACVR1 | 1 |
| AR | ADGRB3 | 1 |
| AR | ADGRL2 | 1 |
| AR | ADGRV1 | 1 |
| AR | AGGF1P1 | 1 |
| AR | AGGF1P2 | 1 |
| AR | AGGF1P3 | 1 |
| AR | ALCAM | 1 |
| AR | AMN | 1 |
| AR | ANKRD50 | 1 |
| AR | ANKRD7 | 1 |
| AR | ANO5 | 1 |
| AR | APCDD1 | 1 |
| AR | ARHGEF9 | 1 |
| AR | ARL10 | 1 |
| AR | ARMCX2 | 1 |
| AR | ARMCX5-GPRASP2 | 1 |
| AR | ATL1 | 1 |
| AR | B3GNT7 | 1 |
| AR | BAALC | 1 |
| AR | BBX | 1 |
| AR | BCL11B | 1 |
| AR | BEX5 | 1 |
| AR | BRWD3 | 1 |
| AR | C1GALT1C1L | 1 |
| AR | C1RL | 1 |
| AR | CACNA2D1 | 1 |
| AR | CARD16 | 1 |
| AR | CARMIL1 | 1 |
| AR | CASP1 | 1 |
| AR | CCDC149 | 1 |
| AR | CCDC181 | 1 |
| AR | CDC42BPA | 1 |

| | | |
|---|---|---|
| AR | CDCA7L | 1 |
| AR | CDH12 | 1 |
| AR | CEP68 | 1 |
| AR | CFLAR | 1 |
| AR | CHCHD10 | 1 |
| AR | CLYBL | 1 |
| AR | CMBL | 1 |
| AR | CNR1 | 1 |
| AR | CNTLN | 1 |
| AR | CPE | 1 |
| AR | CPNE7 | 1 |
| AR | CRNDE | 1 |
| AR | CTBP2 | 1 |
| AR | CXXC4 | 1 |
| AR | DANT2 | 1 |
| AR | DAPL1 | 1 |
| AR | DHRS4-AS1 | 1 |
| AR | DISC1 | 1 |
| AR | DMRTA1 | 1 |
| AR | DPY19L2P4 | 1 |
| AR | DPYSL3 | 1 |
| AR | DSC3 | 1 |
| AR | DTNA | 1 |
| AR | DTWD1 | 1 |
| AR | DYRK4 | 1 |
| AR | EBF4 | 1 |
| AR | EDA | 1 |
| AR | EFNB2 | 1 |
| AR | EIF4EBP3 | 1 |
| AR | ENPP5 | 1 |
| AR | ERC2 | 1 |
| AR | ESRRG | 1 |
| AR | ETNK2 | 1 |
| AR | EVC2 | 1 |
| AR | FAHD2B | 1 |
| AR | FAM155B | 1 |
| AR | FAM81B | 1 |
| AR | FAT2 | 1 |
| AR | FBLIM1 | 1 |
| AR | GALNT12 | 1 |
| AR | GALNT5 | 1 |
| AR | GAREM2 | 1 |
| AR | GDF11 | 1 |
| AR | GDPD2 | 1 |
| AR | GPR155 | 1 |
| AR | GPR63 | 1 |
| AR | GSPT2 | 1 |
| AR | GSTA4 | 1 |
| AR | HDX | 1 |
| AR | HIPK2 | 1 |

| | | |
|---|---|---|
| AR | HMGN5 | 1 |
| AR | HOXD3 | 1 |
| AR | HTR1B | 1 |
| AR | IFI16 | 1 |
| AR | IPCEF1 | 1 |
| AR | IQCA1 | 1 |
| AR | IRAK1BP1 | 1 |
| AR | IRAK3 | 1 |
| AR | IRS1 | 1 |
| AR | IRX3 | 1 |
| AR | IRX5 | 1 |
| AR | JAZF1 | 1 |
| AR | KATNAL2 | 1 |
| AR | KCTD1 | 1 |
| AR | KIAA0825 | 1 |
| AR | KIAA1147 | 1 |
| AR | KLF12 | 1 |
| AR | KLF3-AS1 | 1 |
| AR | KLF5 | 1 |
| AR | KRT13 | 1 |
| AR | KRT15 | 1 |
| AR | KRT18P58 | 1 |
| AR | KRT5 | 1 |
| AR | L3MBTL3 | 1 |
| AR | LANCL3 | 1 |
| AR | LDLRAD4 | 1 |
| AR | LGALS9 | 1 |
| AR | LINC00365 | 1 |
| AR | LINC00865 | 1 |
| AR | LINC00960 | 1 |
| AR | LINC01410 | 1 |
| AR | LINC01515 | 1 |
| AR | LINC01550 | 1 |
| AR | LINC01725 | 1 |
| AR | LINC02198 | 1 |
| AR | LMCD1-AS1 | 1 |
| AR | LNPEP | 1 |
| AR | LPAL2 | 1 |
| AR | LPP-AS2 | 1 |
| AR | LRP12 | 1 |
| AR | LRP1B | 1 |
| AR | LRRC37A7P | 1 |
| AR | LRRC4C | 1 |
| AR | LRRN1 | 1 |
| AR | LSP1 | 1 |
| AR | LYPD6 | 1 |
| AR | MAGED4 | 1 |
| AR | MAGED4B | 1 |
| AR | MAGEE1 | 1 |
| AR | MEIS1 | 1 |

| | | |
|---|---|---|
| AR | MIA3 | 1 |
| AR | MROH2A | 1 |
| AR | MRPS30-DT | 1 |
| AR | NAP1L2 | 1 |
| AR | NFE2L2 | 1 |
| AR | NFIA | 1 |
| AR | NKAIN2 | 1 |
| AR | NKAIN3-IT1 | 1 |
| AR | NR3C1 | 1 |
| AR | NR3C2 | 1 |
| AR | NSG1 | 1 |
| AR | NSUN7 | 1 |
| AR | OR56A5 | 1 |
| AR | PALM | 1 |
| AR | PAM | 1 |
| AR | PAPLN | 1 |
| AR | PBX3 | 1 |
| AR | PCDHB12 | 1 |
| AR | PCDHB16 | 1 |
| AR | PCDHB9 | 1 |
| AR | PDE4A | 1 |
| AR | PDIA3P1 | 1 |
| AR | PER3 | 1 |
| AR | PHYHIP | 1 |
| AR | PIAS2 | 1 |
| AR | PIK3R1 | 1 |
| AR | PKP4 | 1 |
| AR | PLAGL1 | 1 |
| AR | PLBD1-AS1 | 1 |
| AR | PLSCR4 | 1 |
| AR | PLTP | 1 |
| AR | PODXL2 | 1 |
| AR | PPIAP10 | 1 |
| AR | PPP1R1C | 1 |
| AR | PQLC2L | 1 |
| AR | PRDM11 | 1 |
| AR | PRDM5 | 1 |
| AR | PRKAA2 | 1 |
| AR | PRKRA-AS1 | 1 |
| AR | PRKX | 1 |
| AR | PRR33 | 1 |
| AR | RAI2 | 1 |
| AR | RAVER2 | 1 |
| AR | RFX3 | 1 |
| AR | RIMKLB | 1 |
| AR | RN7SKP184 | 1 |
| AR | RN7SKP251 | 1 |
| AR | RNF180 | 1 |
| AR | ROBO1 | 1 |
| AR | ROBO3 | 1 |

| | | |
|---|---|---|
| AR | RPSAP71 | 1 |
| AR | RRN3P1 | 1 |
| AR | SAMD12 | 1 |
| AR | SAMD5 | 1 |
| AR | SCD5 | 1 |
| AR | SCGB3A1 | 1 |
| AR | SDR16C5 | 1 |
| AR | SGCB | 1 |
| AR | SHF | 1 |
| AR | SLC16A2 | 1 |
| AR | SLC22A17 | 1 |
| AR | SLC25A12 | 1 |
| AR | SLC43A1 | 1 |
| AR | SLC5A12 | 1 |
| AR | SLITRK4 | 1 |
| AR | SMIM10L2A | 1 |
| AR | STK33 | 1 |
| AR | STPG2 | 1 |
| AR | TBC1D4 | 1 |
| AR | TBL1X | 1 |
| AR | TBX20 | 1 |
| AR | TLK1 | 1 |
| AR | TMEM246 | 1 |
| AR | TP63 | 1 |
| AR | TPTEP1 | 1 |
| AR | TRAM1L1 | 1 |
| AR | TRIM14 | 1 |
| AR | TRIM7 | 1 |
| AR | TSHZ2 | 1 |
| AR | TTC3P1 | 1 |
| AR | TTC6 | 1 |
| AR | UNC5B-AS1 | 1 |
| AR | USP13 | 1 |
| AR | USP44 | 1 |
| AR | USP51 | 1 |
| AR | VSIG1 | 1 |
| AR | VWDE | 1 |
| AR | WDR41 | 1 |
| AR | WDR72 | 1 |
| AR | WIPF3 | 1 |
| AR | WISP3 | 1 |
| AR | WNT5A | 1 |
| AR | XKR6 | 1 |
| AR | ZBED3 | 1 |
| AR | ZBTB16 | 1 |
| AR | ZC3HAV1 | 1 |
| AR | ZDHHC13 | 1 |
| AR | ZDHHC15 | 1 |
| AR | ZFAND4 | 1 |
| AR | ZFHX4 | 1 |

| | | |
|---|---|---|
| AR | ZMAT1 | 1 |
| AR | ZNF204P | 1 |
| AR | ZNF391 | 1 |
| AR | ZNF462 | 1 |
| AR | ZNF516 | 1 |
| AR | ZNF532 | 1 |
| AR | ZNF608 | 1 |
| AR | ZNF711 | 1 |
| AR | ZSCAN31 | 1 |
